# Transgelin: A New Gene Involved in LDL Endocytosis Identified by a Genome-wide CRISPR-Cas9 Screen

**DOI:** 10.1101/2020.12.23.424170

**Authors:** Diego Lucero, Ozan Dikilitas, Michael M. Mendelson, Promotto Islam, Edward B. Neufeld, Aruna T. Bansal, Lita A. Freeman, Boris Vaisman, Jingrong Tang, Christian A. Combs, Yuesheng Li, Szilard Voros, Iftikhar J. Kullo, Alan T. Remaley

## Abstract

To identify new genes involved in the cellular uptake of low-density lipoprotein (LDL), we applied a novel whole genome CRISPR/Cas9 knockout-screen on HepG2 cell lines. We identified *TAGLN* (transgelin), an actin-binding protein, as a new gene involved in LDL endocytosis. *In silico* validation demonstrated that genetically predicted differences in expression of *TAGLN* in human populations were associated with plasma lipids (triglycerides, total cholesterol, HDL, and LDL cholesterol) in the Global Lipids Genetics Consortium and lipid-related phenotypes in the UK Biobank. Decreased cellular LDL uptake observed in *TAGLN*-knockout cells due to decreased LDL receptor internalization, led to alterations in cellular cholesterol content and compensatory changes in cholesterol biosynthesis. Transgelin was also shown to be involved in the actin-dependent phase of clathrin-mediated endocytosis of other cargo besides LDL. The identification of novel genes involved in LDL uptake may improve the diagnosis of hypercholesterolemia and provide future therapeutic targets for the prevention of cardiovascular disease.

## INTRODUCTION

Epidemiological and Mendelian randomization studies have clearly demonstrated a causal role of low-density lipoprotein cholesterol (LDL-C) in atherosclerotic cardiovascular disease (ASCVD) [1]. Patients with genetically elevated LDL-C are at an especially high risk for ASCVD [1], due to their life-long elevations.

Familial hypercholesterolemia (FH), affecting between 1:250 to 1:300 individuals world-wide, is caused mainly by loss-of-function mutations in *LDLR* (LDL receptor), and less commonly in *APOB* and by gain-of-function mutations in *PCSK9* [2]. Other rare causative mutations in *STAP1* and *LDLRAP1* have also been described [2]; however, a known genetic mutation is not identified in 20 to 40% of patients with clear clinical criteria for FH [3–5]. In addition to monogenic FH, genome-wide association studies have not completely identified all the contributors to the heritability of circulating LDL-C in the general population, suggesting that additional genes could be involved in LDL plasma clearance.

Recent advances in genome editing technologies, especially the application of clustered regularly interspaced short palindromic repeats (CRISPR)/Cas9 technology, have allowed great progress in our understanding of gene function in eukaryotic cells [6]. A single guiding RNA (sgRNA) recognizes a specific genomic region, directing Cas9 nuclease to perform a double stranded break in DNA that is then mainly repaired by the imperfect non-homologous end joining process, generating random insertions and deletions (indels) at the cleavage site, leading to disruption of gene function [7]. Genome-wide pooled CRISPR/Cas9 knockout screens have recently been developed to identify new genes involved in different metabolic cellular pathways, such as in cancer drug-resistance [8], signal transduction [9], and cell fitness and viability [8, 10].

Our aim was to identify new genes involved in hepatic LDL uptake by applying a whole genome CRISPR/Cas9 knockout screen on a liver cell line (HepG2). In the present study, we identified *TAGLN*, an actin-binding protein, as a novel gene involved in cellular LDL uptake. Furthermore, we found that single nucleotide variants (SNV) in the human *TAGLN* locus, as well as genetically predicted expression levels of *TAGLN* are associated with elevated plasma lipids, thus validating our approach by revealing an important role for *TAGLN* in human lipoprotein metabolism. Moreover, we go on to demonstrate the cellular mechanism underlying the role of *TAGLN* in LDL metabolism. Our results potentially have both diagnostic and therapeutic implications for the prevention of ASCVD.

## MATERIALS AND METHODS

### Reagents

Stably Cas9-expressing HepG2 cell line (Cat# T3256), lentiviral sgRNA targeting human LDLR (Cat# 264181110204) and non-viral plasmids containing sgRNA targeting individual candidate genes were purchased from ABM Goods, Inc. (Canada). Whole-genome lentiviral pooled CRISPR/Cas9 knockout library (Cat# 73178-LV) was obtained from Addgene (Cambridge, MA). Alexa Fluor^TM^ 568 carboxylic acid succinyl ester, Alexa Fluor™ 488-Human Transferrin conjugate (Cat# T13342) and DiI–human LDL conjugate (Cat# L3482) were obtained from Molecular Probes (Eugene, OR). Primer kits and Master Mix for RT-qPCR were purchased from Applied Biosystems (Foster City, CA). Monensin, sodium salt, (Cat# M5273) was purchased from Sigma-Aldrich (St. Louis, MO). Anti-LDLR, antibody (Cat# 3839) was purchased from BioVision (Milpitas, CA), anti-TAGLN antibody (Cat# GT336) from Invitrogen (Carlsbad, CA), and anti-PCSK7 antibody (Cat# 19346S) from Cell Signaling Technology (Danvers, MA).

### Cell culture

Stably Cas9-expressing HepG2 cells (ABM Goods, Canada; Cat# T3256) were cultured in growing media: Dulbecco’s Modified Eagle Medium (DMEM) supplemented with 100 IU/ml of penicillin G, 100 μg/ml streptomycin and 10% (vol/vol) FBS. Cells were incubated in a humidified incubator with 5% CO2/95% air and split every 2 or 3 days.

### Preparation and fluorescent labeling of lipoproteins

Very low-density lipoproteins (VLDL) (*d*: 0.940 – 1.006 g/mL) and LDL (*d*: 1.019 – 1.064 g/mL) were isolated from healthy human plasma by preparative sequential KBr density gradient ultracentrifugation (330,000 × g) followed by extensive dialysis at 4°C against PBS to remove KBr. Purified VLDL or LDL (3 mg protein) were then incubated with Alexa Fluor^TM^ 568 carboxylic acid, succinyl ester (Molecular Probes, Eugene, OR) for 1 hour at room temperature in the dark. Free unbound dye was separated from lipoprotein-bound dye by preparative Fast Protein Liquid Chromatography (FPLC), using a HiTrap Desalting column (GE Healthcare, Chicago, IL). Labeled lipoproteins were collected in the void volume.

### Generation of LDLR knockout (LDLR-KO) HepG2 Cells

Stably Cas9-expressing HepG2 cells were transduced at a multiplicity of infection (MOI) of 1.5 with lentivirus containing a sgRNA targeting human LDLR (ABM Goods, Inc. Canada, Cat# 264181110204) and a neomycin selection marker. Transduced cells were selected with neomycin (1 mg/ml) for 14 days. After selection, cells were sorted into 96-well plates (one cell per well) in an BD FACSAria™ III Cell Sorter. Clones were expanded, and loss of LDLR was tested at the mRNA and protein levels by RT-qPCR and Western blotting respectively (Figure 2A). Additionally, phenotype was confirmed by fluorescent LDL uptake, showing ~75% reduction in LDL uptake (Supplementary Figure 1B).

**Figure 1:**
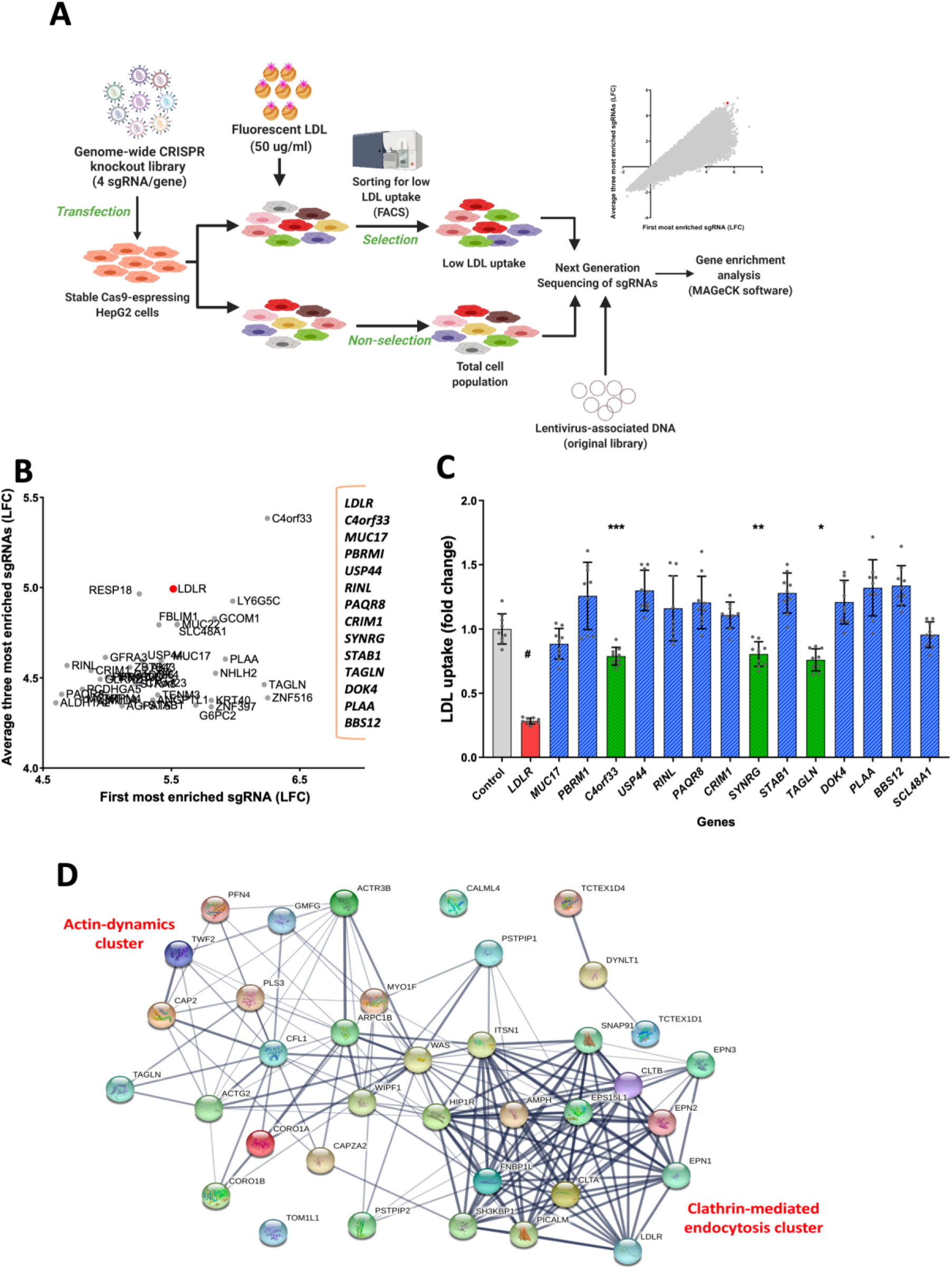
CRISPR-CAs9 screen. (A). Schematic for genome-wide CRISPR screen. HepG2 cells with stable Cas9 expression were transformed with lentivirus containing a genome-wide sgRNA library (76,441 sgRNAs, targeting 19,114 genes; 4 sgRNA/gene). A fraction of transformed cells were not selected and were frozen for further analysis. The remainder of transformed cells were incubated with 50 ug/ml fluorescent LDL for 4 h and sorted for low LDL uptake by flow cytometry. Next generation sequencing was used to detect sgRNA in each sample. Enrichment of sgRNA in cells and library was determined with MAGeCK software. Gene were ranked by the average of log fold change (LFC) of the three most enriched sgRNA for each gene compared to the non-selected cell population. (B) Plot of LFC of average for the three best sgRNA vs. LFC of the most enriched single sgRNA of the top ranked 40 genes. Selected candidate genes indicated on the right. (C) LDL uptake in CRISPR-generated stable knockout clones of the 15 candidates in B. (D) Interactome obtained with endocytosis genes that showed at least 1.5 LFC in the genome-wide CRISPR knock-out screen. Interactome was constructed using STRING database (http://string-db.org/) with moderate confidence (0.40). #p<0.001 vs Control. *p=0.001 vs. control. **p=0.013 vs. control. ***p=0.007 vs control. One-Way ANOVA.

**Figure 2:**
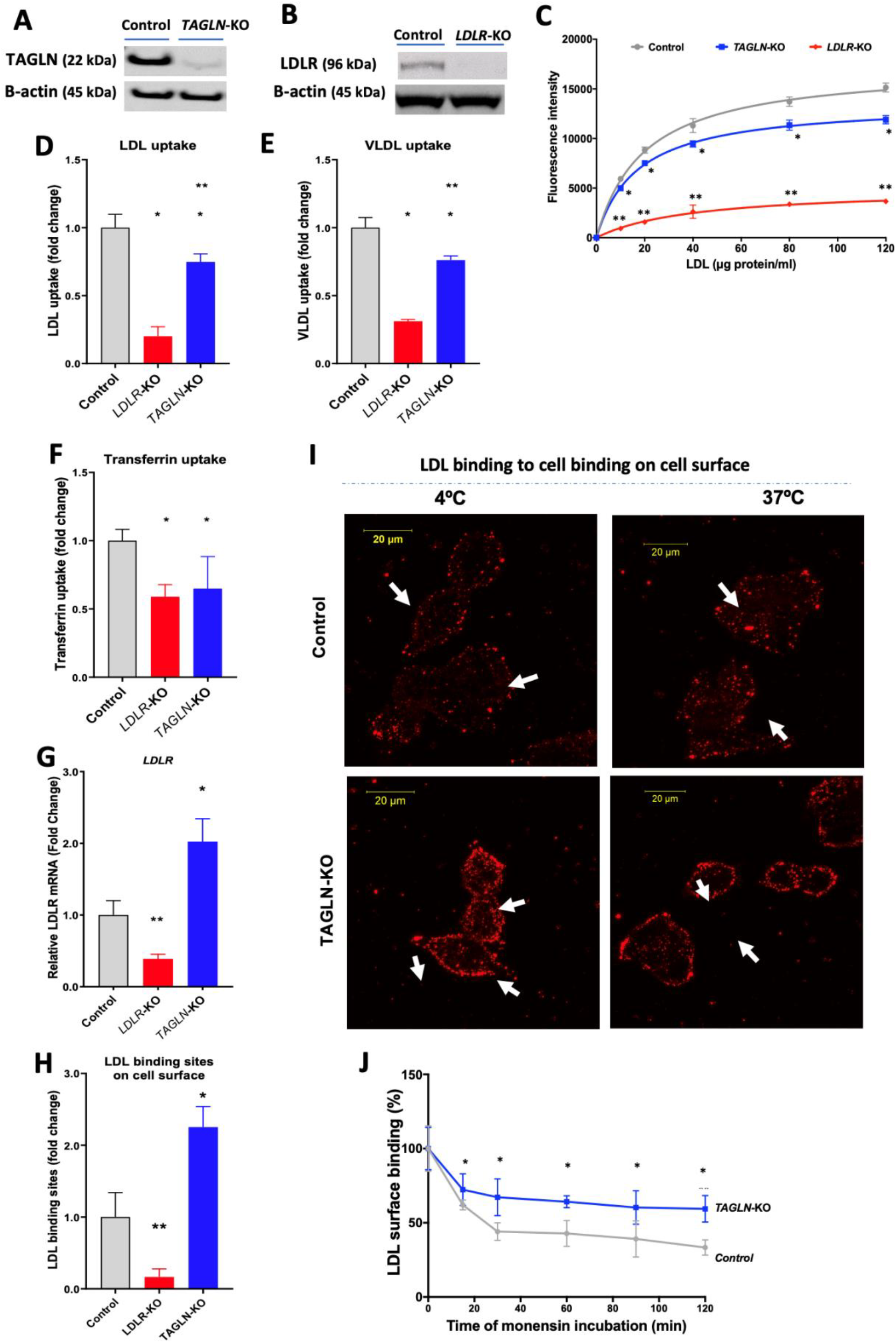
Reduced uptake of low-density lipoprotein (LDL), very low-density lipoprotein (VLDL) and transferrin and reduced LDLR internalization in TAGLN-KO cells. (A) and (B) Western blots for transgelin (TAGLN) and LDL receptor (LDLR) in HepG2 cells, respectively. Cells were transfected with scrambled sgRNA (Control) and sgRNA targeting *TAGLN* and *LDLR*. (C) Control, *TAGLN* knockout (*TAGLN*-KO) and *LDLR* knockout (*LDLR*-KO) HepG2 cells were incubated with increasing concentration of fluorescent LDL. Fluorescence intensity, proportional to LDL internalization, was determined by FACS. (D - F) Control, *LDLR*-KO, and *TAGLN*-KO HepG2 cells were incubated with: (D), fluorescent LDL (40 μg/mL); (E), fluorescent VLDL (50 μg/mL); and (F), transferrin (25 μg/mL). Fluorescence intensity, proportional to probe internalization, was determined by FACS. (G) mRNA expression levels of LDLR were determined in Control, *TAGLN*-KO and *LDLR*-KO HepG2 cells by RT-PCR. (H) Cells were incubated with 10 μg/ml DiI-LDL for 4 h at 4ºC and cell-associated fluorescence was then quantified by FACS. (I) Control and *TAGLN*-KO cells were incubated with 10 μg/ml DiI-LDL for 4 h at 4ºC and then, after thorough washes with cold PBS, imaged by confocal microscopy (upper and lower left panels). After cold PBS washes, a batch of control and *TAGLN*-KO cells was allowed to warm up to 37ºC for 2 h to allow LDL/LDLR internalization and then imaged by confocal microscopy (upper and lower right panels). Images represent the maximum intensity projection of the z-stack and are representative of three independent experiments. White arrows indicate LDL on plasma membrane. (J) Control and *TAGLN*-KO cells were incubated with 30 μM monensin at the indicated time points to block LDLR recycling and were then incubated with 10 μg/ml DiI-LDL for 4 h at 4ºC. After thorough washes with cold PBS, cells were then dissociated with non-enzymatic buffer and cell-associated fluorescence was quantified by FACS. *p<0.05 vs control. ** p<0.05 vs control and TAGLN-KO. Student’s t-test or one-way ANOVA as appropriate.

### Pooled CRISPR/Cas9 knockout screen

Stably Cas9-expressing HepG2 cells were transduced with genome-wide lentiviral pooled CRISPR/Cas9 knockout library (Addgene #73178) (Brunello; Cat# 73178-LV) (Addgene, Inc., Watertown, MA) containing 76,441 sgRNAs, targeting 19,114 (4 sgRNA/gene) [11]. Approximately 10^8^ cells were transduced with lentiviral library, considering that each sgRNA is represented by 400 cells, and at a low MOI (~0.3) to ensure that majority of cells were infected by a single viral particle [12]. After overnight infection, media was changed for fresh growing media for 24 h. Transduced cells were then selected in puromycin (4 μg/ml) for 7 days. Finally, cells were incubated 4 h with fluorescent LDL at 37°C, washed and dissociated by trypsinization. One portion of unsorted cells was kept as non-selected cells, and the rest were sorted according to LDL uptake in an BD FACSAria™ III Cell Sorter (Figure 1A). Fluorescent LDL signal associated with wild type and *LDLR*-KO cells was used as reference to establish laser gates. Cells with 5% lower LDL uptake were collected as those with reduced LDL uptake.

After sorting, cells were cultured for 72 h, and genomic DNA was extracted using a Maxwell® RSC Cell DNA Purification Kit (Promega, Madison, WI), following manufacturer’s instructions. SgRNAs integrated into the genome were PCR-amplified using primers harboring the Illumina TruSeq adapters, P5 and P7. PCR products were separated by preparative gel electrophoresis (1% agarose). Specific bands with the expected amplicon size (354 nt) were purified from gel by Zymoclean Large Fragment DNA Recovery, following manufacturer’s instructions. Purified fragments, containing sgRNA sequences, were sequenced in a HiSeq 2500 (Illumina, San Diego, CA). Fastq files were processed for sgRNA and gene enrichment using MAGeCK software [13]. Genes were ranked according to the average log fold change of the best three sgRNAs targeting each gene (Figure 1B).

### Validation of candidates

Each candidate was validated by generating individual stable knockouts and re-testing LDL uptake. Stably Cas9-expressing HepG2 cells were transfected with plasmids containing sgRNA sequence (Supplementary Table 3) and green fluorescent protein (GFP) as a selection marker, using Lipofectamine 3000 (ThermoFisher, Waltham, MA), following manufacturer’s instructions. After 72 h, transfected cells (GFP positive) were sorted into 96-well plates, one cell per well. Clones were expanded and loss of each gene expression was tested by RT-qPCR. Only those clones with significant reduction of mRNA levels were considered for further analysis. Clones for each gene were individually evaluated for LDL uptake as indicated below.

### Association of lipid traits with genetically predicted expression of TAGLN, SYNRG, and C4orf33

We performed association analysis between genetically predicted expression of *TAGLN, SYNRG, C4orf33* and the lipid traits LDL-C, HDL-C, TC, and TG. We used MASHR-based transcriptome prediction models trained on 49 tissues in the GTEx v8 release data [14] which are available online (http://predictdb.org/). We obtained genome-wide association summary statistics from Global Lipids Genetics Consortium [15] for these lipid traits and imputed them to include all variants present within GTEx reference. We performed tissue-stratified associations using S-PrediXcan [16] for each gene-lipid trait pair and aggregated them in S-MultiXcan [17], which jointly fits the trait on predicted gene expression across multiple tissues to leverage substantial sharing of eQTLs between them. Gene-lipid trait associations were deemed significant at a Bonferroni threshold of two-sided *p-value* <4.2×10^-3^ (0.05/[number of genes tested × number of traits tested]). Harmonization, imputation of summary statistics, and association testing were all performed using the collection of tools available within MetaXcan following their recommended best practice guidelines (https://github.com/hakyimlab/MetaXcan).

### Search for SNV associations at candidate gene loci in previous GWAS

We interrogated previously reported GWAS for SNVs mapped to the three candidate genes identified in the CRISPR screen (C4orf33, *SYNRG* and *TAGLN*) using the National Heart, Lung and Blood Institute (NHLBI) Genome-Wide Repository of Associations Between SNPs and Phenotypes (GRASP v2.0.0.0) [18]. The GRASP v2.0.0.0 catalog includes 8,872,472 genotype-phenotype results at a nominal p-value <0.05 comprising 186,362 unique phenotypes from 177 broad phenotype categories as reported in 2,082 GWAS studies. Our primary search focused on lipid-related phenotypes and coronary artery disease. Our secondary search broadened to all reported phenotypes.

### Phenome-wide scan using genetically predicted expression of TAGLN

To evaluate potential pleiotropic effects of genetically predicted expression of *TAGLN*, we used PhenomeXcan database [Ref https://www.biorxiv.org/content/10.1101/833210v3 and https://www.biorxiv.org/content/10.1101/814350v2, which synthesizes ~8.8 million variants from GWAS on 4,091 traits with gene expression regulation data from GTEx v8 into a transcriptome-wide association resource including 22,255 genes. We filtered pre-computed S-MultiXcan association statistics of *TAGLN* to include available clinical phenotypes ascertained through self-report, laboratory measurements, medication data, and EHR data in the UK Biobank cohort. We deemed associations significant at a Bonferroni threshold of two-sided *p-value* <1×10^-5^.

### Cellular lipoprotein uptake

Control, *TAGLN*-KO and *LDLR*-KO cells were plated in 96-well plates, at a density 20,000 cells/well, and incubated for 48 h in DMEM supplemented with 100 IU/ml of penicillin G, 100 μg/ml streptomycin and 10% (vol/vol) FBS to allow cells to recover the expression of surface receptors. Cells were then incubated with the indicated concentrations of fluorescently labeled LDL or VLDL in serum-free media containing 0.1% BSA (Sigma-Aldrich, St. Louis, MO) for 4 h at 37°C. Cells were then washed with PBS, dissociated by trypsinization, and resuspended in ice-cold PBS containing 0.5% BSA/2.5 μM EDTA. Uptake, proportional to mean fluorescence intensity, was then quantified by fluorescence activated cell sorter (FACS) flow cytometry in a BD LSRFortessa (BD Biosciences, Franklin Lakes, NJ). Typically, the number of counted events was at least 10,000.

### Cellular transferrin uptake

Control, *TAGLN*-KO and *LDLR*-KO cells were seeded onto 12-well plates, and then incubated for 48 hours to allow for recovery of transferrin receptors prior to the experiment. After washing with warm PBS, and incubation with DMEM, containing 0.1% BSA and 25 mM HEPES for 30 minutes, cells were then incubated with 25 μg/ml transferrin in DMEM, 0.1% BSA, 25 mM HEPES for 45 minutes at 37°C. To avoid transferrin re-secretion, cells were placed on ice, washed 3 times with ice-cold PBS, dissociated by brief trypsinization, and then fixed with 4% paraformaldehyde. Uptake, proportional to mean fluorescence intensity, was then quantified by FACS flow cytometry in a BD LSRFortessa (BD Biosciences, Franklin Lakes, NJ). At least 10,000 events were counted.

### Immunoblot analysis

Protein lysates were prepared from control, *LDLR*-KO and *TAGLN*-KO cells by incubation with RIPA buffer in the presence of Halt^TM^ protease and Halt^TM^ phosphatase inhibitors (Thermo Fisher). Western blots were performed as previously described [19] using rabbit anti-human LDLR (1:500) (BioVision, Inc., Milpitas, CA), TAGLN (1:1000), PCSK7 (1:1000) (Cell Signaling Technology, Inc., Danvers, MA), and beta-actin (1:10,000) antibodies. The secondary antibody was donkey anti-rabbit antibody conjugated to HRP (Abcam, Cambridge, UK).

### RNA isolation, expression analysis and RNA-sequencing

Control, *TAGLN*-KO and *LDLR*-KO cells were preincubated 72 h in DMEM supplemented with 10% lipoprotein depleted serum, containing 50 μg/ml of fresh human LDL. Then RNA was extracted using a Qiagen RNeasy® Mini Kit (Qiagen, Hilden, Germany), according to the manufacturer’s instructions. Total RNA concentration was measured using a Nanodrop (ThermoFisher, St. Louis, MO). To quantitate expression of selected genes by RT-qPCR, cDNA was first synthesized from 200 ng of total RNA, using the QuantiTect Reverse Transcription Kit (Qiagen, Hilden, Germany). RT-qPCR was carried out using TaqMan Gene Expression Assays and TaqMan Master Mix (Applied Biosystems, Foster City, CA). 18S (Hs03003631_g1) was used as an internal control. The primer kits used for RT-qPCR can be found in Supplementary table 4.

### Cellular cholesterol assay

For lipid extraction, 3 × 10^6^ cells were plated into a 10 mm dish. Following overnight incubation for cell adherence, cells were washed twice with warm PBS and preincubated for 72 h in DMEM supplemented with 10% lipoprotein depleted serum, containing 50 μg/ml of LDL. Cells were then washed twice with ice-cold PBS, and total lipids were extracted into hexane:isopropanol (3:2) for 30 minutes at RT. The extraction procedure was repeated three times. The combined solvents containing extracted lipids was transferred to an Eppendorf tube and dried under nitrogen. The lipid residue was resuspended in 200 μL of 5% Triton X-100. Total cholesterol was measured by enzymatic colorimetric method (Wako Cholesterol E; Wako, Japan), subtracting a 5% Triton X-100 blank to eliminate non-specific absorbance. Protein residue, precipitated after lipid, extraction, was solubilized with 0.1 N NaOH and protein concentration was determined using Pierce™ BCA Protein Assay Kit (Thermo Fisher, Waltham, MA). Cellular cholesterol content data was expressed as mg cholesterol/mg of protein, and data was normalized to that of the control. Each extraction was performed in triplicate and each experiment was repeated independently at least 3 times.

### Evaluation of LDL binding sites on cell surface

Control and *TAGLN*-KO cells were plated in glass coverslip-bottomed plastic dishes (Mat Tek Co, Ashland, MA) containing 2.5 × 10^5^ cells/dish and grown in complete growth media for 48 h to allow recovery of LDLR on the cell surface. Cells were then washed twice with ice cold PBS and incubated for 4 h in pre-chilled DMEM supplemented with 0.1% BSA at 4°C with 50 μg/ml LDL containing 20% DiI-LDL (Molecular Probes, Eugene, OR). Next, cells were thoroughly washed with ice-cold PBS and fluorescent LDL bound on the cell surface was imaged with a Zeiss 880 confocal microscope (Jena, Germany) using a Zeiss 40x Plan-Apochromat objective lens (N.A. 1.3). DiI-LDL (Molecular Probes, Eugene, OR) fluorescence in labelled cells was acquired using 561nm excitation, an emission bandwidth of 565-700nm, a pinhole set to 1 A.U., a pixel format that varied with optical zoom to provide a lateral pixel size of 110nm, and an interslice thickness of 640nm (total slice number/ stack varied with cell height). In addition, a portion of the cells, after thorough washing with cold PBS, were incubated at 37°C for 2 hours in complete growth media to allow LDL internalization, and then imaged as described above.

### Monensin treatment and analysis of LDL binding sites on the cell surface

Control and *TAGLN*-KO cells were plated in 96-well plates at a density of 20,000 cells/well and cultured for 48 h to allow expression of LDLR on cell surface to recover. Cells were treated with 30 μM monensin for 0, 15, 30, 60, 90 or 120 minutes, and then treated with DiI-LDL in the cold, as described above (*Evaluation of LDL binding sites on cell surface*). Cells were then dissociated with an enzyme-free cell dissociation buffer (Gibco, Gaithersburg, MD) and prepared for flow cytometry. Cell surface-associated fluorescence was quantified by FACS flow cytometry in a BD LSRFortessa (BD Biosciences, Franklin Lakes, NJ). The number of counted events was at least 10,000. The amount of fluorescence associated with cell surface was expressed as % of the fluorescence associated with non-treated cells (no monensin).

### Statistical analysis

Data is presented as mean ± SD. Differences between groups were tested by Student’s t-test and one-way ANOVA, followed by Dunnett’s test, depending on each experiment design and two-sided *p-values* <0.05 were considered statistically significant.

## RESULTS

### Whole genome CRISPR/Cas9 knockout screen of hepatic cellular LDL uptake

To identify new genes involved in LDL uptake in liver cells, we applied a genome-wide knockout CRISPR screen on HepG2 cells, using a lentiviral genome-scale CRISPR loss-of-function library (Brunello; Addgene, Watertown, MA), containing 76,441 sgRNAs, targeting 19,114 genes (4 sgRNA/gene), and 1000 non-targeting control sgRNAs [11]. Transduced cells were incubated for 4 hours with fluorescent LDL. A fraction of unsorted, cells was kept as non-selected cells, while the rest were sorted according to the degree of cellular LDL uptake. Cells with less than the 5^th^ percentile of fluorescence LDL uptake were selected. Cut-point for cell selection was determined based on the LDL fluorescence in wild type HepG2 cells and in HepG2 cells in which the LDL-receptor (LDLR) was specifically knocked out with a sgRNA targeting LDLR (Supplementary Figure 1), which typically led to a ~75% reduction in LDL uptake. Subsequently, deep sequencing was applied to determine the sgRNA representation in both cell groups (low LDL uptake selected cells and non-selected cell population) and in the original sgRNA library. An overall general outline of our screening approach can be found in Figure 1A.

If a gene influences a phenotype, its sgRNAs should be among the most enriched in a CRISPR screening of any selected cell [20]. Since LDLR clearly affects cellular LDL uptake, we examined the enrichment of its 4 sgRNAs that were designed *in silico* in the construction of the original library. Three of the 4 sgRNAs in the library were among the most enriched (above 95^th^ percentile) in the group of cells with lower LDL uptake (Supplementary Figure 2A). In contrast, for the non-sorted control cells, sgRNAs targeting *LDLR* were randomly distributed among all sgRNAs (Supplementary Figure 2B).

Enriched genes from the screen were ranked by averaging the log-fold change (LFC) of their three most enriched sgRNAs to minimize the possible influence of off-target effects on gene classification and because of the known variability in the effectiveness of the individual sgRNAs in the library for their target (Supplementary Table 1). Figure 1B shows the plot of average of LFC of the 3 best sgRNA for each gene versus the LFC of most enriched sgRNA. *LDLR* was on top of the list (Supplementary Figure 2C). Possible other candidate genes involved in LDL uptake were selected by searching for those genes, previously associated with LDL metabolism or were biologically plausible based on their known function. Genes associated with cell growth or apoptosis could potentially be confounders and were not considered. A total of 15 candidate genes were selected for further validation (Figure 1B) after confirmation of their enrichment in an independent replicate screen (Supplementary Figure 2 D and E).

Cellular uptake of fluorescent LDL was tested by flow cytometry in stable knockout cells for each individual candidate gene (Figure 1C). As expected, *LDLR* showed the most robust effect, with a reduction of approximately 80% in cellular LDL uptake (p=0.0002) when compared to control cells (wild type). Although small reductions were observed for several of the candidate genes, consistent reductions in cellular LDL uptake were only observed for the following 3 candidate genes: *SYNRG* (-19%, p=0.013), *C4orf33* (-21%, p=0.007), and *TAGLN* (-25%, p=0.001).

### In silico validation of lipid phenotypes in human populations

To further validate our *in vitro* findings, we used a computational approach where we performed an association analysis between genetically predicted expression of these three candidate genes, i.e. genetic component of the expression levels predicted using common cis-eQTLs, and lipid traits, namely LDL-C, high-density-lipoprotein cholesterol (HDL-C), total cholesterol (TC), and triglycerides (TG). This was done using the Global Lipids Genetics Consortium GWAS summary statistics [15] and transcriptome prediction models trained on Genotype-Tissue Expression (GTEx) v8 [14] data (see Methods). Genetically predicted expression levels of *TAGLN* were negatively and significantly associated with total and LDL cholesterol, and triglycerides at the Bonferroni threshold (*pLDL-C*=2.8×10^-5^, *pTC*=1.2×10^-19^, *pTG*=2.0×10^-57^), and positively with HDL-C (*pHDL-C*=2.3×10^-14^). We did not detect significant associations between *SYNRG* and *C4orf33* for any of the lipid traits examined (*p*>0.18) (Table 1). Additionally, we examined the association of SNV at the 3 *in vitro* validated gene loci (*SYNRG*, *C4orf33*, and *TAGLN*) with plasma lipids in large human populations, using summary statistics from genome-wide association studies (GWAS) (Table 2). Of our 3 candidate genes, SNVs mapped in the *TAGLN* locus (rs17120434, rs2075547, rs2269397, rs508487 and rs641620) had significant associations with elevated triglycerides, total cholesterol and LDL-C (Table 2). Additionally, three SNV in the *TAGLN* locus also had reported associations with soluble transferrin receptor (sTfR): rs508487 (*p*=3×10^-20^), rs641620 (*p*=9.8×10^-15^) and rs664971 (*p*=8.5×10^-17^), indicating that, in addition to lipoprotein metabolism, this locus might also be involved in transferrin/iron metabolism. No significant associations were observed for the other two candidate genes for any lipid variable.

**Table 1.** Association of lipid traits with genetically predicted expression of *TAGLN*,*SYNRG*, and *C4orf33*.

| Lipid trait | Gene | Gene Band | N<sub>tissues</sub> | N<sub>indep</sub> | z-score, mean (SD)* | p-value** |
| --- | --- | --- | --- | --- | --- | --- |
| LDL-C | *TAGLN* | 11q23.3 | 47 | 5 | -0.177 (1.844) | **2.8x10<sup>-5</sup>** |
| HDL-C | *TAGLN* | 11q23.3 | 47 | 5 | 0.082 (4.590) | **2.3x10<sup>-14</sup>** |
| Total cholesterol | *TAGLN* | 11q23.3 | 47 | 5 | -0.545 (3.476) | **1.2x10<sup>-19</sup>** |
| Triglycerides | *TAGLN* | 11q23.3 | 47 | 5 | -1.814 (5.529) | **2.0x10<sup>-57</sup>** |
| LDL-C | *SYNRG* | 17q12 | 47 | 3 | 0.610 (0.695) | 0.328 |
| HDL-C | *SYNRG* | 17q12 | 47 | 3 | 0.862 (0.438) | 0.691 |
| Total cholesterol | *SYNRG* | 17q12 | 47 | 3 | 1.571 (0.284) | 0.181 |
| Triglycerides | *SYNRG* | 17q12 | 47 | 3 | 0.631 (0.420) | 0.770 |
| LDL-C | *C4orf33* | 4q28.2 | 48 | 3 | 0.778 (0.685) | 0.386 |
| HDL-C | *C4orf33* | 4q28.2 | 48 | 3 | -0.098 (0.482) | 0.456 |
| Total cholesterol | *C4orf33* | 4q28.2 | 48 | 3 | 0.367 (0.351) | 0.852 |
| Triglycerides | *C4orf33* | 4q28.2 | 48 | 3 | -0.306 (0.666) | 0.408 |
| N<sub>tissues</sub> – number of tissues available to S-MultiXcan when computing significance for a gene; N<sub>indep</sub> – number of independent components of variations among N<sub>tissues</sub> |  |  |  |  |  |  |
| * Mean z-score and its standard deviation (SD) among single-tissue S-PrediXcan associations |  |  |  |  |  |  |
| ** P-value of S-MultiXcan association, significant associations are bolded. |  |  |  |  |  |  |

**Table 2:**
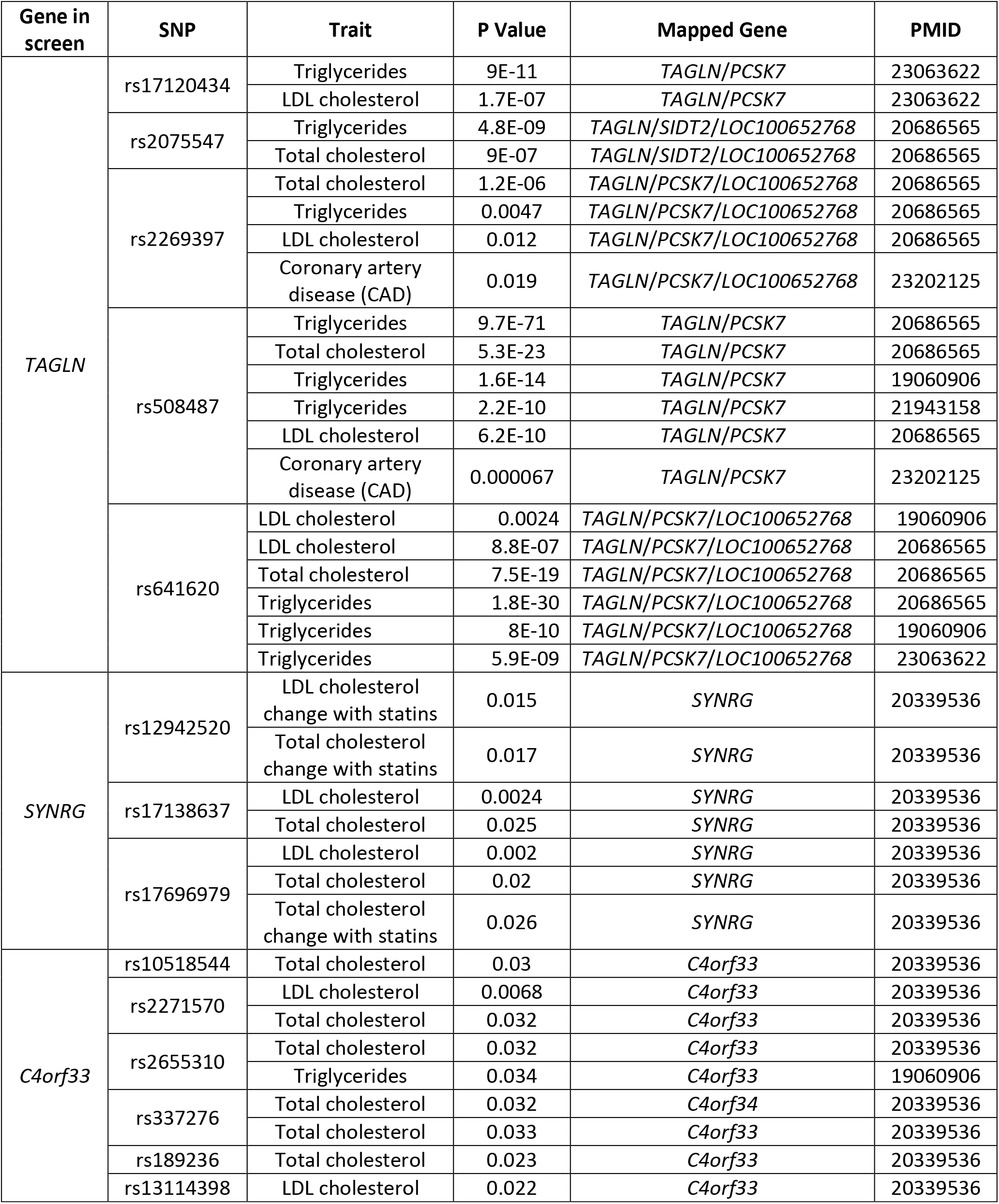
Associations of single nucleotide variants in *LDLR*,*TAGLN*,*C4orf33* and *SYNRG* with plasma lipids, obtained in data analysis from previous genome-wide analysis studies (GWAS).

| Gene in screen | SNP | Trait | P Value | Mapped Gene | PMID |
| --- | --- | --- | --- | --- | --- |
| TAGLN | rs17120434 | Triglycerides | 9E-11 | TAGLN/PCSK7 | 23063622 |
|  |  | LDL cholesterol | 1.7E-07 | TAGLN/PCSK7 | 23063622 |
|  | rs2075547 | Triglycerides | 4.8E-09 | TAGLN/SIDT2/LOC100652768 | 20686565 |
|  |  | Total cholesterol | 9E-07 | TAGLN/SIDT2/LOC100652768 | 20686565 |
|  | rs2269397 | Total cholesterol | 1.2E-06 | TAGLN/PCSK7/LOC100652768 | 20686565 |
|  |  | Triglycerides | 0.0047 | TAGLN/PCSK7/LOC100652768 | 20686565 |
|  |  | LDL cholesterol | 0.012 | TAGLN/PCSK7/LOC100652768 | 20686565 |
|  |  | Coronary artery disease (CAD) | 0.019 | TAGLN/PCSK7/LOC100652768 | 23202125 |
|  | rs508487 | Triglycerides | 9.7E-71 | TAGLN/PCSK7 | 20686565 |
|  |  | Total cholesterol | 5.3E-23 | TAGLN/PCSK7 | 20686565 |
|  |  | Triglycerides | 1.6E-14 | TAGLN/PCSK7 | 19060906 |
|  |  | Triglycerides | 2.2E-10 | TAGLN/PCSK7 | 21943158 |
|  |  | LDL cholesterol | 6.2E-10 | TAGLN/PCSK7 | 20686565 |
|  |  | Coronary artery disease (CAD) | 0.000067 | TAGLN/PCSK7 | 23202125 |
|  | rs641620 | LDL cholesterol | 0.0024 | TAGLN/PCSK7/LOC100652768 | 19060906 |
|  |  | LDL cholesterol | 8.8E-07 | TAGLN/PCSK7/LOC100652768 | 20686565 |
|  |  | Total cholesterol | 7.5E-19 | TAGLN/PCSK7/LOC100652768 | 20686565 |
|  |  | Triglycerides | 1.8E-30 | TAGLN/PCSK7/LOC100652768 | 20686565 |
|  |  | Triglycerides | 8E-10 | TAGLN/PCSK7/LOC100652768 | 19060906 |
|  |  | Triglycerides | 5.9E-09 | TAGLN/PCSK7/LOC100652768 | 23063622 |
| SYNRG | rs12942520 | LDL cholesterol change with statins | 0.015 | SYNRG | 20339536 |
|  |  | Total cholesterol change with statins | 0.017 | SYNRG | 20339536 |
|  | rs17138637 | LDL cholesterol | 0.0024 | SYNRG | 20339536 |
|  |  | Total cholesterol | 0.025 | SYNRG | 20339536 |
|  | rs17696979 | LDL cholesterol | 0.002 | SYNRG | 20339536 |
|  |  | Total cholesterol | 0.02 | SYNRG | 20339536 |
|  |  | Total cholesterol change with statins | 0.026 | SYNRG | 20339536 |
| C4orf33 | rs10518544 | Total cholesterol | 0.03 | C4orf33 | 20339536 |
|  | rs2271570 | LDL cholesterol | 0.0068 | C4orf33 | 20339536 |
|  |  | Total cholesterol | 0.032 | C4orf33 | 20339536 |
|  | rs2655310 | Total cholesterol | 0.032 | C4orf33 | 20339536 |
|  |  | Triglycerides | 0.034 | C4orf33 | 19060906 |
|  | rs337276 | Total cholesterol | 0.032 | C4orf34 | 20339536 |
|  |  | Total cholesterol | 0.033 | C4orf33 | 20339536 |
|  | rs189236 | Total cholesterol | 0.023 | C4orf33 | 20339536 |
|  | rs13114398 | LDL cholesterol | 0.022 | C4orf33 | 20339536 |

It is important to note that the *TAGLN* sequence partially overlaps tail-to-tail with the antisense *PCSK7*, raising the possibility that some of the above-mentioned SNV may be related to *PCSK7*. We, therefore, analyzed whether the sgRNAs targeting *TAGLN* in the library also targeted *PCSK7*. The sgRNAs for *TAGLN* targeted its exons 2 and 3, as well as *PCSK7* on its terminal Intron 19 in its 3’-UTR region (Supplementary Figure 3). In contrast, none of the sgRNAs targeting *PCSK7* were mapped close to the coding region of *TAGLN*. Their targeting sequences are all about 10 Kb downstream from *TAGLN*’s 3’-UTR (Supplementary Figure 3). Furthermore, none of these *PCSK7* sgRNAs showed enrichment in the screen (LFC= 3.59; -1.04; -2.50 and -3.078). To generate stable *TAGLN* knockout (*TAGLN*-KO) cells a fifth independent sgRNA, different to those in the screen, library, was used. Its targeted sequence (CACTTCGCGGCTCATGCCAT) maps to *TAGLN* on Exon 2 and on *PCSK7* to its terminal Intron 19 in the 3’-UTR region. Using this fifth *TAGLN* sgRNA, we found that PCSK7 protein expression was unchanged or, in fact, slightly increased in *TAGLN*-KO cells (Supplementary Figure 4), indicating that the observed reduction in cellular LDL uptake depends upon *TAGLN* and not on *PCSK7*.

Since *TAGLN* was the only one among the three candidates to show significant associations with plasma lipid levels at both the gene-level and SNV-level analyses, we further evaluated its association of genetically predicted expression of *TAGLN* at a phenome-wide scale (PheWAS) in the UK Biobank (UKBB), using PhenomeXcan database (see Methods). Below the Bonferroni significance threshold, predicted expression of *TAGLN* was associated with 18 phenotypes in the UKBB. Of the 18 associations, 10 were related to abnormal lipid metabolism and related phenotypes, such as high cholesterol, lipid lowering medication use, waist circumference, body fat percentage, and body mass index (Table 3). On the other hand, the 8 remaining associations were related to hematologic traits, such as platelet count, hematocrit, red blood cell and platelet distribution widths, and eosinophil count (Table 3). These associations suggest that this locus (11q23.3) could additionally affect erythrocytes, beyond just sTfR metabolism.

**Table 3.** Phenome-wide association of genetically predicted expression of *TAGLN* in the UK Biobank.

| Phenotype | N<sub>tissues</sub> | N<sub>indep</sub> | Best sign | Consensus sign | *p*-value* |
| --- | --- | --- | --- | --- | --- |
| Platelet count | 47 | 5 | + | + | 3.5x10<sup>-16</sup> |
| Platelet distribution width | 47 | 5 | - | - | 3.5x10<sup>-13</sup> |
| Eosinophil count | 47 | 5 | + | - | 1.9x10<sup>-10</sup> |
| Self-reported high cholesterol | 47 | 5 | - | + | 1.3x10<sup>-9</sup> |
| Red blood cell (erythrocyte) distribution width | 47 | 5 | + | - | 2.8x10<sup>-9</sup> |
| Platelet crit | 47 | 5 | + | + | 1.4x10<sup>-8</sup> |
| Leg fat percentage (right) | 47 | 5 | - | - | 5.2x10<sup>-8</sup> |
| Mean platelet (thrombocyte) volume | 47 | 5 | - | - | 5.9x10<sup>-8</sup> |
| Eosinophil percentage | 47 | 5 | + | - | 6.5x10<sup>-8</sup> |
| Leg fat percentage (left) | 47 | 5 | - | - | 1.6x10<sup>-7</sup> |
| Cholesterol lowering medication use | 47 | 5 | - | - | 1.0x10<sup>-6</sup> |
| Waist circumference | 47 | 5 | - | - | 1.1x10<sup>-6</sup> |
| Body fat percentage | 47 | 5 | - | - | 2.3x10<sup>-6</sup> |
| Leg fat mass (right) | 47 | 5 | - | - | 2.7x10<sup>-6</sup> |
| Haematocrit percentage | 47 | 5 | + | + | 3.0x10<sup>-6</sup> |
| Treatment/medication code: fenofibrate | 47 | 5 | - | - | 3.4x10<sup>-6</sup> |
| Body mass index (BMI) | 47 | 5 | - | - | 4.2x10<sup>-6</sup> |
| Leg fat mass (left) | 47 | 5 | - | - | 4.5x10<sup>-6</sup> |
| N<sub>tissues</sub> – number of tissues available to S-MultiXcan when computing significance for a gene; N<sub>indep</sub> – number of independent components of variations among N<sub>tissues</sub>; Best sign – the sign of effect of the most significant tissue where ‘+’ (‘-’) means higher (lower) expression is associated with higher risk or higher value of the phenotype; Consensus sign – the sign of effect of the consensus among those tissues with *p*-value <1x10<sup>-4</sup> where ‘+’ (‘-’) means the same as in the ‘Best sign’ column<br>* Obtained using S-MultiXcan on GWAS summary statistics of UK Biobank phenotypes |  |  |  |  |  |

### Role of endocytosis-related genes in LDL uptake

Internalization of LDLR occurs through clathrin-mediated endocytosis (CME) [21], which is a complex process that comprises not only proteins associated with the formation of the clathrin-coated pit, but also scission proteins and actin filament-associated proteins, [22]. We, therefore, investigated whether endocytosis-related genes, in general, were significantly enriched in our screen and consequently associated with reduced cellular LDL uptake in HepG2 cells. We first used the list of endocytosis-related genes compiled by Lacy *et al* [23], including 105 genes related to each step in the endocytic pathway. Thirty-seven of these genes showed at least 1.5 LFC in our screen (Supplementary Table 2). Figure 1D shows the interactome between these genes and *LDLR*, showing two interconnected clusters: CME and actin-dynamics. Numerous studies have demonstrated that the coordinated function of both clusters is essential to ensure LDL internalization [24–26]. Among all endocytosis-related genes, *TAGLN*, however, showed the highest average enrichment (Average LFC: 4.39).

### Validation of the functional role of TAGLN in LDL internalization

To investigate the biochemical role of *TAGLN* in LDL metabolism, we utilized a stable clone of *TAGLN*-KO HepG2 cells (Figure 2A). Positive and negative controls cells were also generated by transfecting HepG2 cells with a *LDLR*-targeting sgRNA and a non-targeting sgRNA, respectively (Figure 2B). We evaluated LDL uptake using a broad concentration range of LDL (from 0 to 120 μg protein of LDL/ml). Cellular LDL uptake in *TAGLN*-KO cells saturated at ~50 μg/ml of LDL, similar to control cells and to previously reported values for LDL uptake in fibroblasts [27]. However, in marked contrast to control cells, LDL uptake in *TAGLN*-KO cells was significantly reduced by ~30% (Figure 2C).

Since prior GWAS and our computational work showed associations between *TAGLN* loci and triglycerides, soluble transferrin receptor, and hematologic traits, we extended our analysis to transferrin and very low-density lipoprotein (VLDL), the main plasma, lipoprotein carrier of triglycerides. Interestingly, uptake of VLDL and transferrin were also reduced compared to control cells (p<0.04) and to a similar degree as LDL (*p*>0.12) (Figure 2D-F). This suggests that *TAGLN* deficiency affects a common mechanism involved in the internalization of these three different cargoes. In contrast, in *LDLR*-KO cells, the defect in uptake was specific for LDL.

To test whether reduced cellular LDL uptake in *TAGLN*-KO cells is due to alterations in *LDLR* expression, we assessed *LDLR* mRNA expression. Surprisingly, *LDLR* mRNA levels were significantly increased by 2-fold in *TAGLN*-KO cells when compared to controls (*p*=0.0002) (Figure 2G). This finding suggests that decreased LDL uptake in *TAGLN*-KO cells may be due to less efficient internalization of LDLR, rather than reduced LDLR expression and that the defect in LDL internalization in *TAGLN*-KO is partially compensated by upregulation of the receptor. This would imply that the amount of LDLR on cell surface of *TAGLN*-KO cells would be increased compared to control cells. To test this hypothesis, we incubated cells with fluorescent LDL at low temperature (4°C) to prevent LDL internalization in order to allow visualization of LDL binding sites on the cell surface. Quantification by flow cytometry indicated a 2-fold increase in LDL binding sites on the cell surface of *TAGLN*-KO cells compared to controls (*p*=0.0001) (Figure 2H), which correlated with LDLR mRNA levels (Figure 2G).

Confocal microscopy (Figure 2I, left panels) confirmed the above flow cytometry findings. The observed accumulation of fluorescent LDL on the surface of *TAGLN*-KO cells incubated at 4°C is consistent with increased LDLR expression on the plasma membrane. After low-temperature incubation, a portion of cells were warmed to 37°C to allow LDL internalization for 2 hours (Figure 2I, right panels). After this treatment, control cells had reduced LDL fluorescence on the cell surface and increased cytoplasmic DiI-fluorescence (Figure 2I; right, upper panel). In contrast, the amount of LDL fluorescence on the cell surface of *TAGLN*-KO cells (Figure 2I; right, lower panel), appeared to be unchanged, consistent with impaired LDL internalization. To evaluate LDLR internalization rate, we incubated cells with monensin for different times, and then evaluated the remaining LDL binding sites on cell surface by FACS sorting (Figure 2J). Monensin is known to block LDLR recycling in mammal cells, promoting LDLR disappearance from cell surface [28]. Similar to others [29], we found, using FACS, that in control cells, monensin incubation decreased cell surface LDLR by 70%. In contrast, in *TAGLN*-KO cells, loss of surface LDLR was significantly decreased (*p*<0.03) by the monensin treatment (Figure 2J). Taken together, these findings indicate that loss of transgelin expression reduces LDLR internalization, resulting in the retention of LDLR at the plasma membrane.

Transgelin is a cytoskeletal protein that binds to actin, stabilizing actin filament bundles that are critical during the invagination of the endosome [30]. Therefore, we analyzed the arrangement of actin filaments in *TAGLN*-KO and control cells by staining cytoplasmic actin filaments with fluorescent phalloidin. Control cells exhibited characteristic cell spreading with linear cytoplasmic and subcortical actin filaments (Supplementary Figure 5A). In marked contrast, *TAGLN*-KO cells were rounded, with disorganization of intracellular actin filaments and the presence of blebs lacking subcortical actin (Supplementary Figure 5B). These studies demonstrate a pivotal role for transgelin in stabilizing actin filaments during not only endocytosis but also in maintaining overall normal cell morphology.

Given the importance of LDL internalization in overall cholesterol cellular balance, we examined alterations in several known cholesterol homeostatic pathways in *TAGLN*-KO cells. Cells were first incubated in media containing LDL as the sole extracellular source of cholesterol. Cellular cholesterol content in *TAGLN*-KO cells was about 60% lower compared to controls (*p*=0.008), which is comparable to the reduced cellular cholesterol content seen in *LDLR*-KO cells (*p*=0.99) (Figure 3A). We next evaluated mRNA expression levels of several critical enzymes involved in cholesterol synthesis (Figure 3). Compared to controls, *TAGLN*-KO cells showed increased expression of two cholesterol biosynthetic enzymes, namely 3-hydroxy-3-methylglutaryl-CoA reductase (*HMGCR*) and mevalonate kinase (*MVK*) (*p*<0.05), comparable to the increased levels seen in *LDLR*-KO cells (*p*>0.905). Additionally, squalene monooxygenase (*SQLE*) mRNA levels also showed an increase of 40% in *TAGLN*-KO (*p*=0.147). Unlike in *LDLR*-KO cells, mRNA levels of *SREBF2* and *SCAP* were not altered in *TAGLN*-KO cells when compared to controls; however, *SREBF1* mRNA levels were increased (*p*=0.044). Overall, these findings are consistent with compensatory activation of cholesterol biosynthesis in *TAGLN*-KO cells due to decreased cellular cholesterol.

**Figure 3:**
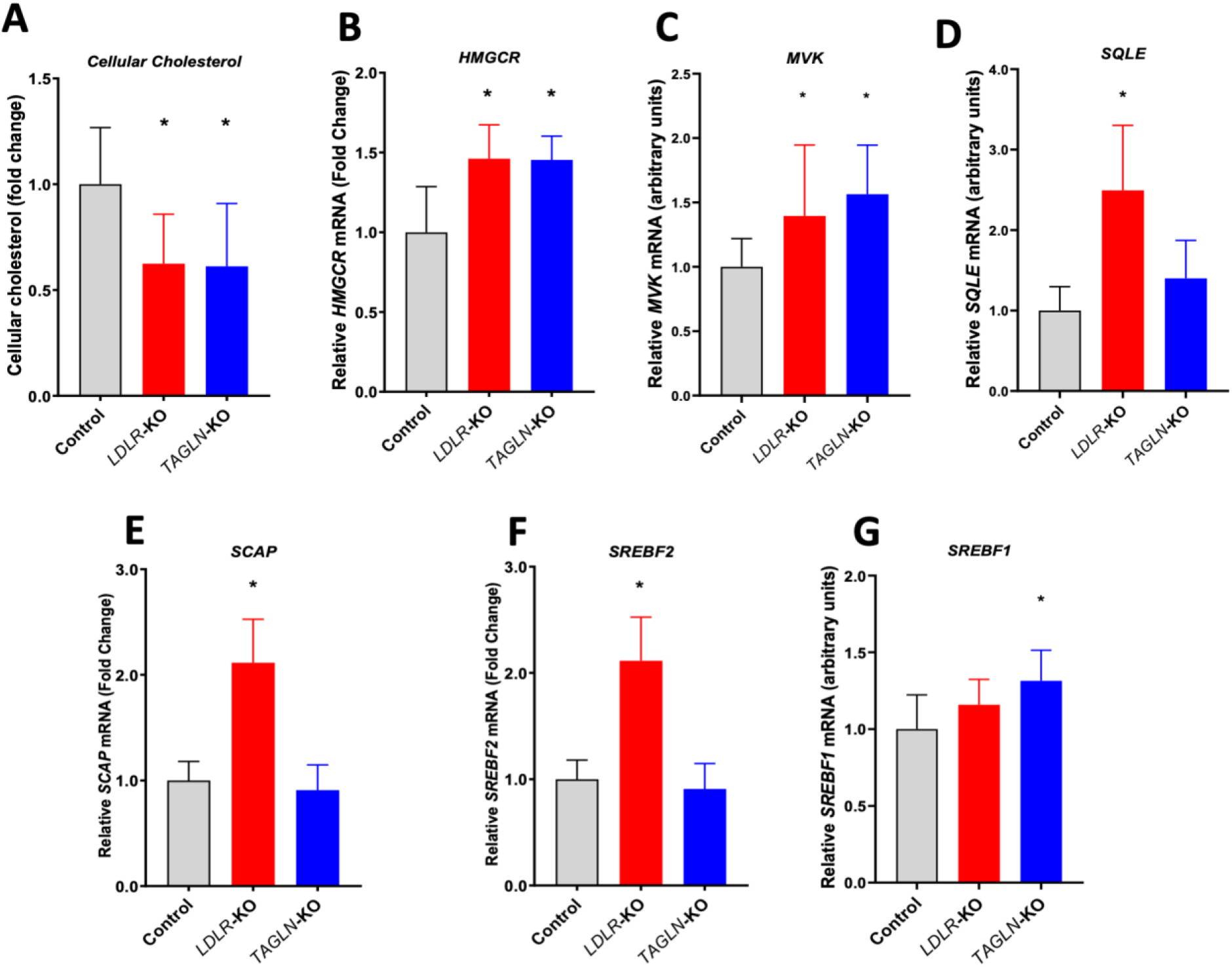
Reduction in cellular cholesterol content in TAGLN-KO cells induces mRNA expression of cholesterol synthesis enzymes. (A) Control, LDLR-KO and TAGLN-KO cells were incubated in serum-free media containing 50 μg/ml LDL for 72 hours. Total lipids were extracted with hexane:isopropanol (3:2) and cholesterol was measured by an enzymatic colorimetric assay. (B – G) Control, LDLR-KO and TAGLN-KO cells were incubated in serum-free media containing 50 μg/ml LDL for 72 hours. Messenger RNA expression levels of 3-Hydroxy-3-Methylglutaryl-CoA Reductase (HMGCR), Mevalonate Kinase (MVK), Squalene Monooxygenase (SQLE), SREBF Chaperone (SCAP), Sterol Regulatory Element Binding Transcription Factor 2 (SREBF2) and SREBF1 were determined by RT-PCR. *p<0.05 vs control. One-way ANOVA.

**Figure 4.**
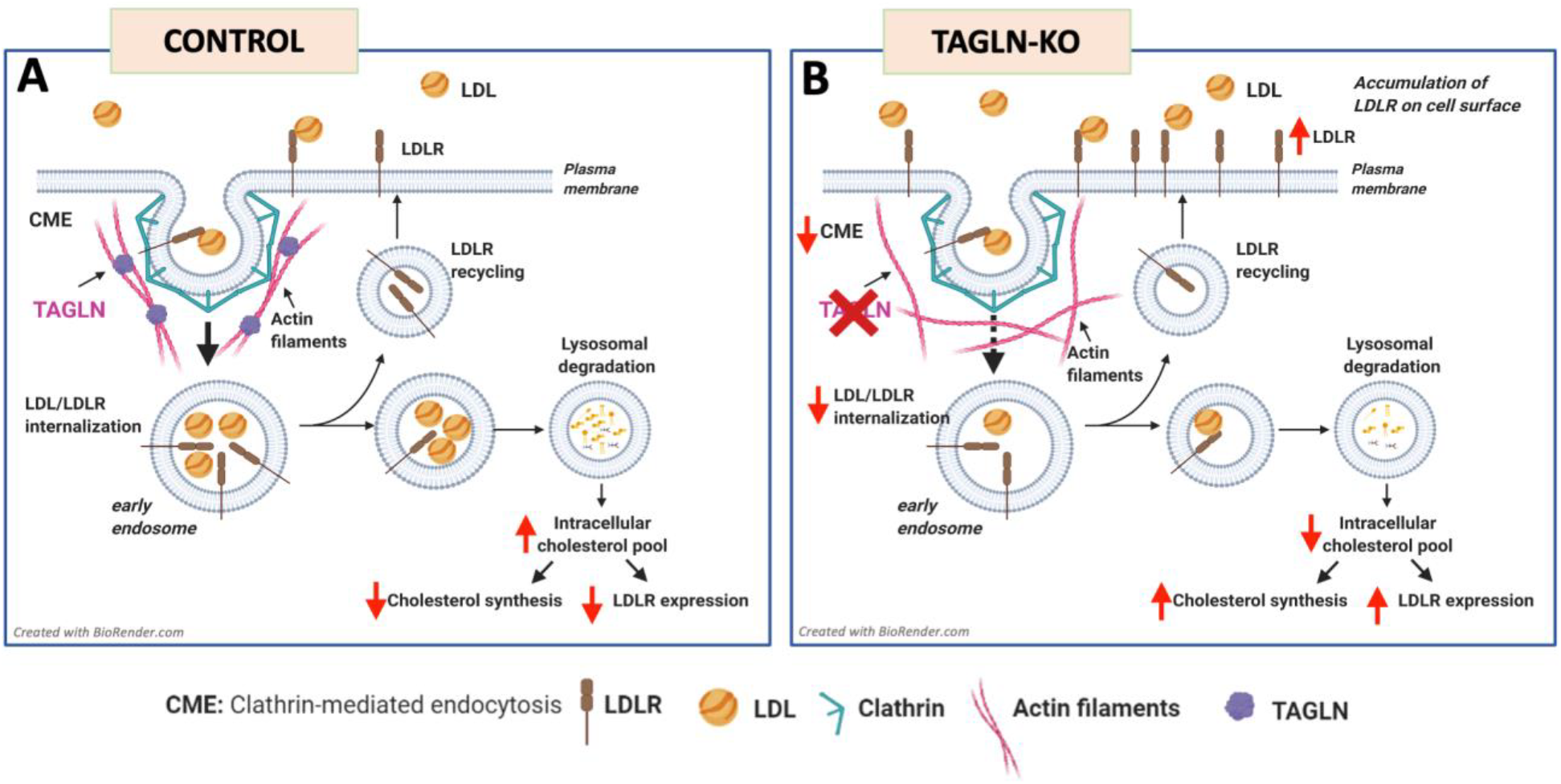
Proposed mechanism for the role of transgelin during endocytosis of low-density lipoprotein (LDL). (A) In control cells, LDL binds to its receptor (LDLR) and the complex is internalized by clathrin-mediated endocytosis (CME). During the actin dependent phase of CME, transgelin (TAGLN) binds to actin filaments, creating actin filament bundles to support plasma membrane bending before endocytic vesicle scission. In early endosomes, LDLR is recycled to plasma membrane, while LDL is directed to lysosomes for degradation. Cholesterol in LDL increases intracellular cholesterol pools, negatively regulating LDLR transcription and suppressing intracellular cholesterol synthesis. (B) In TAGLN-KO cells, a failure in the formation of actin bundles leads to inefficient CME, reducing the internalization of LDLR. Therefore, cellular LDL is decreased and LDLR accumulates on plasma membrane. Lower cellular availability of LDL depletes intracellular cholesterol pools and thus induces expression of LDLR and cholesterol synthesis enzymes.

## DISCUSSION

In the present study, we implemented a new genome-wide CRISPR/Cas9 knockout screen in a liver cell line, using cellular LDL uptake as the basis for selection, in order to identify and characterize new genes involved in LDL internalization. We identified transgelin (*TAGLN*), an actin-binding protein, as a novel gene participating in LDL internalization in liver cells. Furthermore, using different genomic association approaches, we found that common variants at the *TAGLN* locus and genetically, predicted expression of *TAGLN* are both associated with elevations in plasma total and LDL cholesterol and triglycerides, as well as with lipid-related phenotypes, such as body-mass index, waist circumference, body fat percentage, dyslipidemia, and lipid lowering medication use. Finally, our biochemical and cell biology analyses suggest that transgelin plays a vital role during LDLR internalization, most likely by binding to actin filaments during endocytosis, thereby facilitating LDL uptake and consequently, affecting cellular cholesterol homeostasis.

Transgelin is a 22 KDa actin-binding protein, a member of the calponin family, known to stabilize actin filaments [30]. In smooth muscle cells, transgelin is strongly associated with actin filaments and represents one of their earliest differentiation markers [30]. Additionally, transgelin dysregulation has been observed in different types of tumors, suggesting a role in tumor proliferation and invasiveness [31, 32]. Our CRISPR/Cas9 screen of a liver cell line revealed that transgelin also plays a role in LDL metabolism. We observed, however, only modest reduction in LDL uptake in *TAGLN*-KO cells, possibly due to compensatory changes. Cholesterol is a critical molecule for cell survival, and low cellular cholesterol content is sensed in the endoplasmic reticulum and induces the expression of LDLR and cholesterol synthesis enzymes [33]. In *TAGLN*-KO cells incubated in media containing LDL as the exclusive source of extracellular cholesterol, reduced LDL internalization depleted cellular cholesterol content, and therefore, upregulated the expression of cholesterol synthesis enzymes and also LDLR. Nevertheless, increased expression of enzymes involved in cholesterol synthesis was not able to fully compensate for the reduction in cellular cholesterol levels.

Genome-wide association studies have identified many possible associations between SNVs and phenotypic traits in large scale human genetic studies. The associations of SNVs at the *TAGLN* locus (11q23.3) and genetically predicted expression of *TAGLN* with plasma lipids support our *in vitro* findings of an important physiologic role for this locus in lipoprotein metabolism. Although at lower significance, SNVs at the *TAGLN* locus have also been associated with coronary artery disease [34]. In a preliminary study, we also observed a reduction of circulating plasma transgelin, as measured by mass spectrometry, in patients with coronary artery disease in comparison to controls (data not shown), but additional studies will be needed to further investigate this issue. Interestingly, transgelin-deficient mice with an *Apoe*-KO background are reported to develop accelerated atherosclerosis compared to control *Apoe*-KO mice [35]. This was attributed to a phenotypic modulation of smooth muscle cells, but no difference in plasma lipids was reported. Because apoE is a major ligand for LDLR, this may have masked the effect of transgelin on lipid levels.

Cellular LDL uptake, mediated by LDLR, occurs via clathrin-mediated endocytosis (CME). CME is mediated by a complex network of endocytosis-related proteins that we found is also significantly altered in our screen (LFC >1.5). The enrichment of the actin-dynamic cluster seen in our screen further highlights the significant role that actin filaments play during CME, particularly at the later stages, before scission of endocytic vesicles [36]. Our data suggest that the coordinated function of the endocytic network is necessary for cellular LDL uptake to occur.

Actin filaments are thought to generate the force to bend the plasma membrane during endocytosis [37]. In this context, the yeast *TAGLN* ortholog, *SCP1*, plays a central, role during CME, localizing with actin filaments and supporting membrane invagination [38]. We, therefore, hypothesized that, if transgelin is involved in the actin-dependent phase during endocytosis, internalization of LDLR would be impaired, and therefore, excess LDLR would be retained on the plasma membrane (Figure 4). Our 4°C LDL binding experiments are consistent with this hypothesis. Furthermore, our monensin experiments demonstrated a decreased rate of internalization of LDLR in *TAGLN*-KO cells. It has previously been demonstrated that *LDLRAP1* deficiency completely blocks LDLR internalization, leading to receptor retention on the cell surface [29]. The partial blockage of receptor internalization observed in transgelin deficiency may be due to functional compensation by other actin-binding proteins, which requires additional investigation. In addition to its importance in CME, transgelin may have other roles [30]. For example, the stabilization of actin-filaments of the cytoskeleton by transgelin likely explains the morphological changes we observed in *TAGLN*-KO cells.

Like LDLR, LRP1, an important hepatic receptor for VLDL, is also internalized by CME [39, 40]. In *TAGLN*-KO cells, uptake of LDL and VLDL is equally affected, suggesting that CME is affected at a common step, presumably downstream of the interaction with adaptor proteins [41], likely during the actin-dependent phase of endocytosis. Based on prior GWAS findings, carriers of variants in the *TAGLN* locus would be expected to have not only increased LDL-C but also triglycerides, which is consistent with our cell culture findings. Patients with this lipid phenotype are often diagnosed as having the Familial combined hyperlipidemia. These patients are known to be at a marked increased risk of coronary artery disease, but a clear monogenic basis for this disorder is yet to be identified [42]. Hypermethylation of *TAGLN* promoter has been, shown to decrease transgelin expression in tumor cells [43]. Besides mutations and polymorphisms in *TAGLN,* epigenetic *TAGLN* modifications related to metabolic disorders could, therefore, also be linked to lipid levels.

Another ligand known to undergo CME and showed decrease uptake in our *TAGLN*-KO cells is transferrin. The transferrin receptor (TfR) is cleaved from the plasma membrane and its plasma soluble form (sTfR) serves as a marker of iron deficiency [44]. Associations reported in previous GWAS showed that different SNVs in *TAGLN* locus were associated with sTfR [45]. This finding could also further explain the associations we found between predicted expression levels of TAGLN and various erythrocyte indices in the UKBB. Alterations in transgelin function leading to destabilization of actin filaments could possibly also be related to red blood cell fragility [46].

In summary, using a genome-wide CRISPR/Cas9 knockout screen we identified *TAGLN* as novel gene involved in LDL endocytosis. The human relevance of our *in vitro* findings was supported by population genetic analysis, demonstrating that SNVs at the *TAGLN* locus and genetically predicted expression of *TAGLN* were associated with plasma lipids and related phenotypes. Furthermore, our biochemical and cell biology data suggest that TAGLN stabilizes actin filaments during the actin-dependent phase of CME, affecting internalization of not only LDLR, but possibly other cargo. We also observed an enrichment of other genes in our screen related to endocytosis, which raises the possibility that polymorphisms in other endocytosis related genes that may only slightly affect function and are still compatible with life, may also affect plasma LDL levels. The identification of new genetic factors involved in lipoprotein metabolism could eventually reveal new diagnostic markers ad therapeutic targets for reducing plasma LDL-cholesterol for the prevention of ASCVD.

## ACKNOWLEDGEMENTS

We would also like to thank to the Flow Cytometry Core and Bioinformatics and Computational Biology Core, NHLBI, at National Institutes of Health, for technical support.

## SOURCE OF FUNDING

This research was supported by the Intramural Research Program of the National Heart, Lung, and Blood Institute (NHLBI) (HL006095) at National Institutes of Health. MMM was supported by the National Heart, Lung, and Blood Institute (NHLBI K99 HL136875).

## AUTHORS’ DISCLOSURES

S.V. salary and significant ownership in Global Genomics Group. M.M.M. is now an employee of Novartis Institutes of Biomedical Research, but he was not an employee when his contribution to the work was conducted. The rest of authors do not have any conflict of interest regarding this publication.

## AUTHORS’ CONTRIBUTIONS

D.L. and A.T.R. conceived the idea and designed the experiments. D.L. carried out the experiments, with the support of P.I. The project was supervised by A.T.R. B.V., E.B.N., J.T., and L.A.F. contributed with valuable technical support and data interpretation. A.T.B. and S.V. contributed with proteomic data analysis. M.M., O.D., and I.J.K. contributed with genomic analysis, data interpretation, and manuscript edition. C.C. and Y.L. contributed with technical support in confocal microscopy experiments and DNA sequencing respectively. D.L. wrote the manuscript with support from E.B.N., L.A.F., and the supervision of A.T.R.

