## Supplemental Figures for "Transgelin: A New Gene Involved in LDL Endocytosis Identified by a Genome-wide CRISPR-Cas9 Screen"

### Supplementary Figure 1

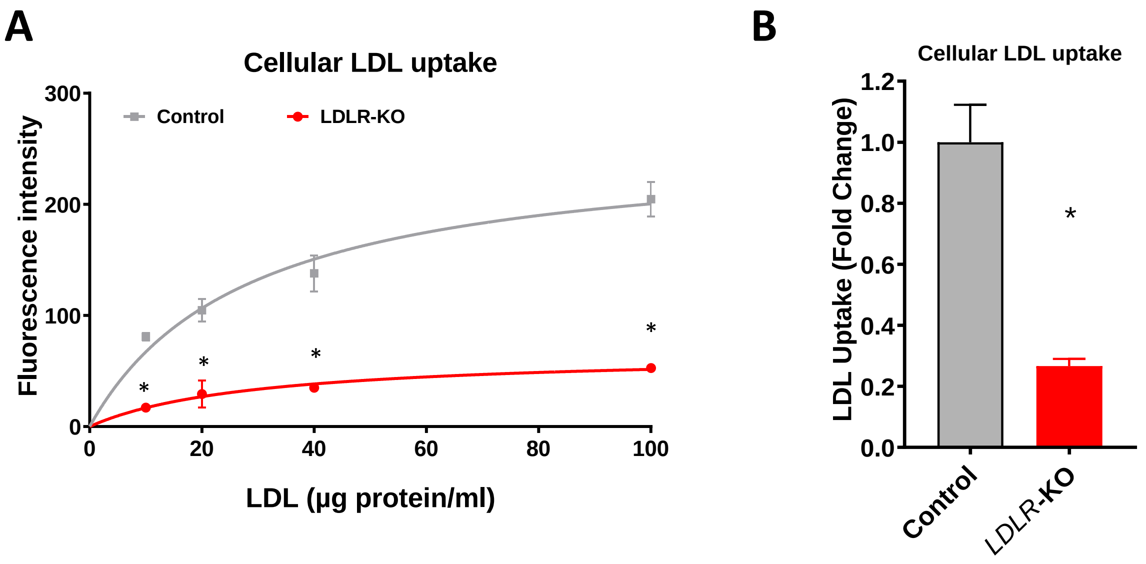


**Supplementary Figure 1. Knocking out low-density lipoprotein receptor (LDLR) in HepG2 cells reduces cellular uptake of LDL by 75%.**

HepG2 cells stably expressing Cas9 were transfected with lentivirus containing a sgRNA targeting human LDLR (ABM Goods, Inc. Canada, Cat# 264181110204). After selection, transfected cells were sorted and clones from individual cells were expanded. Clones with negative LDLR expression by western blot were selected for further studies (*LDLR*-KO). Control cells were generated by transfection of HepG2 cells with a scrambled non-targeting sgRNA. (A) Control and *LDLR*-KO cells were incubated with the indicated concentrations of AlexaFluor$^{TM}$ 568-LDL for 4 h at 37°C. Fluorescence, proportional to LDL internalization, was determined by FACS. (B) Cellular LDL uptake at 40 µg/ml of LDL. *p<0.001. Student's t-test.

### Supplementary Figure 2

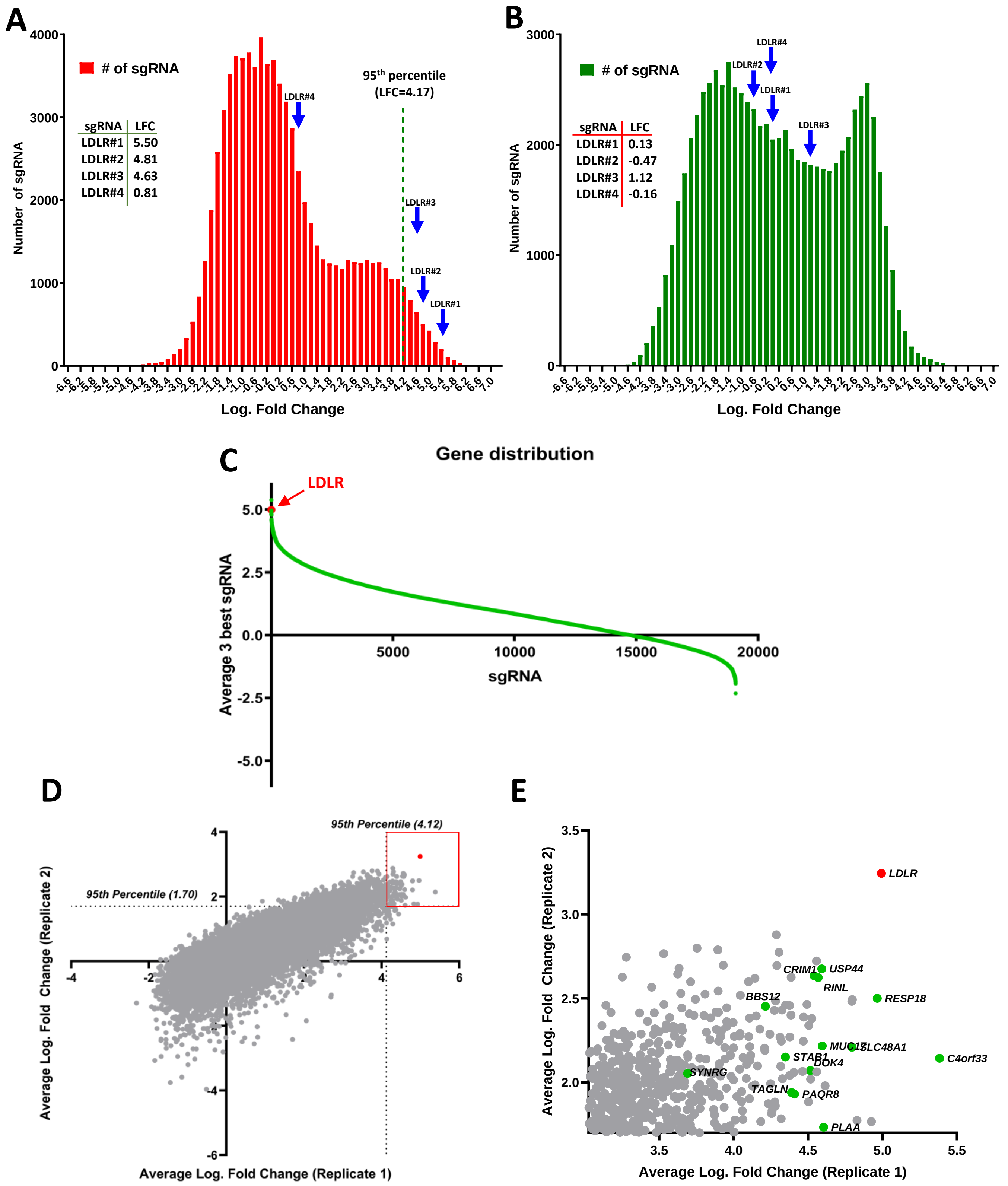


**Supplementary Figure 2: CRISPR/Cas9 screen for cellular LDL uptake genes.** (A and B) Frequency distribution plots of sgRNA enrichment distribution in cells with low LDL uptake and in non-sorted cells respectively. Arrows indicate the enrichment of the sgRNAs targeting *LDLR* (low-density lipoprotein receptor) in each case. Data indicates Log Fold Change (LFC) for each sgRNA. (C) Plot of the 19,114 genes targeted in the screen, ranked by average of their best 3 sgRNAs. Red dot indicates *LDLR*. (D) Scatter plot of the results from the 2 independent screens. Top enriched genes (Average LFC above 95$^{th}$ percentile in both screens), marked in the red square, are amplified in panel (E). Data represents the average of LFC of the 3 best sgRNA for each gene in each independent experiment. Red dot shows LDLR. Green dots, selected candidate genes.

### Supplementary Figure 3

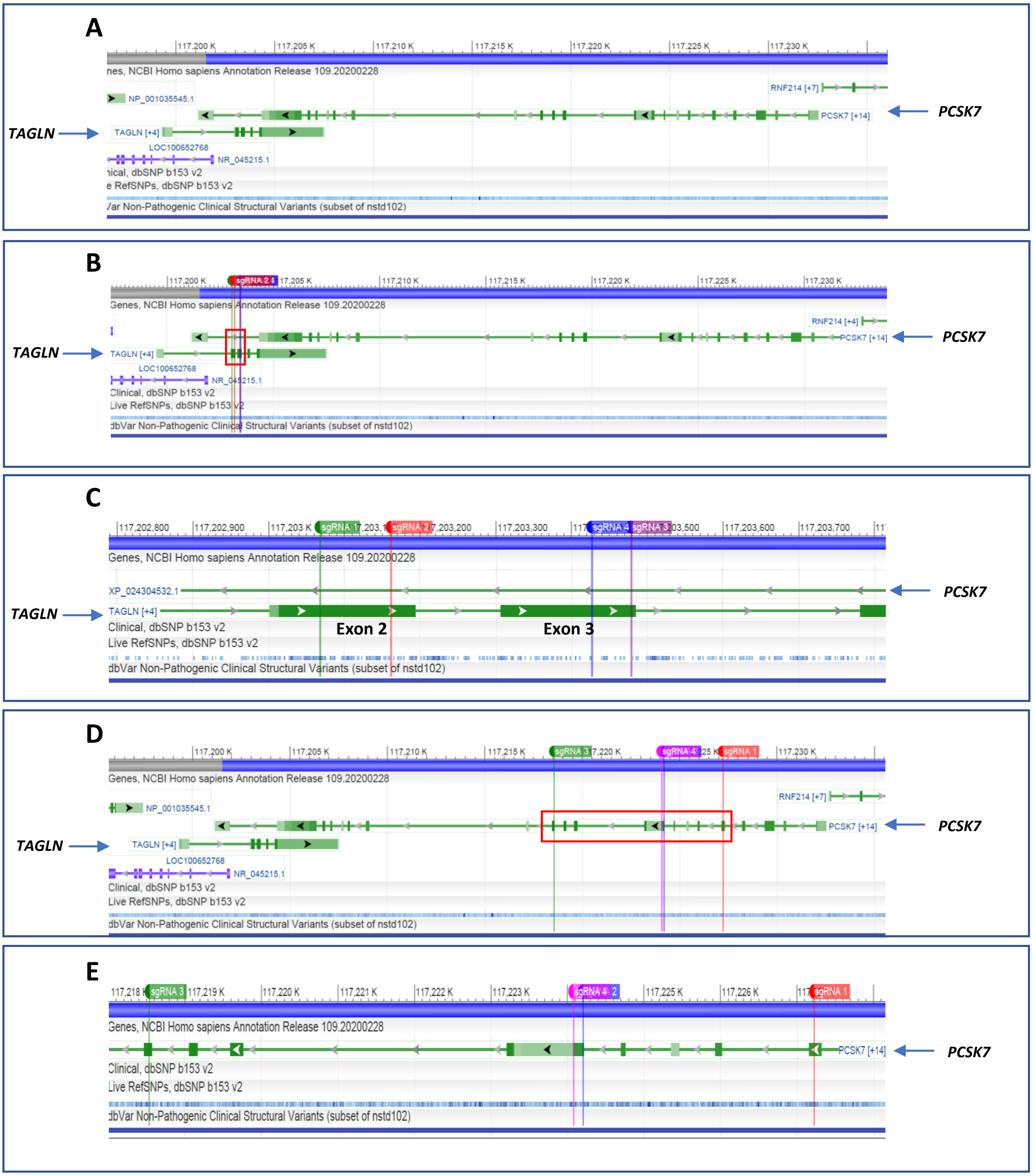


**Supplementary Figure 3: *TAGLN* overlaps tail-to-tail with *PCSK7* and *TAGLN* sgRNA target sites overlap with intron 19 in *PCSK7* 3'-UTR but targeted sites in *PCSK7* do not overlap with *TAGLN* sequence.** (A) *TAGLN* and *PCSK7* (locus 11q23.3) are displayed with National Center for Biotechnology Information (NCBI) Sequence Viewer. Exhibited region: *Homo sapiens* chromosome 11, GRCh38.p13. NCBI Reference Sequence: NC_000011.10 (between 117195814 to 117236537). (B) Targeted sequences on *TAGLN* for each sgRNA in CRISPR library were mapped using NCBI Sequence Viewer: sgRNA 1 (AATCGAGAAGAAGTATGACG), sgRNA 2 (CCTGGAAGCCCAAGCGCCCA), sgRNA 3 (CTTCCTCTCTACCTTCAAAG), and sgRNA 4 (GAAGGCGGCTGAGGACTATG). Targeted sites are in *TAGLN*'s Exons 2 and 3, overlapping with intron 19 in *PCSK7* 3'-UTR. (C) Amplified region indicated in the red square in (B). (D) Targeted sequences on *PCSK7* for each sgRNA in CRISPR library were mapped using NCBI Sequence Viewer: sgRNA 1 (GCGATGTGCAGGAGAGATCG), sgRNA 2 (TCAGGCTGCCTTACAACATG), sgRNA 3 (TGACGTAGGATGCTAAGTAA), and sgRNA 4 (TGGGTGACTACCTATGGTGA). Targeted sequences are not in overlapping regions with *TAGLN* and are in *PCSK7*'s Exons 5, 9, and 12. (E) Amplified region indicated in the red square in (D).

### Supplementary Figure 4

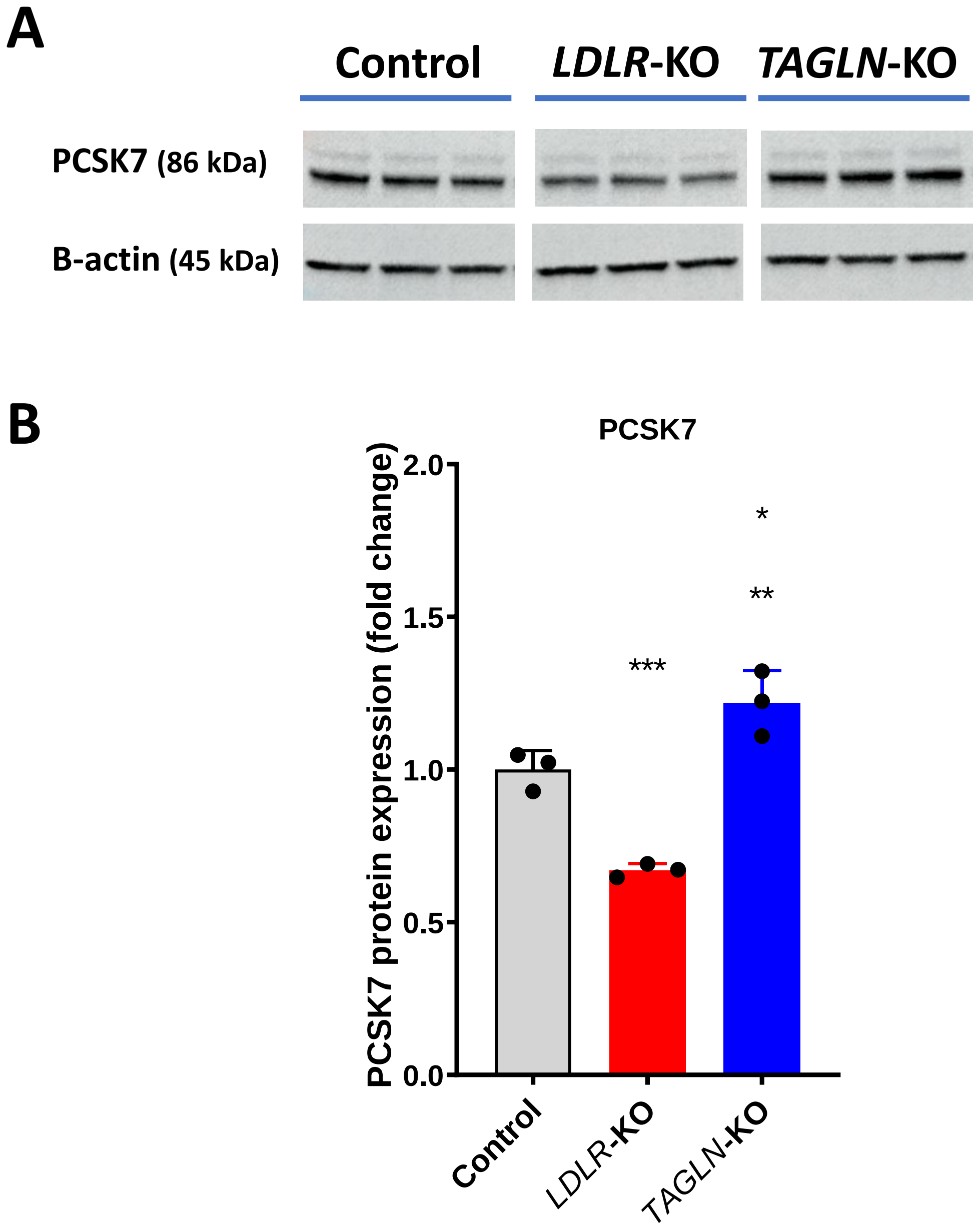


**Supplementary Figure 4: Proprotein Convertase Subtilisin/Kexin Type 7 (PCSK7) expression levels are unchanged or slightly increased in *TAGLN*-KO cells, while it was decreased in *LDLR*-KO cells.**

Control, TAGLN-KO and LDLR-KO cells were plated in 10-mm dishes and incubated in serum-free media containing 50 µg/ml LDL for 72 hours. Cells were then lysed with RIPA buffer (600 ul/dish) and incubated with 4X LDS sample loading buffer (Invitrogen, Carlsbad, CA) at a final concentration of 1X at room temperature for 1 hour. Electrophoresis and Western blotting were as described in Materials and Methods. Antibodies for PCSK7 (86 kDa) were purchased from Cell Signaling Technology (Danvers, MA). (A) Representative blot of three independent runs. Each lane represents and independent sample. (B) Quantification of PCSK7 expression. Each point represents the average of the normalized density for each sample run in three independent runs. *p=0.03 vs control. **p=0.0002 vs LDLR-KO. ***p=0.003 vs control.

#### Supplementary Figure 5:

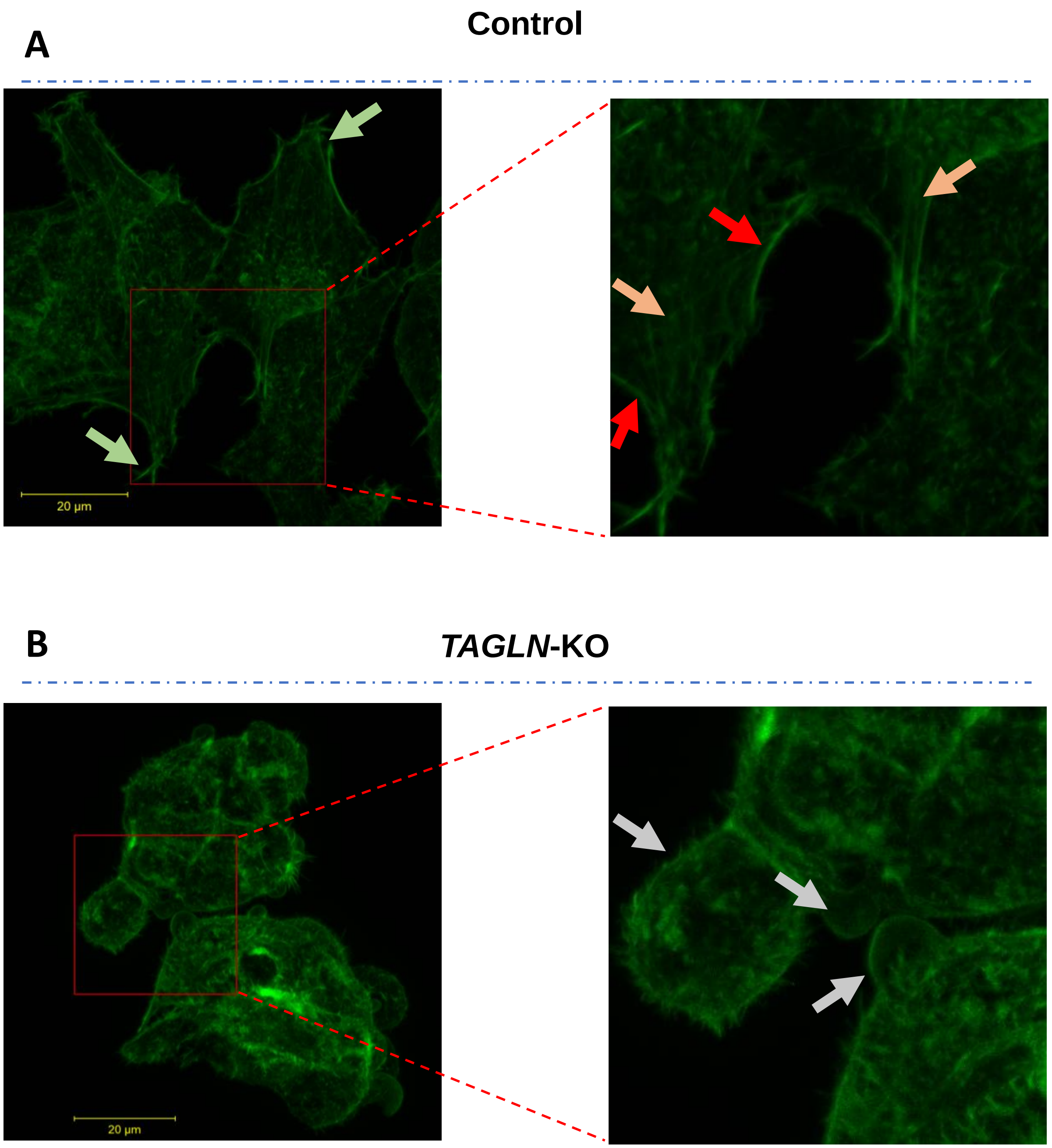


**Supplementary Figure 5. Transgelin deficiency is associated with morphological changes in HepG2 cells, associated with changes in actin cytoskeleton.**

Control and *TAGLN*-KO cells were plated in cell culture dishes containing collagen-coated glass coverslip bottoms. After 48 h, cells were fixed with 4% paraformaldehyde, permeabilized with 0.1% Triton X-100, and stained for 30 minutes with Alexa Fluor® 488 Phalloidin (Molecular Probes, Eugene, OR). Image stacks of cells labelled with Alexa Fluor® 488 Phalloidin (Molecular Probes, Eugene, OR) were acquired on a Zeiss 880 confocal microscope (Jena, Germany) using a Zeiss 40x Plan-Apochromat objective lens (N.A. 1.3) and 488nm excitation, the emission bandwidth set to 490-555nm, the lateral pixel sizes set to 90nm, and the interslice thickness set to 440nm. Maximum projection images as displayed in the figures were generated using the Zeiss Zen software. (A) Control cells, showing characteristic spread shape (green arrows), cortical actin filaments (red arrows), and intracellular actin filaments (orange arrows). (B) *TAGLN*-KO cells were rounder with the presence of blebs (grey arrows). Images represent the maximum intensity projection of the z-stack. Scale bar indicates 20 µm.
