## Supplemental Tables for "Transgelin: A New Gene Involved in LDL Endocytosis Identified by a Genome-wide CRISPR-Cas9 Screen"

**Supplementary Table 1. Whole list of targeted genes (n=19,114) by Brunello genome-wide CRISPR/Cas9 knockout library (Doench, J.G., et al. Nat Biotechnol, 2016).** Each gene was targeted by four single guiding RNA (sgRNA). Genes were ranked by the average of the three most enriched sgRNA.

| Rank | Gene | Log. Fold Change of sgRNAs | | | | |
|---|---|---|---|---|---|---|
| | | 1st | 2nd | 3rd | 4th | Average of 3 most enriched |
| 1 | *C4orf33* | 6.235 | 5.3319 | 4.5448 | 0.17582 | 5.37 |
| 2 | *LDLR* | 5.5014 | 4.8066 | 4.6289 | 0.86656 | 4.98 |
| 3 | *RESP18* | 5.2364 | 5.0818 | 4.5366 | 4.4184 | 4.95 |
| 4 | *LY6G5C* | 5.9644 | 4.5194 | 4.2512 | -0.13231 | 4.91 |
| 5 | *GCOM1* | 5.8228 | 4.5281 | 4.0937 | -1.0997 | 4.81 |
| 6 | *MUC22* | 5.5336 | 4.7189 | 4.0977 | 4.0116 | 4.78 |
| 7 | *SLC48A1* | 5.5297 | 4.5023 | 4.3131 | 0.16173 | 4.78 |
| 8 | *FBLIM1* | 5.3896 | 4.7012 | 4.2504 | 0.58258 | 4.78 |
| 9 | *GFRA3* | 4.9731 | 4.8064 | 4.0208 | 2.7129 | 4.60 |
| 10 | *PLAA* | 5.9071 | 4.3885 | 3.4763 | 1.6061 | 4.59 |
| 11 | *MUC17* | 5.4071 | 4.3603 | 3.9755 | -1.2817 | 4.58 |
| 12 | *USP44* | 5.2643 | 4.2724 | 4.1975 | -0.68741 | 4.58 |
| 13 | *RINL* | 4.6738 | 4.5229 | 4.4673 | 0.42034 | 4.55 |
| 14 | *ZBTB43* | 5.1629 | 4.441 | 4.0296 | -0.31075 | 4.54 |
| 15 | *AK7* | 5.2913 | 4.9282 | 3.4104 | -0.99059 | 4.54 |
| 16 | *CRIM1* | 4.8607 | 4.573 | 4.1477 | 0.062547 | 4.53 |
| 17 | *NHLH2* | 5.8302 | 4.0292 | 3.6763 | -1.9259 | 4.51 |
| 18 | *C1orf86* | 5.1582 | 4.7753 | 3.5778 | -1.9874 | 4.50 |
| 19 | *DOK4* | 5.2721 | 4.2456 | 3.9867 | -0.05054 | 4.50 |
| 20 | *PPP1R3D* | 4.9881 | 4.7152 | 3.7704 | 3.1948 | 4.49 |
| 21 | *IFNA7* | 5.0347 | 4.2577 | 4.176 | 0.9264 | 4.49 |
| 22 | *GLRX2* | 4.9321 | 4.3587 | 4.1456 | -0.44661 | 4.48 |
| 23 | *CXorf23* | 5.2504 | 4.5961 | 3.5244 | -0.09731 | 4.46 |
| 24 | *STRA8* | 5.2058 | 4.1191 | 4.0362 | 0.14017 | 4.45 |
| 25 | *ZNF516* | 6.2097 | 5.7961 | 1.3418 | -2.7736 | 4.45 |
| 26 | *PCDHGA5* | 4.7926 | 4.6757 | 3.8012 | 0.14337 | 4.42 |
| 27 | *PAQR8* | 4.6322 | 4.5894 | 3.9656 | -3.3058 | 4.40 |
| 28 | *TENM3* | 5.3796 | 4.223 | 3.5706 | 1.4592 | 4.39 |
| 29 | *TAGLN* | 6.237 | 3.6013 | 3.2889 | -0.27744 | 4.38 |
| 30 | *MGMT* | 4.791 | 4.5744 | 3.7511 | -0.72666 | 4.37 |
| 31 | *TRPM4* | 4.9192 | 4.6103 | 3.5859 | -0.72973 | 4.37 |
| 32 | *ANGPTL1* | 5.3419 | 4.5496 | 3.2004 | -0.87328 | 4.36 |
| 33 | *JMJD4* | 4.9126 | 4.8843 | 3.2905 | -0.99694 | 4.36 |
| 34 | *KRT40* | 5.7973 | 4.2595 | 3.0288 | 1.1047 | 4.36 |
| 35 | *ALDH1A2* | 4.5863 | 4.4708 | 3.987 | 0.18682 | 4.35 |
| 36 | *G6PC2* | 5.6759 | 4.6681 | 2.6616 | -0.88877 | 4.34 |
| 37 | *STAB1* | 5.2723 | 4.6737 | 3.0566 | 2.273 | 4.33 |
| 38 | *AGPAT5* | 5.1024 | 4.6991 | 3.1897 | 2.8237 | 4.33 |
| 39 | *ZNF397* | 5.798 | 4.3352 | 2.844 | 1.2626 | 4.33 |

| | | | | | | |
|---|---|---|---|---|---|---|
| 40 | *PNLDC1* | 4.4094 | 4.2965 | 4.2362 | -1.8761 | 4.31 |
| 41 | *CYP11B1* | 5.1665 | 4.8214 | 2.946 | -0.64186 | 4.31 |
| 42 | *AP1S1* | 4.9854 | 4.4895 | 3.4433 | 1.3216 | 4.31 |
| 43 | *AHNAK* | 5.065 | 4.2752 | 3.5771 | 2.1505 | 4.31 |
| 44 | *PPTC7* | 5.289 | 4.1202 | 3.4983 | 1.0694 | 4.30 |
| 45 | *ALDH6A1* | 5.2942 | 4.3043 | 3.296 | 0.37781 | 4.30 |
| 46 | *CTRL* | 4.4165 | 4.3068 | 4.151 | -1.0121 | 4.29 |
| 47 | *PSPH* | 5.28 | 3.8804 | 3.6686 | 3.1005 | 4.28 |
| 48 | *PRRC1* | 5.0229 | 4.187 | 3.6123 | 2.4157 | 4.27 |
| 49 | *TPPP3* | 4.365 | 4.2829 | 4.1723 | -1.8735 | 4.27 |
| 50 | *COX19* | 5.362 | 4.157 | 3.2847 | -1.2208 | 4.27 |
| 51 | *MYLPF* | 4.9827 | 4.3594 | 3.459 | -1.4473 | 4.27 |
| 52 | *FAAH2* | 4.923 | 4.5789 | 3.2939 | 2.4812 | 4.27 |
| 53 | *SLC9A9* | 4.4306 | 4.3575 | 4.0066 | -1.1439 | 4.26 |
| 54 | *TNIP1* | 6.1302 | 5.2052 | 1.4539 | -1.1991 | 4.26 |
| 55 | *FUT9* | 4.9041 | 4.7301 | 3.1435 | -0.45029 | 4.26 |
| 56 | *CBY3* | 4.7575 | 4.1568 | 3.8169 | -0.56851 | 4.24 |
| 57 | *ADAM2* | 5.4378 | 3.912 | 3.3745 | -0.10686 | 4.24 |
| 58 | *PLEKHA4* | 4.923 | 4.2332 | 3.5478 | -0.568 | 4.23 |
| 59 | *TANC1* | 5.084 | 4.7594 | 2.8449 | -0.75192 | 4.23 |
| 60 | *KRTAP10-1* | 5.1425 | 4.102 | 3.414 | 0.002584 | 4.22 |
| 61 | *FAM228A* | 5.2795 | 4.1275 | 3.2414 | 1.2437 | 4.22 |
| 62 | *BBS12* | 5.5819 | 4.5587 | 2.4599 | -0.49115 | 4.20 |
| 63 | *NPY4R* | 5.2821 | 3.7334 | 3.5737 | 3.0819 | 4.20 |
| 64 | *TMCC2* | 5.0495 | 4.1247 | 3.4089 | -0.59622 | 4.19 |
| 65 | *GALNT11* | 5.2184 | 4.8919 | 2.4595 | 0.74648 | 4.19 |
| 66 | *PBRM1* | 4.6018 | 4.1477 | 3.8097 | -1.7068 | 4.19 |
| 67 | *GDF7* | 5.11 | 4.3092 | 3.0924 | -1.1442 | 4.17 |
| 68 | *CNTNAP1* | 4.5666 | 4.5029 | 3.4359 | -1.2505 | 4.17 |
| 69 | *NME5* | 4.5095 | 4.2548 | 3.7389 | 1.856 | 4.17 |
| 70 | *RPL22L1* | 5.0493 | 4.3215 | 3.1184 | 0.2484 | 4.16 |
| 71 | *KIAA0196* | 5.2804 | 4.6454 | 2.5203 | -2.0952 | 4.15 |
| 72 | *IFITM1* | 5.8601 | 4.9036 | 1.678 | -1.8341 | 4.15 |
| 73 | *IL13RA1* | 4.8932 | 4.1375 | 3.4089 | -0.83963 | 4.15 |
| 74 | *ASB7* | 4.8313 | 3.8641 | 3.7186 | 2.4343 | 4.14 |
| 75 | *HBE1* | 4.8198 | 4.8095 | 2.7596 | 1.7425 | 4.13 |
| 76 | *PLIN2* | 4.8434 | 4.295 | 3.2483 | 0.12893 | 4.13 |
| 77 | *LRRCC1* | 4.275 | 4.1979 | 3.9093 | 0.4622 | 4.13 |
| 78 | *SAA2* | 5.1925 | 4.3382 | 2.8332 | -0.85194 | 4.12 |
| 79 | *CEP70* | 4.6155 | 3.9294 | 3.8113 | 3.042 | 4.12 |
| 80 | *KRTAP5-2* | 5.0238 | 4.6927 | 2.6276 | -0.79596 | 4.11 |
| 81 | *RNF219* | 5.104 | 3.9672 | 3.2699 | 0.081756 | 4.11 |
| 82 | *TXNL1* | 5.0798 | 3.9237 | 3.2966 | -0.87739 | 4.10 |
| 83 | *HSPA1B* | 5.5324 | 4.8918 | 1.8623 | -0.39924 | 4.10 |
| 84 | *IGLL5* | 5.2724 | 3.9211 | 3.0916 | -0.67149 | 4.10 |
| 85 | *CDH11* | 4.4266 | 4.2785 | 3.5774 | -0.35774 | 4.09 |

| | | | | | | |
|---|---|---|---|---|---|---|
| 86 | *FERD3L* | 4.5885 | 4.3652 | 3.3234 | 0.85755 | 4.09 |
| 87 | *MYH2* | 4.8283 | 4.1251 | 3.3236 | -0.91862 | 4.09 |
| 88 | *IGDCC4* | 4.9772 | 3.9224 | 3.3636 | 1.3734 | 4.09 |
| 89 | *CAPN6* | 4.78 | 4.0612 | 3.4166 | 2.9261 | 4.09 |
| 90 | *SSBP3* | 5.7751 | 4.792 | 1.6586 | -0.86459 | 4.08 |
| 91 | *SUMO3* | 4.4642 | 4.4223 | 3.3347 | 2.8821 | 4.07 |
| 92 | *DIP2C* | 4.9133 | 4.1297 | 3.1678 | 2.7009 | 4.07 |
| 93 | *DPP6* | 4.6649 | 4.6281 | 2.8934 | 0.95594 | 4.06 |
| 94 | *RIC1* | 4.6196 | 3.8528 | 3.705 | -0.5204 | 4.06 |
| 95 | *VEGFB* | 5.8439 | 4.5779 | 1.7266 | 0.77351 | 4.05 |
| 96 | *METTL24* | 5.5065 | 3.8359 | 2.8032 | -1.0761 | 4.05 |
| 97 | *MAP3K1* | 5.3953 | 4.1643 | 2.5748 | 0.94811 | 4.04 |
| 98 | *CABLES2* | 5.3805 | 5.1354 | 1.605 | -1.7971 | 4.04 |
| 99 | *COL16A1* | 5.5757 | 4.4375 | 2.1047 | -1.0205 | 4.04 |
| 100 | *ARMC8* | 4.8313 | 4.7447 | 2.5174 | 0.96941 | 4.03 |
| 101 | *SOSTDC1* | 4.6844 | 4.4416 | 2.9632 | -0.72975 | 4.03 |
| 102 | *STRADB* | 4.9394 | 4.658 | 2.4819 | -1.5552 | 4.03 |
| 103 | *HEATR5B* | 4.6382 | 4.2913 | 3.1445 | -0.53356 | 4.02 |
| 104 | *SPATA31D1* | 4.5179 | 3.9469 | 3.6051 | -0.10857 | 4.02 |
| 105 | *ZNF136* | 4.5762 | 4.0287 | 3.4192 | 1.272 | 4.01 |
| 106 | *CLEC2D* | 4.4325 | 4.1176 | 3.4727 | 0.74176 | 4.01 |
| 107 | *CT45A1* | 4.018 | 3.9906 | | | 4.00 |
| 108 | *USF2* | 5.0679 | 4.6309 | 2.2964 | -1.3268 | 4.00 |
| 109 | *CXorf38* | 5.1174 | 4.4278 | 2.4418 | -1.1426 | 4.00 |
| 110 | *ARFGAP2* | 4.701 | 4.1956 | 3.0768 | -0.67791 | 3.99 |
| 111 | *AHSA1* | 4.1657 | 3.9415 | 3.8603 | -0.84501 | 3.99 |
| 112 | *AXL* | 4.6662 | 4.4846 | 2.8061 | 1.5157 | 3.99 |
| 113 | *TMPRSS15* | 4.3654 | 4.273 | 3.3175 | 0.14105 | 3.99 |
| 114 | *FAM101A* | 5.7959 | 4.7555 | 1.3953 | 0.11747 | 3.98 |
| 115 | *AASDH* | 5.0993 | 4.1903 | 2.6394 | -2.3805 | 3.98 |
| 116 | *NSUN7* | 5.2859 | 5.2711 | 1.3665 | 0.42638 | 3.97 |
| 117 | *MMP26* | 5.0287 | 4.7725 | 2.1061 | -1.8297 | 3.97 |
| 118 | *ZNF248* | 4.4461 | 4.2346 | 3.2098 | 2.7641 | 3.96 |
| 119 | *AP1S2* | 4.3999 | 4.3914 | 3.0987 | -1.4909 | 3.96 |
| 120 | *GTPBP1* | 4.9794 | 4.124 | 2.7841 | -0.78443 | 3.96 |
| 121 | *SPSB2* | 4.8504 | 4.3342 | 2.6717 | -1.8275 | 3.95 |
| 122 | *OCRL* | 5.2781 | 3.8237 | 2.7182 | -0.83784 | 3.94 |
| 123 | *PPP1R3F* | 4.6303 | 4.0558 | 3.1321 | -0.99508 | 3.94 |
| 124 | *SEC63* | 5.3842 | 3.572 | 2.8606 | -1.1576 | 3.94 |
| 125 | *MREG* | 5.4869 | 3.3486 | 2.9801 | -1.8404 | 3.94 |
| 126 | *CYP4A22* | 4.5208 | 3.9189 | 3.3729 | -1.2114 | 3.94 |
| 127 | *KCND1* | 5.4908 | 4.6852 | 1.6078 | -2.8003 | 3.93 |
| 128 | *AHR* | 4.4197 | 3.8891 | 3.4736 | 2.8619 | 3.93 |
| 129 | *OR5H2* | 4.9978 | 4.0713 | 2.7131 | 2.4183 | 3.93 |
| 130 | *UCK2* | 4.1184 | 4.0061 | 3.6492 | -1.4873 | 3.92 |
| 131 | *NOS1* | 4.0834 | 3.9341 | 3.7502 | -0.05655 | 3.92 |

| | | | | | | |
|---|---|---|---|---|---|---|
| 132 | *ZFR* | 4.6188 | 4.4014 | 2.7454 | -1.7963 | 3.92 |
| 133 | *FAM216B* | 4.2992 | 4.0052 | 3.4435 | 3.42 | 3.92 |
| 134 | *HIST1H1C* | 5.7892 | 4.9451 | 1.0131 | -1.344 | 3.92 |
| 135 | *SLC45A1* | 5.6736 | 3.4294 | 2.6299 | 0.12004 | 3.91 |
| 136 | *AMDHD1* | 5.293 | 3.226 | 3.2135 | -0.33363 | 3.91 |
| 137 | *KCNA1* | 4.3689 | 4.3376 | 3.0146 | -2.1807 | 3.91 |
| 138 | *TM6SF2* | 4.9793 | 4.1965 | 2.5354 | -0.17591 | 3.90 |
| 139 | *MAD2L1BP* | 4.4302 | 4.0635 | 3.2022 | 0.95622 | 3.90 |
| 140 | *RSPO4* | 4.4567 | 4.0237 | 3.2144 | 2.396 | 3.90 |
| 141 | *CASP4* | 4.9336 | 3.8257 | 2.9318 | 2.0211 | 3.90 |
| 142 | *MFAP2* | 4.4461 | 4.0781 | 3.1603 | -1.1361 | 3.89 |
| 143 | *ISL1* | 5.7569 | 3.4745 | 2.4393 | 1.7808 | 3.89 |
| 144 | *CLEC5A* | 4.4479 | 3.6258 | 3.5949 | -0.07487 | 3.89 |
| 145 | *THY1* | 5.0076 | 4.0083 | 2.6419 | -0.24912 | 3.89 |
| 146 | *LOC101927844* | 5.4512 | 3.9484 | 2.2578 | -0.19834 | 3.89 |
| 147 | *RBM3* | 4.2465 | 4.0488 | 3.3606 | -1.4234 | 3.89 |
| 148 | *NKX6-2* | 4.3644 | 3.7146 | 3.5738 | -0.68299 | 3.88 |
| 149 | *TRIM38* | 4.1516 | 4.0803 | 3.4139 | 3.2508 | 3.88 |
| 150 | *CCT6B* | 4.2109 | 3.7778 | 3.6524 | -0.14448 | 3.88 |
| 151 | *MYOM1* | 5.5102 | 3.7391 | 2.3786 | -0.42567 | 3.88 |
| 152 | *CPA5* | 4.4892 | 3.9364 | 3.2005 | -0.21175 | 3.88 |
| 153 | *EHMT1* | 4.3348 | 3.7943 | 3.4818 | 2.5072 | 3.87 |
| 154 | *TMEM71* | 4.4579 | 4.1817 | 2.9647 | -1.9713 | 3.87 |
| 155 | *FAM58A* | 4.0679 | 4.0608 | 3.4473 | -0.12333 | 3.86 |
| 156 | *ATP6V1G3* | 4.6434 | 4.3525 | 2.58 | -1.5817 | 3.86 |
| 157 | *ZNF391* | 4.7901 | 3.3956 | 3.3877 | -1.1945 | 3.86 |
| 158 | *ALG8* | 3.9836 | 3.9737 | 3.6145 | -0.72391 | 3.86 |
| 159 | *SLC38A9* | 4.4199 | 3.7376 | 3.3995 | -0.44318 | 3.85 |
| 160 | *OR4F6* | 5.094 | 4.1351 | 2.3263 | 1.0816 | 3.85 |
| 161 | *UIMC1* | 4.5458 | 3.7623 | 3.2464 | 0.40295 | 3.85 |
| 162 | *CDYL* | 5.4713 | 5.29 | 0.78595 | -2.6062 | 3.85 |
| 163 | *BOD1L2* | 4.4663 | 3.5464 | 3.5338 | -0.05622 | 3.85 |
| 164 | *CLUL1* | 4.4669 | 4.3251 | 2.7544 | -1.7024 | 3.85 |
| 165 | *ARHGEF12* | 4.9361 | 3.5427 | 3.0669 | -0.45569 | 3.85 |
| 166 | *C20orf62* | 4.423 | 4.2043 | 2.9021 | -2.0198 | 3.84 |
| 167 | *UBE2E2* | 5.2963 | 4.6647 | 1.5677 | -1.7908 | 3.84 |
| 168 | *SULT1A2* | 4.3832 | 3.8205 | 3.3235 | 1.6898 | 3.84 |
| 169 | *RECQL* | 5.2763 | 3.8791 | 2.3699 | -2.2223 | 3.84 |
| 170 | *SON* | 4.3024 | 4.0271 | 3.1834 | -1.695 | 3.84 |
| 171 | *ZNF729* | 4.168 | 3.892 | 3.4315 | -1.5343 | 3.83 |
| 172 | *FAM156B* | 4.9578 | 4.4239 | 2.1077 | 0.58702 | 3.83 |
| 173 | *ADAMTS4* | 4.1236 | 3.9162 | 3.4493 | -1.1492 | 3.83 |
| 174 | *MLLT4* | 5.5172 | 2.9989 | 2.9656 | 0.47288 | 3.83 |
| 175 | *SPRN* | 5.0432 | 4.0957 | 2.3399 | 2.3384 | 3.83 |
| 176 | *RAP2A* | 4.1536 | 3.9169 | 3.3811 | 1.0712 | 3.82 |
| 177 | *SLC32A1* | 5.1432 | 4.3006 | 2.0075 | 0.035082 | 3.82 |

| | | | | | | |
|---|---|---|---|---|---|---|
| 178 | *ADAMTS8* | 4.1335 | 3.8093 | 3.4675 | 2.8758 | 3.80 |
| 179 | *RHOH* | 5.6095 | 4.1861 | 1.6102 | -1.5268 | 3.80 |
| 180 | *BBOX1* | 3.9436 | 3.9193 | 3.5388 | 3.3045 | 3.80 |
| 181 | *ANKS1A* | 4.3916 | 4.0973 | 2.8972 | 1.5482 | 3.80 |
| 182 | *C1orf158* | 4.3761 | 3.8949 | 3.1098 | 1.2217 | 3.79 |
| 183 | *TESC* | 4.3804 | 3.8566 | 3.1384 | 0.60831 | 3.79 |
| 184 | *SENP5* | 5.0025 | 4.9347 | 1.4368 | -0.3797 | 3.79 |
| 185 | *STAC* | 4.2766 | 4.2572 | 2.8283 | 2.1212 | 3.79 |
| 186 | *FEM1A* | 4.7649 | 4.3984 | 2.1879 | 0.89897 | 3.78 |
| 187 | *GSTM1* | 4.4145 | 4.1794 | 2.7196 | 1.812 | 3.77 |
| 188 | *LILRA1* | 4.1735 | 4.0122 | 3.117 | -0.20462 | 3.77 |
| 189 | *DCD* | 4.298 | 3.7879 | 3.2083 | -1.6 | 3.76 |
| 190 | *CELSR1* | 4.2185 | 3.7462 | 3.317 | -1.5118 | 3.76 |
| 191 | *ARL1* | 5.4235 | 4.7955 | 1.056 | -1.9203 | 3.76 |
| 192 | *OR4X1* | 4.3538 | 4.1593 | 2.759 | -1.6187 | 3.76 |
| 193 | *ATP6V1G2* | 4.8136 | 4.3466 | 2.1101 | -1.3542 | 3.76 |
| 194 | *MRC1* | 4.4094 | 3.9597 | 2.8946 | 1.9157 | 3.75 |
| 195 | *SMAD2* | 4.0393 | 3.959 | 3.2221 | 0.71975 | 3.74 |
| 196 | *ZNF529* | 5.4944 | 4.8033 | 0.92106 | 0.63291 | 3.74 |
| 197 | *RAB8B* | 4.1232 | 4.0052 | 3.09 | 0.3977 | 3.74 |
| 198 | *NAA20* | 4.2891 | 3.6187 | 3.3063 | -0.49045 | 3.74 |
| 199 | *OTOA* | 5.44 | 2.9777 | 2.7866 | -1.2975 | 3.73 |
| 200 | *PIDD1* | 4.6579 | 3.3479 | 3.1883 | 1.5642 | 3.73 |
| 201 | *CREB3L3* | 4.3092 | 3.6519 | 3.2262 | -1.267 | 3.73 |
| 202 | *C3orf36* | 3.9935 | 3.87 | 3.3111 | -0.83436 | 3.72 |
| 203 | *C1orf115* | 4.3825 | 4.1988 | 2.5899 | 1.9494 | 3.72 |
| 204 | *MANBA* | 5.0396 | 3.0953 | 3.0308 | -1.3526 | 3.72 |
| 205 | *LAPTM4A* | 5.1802 | 4.9113 | 1.0692 | -0.02799 | 3.72 |
| 206 | *COL9A1* | 5.3821 | 4.0285 | 1.7477 | 0.71692 | 3.72 |
| 207 | *RAB39B* | 5.1753 | 4.2734 | 1.7078 | 0.063013 | 3.72 |
| 208 | *FASTKD1* | 4.335 | 3.6384 | 3.1682 | 1.2526 | 3.71 |
| 209 | *CDKN2B* | 5.423 | 3.2153 | 2.5005 | 1.6142 | 3.71 |
| 210 | *FNBP1L* | 4.7845 | 4.4528 | 1.9014 | -1.3291 | 3.71 |
| 211 | *FOXA3* | 4.1915 | 3.6112 | 3.3321 | 2.1941 | 3.71 |
| 212 | *CLEC10A* | 5.2276 | 3.257 | 2.6471 | 0.39232 | 3.71 |
| 213 | *ZSCAN21* | 3.8366 | 3.7805 | 3.5098 | 3.0648 | 3.71 |
| 214 | *ZNF382* | 4.8677 | 3.8527 | 2.4057 | 0.25548 | 3.71 |
| 215 | *H2AFV* | 4.352 | 3.813 | 2.9518 | -0.5961 | 3.71 |
| 216 | *ZC3H8* | 5.3525 | 3.0794 | 2.6805 | -0.55074 | 3.70 |
| 217 | *PCSK6* | 3.9129 | 3.6088 | 3.589 | -0.00112 | 3.70 |
| 218 | *KLHL31* | 5.0027 | 4.0619 | 2.0352 | -0.30887 | 3.70 |
| 219 | *GAL3ST3* | 4.6246 | 3.3763 | 3.0901 | -2.3118 | 3.70 |
| 220 | *ERICH4* | 5.0037 | 4.1723 | 1.9112 | 0.87484 | 3.70 |
| 221 | *GCH1* | 4.7787 | 4.1711 | 2.1356 | -1.8238 | 3.70 |
| 222 | *ADI1* | 5.389 | 4.6667 | 1.028 | -0.57304 | 3.69 |
| 223 | *LRWD1* | 3.9191 | 3.6535 | 3.5056 | 1.6397 | 3.69 |

| | | | | | | |
|---|---|---|---|---|---|---|
| 224 | *PCDH9* | 5.1805 | 3.5709 | 2.323 | -0.84337 | 3.69 |
| 225 | *MIA3* | 5.9246 | 4.3478 | 0.79784 | 0.64504 | 3.69 |
| 226 | *GABRB1* | 4.0898 | 3.9709 | 3.0058 | 2.1828 | 3.69 |
| 227 | *CHRM1* | 4.6908 | 4.4163 | 1.9553 | -0.32327 | 3.69 |
| 228 | *ASCL5* | 3.7728 | 3.7511 | 3.5125 | 0.11917 | 3.68 |
| 229 | *CHRM2* | 4.461 | 4.4196 | 2.1556 | 0.25134 | 3.68 |
| 230 | *MED12L* | 4.1437 | 3.6606 | 3.2281 | 0.039331 | 3.68 |
| 231 | *ZBTB46* | 3.8267 | 3.6601 | 3.5449 | -1.5719 | 3.68 |
| 232 | *ZBTB16* | 5.0189 | 4.084 | 1.9159 | -0.86412 | 3.67 |
| 233 | *HIST2H4A* | 3.8861 | 3.6594 | 3.4722 | 1.7769 | 3.67 |
| 234 | *KLF5* | 4.4647 | 3.5953 | 2.9534 | -0.89822 | 3.67 |
| 235 | *OR2Y1* | 4.6846 | 3.3511 | 2.959 | 0.036582 | 3.66 |
| 236 | *HIST1H2BF* | 5.0862 | 3.4436 | 2.4432 | -0.39171 | 3.66 |
| 237 | *DYNC1I2* | 5.2307 | 3.0128 | 2.724 | 0.13241 | 3.66 |
| 238 | *ZNF276* | 4.0256 | 3.8674 | 3.0729 | -0.43305 | 3.66 |
| 239 | *DNM3* | 3.9728 | 3.7264 | 3.2612 | -0.28961 | 3.65 |
| 240 | *SPRED3* | 4.2299 | 3.4113 | 3.318 | -1.9519 | 3.65 |
| 241 | *H1FOO* | 4.7805 | 4.1854 | 1.9932 | -1.0419 | 3.65 |
| 242 | *ZNF554* | 4.7511 | 4.0173 | 2.1899 | -1.8106 | 3.65 |
| 243 | *CYR61* | 4.2363 | 3.8182 | 2.9024 | 0.51492 | 3.65 |
| 244 | *SULT1E1* | 4.8399 | 4.0104 | 2.106 | -2.1321 | 3.65 |
| 245 | *PROCA1* | 4.0883 | 3.6076 | 3.2362 | 0.83433 | 3.64 |
| 246 | *PPIP5K2* | 4.7936 | 3.6488 | 2.4825 | -0.39192 | 3.64 |
| 247 | *LRCH2* | 4.9332 | 4.5919 | 1.3994 | 0.81245 | 3.64 |
| 248 | *ROBO1* | 4.5119 | 3.4547 | 2.9576 | -2.3487 | 3.64 |
| 249 | *TRIM73* | 4.2895 | 4.1971 | 2.4359 | 2.0096 | 3.64 |
| 250 | *TMEM2* | 4.9873 | 3.1194 | 2.8112 | -1.0898 | 3.64 |
| 251 | *SPHK1* | 3.799 | 3.7064 | 3.4112 | -1.4064 | 3.64 |
| 252 | *ZAR1L* | 4.3078 | 3.8102 | 2.7961 | -0.76621 | 3.64 |
| 253 | *SYNRG* | 5.6145 | 5.3887 | -0.09054 | -1.3114 | 3.64 |
| 254 | *TMTC2* | 4.367 | 3.6014 | 2.944 | -1.6302 | 3.64 |
| 255 | *MYO1G* | 4.6967 | 3.397 | 2.8173 | 0.38387 | 3.64 |
| 256 | *MPRIP* | 4.6676 | 3.3311 | 2.9095 | 0.018783 | 3.64 |
| 257 | *LMOD2* | 4.4455 | 3.2719 | 3.1848 | -0.06266 | 3.63 |
| 258 | *IL1RAPL1* | 4.475 | 3.2449 | 3.1819 | 1.3901 | 3.63 |
| 259 | *SAMSN1* | 5.3784 | 4.4674 | 1.0462 | -1.165 | 3.63 |
| 260 | *GNS* | 5.1842 | 3.1029 | 2.6005 | -2.0725 | 3.63 |
| 261 | *SLC7A1* | 4.5507 | 3.4575 | 2.8704 | -1.0702 | 3.63 |
| 262 | *IGFL2* | 5.4422 | 4.8722 | 0.55279 | -1.0582 | 3.62 |
| 263 | *MDH1B* | 4.5141 | 3.2375 | 3.1125 | 1.5365 | 3.62 |
| 264 | *BTNL3* | 4.5465 | 3.576 | 2.7413 | 2.6827 | 3.62 |
| 265 | *KNOP1* | 5.0373 | 3.0412 | 2.7814 | -0.73565 | 3.62 |
| 266 | *PGM1* | 5.7068 | 4.614 | 0.53673 | 0.43624 | 3.62 |
| 267 | *PCDHGB2* | 3.9716 | 3.8068 | 3.0783 | -0.42697 | 3.62 |
| 268 | *TMEM128* | 4.9902 | 4.7318 | 1.1321 | 0.87572 | 3.62 |
| 269 | *KCNC3* | 4.3413 | 3.9014 | 2.6035 | 0.14329 | 3.62 |

| | | | | | | |
|---|---|---|---|---|---|---|
| 270 | *ELOVL4* | 5.4319 | 3.6107 | 1.8024 | -0.89866 | 3.62 |
| 271 | *ICAM2* | 4.6549 | 4.3302 | 1.8562 | -1.5382 | 3.61 |
| 272 | *LPPR2* | 5.3461 | 3.6675 | 1.8267 | -0.93489 | 3.61 |
| 273 | *RHOXF2* | 4.4241 | 3.2598 | 3.148 | -0.69626 | 3.61 |
| 274 | *HES1* | 5.7656 | 2.862 | 2.204 | -0.90229 | 3.61 |
| 275 | *TBX1* | 5.186 | 3.1215 | 2.5213 | 2.4144 | 3.61 |
| 276 | *DEFB126* | 4.5878 | 4.4239 | 1.8139 | -0.13094 | 3.61 |
| 277 | *SEC14L1* | 4.1372 | 3.3929 | 3.2857 | -0.2889 | 3.61 |
| 278 | *PTN* | 4.9353 | 2.9729 | 2.8993 | 2.6056 | 3.60 |
| 279 | *LBP* | 3.844 | 3.5308 | 3.4316 | -1.7134 | 3.60 |
| 280 | *HIST1H1T* | 5.7922 | 5.5653 | -0.55554 | -0.96661 | 3.60 |
| 281 | *EFHB* | 4.5488 | 3.6679 | 2.5799 | 0.15458 | 3.60 |
| 282 | *TAS2R41* | 4.8047 | 3.3964 | 2.5809 | -1.2202 | 3.59 |
| 283 | *OVCH1* | 4.4431 | 3.197 | 3.1383 | 2.0964 | 3.59 |
| 284 | *BSPH1* | 5.183 | 4.2216 | 1.3728 | 0.19797 | 3.59 |
| 285 | *TMCO6* | 4.5116 | 3.6107 | 2.6474 | -1.3242 | 3.59 |
| 286 | *LUC7L2* | 3.8557 | 3.46 | 3.4525 | 1.106 | 3.59 |
| 287 | *TCEAL1* | 4.1337 | 3.5769 | 3.0565 | 1.5708 | 3.59 |
| 288 | *MPHOSPH6* | 4.9843 | 2.8934 | 2.8759 | 1.4317 | 3.58 |
| 289 | *ARVCF* | 4.6714 | 4.0747 | 2.0064 | -1.6711 | 3.58 |
| 290 | *SYTL1* | 4.2648 | 3.5297 | 2.9502 | 1.2424 | 3.58 |
| 291 | *RCBTB1* | 4.2788 | 3.6181 | 2.8445 | -0.9703 | 3.58 |
| 292 | *GOLGA7B* | 5.0016 | 4.2615 | 1.475 | -0.33174 | 3.58 |
| 293 | *TSSC1* | 4.8976 | 3.3711 | 2.4632 | -1.1585 | 3.58 |
| 294 | *TNFAIP6* | 5.5855 | 3.1045 | 2.0334 | 0.15554 | 3.57 |
| 295 | *SPARCL1* | 5.491 | 3.0741 | 2.1558 | -0.56769 | 3.57 |
| 296 | *TUBE1* | 4.3943 | 3.8627 | 2.4574 | 0.53549 | 3.57 |
| 297 | *LIPM* | 4.1984 | 3.7047 | 2.8089 | 2.5867 | 3.57 |
| 298 | *BEX5* | 4.3026 | 3.6585 | 2.7465 | 1.0399 | 3.57 |
| 299 | *PPP4R4* | 4.8962 | 3.7814 | 2.0278 | 1.7767 | 3.57 |
| 300 | *MMACHC* | 4.0969 | 4.0409 | 2.5612 | -0.03924 | 3.57 |
| 301 | *CABLES1* | 5.4833 | 4.2573 | 0.95819 | -0.07166 | 3.57 |
| 302 | *C9orf172* | 4.8349 | 4.4789 | 1.3831 | -0.45786 | 3.57 |
| 303 | *MUC12* | 5.2903 | 3.0405 | 2.3622 | -3.3136 | 3.56 |
| 304 | *HOXB6* | 4.2078 | 3.6954 | 2.7859 | -0.40396 | 3.56 |
| 305 | *LGALS12* | 5.4889 | 2.6488 | 2.5486 | -0.93727 | 3.56 |
| 306 | *SCN8A* | 4.3176 | 4.1588 | 2.2096 | 0.064672 | 3.56 |
| 307 | *SRI* | 4.6347 | 4.3797 | 1.6714 | 0.3432 | 3.56 |
| 308 | *MAATS1* | 4.7805 | 4.3135 | 1.5802 | 1.059 | 3.56 |
| 309 | *IFNA4* | 3.7072 | 3.4927 | 3.4699 | 0.00091 | 3.56 |
| 310 | *GPR75-ASB3* | 5.4004 | 3.009 | 2.2593 | 0.16815 | 3.56 |
| 311 | *DEFB1* | 4.4269 | 3.8116 | 2.43 | -1.5059 | 3.56 |
| 312 | *SCG3* | 4.1382 | 3.4448 | 3.0824 | -0.39494 | 3.56 |
| 313 | *ESPNL* | 4.4426 | 3.3266 | 2.8864 | -0.28752 | 3.55 |
| 314 | *SEPHS1* | 4.4256 | 3.8935 | 2.3315 | -1.4171 | 3.55 |
| 315 | *ARHGEF10L* | 4.2295 | 3.8801 | 2.54 | 0.6301 | 3.55 |

| | | | | | | |
|---|---|---|---|---|---|---|
| 316 | *SERPINA9* | 3.9031 | 3.5319 | 3.2091 | 0.39144 | 3.55 |
| 317 | *GNG7* | 4.9265 | 3.9092 | 1.8058 | -0.84409 | 3.55 |
| 318 | *TBC1D24* | 4.4056 | 4.2296 | 2.0037 | -0.22016 | 3.55 |
| 319 | *HTR1A* | 4.6258 | 4.1067 | 1.9024 | 0.31525 | 3.54 |
| 320 | *DAZL* | 5.0951 | 4.1225 | 1.4081 | 0.4969 | 3.54 |
| 321 | *GAN* | 5.2823 | 4.1078 | 1.2332 | -1.8478 | 3.54 |
| 322 | *TCTN2* | 4.0188 | 3.4492 | 3.1446 | -0.21159 | 3.54 |
| 323 | *IGSF11* | 3.9609 | 3.6296 | 3.015 | 0.27793 | 3.54 |
| 324 | *TMEM14B* | 4.8303 | 3.5921 | 2.1817 | -0.20928 | 3.53 |
| 325 | *RNF183* | 5.2532 | 4.9204 | 0.4304 | -1.5369 | 3.53 |
| 326 | *C4orf51* | 4.8541 | 2.8851 | 2.8644 | 0.15299 | 3.53 |
| 327 | *TGFB3* | 5.5284 | 3.4284 | 1.6434 | -2.5153 | 3.53 |
| 328 | *MAGIX* | 4.6406 | 3.2931 | 2.6604 | 2.5944 | 3.53 |
| 329 | *SLC9A5* | 4.334 | 3.3023 | 2.9545 | -2.817 | 3.53 |
| 330 | *TMEM192* | 4.4851 | 3.5966 | 2.5083 | 0.80493 | 3.53 |
| 331 | *SPAG11A* | 4.9042 | 4.7886 | 0.89563 | 0.52995 | 3.53 |
| 332 | *PGF* | 4.2615 | 3.2359 | 3.0814 | 1.7287 | 3.53 |
| 333 | *VSIG10* | 3.6207 | 3.4832 | 3.4738 | -0.93506 | 3.53 |
| 334 | *CC2D1A* | 4.14 | 4.0393 | 2.3906 | -1.3766 | 3.52 |
| 335 | *TCEAL2* | 5.241 | 5.2207 | 0.10802 | -1.6075 | 3.52 |
| 336 | *RWDD3* | 5.0796 | 3.0775 | 2.4112 | 0.11365 | 3.52 |
| 337 | *SLC25A31* | 5.6173 | 2.8497 | 2.0999 | -1.3443 | 3.52 |
| 338 | *ZNF577* | 5.7169 | 4.1666 | 0.68288 | -1.1046 | 3.52 |
| 339 | *ONECUT1* | 4.8599 | 2.9245 | 2.7796 | 0.31049 | 3.52 |
| 340 | *IRX4* | 5.465 | 4.9693 | 0.12418 | -0.40692 | 3.52 |
| 341 | *CA2* | 4.2584 | 4.1241 | 2.1746 | 2.1307 | 3.52 |
| 342 | *DYNLT1* | 4.0375 | 3.9498 | 2.5672 | -0.51673 | 3.52 |
| 343 | *C17orf112* | 3.666 | 3.6207 | 3.2676 | 2.9027 | 3.52 |
| 344 | *LECT1* | 4.2856 | 3.6107 | 2.6565 | -0.45973 | 3.52 |
| 345 | *ASAP1* | 4.9689 | 3.8067 | 1.7754 | 0.11212 | 3.52 |
| 346 | *SLFN12L* | 5.5471 | 4.1127 | 0.88736 | -1.5128 | 3.52 |
| 347 | *PPP1R3C* | 5.2374 | 3.2357 | 2.0712 | 0.40775 | 3.51 |
| 348 | *ZNF821* | 5.5966 | 5.3648 | -0.4191 | -1.9305 | 3.51 |
| 349 | *TSTD2* | 4.1312 | 3.491 | 2.9114 | -1.7931 | 3.51 |
| 350 | *TPSG1* | 4.6074 | 4.2194 | 1.7054 | -1.0599 | 3.51 |
| 351 | *OPALIN* | 4.6143 | 4.519 | 1.3985 | -0.4082 | 3.51 |
| 352 | *XAGE1B* | 4.1691 | 3.6874 | 2.6737 | -0.0401 | 3.51 |
| 353 | *NDFIP2* | 5.025 | 3.9994 | 1.4987 | -0.781 | 3.51 |
| 354 | *ATAD3A* | 5.3535 | 5.3378 | -0.17042 | -1.916 | 3.51 |
| 355 | *HTN1* | 4.4866 | 4.1614 | 1.8721 | 1.3968 | 3.51 |
| 356 | *TGM3* | 4.7319 | 3.2125 | 2.5724 | -2.0375 | 3.51 |
| 357 | *OAZ2* | 3.6803 | 3.6654 | 3.1599 | 0.93382 | 3.50 |
| 358 | *CRTC1* | 4.4597 | 3.1697 | 2.8706 | -2.2078 | 3.50 |
| 359 | *ZNF202* | 5.2272 | 3.8658 | 1.4017 | -1.3287 | 3.50 |
| 360 | *PAPSS2* | 3.8417 | 3.4444 | 3.2021 | -2.3517 | 3.50 |
| 361 | *TECPR2* | 4.5128 | 3.3211 | 2.6542 | -1.072 | 3.50 |

| | | | | | | |
|---|---|---|---|---|---|---|
| 362 | *CLEC18B* | 4.5869 | 3.3354 | 2.563 | -1.0901 | 3.50 |
| 363 | *C18orf42* | 4.5756 | 3.1114 | 2.7965 | 1.7997 | 3.49 |
| 364 | *ERVMER34-1* | 5.382 | 3.8141 | 1.2837 | -0.53713 | 3.49 |
| 365 | *NOMO1* | 4.8256 | 4.386 | 1.2617 | -0.8715 | 3.49 |
| 366 | *PDP1* | 4.7326 | 3.0177 | 2.7217 | 2.1548 | 3.49 |
| 367 | *PARPBP* | 3.8042 | 3.6469 | 3.0196 | -1.6917 | 3.49 |
| 368 | *MPZ* | 4.1294 | 3.2878 | 3.0487 | 0.28201 | 3.49 |
| 369 | *WAS* | 3.9411 | 3.684 | 2.8318 | -1.8157 | 3.49 |
| 370 | *GABBR2* | 4.5656 | 4.0415 | 1.8477 | -1.536 | 3.48 |
| 371 | *FAM198B* | 4.5412 | 3.5444 | 2.3691 | 0.14895 | 3.48 |
| 372 | *RCN2* | 4.8834 | 3.0374 | 2.5337 | 0.033945 | 3.48 |
| 373 | *DNAH10* | 5.1493 | 4.4813 | 0.81863 | -2.6476 | 3.48 |
| 374 | *CCNG2* | 4.87 | 2.8532 | 2.7256 | -2.7274 | 3.48 |
| 375 | *BLOC1S2* | 5.4031 | 3.0709 | 1.9674 | 0.26985 | 3.48 |
| 376 | *ARPC1B* | 4.9398 | 4.5636 | 0.9361 | 0.67857 | 3.48 |
| 377 | *TRIM39* | 4.8098 | 4.1512 | 1.4756 | -0.72877 | 3.48 |
| 378 | *G3BP2* | 4.8203 | 4.5125 | 1.0997 | -1.2411 | 3.48 |
| 379 | *ZNF383* | 4.526 | 3.4762 | 2.4296 | -1.6033 | 3.48 |
| 380 | *HACD4* | 4.6296 | 4.4224 | 1.3782 | -2.0593 | 3.48 |
| 381 | *MAPK14* | 3.8394 | 3.5603 | 3.027 | 0.94996 | 3.48 |
| 382 | *KMT2C* | 3.9948 | 3.7761 | 2.6518 | -0.71919 | 3.47 |
| 383 | *ASGR1* | 4.3733 | 3.1633 | 2.8828 | -0.93727 | 3.47 |
| 384 | *PCDHGB4* | 4.5118 | 4.178 | 1.727 | 0.24364 | 3.47 |
| 385 | *LCE2C* | 4.4069 | 4.2412 | 1.7679 | -0.60604 | 3.47 |
| 386 | *CNTROB* | 4.8781 | 4.8066 | 0.73023 | 0.035999 | 3.47 |
| 387 | *UBB* | 4.6628 | 4.3789 | 1.3663 | 1.0461 | 3.47 |
| 388 | *CPNE8* | 4.3609 | 3.2918 | 2.7534 | 1.2786 | 3.47 |
| 389 | *TRIL* | 5.4459 | 3.3632 | 1.5937 | -0.82366 | 3.47 |
| 390 | *CASQ2* | 5.4479 | 3.3253 | 1.6268 | -0.16451 | 3.47 |
| 391 | *B9D2* | 3.971 | 3.2944 | 3.1323 | 0.92068 | 3.47 |
| 392 | *OPN1MW* | 4.6396 | 4.2761 | 1.4705 | -0.8202 | 3.46 |
| 393 | *FAM78A* | 4.0901 | 3.2149 | 3.0805 | 0.078506 | 3.46 |
| 394 | *ANKRD9* | 4.7555 | 3.2698 | 2.3533 | 0.90985 | 3.46 |
| 395 | *PDCL2* | 3.8122 | 3.8116 | 2.7543 | -0.22609 | 3.46 |
| 396 | *CCR7* | 4.167 | 3.232 | 2.9789 | 0.17412 | 3.46 |
| 397 | *MATN2* | 4.7509 | 3.7009 | 1.9241 | -0.51539 | 3.46 |
| 398 | *SCIMP* | 4.2289 | 3.7073 | 2.437 | 0.62977 | 3.46 |
| 399 | *DMRTC2* | 4.5366 | 2.9572 | 2.8786 | 0.30333 | 3.46 |
| 400 | *SNX1* | 5.2903 | 5.0169 | 0.06449 | -1.3275 | 3.46 |
| 401 | *UBP1* | 4.8801 | 2.9887 | 2.5028 | -1.1325 | 3.46 |
| 402 | *ADSSL1* | 4.7243 | 3.1956 | 2.4508 | -1.1904 | 3.46 |
| 403 | *TMEM64* | 4.7121 | 3.9613 | 1.6907 | 0.57975 | 3.45 |
| 404 | *NMNAT3* | 4.6235 | 3.6173 | 2.1189 | 1.6763 | 3.45 |
| 405 | *WFDC3* | 4.6284 | 3.1583 | 2.572 | 0.54388 | 3.45 |
| 406 | *ERICH6B* | 3.9651 | 3.3815 | 3.0116 | 0.060481 | 3.45 |
| 407 | *KRTAP9-8* | 4.3765 | 3.5604 | 2.4204 | 0.18521 | 3.45 |

| | | | | | | |
|---|---|---|---|---|---|---|
| 408 | *GYPA* | 5.1039 | 3.868 | 1.383 | -0.86517 | 3.45 |
| 409 | *PLCD3* | 4.9021 | 4.3707 | 1.0767 | -0.29602 | 3.45 |
| 410 | *DIABLO* | 4.7667 | 4.401 | 1.1801 | -0.08298 | 3.45 |
| 411 | *GFRAL* | 4.6572 | 4.331 | 1.3531 | -0.45965 | 3.45 |
| 412 | *OR56A3* | 4.4217 | 4.1164 | 1.7937 | -1.8366 | 3.44 |
| 413 | *TSACC* | 5.7141 | 2.8503 | 1.7668 | -0.01483 | 3.44 |
| 414 | *CNN2* | 4.1443 | 3.7894 | 2.3959 | 0.57904 | 3.44 |
| 415 | *GLIS1* | 4.0373 | 3.5865 | 2.7007 | 0.088476 | 3.44 |
| 416 | *OR4C6* | 4.4219 | 3.2883 | 2.6115 | 2.0609 | 3.44 |
| 417 | *TMEM114* | 4.1892 | 3.6122 | 2.5185 | -0.0994 | 3.44 |
| 418 | *GPR151* | 4.6739 | 3.3977 | 2.2425 | 0.58233 | 3.44 |
| 419 | *CD28* | 4.6614 | 3.8878 | 1.7601 | -2.0218 | 3.44 |
| 420 | *OR2W1* | 4.3767 | 3.0244 | 2.9062 | 2.1939 | 3.44 |
| 421 | *SORCS1* | 4.9176 | 3.1063 | 2.2786 | 1.4651 | 3.43 |
| 422 | *RNF165* | 4.8326 | 3.7422 | 1.7229 | -2.0483 | 3.43 |
| 423 | *MAB21L2* | 4.982 | 4.9344 | 0.37779 | -1.5611 | 3.43 |
| 424 | *TXNIP* | 4.2879 | 3.2374 | 2.7616 | 1.2519 | 3.43 |
| 425 | *IRS4* | 4.4587 | 3.5932 | 2.2348 | -2.4386 | 3.43 |
| 426 | *PDE11A* | 5.0447 | 2.9212 | 2.3196 | -0.96407 | 3.43 |
| 427 | *FXYD6-FXYD2* | 5.0169 | 4.1943 | 1.0686 | -0.41131 | 3.43 |
| 428 | *PRR18* | 5.3369 | 4.0594 | 0.87366 | -0.46956 | 3.42 |
| 429 | *RLBP1* | 5.6371 | 3.2202 | 1.4107 | 0.028035 | 3.42 |
| 430 | *SNAI2* | 3.9655 | 3.6686 | 2.626 | -1.0216 | 3.42 |
| 431 | *PPP1R2* | 5.1235 | 4.5649 | 0.57144 | -2.5478 | 3.42 |
| 432 | *CRCT1* | 4.763 | 2.9658 | 2.5235 | 0.67866 | 3.42 |
| 433 | *PRSS33* | 4.9082 | 3.611 | 1.7296 | 0.41418 | 3.42 |
| 434 | *CERS6* | 4.8599 | 3.6508 | 1.737 | -1.8212 | 3.42 |
| 435 | *ZNF107* | 4.1654 | 3.7886 | 2.2926 | -0.64519 | 3.42 |
| 436 | *MOB3A* | 4.9509 | 2.7863 | 2.5076 | 1.1202 | 3.41 |
| 437 | *C1orf112* | 3.988 | 3.3943 | 2.861 | -0.32552 | 3.41 |
| 438 | *TMEM72* | 4.1953 | 3.637 | 2.4095 | -1.3955 | 3.41 |
| 439 | *OR10W1* | 4.5597 | 4.515 | 1.1657 | 0.1562 | 3.41 |
| 440 | *RSBN1L* | 4.3282 | 3.0782 | 2.8327 | -0.85586 | 3.41 |
| 441 | *CARD17* | 4.8431 | 4.358 | 1.0267 | 0.27203 | 3.41 |
| 442 | *NPFFR1* | 3.6895 | 3.3285 | 3.2075 | 0.091234 | 3.41 |
| 443 | *KPNA5* | 4.1496 | 3.0925 | 2.9817 | -0.78313 | 3.41 |
| 444 | *TDRD7* | 4.913 | 4.8798 | 0.42976 | 0.42427 | 3.41 |
| 445 | *ZNF230* | 4.2575 | 3.224 | 2.7345 | -0.30038 | 3.41 |
| 446 | *TTC12* | 4.364 | 4.0653 | 1.781 | -0.49034 | 3.40 |
| 447 | *GCNT7* | 5.5777 | 4.675 | -0.04444 | -1.7188 | 3.40 |
| 448 | *UBQLNL* | 4.9301 | 3.9683 | 1.3018 | 0.74561 | 3.40 |
| 449 | *PBDC1* | 4.0363 | 3.2874 | 2.8708 | 0.88384 | 3.40 |
| 450 | *ZNF845* | 4.8514 | 3.1987 | 2.1383 | 1.5933 | 3.40 |
| 451 | *DCLRE1C* | 4.1249 | 3.8065 | 2.2551 | -2.5478 | 3.40 |
| 452 | *ZNF787* | 4.3755 | 3.3253 | 2.4828 | -1.9305 | 3.39 |
| 453 | *KIF20B* | 4.5971 | 3.3886 | 2.1931 | -1.8963 | 3.39 |

| | | | | | | |
|---|---|---|---|---|---|---|
| 454 | *C4orf45* | 5.2605 | 4.2683 | 0.64613 | -0.86793 | 3.39 |
| 455 | *LRRC75A* | 4.0025 | 3.1671 | 3.0021 | -0.46711 | 3.39 |
| 456 | *GAMT* | 4.3099 | 2.9835 | 2.8774 | -1.3324 | 3.39 |
| 457 | *A3GALT2* | 4.5561 | 3.2834 | 2.33 | -0.87642 | 3.39 |
| 458 | *HMG20A* | 4.6504 | 3.1866 | 2.3316 | -0.45484 | 3.39 |
| 459 | *ZNF184* | 4.7927 | 2.938 | 2.437 | 1.9554 | 3.39 |
| 460 | *YWHAQ* | 3.7551 | 3.4334 | 2.9785 | -0.31091 | 3.39 |
| 461 | *AMFR* | 3.9535 | 3.3416 | 2.8716 | 1.3708 | 3.39 |
| 462 | *KRTAP25-1* | 5.3975 | 4.2581 | 0.51092 | 0.091323 | 3.39 |
| 463 | *OXTR* | 3.7562 | 3.3206 | 3.0843 | 0.62615 | 3.39 |
| 464 | *SDR16C5* | 5.9577 | 4.4047 | -0.2075 | -1.5869 | 3.38 |
| 465 | *DGKZ* | 4.1375 | 3.0879 | 2.9287 | 0.60294 | 3.38 |
| 466 | *TNF* | 4.8549 | 3.6501 | 1.649 | -0.87837 | 3.38 |
| 467 | *CELF2* | 3.8446 | 3.5552 | 2.7539 | -0.05571 | 3.38 |
| 468 | *STEAP2* | 4.6743 | 3.0299 | 2.4472 | 1.4592 | 3.38 |
| 469 | *MPP4* | 4.3579 | 3.6616 | 2.1307 | -1.7876 | 3.38 |
| 470 | *GFOD1* | 4.3358 | 3.3435 | 2.4703 | -2.7705 | 3.38 |
| 471 | *ASIC3* | 4.7748 | 4.2708 | 1.1012 | 0.12142 | 3.38 |
| 472 | *ABCB10* | 3.98 | 3.4024 | 2.7595 | 0.91511 | 3.38 |
| 473 | *TNS2* | 4.3989 | 3.8706 | 1.8701 | 0.659 | 3.38 |
| 474 | *SPINK13* | 4.9793 | 4.0475 | 1.1118 | -2.075 | 3.38 |
| 475 | *KDELC1* | 4.3553 | 3.8651 | 1.9162 | -1.066 | 3.38 |
| 476 | *KIAA1671* | 3.6139 | 3.5007 | 3.0089 | 0.70341 | 3.37 |
| 477 | *DUSP10* | 3.8784 | 3.3466 | 2.8978 | -1.2246 | 3.37 |
| 478 | *CLEC19A* | 4.3877 | 3.9243 | 1.81 | -1.4078 | 3.37 |
| 479 | *DMTN* | 4.333 | 2.9399 | 2.8465 | -1.4894 | 3.37 |
| 480 | *EVC2* | 4.9032 | 3.7159 | 1.4974 | 0.012816 | 3.37 |
| 481 | *TRIM72* | 5.0085 | 3.9247 | 1.1803 | -0.30611 | 3.37 |
| 482 | *C3orf56* | 4.9806 | 4.3031 | 0.82809 | -0.59634 | 3.37 |
| 483 | *RAP1GDS1* | 4.3923 | 3.3654 | 2.3435 | -0.06975 | 3.37 |
| 484 | *SPI1* | 4.5384 | 3.833 | 1.7288 | 0.45098 | 3.37 |
| 485 | *SLC25A35* | 4.1665 | 3.5818 | 2.3501 | -1.0315 | 3.37 |
| 486 | *CMA1* | 4.5607 | 3.2519 | 2.2836 | 0.20268 | 3.37 |
| 487 | *COG5* | 5.7012 | 3.534 | 0.85953 | 0.8042 | 3.36 |
| 488 | *APBA3* | 4.2746 | 3.8926 | 1.9262 | -0.63643 | 3.36 |
| 489 | *PPP1R14D* | 4.2865 | 3.2395 | 2.5647 | -2.2443 | 3.36 |
| 490 | *ATXN3L* | 4.6471 | 4.162 | 1.2802 | 0.62467 | 3.36 |
| 491 | *SHD* | 3.9758 | 3.2267 | 2.8772 | -0.79108 | 3.36 |
| 492 | *CACNA1I* | 5.02 | 4.1806 | 0.87562 | -1.3496 | 3.36 |
| 493 | *OR5T2* | 4.8094 | 3.6908 | 1.5757 | 0.2573 | 3.36 |
| 494 | *ASRGL1* | 5.7501 | 4.6803 | -0.3579 | -0.73668 | 3.36 |
| 495 | *WFDC5* | 4.5484 | 3.0884 | 2.4353 | -1.0564 | 3.36 |
| 496 | *COX7B2* | 4.1424 | 3.8727 | 2.0544 | -1.274 | 3.36 |
| 497 | *HSPA4* | 3.7336 | 3.1775 | 3.1576 | 2.8072 | 3.36 |
| 498 | *TLR1* | 4.125 | 3.2235 | 2.7131 | -1.1442 | 3.35 |
| 499 | *COX8C* | 4.1417 | 3.086 | 2.8338 | 0.37321 | 3.35 |

| | | | | | | |
|---|---|---|---|---|---|---|
| 500 | *OLFML1* | 3.9896 | 3.8161 | 2.2492 | -1.1704 | 3.35 |
| 501 | *TMEM107* | 4.6486 | 3.8119 | 1.5906 | 0.093927 | 3.35 |
| 502 | *DAZ4* | 5.5221 | 3.9229 | 0.60282 | -0.85018 | 3.35 |
| 503 | *RAB5B* | 4.3906 | 3.9136 | 1.7421 | -0.39079 | 3.35 |
| 504 | *ZNF793* | 4.2827 | 3.9447 | 1.8173 | 0.25544 | 3.35 |
| 505 | *TPRG1* | 4.4291 | 3.8882 | 1.7254 | -0.59195 | 3.35 |
| 506 | *AFAP1L2* | 3.9265 | 3.2234 | 2.8902 | -1.1178 | 3.35 |
| 507 | *KLHL13* | 3.9574 | 3.1534 | 2.9219 | -1.0088 | 3.34 |
| 508 | *LELP1* | 4.2185 | 3.5843 | 2.2209 | -0.62792 | 3.34 |
| 509 | *HSFX2* | 5.2203 | 4.6666 | 0.13639 | -0.45989 | 3.34 |
| 510 | *NPTX2* | 4.9264 | 3.3902 | 1.7046 | 0.026096 | 3.34 |
| 511 | *TMPO* | 3.814 | 3.4658 | 2.7367 | 1.2343 | 3.34 |
| 512 | *ATP11AUN* | 4.2564 | 4.1914 | 1.5628 | 0.42428 | 3.34 |
| 513 | *TIA1* | 4.5357 | 4.0745 | 1.3997 | 0.45049 | 3.34 |
| 514 | *KIAA0408* | 4.5899 | 3.1701 | 2.2416 | 1.5006 | 3.33 |
| 515 | *FKBP3* | 3.9381 | 3.4222 | 2.6404 | -1.6731 | 3.33 |
| 516 | *NKD1* | 5.3084 | 4.453 | 0.23777 | -0.37913 | 3.33 |
| 517 | *SEMG1* | 3.8335 | 3.1662 | 2.9991 | 1.5464 | 3.33 |
| 518 | *APCS* | 4.1429 | 4.1266 | 1.7222 | -0.89438 | 3.33 |
| 519 | *CDCA7L* | 4.4685 | 4.4277 | 1.0949 | -0.95747 | 3.33 |
| 520 | *HTR1E* | 4.1004 | 3.2762 | 2.611 | 0.47111 | 3.33 |
| 521 | *GGA1* | 4.8399 | 3.2619 | 1.8838 | -2.0351 | 3.33 |
| 522 | *RANBP3* | 5.8102 | 5.0392 | -0.87056 | -2.2094 | 3.33 |
| 523 | *C22orf31* | 5.4644 | 4.7075 | -0.19735 | -1.127 | 3.32 |
| 524 | *MFSD4* | 5.0982 | 4.7578 | 0.11456 | -2.3222 | 3.32 |
| 525 | *KRT76* | 5.016 | 3.7102 | 1.2372 | -1.5908 | 3.32 |
| 526 | *PTPRM* | 4.1377 | 2.9762 | 2.8491 | 2.1758 | 3.32 |
| 527 | *FAM50B* | 4.3775 | 3.683 | 1.9024 | -1.2864 | 3.32 |
| 528 | *DYRK2* | 5.2386 | 4.4146 | 0.3084 | 0.13194 | 3.32 |
| 529 | *PSMB10* | 4.5592 | 4.1688 | 1.2241 | -0.44353 | 3.32 |
| 530 | *UPK3BL* | 4.8198 | 2.9035 | 2.2222 | 0.009975 | 3.32 |
| 531 | *NKX3-1* | 4.4489 | 2.8255 | 2.671 | -1.1012 | 3.32 |
| 532 | *ZNF613* | 4.8253 | 3.7249 | 1.3938 | -1.2719 | 3.31 |
| 533 | *ZNF428* | 4.011 | 3.4784 | 2.4525 | -1.4983 | 3.31 |
| 534 | *ANGPTL6* | 4.9233 | 4.4037 | 0.61081 | -1.0997 | 3.31 |
| 535 | *FEZF1* | 5.2374 | 4.2681 | 0.4314 | 0.37368 | 3.31 |
| 536 | *MPPED2* | 3.8424 | 3.84 | 2.2498 | -1.265 | 3.31 |
| 537 | *CACYBP* | 4.8715 | 4.3186 | 0.7417 | 0.22581 | 3.31 |
| 538 | *PITHD1* | 5.165 | 4.2773 | 0.48089 | -0.59272 | 3.31 |
| 539 | *TMEM189-UBE2V1* | 5.0669 | 4.8015 | 0.05283 | -1.0196 | 3.31 |
| 540 | *TRIB3* | 4.0959 | 3.4799 | 2.3421 | -0.24661 | 3.31 |
| 541 | *ANKAR* | 5.0443 | 4.069 | 0.80282 | -2.2811 | 3.31 |
| 542 | *LOC728392* | 4.8604 | 3.8314 | 1.2226 | -0.63643 | 3.30 |
| 543 | *DCAF8L2* | 4.2346 | 3.8883 | 1.7899 | -0.688 | 3.30 |
| 544 | *PEX5* | 4.5729 | 2.7409 | 2.596 | -1.984 | 3.30 |
| 545 | *LIMS1* | 4.2568 | 4.1034 | 1.5444 | -0.80727 | 3.30 |

| | | | | | | |
|---|---|---|---|---|---|---|
| 546 | *AP4B1* | 4.9076 | 3.4323 | 1.5625 | -0.38408 | 3.30 |
| 547 | *C11orf70* | 4.0095 | 3.2264 | 2.6664 | -0.15263 | 3.30 |
| 548 | *SLC35A1* | 4.0941 | 3.4657 | 2.337 | 1.0432 | 3.30 |
| 549 | *KLRK1* | 4.8391 | 4.7262 | 0.32263 | -0.1693 | 3.30 |
| 550 | *ZP1* | 4.3732 | 4.2485 | 1.2651 | -0.27982 | 3.30 |
| 551 | *RASA1* | 5.8683 | 2.9668 | 1.0516 | -1.1333 | 3.30 |
| 552 | *FAM46A* | 5.1808 | 4.6675 | 0.03698 | -0.54934 | 3.30 |
| 553 | *GAST* | 3.8258 | 3.1351 | 2.9218 | -0.3952 | 3.29 |
| 554 | *CHL1* | 5.5151 | 3.4005 | 0.96163 | -0.46724 | 3.29 |
| 555 | *MEF2BNB* | 4.3904 | 2.9164 | 2.5698 | -1.245 | 3.29 |
| 556 | *MIER3* | 4.5912 | 3.229 | 2.0507 | 1.115 | 3.29 |
| 557 | *TMEM115* | 3.7939 | 3.5932 | 2.4817 | 2.0159 | 3.29 |
| 558 | *EDEM3* | 3.6995 | 3.2694 | 2.8988 | 2.7863 | 3.29 |
| 559 | *TINAGL1* | 5.7078 | 2.921 | 1.2369 | -1.843 | 3.29 |
| 560 | *C18orf54* | 3.7974 | 3.6777 | 2.39 | 0.60022 | 3.29 |
| 561 | *MARCH1* | 5.2726 | 2.8595 | 1.7325 | -1.8714 | 3.29 |
| 562 | *GJA10* | 3.906 | 3.3301 | 2.6195 | 0.17066 | 3.29 |
| 563 | *ARSB* | 5.3191 | 3.7608 | 0.77552 | 0.20444 | 3.29 |
| 564 | *ZNF782* | 4.5716 | 4.0396 | 1.2421 | 0.57978 | 3.28 |
| 565 | *DNAJC24* | 3.4565 | 3.2579 | 3.1369 | -2.2886 | 3.28 |
| 566 | *RAB33A* | 3.6235 | 3.4513 | 2.7742 | 0.10609 | 3.28 |
| 567 | *OR3A1* | 5.042 | 3.63 | 1.1743 | -0.9304 | 3.28 |
| 568 | *OR10K1* | 5.095 | 4.069 | 0.6814 | -0.59793 | 3.28 |
| 569 | *CEP57L1* | 3.9672 | 3.6915 | 2.1863 | -1.3604 | 3.28 |
| 570 | *FCRL3* | 4.6302 | 3.2367 | 1.9779 | 1.4641 | 3.28 |
| 571 | *BICD1* | 3.6054 | 3.4529 | 2.7864 | -1.4241 | 3.28 |
| 572 | *OR2H1* | 4.9022 | 4.2807 | 0.66176 | -1.6711 | 3.28 |
| 573 | *KLRG2* | 4.5627 | 3.3939 | 1.8841 | 0.073736 | 3.28 |
| 574 | *FXYD2* | 4.257 | 2.9635 | 2.6179 | 2.1489 | 3.28 |
| 575 | *B3GALT4* | 4.6916 | 4.2893 | 0.85669 | -0.85059 | 3.28 |
| 576 | *YIPF4* | 3.6153 | 3.2059 | 3.0128 | -0.67352 | 3.28 |
| 577 | *MRPL16* | 4.8699 | 2.7984 | 2.163 | 0.31617 | 3.28 |
| 578 | *CFDP1* | 4.2156 | 4.0723 | 1.538 | 0.19565 | 3.28 |
| 579 | *UGT2A3* | 4.2256 | 3.6261 | 1.9701 | -0.15279 | 3.27 |
| 580 | *F2RL2* | 5.2369 | 3.5203 | 1.064 | -1.0212 | 3.27 |
| 581 | *TSHZ3* | 3.6564 | 3.4497 | 2.7138 | 0.94387 | 3.27 |
| 582 | *XKRY2* | 4.6635 | 3.5162 | 1.6348 | -0.43473 | 3.27 |
| 583 | *MACC1* | 3.516 | 3.3543 | 2.9436 | -1.8686 | 3.27 |
| 584 | *CLIP4* | 4.8399 | 3.0531 | 1.9205 | 0.33336 | 3.27 |
| 585 | *SLC25A36* | 3.625 | 3.2768 | 2.9114 | 0.17861 | 3.27 |
| 586 | *CRIPT* | 5.4516 | 4.612 | -0.2505 | -5.5233 | 3.27 |
| 587 | *TCEAL5* | 4.6371 | 3.2702 | 1.8981 | -0.2697 | 3.27 |
| 588 | *TBX19* | 4.7427 | 4.7321 | 0.32923 | -0.72102 | 3.27 |
| 589 | *COG2* | 5.235 | 4.9924 | -0.42353 | -0.90386 | 3.27 |
| 590 | *FAM103A1* | 5.265 | 2.7754 | 1.7625 | -2.0249 | 3.27 |
| 591 | *LOC643802* | 5.1526 | 3.262 | 1.3877 | -0.59696 | 3.27 |

| | | | | | | |
|---|---|---|---|---|---|---|
| 592 | *RELN* | 4.2261 | 3.5155 | 2.0581 | 0.37386 | 3.27 |
| 593 | *NLRC4* | 4.5434 | 4.3368 | 0.91849 | -1.4112 | 3.27 |
| 594 | *ECHS1* | 4.0681 | 3.1444 | 2.5855 | -2.2943 | 3.27 |
| 595 | *PYROXD2* | 5.0498 | 2.9665 | 1.7799 | 0.66509 | 3.27 |
| 596 | *KCNK5* | 4.3411 | 3.4486 | 2.0044 | 0.52216 | 3.26 |
| 597 | *CXCL9* | 4.7472 | 3.5447 | 1.502 | 0.49055 | 3.26 |
| 598 | *PABPC4* | 4.4492 | 3.2985 | 2.046 | -1.5908 | 3.26 |
| 599 | *PTH* | 4.2495 | 3.6688 | 1.8752 | -0.20695 | 3.26 |
| 600 | *PSG9* | 5.0204 | 4.4042 | 0.36759 | -1.2908 | 3.26 |
| 601 | *FAM168B* | 4.5195 | 3.0559 | 2.2163 | -1.8909 | 3.26 |
| 602 | *MBD1* | 3.6889 | 3.1418 | 2.9585 | -0.71493 | 3.26 |
| 603 | *USP16* | 3.668 | 3.3712 | 2.7475 | 0.78458 | 3.26 |
| 604 | *C11orf53* | 5.2056 | 3.3247 | 1.2549 | -0.71753 | 3.26 |
| 605 | *RPA4* | 4.7354 | 4.6407 | 0.40878 | -0.23528 | 3.26 |
| 606 | *ZXDC* | 4.1688 | 3.6094 | 2.0021 | -1.5707 | 3.26 |
| 607 | *ANXA1* | 3.939 | 3.7584 | 2.0808 | -2.3293 | 3.26 |
| 608 | *SFMBT2* | 5.2888 | 4.3049 | 0.1829 | -2.0134 | 3.26 |
| 609 | *MCOLN1* | 4.9319 | 4.2268 | 0.61587 | -1.8792 | 3.26 |
| 610 | *USP50* | 3.9653 | 3.1511 | 2.6486 | -0.66425 | 3.26 |
| 611 | *ADAM20* | 4.4558 | 3.6495 | 1.657 | -0.52276 | 3.25 |
| 612 | *DEFB106A* | 4.8338 | 3.7535 | 1.1742 | 0.45681 | 3.25 |
| 613 | *GCSAML* | 4.5032 | 3.5192 | 1.7383 | 0.93904 | 3.25 |
| 614 | *GNG4* | 4.1943 | 3.434 | 2.1306 | -0.23517 | 3.25 |
| 615 | *PLEKHG1* | 3.3598 | 3.2048 | 3.1894 | -2.007 | 3.25 |
| 616 | *WFDC10A* | 3.6785 | 3.133 | 2.9418 | -0.12627 | 3.25 |
| 617 | *SESTD1* | 4.7012 | 4.5229 | 0.52602 | -0.22492 | 3.25 |
| 618 | *TLL2* | 4.1475 | 3.3247 | 2.2755 | 0.31574 | 3.25 |
| 619 | *PPP1R14A* | 5.3828 | 5.0104 | -0.64748 | -2.0384 | 3.25 |
| 620 | *CLHC1* | 5.1397 | 2.8828 | 1.7232 | -0.34456 | 3.25 |
| 621 | *SECISBP2L* | 4.8752 | 3.7588 | 1.1097 | -2.4018 | 3.25 |
| 622 | *NLRP10* | 5.1445 | 2.3828 | 2.2133 | 1.4508 | 3.25 |
| 623 | *DUS4L* | 3.5056 | 3.2519 | 2.9803 | -1.1737 | 3.25 |
| 624 | *CUEDC1* | 3.3937 | 3.2104 | 3.1336 | -0.67378 | 3.25 |
| 625 | *HAUS6* | 3.7566 | 3.2171 | 2.7628 | -1.7171 | 3.25 |
| 626 | *TFAP2A* | 5.3167 | 3.5999 | 0.8183 | -0.72085 | 3.24 |
| 627 | *MEIS1* | 4.1329 | 3.2881 | 2.3135 | 1.674 | 3.24 |
| 628 | *FARP2* | 5.0048 | 3.7213 | 1.0043 | -1.0704 | 3.24 |
| 629 | *VCX3B* | 4.6184 | 4.1636 | 0.94507 | -1.2569 | 3.24 |
| 630 | *CTIF* | 5.8259 | 2.3531 | 1.5398 | -1.8587 | 3.24 |
| 631 | *KIAA1033* | 3.5931 | 3.2696 | 2.8512 | -0.32101 | 3.24 |
| 632 | *ARL11* | 4.3787 | 4.0364 | 1.2984 | 0.4084 | 3.24 |
| 633 | *HIST1H2AC* | 4.5233 | 3.229 | 1.9587 | -2.5663 | 3.24 |
| 634 | *DPYSL4* | 3.3283 | 3.2014 | 3.1796 | -1.5687 | 3.24 |
| 635 | *CMSS1* | 4.1129 | 3.6502 | 1.9451 | 0.29285 | 3.24 |
| 636 | *AKR7A3* | 4.0068 | 3.3047 | 2.3946 | 0.39969 | 3.24 |
| 637 | *ARHGEF15* | 3.8049 | 3.6821 | 2.2163 | -1.4969 | 3.23 |

| | | | | | | |
|---|---|---|---|---|---|---|
| 638 | *FAM167A* | 3.8337 | 3.1033 | 2.7662 | 1.0791 | 3.23 |
| 639 | *CD200* | 4.4508 | 3.1684 | 2.0812 | 1.9874 | 3.23 |
| 640 | *NDUFA3* | 5.2263 | 2.484 | 1.9878 | -1.3097 | 3.23 |
| 641 | *BCAR1* | 3.5161 | 3.2622 | 2.9194 | -2.6024 | 3.23 |
| 642 | *DNMT3L* | 5.0187 | 2.5613 | 2.1129 | -1.1186 | 3.23 |
| 643 | *MPG* | 3.6044 | 3.2254 | 2.8617 | -0.03472 | 3.23 |
| 644 | *TTC9C* | 4.4477 | 3.1581 | 2.0856 | -1.1955 | 3.23 |
| 645 | *F2R* | 3.9146 | 3.4933 | 2.2834 | -0.72157 | 3.23 |
| 646 | *SPATS1* | 4.4356 | 3.3842 | 1.8713 | 0.15281 | 3.23 |
| 647 | *IL4* | 3.7415 | 3.091 | 2.8581 | 2.6592 | 3.23 |
| 648 | *PPIAL4A* | 4.9499 | 4.4487 | 0.29117 | -3.5396 | 3.23 |
| 649 | *FAM221B* | 4.4917 | 3.4668 | 1.7249 | -1.9164 | 3.23 |
| 650 | *TGFB2* | 5.8245 | 2.8866 | 0.97199 | 0.53842 | 3.23 |
| 651 | *OR52A1* | 4.0758 | 4.0022 | 1.6015 | -2.3776 | 3.23 |
| 652 | *ZNF667* | 4.5688 | 4.1543 | 0.9554 | 0.36987 | 3.23 |
| 653 | *PIP5K1B* | 4.3844 | 3.0879 | 2.1956 | -0.05066 | 3.22 |
| 654 | *SAFB* | 4.7629 | 3.7033 | 1.2007 | 0.8235 | 3.22 |
| 655 | *GTF2E1* | 4.9004 | 3.3794 | 1.3796 | 0.24871 | 3.22 |
| 656 | *TRIP12* | 4.654 | 4.0422 | 0.95827 | -0.72771 | 3.22 |
| 657 | *PKIA* | 4.7039 | 3.7472 | 1.2031 | 0.65454 | 3.22 |
| 658 | *ZC3HAV1L* | 3.7425 | 3.637 | 2.2722 | 1.2152 | 3.22 |
| 659 | *PGP* | 4.7481 | 2.9876 | 1.9159 | 0.41821 | 3.22 |
| 660 | *G6PC* | 5.0118 | 4.1718 | 0.46635 | -2.6675 | 3.22 |
| 661 | *M6PR* | 4.7172 | 3.4416 | 1.4903 | -1.6275 | 3.22 |
| 662 | *CPE* | 4.4813 | 2.7938 | 2.3723 | -1.8583 | 3.22 |
| 663 | *RNF182* | 5.159 | 3.4285 | 1.0582 | -1.8642 | 3.22 |
| 664 | *UAP1* | 3.913 | 3.5978 | 2.1265 | -0.34255 | 3.21 |
| 665 | *MIEF2* | 3.778 | 2.9951 | 2.8639 | 0.67989 | 3.21 |
| 666 | *TRAPPC5* | 5.038 | 3.7989 | 0.79432 | -1.6682 | 3.21 |
| 667 | *NLRP6* | 4.7896 | 2.976 | 1.8651 | 1.3407 | 3.21 |
| 668 | *PLEC* | 5.9129 | 2.6943 | 1.0215 | 0.35076 | 3.21 |
| 669 | *GSTZ1* | 4.2137 | 3.6178 | 1.7925 | 1.2124 | 3.21 |
| 670 | *OR10G3* | 4.1855 | 4.1237 | 1.3107 | -1.4926 | 3.21 |
| 671 | *LY6K* | 4.1176 | 2.8174 | 2.6844 | -0.06015 | 3.21 |
| 672 | *CARF* | 4.5064 | 4.2721 | 0.84074 | -1.2987 | 3.21 |
| 673 | *CALML6* | 5.2246 | 4.7857 | -0.39318 | -0.92217 | 3.21 |
| 674 | *PPP2CA* | 3.4228 | 3.1749 | 3.0173 | 2.9739 | 3.21 |
| 675 | *CHDH* | 4.8674 | 2.8365 | 1.9104 | -1.3203 | 3.20 |
| 676 | *TMEM255A* | 5.7505 | 3.7382 | 0.12492 | -0.11186 | 3.20 |
| 677 | *SEMA3A* | 3.9311 | 3.0961 | 2.5862 | 0.23008 | 3.20 |
| 678 | *DOCK9* | 5.3199 | 3.3548 | 0.93869 | -1.4173 | 3.20 |
| 679 | *SEC62* | 4.4738 | 3.2881 | 1.8475 | -0.88834 | 3.20 |
| 680 | *UNCX* | 4.855 | 2.5722 | 2.1821 | -1.7364 | 3.20 |
| 681 | *MYH13* | 4.4669 | 3.4457 | 1.6936 | -2.0141 | 3.20 |
| 682 | *ARC* | 3.9569 | 2.9279 | 2.7181 | 0.93991 | 3.20 |
| 683 | *GDAP1* | 3.6336 | 3.3859 | 2.5831 | -3.873 | 3.20 |

| | | | | | | |
|---|---|---|---|---|---|---|
| 684 | *CENPE* | 3.7345 | 3.4783 | 2.3893 | -0.17714 | 3.20 |
| 685 | *STAG3* | 5.1496 | 4.5107 | -0.06139 | -2.4094 | 3.20 |
| 686 | *DAZ1* | 5.4041 | 4.2087 | -0.01439 | -1.9065 | 3.20 |
| 687 | *SUGP1* | 5.4876 | 4.4909 | -0.39916 | -1.5744 | 3.19 |
| 688 | *STYXL1* | 4.9725 | 3.2433 | 1.3602 | 0.28279 | 3.19 |
| 689 | *TRIM51* | 4.5877 | 3.0442 | 1.9411 | 1.3397 | 3.19 |
| 690 | *THAP2* | 5.122 | 3.8177 | 0.63277 | -0.59187 | 3.19 |
| 691 | *RNF24* | 4.8493 | 3.6209 | 1.102 | -1.2919 | 3.19 |
| 692 | *PRR16* | 5.5125 | 2.4514 | 1.5982 | -0.26128 | 3.19 |
| 693 | *INPPL1* | 4.9161 | 3.8074 | 0.83829 | -0.62694 | 3.19 |
| 694 | *UROS* | 4.2065 | 3.1162 | 2.2383 | -1.2342 | 3.19 |
| 695 | *MDGA1* | 3.9908 | 2.7857 | 2.7842 | 1.3341 | 3.19 |
| 696 | *SLAIN1* | 4.9889 | 3.0932 | 1.4766 | 1.3887 | 3.19 |
| 697 | *ARSH* | 4.5396 | 3.2989 | 1.7196 | -0.41291 | 3.19 |
| 698 | *DSCAM* | 3.3381 | 3.1528 | 3.0654 | -0.72916 | 3.19 |
| 699 | *CDC42EP2* | 4.8804 | 4.3416 | 0.33426 | 0.16186 | 3.19 |
| 700 | *HORMAD1* | 4.1582 | 3.9178 | 1.4782 | -2.1312 | 3.18 |
| 701 | *IGSF21* | 4.0191 | 2.995 | 2.5384 | 0.861 | 3.18 |
| 702 | *KIAA0226L* | 4.8219 | 4.5167 | 0.21027 | -0.4521 | 3.18 |
| 703 | *CLTB* | 4.7656 | 4.4864 | 0.29654 | 0.11372 | 3.18 |
| 704 | *DST* | 4.4192 | 4.3549 | 0.77215 | -1.7595 | 3.18 |
| 705 | *SRPK3* | 3.8639 | 2.9472 | 2.7351 | 0.3289 | 3.18 |
| 706 | *C19orf54* | 4.2217 | 3.5495 | 1.772 | 1.0009 | 3.18 |
| 707 | *CDKL3* | 5.3959 | 3.6204 | 0.52665 | -3.0687 | 3.18 |
| 708 | *SLC39A4* | 3.8926 | 3.6505 | 1.9954 | -0.83566 | 3.18 |
| 709 | *NPSR1* | 5.4579 | 3.2244 | 0.85067 | 0.56815 | 3.18 |
| 710 | *APOA5* | 4.2399 | 2.6839 | 2.609 | 1.0231 | 3.18 |
| 711 | *TMEM121* | 3.7475 | 3.2054 | 2.5793 | -1.7322 | 3.18 |
| 712 | *SPINT1* | 4.4246 | 3.5967 | 1.5101 | -0.99059 | 3.18 |
| 713 | *C16orf74* | 4.5231 | 2.9235 | 2.0815 | -0.65705 | 3.18 |
| 714 | *SMCO2* | 4.566 | 3.6434 | 1.3186 | -0.1672 | 3.18 |
| 715 | *ADRB3* | 5.7677 | 2.3855 | 1.3745 | -0.25832 | 3.18 |
| 716 | *ZMYND10* | 5.5989 | 3.9975 | -0.07032 | -0.71032 | 3.18 |
| 717 | *ODC1* | 4.2246 | 3.4522 | 1.8492 | -1.2639 | 3.18 |
| 718 | *ZBTB8B* | 4.2297 | 3.3979 | 1.8949 | 0.051568 | 3.17 |
| 719 | *MESDC2* | 3.3955 | 3.1769 | 2.9487 | -0.691 | 3.17 |
| 720 | *LEPROTL1* | 4.1258 | 4.1179 | 1.2753 | -0.2099 | 3.17 |
| 721 | *FLRT2* | 3.811 | 2.8854 | 2.8211 | 1.1673 | 3.17 |
| 722 | *ANKRD34C* | 4.8642 | 3.1609 | 1.4916 | -0.72488 | 3.17 |
| 723 | *SLX4IP* | 4.3167 | 3.1947 | 2.0038 | -0.08458 | 3.17 |
| 724 | *DOCK5* | 3.9681 | 3.3311 | 2.2156 | -1.2649 | 3.17 |
| 725 | *HS3ST1* | 4.3089 | 3.8906 | 1.3116 | -0.95184 | 3.17 |
| 726 | *RAD54L2* | 3.3659 | 3.2473 | 2.8971 | 0.10278 | 3.17 |
| 727 | *CORO1A* | 4.7969 | 3.9305 | 0.78273 | -1.3153 | 3.17 |
| 728 | *SPIN2B* | 4.1337 | 3.0296 | 2.3458 | -0.49941 | 3.17 |
| 729 | *SH3RF3* | 4.3257 | 3.0707 | 2.1121 | -1.0497 | 3.17 |

| | | | | | | |
|---|---|---|---|---|---|---|
| 730 | *PLCH1* | 4.9159 | 4.1834 | 0.40817 | -1.6604 | 3.17 |
| 731 | *SRSF5* | 4.7171 | 3.9294 | 0.86 | 0.84005 | 3.17 |
| 732 | *ASB17* | 4.6805 | 4.2805 | 0.54055 | -0.96878 | 3.17 |
| 733 | *ARF3* | 3.8628 | 3.8211 | 1.8109 | 0.69118 | 3.16 |
| 734 | *CCDC134* | 4.2594 | 2.8242 | 2.4106 | -1.059 | 3.16 |
| 735 | *CCBL2* | 4.596 | 3.7898 | 1.1075 | -0.76358 | 3.16 |
| 736 | *C1orf122* | 4.9423 | 4.001 | 0.54907 | 0.082627 | 3.16 |
| 737 | *CD109* | 5.3696 | 4.1534 | -0.03148 | -0.27796 | 3.16 |
| 738 | *ZC3H14* | 4.6849 | 3.7769 | 1.0265 | -1.632 | 3.16 |
| 739 | *AKR1C3* | 5.1286 | 3.7571 | 0.60079 | -2.5865 | 3.16 |
| 740 | *C14orf132* | 6.0317 | 2.1613 | 1.2932 | -1.8033 | 3.16 |
| 741 | *MAGEA12* | 4.7215 | 4.5903 | 0.1739 | -0.44353 | 3.16 |
| 742 | *GNAS* | 4.4487 | 3.8568 | 1.1784 | -0.30355 | 3.16 |
| 743 | *SULT1A1* | 3.687 | 3.1808 | 2.6152 | -1.0643 | 3.16 |
| 744 | *GAS8* | 4.1545 | 3.7855 | 1.5416 | -1.4339 | 3.16 |
| 745 | *RASSF4* | 4.0468 | 3.0157 | 2.4191 | 0.50787 | 3.16 |
| 746 | *GNAI2* | 3.7868 | 3.3248 | 2.3694 | 1.0419 | 3.16 |
| 747 | *MANF* | 4.945 | 4.6577 | -0.12565 | -2.1018 | 3.16 |
| 748 | *GYS2* | 5.1997 | 3.5553 | 0.72185 | 0.5297 | 3.16 |
| 749 | *NADK* | 4.9226 | 4.2538 | 0.29321 | -5.1224 | 3.16 |
| 750 | *KPNA1* | 4.89 | 4.0043 | 0.57295 | 0.030847 | 3.16 |
| 751 | *GDPD2* | 3.9589 | 3.1701 | 2.3376 | 1.5728 | 3.16 |
| 752 | *OTULIN* | 4.6613 | 4.6399 | 0.16234 | -1.3694 | 3.15 |
| 753 | *PECR* | 4.1674 | 2.9127 | 2.3797 | 1.0417 | 3.15 |
| 754 | *CLCA4* | 5.4851 | 3.0381 | 0.93577 | -0.17318 | 3.15 |
| 755 | *FGGY* | 4.6971 | 2.9377 | 1.8147 | -2.6157 | 3.15 |
| 756 | *SLC26A5* | 4.6247 | 4.098 | 0.72264 | -1.571 | 3.15 |
| 757 | *GPR61* | 5.8813 | 4.6447 | -1.0826 | -1.9581 | 3.15 |
| 758 | *PHLDA1* | 4.1637 | 2.6637 | 2.6069 | 1.9934 | 3.14 |
| 759 | *PLOD3* | 3.5759 | 3.0336 | 2.8227 | -0.32894 | 3.14 |
| 760 | *ULK4* | 5.0465 | 3.354 | 1.0299 | -1.0858 | 3.14 |
| 761 | *PPIAL4C* | 4.027 | 3.1957 | 2.2045 | -0.20924 | 3.14 |
| 762 | *CSNK1G2* | 4.0106 | 2.8199 | 2.5961 | -2.2167 | 3.14 |
| 763 | *TRIM23* | 4.7651 | 4.5012 | 0.15492 | -0.56382 | 3.14 |
| 764 | *CHPF2* | 4.0655 | 3.1018 | 2.2536 | 0.053863 | 3.14 |
| 765 | *SURF2* | 5.9923 | 1.9177 | 1.5097 | -1.6876 | 3.14 |
| 766 | *ORMDL1* | 4.5025 | 3.2059 | 1.7108 | -1.153 | 3.14 |
| 767 | *MX2* | 5.3561 | 4.8824 | -0.82 | -2.071 | 3.14 |
| 768 | *KCNJ8* | 4.6885 | 2.572 | 2.1551 | 1.2948 | 3.14 |
| 769 | *PRSS27* | 4.8397 | 4.0038 | 0.56964 | 0.13197 | 3.14 |
| 770 | *PRM3* | 3.9554 | 3.3783 | 2.0787 | 0.74505 | 3.14 |
| 771 | *WNT16* | 3.6667 | 3.5251 | 2.2176 | 0.78047 | 3.14 |
| 772 | *CCDC8* | 5.7511 | 4.3688 | -0.71051 | -1.4183 | 3.14 |
| 773 | *ANXA2* | 5.3569 | 4.7231 | -0.67413 | -0.92046 | 3.14 |
| 774 | *ZNF77* | 3.4673 | 3.2653 | 2.6682 | 1.2514 | 3.13 |
| 775 | *MBOAT1* | 4.2865 | 3.8229 | 1.2892 | -1.0145 | 3.13 |

| | | | | | | |
|---|---|---|---|---|---|---|
| 776 | *MCCC2* | 3.8336 | 3.5617 | 2.0026 | -1.0757 | 3.13 |
| 777 | *MICA* | 5.4906 | 4.2794 | -0.37461 | -1.525 | 3.13 |
| 778 | *KITLG* | 4.5731 | 2.8467 | 1.9716 | -0.10888 | 3.13 |
| 779 | *RBM11* | 4.5401 | 2.5854 | 2.2657 | 0.45315 | 3.13 |
| 780 | *ALG6* | 3.786 | 3.6644 | 1.9373 | -0.75053 | 3.13 |
| 781 | *DCUN1D2* | 3.2927 | 3.2304 | 2.8595 | -0.85686 | 3.13 |
| 782 | *C1QTNF9* | 4.9773 | 2.8626 | 1.5399 | -1.7526 | 3.13 |
| 783 | *QRFPR* | 4.2149 | 2.7203 | 2.4441 | 1.1126 | 3.13 |
| 784 | *LAMTOR3* | 5.5723 | 2.0439 | 1.7624 | -1.5082 | 3.13 |
| 785 | *KCNJ11* | 3.9539 | 3.5222 | 1.9019 | -0.53417 | 3.13 |
| 786 | *TDRD12* | 7.0726 | 1.7051 | 0.59944 | -1.927 | 3.13 |
| 787 | *WWC2* | 4.6166 | 3.0309 | 1.7295 | -0.80771 | 3.13 |
| 788 | *HPS6* | 4.04 | 2.7584 | 2.5786 | -0.70417 | 3.13 |
| 789 | *FGFR1OP2* | 5.1578 | 2.8795 | 1.3385 | -1.2666 | 3.13 |
| 790 | *HIBADH* | 4.3975 | 3.3702 | 1.6056 | -1.5579 | 3.12 |
| 791 | *NTMT1* | 3.9878 | 3.9646 | 1.4205 | -0.57896 | 3.12 |
| 792 | *SHBG* | 5.0877 | 3.2797 | 1.0044 | 0.096888 | 3.12 |
| 793 | *LPAR4* | 3.9107 | 3.8757 | 1.5823 | -2.1162 | 3.12 |
| 794 | *FUK* | 4.6196 | 4.1209 | 0.62701 | 0.049057 | 3.12 |
| 795 | *ZNF606* | 3.9094 | 3.4892 | 1.9627 | -0.89699 | 3.12 |
| 796 | *GRK6* | 4.9573 | 3.2532 | 1.1487 | -0.14543 | 3.12 |
| 797 | *OR8D2* | 4.2815 | 4.2185 | 0.85826 | -1.5176 | 3.12 |
| 798 | *CCDC66* | 4.5097 | 4.2617 | 0.58665 | -1.2328 | 3.12 |
| 799 | *ACOT12* | 3.9819 | 3.6498 | 1.725 | -2.5647 | 3.12 |
| 800 | *GOLGA1* | 5.344 | 3.2279 | 0.78169 | -0.16122 | 3.12 |
| 801 | *TSPAN8* | 4.9528 | 4.0061 | 0.39457 | -0.50246 | 3.12 |
| 802 | *MED1* | 4.1259 | 3.4569 | 1.7643 | -1.38 | 3.12 |
| 803 | *ZNF37A* | 5.6403 | 2.2003 | 1.5061 | -0.42014 | 3.12 |
| 804 | *XRRA1* | 4.5446 | 3.3091 | 1.4924 | -0.20142 | 3.12 |
| 805 | *LMX1B* | 5.3732 | 3.1546 | 0.81339 | 0.12611 | 3.11 |
| 806 | *NYX* | 3.9999 | 3.944 | 1.3915 | -0.12742 | 3.11 |
| 807 | *IFFO1* | 5.0933 | 4.3653 | -0.12428 | -0.15596 | 3.11 |
| 808 | *SLIRP* | 4.5968 | 3.8252 | 0.90715 | -1.507 | 3.11 |
| 809 | *KSR2* | 4.7439 | 4.1135 | 0.47073 | -2.8044 | 3.11 |
| 810 | *NPRL2* | 5.0855 | 4.8957 | -0.6548 | -1.8402 | 3.11 |
| 811 | *CUX1* | 4.4082 | 2.9331 | 1.984 | 1.2744 | 3.11 |
| 812 | *S100A1* | 4.4666 | 4.0112 | 0.84658 | -0.27982 | 3.11 |
| 813 | *ZNF700* | 4.1924 | 2.692 | 2.4353 | -1.009 | 3.11 |
| 814 | *AASS* | 4.2526 | 3.3259 | 1.7402 | 1.7 | 3.11 |
| 815 | *LURAP1L* | 4.7229 | 3.4036 | 1.1897 | -0.71132 | 3.11 |
| 816 | *CDR1* | 4.619 | 3.4937 | 1.2025 | -0.43264 | 3.11 |
| 817 | *PSD* | 3.8632 | 2.9874 | 2.462 | -0.17025 | 3.10 |
| 818 | *DAPP1* | 3.6831 | 3.2667 | 2.3626 | -1.2027 | 3.10 |
| 819 | *MEGF6* | 3.7689 | 2.8383 | 2.7051 | -0.74086 | 3.10 |
| 820 | *NRG1* | 3.6967 | 3.3823 | 2.2315 | -1.729 | 3.10 |
| 821 | *GPX3* | 4.5253 | 4.2732 | 0.50891 | -1.5538 | 3.10 |

| | | | | | | |
|---|---|---|---|---|---|---|
| 822 | *KRTAP10-8* | 4.0098 | 2.6691 | 2.6275 | 0.44775 | 3.10 |
| 823 | *C2orf40* | 4.3887 | 3.2474 | 1.6682 | 1.4073 | 3.10 |
| 824 | *SCGN* | 4.3981 | 4.3788 | 0.51957 | 0.39116 | 3.10 |
| 825 | *PPAPDC1B* | 3.6444 | 3.392 | 2.2598 | 0.31761 | 3.10 |
| 826 | *MYADML2* | 4.4047 | 2.9479 | 1.9428 | -2.2951 | 3.10 |
| 827 | *LAMP1* | 3.5752 | 3.462 | 2.2581 | 1.2512 | 3.10 |
| 828 | *POLK* | 4.8341 | 4.2939 | 0.16383 | -0.31475 | 3.10 |
| 829 | *ZNF449* | 4.2956 | 4.2263 | 0.76881 | 0.29782 | 3.10 |
| 830 | *OTUD4* | 5.0264 | 4.5138 | -0.25004 | -0.93057 | 3.10 |
| 831 | *KPNA3* | 4.9721 | 4.001 | 0.31294 | -1.0313 | 3.10 |
| 832 | *OR2T2* | 4.2355 | 4.1781 | 0.87135 | -0.58404 | 3.09 |
| 833 | *AGTR2* | 3.6967 | 3.2966 | 2.2909 | 0.28143 | 3.09 |
| 834 | *FYB* | 5.8263 | 3.3842 | 0.07224 | -0.07903 | 3.09 |
| 835 | *CAPRIN1* | 5.3791 | 4.1644 | -0.26203 | -0.84419 | 3.09 |
| 836 | *GALNS* | 5.9109 | 2.2126 | 1.1576 | 0.009703 | 3.09 |
| 837 | *VCY1B* | 4.4537 | 2.5061 | 2.3205 | -2.3949 | 3.09 |
| 838 | *FAM57B* | 3.6831 | 3.5645 | 2.0311 | -2.5743 | 3.09 |
| 839 | *ZNRF1* | 5.1603 | 3.1957 | 0.91407 | 0.4966 | 3.09 |
| 840 | *ZNF732* | 4.8359 | 2.9929 | 1.4387 | -0.41762 | 3.09 |
| 841 | *TEX37* | 4.6944 | 2.4207 | 2.152 | -0.34819 | 3.09 |
| 842 | *NCAPD2* | 4.689 | 4.0369 | 0.54017 | -0.99387 | 3.09 |
| 843 | *FAM179A* | 4.3498 | 3.1088 | 1.8063 | -1.3393 | 3.09 |
| 844 | *PLA2G2A* | 4.848 | 3.4959 | 0.91085 | -1.8253 | 3.08 |
| 845 | *GNA13* | 3.8521 | 2.7663 | 2.6302 | 1.2351 | 3.08 |
| 846 | *HCAR2* | 4.0118 | 3.3485 | 1.8869 | 0.080928 | 3.08 |
| 847 | *MAGEA10-MAGEA[illegible]* | 4.995 | 2.9263 | 1.3235 | -0.26207 | 3.08 |
| 848 | *HIF1AN* | 4.3314 | 3.1314 | 1.7773 | 0.12471 | 3.08 |
| 849 | *VHLL* | 4.2258 | 3.6724 | 1.341 | -0.44884 | 3.08 |
| 850 | *EHD2* | 4.3164 | 4.0063 | 0.91429 | -1.6288 | 3.08 |
| 851 | *ZCCHC5* | 4.8836 | 3.8485 | 0.50078 | -0.37215 | 3.08 |
| 852 | *ANKHD1* | 3.8327 | 3.1364 | 2.2626 | -1.7945 | 3.08 |
| 853 | *C1orf100* | 4.705 | 2.8323 | 1.6934 | 0.14603 | 3.08 |
| 854 | *EEFSEC* | 3.185 | 3.0529 | 2.9906 | -0.78793 | 3.08 |
| 855 | *KCTD6* | 4.6776 | 3.442 | 1.1081 | -0.61119 | 3.08 |
| 856 | *CLLU1OS* | 5.4441 | 3.2766 | 0.50492 | 0.38838 | 3.08 |
| 857 | *DYRK4* | 4.3435 | 2.8997 | 1.9811 | -0.54663 | 3.07 |
| 858 | *ESAM* | 3.5847 | 3.1187 | 2.5154 | 1.5361 | 3.07 |
| 859 | *FANK1* | 4.4184 | 3.9859 | 0.81204 | 0.62542 | 3.07 |
| 860 | *CCDC6* | 4.7996 | 2.3583 | 2.0579 | 0.020029 | 3.07 |
| 861 | *SH2D3C* | 4.0568 | 4.0416 | 1.1162 | 0.049826 | 3.07 |
| 862 | *FAM83C* | 4.9449 | 4.4549 | -0.18898 | -1.8851 | 3.07 |
| 863 | *IL7* | 3.4972 | 2.8899 | 2.8222 | 0.01556 | 3.07 |
| 864 | *C11orf1* | 4.955 | 2.5151 | 1.7389 | -0.32446 | 3.07 |
| 865 | *LCNL1* | 3.6162 | 2.8641 | 2.7278 | 0.85583 | 3.07 |
| 866 | *TGM6* | 3.3126 | 3.1493 | 2.7462 | 1.2105 | 3.07 |
| 867 | *AVPI1* | 4.222 | 3.4708 | 1.5141 | 0.17324 | 3.07 |

| | | | | | | |
|---|---|---|---|---|---|---|
| 868 | *KLRF1* | 3.6468 | 2.904 | 2.6556 | -1.547 | 3.07 |
| 869 | *RNASE1* | 3.8501 | 2.7177 | 2.6381 | 1.6948 | 3.07 |
| 870 | *ENG* | 4.6833 | 4.0906 | 0.42787 | -0.51752 | 3.07 |
| 871 | *CDSN* | 5.1466 | 2.2781 | 1.7768 | -1.5971 | 3.07 |
| 872 | *B3GNT2* | 4.9536 | 3.4098 | 0.83536 | -1.0095 | 3.07 |
| 873 | *ZNF844* | 3.7339 | 3.0372 | 2.4245 | 0.36126 | 3.07 |
| 874 | *NEFM* | 4.8244 | 2.789 | 1.5812 | -1.9326 | 3.06 |
| 875 | *OR4F3* | 4.3592 | 3.7779 | 1.0546 | -0.82053 | 3.06 |
| 876 | *BTNL8* | 3.8878 | 2.8824 | 2.4203 | -0.03648 | 3.06 |
| 877 | *FAM167B* | 4.3915 | 3.7452 | 1.0528 | -0.37495 | 3.06 |
| 878 | *KCNE3* | 4.0719 | 2.8149 | 2.3017 | 0.84712 | 3.06 |
| 879 | *AOAH* | 4.6698 | 3.4097 | 1.1086 | -0.95565 | 3.06 |
| 880 | *DYNC2H1* | 4.376 | 2.7917 | 2.0159 | -1.1733 | 3.06 |
| 881 | *NUP37* | 4.5294 | 4.0333 | 0.62018 | -0.23116 | 3.06 |
| 882 | *SLC4A8* | 3.7551 | 3.2074 | 2.2197 | -0.69286 | 3.06 |
| 883 | *KRTAP19-4* | 4.7089 | 2.5419 | 1.9303 | -0.05 | 3.06 |
| 884 | *PHPT1* | 3.4469 | 3.3965 | 2.3376 | -1.3407 | 3.06 |
| 885 | *OAS3* | 3.9044 | 3.5192 | 1.7539 | -2.3294 | 3.06 |
| 886 | *LOC100130370* | 4.9562 | 4.048 | 0.1727 | -0.4264 | 3.06 |
| 887 | *HERC4* | 4.0947 | 3.8166 | 1.2597 | 0.30815 | 3.06 |
| 888 | *PSEN2* | 4.9159 | 4.644 | -0.38945 | -1.3696 | 3.06 |
| 889 | *DDC* | 3.9913 | 3.135 | 2.0435 | -1.3208 | 3.06 |
| 890 | *SLC24A2* | 5.5315 | 4.4241 | -0.78588 | -1.5121 | 3.06 |
| 891 | *IPMK* | 4.8758 | 4.3591 | -0.0665 | -0.36196 | 3.06 |
| 892 | *DIAPH3* | 3.8697 | 3.4148 | 1.8832 | 0.14641 | 3.06 |
| 893 | *ARHGEF1* | 3.9446 | 3.1118 | 2.111 | -2.6257 | 3.06 |
| 894 | *UHMK1* | 4.5716 | 4.1558 | 0.43849 | -0.55047 | 3.06 |
| 895 | *SYCE1* | 3.5242 | 3.4561 | 2.1847 | 0.81769 | 3.06 |
| 896 | *C14orf180* | 4.3192 | 4.1375 | 0.70551 | -0.05804 | 3.05 |
| 897 | *SH3BGR* | 4.6024 | 3.4848 | 1.0747 | 0.43719 | 3.05 |
| 898 | *MYOF* | 3.8458 | 3.4991 | 1.8138 | 1.2541 | 3.05 |
| 899 | *C16orf87* | 3.6759 | 3.0023 | 2.479 | -1.6617 | 3.05 |
| 900 | *SP7* | 4.4457 | 2.7893 | 1.9214 | 1.7623 | 3.05 |
| 901 | *METRN* | 4.9691 | 3.781 | 0.40322 | -1.5752 | 3.05 |
| 902 | *RERGL* | 5.4132 | 4.1536 | -0.41469 | -2.0476 | 3.05 |
| 903 | *ZNF548* | 4.8858 | 3.3679 | 0.89741 | -1.0564 | 3.05 |
| 904 | *TMEM176B* | 5.2063 | 3.4025 | 0.54228 | -0.93764 | 3.05 |
| 905 | *VWF* | 4.069 | 2.7211 | 2.3598 | 0.72362 | 3.05 |
| 906 | *TULP4* | 3.7709 | 3.1226 | 2.2542 | 1.5469 | 3.05 |
| 907 | *SLC15A4* | 5.5275 | 3.3999 | 0.2169 | -0.82959 | 3.05 |
| 908 | *HIST1H1D* | 4.4895 | 4.0775 | 0.57685 | -1.7549 | 3.05 |
| 909 | *ROCK1* | 4.8016 | 4.2547 | 0.08598 | -2.1217 | 3.05 |
| 910 | *CYP7A1* | 3.9875 | 3.7627 | 1.3875 | -0.46384 | 3.05 |
| 911 | *WDR31* | 3.7043 | 2.8285 | 2.6047 | -1.7061 | 3.05 |
| 912 | *ACER2* | 4.5383 | 3.1615 | 1.4344 | 0.40931 | 3.04 |
| 913 | *PTDSS2* | 5.1545 | 2.8626 | 1.1098 | -0.33174 | 3.04 |

| | | | | | | |
|---|---|---|---|---|---|---|
| 914 | *ASMTL* | 4.9635 | 3.7366 | 0.42619 | -1.8583 | 3.04 |
| 915 | *ZNF316* | 4.4984 | 3.9545 | 0.6695 | -0.10928 | 3.04 |
| 916 | *PEX19* | 3.8114 | 3.3287 | 1.9822 | -1.2955 | 3.04 |
| 917 | *CCDC22* | 3.8468 | 3.1426 | 2.1244 | -1.8816 | 3.04 |
| 918 | *ITFG3* | 3.9603 | 3.7904 | 1.3566 | -1.0117 | 3.04 |
| 919 | *ADCY2* | 4.4018 | 3.5456 | 1.1597 | -0.09235 | 3.04 |
| 920 | *PON3* | 4.8316 | 3.6027 | 0.67277 | -0.81205 | 3.04 |
| 921 | *RAP2C* | 5.383 | 4.127 | -0.40319 | -1.8123 | 3.04 |
| 922 | *KLF1* | 4.297 | 3.0909 | 1.7167 | 0.26476 | 3.03 |
| 923 | *PLIN1* | 4.4303 | 2.4241 | 2.246 | 0.73119 | 3.03 |
| 924 | *OR4N5* | 3.6283 | 3.2866 | 2.1835 | -1.8076 | 3.03 |
| 925 | *ARHGAP19* | 4.8579 | 3.9939 | 0.24633 | -1.2867 | 3.03 |
| 926 | *ZSCAN32* | 4.7328 | 4.3655 | -0.00066 | -3.1229 | 3.03 |
| 927 | *DUS3L* | 4.2455 | 2.5617 | 2.288 | 1.25 | 3.03 |
| 928 | *PPP1R3A* | 4.1418 | 4.0174 | 0.93548 | -2.3849 | 3.03 |
| 929 | *FAM132A* | 3.9052 | 2.7243 | 2.4647 | -0.11906 | 3.03 |
| 930 | *TXNDC15* | 3.6207 | 2.7616 | 2.7097 | -1.1247 | 3.03 |
| 931 | *HNRNPUL1* | 3.4594 | 3.1746 | 2.4546 | -0.51259 | 3.03 |
| 932 | *IL22* | 4.4264 | 3.1653 | 1.4952 | -0.39856 | 3.03 |
| 933 | *CALY* | 5.1745 | 3.3725 | 0.53888 | -1.0221 | 3.03 |
| 934 | *TRIM37* | 3.4001 | 2.8829 | 2.8011 | -0.06671 | 3.03 |
| 935 | *AANAT* | 3.7291 | 3.3899 | 1.9629 | -1.1313 | 3.03 |
| 936 | *OR4A5* | 3.4268 | 2.8769 | 2.7747 | -0.15733 | 3.03 |
| 937 | *NFIC* | 4.5909 | 3.3292 | 1.1567 | -1.1338 | 3.03 |
| 938 | *MRPS9* | 4.0872 | 2.6565 | 2.3321 | 0.4688 | 3.03 |
| 939 | *RNF152* | 4.4977 | 2.4912 | 2.0866 | 1.2117 | 3.03 |
| 940 | *PIWIL1* | 4.2867 | 2.7237 | 2.0611 | -1.3713 | 3.02 |
| 941 | *TEX11* | 3.7475 | 3.1943 | 2.1254 | -1.0063 | 3.02 |
| 942 | *SFRP4* | 5.2362 | 3.7208 | 0.10924 | -1.2234 | 3.02 |
| 943 | *RBMY1B* | 4.2014 | 3.7717 | 1.0887 | -0.61092 | 3.02 |
| 944 | *PALLD* | 5.7063 | 3.4285 | -0.07335 | -3.0444 | 3.02 |
| 945 | *TAPBP* | 4.3817 | 3.4804 | 1.1989 | 0.001973 | 3.02 |
| 946 | *THAP11* | 4.9398 | 2.5991 | 1.522 | 0.56658 | 3.02 |
| 947 | *EBF3* | 5.2987 | 2.0728 | 1.683 | 1.363 | 3.02 |
| 948 | *MMP15* | 4.0063 | 2.5653 | 2.4801 | -1.138 | 3.02 |
| 949 | *PSG8* | 5.0845 | 3.1545 | 0.81141 | -0.86821 | 3.02 |
| 950 | *LOC100505549* | 3.3143 | 3.0985 | 2.6346 | 1.2898 | 3.02 |
| 951 | *GSTA1* | 4.8385 | 4.1545 | 0.05428 | -0.198 | 3.02 |
| 952 | *AK4* | 4.3883 | 4.1481 | 0.50936 | -0.42675 | 3.02 |
| 953 | *NBPF9* | 4.7507 | 4.0336 | 0.25952 | -1.449 | 3.01 |
| 954 | *SDR42E1* | 3.3902 | 3.1335 | 2.518 | 1.4349 | 3.01 |
| 955 | *PITPNM3* | 3.8883 | 3.3315 | 1.8194 | -1.3781 | 3.01 |
| 956 | *OTUD5* | 3.1772 | 3.1619 | 2.6961 | -1.0089 | 3.01 |
| 957 | *EBF1* | 4.1578 | 3.5487 | 1.3277 | 0.16456 | 3.01 |
| 958 | *PRR5-ARHGAP8* | 3.6877 | 2.7245 | 2.6202 | -0.72969 | 3.01 |
| 959 | *FAM91A1* | 4.3102 | 4.2891 | 0.43054 | 0.31699 | 3.01 |

| | | | | | | |
|---|---|---|---|---|---|---|
| 960 | *GPR84* | 5.6612 | 3.0098 | 0.35707 | -0.58736 | 3.01 |
| 961 | *TBX10* | 5.8211 | 3.9839 | -0.77895 | -1.3523 | 3.01 |
| 962 | *ZNF200* | 5.3335 | 2.2676 | 1.4244 | 0.038662 | 3.01 |
| 963 | *GPR143* | 4.0785 | 2.8732 | 2.0726 | -1.0444 | 3.01 |
| 964 | *WDFY3* | 4.2598 | 2.7475 | 2.0168 | -2.3496 | 3.01 |
| 965 | *GPNMB* | 4.8169 | 2.4621 | 1.745 | -1.0528 | 3.01 |
| 966 | *VCL* | 4.5982 | 2.4265 | 1.9993 | 0.50522 | 3.01 |
| 967 | *CLEC17A* | 4.4605 | 4.2392 | 0.32334 | 0.19285 | 3.01 |
| 968 | *OR51L1* | 5.1934 | 3.5184 | 0.31065 | -1.3098 | 3.01 |
| 969 | *EMID1* | 5.0621 | 2.559 | 1.4013 | 0.69052 | 3.01 |
| 970 | *BHLHE40* | 3.7367 | 3.5311 | 1.7514 | 0.55151 | 3.01 |
| 971 | *EEPD1* | 4.0296 | 2.5007 | 2.4885 | 0.6291 | 3.01 |
| 972 | *LYNX1* | 4.4295 | 2.463 | 2.1259 | -0.36148 | 3.01 |
| 973 | *METTL21B* | 4.2331 | 3.3939 | 1.3892 | 0.76725 | 3.01 |
| 974 | *PHKB* | 4.5709 | 2.2821 | 2.1623 | -0.714 | 3.01 |
| 975 | *TIGD3* | 4.985 | 3.8919 | 0.13796 | -0.09541 | 3.00 |
| 976 | *GLP1R* | 4.3962 | 4.3133 | 0.30472 | -1.707 | 3.00 |
| 977 | *CEMIP* | 5.2751 | 2.8104 | 0.92641 | -0.61164 | 3.00 |
| 978 | *FLNC* | 3.8173 | 3.1064 | 2.0855 | 1.0684 | 3.00 |
| 979 | *F13A1* | 4.53 | 3.1447 | 1.3337 | -0.47183 | 3.00 |
| 980 | *TNRC18* | 4.3078 | 2.7608 | 1.9391 | 0.29241 | 3.00 |
| 981 | *VLDLR* | 5.5502 | 1.8964 | 1.5589 | -1.7652 | 3.00 |
| 982 | *OSR2* | 4.6328 | 4.159 | 0.21256 | -1.4935 | 3.00 |
| 983 | *OR14C36* | 4.0925 | 4.0796 | 0.82971 | -1.3035 | 3.00 |
| 984 | *GSK3A* | 4.6661 | 4.6035 | -0.26854 | -1.6663 | 3.00 |
| 985 | *CEP85L* | 3.3404 | 2.9581 | 2.6994 | -1.3457 | 3.00 |
| 986 | *TIMP4* | 4.1422 | 3.9017 | 0.95279 | 0.74742 | 3.00 |
| 987 | *TVP23B* | 3.0889 | 3.0422 | 2.8633 | -0.80771 | 3.00 |
| 988 | *FADS3* | 4.308 | 3.4152 | 1.2674 | 0.37779 | 3.00 |
| 989 | *DKK2* | 4.5766 | 2.496 | 1.912 | -1.9821 | 2.99 |
| 990 | *DHX29* | 4.3134 | 2.9681 | 1.7008 | -1.8946 | 2.99 |
| 991 | *ARRDC5* | 4.2483 | 4.2074 | 0.52335 | -1.1573 | 2.99 |
| 992 | *CCDC172* | 3.9622 | 3.8347 | 1.1817 | -0.57824 | 2.99 |
| 993 | *CDC25A* | 4.7498 | 4.6054 | -0.37898 | -1.1421 | 2.99 |
| 994 | *DYDC2* | 3.6318 | 3.1148 | 2.2244 | 0.28506 | 2.99 |
| 995 | *IFT80* | 3.9593 | 2.9829 | 2.0277 | -0.25892 | 2.99 |
| 996 | *CD19* | 4.9432 | 4.5032 | -0.47777 | -1.0626 | 2.99 |
| 997 | *C12orf50* | 4.0539 | 3.751 | 1.1637 | -0.55779 | 2.99 |
| 998 | *ART1* | 5.4914 | 4.5256 | -1.0564 | -1.3358 | 2.99 |
| 999 | *ASZ1* | 4.5816 | 2.6415 | 1.736 | -0.49989 | 2.99 |
| 1000 | *CD47* | 4.0529 | 3.6732 | 1.2327 | -1.1927 | 2.99 |
| 1001 | *LIN7A* | 5.3074 | 4.0794 | -0.43077 | -1.7493 | 2.99 |
| 1002 | *ZNF292* | 3.7714 | 3.3871 | 1.7966 | -0.47023 | 2.99 |
| 1003 | *PRORY* | 4.6632 | 3.4573 | 0.83398 | -0.5395 | 2.98 |
| 1004 | *ZSWIM4* | 5.0844 | 4.2459 | -0.3791 | -1.0324 | 2.98 |
| 1005 | *CDKN1A* | 3.9051 | 2.5486 | 2.4952 | -2.6072 | 2.98 |

| | | | | | | |
|---|---|---|---|---|---|---|
| 1006 | *PRAF2* | 4.204 | 3.2615 | 1.4825 | 0.33089 | 2.98 |
| 1007 | *RPH3AL* | 3.5064 | 3.0244 | 2.414 | -1.8889 | 2.98 |
| 1008 | *FAM98A* | 3.9243 | 3.908 | 1.1098 | -0.12837 | 2.98 |
| 1009 | *NR2F6* | 3.621 | 3.3008 | 2.0187 | -0.10911 | 2.98 |
| 1010 | *OR4A16* | 4.7041 | 3.3679 | 0.86615 | 0.70665 | 2.98 |
| 1011 | *RWDD2B* | 3.3402 | 3.008 | 2.5897 | -0.32495 | 2.98 |
| 1012 | *SERPINB2* | 4.0723 | 2.6513 | 2.2134 | -1.6526 | 2.98 |
| 1013 | *TC2N* | 4.4602 | 4.2863 | 0.19018 | -3.2582 | 2.98 |
| 1014 | *GMPR* | 3.768 | 2.8664 | 2.3017 | 0.06209 | 2.98 |
| 1015 | *YWHAG* | 3.4364 | 2.7709 | 2.7287 | -0.8903 | 2.98 |
| 1016 | *PPP1R12C* | 5.1792 | 3.1227 | 0.63103 | -0.54501 | 2.98 |
| 1017 | *MAN2B1* | 4.5562 | 2.5958 | 1.7805 | 0.69304 | 2.98 |
| 1018 | *GCSAM* | 4.0758 | 3.1622 | 1.6943 | -0.43203 | 2.98 |
| 1019 | *RPL3L* | 4.3553 | 3.4934 | 1.0821 | -2.0164 | 2.98 |
| 1020 | *GALNT1* | 3.2604 | 2.862 | 2.8081 | 0.5143 | 2.98 |
| 1021 | *BTN3A1* | 3.3194 | 3.2457 | 2.364 | 0.7829 | 2.98 |
| 1022 | *ZNF627* | 3.8564 | 2.9785 | 2.0914 | 1.3754 | 2.98 |
| 1023 | *USP17L1* | 4.2616 | 4.2601 | 0.4045 | -1.4241 | 2.98 |
| 1024 | *RASGRF2* | 5.4734 | 3.4232 | 0.02935 | -1.4376 | 2.98 |
| 1025 | *LEUTX* | 3.8871 | 3.1416 | 1.8945 | -0.46749 | 2.97 |
| 1026 | *SVIP* | 3.9851 | 2.7407 | 2.1942 | -1.8342 | 2.97 |
| 1027 | *ZNF418* | 4.016 | 3.9994 | 0.90157 | 0.32028 | 2.97 |
| 1028 | *TIPARP* | 3.427 | 3.226 | 2.2635 | -0.57921 | 2.97 |
| 1029 | *METTL6* | 4.3596 | 4.3406 | 0.21608 | 0.10942 | 2.97 |
| 1030 | *FBLN1* | 4.5847 | 4.1543 | 0.17591 | -0.92581 | 2.97 |
| 1031 | *UNC80* | 4.7912 | 3.348 | 0.77541 | -0.01594 | 2.97 |
| 1032 | *CLCN5* | 4.7276 | 2.9645 | 1.2223 | 0.57821 | 2.97 |
| 1033 | *OR6Q1* | 3.6787 | 2.9792 | 2.2553 | -0.35089 | 2.97 |
| 1034 | *FIS1* | 5.2375 | 2.8052 | 0.8702 | -0.96072 | 2.97 |
| 1035 | *BCKDHA* | 5.2346 | 3.6763 | 0.00116 | -0.3859 | 2.97 |
| 1036 | *KIAA1324L* | 5.2128 | 3.723 | -0.02429 | -1.8282 | 2.97 |
| 1037 | *EPS15L1* | 4.764 | 2.7912 | 1.3561 | 1.1738 | 2.97 |
| 1038 | *ZXDA* | 5.1578 | 3.9317 | -0.18153 | -0.2728 | 2.97 |
| 1039 | *KLF10* | 4.0168 | 3.9075 | 0.98354 | 0.38526 | 2.97 |
| 1040 | *OR1C1* | 4.7927 | 2.5767 | 1.537 | 1.4043 | 2.97 |
| 1041 | *BRPF1* | 3.6324 | 2.9033 | 2.3705 | 0.13294 | 2.97 |
| 1042 | *ITM2A* | 4.4695 | 3.5613 | 0.87508 | -0.83101 | 2.97 |
| 1043 | *VIL1* | 4.7078 | 4.2752 | -0.07731 | -0.99033 | 2.97 |
| 1044 | *EPHA3* | 5.0056 | 3.2165 | 0.68229 | -1.677 | 2.97 |
| 1045 | *MAST3* | 4.806 | 2.3118 | 1.7853 | -1.894 | 2.97 |
| 1046 | *KCNAB2* | 4.518 | 3.5759 | 0.80876 | 0.43849 | 2.97 |
| 1047 | *RANBP3L* | 5.048 | 2.0945 | 1.7598 | 0.50421 | 2.97 |
| 1048 | *LPAR3* | 3.5447 | 3.1093 | 2.248 | -0.86229 | 2.97 |
| 1049 | *ARAP1* | 5.2021 | 4.7045 | -1.0093 | -1.3282 | 2.97 |
| 1050 | *MRPL24* | 4.2721 | 2.561 | 2.0628 | -1.4375 | 2.97 |
| 1051 | *ZSWIM7* | 4.96 | 3.1804 | 0.74827 | 0.19567 | 2.96 |

| | | | | | | |
|---|---|---|---|---|---|---|
| 1052 | *RAP2B* | 4.8822 | 3.7376 | 0.26815 | -0.76593 | 2.96 |
| 1053 | *SOCS1* | 4.9904 | 2.9019 | 0.99547 | -1.6524 | 2.96 |
| 1054 | *ZNF536* | 4.8972 | 4.606 | -0.61687 | -1.0185 | 2.96 |
| 1055 | *LRPAP1* | 4.4375 | 3.1643 | 1.2806 | -0.86403 | 2.96 |
| 1056 | *PCDHA12* | 4.0796 | 3.6654 | 1.1357 | -0.53657 | 2.96 |
| 1057 | *DEGS1* | 4.2238 | 3.9985 | 0.65767 | -1.1927 | 2.96 |
| 1058 | *RGS12* | 4.8019 | 4.4354 | -0.35924 | -1.2329 | 2.96 |
| 1059 | *TCP11* | 4.5644 | 4.1437 | 0.16896 | -0.05159 | 2.96 |
| 1060 | *DNAJB9* | 4.6955 | 3.4227 | 0.75872 | -1.3617 | 2.96 |
| 1061 | *FAM53C* | 5.3849 | 3.0131 | 0.47848 | -0.03775 | 2.96 |
| 1062 | *C1orf95* | 4.9006 | 4.0868 | -0.11666 | -0.44353 | 2.96 |
| 1063 | *ZNF774* | 4.5268 | 3.9719 | 0.37026 | -0.73362 | 2.96 |
| 1064 | *CRTC3* | 3.3617 | 3.0305 | 2.4761 | 1.0958 | 2.96 |
| 1065 | *PCIF1* | 4.7788 | 3.0881 | 1.0013 | -0.09893 | 2.96 |
| 1066 | *HIST1H3F* | 4.8404 | 3.5337 | 0.49389 | -2.0476 | 2.96 |
| 1067 | *HMCN2* | 3.7218 | 2.8116 | 2.333 | 1.1733 | 2.96 |
| 1068 | *GZMH* | 3.6475 | 3.3091 | 1.9093 | 1.2681 | 2.96 |
| 1069 | *GAA* | 5.068 | 3.0119 | 0.77904 | -1.2576 | 2.95 |
| 1070 | *ZDHHC15* | 3.2882 | 3.1886 | 2.3819 | -0.63139 | 2.95 |
| 1071 | *ZNF705A* | 4.5096 | 2.494 | 1.8533 | -0.23286 | 2.95 |
| 1072 | *REC114* | 4.2929 | 3.7384 | 0.82264 | 0.5635 | 2.95 |
| 1073 | *KRT72* | 3.8711 | 3.0127 | 1.9694 | 0.64312 | 2.95 |
| 1074 | *ADAD1* | 4.5265 | 3.3665 | 0.95926 | -0.5468 | 2.95 |
| 1075 | *SETMAR* | 4.1541 | 2.3563 | 2.3418 | -0.46205 | 2.95 |
| 1076 | *TDRD1* | 4.4534 | 4.1634 | 0.23329 | -0.20142 | 2.95 |
| 1077 | *MCC* | 3.5508 | 3.5431 | 1.7561 | 0.82785 | 2.95 |
| 1078 | *HAND2* | 4.3961 | 3.7181 | 0.73398 | -0.76405 | 2.95 |
| 1079 | *DHCR7* | 3.9341 | 2.6976 | 2.2122 | -0.47199 | 2.95 |
| 1080 | *RPUSD3* | 4.3414 | 2.5784 | 1.9229 | -2.2352 | 2.95 |
| 1081 | *ZNF79* | 5.27 | 2.8724 | 0.69967 | -0.443 | 2.95 |
| 1082 | *PRAMEF4* | 4.1471 | 3.4171 | 1.2763 | -2.0016 | 2.95 |
| 1083 | *AGO3* | 5.5504 | 3.7461 | -0.45867 | -1.3683 | 2.95 |
| 1084 | *NUCKS1* | 5.0009 | 2.9873 | 0.84087 | -1.0296 | 2.94 |
| 1085 | *MMAB* | 3.4684 | 2.9129 | 2.4449 | -0.82654 | 2.94 |
| 1086 | *CTSG* | 4.4699 | 3.2102 | 1.146 | -2.3282 | 2.94 |
| 1087 | *BRIP1* | 5.741 | 2.8288 | 0.25593 | -0.86712 | 2.94 |
| 1088 | *STT3A* | 3.9241 | 2.7018 | 2.1981 | -0.70665 | 2.94 |
| 1089 | *NOTCH2NL* | 4.8051 | 3.2884 | 0.72968 | -0.44353 | 2.94 |
| 1090 | *ABCD2* | 4.6386 | 3.6591 | 0.52504 | -1.3951 | 2.94 |
| 1091 | *CTSW* | 4.9754 | 3.2011 | 0.64535 | -1.349 | 2.94 |
| 1092 | *SPRR2G* | 4.1826 | 3.1833 | 1.4556 | -0.35182 | 2.94 |
| 1093 | *SSX2B* | 6.1791 | 3.5233 | -0.88159 | -1.2567 | 2.94 |
| 1094 | *LINC00452* | 4.7856 | 4.1109 | -0.07659 | -0.47481 | 2.94 |
| 1095 | *TNFSF18* | 3.7726 | 3.7624 | 1.2836 | 0.78283 | 2.94 |
| 1096 | *KRTAP12-4* | 4.3989 | 2.9671 | 1.452 | -0.23203 | 2.94 |
| 1097 | *RP1* | 4.7309 | 3.1755 | 0.9111 | 0.84999 | 2.94 |

| | | | | | | |
|---|---|---|---|---|---|---|
| 1098 | *TNPO2* | 3.6145 | 3.2661 | 1.9368 | -0.82056 | 2.94 |
| 1099 | *EPOR* | 5.4961 | 2.2852 | 1.0352 | -0.55288 | 2.94 |
| 1100 | *UGT1A1* | 3.79 | 3.1636 | 1.8622 | -0.26519 | 2.94 |
| 1101 | *TYR* | 3.7193 | 2.6991 | 2.3968 | 0.062078 | 2.94 |
| 1102 | *OR10P1* | 3.0758 | 2.8727 | 2.8659 | 1.3206 | 2.94 |
| 1103 | *MTRR* | 5.1734 | 2.6217 | 1.0183 | -1.7766 | 2.94 |
| 1104 | *OR5H15* | 3.7923 | 2.954 | 2.0654 | -2.3372 | 2.94 |
| 1105 | *MYOCD* | 3.9387 | 3.0625 | 1.8082 | 0.58461 | 2.94 |
| 1106 | *TAOK2* | 4.5718 | 2.5023 | 1.7331 | 1.1449 | 2.94 |
| 1107 | *EPPIN-WFDC6* | 4.6188 | 2.5458 | 1.6424 | -1.7493 | 2.94 |
| 1108 | *PRELID2* | 4.0016 | 3.4987 | 1.3057 | -0.27982 | 2.94 |
| 1109 | *CEP112* | 4.1141 | 3.85 | 0.8389 | -1.2867 | 2.93 |
| 1110 | *SENP1* | 4.2278 | 3.6037 | 0.97037 | -1.3117 | 2.93 |
| 1111 | *RABL3* | 3.3878 | 3.2089 | 2.2046 | -1.0453 | 2.93 |
| 1112 | *FTH1P18* | 3.0823 | 3.0363 | 2.6821 | 0.30568 | 2.93 |
| 1113 | *ANAPC13* | 4.4564 | 4.2447 | 0.0995 | -0.58959 | 2.93 |
| 1114 | *TJP1* | 5.2753 | 4.7364 | -1.2114 | -3.0595 | 2.93 |
| 1115 | *FBXL5* | 3.5885 | 3.2524 | 1.9588 | 1.5088 | 2.93 |
| 1116 | *IDH1* | 4.135 | 3.7324 | 0.93217 | -0.57042 | 2.93 |
| 1117 | *PODN* | 4.5425 | 2.9266 | 1.3303 | -0.83883 | 2.93 |
| 1118 | *LRRC40* | 4.0816 | 3.4724 | 1.2446 | -0.02523 | 2.93 |
| 1119 | *TMEM38B* | 5.4802 | 2.1784 | 1.1372 | -0.53929 | 2.93 |
| 1120 | *LOC100129520* | 4.1716 | 3.5212 | 1.1024 | -0.26315 | 2.93 |
| 1121 | *GID8* | 4.364 | 4.001 | 0.42975 | -2.2456 | 2.93 |
| 1122 | *NLRP5* | 4.3029 | 2.7724 | 1.7175 | -1.2246 | 2.93 |
| 1123 | *PRAMEF17* | 4.8436 | 4.0498 | -0.1042 | -0.44137 | 2.93 |
| 1124 | *GUSB* | 4.0694 | 2.4131 | 2.304 | 0.19372 | 2.93 |
| 1125 | *GRID1* | 5.0541 | 3.7062 | 0.0251 | -0.97138 | 2.93 |
| 1126 | *SLC35F2* | 3.9631 | 3.3919 | 1.4289 | -1.535 | 2.93 |
| 1127 | *GP6* | 3.91 | 3.8118 | 1.0613 | -1.4864 | 2.93 |
| 1128 | *BTBD16* | 5.1672 | 3.4878 | 0.12703 | -1.4684 | 2.93 |
| 1129 | *EPGN* | 4.7327 | 3.8423 | 0.20381 | -0.98062 | 2.93 |
| 1130 | *DRC7* | 4.6641 | 4.4112 | -0.29752 | -1.3664 | 2.93 |
| 1131 | *RNF130* | 4.9795 | 3.556 | 0.23983 | -0.22048 | 2.93 |
| 1132 | *PUM1* | 4.9991 | 3.3556 | 0.41853 | -1.5244 | 2.92 |
| 1133 | *HIST1H3G* | 4.4766 | 3.9776 | 0.31771 | 0.24995 | 2.92 |
| 1134 | *TMEM167B* | 5.0909 | 3.8581 | -0.18006 | -0.8952 | 2.92 |
| 1135 | *GABRQ* | 4.1085 | 3.0268 | 1.6321 | 0.86523 | 2.92 |
| 1136 | *SIPA1* | 3.9524 | 2.5444 | 2.2672 | -0.08614 | 2.92 |
| 1137 | *C1orf174* | 4.4265 | 3.9783 | 0.3582 | -1.9054 | 2.92 |
| 1138 | *MARCH3* | 5.3339 | 3.2207 | 0.20193 | -0.80636 | 2.92 |
| 1139 | *GALNT4* | 5.0701 | 3.9388 | -0.25253 | -1.1169 | 2.92 |
| 1140 | *BST2* | 4.1854 | 3.4182 | 1.1523 | -0.85794 | 2.92 |
| 1141 | *HAAO* | 4.8631 | 2.2106 | 1.6811 | 0.83672 | 2.92 |
| 1142 | *TECTA* | 4.7847 | 2.9491 | 1.0205 | -0.7512 | 2.92 |
| 1143 | *CASP8* | 4.0207 | 2.7323 | 1.9964 | -0.40112 | 2.92 |

| | | | | | | |
|---|---|---|---|---|---|---|
| 1144 | *MUC4* | 4.2394 | 3.3048 | 1.2034 | 0.9368 | 2.92 |
| 1145 | *DNAH14* | 4.0239 | 3.7823 | 0.9403 | -2.1342 | 2.92 |
| 1146 | *ADNP2* | 4.2209 | 3.5921 | 0.93283 | -1.5631 | 2.92 |
| 1147 | *TOP3B* | 3.579 | 3.4002 | 1.7658 | 0.011438 | 2.92 |
| 1148 | *CD209* | 5.7087 | 3.5215 | -0.48756 | -0.5101 | 2.91 |
| 1149 | *IFT88* | 3.5045 | 2.6296 | 2.6034 | 1.1922 | 2.91 |
| 1150 | *SRR* | 4.3387 | 4.1324 | 0.26451 | -1.4962 | 2.91 |
| 1151 | *ADAM7* | 4.045 | 4.0323 | 0.65411 | 0.24553 | 2.91 |
| 1152 | *GSE1* | 4.6714 | 4.3716 | -0.31191 | -0.31232 | 2.91 |
| 1153 | *CHD9* | 4.2157 | 3.1526 | 1.3599 | -0.24191 | 2.91 |
| 1154 | *FLT3LG* | 5.1714 | 2.7314 | 0.82421 | -1.2569 | 2.91 |
| 1155 | *C1orf186* | 4.1236 | 2.6243 | 1.9781 | 1.0849 | 2.91 |
| 1156 | *TREX2* | 5.1681 | 3.1939 | 0.36019 | 0.20959 | 2.91 |
| 1157 | *RASSF7* | 4.6482 | 2.1542 | 1.9181 | 0.68157 | 2.91 |
| 1158 | *TNIP3* | 3.9534 | 3.0639 | 1.7012 | -0.04425 | 2.91 |
| 1159 | *BCL11B* | 5.0381 | 2.7461 | 0.93044 | -0.27541 | 2.90 |
| 1160 | *GAP43* | 3.7123 | 3.0737 | 1.9268 | -2.0059 | 2.90 |
| 1161 | *PACSIN3* | 3.5093 | 2.7181 | 2.4849 | 0.47147 | 2.90 |
| 1162 | *ZNF726* | 4.423 | 4.3931 | -0.10676 | -0.58957 | 2.90 |
| 1163 | *C9orf173* | 4.075 | 3.1419 | 1.4916 | 0.91604 | 2.90 |
| 1164 | *CES5A* | 3.5822 | 3.4474 | 1.6778 | -1.5533 | 2.90 |
| 1165 | *TRAF5* | 3.7426 | 3.7098 | 1.255 | -1.5552 | 2.90 |
| 1166 | *PAGE2* | 3.6673 | 2.9186 | 2.1184 | 0.94917 | 2.90 |
| 1167 | *TFDP3* | 5.6809 | 3.418 | -0.39506 | -2.4394 | 2.90 |
| 1168 | *ZNF26* | 4.5689 | 2.8056 | 1.3283 | -1.2509 | 2.90 |
| 1169 | *LIF* | 5.3015 | 2.8296 | 0.57159 | -2.2165 | 2.90 |
| 1170 | *SLC2A11* | 5.5485 | 3.644 | -0.49247 | -1.5765 | 2.90 |
| 1171 | *TRH* | 4.4138 | 3.7811 | 0.50496 | -0.82326 | 2.90 |
| 1172 | *TARBP2* | 3.2086 | 3.0846 | 2.4062 | -0.74913 | 2.90 |
| 1173 | *FAM214A* | 4.2329 | 2.8656 | 1.5994 | 0.34505 | 2.90 |
| 1174 | *LGI4* | 4.0663 | 3.8502 | 0.78012 | -0.81512 | 2.90 |
| 1175 | *RAB20* | 3.5257 | 2.6628 | 2.5081 | 0.9037 | 2.90 |
| 1176 | *UMAD1* | 5.6086 | 4.5717 | -1.4841 | -1.8157 | 2.90 |
| 1177 | *RASAL3* | 4.0655 | 3.3435 | 1.287 | -1.7365 | 2.90 |
| 1178 | *FBXL6* | 3.876 | 3.7664 | 1.0484 | 0.63235 | 2.90 |
| 1179 | *MFGE8* | 5.0259 | 4.2057 | -0.54391 | -1.8851 | 2.90 |
| 1180 | *CTCF* | 3.9009 | 2.6273 | 2.1583 | 0.030912 | 2.90 |
| 1181 | *ATXN7* | 5.2068 | 2.0996 | 1.3796 | 0.14155 | 2.90 |
| 1182 | *FNIP2* | 4.8164 | 3.401 | 0.46662 | -0.275 | 2.89 |
| 1183 | *KCNJ18* | 3.9283 | 2.6443 | 2.1111 | 2.0277 | 2.89 |
| 1184 | *SMKR1* | 3.5192 | 2.9383 | 2.2254 | -4.8437 | 2.89 |
| 1185 | *GALE* | 4.6078 | 3.5179 | 0.55475 | -0.44353 | 2.89 |
| 1186 | *NOL8* | 3.3045 | 2.7793 | 2.5929 | -1.7528 | 2.89 |
| 1187 | *RFX3* | 5.1885 | 2.2917 | 1.1952 | -1.7277 | 2.89 |
| 1188 | *ZDBF2* | 4.8695 | 4.0574 | -0.25605 | -1.6885 | 2.89 |
| 1189 | *HDGFL1* | 4.8364 | 4.2593 | -0.42615 | -1.9919 | 2.89 |

| | | | | | | |
|---|---|---|---|---|---|---|
| 1190 | *ZRANB3* | 3.3219 | 2.7865 | 2.56 | 0.072941 | 2.89 |
| 1191 | *SPINK2* | 4.1119 | 3.4657 | 1.0892 | -0.67853 | 2.89 |
| 1192 | *PHACTR4* | 3.7822 | 3.2151 | 1.6667 | -1.42 | 2.89 |
| 1193 | *ITGAD* | 5.1368 | 4.0171 | -0.49127 | -0.85565 | 2.89 |
| 1194 | *SMIM3* | 4.2499 | 3.1964 | 1.2163 | 0.061214 | 2.89 |
| 1195 | *FAM214B* | 4.5384 | 3.4723 | 0.65175 | 0.41875 | 2.89 |
| 1196 | *FEM1B* | 4.6708 | 3.9851 | 0.00589 | -0.60113 | 2.89 |
| 1197 | *OPRL1* | 5.3667 | 3.1156 | 0.17807 | -0.89699 | 2.89 |
| 1198 | *CCND2* | 3.8015 | 2.5993 | 2.2565 | -1.3186 | 2.89 |
| 1199 | *LDHD* | 4.0835 | 2.7488 | 1.8192 | -0.01493 | 2.88 |
| 1200 | *PCSK4* | 4.3427 | 2.4198 | 1.8885 | -1.3028 | 2.88 |
| 1201 | *SORT1* | 4.8838 | 2.6705 | 1.0964 | -1.9381 | 2.88 |
| 1202 | *GID4* | 3.3905 | 2.7353 | 2.5224 | -0.68869 | 2.88 |
| 1203 | *CCNT2* | 4.5132 | 3.7629 | 0.37208 | -1.7229 | 2.88 |
| 1204 | *PCDHB12* | 4.3416 | 3.1542 | 1.1509 | -2.2075 | 2.88 |
| 1205 | *NFIL3* | 3.994 | 3.897 | 0.75464 | -0.87655 | 2.88 |
| 1206 | *PIN1* | 3.2757 | 3.2454 | 2.1208 | 0.2288 | 2.88 |
| 1207 | *SRD5A2* | 4.0375 | 2.3342 | 2.2702 | 0.89431 | 2.88 |
| 1208 | *STH* | 4.6276 | 2.8879 | 1.1236 | -0.51767 | 2.88 |
| 1209 | *HOXD9* | 3.8217 | 2.7078 | 2.1087 | -0.17238 | 2.88 |
| 1210 | *CCDC122* | 3.5945 | 3.5298 | 1.5139 | -0.19718 | 2.88 |
| 1211 | *FASTKD2* | 3.2988 | 3.0667 | 2.2686 | -0.14091 | 2.88 |
| 1212 | *NRK* | 3.9294 | 3.8579 | 0.84579 | -0.45757 | 2.88 |
| 1213 | *PCCA* | 3.7561 | 3.2357 | 1.6401 | 0.44709 | 2.88 |
| 1214 | *MTAP* | 4.0849 | 3.2223 | 1.3244 | -1.2716 | 2.88 |
| 1215 | *IRAK1* | 3.4585 | 3.08 | 2.0924 | -0.64967 | 2.88 |
| 1216 | *SMIM8* | 4.2484 | 3.6562 | 0.72413 | -0.61818 | 2.88 |
| 1217 | *RHCG* | 3.2744 | 3.1176 | 2.2366 | -1.6735 | 2.88 |
| 1218 | *TBX22* | 3.6303 | 3.2497 | 1.7468 | -0.1536 | 2.88 |
| 1219 | *AGBL2* | 4.2784 | 3.715 | 0.62861 | 0.060969 | 2.87 |
| 1220 | *CHST8* | 3.6988 | 2.7624 | 2.1607 | -1.5996 | 2.87 |
| 1221 | *PGPEP1* | 4.3938 | 2.7655 | 1.4617 | -2.3358 | 2.87 |
| 1222 | *ACTL8* | 4.416 | 2.7603 | 1.4439 | -1.8972 | 2.87 |
| 1223 | *KIAA1217* | 4.8495 | 3.5308 | 0.23715 | -1.4996 | 2.87 |
| 1224 | *MRGPRG* | 3.5111 | 2.7983 | 2.3062 | 0.65057 | 2.87 |
| 1225 | *SDR9C7* | 4.9411 | 2.5212 | 1.1528 | -1.8623 | 2.87 |
| 1226 | *NAALADL2* | 4.75 | 3.47 | 0.39313 | 0.22925 | 2.87 |
| 1227 | *CDC42EP1* | 4.7866 | 3.6596 | 0.16456 | -2.1155 | 2.87 |
| 1228 | *HNRNPDL* | 4.363 | 4.3616 | -0.12092 | -0.35328 | 2.87 |
| 1229 | *MBD4* | 4.5859 | 2.2304 | 1.7845 | -0.93112 | 2.87 |
| 1230 | *USP49* | 4.7424 | 3.46 | 0.39528 | -0.03976 | 2.87 |
| 1231 | *KCNJ5* | 3.8815 | 3.5064 | 1.2069 | -0.93892 | 2.86 |
| 1232 | *OR56B4* | 3.8056 | 3.0258 | 1.7628 | -1.9548 | 2.86 |
| 1233 | *GRM5* | 4.2259 | 4.0409 | 0.31613 | -1.5073 | 2.86 |
| 1234 | *PLEKHH2* | 5.0142 | 2.8408 | 0.72458 | -2.1669 | 2.86 |
| 1235 | *ZCCHC17* | 5.1517 | 2.7591 | 0.66646 | -1.4782 | 2.86 |

| | | | | | | |
|---|---|---|---|---|---|---|
| 1236 | *TMEM60* | 3.389 | 2.8146 | 2.3717 | 0.54766 | 2.86 |
| 1237 | *CALCR* | 4.0385 | 2.5096 | 2.0257 | 0.18696 | 2.86 |
| 1238 | *MAST2* | 4.1309 | 2.9834 | 1.459 | -0.73145 | 2.86 |
| 1239 | *USP43* | 3.5652 | 2.6289 | 2.3774 | -1.1858 | 2.86 |
| 1240 | *TAF7* | 4.0101 | 3.9714 | 0.58878 | 0.49222 | 2.86 |
| 1241 | *CSF2RB* | 4.513 | 3.5121 | 0.54515 | 0.22531 | 2.86 |
| 1242 | *RAD18* | 3.6179 | 3.4508 | 1.5003 | 0.60507 | 2.86 |
| 1243 | *COL8A2* | 3.5079 | 3.3123 | 1.748 | -0.45373 | 2.86 |
| 1244 | *IRAK1BP1* | 4.4106 | 4.3294 | -0.17696 | -2.4333 | 2.85 |
| 1245 | *AOC2* | 5.047 | 2.1162 | 1.3983 | -1.0733 | 2.85 |
| 1246 | *TMEM117* | 4.4458 | 3.8287 | 0.28648 | 0.25825 | 2.85 |
| 1247 | *SNX2* | 3.7512 | 3.4984 | 1.3104 | 0.92154 | 2.85 |
| 1248 | *ZNF277* | 4.6095 | 3.3117 | 0.63682 | 0.32367 | 2.85 |
| 1249 | *PDE7A* | 5.4921 | 2.3174 | 0.74748 | -0.69399 | 2.85 |
| 1250 | *CEACAM4* | 5.0818 | 2.6774 | 0.79458 | 0.001501 | 2.85 |
| 1251 | *CAB39L* | 4.8941 | 3.7669 | -0.10857 | -1.2399 | 2.85 |
| 1252 | *SLC9A7* | 5.2914 | 1.8285 | 1.4319 | -0.03501 | 2.85 |
| 1253 | *NRG4* | 4.4828 | 4.2616 | -0.19364 | -1.7766 | 2.85 |
| 1254 | *SLC5A9* | 4.9581 | 3.4007 | 0.18807 | -0.11483 | 2.85 |
| 1255 | *ZNF800* | 3.6462 | 3.1437 | 1.7568 | -1.1523 | 2.85 |
| 1256 | *C17orf96* | 5.1686 | 4.2416 | -0.86358 | -1.1263 | 2.85 |
| 1257 | *VAX1* | 5.353 | 1.7374 | 1.4561 | -0.27023 | 2.85 |
| 1258 | *CAPN13* | 5.2067 | 2.3913 | 0.94832 | -0.03648 | 2.85 |
| 1259 | *PDGFD* | 4.7947 | 2.9079 | 0.8417 | -0.80432 | 2.85 |
| 1260 | *TGDS* | 4.8199 | 4.4643 | -0.74279 | -1.3272 | 2.85 |
| 1261 | *PRKRIR* | 5.4539 | 3.3195 | -0.23264 | -1.2797 | 2.85 |
| 1262 | *B3GNT5* | 4.7656 | 2.7517 | 1.022 | -1.9062 | 2.85 |
| 1263 | *ZKSCAN1* | 4.4728 | 2.4454 | 1.6205 | -1.439 | 2.85 |
| 1264 | *FAM171A1* | 3.6915 | 3.1351 | 1.7113 | 1.6846 | 2.85 |
| 1265 | *CELSR3* | 4.1713 | 3.9408 | 0.42579 | -0.98099 | 2.85 |
| 1266 | *TMEM155* | 4.451 | 3.7448 | 0.34011 | -1.1692 | 2.85 |
| 1267 | *ODF3L1* | 4.4334 | 2.6782 | 1.4233 | -1.237 | 2.84 |
| 1268 | *CPO* | 5.0489 | 2.4395 | 1.0447 | 0.32798 | 2.84 |
| 1269 | *OSCP1* | 4.2926 | 3.4548 | 0.78479 | -0.27771 | 2.84 |
| 1270 | *SMLR1* | 5.4296 | 1.9785 | 1.1229 | 0.82871 | 2.84 |
| 1271 | *PCCB* | 4.6524 | 2.6731 | 1.2025 | -1.146 | 2.84 |
| 1272 | *GCM2* | 4.9558 | 3.5614 | 0.0091 | -0.69961 | 2.84 |
| 1273 | *TXNRD3* | 5.7086 | 2.7338 | 0.08159 | -0.62125 | 2.84 |
| 1274 | *FBXO25* | 4.4386 | 3.5576 | 0.52708 | -0.81963 | 2.84 |
| 1275 | *MSRB3* | 4.6591 | 1.9853 | 1.8778 | 0.78652 | 2.84 |
| 1276 | *WIPF1* | 5.3792 | 3.8415 | -0.70193 | -1.9099 | 2.84 |
| 1277 | *SLC1A1* | 4.8539 | 3.9534 | -0.28901 | -0.85345 | 2.84 |
| 1278 | *ZSCAN18* | 4.5094 | 3.0808 | 0.92731 | 0.91889 | 2.84 |
| 1279 | *NIPAL1* | 4.2196 | 2.9623 | 1.3352 | -4.897 | 2.84 |
| 1280 | *COL4A1* | 4.3944 | 2.1502 | 1.9694 | 0.46873 | 2.84 |
| 1281 | *SBSPON* | 4.6662 | 3.3753 | 0.47145 | -0.80213 | 2.84 |

| | | | | | | |
|---|---|---|---|---|---|---|
| 1282 | *CCDC132* | 3.4168 | 2.9147 | 2.1752 | 0.50266 | 2.84 |
| 1283 | *P2RX6* | 4.7916 | 3.0261 | 0.68874 | -0.38638 | 2.84 |
| 1284 | *C5AR1* | 4.1166 | 2.7594 | 1.6296 | 0.40518 | 2.84 |
| 1285 | *CYSTM1* | 4.566 | 2.9097 | 1.0298 | -1.6594 | 2.84 |
| 1286 | *CLPTM1* | 3.0272 | 2.982 | 2.4924 | 2.3659 | 2.83 |
| 1287 | *SOX2* | 4.0187 | 3.0091 | 1.4691 | 0.031644 | 2.83 |
| 1288 | *ERCC5* | 5.3576 | 2.246 | 0.88995 | 0.65005 | 2.83 |
| 1289 | *HOSPHO2-KLHL2* | 5.4069 | 3.2135 | -0.12728 | -1.0021 | 2.83 |
| 1290 | *CYB5D1* | 3.357 | 2.9614 | 2.1725 | -0.88667 | 2.83 |
| 1291 | *IQCF2* | 3.5237 | 2.8517 | 2.112 | -0.27877 | 2.83 |
| 1292 | *C9orf106* | 4.5164 | 3.0074 | 0.96042 | -1.928 | 2.83 |
| 1293 | *CARM1* | 3.5831 | 2.7119 | 2.1881 | -1.4864 | 2.83 |
| 1294 | *TSPO2* | 3.8161 | 2.502 | 2.162 | 0.81991 | 2.83 |
| 1295 | *KDM5B* | 5.1148 | 3.7088 | -0.34553 | -0.67417 | 2.83 |
| 1296 | *NID2* | 3.3151 | 2.8767 | 2.2859 | 0.1789 | 2.83 |
| 1297 | *DCLK3* | 4.1385 | 3.2044 | 1.1347 | -1.7235 | 2.83 |
| 1298 | *CDC20B* | 5.184 | 3.7698 | -0.47777 | -2.1921 | 2.83 |
| 1299 | *GRIA1* | 5.0919 | 2.31 | 1.0735 | -0.08301 | 2.83 |
| 1300 | *ZFAT* | 5.2738 | 3.3665 | -0.16622 | -1.9693 | 2.82 |
| 1301 | *LIPC* | 4.8323 | 2.4576 | 1.184 | -1.2864 | 2.82 |
| 1302 | *NEK2* | 4.3853 | 3.0063 | 1.0821 | -0.65417 | 2.82 |
| 1303 | *IZUMO3* | 5.4185 | 1.9225 | 1.1322 | 1.0872 | 2.82 |
| 1304 | *EPHA7* | 4.113 | 3.7221 | 0.63521 | -1.622 | 2.82 |
| 1305 | *CTRB2* | 4.351 | 3.5716 | 0.54694 | -0.58349 | 2.82 |
| 1306 | *COMMD9* | 2.9833 | 2.9114 | 2.5748 | -1.9295 | 2.82 |
| 1307 | *CLEC9A* | 3.4066 | 3.0792 | 1.9836 | -0.67458 | 2.82 |
| 1308 | *HERPUD1* | 4.8498 | 3.8271 | -0.2098 | -0.57133 | 2.82 |
| 1309 | *MAPT* | 3.6966 | 2.6792 | 2.0907 | -1.1629 | 2.82 |
| 1310 | *CEP63* | 3.0465 | 2.8278 | 2.592 | -2.372 | 2.82 |
| 1311 | *MCM3AP* | 4.8698 | 3.3412 | 0.25501 | -0.95579 | 2.82 |
| 1312 | *TPRA1* | 4.2317 | 3.8951 | 0.33384 | -0.19793 | 2.82 |
| 1313 | *PHKA2* | 4.9085 | 4.0662 | -0.5144 | -1.7994 | 2.82 |
| 1314 | *PEX7* | 4.5146 | 2.9018 | 1.0428 | -0.71421 | 2.82 |
| 1315 | *ZNF677* | 4.4579 | 4.0742 | -0.07436 | -1.4936 | 2.82 |
| 1316 | *RAB27B* | 4.4394 | 2.6883 | 1.3298 | -1.2342 | 2.82 |
| 1317 | *TMBIM6* | 4.5156 | 2.5037 | 1.4356 | -1.8357 | 2.82 |
| 1318 | *NECAB3* | 3.2286 | 2.8672 | 2.3585 | -0.51713 | 2.82 |
| 1319 | *P3H1* | 5.3196 | 2.229 | 0.90514 | -0.66488 | 2.82 |
| 1320 | *ALDH3B1* | 5.394 | 3.9479 | -0.8895 | -1.7696 | 2.82 |
| 1321 | *EPB41L2* | 4.1973 | 3.8499 | 0.40435 | -1.8842 | 2.82 |
| 1322 | *ENDOG* | 4.6211 | 4.0178 | -0.18783 | -2.7616 | 2.82 |
| 1323 | *GSPT2* | 3.4412 | 3.167 | 1.8424 | -1.773 | 2.82 |
| 1324 | *PPP2R2A* | 4.3302 | 2.5245 | 1.5933 | 1.2979 | 2.82 |
| 1325 | *NR2F1* | 4.4606 | 3.6647 | 0.32236 | -1.4607 | 2.82 |
| 1326 | *PNMA3* | 3.8525 | 3.7211 | 0.87394 | -0.79678 | 2.82 |
| 1327 | *CLEC12A* | 3.5417 | 2.6411 | 2.2643 | 1.9213 | 2.82 |

| | | | | | | |
|---|---|---|---|---|---|---|
| 1328 | *KLRC4* | 4.9605 | 3.1974 | 0.28605 | -1.683 | 2.81 |
| 1329 | *APPL2* | 4.0112 | 2.4863 | 1.9461 | -0.72819 | 2.81 |
| 1330 | *ANAPC7* | 3.1499 | 2.9025 | 2.3901 | 0.50427 | 2.81 |
| 1331 | *HCRTR2* | 3.4579 | 2.7287 | 2.2559 | -0.26575 | 2.81 |
| 1332 | *RIPK4* | 3.9511 | 2.8588 | 1.6319 | -0.42356 | 2.81 |
| 1333 | *ZAR1* | 4.0651 | 3.6147 | 0.76029 | -1.2981 | 2.81 |
| 1334 | *STEAP1* | 3.4538 | 2.8569 | 2.1256 | -0.42276 | 2.81 |
| 1335 | *PRR4* | 4.6279 | 3.4663 | 0.34044 | 0.22469 | 2.81 |
| 1336 | *SEC14L2* | 5.0485 | 3.0477 | 0.33838 | -1.6926 | 2.81 |
| 1337 | *DPP10* | 3.9781 | 2.6682 | 1.7882 | 0.68145 | 2.81 |
| 1338 | *CD24* | 4.4206 | 2.7263 | 1.2862 | 0.38204 | 2.81 |
| 1339 | *LAMA1* | 5.2083 | 4.3321 | -1.1075 | -1.9558 | 2.81 |
| 1340 | *KIAA1586* | 4.3785 | 3.1965 | 0.85386 | -0.8472 | 2.81 |
| 1341 | *PCYOX1L* | 4.2162 | 2.4085 | 1.8021 | 1.4331 | 2.81 |
| 1342 | *GNRH2* | 4.8808 | 3.2895 | 0.24705 | -0.12998 | 2.81 |
| 1343 | *SASS6* | 5.166 | 2.3808 | 0.8688 | -0.88818 | 2.81 |
| 1344 | *CXCR6* | 4.3343 | 2.4717 | 1.6091 | -0.42181 | 2.81 |
| 1345 | *TRIM56* | 4.5218 | 3.0098 | 0.88094 | -0.71093 | 2.80 |
| 1346 | *RAC1* | 4.014 | 3.4441 | 0.95209 | -0.77418 | 2.80 |
| 1347 | *FGD3* | 4.4766 | 3.1629 | 0.77066 | -1.2571 | 2.80 |
| 1348 | *ZNF573* | 4.0419 | 3.642 | 0.72381 | -1.1953 | 2.80 |
| 1349 | *OR5M10* | 4.6001 | 2.9024 | 0.90518 | -1.2662 | 2.80 |
| 1350 | *EPYC* | 3.877 | 3.3796 | 1.1509 | 0.28289 | 2.80 |
| 1351 | *TDRD5* | 5.2106 | 4.0887 | -0.89219 | -0.89988 | 2.80 |
| 1352 | *WFDC8* | 4.14 | 3.3624 | 0.90312 | 0.37605 | 2.80 |
| 1353 | *FUT3* | 4.8411 | 4.512 | -0.94903 | -1.6374 | 2.80 |
| 1354 | *PDHA2* | 3.953 | 2.5872 | 1.8638 | -0.52042 | 2.80 |
| 1355 | *ELAVL4* | 4.8626 | 4.2346 | -0.69678 | -1.0388 | 2.80 |
| 1356 | *CCNE2* | 5.4593 | 3.4115 | -0.47135 | -0.85194 | 2.80 |
| 1357 | *RNF103* | 5.2594 | 2.4022 | 0.73769 | -0.58597 | 2.80 |
| 1358 | *ZFP82* | 5.0217 | 3.0565 | 0.31975 | -0.26383 | 2.80 |
| 1359 | *PDZRN3* | 3.7257 | 3.588 | 1.0839 | 0.1409 | 2.80 |
| 1360 | *PCYOX1* | 6.0076 | 2.5931 | -0.20648 | -0.52732 | 2.80 |
| 1361 | *PCBD2* | 3.0564 | 2.967 | 2.3705 | 0.45236 | 2.80 |
| 1362 | *NXPH2* | 4.2397 | 3.4905 | 0.66302 | 0.5806 | 2.80 |
| 1363 | *HSPBP1* | 3.8365 | 3.6208 | 0.93487 | 0.86057 | 2.80 |
| 1364 | *CD6* | 4.8978 | 4.2082 | -0.71467 | -1.7753 | 2.80 |
| 1365 | *S100G* | 4.6102 | 2.1729 | 1.6036 | 0.94601 | 2.80 |
| 1366 | *ERBB2IP* | 5.3468 | 2.1246 | 0.91528 | -1.8961 | 2.80 |
| 1367 | *OR2T8* | 4.3014 | 2.6993 | 1.384 | 0.98094 | 2.79 |
| 1368 | *BPY2C* | 5.1925 | 4.1826 | -0.99059 | -1.3038 | 2.79 |
| 1369 | *OGFRL1* | 5.5437 | 3.0388 | -0.19943 | -0.42153 | 2.79 |
| 1370 | *CNGB1* | 4.9353 | 3.9479 | -0.50053 | -1.5485 | 2.79 |
| 1371 | *FCAMR* | 2.8954 | 2.8111 | 2.6749 | 2.0581 | 2.79 |
| 1372 | *NXF5* | 4.2545 | 3.8788 | 0.24698 | -1.1591 | 2.79 |
| 1373 | *CRYGB* | 4.26 | 2.4257 | 1.6921 | -2.3778 | 2.79 |

| | | | | | | |
|---|---|---|---|---|---|---|
| 1374 | *TPBG* | 5.6164 | 2.8756 | -0.11452 | -0.43251 | 2.79 |
| 1375 | *NXPE3* | 4.8797 | 2.2295 | 1.2642 | 0.35003 | 2.79 |
| 1376 | *RAI1* | 3.9016 | 3.8814 | 0.58992 | -0.0937 | 2.79 |
| 1377 | *OLA1* | 3.4264 | 3.018 | 1.9276 | -1.0147 | 2.79 |
| 1378 | *POMC* | 4.2731 | 2.1651 | 1.9318 | 1.8788 | 2.79 |
| 1379 | *NTS* | 5.5041 | 1.869 | 0.9942 | -1.1598 | 2.79 |
| 1380 | *USF1* | 4.3312 | 2.4149 | 1.6211 | -0.07266 | 2.79 |
| 1381 | *FXYD7* | 3.8342 | 3.8256 | 0.70739 | -1.3844 | 2.79 |
| 1382 | *SCAMP3* | 4.2334 | 3.2108 | 0.92165 | -1.988 | 2.79 |
| 1383 | *ROGDI* | 4.6624 | 3.3687 | 0.33301 | -1.5744 | 2.79 |
| 1384 | *THAP5* | 3.6323 | 3.0806 | 1.6496 | 1.0586 | 2.79 |
| 1385 | *CCDC60* | 4.2551 | 3.5873 | 0.51984 | -2.2165 | 2.79 |
| 1386 | *SKA1* | 3.6656 | 3.5467 | 1.1478 | -1.6352 | 2.79 |
| 1387 | *ENOPH1* | 3.2524 | 2.6192 | 2.4876 | -0.72936 | 2.79 |
| 1388 | *LRRC8C* | 4.6322 | 2.7106 | 1.0159 | -0.03346 | 2.79 |
| 1389 | *SLC35F6* | 5.0765 | 3.7351 | -0.45296 | -0.61222 | 2.79 |
| 1390 | *GALNT8* | 4.182 | 2.3117 | 1.8597 | 0.13609 | 2.78 |
| 1391 | *SPINK14* | 4.0886 | 2.2766 | 1.9875 | -0.78959 | 2.78 |
| 1392 | *PDZRN4* | 3.4594 | 2.61 | 2.282 | -0.7763 | 2.78 |
| 1393 | *TLR10* | 4.6269 | 3.2775 | 0.44601 | -1.4123 | 2.78 |
| 1394 | *TMUB2* | 5.0108 | 2.3063 | 1.0311 | -2.0258 | 2.78 |
| 1395 | *FAM89B* | 4.4146 | 4.2645 | -0.33098 | -2.0871 | 2.78 |
| 1396 | *PCED1A* | 5.4351 | 2.699 | 0.21153 | -1.2917 | 2.78 |
| 1397 | *CENPQ* | 3.0037 | 2.7553 | 2.585 | 0.052809 | 2.78 |
| 1398 | *BEX2* | 4.5575 | 3.5197 | 0.26607 | 0.006223 | 2.78 |
| 1399 | *ZNF84* | 3.4105 | 3.0534 | 1.8793 | 0.65684 | 2.78 |
| 1400 | *PRKAA1* | 4.4865 | 3.2193 | 0.63592 | -0.55492 | 2.78 |
| 1401 | *MEF2BNB-MEF2B* | 4.6151 | 2.0874 | 1.6372 | -0.34051 | 2.78 |
| 1402 | *SUCNR1* | 4.1469 | 2.3184 | 1.8733 | -0.42899 | 2.78 |
| 1403 | *SOX14* | 4.1246 | 3.8289 | 0.38458 | -1.2676 | 2.78 |
| 1404 | *CYP46A1* | 3.3407 | 3.1724 | 1.8227 | 0.37505 | 2.78 |
| 1405 | *SLC6A5* | 5.5583 | 1.4878 | 1.2853 | -1.4681 | 2.78 |
| 1406 | *SYNE1* | 4.4157 | 3.8159 | 0.09878 | -0.90092 | 2.78 |
| 1407 | *EPCAM* | 3.8522 | 3.4337 | 1.0426 | -0.03524 | 2.78 |
| 1408 | *SPOCK3* | 4.0412 | 3.4591 | 0.82643 | -0.00546 | 2.78 |
| 1409 | *SPRR2D* | 4.4736 | 3.8793 | -0.02643 | -1.28 | 2.78 |
| 1410 | *SVOP* | 4.424 | 2.6917 | 1.21 | -0.29562 | 2.78 |
| 1411 | *CR1* | 2.9317 | 2.8692 | 2.5242 | 1.145 | 2.78 |
| 1412 | *TMEM247* | 3.7351 | 3.5033 | 1.0866 | -1.5671 | 2.78 |
| 1413 | *MLLT11* | 5.539 | 3.1512 | -0.36569 | -1.1825 | 2.77 |
| 1414 | *FOXL2* | 4.2046 | 2.7024 | 1.4171 | 1.0621 | 2.77 |
| 1415 | *ARL6* | 3.1736 | 2.7699 | 2.3795 | -1.1699 | 2.77 |
| 1416 | *GADD45B* | 4.6501 | 2.3365 | 1.3354 | -0.31317 | 2.77 |
| 1417 | *FAM208B* | 4.1013 | 2.6519 | 1.5657 | -1.3952 | 2.77 |
| 1418 | *ALDH3A1* | 5.1778 | 3.7609 | -0.62005 | -0.62833 | 2.77 |
| 1419 | *GABRB2* | 5.4508 | 2.1427 | 0.72495 | 0.14269 | 2.77 |

| | | | | | | |
|---|---|---|---|---|---|---|
| 1420 | *TRABD* | 5.0215 | 2.4368 | 0.85932 | -1.4541 | 2.77 |
| 1421 | *RAD51AP1* | 3.2822 | 2.6131 | 2.4223 | -2.3958 | 2.77 |
| 1422 | *CDC25C* | 3.6867 | 3.1337 | 1.4949 | 0.10279 | 2.77 |
| 1423 | *C9orf64* | 4.3322 | 2.9621 | 1.0184 | -0.71988 | 2.77 |
| 1424 | *SPATA7* | 4.6121 | 2.3532 | 1.3457 | -0.7722 | 2.77 |
| 1425 | *VSIG2* | 4.3328 | 2.1091 | 1.8652 | 0.63802 | 2.77 |
| 1426 | *LHX6* | 4.0632 | 3.4664 | 0.77367 | -1.0516 | 2.77 |
| 1427 | *GM2A* | 4.3299 | 2.5805 | 1.3901 | 0.51513 | 2.77 |
| 1428 | *TBC1D28* | 4.4214 | 3.6466 | 0.23173 | -0.59816 | 2.77 |
| 1429 | *PTBP2* | 3.5308 | 3.416 | 1.352 | -0.84248 | 2.77 |
| 1430 | *OLIG1* | 3.7047 | 3.5762 | 1.0163 | -0.158 | 2.77 |
| 1431 | *RHBDD3* | 3.4976 | 2.591 | 2.2066 | -0.37215 | 2.77 |
| 1432 | *OR1N2* | 3.7794 | 3.6225 | 0.89246 | -0.90296 | 2.76 |
| 1433 | *EPHB6* | 4.2718 | 3.3441 | 0.67678 | -1.2086 | 2.76 |
| 1434 | *PRCC* | 4.6693 | 2.0919 | 1.5285 | 0.52501 | 2.76 |
| 1435 | *ZMYM6NB* | 4.6058 | 3.0984 | 0.58544 | -0.3091 | 2.76 |
| 1436 | *OR2Z1* | 5.1908 | 1.6001 | 1.4976 | -1.8547 | 2.76 |
| 1437 | *BHLHA15* | 2.9692 | 2.8096 | 2.5079 | -0.6714 | 2.76 |
| 1438 | *KCNN1* | 3.6702 | 3.2059 | 1.409 | -0.67913 | 2.76 |
| 1439 | *DNASE1L3* | 4.0443 | 3.7027 | 0.53765 | -1.8775 | 2.76 |
| 1440 | *ZNF878* | 3.6511 | 3.0912 | 1.5421 | -0.68757 | 2.76 |
| 1441 | *E2F4* | 4.5022 | 3.614 | 0.1674 | -0.63845 | 2.76 |
| 1442 | *C3orf58* | 5.256 | 1.6273 | 1.398 | -1.2267 | 2.76 |
| 1443 | *GLRA1* | 4.2667 | 3.5244 | 0.48998 | -1.9675 | 2.76 |
| 1444 | *GPM6B* | 4.4201 | 4.2234 | -0.36315 | -0.61663 | 2.76 |
| 1445 | *TNR* | 4.1678 | 3.7344 | 0.37721 | -0.22962 | 2.76 |
| 1446 | *SMIM21* | 4.7128 | 3.4943 | 0.07209 | -1.2208 | 2.76 |
| 1447 | *PDILT* | 5.0654 | 4.0896 | -0.87817 | -1.7662 | 2.76 |
| 1448 | *OR2T5* | 3.9799 | 3.2483 | 1.0461 | 0.10959 | 2.76 |
| 1449 | *OXR1* | 4.3467 | 3.1132 | 0.81408 | -0.13823 | 2.76 |
| 1450 | *FAXC* | 4.1946 | 2.8641 | 1.2147 | 0.24958 | 2.76 |
| 1451 | *DNAJC4* | 3.3418 | 2.9007 | 2.0261 | -2.1151 | 2.76 |
| 1452 | *ZNF500* | 4.1709 | 3.4386 | 0.65887 | -1.1209 | 2.76 |
| 1453 | *CTSV* | 3.8667 | 3.7265 | 0.67455 | 0.40785 | 2.76 |
| 1454 | *TAZ* | 3.0226 | 2.7064 | 2.533 | 1.9525 | 2.75 |
| 1455 | *KIAA1257* | 2.9372 | 2.8613 | 2.4622 | -0.09762 | 2.75 |
| 1456 | *AMZ2* | 3.6711 | 3.3007 | 1.2875 | -0.44134 | 2.75 |
| 1457 | *ACKR4* | 4.9048 | 3.2031 | 0.14969 | 0.071915 | 2.75 |
| 1458 | *PLA2G4A* | 3.713 | 2.8545 | 1.6896 | 0.35324 | 2.75 |
| 1459 | *MFSD7* | 5.3812 | 2.8146 | 0.05963 | -0.78429 | 2.75 |
| 1460 | *ENTPD2* | 4.9631 | 3.1317 | 0.15808 | -0.5206 | 2.75 |
| 1461 | *PUF60* | 3.2192 | 2.7291 | 2.3045 | -0.23653 | 2.75 |
| 1462 | *TSGA10IP* | 4.0561 | 2.2631 | 1.9332 | -0.42349 | 2.75 |
| 1463 | *SPTLC3* | 3.2415 | 2.962 | 2.0481 | 1.0363 | 2.75 |
| 1464 | *CKLF* | 4.7547 | 3.5675 | -0.07138 | -0.46372 | 2.75 |
| 1465 | *OC90* | 4.9374 | 2.9385 | 0.37296 | -1.9748 | 2.75 |

| | | | | | | |
|---|---|---|---|---|---|---|
| 1466 | *GRIA3* | 4.1808 | 2.6983 | 1.3674 | 0.34861 | 2.75 |
| 1467 | *KCNN2* | 3.3913 | 3.109 | 1.7433 | -1.2426 | 2.75 |
| 1468 | *SEMA4B* | 3.3535 | 2.7446 | 2.1454 | 1.9195 | 2.75 |
| 1469 | *UGT1A4* | 3.523 | 3.1532 | 1.5637 | -1.4749 | 2.75 |
| 1470 | *VPS4B* | 4.3854 | 3.6583 | 0.19594 | -2.2951 | 2.75 |
| 1471 | *DIDO1* | 5.2027 | 2.3244 | 0.71144 | -0.40097 | 2.75 |
| 1472 | *IRF8* | 4.4528 | 3.6245 | 0.16049 | -1.7552 | 2.75 |
| 1473 | *SRM* | 4.7595 | 3.6755 | -0.19943 | -0.94113 | 2.75 |
| 1474 | *NAGPA* | 4.5644 | 2.486 | 1.1842 | 0.64483 | 2.74 |
| 1475 | *UBE2W* | 3.131 | 3.1078 | 1.9953 | 0.70365 | 2.74 |
| 1476 | *PDRG1* | 3.6291 | 3.5344 | 1.0705 | -0.33152 | 2.74 |
| 1477 | *PNLIP* | 3.7207 | 2.9862 | 1.5237 | 0.30216 | 2.74 |
| 1478 | *DNMT3B* | 4.684 | 3.5634 | -0.01699 | -0.91269 | 2.74 |
| 1479 | *ZIC1* | 3.9849 | 2.1641 | 2.0814 | 0.18044 | 2.74 |
| 1480 | *SPATA6* | 4.3547 | 2.3832 | 1.4919 | 1.2313 | 2.74 |
| 1481 | *TRMT5* | 3.2543 | 2.5314 | 2.4421 | 2.3176 | 2.74 |
| 1482 | *TCTEX1D4* | 5.4376 | 2.7664 | 0.02334 | -1.3139 | 2.74 |
| 1483 | *SAT2* | 4.4782 | 3.4213 | 0.32523 | -1.5116 | 2.74 |
| 1484 | *TAAR1* | 3.5476 | 2.8525 | 1.8245 | -0.32122 | 2.74 |
| 1485 | *A4GNT* | 5.3949 | 3.9912 | -1.162 | -2.0337 | 2.74 |
| 1486 | *ALPK2* | 4.6706 | 4.4304 | -0.87898 | -1.917 | 2.74 |
| 1487 | *EXD3* | 4.5957 | 2.2215 | 1.4019 | -0.56634 | 2.74 |
| 1488 | *FOXL2NB* | 5.0109 | 3.2225 | -0.01548 | -2.6188 | 2.74 |
| 1489 | *HHLA3* | 4.054 | 2.0978 | 2.0661 | -0.96463 | 2.74 |
| 1490 | *FOXRED2* | 5.2565 | 2.2307 | 0.73007 | 0.25632 | 2.74 |
| 1491 | *TMEM196* | 3.99 | 3.6065 | 0.61939 | -0.32537 | 2.74 |
| 1492 | *MRPL15* | 3.8726 | 3.403 | 0.94 | -1.7592 | 2.74 |
| 1493 | *EGFR* | 3.8422 | 3.6305 | 0.74126 | -0.26711 | 2.74 |
| 1494 | *REEP5* | 4.2373 | 2.186 | 1.7897 | 0.37134 | 2.74 |
| 1495 | *CYS1* | 2.9117 | 2.7733 | 2.526 | 0.40961 | 2.74 |
| 1496 | *TECPR1* | 3.9643 | 3.1865 | 1.06 | 0.30312 | 2.74 |
| 1497 | *LOXL1* | 4.1872 | 4.1767 | -0.15385 | -0.35694 | 2.74 |
| 1498 | *OGG1* | 3.9137 | 3.7554 | 0.54046 | -0.66604 | 2.74 |
| 1499 | *BBC3* | 5.0448 | 2.0652 | 1.0983 | -0.89593 | 2.74 |
| 1500 | *TMEM74B* | 5.0654 | 2.6504 | 0.49197 | -1.4529 | 2.74 |
| 1501 | *KIF13A* | 3.7945 | 2.9378 | 1.475 | -0.59025 | 2.74 |
| 1502 | *CRYGD* | 5.5225 | 4.3463 | -1.6635 | -2.4157 | 2.74 |
| 1503 | *CCR1* | 3.4802 | 3.4542 | 1.2691 | -1.5876 | 2.73 |
| 1504 | *SRF* | 4.42 | 2.4434 | 1.3392 | -1.9252 | 2.73 |
| 1505 | *PRSS36* | 3.563 | 3.4453 | 1.1936 | -0.95573 | 2.73 |
| 1506 | *QSOX1* | 4.2678 | 3.1164 | 0.81744 | -1.638 | 2.73 |
| 1507 | *CEP192* | 3.8976 | 2.8897 | 1.4143 | -2.1445 | 2.73 |
| 1508 | *DPCD* | 4.4906 | 2.5808 | 1.1294 | -0.1855 | 2.73 |
| 1509 | *FCRL2* | 4.3651 | 3.8576 | -0.0249 | -0.09906 | 2.73 |
| 1510 | *MTMR14* | 4.3312 | 2.5675 | 1.2956 | -0.56098 | 2.73 |
| 1511 | *DPPA5* | 5.1715 | 2.2434 | 0.77879 | -0.60065 | 2.73 |

| | | | | | | |
|---|---|---|---|---|---|---|
| 1512 | *ANP32E* | 3.8983 | 2.9283 | 1.3666 | -1.7049 | 2.73 |
| 1513 | *CLEC16A* | 5.445 | 1.4434 | 1.3036 | -0.85551 | 2.73 |
| 1514 | *BSN* | 3.9011 | 3.3599 | 0.9297 | 0.65767 | 2.73 |
| 1515 | *TTLL3* | 5.185 | 2.7836 | 0.22099 | -0.85739 | 2.73 |
| 1516 | *ZNF799* | 3.6319 | 2.4586 | 2.0975 | -1.8484 | 2.73 |
| 1517 | *NMRK1* | 4.2095 | 4.0311 | -0.05387 | -0.55359 | 2.73 |
| 1518 | *CHRNA7* | 4.285 | 2.482 | 1.4192 | 0.80859 | 2.73 |
| 1519 | *MTO1* | 4.1626 | 3.4753 | 0.5469 | -1.6711 | 2.73 |
| 1520 | *SUV39H2* | 4.9112 | 2.3641 | 0.90894 | -1.4597 | 2.73 |
| 1521 | *PCOLCE2* | 3.7377 | 2.7833 | 1.6605 | -1.0028 | 2.73 |
| 1522 | *SUMO4* | 4.9533 | 1.8204 | 1.4074 | -1.0932 | 2.73 |
| 1523 | *TRAPPC13* | 4.4319 | 3.0784 | 0.66958 | 0.27172 | 2.73 |
| 1524 | *OR11L1* | 3.3798 | 3.271 | 1.5239 | -1.0373 | 2.72 |
| 1525 | *RIOK1* | 5.1222 | 2.9546 | 0.09624 | -1.7268 | 2.72 |
| 1526 | *TRIM24* | 4.5346 | 2.4808 | 1.1569 | 0.85373 | 2.72 |
| 1527 | *ESX1* | 4.0221 | 4.0174 | 0.12923 | -1.3387 | 2.72 |
| 1528 | *HK1* | 4.695 | 2.0095 | 1.4591 | -1.0529 | 2.72 |
| 1529 | *LPPR5* | 3.7576 | 2.8793 | 1.5244 | -1.1709 | 2.72 |
| 1530 | *ALPI* | 5.2302 | 3.4634 | -0.53332 | -1.1545 | 2.72 |
| 1531 | *PTH1R* | 4.5869 | 3.8687 | -0.29638 | -1.2277 | 2.72 |
| 1532 | *RHOBTB3* | 5.1454 | 2.3289 | 0.6828 | 0.35637 | 2.72 |
| 1533 | *CCDC106* | 5.2636 | 3.2218 | -0.32849 | -0.95036 | 2.72 |
| 1534 | *STK17B* | 4.9495 | 2.8009 | 0.40276 | -0.53254 | 2.72 |
| 1535 | *FMN1* | 3.7811 | 3.0507 | 1.3191 | 0.49269 | 2.72 |
| 1536 | *RAB23* | 4.2238 | 4.0326 | -0.10619 | -0.42361 | 2.72 |
| 1537 | *OR1G1* | 4.1745 | 3.0766 | 0.89846 | -0.73259 | 2.72 |
| 1538 | *KRTAP5-6* | 4.4308 | 3.7045 | 0.01374 | -0.26302 | 2.72 |
| 1539 | *GSTO2* | 3.5282 | 3.1812 | 1.4388 | 0.89403 | 2.72 |
| 1540 | *CNN1* | 3.539 | 2.8849 | 1.722 | -1.5672 | 2.72 |
| 1541 | *KCNK16* | 5.9073 | 2.7392 | -0.50062 | -0.72168 | 2.72 |
| 1542 | *FBXO32* | 4.103 | 2.0764 | 1.9655 | 0.26709 | 2.71 |
| 1543 | *CEACAM8* | 3.6445 | 3.4803 | 1.02 | 0.87156 | 2.71 |
| 1544 | *C12orf57* | 2.9718 | 2.6216 | 2.5493 | 2.2731 | 2.71 |
| 1545 | *AMPH* | 4.8977 | 1.9093 | 1.3351 | -1.5095 | 2.71 |
| 1546 | *OSBP* | 4.7544 | 2.7001 | 0.68723 | -0.54515 | 2.71 |
| 1547 | *STIL* | 3.9181 | 2.4127 | 1.8103 | -1.9563 | 2.71 |
| 1548 | *AGR2* | 4.8379 | 3.4284 | -0.12651 | -0.79149 | 2.71 |
| 1549 | *C8orf4* | 3.9493 | 2.6492 | 1.5411 | -2.1983 | 2.71 |
| 1550 | *TOR2A* | 5.3211 | 1.892 | 0.92478 | -1.6741 | 2.71 |
| 1551 | *VCX3A* | 4.2343 | 3.7514 | 0.1517 | -0.89474 | 2.71 |
| 1552 | *TSPYL1* | 4.8135 | 3.2261 | 0.09568 | -0.19842 | 2.71 |
| 1553 | *GJB1* | 4.4758 | 3.0234 | 0.6356 | -1.4473 | 2.71 |
| 1554 | *AGAP3* | 3.5723 | 2.5575 | 2.0049 | -1.1964 | 2.71 |
| 1555 | *GOLGA6L2* | 3.612 | 2.551 | 1.9699 | 1.848 | 2.71 |
| 1556 | *PCSK1* | 3.1594 | 3.0362 | 1.9372 | 0.88166 | 2.71 |
| 1557 | *SPDYE4* | 4.033 | 3.2424 | 0.85729 | -0.71393 | 2.71 |

| | | | | | | |
|---|---|---|---|---|---|---|
| 1558 | *RER1* | 3.039 | 2.8069 | 2.283 | -0.49155 | 2.71 |
| 1559 | *OR8K1* | 3.7137 | 3.275 | 1.1385 | -1.5984 | 2.71 |
| 1560 | *MYO3B* | 4.0381 | 3.0398 | 1.0431 | 0.28497 | 2.71 |
| 1561 | *SLC17A6* | 4.5082 | 2.2162 | 1.3946 | -0.58274 | 2.71 |
| 1562 | *SEC24B* | 3.8517 | 2.6013 | 1.6659 | -0.79149 | 2.71 |
| 1563 | *CPLX2* | 3.6799 | 2.9398 | 1.4983 | 1.1743 | 2.71 |
| 1564 | *TMEM106A* | 4.1249 | 3.7567 | 0.23424 | -1.7725 | 2.71 |
| 1565 | *CDC169-SOHLH2* | 4.727 | 3.4397 | -0.05133 | -1.4142 | 2.71 |
| 1566 | *NPB* | 3.5152 | 2.9454 | 1.6492 | 1.2316 | 2.70 |
| 1567 | *C8orf59* | 3.7162 | 2.8708 | 1.522 | -0.87997 | 2.70 |
| 1568 | *NLGN1* | 4.2287 | 3.1332 | 0.74684 | -2.7442 | 2.70 |
| 1569 | *CBS* | 3.897 | 2.216 | 1.9952 | -1.6796 | 2.70 |
| 1570 | *NIF3L1* | 2.9967 | 2.6109 | 2.4979 | -0.95505 | 2.70 |
| 1571 | *MORF4L1* | 3.6808 | 2.6065 | 1.8169 | -0.63097 | 2.70 |
| 1572 | *TBC1D19* | 3.3486 | 2.8889 | 1.8658 | -2.0267 | 2.70 |
| 1573 | *TSTD1* | 5.2369 | 3.1699 | -0.30424 | -1.8154 | 2.70 |
| 1574 | *ESPL1* | 3.1387 | 2.5246 | 2.4385 | 2.0758 | 2.70 |
| 1575 | *WDR38* | 4.9502 | 4.4774 | -1.3288 | -2.0574 | 2.70 |
| 1576 | *CP* | 5.0152 | 2.4516 | 0.63111 | 0.57403 | 2.70 |
| 1577 | *SLC25A11* | 3.9664 | 3.7502 | 0.38081 | -0.99892 | 2.70 |
| 1578 | *FMN2* | 3.3374 | 3.3361 | 1.4226 | -0.57766 | 2.70 |
| 1579 | *NUTM2G* | 3.4662 | 3.4467 | 1.1821 | -1.4747 | 2.70 |
| 1580 | *TCEAL3* | 3.7576 | 2.2081 | 2.129 | -0.32811 | 2.70 |
| 1581 | *PDCL* | 3.8831 | 3.5477 | 0.66088 | -1.1346 | 2.70 |
| 1582 | *SLAMF6* | 4.471 | 3.3615 | 0.25714 | -0.69972 | 2.70 |
| 1583 | *ZNF33A* | 3.7272 | 3.4188 | 0.94361 | 0.65601 | 2.70 |
| 1584 | *GCLC* | 4.1087 | 4.034 | -0.05369 | -1.1501 | 2.70 |
| 1585 | *IFT20* | 4.2822 | 3.1235 | 0.68273 | -0.25605 | 2.70 |
| 1586 | *DMD* | 3.3774 | 2.4249 | 2.2834 | 0.58032 | 2.70 |
| 1587 | *FGFR1OP* | 2.9978 | 2.7283 | 2.3586 | 0.33236 | 2.69 |
| 1588 | *B3GNT9* | 4.6023 | 2.7449 | 0.73377 | -0.02412 | 2.69 |
| 1589 | *FAM19A1* | 4.1094 | 3.4239 | 0.5469 | -1.2437 | 2.69 |
| 1590 | *MLKL* | 4.5947 | 3.4841 | -0.00032 | -1.0832 | 2.69 |
| 1591 | *SOCS3* | 4.4225 | 2.8245 | 0.83078 | -0.71645 | 2.69 |
| 1592 | *CNTNAP2* | 5.1814 | 1.4899 | 1.4028 | -1.2455 | 2.69 |
| 1593 | *OR5H6* | 5.1603 | 4.0801 | -1.1665 | -1.843 | 2.69 |
| 1594 | *ZDHHC24* | 4.285 | 4.0031 | -0.21434 | -0.23105 | 2.69 |
| 1595 | *CORO1B* | 4.453 | 3.9732 | -0.35367 | -1.3161 | 2.69 |
| 1596 | *CST4* | 4.1266 | 3.3093 | 0.63448 | 0.15099 | 2.69 |
| 1597 | *UST* | 3.7505 | 2.5884 | 1.7292 | -1.714 | 2.69 |
| 1598 | *OR4C15* | 3.7677 | 3.007 | 1.2927 | -1.7438 | 2.69 |
| 1599 | *ERAP1* | 3.6035 | 3.5743 | 0.88735 | -0.3791 | 2.69 |
| 1600 | *LRRC37A3* | 4.1083 | 3.4173 | 0.53898 | -1.7994 | 2.69 |
| 1601 | *HNRNPH2* | 4.4353 | 2.3267 | 1.3021 | -0.47407 | 2.69 |
| 1602 | *SRSF9* | 3.6116 | 3.1241 | 1.3251 | 0.75933 | 2.69 |
| 1603 | *CYBRD1* | 4.3609 | 1.8669 | 1.8323 | -0.25493 | 2.69 |

| | | | | | | |
|---|---|---|---|---|---|---|
| 1604 | *LY6G5B* | 4.8234 | 1.6592 | 1.5761 | -0.30692 | 2.69 |
| 1605 | *MS4A3* | 5.0366 | 3.8677 | -0.84652 | -1.0914 | 2.69 |
| 1606 | *KANSL1L* | 4.058 | 2.241 | 1.7573 | 0.2293 | 2.69 |
| 1607 | *HIST4H4* | 4.1547 | 2.4565 | 1.4421 | -0.22269 | 2.68 |
| 1608 | *GPRASP2* | 4.2502 | 3.4073 | 0.39196 | -1.3784 | 2.68 |
| 1609 | *HES3* | 4.3011 | 2.3552 | 1.392 | 0.29116 | 2.68 |
| 1610 | *ITM2B* | 4.7955 | 3.6117 | -0.36057 | -0.4139 | 2.68 |
| 1611 | *SMPDL3A* | 3.9434 | 2.9168 | 1.1858 | 0.1919 | 2.68 |
| 1612 | *CCDC144A* | 5.3128 | 2.0264 | 0.70596 | -1.5635 | 2.68 |
| 1613 | *SHMT2* | 4.1292 | 3.188 | 0.7269 | -0.61132 | 2.68 |
| 1614 | *CCDC14* | 4.5502 | 4.4718 | -0.98062 | -1.8016 | 2.68 |
| 1615 | *SEPT14* | 4.2525 | 3.8011 | -0.01245 | -0.95722 | 2.68 |
| 1616 | *ITGA5* | 4.1672 | 3.0995 | 0.7744 | -1.3533 | 2.68 |
| 1617 | *KIF5B* | 4.5299 | 2.6159 | 0.89528 | -1.1077 | 2.68 |
| 1618 | *DDX21* | 5.0098 | 3.6902 | -0.66008 | -1.3795 | 2.68 |
| 1619 | *ZDHHC1* | 5.7836 | 1.6347 | 0.6216 | -1.7991 | 2.68 |
| 1620 | *SYCE3* | 3.9501 | 3.1578 | 0.9318 | -1.671 | 2.68 |
| 1621 | *SULT1B1* | 4.7505 | 2.9398 | 0.34851 | -0.80754 | 2.68 |
| 1622 | *ARL5B* | 5.4221 | 4.406 | -1.7945 | -2.4322 | 2.68 |
| 1623 | *CHP2* | 3.8619 | 3.6458 | 0.52471 | -2.9417 | 2.68 |
| 1624 | *HDHD1* | 3.5105 | 2.3061 | 2.2143 | 2.0325 | 2.68 |
| 1625 | *CERS3* | 5.3852 | 3.5162 | -0.87208 | -1.2193 | 2.68 |
| 1626 | *C7orf65* | 3.3816 | 2.5119 | 2.1357 | 0.91529 | 2.68 |
| 1627 | *PRH2* | 4.9843 | 2.5057 | 0.53795 | -0.14019 | 2.68 |
| 1628 | *NUP62CL* | 3.4121 | 2.5797 | 2.0338 | 0.34077 | 2.68 |
| 1629 | *TRIB1* | 6.0406 | 3.2836 | -1.2988 | -3.7834 | 2.68 |
| 1630 | *PHLDB3* | 4.7924 | 3.5077 | -0.27663 | -2.2809 | 2.67 |
| 1631 | *OR8K3* | 4.9193 | 2.0665 | 1.0327 | -0.67235 | 2.67 |
| 1632 | *RIPK2* | 3.643 | 3.4346 | 0.94035 | -1.061 | 2.67 |
| 1633 | *TRIM49* | 3.6981 | 2.5922 | 1.7274 | -2.1239 | 2.67 |
| 1634 | *THAP4* | 4.1282 | 2.9554 | 0.93222 | 0.57874 | 2.67 |
| 1635 | *ZNF430* | 3.7983 | 2.752 | 1.4627 | -3.2089 | 2.67 |
| 1636 | *SEC16B* | 4.5208 | 3.1145 | 0.37723 | -0.70187 | 2.67 |
| 1637 | *MAS1* | 4.9824 | 2.2457 | 0.78415 | 0.54456 | 2.67 |
| 1638 | *LHX9* | 3.2969 | 3.203 | 1.508 | -0.6501 | 2.67 |
| 1639 | *TSPAN6* | 5.1156 | 4.3748 | -1.4842 | -2.2669 | 2.67 |
| 1640 | *C1QTNF6* | 4.5192 | 2.5793 | 0.9075 | -1.5375 | 2.67 |
| 1641 | *TERF2IP* | 3.8445 | 2.9909 | 1.1701 | -1.6347 | 2.67 |
| 1642 | *AEBP1* | 4.7014 | 2.5521 | 0.75163 | -0.80248 | 2.67 |
| 1643 | *CLEC2A* | 2.9928 | 2.9647 | 2.0474 | 0.7832 | 2.67 |
| 1644 | *ERI1* | 4.1647 | 2.8185 | 1.0214 | -3.4433 | 2.67 |
| 1645 | *MAML2* | 3.769 | 3.1421 | 1.0927 | -0.24488 | 2.67 |
| 1646 | *NME1* | 3.9996 | 3.4122 | 0.58825 | -0.8303 | 2.67 |
| 1647 | *RUNX1T1* | 4.101 | 4.0673 | -0.1685 | -1.5304 | 2.67 |
| 1648 | *SEMA3B* | 4.5208 | 4.2266 | -0.74815 | -1.35 | 2.67 |
| 1649 | *TOR3A* | 4.8939 | 3.8166 | -0.71268 | -0.76965 | 2.67 |

| | | | | | | |
|---|---|---|---|---|---|---|
| 1650 | *KCNJ16* | 2.9869 | 2.7383 | 2.2718 | 0.86274 | 2.67 |
| 1651 | *SPANXB1* | 4.3997 | 3.1733 | 0.42339 | -0.95068 | 2.67 |
| 1652 | *TUBA8* | 4.3505 | 3.2511 | 0.39392 | 0.29569 | 2.67 |
| 1653 | *MKRN2OS* | 4.0344 | 3.6364 | 0.3228 | -2.5048 | 2.66 |
| 1654 | *TMEM225* | 3.9988 | 3.0529 | 0.94015 | -1.6027 | 2.66 |
| 1655 | *VASH2* | 3.0373 | 2.7749 | 2.1771 | 0.65861 | 2.66 |
| 1656 | *DDRGK1* | 4.7775 | 4.3161 | -1.1053 | -1.2989 | 2.66 |
| 1657 | *SIN3B* | 4.1192 | 3.8951 | -0.02602 | -1.0175 | 2.66 |
| 1658 | *TMPRSS11D* | 3.8107 | 3.216 | 0.96023 | -0.61839 | 2.66 |
| 1659 | *LMO7* | 4.6597 | 2.707 | 0.6201 | -1.6907 | 2.66 |
| 1660 | *GAK* | 3.2293 | 3.2058 | 1.5511 | -0.52825 | 2.66 |
| 1661 | *RIBC1* | 3.1952 | 2.6694 | 2.1205 | 0.59638 | 2.66 |
| 1662 | *ANKFN1* | 4.5797 | 2.0095 | 1.3956 | 0.42942 | 2.66 |
| 1663 | *FBXO18* | 4.0969 | 2.2804 | 1.6048 | 0.54427 | 2.66 |
| 1664 | *OR1J1* | 4.155 | 2.0908 | 1.735 | -0.33524 | 2.66 |
| 1665 | *PPP2R5C* | 5.2502 | 1.7584 | 0.96984 | 0.5847 | 2.66 |
| 1666 | *C4A* | 4.4247 | 2.835 | 0.71813 | -1.4685 | 2.66 |
| 1667 | *HOXA6* | 4.0024 | 2.7405 | 1.2346 | -0.3306 | 2.66 |
| 1668 | *TMED8* | 4.3429 | 3.6063 | 0.02805 | -0.00832 | 2.66 |
| 1669 | *MTSS1L* | 4.1858 | 2.8558 | 0.93345 | -0.43974 | 2.66 |
| 1670 | *RWDD1* | 3.9609 | 3.078 | 0.93552 | -0.68144 | 2.66 |
| 1671 | *RBM12B* | 3.8843 | 2.8992 | 1.1908 | -1.2537 | 2.66 |
| 1672 | *MAPK15* | 3.5727 | 3.4714 | 0.9268 | -2.3577 | 2.66 |
| 1673 | *MYO1H* | 4.9682 | 3.8145 | -0.81193 | -2.3602 | 2.66 |
| 1674 | *ANKRD1* | 4.8209 | 2.8666 | 0.28169 | -1.0005 | 2.66 |
| 1675 | *MAP7D1* | 3.3976 | 2.7709 | 1.7991 | 1.3904 | 2.66 |
| 1676 | *SMIM19* | 3.5841 | 3.3383 | 1.045 | -0.32779 | 2.66 |
| 1677 | *DHRS4* | 3.954 | 3.4574 | 0.55396 | -1.0458 | 2.66 |
| 1678 | *KCNMB3* | 3.8579 | 2.0986 | 2.007 | -0.27761 | 2.65 |
| 1679 | *TSPAN4* | 3.9529 | 2.308 | 1.7018 | 0.51807 | 2.65 |
| 1680 | *HDAC4* | 3.3499 | 2.6552 | 1.9553 | -0.0398 | 2.65 |
| 1681 | *FHDC1* | 4.3348 | 2.6154 | 1.0095 | 0.73531 | 2.65 |
| 1682 | *SYT9* | 3.3168 | 2.3933 | 2.2493 | -0.31411 | 2.65 |
| 1683 | *GABRD* | 3.784 | 2.629 | 1.5462 | 0.78273 | 2.65 |
| 1684 | *BHMT2* | 3.5341 | 3.1847 | 1.2391 | -1.6076 | 2.65 |
| 1685 | *SEMA3C* | 5.4727 | 1.9377 | 0.54647 | -1.1027 | 2.65 |
| 1686 | *SCD5* | 4.9258 | 3.0008 | 0.02904 | -0.66635 | 2.65 |
| 1687 | *TNRC6B* | 4.3799 | 3.3461 | 0.22814 | -0.02683 | 2.65 |
| 1688 | *ZBTB25* | 4.1303 | 3.1789 | 0.64197 | -0.61963 | 2.65 |
| 1689 | *MTCP1* | 4.4978 | 4.0091 | -0.5567 | -1.1156 | 2.65 |
| 1690 | *OR2F2* | 2.6867 | 2.6601 | 2.6033 | -1.2305 | 2.65 |
| 1691 | *MTERF4* | 3.7198 | 3.3823 | 0.8464 | -1.5848 | 2.65 |
| 1692 | *WISP2* | 4.8921 | 2.21 | 0.84594 | -0.65971 | 2.65 |
| 1693 | *DCBLD2* | 4.0081 | 2.2126 | 1.7271 | -0.42573 | 2.65 |
| 1694 | *SMIM10* | 4.0028 | 2.6472 | 1.2977 | -0.93967 | 2.65 |
| 1695 | *TBC1D1* | 5.6843 | 1.7094 | 0.55371 | -1.8022 | 2.65 |

| | | | | | | |
|---|---|---|---|---|---|---|
| 1696 | *STPG1* | 3.1955 | 2.6093 | 2.1391 | -0.9382 | 2.65 |
| 1697 | *SAA2-SAA4* | 4.669 | 3.4803 | -0.20721 | -0.3433 | 2.65 |
| 1698 | *EXPH5* | 4.3615 | 2.5968 | 0.98351 | 0.34821 | 2.65 |
| 1699 | *PYROXD1* | 3.8075 | 3.034 | 1.098 | -0.95701 | 2.65 |
| 1700 | *HAS1* | 4.4024 | 2.9489 | 0.58535 | -1.5237 | 2.65 |
| 1701 | *PRR29* | 4.272 | 3.4769 | 0.18588 | -0.27651 | 2.64 |
| 1702 | *BCL9* | 4.968 | 2.7429 | 0.22243 | -1.0564 | 2.64 |
| 1703 | *S1PR1* | 4.2646 | 2.1768 | 1.4905 | -1.4584 | 2.64 |
| 1704 | *TMEM217* | 4.2238 | 3.3874 | 0.31759 | -0.95918 | 2.64 |
| 1705 | *CBLN3* | 3.9871 | 2.167 | 1.7738 | -2.3452 | 2.64 |
| 1706 | *ANKRD62* | 3.1361 | 2.5959 | 2.1953 | -0.41835 | 2.64 |
| 1707 | *HMGXB4* | 4.1857 | 3.8203 | -0.07895 | -1.1995 | 2.64 |
| 1708 | *GDNF-AS1* | 4.2777 | 2.9267 | 0.72251 | -1.7766 | 2.64 |
| 1709 | *CPXM1* | 4.3461 | 2.8487 | 0.73137 | -0.1136 | 2.64 |
| 1710 | *G0S2* | 4.6971 | 1.6889 | 1.5393 | 1.4797 | 2.64 |
| 1711 | *HCAR3* | 4.267 | 2.9982 | 0.65987 | -0.19423 | 2.64 |
| 1712 | *TMEM256* | 4.4791 | 2.075 | 1.3692 | -1.1201 | 2.64 |
| 1713 | *TMEM183B* | 4.2143 | 3.2844 | 0.42153 | -0.76169 | 2.64 |
| 1714 | *OR2A42* | 3.1914 | 3.1512 | 1.5758 | 0.77768 | 2.64 |
| 1715 | *SLC9C1* | 4.1882 | 3.4664 | 0.2614 | 0.23329 | 2.64 |
| 1716 | *C5orf52* | 3.8356 | 3.1449 | 0.93523 | -1.3176 | 2.64 |
| 1717 | *SLC35G1* | 2.993 | 2.605 | 2.3176 | -0.34998 | 2.64 |
| 1718 | *RDH14* | 4.2936 | 2.7264 | 0.89402 | 0.80623 | 2.64 |
| 1719 | *KLK15* | 3.5436 | 2.7933 | 1.5771 | -1.5663 | 2.64 |
| 1720 | *C5orf30* | 4.3914 | 3.1386 | 0.38066 | -0.4102 | 2.64 |
| 1721 | *PTCHD2* | 4.8963 | 2.5491 | 0.46373 | -0.78439 | 2.64 |
| 1722 | *GBP4* | 3.6761 | 3.2671 | 0.96447 | -0.12601 | 2.64 |
| 1723 | *GGTLC1* | 4.4604 | 2.6037 | 0.84349 | 0.036324 | 2.64 |
| 1724 | *DLGAP5* | 4.261 | 2.8667 | 0.77594 | -1.1027 | 2.63 |
| 1725 | *GABRR1* | 3.5613 | 3.1625 | 1.1795 | -0.4158 | 2.63 |
| 1726 | *SH3BP2* | 4.3474 | 3.2772 | 0.27699 | -2.3696 | 2.63 |
| 1727 | *FHOD1* | 4.3457 | 3.0595 | 0.4959 | 0.12927 | 2.63 |
| 1728 | *GTF2A1L* | 4.8502 | 3.4993 | -0.4488 | -1.5383 | 2.63 |
| 1729 | *NINJ1* | 3.8909 | 3.5481 | 0.46124 | -1.7757 | 2.63 |
| 1730 | *FMNL3* | 3.9127 | 3.9017 | 0.08381 | -0.59061 | 2.63 |
| 1731 | *HTRA2* | 4.9849 | 3.1897 | -0.2765 | -1.6417 | 2.63 |
| 1732 | *ZNF716* | 4.1027 | 2.5679 | 1.2274 | 0.97075 | 2.63 |
| 1733 | *HES7* | 3.7173 | 3.6432 | 0.5365 | -0.89866 | 2.63 |
| 1734 | *ATOH8* | 3.9522 | 3.1866 | 0.75794 | 0.41567 | 2.63 |
| 1735 | *GTDC1* | 4.7159 | 2.9041 | 0.27454 | 0.098257 | 2.63 |
| 1736 | *IL5RA* | 4.2272 | 2.5294 | 1.1371 | -0.52968 | 2.63 |
| 1737 | *IGFBP1* | 5.4159 | 3.0276 | -0.55184 | -1.8884 | 2.63 |
| 1738 | *CATSPERG* | 4.9844 | 1.824 | 1.0823 | -0.54786 | 2.63 |
| 1739 | *CRYBB2* | 4.1271 | 3.7484 | 0.01415 | -1.3287 | 2.63 |
| 1740 | *PICK1* | 4.5417 | 2.6268 | 0.71885 | -1.5635 | 2.63 |
| 1741 | *TUBGCP3* | 5.1924 | 3.3469 | -0.65198 | -1.6877 | 2.63 |

| | | | | | | |
|---|---|---|---|---|---|---|
| 1742 | *ZNF705B* | 3.5275 | 3.1235 | 1.236 | -1.0599 | 2.63 |
| 1743 | *PGLYRP2* | 3.6503 | 3.3419 | 0.89416 | -0.22378 | 2.63 |
| 1744 | *ITGA2B* | 4.2186 | 2.8224 | 0.84507 | 0.23912 | 2.63 |
| 1745 | *FOXQ1* | 5.0543 | 3.439 | -0.60752 | -1.3042 | 2.63 |
| 1746 | *LRRC3B* | 3.9608 | 3.7256 | 0.19646 | -1.4192 | 2.63 |
| 1747 | *IL1RAPL2* | 3.6931 | 2.7228 | 1.4657 | -1.0477 | 2.63 |
| 1748 | *OR4K5* | 3.3688 | 2.5943 | 1.9181 | -0.89384 | 2.63 |
| 1749 | *OR5C1* | 4.2067 | 2.6827 | 0.99069 | -1.0221 | 2.63 |
| 1750 | *C15orf48* | 5.7195 | 3.3734 | -1.2152 | -1.5059 | 2.63 |
| 1751 | *CFAP69* | 4.0608 | 2.5086 | 1.308 | -0.71526 | 2.63 |
| 1752 | *CRP* | 3.5432 | 2.3377 | 1.9964 | -0.71363 | 2.63 |
| 1753 | *ZNF585B* | 3.4493 | 3.1472 | 1.2774 | 0.25994 | 2.62 |
| 1754 | *IGFBPL1* | 4.4856 | 1.8195 | 1.5686 | -1.1585 | 2.62 |
| 1755 | *FAM72D* | 4.3293 | 1.9205 | 1.6232 | -1.7257 | 2.62 |
| 1756 | *CEACAM21* | 5.443 | 1.4698 | 0.9562 | 0.46987 | 2.62 |
| 1757 | *PIP5K1A* | 5.3874 | 2.1508 | 0.32923 | -0.33638 | 2.62 |
| 1758 | *OR52I2* | 4.6763 | 3.2668 | -0.07644 | -0.95802 | 2.62 |
| 1759 | *FAM131B* | 4.2597 | 1.8218 | 1.784 | 0.4557 | 2.62 |
| 1760 | *TRMT61A* | 4.9555 | 4.0323 | -1.1223 | -1.3139 | 2.62 |
| 1761 | *SLC30A10* | 5.0545 | 1.9741 | 0.83488 | -0.99175 | 2.62 |
| 1762 | *SEC23A* | 4.4645 | 3.2579 | 0.13956 | -0.89294 | 2.62 |
| 1763 | *EEF1E1* | 4.0799 | 2.4685 | 1.3107 | -0.34111 | 2.62 |
| 1764 | *SLC17A7* | 4.135 | 3.221 | 0.5013 | -0.73979 | 2.62 |
| 1765 | *TMEM102* | 3.6347 | 2.9437 | 1.2787 | -0.55975 | 2.62 |
| 1766 | *VN1R5* | 4.3285 | 1.9882 | 1.5384 | -0.02451 | 2.62 |
| 1767 | *ITGB8* | 4.4635 | 3.4054 | -0.01399 | -0.14196 | 2.62 |
| 1768 | *GABRA5* | 3.9914 | 2.4297 | 1.4319 | 0.23824 | 2.62 |
| 1769 | *CCL1* | 4.8534 | 1.9002 | 1.0985 | -1.8462 | 2.62 |
| 1770 | *NCK1* | 5.456 | 1.6145 | 0.781 | -1.6597 | 2.62 |
| 1771 | *ANO6* | 5.3498 | 1.2887 | 1.2104 | 0.59175 | 2.62 |
| 1772 | *MAPK6* | 6.2181 | 2.4437 | -0.81329 | -1.0019 | 2.62 |
| 1773 | *SLC23A2* | 3.4779 | 2.6135 | 1.7571 | -1.2241 | 2.62 |
| 1774 | *SGCG* | 3.5633 | 2.8971 | 1.3879 | -0.17048 | 2.62 |
| 1775 | *TGFB1I1* | 5.0439 | 1.4604 | 1.3438 | 1.2807 | 2.62 |
| 1776 | *CITED2* | 2.8735 | 2.788 | 2.1848 | 1.0236 | 2.62 |
| 1777 | *BTAF1* | 3.6089 | 2.8233 | 1.4137 | -0.47864 | 2.62 |
| 1778 | *FKBP1B* | 2.9673 | 2.7617 | 2.1114 | 1.2803 | 2.61 |
| 1779 | *CHADL* | 5.7516 | 1.9147 | 0.17372 | -2.4172 | 2.61 |
| 1780 | *OR10S1* | 4.6259 | 3.6048 | -0.39313 | -0.62017 | 2.61 |
| 1781 | *NEDD4* | 5.217 | 2.5255 | 0.09472 | -0.76366 | 2.61 |
| 1782 | *INHBB* | 3.9925 | 2.7987 | 1.045 | -0.67691 | 2.61 |
| 1783 | *TM4SF4* | 4.3579 | 2.5115 | 0.96666 | -1.1576 | 2.61 |
| 1784 | *ST7* | 3.1981 | 2.9024 | 1.7353 | -0.35699 | 2.61 |
| 1785 | *CASP10* | 5.4336 | 2.0469 | 0.35343 | -1.7994 | 2.61 |
| 1786 | *C2orf91* | 3.8893 | 2.4834 | 1.4607 | -1.2169 | 2.61 |
| 1787 | *TFF1* | 3.2883 | 2.6976 | 1.8474 | -0.27194 | 2.61 |

| | | | | | | |
|---|---|---|---|---|---|---|
| 1788 | *BMP7* | 4.7022 | 2.3394 | 0.7902 | -0.63242 | 2.61 |
| 1789 | *OR6K2* | 4.6845 | 2.219 | 0.92709 | -1.3407 | 2.61 |
| 1790 | *SLC28A1* | 3.6067 | 3.5501 | 0.67222 | -0.91644 | 2.61 |
| 1791 | *CALHM1* | 3.2122 | 2.3136 | 2.303 | -0.28661 | 2.61 |
| 1792 | *NABP1* | 4.4656 | 2.332 | 1.0307 | -1.3174 | 2.61 |
| 1793 | *ZNF480* | 4.2711 | 3.9677 | -0.41123 | -0.89218 | 2.61 |
| 1794 | *KDM5A* | 3.5027 | 3.149 | 1.1757 | -1.047 | 2.61 |
| 1795 | *F8* | 3.841 | 3.2926 | 0.69327 | 0.16704 | 2.61 |
| 1796 | *B3GNT8* | 4.1337 | 2.9687 | 0.72251 | -1.227 | 2.61 |
| 1797 | *CELA1* | 2.9116 | 2.5894 | 2.3219 | -2.3904 | 2.61 |
| 1798 | *HOXA11* | 4.7559 | 3.2165 | -0.14969 | -0.17042 | 2.61 |
| 1799 | *SCG2* | 5.1785 | 1.4104 | 1.2325 | -1.5763 | 2.61 |
| 1800 | *LPXN* | 3.1008 | 2.4545 | 2.2647 | 0.59448 | 2.61 |
| 1801 | *PRKD2* | 5.096 | 2.8255 | -0.10176 | -0.84229 | 2.61 |
| 1802 | *C1orf64* | 4.5261 | 3.5826 | -0.28963 | -0.47665 | 2.61 |
| 1803 | *SCN2A* | 4.1085 | 1.8815 | 1.8288 | -0.99736 | 2.61 |
| 1804 | *CDK14* | 3.3357 | 2.7332 | 1.7496 | -1.0436 | 2.61 |
| 1805 | *PPIB* | 4.1971 | 2.9426 | 0.67721 | 0.031969 | 2.61 |
| 1806 | *ZDHHC19* | 3.5285 | 3.5233 | 0.76041 | -0.8929 | 2.60 |
| 1807 | *MRI1* | 4.8722 | 4.0551 | -1.1189 | -1.4258 | 2.60 |
| 1808 | *CYP7B1* | 4.1595 | 3.0533 | 0.59497 | -1.5582 | 2.60 |
| 1809 | *APOM* | 4.3194 | 3.8081 | -0.32061 | -0.43505 | 2.60 |
| 1810 | *PLA2G4D* | 5.1957 | 2.7235 | -0.11411 | -1.95 | 2.60 |
| 1811 | *GAPVD1* | 3.8074 | 3.3653 | 0.63031 | -1.2009 | 2.60 |
| 1812 | *HMG20B* | 3.5241 | 3.3617 | 0.91478 | 0.16024 | 2.60 |
| 1813 | *PPARD* | 3.1361 | 2.8832 | 1.7801 | -0.97064 | 2.60 |
| 1814 | *LRRC15* | 4.661 | 3.2831 | -0.1449 | -1.4682 | 2.60 |
| 1815 | *TMEM31* | 4.384 | 2.6835 | 0.72983 | -1.0654 | 2.60 |
| 1816 | *MIP* | 5.0714 | 3.251 | -0.52741 | -1.7589 | 2.60 |
| 1817 | *SYT17* | 4.2995 | 4.1237 | -0.62875 | -0.72674 | 2.60 |
| 1818 | *SLC10A5* | 4.8874 | 3.0877 | -0.18146 | -1.1223 | 2.60 |
| 1819 | *MBD2* | 3.8966 | 2.0117 | 1.8853 | 1.2066 | 2.60 |
| 1820 | *DPM2* | 4.6288 | 2.9057 | 0.25866 | -1.6755 | 2.60 |
| 1821 | *ST14* | 3.5716 | 3.1945 | 1.0267 | 0.12137 | 2.60 |
| 1822 | *SERTAD4* | 5.2015 | 3.5348 | -0.94583 | -1.9678 | 2.60 |
| 1823 | *KCNJ13* | 3.6584 | 2.313 | 1.819 | -1.3941 | 2.60 |
| 1824 | *PDE5A* | 4.0978 | 2.377 | 1.3147 | 0.72992 | 2.60 |
| 1825 | *VEPH1* | 4.179 | 3.6689 | -0.06095 | -1.7884 | 2.60 |
| 1826 | *LOX* | 3.7497 | 3.3495 | 0.6871 | -0.88839 | 2.60 |
| 1827 | *FAM219B* | 5.1274 | 1.5848 | 1.0721 | -1.3818 | 2.59 |
| 1828 | *SH2D4B* | 4.7057 | 2.7261 | 0.3521 | -1.3481 | 2.59 |
| 1829 | *ING2* | 4.7701 | 3.0474 | -0.0343 | -0.12189 | 2.59 |
| 1830 | *SLC26A9* | 4.149 | 3.8538 | -0.22131 | -0.63196 | 2.59 |
| 1831 | *ARHGEF3* | 4.0608 | 2.8935 | 0.82678 | -1.3371 | 2.59 |
| 1832 | *GBX2* | 3.5282 | 3.4063 | 0.8465 | 0.38164 | 2.59 |
| 1833 | *JAK2* | 3.6571 | 2.2608 | 1.8625 | -1.7852 | 2.59 |

| | | | | | | |
|---|---|---|---|---|---|---|
| 1834 | *ZNF34* | 4.1757 | 1.9293 | 1.6744 | 1.1214 | 2.59 |
| 1835 | *TINAG* | 4.1593 | 3.2373 | 0.38198 | -0.30293 | 2.59 |
| 1836 | *RPS6KA2* | 3.6549 | 3.0018 | 1.1204 | -0.60162 | 2.59 |
| 1837 | *TMEM27* | 2.8407 | 2.6821 | 2.254 | 1.0859 | 2.59 |
| 1838 | *FAM43A* | 4.1779 | 3.3179 | 0.27973 | -0.85471 | 2.59 |
| 1839 | *CES1* | 4.1579 | 2.5533 | 1.0642 | -1.2301 | 2.59 |
| 1840 | *EDRF1* | 4.6873 | 2.9101 | 0.17513 | -0.61767 | 2.59 |
| 1841 | *TIMM8B* | 3.8445 | 2.8041 | 1.1222 | 0.020906 | 2.59 |
| 1842 | *AHNAK2* | 4.768 | 4.5903 | -1.5894 | -2.1709 | 2.59 |
| 1843 | *CBX2* | 5.2224 | 3.2554 | -0.70976 | -1.3472 | 2.59 |
| 1844 | *ZNF578* | 4.6926 | 3.7865 | -0.71291 | -1.439 | 2.59 |
| 1845 | *GPN3* | 4.7027 | 3.387 | -0.32378 | -0.44353 | 2.59 |
| 1846 | *TSPO* | 3.9515 | 2.8085 | 1.0055 | 0.78012 | 2.59 |
| 1847 | *TDRD3* | 5.5057 | 2.289 | -0.02993 | -1.882 | 2.59 |
| 1848 | *ZNF621* | 3.3535 | 2.4055 | 2.0052 | -1.3978 | 2.59 |
| 1849 | *TACO1* | 3.7508 | 3.0538 | 0.95925 | -0.74442 | 2.59 |
| 1850 | *P2RY8* | 4.3209 | 1.844 | 1.5988 | 0.56084 | 2.59 |
| 1851 | *LYRM9* | 4.0407 | 4.0108 | -0.28823 | -1.3184 | 2.59 |
| 1852 | *FSTL3* | 3.9687 | 2.5088 | 1.2852 | -0.86694 | 2.59 |
| 1853 | *CLMN* | 4.722 | 2.4538 | 0.58443 | 0.49493 | 2.59 |
| 1854 | *TSPEAR* | 3.9947 | 2.5262 | 1.2379 | -0.64164 | 2.59 |
| 1855 | *NFE2* | 3.3319 | 2.8639 | 1.562 | -0.76074 | 2.59 |
| 1856 | *FER* | 4.4597 | 3.9499 | -0.65389 | -0.70927 | 2.59 |
| 1857 | *MT3* | 3.5559 | 2.4867 | 1.7118 | -1.8975 | 2.58 |
| 1858 | *RARRES2* | 3.0903 | 2.4314 | 2.2326 | 1.0804 | 2.58 |
| 1859 | *TOP2B* | 4.2924 | 2.5446 | 0.91584 | 0.14574 | 2.58 |
| 1860 | *ADGRD1* | 4.9427 | 3.2515 | -0.44353 | -1.6525 | 2.58 |
| 1861 | *PFN1* | 3.5824 | 2.8903 | 1.2775 | -0.52601 | 2.58 |
| 1862 | *MND1* | 4.1867 | 3.7854 | -0.222 | -1.2342 | 2.58 |
| 1863 | *OR10A4* | 3.9604 | 2.9224 | 0.86725 | -2.1745 | 2.58 |
| 1864 | *CTBP2* | 4.8391 | 3.1671 | -0.25827 | -1.1621 | 2.58 |
| 1865 | *MTNR1A* | 5.6214 | 2.6353 | -0.50978 | -1.1605 | 2.58 |
| 1866 | *KRTAP10-4* | 5.0187 | 4.4516 | -1.7247 | -1.8714 | 2.58 |
| 1867 | *LIPT1* | 4.7148 | 2.1237 | 0.90532 | 0.10522 | 2.58 |
| 1868 | *SYT10* | 4.4514 | 2.431 | 0.85976 | -0.77241 | 2.58 |
| 1869 | *PGC* | 3.9413 | 1.9778 | 1.8213 | 1.7266 | 2.58 |
| 1870 | *CA9* | 3.712 | 2.6392 | 1.3869 | -0.95322 | 2.58 |
| 1871 | *HS3ST6* | 3.6363 | 3.4971 | 0.60406 | -1.116 | 2.58 |
| 1872 | *SIAH3* | 4.1282 | 2.3204 | 1.2875 | 0.63161 | 2.58 |
| 1873 | *ANKS1B* | 2.7074 | 2.6755 | 2.3521 | -0.60183 | 2.58 |
| 1874 | *PARP11* | 5.3886 | 2.0455 | 0.29711 | -0.79711 | 2.58 |
| 1875 | *MAP3K14* | 3.3517 | 3.2927 | 1.0848 | -0.22309 | 2.58 |
| 1876 | *SMPDL3B* | 4.9176 | 1.755 | 1.0555 | -0.46265 | 2.58 |
| 1877 | *ZNF112* | 3.4584 | 3.2978 | 0.97132 | -0.04361 | 2.58 |
| 1878 | *TMEM202* | 4.8881 | 3.3658 | -0.52731 | -1.9673 | 2.58 |
| 1879 | *ZBTB49* | 3.9341 | 3.6209 | 0.17112 | -0.1134 | 2.58 |

| | | | | | | |
|---|---|---|---|---|---|---|
| 1880 | *SYCP1* | 3.5241 | 3.501 | 0.70084 | 0.12597 | 2.58 |
| 1881 | *TACC2* | 3.2226 | 2.4158 | 2.0866 | -1.7297 | 2.58 |
| 1882 | *PDGFRB* | 3.3231 | 3.287 | 1.1131 | -1.7075 | 2.57 |
| 1883 | *DENND5A* | 5.3165 | 3.2793 | -0.87482 | -1.5978 | 2.57 |
| 1884 | *SPO11* | 4.3337 | 3.2347 | 0.15236 | -0.65197 | 2.57 |
| 1885 | *AGR3* | 4.8076 | 3.7171 | -0.80442 | -1.2171 | 2.57 |
| 1886 | *RALBP1* | 3.4226 | 2.325 | 1.972 | 0.041724 | 2.57 |
| 1887 | *FAM46D* | 3.1701 | 2.9804 | 1.5685 | -0.42999 | 2.57 |
| 1888 | *ZNF530* | 5.1767 | 2.3744 | 0.16742 | -0.13028 | 2.57 |
| 1889 | *SPTA1* | 3.8708 | 3.467 | 0.37957 | -0.45048 | 2.57 |
| 1890 | *PPM1B* | 4.3415 | 3.3077 | 0.0669 | -0.63112 | 2.57 |
| 1891 | *SLC25A34* | 3.9763 | 2.2494 | 1.4886 | -1.9668 | 2.57 |
| 1892 | *TMEM233* | 4.1825 | 2.331 | 1.2007 | -0.75388 | 2.57 |
| 1893 | *ZNF181* | 3.9205 | 2.4259 | 1.3666 | 0.34162 | 2.57 |
| 1894 | *TRIQK* | 4.7955 | 2.1612 | 0.756 | -2.6768 | 2.57 |
| 1895 | *CD164L2* | 2.7893 | 2.6488 | 2.2735 | -2.0283 | 2.57 |
| 1896 | *OSBPL6* | 4.3644 | 2.6868 | 0.65699 | 0.030554 | 2.57 |
| 1897 | *UNC79* | 4.0667 | 2.2666 | 1.3744 | -0.20942 | 2.57 |
| 1898 | *NFATC3* | 3.7104 | 2.5428 | 1.4544 | -2.7379 | 2.57 |
| 1899 | *SAMD4A* | 3.6359 | 3.4798 | 0.59132 | -1.917 | 2.57 |
| 1900 | *DNASE2* | 4.3944 | 3.3814 | -0.07058 | -1.399 | 2.57 |
| 1901 | *CDH3* | 3.9932 | 2.3748 | 1.337 | -0.26161 | 2.57 |
| 1902 | *ZNF233* | 4.3186 | 3.3376 | 0.0473 | -0.02523 | 2.57 |
| 1903 | *PLA2G5* | 5.2431 | 1.9463 | 0.51379 | -2.3222 | 2.57 |
| 1904 | *FAM47E-STBD1* | 3.9185 | 2.562 | 1.2222 | -0.7623 | 2.57 |
| 1905 | *FHIT* | 4.8593 | 1.4679 | 1.3753 | -0.67461 | 2.57 |
| 1906 | *ADCY1* | 4.3225 | 3.9428 | -0.5628 | -1.7364 | 2.57 |
| 1907 | *KRTAP5-3* | 4.406 | 3.1577 | 0.13738 | -1.0903 | 2.57 |
| 1908 | *NIPAL2* | 3.7036 | 2.3597 | 1.6376 | -0.64458 | 2.57 |
| 1909 | *CCDC50* | 5.3058 | 1.5759 | 0.81481 | -0.91061 | 2.57 |
| 1910 | *CLDN7* | 5.0392 | 2.3358 | 0.32102 | -0.49941 | 2.57 |
| 1911 | *ZNF254* | 3.3928 | 2.3361 | 1.9651 | -0.86691 | 2.56 |
| 1912 | *LPAR6* | 3.4728 | 2.5976 | 1.6233 | -1.4151 | 2.56 |
| 1913 | *BEX1* | 4.3586 | 2.316 | 1.018 | -1.7644 | 2.56 |
| 1914 | *GRP* | 4.099 | 3.1376 | 0.45591 | -1.8038 | 2.56 |
| 1915 | *OR2V1* | 3.962 | 3.5718 | 0.15535 | -0.43203 | 2.56 |
| 1916 | *CDH15* | 3.6423 | 2.3283 | 1.7185 | 0.69415 | 2.56 |
| 1917 | *NAPB* | 4.2006 | 3.869 | -0.38384 | -0.96719 | 2.56 |
| 1918 | *CHM* | 5.0362 | 1.8321 | 0.81592 | -1.3296 | 2.56 |
| 1919 | *C15orf40* | 5.7123 | 3.2344 | -1.2628 | -1.7034 | 2.56 |
| 1920 | *SLITRK2* | 4.4612 | 2.8415 | 0.38094 | -0.46728 | 2.56 |
| 1921 | *ZMYND19* | 3.5057 | 2.3754 | 1.8008 | 0.22057 | 2.56 |
| 1922 | *DGUOK* | 4.8047 | 1.771 | 1.1045 | -3.2662 | 2.56 |
| 1923 | *GPR142* | 4.1496 | 3.7896 | -0.26009 | -1.1448 | 2.56 |
| 1924 | *GCDH* | 4.4241 | 2.2871 | 0.96721 | 0.38227 | 2.56 |
| 1925 | *ERICH2* | 3.516 | 2.5108 | 1.6486 | -2.2381 | 2.56 |

| | | | | | | |
|---|---|---|---|---|---|---|
| 1926 | *GPR101* | 4.8485 | 3.446 | -0.61967 | -1.6943 | 2.56 |
| 1927 | *FAM109A* | 4.8126 | 2.9538 | -0.09663 | -0.75889 | 2.56 |
| 1928 | *ARHGAP25* | 4.0223 | 3.6261 | 0.01846 | -0.01399 | 2.56 |
| 1929 | *DEFB4A* | 3.5517 | 2.0813 | 2.0333 | -0.44353 | 2.56 |
| 1930 | *OR1E2* | 3.5297 | 2.8373 | 1.2963 | -2.7486 | 2.55 |
| 1931 | *COQ10B* | 4.9825 | 1.7633 | 0.91724 | -1.5376 | 2.55 |
| 1932 | *FAM213B* | 4.5052 | 3.1123 | 0.04374 | -0.60465 | 2.55 |
| 1933 | *EPPIN* | 3.3438 | 2.663 | 1.6518 | -2.0622 | 2.55 |
| 1934 | *RNASE11* | 3.3287 | 2.7402 | 1.5893 | -0.79108 | 2.55 |
| 1935 | *DCUN1D5* | 4.154 | 2.5054 | 0.99866 | -1.7882 | 2.55 |
| 1936 | *TMPRSS13* | 3.4747 | 3.2924 | 0.88915 | -0.02187 | 2.55 |
| 1937 | *AFP* | 3.9351 | 2.9311 | 0.78924 | 0.7006 | 2.55 |
| 1938 | *TMEM229B* | 4.3745 | 2.9138 | 0.36684 | -0.94341 | 2.55 |
| 1939 | *PDCD10* | 3.143 | 2.8273 | 1.6839 | -0.25971 | 2.55 |
| 1940 | *RELB* | 4.7333 | 1.5467 | 1.3742 | -0.38248 | 2.55 |
| 1941 | *PFDN4* | 4.0319 | 2.4244 | 1.1946 | 0.27203 | 2.55 |
| 1942 | *CNR2* | 4.2222 | 3.9111 | -0.48652 | -2.1648 | 2.55 |
| 1943 | *IL4I1* | 4.1063 | 3.9751 | -0.43561 | -1.0178 | 2.55 |
| 1944 | *HCLS1* | 3.5644 | 2.4601 | 1.6211 | 0.11496 | 2.55 |
| 1945 | *ZNF567* | 3.7087 | 2.2297 | 1.7066 | -1.2858 | 2.55 |
| 1946 | *HMGB4* | 5.3437 | 1.3256 | 0.97516 | -3.3077 | 2.55 |
| 1947 | *PACSIN2* | 4.3814 | 3.6491 | -0.38664 | -0.84261 | 2.55 |
| 1948 | *SLC10A6* | 3.7065 | 2.9321 | 1.003 | -1.6906 | 2.55 |
| 1949 | *MTR* | 3.699 | 2.0729 | 1.8684 | -0.70697 | 2.55 |
| 1950 | *SMIM15* | 4.5909 | 1.8816 | 1.1669 | 0.51453 | 2.55 |
| 1951 | *RNF25* | 4.5336 | 3.138 | -0.03297 | -0.51785 | 2.55 |
| 1952 | *TUBAL3* | 6.2265 | 1.5375 | -0.1284 | -0.44353 | 2.55 |
| 1953 | *AIM2* | 4.8059 | 3.7277 | -0.90001 | -1.7434 | 2.54 |
| 1954 | *HMP19* | 3.5492 | 3.2066 | 0.87718 | -0.86506 | 2.54 |
| 1955 | *LCE3C* | 4.9742 | 1.7802 | 0.87831 | -1.1553 | 2.54 |
| 1956 | *CCPG1* | 4.3058 | 3.5864 | -0.26018 | -0.82897 | 2.54 |
| 1957 | *PRDX4* | 3.3829 | 2.8949 | 1.3526 | 0.75932 | 2.54 |
| 1958 | *AKIP1* | 4.5863 | 2.0104 | 1.0318 | -0.52552 | 2.54 |
| 1959 | *GNPDA2* | 3.8902 | 3.8467 | -0.10861 | -0.13837 | 2.54 |
| 1960 | *LCN8* | 4.311 | 3.4021 | -0.08496 | -1.1253 | 2.54 |
| 1961 | *OR5M1* | 3.8752 | 3.2843 | 0.46706 | -0.02804 | 2.54 |
| 1962 | *PI4K2A* | 4.0103 | 3.7052 | -0.08937 | -2.1817 | 2.54 |
| 1963 | *TMEM130* | 4.5762 | 4.4754 | -1.4258 | -2.6753 | 2.54 |
| 1964 | *C6orf99* | 4.7315 | 1.7859 | 1.1045 | 0.47011 | 2.54 |
| 1965 | *ZNF83* | 4.3959 | 3.2728 | -0.04876 | -2.101 | 2.54 |
| 1966 | *CDK5RAP2* | 4.0214 | 1.8656 | 1.7326 | -0.43068 | 2.54 |
| 1967 | *FBXO31* | 5.2473 | 3.0582 | -0.68869 | -1.583 | 2.54 |
| 1968 | *SIT1* | 4.0309 | 2.7237 | 0.86091 | -0.54972 | 2.54 |
| 1969 | *SLC22A2* | 4.5833 | 2.7528 | 0.27509 | -1.4082 | 2.54 |
| 1970 | *PACS2* | 3.3448 | 3.3391 | 0.92699 | 0.78696 | 2.54 |
| 1971 | *ARFGEF3* | 3.9858 | 3.6996 | -0.07656 | -1.1885 | 2.54 |

| | | | | | | |
|---|---|---|---|---|---|---|
| 1972 | *MID1IP1* | 3.1457 | 3.1058 | 1.355 | 1.1655 | 2.54 |
| 1973 | *ABCA2* | 4.5452 | 2.0337 | 1.0257 | -1.6343 | 2.53 |
| 1974 | *BSND* | 5.2616 | 1.284 | 1.0578 | -1.7567 | 2.53 |
| 1975 | *CD1B* | 4.4121 | 3.3788 | -0.18847 | -1.0903 | 2.53 |
| 1976 | *HMGCLL1* | 5.0613 | 2.9846 | -0.44353 | -1.8041 | 2.53 |
| 1977 | *ZNF146* | 4.3349 | 2.2334 | 1.032 | 0.40934 | 2.53 |
| 1978 | *CACNA2D1* | 4.822 | 1.5758 | 1.2023 | -0.53086 | 2.53 |
| 1979 | *ZNF346* | 3.9914 | 3.4266 | 0.18078 | -1.7646 | 2.53 |
| 1980 | *LBX2* | 4.8748 | 2.0536 | 0.66956 | -1.5401 | 2.53 |
| 1981 | *RUNX3* | 3.3876 | 3.1409 | 1.0691 | -0.33191 | 2.53 |
| 1982 | *OPA3* | 3.5237 | 2.1596 | 1.9115 | -0.26978 | 2.53 |
| 1983 | *CACNA1F* | 3.618 | 3.4533 | 0.52193 | -1.0089 | 2.53 |
| 1984 | *GLMN* | 3.9116 | 3.5269 | 0.15419 | -1.9817 | 2.53 |
| 1985 | *BAIAP2L2* | 4.5879 | 2.6027 | 0.40178 | -0.81714 | 2.53 |
| 1986 | *NBPF7* | 3.4253 | 3.2547 | 0.91197 | -1.6447 | 2.53 |
| 1987 | *SIGLEC5* | 4.329 | 1.8761 | 1.3861 | 0.70578 | 2.53 |
| 1988 | *CCDC167* | 5.7045 | 1.7423 | 0.14422 | -0.26093 | 2.53 |
| 1989 | *COLGALT2* | 2.9181 | 2.601 | 2.0719 | 1.4164 | 2.53 |
| 1990 | *OR1S1* | 3.0325 | 2.475 | 2.0827 | 0.26002 | 2.53 |
| 1991 | *ZSCAN4* | 4.4336 | 3.618 | -0.46248 | -1.2683 | 2.53 |
| 1992 | *NABP2* | 5.0652 | 2.6502 | -0.12681 | -1.5383 | 2.53 |
| 1993 | *TGFA* | 3.9289 | 2.5936 | 1.0644 | 0.19946 | 2.53 |
| 1994 | *DPT* | 5.2745 | 2.6444 | -0.33437 | -0.74551 | 2.53 |
| 1995 | *SLC45A3* | 3.9741 | 3.0614 | 0.54726 | 0.22102 | 2.53 |
| 1996 | *GNB2* | 3.4648 | 3.1271 | 0.98856 | 0.39882 | 2.53 |
| 1997 | *AMN* | 4.3049 | 3.2139 | 0.05795 | -0.49704 | 2.53 |
| 1998 | *RCAN1* | 4.2242 | 3.003 | 0.34895 | -0.65889 | 2.53 |
| 1999 | *YIPF7* | 4.5485 | 3.3903 | -0.36357 | -0.68109 | 2.53 |
| 2000 | *KANK1* | 4.1213 | 2.7828 | 0.66961 | -2.0542 | 2.52 |
| 2001 | *TOX3* | 3.9605 | 3.3287 | 0.284 | -1.6295 | 2.52 |
| 2002 | *TTC26* | 4.2405 | 3.3059 | 0.02667 | -2.2403 | 2.52 |
| 2003 | *ARSD* | 3.7303 | 3.0297 | 0.81197 | -0.78096 | 2.52 |
| 2004 | *CETP* | 3.6732 | 2.5529 | 1.3447 | -1.8929 | 2.52 |
| 2005 | *DUSP19* | 4.6466 | 3.3643 | -0.44353 | -1.0386 | 2.52 |
| 2006 | *ADGRB2* | 3.9346 | 3.1613 | 0.46968 | -1.4893 | 2.52 |
| 2007 | *OR8B4* | 3.4248 | 3.0891 | 1.0513 | -1.5954 | 2.52 |
| 2008 | *RUNDC3B* | 3.8622 | 2.2153 | 1.4871 | 0.44356 | 2.52 |
| 2009 | *ACOT4* | 4.6558 | 3.5364 | -0.6283 | -1.7462 | 2.52 |
| 2010 | *ZNF331* | 3.8481 | 2.0331 | 1.6813 | 1.4686 | 2.52 |
| 2011 | *LIX1L* | 3.4868 | 2.972 | 1.1016 | -1.0204 | 2.52 |
| 2012 | *KIF9* | 3.454 | 3.1629 | 0.94319 | -0.72936 | 2.52 |
| 2013 | *ZNF74* | 3.7699 | 3.5663 | 0.22383 | -0.87719 | 2.52 |
| 2014 | *FRMD7* | 4.6217 | 4.5549 | -1.6174 | -2.212 | 2.52 |
| 2015 | *CXCL10* | 3.7916 | 2.9824 | 0.78193 | -0.89957 | 2.52 |
| 2016 | *SHISA3* | 3.9157 | 3.6063 | 0.03295 | -1.3465 | 2.52 |
| 2017 | *APOBEC3F* | 4.7294 | 3.0481 | -0.22301 | -1.9987 | 2.52 |

| | | | | | | |
|---|---|---|---|---|---|---|
| 2018 | *ME2* | 4.3065 | 3.265 | -0.01788 | -0.22133 | 2.52 |
| 2019 | *C17orf70* | 3.7346 | 3.4266 | 0.39025 | -1.8649 | 2.52 |
| 2020 | *CPD* | 4.4524 | 2.5416 | 0.55692 | -2.6062 | 2.52 |
| 2021 | *AAAS* | 3.6811 | 3.4919 | 0.3765 | -2.6157 | 2.52 |
| 2022 | *TRIM62* | 4.1469 | 2.7992 | 0.60277 | -0.66061 | 2.52 |
| 2023 | *SNAPC2* | 4.5456 | 3.6286 | -0.62548 | -1.8016 | 2.52 |
| 2024 | *PTPRCAP* | 4.8375 | 2.8685 | -0.15733 | -1.9622 | 2.52 |
| 2025 | *FAM159A* | 3.0797 | 2.7367 | 1.7312 | 0.022848 | 2.52 |
| 2026 | *OR51D1* | 3.22 | 2.9382 | 1.3893 | -1.8227 | 2.52 |
| 2027 | *ZNF790* | 3.7348 | 3.442 | 0.36967 | -1.0711 | 2.52 |
| 2028 | *MBTPS1* | 3.2315 | 2.9627 | 1.3514 | 0.79448 | 2.52 |
| 2029 | *TNFRSF13C* | 3.0082 | 2.4001 | 2.1371 | -1.2252 | 2.52 |
| 2030 | *OR2C1* | 3.8947 | 3.1009 | 0.54961 | -1.0726 | 2.52 |
| 2031 | *C12orf74* | 3.6885 | 2.121 | 1.7356 | -0.35118 | 2.52 |
| 2032 | *FLYWCH2* | 4.0854 | 3.7728 | -0.31487 | -3.2312 | 2.51 |
| 2033 | *S100A11* | 2.9661 | 2.3886 | 2.1886 | 2.0281 | 2.51 |
| 2034 | *XIAP* | 4.296 | 2.4339 | 0.81301 | -0.28712 | 2.51 |
| 2035 | *HOXA1* | 5.1256 | 1.8761 | 0.54063 | -2.2313 | 2.51 |
| 2036 | *ADORA3* | 4.9348 | 1.8233 | 0.78382 | -1.2605 | 2.51 |
| 2037 | *RPRD1A* | 4.3816 | 3.8522 | -0.69287 | -1.2936 | 2.51 |
| 2038 | *SLTM* | 3.9001 | 3.1657 | 0.47395 | -3.4483 | 2.51 |
| 2039 | *C1RL* | 4.9343 | 1.5125 | 1.0928 | 0.33766 | 2.51 |
| 2040 | *HACD1* | 4.625 | 1.8748 | 1.0393 | -1.3536 | 2.51 |
| 2041 | *ASH2L* | 3.2244 | 2.9241 | 1.3895 | -0.49803 | 2.51 |
| 2042 | *KRTAP21-2* | 4.464 | 3.7355 | -0.66169 | -0.8004 | 2.51 |
| 2043 | *SFXN5* | 5.1542 | 2.5835 | -0.20302 | -0.76106 | 2.51 |
| 2044 | *HTR1D* | 3.9035 | 3.3521 | 0.27856 | -1.666 | 2.51 |
| 2045 | *MKL2* | 4.226 | 2.9889 | 0.31824 | -0.87644 | 2.51 |
| 2046 | *BTRC* | 3.7138 | 3.0711 | 0.74747 | -1.2489 | 2.51 |
| 2047 | *ADAM21* | 3.3236 | 3.2668 | 0.94183 | 0.38075 | 2.51 |
| 2048 | *PNPLA5* | 4.3488 | 2.8581 | 0.32513 | -1.7646 | 2.51 |
| 2049 | *PDGFRL* | 3.9218 | 3.2317 | 0.3773 | 0.37252 | 2.51 |
| 2050 | *PIGN* | 3.658 | 3.236 | 0.63535 | -0.42361 | 2.51 |
| 2051 | *ATP13A2* | 4.4121 | 2.4875 | 0.62692 | -0.88965 | 2.51 |
| 2052 | *C2CD2L* | 4.6096 | 2.9892 | -0.07496 | -2.0161 | 2.51 |
| 2053 | *PPA1* | 4.7599 | 1.961 | 0.80239 | -0.0649 | 2.51 |
| 2054 | *NAT14* | 3.8472 | 3.4013 | 0.27373 | -0.2765 | 2.51 |
| 2055 | *KRTAP15-1* | 3.4295 | 2.5495 | 1.5418 | -0.14649 | 2.51 |
| 2056 | *TMEM204* | 4.8618 | 3.5961 | -0.93727 | -1.1267 | 2.51 |
| 2057 | *TRIM2* | 3.2861 | 2.9094 | 1.325 | 0.078477 | 2.51 |
| 2058 | *OMG* | 3.8689 | 3.4671 | 0.18355 | -1.7043 | 2.51 |
| 2059 | *FAM127B* | 4.3409 | 2.055 | 1.1232 | -0.65184 | 2.51 |
| 2060 | *ZNF814* | 3.2439 | 2.6042 | 1.6706 | 0.28213 | 2.51 |
| 2061 | *ERC1* | 3.343 | 2.7477 | 1.4276 | -0.93489 | 2.51 |
| 2062 | *BCL11A* | 4.5446 | 2.7323 | 0.24096 | -1.6891 | 2.51 |
| 2063 | *XYLB* | 3.6497 | 3.2498 | 0.61636 | -0.44711 | 2.51 |

| | | | | | | |
|---|---|---|---|---|---|---|
| 2064 | *ACSM4* | 3.7287 | 2.3249 | 1.4608 | -1.5393 | 2.50 |
| 2065 | *BASP1* | 5.4449 | 1.056 | 1.0131 | -0.22538 | 2.50 |
| 2066 | *MRGPRX3* | 5.5247 | 2.7032 | -0.71443 | -0.76015 | 2.50 |
| 2067 | *ZNF761* | 3.6883 | 3.3099 | 0.51517 | -1.1209 | 2.50 |
| 2068 | *TMEM14C* | 4.3927 | 2.9673 | 0.15229 | -1.1739 | 2.50 |
| 2069 | *ZXDB* | 3.1398 | 2.5388 | 1.8324 | -0.09072 | 2.50 |
| 2070 | *DACH2* | 4.106 | 2.8493 | 0.55323 | -0.30711 | 2.50 |
| 2071 | *ZNF134* | 4.2863 | 3.5323 | -0.3105 | -0.3922 | 2.50 |
| 2072 | *RBAK-RBAKDN* | 3.5406 | 2.0786 | 1.8889 | -0.33131 | 2.50 |
| 2073 | *NEK11* | 4.0021 | 2.0482 | 1.4543 | -0.24973 | 2.50 |
| 2074 | *VSIG8* | 4.5204 | 4.4704 | -1.4864 | -2.5226 | 2.50 |
| 2075 | *C6orf62* | 4.3832 | 4.2367 | -1.1159 | -1.4289 | 2.50 |
| 2076 | *UQCRC1* | 3.3126 | 2.3686 | 1.8187 | -0.81178 | 2.50 |
| 2077 | *KCNK3* | 4.8459 | 1.9462 | 0.70671 | -1.5684 | 2.50 |
| 2078 | *KNG1* | 4.4318 | 1.8527 | 1.2128 | -2.8823 | 2.50 |
| 2079 | *MDFIC* | 3.7695 | 2.1504 | 1.573 | 1.3057 | 2.50 |
| 2080 | *KRTDAP* | 3.1595 | 2.6127 | 1.7199 | -2.2381 | 2.50 |
| 2081 | *MAD1L1* | 4.9466 | 1.8257 | 0.71818 | -2.075 | 2.50 |
| 2082 | *C4orf17* | 4.4278 | 2.491 | 0.57061 | -1.5039 | 2.50 |
| 2083 | *SPRYD4* | 4.3221 | 3.0812 | 0.08403 | -1.3675 | 2.50 |
| 2084 | *RRP8* | 4.707 | 2.372 | 0.4082 | -0.98138 | 2.50 |
| 2085 | *MRAP2* | 4.8127 | 2.3941 | 0.27849 | -0.81031 | 2.50 |
| 2086 | *COA1* | 3.6965 | 2.3102 | 1.4781 | 0.93632 | 2.49 |
| 2087 | *SIVA1* | 3.7084 | 2.2442 | 1.5321 | -0.37692 | 2.49 |
| 2088 | *SH3KBP1* | 4.1292 | 3.0958 | 0.25951 | -0.49028 | 2.49 |
| 2089 | *ERGIC3* | 3.9906 | 3.282 | 0.21175 | -1.0036 | 2.49 |
| 2090 | *ZNF683* | 4.7699 | 3.105 | -0.39079 | -1.1218 | 2.49 |
| 2091 | *SLC39A2* | 3.764 | 3.5485 | 0.17128 | -3.0713 | 2.49 |
| 2092 | *PPIL1* | 3.9064 | 3.7668 | -0.18991 | -1.2771 | 2.49 |
| 2093 | *STYX* | 5.1964 | 1.7881 | 0.49798 | -1.0898 | 2.49 |
| 2094 | *WDR91* | 5.012 | 3.199 | -0.72882 | -1.1325 | 2.49 |
| 2095 | *KRTAP13-2* | 3.5336 | 2.0646 | 1.8833 | 0.70262 | 2.49 |
| 2096 | *YTHDF3* | 4.5933 | 3.1694 | -0.28176 | -1.0312 | 2.49 |
| 2097 | *TREML4* | 4.4503 | 3.9938 | -0.96349 | -1.0773 | 2.49 |
| 2098 | *TLDC1* | 4.0223 | 2.9014 | 0.55629 | 0.11567 | 2.49 |
| 2099 | *BHLHE41* | 5.3756 | 4.08 | -1.9758 | -2.3681 | 2.49 |
| 2100 | *DCAF12L1* | 3.6532 | 2.0215 | 1.8033 | 0.95048 | 2.49 |
| 2101 | *RAD9B* | 4.2688 | 2.9472 | 0.26036 | -2.0838 | 2.49 |
| 2102 | *NPPA* | 3.3931 | 2.8201 | 1.2617 | -0.27356 | 2.49 |
| 2103 | *SPANXA1* | 3.0054 | 2.8011 | 1.6674 | 0.098389 | 2.49 |
| 2104 | *ATP6V1H* | 4.2455 | 4.0189 | -0.79083 | -1.3988 | 2.49 |
| 2105 | *TCEAL4* | 3.8653 | 2.4295 | 1.1756 | -1.2779 | 2.49 |
| 2106 | *C9orf163* | 4.1099 | 3.1205 | 0.23877 | -1.2866 | 2.49 |
| 2107 | *CAGE1* | 3.2936 | 2.9868 | 1.1885 | 0.5805 | 2.49 |
| 2108 | *FUT6* | 4.4157 | 3.5526 | -0.49953 | -1.2619 | 2.49 |
| 2109 | *IKZF5* | 3.7598 | 3.7159 | -0.00828 | -0.91696 | 2.49 |

| | | | | | | |
|---|---|---|---|---|---|---|
| 2110 | *LMF2* | 3.9639 | 2.4916 | 1.0119 | 0.061768 | 2.49 |
| 2111 | *PTBP3* | 3.5227 | 3.2955 | 0.64843 | -1.3907 | 2.49 |
| 2112 | *RND1* | 5.2432 | 2.1134 | 0.10729 | -1.4102 | 2.49 |
| 2113 | *SERPINA3* | 4.4862 | 1.6936 | 1.284 | -0.54142 | 2.49 |
| 2114 | *LCP2* | 2.9477 | 2.2672 | 2.2479 | 0.1832 | 2.49 |
| 2115 | *RETN* | 3.7789 | 2.4454 | 1.2383 | 0.29117 | 2.49 |
| 2116 | *MARVELD1* | 3.6094 | 3.5618 | 0.29118 | -1.9417 | 2.49 |
| 2117 | *CFP* | 5.5543 | 1.4941 | 0.41381 | 0.12053 | 2.49 |
| 2118 | *ZNF285* | 3.3699 | 3.162 | 0.93021 | -0.19984 | 2.49 |
| 2119 | *IL20* | 3.1516 | 2.5428 | 1.7677 | -0.44986 | 2.49 |
| 2120 | *ADGRE3* | 3.1226 | 2.6318 | 1.7065 | 0.46797 | 2.49 |
| 2121 | *IGF1* | 4.2768 | 4.262 | -1.0793 | -1.8695 | 2.49 |
| 2122 | *KCNMA1* | 3.6519 | 2.4467 | 1.3606 | -1.6597 | 2.49 |
| 2123 | *VN1R4* | 4.0861 | 3.0451 | 0.32727 | 0.31672 | 2.49 |
| 2124 | *R3HCC1L* | 3.969 | 2.3695 | 1.1198 | -1.5313 | 2.49 |
| 2125 | *ASF1B* | 5.0745 | 1.8477 | 0.53309 | 0.49883 | 2.49 |
| 2126 | *PHF6* | 3.552 | 2.5411 | 1.3621 | 0.40997 | 2.49 |
| 2127 | *ITGA9* | 3.9025 | 2.5166 | 1.0349 | -1.1706 | 2.48 |
| 2128 | *MPHOSPH9* | 6.0937 | 2.2001 | -0.84001 | -0.95017 | 2.48 |
| 2129 | *TEKT1* | 3.9986 | 3.0748 | 0.37516 | -1.7512 | 2.48 |
| 2130 | *CST9* | 4.0683 | 1.9193 | 1.4603 | 0.63445 | 2.48 |
| 2131 | *LCMT2* | 4.498 | 2.4578 | 0.49083 | -0.08206 | 2.48 |
| 2132 | *PLA2G12A* | 3.9759 | 3.4019 | 0.06868 | -1.3951 | 2.48 |
| 2133 | *CCDC158* | 3.1539 | 2.6197 | 1.6726 | 0.47319 | 2.48 |
| 2134 | *TMEM184C* | 4.2788 | 3.9779 | -0.8118 | -1.195 | 2.48 |
| 2135 | *CCDC65* | 3.5103 | 2.5739 | 1.3583 | -1.9172 | 2.48 |
| 2136 | *R3HDM1* | 3.9105 | 2.8505 | 0.68026 | 0.35076 | 2.48 |
| 2137 | *OR2T12* | 3.5933 | 3.4851 | 0.36265 | -1.9563 | 2.48 |
| 2138 | *KLHL34* | 4.4539 | 3.6755 | -0.6886 | -1.7285 | 2.48 |
| 2139 | *GANAB* | 4.4507 | 3.0951 | -0.10525 | -0.25741 | 2.48 |
| 2140 | *B9D1* | 3.3233 | 2.3259 | 1.7913 | -1.1588 | 2.48 |
| 2141 | *UEVLD* | 3.3345 | 2.9816 | 1.1228 | -1.1908 | 2.48 |
| 2142 | *PKD1* | 4.476 | 3.8731 | -0.91192 | -1.3542 | 2.48 |
| 2143 | *GIMAP7* | 3.9238 | 2.5308 | 0.98231 | -1.9351 | 2.48 |
| 2144 | *MID2* | 3.1329 | 2.7704 | 1.5331 | 1.1841 | 2.48 |
| 2145 | *CRBN* | 3.3156 | 2.2911 | 1.8249 | 0.11201 | 2.48 |
| 2146 | *PTGFRN* | 4.5124 | 2.2699 | 0.64765 | -0.01323 | 2.48 |
| 2147 | *OR5L1* | 3.673 | 3.6077 | 0.1486 | -1.3769 | 2.48 |
| 2148 | *CFAP45* | 5.0472 | 1.816 | 0.56476 | -0.06364 | 2.48 |
| 2149 | *SVOPL* | 4.476 | 2.1857 | 0.76425 | 0.00025 | 2.48 |
| 2150 | *CARD18* | 4.2806 | 2.135 | 1.009 | -0.88804 | 2.47 |
| 2151 | *NUDT5* | 3.8717 | 2.0891 | 1.4638 | 1.3561 | 2.47 |
| 2152 | *CAPN9* | 4.7003 | 1.7111 | 1.0124 | -2.0838 | 2.47 |
| 2153 | *PPFIA4* | 3.8964 | 3.3453 | 0.18039 | -1.4611 | 2.47 |
| 2154 | *KLF6* | 5.0664 | 3.8048 | -1.4493 | -1.6975 | 2.47 |
| 2155 | *IFI27* | 4.7258 | 1.3888 | 1.3071 | -0.29986 | 2.47 |

| | | | | | | |
|---|---|---|---|---|---|---|
| 2156 | *TTLL9* | 4.16 | 2.0383 | 1.2228 | -1.3787 | 2.47 |
| 2157 | *GLO1* | 3.8463 | 3.356 | 0.21634 | -1.7507 | 2.47 |
| 2158 | *SERPINF1* | 3.6692 | 2.2444 | 1.5037 | -0.2475 | 2.47 |
| 2159 | *MIF* | 5.1756 | 1.903 | 0.33664 | -0.67981 | 2.47 |
| 2160 | *MYC* | 4.1961 | 1.9024 | 1.3144 | -1.7107 | 2.47 |
| 2161 | *ANKUB1* | 4.1231 | 1.6789 | 1.6105 | -1.7602 | 2.47 |
| 2162 | *KLHL6* | 4.3161 | 3.2383 | -0.14198 | -0.26653 | 2.47 |
| 2163 | *SBK2* | 3.9289 | 2.1882 | 1.2943 | -1.0743 | 2.47 |
| 2164 | *ING4* | 4.6709 | 2.5905 | 0.14964 | -1.7935 | 2.47 |
| 2165 | *NLRP7* | 4.0997 | 3.3167 | -0.00555 | -0.17446 | 2.47 |
| 2166 | *ANKRD34A* | 5.367 | 1.1966 | 0.84714 | -1.3443 | 2.47 |
| 2167 | *MICAL1* | 4.7451 | 2.5876 | 0.07718 | -1.3111 | 2.47 |
| 2168 | *FLJ45513* | 3.3781 | 3.0505 | 0.98008 | -1.7079 | 2.47 |
| 2169 | *STRIP1* | 3.5578 | 2.4328 | 1.4177 | -1.8598 | 2.47 |
| 2170 | *SLC9C2* | 3.8091 | 3.3477 | 0.24896 | 0.12995 | 2.47 |
| 2171 | *OR6C4* | 3.8833 | 1.8199 | 1.7008 | -3.0028 | 2.47 |
| 2172 | *OR5J2* | 3.7069 | 2.0716 | 1.6252 | -1.2357 | 2.47 |
| 2173 | *CCKAR* | 3.8384 | 1.9872 | 1.5781 | -2.2078 | 2.47 |
| 2174 | *CATSPER2* | 3.9521 | 2.4413 | 1.0095 | -1.1426 | 2.47 |
| 2175 | *MAZ* | 3.5707 | 2.7344 | 1.0961 | -0.61655 | 2.47 |
| 2176 | *PATE2* | 3.8246 | 3.438 | 0.13738 | 0.054968 | 2.47 |
| 2177 | *TIFAB* | 3.3496 | 2.5804 | 1.4688 | -0.7372 | 2.47 |
| 2178 | *KPNA7* | 4.4552 | 2.2075 | 0.73441 | -2.3095 | 2.47 |
| 2179 | *PORCN* | 2.6392 | 2.4956 | 2.2621 | -1.068 | 2.47 |
| 2180 | *DPRX* | 3.9899 | 3.9694 | -0.56272 | -1.149 | 2.47 |
| 2181 | *ALDH1B1* | 3.6371 | 2.7858 | 0.97266 | -0.54028 | 2.47 |
| 2182 | *DPY19L4* | 4.3121 | 3.3511 | -0.26832 | -1.3128 | 2.46 |
| 2183 | *GABPB2* | 3.7603 | 2.1181 | 1.5151 | 1.2415 | 2.46 |
| 2184 | *APPL1* | 3.5823 | 3.1805 | 0.62804 | 0.068285 | 2.46 |
| 2185 | *SAT1* | 4.3217 | 1.9547 | 1.1138 | -1.505 | 2.46 |
| 2186 | *C17orf107* | 4.0444 | 3.3643 | -0.01938 | -0.36571 | 2.46 |
| 2187 | *RNF223* | 3.6699 | 2.446 | 1.2726 | 0.53163 | 2.46 |
| 2188 | *UGT1A3* | 4.7065 | 3.5871 | -0.90663 | -1.4997 | 2.46 |
| 2189 | *CLDN4* | 4.1405 | 2.5761 | 0.66966 | -0.4264 | 2.46 |
| 2190 | *GSTA4* | 3.5406 | 2.0346 | 1.8099 | 1.2057 | 2.46 |
| 2191 | *NFATC4* | 4.9242 | 3.0366 | -0.57713 | -0.84872 | 2.46 |
| 2192 | *ZC3HC1* | 3.9567 | 2.0256 | 1.3999 | -0.24239 | 2.46 |
| 2193 | *ORAI3* | 4.5706 | 1.7638 | 1.045 | -0.90149 | 2.46 |
| 2194 | *RDX* | 3.4172 | 2.392 | 1.5676 | -1.6502 | 2.46 |
| 2195 | *KLHL35* | 3.6592 | 2.205 | 1.5123 | -1.0852 | 2.46 |
| 2196 | *AKAP10* | 3.8365 | 1.7906 | 1.7493 | -1.9915 | 2.46 |
| 2197 | *EQTN* | 3.6924 | 2.0843 | 1.5987 | -3.3806 | 2.46 |
| 2198 | *RTN2* | 3.8372 | 3.618 | -0.08064 | -0.49286 | 2.46 |
| 2199 | *ZNF518B* | 3.769 | 3.2749 | 0.32816 | -0.95224 | 2.46 |
| 2200 | *GDNF* | 2.7524 | 2.7467 | 1.8722 | -1.4125 | 2.46 |
| 2201 | *ARRDC1* | 4.9551 | 4.2108 | -1.7963 | -1.9713 | 2.46 |

| | | | | | | |
|---|---|---|---|---|---|---|
| 2202 | *ENPP3* | 3.8436 | 3.4133 | 0.11185 | -0.58289 | 2.46 |
| 2203 | *LAMA2* | 4.8872 | 1.9622 | 0.51893 | -0.44353 | 2.46 |
| 2204 | *FOLR1* | 4.7769 | 3.7423 | -1.151 | -2.1921 | 2.46 |
| 2205 | *CAPN12* | 4.5365 | 2.368 | 0.46328 | -0.37417 | 2.46 |
| 2206 | *DPP7* | 5.5093 | 1.9087 | -0.05066 | -1.5848 | 2.46 |
| 2207 | *PSD3* | 4.0399 | 3.554 | -0.22757 | -0.39872 | 2.46 |
| 2208 | *ENDOD1* | 4.3932 | 2.4544 | 0.51575 | -1.049 | 2.45 |
| 2209 | *FAM102A* | 3.5059 | 2.4666 | 1.3897 | 0.53106 | 2.45 |
| 2210 | *ABRA* | 3.3031 | 2.6931 | 1.3648 | -0.04135 | 2.45 |
| 2211 | *CCL11* | 4.085 | 3.7881 | -0.51221 | -1.3835 | 2.45 |
| 2212 | *C11orf96* | 4.237 | 3.7445 | -0.62065 | -1.0861 | 2.45 |
| 2213 | *B4GALT3* | 5.0188 | 2.5131 | -0.17629 | -1.4325 | 2.45 |
| 2214 | *NOS1AP* | 3.5837 | 1.8925 | 1.8779 | -1.4811 | 2.45 |
| 2215 | *CYP3A43* | 3.7314 | 3.5556 | 0.06617 | -0.79825 | 2.45 |
| 2216 | *C1orf21* | 5.4341 | 2.4771 | -0.55848 | -0.79663 | 2.45 |
| 2217 | *PNMA1* | 3.0089 | 2.6949 | 1.6467 | -1.5923 | 2.45 |
| 2218 | *RS1* | 4.2858 | 3.0416 | 0.02027 | -0.33304 | 2.45 |
| 2219 | *FCRLA* | 4.8318 | 3.3494 | -0.83459 | -1.3033 | 2.45 |
| 2220 | *OR6K6* | 4.0505 | 3.2366 | 0.05775 | -1.206 | 2.45 |
| 2221 | *NFIX* | 4.065 | 2.3578 | 0.9183 | 0.71966 | 2.45 |
| 2222 | *SSX1* | 4.0871 | 2.5707 | 0.68287 | -0.58894 | 2.45 |
| 2223 | *SLC25A52* | 4.5701 | 1.7165 | 1.0538 | -0.94472 | 2.45 |
| 2224 | *ZNF253* | 5.5399 | 1.5648 | 0.23528 | -1.6315 | 2.45 |
| 2225 | *CABYR* | 4.2642 | 2.027 | 1.0468 | -0.57078 | 2.45 |
| 2226 | *P4HA2* | 4.0313 | 3.8585 | -0.55181 | -0.9414 | 2.45 |
| 2227 | *ATXN3* | 4.4581 | 2.8375 | 0.04176 | -0.03588 | 2.45 |
| 2228 | *PHF11* | 5.3657 | 1.5133 | 0.45807 | -1.2679 | 2.45 |
| 2229 | *OR1S2* | 4.1374 | 2.8781 | 0.32041 | -0.29414 | 2.45 |
| 2230 | *TADA2B* | 3.9694 | 3.0456 | 0.32043 | -1.1925 | 2.45 |
| 2231 | *RAB13* | 4.9705 | 3.1372 | -0.77284 | -2.0113 | 2.44 |
| 2232 | *TMEM183A* | 3.5512 | 1.9848 | 1.7981 | -0.1938 | 2.44 |
| 2233 | *XG* | 3.9407 | 3.4837 | -0.09032 | -1.2985 | 2.44 |
| 2234 | *SPTY2D1* | 3.8845 | 1.7403 | 1.7088 | -0.54329 | 2.44 |
| 2235 | *EGFL6* | 4.568 | 2.56 | 0.20391 | -1.8724 | 2.44 |
| 2236 | *ATP1A4* | 3.2455 | 2.5587 | 1.5276 | 1.4833 | 2.44 |
| 2237 | *ASTE1* | 5.3443 | 1.6071 | 0.38031 | -1.479 | 2.44 |
| 2238 | *PCDHA1* | 4.1259 | 3.1529 | 0.05181 | -2.6438 | 2.44 |
| 2239 | *LRP12* | 3.8324 | 2.8725 | 0.62444 | 0.12879 | 2.44 |
| 2240 | *ZNF850* | 3.9311 | 2.4438 | 0.95268 | -1.5105 | 2.44 |
| 2241 | *VAMP4* | 5.9385 | 0.7189 | 0.66945 | -1.3544 | 2.44 |
| 2242 | *MARCH6* | 5.1982 | 1.2352 | 0.89321 | -1.8999 | 2.44 |
| 2243 | *SLC2A5* | 3.5961 | 2.0434 | 1.6854 | 1.4669 | 2.44 |
| 2244 | *SMTNL2* | 4.7915 | 1.6002 | 0.93207 | -1.4526 | 2.44 |
| 2245 | *MOXD1* | 3.5402 | 3.3616 | 0.42022 | -1.1287 | 2.44 |
| 2246 | *NUAK2* | 4.3428 | 3.4898 | -0.51117 | -1.3469 | 2.44 |
| 2247 | *TRIM40* | 3.5346 | 3.2522 | 0.53086 | -0.41548 | 2.44 |

| | | | | | | |
|---|---|---|---|---|---|---|
| 2248 | *FAM184A* | 3.7774 | 3.5464 | -0.00662 | -0.81546 | 2.44 |
| 2249 | *COQ3* | 5.1231 | 2.4966 | -0.30528 | -1.5304 | 2.44 |
| 2250 | *EVC* | 3.6502 | 2.9036 | 0.76022 | -0.03924 | 2.44 |
| 2251 | *SUN1* | 4.0341 | 2.3937 | 0.88332 | -0.59581 | 2.44 |
| 2252 | *SPAG11B* | 3.7756 | 2.362 | 1.1729 | -1.2342 | 2.44 |
| 2253 | *LOC729159* | 3.106 | 2.9046 | 1.2987 | -0.63502 | 2.44 |
| 2254 | *TCP11L1* | 5.8601 | 2.1928 | -0.7443 | -1.7235 | 2.44 |
| 2255 | *ZNF610* | 2.7122 | 2.6289 | 1.9665 | -0.24142 | 2.44 |
| 2256 | *ABHD14A* | 3.5004 | 2.204 | 1.6021 | -1.4545 | 2.44 |
| 2257 | *DCAF5* | 4.6611 | 4.4312 | -1.7865 | -2.4286 | 2.44 |
| 2258 | *TIMM44* | 5.5623 | 2.1867 | -0.44353 | -0.83277 | 2.44 |
| 2259 | *ZNF41* | 5.035 | 1.9912 | 0.27224 | -1.5996 | 2.43 |
| 2260 | *LRP2BP* | 5.0261 | 4.2751 | -2.0042 | -2.6973 | 2.43 |
| 2261 | *RNF26* | 3.1246 | 2.1373 | 2.0336 | -0.33234 | 2.43 |
| 2262 | *WBSCR17* | 3.8644 | 2.8694 | 0.56046 | -0.37619 | 2.43 |
| 2263 | *SMIM6* | 5.186 | 1.505 | 0.60293 | -1.1021 | 2.43 |
| 2264 | *VAMP5* | 4.1414 | 2.4143 | 0.73823 | -1.8979 | 2.43 |
| 2265 | *WWP2* | 3.3133 | 3.1362 | 0.84357 | 0.013351 | 2.43 |
| 2266 | *OR10K2* | 4.3162 | 1.6199 | 1.356 | 0.096843 | 2.43 |
| 2267 | *USP17L15* | 4.0256 | 3.5495 | -0.28317 | -2.775 | 2.43 |
| 2268 | *COMMD4* | 4.0351 | 3.1295 | 0.12279 | 0.065382 | 2.43 |
| 2269 | *MGAT4B* | 5.7154 | 3.3185 | -1.7466 | -1.99 | 2.43 |
| 2270 | *TMEM262* | 4.7345 | 1.542 | 1.0068 | -1.031 | 2.43 |
| 2271 | *GRHL1* | 4.064 | 2.0604 | 1.158 | -0.24704 | 2.43 |
| 2272 | *ST3GAL6* | 3.8613 | 3.8573 | -0.43634 | -1.6891 | 2.43 |
| 2273 | *L3MBTL1* | 4.8117 | 2.1438 | 0.32624 | -0.48085 | 2.43 |
| 2274 | *OR9G1* | 4.5208 | 3.3398 | -0.57896 | -0.81293 | 2.43 |
| 2275 | *TSPAN19* | 4.5113 | 3.1224 | -0.35458 | -0.40772 | 2.43 |
| 2276 | *LUZP6* | 3.2922 | 2.2986 | 1.6883 | 0.34091 | 2.43 |
| 2277 | *NUMA1* | 5.2108 | 1.7513 | 0.31611 | -2.0208 | 2.43 |
| 2278 | *EPHB1* | 3.4122 | 2.3188 | 1.5471 | 0.15092 | 2.43 |
| 2279 | *COMMD6* | 3.7529 | 3.47 | 0.05454 | -1.257 | 2.43 |
| 2280 | *PTPN4* | 4.0276 | 2.2169 | 1.0321 | -0.55774 | 2.43 |
| 2281 | *SREK1* | 4.5143 | 2.0024 | 0.75943 | -0.41687 | 2.43 |
| 2282 | *PDIA4* | 3.635 | 2.6316 | 1.0084 | -1.5894 | 2.43 |
| 2283 | *PJA1* | 4.4755 | 3.0199 | -0.22048 | -1.0946 | 2.42 |
| 2284 | *PHF21B* | 4.3836 | 2.1633 | 0.72734 | -1.7403 | 2.42 |
| 2285 | *KLHL23* | 4.4772 | 2.4814 | 0.31541 | -1.4536 | 2.42 |
| 2286 | *PRPSAP2* | 4.1516 | 1.9556 | 1.1667 | -1.0868 | 2.42 |
| 2287 | *BPIFB1* | 4.1327 | 3.5014 | -0.36047 | -0.54019 | 2.42 |
| 2288 | *WDYHV1* | 4.3079 | 3.8695 | -0.90386 | -1.1066 | 2.42 |
| 2289 | *CCL15* | 4.6857 | 2.2487 | 0.33799 | -0.2372 | 2.42 |
| 2290 | *TNFSF9* | 5.8157 | 0.93409 | 0.52108 | -1.0041 | 2.42 |
| 2291 | *KIAA1524* | 4.4076 | 1.9578 | 0.90406 | -0.5018 | 2.42 |
| 2292 | *FAM192A* | 4.7555 | 2.862 | -0.34906 | -1.5722 | 2.42 |
| 2293 | *KIAA0100* | 3.8878 | 2.1167 | 1.2634 | -2.0382 | 2.42 |

| | | | | | | |
|---|---|---|---|---|---|---|
| 2294 | *GCC2* | 3.2061 | 3.2009 | 0.86053 | 0.61385 | 2.42 |
| 2295 | *USH2A* | 4.9153 | 2.1798 | 0.17147 | -1.2867 | 2.42 |
| 2296 | *LAYN* | 2.7249 | 2.6258 | 1.9153 | 0.51421 | 2.42 |
| 2297 | *ACAP3* | 3.8079 | 3.1143 | 0.34328 | -1.2519 | 2.42 |
| 2298 | *CECR2* | 2.8456 | 2.3273 | 2.0924 | 1.895 | 2.42 |
| 2299 | *UBE3B* | 4.6601 | 3.1042 | -0.49941 | -2.0116 | 2.42 |
| 2300 | *KIF7* | 4.3635 | 2.3144 | 0.58585 | -0.34376 | 2.42 |
| 2301 | *TMEM194A* | 3.7524 | 3.5451 | -0.03456 | -2.1354 | 2.42 |
| 2302 | *MAP3K2* | 4.7726 | 3.0782 | -0.58851 | -0.63939 | 2.42 |
| 2303 | *JPH2* | 3.5846 | 2.9975 | 0.68011 | -1.0186 | 2.42 |
| 2304 | *THADA* | 3.8694 | 3.7295 | -0.33844 | -0.41308 | 2.42 |
| 2305 | *GLB1L3* | 3.3861 | 2.7788 | 1.0947 | -1.0978 | 2.42 |
| 2306 | *STARD3* | 4.7791 | 2.5002 | -0.01973 | -0.0777 | 2.42 |
| 2307 | *FMO3* | 3.8496 | 1.8337 | 1.5761 | 1.4245 | 2.42 |
| 2308 | *CNBD2* | 4.8488 | 4.025 | -1.6146 | -2.3005 | 2.42 |
| 2309 | *HOXC6* | 3.8265 | 2.8613 | 0.57124 | -0.77997 | 2.42 |
| 2310 | *PDP2* | 3.9352 | 2.1141 | 1.2091 | -1.25 | 2.42 |
| 2311 | *SLPI* | 4.0335 | 3.8494 | -0.62476 | -1.9292 | 2.42 |
| 2312 | *EED* | 4.9755 | 1.499 | 0.78274 | -0.62747 | 2.42 |
| 2313 | *PICALM* | 4.8185 | 1.8296 | 0.6088 | -1.1105 | 2.42 |
| 2314 | *SCARB1* | 4.6665 | 2.6874 | -0.09709 | -0.67141 | 2.42 |
| 2315 | *THSD7A* | 4.2029 | 3.5343 | -0.48156 | -1.631 | 2.42 |
| 2316 | *FAM127A* | 4.0272 | 2.6613 | 0.5659 | -1.6597 | 2.42 |
| 2317 | *IGDCC3* | 3.2584 | 2.2096 | 1.7861 | -0.76074 | 2.42 |
| 2318 | *ZNF728* | 4.3697 | 2.2131 | 0.66944 | -0.44353 | 2.42 |
| 2319 | *ABCA13* | 4.9036 | 1.5257 | 0.821 | 0.081017 | 2.42 |
| 2320 | *FAHD1* | 5.0354 | 1.5927 | 0.62183 | -1.4909 | 2.42 |
| 2321 | *PPP2R3B* | 4.212 | 3.1828 | -0.1449 | -1.0323 | 2.42 |
| 2322 | *LPCAT4* | 4.0472 | 2.6712 | 0.53149 | 0.38934 | 2.42 |
| 2323 | *TRMT10B* | 3.4219 | 2.9191 | 0.90845 | -2.1523 | 2.42 |
| 2324 | *ZNF461* | 2.8142 | 2.6667 | 1.7679 | -1.0384 | 2.42 |
| 2325 | *FCRL5* | 3.5876 | 1.9156 | 1.7441 | -0.15226 | 2.42 |
| 2326 | *TAS1R1* | 3.9385 | 2.6545 | 0.65303 | -2.2254 | 2.42 |
| 2327 | *PROS1* | 4.1069 | 2.4027 | 0.73603 | -0.88278 | 2.42 |
| 2328 | *NPY5R* | 4.1541 | 2.3865 | 0.70487 | 0.17026 | 2.42 |
| 2329 | *C2orf83* | 3.4569 | 2.3422 | 1.4438 | -1.8049 | 2.41 |
| 2330 | *CTNNAL1* | 5.196 | 1.3665 | 0.67885 | -0.97982 | 2.41 |
| 2331 | *KIAA1683* | 4.9243 | 1.2823 | 1.0337 | -0.13183 | 2.41 |
| 2332 | *SLC44A5* | 3.7258 | 3.0288 | 0.4856 | -2.058 | 2.41 |
| 2333 | *KIF4B* | 3.8764 | 2.3018 | 1.0613 | -0.36087 | 2.41 |
| 2334 | *CXCL16* | 4.081 | 2.8902 | 0.26709 | -1.3419 | 2.41 |
| 2335 | *CNNM2* | 4.9237 | 3.2034 | -0.88927 | -1.0386 | 2.41 |
| 2336 | *DSG1* | 3.4442 | 2.9901 | 0.80248 | -1.1958 | 2.41 |
| 2337 | *LOC643355* | 3.9664 | 3.1353 | 0.13453 | -0.44744 | 2.41 |
| 2338 | *TWF2* | 3.3152 | 2.1061 | 1.8107 | -1.0106 | 2.41 |
| 2339 | *RCSD1* | 3.564 | 3.5177 | 0.14885 | -0.58806 | 2.41 |

| | | | | | | |
|---|---|---|---|---|---|---|
| 2340 | *SPINK6* | 5.5043 | 1.7862 | -0.06072 | -0.08199 | 2.41 |
| 2341 | *TUSC5* | 4.1319 | 2.511 | 0.58678 | -1.4474 | 2.41 |
| 2342 | *NR2C1* | 3.474 | 3.3001 | 0.45421 | -0.9612 | 2.41 |
| 2343 | *SPATA31C1* | 3.9071 | 3.7646 | -0.44353 | -0.60727 | 2.41 |
| 2344 | *GAREM* | 3.0953 | 2.1471 | 1.9837 | -0.24578 | 2.41 |
| 2345 | *ARF5* | 4.1032 | 2.848 | 0.27433 | -2.5298 | 2.41 |
| 2346 | *UPK3A* | 5.2193 | 1.6429 | 0.36147 | 0.14277 | 2.41 |
| 2347 | *NAF1* | 4.138 | 3.8555 | -0.77162 | -1.6262 | 2.41 |
| 2348 | *LDLRAP1* | 3.9251 | 1.771 | 1.5243 | -0.87195 | 2.41 |
| 2349 | *VMO1* | 3.719 | 2.9864 | 0.5094 | -0.66215 | 2.40 |
| 2350 | *MORN2* | 4.3735 | 2.7615 | 0.07822 | -0.64793 | 2.40 |
| 2351 | *PABPC4L* | 2.6353 | 2.4771 | 2.1004 | -0.95682 | 2.40 |
| 2352 | *FAM205A* | 5.4646 | 1.9174 | -0.17026 | -2.0383 | 2.40 |
| 2353 | *FABP2* | 4.1513 | 2.6853 | 0.37511 | -2.1984 | 2.40 |
| 2354 | *G6PC3* | 3.6297 | 2.7397 | 0.84205 | -1.0172 | 2.40 |
| 2355 | *POM121L12* | 4.1298 | 2.7727 | 0.30799 | -1.0696 | 2.40 |
| 2356 | *HDAC5* | 2.9761 | 2.46 | 1.7741 | -1.2649 | 2.40 |
| 2357 | *OR4C46* | 4.6442 | 3.5625 | -0.99699 | -1.5619 | 2.40 |
| 2358 | *PRB2* | 5.0689 | 2.134 | 0.00631 | -0.13293 | 2.40 |
| 2359 | *HRH2* | 2.9188 | 2.2369 | 2.0515 | 0.93074 | 2.40 |
| 2360 | *PMAIP1* | 5.2834 | 2.3233 | -0.39971 | -2.4168 | 2.40 |
| 2361 | *RRBP1* | 3.8786 | 2.4372 | 0.89116 | -0.7361 | 2.40 |
| 2362 | *NOTCH3* | 4.1543 | 3.3924 | -0.34014 | -0.74141 | 2.40 |
| 2363 | *MAGEB2* | 4.0602 | 3.4347 | -0.28926 | -2.3287 | 2.40 |
| 2364 | *ZCCHC6* | 4.4831 | 2.5589 | 0.16218 | -1.3055 | 2.40 |
| 2365 | *RGPD3* | 3.4644 | 2.0367 | 1.7027 | 0.77812 | 2.40 |
| 2366 | *WNT9A* | 4.4956 | 2.0146 | 0.69034 | -0.7189 | 2.40 |
| 2367 | *FAM19A3* | 3.1477 | 2.8323 | 1.2204 | 0.66504 | 2.40 |
| 2368 | *EVI2B* | 4.8068 | 1.8858 | 0.50464 | -0.47857 | 2.40 |
| 2369 | *GCNT1* | 2.8813 | 2.7532 | 1.5627 | -0.02187 | 2.40 |
| 2370 | *GPR157* | 4.2789 | 2.7587 | 0.15915 | -1.0284 | 2.40 |
| 2371 | *FNBP4* | 3.4305 | 2.0565 | 1.7084 | -1.4624 | 2.40 |
| 2372 | *PDE4B* | 3.9444 | 1.9557 | 1.2941 | 1.1886 | 2.40 |
| 2373 | *HRNR* | 4.6734 | 1.8709 | 0.64725 | 0.30293 | 2.40 |
| 2374 | *AAMDC* | 3.3454 | 2.5911 | 1.2548 | -1.9019 | 2.40 |
| 2375 | *LOC79999* | 4.1602 | 3.6998 | -0.66992 | -2.8628 | 2.40 |
| 2376 | *KBTBD7* | 3.7344 | 3.2441 | 0.21027 | -1.8016 | 2.40 |
| 2377 | *SPATA31A5* | 4.2255 | 4.1528 | -1.1898 | -2.5392 | 2.40 |
| 2378 | *CT47A5* | 4.1543 | 3.5871 | -0.55379 | -1.2055 | 2.40 |
| 2379 | *C8orf58* | 4.5145 | 2.9762 | -0.30336 | -1.0291 | 2.40 |
| 2380 | *TPM3* | 3.1766 | 3.0978 | 0.91013 | 0.65059 | 2.39 |
| 2381 | *CRMP1* | 5.4676 | 1.1327 | 0.58355 | 0.53406 | 2.39 |
| 2382 | *ATP6V0A4* | 4.2991 | 3.9203 | -1.0359 | -1.3507 | 2.39 |
| 2383 | *PHYKPL* | 3.5745 | 2.8546 | 0.75321 | -2.9054 | 2.39 |
| 2384 | *PCDHA2* | 4.4251 | 1.3843 | 1.3707 | 1.2214 | 2.39 |
| 2385 | *SUSD6* | 3.7516 | 3.0325 | 0.39467 | -1.584 | 2.39 |

| | | | | | | |
|---|---|---|---|---|---|---|
| 2386 | *SLC10A7* | 4.9665 | 1.4863 | 0.72543 | 0.70596 | 2.39 |
| 2387 | *ANAPC4* | 4.0014 | 3.9142 | -0.7385 | -1.6324 | 2.39 |
| 2388 | *PTPN7* | 4.4319 | 2.6657 | 0.07904 | -1.1189 | 2.39 |
| 2389 | *MAPK3* | 4.3102 | 1.6857 | 1.1772 | 0.49084 | 2.39 |
| 2390 | *MAN1C1* | 3.6868 | 2.2646 | 1.2206 | 0.51273 | 2.39 |
| 2391 | *PRB1* | 3.3483 | 3.1092 | 0.71407 | -0.78229 | 2.39 |
| 2392 | *MRS2* | 5.2572 | 2.5481 | -0.63575 | -1.3407 | 2.39 |
| 2393 | *POU4F1* | 4.4852 | 3.1147 | -0.43101 | -1.248 | 2.39 |
| 2394 | *DGAT2* | 3.3059 | 2.9942 | 0.86844 | -1.2603 | 2.39 |
| 2395 | *RNF44* | 3.581 | 3.5283 | 0.05848 | -1.1684 | 2.39 |
| 2396 | *CUTC* | 3.0461 | 2.4174 | 1.703 | -0.98486 | 2.39 |
| 2397 | *GPR171* | 3.7267 | 3.1825 | 0.2573 | -0.7064 | 2.39 |
| 2398 | *SYNGAP1* | 4.02 | 1.6936 | 1.4527 | -1.2288 | 2.39 |
| 2399 | *ITGAX* | 4.6197 | 1.416 | 1.1302 | -1.5441 | 2.39 |
| 2400 | *SLC22A5* | 3.8796 | 2.692 | 0.5941 | 0.072125 | 2.39 |
| 2401 | *LOC286238* | 3.7662 | 3.1652 | 0.2336 | -2.2413 | 2.39 |
| 2402 | *ZNF678* | 2.6086 | 2.5459 | 2.009 | 1.1296 | 2.39 |
| 2403 | *AVPR2* | 3.5734 | 2.3727 | 1.2173 | -1.5965 | 2.39 |
| 2404 | *TRIM45* | 3.9008 | 2.6249 | 0.6351 | -0.0442 | 2.39 |
| 2405 | *TMEM219* | 3.2542 | 2.0408 | 1.8657 | -0.03506 | 2.39 |
| 2406 | *FAM104B* | 3.5841 | 3.3359 | 0.23956 | -0.98459 | 2.39 |
| 2407 | *TMPRSS11A* | 3.7898 | 3.3082 | 0.06002 | -0.27016 | 2.39 |
| 2408 | *CEP170B* | 4.3117 | 2.9856 | -0.13943 | -0.90954 | 2.39 |
| 2409 | *DGKD* | 4.4362 | 4.3008 | -1.5795 | -2.08 | 2.39 |
| 2410 | *OR2T10* | 4.6791 | 2.7809 | -0.30268 | -1.4576 | 2.39 |
| 2411 | *RGS7BP* | 3.1028 | 2.5903 | 1.4634 | -0.04135 | 2.39 |
| 2412 | *MAST1* | 4.2119 | 3.2776 | -0.33312 | -2.1848 | 2.39 |
| 2413 | *TSPAN1* | 3.7886 | 1.9179 | 1.4496 | -0.42817 | 2.39 |
| 2414 | *CCDC121* | 4.2306 | 1.8335 | 1.0914 | 0.80953 | 2.39 |
| 2415 | *RAB5C* | 5.2372 | 2.8317 | -0.91363 | -2.0613 | 2.39 |
| 2416 | *CPM* | 3.7861 | 2.9089 | 0.45892 | 0.40342 | 2.38 |
| 2417 | *GAS2L2* | 3.7182 | 2.9411 | 0.49347 | -1.6763 | 2.38 |
| 2418 | *ABLIM3* | 3.802 | 2.8191 | 0.53135 | 0.054336 | 2.38 |
| 2419 | *CFL1* | 3.0905 | 2.3667 | 1.6951 | -1.5402 | 2.38 |
| 2420 | *ABRACL* | 3.6776 | 3.266 | 0.20851 | -1.7577 | 2.38 |
| 2421 | *NUDT11* | 4.2517 | 3.4094 | -0.50957 | -2.6188 | 2.38 |
| 2422 | *HIST1H2BO* | 2.8152 | 2.4766 | 1.8596 | 0.069589 | 2.38 |
| 2423 | *MED14OS* | 4.5238 | 3.1701 | -0.54604 | -1.5125 | 2.38 |
| 2424 | *TTC30B* | 4.4257 | 1.5158 | 1.2063 | 0.47245 | 2.38 |
| 2425 | *ARSE* | 3.5217 | 2.4719 | 1.1535 | -2.5795 | 2.38 |
| 2426 | *PER2* | 3.8627 | 3.3749 | -0.09062 | -0.17241 | 2.38 |
| 2427 | *OR2T11* | 2.736 | 2.4907 | 1.9197 | 1.1803 | 2.38 |
| 2428 | *CPNE1* | 4.7161 | 1.3823 | 1.0471 | -1.1644 | 2.38 |
| 2429 | *JUP* | 4.8035 | 2.5085 | -0.16814 | -0.75548 | 2.38 |
| 2430 | *NT5M* | 3.2946 | 3.2043 | 0.64459 | -0.5667 | 2.38 |
| 2431 | *NME4* | 3.6196 | 2.465 | 1.0584 | 0.89315 | 2.38 |

| | | | | | | |
|---|---|---|---|---|---|---|
| 2432 | *ARHGEF11* | 4.5535 | 2.0046 | 0.58403 | -0.57732 | 2.38 |
| 2433 | *ANKRD61* | 4.091 | 3.2228 | -0.17243 | -0.84001 | 2.38 |
| 2434 | *GRIN2D* | 6.092 | 0.73457 | 0.31401 | -2.3062 | 2.38 |
| 2435 | *PRRG4* | 3.2047 | 2.6624 | 1.2728 | 0.58455 | 2.38 |
| 2436 | *MKRN2* | 2.8291 | 2.2561 | 2.0544 | -2.7223 | 2.38 |
| 2437 | *KRTAP12-1* | 3.6159 | 2.0185 | 1.5047 | 0.9039 | 2.38 |
| 2438 | *BRS3* | 3.9977 | 2.6906 | 0.45067 | -0.45119 | 2.38 |
| 2439 | *KIF13B* | 3.351 | 2.3404 | 1.4475 | 0.88805 | 2.38 |
| 2440 | *C19orf38* | 3.4705 | 1.9995 | 1.6675 | 0.003685 | 2.38 |
| 2441 | *CBR1* | 5.19 | 2.5052 | -0.56036 | -2.8398 | 2.38 |
| 2442 | *MLLT10* | 5.1279 | 2.8216 | -0.81491 | -0.81863 | 2.38 |
| 2443 | *CPZ* | 3.5524 | 1.8998 | 1.6814 | -1.6586 | 2.38 |
| 2444 | *ARL5A* | 3.4892 | 2.6889 | 0.95539 | 0.284 | 2.38 |
| 2445 | *THEM5* | 3.5152 | 3.0195 | 0.59683 | -0.03924 | 2.38 |
| 2446 | *TBL1XR1* | 4.0969 | 2.8034 | 0.23006 | -2.2713 | 2.38 |
| 2447 | *APBA1* | 4.2718 | 3.1925 | -0.33601 | -0.79558 | 2.38 |
| 2448 | *MYF6* | 4.1776 | 3.0281 | -0.07743 | -1.2535 | 2.38 |
| 2449 | *PTRHD1* | 4.2573 | 2.762 | 0.10857 | -0.19889 | 2.38 |
| 2450 | *NASP* | 4.0717 | 3.2598 | -0.20419 | -1.1891 | 2.38 |
| 2451 | *ARL6IP4* | 2.816 | 2.8015 | 1.5097 | 0.35851 | 2.38 |
| 2452 | *HLA-A* | 4.9368 | 2.2164 | -0.02694 | -0.60113 | 2.38 |
| 2453 | *FSD2* | 3.5636 | 1.8157 | 1.7468 | -1.4764 | 2.38 |
| 2454 | *TOMM40L* | 4.7453 | 2.7184 | -0.33821 | -0.8462 | 2.38 |
| 2455 | *AEBP2* | 3.7859 | 2.8489 | 0.49055 | -1.8253 | 2.38 |
| 2456 | *ANKRD30B* | 3.1278 | 2.9886 | 1.0082 | -0.48991 | 2.37 |
| 2457 | *CRNN* | 4.1409 | 2.6846 | 0.2982 | 0.033383 | 2.37 |
| 2458 | *LY75* | 4.4254 | 2.2312 | 0.46631 | 0.27373 | 2.37 |
| 2459 | *OR10A2* | 3.7973 | 2.2245 | 1.0989 | 0.26509 | 2.37 |
| 2460 | *C19orf47* | 4.6626 | 1.8673 | 0.58868 | -0.86923 | 2.37 |
| 2461 | *CEACAM18* | 3.8812 | 1.6955 | 1.5418 | -1.0451 | 2.37 |
| 2462 | *TMEM127* | 3.442 | 3.0325 | 0.64351 | -0.55516 | 2.37 |
| 2463 | *AQP3* | 3.5338 | 3.0565 | 0.52748 | 0.19028 | 2.37 |
| 2464 | *PIK3CG* | 4.6634 | 3.3893 | -0.93555 | -1.2935 | 2.37 |
| 2465 | *MEX3B* | 3.642 | 3.5473 | -0.07219 | -0.29292 | 2.37 |
| 2466 | *AIM1* | 4.03 | 2.3024 | 0.78393 | -1.7649 | 2.37 |
| 2467 | *EHD4* | 5.4343 | 1.2398 | 0.44192 | -1.0412 | 2.37 |
| 2468 | *ADCY10* | 4.2036 | 1.5076 | 1.4047 | -1.6235 | 2.37 |
| 2469 | *ASB1* | 3.495 | 2.3629 | 1.2578 | -1.7079 | 2.37 |
| 2470 | *C4orf19* | 4.6058 | 2.0539 | 0.45432 | 0.33187 | 2.37 |
| 2471 | *FYCO1* | 4.2232 | 1.5136 | 1.3763 | 0.24233 | 2.37 |
| 2472 | *AIDA* | 4.4736 | 1.3624 | 1.2727 | -1.4174 | 2.37 |
| 2473 | *SLFN14* | 2.7691 | 2.2171 | 2.1223 | 1.1669 | 2.37 |
| 2474 | *SREBF2* | 5.5392 | 0.95143 | 0.61711 | 0.22088 | 2.37 |
| 2475 | *PDE6A* | 3.6954 | 1.7136 | 1.6986 | 0.22031 | 2.37 |
| 2476 | *PTAFR* | 3.5447 | 3.2884 | 0.27343 | -1.8181 | 2.37 |
| 2477 | *TUBB1* | 4.2227 | 3.1238 | -0.24037 | -0.43313 | 2.37 |

| | | | | | | |
|---|---|---|---|---|---|---|
| 2478 | *SPSB3* | 3.6195 | 2.386 | 1.1004 | -0.05744 | 2.37 |
| 2479 | *PLCB3* | 3.4761 | 1.8466 | 1.7819 | -2.0664 | 2.37 |
| 2480 | *DUSP11* | 5.1483 | 1.3405 | 0.61328 | -5.7233 | 2.37 |
| 2481 | *CFAP221* | 4.4544 | 2.6522 | -0.00551 | -2.0586 | 2.37 |
| 2482 | *MXRA8* | 3.7377 | 2.4426 | 0.92019 | 0.54218 | 2.37 |
| 2483 | *SERP2* | 4.1232 | 3.1761 | -0.19911 | -1.6984 | 2.37 |
| 2484 | *ASPA* | 4.173 | 3.1661 | -0.24126 | -1.8435 | 2.37 |
| 2485 | *C6orf229* | 3.5508 | 2.5993 | 0.94747 | 0.53922 | 2.37 |
| 2486 | *STAT3* | 4.8299 | 1.5019 | 0.76425 | -1.6676 | 2.37 |
| 2487 | *ITGBL1* | 4.0963 | 2.2861 | 0.71179 | -0.73243 | 2.36 |
| 2488 | *CXXC5* | 4.2685 | 3.7461 | -0.92186 | -1.8941 | 2.36 |
| 2489 | *OR4F15* | 4.353 | 1.4248 | 1.3134 | -1.8157 | 2.36 |
| 2490 | *ARHGEF18* | 3.5437 | 3.0673 | 0.4797 | -2.4853 | 2.36 |
| 2491 | *BMP8A* | 3.6974 | 2.0741 | 1.319 | -2.716 | 2.36 |
| 2492 | *CIT* | 3.0336 | 2.7771 | 1.2796 | -1.8478 | 2.36 |
| 2493 | *C11orf74* | 3.6191 | 2.7483 | 0.7223 | -2.1108 | 2.36 |
| 2494 | *MYO7A* | 5.0212 | 1.4416 | 0.62659 | -0.3783 | 2.36 |
| 2495 | *HEXDC* | 4.1735 | 1.522 | 1.3933 | 0.17427 | 2.36 |
| 2496 | *HNRNPA0* | 4.546 | 4.1628 | -1.6213 | -1.63 | 2.36 |
| 2497 | *CSF2* | 3.2159 | 3.058 | 0.81265 | -0.20663 | 2.36 |
| 2498 | *AIP* | 3.6033 | 3.072 | 0.40913 | -0.85799 | 2.36 |
| 2499 | *SYP* | 3.5835 | 2.5891 | 0.91083 | 0.55001 | 2.36 |
| 2500 | *SLC13A2* | 2.7986 | 2.7696 | 1.5121 | -0.68422 | 2.36 |
| 2501 | *OR2T34* | 3.7202 | 3.1987 | 0.16091 | -0.44353 | 2.36 |
| 2502 | *STAT5B* | 2.4582 | 2.3944 | 2.2267 | -0.85324 | 2.36 |
| 2503 | *COL11A1* | 3.8156 | 1.9708 | 1.2927 | -1.1096 | 2.36 |
| 2504 | *PGBD2* | 4.2969 | 3.2919 | -0.5113 | -0.63826 | 2.36 |
| 2505 | *CSGALNACT1* | 3.8357 | 3.2941 | -0.0525 | -0.91701 | 2.36 |
| 2506 | *ZNF222* | 3.3969 | 2.3847 | 1.2934 | -3.8595 | 2.36 |
| 2507 | *RNF32* | 3.5581 | 2.0484 | 1.4664 | -0.76184 | 2.36 |
| 2508 | *PCDHGC3* | 3.4135 | 3.0653 | 0.59377 | -1.7766 | 2.36 |
| 2509 | *ZFP14* | 4.8688 | 1.9148 | 0.28885 | -0.83187 | 2.36 |
| 2510 | *HTR3C* | 5.3987 | 2.4169 | -0.7435 | -1.1836 | 2.36 |
| 2511 | *ABCD1* | 4.5089 | 2.6207 | -0.05801 | -0.54775 | 2.36 |
| 2512 | *KCNH2* | 2.9709 | 2.9621 | 1.1381 | -0.51664 | 2.36 |
| 2513 | *CD59* | 4.2135 | 2.4387 | 0.41717 | -0.14849 | 2.36 |
| 2514 | *AKR1E2* | 3.1361 | 2.5704 | 1.3627 | -0.80436 | 2.36 |
| 2515 | *LOC101928841* | 4.1355 | 2.7598 | 0.17356 | -0.52372 | 2.36 |
| 2516 | *LIPA* | 4.5301 | 1.4789 | 1.0595 | -0.82833 | 2.36 |
| 2517 | *GRB10* | 4.8331 | 1.1721 | 1.0619 | -0.40214 | 2.36 |
| 2518 | *RPS21* | 5.8381 | 2.1353 | -0.90854 | -1.0134 | 2.35 |
| 2519 | *ZNF730* | 3.6347 | 2.1542 | 1.2751 | -0.38945 | 2.35 |
| 2520 | *METTL21A* | 4.6558 | 1.9634 | 0.44324 | -1.632 | 2.35 |
| 2521 | *ZNF214* | 3.619 | 1.8272 | 1.6139 | 0.31183 | 2.35 |
| 2522 | *ADAMTS10* | 4.0141 | 2.4026 | 0.64207 | -1.2884 | 2.35 |
| 2523 | *ALDH1A1* | 4.0494 | 2.5958 | 0.41346 | -1.3495 | 2.35 |

| | | | | | | |
|---|---|---|---|---|---|---|
| 2524 | *NRD1* | 4.1544 | 2.114 | 0.79022 | -0.1611 | 2.35 |
| 2525 | *NDST3* | 3.2703 | 2.3419 | 1.4449 | -0.49727 | 2.35 |
| 2526 | *TSPAN13* | 2.8631 | 2.1924 | 2.0011 | -1.6776 | 2.35 |
| 2527 | *MAFG* | 3.9616 | 3.3646 | -0.26968 | -1.9935 | 2.35 |
| 2528 | *MOB1B* | 5.3287 | 2.0716 | -0.34486 | -1.6934 | 2.35 |
| 2529 | *HTR3D* | 4.1805 | 3.6922 | -0.8175 | -0.9562 | 2.35 |
| 2530 | *TMEM168* | 4.3665 | 1.8856 | 0.80062 | -2.1058 | 2.35 |
| 2531 | *PLXNA2* | 2.874 | 2.5704 | 1.6066 | -1.8538 | 2.35 |
| 2532 | *SEZ6* | 3.531 | 2.9727 | 0.54673 | -1.2412 | 2.35 |
| 2533 | *TBRG1* | 5.0332 | 1.8243 | 0.19121 | -0.84992 | 2.35 |
| 2534 | *MSANTD1* | 3.6784 | 1.8886 | 1.4814 | -1.2692 | 2.35 |
| 2535 | *CES3* | 2.9633 | 2.8111 | 1.2736 | -0.08283 | 2.35 |
| 2536 | *PSMC6* | 4.1918 | 3.2542 | -0.39838 | -1.7153 | 2.35 |
| 2537 | *LMAN2L* | 5.0462 | 1.1995 | 0.80184 | -1.4684 | 2.35 |
| 2538 | *ADAMTS19* | 4.0137 | 3.6967 | -0.66353 | -0.78345 | 2.35 |
| 2539 | *PCDH1* | 3.5325 | 3.3053 | 0.20893 | -0.62875 | 2.35 |
| 2540 | *GBP5* | 4.2998 | 2.9798 | -0.23309 | -0.52154 | 2.35 |
| 2541 | *GJA3* | 2.4993 | 2.4074 | 2.1382 | -0.49001 | 2.35 |
| 2542 | *IFNA6* | 4.8633 | 1.6013 | 0.57427 | -0.36705 | 2.35 |
| 2543 | *C4BPA* | 2.7482 | 2.1577 | 2.1323 | -0.61429 | 2.35 |
| 2544 | *RAB28* | 3.9356 | 2.4685 | 0.63314 | -0.27707 | 2.35 |
| 2545 | *ZNF697* | 2.738 | 2.4833 | 1.8154 | -1.2118 | 2.35 |
| 2546 | *FNDC4* | 4.3827 | 3.7543 | -1.1006 | -1.8272 | 2.35 |
| 2547 | *OTOP1* | 4.1805 | 3.3497 | -0.49389 | -0.912 | 2.35 |
| 2548 | *OR2J2* | 3.0654 | 2.0585 | 1.9121 | 1.0927 | 2.35 |
| 2549 | *MKLN1* | 4.4243 | 1.814 | 0.79654 | -1.7217 | 2.34 |
| 2550 | *PABPC3* | 2.973 | 2.6799 | 1.3789 | -0.53856 | 2.34 |
| 2551 | *GBP1* | 4.1433 | 3.9482 | -1.0613 | -1.0739 | 2.34 |
| 2552 | *CALB1* | 3.8837 | 2.0977 | 1.0484 | -0.11514 | 2.34 |
| 2553 | *ZNF302* | 2.5661 | 2.3435 | 2.1199 | -1.2936 | 2.34 |
| 2554 | *PKP2* | 3.9064 | 3.2434 | -0.12053 | -0.36295 | 2.34 |
| 2555 | *WWTR1* | 3.5886 | 3.5068 | -0.06657 | -0.92868 | 2.34 |
| 2556 | *OR10H2* | 4.1562 | 2.773 | 0.09874 | -1.3371 | 2.34 |
| 2557 | *NXT2* | 5.3852 | 0.90318 | 0.73816 | -0.16749 | 2.34 |
| 2558 | *EXOSC3* | 4.7991 | 1.4668 | 0.7603 | -0.07903 | 2.34 |
| 2559 | *CCDC64B* | 4.9432 | 1.2379 | 0.84157 | -0.11902 | 2.34 |
| 2560 | *SH3BP4* | 4.1585 | 2.3859 | 0.47778 | -0.50809 | 2.34 |
| 2561 | *SMARCE1* | 3.6983 | 2.4191 | 0.90347 | 0.6891 | 2.34 |
| 2562 | *ZNF695* | 4.7366 | 1.4812 | 0.79954 | -0.10753 | 2.34 |
| 2563 | *VTCN1* | 3.3791 | 2.7107 | 0.92685 | -1.7505 | 2.34 |
| 2564 | *LRRC17* | 3.3591 | 1.9709 | 1.6822 | 0.78117 | 2.34 |
| 2565 | *FBXO27* | 3.2252 | 2.9374 | 0.84897 | 0.30181 | 2.34 |
| 2566 | *GFRA1* | 4.6682 | 1.2788 | 1.0644 | 0.24059 | 2.34 |
| 2567 | *SPANXN1* | 3.7202 | 1.7023 | 1.5886 | 0.069884 | 2.34 |
| 2568 | *OR14A16* | 5.1847 | 0.95637 | 0.86844 | 0.60637 | 2.34 |
| 2569 | *DSCR3* | 2.9934 | 2.476 | 1.5401 | 0.48579 | 2.34 |

| | | | | | | |
|---|---|---|---|---|---|---|
| 2570 | *HIST2H2AC* | 3.3736 | 3.2757 | 0.35928 | -0.78098 | 2.34 |
| 2571 | *CTHRC1* | 2.6644 | 2.3368 | 2.0069 | -1.295 | 2.34 |
| 2572 | *RLN1* | 3.7156 | 2.484 | 0.80838 | 0.49314 | 2.34 |
| 2573 | *GNAQ* | 4.9715 | 1.3944 | 0.64091 | -0.67814 | 2.34 |
| 2574 | *SERPINE2* | 2.8711 | 2.3278 | 1.8079 | -0.41754 | 2.34 |
| 2575 | *CYP26B1* | 4.372 | 2.1467 | 0.48753 | 0.06222 | 2.34 |
| 2576 | *IL1RAP* | 3.4634 | 2.0092 | 1.5329 | 1.3255 | 2.34 |
| 2577 | *C8orf87* | 4.103 | 2.7389 | 0.15971 | -0.00014 | 2.33 |
| 2578 | *MUC7* | 3.0251 | 2.7092 | 1.2665 | 1.0576 | 2.33 |
| 2579 | *FGB* | 4.0113 | 3.2417 | -0.25225 | -1.4256 | 2.33 |
| 2580 | *CCDC144NL* | 5.0334 | 1.1931 | 0.7741 | -1.5517 | 2.33 |
| 2581 | *BACH2* | 3.7188 | 3.481 | -0.20256 | -0.27771 | 2.33 |
| 2582 | *SPRR4* | 5.0586 | 1.5304 | 0.40794 | -2.3145 | 2.33 |
| 2583 | *ARV1* | 4.3773 | 2.6279 | -0.00828 | -1.7297 | 2.33 |
| 2584 | *GLIS2* | 3.1426 | 2.7601 | 1.0938 | -0.17081 | 2.33 |
| 2585 | *PODNL1* | 2.9653 | 2.4771 | 1.5535 | -0.07496 | 2.33 |
| 2586 | *CXCL6* | 4.1077 | 1.4816 | 1.4056 | 1.0734 | 2.33 |
| 2587 | *PTPN2* | 4.6945 | 1.8082 | 0.49202 | -1.3837 | 2.33 |
| 2588 | *INPP5F* | 3.9236 | 2.5646 | 0.50498 | -1.4981 | 2.33 |
| 2589 | *RNF4* | 3.9458 | 2.5725 | 0.47313 | -0.55509 | 2.33 |
| 2590 | *KRTAP10-10* | 3.5398 | 2.5749 | 0.8766 | -1.4469 | 2.33 |
| 2591 | *TCAIM* | 4.8683 | 2.59 | -0.4676 | -0.75669 | 2.33 |
| 2592 | *DNAJB11* | 3.5538 | 2.5389 | 0.89701 | -1.9582 | 2.33 |
| 2593 | *APBA2* | 5.2689 | 1.4973 | 0.22314 | -0.82166 | 2.33 |
| 2594 | *LOXL2* | 3.6522 | 2.4546 | 0.88172 | -1.2013 | 2.33 |
| 2595 | *GJD3* | 3.7828 | 3.2457 | -0.04174 | -0.34535 | 2.33 |
| 2596 | *ACD* | 2.6127 | 2.4614 | 1.912 | -0.55418 | 2.33 |
| 2597 | *VGLL3* | 4.3751 | 2.1253 | 0.48567 | -0.14568 | 2.33 |
| 2598 | *AK3* | 3.7661 | 1.8182 | 1.4015 | -1.9728 | 2.33 |
| 2599 | *GALK1* | 5.0528 | 2.4733 | -0.54155 | -0.8604 | 2.33 |
| 2600 | *DECR1* | 3.8899 | 2.8998 | 0.19425 | -0.10289 | 2.33 |
| 2601 | *ZNF559* | 3.6214 | 1.8312 | 1.5305 | -0.43077 | 2.33 |
| 2602 | *OR6Y1* | 3.8637 | 3.616 | -0.49746 | -0.61334 | 2.33 |
| 2603 | *TNFSF14* | 3.0078 | 2.955 | 1.0177 | -1.2086 | 2.33 |
| 2604 | *ISM2* | 4.5552 | 1.7852 | 0.63918 | -1.2395 | 2.33 |
| 2605 | *MTX2* | 3.8415 | 2.4499 | 0.68801 | -0.45663 | 2.33 |
| 2606 | *CCDC168* | 5.8236 | 2.4 | -1.246 | -2.1687 | 2.33 |
| 2607 | *KRT13* | 4.1852 | 1.5975 | 1.1934 | -0.31934 | 2.33 |
| 2608 | *PPP1R16A* | 4.9564 | 2.0833 | -0.06539 | -1.7194 | 2.32 |
| 2609 | *SLC34A3* | 3.9264 | 3.4813 | -0.43365 | -1.0528 | 2.32 |
| 2610 | *MRPL39* | 4.071 | 2.7257 | 0.17695 | -0.73944 | 2.32 |
| 2611 | *CCDC169* | 4.4484 | 2.558 | -0.03289 | -0.0801 | 2.32 |
| 2612 | *STBD1* | 3.7707 | 3.4014 | -0.19943 | -1.4736 | 2.32 |
| 2613 | *SIRT7* | 4.6483 | 1.6284 | 0.69513 | -1.7001 | 2.32 |
| 2614 | *LCE6A* | 3.6915 | 2.7463 | 0.53265 | -1.215 | 2.32 |
| 2615 | *ZNF860* | 4.0565 | 1.9722 | 0.94162 | -1.6075 | 2.32 |

| | | | | | | |
|---|---|---|---|---|---|---|
| 2616 | *CPT2* | 2.9603 | 2.9063 | 1.1036 | -2.0171 | 2.32 |
| 2617 | *TPCN2* | 2.961 | 2.0534 | 1.9557 | 1.8451 | 2.32 |
| 2618 | *RUSC1* | 4.0261 | 2.9198 | 0.02335 | -0.53031 | 2.32 |
| 2619 | *PTPRT* | 5.3814 | 2.2568 | -0.66914 | -1.2846 | 2.32 |
| 2620 | *HTRA1* | 3.553 | 3.3946 | 0.02124 | -1.3371 | 2.32 |
| 2621 | *DNAJC10* | 2.6654 | 2.4656 | 1.8343 | -1.0007 | 2.32 |
| 2622 | *RDH11* | 4.451 | 2.1833 | 0.3307 | -1.4415 | 2.32 |
| 2623 | *CHRD* | 4.1733 | 1.65 | 1.1411 | -1.9996 | 2.32 |
| 2624 | *GNG11* | 3.8142 | 3.36 | -0.20983 | -1.0099 | 2.32 |
| 2625 | *FBXO40* | 4.5537 | 2.5207 | -0.11076 | -0.80897 | 2.32 |
| 2626 | *PPP6R1* | 4.6496 | 1.4233 | 0.88999 | -0.5605 | 2.32 |
| 2627 | *KRTAP19-7* | 3.3033 | 2.9324 | 0.72714 | -0.95321 | 2.32 |
| 2628 | *TFF2* | 4.8823 | 2.0532 | 0.02701 | -0.97611 | 2.32 |
| 2629 | *ADAMTS15* | 4.5057 | 2.8112 | -0.3554 | -0.41227 | 2.32 |
| 2630 | *ZNF670* | 4.2496 | 1.8628 | 0.84899 | -0.78229 | 2.32 |
| 2631 | *C1QTNF5* | 2.9445 | 2.9171 | 1.0995 | -1.1223 | 2.32 |
| 2632 | *NCAN* | 2.8659 | 2.2191 | 1.8752 | 0.18375 | 2.32 |
| 2633 | *RFX2* | 4.1982 | 2.2257 | 0.53539 | -2.189 | 2.32 |
| 2634 | *HOOK2* | 3.6481 | 2.6223 | 0.68701 | -0.07847 | 2.32 |
| 2635 | *IL4R* | 4.3094 | 2.1922 | 0.45378 | -2.1873 | 2.32 |
| 2636 | *HOXC11* | 3.9097 | 3.2937 | -0.24966 | -1.3976 | 2.32 |
| 2637 | *EPHX2* | 4.3188 | 3.6143 | -0.97952 | -1.7078 | 2.32 |
| 2638 | *PCDHA4* | 4.0905 | 1.7611 | 1.1019 | -1.7836 | 2.32 |
| 2639 | *DHRS12* | 3.6669 | 3.4429 | -0.15787 | -1.9008 | 2.32 |
| 2640 | *C9* | 3.6701 | 2.4661 | 0.81502 | 0.23092 | 2.32 |
| 2641 | *SLC22A18* | 4.3254 | 1.6903 | 0.93499 | -0.01337 | 2.32 |
| 2642 | *DEFB129* | 3.9973 | 2.0661 | 0.88699 | -1.4606 | 2.32 |
| 2643 | *PPP1R3G* | 3.9942 | 2.2822 | 0.67348 | -0.33933 | 2.32 |
| 2644 | *FFAR3* | 5.0604 | 1.8687 | 0.02021 | -0.15455 | 2.32 |
| 2645 | *CCDC179* | 3.9808 | 2.3822 | 0.58593 | -0.95258 | 2.32 |
| 2646 | *RBP5* | 3.4633 | 2.0616 | 1.424 | -2.3949 | 2.32 |
| 2647 | *ELMO2* | 4.0956 | 1.4659 | 1.3866 | -0.86793 | 2.32 |
| 2648 | *ZNF211* | 4.2249 | 2.4855 | 0.2372 | -1.3126 | 2.32 |
| 2649 | *CECR1* | 4.1376 | 3.7028 | -0.89699 | -1.8382 | 2.31 |
| 2650 | *KRTAP19-8* | 3.7758 | 2.9937 | 0.17389 | -0.5287 | 2.31 |
| 2651 | *CFAP97* | 3.0821 | 2.7825 | 1.0787 | -1.3554 | 2.31 |
| 2652 | *SUSD5* | 4.2947 | 2.9022 | -0.25403 | -1.1069 | 2.31 |
| 2653 | *CYP2C9* | 5.0588 | 2.5471 | -0.66445 | -1.0544 | 2.31 |
| 2654 | *LRRC8A* | 4.6143 | 2.2928 | 0.03422 | -2.0436 | 2.31 |
| 2655 | *RBM6* | 3.3969 | 3.3639 | 0.17894 | -0.20673 | 2.31 |
| 2656 | *PPIG* | 5.1261 | 1.5662 | 0.24682 | -1.2845 | 2.31 |
| 2657 | *TAT* | 3.1844 | 2.6879 | 1.0663 | -0.1059 | 2.31 |
| 2658 | *NGF* | 5.2549 | 1.1046 | 0.57803 | -2.1983 | 2.31 |
| 2659 | *TARS2* | 4.9561 | 2.2627 | -0.28159 | -1.0792 | 2.31 |
| 2660 | *AQP10* | 2.7912 | 2.4039 | 1.7413 | -2.8478 | 2.31 |
| 2661 | *PLSCR5* | 4.4743 | 2.4581 | 0.00309 | -3.3806 | 2.31 |

| | | | | | | |
|---|---|---|---|---|---|---|
| 2662 | *ZBTB34* | 3.0364 | 1.9873 | 1.9102 | 0.25651 | 2.31 |
| 2663 | *ASPH* | 3.9701 | 2.8387 | 0.12122 | -1.3666 | 2.31 |
| 2664 | *SLC1A7* | 2.8879 | 2.6499 | 1.3918 | -1.624 | 2.31 |
| 2665 | *C17orf98* | 4.3605 | 2.4361 | 0.13298 | -0.12565 | 2.31 |
| 2666 | *TCEAL7* | 4.0029 | 3.5146 | -0.58806 | -1.1174 | 2.31 |
| 2667 | *SIRPB1* | 3.1309 | 2.6846 | 1.113 | -1.3694 | 2.31 |
| 2668 | *ZKSCAN7* | 4.5625 | 3.971 | -1.6058 | -1.9073 | 2.31 |
| 2669 | *KLC4* | 3.5248 | 2.1136 | 1.2867 | -0.55117 | 2.31 |
| 2670 | *UNC45A* | 4.7013 | 3.0495 | -0.82582 | -1.8909 | 2.31 |
| 2671 | *OR5T3* | 4.0256 | 2.8297 | 0.0695 | -1.164 | 2.31 |
| 2672 | *TRIP4* | 3.9074 | 3.4603 | -0.44374 | -1.3755 | 2.31 |
| 2673 | *URI1* | 3.9269 | 3.5261 | -0.52978 | -1.2567 | 2.31 |
| 2674 | *CLDN17* | 3.4435 | 3.4299 | 0.04956 | -0.735 | 2.31 |
| 2675 | *LOC403312* | 3.6677 | 2.2052 | 1.0498 | -0.35379 | 2.31 |
| 2676 | *PRL* | 5.0734 | 2.7722 | -0.92326 | -1.0669 | 2.31 |
| 2677 | *DBH* | 5.8476 | 1.3634 | -0.2889 | -1.4168 | 2.31 |
| 2678 | *LY96* | 3.0511 | 2.6884 | 1.1823 | -1.9567 | 2.31 |
| 2679 | *C9orf117* | 3.7505 | 2.8328 | 0.33744 | -0.82631 | 2.31 |
| 2680 | *SNX20* | 4.422 | 1.623 | 0.87543 | -0.73198 | 2.31 |
| 2681 | *PKP3* | 4.8694 | 1.3612 | 0.68934 | -0.4778 | 2.31 |
| 2682 | *RAET1G* | 2.7965 | 2.0801 | 2.0424 | -1.5884 | 2.31 |
| 2683 | *EDA* | 3.9191 | 1.7427 | 1.2564 | 0.91314 | 2.31 |
| 2684 | *C19orf24* | 4.3721 | 2.6998 | -0.15499 | -1.0221 | 2.31 |
| 2685 | *CEP104* | 4.838 | 1.1778 | 0.90056 | 0.50176 | 2.31 |
| 2686 | *YDJC* | 3.8134 | 1.7529 | 1.3494 | -0.74604 | 2.31 |
| 2687 | *TTC36* | 5.0516 | 1.2608 | 0.60136 | -2.2495 | 2.30 |
| 2688 | *OR2B11* | 3.2467 | 2.4897 | 1.1756 | 0.87825 | 2.30 |
| 2689 | *SLC35A5* | 4.3669 | 3.0023 | -0.45893 | -0.82573 | 2.30 |
| 2690 | *ZHX1* | 4.0746 | 2.0593 | 0.77557 | 0.066534 | 2.30 |
| 2691 | *CTAGE6* | 3.6786 | 2.5899 | 0.63921 | -1.5848 | 2.30 |
| 2692 | *ZNF395* | 4.575 | 2.3818 | -0.04968 | -0.84523 | 2.30 |
| 2693 | *ZNF600* | 4.5392 | 3.7515 | -1.3837 | -1.4893 | 2.30 |
| 2694 | *TRPC6* | 3.9521 | 2.58 | 0.37386 | -0.50048 | 2.30 |
| 2695 | *MAP4K2* | 4.1764 | 3.576 | -0.84731 | -0.86007 | 2.30 |
| 2696 | *GKN1* | 3.5223 | 2.2329 | 1.1482 | -1.5737 | 2.30 |
| 2697 | *GEM* | 4.3658 | 1.4958 | 1.0401 | -0.45069 | 2.30 |
| 2698 | *HES5* | 3.5605 | 2.478 | 0.86202 | -1.729 | 2.30 |
| 2699 | *NKX2-8* | 3.6264 | 2.2714 | 1.0027 | -0.68514 | 2.30 |
| 2700 | *SLCO2A1* | 4.6677 | 2.163 | 0.06892 | -1.0458 | 2.30 |
| 2701 | *ALG9* | 3.8589 | 2.1525 | 0.88743 | -1.2174 | 2.30 |
| 2702 | *C12orf66* | 4.7702 | 2.3288 | -0.20194 | -0.9728 | 2.30 |
| 2703 | *NEK5* | 3.0242 | 2.4373 | 1.4345 | 1.4129 | 2.30 |
| 2704 | *HTATIP2* | 4.0942 | 1.7197 | 1.0816 | -1.7203 | 2.30 |
| 2705 | *MBLAC2* | 5.1127 | 2.2704 | -0.48821 | -0.78279 | 2.30 |
| 2706 | *HESX1* | 3.7125 | 2.3359 | 0.84631 | -1.4856 | 2.30 |
| 2707 | *GNAI1* | 2.7407 | 2.2976 | 1.8541 | -1.0626 | 2.30 |

| | | | | | | |
|---|---|---|---|---|---|---|
| 2708 | *BIRC6* | 3.147 | 2.5662 | 1.1775 | -3.1673 | 2.30 |
| 2709 | *IL33* | 4.4835 | 2.008 | 0.39643 | -1.7675 | 2.30 |
| 2710 | *LRRC71* | 4.8108 | 2.0702 | 0.00665 | -1.6453 | 2.30 |
| 2711 | *RBM43* | 4.3286 | 1.7574 | 0.8015 | 0.22658 | 2.30 |
| 2712 | *FAM213A* | 2.6145 | 2.484 | 1.7883 | 0.77673 | 2.30 |
| 2713 | *MFAP3* | 3.4545 | 2.4743 | 0.95785 | -1.4789 | 2.30 |
| 2714 | *CORO6* | 3.882 | 2.5821 | 0.42162 | -1.308 | 2.30 |
| 2715 | *ATP11C* | 3.6787 | 2.6805 | 0.526 | -1.6684 | 2.30 |
| 2716 | *PAPD7* | 2.6379 | 2.5287 | 1.7184 | -0.72635 | 2.30 |
| 2717 | *RUNX1* | 4.3414 | 2.2351 | 0.30692 | -0.04053 | 2.29 |
| 2718 | *CC2D2A* | 3.6684 | 2.5447 | 0.66966 | 0.28842 | 2.29 |
| 2719 | *BCL2L2* | 3.794 | 3.1633 | -0.07689 | -0.79389 | 2.29 |
| 2720 | *TRPM8* | 4.3866 | 2.717 | -0.22669 | -0.23418 | 2.29 |
| 2721 | *UBTD2* | 3.7518 | 2.3514 | 0.77362 | -2.0952 | 2.29 |
| 2722 | *PSMB8* | 4.3672 | 3.6134 | -1.1053 | -1.6664 | 2.29 |
| 2723 | *TMEM11* | 3.9764 | 3.1258 | -0.22758 | -0.57573 | 2.29 |
| 2724 | *A2ML1* | 5.1336 | 2.0429 | -0.30218 | -0.81034 | 2.29 |
| 2725 | *EPHA6* | 4.9522 | 1.2183 | 0.70302 | 0.047775 | 2.29 |
| 2726 | *PPIL6* | 3.6589 | 3.1974 | 0.01511 | -1.5386 | 2.29 |
| 2727 | *PRKAR1A* | 3.3935 | 3.3442 | 0.13368 | -2.4205 | 2.29 |
| 2728 | *ASPN* | 3.6154 | 2.4491 | 0.80583 | -0.77184 | 2.29 |
| 2729 | *TRAF4* | 4.0579 | 1.7879 | 1.0245 | -0.28164 | 2.29 |
| 2730 | *ASCL4* | 3.9733 | 1.5994 | 1.2972 | 0.43635 | 2.29 |
| 2731 | *TMPRSS5* | 4.2108 | 2.0899 | 0.56845 | -0.80069 | 2.29 |
| 2732 | *PET100* | 5.0128 | 1.2438 | 0.61161 | 0.091077 | 2.29 |
| 2733 | *KLRC4-KLRK1* | 4.6691 | 2.0803 | 0.11542 | -2.5442 | 2.29 |
| 2734 | *TRIM36* | 3.7027 | 3.6371 | -0.47739 | -0.95717 | 2.29 |
| 2735 | *DHX36* | 3.0282 | 2.6844 | 1.1491 | 0.098411 | 2.29 |
| 2736 | *OR5P3* | 3.2043 | 3.0662 | 0.59007 | -0.0201 | 2.29 |
| 2737 | *HAL* | 3.4556 | 2.522 | 0.88288 | -0.26518 | 2.29 |
| 2738 | *CD276* | 3.3214 | 2.1506 | 1.3869 | -0.99175 | 2.29 |
| 2739 | *CD163* | 4.2243 | 1.8322 | 0.80154 | -0.15394 | 2.29 |
| 2740 | *PKD2L2* | 3.595 | 2.2054 | 1.0465 | 0.82453 | 2.28 |
| 2741 | *IQGAP1* | 3.217 | 2.3999 | 1.2278 | -0.55473 | 2.28 |
| 2742 | *KLHL17* | 4.3251 | 1.42 | 1.0995 | 0.52108 | 2.28 |
| 2743 | *C6orf222* | 4.2696 | 1.4438 | 1.1311 | 0.61322 | 2.28 |
| 2744 | *ABCD3* | 2.8557 | 2.4076 | 1.581 | 0.46801 | 2.28 |
| 2745 | *GKN2* | 3.7347 | 1.9652 | 1.144 | -0.10484 | 2.28 |
| 2746 | *PSTK* | 2.5264 | 2.166 | 2.1509 | -1.3888 | 2.28 |
| 2747 | *SV2C* | 3.2428 | 1.909 | 1.6906 | 0.4766 | 2.28 |
| 2748 | *PCDHGB1* | 5.2491 | 1.386 | 0.20588 | -0.55441 | 2.28 |
| 2749 | *FCRL4* | 4.2751 | 3.0707 | -0.50503 | -0.74069 | 2.28 |
| 2750 | *MAGEB1* | 5.5725 | 1.449 | -0.18124 | -2.071 | 2.28 |
| 2751 | *CHD1* | 3.0461 | 3.0148 | 0.77802 | -1.7278 | 2.28 |
| 2752 | *IL36B* | 3.6491 | 2.0503 | 1.1388 | -1.1646 | 2.28 |
| 2753 | *COLQ* | 2.8726 | 2.694 | 1.2715 | 0.15898 | 2.28 |

| | | | | | | |
|---|---|---|---|---|---|---|
| 2754 | *RUSC2* | 3.9276 | 3.8174 | -0.90835 | -1.3997 | 2.28 |
| 2755 | *PFN3* | 4.6666 | 1.7922 | 0.37684 | -0.98031 | 2.28 |
| 2756 | *GJD4* | 4.6303 | 2.0139 | 0.19132 | -0.44084 | 2.28 |
| 2757 | *RARRES3* | 3.887 | 2.972 | -0.02371 | -0.36315 | 2.28 |
| 2758 | *CUL4B* | 4.1932 | 2.0614 | 0.57945 | -1.631 | 2.28 |
| 2759 | *TEX19* | 4.6417 | 1.2079 | 0.98421 | -1.7845 | 2.28 |
| 2760 | *EIF3K* | 3.5946 | 2.301 | 0.93794 | -0.578 | 2.28 |
| 2761 | *C12orf49* | 2.8014 | 2.1248 | 1.9062 | -0.2531 | 2.28 |
| 2762 | *SMIM13* | 4.0111 | 2.4525 | 0.36853 | -1.1043 | 2.28 |
| 2763 | *PTPN14* | 4.6005 | 2.5237 | -0.29227 | -1.4031 | 2.28 |
| 2764 | *LONRF2* | 3.055 | 2.4948 | 1.2793 | -1.2896 | 2.28 |
| 2765 | *ADRA1A* | 4.91 | 1.9207 | -0.0027 | -1.2617 | 2.28 |
| 2766 | *CHRNB4* | 3.3993 | 2.5485 | 0.87878 | -0.00823 | 2.28 |
| 2767 | *MAP1LC3B* | 3.8924 | 2.9253 | 0.00788 | -1.0573 | 2.28 |
| 2768 | *GATA4* | 3.6447 | 2.1747 | 1.0059 | 0.019931 | 2.28 |
| 2769 | *GPC6* | 4.6872 | 2.7372 | -0.60267 | -0.9703 | 2.27 |
| 2770 | *TMEM81* | 2.7971 | 2.2735 | 1.7502 | -0.58534 | 2.27 |
| 2771 | *PNPLA4* | 4.2475 | 2.7244 | -0.15203 | -1.5255 | 2.27 |
| 2772 | *C5orf66* | 4.4262 | 1.9278 | 0.46462 | 0.44056 | 2.27 |
| 2773 | *STOML3* | 3.473 | 2.7884 | 0.55662 | -0.70014 | 2.27 |
| 2774 | *IL26* | 4.4176 | 3.3597 | -0.9606 | -2.5795 | 2.27 |
| 2775 | *C16orf93* | 4.2941 | 2.919 | -0.39664 | -2.5546 | 2.27 |
| 2776 | *CXorf58* | 3.3695 | 2.7237 | 0.72144 | 0.40396 | 2.27 |
| 2777 | *TMEM265* | 3.4494 | 2.3595 | 1.0026 | -0.12428 | 2.27 |
| 2778 | *SAAL1* | 5.6644 | 2.4231 | -1.2766 | -1.4325 | 2.27 |
| 2779 | *FBXL2* | 4.0137 | 3.0934 | -0.29632 | -0.74833 | 2.27 |
| 2780 | *LYZL4* | 5.294 | 2.4982 | -0.98159 | -1.3504 | 2.27 |
| 2781 | *APEX2* | 3.5432 | 2.5178 | 0.74552 | 0.36493 | 2.27 |
| 2782 | *SERPINE1* | 5.4594 | 1.4088 | -0.06199 | -0.69245 | 2.27 |
| 2783 | *DCAF4* | 4.1312 | 3.0777 | -0.40292 | -1.9959 | 2.27 |
| 2784 | *INIP* | 4.3393 | 2.0955 | 0.37008 | 0.33331 | 2.27 |
| 2785 | *LHCGR* | 2.9006 | 2.4458 | 1.4569 | -0.23375 | 2.27 |
| 2786 | *MAGEA3* | 3.8956 | 2.6635 | 0.24414 | -1.8963 | 2.27 |
| 2787 | *UTS2R* | 3.7633 | 2.5055 | 0.53325 | -1.468 | 2.27 |
| 2788 | *TAS2R20* | 3.8439 | 3.4367 | -0.47884 | -1.2118 | 2.27 |
| 2789 | *DBX1* | 3.6151 | 1.9768 | 1.2096 | -0.49647 | 2.27 |
| 2790 | *CAP2* | 3.4496 | 3.22 | 0.13093 | -0.42195 | 2.27 |
| 2791 | *ITIH1* | 3.8667 | 2.6371 | 0.29191 | 0.25364 | 2.27 |
| 2792 | *COL7A1* | 5.0016 | 2.8595 | -1.0654 | -2.3885 | 2.27 |
| 2793 | *ADCY8* | 3.6523 | 3.557 | -0.41395 | -1.6648 | 2.27 |
| 2794 | *FOXR2* | 4.9849 | 3.0085 | -1.2015 | -1.2572 | 2.26 |
| 2795 | *SMAD3* | 4.0106 | 3.415 | -0.63415 | -1.4311 | 2.26 |
| 2796 | *MKRN1* | 4.2439 | 1.3779 | 1.1689 | 0.9534 | 2.26 |
| 2797 | *NOX5* | 3.9087 | 3.521 | -0.6399 | -0.71669 | 2.26 |
| 2798 | *SCGB2A2* | 3.5012 | 2.7682 | 0.52022 | -0.63709 | 2.26 |
| 2799 | *PAGE3* | 4.003 | 2.5852 | 0.20127 | -1.2951 | 2.26 |

| | | | | | | |
|---|---|---|---|---|---|---|
| 2800 | *CDRT1* | 4.3512 | 2.5235 | -0.08601 | -2.4661 | 2.26 |
| 2801 | *CHMP2B* | 4.5777 | 3.615 | -1.4055 | -1.4614 | 2.26 |
| 2802 | *ZNF749* | 4.2712 | 3.1752 | -0.65946 | -0.86787 | 2.26 |
| 2803 | *TRABD2B* | 2.5747 | 2.2783 | 1.9324 | -0.91427 | 2.26 |
| 2804 | *KIF23* | 3.3515 | 3.1239 | 0.30673 | -2.0614 | 2.26 |
| 2805 | *SYDE1* | 3.1831 | 1.8714 | 1.7273 | 1.1238 | 2.26 |
| 2806 | *DYRK1A* | 4.55 | 2.8036 | -0.57308 | -1.2277 | 2.26 |
| 2807 | *SOX10* | 3.3186 | 1.8114 | 1.6486 | -0.12474 | 2.26 |
| 2808 | *BTLA* | 4.9159 | 3.5618 | -1.6992 | -2.9931 | 2.26 |
| 2809 | *SEMA4C* | 4.3053 | 1.5802 | 0.89273 | -0.28258 | 2.26 |
| 2810 | *BEND6* | 3.346 | 2.7983 | 0.63356 | -1.1766 | 2.26 |
| 2811 | *GUCA1B* | 4.6967 | 1.839 | 0.24169 | -2.5593 | 2.26 |
| 2812 | *PMP2* | 2.4724 | 2.3185 | 1.9863 | 1.027 | 2.26 |
| 2813 | *STX11* | 3.313 | 2.816 | 0.64637 | 0.071769 | 2.26 |
| 2814 | *FAM107A* | 2.849 | 2.0882 | 1.8341 | -5.3303 | 2.26 |
| 2815 | *ENOSF1* | 2.9748 | 1.9052 | 1.8905 | -1.5091 | 2.26 |
| 2816 | *CHMP1A* | 3.0156 | 2.6063 | 1.1485 | 0.6339 | 2.26 |
| 2817 | *ENDOV* | 2.4417 | 2.2696 | 2.0591 | -0.36281 | 2.26 |
| 2818 | *FANCB* | 4.2698 | 1.8769 | 0.62175 | 0.22862 | 2.26 |
| 2819 | *LGALS9C* | 3.3316 | 1.9134 | 1.5219 | -1.4921 | 2.26 |
| 2820 | *PAK1* | 3.8373 | 3.819 | -0.8897 | -1.9023 | 2.26 |
| 2821 | *CCDC117* | 4.3937 | 2.4954 | -0.12259 | -0.40231 | 2.26 |
| 2822 | *GPR135* | 5.376 | 0.72478 | 0.66404 | 0.24793 | 2.25 |
| 2823 | *BACE1* | 2.7923 | 2.113 | 1.8595 | -0.8338 | 2.25 |
| 2824 | *GJB2* | 3.3095 | 2.5607 | 0.89421 | -0.25822 | 2.25 |
| 2825 | *LYPD5* | 3.5959 | 2.7422 | 0.42579 | -1.0149 | 2.25 |
| 2826 | *ARF6* | 3.3497 | 2.9738 | 0.43727 | -1.7119 | 2.25 |
| 2827 | *PTX3* | 4.3858 | 2.2198 | 0.15514 | -1.236 | 2.25 |
| 2828 | *LLGL1* | 3.1852 | 2.0557 | 1.5196 | -0.60032 | 2.25 |
| 2829 | *ICOSLG* | 4.8041 | 1.8623 | 0.09301 | -0.69287 | 2.25 |
| 2830 | *LTBP4* | 2.6785 | 2.3355 | 1.745 | -0.26747 | 2.25 |
| 2831 | *FAM47C* | 4.4245 | 3.5214 | -1.1879 | -1.5165 | 2.25 |
| 2832 | *WI2-2373I1.2* | 5.1626 | 1.6186 | -0.0236 | -1.3183 | 2.25 |
| 2833 | *SPTBN2* | 4.9517 | 1.7803 | 0.0251 | -2.0059 | 2.25 |
| 2834 | *PRR27* | 2.9715 | 1.9385 | 1.8468 | 1.0352 | 2.25 |
| 2835 | *UBC* | 3.5385 | 1.6341 | 1.5837 | 0.78181 | 2.25 |
| 2836 | *PANX1* | 3.4365 | 2.5946 | 0.72441 | -1.8297 | 2.25 |
| 2837 | *ORM2* | 3.5711 | 2.8121 | 0.37157 | -1.6086 | 2.25 |
| 2838 | *APH1A* | 3.8187 | 2.7683 | 0.16742 | 0.083808 | 2.25 |
| 2839 | *NFKBID* | 4.6687 | 1.732 | 0.35364 | 0.099866 | 2.25 |
| 2840 | *KIAA1522* | 3.0109 | 2.6388 | 1.1042 | -0.73155 | 2.25 |
| 2841 | *PKP4* | 4.2968 | 2.968 | -0.51146 | -1.5204 | 2.25 |
| 2842 | *PFKFB3* | 5.2481 | 2.9868 | -1.4829 | -1.6913 | 2.25 |
| 2843 | *ANKRD18A* | 3.6809 | 2.9364 | 0.13166 | -0.84713 | 2.25 |
| 2844 | *ASB2* | 4.1045 | 2.1643 | 0.47991 | -0.33524 | 2.25 |
| 2845 | *ARSJ* | 3.779 | 3.183 | -0.21348 | -1.8154 | 2.25 |

| | | | | | | |
|---|---|---|---|---|---|---|
| 2846 | *PLA2G2F* | 3.8515 | 3.5486 | -0.65169 | -0.78229 | 2.25 |
| 2847 | *S1PR2* | 3.355 | 3.2945 | 0.09832 | -0.55554 | 2.25 |
| 2848 | *SLC8A2* | 5.983 | 0.44198 | 0.32263 | 0.14975 | 2.25 |
| 2849 | *SPRYD7* | 4.4892 | 2.3001 | -0.04343 | -0.64467 | 2.25 |
| 2850 | *CCNI* | 3.3874 | 3.2315 | 0.1264 | -0.38589 | 2.25 |
| 2851 | *LINS* | 5.6091 | 0.90635 | 0.22738 | -0.20758 | 2.25 |
| 2852 | *GPRC5D* | 3.699 | 2.5986 | 0.44429 | -2.0042 | 2.25 |
| 2853 | *MAB21L1* | 3.2961 | 2.0296 | 1.4161 | -1.3296 | 2.25 |
| 2854 | *IL17RD* | 2.8317 | 2.5984 | 1.3095 | -0.35325 | 2.25 |
| 2855 | *SLC30A3* | 5.2126 | 2.7084 | -1.1815 | -3.8687 | 2.25 |
| 2856 | *PLD6* | 5.588 | 1.2832 | -0.13185 | -1.0283 | 2.25 |
| 2857 | *TMEM184A* | 3.5182 | 3.2517 | -0.03153 | -0.97585 | 2.25 |
| 2858 | *FAM155B* | 3.757 | 2.4513 | 0.52983 | -0.811 | 2.25 |
| 2859 | *PARP14* | 3.991 | 2.9222 | -0.17566 | -1.0797 | 2.25 |
| 2860 | *LPAR5* | 2.5367 | 2.364 | 1.8367 | -0.74428 | 2.25 |
| 2861 | *ETFDH* | 3.4616 | 2.9077 | 0.3679 | -2.1066 | 2.25 |
| 2862 | *PLIN4* | 3.6036 | 3.0056 | 0.12787 | -0.42661 | 2.25 |
| 2863 | *FREM2* | 3.976 | 3.4452 | -0.68466 | -1.2291 | 2.25 |
| 2864 | *CCSER1* | 3.8937 | 3.5964 | -0.75369 | -1.0014 | 2.25 |
| 2865 | *SLC25A44* | 3.1285 | 2.1005 | 1.5073 | 1.4622 | 2.25 |
| 2866 | *CYP2D6* | 3.7051 | 2.9198 | 0.11095 | -1.0704 | 2.25 |
| 2867 | *SSX7* | 3.7215 | 1.7296 | 1.2832 | -2.401 | 2.24 |
| 2868 | *C9orf153* | 3.719 | 3.272 | -0.25854 | -0.75173 | 2.24 |
| 2869 | *MAGEA5* | 4.0851 | 2.5767 | 0.07047 | -1.9624 | 2.24 |
| 2870 | *OR1L3* | 3.1822 | 2.8513 | 0.69842 | -0.00317 | 2.24 |
| 2871 | *FAM174A* | 3.278 | 2.2209 | 1.2329 | -1.0139 | 2.24 |
| 2872 | *CGB7* | 4.4641 | 1.4263 | 0.84021 | -2.0384 | 2.24 |
| 2873 | *SUPT20H* | 3.5608 | 2.7452 | 0.42438 | 0.21115 | 2.24 |
| 2874 | *FAM195A* | 3.2586 | 3.1712 | 0.30034 | -1.8851 | 2.24 |
| 2875 | *GATA1* | 4.9577 | 3.5864 | -1.8154 | -2.224 | 2.24 |
| 2876 | *ECM2* | 3.2868 | 1.9867 | 1.4547 | 0.33169 | 2.24 |
| 2877 | *SPHAR* | 5.1119 | -0.62666 | | | 2.24 |
| 2878 | *OR4D1* | 2.6392 | 2.3775 | 1.7104 | -0.97667 | 2.24 |
| 2879 | *WDSUB1* | 4.1818 | 2.334 | 0.2113 | -1.9495 | 2.24 |
| 2880 | *ADAM28* | 3.9189 | 2.7311 | 0.07667 | -0.66462 | 2.24 |
| 2881 | *ACP2* | 4.4057 | 4.2028 | -1.882 | -5.3303 | 2.24 |
| 2882 | *CLMP* | 3.4438 | 3.1143 | 0.16765 | -1.237 | 2.24 |
| 2883 | *LAX1* | 4.3913 | 2.434 | -0.09965 | -1.0662 | 2.24 |
| 2884 | *AMELX* | 4.2872 | 1.3563 | 1.0819 | 0.24364 | 2.24 |
| 2885 | *MOB4* | 3.9278 | 2.7053 | 0.09207 | -0.31145 | 2.24 |
| 2886 | *PDK1* | 4.185 | 1.5123 | 1.0277 | -0.14425 | 2.24 |
| 2887 | *EDN1* | 5.3522 | 1.0295 | 0.34077 | 0.33438 | 2.24 |
| 2888 | *VIPAS39* | 3.5604 | 1.6357 | 1.5244 | -0.71936 | 2.24 |
| 2889 | *CT47B1* | 3.1091 | 2.9189 | 0.69213 | 0.033607 | 2.24 |
| 2890 | *TTC39A* | 4.66 | 2.1407 | -0.08232 | -1.3916 | 2.24 |
| 2891 | *TRPM2* | 3.9497 | 2.9833 | -0.21532 | -1.0087 | 2.24 |

| | | | | | | |
|---|---|---|---|---|---|---|
| 2892 | *NLGN4X* | 3.4407 | 2.3454 | 0.9314 | 0.83177 | 2.24 |
| 2893 | *CDH6* | 3.0721 | 2.8036 | 0.84066 | -1.6559 | 2.24 |
| 2894 | *TM2D2* | 2.9034 | 2.5638 | 1.2484 | -2.1523 | 2.24 |
| 2895 | *SEC14L4* | 5.1337 | 1.4784 | 0.10337 | -0.27451 | 2.24 |
| 2896 | *LRP5* | 2.7033 | 2.5008 | 1.5109 | 0.39741 | 2.24 |
| 2897 | *PLXNB3* | 4.3104 | 3.295 | -0.89194 | -0.97227 | 2.24 |
| 2898 | *ASPM* | 5.1938 | 2.3585 | -0.83912 | -1.5031 | 2.24 |
| 2899 | *RAET1L* | 3.6549 | 2.4008 | 0.65618 | -0.47186 | 2.24 |
| 2900 | *DEFB107B* | 2.2371 | | | | 2.24 |
| 2901 | *ZNF273* | 3.7936 | 1.742 | 1.1733 | -0.4013 | 2.24 |
| 2902 | *ADGRL4* | 2.6693 | 2.07 | 1.9687 | -1.9624 | 2.24 |
| 2903 | *ACSF3* | 2.7238 | 2.4431 | 1.5405 | -0.40785 | 2.24 |
| 2904 | *CRLS1* | 4.7157 | 3.0445 | -1.053 | -1.8575 | 2.24 |
| 2905 | *TRMT1L* | 3.7541 | 2.8201 | 0.13231 | -1.4233 | 2.24 |
| 2906 | *THEG5* | 5.0315 | 1.0778 | 0.59644 | -1.2284 | 2.24 |
| 2907 | *MUM1L1* | 3.2704 | 1.8538 | 1.5812 | -1.2417 | 2.24 |
| 2908 | *TYW5* | 2.713 | 2.5111 | 1.4812 | -0.77196 | 2.24 |
| 2909 | *OR51I2* | 3.4322 | 1.9168 | 1.3546 | -0.07232 | 2.23 |
| 2910 | *MAT2A* | 3.6556 | 2.5094 | 0.53703 | -0.95064 | 2.23 |
| 2911 | *GALNT3* | 4.6717 | 2.8532 | -0.82368 | -1.7512 | 2.23 |
| 2912 | *AKR1C1* | 5.8028 | 0.64061 | 0.25776 | -2.5007 | 2.23 |
| 2913 | *HMGB3* | 5.6489 | 0.72379 | 0.32753 | -1.1233 | 2.23 |
| 2914 | *HIC1* | 4.3879 | 3.1348 | -0.82272 | -1.2912 | 2.23 |
| 2915 | *ETV3L* | 3.597 | 3.4194 | -0.31815 | -2.0978 | 2.23 |
| 2916 | *OXT* | 3.3259 | 2.0758 | 1.2964 | -1.1211 | 2.23 |
| 2917 | *ACCS* | 3.0802 | 2.8085 | 0.8092 | -0.68565 | 2.23 |
| 2918 | *RNF220* | 3.2907 | 2.2256 | 1.1814 | -0.38163 | 2.23 |
| 2919 | *C20orf85* | 2.4985 | 2.479 | 1.7199 | -1.3593 | 2.23 |
| 2920 | *RNF17* | 4.9221 | 1.5917 | 0.18297 | -1.0633 | 2.23 |
| 2921 | *ERCC6L* | 3.7929 | 2.5984 | 0.30419 | -0.53437 | 2.23 |
| 2922 | *HIGD1B* | 3.975 | 1.5526 | 1.1674 | -1.2456 | 2.23 |
| 2923 | *LOC101928093* | 4.1719 | 2.8625 | -0.34093 | -1.5739 | 2.23 |
| 2924 | *PNPLA7* | 5.3939 | 2.2353 | -0.93789 | -2.7214 | 2.23 |
| 2925 | *UBE3C* | 4.5007 | 2.8324 | -0.64229 | -1.2691 | 2.23 |
| 2926 | *PELP1* | 5.3609 | 1.7477 | -0.41803 | -0.68606 | 2.23 |
| 2927 | *HRH3* | 3.3198 | 2.0529 | 1.3164 | -1.0189 | 2.23 |
| 2928 | *XKRX* | 5.2407 | 1.3529 | 0.09422 | -0.87662 | 2.23 |
| 2929 | *C16orf78* | 3.204 | 2.487 | 0.99541 | 0.19092 | 2.23 |
| 2930 | *FOXI1* | 3.1128 | 2.6812 | 0.89213 | 0.63025 | 2.23 |
| 2931 | *THNSL2* | 3.4615 | 2.7888 | 0.43502 | -0.38677 | 2.23 |
| 2932 | *LOC100130451* | 3.9784 | 2.2624 | 0.44385 | 0.42014 | 2.23 |
| 2933 | *MICU1* | 5.3741 | 0.71749 | 0.5922 | -0.80379 | 2.23 |
| 2934 | *SLC25A29* | 4.053 | 2.7754 | -0.1449 | -1.8318 | 2.23 |
| 2935 | *CNKSR1* | 4.4594 | 2.4574 | -0.23445 | -0.47726 | 2.23 |
| 2936 | *DNAJC5G* | 4.3817 | 1.5814 | 0.71918 | -0.53406 | 2.23 |
| 2937 | *LAMTOR2* | 2.9882 | 2.1877 | 1.5058 | 1.094 | 2.23 |

| | | | | | | |
|---|---|---|---|---|---|---|
| 2938 | *FNDC3B* | 3.8915 | 2.8404 | -0.05104 | -0.53036 | 2.23 |
| 2939 | *SCYL2* | 2.8305 | 2.0834 | 1.7663 | 1.5354 | 2.23 |
| 2940 | *C4orf29* | 5.6751 | 0.65926 | 0.34578 | -2.3517 | 2.23 |
| 2941 | *CXorf66* | 3.2785 | 2.5967 | 0.80183 | -1.0948 | 2.23 |
| 2942 | *CPPED1* | 3.6745 | 1.6303 | 1.3709 | -1.4745 | 2.23 |
| 2943 | *SLC44A1* | 4.5892 | 3.9237 | -1.8376 | -2.0062 | 2.23 |
| 2944 | *FBXO46* | 3.2942 | 1.8321 | 1.5477 | -0.91134 | 2.22 |
| 2945 | *FAM168A* | 2.9744 | 2.3234 | 1.3754 | -0.00823 | 2.22 |
| 2946 | *LYSMD1* | 5.279 | 1.7789 | -0.38611 | -0.97252 | 2.22 |
| 2947 | *RBAK* | 2.9097 | 2.8122 | 0.94988 | 0.4969 | 2.22 |
| 2948 | *CCDC85A* | 3.2856 | 2.6361 | 0.74769 | 0.68386 | 2.22 |
| 2949 | *STMN1* | 5.5804 | 0.60294 | 0.48494 | -1.2301 | 2.22 |
| 2950 | *SEC24C* | 5.9597 | 0.64923 | 0.05766 | -0.18568 | 2.22 |
| 2951 | *ANKRD6* | 3.7638 | 2.5288 | 0.37314 | -1.4401 | 2.22 |
| 2952 | *ARSK* | 3.4223 | 3.3347 | -0.09253 | -0.5018 | 2.22 |
| 2953 | *CRYZL1* | 4.1079 | 2.1854 | 0.37085 | -0.60082 | 2.22 |
| 2954 | *SDCBP2* | 4.456 | 2.6099 | -0.40577 | -0.65557 | 2.22 |
| 2955 | *CYBA* | 3.6515 | 3.0901 | -0.08234 | -2.058 | 2.22 |
| 2956 | *DENND6B* | 3.6318 | 3.081 | -0.05603 | -2.1418 | 2.22 |
| 2957 | *FZD6* | 3.4519 | 2.9047 | 0.29904 | -1.7662 | 2.22 |
| 2958 | *DIRAS1* | 5.0422 | 0.88303 | 0.7291 | 0.43955 | 2.22 |
| 2959 | *ZNF534* | 5.0523 | 1.4637 | 0.13811 | 0.053745 | 2.22 |
| 2960 | *OR7C2* | 4.6561 | 1.4071 | 0.59087 | 0.39993 | 2.22 |
| 2961 | *FUNDC1* | 3.3212 | 2.7707 | 0.56196 | -0.98717 | 2.22 |
| 2962 | *EDF1* | 3.6095 | 2.8992 | 0.14491 | -3.2979 | 2.22 |
| 2963 | *C6orf106* | 3.8779 | 3.4435 | -0.66787 | -1.3148 | 2.22 |
| 2964 | *ACKR3* | 4.4758 | 1.7987 | 0.37887 | 0.099654 | 2.22 |
| 2965 | *TLE1* | 3.0589 | 1.9355 | 1.6588 | -0.7829 | 2.22 |
| 2966 | *CRIP2* | 2.8989 | 2.0003 | 1.7523 | -1.5744 | 2.22 |
| 2967 | *CHURC1-FNTB* | 3.823 | 3.2711 | -0.44353 | -2.1547 | 2.22 |
| 2968 | *SERGEF* | 3.0403 | 1.9516 | 1.6578 | -0.89851 | 2.22 |
| 2969 | *ZNF560* | 3.1934 | 1.7729 | 1.6817 | 1.5276 | 2.22 |
| 2970 | *ZSCAN30* | 3.8543 | 3.6817 | -0.89 | -2.485 | 2.22 |
| 2971 | *TMEM255B* | 4.6106 | 2.7094 | -0.67504 | -0.77627 | 2.21 |
| 2972 | *APIP* | 5.5515 | 0.75238 | 0.34102 | -1.0198 | 2.21 |
| 2973 | *ZNF544* | 5.0746 | 1.8552 | -0.28761 | -0.77934 | 2.21 |
| 2974 | *APBB3* | 4.5868 | 1.6806 | 0.37475 | -1.4291 | 2.21 |
| 2975 | *LVRN* | 3.6579 | 2.6495 | 0.33377 | -0.74505 | 2.21 |
| 2976 | *ZNF714* | 4.9497 | 0.9731 | 0.71807 | -1.5876 | 2.21 |
| 2977 | *EZH1* | 3.9161 | 2.2846 | 0.43954 | -0.19834 | 2.21 |
| 2978 | *ZNF80* | 3.9936 | 3.8464 | -1.1998 | -2.8553 | 2.21 |
| 2979 | *TMEM189* | 4.0294 | 3.2635 | -0.65545 | -3.0312 | 2.21 |
| 2980 | *CA11* | 4.5531 | 1.0975 | 0.98576 | -0.67853 | 2.21 |
| 2981 | *LGALS1* | 4.1607 | 1.7749 | 0.70052 | 0.35737 | 2.21 |
| 2982 | *ABCA8* | 4.2631 | 3.4157 | -1.0433 | -1.714 | 2.21 |
| 2983 | *FGF1* | 4.0281 | 3.7002 | -1.0951 | -1.6939 | 2.21 |

| | | | | | | |
|---|---|---|---|---|---|---|
| 2984 | *SUDS3* | 4.3316 | 3.337 | -1.0359 | -1.3565 | 2.21 |
| 2985 | *KRR1* | 3.9151 | 2.7789 | -0.062 | -1.7705 | 2.21 |
| 2986 | *KRTAP20-3* | 4.2729 | 3.4314 | -1.0726 | -1.4926 | 2.21 |
| 2987 | *THYN1* | 4.4267 | 2.4619 | -0.25753 | -0.27491 | 2.21 |
| 2988 | *LYPD3* | 2.7845 | 2.352 | 1.4924 | -0.01111 | 2.21 |
| 2989 | *GPR139* | 3.3747 | 2.3837 | 0.86834 | -1.1306 | 2.21 |
| 2990 | *ZNF771* | 4.4581 | 2.6119 | -0.44353 | -1.195 | 2.21 |
| 2991 | *FLRT3* | 5.863 | 1.5029 | -0.73961 | -1.95 | 2.21 |
| 2992 | *ME1* | 3.9143 | 1.5892 | 1.1224 | -1.5791 | 2.21 |
| 2993 | *HMGN3* | 4.2834 | 2.809 | -0.4677 | -0.92623 | 2.21 |
| 2994 | *OR2T33* | 4.7379 | 1.2906 | 0.59588 | 0.17958 | 2.21 |
| 2995 | *ACRBP* | 3.7776 | 2.9091 | -0.06337 | -2.7245 | 2.21 |
| 2996 | *COL22A1* | 3.4267 | 1.8875 | 1.3083 | -0.6726 | 2.21 |
| 2997 | *LILRB3* | 4.42 | 2.3077 | -0.1059 | -1.1021 | 2.21 |
| 2998 | *LYPD4* | 3.2074 | 1.942 | 1.4722 | 0.4371 | 2.21 |
| 2999 | *MAGOHB* | 4.593 | 1.3972 | 0.63011 | 0.57106 | 2.21 |
| 3000 | *CYP4F22* | 5.34 | 1.8649 | -0.58495 | -1.2572 | 2.21 |
| 3001 | *AGER* | 3.4532 | 2.043 | 1.1207 | -1.2668 | 2.21 |
| 3002 | *FCER1A* | 3.1815 | 1.9301 | 1.503 | 1.29 | 2.20 |
| 3003 | *CA8* | 3.9644 | 1.5507 | 1.0991 | -2.0851 | 2.20 |
| 3004 | *CSRP2BP* | 4.1539 | 1.5204 | 0.93858 | 0.49511 | 2.20 |
| 3005 | *GAPDHS* | 4.3547 | 3.0823 | -0.8248 | -1.0691 | 2.20 |
| 3006 | *C1QC* | 4.1894 | 1.7235 | 0.69816 | -0.14957 | 2.20 |
| 3007 | *AZIN2* | 4.2286 | 2.1303 | 0.25152 | -0.62153 | 2.20 |
| 3008 | *ARHGAP35* | 2.927 | 2.3492 | 1.3338 | -1.6176 | 2.20 |
| 3009 | *CALD1* | 4.1148 | 3.1363 | -0.6421 | -1.0046 | 2.20 |
| 3010 | *C12orf29* | 3.7694 | 1.9915 | 0.84804 | -2.5878 | 2.20 |
| 3011 | *OLFML2B* | 2.8516 | 2.4626 | 1.2929 | 1.1299 | 2.20 |
| 3012 | *GINS1* | 4.2654 | 3.3218 | -0.98032 | -1.377 | 2.20 |
| 3013 | *OAT* | 3.7451 | 3.4391 | -0.57927 | -2.1404 | 2.20 |
| 3014 | *EZH2* | 4.587 | 2.4271 | -0.40976 | -1.7591 | 2.20 |
| 3015 | *HAGHL* | 4.6692 | 2.0741 | -0.13928 | -1.1055 | 2.20 |
| 3016 | *CAPN15* | 4.3446 | 1.6959 | 0.56328 | -0.30971 | 2.20 |
| 3017 | *LOC100132146* | 4.0602 | 3.4701 | -0.92764 | -1.0868 | 2.20 |
| 3018 | *TSSC4* | 5.0991 | 1.7714 | -0.26832 | -1.5848 | 2.20 |
| 3019 | *LRPPRC* | 3.7626 | 2.595 | 0.24438 | -0.36536 | 2.20 |
| 3020 | *SIRPD* | 3.1687 | 2.9591 | 0.47336 | -1.9622 | 2.20 |
| 3021 | *SPRR2E* | 3.7479 | 2.4305 | 0.4223 | -2.6343 | 2.20 |
| 3022 | *ERP29* | 3.6863 | 3.0999 | -0.18667 | -1.8348 | 2.20 |
| 3023 | *KCNIP4* | 4.6374 | 2.3666 | -0.40467 | -1.2633 | 2.20 |
| 3024 | *PTPRO* | 2.9977 | 2.9415 | 0.65812 | -0.04332 | 2.20 |
| 3025 | *MCAM* | 3.6799 | 3.3842 | -0.46705 | -0.60863 | 2.20 |
| 3026 | *NYAP2* | 5.3687 | 0.75048 | 0.47728 | 0.17607 | 2.20 |
| 3027 | *ATRX* | 4.5239 | 3.2183 | -1.146 | -1.8073 | 2.20 |
| 3028 | *H2BFWT* | 4.4824 | 2.1438 | -0.03089 | -0.11566 | 2.20 |
| 3029 | *USP34* | 3.4221 | 3.2198 | -0.04736 | -1.3574 | 2.20 |

| | | | | | | |
|---|---|---|---|---|---|---|
| 3030 | *CLEC2B* | 5.0557 | 2.7959 | -1.2574 | -1.8649 | 2.20 |
| 3031 | *DUSP28* | 3.287 | 3.0034 | 0.30301 | -0.02618 | 2.20 |
| 3032 | *CACNA1H* | 2.5908 | 2.5584 | 1.4436 | -1.1575 | 2.20 |
| 3033 | *SLC4A1AP* | 5.2417 | 0.69496 | 0.65491 | -0.69683 | 2.20 |
| 3034 | *RPS3A* | 4.5128 | 2.2397 | -0.16128 | -0.794 | 2.20 |
| 3035 | *NETO1* | 4.6972 | 1.9015 | -0.00824 | -0.85222 | 2.20 |
| 3036 | *CYTH2* | 3.66 | 2.3148 | 0.61561 | -1.6413 | 2.20 |
| 3037 | *NIN* | 3.5086 | 1.8126 | 1.2691 | -0.15079 | 2.20 |
| 3038 | *MINPP1* | 4.9864 | 1.3227 | 0.2809 | -1.3197 | 2.20 |
| 3039 | *GOLGA2* | 3.8227 | 3.76 | -0.99418 | -2.2117 | 2.20 |
| 3040 | *LOC730183* | 5.2615 | 1.8899 | -0.5639 | -2.3623 | 2.20 |
| 3041 | *LARGE* | 4.6811 | 1.9079 | -0.00197 | -0.59838 | 2.20 |
| 3042 | *UNC13B* | 2.46 | 2.3391 | 1.7871 | 1.4941 | 2.20 |
| 3043 | *ANO2* | 3.4435 | 2.0112 | 1.1303 | -1.2276 | 2.20 |
| 3044 | *OPN1MW2* | 2.5919 | 2.2205 | 1.7722 | -1.584 | 2.19 |
| 3045 | *S100B* | 2.7231 | 1.9348 | 1.9256 | 0.033516 | 2.19 |
| 3046 | *ZNF718* | 4.1292 | 1.7488 | 0.70468 | -0.54307 | 2.19 |
| 3047 | *ARSG* | 3.5492 | 3.4168 | -0.38378 | -1.6444 | 2.19 |
| 3048 | *TUB* | 3.2004 | 2.2744 | 1.1073 | -1.2753 | 2.19 |
| 3049 | *SLC31A2* | 3.453 | 2.6378 | 0.48995 | -0.36861 | 2.19 |
| 3050 | *ZNF423* | 3.4545 | 1.9065 | 1.2193 | 0.25863 | 2.19 |
| 3051 | *SMARCA2* | 3.0534 | 2.7182 | 0.80786 | -2.1541 | 2.19 |
| 3052 | *JMY* | 2.8769 | 2.6477 | 1.0544 | 0.37483 | 2.19 |
| 3053 | *TACC1* | 4.9867 | 1.3594 | 0.23176 | -0.05035 | 2.19 |
| 3054 | *GLP2R* | 4.3652 | 1.9028 | 0.30977 | 0.031433 | 2.19 |
| 3055 | *RABL2B* | 3.6784 | 2.4452 | 0.45396 | -0.19566 | 2.19 |
| 3056 | *ZAK* | 3.8415 | 1.9937 | 0.74157 | -1.2051 | 2.19 |
| 3057 | *PHB2* | 3.0367 | 2.817 | 0.72192 | -2.2433 | 2.19 |
| 3058 | *LARP1B* | 3.5174 | 3.2189 | -0.16195 | -1.8219 | 2.19 |
| 3059 | *DNAJB8* | 3.4665 | 3.1378 | -0.0304 | -0.23196 | 2.19 |
| 3060 | *MCM9* | 3.6992 | 3.6313 | -0.75762 | -2.2948 | 2.19 |
| 3061 | *PUS1* | 3.5531 | 2.1334 | 0.8845 | -0.50775 | 2.19 |
| 3062 | *ZC3HAV1* | 3.4928 | 1.7699 | 1.3068 | 0.44713 | 2.19 |
| 3063 | *KRTAP19-6* | 5.3939 | 1.2121 | -0.03724 | -1.5212 | 2.19 |
| 3064 | *OR7D2* | 4.8355 | 3.0229 | -1.2903 | -1.7804 | 2.19 |
| 3065 | *MYOM2* | 3.4633 | 3.4373 | -0.33266 | -1.9172 | 2.19 |
| 3066 | *C18orf65* | 3.6994 | 3.0145 | -0.14627 | -0.18896 | 2.19 |
| 3067 | *LHX1* | 3.4532 | 2.7736 | 0.33886 | -0.44353 | 2.19 |
| 3068 | *BRD1* | 3.3901 | 3.0077 | 0.16772 | -0.41502 | 2.19 |
| 3069 | *IPPK* | 3.7399 | 2.1824 | 0.64312 | -0.41567 | 2.19 |
| 3070 | *HELT* | 3.9654 | 1.3946 | 1.2044 | -0.61027 | 2.19 |
| 3071 | *CD84* | 3.9193 | 1.5496 | 1.0945 | -0.20594 | 2.19 |
| 3072 | *SYNPO* | 2.8966 | 2.1051 | 1.561 | 0.60261 | 2.19 |
| 3073 | *KIAA1407* | 4.3981 | 1.3932 | 0.77129 | -0.92025 | 2.19 |
| 3074 | *CDKN1C* | 3.3262 | 3.029 | 0.20665 | -1.9166 | 2.19 |
| 3075 | *TBC1D25* | 3.1815 | 2.9642 | 0.41564 | -1.6213 | 2.19 |

| | | | | | | |
|---|---|---|---|---|---|---|
| 3076 | *SLC39A12* | 4.252 | 2.2938 | 0.01504 | -1.4682 | 2.19 |
| 3077 | *RAB1B* | 3.3388 | 2.7554 | 0.46469 | -1.0148 | 2.19 |
| 3078 | *BMP5* | 3.696 | 3.1646 | -0.30258 | -0.34227 | 2.19 |
| 3079 | *DUS2* | 3.9074 | 1.4764 | 1.1739 | -1.3523 | 2.19 |
| 3080 | *TMEM132E* | 3.8945 | 2.1709 | 0.49221 | -1.6346 | 2.19 |
| 3081 | *C10orf62* | 3.2522 | 2.6021 | 0.70287 | -1.0544 | 2.19 |
| 3082 | *ALDOA* | 2.8432 | 2.2983 | 1.415 | -0.64037 | 2.19 |
| 3083 | *CCDC91* | 3.1194 | 1.7716 | 1.6629 | 0.43843 | 2.18 |
| 3084 | *VSIG4* | 3.8252 | 1.5191 | 1.2083 | -0.7853 | 2.18 |
| 3085 | *CDC42BPG* | 4.2559 | 1.7606 | 0.53427 | -0.41703 | 2.18 |
| 3086 | *TUBB2B* | 3.1471 | 3.0327 | 0.36899 | -1.6824 | 2.18 |
| 3087 | *ZNF605* | 5.3772 | 0.88613 | 0.28491 | -0.42899 | 2.18 |
| 3088 | *TPP1* | 3.1314 | 2.5608 | 0.85534 | 0.26891 | 2.18 |
| 3089 | *NUTF2* | 4.8968 | 1.47 | 0.18033 | -1.035 | 2.18 |
| 3090 | *PDGFRA* | 4.1805 | 2.6471 | -0.28384 | -2.7139 | 2.18 |
| 3091 | *MEMO1* | 4.7972 | 1.8928 | -0.14649 | -0.25056 | 2.18 |
| 3092 | *KRIT1* | 5.0059 | 1.7619 | -0.22611 | -1.2008 | 2.18 |
| 3093 | *SIDT1* | 5.3747 | 1.6037 | -0.43674 | -0.88584 | 2.18 |
| 3094 | *ZFAND2A* | 4.3508 | 2.3483 | -0.15963 | -1.6804 | 2.18 |
| 3095 | *TCL1B* | 2.807 | 2.481 | 1.2501 | -0.45544 | 2.18 |
| 3096 | *PPAP2C* | 3.7719 | 3.5931 | -0.83101 | -1.416 | 2.18 |
| 3097 | *KL* | 4.5782 | 2.5582 | -0.60267 | -0.85002 | 2.18 |
| 3098 | *LUM* | 4.2915 | 2.1805 | 0.06165 | -1.7466 | 2.18 |
| 3099 | *RNF144A* | 4.8473 | 0.8516 | 0.8346 | 0.62248 | 2.18 |
| 3100 | *ARL5C* | 3.3639 | 2.6745 | 0.4944 | 0.084092 | 2.18 |
| 3101 | *PCDHGA11* | 4.7123 | 1.8002 | 0.01988 | -0.98194 | 2.18 |
| 3102 | *UBR2* | 3.2118 | 2.5574 | 0.76069 | 0.6088 | 2.18 |
| 3103 | *TUBB3* | 3.8079 | 1.8157 | 0.90472 | -0.09645 | 2.18 |
| 3104 | *PCDHGB3* | 3.8793 | 2.0695 | 0.57936 | 0.07696 | 2.18 |
| 3105 | *PRTFDC1* | 2.5672 | 2.284 | 1.6768 | 0.71705 | 2.18 |
| 3106 | *ODAM* | 3.3303 | 2.1699 | 1.0277 | -0.68388 | 2.18 |
| 3107 | *LCA5* | 4.9442 | 1.2066 | 0.37603 | -0.53481 | 2.18 |
| 3108 | *IRF4* | 3.2574 | 3.2207 | 0.04858 | -1.5304 | 2.18 |
| 3109 | *GSTA2* | 3.5992 | 3.1335 | -0.20606 | -1.2993 | 2.18 |
| 3110 | *SMIM1* | 4.5687 | 1.4614 | 0.49584 | -0.34014 | 2.18 |
| 3111 | *DOK6* | 4.0833 | 1.3583 | 1.0841 | -0.31633 | 2.18 |
| 3112 | *CHCHD7* | 4.6266 | 2.4778 | -0.5794 | -0.71561 | 2.18 |
| 3113 | *C15orf26* | 4.9638 | 0.86799 | 0.69116 | -1.6776 | 2.17 |
| 3114 | *POLR2J2* | 4.3351 | 1.3736 | 0.81187 | -1.8532 | 2.17 |
| 3115 | *WT1* | 3.1163 | 3.0487 | 0.35396 | -0.96407 | 2.17 |
| 3116 | *NKX2-3* | 3.1491 | 2.9301 | 0.43916 | -0.8828 | 2.17 |
| 3117 | *TRIM9* | 3.4132 | 3.0581 | 0.04597 | -0.3064 | 2.17 |
| 3118 | *KRTAP6-2* | 3.1949 | 1.7207 | 1.6009 | 0.15863 | 2.17 |
| 3119 | *MNX1* | 3.9389 | 2.4082 | 0.16852 | -1.0288 | 2.17 |
| 3120 | *IFI27L2* | 4.8358 | 1.1557 | 0.52293 | -0.55117 | 2.17 |
| 3121 | *TEX36* | 3.6162 | 3.0753 | -0.17772 | -0.26798 | 2.17 |

| | | | | | | |
|---|---|---|---|---|---|---|
| 3122 | *FAM71F2* | 3.4644 | 2.3263 | 0.72294 | -0.39004 | 2.17 |
| 3123 | *UNG* | 3.8423 | 3.0847 | -0.41738 | -1.1845 | 2.17 |
| 3124 | *ZNF789* | 4.4057 | 1.6659 | 0.43769 | -1.3617 | 2.17 |
| 3125 | *BSCL2* | 4.3096 | 3.7886 | -1.589 | -2.0613 | 2.17 |
| 3126 | *UGT2B4* | 2.5832 | 2.3542 | 1.5718 | 0.97673 | 2.17 |
| 3127 | *CCR5* | 4.7985 | 2.9453 | -1.2371 | -3.038 | 2.17 |
| 3128 | *SPRR2F* | 4.6285 | 0.94463 | 0.93279 | -0.19661 | 2.17 |
| 3129 | *RABEPK* | 5.2937 | 2.3789 | -1.1667 | -1.2457 | 2.17 |
| 3130 | *BCL2A1* | 3.3999 | 2.0307 | 1.0744 | 0.98887 | 2.17 |
| 3131 | *ATP6V0A2* | 4.5457 | 2.6311 | -0.6728 | -1.28 | 2.17 |
| 3132 | *FAM81B* | 4.5036 | 1.8466 | 0.15352 | -1.0596 | 2.17 |
| 3133 | *PRPSAP1* | 4.1203 | 3.0167 | -0.63399 | -2.0952 | 2.17 |
| 3134 | *DAZAP2* | 4.3551 | 2.4888 | -0.34118 | -0.4055 | 2.17 |
| 3135 | *ZIC4* | 5.3999 | 2.424 | -1.3218 | -1.8318 | 2.17 |
| 3136 | *ZNF641* | 4.4716 | 1.5101 | 0.51921 | -0.81524 | 2.17 |
| 3137 | *FKBP8* | 3.953 | 1.6314 | 0.91428 | -0.57797 | 2.17 |
| 3138 | *LOC554223* | 4.0919 | 1.3563 | 1.0503 | -1.232 | 2.17 |
| 3139 | *SLC16A6* | 4.3887 | 2.8282 | -0.71945 | -0.87211 | 2.17 |
| 3140 | *HDX* | 5.0143 | 0.88265 | 0.5997 | 0.14429 | 2.17 |
| 3141 | *OR5B3* | 4.456 | 2.8151 | -0.77562 | -1.0435 | 2.17 |
| 3142 | *TRABD2A* | 4.8948 | 1.4015 | 0.19802 | -0.14663 | 2.16 |
| 3143 | *ZNF675* | 3.4601 | 3.2048 | -0.1715 | -1.8076 | 2.16 |
| 3144 | *GPR35* | 3.2372 | 3.0601 | 0.19565 | 0.13811 | 2.16 |
| 3145 | *UPRT* | 6.1376 | 0.23591 | 0.11859 | -0.86494 | 2.16 |
| 3146 | *TIGAR* | 5.2501 | 0.84514 | 0.39643 | -0.78229 | 2.16 |
| 3147 | *OR8S1* | 4.6466 | 2.8367 | -0.99175 | -1.127 | 2.16 |
| 3148 | *C15orf27* | 4.6622 | 1.1004 | 0.72814 | -0.64096 | 2.16 |
| 3149 | *HSPB2* | 4.914 | 1.913 | -0.33821 | -0.95321 | 2.16 |
| 3150 | *CHD1L* | 4.3223 | 1.1225 | 1.0436 | -0.93592 | 2.16 |
| 3151 | *PRDM5* | 3.5033 | 2.7582 | 0.22661 | -0.17526 | 2.16 |
| 3152 | *RBMX2* | 3.4731 | 1.5744 | 1.4406 | -0.86482 | 2.16 |
| 3153 | *ABCC6* | 4.5836 | 0.98354 | 0.92007 | -0.32806 | 2.16 |
| 3154 | *PSPN* | 4.1725 | 1.4138 | 0.90022 | -0.3701 | 2.16 |
| 3155 | *DPP3* | 2.9704 | 2.706 | 0.80891 | -0.81532 | 2.16 |
| 3156 | *SPECC1L* | 3.9732 | 1.6559 | 0.85583 | -0.74014 | 2.16 |
| 3157 | *CIRBP* | 4.5626 | 1.2945 | 0.62766 | -1.2993 | 2.16 |
| 3158 | *JMJD6* | 6.0008 | 2.3303 | -1.8469 | -3.9921 | 2.16 |
| 3159 | *PIK3C2B* | 4.1845 | 1.6097 | 0.68947 | -0.48085 | 2.16 |
| 3160 | *STYK1* | 3.8446 | 2.0205 | 0.61791 | -0.95708 | 2.16 |
| 3161 | *ST8SIA2* | 5.1146 | 1.629 | -0.26314 | -2.3557 | 2.16 |
| 3162 | *CORO2B* | 3.7373 | 2.9207 | -0.17755 | -0.64918 | 2.16 |
| 3163 | *PRAMEF6* | 4.5644 | 0.99389 | 0.92092 | -0.99694 | 2.16 |
| 3164 | *ANKRD37* | 3.6166 | 3.5044 | -0.64362 | -1.0261 | 2.16 |
| 3165 | *ABTB2* | 2.689 | 2.3448 | 1.4429 | 0.016206 | 2.16 |
| 3166 | *SPINT4* | 3.9734 | 2.4005 | 0.10263 | -1.4717 | 2.16 |
| 3167 | *EGFL8* | 4.7647 | 1.6412 | 0.06717 | -0.00333 | 2.16 |

| | | | | | | |
|---|---|---|---|---|---|---|
| 3168 | *LIPJ* | 2.9553 | 2.5397 | 0.97788 | -1.3431 | 2.16 |
| 3169 | *AKIRIN1* | 5.5685 | 1.5414 | -0.63746 | -0.8531 | 2.16 |
| 3170 | *TRMT112* | 5.2463 | 2.1035 | -0.87788 | -1.5918 | 2.16 |
| 3171 | *TAS1R2* | 2.6968 | 2.5402 | 1.2343 | -1.4683 | 2.16 |
| 3172 | *SMPD4* | 3.2679 | 3.0549 | 0.1483 | -0.70544 | 2.16 |
| 3173 | *ZCWPW2* | 3.9442 | 3.7407 | -1.2152 | -1.3162 | 2.16 |
| 3174 | *ZBTB6* | 3.5915 | 1.6162 | 1.2616 | -1.4284 | 2.16 |
| 3175 | *C5orf46* | 4.4872 | 1.0244 | 0.95752 | 0.66656 | 2.16 |
| 3176 | *ZNF652* | 3.9601 | 3.6932 | -1.1843 | -1.5633 | 2.16 |
| 3177 | *OR5AK2* | 3.6219 | 1.4236 | 1.423 | -1.2505 | 2.16 |
| 3178 | *BHLHB9* | 3.6753 | 3.4457 | -0.6548 | -1.882 | 2.16 |
| 3179 | *WDR27* | 5.1127 | 0.89951 | 0.4533 | -2.2297 | 2.16 |
| 3180 | *NKX2-6* | 4.5175 | 1.6707 | 0.27668 | 0.16167 | 2.15 |
| 3181 | *USP35* | 3.6217 | 2.5024 | 0.34067 | -0.92764 | 2.15 |
| 3182 | *PARP2* | 4.2844 | 1.3328 | 0.84491 | -0.54775 | 2.15 |
| 3183 | *CUBN* | 4.1546 | 3.1496 | -0.84261 | -1.529 | 2.15 |
| 3184 | *CHRNA5* | 4.2312 | 1.2343 | 0.99582 | -1.2968 | 2.15 |
| 3185 | *NT5DC2* | 3.157 | 2.7169 | 0.58702 | 0.53982 | 2.15 |
| 3186 | *ACPP* | 4.4782 | 1.8055 | 0.17647 | -1.0201 | 2.15 |
| 3187 | *CPSF4* | 2.9575 | 2.108 | 1.3934 | -1.683 | 2.15 |
| 3188 | *IFNAR2* | 2.6473 | 2.444 | 1.3668 | 0.56522 | 2.15 |
| 3189 | *CLASP1* | 3.528 | 3.291 | -0.36102 | -0.45544 | 2.15 |
| 3190 | *NAT1* | 3.3659 | 2.8402 | 0.24958 | -0.13837 | 2.15 |
| 3191 | *EIF4A1* | 4.7179 | 2.4233 | -0.68606 | -0.8792 | 2.15 |
| 3192 | *KCNE1* | 3.8307 | 1.4718 | 1.1521 | 0.81091 | 2.15 |
| 3193 | *ANTXR2* | 4.4768 | 2.009 | -0.03316 | -1.4504 | 2.15 |
| 3194 | *CD74* | 4.536 | 1.1218 | 0.79469 | -0.62774 | 2.15 |
| 3195 | *KIAA0753* | 3.4778 | 2.5145 | 0.45699 | -0.5287 | 2.15 |
| 3196 | *SCMH1* | 3.3593 | 2.1101 | 0.97985 | -0.08097 | 2.15 |
| 3197 | *YAP1* | 3.1042 | 2.6755 | 0.66955 | -0.43113 | 2.15 |
| 3198 | *ADGRG2* | 3.698 | 1.9818 | 0.76887 | -1.1801 | 2.15 |
| 3199 | *APOBEC4* | 3.997 | 1.6355 | 0.81574 | -0.16781 | 2.15 |
| 3200 | *SCML4* | 3.5576 | 1.9972 | 0.89291 | 0.49265 | 2.15 |
| 3201 | *ITFG1* | 4.0746 | 1.5941 | 0.77753 | -0.91165 | 2.15 |
| 3202 | *LDHC* | 3.4474 | 2.0012 | 0.99575 | -1.5965 | 2.15 |
| 3203 | *OR6M1* | 3.7632 | 1.6569 | 1.024 | 0.98239 | 2.15 |
| 3204 | *PRICKLE2* | 3.4501 | 2.6685 | 0.32481 | -0.10374 | 2.15 |
| 3205 | *AIMP1* | 3.5879 | 2.6337 | 0.22171 | -3.475 | 2.15 |
| 3206 | *PRR15L* | 3.9317 | 3.2249 | -0.71443 | -0.75537 | 2.15 |
| 3207 | *UBR7* | 3.3274 | 2.7233 | 0.39018 | -0.55154 | 2.15 |
| 3208 | *SOAT1* | 3.7178 | 2.9956 | -0.27263 | -1.2754 | 2.15 |
| 3209 | *OR6A2* | 5.1456 | 1.3667 | -0.07199 | -0.73814 | 2.15 |
| 3210 | *DIO1* | 3.0733 | 2.1572 | 1.2098 | -0.75099 | 2.15 |
| 3211 | *BCAT2* | 2.846 | 2.0102 | 1.5839 | -0.18402 | 2.15 |
| 3212 | *KRTAP11-1* | 5.6701 | 0.81566 | -0.04984 | -1.4747 | 2.15 |
| 3213 | *SHFM1* | 3.8685 | 2.432 | 0.13535 | -0.54093 | 2.15 |

| | | | | | | |
|---|---|---|---|---|---|---|
| 3214 | *LNP1* | 3.8567 | 3.3864 | -0.80774 | -1.9619 | 2.15 |
| 3215 | *SDC4* | 3.3002 | 1.6432 | 1.4907 | -3.3966 | 2.14 |
| 3216 | *SPP1* | 4.1177 | 2.4484 | -0.13307 | -0.97093 | 2.14 |
| 3217 | *POLR2K* | 3.3069 | 2.4996 | 0.62632 | 0.47537 | 2.14 |
| 3218 | *OR2H2* | 5.1961 | 0.69953 | 0.53636 | 0.34151 | 2.14 |
| 3219 | *SEC31A* | 3.5857 | 2.6648 | 0.17903 | -0.20512 | 2.14 |
| 3220 | *TMEM144* | 3.6459 | 2.767 | 0.0154 | -0.8828 | 2.14 |
| 3221 | *SLC25A46* | 3.0098 | 2.0054 | 1.412 | -1.338 | 2.14 |
| 3222 | *DRGX* | 3.3921 | 2.7647 | 0.26896 | -0.83434 | 2.14 |
| 3223 | *SPATA31A6* | 3.7708 | 2.8723 | -0.21974 | -0.49414 | 2.14 |
| 3224 | *HSF1* | 4.062 | 2.8219 | -0.46076 | -0.77141 | 2.14 |
| 3225 | *TMEM161B* | 3.246 | 2.6839 | 0.49311 | -1.8921 | 2.14 |
| 3226 | *TYRP1* | 3.5021 | 1.5773 | 1.3417 | -0.8756 | 2.14 |
| 3227 | *TSHR* | 3.1517 | 2.0962 | 1.1711 | 0.061648 | 2.14 |
| 3228 | *LXN* | 3.4736 | 3.165 | -0.22006 | -0.72882 | 2.14 |
| 3229 | *PDLIM2* | 3.021 | 2.0458 | 1.3512 | 0.97236 | 2.14 |
| 3230 | *BIRC3* | 3.873 | 3.1418 | -0.59683 | -1.3132 | 2.14 |
| 3231 | *AMBRA1* | 2.6308 | 2.6097 | 1.1758 | 1.1177 | 2.14 |
| 3232 | *LRAT* | 3.1483 | 2.686 | 0.58147 | 0.12633 | 2.14 |
| 3233 | *SF3B6* | 3.7636 | 2.1292 | 0.51986 | -1.5894 | 2.14 |
| 3234 | *OOSP2* | 4.3543 | 2.2343 | -0.17614 | -1.3835 | 2.14 |
| 3235 | *ANKRD12* | 3.6014 | 1.4904 | 1.3199 | -0.59223 | 2.14 |
| 3236 | *MEST* | 2.7585 | 2.562 | 1.0909 | 0.14017 | 2.14 |
| 3237 | *SRFBP1* | 3.3712 | 2.4115 | 0.62851 | -0.68153 | 2.14 |
| 3238 | *SIL1* | 3.0361 | 3.0081 | 0.36594 | -0.79663 | 2.14 |
| 3239 | *RNF112* | 4.5484 | 1.2076 | 0.65341 | -0.29186 | 2.14 |
| 3240 | *MFN1* | 3.4057 | 2.763 | 0.23966 | -0.08059 | 2.14 |
| 3241 | *SLC9A3R2* | 3.8363 | 1.4703 | 1.0997 | -1.3675 | 2.14 |
| 3242 | *MVD* | 3.9995 | 1.5273 | 0.87907 | 0.62977 | 2.14 |
| 3243 | *C1QTNF1* | 4.6557 | 3.0226 | -1.2728 | -1.8557 | 2.14 |
| 3244 | *PIAS2* | 3.9028 | 2.281 | 0.22167 | -1.5857 | 2.14 |
| 3245 | *HFE* | 3.5538 | 2.4042 | 0.44746 | -0.00436 | 2.14 |
| 3246 | *OR5AC2* | 4.5313 | 2.5839 | -0.70988 | -1.368 | 2.14 |
| 3247 | *ABCA3* | 2.8977 | 2.5357 | 0.96946 | -1.292 | 2.13 |
| 3248 | *ZAN* | 4.1038 | 2.1098 | 0.18841 | -0.49857 | 2.13 |
| 3249 | *C1QTNF8* | 3.3483 | 1.7239 | 1.3294 | -1.5235 | 2.13 |
| 3250 | *PLEKHM3* | 4.9192 | 0.89296 | 0.58859 | 0.013933 | 2.13 |
| 3251 | *IVD* | 3.5274 | 2.25 | 0.62293 | -0.02963 | 2.13 |
| 3252 | *KRBA1* | 3.4994 | 2.5893 | 0.31146 | 0.086364 | 2.13 |
| 3253 | *KLRC2* | 3.7034 | 2.1622 | 0.53439 | -0.2677 | 2.13 |
| 3254 | *SLC4A11* | 2.6231 | 2.3437 | 1.4319 | 0.87952 | 2.13 |
| 3255 | *NDUFA6* | 4.1191 | 2.3251 | -0.04728 | -1.1156 | 2.13 |
| 3256 | *BORA* | 4.5931 | 1.3468 | 0.45681 | -1.4572 | 2.13 |
| 3257 | *OAS2* | 5.0237 | 1.9294 | -0.55678 | -0.87728 | 2.13 |
| 3258 | *ALKBH8* | 5.5152 | 0.52416 | 0.3563 | -2.0778 | 2.13 |
| 3259 | *ZNF692* | 3.923 | 1.8036 | 0.66806 | -1.0564 | 2.13 |

| | | | | | | |
|---|---|---|---|---|---|---|
| 3260 | *FBLN5* | 3.0948 | 2.9963 | 0.3027 | -0.49631 | 2.13 |
| 3261 | *OR4F17* | 3.3511 | 1.8359 | 1.2064 | 0.62 | 2.13 |
| 3262 | *PLXNB1* | 4.0514 | 1.5858 | 0.75517 | -1.525 | 2.13 |
| 3263 | *CYP4A11* | 3.1823 | 1.7239 | 1.4852 | 0.82277 | 2.13 |
| 3264 | *SLC25A23* | 3.1178 | 1.6909 | 1.5825 | -0.12852 | 2.13 |
| 3265 | *COG1* | 3.1943 | 1.8491 | 1.3477 | -0.2032 | 2.13 |
| 3266 | *MS4A8* | 4.8716 | 1.2674 | 0.252 | -2.4709 | 2.13 |
| 3267 | *WNT1* | 3.8535 | 2.5175 | 0.01976 | -0.19897 | 2.13 |
| 3268 | *FOXJ1* | 4.1765 | 2.3473 | -0.13314 | -1.7745 | 2.13 |
| 3269 | *ASB4* | 4.0267 | 2.1185 | 0.24498 | -0.53767 | 2.13 |
| 3270 | *RHOXF1* | 3.3298 | 3.0863 | -0.02598 | -1.1752 | 2.13 |
| 3271 | *GNLY* | 3.3718 | 2.2148 | 0.80277 | -0.20375 | 2.13 |
| 3272 | *ADGRF1* | 3.3586 | 3.1639 | -0.1358 | -0.48801 | 2.13 |
| 3273 | *MTUS2* | 3.085 | 1.867 | 1.4331 | 0.015704 | 2.13 |
| 3274 | *CD99L2* | 3.4329 | 3.4291 | -0.47704 | -1.7538 | 2.13 |
| 3275 | *DAPL1* | 3.6759 | 2.2558 | 0.45325 | 0.017898 | 2.13 |
| 3276 | *PIGL* | 2.9541 | 2.4122 | 1.018 | -2.2712 | 2.13 |
| 3277 | *ADGRB3* | 3.0081 | 2.4634 | 0.91238 | -2.1616 | 2.13 |
| 3278 | *PTPDC1* | 4.7179 | 2.4383 | -0.77308 | -2.4018 | 2.13 |
| 3279 | *SLC37A2* | 4.903 | 2.0216 | -0.54177 | -2.374 | 2.13 |
| 3280 | *BPHL* | 4.6886 | 1.2315 | 0.46243 | -0.63071 | 2.13 |
| 3281 | *RIN1* | 4.2787 | 2.3811 | -0.27775 | -1.5947 | 2.13 |
| 3282 | *PTPN9* | 3.2237 | 2.8341 | 0.32255 | -1.6597 | 2.13 |
| 3283 | *MERTK* | 3.6313 | 2.596 | 0.15206 | -1.1415 | 2.13 |
| 3284 | *MAMLD1* | 4.5103 | 1.635 | 0.23379 | -0.93764 | 2.13 |
| 3285 | *CEP126* | 2.7827 | 2.6387 | 0.95685 | -0.45663 | 2.13 |
| 3286 | *SH3GLB1* | 4.1544 | 2.2462 | -0.02312 | -0.48231 | 2.13 |
| 3287 | *PEF1* | 4.0558 | 1.2735 | 1.0477 | -2.1472 | 2.13 |
| 3288 | *OR52D1* | 5.4475 | 0.69121 | 0.23778 | -0.98504 | 2.13 |
| 3289 | *ACSS3* | 3.8123 | 1.6577 | 0.90583 | -0.21998 | 2.13 |
| 3290 | *TMEM19* | 4.7015 | 2.3298 | -0.65553 | -1.4607 | 2.13 |
| 3291 | *NDUFAF6* | 3.3737 | 3.3246 | -0.32344 | -1.6523 | 2.12 |
| 3292 | *PHC3* | 4.3087 | 3.0127 | -0.94747 | -1.6306 | 2.12 |
| 3293 | *PGA3* | 3.6936 | 2.7544 | -0.07422 | -0.72349 | 2.12 |
| 3294 | *TMEM201* | 3.8615 | 3.4359 | -0.92629 | -1.551 | 2.12 |
| 3295 | *TEX26* | 3.946 | 2.1131 | 0.31179 | 0.21507 | 2.12 |
| 3296 | *BTBD2* | 3.7222 | 1.5294 | 1.1169 | -1.5121 | 2.12 |
| 3297 | *DIO3* | 2.8114 | 2.0515 | 1.5055 | -2.0346 | 2.12 |
| 3298 | *EPM2AIP1* | 5.0363 | 2.2413 | -0.90922 | -1.0884 | 2.12 |
| 3299 | *MCCD1* | 3.3759 | 3.2643 | -0.27379 | -1.4143 | 2.12 |
| 3300 | *TMIGD3* | 3.7742 | 3.0071 | -0.41548 | -2.1724 | 2.12 |
| 3301 | *PBXIP1* | 2.5637 | 1.9419 | 1.8601 | 1.5488 | 2.12 |
| 3302 | *ZNF784* | 3.0407 | 2.9612 | 0.36179 | -0.58285 | 2.12 |
| 3303 | *FCHSD2* | 4.135 | 1.592 | 0.63628 | -0.48232 | 2.12 |
| 3304 | *FGF18* | 3.8082 | 1.3199 | 1.2328 | -1.03 | 2.12 |
| 3305 | *KLK7* | 4.2913 | 3.2441 | -1.1756 | -1.2249 | 2.12 |

| | | | | | | |
|---|---|---|---|---|---|---|
| 3306 | *RAI14* | 2.523 | 2.0638 | 1.7724 | -0.10176 | 2.12 |
| 3307 | *ANGPTL5* | 4.892 | 2.809 | -1.3419 | -1.7845 | 2.12 |
| 3308 | *QRFP* | 3.4681 | 2.6552 | 0.23533 | -4.0089 | 2.12 |
| 3309 | *RAI2* | 3.2684 | 2.3349 | 0.75512 | 0.41415 | 2.12 |
| 3310 | *KIAA0040* | 3.8384 | 2.2831 | 0.23613 | -0.50872 | 2.12 |
| 3311 | *TMEM241* | 5.3309 | 0.96727 | 0.05942 | -0.72798 | 2.12 |
| 3312 | *HOXB9* | 4.0612 | 2.7856 | -0.48982 | -0.69678 | 2.12 |
| 3313 | *SNURF* | 4.4791 | 1.2456 | 0.63122 | -0.36677 | 2.12 |
| 3314 | *RMND5A* | 3.8322 | 3.6883 | -1.1676 | -1.2277 | 2.12 |
| 3315 | *KCNF1* | 4.0873 | 2.9545 | -0.68934 | -1.3728 | 2.12 |
| 3316 | *LPIN2* | 4.1095 | 3.0197 | -0.7771 | -1.4572 | 2.12 |
| 3317 | *HACL1* | 2.8424 | 2.8146 | 0.69425 | -0.1453 | 2.12 |
| 3318 | *PMFBP1* | 3.2844 | 1.9984 | 1.0665 | 0.51635 | 2.12 |
| 3319 | *EMC10* | 4.9802 | 1.0757 | 0.29321 | -0.994 | 2.12 |
| 3320 | *SERPINE3* | 3.5234 | 2.1339 | 0.6915 | -0.4873 | 2.12 |
| 3321 | *MUC13* | 4.3723 | 1.0242 | 0.94907 | -0.90947 | 2.12 |
| 3322 | *IFIT1* | 3.4672 | 1.628 | 1.2503 | -0.05937 | 2.12 |
| 3323 | *CCDC171* | 3.1832 | 1.6616 | 1.4982 | -0.76728 | 2.11 |
| 3324 | *ALAS2* | 4.8681 | 1.1422 | 0.33252 | 0.25848 | 2.11 |
| 3325 | *GALM* | 3.8338 | 3.7013 | -1.1927 | -1.8102 | 2.11 |
| 3326 | *PSTPIP2* | 3.9571 | 1.7151 | 0.66965 | -0.55699 | 2.11 |
| 3327 | *MTHFD2* | 3.1298 | 2.9257 | 0.28617 | -1.2782 | 2.11 |
| 3328 | *UBN2* | 3.5326 | 3.3895 | -0.58054 | -1.2968 | 2.11 |
| 3329 | *LPCAT1* | 3.7407 | 1.5698 | 1.0299 | -0.00825 | 2.11 |
| 3330 | *NEO1* | 2.943 | 2.9194 | 0.47738 | -1.5225 | 2.11 |
| 3331 | *RAG2* | 3.7314 | 3.3819 | -0.77368 | -0.8198 | 2.11 |
| 3332 | *HELQ* | 4.0185 | 1.2002 | 1.1186 | -1.2975 | 2.11 |
| 3333 | *SS18* | 3.945 | 3.2595 | -0.86829 | -1.3665 | 2.11 |
| 3334 | *ANPEP* | 4.8483 | 0.88802 | 0.59669 | -1.3358 | 2.11 |
| 3335 | *AMOT* | 3.5702 | 3.288 | -0.52574 | -1.1585 | 2.11 |
| 3336 | *CFHR3* | 4.1616 | 1.9295 | 0.24009 | -1.1712 | 2.11 |
| 3337 | *USP30* | 3.432 | 2.8208 | 0.07739 | -0.0614 | 2.11 |
| 3338 | *DNAJC1* | 3.8284 | 1.6871 | 0.81405 | -1.5039 | 2.11 |
| 3339 | *RASAL1* | 4.0332 | 1.8181 | 0.47818 | 0.006715 | 2.11 |
| 3340 | *C15orf32* | 3.4483 | 3.1822 | -0.30166 | -1.1546 | 2.11 |
| 3341 | *TMEM210* | 3.8944 | 2.3082 | 0.12545 | -1.8107 | 2.11 |
| 3342 | *APITD1* | 4.7863 | 1.4966 | 0.04374 | -0.12258 | 2.11 |
| 3343 | *TAS2R19* | 3.5117 | 1.7446 | 1.0688 | -2.1956 | 2.11 |
| 3344 | *CREB1* | 3.8257 | 1.9123 | 0.58664 | -0.05385 | 2.11 |
| 3345 | *EPHX1* | 3.9542 | 1.4296 | 0.94084 | -3.051 | 2.11 |
| 3346 | *ZNF611* | 4.7852 | 0.9261 | 0.61325 | -0.18187 | 2.11 |
| 3347 | *WEE2* | 4.5502 | 1.8035 | -0.0292 | -0.83133 | 2.11 |
| 3348 | *CNST* | 4.9801 | 1.2223 | 0.11808 | -1.4081 | 2.11 |
| 3349 | *IGFBP7* | 3.7861 | 3.7069 | -1.1729 | -1.7599 | 2.11 |
| 3350 | *DPYSL2* | 3.2875 | 2.6307 | 0.39935 | -0.86007 | 2.11 |
| 3351 | *PDZD3* | 3.4551 | 1.6264 | 1.2358 | 0.87907 | 2.11 |

| | | | | | | |
|---|---|---|---|---|---|---|
| 3352 | *NKX2-2* | 3.3711 | 2.8357 | 0.1092 | -2.0902 | 2.11 |
| 3353 | *RAPGEF2* | 4.0325 | 3.3156 | -1.0333 | -2.5865 | 2.10 |
| 3354 | *KRTAP9-6* | 4.2803 | 2.6677 | -0.63405 | -0.92484 | 2.10 |
| 3355 | *MOSPD1* | 3.3697 | 3.0589 | -0.11567 | -1.106 | 2.10 |
| 3356 | *HSD3B2* | 5.162 | 0.58355 | 0.56737 | -1.761 | 2.10 |
| 3357 | *MRGPRX4* | 5.6206 | 0.92186 | -0.2298 | -0.27596 | 2.10 |
| 3358 | *PLP2* | 4.7471 | 1.5719 | -0.00832 | -0.26446 | 2.10 |
| 3359 | *FRMD4A* | 4.6708 | 1.2307 | 0.40897 | -1.864 | 2.10 |
| 3360 | *BRD2* | 4.403 | 3.2966 | -1.3892 | -1.7247 | 2.10 |
| 3361 | *SMR3A* | 3.8709 | 2.2404 | 0.19908 | -1.6805 | 2.10 |
| 3362 | *NR4A2* | 2.7188 | 2.3682 | 1.2224 | 0.27066 | 2.10 |
| 3363 | *TMCC3* | 4.9669 | 0.78415 | 0.5547 | -1.3823 | 2.10 |
| 3364 | *EPS8* | 4.7341 | 1.2174 | 0.3523 | -2.1496 | 2.10 |
| 3365 | *IER2* | 3.3383 | 2.317 | 0.64634 | -0.28421 | 2.10 |
| 3366 | *HIST1H2BN* | 4.5911 | 2.1345 | -0.42417 | -1.3409 | 2.10 |
| 3367 | *CHRNA10* | 2.2457 | 2.2143 | 1.8414 | -1.5956 | 2.10 |
| 3368 | *MEX3A* | 4.0114 | 1.4913 | 0.79764 | -1.1283 | 2.10 |
| 3369 | *AZU1* | 4.0955 | 3.2139 | -1.0095 | -1.1297 | 2.10 |
| 3370 | *ECEL1* | 3.5922 | 3.379 | -0.67153 | -1.7407 | 2.10 |
| 3371 | *PIK3C2G* | 3.2249 | 2.0444 | 1.0298 | 0.38951 | 2.10 |
| 3372 | *FAIM* | 5.0117 | 0.77753 | 0.50438 | -0.04968 | 2.10 |
| 3373 | *GPR141* | 3.9102 | 2.6318 | -0.24856 | -0.5597 | 2.10 |
| 3374 | *NRAP* | 4.5663 | 1.3611 | 0.3659 | 0.22103 | 2.10 |
| 3375 | *BCL2L2-PABPN1* | 4.1888 | 2.5038 | -0.39955 | -0.94777 | 2.10 |
| 3376 | *ZSCAN16* | 2.7255 | 2.3669 | 1.1984 | -2.1444 | 2.10 |
| 3377 | *LRRC7* | 2.3347 | 2.2404 | 1.7131 | -1.344 | 2.10 |
| 3378 | *PDCD4* | 3.5629 | 1.5932 | 1.1317 | -1.688 | 2.10 |
| 3379 | *HTR3A* | 4.5652 | 1.6622 | 0.05958 | -1.6664 | 2.10 |
| 3380 | *USP9Y* | 5.684 | 0.59098 | 0.00829 | -0.95388 | 2.09 |
| 3381 | *ERVV-1* | 2.9547 | 2.7466 | 0.57972 | -0.96083 | 2.09 |
| 3382 | *CCDC154* | 2.499 | 1.9566 | 1.8252 | 0.01825 | 2.09 |
| 3383 | *ARMC5* | 5.5451 | 1.2736 | -0.53815 | -2.0147 | 2.09 |
| 3384 | *YIPF5* | 2.9126 | 2.5125 | 0.85478 | -1.1005 | 2.09 |
| 3385 | *EIF2D* | 3.8377 | 1.6881 | 0.75389 | -1.1421 | 2.09 |
| 3386 | *S100A14* | 4.5783 | 1.3179 | 0.38184 | -1.7935 | 2.09 |
| 3387 | *TAS2R40* | 4.3871 | 2.8315 | -0.94113 | -2.8823 | 2.09 |
| 3388 | *OR8B12* | 2.944 | 1.914 | 1.4186 | 0.70012 | 2.09 |
| 3389 | *KCNU1* | 4.1639 | 1.5061 | 0.60397 | -0.22023 | 2.09 |
| 3390 | *GSC2* | 3.4318 | 3.1863 | -0.34431 | -2.3636 | 2.09 |
| 3391 | *BST1* | 4.0657 | 2.7434 | -0.53584 | -1.3697 | 2.09 |
| 3392 | *INSC* | 4.7023 | 1.0076 | 0.56191 | -1.1472 | 2.09 |
| 3393 | *SLC2A4* | 4.064 | 3.6487 | -1.4414 | -2.2329 | 2.09 |
| 3394 | *CRISPLD1* | 2.763 | 2.0565 | 1.4512 | -0.78026 | 2.09 |
| 3395 | *DHRS4L1* | 2.9968 | 1.6891 | 1.5842 | 0.075504 | 2.09 |
| 3396 | *RYR2* | 4.5953 | 1.2888 | 0.3849 | -0.35111 | 2.09 |
| 3397 | *KCNK1* | 2.9947 | 2.5489 | 0.72355 | 0.43235 | 2.09 |

| | | | | | | |
|---|---|---|---|---|---|---|
| 3398 | *ASNA1* | 4.4583 | 1.4848 | 0.32247 | -1.0128 | 2.09 |
| 3399 | *STRADA* | 3.2591 | 3.0122 | -0.00628 | -0.10716 | 2.09 |
| 3400 | *TLX3* | 3.5704 | 2.1417 | 0.5526 | -0.55274 | 2.09 |
| 3401 | *SEMA4G* | 5.0746 | 1.5284 | -0.33881 | -1.0977 | 2.09 |
| 3402 | *MC1R* | 4.125 | 2.0282 | 0.10939 | -0.74744 | 2.09 |
| 3403 | *ELF1* | 3.9061 | 2.5581 | -0.2038 | -2.2877 | 2.09 |
| 3404 | *PLAC1* | 3.032 | 1.9038 | 1.3238 | 0.17688 | 2.09 |
| 3405 | *KRTAP4-9* | 5.1693 | 2.2714 | -1.1815 | -1.2985 | 2.09 |
| 3406 | *USP21* | 5.7636 | 1.6082 | -1.1126 | -1.8557 | 2.09 |
| 3407 | *PRDM15* | 3.5594 | 2.0302 | 0.66953 | -2.3154 | 2.09 |
| 3408 | *OSMR* | 4.0406 | 1.1885 | 1.0299 | -0.00196 | 2.09 |
| 3409 | *CHML* | 3.6548 | 1.7511 | 0.85308 | -0.95964 | 2.09 |
| 3410 | *TAS2R14* | 4.4691 | 1.5276 | 0.26218 | -1.0123 | 2.09 |
| 3411 | *NXPE1* | 3.0292 | 2.4202 | 0.80915 | 0.059745 | 2.09 |
| 3412 | *GHR* | 4.9747 | 0.89306 | 0.39076 | -0.72348 | 2.09 |
| 3413 | *ZFR2* | 3.3322 | 2.5515 | 0.37426 | -0.38163 | 2.09 |
| 3414 | *DNAJC9* | 4.3726 | 1.7694 | 0.11561 | -1.011 | 2.09 |
| 3415 | *DGKH* | 4.0834 | 2.9577 | -0.78444 | -1.0197 | 2.09 |
| 3416 | *C12orf71* | 3.7079 | 1.6119 | 0.93573 | -0.60976 | 2.09 |
| 3417 | *CCDC62* | 4.0848 | 2.3265 | -0.15579 | -0.84326 | 2.09 |
| 3418 | *ZNF737* | 2.6597 | 1.8815 | 1.713 | -0.58289 | 2.08 |
| 3419 | *FPR2* | 2.911 | 2.6929 | 0.64923 | -0.64982 | 2.08 |
| 3420 | *NTRK3* | 4.2697 | 2.0504 | -0.06938 | -1.6881 | 2.08 |
| 3421 | *FGF13* | 3.2726 | 1.9478 | 1.0301 | 0.29068 | 2.08 |
| 3422 | *TTC29* | 5.4204 | 1.118 | -0.28819 | -1.1303 | 2.08 |
| 3423 | *LRRC48* | 2.834 | 2.0703 | 1.3456 | 1.2321 | 2.08 |
| 3424 | *GATA6* | 2.7412 | 2.1232 | 1.3853 | -0.36246 | 2.08 |
| 3425 | *PLEK2* | 4.0664 | 2.372 | -0.19182 | -0.49623 | 2.08 |
| 3426 | *DPEP2* | 2.5499 | 2.0112 | 1.6851 | 0.065751 | 2.08 |
| 3427 | *INSL5* | 3.7225 | 2.5313 | -0.00829 | -0.25893 | 2.08 |
| 3428 | *TLR9* | 3.6978 | 3.6758 | -1.1282 | -1.2092 | 2.08 |
| 3429 | *EPB41L4B* | 3.368 | 1.8895 | 0.98788 | 0.20921 | 2.08 |
| 3430 | *DC400927-CSNK1* | 4.1581 | 1.155 | 0.93183 | -0.38901 | 2.08 |
| 3431 | *PSMG1* | 3.7273 | 2.4777 | 0.03984 | -0.31584 | 2.08 |
| 3432 | *MYO16* | 3.3679 | 2.3788 | 0.49761 | 0.30616 | 2.08 |
| 3433 | *GIGYF1* | 4.1617 | 2.4075 | -0.32586 | -1.3078 | 2.08 |
| 3434 | *FAM111B* | 4.9965 | 1.3209 | -0.07438 | -1.7364 | 2.08 |
| 3435 | *KLHL25* | 3.7629 | 1.5439 | 0.93587 | -1.1303 | 2.08 |
| 3436 | *ALDH18A1* | 4.4223 | 1.9872 | -0.16693 | -4.0255 | 2.08 |
| 3437 | *MYB* | 2.5794 | 2.1653 | 1.4967 | 1.1784 | 2.08 |
| 3438 | *GJA8* | 4.104 | 1.6023 | 0.53464 | -1.2109 | 2.08 |
| 3439 | *PCDHB9* | 3.0421 | 2.4189 | 0.77921 | -0.83684 | 2.08 |
| 3440 | *CSNK2A2* | 3.9319 | 2.2068 | 0.10146 | -1.6128 | 2.08 |
| 3441 | *LIMA1* | 2.3616 | 2.1577 | 1.7208 | -0.07158 | 2.08 |
| 3442 | *TNFRSF1B* | 3.0555 | 2.212 | 0.97198 | -1.2863 | 2.08 |
| 3443 | *IFIH1* | 5.2288 | 0.90934 | 0.10106 | -0.82307 | 2.08 |

| | | | | | | |
|---|---|---|---|---|---|---|
| 3444 | *C2orf42* | 3.1323 | 2.6128 | 0.49398 | -1.8251 | 2.08 |
| 3445 | *NR1I2* | 3.5043 | 2.0472 | 0.68698 | -0.59793 | 2.08 |
| 3446 | *HOXB3* | 3.3112 | 2.248 | 0.67922 | -0.5254 | 2.08 |
| 3447 | *AES* | 5.2377 | 3.0551 | -2.0545 | -2.1419 | 2.08 |
| 3448 | *KBTBD3* | 3.7116 | 3.0424 | -0.51632 | -1.2928 | 2.08 |
| 3449 | *YES1* | 3.2599 | 3.0364 | -0.05867 | -0.45847 | 2.08 |
| 3450 | *ROBO2* | 3.9244 | 1.2001 | 1.1131 | -0.55134 | 2.08 |
| 3451 | *PHKG1* | 3.2216 | 1.9717 | 1.043 | -0.24904 | 2.08 |
| 3452 | *DEPDC7* | 2.7051 | 2.5342 | 0.99642 | -1.4518 | 2.08 |
| 3453 | *C11orf71* | 4.7862 | 2.1562 | -0.70712 | -1.5261 | 2.08 |
| 3454 | *TRDMT1* | 2.8621 | 1.8307 | 1.5409 | 0.21699 | 2.08 |
| 3455 | *E2F8* | 3.8388 | 1.4199 | 0.97471 | -0.88094 | 2.08 |
| 3456 | *PCDHA6* | 3.8167 | 1.678 | 0.73869 | -1.9794 | 2.08 |
| 3457 | *ZNF649* | 3.5514 | 3.4278 | -0.74604 | -1.1576 | 2.08 |
| 3458 | *GIMAP1-GIMAP5* | 3.8344 | 2.6655 | -0.26713 | -0.46372 | 2.08 |
| 3459 | *KDF1* | 4.0492 | 1.1676 | 1.0159 | -1.7675 | 2.08 |
| 3460 | *COLEC12* | 3.5277 | 1.6943 | 1.01 | -0.8606 | 2.08 |
| 3461 | *RORB* | 5.0305 | 1.6517 | -0.45176 | -0.59055 | 2.08 |
| 3462 | *CYP21A2* | 2.8748 | 1.8131 | 1.5424 | 0.055383 | 2.08 |
| 3463 | *BRDT* | 3.8961 | 2.3752 | -0.04235 | -1.2619 | 2.08 |
| 3464 | *GPR3* | 3.2708 | 1.6045 | 1.3522 | -0.7067 | 2.08 |
| 3465 | *GAL3ST2* | 4.8797 | 2.4546 | -1.1084 | -1.4881 | 2.08 |
| 3466 | *DNASE1L1* | 3.0756 | 2.4358 | 0.71415 | -2.7974 | 2.08 |
| 3467 | *IQSEC3* | 3.1579 | 2.3907 | 0.67647 | -0.10851 | 2.08 |
| 3468 | *SLC7A14* | 4.8196 | 1.1506 | 0.25443 | -0.52757 | 2.07 |
| 3469 | *MS4A14* | 2.9165 | 2.1672 | 1.1409 | -0.21752 | 2.07 |
| 3470 | *THAP10* | 3.7063 | 1.414 | 1.1033 | -1.8157 | 2.07 |
| 3471 | *GML* | 3.3098 | 2.8117 | 0.10195 | -0.74011 | 2.07 |
| 3472 | *ZNF570* | 3.398 | 1.5332 | 1.292 | -0.37942 | 2.07 |
| 3473 | *CACNA2D2* | 3.5175 | 2.769 | -0.06539 | -1.1492 | 2.07 |
| 3474 | *MSI1* | 3.7492 | 2.9688 | -0.49803 | -1.3023 | 2.07 |
| 3475 | *CC2D2B* | 3.7131 | 2.8641 | -0.35735 | -1.1241 | 2.07 |
| 3476 | *DOCK4* | 4.0365 | 3.0055 | -0.82368 | -1.4512 | 2.07 |
| 3477 | *TATDN3* | 3.8075 | 3.5475 | -1.1382 | -1.9041 | 2.07 |
| 3478 | *NUDT8* | 3.3448 | 2.2156 | 0.65491 | -0.69886 | 2.07 |
| 3479 | *CKAP2L* | 5.0896 | 1.2304 | -0.10483 | -0.53815 | 2.07 |
| 3480 | *TAL1* | 3.5614 | 2.3968 | 0.25591 | -0.66194 | 2.07 |
| 3481 | *ITSN1* | 3.9755 | 1.9877 | 0.25016 | -0.44713 | 2.07 |
| 3482 | *EPHA8* | 4.1047 | 1.185 | 0.92356 | 0.005872 | 2.07 |
| 3483 | *MYT1L* | 3.3139 | 3.0634 | -0.16631 | -1.1821 | 2.07 |
| 3484 | *TAF1L* | 3.2944 | 3.1918 | -0.27609 | -0.55068 | 2.07 |
| 3485 | *KIAA1324* | 3.0551 | 2.0828 | 1.0722 | -0.71371 | 2.07 |
| 3486 | *CDH5* | 3.7178 | 3.1291 | -0.63757 | -0.79821 | 2.07 |
| 3487 | *PDCD5* | 4.4763 | 2.0312 | -0.29854 | -0.43114 | 2.07 |
| 3488 | *PPP1R12B* | 3.4897 | 2.291 | 0.4282 | -0.91269 | 2.07 |
| 3489 | *E2F7* | 3.1139 | 2.1858 | 0.90745 | 0.092283 | 2.07 |

| | | | | | | |
|---|---|---|---|---|---|---|
| 3490 | *TMEM132D* | 5.1716 | 1.8559 | -0.82044 | -1.8341 | 2.07 |
| 3491 | *DCAF6* | 3.6252 | 1.887 | 0.6932 | -1.5252 | 2.07 |
| 3492 | *ACTR6* | 5.3715 | 1.2922 | -0.45867 | -2.0778 | 2.07 |
| 3493 | *OR2B3* | 4.3568 | 2.9619 | -1.1152 | -1.1476 | 2.07 |
| 3494 | *MSH2* | 4.4756 | 1.4224 | 0.30523 | -0.29054 | 2.07 |
| 3495 | *LAT* | 4.8984 | 1.2464 | 0.05782 | -1.3382 | 2.07 |
| 3496 | *ATP6AP1L* | 3.2929 | 2.9558 | -0.04793 | -0.93683 | 2.07 |
| 3497 | *CCL23* | 3.2481 | 2.3528 | 0.59948 | -1.9013 | 2.07 |
| 3498 | *CYP1B1* | 4.4939 | 1.1889 | 0.51575 | -0.75711 | 2.07 |
| 3499 | *GPR173* | 4.76 | 2.805 | -1.3665 | -1.7065 | 2.07 |
| 3500 | *TNP2* | 4.6594 | 2.0945 | -0.55586 | -1.0742 | 2.07 |
| 3501 | *MAGT1* | 3.7355 | 2.8517 | -0.39017 | -1.9326 | 2.07 |
| 3502 | *RPS4Y2* | 3.7951 | 1.4488 | 0.95182 | 0.01812 | 2.07 |
| 3503 | *BLK* | 4.7362 | 1.1344 | 0.32485 | -2.1187 | 2.07 |
| 3504 | *PFKFB4* | 4.3334 | 3.3307 | -1.4688 | -2.8922 | 2.07 |
| 3505 | *CYB561* | 5.201 | 0.53052 | 0.46329 | -1.2167 | 2.06 |
| 3506 | *FAM71D* | 3.5926 | 2.5143 | 0.08783 | -1.0359 | 2.06 |
| 3507 | *CDCA4* | 4.1044 | 2.396 | -0.30667 | -1.7577 | 2.06 |
| 3508 | *CEBPB* | 4.2985 | 1.6891 | 0.20469 | -1.7188 | 2.06 |
| 3509 | *CRTC2* | 3.1291 | 2.8369 | 0.22608 | -1.9284 | 2.06 |
| 3510 | *PLN* | 3.3363 | 1.9007 | 0.955 | 0.02195 | 2.06 |
| 3511 | *LCN2* | 3.1854 | 2.6082 | 0.39778 | -0.93287 | 2.06 |
| 3512 | *CMC2* | 4.5735 | 1.385 | 0.23246 | 0.1072 | 2.06 |
| 3513 | *MED22* | 3.9318 | 3.5494 | -1.2914 | -2.3529 | 2.06 |
| 3514 | *OR56A5* | 4.3462 | 2.5467 | -0.70463 | -1.4299 | 2.06 |
| 3515 | *PF4* | 2.7326 | 2.1591 | 1.2962 | 0.32075 | 2.06 |
| 3516 | *RAVER1* | 3.9332 | 3.302 | -1.0475 | -1.2985 | 2.06 |
| 3517 | *OBSL1* | 4.7253 | 1.2977 | 0.16338 | -0.32528 | 2.06 |
| 3518 | *ESRP2* | 5.6156 | 0.32699 | 0.24375 | -1.7819 | 2.06 |
| 3519 | *HS2ST1* | 3.1291 | 2.1004 | 0.95659 | -2.1572 | 2.06 |
| 3520 | *ZNF300* | 3.1872 | 1.5386 | 1.459 | -0.68917 | 2.06 |
| 3521 | *TMED4* | 5.239 | 1.0565 | -0.11263 | -0.22044 | 2.06 |
| 3522 | *TUBB4A* | 5.5214 | 0.96637 | -0.30608 | -2.8917 | 2.06 |
| 3523 | *CYTIP* | 3.8912 | 2.6107 | -0.32117 | -1.344 | 2.06 |
| 3524 | *WDR75* | 3.9428 | 1.629 | 0.60858 | -1.1852 | 2.06 |
| 3525 | *N4BP3* | 2.5486 | 2.5174 | 1.1143 | 0.50218 | 2.06 |
| 3526 | *SEL1L* | 3.6306 | 3.1252 | -0.5765 | -0.61694 | 2.06 |
| 3527 | *DYNLRB2* | 4.6581 | 0.85487 | 0.66502 | -0.96962 | 2.06 |
| 3528 | *FZR1* | 3.4925 | 2.361 | 0.32441 | -0.7912 | 2.06 |
| 3529 | *ZNF781* | 3.1751 | 2.4401 | 0.56187 | -1.2893 | 2.06 |
| 3530 | *AQP12A* | 4.2445 | 3.4285 | -1.4998 | -2.581 | 2.06 |
| 3531 | *C11orf65* | 3.3169 | 2.3529 | 0.50326 | 0.18946 | 2.06 |
| 3532 | *PIM1* | 4.0391 | 1.3917 | 0.74161 | -0.48816 | 2.06 |
| 3533 | *IL11RA* | 3.7458 | 1.764 | 0.66258 | -0.7342 | 2.06 |
| 3534 | *PTCD2* | 3.6306 | 1.749 | 0.79262 | 0.2556 | 2.06 |
| 3535 | *CYP4F12* | 3.87 | 1.3293 | 0.97258 | 0.05938 | 2.06 |

| | | | | | | |
|---|---|---|---|---|---|---|
| 3536 | *KRTAP9-9* | 3.8546 | 3.2468 | -0.93011 | -1.257 | 2.06 |
| 3537 | *MEGF9* | 4.0258 | 1.972 | 0.1734 | -0.84158 | 2.06 |
| 3538 | *APOOL* | 2.658 | 2.089 | 1.424 | 1.1806 | 2.06 |
| 3539 | *PXDN* | 3.2348 | 1.6475 | 1.288 | -0.75199 | 2.06 |
| 3540 | *TBPL2* | 4.6267 | 2.1153 | -0.57224 | -2.2385 | 2.06 |
| 3541 | *DMC1* | 4.4901 | 2.0251 | -0.34597 | -1.5352 | 2.06 |
| 3542 | *NMBR* | 5.5643 | 0.44994 | 0.15463 | -0.60845 | 2.06 |
| 3543 | *IFNA10* | 4.7521 | 0.74577 | 0.66934 | 0.28367 | 2.06 |
| 3544 | *HIST1H2AB* | 2.2522 | 2.0741 | 1.8407 | -1.2726 | 2.06 |
| 3545 | *GNG10* | 4.8094 | 2.6931 | -1.3358 | -1.3542 | 2.06 |
| 3546 | *SLFN5* | 3.9077 | 1.6543 | 0.60444 | -0.95017 | 2.06 |
| 3547 | *PGRMC1* | 2.7739 | 2.2065 | 1.1859 | -1.2676 | 2.06 |
| 3548 | *ITPKB* | 3.0566 | 2.6053 | 0.50271 | 0.38329 | 2.05 |
| 3549 | *PMS2* | 3.0283 | 1.9329 | 1.2026 | -1.2995 | 2.05 |
| 3550 | *SLC44A4* | 3.7698 | 1.4171 | 0.97671 | -0.32895 | 2.05 |
| 3551 | *PATE4* | 4.0342 | 1.297 | 0.82994 | 0.45306 | 2.05 |
| 3552 | *IRX6* | 3.816 | 2.5575 | -0.21291 | -2.914 | 2.05 |
| 3553 | *NTSR1* | 3.6926 | 2.4431 | 0.02489 | -0.68761 | 2.05 |
| 3554 | *UBXN6* | 3.1548 | 2.4501 | 0.5517 | -1.4036 | 2.05 |
| 3555 | *IFNL1* | 3.7424 | 1.8296 | 0.58187 | 0.042565 | 2.05 |
| 3556 | *UNC5A* | 3.3385 | 1.4832 | 1.3316 | 0.15802 | 2.05 |
| 3557 | *COL4A4* | 3.5381 | 1.9945 | 0.62056 | -0.95105 | 2.05 |
| 3558 | *ATP13A4* | 3.9681 | 1.361 | 0.82336 | -0.82524 | 2.05 |
| 3559 | *IRAK4* | 3.9515 | 1.5308 | 0.66955 | -0.77667 | 2.05 |
| 3560 | *THNSL1* | 4.3633 | 1.3384 | 0.44981 | -1.5212 | 2.05 |
| 3561 | *SLITRK6* | 2.8499 | 2.5361 | 0.76484 | 0.16938 | 2.05 |
| 3562 | *SCARA5* | 3.1517 | 2.3996 | 0.59946 | -0.44705 | 2.05 |
| 3563 | *MFHAS1* | 3.7463 | 3.6191 | -1.2159 | -2.9834 | 2.05 |
| 3564 | *SCML2* | 3.9179 | 1.9952 | 0.23572 | -0.78236 | 2.05 |
| 3565 | *SERPINA5* | 4.0968 | 2.8829 | -0.832 | -1.1945 | 2.05 |
| 3566 | *PGAM5* | 2.1939 | 2.0616 | 1.8921 | -0.71363 | 2.05 |
| 3567 | *PDZD8* | 3.6035 | 1.8062 | 0.73745 | 0.32514 | 2.05 |
| 3568 | *CACHD1* | 4.035 | 3.1561 | -1.046 | -1.1815 | 2.05 |
| 3569 | *CHRNA1* | 3.0969 | 1.8979 | 1.1497 | -1.4739 | 2.05 |
| 3570 | *NEK3* | 3.538 | 2.6544 | -0.04817 | -2.2264 | 2.05 |
| 3571 | *CNTN4* | 3.8976 | 2.957 | -0.71093 | -1.4415 | 2.05 |
| 3572 | *TET1* | 3.9058 | 1.6091 | 0.62764 | -0.87934 | 2.05 |
| 3573 | *VWCE* | 3.0082 | 2.2988 | 0.83488 | -0.15253 | 2.05 |
| 3574 | *SURF4* | 5.0767 | 0.62682 | 0.43717 | -1.8698 | 2.05 |
| 3575 | *RAB5A* | 3.0103 | 1.9559 | 1.1744 | 0.26309 | 2.05 |
| 3576 | *H1F0* | 3.7019 | 1.7895 | 0.64843 | -2.6072 | 2.05 |
| 3577 | *LOC100505841* | 2.8046 | 2.1124 | 1.222 | -1.489 | 2.05 |
| 3578 | *MAGI3* | 3.2244 | 2.0009 | 0.91333 | -0.87318 | 2.05 |
| 3579 | *MAGED2* | 3.2089 | 1.6102 | 1.3187 | 0.87741 | 2.05 |
| 3580 | *NAV3* | 3.4343 | 1.7179 | 0.98525 | 0.5639 | 2.05 |
| 3581 | *KHDRBS1* | 3.9344 | 3.2559 | -1.0529 | -1.5026 | 2.05 |

| | | | | | | |
|---|---|---|---|---|---|---|
| 3582 | *CIDEA* | 3.2983 | 2.4413 | 0.39597 | -0.57379 | 2.05 |
| 3583 | *ASCL3* | 4.5396 | 2.7242 | -1.1316 | -1.6288 | 2.04 |
| 3584 | *ZNF804B* | 3.2765 | 2.6779 | 0.17752 | -1.5525 | 2.04 |
| 3585 | *BMPR1B* | 2.1937 | 2.056 | 1.8821 | -1.696 | 2.04 |
| 3586 | *FAM3D* | 4.6055 | 2.1042 | -0.57974 | -1.0589 | 2.04 |
| 3587 | *OR52B4* | 3.4782 | 2.6201 | 0.03164 | -1.0751 | 2.04 |
| 3588 | *IL17RC* | 3.1107 | 2.46 | 0.55896 | -1.3279 | 2.04 |
| 3589 | *GATM* | 3.2163 | 1.9626 | 0.95025 | 0.72749 | 2.04 |
| 3590 | *TMEM50B* | 2.5803 | 2.4766 | 1.0706 | -0.00112 | 2.04 |
| 3591 | *LOC100287036* | 3.3473 | 2.0895 | 0.68886 | 0.67225 | 2.04 |
| 3592 | *PRSS21* | 2.8985 | 1.9706 | 1.2563 | -1.748 | 2.04 |
| 3593 | *APOBR* | 2.9733 | 2.8694 | 0.28258 | -0.05413 | 2.04 |
| 3594 | *TCP11X2* | 3.0265 | 2.9004 | 0.19713 | -1.3591 | 2.04 |
| 3595 | *SLC4A3* | 4.1445 | 2.3294 | -0.35118 | -0.89934 | 2.04 |
| 3596 | *GCLM* | 3.9873 | 1.6744 | 0.46095 | 0.066753 | 2.04 |
| 3597 | *HNRNPAB* | 5.005 | 0.61394 | 0.50358 | -1.4749 | 2.04 |
| 3598 | *TTF2* | 4.3948 | 1.0994 | 0.62805 | -1.5152 | 2.04 |
| 3599 | *CYB561A3* | 2.867 | 2.1033 | 1.151 | 0.17003 | 2.04 |
| 3600 | *PENK* | 3.0939 | 2.813 | 0.21168 | -1.4524 | 2.04 |
| 3601 | *OR56A1* | 3.9651 | 3.1264 | -0.97303 | -1.5821 | 2.04 |
| 3602 | *TIMM10B* | 3.6671 | 2.7643 | -0.31295 | -0.93415 | 2.04 |
| 3603 | *ZIC3* | 2.0851 | 2.0511 | 1.9808 | -1.9036 | 2.04 |
| 3604 | *CTSO* | 3.5327 | 1.5952 | 0.98905 | -1.4511 | 2.04 |
| 3605 | *MEF2D* | 3.7726 | 2.5576 | -0.2143 | -0.89178 | 2.04 |
| 3606 | *GPR1* | 3.4019 | 2.1566 | 0.5573 | -0.18502 | 2.04 |
| 3607 | *PER3* | 3.7571 | 1.4413 | 0.91721 | -0.03251 | 2.04 |
| 3608 | *MAX* | 3.8623 | 1.9135 | 0.33948 | -0.84542 | 2.04 |
| 3609 | *EPHA5* | 4.2759 | 1.2569 | 0.58191 | 0.45747 | 2.04 |
| 3610 | *SLC25A20* | 5.053 | 1.3991 | -0.33758 | -0.37185 | 2.04 |
| 3611 | *KNDC1* | 3.1742 | 1.9913 | 0.94896 | -0.76106 | 2.04 |
| 3612 | *RAB40C* | 3.8805 | 2.7372 | -0.50394 | -0.54404 | 2.04 |
| 3613 | *NFE2L1* | 4.5794 | 0.882 | 0.65218 | -0.00448 | 2.04 |
| 3614 | *UCP1* | 3.0472 | 2.4761 | 0.58901 | -1.4323 | 2.04 |
| 3615 | *SOX21* | 2.7674 | 2.6077 | 0.73696 | 0.34848 | 2.04 |
| 3616 | *TTC22* | 4.263 | 1.5047 | 0.34426 | 0.30635 | 2.04 |
| 3617 | *SLAMF7* | 3.3368 | 3.2142 | -0.43956 | -0.65312 | 2.04 |
| 3618 | *ACHE* | 2.4641 | 2.3588 | 1.287 | -2.5707 | 2.04 |
| 3619 | *CISD2* | 3.6906 | 1.6875 | 0.73029 | 0.69767 | 2.04 |
| 3620 | *C17orf51* | 3.4305 | 3.1563 | -0.47843 | -1.0708 | 2.04 |
| 3621 | *MAPK8IP1* | 5.08 | 0.88226 | 0.14567 | 0.03447 | 2.04 |
| 3622 | *DENND1A* | 4.7322 | 2.6213 | -1.248 | -1.5563 | 2.04 |
| 3623 | *PITX3* | 4.5425 | 2.1184 | -0.55568 | -1.3296 | 2.04 |
| 3624 | *SLC9A1* | 2.7205 | 2.1031 | 1.2812 | -0.05504 | 2.03 |
| 3625 | *ZBED4* | 4.5715 | 1.2239 | 0.30896 | -0.98129 | 2.03 |
| 3626 | *CLCN7* | 3.236 | 2.9559 | -0.0877 | -1.0134 | 2.03 |
| 3627 | *CCDC96* | 3.9173 | 2.2652 | -0.07919 | -1.7879 | 2.03 |

| | | | | | | |
|---|---|---|---|---|---|---|
| 3628 | *IDH2* | 3.946 | 2.1726 | -0.01696 | -2.5206 | 2.03 |
| 3629 | *RECQL5* | 5.2554 | 1.494 | -0.64918 | -2.1351 | 2.03 |
| 3630 | *SPTBN4* | 4.5293 | 1.991 | -0.42012 | -0.71161 | 2.03 |
| 3631 | *LMAN1L* | 3.5522 | 2.6704 | -0.12259 | -1.9746 | 2.03 |
| 3632 | *BZW2* | 5.3509 | 0.5892 | 0.15909 | -0.73462 | 2.03 |
| 3633 | *IGSF10* | 3.8633 | 1.6932 | 0.54196 | -0.92629 | 2.03 |
| 3634 | *ANKS4B* | 2.2342 | 2.0273 | 1.8365 | -1.0763 | 2.03 |
| 3635 | *TAAR6* | 4.1152 | 1.5092 | 0.47351 | -0.07895 | 2.03 |
| 3636 | *ANKRD23* | 2.547 | 2.3708 | 1.1799 | -0.55275 | 2.03 |
| 3637 | *PNMAL1* | 4.1403 | 1.7474 | 0.20971 | -0.59647 | 2.03 |
| 3638 | *MEF2A* | 2.2811 | 1.9281 | 1.8882 | 0.34617 | 2.03 |
| 3639 | *DCDC2B* | 4.0309 | 3.5337 | -1.4689 | -2.0542 | 2.03 |
| 3640 | *SPACA1* | 5.1966 | 0.87217 | 0.02688 | -0.64918 | 2.03 |
| 3641 | *PPOX* | 4.6478 | 0.76009 | 0.68763 | -0.40534 | 2.03 |
| 3642 | *PDGFA* | 4.2461 | 1.4538 | 0.39528 | -0.1958 | 2.03 |
| 3643 | *CD300A* | 3.7454 | 3.6929 | -1.3439 | -2.4244 | 2.03 |
| 3644 | *FAM73B* | 2.1699 | 2.0063 | 1.918 | -1.4823 | 2.03 |
| 3645 | *CRAMP1L* | 3.4145 | 1.8459 | 0.83358 | -0.38025 | 2.03 |
| 3646 | *TBC1D17* | 4.2818 | 1.6965 | 0.11544 | -0.58743 | 2.03 |
| 3647 | *PLEKHG7* | 5.1098 | 0.94815 | 0.03503 | -1.696 | 2.03 |
| 3648 | *MZB1* | 6.0936 | 1.5349 | -1.536 | -2.0242 | 2.03 |
| 3649 | *LCN1* | 4.2568 | 1.1246 | 0.71068 | -0.28888 | 2.03 |
| 3650 | *KCNK9* | 3.0406 | 2.4372 | 0.61421 | 0.17727 | 2.03 |
| 3651 | *GPR155* | 3.0971 | 2.9449 | 0.04999 | -1.0669 | 2.03 |
| 3652 | *STAU2* | 3.9987 | 3.3311 | -1.2397 | -1.9623 | 2.03 |
| 3653 | *CCS* | 4.1073 | 1.7911 | 0.19067 | -0.08529 | 2.03 |
| 3654 | *LRRC31* | 3.6761 | 1.8751 | 0.53784 | -1.1011 | 2.03 |
| 3655 | *POMGNT1* | 4.5211 | 0.92736 | 0.64045 | -0.57042 | 2.03 |
| 3656 | *GPM6A* | 2.3127 | 2.2099 | 1.5661 | -0.96161 | 2.03 |
| 3657 | *XRCC2* | 4.5491 | 0.95136 | 0.58812 | -1.456 | 2.03 |
| 3658 | *CLC* | 3.578 | 2.7544 | -0.24474 | -0.26575 | 2.03 |
| 3659 | *GRTP1* | 2.963 | 2.1879 | 0.93608 | -1.9028 | 2.03 |
| 3660 | *SCN1B* | 2.9558 | 2.8474 | 0.28345 | -0.66914 | 2.03 |
| 3661 | *CXXC1* | 3.6987 | 2.7501 | -0.36246 | -0.55007 | 2.03 |
| 3662 | *CNTFR* | 3.3105 | 2.1426 | 0.63315 | -0.37636 | 2.03 |
| 3663 | *KIAA0226* | 3.6354 | 1.5201 | 0.92988 | 0.14678 | 2.03 |
| 3664 | *GAS2* | 3.1206 | 2.8987 | 0.06583 | -0.34915 | 2.03 |
| 3665 | *SLC38A3* | 3.5651 | 1.6757 | 0.84431 | -0.01854 | 2.03 |
| 3666 | *COL5A1* | 3.0208 | 1.6145 | 1.4494 | -0.39004 | 2.03 |
| 3667 | *AVIL* | 2.9585 | 2.9406 | 0.18552 | -1.0921 | 2.03 |
| 3668 | *GNAZ* | 3.5769 | 2.7653 | -0.25767 | -0.44353 | 2.03 |
| 3669 | *OXER1* | 4.2472 | 1.7388 | 0.09689 | -0.13743 | 2.03 |
| 3670 | *HAVCR2* | 3.5795 | 1.3546 | 1.1487 | -1.4064 | 2.03 |
| 3671 | *GIT1* | 5.9519 | 0.35701 | -0.22673 | -0.53304 | 2.03 |
| 3672 | *DCAF8L1* | 4.3995 | 2.1047 | -0.42378 | -1.3851 | 2.03 |
| 3673 | *DSP* | 3.3232 | 1.6989 | 1.0581 | 0.46222 | 2.03 |

| | | | | | | |
|---|---|---|---|---|---|---|
| 3674 | *SEC11C* | 3.9748 | 1.4375 | 0.66779 | -1.5099 | 2.03 |
| 3675 | *OSBPL2* | 3.6219 | 1.7557 | 0.7014 | -0.48415 | 2.03 |
| 3676 | *DHX15* | 3.3862 | 2.2457 | 0.44703 | -2.3452 | 2.03 |
| 3677 | *APOBEC3B* | 4.4772 | 2.4558 | -0.85686 | -1.208 | 2.03 |
| 3678 | *IL36G* | 3.8481 | 3.0331 | -0.80594 | -1.7743 | 2.03 |
| 3679 | *DIP2A* | 3.2862 | 2.6059 | 0.18292 | -2.2432 | 2.03 |
| 3680 | *ETS2* | 2.4762 | 2.0755 | 1.5209 | -1.7158 | 2.02 |
| 3681 | *UBL4A* | 3.819 | 2.9061 | -0.6536 | -0.80144 | 2.02 |
| 3682 | *KIAA2018* | 2.5825 | 2.4081 | 1.0804 | 0.092069 | 2.02 |
| 3683 | *ATP1B2* | 4.0022 | 2.2204 | -0.15313 | -1.4709 | 2.02 |
| 3684 | *MAPK8* | 2.9296 | 2.2977 | 0.84165 | -1.6014 | 2.02 |
| 3685 | *EGLN3* | 3.003 | 2.325 | 0.74036 | -2.1217 | 2.02 |
| 3686 | *EPHB4* | 2.9017 | 2.2763 | 0.89035 | -2.6289 | 2.02 |
| 3687 | *MRAP* | 5.1688 | 1.6354 | -0.73708 | -2.5795 | 2.02 |
| 3688 | *PDX1* | 4.3171 | 1.2364 | 0.51174 | -0.61169 | 2.02 |
| 3689 | *DSCAML1* | 4.7778 | 1.6849 | -0.39865 | -0.62059 | 2.02 |
| 3690 | *GOLGA6B* | 4.5063 | 2.8438 | -1.2864 | -2.3222 | 2.02 |
| 3691 | *AFF1* | 4.4055 | 2.9246 | -1.2676 | -2.1674 | 2.02 |
| 3692 | *TAF1C* | 3.4932 | 3.0158 | -0.44727 | -0.83146 | 2.02 |
| 3693 | *RANBP17* | 4.8521 | 0.98029 | 0.2284 | 0.20413 | 2.02 |
| 3694 | *EMILIN3* | 3.3418 | 3.0836 | -0.36528 | -0.47473 | 2.02 |
| 3695 | *MTFP1* | 5.0152 | 1.9815 | -0.93661 | -1.1415 | 2.02 |
| 3696 | *MGST3* | 2.9277 | 2.6897 | 0.44245 | -0.22854 | 2.02 |
| 3697 | *C6orf1* | 2.8986 | 2.7252 | 0.43571 | -1.176 | 2.02 |
| 3698 | *CROCC* | 3.8728 | 1.8934 | 0.29307 | -0.51002 | 2.02 |
| 3699 | *FGF6* | 4.1423 | 0.96932 | 0.94737 | -1.9912 | 2.02 |
| 3700 | *SPIN4* | 3.4182 | 1.3489 | 1.2901 | -1.2571 | 2.02 |
| 3701 | *NATD1* | 3.3894 | 1.7147 | 0.9487 | -1.8484 | 2.02 |
| 3702 | *SPANXN2* | 2.9883 | 2.2402 | 0.82316 | -0.26105 | 2.02 |
| 3703 | *THRB* | 2.8903 | 2.2522 | 0.90916 | -1.1126 | 2.02 |
| 3704 | *CISD1* | 3.8209 | 2.7769 | -0.5497 | -2.2955 | 2.02 |
| 3705 | *ZDHHC8* | 2.7604 | 2.4789 | 0.80653 | -0.35232 | 2.02 |
| 3706 | *MCEE* | 2.786 | 2.7055 | 0.55179 | -0.2426 | 2.01 |
| 3707 | *C10orf95* | 4.8908 | 1.4404 | -0.28877 | -1.3205 | 2.01 |
| 3708 | *SPPL2A* | 2.3054 | 2.026 | 1.7107 | -0.02943 | 2.01 |
| 3709 | *C9orf72* | 3.9069 | 1.7706 | 0.36397 | -0.81152 | 2.01 |
| 3710 | *LAMP5* | 3.5916 | 1.7617 | 0.6875 | -1.4414 | 2.01 |
| 3711 | *CKMT1B* | 2.8877 | 2.1615 | 0.98914 | 0.41321 | 2.01 |
| 3712 | *GRIP2* | 3.6394 | 1.8318 | 0.56702 | -0.34006 | 2.01 |
| 3713 | *S100P* | 3.9478 | 1.411 | 0.67802 | -2.5307 | 2.01 |
| 3714 | *NUFIP1* | 3.9481 | 1.7418 | 0.34398 | 0.093012 | 2.01 |
| 3715 | *TEX30* | 3.6382 | 1.5311 | 0.86403 | -2.1873 | 2.01 |
| 3716 | *NEFH* | 4.1837 | 2.8404 | -0.99217 | -1.4402 | 2.01 |
| 3717 | *AKAP13* | 3.6036 | 1.3197 | 1.1081 | -0.0083 | 2.01 |
| 3718 | *MORN5* | 3.9796 | 2.9714 | -0.92078 | -1.2884 | 2.01 |
| 3719 | *THRAP3* | 2.7761 | 2.3419 | 0.91158 | -1.4317 | 2.01 |

| | | | | | | |
|---|---|---|---|---|---|---|
| 3720 | *TMEM68* | 4.7251 | 1.704 | -0.40121 | -1.606 | 2.01 |
| 3721 | *TMTC4* | 6.0367 | 0.13513 | -0.14427 | -1.9833 | 2.01 |
| 3722 | *CDCP1* | 4.9587 | 1.0281 | 0.04071 | -0.84291 | 2.01 |
| 3723 | *JAKMIP2* | 4.4585 | 1.4287 | 0.1385 | -0.8737 | 2.01 |
| 3724 | *CD63* | 2.956 | 2.4083 | 0.66127 | -1.4864 | 2.01 |
| 3725 | *ADHFE1* | 3.8673 | 1.8669 | 0.29119 | -2.3251 | 2.01 |
| 3726 | *BAIAP3* | 4.0703 | 2.067 | -0.11303 | -0.44283 | 2.01 |
| 3727 | *IFIT5* | 4.65 | 1.5333 | -0.15913 | -1.962 | 2.01 |
| 3728 | *GAB4* | 2.5458 | 2.2738 | 1.2035 | -0.67865 | 2.01 |
| 3729 | *KCNMB4* | 3.9074 | 1.2329 | 0.88146 | 0.26777 | 2.01 |
| 3730 | *OR14I1* | 3.416 | 2.7026 | -0.09709 | -0.52847 | 2.01 |
| 3731 | *TLR8* | 5.0406 | 1.3314 | -0.35059 | -0.79365 | 2.01 |
| 3732 | *ANKLE1* | 3.3752 | 1.9417 | 0.70399 | -0.05 | 2.01 |
| 3733 | *HSBP1L1* | 5.0243 | 2.824 | -1.8275 | -1.9806 | 2.01 |
| 3734 | *F11R* | 3.7762 | 2.0081 | 0.23634 | -0.34767 | 2.01 |
| 3735 | *FOSL2* | 5.2736 | 2.4646 | -1.7179 | -2.3017 | 2.01 |
| 3736 | *PRR12* | 3.904 | 1.3693 | 0.74587 | -0.75388 | 2.01 |
| 3737 | *ZFP69* | 3.072 | 2.6409 | 0.30592 | -0.08354 | 2.01 |
| 3738 | *P2RY1* | 3.1957 | 2.9798 | -0.15766 | -1.3141 | 2.01 |
| 3739 | *FUT8* | 4.4164 | 1.5488 | 0.05164 | -2.0242 | 2.01 |
| 3740 | *CTF1* | 3.2391 | 2.7899 | -0.01337 | -1.0846 | 2.01 |
| 3741 | *FBP2* | 4.0648 | 1.901 | 0.04944 | -0.18263 | 2.01 |
| 3742 | *FAM178A* | 4.2818 | 2.7553 | -1.0221 | -1.7065 | 2.01 |
| 3743 | *GGN* | 3.5549 | 3.3367 | -0.87826 | -0.90523 | 2.00 |
| 3744 | *POU6F2* | 4.1852 | 1.9562 | -0.1283 | -2.0851 | 2.00 |
| 3745 | *SLC25A5* | 4.5485 | 1.604 | -0.14091 | -0.96161 | 2.00 |
| 3746 | *PTPN22* | 3.7111 | 2.981 | -0.68122 | -1.1727 | 2.00 |
| 3747 | *ITGA8* | 3.1759 | 2.5369 | 0.29731 | 0.20167 | 2.00 |
| 3748 | *ALKBH4* | 3.4283 | 2.3252 | 0.25556 | -0.20485 | 2.00 |
| 3749 | *PPFIBP1* | 3.5294 | 3.1577 | -0.6799 | -0.78229 | 2.00 |
| 3750 | *FBLN7* | 3.3241 | 1.3686 | 1.3144 | -2.4735 | 2.00 |
| 3751 | *IRGM* | 5.0002 | 0.99449 | 0.01212 | -0.96572 | 2.00 |
| 3752 | *MRPS23* | 3.339 | 2.6484 | 0.0192 | -0.15216 | 2.00 |
| 3753 | *OR51T1* | 2.8497 | 2.5111 | 0.64558 | -0.36876 | 2.00 |
| 3754 | *ARMC9* | 3.6681 | 2.6009 | -0.26347 | -0.82453 | 2.00 |
| 3755 | *DNAJB13* | 5.886 | 0.43602 | -0.31674 | -2.2951 | 2.00 |
| 3756 | *SPRED1* | 4.1199 | 1.3486 | 0.53613 | 0.29777 | 2.00 |
| 3757 | *RSPH1* | 2.8345 | 2.6074 | 0.56217 | -1.7303 | 2.00 |
| 3758 | *FHL1* | 3.4486 | 1.731 | 0.82234 | 0.76074 | 2.00 |
| 3759 | *C7orf43* | 3.5433 | 2.7222 | -0.26357 | -0.30878 | 2.00 |
| 3760 | *CBX7* | 2.9083 | 2.365 | 0.72823 | -0.4433 | 2.00 |
| 3761 | *DRAXIN* | 4.122 | 2.9006 | -1.0211 | -1.2163 | 2.00 |
| 3762 | *SERPINA1* | 4.4692 | 1.2586 | 0.27291 | -0.73911 | 2.00 |
| 3763 | *SLC18A1* | 3.2547 | 1.3744 | 1.3715 | -1.2195 | 2.00 |
| 3764 | *RNF181* | 3.742 | 2.5035 | -0.24569 | -1.5125 | 2.00 |
| 3765 | *PPIF* | 4.9954 | 0.70375 | 0.30021 | -2.599 | 2.00 |

| | | | | | | |
|---|---|---|---|---|---|---|
| 3766 | *TNS4* | 3.387 | 1.59 | 1.022 | -0.69866 | 2.00 |
| 3767 | *SLITRK1* | 4.1716 | 1.5442 | 0.28261 | 0.071465 | 2.00 |
| 3768 | *ZYG11B* | 4.1726 | 1.013 | 0.81109 | -1.3205 | 2.00 |
| 3769 | *ABHD15* | 3.4443 | 2.3908 | 0.15966 | -1.3961 | 2.00 |
| 3770 | *CSDE1* | 4.1238 | 0.9548 | 0.9133 | -1.2619 | 2.00 |
| 3771 | *GIMAP4* | 2.4763 | 1.8209 | 1.694 | -0.41406 | 2.00 |
| 3772 | *SOS2* | 3.0734 | 2.2346 | 0.68273 | -0.35676 | 2.00 |
| 3773 | *HAT1* | 3.0182 | 2.7271 | 0.24369 | -1.1709 | 2.00 |
| 3774 | *KRT39* | 4.106 | 2.136 | -0.25358 | -2.425 | 2.00 |
| 3775 | *KRTAP4-4* | 3.9945 | 1.7395 | 0.25227 | -2.4661 | 2.00 |
| 3776 | *APP* | 4.3335 | 0.96898 | 0.68365 | -0.29906 | 2.00 |
| 3777 | *EFCAB1* | 2.9672 | 2.7552 | 0.26323 | -0.44353 | 2.00 |
| 3778 | *JAK1* | 2.7509 | 2.5717 | 0.66275 | -1.8816 | 2.00 |
| 3779 | *BPIFC* | 5.5848 | 0.91268 | -0.5133 | -1.2705 | 1.99 |
| 3780 | *ZBED3* | 3.8493 | 2.4391 | -0.3052 | -1.241 | 1.99 |
| 3781 | *MTRNR2L9* | 3.8563 | 2.7918 | -0.66535 | -2.2607 | 1.99 |
| 3782 | *SLC22A25* | 2.5532 | 2.4733 | 0.95617 | -0.93474 | 1.99 |
| 3783 | *PKN2* | 3.7114 | 1.4256 | 0.84514 | -1.6525 | 1.99 |
| 3784 | *SPRR1A* | 3.6624 | 2.1362 | 0.183 | -1.0197 | 1.99 |
| 3785 | *MED27* | 4.7453 | 1.6355 | -0.39967 | -0.55409 | 1.99 |
| 3786 | *OR2AK2* | 5.4246 | 0.57698 | -0.02101 | -0.31411 | 1.99 |
| 3787 | *RAB14* | 3.9189 | 1.8608 | 0.20058 | -0.63976 | 1.99 |
| 3788 | *CLEC4E* | 3.4066 | 1.5321 | 1.0412 | -1.338 | 1.99 |
| 3789 | *SGCD* | 3.5929 | 1.827 | 0.5599 | -2.7579 | 1.99 |
| 3790 | *ALS2CR11* | 4.0858 | 2.78 | -0.88784 | -1.3892 | 1.99 |
| 3791 | *SAG* | 3.3943 | 2.145 | 0.43614 | 0.020306 | 1.99 |
| 3792 | *NPIPA7* | 3.6496 | 3.0627 | -0.73814 | -1.1364 | 1.99 |
| 3793 | *CST7* | 3.5424 | 2.4338 | -0.00464 | -0.77748 | 1.99 |
| 3794 | *PIP5K1C* | 3.4609 | 1.3982 | 1.1119 | -1.3226 | 1.99 |
| 3795 | *SLC30A7* | 2.741 | 2.2006 | 1.0289 | -0.20877 | 1.99 |
| 3796 | *TM9SF4* | 3.6988 | 3.1149 | -0.84337 | -2.2473 | 1.99 |
| 3797 | *ATF6B* | 3.4875 | 2.555 | -0.07238 | -1.0821 | 1.99 |
| 3798 | *ANKRD31* | 4.7001 | 0.71584 | 0.55401 | -0.03223 | 1.99 |
| 3799 | *KIAA0586* | 3.5724 | 1.3082 | 1.0877 | -2.1947 | 1.99 |
| 3800 | *C1orf234* | 5.7698 | 0.87678 | -0.67887 | -1.258 | 1.99 |
| 3801 | *BRICD5* | 2.9489 | 2.2038 | 0.81428 | -0.07796 | 1.99 |
| 3802 | *AMH* | 4.7666 | 0.8308 | 0.36935 | 0.035999 | 1.99 |
| 3803 | *HNRNPCL2* | 5.4062 | 0.30616 | 0.25411 | -1.3637 | 1.99 |
| 3804 | *FKTN* | 3.2201 | 2.6946 | 0.0517 | 0.037191 | 1.99 |
| 3805 | *VAV1* | 3.4302 | 2.0351 | 0.50097 | -1.146 | 1.99 |
| 3806 | *TCEAL6* | 3.0276 | 2.6411 | 0.29629 | -1.3255 | 1.99 |
| 3807 | *LTF* | 2.2588 | 1.86 | 1.8458 | -1.5593 | 1.99 |
| 3808 | *APBB2* | 5.3001 | 2.3004 | -1.6362 | -1.7585 | 1.99 |
| 3809 | *C10orf105* | 4.5997 | 1.1591 | 0.20528 | -1.9766 | 1.99 |
| 3810 | *BCL2L14* | 4.6841 | 2.845 | -1.5658 | -1.7445 | 1.99 |
| 3811 | *ZDHHC17* | 4.7146 | 1.5492 | -0.30086 | -2.0218 | 1.99 |

| | | | | | | |
|---|---|---|---|---|---|---|
| 3812 | *SLC39A8* | 4.1402 | 1.0577 | 0.76478 | -0.74638 | 1.99 |
| 3813 | *OTUD6A* | 3.7474 | 1.1474 | 1.067 | -1.0707 | 1.99 |
| 3814 | *RPF1* | 3.3175 | 2.2874 | 0.35584 | -0.69542 | 1.99 |
| 3815 | *ZNF620* | 5.22 | 0.37868 | 0.36079 | -1.5984 | 1.99 |
| 3816 | *MICB* | 3.755 | 3.224 | -1.0223 | -1.2676 | 1.99 |
| 3817 | *CUZD1* | 3.0879 | 2.5767 | 0.29117 | -0.89866 | 1.99 |
| 3818 | *GJB7* | 5.4402 | 0.47464 | 0.04023 | -0.64234 | 1.99 |
| 3819 | *E4F1* | 3.3633 | 2.6616 | -0.06987 | -1.0451 | 1.99 |
| 3820 | *NUDT14* | 4.1207 | 2.5977 | -0.76361 | -1.6799 | 1.98 |
| 3821 | *CAMK2B* | 3.0377 | 2.2758 | 0.64067 | -0.44353 | 1.98 |
| 3822 | *IL11* | 5.1601 | 2.7993 | -2.0059 | -2.0337 | 1.98 |
| 3823 | *UFSP1* | 3.485 | 1.2649 | 1.2036 | -1.6098 | 1.98 |
| 3824 | *IFI44L* | 4.1464 | 1.9516 | -0.1449 | -1.3291 | 1.98 |
| 3825 | *MLC1* | 4.9587 | 0.62931 | 0.36374 | -0.36917 | 1.98 |
| 3826 | *ATPAF1* | 5.4424 | 2.1196 | -1.6129 | -2.1282 | 1.98 |
| 3827 | *DHRS9* | 3.3351 | 2.2836 | 0.32826 | 0.092952 | 1.98 |
| 3828 | *PRR15* | 3.0655 | 2.2419 | 0.63937 | 0.11317 | 1.98 |
| 3829 | *CGB2* | 3.6353 | 2.5238 | -0.21242 | -1.5752 | 1.98 |
| 3830 | *KLF3* | 3.0697 | 1.816 | 1.0609 | -2.5057 | 1.98 |
| 3831 | *LOC653602* | 4.1617 | 2.5394 | -0.75529 | -1.6411 | 1.98 |
| 3832 | *MSH3* | 4.0367 | 1.4151 | 0.49307 | -1.2845 | 1.98 |
| 3833 | *RGL3* | 3.1036 | 2.1173 | 0.72264 | -0.67874 | 1.98 |
| 3834 | *TACSTD2* | 3.6933 | 1.6699 | 0.5799 | -0.75711 | 1.98 |
| 3835 | *TATDN1* | 3.2772 | 2.6324 | 0.03342 | -1.2577 | 1.98 |
| 3836 | *APEX1* | 4.0969 | 2.0576 | -0.2122 | -0.67352 | 1.98 |
| 3837 | *SCLT1* | 4.3612 | 0.80144 | 0.77904 | 0.18277 | 1.98 |
| 3838 | *GALNT15* | 3.965 | 1.5914 | 0.38494 | -2.382 | 1.98 |
| 3839 | *CDK5R2* | 3.2071 | 2.2804 | 0.45215 | -0.90772 | 1.98 |
| 3840 | *ANAPC16* | 3.297 | 1.7472 | 0.89532 | -1.2728 | 1.98 |
| 3841 | *TM4SF20* | 5.1212 | 1.0095 | -0.19422 | -1.6218 | 1.98 |
| 3842 | *DAOA* | 2.5645 | 2.3452 | 1.0249 | -1.0583 | 1.98 |
| 3843 | *SLC22A3* | 3.817 | 2.2072 | -0.09036 | -0.5853 | 1.98 |
| 3844 | *SERINC5* | 4.963 | 1.3492 | -0.37888 | -0.47472 | 1.98 |
| 3845 | *SOX17* | 5.2049 | 0.62795 | 0.09965 | -0.90256 | 1.98 |
| 3846 | *MAGEB18* | 3.2934 | 2.0885 | 0.55029 | -0.23353 | 1.98 |
| 3847 | *BANK1* | 2.665 | 1.9282 | 1.3388 | -0.34001 | 1.98 |
| 3848 | *BMX* | 3.7181 | 1.2383 | 0.97528 | 0.46199 | 1.98 |
| 3849 | *CHST1* | 4.0659 | 1.2272 | 0.63732 | -0.85928 | 1.98 |
| 3850 | *PTGES2* | 3.8257 | 1.9109 | 0.19246 | -0.38471 | 1.98 |
| 3851 | *USP36* | 3.6812 | 1.8932 | 0.35351 | -0.91944 | 1.98 |
| 3852 | *PAK2* | 2.3301 | 1.9729 | 1.6229 | 0.77174 | 1.98 |
| 3853 | *B3GNT7* | 3.1636 | 2.3855 | 0.37607 | -0.14163 | 1.98 |
| 3854 | *RFC3* | 4.4362 | 0.91832 | 0.56958 | -0.3866 | 1.97 |
| 3855 | *KIFAP3* | 4.7482 | 1.1249 | 0.0504 | -0.01346 | 1.97 |
| 3856 | *SLFN12* | 2.9124 | 2.6004 | 0.40853 | -1.2284 | 1.97 |
| 3857 | *TRHDE* | 2.7221 | 2.4865 | 0.71096 | -0.154 | 1.97 |

| | | | | | | |
|---|---|---|---|---|---|---|
| 3858 | *NPR2* | 3.8741 | 2.5697 | -0.52426 | -1.3767 | 1.97 |
| 3859 | *SMOC2* | 3.3253 | 3.1525 | -0.55849 | -0.71363 | 1.97 |
| 3860 | *CDC26* | 5.8599 | 0.07319 | -0.01531 | -1.4998 | 1.97 |
| 3861 | *HMGN5* | 5.8142 | 0.70752 | -0.60619 | -0.62213 | 1.97 |
| 3862 | *FAM83B* | 4.7272 | 0.85887 | 0.32887 | -0.89258 | 1.97 |
| 3863 | *GPX7* | 3.6288 | 2.0472 | 0.23854 | -0.47033 | 1.97 |
| 3864 | *EDA2R* | 4.6325 | 1.2052 | 0.07616 | -0.12773 | 1.97 |
| 3865 | *HAPLN3* | 3.1149 | 2.9394 | -0.14048 | -0.86756 | 1.97 |
| 3866 | *LMAN2* | 5.1098 | 1.2438 | -0.44134 | -0.56195 | 1.97 |
| 3867 | *CYSLTR2* | 4.1788 | 1.2671 | 0.46539 | 0.36838 | 1.97 |
| 3868 | *NPAS2* | 5.189 | 0.75378 | -0.03161 | -2.2624 | 1.97 |
| 3869 | *BOK* | 3.8504 | 2.0364 | 0.02358 | -1.5511 | 1.97 |
| 3870 | *GSS* | 4.8252 | 1.8201 | -0.73511 | -1.0529 | 1.97 |
| 3871 | *TM6SF1* | 2.9789 | 2.6715 | 0.25899 | -2.0838 | 1.97 |
| 3872 | *GJB4* | 4.1126 | 3.4427 | -1.6463 | -2.3795 | 1.97 |
| 3873 | *PITX2* | 3.4752 | 1.5781 | 0.85497 | -0.57261 | 1.97 |
| 3874 | *MBD3L3* | 3.5636 | 2.9159 | -0.57134 | -0.80607 | 1.97 |
| 3875 | *SCAF4* | 2.5868 | 1.7339 | 1.5871 | 0.17647 | 1.97 |
| 3876 | *CEP164* | 4.1532 | 1.3027 | 0.451 | -1.0641 | 1.97 |
| 3877 | *CDH17* | 3.3959 | 2.1453 | 0.36466 | -0.96751 | 1.97 |
| 3878 | *KRTAP5-1* | 4.4746 | 0.92033 | 0.51055 | 0.12402 | 1.97 |
| 3879 | *ZBTB26* | 2.7389 | 1.8619 | 1.3029 | -0.10473 | 1.97 |
| 3880 | *MGAT5B* | 4.0628 | 1.2435 | 0.59723 | -0.49481 | 1.97 |
| 3881 | *CYLC1* | 4.3807 | 2.6431 | -1.1209 | -1.6047 | 1.97 |
| 3882 | *MAGEH1* | 4.3688 | 1.8729 | -0.34111 | -1.0099 | 1.97 |
| 3883 | *OR52E8* | 3.3271 | 2.5122 | 0.06045 | -1.7338 | 1.97 |
| 3884 | *FPR3* | 4.7015 | 2.7325 | -1.535 | -2.3488 | 1.97 |
| 3885 | *C14orf79* | 4.652 | 1.0107 | 0.23618 | -1.2418 | 1.97 |
| 3886 | *SAP30* | 4.538 | 2.0504 | -0.69146 | -0.77141 | 1.97 |
| 3887 | *ARHGEF6* | 4.2992 | 2.1069 | -0.51141 | -1.0297 | 1.96 |
| 3888 | *DCDC5* | 3.9809 | 1.5561 | 0.35737 | -0.62808 | 1.96 |
| 3889 | *CNDP2* | 4.1143 | 3.1146 | -1.3361 | -1.3808 | 1.96 |
| 3890 | *SKAP1* | 3.9618 | 1.5646 | 0.36569 | 0.20523 | 1.96 |
| 3891 | *TMEM212* | 4.3907 | 1.7392 | -0.23883 | -0.98208 | 1.96 |
| 3892 | *ACOX2* | 3.0712 | 2.685 | 0.13202 | -1.2728 | 1.96 |
| 3893 | *PLEKHA2* | 4.8589 | 1.246 | -0.21809 | -1.4764 | 1.96 |
| 3894 | *RHEBL1* | 3.3756 | 1.9743 | 0.53686 | -1.0712 | 1.96 |
| 3895 | *CEP95* | 3.0817 | 2.9617 | -0.15681 | -0.91757 | 1.96 |
| 3896 | *CDKN2AIP* | 4.6809 | 1.0052 | 0.2001 | -0.7687 | 1.96 |
| 3897 | *DNAJB4* | 2.3633 | 2.0431 | 1.4791 | 1.1598 | 1.96 |
| 3898 | *FIGLA* | 3.7732 | 3.2477 | -1.1364 | -1.5101 | 1.96 |
| 3899 | *OR8J1* | 3.6978 | 1.9782 | 0.20845 | -1.7547 | 1.96 |
| 3900 | *HOXB8* | 4.0782 | 1.6373 | 0.16888 | -1.448 | 1.96 |
| 3901 | *ARL9* | 3.9542 | 2.4545 | -0.52547 | -1.1637 | 1.96 |
| 3902 | *CARD11* | 3.45 | 2.9608 | -0.52858 | -2.1599 | 1.96 |
| 3903 | *MAPKAP1* | 3.0834 | 1.8272 | 0.9713 | -0.42356 | 1.96 |

| | | | | | | |
|---|---|---|---|---|---|---|
| 3904 | *KRT10* | 3.4322 | 2.2489 | 0.19862 | -1.4507 | 1.96 |
| 3905 | *RAB1A* | 2.8687 | 2.8226 | 0.18761 | -0.64632 | 1.96 |
| 3906 | *RNF170* | 2.2053 | 2.1088 | 1.5646 | 1.4953 | 1.96 |
| 3907 | *IFI6* | 2.7979 | 1.9665 | 1.1142 | 1.0709 | 1.96 |
| 3908 | *PPP1R13B* | 4.2245 | 0.88539 | 0.76859 | -1.4717 | 1.96 |
| 3909 | *PAX6* | 3.2253 | 1.9341 | 0.71905 | -0.6602 | 1.96 |
| 3910 | *TTLL2* | 3.5606 | 2.8646 | -0.54972 | -1.1307 | 1.96 |
| 3911 | *DEFB124* | 2.8145 | 2.4395 | 0.62092 | -0.89313 | 1.96 |
| 3912 | *TEX40* | 3.8926 | 2.5786 | -0.59634 | -1.5429 | 1.96 |
| 3913 | *KIAA1551* | 4.3722 | 1.4139 | 0.08858 | -2.5153 | 1.96 |
| 3914 | *RALYL* | 4.9868 | 1.4641 | -0.57704 | -0.9673 | 1.96 |
| 3915 | *TRMT10A* | 3.017 | 1.9862 | 0.87059 | -0.63009 | 1.96 |
| 3916 | *APOA4* | 4.0429 | 1.5211 | 0.30954 | -1.6274 | 1.96 |
| 3917 | *KAAG1* | 2.6777 | 1.7452 | 1.45 | -2.5519 | 1.96 |
| 3918 | *ADPRM* | 2.8313 | 2.2436 | 0.79706 | -1.3053 | 1.96 |
| 3919 | *NUDT3* | 3.1307 | 2.1904 | 0.54834 | -1.3705 | 1.96 |
| 3920 | *FAM118A* | 3.2054 | 2.346 | 0.31329 | 0.16099 | 1.95 |
| 3921 | *NEDD4L* | 2.7218 | 2.4876 | 0.65529 | -1.8095 | 1.95 |
| 3922 | *CAMK2A* | 4.4131 | 1.2433 | 0.20767 | 0.10415 | 1.95 |
| 3923 | *DEFA1B* | 2.3613 | 2.3221 | 1.1795 | -3.3058 | 1.95 |
| 3924 | *DOPEY1* | 2.8644 | 2.205 | 0.79305 | -0.47423 | 1.95 |
| 3925 | *STX2* | 3.3595 | 3.1277 | -0.62489 | -1.5015 | 1.95 |
| 3926 | *NAV2* | 3.0984 | 2.7108 | 0.05194 | -1.7619 | 1.95 |
| 3927 | *TBC1D15* | 3.8449 | 1.4462 | 0.56712 | -0.25163 | 1.95 |
| 3928 | *YTHDF1* | 2.8576 | 1.8162 | 1.184 | -1.5524 | 1.95 |
| 3929 | *RNASE12* | 4.6649 | 1.6949 | -0.50227 | -2.4491 | 1.95 |
| 3930 | *DNAH8* | 3.5779 | 2.482 | -0.20242 | -0.39817 | 1.95 |
| 3931 | *REEP4* | 4.5875 | 1.7059 | -0.43694 | -1.7188 | 1.95 |
| 3932 | *ROPN1B* | 2.916 | 2.7818 | 0.15863 | -1.3701 | 1.95 |
| 3933 | *GTF2IRD1* | 3.4215 | 1.2951 | 1.1396 | -0.64149 | 1.95 |
| 3934 | *GIMAP1* | 3.0508 | 2.7457 | 0.05866 | -2.6322 | 1.95 |
| 3935 | *TSPAN32* | 3.8608 | 1.708 | 0.2859 | -0.42588 | 1.95 |
| 3936 | *NAP1L2* | 3.1867 | 1.5967 | 1.0705 | -0.00828 | 1.95 |
| 3937 | *ABCC3* | 4.4244 | 2.7174 | -1.2884 | -2.4015 | 1.95 |
| 3938 | *PPT1* | 3.1331 | 2.5802 | 0.1395 | -1.2651 | 1.95 |
| 3939 | *SDC1* | 4.1375 | 0.91273 | 0.80203 | 0.31834 | 1.95 |
| 3940 | *FREM1* | 4.4628 | 0.75104 | 0.63841 | -1.5467 | 1.95 |
| 3941 | *WBSCR27* | 3.6248 | 3.4342 | -1.2075 | -1.4373 | 1.95 |
| 3942 | *RGS8* | 4.3822 | 1.9109 | -0.44236 | -1.4819 | 1.95 |
| 3943 | *PVRL4* | 2.6124 | 1.7641 | 1.4738 | -1.7184 | 1.95 |
| 3944 | *NUCB2* | 4.2954 | 0.88489 | 0.66961 | -1.0695 | 1.95 |
| 3945 | *CCDC39* | 4.4506 | 1.8357 | -0.43694 | -1.2358 | 1.95 |
| 3946 | *FOXD4L3* | 4.2079 | 0.89321 | 0.74794 | -2.5934 | 1.95 |
| 3947 | *PIH1D3* | 4.6622 | 0.78255 | 0.4024 | -1.1016 | 1.95 |
| 3948 | *KLC2* | 3.435 | 2.2094 | 0.20213 | -1.1823 | 1.95 |
| 3949 | *DPEP3* | 4.6758 | 1.1533 | 0.01701 | -1.2486 | 1.95 |

| | | | | | | |
|---|---|---|---|---|---|---|
| 3950 | *EPHX4* | 3.8663 | 2.7644 | -0.78506 | -1.0386 | 1.95 |
| 3951 | *ZNF350* | 3.5608 | 1.7106 | 0.57397 | -2.0838 | 1.95 |
| 3952 | *TEX33* | 2.797 | 2.1324 | 0.91549 | -0.44079 | 1.95 |
| 3953 | *KRT84* | 4.7565 | 0.80437 | 0.2836 | -1.1964 | 1.95 |
| 3954 | *TGIF2LY* | 3.3676 | 2.0337 | 0.44307 | -0.58131 | 1.95 |
| 3955 | *C6orf48* | 2.7669 | 2.4866 | 0.59052 | -0.51689 | 1.95 |
| 3956 | *SLC13A5* | 2.7047 | 2.3301 | 0.80823 | -0.16266 | 1.95 |
| 3957 | *MECOM* | 2.8331 | 2.0769 | 0.9328 | -0.15226 | 1.95 |
| 3958 | *GNAT2* | 4.339 | 1.7645 | -0.26125 | -1.4509 | 1.95 |
| 3959 | *ATAD1* | 3.7726 | 2.8104 | -0.74086 | -1.683 | 1.95 |
| 3960 | *RORA* | 4.3735 | 1.185 | 0.28355 | 0.13558 | 1.95 |
| 3961 | *PIGX* | 4.186 | 1.9087 | -0.25425 | -1.1316 | 1.95 |
| 3962 | *SOWAHA* | 3.5738 | 1.3221 | 0.94376 | 0.28124 | 1.95 |
| 3963 | *TMOD1* | 4.8452 | 0.85727 | 0.1365 | -1.707 | 1.95 |
| 3964 | *ZNF491* | 3.7138 | 1.8387 | 0.28566 | -1.0922 | 1.95 |
| 3965 | *ZNF264* | 3.5821 | 1.3122 | 0.94338 | 0.38037 | 1.95 |
| 3966 | *GSDMD* | 4.3576 | 1.6099 | -0.13025 | -0.18636 | 1.95 |
| 3967 | *KIAA0930* | 3.5356 | 2.0304 | 0.26771 | -1.2608 | 1.94 |
| 3968 | *ZNF775* | 4.938 | 1.143 | -0.24794 | -0.39078 | 1.94 |
| 3969 | *ATP8B1* | 2.5914 | 1.909 | 1.3314 | 0.76235 | 1.94 |
| 3970 | *TP73* | 2.9501 | 1.7081 | 1.1723 | -1.6781 | 1.94 |
| 3971 | *COL12A1* | 3.468 | 2.921 | -0.55929 | -1.3583 | 1.94 |
| 3972 | *NAPSA* | 3.5053 | 1.8351 | 0.48897 | -0.83753 | 1.94 |
| 3973 | *DOK3* | 2.1625 | 2.0659 | 1.6006 | -1.0548 | 1.94 |
| 3974 | *SLC41A2* | 4.623 | 0.66956 | 0.53539 | -0.29046 | 1.94 |
| 3975 | *CCDC127* | 4.5744 | 3.2564 | -2.003 | -2.4557 | 1.94 |
| 3976 | *OSBPL10* | 3.0452 | 1.9226 | 0.85974 | 0.44828 | 1.94 |
| 3977 | *RBM5* | 3.7836 | 2.3855 | -0.34536 | -0.3701 | 1.94 |
| 3978 | *TLE4* | 3.4339 | 2.5211 | -0.1314 | -0.64219 | 1.94 |
| 3979 | *COL4A5* | 3.9364 | 0.99705 | 0.88837 | -1.2985 | 1.94 |
| 3980 | *BTN1A1* | 4.0929 | 2.2686 | -0.53969 | -0.97066 | 1.94 |
| 3981 | *CRISP3* | 4.4425 | 1.4633 | -0.08437 | -0.29929 | 1.94 |
| 3982 | *TRIM74* | 3.6862 | 1.3335 | 0.79888 | 0.32027 | 1.94 |
| 3983 | *C1orf50* | 3.5745 | 1.1841 | 1.0589 | -1.9513 | 1.94 |
| 3984 | *CTNNB1* | 4.9038 | 1.3795 | -0.46733 | -1.8937 | 1.94 |
| 3985 | *ADRBK2* | 2.8826 | 2.5563 | 0.3765 | -1.2328 | 1.94 |
| 3986 | *LY86* | 3.2984 | 2.1367 | 0.38004 | -0.2613 | 1.94 |
| 3987 | *LRFN3* | 3.3458 | 1.7991 | 0.66953 | -0.59008 | 1.94 |
| 3988 | *CUEDC2* | 4.2181 | 1.5409 | 0.05454 | -1.1747 | 1.94 |
| 3989 | *NPPB* | 2.0268 | 1.982 | 1.8047 | -1.7364 | 1.94 |
| 3990 | *USP9X* | 5.4345 | 0.70578 | -0.32689 | -0.60658 | 1.94 |
| 3991 | *STRN4* | 2.9458 | 1.4594 | 1.4075 | -0.93726 | 1.94 |
| 3992 | *KRTAP13-3* | 3.332 | 1.5037 | 0.97699 | 0.1111 | 1.94 |
| 3993 | *OPRK1* | 3.5047 | 1.7833 | 0.52357 | -0.0576 | 1.94 |
| 3994 | *UGDH* | 3.9282 | 0.97082 | 0.912 | -0.00827 | 1.94 |
| 3995 | *FBXO42* | 4.2095 | 1.3661 | 0.23442 | -0.87673 | 1.94 |

| | | | | | | |
|---|---|---|---|---|---|---|
| 3996 | *ID4* | 3.427 | 2.7676 | -0.38471 | -2.3615 | 1.94 |
| 3997 | *PDE6H* | 4.3718 | 1.8409 | -0.40348 | -0.41643 | 1.94 |
| 3998 | *SIAE* | 5.5523 | 0.35443 | -0.09922 | -1.5608 | 1.94 |
| 3999 | *B4GALT1* | 2.8148 | 1.921 | 1.0709 | -1.8348 | 1.94 |
| 4000 | *TAS2R16* | 3.9291 | 1.0899 | 0.7869 | 0.61626 | 1.94 |
| 4001 | *DNASE2B* | 2.7758 | 2.5818 | 0.44637 | -0.31475 | 1.93 |
| 4002 | *PRSS12* | 3.705 | 1.1133 | 0.98511 | -1.4089 | 1.93 |
| 4003 | *HDAC1* | 3.2924 | 1.7205 | 0.79047 | 0.7274 | 1.93 |
| 4004 | *KIZ* | 5.2272 | 0.52922 | 0.04637 | -0.05538 | 1.93 |
| 4005 | *TNFRSF11B* | 4.5971 | 0.616 | 0.58934 | -0.00831 | 1.93 |
| 4006 | *SLC4A2* | 3.5555 | 2.3324 | -0.0862 | -0.23857 | 1.93 |
| 4007 | *TRAF6* | 4.6589 | 1.3526 | -0.20989 | -1.748 | 1.93 |
| 4008 | *PADI3* | 2.7217 | 2.0895 | 0.98902 | -2.716 | 1.93 |
| 4009 | *FAM21C* | 2.7456 | 1.7722 | 1.2819 | 0.11502 | 1.93 |
| 4010 | *CCR8* | 3.3042 | 1.8907 | 0.60448 | -0.41064 | 1.93 |
| 4011 | *TMBIM1* | 4.2393 | 1.2102 | 0.34895 | -0.99833 | 1.93 |
| 4012 | *CT45A5* | 3.5117 | 3.0901 | -0.80394 | -1.6799 | 1.93 |
| 4013 | *OBP2B* | 5.0286 | 1.0777 | -0.30863 | -1.951 | 1.93 |
| 4014 | *DNAH11* | 3.3408 | 2.6435 | -0.18689 | -1.8462 | 1.93 |
| 4015 | *CTAG1A* | 5.7469 | 0.1717 | -0.12213 | -0.88274 | 1.93 |
| 4016 | *LCE3E* | 2.36 | 2.2433 | 1.193 | -1.4968 | 1.93 |
| 4017 | *IVNS1ABP* | 4.9633 | 0.61907 | 0.21366 | -1.8311 | 1.93 |
| 4018 | *SDC3* | 2.6909 | 2.381 | 0.72413 | -0.17305 | 1.93 |
| 4019 | *HS1BP3* | 2.8638 | 2.43 | 0.50214 | -0.69564 | 1.93 |
| 4020 | *S100A7L2* | 3.6115 | 2.8842 | -0.70106 | -1.6663 | 1.93 |
| 4021 | *PNOC* | 4.8353 | 0.81815 | 0.13972 | -0.95136 | 1.93 |
| 4022 | *MSTN* | 2.9891 | 2.4584 | 0.34489 | -1.5959 | 1.93 |
| 4023 | *ADAMTS18* | 3.9114 | 1.137 | 0.74305 | -1.0547 | 1.93 |
| 4024 | *SGSH* | 3.7976 | 2.1693 | -0.1771 | -0.9952 | 1.93 |
| 4025 | *GRWD1* | 5.6755 | 0.37511 | -0.26125 | -1.5082 | 1.93 |
| 4026 | *SAC3D1* | 4.1934 | 0.93868 | 0.65625 | -0.45403 | 1.93 |
| 4027 | *TCP1* | 4.1917 | 1.578 | 0.01837 | -0.18955 | 1.93 |
| 4028 | *BSPRY* | 3.3823 | 1.4435 | 0.96221 | -0.70591 | 1.93 |
| 4029 | *SERPINB7* | 4.8065 | 0.78556 | 0.19546 | -1.3686 | 1.93 |
| 4030 | *METTL1* | 3.4789 | 2.8925 | -0.5842 | -1.1821 | 1.93 |
| 4031 | *MAOB* | 2.802 | 2.0907 | 0.89401 | -1.2331 | 1.93 |
| 4032 | *PADI6* | 4.0861 | 1.3147 | 0.38487 | -1.5026 | 1.93 |
| 4033 | *CSTB* | 4.4756 | 0.68368 | 0.62632 | -0.72479 | 1.93 |
| 4034 | *SPX* | 4.2347 | 2.9762 | -1.4269 | -1.9446 | 1.93 |
| 4035 | *SLIT2* | 3.314 | 1.5895 | 0.88024 | -0.03957 | 1.93 |
| 4036 | *FAM196B* | 2.9987 | 2.0407 | 0.74419 | -0.5911 | 1.93 |
| 4037 | *CHAMP1* | 2.4803 | 2.0204 | 1.2823 | 0.079048 | 1.93 |
| 4038 | *SLC28A2* | 3.7283 | 1.191 | 0.86333 | 0.32942 | 1.93 |
| 4039 | *GLRX* | 3.4158 | 1.9453 | 0.42124 | -0.06464 | 1.93 |
| 4040 | *TRPC4AP* | 4.4629 | 1.6578 | -0.33886 | -1.9013 | 1.93 |
| 4041 | *TANK* | 4.0776 | 3.1638 | -1.4596 | -1.5672 | 1.93 |

| | | | | | | |
|---|---|---|---|---|---|---|
| 4042 | *RFK* | 3.9863 | 2.8607 | -1.0653 | -2.0276 | 1.93 |
| 4043 | *CRTAM* | 4.3636 | 1.4209 | -0.00369 | -1.0809 | 1.93 |
| 4044 | *PKD1L1* | 5.0096 | 0.44563 | 0.32406 | -0.07909 | 1.93 |
| 4045 | *HERC1* | 4.7911 | 2.2322 | -1.2463 | -1.5268 | 1.93 |
| 4046 | *KLHDC9* | 4.4292 | 0.68476 | 0.66132 | -0.51304 | 1.93 |
| 4047 | *SLC15A1* | 4.1272 | 1.1308 | 0.51635 | -2.5292 | 1.92 |
| 4048 | *IGSF9* | 2.7451 | 2.132 | 0.89644 | 0.080286 | 1.92 |
| 4049 | *ZNF394* | 4.0627 | 1.526 | 0.18069 | 0.1421 | 1.92 |
| 4050 | *MALT1* | 3.8778 | 1.1791 | 0.71191 | -1.1747 | 1.92 |
| 4051 | *MTMR8* | 5.5039 | 0.16693 | 0.0973 | -0.91317 | 1.92 |
| 4052 | *TMEM214* | 4.53 | 0.92279 | 0.31528 | -1.28 | 1.92 |
| 4053 | *SYNPO2L* | 6.1852 | 0.79432 | -1.212 | -1.53 | 1.92 |
| 4054 | *SLC28A3* | 3.0598 | 2.1278 | 0.57918 | -0.34241 | 1.92 |
| 4055 | *CHODL* | 3.2125 | 1.6482 | 0.90574 | -1.2427 | 1.92 |
| 4056 | *DCUN1D1* | 4.6874 | 1.1022 | -0.02341 | -0.64083 | 1.92 |
| 4057 | *TEX12* | 5.4459 | 0.42422 | -0.10528 | -2.5198 | 1.92 |
| 4058 | *WASF1* | 3.3917 | 1.7637 | 0.60849 | 0.36853 | 1.92 |
| 4059 | *CFAP70* | 2.5327 | 1.8387 | 1.3923 | -0.05504 | 1.92 |
| 4060 | *MYBPH* | 3.7317 | 1.1782 | 0.85163 | -0.14151 | 1.92 |
| 4061 | *STX19* | 4.4392 | 2.2274 | -0.90549 | -1.6826 | 1.92 |
| 4062 | *MDK* | 3.7819 | 2.338 | -0.35893 | -1.0807 | 1.92 |
| 4063 | *MTRNR2L8* | 5.329 | 0.53752 | -0.1057 | -1.2186 | 1.92 |
| 4064 | *DENND4C* | 3.1076 | 2.399 | 0.25407 | -1.3023 | 1.92 |
| 4065 | *RNASE4* | 5.6882 | 0.32159 | -0.24942 | -0.55904 | 1.92 |
| 4066 | *OR7G2* | 3.3165 | 1.4146 | 1.028 | -0.65378 | 1.92 |
| 4067 | *MAP4K4* | 4.3466 | 1.1669 | 0.24437 | -0.04287 | 1.92 |
| 4068 | *DCTD* | 4.5849 | 2.211 | -1.0385 | -1.6157 | 1.92 |
| 4069 | *GPR89B* | 3.1217 | 2.0125 | 0.62248 | -0.94558 | 1.92 |
| 4070 | *IL12A* | 2.3287 | 1.7943 | 1.6331 | -1.7466 | 1.92 |
| 4071 | *ATF2* | 5.1374 | 0.5432 | 0.07542 | -1.2987 | 1.92 |
| 4072 | *RAX* | 2.93 | 1.6281 | 1.1965 | 0.50245 | 1.92 |
| 4073 | *TCHH* | 4.9738 | 1.0298 | -0.25004 | -0.31787 | 1.92 |
| 4074 | *PLIN5* | 3.7681 | 1.7211 | 0.26292 | -1.3377 | 1.92 |
| 4075 | *TMEM170B* | 3.9231 | 1.9244 | -0.09656 | -0.38119 | 1.92 |
| 4076 | *IP6K1* | 3.4933 | 1.8725 | 0.38493 | 0.22057 | 1.92 |
| 4077 | *FAM46B* | 3.7175 | 1.439 | 0.59408 | -1.3783 | 1.92 |
| 4078 | *PNRC2* | 4.6922 | 0.91493 | 0.14305 | -0.96144 | 1.92 |
| 4079 | *ELOVL6* | 3.6065 | 1.2108 | 0.93279 | 0.43428 | 1.92 |
| 4080 | *DDX43* | 4.2805 | 1.0422 | 0.4261 | -0.90161 | 1.92 |
| 4081 | *FOXM1* | 2.5227 | 1.7955 | 1.4302 | -0.39722 | 1.92 |
| 4082 | *MYL10* | 2.8738 | 2.0444 | 0.82951 | -0.69794 | 1.92 |
| 4083 | *CAT* | 3.5806 | 3.1888 | -1.0224 | -1.0639 | 1.92 |
| 4084 | *TMEM33* | 3.6758 | 2.3191 | -0.24849 | -0.8183 | 1.92 |
| 4085 | *SETD5* | 5.0573 | 0.37331 | 0.31479 | -1.8842 | 1.92 |
| 4086 | *ACOT7* | 5.1593 | 0.67443 | -0.08851 | -1.0576 | 1.92 |
| 4087 | *GBP3* | 3.4338 | 2.129 | 0.18168 | -0.46351 | 1.91 |

| | | | | | | |
|---|---|---|---|---|---|---|
| 4088 | *TRIM16L* | 3.2101 | 2.0138 | 0.52046 | -1.8314 | 1.91 |
| 4089 | *ZNF707* | 4.4108 | 1.125 | 0.20832 | -1.2448 | 1.91 |
| 4090 | *MGST2* | 3.0594 | 2.6553 | 0.02784 | -2.1984 | 1.91 |
| 4091 | *CFAP52* | 4.2626 | 2.5744 | -1.0949 | -1.3892 | 1.91 |
| 4092 | *KLHL21* | 2.5389 | 2.0435 | 1.1585 | -1.9678 | 1.91 |
| 4093 | *TAF4* | 3.0029 | 2.3672 | 0.36965 | 0.078586 | 1.91 |
| 4094 | *QSER1* | 3.7738 | 2.6428 | -0.67751 | -1.4798 | 1.91 |
| 4095 | *IL17RB* | 2.7409 | 1.5615 | 1.4363 | -1.8572 | 1.91 |
| 4096 | *GIGYF2* | 3.0548 | 1.4452 | 1.2383 | -0.07172 | 1.91 |
| 4097 | *THUMPD3* | 2.5499 | 2.3669 | 0.82132 | -1.9766 | 1.91 |
| 4098 | *CCDC68* | 3.4768 | 1.5826 | 0.67802 | -1.5526 | 1.91 |
| 4099 | *GPR137* | 4.5285 | 0.66453 | 0.54271 | 0.24439 | 1.91 |
| 4100 | *HLA-DOB* | 2.9016 | 2.5767 | 0.25569 | -1.9375 | 1.91 |
| 4101 | *SLC47A1* | 4.1575 | 2.7332 | -1.1571 | -1.8932 | 1.91 |
| 4102 | *TSGA13* | 4.7515 | 0.66965 | 0.31154 | -1.7065 | 1.91 |
| 4103 | *GOSR1* | 5.242 | 1.6181 | -1.1283 | -1.8335 | 1.91 |
| 4104 | *LAIR1* | 3.2275 | 1.3341 | 1.1701 | -1.7077 | 1.91 |
| 4105 | *SUCO* | 3.5913 | 3.4405 | -1.3023 | -1.6431 | 1.91 |
| 4106 | *LOC149373* | 2.3556 | 2.2168 | 1.1571 | -0.81505 | 1.91 |
| 4107 | *XPNPEP1* | 5.319 | 0.4644 | -0.0566 | -0.77333 | 1.91 |
| 4108 | *ADCK4* | 2.6759 | 2.2441 | 0.80676 | 0.21455 | 1.91 |
| 4109 | *SNX5* | 3.2012 | 1.5384 | 0.98692 | -1.6276 | 1.91 |
| 4110 | *TM9SF2* | 3.8841 | 2.8319 | -0.99079 | -2.4067 | 1.91 |
| 4111 | *CD79A* | 3.7621 | 1.2269 | 0.73578 | 0.47341 | 1.91 |
| 4112 | *SH2B3* | 5.2752 | 0.81574 | -0.36654 | -2.0283 | 1.91 |
| 4113 | *RAPGEFL1* | 3.4256 | 1.8541 | 0.44229 | -1.4762 | 1.91 |
| 4114 | *POSTN* | 4.7899 | 0.70965 | 0.22191 | -2.9354 | 1.91 |
| 4115 | *ZNF829* | 2.76 | 2.4456 | 0.51527 | -0.44353 | 1.91 |
| 4116 | *MS4A5* | 3.7747 | 2.0018 | -0.05637 | -0.97595 | 1.91 |
| 4117 | *CLINT1* | 4.5197 | 0.60457 | 0.5935 | -1.0091 | 1.91 |
| 4118 | *SYT6* | 5.0401 | 0.90557 | -0.22803 | -0.57018 | 1.91 |
| 4119 | *HLA-C* | 4.6061 | 1.1801 | -0.06874 | -0.87817 | 1.91 |
| 4120 | *ANXA10* | 3.6687 | 1.1091 | 0.93894 | -0.97093 | 1.91 |
| 4121 | *WDPCP* | 3.7364 | 2.8877 | -0.908 | -1.1027 | 1.91 |
| 4122 | *GPR148* | 4.3779 | 1.5895 | -0.25135 | -1.6274 | 1.91 |
| 4123 | *TCTEX1D1* | 3.4506 | 1.9289 | 0.33545 | 0.016531 | 1.90 |
| 4124 | *STKLD1* | 3.7206 | 1.9481 | 0.045 | -0.04792 | 1.90 |
| 4125 | *ZNF439* | 3.6867 | 2.9043 | -0.87817 | -1.773 | 1.90 |
| 4126 | *ISCA1* | 3.3808 | 1.9488 | 0.38204 | -1.0391 | 1.90 |
| 4127 | *FLVCR1* | 3.0334 | 1.5389 | 1.1373 | -2.0197 | 1.90 |
| 4128 | *GGA2* | 2.9235 | 2.8858 | -0.09982 | -1.3797 | 1.90 |
| 4129 | *PADI4* | 2.5366 | 2.0119 | 1.1605 | -1.5736 | 1.90 |
| 4130 | *DYNC1LI2* | 3.9177 | 0.903 | 0.88827 | 0.28694 | 1.90 |
| 4131 | *PFKFB1* | 4.0519 | 1.105 | 0.55205 | 0.15749 | 1.90 |
| 4132 | *ZDHHC13* | 3.5187 | 2.8634 | -0.67521 | -0.84872 | 1.90 |
| 4133 | *CSRNP1* | 4.5637 | 1.2289 | -0.08601 | -0.51859 | 1.90 |

| | | | | | | |
|---|---|---|---|---|---|---|
| 4134 | *ZNF347* | 3.575 | 3.1651 | -1.0336 | -1.0954 | 1.90 |
| 4135 | *GIPC3* | 4.1909 | 2.5778 | -1.0623 | -1.4682 | 1.90 |
| 4136 | *CASZ1* | 2.7214 | 2.3772 | 0.60749 | -1.3498 | 1.90 |
| 4137 | *CXorf67* | 4.6512 | 0.79371 | 0.26076 | -0.42658 | 1.90 |
| 4138 | *NPNT* | 4.6406 | 1.4536 | -0.38937 | -0.65924 | 1.90 |
| 4139 | *PDPR* | 2.3993 | 2.3629 | 0.94176 | 0.60858 | 1.90 |
| 4140 | *SMIM5* | 3.1993 | 2.7841 | -0.27964 | -0.76879 | 1.90 |
| 4141 | *PLCE1* | 4.1853 | 1.4078 | 0.10968 | -0.29865 | 1.90 |
| 4142 | *JOSD1* | 5.8731 | 0.27314 | -0.44575 | -2.4018 | 1.90 |
| 4143 | *C9orf84* | 4.7781 | 0.46507 | 0.45695 | -2.1496 | 1.90 |
| 4144 | *AQP6* | 3.2497 | 2.6127 | -0.16295 | -2.6456 | 1.90 |
| 4145 | *CLN8* | 2.6195 | 2.0063 | 1.0729 | -1.1651 | 1.90 |
| 4146 | *SLK* | 2.1765 | 1.9434 | 1.5773 | -0.79798 | 1.90 |
| 4147 | *DUSP18* | 4.2894 | 1.0439 | 0.36293 | -0.43841 | 1.90 |
| 4148 | *E2F2* | 4.0753 | 3.0362 | -1.4183 | -2.7628 | 1.90 |
| 4149 | *ALG12* | 3.3515 | 1.6646 | 0.67707 | -0.40125 | 1.90 |
| 4150 | *VRTN* | 4.2332 | 1.1382 | 0.32059 | -1.7865 | 1.90 |
| 4151 | *GNA12* | 2.4621 | 1.9076 | 1.322 | 0.87408 | 1.90 |
| 4152 | *OSTM1* | 3.8327 | 2.1025 | -0.24431 | -1.0509 | 1.90 |
| 4153 | *SLC26A7* | 3.1123 | 2.1046 | 0.4739 | -0.1601 | 1.90 |
| 4154 | *ARHGAP36* | 4.9904 | 0.99667 | -0.29681 | -1.1685 | 1.90 |
| 4155 | *ATP4A* | 2.9393 | 2.2029 | 0.54804 | 0.095053 | 1.90 |
| 4156 | *MRFAP1L1* | 4.2238 | 1.8963 | -0.42989 | -1.1477 | 1.90 |
| 4157 | *INAFM2* | 2.8386 | 2.1493 | 0.70225 | -0.09 | 1.90 |
| 4158 | *CDIP1* | 2.2581 | 2.0161 | 1.4154 | -0.31164 | 1.90 |
| 4159 | *OR4A47* | 5.0173 | 0.7708 | -0.09892 | -1.1795 | 1.90 |
| 4160 | *HBG2* | 4.5839 | 0.97762 | 0.1276 | -1.3513 | 1.90 |
| 4161 | *ZNF587* | 3.5435 | 1.8457 | 0.29991 | -1.6604 | 1.90 |
| 4162 | *GRM3* | 4.1778 | 1.8883 | -0.37727 | -0.38815 | 1.90 |
| 4163 | *RCHY1* | 4.5738 | 1.0416 | 0.07273 | -0.45246 | 1.90 |
| 4164 | *NDRG3* | 4.3344 | 0.88385 | 0.46795 | -0.11381 | 1.90 |
| 4165 | *DENND2A* | 3.5744 | 1.8898 | 0.22141 | -0.97526 | 1.90 |
| 4166 | *EMD* | 3.6692 | 1.048 | 0.96819 | -1.2907 | 1.90 |
| 4167 | *TPPP2* | 3.7739 | 2.4341 | -0.52332 | -1.1089 | 1.89 |
| 4168 | *OR2M7* | 4.1275 | 2.5209 | -0.96407 | -1.4881 | 1.89 |
| 4169 | *TMED1* | 4.1271 | 1.8469 | -0.29194 | -0.5898 | 1.89 |
| 4170 | *FSCB* | 5.0305 | 0.67393 | -0.0227 | -1.9573 | 1.89 |
| 4171 | *WDR89* | 4.0437 | 1.382 | 0.25504 | -0.52679 | 1.89 |
| 4172 | *HDAC10* | 2.8729 | 2.7562 | 0.05148 | -1.3296 | 1.89 |
| 4173 | *RAD51AP2* | 2.5335 | 2.3799 | 0.76684 | -1.3074 | 1.89 |
| 4174 | *SLC18A2* | 4.0131 | 2.1045 | -0.43776 | -1.126 | 1.89 |
| 4175 | *DAZ3* | 2.7859 | 2.4832 | 0.41015 | 0.39006 | 1.89 |
| 4176 | *LOC400863* | 3.7634 | 1.0431 | 0.87266 | -0.44134 | 1.89 |
| 4177 | *PRY2* | 4.1844 | 1.3132 | 0.18083 | -1.725 | 1.89 |
| 4178 | *RRNAD1* | 3.2287 | 2.3451 | 0.10414 | -1.3237 | 1.89 |
| 4179 | *CDH20* | 3.297 | 2.8926 | -0.51278 | -0.56737 | 1.89 |

| | | | | | | |
|---|---|---|---|---|---|---|
| 4180 | *MMEL1* | 5.9459 | 0.74882 | -1.0191 | -2.0791 | 1.89 |
| 4181 | *BAG5* | 4.3961 | 1.5906 | -0.31164 | -2.2313 | 1.89 |
| 4182 | *DEFB113* | 4.795 | 0.61572 | 0.26415 | -1.5415 | 1.89 |
| 4183 | *SH2D3A* | 3.4365 | 1.9341 | 0.30403 | -1.0764 | 1.89 |
| 4184 | *VAMP1* | 5.1363 | 0.39188 | 0.14638 | -1.0359 | 1.89 |
| 4185 | *PAN3* | 3.9242 | 3.277 | -1.5268 | -1.748 | 1.89 |
| 4186 | *ZNF14* | 5.0988 | 2.0477 | -1.4722 | -2.3885 | 1.89 |
| 4187 | *DYNC1LI1* | 3.5187 | 2.0436 | 0.11192 | -0.43276 | 1.89 |
| 4188 | *AKNAD1* | 3.2139 | 1.5769 | 0.88332 | -1.3668 | 1.89 |
| 4189 | *GOLGA6A* | 3.7834 | 3.0453 | -1.1553 | -1.1845 | 1.89 |
| 4190 | *C6orf15* | 3.9457 | 1.845 | -0.11745 | -0.59061 | 1.89 |
| 4191 | *CALML4* | 3.1345 | 1.8361 | 0.70264 | -1.7988 | 1.89 |
| 4192 | *ZFY* | 5.2511 | 0.54868 | -0.12681 | -0.7757 | 1.89 |
| 4193 | *TMEM70* | 4.8986 | 0.65386 | 0.11986 | -0.75762 | 1.89 |
| 4194 | *DHDH* | 3.0651 | 1.7718 | 0.83495 | -0.42615 | 1.89 |
| 4195 | *RBMY1E* | 3.5632 | 1.8525 | 0.25544 | -0.27421 | 1.89 |
| 4196 | *STAP1* | 3.3084 | 1.8686 | 0.49269 | -0.01218 | 1.89 |
| 4197 | *S100A6* | 4.377 | 0.72847 | 0.56396 | 0.459 | 1.89 |
| 4198 | *OR52M1* | 3.1306 | 1.6946 | 0.84414 | -0.49684 | 1.89 |
| 4199 | *CRLF3* | 3.1083 | 2.501 | 0.05963 | -1.0185 | 1.89 |
| 4200 | *CRYBB3* | 3.7183 | 2.3812 | -0.43057 | -1.3255 | 1.89 |
| 4201 | *MLPH* | 3.7635 | 1.907 | -0.00256 | -0.52494 | 1.89 |
| 4202 | *MAP2K7* | 3.3421 | 2.5642 | -0.23846 | -0.93489 | 1.89 |
| 4203 | *PLEK* | 3.6496 | 1.8265 | 0.1915 | -0.37282 | 1.89 |
| 4204 | *PTPRN2* | 4.0464 | 1.5081 | 0.11287 | -1.1272 | 1.89 |
| 4205 | *CDC27* | 4.073 | 1.7618 | -0.16749 | -2.3552 | 1.89 |
| 4206 | *TTPA* | 4.2297 | 1.1964 | 0.24016 | -0.9879 | 1.89 |
| 4207 | *MYBPC1* | 4.1132 | 2.2468 | -0.69382 | -1.7547 | 1.89 |
| 4208 | *TPCN1* | 4.5913 | 1.2178 | -0.14312 | -0.79962 | 1.89 |
| 4209 | *CD34* | 3.3106 | 2.2064 | 0.14819 | -1.2195 | 1.89 |
| 4210 | *FAM199X* | 4.0785 | 1.321 | 0.26546 | 0.18914 | 1.89 |
| 4211 | *LILRA5* | 3.084 | 2.8614 | -0.28169 | -1.6089 | 1.89 |
| 4212 | *PRR7* | 5.0103 | 0.34578 | 0.30757 | -0.04796 | 1.89 |
| 4213 | *ATM* | 4.1702 | 2.6029 | -1.1117 | -1.3568 | 1.89 |
| 4214 | *ZNF592* | 3.0198 | 1.7366 | 0.90495 | -1.5502 | 1.89 |
| 4215 | *ZNF689* | 3.4615 | 1.4602 | 0.73945 | -0.7329 | 1.89 |
| 4216 | *PHF21A* | 3.5742 | 1.995 | 0.09128 | -0.72488 | 1.89 |
| 4217 | *HSPA8* | 3.2636 | 1.9625 | 0.43381 | -1.0441 | 1.89 |
| 4218 | *GSDMB* | 2.6556 | 2.2214 | 0.78047 | -0.54008 | 1.89 |
| 4219 | *HBP1* | 2.5046 | 1.9258 | 1.2265 | -0.02105 | 1.89 |
| 4220 | *SLC5A7* | 2.3048 | 1.7848 | 1.5659 | 0.2664 | 1.89 |
| 4221 | *KCNJ9* | 2.7721 | 2.48 | 0.40277 | 0.30546 | 1.88 |
| 4222 | *DBN1* | 2.7155 | 2.4516 | 0.4876 | -0.48905 | 1.88 |
| 4223 | *C1QL2* | 2.6316 | 2.3652 | 0.65788 | -2.2599 | 1.88 |
| 4224 | *RNF19B* | 3.1238 | 1.4721 | 1.0578 | -0.76541 | 1.88 |
| 4225 | *SEC11A* | 4.1473 | 2.9181 | -1.4123 | -1.7787 | 1.88 |

| | | | | | | |
|---|---|---|---|---|---|---|
| 4226 | *CAPZA2* | 4.4102 | 1.153 | 0.0896 | -0.78588 | 1.88 |
| 4227 | *DOT1L* | 4.4481 | 2.5383 | -1.3338 | -1.5817 | 1.88 |
| 4228 | *OR4C11* | 3.4905 | 2.1193 | 0.04269 | -0.30559 | 1.88 |
| 4229 | *GMIP* | 4.5171 | 1.4374 | -0.30279 | -0.85829 | 1.88 |
| 4230 | *VPS54* | 3.2696 | 1.4788 | 0.90318 | 0.85108 | 1.88 |
| 4231 | *GPR32* | 3.3551 | 2.6289 | -0.33246 | -0.49867 | 1.88 |
| 4232 | *TBC1D8B* | 3.3979 | 3.2186 | -0.96632 | -1.3637 | 1.88 |
| 4233 | *SCN4B* | 4.2742 | 2.3666 | -0.99088 | -1.4407 | 1.88 |
| 4234 | *IL10RA* | 3.5709 | 2.1703 | -0.09161 | -0.50041 | 1.88 |
| 4235 | *LRRC63* | 3.1519 | 2.3396 | 0.15768 | -0.38671 | 1.88 |
| 4236 | *DNAJC30* | 3.5619 | 1.2469 | 0.83845 | -1.1641 | 1.88 |
| 4237 | *NEU1* | 4.9589 | 0.48719 | 0.20086 | -0.91842 | 1.88 |
| 4238 | *DDX26B* | 3.961 | 0.90769 | 0.77753 | -0.77038 | 1.88 |
| 4239 | *ABHD3* | 3.6776 | 1.9159 | 0.05267 | -0.54579 | 1.88 |
| 4240 | *COL6A6* | 4.9756 | 0.66103 | 0.00926 | -0.36192 | 1.88 |
| 4241 | *ZMIZ2* | 3.3861 | 2.8386 | -0.57899 | -2.4904 | 1.88 |
| 4242 | *CPOX* | 4.4489 | 1.6178 | -0.42195 | -2.1429 | 1.88 |
| 4243 | *ICE2* | 5.2528 | 0.57972 | -0.18819 | -0.95708 | 1.88 |
| 4244 | *ANKS6* | 2.9951 | 2.5596 | 0.08594 | -2.0766 | 1.88 |
| 4245 | *ZNF705G* | 4.3157 | 1.7483 | -0.4235 | -0.71093 | 1.88 |
| 4246 | *ZNF24* | 2.7746 | 1.6385 | 1.2269 | 0.2536 | 1.88 |
| 4247 | *TSPAN5* | 4.3596 | 1.1685 | 0.11177 | -1.7232 | 1.88 |
| 4248 | *PBX4* | 4.0496 | 1.7345 | -0.1449 | -0.25711 | 1.88 |
| 4249 | *C15orf59* | 2.4409 | 1.8838 | 1.3141 | -1.2287 | 1.88 |
| 4250 | *C2CD4B* | 5.5582 | 0.096888 | -0.01766 | -1.136 | 1.88 |
| 4251 | *LRRC23* | 5.3654 | 1.3731 | -1.1021 | -2.2455 | 1.88 |
| 4252 | *NACC1* | 3.9439 | 1.017 | 0.67333 | -1.2084 | 1.88 |
| 4253 | *APOBEC1* | 2.0372 | 1.8053 | 1.7899 | -0.75512 | 1.88 |
| 4254 | *LDOC1L* | 2.5989 | 2.474 | 0.55945 | -0.20687 | 1.88 |
| 4255 | *GATB* | 4.2666 | 2.7576 | -1.392 | -2.6491 | 1.88 |
| 4256 | *DLAT* | 3.5703 | 2.3631 | -0.30135 | -0.99876 | 1.88 |
| 4257 | *MYO15A* | 4.7224 | 0.74251 | 0.16601 | -1.2071 | 1.88 |
| 4258 | *TGIF1* | 4.4723 | 0.71569 | 0.44282 | -0.43875 | 1.88 |
| 4259 | *DUSP16* | 3.1825 | 2.268 | 0.178 | -0.24356 | 1.88 |
| 4260 | *CD58* | 3.5775 | 2.2197 | -0.16927 | -1.0553 | 1.88 |
| 4261 | *DUSP15* | 4.4473 | 1.1121 | 0.06635 | -1.1308 | 1.88 |
| 4262 | *GUCA1A* | 4.3745 | 0.62921 | 0.62181 | -1.4633 | 1.88 |
| 4263 | *NEB* | 3.478 | 2.8579 | -0.71051 | -0.76371 | 1.88 |
| 4264 | *KPTN* | 3.9973 | 1.3638 | 0.26393 | -2.4858 | 1.88 |
| 4265 | *TGM4* | 3.3856 | 1.3023 | 0.93707 | 0.9038 | 1.87 |
| 4266 | *TMEM56-RWDD3* | 3.3702 | 2.6519 | -0.39838 | -1.4557 | 1.87 |
| 4267 | *TMEM62* | 3.0738 | 2.8071 | -0.25734 | -1.6673 | 1.87 |
| 4268 | *RAPGEF3* | 2.8998 | 2.1702 | 0.55312 | -0.03164 | 1.87 |
| 4269 | *PRH1* | 4.4356 | 0.82326 | 0.36378 | -0.6799 | 1.87 |
| 4270 | *FAF1* | 3.6176 | 2.1398 | -0.13488 | -0.17712 | 1.87 |
| 4271 | *METTL23* | 4.3783 | 0.62309 | 0.62065 | -0.6951 | 1.87 |

| | | | | | | |
|---|---|---|---|---|---|---|
| 4272 | *ZNF473* | 3.2565 | 1.8656 | 0.4991 | -0.9554 | 1.87 |
| 4273 | *RASA4* | 4.0626 | 2.7839 | -1.2257 | -1.5875 | 1.87 |
| 4274 | *LRRN4CL* | 4.8428 | 1.3914 | -0.61374 | -0.72776 | 1.87 |
| 4275 | *GTSCR1* | 4.2394 | 0.76817 | 0.6116 | 0.047153 | 1.87 |
| 4276 | *DSG4* | 2.508 | 1.7509 | 1.3597 | -0.06641 | 1.87 |
| 4277 | *KCNJ12* | 3.3182 | 2.1713 | 0.12799 | -0.11602 | 1.87 |
| 4278 | *ERICH6* | 3.4517 | 2.2992 | -0.1339 | -0.67846 | 1.87 |
| 4279 | *ACBD3* | 4.8646 | 0.70976 | 0.04192 | -2.4825 | 1.87 |
| 4280 | *TTC32* | 4.3788 | 0.68035 | 0.55585 | -0.2765 | 1.87 |
| 4281 | *SLC12A4* | 3.9863 | 2.4671 | -0.83851 | -1.8335 | 1.87 |
| 4282 | *IFI30* | 4.8709 | 0.95321 | -0.20958 | -1.8866 | 1.87 |
| 4283 | *MMP17* | 5.2001 | 1.0893 | -0.67688 | -1.917 | 1.87 |
| 4284 | *CCNL1* | 3.1361 | 1.7785 | 0.69763 | 0.057818 | 1.87 |
| 4285 | *MORC1* | 4.0307 | 2.3775 | -0.79632 | -1.1357 | 1.87 |
| 4286 | *HN1L* | 3.8257 | 2.2772 | -0.49127 | -1.7496 | 1.87 |
| 4287 | *FUZ* | 4.5166 | 1.5216 | -0.42938 | -0.63483 | 1.87 |
| 4288 | *FAM49A* | 4.1505 | 1.6799 | -0.22191 | -1.7789 | 1.87 |
| 4289 | *MEX3D* | 3.3028 | 3.1236 | -0.81838 | -0.88661 | 1.87 |
| 4290 | *VCX* | 4.3145 | 1.1257 | 0.1676 | -0.09468 | 1.87 |
| 4291 | *C6orf118* | 4.4882 | 0.92815 | 0.19099 | -2.1745 | 1.87 |
| 4292 | *MT1G* | 3.2849 | 1.5002 | 0.82109 | -1.8081 | 1.87 |
| 4293 | *MAP2K4* | 5.0295 | 1.0752 | -0.49902 | -2.7277 | 1.87 |
| 4294 | *PTGDR* | 4.4163 | 1.5852 | -0.39603 | -1.4688 | 1.87 |
| 4295 | *ST8SIA6* | 3.7151 | 2.5287 | -0.63912 | -0.7819 | 1.87 |
| 4296 | *OR1K1* | 4.2665 | 1.6153 | -0.27804 | -1.8516 | 1.87 |
| 4297 | *HSPBAP1* | 4.0009 | 1.3557 | 0.24672 | -0.74604 | 1.87 |
| 4298 | *NCOA6* | 4.6244 | 1.0984 | -0.11955 | -0.88279 | 1.87 |
| 4299 | *SAMM50* | 5.0567 | 0.6839 | -0.13746 | -1.0949 | 1.87 |
| 4300 | *VPS37C* | 3.4994 | 2.2024 | -0.09906 | -1.451 | 1.87 |
| 4301 | *ZNF616* | 2.7435 | 1.4413 | 1.4152 | -1.0123 | 1.87 |
| 4302 | *GNAI3* | 3.7544 | 1.8661 | -0.02056 | -0.38011 | 1.87 |
| 4303 | *SPAG4* | 3.1206 | 1.3921 | 1.087 | -1.5502 | 1.87 |
| 4304 | *KCTD16* | 3.1379 | 1.4652 | 0.99569 | -0.88418 | 1.87 |
| 4305 | *TJAP1* | 4.9012 | 1.1125 | -0.41568 | -1.8049 | 1.87 |
| 4306 | *GREM2* | 3.0659 | 1.9024 | 0.62913 | 0.33767 | 1.87 |
| 4307 | *S100A4* | 3.8436 | 1.2152 | 0.5375 | 0.52635 | 1.87 |
| 4308 | *CYP19A1* | 3.4636 | 1.8577 | 0.27462 | -0.05181 | 1.87 |
| 4309 | *AFAP1* | 3.9184 | 3.3608 | -1.6847 | -2.0351 | 1.86 |
| 4310 | *NTNG1* | 2.8206 | 1.9296 | 0.84357 | 0.1368 | 1.86 |
| 4311 | *IP6K2* | 3.1938 | 2.3611 | 0.03873 | -0.70255 | 1.86 |
| 4312 | *CCRN4L* | 3.4865 | 2.8494 | -0.74308 | -1.296 | 1.86 |
| 4313 | *LGALS7* | 3.4357 | 3.1607 | -1.0039 | -2.6549 | 1.86 |
| 4314 | *GAB3* | 3.0808 | 1.9468 | 0.56456 | -0.35894 | 1.86 |
| 4315 | *ARHGAP29* | 6.0891 | 0.15856 | -0.65707 | -2.0429 | 1.86 |
| 4316 | *OR13C2* | 2.8105 | 2.6551 | 0.12492 | -0.74179 | 1.86 |
| 4317 | *PROB1* | 3.0344 | 2.1262 | 0.42954 | 0.10343 | 1.86 |

| | | | | | | |
|---|---|---|---|---|---|---|
| 4318 | *ANO9* | 3.5898 | 2.0617 | -0.062 | -1.0216 | 1.86 |
| 4319 | *C16orf70* | 2.154 | 1.8965 | 1.5389 | -0.61682 | 1.86 |
| 4320 | *KDM4E* | 4.335 | 0.96148 | 0.29117 | -0.37213 | 1.86 |
| 4321 | *VCAM1* | 3.6946 | 2.6228 | -0.73051 | -2.2445 | 1.86 |
| 4322 | *TBK1* | 3.4584 | 1.9358 | 0.19215 | -0.33709 | 1.86 |
| 4323 | *CT45A6* | 3.6814 | 1.2383 | 0.66434 | -1.9251 | 1.86 |
| 4324 | *PLA2G3* | 5.304 | 0.25544 | 0.02442 | -1.3505 | 1.86 |
| 4325 | *TTC21A* | 3.1746 | 2.9645 | -0.55544 | -0.73014 | 1.86 |
| 4326 | *HAMP* | 2.6786 | 2.2411 | 0.6633 | -2.0423 | 1.86 |
| 4327 | *SSX2IP* | 3.3762 | 2.1836 | 0.0224 | -0.66493 | 1.86 |
| 4328 | *SWAP70* | 4.0416 | 0.79797 | 0.74226 | -1.2154 | 1.86 |
| 4329 | *HOXA5* | 2.9029 | 2.2047 | 0.47286 | -2.1802 | 1.86 |
| 4330 | *KLHL7* | 4.478 | 1.3946 | -0.29273 | -2.1239 | 1.86 |
| 4331 | *BSX* | 3.3475 | 2.442 | -0.20988 | -0.94119 | 1.86 |
| 4332 | *OR2A7* | 3.7834 | 1.2588 | 0.53705 | -0.39011 | 1.86 |
| 4333 | *PDLIM1* | 2.6438 | 2.1463 | 0.78909 | -0.19618 | 1.86 |
| 4334 | *GADL1* | 3.2699 | 2.0721 | 0.23712 | -1.5883 | 1.86 |
| 4335 | *EDNRB* | 2.6518 | 1.8854 | 1.0415 | 0.78462 | 1.86 |
| 4336 | *EAF2* | 3.1306 | 2.7342 | -0.28632 | -1.1898 | 1.86 |
| 4337 | *IQCE* | 3.5022 | 2.9177 | -0.84151 | -1.0933 | 1.86 |
| 4338 | *HSPA13* | 2.7602 | 2.0294 | 0.78805 | -1.5352 | 1.86 |
| 4339 | *CHEK2* | 3.7614 | 1.1933 | 0.62265 | 0.031866 | 1.86 |
| 4340 | *H6PD* | 3.9589 | 2.4853 | -0.86744 | -1.3258 | 1.86 |
| 4341 | *FZD4* | 3.4271 | 1.3235 | 0.82607 | 0.30419 | 1.86 |
| 4342 | *CPT1B* | 3.3587 | 1.2574 | 0.96019 | 5.25E-05 | 1.86 |
| 4343 | *SPECC1* | 5.4397 | 0.13285 | 0.00293 | -1.4859 | 1.86 |
| 4344 | *OR8J3* | 5.3879 | 0.9219 | -0.73483 | -1.4842 | 1.86 |
| 4345 | *AS3MT* | 2.3257 | 2.1778 | 1.0704 | -2.7863 | 1.86 |
| 4346 | *FAM53B* | 3.5228 | 1.8486 | 0.20174 | -1.2317 | 1.86 |
| 4347 | *OR8I2* | 3.7653 | 1.2498 | 0.55758 | -1.4312 | 1.86 |
| 4348 | *LRCH3* | 3.37 | 2.8185 | -0.61632 | -1.7649 | 1.86 |
| 4349 | *FOXB1* | 4.5132 | 0.9031 | 0.15586 | -0.31299 | 1.86 |
| 4350 | *SIM2* | 3.3703 | 2.3536 | -0.15201 | -2.4481 | 1.86 |
| 4351 | *PF4V1* | 4.253 | 0.97053 | 0.34829 | -0.29953 | 1.86 |
| 4352 | *TUBA1C* | 4.6811 | 1.4376 | -0.54826 | -1.7931 | 1.86 |
| 4353 | *PKNOX1* | 3.1094 | 2.6397 | -0.17952 | -0.49867 | 1.86 |
| 4354 | *FSHB* | 2.6397 | 1.9186 | 1.0103 | -0.6602 | 1.86 |
| 4355 | *C10orf53* | 3.4649 | 2.5223 | -0.4188 | -0.62585 | 1.86 |
| 4356 | *NBPF8* | 4.3409 | 1.1609 | 0.06603 | -1.4048 | 1.86 |
| 4357 | *HOOK3* | 3.6806 | 2.0729 | -0.18633 | -0.531 | 1.86 |
| 4358 | *TMEM134* | 3.6892 | 1.5735 | 0.30413 | -0.57567 | 1.86 |
| 4359 | *MSL1* | 3.7714 | 1.6391 | 0.15417 | -1.5965 | 1.85 |
| 4360 | *WDR45* | 4.5004 | 1.2076 | -0.14391 | -0.66689 | 1.85 |
| 4361 | *AP5S1* | 2.509 | 1.7346 | 1.3181 | 0.14155 | 1.85 |
| 4362 | *GPR119* | 4.9855 | 1.9874 | -1.4112 | -2.7254 | 1.85 |
| 4363 | *POU2F2* | 3.5016 | 2.6998 | -0.64278 | -0.79392 | 1.85 |

| | | | | | | |
|---|---|---|---|---|---|---|
| 4364 | *IQCA1L* | 4.4329 | 0.56529 | 0.56025 | -0.48911 | 1.85 |
| 4365 | *LUZP4* | 5.7216 | 0.68717 | -0.85059 | -1.6368 | 1.85 |
| 4366 | *TMSB15B* | 3.7607 | 1.9701 | -0.17331 | -0.86358 | 1.85 |
| 4367 | *ZNF317* | 5.3138 | 0.33366 | -0.09022 | -0.35946 | 1.85 |
| 4368 | *DPY19L2* | 4.8737 | 0.34486 | 0.33826 | -0.0171 | 1.85 |
| 4369 | *TBC1D8* | 3.4209 | 1.9101 | 0.22562 | -0.42369 | 1.85 |
| 4370 | *APBB1IP* | 5.0784 | 0.61545 | -0.13748 | -0.97237 | 1.85 |
| 4371 | *CORIN* | 3.6999 | 2.6834 | -0.82705 | -1.2447 | 1.85 |
| 4372 | *TIMD4* | 3.7685 | 1.2917 | 0.49592 | 0.18216 | 1.85 |
| 4373 | *C10orf35* | 3.7863 | 1.6906 | 0.07914 | -0.6622 | 1.85 |
| 4374 | *B3GALTL* | 4.6066 | 1.0113 | -0.06271 | -1.0416 | 1.85 |
| 4375 | *SLC22A1* | 4.2826 | 0.71509 | 0.55662 | -0.22803 | 1.85 |
| 4376 | *TMEM213* | 3.0169 | 2.3172 | 0.21879 | -1.2249 | 1.85 |
| 4377 | *HAND1* | 3.3235 | 2.2658 | -0.03648 | -0.33967 | 1.85 |
| 4378 | *RBFOX1* | 3.9153 | 1.0457 | 0.59119 | -1.0344 | 1.85 |
| 4379 | *LYRM1* | 2.8386 | 2.3833 | 0.32941 | 0.067859 | 1.85 |
| 4380 | *UBE2L3* | 4.4613 | 0.72375 | 0.36615 | -1.9358 | 1.85 |
| 4381 | *OSBPL11* | 3.502 | 1.771 | 0.27723 | -1.183 | 1.85 |
| 4382 | *B3GALT2* | 4.4073 | 1.3073 | -0.16468 | -1.9099 | 1.85 |
| 4383 | *TAP1* | 4.6362 | 0.48648 | 0.4271 | -1.3297 | 1.85 |
| 4384 | *LGR5* | 2.7071 | 2.5842 | 0.25825 | -2.3678 | 1.85 |
| 4385 | *ERCC6-PGBD3* | 3.4253 | 1.1965 | 0.92716 | -0.99137 | 1.85 |
| 4386 | *PLS3* | 3.6316 | 1.7694 | 0.14607 | -0.73546 | 1.85 |
| 4387 | *GINM1* | 4.4 | 1.0784 | 0.06861 | -0.28819 | 1.85 |
| 4388 | *PON2* | 3.9377 | 2.9188 | -1.3114 | -1.7911 | 1.85 |
| 4389 | *PGM2* | 5.4728 | 0.33464 | -0.26237 | -1.6826 | 1.85 |
| 4390 | *RAB24* | 3.1879 | 2.1521 | 0.2036 | -0.32625 | 1.85 |
| 4391 | *MBL2* | 3.5542 | 2.9176 | -0.92861 | -1.761 | 1.85 |
| 4392 | *FAM64A* | 2.5832 | 2.5509 | 0.40862 | -0.24887 | 1.85 |
| 4393 | *UNC93B1* | 3.5253 | 2.0484 | -0.03102 | -1.356 | 1.85 |
| 4394 | *SOX8* | 3.575 | 1.4246 | 0.54305 | 0.49463 | 1.85 |
| 4395 | *LSR* | 2.5452 | 2.1828 | 0.81297 | -0.40482 | 1.85 |
| 4396 | *AQP9* | 2.8568 | 2.5729 | 0.1087 | -1.9052 | 1.85 |
| 4397 | *MPV17L2* | 3.2499 | 2.4535 | -0.16651 | -0.4246 | 1.85 |
| 4398 | *MIXL1* | 4.042 | 2.4257 | -0.93114 | -1.5408 | 1.85 |
| 4399 | *SCN1A* | 3.2097 | 1.5709 | 0.75466 | 0.51723 | 1.85 |
| 4400 | *UHRF1* | 4.5062 | 0.68863 | 0.34032 | -0.26357 | 1.85 |
| 4401 | *TACR2* | 2.6655 | 1.6409 | 1.2284 | 0.29118 | 1.84 |
| 4402 | *IMPG2* | 4.9837 | 0.50854 | 0.04244 | -0.94747 | 1.84 |
| 4403 | *TRPC1* | 4.5766 | 0.48199 | 0.47594 | -0.95964 | 1.84 |
| 4404 | *DYM* | 4.2438 | 0.85492 | 0.43515 | -1.1761 | 1.84 |
| 4405 | *MUT* | 2.7304 | 1.5965 | 1.2051 | 1.1252 | 1.84 |
| 4406 | *CEP152* | 4.4055 | 0.72088 | 0.4055 | 0.17543 | 1.84 |
| 4407 | *ZNF132* | 4.0423 | 1.4611 | 0.02759 | -1.8306 | 1.84 |
| 4408 | *RAMP3* | 3.1194 | 1.7718 | 0.63936 | 0.43058 | 1.84 |
| 4409 | *WNT3A* | 2.9588 | 1.386 | 1.1852 | 0.11018 | 1.84 |

| | | | | | | |
|---|---|---|---|---|---|---|
| 4410 | *COX6A2* | 3.2949 | 2.2309 | 0.00408 | -0.79573 | 1.84 |
| 4411 | *ETV6* | 3.8562 | 1.8746 | -0.20105 | -0.20626 | 1.84 |
| 4412 | *LALBA* | 4.4252 | 2.1743 | -1.0706 | -1.5512 | 1.84 |
| 4413 | *FAM124A* | 6.0557 | 0.31225 | -0.84001 | -3.1023 | 1.84 |
| 4414 | *ACBD6* | 3.3764 | 1.3339 | 0.81698 | -0.54126 | 1.84 |
| 4415 | *C22orf29* | 4.023 | 2.6925 | -1.1891 | -1.4811 | 1.84 |
| 4416 | *CDPF1* | 3.3285 | 1.564 | 0.63342 | -0.84841 | 1.84 |
| 4417 | *MATK* | 3.2672 | 2.3412 | -0.08269 | -0.42701 | 1.84 |
| 4418 | *IGF1R* | 3.2198 | 2.7397 | -0.43398 | -1.7072 | 1.84 |
| 4419 | *GFI1B* | 3.8094 | 2.0634 | -0.34809 | -0.40863 | 1.84 |
| 4420 | *MAGEB10* | 2.4618 | 1.8218 | 1.241 | 0.64075 | 1.84 |
| 4421 | *NPW* | 3.1439 | 2.0197 | 0.36064 | -1.1108 | 1.84 |
| 4422 | *RBM4B* | 4.1066 | 1.9235 | -0.50685 | -1.3912 | 1.84 |
| 4423 | *DHRS7B* | 4.1625 | 0.78842 | 0.57188 | -1.1715 | 1.84 |
| 4424 | *GNMT* | 2.6818 | 1.5467 | 1.2932 | -1.7753 | 1.84 |
| 4425 | *H2AFY* | 5.3269 | 0.64059 | -0.44661 | -4.0089 | 1.84 |
| 4426 | *ALLC* | 2.5595 | 2.4621 | 0.49881 | -1.8293 | 1.84 |
| 4427 | *TRIP11* | 2.9001 | 2.3521 | 0.26807 | -0.04917 | 1.84 |
| 4428 | *AADACL3* | 5.0678 | 0.52531 | -0.0729 | -1.4104 | 1.84 |
| 4429 | *IL1RL2* | 3.993 | 2.0761 | -0.54914 | -2.012 | 1.84 |
| 4430 | *ENPP7* | 2.6019 | 1.892 | 1.0255 | -1.1681 | 1.84 |
| 4431 | *NDUFB1* | 4.1812 | 0.93545 | 0.40171 | -1.1144 | 1.84 |
| 4432 | *ADH6* | 3.6521 | 1.9891 | -0.124 | -0.77794 | 1.84 |
| 4433 | *GOLGA6D* | 3.0114 | 2.8211 | -0.31624 | -1.525 | 1.84 |
| 4434 | *LAS1L* | 3.6437 | 2.3951 | -0.52269 | -1.2643 | 1.84 |
| 4435 | *ABCG1* | 4.0963 | 1.4561 | -0.03683 | -1.5073 | 1.84 |
| 4436 | *DSC3* | 2.336 | 1.7517 | 1.4265 | 0.39603 | 1.84 |
| 4437 | *RBM14-RBM4* | 2.9325 | 2.5563 | 0.0251 | -0.42877 | 1.84 |
| 4438 | *MAP10* | 2.8889 | 2.1453 | 0.47908 | 0.33208 | 1.84 |
| 4439 | *REN* | 3.2948 | 1.8138 | 0.40461 | -0.67556 | 1.84 |
| 4440 | *KRTAP3-1* | 2.6498 | 1.9187 | 0.94468 | -1.4043 | 1.84 |
| 4441 | *PCDHGA8* | 3.5692 | 1.8951 | 0.04877 | -1.8503 | 1.84 |
| 4442 | *ENO4* | 3.9871 | 0.91266 | 0.61192 | -0.36285 | 1.84 |
| 4443 | *CALML3* | 4.7304 | 1.8476 | -1.0669 | -1.526 | 1.84 |
| 4444 | *CDKN2C* | 3.9664 | 1.0952 | 0.4488 | -1.1692 | 1.84 |
| 4445 | *PQLC3* | 4.4715 | 0.86991 | 0.16852 | -1.1507 | 1.84 |
| 4446 | *NUDT12* | 3.248 | 1.5534 | 0.70812 | -0.33788 | 1.84 |
| 4447 | *TNFRSF12A* | 4.8622 | 1.168 | -0.52105 | -2.2844 | 1.84 |
| 4448 | *KCNQ4* | 3.013 | 2.8205 | -0.32443 | -2.4338 | 1.84 |
| 4449 | *TEX101* | 3.5334 | 1.2337 | 0.74016 | -1.1724 | 1.84 |
| 4450 | *MIPOL1* | 4.9079 | 0.4811 | 0.11761 | -1.9654 | 1.84 |
| 4451 | *TRAK2* | 4.1829 | 1.83 | -0.50637 | -0.65895 | 1.84 |
| 4452 | *RAB6A* | 2.9037 | 1.4569 | 1.1456 | 0.7603 | 1.84 |
| 4453 | *CALU* | 4.0156 | 1.9848 | -0.49425 | -0.68627 | 1.84 |
| 4454 | *NCR3* | 3.2875 | 1.8748 | 0.34341 | -1.0623 | 1.84 |
| 4455 | *CLN6* | 3.3591 | 3.0183 | -0.87418 | -2.2473 | 1.83 |

| | | | | | | |
|---|---|---|---|---|---|---|
| 4456 | *SNTG1* | 3.9513 | 1.5968 | -0.04542 | -0.62307 | 1.83 |
| 4457 | *FAM160A1* | 4.1638 | 1.9992 | -0.66164 | -0.9481 | 1.83 |
| 4458 | *TAF1D* | 4.312 | 0.98486 | 0.20227 | -0.56481 | 1.83 |
| 4459 | *KCNQ3* | 3.5551 | 2.4001 | -0.45669 | -1.3085 | 1.83 |
| 4460 | *HTRA3* | 2.5207 | 1.8477 | 1.1294 | 1.0782 | 1.83 |
| 4461 | *MTHFR* | 3.1408 | 2.1508 | 0.20432 | -1.7027 | 1.83 |
| 4462 | *RASA3* | 2.3404 | 2.3086 | 0.8469 | -1.626 | 1.83 |
| 4463 | *ASIC4* | 3.7631 | 1.9683 | -0.23802 | -1.8899 | 1.83 |
| 4464 | *CLIP1* | 3.9569 | 0.92224 | 0.61408 | 0.14404 | 1.83 |
| 4465 | *LASP1* | 4.1819 | 2.8086 | -1.4976 | -1.5401 | 1.83 |
| 4466 | *MCUR1* | 4.1825 | 0.79518 | 0.51497 | -1.6288 | 1.83 |
| 4467 | *ITPR1* | 5.0103 | 1.0964 | -0.6141 | -1.7675 | 1.83 |
| 4468 | *FAM9C* | 2.7779 | 1.452 | 1.2598 | -0.19851 | 1.83 |
| 4469 | *RCCD1* | 4.8367 | 0.40272 | 0.25015 | -0.18718 | 1.83 |
| 4470 | *CEACAM6* | 2.65 | 1.8553 | 0.98304 | -1.7109 | 1.83 |
| 4471 | *C17orf80* | 4.5766 | 0.46733 | 0.44389 | -0.94777 | 1.83 |
| 4472 | *FCGR2B* | 5.488 | 0.33139 | -0.33179 | -1.4798 | 1.83 |
| 4473 | *SASH3* | 3.374 | 1.1919 | 0.92166 | -1.6952 | 1.83 |
| 4474 | *CCNL2* | 4.0577 | 0.80993 | 0.61894 | -1.5415 | 1.83 |
| 4475 | *ACSL4* | 3.132 | 1.499 | 0.85523 | -1.25 | 1.83 |
| 4476 | *SMIM9* | 3.1771 | 1.2695 | 1.0394 | 0.41367 | 1.83 |
| 4477 | *SEC22C* | 2.3647 | 1.6087 | 1.5124 | -0.50286 | 1.83 |
| 4478 | *LRRC36* | 3.3631 | 1.5467 | 0.5747 | -0.12428 | 1.83 |
| 4479 | *VILL* | 3.2912 | 2.2825 | -0.09082 | -3.629 | 1.83 |
| 4480 | *GPALPP1* | 3.3411 | 1.1554 | 0.98629 | -0.48959 | 1.83 |
| 4481 | *POLM* | 2.5896 | 1.7331 | 1.1581 | 0.43394 | 1.83 |
| 4482 | *ZNF451* | 4.1945 | 0.91095 | 0.37508 | -3.8991 | 1.83 |
| 4483 | *TMEM235* | 2.9653 | 2.6917 | -0.17663 | -2.6476 | 1.83 |
| 4484 | *GCHFR* | 4.8341 | 0.81424 | -0.16962 | -1.8033 | 1.83 |
| 4485 | *SPATA33* | 4.6412 | 2.2848 | -1.4473 | -1.7945 | 1.83 |
| 4486 | *PRR13* | 4.5571 | 0.83087 | 0.08711 | -0.76099 | 1.83 |
| 4487 | *OR10AD1* | 2.9847 | 2.5695 | -0.08162 | -0.34017 | 1.82 |
| 4488 | *CLIP2* | 3.2009 | 1.5295 | 0.74083 | 0.33579 | 1.82 |
| 4489 | *KIF5C* | 4.3916 | 0.91153 | 0.16792 | -1.0955 | 1.82 |
| 4490 | *ADGRF3* | 4.3886 | 1.0676 | 0.01483 | -1.4983 | 1.82 |
| 4491 | *ZNF598* | 4.1336 | 0.88229 | 0.45485 | -0.17746 | 1.82 |
| 4492 | *PLD1* | 5.4087 | 0.38031 | -0.31916 | -1.9965 | 1.82 |
| 4493 | *ASL* | 4.805 | 1.7578 | -1.094 | -2.4642 | 1.82 |
| 4494 | *NBEAL2* | 2.9885 | 2.4062 | 0.07189 | -1.52 | 1.82 |
| 4495 | *CAMK1* | 2.2193 | 2.0664 | 1.1795 | -1.6089 | 1.82 |
| 4496 | *CCL25* | 3.229 | 1.1346 | 1.1005 | -2.8019 | 1.82 |
| 4497 | *TAPT1* | 3.4235 | 1.0912 | 0.94909 | -0.42938 | 1.82 |
| 4498 | *TPRX1* | 2.8909 | 1.8484 | 0.72332 | 0.61378 | 1.82 |
| 4499 | *OSCAR* | 3.3948 | 1.9333 | 0.13431 | -4.247 | 1.82 |
| 4500 | *ACADS* | 3.3079 | 1.4573 | 0.69682 | -0.9472 | 1.82 |
| 4501 | *KIRREL* | 3.5306 | 2.4735 | -0.54229 | -0.57342 | 1.82 |

| | | | | | | |
|---|---|---|---|---|---|---|
| 4502 | *CT45A2* | 4.1199 | -0.47913 | | | 1.82 |
| 4503 | *SFTA2* | 2.3271 | 2.014 | 1.12 | -0.61281 | 1.82 |
| 4504 | *ZNF497* | 3.844 | 1.0197 | 0.59737 | -1.3082 | 1.82 |
| 4505 | *CEP128* | 4.1173 | 2.4947 | -1.1511 | -2.8645 | 1.82 |
| 4506 | *DHFRL1* | 2.5111 | 2.2796 | 0.66964 | -0.84337 | 1.82 |
| 4507 | *GABRR3* | 3.0776 | 2.0356 | 0.3463 | -0.86261 | 1.82 |
| 4508 | *PLVAP* | 4.4033 | 1.3076 | -0.25198 | -2.2879 | 1.82 |
| 4509 | *SEMA7A* | 4.2026 | 2.0491 | -0.79422 | -2.7427 | 1.82 |
| 4510 | *CAPN3* | 4.0066 | 2.5452 | -1.0949 | -1.5912 | 1.82 |
| 4511 | *GABARAPL1* | 4.5431 | 1.1105 | -0.1967 | -0.99463 | 1.82 |
| 4512 | *FAM131A* | 2.9735 | 1.4686 | 1.0143 | -1.6575 | 1.82 |
| 4513 | *NUDT7* | 4.1481 | 0.67592 | 0.63229 | -1.3946 | 1.82 |
| 4514 | *ACP1* | 3.5573 | 1.6299 | 0.26835 | -1.448 | 1.82 |
| 4515 | *POTEB3* | 4.6336 | 0.48128 | 0.34029 | -1.9431 | 1.82 |
| 4516 | *CCDC160* | 4.4297 | 0.61036 | 0.41434 | -2.0116 | 1.82 |
| 4517 | *HIF3A* | 3.3559 | 2.3268 | -0.2283 | -2.5452 | 1.82 |
| 4518 | *PRR25* | 4.0541 | 1.0984 | 0.3004 | -1.8095 | 1.82 |
| 4519 | *RSPH9* | 3.8372 | 1.9046 | -0.28952 | -1.1075 | 1.82 |
| 4520 | *ARAF* | 3.6694 | 2.2832 | -0.50221 | -1.3184 | 1.82 |
| 4521 | *BDKRB1* | 3.0455 | 1.2686 | 1.1349 | 0.16498 | 1.82 |
| 4522 | *CYP2A7* | 3.8588 | 1.1817 | 0.40785 | 0.094924 | 1.82 |
| 4523 | *KRT28* | 5.251 | 0.17256 | 0.02418 | -0.2273 | 1.82 |
| 4524 | *MYCT1* | 2.8064 | 2.1529 | 0.48477 | -0.40223 | 1.81 |
| 4525 | *ZNF100* | 2.9114 | 1.3421 | 1.1896 | -1.4852 | 1.81 |
| 4526 | *SH2D2A* | 3.2288 | 1.8415 | 0.37258 | -0.34144 | 1.81 |
| 4527 | *NEIL2* | 2.9461 | 1.935 | 0.56161 | -1.2301 | 1.81 |
| 4528 | *SOCS2* | 4.9258 | 0.33218 | 0.18366 | 0.074008 | 1.81 |
| 4529 | *DRG2* | 3.1105 | 1.5056 | 0.8254 | 0.49507 | 1.81 |
| 4530 | *OR6B2* | 3.1964 | 1.5595 | 0.68482 | -0.06857 | 1.81 |
| 4531 | *KCTD9* | 4.3579 | 1.5062 | -0.42411 | -1.2758 | 1.81 |
| 4532 | *FAM19A2* | 2.9226 | 1.6751 | 0.84084 | -1.5663 | 1.81 |
| 4533 | *ECHDC2* | 4.5399 | 0.72495 | 0.1733 | -0.72468 | 1.81 |
| 4534 | *DCT* | 3.1433 | 1.3945 | 0.90025 | -1.0304 | 1.81 |
| 4535 | *KMT2B* | 2.3152 | 1.7441 | 1.3785 | -0.13224 | 1.81 |
| 4536 | *RNF175* | 1.9208 | 1.7889 | 1.7279 | 1.4211 | 1.81 |
| 4537 | *CELF6* | 5.0404 | 0.71818 | -0.32121 | -1.4736 | 1.81 |
| 4538 | *GPR176* | 3.2019 | 3.0518 | -0.81746 | -1.1465 | 1.81 |
| 4539 | *EGLN1* | 3.2476 | 1.9779 | 0.20644 | -1.2277 | 1.81 |
| 4540 | *NRROS* | 3.2398 | 1.1702 | 1.0212 | -1.57 | 1.81 |
| 4541 | *OR2G6* | 3.112 | 2.4778 | -0.16091 | -0.95474 | 1.81 |
| 4542 | *TMPRSS11B* | 2.5804 | 2.4461 | 0.40117 | 0.35867 | 1.81 |
| 4543 | *FAM186B* | 2.8959 | 1.3389 | 1.1903 | -1.2411 | 1.81 |
| 4544 | *GPR63* | 4.0797 | 0.97793 | 0.36744 | -1.2094 | 1.81 |
| 4545 | *C8orf46* | 4.4025 | 0.74095 | 0.28061 | -1.3056 | 1.81 |
| 4546 | *IL9R* | 3.8706 | 1.5689 | -0.01595 | -1.4269 | 1.81 |
| 4547 | *PAQR9* | 3.7511 | 2.1572 | -0.48973 | -2.005 | 1.81 |

| | | | | | | |
|---|---|---|---|---|---|---|
| 4548 | *IRF3* | 3.6116 | 2.1053 | -0.29835 | -1.1486 | 1.81 |
| 4549 | *MAPK10* | 3.3926 | 2.7366 | -0.71119 | -1.371 | 1.81 |
| 4550 | *HLA-DRB5* | 3.4596 | 1.5142 | 0.44404 | 0.052351 | 1.81 |
| 4551 | *OR1F1* | 2.3676 | 2.3253 | 0.72493 | -1.0852 | 1.81 |
| 4552 | *RGS19* | 4.3546 | 2.5875 | -1.5243 | -4.0338 | 1.81 |
| 4553 | *SRL* | 5.0704 | 0.37537 | -0.02943 | -0.21797 | 1.81 |
| 4554 | *ZDHHC16* | 3.2024 | 2.3264 | -0.1126 | -0.98842 | 1.81 |
| 4555 | *IGF2R* | 4.9715 | 2.4129 | -1.9686 | -1.9938 | 1.81 |
| 4556 | *FRG2* | 4.5464 | 2.1423 | -1.2736 | -2.0249 | 1.81 |
| 4557 | *TYMSOS* | 4.4998 | 0.86038 | 0.05417 | -1.7513 | 1.80 |
| 4558 | *MYL9* | 5.8886 | 0.013161 | -0.48756 | -1.5458 | 1.80 |
| 4559 | *CARD10* | 3.6578 | 2.7891 | -1.0329 | -1.2195 | 1.80 |
| 4560 | *BTC* | 3.9237 | 1.4258 | 0.06425 | -1.4509 | 1.80 |
| 4561 | *HACE1* | 3.1337 | 2.5635 | -0.28581 | -0.39127 | 1.80 |
| 4562 | *TEX15* | 2.927 | 1.3493 | 1.135 | -0.33444 | 1.80 |
| 4563 | *SLC6A8* | 5.2097 | 1.5211 | -1.3209 | -3.7932 | 1.80 |
| 4564 | *POTEB2* | 4.8714 | 1.7131 | -1.1747 | -1.5894 | 1.80 |
| 4565 | *ZNF404* | 3.117 | 2.9504 | -0.65811 | -0.85977 | 1.80 |
| 4566 | *GIMD1* | 4.1241 | 3.2055 | -1.9213 | -2.0542 | 1.80 |
| 4567 | *RPL8* | 4.3649 | 2.3004 | -1.257 | -2.4762 | 1.80 |
| 4568 | *PRSS37* | 2.7671 | 1.8835 | 0.75699 | -0.92695 | 1.80 |
| 4569 | *TMEM18* | 4.0183 | 0.97691 | 0.41106 | -1.6867 | 1.80 |
| 4570 | *FUBP1* | 3.3555 | 2.4852 | -0.43467 | -2.0102 | 1.80 |
| 4571 | *NANS* | 4.2434 | 1.4143 | -0.25332 | -1.0221 | 1.80 |
| 4572 | *ASB16* | 4.207 | 2.4085 | -1.2114 | -1.7789 | 1.80 |
| 4573 | *CCL7* | 4.2593 | 0.87196 | 0.27109 | -0.53009 | 1.80 |
| 4574 | *KCTD4* | 4.4711 | 1.5566 | -0.6254 | -1.0884 | 1.80 |
| 4575 | *SNED1* | 3.3388 | 2.2088 | -0.14556 | -1.4541 | 1.80 |
| 4576 | *PDF* | 4.1244 | 2.0628 | -0.78593 | -1.7963 | 1.80 |
| 4577 | *TBC1D32* | 2.6749 | 1.7188 | 1.0068 | -1.9523 | 1.80 |
| 4578 | *CCDC129* | 3.0841 | 1.8106 | 0.50457 | 0.48352 | 1.80 |
| 4579 | *DNAH7* | 3.4983 | 2.6835 | -0.78329 | -1.1764 | 1.80 |
| 4580 | *BMP3* | 3.8428 | 1.3075 | 0.24759 | -0.20079 | 1.80 |
| 4581 | *RGS14* | 3.3772 | 1.4923 | 0.52744 | -0.71723 | 1.80 |
| 4582 | *COL9A2* | 3.9082 | 0.8509 | 0.63773 | -0.87725 | 1.80 |
| 4583 | *LOC102724279* | 5.4371 | 0.50701 | -0.54743 | -1.1062 | 1.80 |
| 4584 | *CIZ1* | 2.3569 | 2.219 | 0.81814 | 0.19038 | 1.80 |
| 4585 | *BMP15* | 4.4689 | 1.3696 | -0.44528 | -0.54786 | 1.80 |
| 4586 | *GMFG* | 3.0507 | 2.5399 | -0.19984 | -0.88684 | 1.80 |
| 4587 | *NEUROD1* | 3.16 | 2.1736 | 0.05705 | -0.34352 | 1.80 |
| 4588 | *EMILIN1* | 3.7462 | 1.2928 | 0.35144 | -1.5138 | 1.80 |
| 4589 | *CNKSR3* | 4.4695 | 0.66081 | 0.2592 | -1.4209 | 1.80 |
| 4590 | *LCE5A* | 3.8432 | 1.9392 | -0.39338 | -0.50718 | 1.80 |
| 4591 | *SULT2A1* | 2.6325 | 2.607 | 0.14942 | -0.63572 | 1.80 |
| 4592 | *BCAN* | 2.9173 | 2.428 | 0.0418 | -1.5289 | 1.80 |
| 4593 | *FBXW12* | 3.7314 | 1.5436 | 0.11164 | -1.0884 | 1.80 |

| | | | | | | |
|---|---|---|---|---|---|---|
| 4594 | *NOX3* | 3.5909 | 1.6507 | 0.14491 | -1.1491 | 1.80 |
| 4595 | *TEP1* | 3.3431 | 3.1629 | -1.1214 | -1.3007 | 1.79 |
| 4596 | *OR10G9* | 3.6087 | 1.0506 | 0.72504 | -0.96407 | 1.79 |
| 4597 | *SLC26A10* | 2.6879 | 1.7922 | 0.90263 | -2.9068 | 1.79 |
| 4598 | *TRIM5* | 3.3243 | 2.1227 | -0.0658 | -1.5911 | 1.79 |
| 4599 | *DDO* | 3.1056 | 2.9255 | -0.65092 | -1.3594 | 1.79 |
| 4600 | *SAMD4B* | 4.3902 | 2.1272 | -1.1392 | -1.478 | 1.79 |
| 4601 | *USP12* | 4.6299 | 1.4761 | -0.72798 | -1.4182 | 1.79 |
| 4602 | *KIAA1429* | 2.9535 | 2.1966 | 0.22426 | -1.6049 | 1.79 |
| 4603 | *GXYLT1* | 4.4611 | 0.50981 | 0.40291 | -2.4229 | 1.79 |
| 4604 | *WDR45B* | 3.3119 | 1.1613 | 0.89991 | -0.7391 | 1.79 |
| 4605 | *MTDH* | 3.7495 | 1.8566 | -0.23404 | -2.3915 | 1.79 |
| 4606 | *PSKH2* | 3.7838 | 1.0686 | 0.51902 | -1.9302 | 1.79 |
| 4607 | *CDY1B* | 3.6587 | 1.4325 | 0.27983 | -0.31362 | 1.79 |
| 4608 | *ATP6V1C2* | 2.8892 | 2.1694 | 0.31225 | -1.6105 | 1.79 |
| 4609 | *FAM86C1* | 3.6891 | 2.2095 | -0.52839 | -1.1637 | 1.79 |
| 4610 | *MTCL1* | 3.7366 | 0.95663 | 0.6763 | -0.03342 | 1.79 |
| 4611 | *FBXO22* | 2.9716 | 2.1728 | 0.22456 | -1.2027 | 1.79 |
| 4612 | *NPC2* | 3.1803 | 1.8816 | 0.30552 | -0.54155 | 1.79 |
| 4613 | *P2RY12* | 4.5283 | 0.71573 | 0.12321 | -0.98653 | 1.79 |
| 4614 | *GSAP* | 2.9269 | 1.8796 | 0.56041 | -1.2732 | 1.79 |
| 4615 | *SNAP91* | 3.9105 | 1.2668 | 0.18868 | -2.0953 | 1.79 |
| 4616 | *S100A2* | 4.2196 | 0.71638 | 0.42947 | -2.0398 | 1.79 |
| 4617 | *SELPLG* | 3.2159 | 2.5344 | -0.38655 | -0.92028 | 1.79 |
| 4618 | *GRID2* | 4.8536 | 1.1567 | -0.64986 | -0.77127 | 1.79 |
| 4619 | *AMY1B* | 2.7254 | 2.277 | 0.35756 | -0.5169 | 1.79 |
| 4620 | *CERS4* | 4.5049 | 1.2763 | -0.42153 | -3.8502 | 1.79 |
| 4621 | *NIPSNAP3B* | 3.9874 | 2.3255 | -0.95388 | -1.4509 | 1.79 |
| 4622 | *ASXL3* | 5.3132 | 1.1057 | -1.0599 | -2.095 | 1.79 |
| 4623 | *KLHL22* | 3.4971 | 1.5607 | 0.30105 | -0.68911 | 1.79 |
| 4624 | *TDP2* | 4.0397 | 1.6454 | -0.3271 | -1.2305 | 1.79 |
| 4625 | *MRM1* | 3.1342 | 2.2871 | -0.06383 | -1.0021 | 1.79 |
| 4626 | *STAC3* | 3.6312 | 0.98335 | 0.7424 | -1.545 | 1.79 |
| 4627 | *PLD5* | 1.9092 | 1.7263 | 1.7208 | 1.1035 | 1.79 |
| 4628 | *SSPN* | 3.5032 | 2.2966 | -0.44353 | -0.86123 | 1.79 |
| 4629 | *FAM83D* | 2.9803 | 2.6983 | -0.32248 | -1.1961 | 1.79 |
| 4630 | *FOLR3* | 3.4089 | 1.6604 | 0.2846 | -1.2826 | 1.78 |
| 4631 | *FSHR* | 4.856 | 1.5232 | -1.0258 | -2.1802 | 1.78 |
| 4632 | *DOCK2* | 4.5625 | 2.4208 | -1.6313 | -2.28 | 1.78 |
| 4633 | *ATG4B* | 2.4771 | 2.457 | 0.4165 | -0.38675 | 1.78 |
| 4634 | *RBM39* | 2.5183 | 2.3435 | 0.48703 | -0.83584 | 1.78 |
| 4635 | *C2* | 2.728 | 2.3625 | 0.25821 | -0.6118 | 1.78 |
| 4636 | *GLRX3* | 3.425 | 2.2175 | -0.29402 | -0.78229 | 1.78 |
| 4637 | *SETD6* | 3.5708 | 3.514 | -1.7365 | -1.8157 | 1.78 |
| 4638 | *NOTCH1* | 4.0386 | 0.81337 | 0.4961 | -0.36246 | 1.78 |
| 4639 | *OCA2* | 3.6414 | 1.0213 | 0.6851 | -0.06182 | 1.78 |

| | | | | | | |
|---|---|---|---|---|---|---|
| 4640 | *PSMD10* | 4.2152 | 0.61617 | 0.51616 | -1.1676 | 1.78 |
| 4641 | *USP17L2* | 3.0139 | 2.7078 | -0.37417 | -1.8822 | 1.78 |
| 4642 | *KIT* | 4.5948 | 1.1242 | -0.37194 | -1.2052 | 1.78 |
| 4643 | *WDR63* | 3.3723 | 1.5491 | 0.42458 | 0.20047 | 1.78 |
| 4644 | *SULT1C4* | 4.682 | 0.6335 | 0.02927 | -0.17439 | 1.78 |
| 4645 | *CD3G* | 4.8061 | 0.89936 | -0.36086 | -0.43472 | 1.78 |
| 4646 | *GZF1* | 3.6224 | 1.3173 | 0.40417 | -1.2827 | 1.78 |
| 4647 | *CPNE2* | 3.1505 | 2.1701 | 0.023 | -0.35272 | 1.78 |
| 4648 | *TOR4A* | 2.9906 | 1.2282 | 1.1244 | 0.58508 | 1.78 |
| 4649 | *DEFB104B* | 3.0964 | 1.6217 | 0.62473 | -1.0029 | 1.78 |
| 4650 | *TFPI2* | 4.7864 | 1.8498 | -1.2935 | -1.4498 | 1.78 |
| 4651 | *L1TD1* | 2.7583 | 1.912 | 0.67208 | -0.08356 | 1.78 |
| 4652 | *NDUFS1* | 3.4638 | 1.9627 | -0.08477 | -0.65889 | 1.78 |
| 4653 | *TCEB3B* | 2.8491 | 2.2498 | 0.24221 | -0.64164 | 1.78 |
| 4654 | *SPEN* | 2.5741 | 1.9402 | 0.82658 | -0.4464 | 1.78 |
| 4655 | *EXOC2* | 4.1906 | 0.67748 | 0.4727 | -1.1756 | 1.78 |
| 4656 | *OXNAD1* | 3.6545 | 1.0047 | 0.68058 | -0.36855 | 1.78 |
| 4657 | *PGK1* | 2.616 | 1.6724 | 1.0505 | -0.93024 | 1.78 |
| 4658 | *DNTT* | 4.61 | 0.45281 | 0.27574 | -3.087 | 1.78 |
| 4659 | *PGBD4* | 3.4954 | 2.6674 | -0.8253 | -1.0486 | 1.78 |
| 4660 | *SHMT1* | 3.0849 | 2.696 | -0.44353 | -2.4193 | 1.78 |
| 4661 | *DENND5B* | 3.2516 | 1.1903 | 0.89509 | -0.97509 | 1.78 |
| 4662 | *CLTA* | 3.2583 | 1.0428 | 1.0357 | 0.33194 | 1.78 |
| 4663 | *SLC1A4* | 4.1617 | 1.703 | -0.52795 | -1.8462 | 1.78 |
| 4664 | *UBLCP1* | 4.5649 | 1.3467 | -0.57625 | -0.91793 | 1.78 |
| 4665 | *SEC61A2* | 2.459 | 2.2953 | 0.58038 | -2.1134 | 1.78 |
| 4666 | *OR12D2* | 3.8493 | 1.1399 | 0.34516 | -1.12 | 1.78 |
| 4667 | *ARHGAP33* | 4.1817 | 0.70819 | 0.44389 | -2.7384 | 1.78 |
| 4668 | *RASSF9* | 3.8081 | 1.9487 | -0.42359 | -1.0226 | 1.78 |
| 4669 | *FAM107B* | 4.9622 | 0.65359 | -0.28301 | -0.9382 | 1.78 |
| 4670 | *KCNN3* | 4.1919 | 2.2169 | -1.0761 | -1.6781 | 1.78 |
| 4671 | *ZNF268* | 3.3772 | 1.2828 | 0.67234 | -0.36913 | 1.78 |
| 4672 | *CPAMD8* | 1.906 | 1.7857 | 1.6406 | 0.96239 | 1.78 |
| 4673 | *TMEM108* | 3.3284 | 1.4865 | 0.51694 | -1.0343 | 1.78 |
| 4674 | *GADD45A* | 3.5984 | 2.0582 | -0.32719 | -1.1258 | 1.78 |
| 4675 | *NR1D1* | 4.8055 | 1.7972 | -1.2735 | -1.9748 | 1.78 |
| 4676 | *HDDC3* | 3.3958 | 1.3677 | 0.56529 | -0.714 | 1.78 |
| 4677 | *NOA1* | 4.357 | 0.97102 | 0.00025 | -1.8858 | 1.78 |
| 4678 | *C17orf97* | 5.3707 | 0.12602 | -0.16868 | -1.2367 | 1.78 |
| 4679 | *SPERT* | 4.1694 | 1.1016 | 0.05557 | -2.672 | 1.78 |
| 4680 | *CD164* | 3.9352 | 2.119 | -0.72772 | -1.8836 | 1.78 |
| 4681 | *B4GALNT1* | 3.7139 | 2.0311 | -0.41917 | -2.0062 | 1.78 |
| 4682 | *ZNF410* | 2.3781 | 1.669 | 1.2778 | 1.0505 | 1.77 |
| 4683 | *SPACA5B* | 2.4474 | 2.4382 | 0.43862 | -0.13635 | 1.77 |
| 4684 | *OR51E1* | 4.2023 | 1.5038 | -0.38215 | -1.0723 | 1.77 |
| 4685 | *PDCD6* | 4.2894 | 1.1182 | -0.0869 | -1.5243 | 1.77 |

| | | | | | | |
|---|---|---|---|---|---|---|
| 4686 | *ZC3H3* | 4.3008 | 0.56359 | 0.45591 | -0.41596 | 1.77 |
| 4687 | *MATN3* | 4.5289 | 0.82402 | -0.03342 | -1.2076 | 1.77 |
| 4688 | *GPATCH3* | 3.68 | 1.6866 | -0.04862 | -0.52786 | 1.77 |
| 4689 | *OR4C45* | 3.3911 | 2.1015 | -0.17498 | -0.3819 | 1.77 |
| 4690 | *DGKA* | 3.5789 | 3.4165 | -1.6791 | -1.8572 | 1.77 |
| 4691 | *PLXNB2* | 2.503 | 1.7172 | 1.0961 | -0.47527 | 1.77 |
| 4692 | *SNPH* | 4.1885 | 2.2915 | -1.1637 | -2.0675 | 1.77 |
| 4693 | *NCS1* | 5.3472 | 0.12509 | -0.15842 | -0.8792 | 1.77 |
| 4694 | *PHF19* | 3.4076 | 1.7167 | 0.18857 | -0.3589 | 1.77 |
| 4695 | *OR10AG1* | 4.5706 | 1.239 | -0.49704 | -1.7945 | 1.77 |
| 4696 | *GAL3ST4* | 3.2993 | 1.1491 | 0.86377 | -0.18353 | 1.77 |
| 4697 | *RAB3IP* | 3.1183 | 2.9396 | -0.74618 | -3.1317 | 1.77 |
| 4698 | *CCDC178* | 4.9133 | 0.92474 | -0.52643 | -1.6186 | 1.77 |
| 4699 | *NGLY1* | 3.9479 | 0.87678 | 0.48647 | -1.0212 | 1.77 |
| 4700 | *PHLPP2* | 2.8875 | 1.6807 | 0.74217 | -1.6386 | 1.77 |
| 4701 | *FOXD3* | 4.6705 | 0.43125 | 0.20829 | -2.4015 | 1.77 |
| 4702 | *LDHB* | 4.0129 | 2.2925 | -0.99608 | -1.0837 | 1.77 |
| 4703 | *SIRPB2* | 2.6044 | 2.4584 | 0.24553 | -1.6766 | 1.77 |
| 4704 | *CHAF1B* | 5.2088 | 0.21956 | -0.12054 | -0.81496 | 1.77 |
| 4705 | *PHYHIP* | 3.0523 | 1.2322 | 1.023 | -1.2975 | 1.77 |
| 4706 | *MGP* | 4.029 | 1.6647 | -0.38675 | -0.57067 | 1.77 |
| 4707 | *EMP3* | 3.6615 | 1.8534 | -0.20849 | -0.72654 | 1.77 |
| 4708 | *OR6B3* | 3.9152 | 0.98298 | 0.40814 | -0.98238 | 1.77 |
| 4709 | *N4BP2* | 4.7197 | 1.8818 | -1.2958 | -1.522 | 1.77 |
| 4710 | *PRSS8* | 2.3528 | 2.2942 | 0.65808 | -0.9519 | 1.77 |
| 4711 | *OR5M11* | 3.1115 | 1.4052 | 0.78827 | -1.6411 | 1.77 |
| 4712 | *COPZ1* | 5.4908 | 0.40306 | -0.58921 | -1.1987 | 1.77 |
| 4713 | *AWAT2* | 2.6283 | 2.0896 | 0.58644 | -1.6731 | 1.77 |
| 4714 | *TRAFD1* | 3.8792 | 0.86789 | 0.55585 | -2.4735 | 1.77 |
| 4715 | *NDUFB7* | 2.2461 | 1.6641 | 1.3897 | 0.70422 | 1.77 |
| 4716 | *HLA-B* | 2.2418 | 1.6882 | 1.3692 | -1.8591 | 1.77 |
| 4717 | *BAK1* | 3.199 | 1.9915 | 0.10865 | 0.076527 | 1.77 |
| 4718 | *ZNF662* | 4.9082 | 2.1995 | -1.8087 | -1.8906 | 1.77 |
| 4719 | *SRXN1* | 2.5599 | 1.378 | 1.3608 | -0.47496 | 1.77 |
| 4720 | *HEXB* | 3.981 | 1.5653 | -0.24946 | -1.4741 | 1.77 |
| 4721 | *DLK2* | 5.1843 | 0.25894 | -0.14649 | -1.3458 | 1.77 |
| 4722 | *TSTD3* | 4.9275 | 0.74561 | -0.37686 | -0.60801 | 1.77 |
| 4723 | *PDE6C* | 3.6387 | 1.2825 | 0.37471 | 0.077067 | 1.77 |
| 4724 | *C5orf22* | 2.1897 | 1.8071 | 1.2991 | -0.15263 | 1.77 |
| 4725 | *GAGE2A* | 4.6054 | 0.45567 | 0.23473 | -0.4482 | 1.77 |
| 4726 | *KIAA1841* | 2.7691 | 1.8296 | 0.69676 | -0.00832 | 1.77 |
| 4727 | *MMP27* | 3.2901 | 2.2073 | -0.20239 | -2.0837 | 1.77 |
| 4728 | *SLFN11* | 4.8674 | 0.94571 | -0.51868 | -1.2832 | 1.76 |
| 4729 | *LRRC16A* | 3.8262 | 1.3322 | 0.13578 | -1.1886 | 1.76 |
| 4730 | *SERPINB1* | 5.1471 | 0.52306 | -0.37619 | -0.96144 | 1.76 |
| 4731 | *UCHL1* | 3.6231 | 1.9113 | -0.24058 | -0.60717 | 1.76 |

| | | | | | | |
|---|---|---|---|---|---|---|
| 4732 | *TPD52L3* | 4.6672 | 1.1476 | -0.52122 | -1.3875 | 1.76 |
| 4733 | *GUCA2B* | 3.2922 | 2.5215 | -0.52017 | -1.9746 | 1.76 |
| 4734 | *SRCIN1* | 2.1866 | 1.7811 | 1.3258 | -1.0898 | 1.76 |
| 4735 | *OR52R1* | 3.0336 | 1.7357 | 0.52383 | -0.20952 | 1.76 |
| 4736 | *NYAP1* | 4.9572 | 0.84886 | -0.51304 | -2.4286 | 1.76 |
| 4737 | *GPR158* | 3.2054 | 1.8158 | 0.27146 | -0.15394 | 1.76 |
| 4738 | *ACOT2* | 4.6245 | 0.74656 | -0.07845 | -1.2194 | 1.76 |
| 4739 | *CNN3* | 2.8212 | 2.23 | 0.24124 | -0.37253 | 1.76 |
| 4740 | *DCSTAMP* | 3.2446 | 1.9753 | 0.07022 | -1.4968 | 1.76 |
| 4741 | *WDR59* | 4.6996 | 0.60079 | -0.01081 | -0.74242 | 1.76 |
| 4742 | *NEK8* | 4.1151 | 2.2388 | -1.0645 | -1.4374 | 1.76 |
| 4743 | *PIWIL3* | 2.7189 | 1.7355 | 0.83445 | -0.11953 | 1.76 |
| 4744 | *SMAD9* | 4.5083 | 1.6604 | -0.88008 | -1.5741 | 1.76 |
| 4745 | *POR* | 2.8943 | 2.0838 | 0.31033 | -0.58295 | 1.76 |
| 4746 | *SLC2A6* | 3.4066 | 1.1242 | 0.75751 | 0.34657 | 1.76 |
| 4747 | *LRRC14B* | 4.3015 | 1.2268 | -0.24035 | -0.38796 | 1.76 |
| 4748 | *BAG3* | 3.5915 | 2.392 | -0.69758 | -2.0629 | 1.76 |
| 4749 | *MBOAT7* | 3.0042 | 1.7866 | 0.49511 | 0.048132 | 1.76 |
| 4750 | *GLTSCR2* | 3.7629 | 2.505 | -0.98239 | -2.329 | 1.76 |
| 4751 | *ITGB3BP* | 5.0738 | 0.30621 | -0.0952 | -0.35038 | 1.76 |
| 4752 | *ERVV-2* | 4.2321 | 1.3337 | -0.28308 | -0.35232 | 1.76 |
| 4753 | *NPIPB9* | 2.3887 | 1.7013 | 1.1917 | -2.4709 | 1.76 |
| 4754 | *ATP8A1* | 4.3616 | 0.88577 | 0.03422 | -1.2954 | 1.76 |
| 4755 | *PRSS46* | 3.0659 | 1.2284 | 0.98694 | -0.2098 | 1.76 |
| 4756 | *UTP14C* | 4.6286 | 1.355 | -0.70255 | -1.0503 | 1.76 |
| 4757 | *ABCF3* | 4.0702 | 1.5472 | -0.33645 | -0.88418 | 1.76 |
| 4758 | *GFY* | 3.8109 | 1.8665 | -0.39676 | -1.776 | 1.76 |
| 4759 | *LIPN* | 3.9847 | 1.6117 | -0.31641 | -1.4561 | 1.76 |
| 4760 | *TCTA* | 4.5894 | 1.0632 | -0.37323 | -1.7647 | 1.76 |
| 4761 | *HIAT1* | 2.8825 | 1.2703 | 1.1265 | 1.0351 | 1.76 |
| 4762 | *ZSCAN2* | 4.0413 | 0.63118 | 0.60573 | -1.5458 | 1.76 |
| 4763 | *SCN4A* | 2.8867 | 2.3169 | 0.07454 | -0.2562 | 1.76 |
| 4764 | *RAB3B* | 3.0973 | 2.0168 | 0.16383 | -0.89699 | 1.76 |
| 4765 | *C3orf33* | 4.957 | 2.013 | -1.6926 | -2.163 | 1.76 |
| 4766 | *SETSIP* | 5.5839 | 0.030302 | -0.33741 | -2.033 | 1.76 |
| 4767 | *ZBED8* | 3.0456 | 2.6135 | -0.38231 | -0.67243 | 1.76 |
| 4768 | *TRIM61* | 2.4249 | 2.3289 | 0.52247 | -0.53689 | 1.76 |
| 4769 | *IWS1* | 3.6619 | 1.6226 | -0.00829 | -1.1287 | 1.76 |
| 4770 | *LGR4* | 3.9193 | 1.2217 | 0.13507 | -0.24508 | 1.76 |
| 4771 | *PPAN* | 4.1652 | 0.64045 | 0.46987 | -1.8371 | 1.76 |
| 4772 | *FAM89A* | 2.8245 | 2.1036 | 0.34664 | -1.3727 | 1.76 |
| 4773 | *WDR48* | 3.5966 | 1.4721 | 0.20547 | -1.0564 | 1.76 |
| 4774 | *LYPLAL1* | 2.9246 | 2.1361 | 0.21312 | -0.63575 | 1.76 |
| 4775 | *C6orf52* | 3.7643 | 1.0862 | 0.4229 | -1.2208 | 1.76 |
| 4776 | *CCDC152* | 2.7755 | 2.3877 | 0.10996 | -0.77114 | 1.76 |
| 4777 | *STK39* | 4.0412 | 1.1434 | 0.08797 | -0.22313 | 1.76 |

| | | | | | | |
|---|---|---|---|---|---|---|
| 4778 | *CCNB3* | 3.5079 | 1.1289 | 0.63569 | 0.3281 | 1.76 |
| 4779 | *OR13D1* | 3.2955 | 1.401 | 0.57567 | -0.22251 | 1.76 |
| 4780 | *STIM2* | 3.5178 | 1.0588 | 0.69473 | -1.1534 | 1.76 |
| 4781 | *LDB1* | 5.9458 | -0.21491 | -0.45972 | -2.1125 | 1.76 |
| 4782 | *LCN6* | 3.6752 | 0.80054 | 0.79535 | -0.92284 | 1.76 |
| 4783 | *RD3L* | 4.814 | 0.29413 | 0.16285 | -0.55586 | 1.76 |
| 4784 | *AMY2B* | 4.179 | 1.1842 | -0.09268 | -1.6524 | 1.76 |
| 4785 | *IQGAP2* | 4.4879 | 0.50788 | 0.27474 | -0.00303 | 1.76 |
| 4786 | *IP6K3* | 3.7008 | 1.8067 | -0.23758 | -1.5667 | 1.76 |
| 4787 | *NRP1* | 2.1671 | 2.1016 | 0.99931 | -0.60347 | 1.76 |
| 4788 | *CASP3* | 3.8618 | 0.72204 | 0.68303 | -2.0146 | 1.76 |
| 4789 | *ZBTB41* | 4.0576 | 1.0265 | 0.1827 | -0.58404 | 1.76 |
| 4790 | *OR6F1* | 2.6157 | 2.4631 | 0.18707 | -0.30574 | 1.76 |
| 4791 | *GLTPD2* | 3.6805 | 2.4859 | -0.90207 | -1.2188 | 1.75 |
| 4792 | *KCTD18* | 2.6846 | 1.7562 | 0.82297 | 0.62749 | 1.75 |
| 4793 | *ZFYVE19* | 2.8452 | 2.498 | -0.07952 | -1.8453 | 1.75 |
| 4794 | *HEY1* | 2.7415 | 1.3376 | 1.1837 | -1.0359 | 1.75 |
| 4795 | *WARS2* | 3.7919 | 1.3916 | 0.07906 | -0.57967 | 1.75 |
| 4796 | *ADAP1* | 4.0635 | 2.6477 | -1.4493 | -1.4842 | 1.75 |
| 4797 | *ANK1* | 3.4987 | 2.8754 | -1.1137 | -1.359 | 1.75 |
| 4798 | *CCDC69* | 2.5173 | 1.6059 | 1.1367 | -0.96647 | 1.75 |
| 4799 | *C1orf123* | 3.7305 | 1.4954 | 0.03357 | -2.1802 | 1.75 |
| 4800 | *MRPS15* | 5.2488 | 0.13368 | -0.12425 | -1.3287 | 1.75 |
| 4801 | *NINJ2* | 2.8577 | 1.2976 | 1.1029 | -0.86756 | 1.75 |
| 4802 | *PTCHD3* | 4.2426 | 2.045 | -1.0303 | -1.9935 | 1.75 |
| 4803 | *WDR26* | 3.8237 | 0.78888 | 0.64447 | -1.2863 | 1.75 |
| 4804 | *FAM71F1* | 2.5816 | 2.2349 | 0.44052 | -0.5271 | 1.75 |
| 4805 | *REEP6* | 2.844 | 2.24 | 0.17212 | -1.5121 | 1.75 |
| 4806 | *FAM24A* | 3.0812 | 1.185 | 0.98936 | -0.52154 | 1.75 |
| 4807 | *PRRT2* | 4.7738 | 0.97445 | -0.49311 | -1.6525 | 1.75 |
| 4808 | *CEP41* | 4.7796 | 1.334 | -0.85871 | -0.98842 | 1.75 |
| 4809 | *TAAR9* | 2.7077 | 2.5943 | -0.04775 | -1.1838 | 1.75 |
| 4810 | *GUCA2A* | 3.0579 | 1.8994 | 0.29636 | -0.71063 | 1.75 |
| 4811 | *SDHAF1* | 3.1443 | 2.3828 | -0.27359 | -1.8179 | 1.75 |
| 4812 | *CACNA1S* | 5.4295 | -0.04425 | -0.1318 | -0.45867 | 1.75 |
| 4813 | *ATOH7* | 3.7812 | 1.6744 | -0.20246 | -1.7024 | 1.75 |
| 4814 | *STX7* | 2.8616 | 2.2291 | 0.16232 | 0.15451 | 1.75 |
| 4815 | *ZNF696* | 3.1538 | 1.577 | 0.52198 | -0.34266 | 1.75 |
| 4816 | *TGIF2LX* | 4.0672 | 1.1182 | 0.06701 | -0.98907 | 1.75 |
| 4817 | *GATSL2* | 4.4018 | 1.6158 | -0.76529 | -1.4404 | 1.75 |
| 4818 | *CNR1* | 3.6052 | 1.9116 | -0.26468 | -0.58603 | 1.75 |
| 4819 | *NES* | 5.7279 | 0.04489 | -0.52154 | -0.97138 | 1.75 |
| 4820 | *GRIK2* | 3.3464 | 2.5725 | -0.66767 | -1.8253 | 1.75 |
| 4821 | *OR52B2* | 2.527 | 1.7249 | 0.99825 | -0.43473 | 1.75 |
| 4822 | *CCR10* | 3.0011 | 1.3174 | 0.93087 | -0.23297 | 1.75 |
| 4823 | *MEA1* | 5.2091 | 0.12358 | -0.08446 | -1.0301 | 1.75 |

| | | | | | | |
|---|---|---|---|---|---|---|
| 4824 | *NME2* | 4.4033 | 2.445 | -1.6011 | -2.4595 | 1.75 |
| 4825 | *CRYBG3* | 2.9335 | 2.7671 | -0.45403 | -1.7967 | 1.75 |
| 4826 | *PUS10* | 3.7435 | 2.1266 | -0.62423 | -1.6181 | 1.75 |
| 4827 | *SLC22A7* | 3.4023 | 1.4629 | 0.38025 | -0.07154 | 1.75 |
| 4828 | *C14orf2* | 3.2266 | 3.0356 | -1.017 | -1.6526 | 1.75 |
| 4829 | *CHAC1* | 4.3931 | 0.49834 | 0.35324 | -1.6246 | 1.75 |
| 4830 | *CBX8* | 2.7742 | 2.5315 | -0.06297 | -0.29121 | 1.75 |
| 4831 | *CLEC3A* | 3.36 | 1.5342 | 0.34848 | 0.27664 | 1.75 |
| 4832 | *TIFA* | 3.5323 | 1.9803 | -0.27044 | -1.0743 | 1.75 |
| 4833 | *BIK* | 3.2908 | 3.0742 | -1.1241 | -1.7364 | 1.75 |
| 4834 | *ARL14EPL* | 4.5704 | 0.68273 | -0.01337 | -0.25227 | 1.75 |
| 4835 | *SMAD7* | 2.9299 | 2.7533 | -0.44353 | -1.8227 | 1.75 |
| 4836 | *THRA* | 2.8779 | 2.4224 | -0.06103 | -0.63912 | 1.75 |
| 4837 | *INF2* | 4.9462 | 1.2181 | -0.92524 | -1.3937 | 1.75 |
| 4838 | *TMEM87B* | 2.7755 | 2.7438 | -0.28047 | -0.81136 | 1.75 |
| 4839 | *TNFRSF14* | 3.7643 | 0.87333 | 0.59983 | 0.35462 | 1.75 |
| 4840 | *SERPIND1* | 3.089 | 2.0748 | 0.07319 | -0.25803 | 1.75 |
| 4841 | *MIDN* | 4.1874 | 0.67533 | 0.37426 | -1.0999 | 1.75 |
| 4842 | *MTHFD1L* | 4.087 | 2.8939 | -1.7445 | -2.4472 | 1.75 |
| 4843 | *MRPS33* | 3.8889 | 1.2333 | 0.11357 | -0.01785 | 1.75 |
| 4844 | *ACSL1* | 5.1089 | 1.0691 | -0.94312 | -1.1805 | 1.74 |
| 4845 | *ZBTB22* | 2.9288 | 1.5808 | 0.72505 | -1.1316 | 1.74 |
| 4846 | *MAP3K11* | 2.6453 | 2.0723 | 0.51667 | 0.43942 | 1.74 |
| 4847 | *ABCG2* | 4.2347 | 1.0594 | -0.06182 | -0.559 | 1.74 |
| 4848 | *MYO1F* | 5.2218 | 0.85335 | -0.84337 | -1.4287 | 1.74 |
| 4849 | *CLCN2* | 4.7026 | 1.3605 | -0.83184 | -2.2996 | 1.74 |
| 4850 | *TGOLN2* | 3.6173 | 1.7858 | -0.17257 | -0.8901 | 1.74 |
| 4851 | *DRG1* | 3.4987 | 1.5644 | 0.16693 | -0.46056 | 1.74 |
| 4852 | *PRMT9* | 3.5642 | 1.122 | 0.54379 | -0.43146 | 1.74 |
| 4853 | *SOST* | 2.6967 | 2.2934 | 0.23955 | -0.62489 | 1.74 |
| 4854 | *CDC42EP5* | 3.0376 | 2.2057 | -0.01399 | -1.3769 | 1.74 |
| 4855 | *JADE1* | 2.8679 | 1.2778 | 1.0807 | -1.0948 | 1.74 |
| 4856 | *MS4A2* | 4.043 | 0.76415 | 0.41894 | 0.071886 | 1.74 |
| 4857 | *COX7A2* | 3.3115 | 1.8396 | 0.07481 | -0.52092 | 1.74 |
| 4858 | *DNAJB7* | 3.5714 | 1.7332 | -0.08008 | -1.8085 | 1.74 |
| 4859 | *GTF2IRD2B* | 2.5144 | 1.531 | 1.1788 | -1.5538 | 1.74 |
| 4860 | *PCDH20* | 3.8228 | 0.96498 | 0.4358 | -1.5765 | 1.74 |
| 4861 | *LRRIQ3* | 3.0639 | 2.6198 | -0.46017 | -2.4011 | 1.74 |
| 4862 | *PPFIA3* | 4.0714 | 0.68097 | 0.47084 | -1.4273 | 1.74 |
| 4863 | *ZNF30* | 3.3267 | 1.2809 | 0.61525 | -1.3523 | 1.74 |
| 4864 | *MED29* | 4.6019 | 1.8301 | -1.2098 | -1.5684 | 1.74 |
| 4865 | *FAM122B* | 3.606 | 2.0918 | -0.4759 | -0.50577 | 1.74 |
| 4866 | *TRO* | 3.07 | 1.2052 | 0.94444 | -2.2064 | 1.74 |
| 4867 | *NANP* | 4.0934 | 1.5519 | -0.42573 | -1.1343 | 1.74 |
| 4868 | *MKI67* | 2.2543 | 1.8239 | 1.1407 | -1.7215 | 1.74 |
| 4869 | *ZNF48* | 2.7027 | 1.7835 | 0.73238 | -1.3948 | 1.74 |

| | | | | | | |
|---|---|---|---|---|---|---|
| 4870 | *BRAT1* | 3.8551 | 2.8307 | -1.4685 | -1.4864 | 1.74 |
| 4871 | *MMP12* | 3.8365 | 1.6376 | -0.25737 | -0.30045 | 1.74 |
| 4872 | *SPATA45* | 3.8272 | 2.329 | -0.9428 | -2.1182 | 1.74 |
| 4873 | *BLVRA* | 4.2205 | 0.66027 | 0.33167 | 0.076188 | 1.74 |
| 4874 | *C11orf30* | 5.5017 | 0.17607 | -0.46721 | -0.8417 | 1.74 |
| 4875 | *BNC1* | 2.8332 | 1.8788 | 0.49816 | -1.2065 | 1.74 |
| 4876 | *ZNF398* | 3.7666 | 1.6272 | -0.18432 | -0.42171 | 1.74 |
| 4877 | *TCTE1* | 3.5729 | 1.373 | 0.26246 | -0.04618 | 1.74 |
| 4878 | *IL3* | 4.0321 | 0.631 | 0.54446 | -0.37546 | 1.74 |
| 4879 | *ZNF583* | 2.7377 | 1.4344 | 1.0342 | 0.23549 | 1.74 |
| 4880 | *UBE2O* | 3.3307 | 1.516 | 0.35833 | -0.98062 | 1.74 |
| 4881 | *CIB2* | 3.8058 | 2.3279 | -0.93099 | -2.2669 | 1.73 |
| 4882 | *TBC1D9* | 3.9834 | 1.5919 | -0.37261 | -0.95796 | 1.73 |
| 4883 | *GRIA4* | 4.7638 | 1.7075 | -1.2694 | -2.0536 | 1.73 |
| 4884 | *ZC3H13* | 3.7965 | 1.7757 | -0.37052 | -1.3247 | 1.73 |
| 4885 | *CCNG1* | 3.8845 | 0.7079 | 0.60902 | -1.1866 | 1.73 |
| 4886 | *GCM1* | 3.5658 | 2.5248 | -0.8897 | -1.9513 | 1.73 |
| 4887 | *RHOJ* | 2.2013 | 2.0868 | 0.91252 | -1.2852 | 1.73 |
| 4888 | *COPS7B* | 4.4129 | 0.48698 | 0.30057 | -1.5368 | 1.73 |
| 4889 | *CDKL5* | 3.1576 | 1.853 | 0.18983 | -0.28355 | 1.73 |
| 4890 | *MMAA* | 2.4973 | 1.6528 | 1.05 | -0.32254 | 1.73 |
| 4891 | *SEMA3G* | 2.7984 | 2.5244 | -0.12326 | -0.47818 | 1.73 |
| 4892 | *SLC30A5* | 4.7201 | 1.0906 | -0.61164 | -1.1138 | 1.73 |
| 4893 | *MPP3* | 3.4365 | 1.3685 | 0.39364 | -0.47083 | 1.73 |
| 4894 | *LILRB2* | 4.1145 | 2.4001 | -1.316 | -1.9227 | 1.73 |
| 4895 | *MXRA7* | 2.8021 | 1.8708 | 0.52528 | -0.82628 | 1.73 |
| 4896 | *PML* | 2.3997 | 1.7608 | 1.0368 | -0.36841 | 1.73 |
| 4897 | *C11orf16* | 4.3379 | 1.1288 | -0.26984 | -0.37787 | 1.73 |
| 4898 | *MAP2K3* | 4.2737 | 1.3671 | -0.44511 | -0.69865 | 1.73 |
| 4899 | *FOXJ3* | 3.0064 | 2.4633 | -0.27443 | -0.53843 | 1.73 |
| 4900 | *CST3* | 3.3733 | 1.2264 | 0.59549 | -0.03316 | 1.73 |
| 4901 | *LZTR1* | 3.644 | 1.9578 | -0.40671 | -2.2794 | 1.73 |
| 4902 | *IDE* | 4.2255 | 0.57882 | 0.39043 | -0.5293 | 1.73 |
| 4903 | *NDC1* | 2.9352 | 1.6468 | 0.61214 | -1.5053 | 1.73 |
| 4904 | *EVI5* | 2.8496 | 2.2017 | 0.14196 | -1.1008 | 1.73 |
| 4905 | *GUCY1B3* | 3.5176 | 1.7388 | -0.06418 | -1.6061 | 1.73 |
| 4906 | *PPEF2* | 5.5018 | -0.1549 | -0.15499 | -0.95933 | 1.73 |
| 4907 | *IER5L* | 3.0544 | 2.018 | 0.11924 | 0.015043 | 1.73 |
| 4908 | *RIMKLB* | 2.6272 | 1.9865 | 0.57708 | 0.46392 | 1.73 |
| 4909 | *CYP2W1* | 2.4351 | 2.1003 | 0.65475 | -1.853 | 1.73 |
| 4910 | *POU3F4* | 3.5072 | 0.84072 | 0.84049 | -0.70438 | 1.73 |
| 4911 | *GLI4* | 3.1637 | 2.2346 | -0.21049 | -1.4504 | 1.73 |
| 4912 | *GPR55* | 3.3584 | 2.5344 | -0.70513 | -0.91685 | 1.73 |
| 4913 | *KLHDC1* | 4.0103 | 0.61689 | 0.5598 | 0.32083 | 1.73 |
| 4914 | *GSDMC* | 4.179 | 0.77651 | 0.23049 | -0.51585 | 1.73 |
| 4915 | *PLSCR4* | 4.8281 | 0.80931 | -0.45185 | -1.4128 | 1.73 |

| | | | | | | |
|---|---|---|---|---|---|---|
| 4916 | *ZNF454* | 2.9739 | 1.5553 | 0.65612 | -0.04111 | 1.73 |
| 4917 | *ZBTB24* | 5.046 | 0.61874 | -0.47964 | -0.97164 | 1.73 |
| 4918 | *UBALD2* | 2.3624 | 2.2384 | 0.58319 | 0.3 | 1.73 |
| 4919 | *PRR5* | 3.5772 | 1.3897 | 0.21695 | -1.0733 | 1.73 |
| 4920 | *SKAP2* | 3.3815 | 1.9794 | -0.17712 | -1.0443 | 1.73 |
| 4921 | *KRT81* | 5.0999 | 0.11164 | -0.03036 | -2.0706 | 1.73 |
| 4922 | *PPP1R9A* | 2.6128 | 2.2087 | 0.35931 | -0.72529 | 1.73 |
| 4923 | *CHST13* | 2.5437 | 1.6864 | 0.9505 | -0.68879 | 1.73 |
| 4924 | *ANKDD1B* | 2.8177 | 2.6011 | -0.23916 | -1.2719 | 1.73 |
| 4925 | *CAMSAP3* | 5.3852 | 0.98986 | -1.197 | -1.2932 | 1.73 |
| 4926 | *KRT31* | 3.9983 | 0.97324 | 0.20649 | -1.9358 | 1.73 |
| 4927 | *ELAVL2* | 4.839 | 0.18778 | 0.15064 | -1.049 | 1.73 |
| 4928 | *ESRP1* | 3.866 | 0.96444 | 0.34693 | 0.32682 | 1.73 |
| 4929 | *AMIGO3* | 3.9176 | 2.2121 | -0.95322 | -1.4491 | 1.73 |
| 4930 | *FOXS1* | 2.6513 | 2.2555 | 0.26956 | -1.3959 | 1.73 |
| 4931 | *KRTAP19-5* | 3.847 | 0.72204 | 0.60721 | -0.96946 | 1.73 |
| 4932 | *OR6B1* | 4.2375 | 0.67486 | 0.2638 | -0.68303 | 1.73 |
| 4933 | *PRKAG2* | 2.5727 | 1.5542 | 1.049 | -0.3282 | 1.73 |
| 4934 | *CACFD1* | 4.3269 | 1.001 | -0.15212 | -2.5302 | 1.73 |
| 4935 | *FLNB* | 4.1101 | 2.7549 | -1.6893 | -2.1282 | 1.73 |
| 4936 | *HIP1R* | 4.0042 | 1.5834 | -0.4119 | -0.67608 | 1.73 |
| 4937 | *ATG101* | 2.3839 | 1.7118 | 1.079 | 1.0335 | 1.72 |
| 4938 | *ZNF607* | 4.7273 | 0.23285 | 0.21428 | -1.2211 | 1.72 |
| 4939 | *PAMR1* | 2.5601 | 1.5631 | 1.0502 | 0.52584 | 1.72 |
| 4940 | *ATP1A3* | 3.4185 | 0.98936 | 0.76532 | -0.7329 | 1.72 |
| 4941 | *GTF2A2* | 4.303 | 1.0092 | -0.13928 | -1.3245 | 1.72 |
| 4942 | *AGPAT4* | 3.3079 | 1.6362 | 0.22797 | -1.5204 | 1.72 |
| 4943 | *C1QL3* | 3.9727 | 0.64417 | 0.55491 | 0.21274 | 1.72 |
| 4944 | *CLN5* | 3.5588 | 1.2111 | 0.40041 | 0.088149 | 1.72 |
| 4945 | *VSTM2L* | 4.4272 | 1.1822 | -0.44182 | -1.4333 | 1.72 |
| 4946 | *RMCX5-GPRASP2* | 3.1143 | 2.1399 | -0.08676 | -0.41575 | 1.72 |
| 4947 | *LINGO4* | 3.3505 | 2.455 | -0.63834 | -1.3218 | 1.72 |
| 4948 | *CABS1* | 3.1539 | 1.134 | 0.87917 | -1.153 | 1.72 |
| 4949 | *NUDT16L1* | 4.1359 | 1.7143 | -0.68409 | -1.8388 | 1.72 |
| 4950 | *ATP5C1* | 4.1703 | 1.1646 | -0.17208 | -1.4683 | 1.72 |
| 4951 | *MRPL42* | 3.624 | 2.6464 | -1.1096 | -2.4496 | 1.72 |
| 4952 | *NEGR1* | 3.3571 | 0.98637 | 0.8165 | 0.051389 | 1.72 |
| 4953 | *USP17L21* | 2.8752 | 1.9762 | 0.30826 | -0.8604 | 1.72 |
| 4954 | *ZNF283* | 4.9073 | 0.15449 | 0.0973 | -0.87198 | 1.72 |
| 4955 | *CEACAM16* | 2.8548 | 1.5082 | 0.79603 | -1.8856 | 1.72 |
| 4956 | *CTDSP1* | 2.3121 | 1.6641 | 1.1825 | -0.24817 | 1.72 |
| 4957 | *AUTS2* | 6.0977 | 1.2542 | -2.1947 | -2.3101 | 1.72 |
| 4958 | *ZNF81* | 3.0664 | 1.6546 | 0.43613 | -0.05863 | 1.72 |
| 4959 | *KRTAP1-4* | 4.7906 | 0.21127 | 0.15516 | -1.714 | 1.72 |
| 4960 | *HIST1H2AK* | 3.2776 | 1.5818 | 0.29707 | -1.1228 | 1.72 |
| 4961 | *PARD3B* | 2.8389 | 1.8515 | 0.46366 | -1.0005 | 1.72 |

| | | | | | | |
|---|---|---|---|---|---|---|
| 4962 | *RFXANK* | 2.3495 | 1.6989 | 1.1048 | -0.62307 | 1.72 |
| 4963 | *OPCML* | 4.8163 | 0.94144 | -0.60465 | -2.2947 | 1.72 |
| 4964 | *OR1B1* | 5.3281 | -0.05911 | -0.11621 | -1.3613 | 1.72 |
| 4965 | *SLAIN2* | 4.2275 | 0.71562 | 0.20951 | -1.2887 | 1.72 |
| 4966 | *MCPH1* | 1.9285 | 1.855 | 1.368 | -0.25459 | 1.72 |
| 4967 | *SHC4* | 3.3809 | 0.93664 | 0.83301 | -1.0288 | 1.72 |
| 4968 | *HCRT* | 4.2207 | 0.50033 | 0.42915 | -0.48572 | 1.72 |
| 4969 | *TXLNA* | 5.2496 | -0.04134 | -0.05851 | -0.35722 | 1.72 |
| 4970 | *LTBP2* | 3.901 | 0.6905 | 0.55745 | -0.55002 | 1.72 |
| 4971 | *TMCO5A* | 2.5316 | 2.3065 | 0.3095 | -0.06351 | 1.72 |
| 4972 | *ADAMTS5* | 3.732 | 0.75092 | 0.66416 | -1.1943 | 1.72 |
| 4973 | *SFT2D2* | 3.0408 | 2.3073 | -0.20166 | -1.2842 | 1.72 |
| 4974 | *ASCC1* | 3.8952 | 1.7013 | -0.4521 | -0.66569 | 1.71 |
| 4975 | *SRRM4* | 4.2094 | 1.9464 | -1.0125 | -1.3627 | 1.71 |
| 4976 | *IFNA21* | 2.671 | 2.2747 | 0.19757 | -0.68169 | 1.71 |
| 4977 | *MAEA* | 3.053 | 1.7298 | 0.3597 | 0.30022 | 1.71 |
| 4978 | *VWA8* | 4.5506 | 1.6516 | -1.06 | -1.7039 | 1.71 |
| 4979 | *POT1* | 4.1157 | 1.2822 | -0.25708 | -1.234 | 1.71 |
| 4980 | *HNF4G* | 4.3825 | 0.57604 | 0.18138 | -1.4452 | 1.71 |
| 4981 | *PRKCH* | 2.6798 | 2.3153 | 0.14384 | -0.18395 | 1.71 |
| 4982 | *PRKCD* | 3.613 | 0.93194 | 0.59379 | -1.5436 | 1.71 |
| 4983 | *GPR18* | 4.9998 | 0.35756 | -0.2192 | -0.25971 | 1.71 |
| 4984 | *ANP32C* | 2.3943 | 1.6491 | 1.0945 | 0.82914 | 1.71 |
| 4985 | *BMP8B* | 3.371 | 1.7803 | -0.01383 | -0.94121 | 1.71 |
| 4986 | *KCTD19* | 3.9902 | 2.1637 | -1.0172 | -1.942 | 1.71 |
| 4987 | *TP53INP1* | 3.4712 | 1.2869 | 0.37811 | -0.9279 | 1.71 |
| 4988 | *ARHGAP30* | 2.0673 | 1.665 | 1.4033 | 0.095265 | 1.71 |
| 4989 | *C1orf189* | 4.0504 | 1.1529 | -0.0677 | -1.5673 | 1.71 |
| 4990 | *RSL24D1* | 4.63 | 1.0827 | -0.57713 | -2.7406 | 1.71 |
| 4991 | *BET1* | 3.5198 | 2.8224 | -1.2067 | -2.1987 | 1.71 |
| 4992 | *CLYBL* | 3.9477 | 1.7716 | -0.58451 | -1.9036 | 1.71 |
| 4993 | *DNAJB6* | 2.7894 | 2.3983 | -0.05341 | -1.2098 | 1.71 |
| 4994 | *NUDT22* | 3.4945 | 1.4034 | 0.23609 | -0.50961 | 1.71 |
| 4995 | *VAMP7* | 3.6327 | 1.4941 | 0.0069 | -1.4998 | 1.71 |
| 4996 | *UCMA* | 2.4005 | 2.3605 | 0.37192 | 0.30283 | 1.71 |
| 4997 | *DMGDH* | 2.8109 | 2.5169 | -0.19494 | -1.3664 | 1.71 |
| 4998 | *AP4E1* | 4.1009 | 1.7042 | -0.67342 | -1.3533 | 1.71 |
| 4999 | *ANXA4* | 2.305 | 1.9998 | 0.82682 | 0.35788 | 1.71 |
| 5000 | *ZNF614* | 4.8436 | 0.28367 | 0.00396 | -0.66997 | 1.71 |
| 5001 | *TMEM244* | 2.5012 | 1.9432 | 0.68667 | -1.4185 | 1.71 |
| 5002 | *TNFRSF11A* | 2.6142 | 2.1593 | 0.35491 | -2.1193 | 1.71 |
| 5003 | *C7orf76* | 2.6019 | 2.4997 | 0.02656 | -1.9623 | 1.71 |
| 5004 | *TBC1D26* | 3.9029 | 0.72659 | 0.49861 | 0.2579 | 1.71 |
| 5005 | *GPRC5B* | 3.0065 | 2.0377 | 0.08211 | -0.75767 | 1.71 |
| 5006 | *CLIC2* | 3.0221 | 1.826 | 0.2764 | -0.87241 | 1.71 |
| 5007 | *LSM4* | 5.1501 | 0.18668 | -0.21291 | -2.0358 | 1.71 |

| | | | | | | |
|---|---|---|---|---|---|---|
| 5008 | *PPP2R2D* | 2.4152 | 1.5087 | 1.1988 | -0.69787 | 1.71 |
| 5009 | *C19orf57* | 5.0808 | 0.28489 | -0.24376 | -1.0278 | 1.71 |
| 5010 | *TGIF2* | 2.3068 | 1.8658 | 0.94904 | -0.31645 | 1.71 |
| 5011 | *GFPT2* | 3.1665 | 1.555 | 0.39963 | -0.6399 | 1.71 |
| 5012 | *KIF5A* | 5.2194 | 0.54867 | -0.64748 | -2.5081 | 1.71 |
| 5013 | *UBXN2B* | 2.7669 | 1.4713 | 0.88198 | 0.50669 | 1.71 |
| 5014 | *AP5M1* | 5.916 | 0.92767 | -1.7235 | -2.2165 | 1.71 |
| 5015 | *OR2AG2* | 3.5746 | 1.3054 | 0.24015 | -1.0992 | 1.71 |
| 5016 | *MYO3A* | 4.9258 | 0.79371 | -0.59961 | -1.9287 | 1.71 |
| 5017 | *RIMBP2* | 2.9462 | 2.2045 | -0.03161 | -0.49716 | 1.71 |
| 5018 | *RBMXL1* | 3.3238 | 1.2748 | 0.51984 | -0.53554 | 1.71 |
| 5019 | *C2orf71* | 2.6315 | 2.4505 | 0.03565 | -2.3602 | 1.71 |
| 5020 | *GSKIP* | 1.9952 | 1.5905 | 1.5308 | 0.56247 | 1.71 |
| 5021 | *PPP1R1B* | 3.2609 | 1.8935 | -0.03826 | -0.59225 | 1.71 |
| 5022 | *PRTG* | 2.4079 | 1.8499 | 0.85789 | 0.84112 | 1.71 |
| 5023 | *ZBTB7C* | 3.8786 | 1.3313 | -0.09431 | -0.26429 | 1.71 |
| 5024 | *PCDHGA9* | 4.0705 | 0.79987 | 0.24514 | -1.4701 | 1.71 |
| 5025 | *CCZ1* | 2.4619 | 1.4537 | 1.1995 | -0.46064 | 1.71 |
| 5026 | *ADAM12* | 3.1527 | 1.9914 | -0.02915 | -1.5684 | 1.70 |
| 5027 | *SPRY2* | 4.0802 | 1.2691 | -0.23481 | -1.4852 | 1.70 |
| 5028 | *OR1J4* | 2.6288 | 1.6791 | 0.80532 | -2.0465 | 1.70 |
| 5029 | *EXOC5* | 5.7029 | 0.025017 | -0.61555 | -2.7384 | 1.70 |
| 5030 | *APOBEC2* | 3.7228 | 2.6565 | -1.2673 | -1.4317 | 1.70 |
| 5031 | *SUB1* | 4.2387 | 0.80653 | 0.06592 | -0.83243 | 1.70 |
| 5032 | *RASL12* | 2.434 | 2.3328 | 0.34431 | -0.60465 | 1.70 |
| 5033 | *EPN3* | 5.0785 | 1.1417 | -1.1098 | -1.1793 | 1.70 |
| 5034 | *KRT35* | 3.3572 | 1.7662 | -0.01327 | -1.175 | 1.70 |
| 5035 | *ZNF177* | 4.6977 | 0.56594 | -0.15366 | -1.1713 | 1.70 |
| 5036 | *MARC1* | 3.9074 | 1.5853 | -0.38319 | -0.44353 | 1.70 |
| 5037 | *TRERF1* | 3.5526 | 1.4116 | 0.14451 | -0.18353 | 1.70 |
| 5038 | *OR9K2* | 3.6261 | 1.7667 | -0.28427 | -0.44353 | 1.70 |
| 5039 | *PIGS* | 3.0103 | 1.2275 | 0.87036 | -1.9623 | 1.70 |
| 5040 | *SNX10* | 3.995 | 0.63803 | 0.47491 | -1.5651 | 1.70 |
| 5041 | *CDCA7* | 3.4038 | 3.2703 | -1.5663 | -1.6208 | 1.70 |
| 5042 | *TRIM16* | 2.4894 | 1.6013 | 1.0162 | -0.74086 | 1.70 |
| 5043 | *SLC4A4* | 4.4587 | 0.60104 | 0.04714 | -1.403 | 1.70 |
| 5044 | *RAP1A* | 2.9578 | 2.5939 | -0.44499 | -0.69319 | 1.70 |
| 5045 | *ZNF576* | 3.8097 | 2.1345 | -0.83784 | -2.1477 | 1.70 |
| 5046 | *C11orf88* | 2.1018 | 2.028 | 0.97636 | -0.09802 | 1.70 |
| 5047 | *RAET1E* | 2.2456 | 1.4706 | 1.3884 | -1.7655 | 1.70 |
| 5048 | *TLN2* | 4.1487 | 1.0795 | -0.12425 | -1.9358 | 1.70 |
| 5049 | *TNFSF4* | 2.2151 | 1.7877 | 1.1008 | -1.9271 | 1.70 |
| 5050 | *SERPINB9* | 3.3799 | 1.411 | 0.31268 | -0.78398 | 1.70 |
| 5051 | *CLSPN* | 2.3618 | 2.3202 | 0.42143 | -0.52655 | 1.70 |
| 5052 | *KBTBD6* | 2.8753 | 2.5822 | -0.35548 | -0.69155 | 1.70 |
| 5053 | *SCAF8* | 2.7634 | 2.1415 | 0.19702 | -2.0545 | 1.70 |

| | | | | | | |
|---|---|---|---|---|---|---|
| 5054 | *TMED9* | 2.8903 | 1.2102 | 0.99834 | -3.0575 | 1.70 |
| 5055 | *INPP5B* | 5.2341 | 0.88301 | -1.0197 | -1.8508 | 1.70 |
| 5056 | *LOC101929372* | 4.5271 | 1.0109 | -0.44134 | -1.3696 | 1.70 |
| 5057 | *H1FX* | 2.4439 | 1.5161 | 1.1365 | 0.33706 | 1.70 |
| 5058 | *RB1* | 2.0388 | 2.0099 | 1.0469 | -0.92114 | 1.70 |
| 5059 | *RNASE6* | 2.8962 | 1.2001 | 0.99893 | 0.16118 | 1.70 |
| 5060 | *IFNE* | 2.5371 | 2.0736 | 0.484 | -0.23053 | 1.70 |
| 5061 | *ZNF69* | 4.69 | 0.79841 | -0.39416 | -0.53931 | 1.70 |
| 5062 | *TVP23C-CDRT4* | 4.1571 | 0.90838 | 0.02859 | -1.6885 | 1.70 |
| 5063 | *SYCE1L* | 3.4975 | 1.2591 | 0.33736 | -0.6536 | 1.70 |
| 5064 | *IZUMO4* | 2.0452 | 1.5416 | 1.5066 | -0.77114 | 1.70 |
| 5065 | *CCL20* | 4.5672 | 0.85457 | -0.32906 | -0.79035 | 1.70 |
| 5066 | *A1CF* | 2.9512 | 2.3032 | -0.16207 | -1.9163 | 1.70 |
| 5067 | *DMWD* | 4.0128 | 0.84586 | 0.23259 | 0.14819 | 1.70 |
| 5068 | *YBX2* | 4.1267 | 1.1521 | -0.18822 | -1.1977 | 1.70 |
| 5069 | *COX14* | 4.718 | 1.0828 | -0.71041 | -1.6765 | 1.70 |
| 5070 | *CCL27* | 3.272 | 2.6366 | -0.81873 | -3.4039 | 1.70 |
| 5071 | *LMX1A* | 3.9717 | 1.0413 | 0.07633 | -1.165 | 1.70 |
| 5072 | *MOAP1* | 4.9964 | 0.73578 | -0.64334 | -2.3357 | 1.70 |
| 5073 | *C1orf101* | 3.5477 | 3.1471 | -1.6072 | -1.8269 | 1.70 |
| 5074 | *SCGB3A1* | 3.794 | 2.6228 | -1.3292 | -1.8582 | 1.70 |
| 5075 | *PAGR1* | 2.9158 | 2.1823 | -0.01264 | -0.96093 | 1.70 |
| 5076 | *DEFB115* | 3.305 | 1.4552 | 0.32509 | -1.2411 | 1.70 |
| 5077 | *GZMK* | 3.5727 | 1.4661 | 0.04435 | -0.44236 | 1.69 |
| 5078 | *IFT172* | 4.1214 | 0.79121 | 0.17014 | -1.0485 | 1.69 |
| 5079 | *CLASRP* | 4.1158 | 1.223 | -0.25618 | -1.3892 | 1.69 |
| 5080 | *MYH3* | 3.1659 | 1.4715 | 0.44515 | 0.32083 | 1.69 |
| 5081 | *NME8* | 2.4718 | 1.3338 | 1.276 | -1.384 | 1.69 |
| 5082 | *CD180* | 3.587 | 1.8213 | -0.32699 | -0.51799 | 1.69 |
| 5083 | *C1QTNF9B* | 2.5888 | 1.7175 | 0.77289 | 0.39232 | 1.69 |
| 5084 | *TMEM74* | 3.3264 | 1.999 | -0.2469 | -1.0798 | 1.69 |
| 5085 | *HK2* | 3.4621 | 0.91486 | 0.7013 | -1.3057 | 1.69 |
| 5086 | *TMEM129* | 3.272 | 1.672 | 0.1342 | -0.15422 | 1.69 |
| 5087 | *SMPD1* | 3.1662 | 1.7429 | 0.16888 | -2.2483 | 1.69 |
| 5088 | *JADE3* | 3.9977 | 2.2676 | -1.1897 | -1.2739 | 1.69 |
| 5089 | *RIPPLY3* | 4.0304 | 0.80889 | 0.23624 | 0.13675 | 1.69 |
| 5090 | *CCDC148* | 3.8082 | 0.70236 | 0.5635 | 0.41384 | 1.69 |
| 5091 | *CXCL13* | 3.8378 | 1.9883 | -0.75338 | -1.7757 | 1.69 |
| 5092 | *GSTT2B* | 3.4302 | 1.4502 | 0.19155 | -1.1923 | 1.69 |
| 5093 | *MTG2* | 3.6695 | 1.768 | -0.36566 | -0.98769 | 1.69 |
| 5094 | *DMRT2* | 4.9425 | 0.69628 | -0.57042 | -1.2699 | 1.69 |
| 5095 | *ADPGK* | 4.2151 | 0.65762 | 0.19522 | -1.5588 | 1.69 |
| 5096 | *DNAH2* | 4.7075 | 0.19443 | 0.16545 | -0.19735 | 1.69 |
| 5097 | *FAM86B2* | 3.5713 | 0.79316 | 0.7024 | -2.2528 | 1.69 |
| 5098 | *ZDHHC14* | 3.5058 | 3.0255 | -1.4652 | -2.4784 | 1.69 |
| 5099 | *USP1* | 3.3158 | 1.0494 | 0.7007 | -3.4683 | 1.69 |

| | | | | | | |
|---|---|---|---|---|---|---|
| 5100 | *PIWIL2* | 3.4987 | 1.7776 | -0.2106 | -2.1926 | 1.69 |
| 5101 | *NKX1-2* | 4.3921 | 1.5404 | -0.86725 | -1.7865 | 1.69 |
| 5102 | *HNRNPLL* | 2.7215 | 1.6637 | 0.6799 | -0.11784 | 1.69 |
| 5103 | *API5* | 3.5535 | 1.4015 | 0.10969 | -0.88492 | 1.69 |
| 5104 | *GOLGB1* | 4.4959 | 0.31183 | 0.25687 | -0.77778 | 1.69 |
| 5105 | *RIPK1* | 2.2154 | 1.5922 | 1.2568 | 0.91805 | 1.69 |
| 5106 | *TRNAU1AP* | 2.7899 | 1.8527 | 0.42156 | -0.88898 | 1.69 |
| 5107 | *DCAF7* | 2.3355 | 2.0709 | 0.65691 | 0.038126 | 1.69 |
| 5108 | *MAGEC3* | 3.1525 | 2.2777 | -0.36761 | -0.60065 | 1.69 |
| 5109 | *CHMP4C* | 3.4685 | 1.6395 | -0.04542 | -1.3565 | 1.69 |
| 5110 | *PFN4* | 2.4943 | 1.8352 | 0.73229 | -0.12228 | 1.69 |
| 5111 | *IFT22* | 3.6378 | 2.0218 | -0.59816 | -0.73708 | 1.69 |
| 5112 | *TCF25* | 3.9559 | 1.3533 | -0.24856 | -1.6417 | 1.69 |
| 5113 | *STAT1* | 3.3366 | 1.3772 | 0.34675 | -1.855 | 1.69 |
| 5114 | *TTL* | 2.9501 | 2.5946 | -0.4847 | -1.8087 | 1.69 |
| 5115 | *TLX2* | 6.0123 | -0.42708 | -0.52552 | -2.3615 | 1.69 |
| 5116 | *USP24* | 2.7655 | 2.7368 | -0.44353 | -0.58534 | 1.69 |
| 5117 | *MEIOB* | 3.4165 | 1.9216 | -0.28052 | -0.71753 | 1.69 |
| 5118 | *MAN1A2* | 2.8354 | 1.341 | 0.8804 | -1.4529 | 1.69 |
| 5119 | *EMC1* | 4.1331 | 1.5729 | -0.64943 | -1.6791 | 1.69 |
| 5120 | *CYB5B* | 4.3089 | 1.553 | -0.80545 | -1.8136 | 1.69 |
| 5121 | *FAM200B* | 4.4017 | 1.6228 | -0.96823 | -2.1496 | 1.69 |
| 5122 | *ADAMTS9* | 3.2312 | 1.1205 | 0.70325 | -0.43082 | 1.68 |
| 5123 | *RNF207* | 3.5404 | 1.1336 | 0.38026 | -1.3525 | 1.68 |
| 5124 | *MAGEB17* | 4.2044 | 0.72679 | 0.12276 | -0.46024 | 1.68 |
| 5125 | *UQCC1* | 4.0124 | 1.0299 | 0.01138 | -0.9999 | 1.68 |
| 5126 | *ARHGEF16* | 4.5853 | 0.91135 | -0.44353 | -0.96788 | 1.68 |
| 5127 | *LOC283710* | 3.8351 | 1.3301 | -0.11212 | -2.4596 | 1.68 |
| 5128 | *ZNF597* | 3.4725 | 1.9481 | -0.36761 | -1.3759 | 1.68 |
| 5129 | *CDC37L1* | 3.1597 | 1.2844 | 0.60858 | -0.82897 | 1.68 |
| 5130 | *FUOM* | 4.6996 | 1.4406 | -1.0886 | -1.695 | 1.68 |
| 5131 | *TANC2* | 3.8004 | 1.4723 | -0.22131 | -0.9619 | 1.68 |
| 5132 | *XXYLT1* | 3.9501 | 1.1378 | -0.03675 | -1.6174 | 1.68 |
| 5133 | *GPR137C* | 2.8651 | 2.0028 | 0.1832 | -0.01401 | 1.68 |
| 5134 | *CCZ1B* | 3.4823 | 2.2552 | -0.68693 | -1.6182 | 1.68 |
| 5135 | *GPD1* | 4.3455 | 0.60948 | 0.09518 | -2.1407 | 1.68 |
| 5136 | *FAM78B* | 4.2587 | 0.43887 | 0.35137 | -0.87332 | 1.68 |
| 5137 | *C10orf54* | 4.7004 | 1.5382 | -1.1898 | -2.4966 | 1.68 |
| 5138 | *GAGE10* | 3.9399 | 3.2506 | -2.1419 | -2.7546 | 1.68 |
| 5139 | *GRIK5* | 2.5549 | 2.0581 | 0.43394 | -0.67536 | 1.68 |
| 5140 | *RASSF6* | 3.4409 | 1.7703 | -0.16455 | -0.93287 | 1.68 |
| 5141 | *STS* | 2.811 | 2.6014 | -0.36685 | -0.44353 | 1.68 |
| 5142 | *TNFRSF10B* | 3.7649 | 0.64864 | 0.63157 | 0.004564 | 1.68 |
| 5143 | *FZD1* | 3.0073 | 1.7916 | 0.24574 | -0.87062 | 1.68 |
| 5144 | *UCK1* | 4.0179 | 2.4508 | -1.4249 | -2.5707 | 1.68 |
| 5145 | *MBTPS2* | 3.3024 | 2.871 | -1.1303 | -2.3017 | 1.68 |

| | | | | | | |
|---|---|---|---|---|---|---|
| 5146 | *C14orf28* | 3.8368 | 2.3631 | -1.1573 | -2.1108 | 1.68 |
| 5147 | *TCEA2* | 4.717 | 1.0353 | -0.71016 | -0.8462 | 1.68 |
| 5148 | *TSKU* | 3.921 | 1.4539 | -0.33312 | -2.1085 | 1.68 |
| 5149 | *OR5K1* | 4.7325 | 1.8218 | -1.5128 | -3.1665 | 1.68 |
| 5150 | *DTL* | 4.4519 | 1.047 | -0.4578 | -0.51799 | 1.68 |
| 5151 | *HMX3* | 3.6203 | 1.9616 | -0.54226 | -1.1221 | 1.68 |
| 5152 | *FAM157A* | 4.6072 | 2.057 | -1.6246 | -1.849 | 1.68 |
| 5153 | *PEAK1* | 4.0339 | 1.6699 | -0.66468 | -1.3478 | 1.68 |
| 5154 | *OR51B2* | 2.9981 | 1.8661 | 0.17482 | -2.458 | 1.68 |
| 5155 | *ATF7IP* | 3.2274 | 1.182 | 0.62953 | 0.58609 | 1.68 |
| 5156 | *CRK* | 3.6053 | 1.8046 | -0.37125 | -1.7001 | 1.68 |
| 5157 | *LSAMP* | 3.6513 | 1.5307 | -0.1438 | -0.27669 | 1.68 |
| 5158 | *LRRTM2* | 2.1133 | 1.8428 | 1.0808 | -0.86572 | 1.68 |
| 5159 | *TRIM59* | 2.6345 | 2.2659 | 0.13621 | -0.67019 | 1.68 |
| 5160 | *ZNF776* | 2.8055 | 2.0484 | 0.1821 | 0.16107 | 1.68 |
| 5161 | *ZSCAN1* | 3.9978 | 1.446 | -0.40785 | -1.6176 | 1.68 |
| 5162 | *ACAA1* | 2.0114 | 1.7505 | 1.273 | -0.74965 | 1.68 |
| 5163 | *TEN1* | 4.3188 | 0.52822 | 0.18787 | -0.96529 | 1.68 |
| 5164 | *SIMC1* | 4.4569 | 0.93319 | -0.35548 | -1.5687 | 1.68 |
| 5165 | *PCNT* | 3.2302 | 1.7572 | 0.04687 | -1.7299 | 1.68 |
| 5166 | *LILRB4* | 3.9318 | 1.1719 | -0.07001 | -0.77105 | 1.68 |
| 5167 | *MELK* | 3.5336 | 2.1321 | -0.63214 | -1.4089 | 1.68 |
| 5168 | *LRRC59* | 2.4039 | 1.9681 | 0.6608 | -0.64556 | 1.68 |
| 5169 | *GUCY2F* | 4.9749 | 0.84716 | -0.78956 | -1.2241 | 1.68 |
| 5170 | *COMMD1* | 4.9113 | 1.0473 | -0.9263 | -1.257 | 1.68 |
| 5171 | *VAC14* | 4.9507 | 0.051371 | 0.02994 | -1.2161 | 1.68 |
| 5172 | *SYNPO2* | 4.026 | 0.94125 | 0.06474 | -2.1732 | 1.68 |
| 5173 | *PTGIS* | 3.489 | 1.2781 | 0.26487 | -2.0187 | 1.68 |
| 5174 | *GNPNAT1* | 4.6884 | 0.35631 | -0.01275 | -0.44353 | 1.68 |
| 5175 | *ADRBK1* | 2.8207 | 1.8745 | 0.33664 | -1.5383 | 1.68 |
| 5176 | *STX10* | 4.2687 | 1.5778 | -0.81509 | -1.2616 | 1.68 |
| 5177 | *PTGES3L-AARSD1* | 3.8953 | 1.5312 | -0.39627 | -0.62423 | 1.68 |
| 5178 | *ZMYM2* | 4.7386 | 1.6458 | -1.3542 | -1.6097 | 1.68 |
| 5179 | *MICAL2* | 4.0781 | 0.76859 | 0.18254 | 0.13956 | 1.68 |
| 5180 | *SLC51A* | 2.7172 | 1.496 | 0.81585 | -0.36412 | 1.68 |
| 5181 | *RNF150* | 4.2178 | 0.63881 | 0.17242 | -0.40291 | 1.68 |
| 5182 | *AKAP9* | 2.4357 | 2.1155 | 0.47667 | -0.91134 | 1.68 |
| 5183 | *SIGMAR1* | 4.2907 | 0.49106 | 0.2448 | -0.56423 | 1.68 |
| 5184 | *EPN1* | 4.3182 | 0.82008 | -0.11216 | -0.83856 | 1.68 |
| 5185 | *TIMM17B* | 3.1381 | 1.1792 | 0.70773 | -1.1401 | 1.68 |
| 5186 | *MATN1* | 4.0934 | 1.2666 | -0.33521 | -0.40547 | 1.67 |
| 5187 | *OSBPL1A* | 3.7483 | 1.2621 | 0.01424 | -1.1084 | 1.67 |
| 5188 | *CDK20* | 2.8858 | 2.3084 | -0.16971 | -2.1187 | 1.67 |
| 5189 | *GATC* | 4.8897 | 0.28826 | -0.15424 | -1.5401 | 1.67 |
| 5190 | *STK31* | 2.4821 | 1.387 | 1.1544 | -0.7532 | 1.67 |
| 5191 | *MAGEB16* | 3.2839 | 1.1873 | 0.5522 | 0.005561 | 1.67 |

| | | | | | | |
|---|---|---|---|---|---|---|
| 5192 | *NAA16* | 3.6569 | 0.95273 | 0.41353 | -0.65512 | 1.67 |
| 5193 | *PHOX2A* | 4.0306 | 0.5347 | 0.45732 | -0.45569 | 1.67 |
| 5194 | *ACO2* | 3.0794 | 2.0665 | -0.12338 | -0.44353 | 1.67 |
| 5195 | *ADH7* | 3.1933 | 3.0748 | -1.2457 | -1.4544 | 1.67 |
| 5196 | *CD226* | 5.2976 | 0.1673 | -0.44353 | -0.69377 | 1.67 |
| 5197 | *KRT74* | 3.9145 | 1.912 | -0.80558 | -1.9844 | 1.67 |
| 5198 | *CELA3B* | 4.5766 | 0.24308 | 0.20113 | -1.1761 | 1.67 |
| 5199 | *TTPAL* | 2.6583 | 2.0493 | 0.31087 | -1.8227 | 1.67 |
| 5200 | *LRRC70* | 3.5638 | 1.5819 | -0.12876 | -1.0905 | 1.67 |
| 5201 | *SLC41A1* | 3.1297 | 1.4378 | 0.44901 | -1.7646 | 1.67 |
| 5202 | *LMNTD2* | 3.9912 | 1.2119 | -0.18718 | -2.2075 | 1.67 |
| 5203 | *AKAP5* | 3.7759 | 1.1489 | 0.09105 | -2.0055 | 1.67 |
| 5204 | *IQCA1* | 2.6109 | 2.1112 | 0.29227 | -1.2827 | 1.67 |
| 5205 | *E2F6* | 3.8363 | 1.6215 | -0.44353 | -1.4841 | 1.67 |
| 5206 | *R3HCC1* | 3.7149 | 0.88074 | 0.41609 | -2.3517 | 1.67 |
| 5207 | *ZNF133* | 3.3404 | 1.3279 | 0.34162 | -0.76728 | 1.67 |
| 5208 | *NPY* | 5.7331 | 0.39934 | -1.1233 | -2.2075 | 1.67 |
| 5209 | *PIP4K2A* | 5.524 | 0.44134 | -0.95645 | -1.095 | 1.67 |
| 5210 | *OBSCN* | 3.8392 | 0.87812 | 0.29114 | -1.462 | 1.67 |
| 5211 | *CMTM7* | 3.6918 | 0.68002 | 0.63621 | -0.91192 | 1.67 |
| 5212 | *SCGB3A2* | 3.8282 | 1.7878 | -0.60806 | -1.813 | 1.67 |
| 5213 | *C1QTNF3* | 4.7841 | 0.55579 | -0.33198 | -1.3385 | 1.67 |
| 5214 | *C9orf24* | 4.0142 | 1.7205 | -0.72728 | -1.9358 | 1.67 |
| 5215 | *STON2* | 2.9644 | 1.963 | 0.07988 | -0.66841 | 1.67 |
| 5216 | *FBF1* | 3.029 | 2.603 | -0.62508 | -1.7372 | 1.67 |
| 5217 | *MTHFD2L* | 2.4202 | 1.4729 | 1.1136 | -1.2917 | 1.67 |
| 5218 | *HMOX1* | 2.7619 | 2.4663 | -0.22244 | -1.0291 | 1.67 |
| 5219 | *B3GALT1* | 2.0592 | 1.9215 | 1.0246 | -1.8348 | 1.67 |
| 5220 | *TRIM15* | 3.1609 | 1.7666 | 0.07695 | -0.93116 | 1.67 |
| 5221 | *ZNF831* | 3.1725 | 1.1036 | 0.72814 | -0.46153 | 1.67 |
| 5222 | *PCLO* | 4.1644 | 0.48305 | 0.35566 | -0.36821 | 1.67 |
| 5223 | *REV1* | 4.9287 | 0.17122 | -0.09723 | -0.8756 | 1.67 |
| 5224 | *PROZ* | 4.1071 | 0.83961 | 0.05569 | 0.046166 | 1.67 |
| 5225 | *COPS7A* | 1.9855 | 1.6403 | 1.3746 | 0.85562 | 1.67 |
| 5226 | *FCER2* | 5.396 | 0.95208 | -1.3492 | -1.5524 | 1.67 |
| 5227 | *OTOS* | 3.4633 | 1.8101 | -0.27501 | -0.88278 | 1.67 |
| 5228 | *ENSA* | 4.6138 | 1.1484 | -0.76389 | -2.2879 | 1.67 |
| 5229 | *TM4SF19* | 2.8637 | 2.089 | 0.04515 | -1.6748 | 1.67 |
| 5230 | *CCDC40* | 4.1297 | 1.1559 | -0.28834 | -0.68298 | 1.67 |
| 5231 | *GUCY1A3* | 4.7796 | 0.12878 | 0.08854 | -0.90382 | 1.67 |
| 5232 | *VCPIP1* | 4.9958 | 0.63861 | -0.63753 | -1.9168 | 1.67 |
| 5233 | *KIAA1468* | 2.3339 | 2.2961 | 0.36684 | -1.4681 | 1.67 |
| 5234 | *UGT1A7* | 3.9024 | 0.65784 | 0.43602 | 0.14156 | 1.67 |
| 5235 | *ERMARD* | 2.2264 | 2.0881 | 0.68176 | 0.45693 | 1.67 |
| 5236 | *ATP13A3* | 4.4491 | 0.34753 | 0.19879 | -0.9472 | 1.67 |
| 5237 | *PTTG2* | 4.1238 | 0.51625 | 0.35534 | -0.00827 | 1.67 |

| | | | | | | |
|---|---|---|---|---|---|---|
| 5238 | *FBXO33* | 3.0005 | 2.3099 | -0.31521 | -0.98834 | 1.67 |
| 5239 | *ITCH* | 5.3942 | 0.25896 | -0.6603 | -2.0452 | 1.66 |
| 5240 | *WDR44* | 4.961 | 0.19103 | -0.15992 | -1.6836 | 1.66 |
| 5241 | *ERCC8* | 3.3632 | 0.92003 | 0.70849 | -0.88775 | 1.66 |
| 5242 | *IL17A* | 2.7913 | 2.3056 | -0.10524 | -0.15965 | 1.66 |
| 5243 | *KCNA10* | 5.6171 | -0.18246 | -0.44353 | -1.6746 | 1.66 |
| 5244 | *NCAPH* | 3.1212 | 2.3009 | -0.43279 | -1.1496 | 1.66 |
| 5245 | *SCAMP1* | 2.1751 | 1.6498 | 1.1617 | -1.8102 | 1.66 |
| 5246 | *SEMA3D* | 4.4047 | 0.31635 | 0.26546 | -2.5006 | 1.66 |
| 5247 | *PPP1R14C* | 3.8992 | 1.2001 | -0.11454 | -1.7606 | 1.66 |
| 5248 | *C1orf137* | 4.4323 | 0.39367 | 0.15744 | -0.07692 | 1.66 |
| 5249 | *C10orf126* | 3.7726 | 1.4598 | -0.24951 | -1.191 | 1.66 |
| 5250 | *NSFL1C* | 4.5107 | 1.1005 | -0.62852 | -1.2301 | 1.66 |
| 5251 | *ARF4* | 4.4199 | 1.6091 | -1.0482 | -1.6007 | 1.66 |
| 5252 | *DHX34* | 4.4846 | 0.46648 | 0.02965 | -0.30611 | 1.66 |
| 5253 | *SMURF2* | 2.9014 | 1.1594 | 0.91961 | -1.1223 | 1.66 |
| 5254 | *CFAP53* | 2.9941 | 1.2979 | 0.68831 | -0.30609 | 1.66 |
| 5255 | *MPEG1* | 2.5632 | 2.4409 | -0.02429 | -1.1964 | 1.66 |
| 5256 | *ARAP2* | 2.8057 | 2.7535 | -0.58054 | -1.6726 | 1.66 |
| 5257 | *C8G* | 3.1061 | 1.6905 | 0.18187 | -2.0989 | 1.66 |
| 5258 | *GSTO1* | 3.4564 | 0.86686 | 0.65506 | -1.0289 | 1.66 |
| 5259 | *KCNA4* | 2.9786 | 2.3709 | -0.37121 | -0.51447 | 1.66 |
| 5260 | *ATXN7L3* | 3.5119 | 1.8191 | -0.35315 | -2.9338 | 1.66 |
| 5261 | *RNASEH1* | 5.1645 | 0.63502 | -0.82337 | -3.9666 | 1.66 |
| 5262 | *ETV1* | 2.6222 | 1.2731 | 1.0805 | -0.92916 | 1.66 |
| 5263 | *CHRNE* | 4.1093 | 1.8827 | -1.0168 | -2.1077 | 1.66 |
| 5264 | *DDX25* | 2.5459 | 1.3594 | 1.0698 | -0.34148 | 1.66 |
| 5265 | *SPARC* | 3.5335 | 2.6091 | -1.1676 | -1.9269 | 1.66 |
| 5266 | *GNB4* | 2.5958 | 1.9039 | 0.47493 | -0.45505 | 1.66 |
| 5267 | *FOXD4L5* | 3.812 | 1.7978 | -0.63551 | -1.5352 | 1.66 |
| 5268 | *ELP2* | 2.0082 | 1.6476 | 1.3172 | 1.2864 | 1.66 |
| 5269 | *ZNF426* | 4.2987 | 1.6583 | -0.98421 | -0.99079 | 1.66 |
| 5270 | *VSTM4* | 4.2666 | 0.85396 | -0.14828 | -0.36914 | 1.66 |
| 5271 | *FGF20* | 3.1089 | 2.7999 | -0.93726 | -1.1328 | 1.66 |
| 5272 | *RSU1* | 3.6832 | 1.9816 | -0.69407 | -1.2097 | 1.66 |
| 5273 | *PRR23A* | 2.7696 | 2.4085 | -0.20742 | -0.35716 | 1.66 |
| 5274 | *NDUFB10* | 3.9387 | 1.8035 | -0.77155 | -1.4959 | 1.66 |
| 5275 | *SLC51B* | 4.2768 | 0.79913 | -0.10561 | -1.761 | 1.66 |
| 5276 | *C18orf21* | 3.7872 | 0.74817 | 0.43386 | -0.39186 | 1.66 |
| 5277 | *NSUN2* | 3.2592 | 0.98989 | 0.71905 | -0.51219 | 1.66 |
| 5278 | *RIBC2* | 2.119 | 1.5876 | 1.2615 | -2.1984 | 1.66 |
| 5279 | *SLC45A2* | 1.6835 | 1.6609 | 1.6229 | 1.0627 | 1.66 |
| 5280 | *CCDC63* | 3.2944 | 2.5437 | -0.87128 | -2.1859 | 1.66 |
| 5281 | *PKIB* | 5.8574 | 0.57373 | -1.4644 | -2.1177 | 1.66 |
| 5282 | *ZNF546* | 3.2097 | 1.1631 | 0.59297 | -0.15649 | 1.66 |
| 5283 | *RLN2* | 3.2247 | 1.3698 | 0.37111 | -1.297 | 1.66 |

| | | | | | | |
|---|---|---|---|---|---|---|
| 5284 | *SLC5A3* | 2.9406 | 1.6664 | 0.35858 | -2.1077 | 1.66 |
| 5285 | *IRAK3* | 3.2328 | 0.8924 | 0.83963 | -0.56687 | 1.65 |
| 5286 | *LRRC42* | 3.7386 | 0.68772 | 0.53731 | -0.10031 | 1.65 |
| 5287 | *MAP3K15* | 2.149 | 1.7853 | 1.0292 | 0.98233 | 1.65 |
| 5288 | *PBLD* | 3.0369 | 1.4486 | 0.47742 | -0.21739 | 1.65 |
| 5289 | *ENTPD3* | 4.4432 | 0.44395 | 0.07554 | -1.191 | 1.65 |
| 5290 | *TMEM59L* | 2.8488 | 1.5835 | 0.53031 | -1.4442 | 1.65 |
| 5291 | *ZSWIM3* | 3.5936 | 2.5883 | -1.2193 | -1.7001 | 1.65 |
| 5292 | *MIS12* | 3.8841 | 0.76704 | 0.31087 | -1.3253 | 1.65 |
| 5293 | *FAM174B* | 5.5466 | 0.92726 | -1.5128 | -2.7042 | 1.65 |
| 5294 | *ITGB4* | 2.4709 | 1.9734 | 0.51667 | 0.18682 | 1.65 |
| 5295 | *SPRY4* | 4.8184 | 0.11254 | 0.02998 | -1.3416 | 1.65 |
| 5296 | *ZCCHC10* | 2.0302 | 1.937 | 0.99269 | -0.31475 | 1.65 |
| 5297 | *SORL1* | 4.1116 | 1.1616 | -0.31413 | -0.56634 | 1.65 |
| 5298 | *SOX5* | 2.6973 | 2.6642 | -0.40504 | -2.508 | 1.65 |
| 5299 | *TPO* | 2.7133 | 2.1744 | 0.06875 | -0.86412 | 1.65 |
| 5300 | *FAM166B* | 2.9454 | 1.7412 | 0.26972 | -0.08851 | 1.65 |
| 5301 | *VTA1* | 4.2616 | 1.6445 | -0.9512 | -1.6913 | 1.65 |
| 5302 | *DDIT4* | 2.0863 | 1.5048 | 1.3632 | 1.0835 | 1.65 |
| 5303 | *ZNF326* | 2.0265 | 1.9914 | 0.93595 | -0.42232 | 1.65 |
| 5304 | *TRIB2* | 4.6364 | 1.3947 | -1.078 | -1.6469 | 1.65 |
| 5305 | *TMEM139* | 3.7342 | 1.2038 | 0.01376 | -0.93329 | 1.65 |
| 5306 | *GPS2* | 4.2441 | 1.8607 | -1.154 | -2.1893 | 1.65 |
| 5307 | *GOT1L1* | 4.2572 | 1.2196 | -0.52601 | -0.60659 | 1.65 |
| 5308 | *CTTNBP2NL* | 2.0747 | 1.955 | 0.91974 | -0.36246 | 1.65 |
| 5309 | *HAGH* | 5.273 | 0.043791 | -0.36761 | -1.6456 | 1.65 |
| 5310 | *TSPAN9* | 3.4875 | 0.87762 | 0.58395 | -0.98031 | 1.65 |
| 5311 | *CDC14B* | 2.9135 | 2.8912 | -0.85739 | -1.2052 | 1.65 |
| 5312 | *PPP1R14B* | 2.9978 | 2.0064 | -0.05702 | -0.86479 | 1.65 |
| 5313 | *RAB43* | 2.674 | 2.5549 | -0.28254 | -1.4116 | 1.65 |
| 5314 | *SLC7A3* | 4.478 | 0.67343 | -0.20552 | -0.94085 | 1.65 |
| 5315 | *LARP6* | 2.8233 | 2.0753 | 0.04637 | -0.83566 | 1.65 |
| 5316 | *SYT16* | 3.241 | 2.4813 | -0.77877 | -0.80061 | 1.65 |
| 5317 | *C7orf71* | 3.6942 | 2.4463 | -1.1973 | -2.0452 | 1.65 |
| 5318 | *TCOF1* | 3.6968 | 1.9246 | -0.67865 | -0.91134 | 1.65 |
| 5319 | *INO80C* | 3.1721 | 1.667 | 0.10289 | -0.81533 | 1.65 |
| 5320 | *BLCAP* | 4.4951 | 1.7033 | -1.2573 | -1.7441 | 1.65 |
| 5321 | *LENEP* | 5.1255 | 1.1508 | -1.3358 | -2.2955 | 1.65 |
| 5322 | *MVB12A* | 2.9686 | 1.6444 | 0.32647 | -1.9372 | 1.65 |
| 5323 | *KRTAP21-3* | 1.7295 | 1.6688 | 1.5403 | -0.18225 | 1.65 |
| 5324 | *SLC39A7* | 4.2262 | 1.46 | -0.74887 | -2.0827 | 1.65 |
| 5325 | *SNX8* | 4.4767 | 1.8441 | -1.3835 | -2.252 | 1.65 |
| 5326 | *TFPT* | 2.7553 | 2.3521 | -0.1715 | -1.0311 | 1.65 |
| 5327 | *CSPG5* | 3.9372 | 2.487 | -1.4893 | -2.1984 | 1.64 |
| 5328 | *CFHR4* | 3.8564 | 1.0393 | 0.03729 | -0.40191 | 1.64 |
| 5329 | *FAM26E* | 3.8505 | 1.6921 | -0.60961 | -1.5719 | 1.64 |

| | | | | | | |
|---|---|---|---|---|---|---|
| 5330 | *C4orf26* | 2.2771 | 1.4717 | 1.1825 | -0.77155 | 1.64 |
| 5331 | *TMEM259* | 2.9394 | 2.3247 | -0.3349 | -0.58479 | 1.64 |
| 5332 | *HMX1* | 3.9067 | 1.196 | -0.17389 | -2.9338 | 1.64 |
| 5333 | *TLDC2* | 3.2337 | 1.6138 | 0.08083 | -0.27857 | 1.64 |
| 5334 | *DISP1* | 3.1101 | 1.945 | -0.12681 | -2.7436 | 1.64 |
| 5335 | *CCDC23* | 4.1263 | 0.55475 | 0.24607 | 0.01812 | 1.64 |
| 5336 | *FMO1* | 5.0472 | 0.32726 | -0.44742 | -1.1566 | 1.64 |
| 5337 | *C3orf52* | 4.4082 | 1.0774 | -0.55927 | -1.0288 | 1.64 |
| 5338 | *PRDM4* | 3.2031 | 1.7378 | -0.01518 | -1.3507 | 1.64 |
| 5339 | *ANO5* | 2.312 | 1.3671 | 1.2453 | 0.35682 | 1.64 |
| 5340 | *PDCD2L* | 3.7101 | 2.0041 | -0.79035 | -1.27 | 1.64 |
| 5341 | *HRAS* | 3.3911 | 2.442 | -0.91045 | -1.2414 | 1.64 |
| 5342 | *FBXO38* | 4.0734 | 0.72361 | 0.1256 | -0.2624 | 1.64 |
| 5343 | *SLC5A10* | 5.3579 | 0.076946 | -0.51363 | -0.94341 | 1.64 |
| 5344 | *PLXND1* | 3.5004 | 2.5158 | -1.095 | -1.8714 | 1.64 |
| 5345 | *KLK2* | 2.7557 | 1.228 | 0.93657 | -1.317 | 1.64 |
| 5346 | *IRGC* | 3.8845 | 0.97587 | 0.05929 | -0.30061 | 1.64 |
| 5347 | *PCDHGA1* | 2.8369 | 1.7162 | 0.36617 | -1.6396 | 1.64 |
| 5348 | *HSD3B7* | 3.9765 | 0.98664 | -0.04543 | -2.1217 | 1.64 |
| 5349 | *KRTAP1-1* | 4.7279 | 0.61801 | -0.42859 | -2.5934 | 1.64 |
| 5350 | *AGPAT9* | 2.7444 | 2.0407 | 0.132 | -0.48821 | 1.64 |
| 5351 | *ITIH2* | 5.2071 | 1.0551 | -1.3457 | -1.8157 | 1.64 |
| 5352 | *PAPOLA* | 4.1424 | 0.52157 | 0.25112 | -0.28391 | 1.64 |
| 5353 | *BCO1* | 2.7025 | 1.2781 | 0.93423 | -2.6705 | 1.64 |
| 5354 | *PRRG3* | 3.4081 | 1.0351 | 0.47083 | 0.41023 | 1.64 |
| 5355 | *HOXB7* | 2.4973 | 1.2478 | 1.1677 | -1.4894 | 1.64 |
| 5356 | *STARD5* | 2.9609 | 1.0041 | 0.94643 | -0.12469 | 1.64 |
| 5357 | *AXIN2* | 2.8493 | 2.5612 | -0.49984 | -1.2996 | 1.64 |
| 5358 | *EFCAB5* | 3.6821 | 0.75557 | 0.47254 | -0.07935 | 1.64 |
| 5359 | *SCN9A* | 3.8394 | 0.91211 | 0.15829 | -0.45786 | 1.64 |
| 5360 | *KLRD1* | 2.4307 | 1.8422 | 0.63621 | 0.10078 | 1.64 |
| 5361 | *FAM204A* | 4.0816 | 1.0148 | -0.18762 | -1.0919 | 1.64 |
| 5362 | *CSPP1* | 3.31 | 1.3493 | 0.24928 | -1.1145 | 1.64 |
| 5363 | *PANK1* | 2.7577 | 1.62 | 0.5297 | -1.8272 | 1.64 |
| 5364 | *CNIH3* | 4.1744 | 0.38561 | 0.34697 | -0.2301 | 1.64 |
| 5365 | *FAM120AOS* | 4.2287 | 0.52879 | 0.14903 | -0.45185 | 1.64 |
| 5366 | *GOLGA8M* | 4.6052 | 1.3526 | -1.052 | -1.8154 | 1.64 |
| 5367 | *SPON1* | 4.6116 | 0.45984 | -0.16614 | -1.8842 | 1.64 |
| 5368 | *SHROOM3* | 3.3223 | 1.9645 | -0.3817 | -1.3264 | 1.64 |
| 5369 | *TACR3* | 3.1676 | 1.7584 | -0.02092 | -0.61836 | 1.64 |
| 5370 | *BSDC1* | 4.7049 | 1.5189 | -1.3189 | -1.6343 | 1.63 |
| 5371 | *HOMER3* | 3.6391 | 1.234 | 0.03166 | -1.4233 | 1.63 |
| 5372 | *TXNDC9* | 4.1306 | 1.193 | -0.4191 | -0.63584 | 1.63 |
| 5373 | *STX8* | 4.5884 | 0.35623 | -0.04036 | -1.4191 | 1.63 |
| 5374 | *TMIGD1* | 2.3058 | 2.0617 | 0.5365 | -1.3836 | 1.63 |
| 5375 | *SLC35B2* | 3.6561 | 0.66185 | 0.58535 | -1.4388 | 1.63 |

| | | | | | | |
|---|---|---|---|---|---|---|
| 5376 | *FAM63A* | 4.8027 | 0.22302 | -0.12278 | -2.1312 | 1.63 |
| 5377 | *LMO2* | 3.4937 | 2.1888 | -0.77963 | -0.80876 | 1.63 |
| 5378 | *C12orf4* | 2.6474 | 1.2773 | 0.97774 | -1.6335 | 1.63 |
| 5379 | *RBBP7* | 2.5208 | 1.6616 | 0.71939 | -0.35441 | 1.63 |
| 5380 | *MAGEA9B* | 3.1857 | 1.2047 | 0.51117 | -0.48884 | 1.63 |
| 5381 | *SFR1* | 2.6034 | 1.67 | 0.62805 | -1.3507 | 1.63 |
| 5382 | *GP1BB* | 4.7946 | 0.46164 | -0.35735 | -2.8702 | 1.63 |
| 5383 | *ELAVL3* | 4.0147 | 1.0977 | -0.21405 | -0.3364 | 1.63 |
| 5384 | *XKR7* | 4.7488 | 0.41393 | -0.26468 | -0.50507 | 1.63 |
| 5385 | *FAM180B* | 3.7714 | 1.8262 | -0.70137 | -0.8504 | 1.63 |
| 5386 | *EYA3* | 3.8193 | 0.8403 | 0.23578 | -0.83912 | 1.63 |
| 5387 | *SLC34A1* | 4.115 | 0.95197 | -0.17199 | -0.42573 | 1.63 |
| 5388 | *ZNF727* | 2.1102 | 1.9188 | 0.86571 | 0.17659 | 1.63 |
| 5389 | *DZANK1* | 5.0786 | 0.65341 | -0.83739 | -1.4823 | 1.63 |
| 5390 | *IGFL1* | 2.9163 | 1.1018 | 0.87648 | -1.9398 | 1.63 |
| 5391 | *KRTAP5-11* | 3.8469 | 1.3604 | -0.31309 | -0.9644 | 1.63 |
| 5392 | *OR5AS1* | 3.7248 | 1.6365 | -0.46751 | -0.77038 | 1.63 |
| 5393 | *C3orf22* | 2.9528 | 1.7676 | 0.17324 | -0.0541 | 1.63 |
| 5394 | *CLDN23* | 3.2126 | 2.15 | -0.47001 | -1.911 | 1.63 |
| 5395 | *S100A3* | 3.8705 | 2.1571 | -1.1353 | -1.6119 | 1.63 |
| 5396 | *CDYL2* | 4.3399 | 0.81599 | -0.26384 | -0.91236 | 1.63 |
| 5397 | *MARCH5* | 4.8085 | 0.28057 | -0.19704 | -0.22638 | 1.63 |
| 5398 | *ODF3L2* | 2.4294 | 2.1238 | 0.33869 | -1.5588 | 1.63 |
| 5399 | *HS3ST5* | 3.2908 | 0.83939 | 0.76116 | -1.0784 | 1.63 |
| 5400 | *SOX7* | 2.8233 | 1.1001 | 0.9677 | 0.2058 | 1.63 |
| 5401 | *HSPB8* | 3.3288 | 1.5518 | 0.00962 | -0.2863 | 1.63 |
| 5402 | *HUNK* | 1.9706 | 1.5415 | 1.3779 | 1.3244 | 1.63 |
| 5403 | *UBE3A* | 4.4943 | 0.97511 | -0.58028 | -0.84419 | 1.63 |
| 5404 | *HIST1H4D* | 4.8741 | 0.67729 | -0.66251 | -2.3389 | 1.63 |
| 5405 | *INSRR* | 3.4576 | 0.8685 | 0.56252 | -0.99832 | 1.63 |
| 5406 | *ZNF205* | 3.9815 | 1.3948 | -0.48795 | -0.72038 | 1.63 |
| 5407 | *PRMT6* | 2.6941 | 1.8981 | 0.29586 | -0.16202 | 1.63 |
| 5408 | *PYURF* | 3.051 | 2.4518 | -0.61494 | -0.72553 | 1.63 |
| 5409 | *RPA2* | 3.8499 | 0.60364 | 0.43401 | -1.0803 | 1.63 |
| 5410 | *GALNT18* | 3.9346 | 0.88982 | 0.06286 | -0.58686 | 1.63 |
| 5411 | *CD3E* | 3.0883 | 2.2842 | -0.48564 | -2.3466 | 1.63 |
| 5412 | *ULBP3* | 3.6934 | 1.0828 | 0.11049 | -0.78632 | 1.63 |
| 5413 | *KRTAP10-7* | 3.4734 | 2.2751 | -0.86182 | -1.2051 | 1.63 |
| 5414 | *TARP* | 2.7514 | 1.1164 | 1.0184 | 0.9593 | 1.63 |
| 5415 | *STXBP5* | 2.0338 | 1.5061 | 1.3457 | 0.66954 | 1.63 |
| 5416 | *EPN2* | 4.3049 | 1.2683 | -0.68909 | -2.071 | 1.63 |
| 5417 | *RBPMS* | 3.7072 | 1.004 | 0.1727 | -2.5935 | 1.63 |
| 5418 | *WDR24* | 3.7706 | 1.1745 | -0.06144 | -0.48263 | 1.63 |
| 5419 | *UBN1* | 3.9752 | 1.7875 | -0.88008 | -2.0113 | 1.63 |
| 5420 | *SNCAIP* | 3.5933 | 0.65552 | 0.63208 | 0.54744 | 1.63 |
| 5421 | *CYP11B2* | 3.72 | 2.4895 | -1.3287 | -1.8714 | 1.63 |

| | | | | | | |
|---|---|---|---|---|---|---|
| 5422 | *HCAR1* | 4.5122 | 0.36192 | 0.00651 | -0.55122 | 1.63 |
| 5423 | *LOC200726* | 2.5619 | 1.7489 | 0.56949 | -1.864 | 1.63 |
| 5424 | *SCN10A* | 3.3111 | 1.7668 | -0.20039 | -1.6243 | 1.63 |
| 5425 | *SELK* | 4.1105 | 0.61339 | 0.15312 | -2.0363 | 1.63 |
| 5426 | *KIAA0513* | 4.1879 | 0.77703 | -0.08832 | -1.3203 | 1.63 |
| 5427 | *FAM135A* | 3.0674 | 1.326 | 0.48301 | -0.17208 | 1.63 |
| 5428 | *RFPL4A* | 3.5581 | 2.7499 | -1.4319 | -1.5431 | 1.63 |
| 5429 | *C10orf88* | 2.2917 | 1.8658 | 0.71843 | -0.56145 | 1.63 |
| 5430 | *TRIM4* | 3.8954 | 0.56999 | 0.41031 | -0.8562 | 1.63 |
| 5431 | *ZBTB12* | 2.9238 | 2.8143 | -0.86367 | -1.4223 | 1.62 |
| 5432 | *CRTAC1* | 3.4325 | 2.696 | -1.2541 | -2.1202 | 1.62 |
| 5433 | *SNX6* | 4.1719 | 2.1177 | -1.4154 | -1.9375 | 1.62 |
| 5434 | *TMEM123* | 2.8332 | 2.1111 | -0.07051 | -0.3703 | 1.62 |
| 5435 | *BTNL9* | 2.3446 | 2.1729 | 0.35415 | -0.27146 | 1.62 |
| 5436 | *IL16* | 3.5755 | 1.5243 | -0.2283 | -0.45569 | 1.62 |
| 5437 | *PABPC1L2A* | 3.0772 | 0.90799 | 0.88599 | 0.44793 | 1.62 |
| 5438 | *KCNS3* | 3.8438 | 0.52363 | 0.50316 | -0.92642 | 1.62 |
| 5439 | *MAP3K12* | 3.9399 | 0.82297 | 0.10662 | -0.44353 | 1.62 |
| 5440 | *MSRB2* | 2.6756 | 1.1881 | 1.0046 | 0.62611 | 1.62 |
| 5441 | *CBLC* | 2.7036 | 2.3367 | -0.17202 | -0.59264 | 1.62 |
| 5442 | *USP25* | 2.9483 | 1.7089 | 0.20954 | -0.77097 | 1.62 |
| 5443 | *RORC* | 2.908 | 1.5904 | 0.36831 | -1.1575 | 1.62 |
| 5444 | *RAB4A* | 2.9138 | 1.9337 | 0.01861 | -0.63368 | 1.62 |
| 5445 | *ADORA2B* | 5.0062 | 0.27934 | -0.42075 | -0.44353 | 1.62 |
| 5446 | *PLIN3* | 3.1716 | 1.1484 | 0.54435 | -1.7908 | 1.62 |
| 5447 | *TMCO1* | 4.3951 | 1.1841 | -0.71522 | -1.8087 | 1.62 |
| 5448 | *IL15* | 2.67 | 1.5619 | 0.62956 | -0.08049 | 1.62 |
| 5449 | *TOM1L1* | 2.3217 | 1.3165 | 1.2227 | 0.49083 | 1.62 |
| 5450 | *EIF1B* | 2.5636 | 1.8361 | 0.46063 | -1.7241 | 1.62 |
| 5451 | *WDR78* | 3.3493 | 0.78652 | 0.72425 | -0.9879 | 1.62 |
| 5452 | *KCTD1* | 2.5743 | 2.519 | -0.23372 | -1.7585 | 1.62 |
| 5453 | *SWT1* | 2.2284 | 1.483 | 1.1481 | -1.7378 | 1.62 |
| 5454 | *CLEC4M* | 5.4807 | -0.10195 | -0.5193 | -0.65169 | 1.62 |
| 5455 | *BDP1* | 3.6981 | 1.4304 | -0.26968 | -1.8073 | 1.62 |
| 5456 | *APELA* | 3.0753 | 1.3332 | 0.4491 | 0.18702 | 1.62 |
| 5457 | *TRIM64* | 4.1288 | 0.65641 | 0.07112 | -1.6801 | 1.62 |
| 5458 | *GJA4* | 3.0767 | 2.5193 | -0.74011 | -1.0663 | 1.62 |
| 5459 | *ART4* | 5.1801 | 0.70097 | -1.0256 | -1.3711 | 1.62 |
| 5460 | *DCTN3* | 4.9087 | 0.26746 | -0.32075 | -2.0697 | 1.62 |
| 5461 | *SCLY* | 3.4885 | 0.68479 | 0.68177 | 0.30034 | 1.62 |
| 5462 | *PTK7* | 3.719 | 1.0955 | 0.04039 | -1.8121 | 1.62 |
| 5463 | *ASXL1* | 4.8617 | 1.3535 | -1.3613 | -2.2449 | 1.62 |
| 5464 | *GPATCH2* | 2.4647 | 2.4502 | -0.06143 | -1.111 | 1.62 |
| 5465 | *H3F3C* | 3.0094 | 1.1851 | 0.65884 | 0.15694 | 1.62 |
| 5466 | *TIGD4* | 4.8064 | 0.038871 | 0.00711 | -0.9382 | 1.62 |
| 5467 | *ZNF485* | 5.0454 | 0.045942 | -0.23922 | -0.34906 | 1.62 |

| | | | | | | |
|---|---|---|---|---|---|---|
| 5468 | *SENP6* | 3.1397 | 0.86236 | 0.84938 | -3.4062 | 1.62 |
| 5469 | *REEP1* | 4.4312 | 0.40731 | 0.01243 | -0.95057 | 1.62 |
| 5470 | *SERPINB8* | 3.9348 | 0.55526 | 0.36065 | -0.56635 | 1.62 |
| 5471 | *ELF4* | 2.435 | 1.7351 | 0.6798 | -1.9874 | 1.62 |
| 5472 | *CTLA4* | 4.5228 | 0.90782 | -0.58096 | -1.5061 | 1.62 |
| 5473 | *TMEM56* | 2.835 | 1.514 | 0.50055 | -0.34987 | 1.62 |
| 5474 | *LILRA6* | 3.4758 | 1.0956 | 0.27812 | -0.93188 | 1.62 |
| 5475 | *SLC35F5* | 4.2133 | 0.53688 | 0.09832 | -1.5153 | 1.62 |
| 5476 | *ASTN2* | 4.1076 | 0.87731 | -0.13669 | -1.4197 | 1.62 |
| 5477 | *ABCB5* | 2.6766 | 1.5458 | 0.62537 | -0.0062 | 1.62 |
| 5478 | *FGD1* | 3.3829 | 2.1497 | -0.68495 | -2.0433 | 1.62 |
| 5479 | *CXCR3* | 3.9729 | 0.9251 | -0.05048 | -0.10341 | 1.62 |
| 5480 | *OR8G2* | 5.3929 | 0.34909 | -0.89474 | -1.6526 | 1.62 |
| 5481 | *PTH2R* | 3.9283 | 1.6711 | -0.75313 | -1.6992 | 1.62 |
| 5482 | *CXCL17* | 2.5277 | 2.3922 | -0.07372 | -0.82875 | 1.62 |
| 5483 | *ARPP19* | 2.7838 | 2.7711 | -0.70984 | -1.4996 | 1.62 |
| 5484 | *PRKAR1B* | 3.5453 | 2.3562 | -1.0567 | -1.5302 | 1.61 |
| 5485 | *CTXN2* | 3.041 | 1.2373 | 0.56623 | -0.01938 | 1.61 |
| 5486 | *OR52J3* | 2.4729 | 1.9915 | 0.37994 | 0.186 | 1.61 |
| 5487 | *PALD1* | 3.8418 | 1.8303 | -0.82833 | -0.83278 | 1.61 |
| 5488 | *JMJD8* | 4.1528 | 0.40753 | 0.28287 | -1.1169 | 1.61 |
| 5489 | *GSTA3* | 2.7495 | 2.48 | -0.38714 | -0.51164 | 1.61 |
| 5490 | *FBXL12* | 4.3777 | 1.3735 | -0.90947 | -1.2396 | 1.61 |
| 5491 | *PSMC1* | 4.0265 | 1.7525 | -0.93781 | -1.2705 | 1.61 |
| 5492 | *RHOU* | 2.4143 | 2.0583 | 0.36788 | -0.78911 | 1.61 |
| 5493 | *L3MBTL2* | 3.9495 | 0.63118 | 0.25913 | -1.927 | 1.61 |
| 5494 | *CHAC2* | 3.3991 | 1.2977 | 0.14284 | -2.2254 | 1.61 |
| 5495 | *TSFM* | 4.2314 | 0.75771 | -0.14987 | -0.8198 | 1.61 |
| 5496 | *PABPN1L* | 2.9636 | 1.0378 | 0.83702 | -0.52896 | 1.61 |
| 5497 | *ERBB3* | 3.0551 | 1.1064 | 0.67638 | -2.1648 | 1.61 |
| 5498 | *HS6ST2* | 4.5492 | 1.6164 | -1.329 | -1.9203 | 1.61 |
| 5499 | *KRTAP8-1* | 3.6903 | 0.73702 | 0.40904 | -0.63687 | 1.61 |
| 5500 | *OR2L8* | 3.5093 | 1.0984 | 0.22748 | -0.43488 | 1.61 |
| 5501 | *ZNF92* | 2.4866 | 1.8234 | 0.52418 | 0.38037 | 1.61 |
| 5502 | *PIGO* | 3.2159 | 1.9637 | -0.34601 | -0.69935 | 1.61 |
| 5503 | *ITGA1* | 2.4273 | 1.4641 | 0.94137 | -2.3795 | 1.61 |
| 5504 | *KRTAP19-1* | 5.803 | 0.21321 | -1.1845 | -1.9431 | 1.61 |
| 5505 | *PRPF3* | 2.548 | 1.9703 | 0.3133 | 0.051282 | 1.61 |
| 5506 | *CREB5* | 3.7576 | 1.0861 | -0.01275 | -0.68992 | 1.61 |
| 5507 | *SLC14A2* | 3.8747 | 0.63198 | 0.32383 | -0.79438 | 1.61 |
| 5508 | *YTHDF2* | 3.2443 | 1.214 | 0.37094 | -1.5956 | 1.61 |
| 5509 | *REV3L* | 3.8382 | 3.1708 | -2.1798 | -3.0833 | 1.61 |
| 5510 | *CYP24A1* | 3.5946 | 0.64758 | 0.58689 | 0.46584 | 1.61 |
| 5511 | *TMEM200B* | 2.5039 | 2.2578 | 0.06698 | -0.26949 | 1.61 |
| 5512 | *ARHGAP17* | 3.4457 | 1.74 | -0.35735 | -1.6626 | 1.61 |
| 5513 | *POMGNT2* | 4.0592 | 1.4107 | -0.64219 | -0.88876 | 1.61 |

| | | | | | | |
|---|---|---|---|---|---|---|
| 5514 | *DMXL1* | 4.124 | 0.52075 | 0.18284 | -1.1084 | 1.61 |
| 5515 | *DNAAF1* | 2.6756 | 1.4485 | 0.7034 | -0.41505 | 1.61 |
| 5516 | *FOXK1* | 4.305 | 0.92244 | -0.40118 | -2.1926 | 1.61 |
| 5517 | *RAB26* | 3.1137 | 1.6922 | 0.01981 | -0.77018 | 1.61 |
| 5518 | *ADCYAP1R1* | 4.5423 | 0.26234 | 0.02079 | -1.2135 | 1.61 |
| 5519 | *GPR20* | 4.7557 | 0.19645 | -0.12701 | -0.4674 | 1.61 |
| 5520 | *TRHR* | 4.6142 | 0.16383 | 0.04651 | -0.17629 | 1.61 |
| 5521 | *TOMM70A* | 4.6407 | 0.48014 | -0.29652 | -1.4913 | 1.61 |
| 5522 | *SNX13* | 4.2164 | 0.3066 | 0.3003 | -0.85214 | 1.61 |
| 5523 | *P4HB* | 3.826 | 2.8311 | -1.8341 | -2.1956 | 1.61 |
| 5524 | *NLRP12* | 5.2141 | 0.1473 | -0.53904 | -2.0382 | 1.61 |
| 5525 | *RBM25* | 3.3385 | 2.0149 | -0.53321 | -0.84141 | 1.61 |
| 5526 | *EXOC6B* | 4.1619 | 0.87025 | -0.21267 | -1.342 | 1.61 |
| 5527 | *SLC25A17* | 3.3141 | 3.1579 | -1.6528 | -2.4405 | 1.61 |
| 5528 | *NCOA3* | 3.5869 | 1.2186 | 0.01352 | -0.23533 | 1.61 |
| 5529 | *SCFD1* | 3.199 | 0.97612 | 0.64372 | -1.0406 | 1.61 |
| 5530 | *IDI2* | 4.8791 | 0.88301 | -0.94333 | -2.5156 | 1.61 |
| 5531 | *SFXN1* | 3.5408 | 0.64629 | 0.63081 | 0.069986 | 1.61 |
| 5532 | *CD52* | 2.4765 | 2.1091 | 0.23173 | 0.067594 | 1.61 |
| 5533 | *MTHFD1* | 1.9155 | 1.6207 | 1.2811 | -1.013 | 1.61 |
| 5534 | *MAN1B1* | 3.7963 | 1.036 | -0.01599 | -0.87396 | 1.61 |
| 5535 | *SGCA* | 2.8176 | 2.3486 | -0.34996 | -1.9878 | 1.61 |
| 5536 | *CUL2* | 4.0602 | 0.59341 | 0.16209 | -0.91253 | 1.61 |
| 5537 | *USP26* | 1.8396 | 1.8098 | 1.1661 | -0.91283 | 1.61 |
| 5538 | *FAM45A* | 2.983 | 1.6725 | 0.15898 | -0.77404 | 1.60 |
| 5539 | *PRDM7* | 3.5725 | 2.3228 | -1.082 | -1.2903 | 1.60 |
| 5540 | *TFIP11* | 3.7195 | 1.6909 | -0.59804 | -0.90588 | 1.60 |
| 5541 | *ZNF812* | 5.5973 | 0.342 | -1.1271 | -1.2125 | 1.60 |
| 5542 | *POMK* | 4.3523 | 1.1162 | -0.65667 | -3.5581 | 1.60 |
| 5543 | *DERL2* | 4.8403 | 0.16107 | -0.19042 | -0.97986 | 1.60 |
| 5544 | *SLCO5A1* | 4.8281 | 1.1896 | -1.2071 | -1.666 | 1.60 |
| 5545 | *CD86* | 3.993 | 1.5564 | -0.73911 | -1.7034 | 1.60 |
| 5546 | *PRAMEF2* | 3.2315 | 2.0257 | -0.44742 | -0.5458 | 1.60 |
| 5547 | *CCDC24* | 3.1449 | 2.6605 | -0.9968 | -3.0219 | 1.60 |
| 5548 | *KCNB2* | 3.6212 | 0.67919 | 0.50728 | 0.34505 | 1.60 |
| 5549 | *FDCSP* | 2.6524 | 1.4998 | 0.65531 | -0.35367 | 1.60 |
| 5550 | *GPX5* | 2.9253 | 2.6717 | -0.79016 | -1.6824 | 1.60 |
| 5551 | *MAB21L3* | 2.7836 | 2.2358 | -0.21349 | -1.0293 | 1.60 |
| 5552 | *PARK7* | 3.7384 | 1.2897 | -0.22242 | -1.7379 | 1.60 |
| 5553 | *PIK3R6* | 4.5133 | 0.61486 | -0.32262 | -0.9858 | 1.60 |
| 5554 | *DNAJC27* | 4.3276 | 1.1913 | -0.71347 | -0.8934 | 1.60 |
| 5555 | *PGK2* | 3.3415 | 1.0001 | 0.46373 | -1.6756 | 1.60 |
| 5556 | *PRKACB* | 2.1187 | 1.5947 | 1.0914 | 0.32079 | 1.60 |
| 5557 | *DAB1* | 3.2462 | 1.6617 | -0.1037 | -0.48995 | 1.60 |
| 5558 | *EVA1C* | 3.3954 | 1.8927 | -0.48419 | -5.7716 | 1.60 |
| 5559 | *DHRS7* | 4.1734 | 1.6307 | -1.0019 | -1.7822 | 1.60 |

| | | | | | | |
|---|---|---|---|---|---|---|
| 5560 | *TEX9* | 2.7018 | 2.0683 | 0.03176 | -1.2056 | 1.60 |
| 5561 | *FAM195B* | 4.6085 | 0.27906 | -0.08607 | -0.80165 | 1.60 |
| 5562 | *BEST1* | 5.0306 | -0.01496 | -0.2146 | -1.7364 | 1.60 |
| 5563 | *MST1R* | 4.0868 | 0.39834 | 0.31512 | -2.0818 | 1.60 |
| 5564 | *TMPRSS7* | 3.0339 | 1.4423 | 0.32337 | -0.63486 | 1.60 |
| 5565 | *SLAMF9* | 4.3313 | 0.6579 | -0.19136 | -0.78823 | 1.60 |
| 5566 | *UBAC2* | 4.7575 | 0.61885 | -0.57899 | -0.7068 | 1.60 |
| 5567 | *HRSP12* | 3.0664 | 1.0441 | 0.68684 | -2.7566 | 1.60 |
| 5568 | *TTC7A* | 3.8738 | 1.3353 | -0.41207 | -0.85935 | 1.60 |
| 5569 | *PCOLCE* | 2.7768 | 2.3618 | -0.34227 | -2.0059 | 1.60 |
| 5570 | *FNIP1* | 3.2564 | 0.8204 | 0.71881 | -0.19793 | 1.60 |
| 5571 | *PDE10A* | 3.9121 | 0.68105 | 0.20224 | -0.37334 | 1.60 |
| 5572 | *KCP* | 4.5887 | 0.94611 | -0.74086 | -2.0952 | 1.60 |
| 5573 | *IL18R1* | 2.9008 | 1.7753 | 0.1177 | -1.2845 | 1.60 |
| 5574 | *CXADR* | 2.5555 | 2.4634 | -0.22673 | -0.63753 | 1.60 |
| 5575 | *ZNF682* | 1.7756 | 1.5114 | 1.5049 | -0.38937 | 1.60 |
| 5576 | *LBHD1* | 3.5145 | 0.97964 | 0.29658 | 0.010355 | 1.60 |
| 5577 | *TMEM185A* | 4.2965 | 1.0909 | -0.59678 | -1.237 | 1.60 |
| 5578 | *EIF5A2* | 3.4486 | 1.9461 | -0.60468 | -0.7614 | 1.60 |
| 5579 | *ARMC3* | 2.6822 | 1.2793 | 0.8281 | -0.91913 | 1.60 |
| 5580 | *IFITM3* | 2.9821 | 2.0519 | -0.24512 | -0.4537 | 1.60 |
| 5581 | *NT5C* | 3.2701 | 1.9632 | -0.44464 | -1.5608 | 1.60 |
| 5582 | *ARR3* | 4.0011 | 1.223 | -0.43683 | -1.0451 | 1.60 |
| 5583 | *LOC100652758* | 3.7059 | 1.2985 | -0.21806 | -1.1463 | 1.60 |
| 5584 | *KDELR2* | 4.2961 | 1.1315 | -0.64268 | -1.1718 | 1.59 |
| 5585 | *CREB3* | 4.4271 | 0.73403 | -0.37698 | -1.307 | 1.59 |
| 5586 | *COL4A6* | 2.6278 | 2.3548 | -0.19858 | -3.0444 | 1.59 |
| 5587 | *MOB2* | 2.853 | 2.2554 | -0.32455 | -0.3576 | 1.59 |
| 5588 | *FANCG* | 4.1178 | 0.7007 | -0.03472 | -0.37215 | 1.59 |
| 5589 | *CH25H* | 2.6762 | 1.129 | 0.97836 | -2.2385 | 1.59 |
| 5590 | *WHAMM* | 3.8438 | 0.98837 | -0.04931 | -0.38667 | 1.59 |
| 5591 | *WDR35* | 4.6344 | 0.40132 | -0.25291 | -0.37656 | 1.59 |
| 5592 | *COL2A1* | 4.3599 | 1.2181 | -0.79663 | -1.5467 | 1.59 |
| 5593 | *CEP162* | 4.191 | 0.38929 | 0.20105 | -1.5908 | 1.59 |
| 5594 | *PIK3IP1* | 3.8358 | 1.9914 | -1.0467 | -1.5695 | 1.59 |
| 5595 | *ZNRF4* | 4.9522 | 1.4853 | -1.6597 | -1.8952 | 1.59 |
| 5596 | *SNX14* | 4.0069 | 0.42299 | 0.34629 | -2.0548 | 1.59 |
| 5597 | *CPT1C* | 3.2888 | 0.99308 | 0.4923 | -1.2985 | 1.59 |
| 5598 | *SLC7A7* | 4.2676 | 0.35108 | 0.1532 | -0.90281 | 1.59 |
| 5599 | *MRPS18C* | 4.0323 | 0.92431 | -0.1849 | -1.3074 | 1.59 |
| 5600 | *KCNE4* | 2.927 | 1.1221 | 0.72224 | -0.91379 | 1.59 |
| 5601 | *B3GAT2* | 3.5089 | 1.1885 | 0.07384 | -1.5039 | 1.59 |
| 5602 | *WBP1* | 2.8386 | 1.7095 | 0.22299 | -0.05054 | 1.59 |
| 5603 | *RPRML* | 4.0354 | 0.54222 | 0.19327 | 0.08196 | 1.59 |
| 5604 | *GPRIN2* | 3.1722 | 1.2549 | 0.34292 | -0.0209 | 1.59 |
| 5605 | *NKX6-1* | 4.5215 | 0.24025 | 0.00711 | -1.4435 | 1.59 |

| | | | | | | |
|---|---|---|---|---|---|---|
| 5606 | *TRIOBP* | 3.5374 | 1.1496 | 0.08176 | -1.5667 | 1.59 |
| 5607 | *APOC2* | 4.2841 | 0.42022 | 0.06348 | -2.1321 | 1.59 |
| 5608 | *TIAM1* | 3.39 | 0.93837 | 0.43888 | -0.18167 | 1.59 |
| 5609 | *EHBP1* | 3.8152 | 0.75886 | 0.19317 | -0.78805 | 1.59 |
| 5610 | *KIF15* | 3.6169 | 2.5475 | -1.3973 | -2.8398 | 1.59 |
| 5611 | *GOLPH3L* | 2.66 | 1.1341 | 0.9708 | 0.8949 | 1.59 |
| 5612 | *SIX3* | 3.0505 | 2.491 | -0.7769 | -1.09 | 1.59 |
| 5613 | *TFR2* | 2.5503 | 1.2523 | 0.96077 | -1.9874 | 1.59 |
| 5614 | *USP28* | 5.1611 | -0.18108 | -0.21692 | -0.71753 | 1.59 |
| 5615 | *C1orf146* | 5.8607 | -0.43276 | -0.66583 | -0.66948 | 1.59 |
| 5616 | *THOC6* | 2.7331 | 1.8187 | 0.21 | -0.34566 | 1.59 |
| 5617 | *CRY2* | 4.2499 | 2.1742 | -1.6636 | -2.4405 | 1.59 |
| 5618 | *DFFB* | 3.3851 | 1.2035 | 0.17047 | -1.6084 | 1.59 |
| 5619 | *AKR1B15* | 3.3244 | 0.88108 | 0.55316 | -0.04588 | 1.59 |
| 5620 | *C1orf204* | 4.2716 | 1.1841 | -0.69748 | -0.83471 | 1.59 |
| 5621 | *PRRX1* | 2.9506 | 1.7632 | 0.04435 | -0.55371 | 1.59 |
| 5622 | *OR8A1* | 2.3673 | 1.2227 | 1.1667 | 0.35826 | 1.59 |
| 5623 | *SPIRE1* | 2.2589 | 1.9581 | 0.53845 | -1.6776 | 1.59 |
| 5624 | *GALNT10* | 3.0314 | 1.5637 | 0.16025 | -0.44353 | 1.59 |
| 5625 | *ALKBH3* | 2.4403 | 1.5733 | 0.73928 | -0.80743 | 1.58 |
| 5626 | *ZNF148* | 3.0898 | 2.2232 | -0.56036 | -1.5552 | 1.58 |
| 5627 | *CYFIP1* | 4.9728 | -0.03695 | -0.18328 | -0.92721 | 1.58 |
| 5628 | *PHLDA3* | 2.9937 | 1.1829 | 0.57566 | -2.1648 | 1.58 |
| 5629 | *DNAJC16* | 3.8468 | 1.409 | -0.50357 | -1.1479 | 1.58 |
| 5630 | *RIPPLY2* | 3.5542 | 0.67683 | 0.52067 | -4.6085 | 1.58 |
| 5631 | *AKAP2* | 4.1484 | 0.72209 | -0.12131 | -0.74804 | 1.58 |
| 5632 | *SLC20A2* | 2.9973 | 2.0376 | -0.28621 | -0.38677 | 1.58 |
| 5633 | *NRIP1* | 1.8339 | 1.5479 | 1.3658 | 0.45962 | 1.58 |
| 5634 | *ABCB4* | 3.9191 | 1.0709 | -0.24411 | -1.5635 | 1.58 |
| 5635 | *PAGE4* | 3.4558 | 1.1775 | 0.11251 | -2.0653 | 1.58 |
| 5636 | *STMN2* | 2.8389 | 1.4603 | 0.44653 | -1.4221 | 1.58 |
| 5637 | *DEPDC1* | 4.0918 | 0.85849 | -0.20669 | -1.1321 | 1.58 |
| 5638 | *TM4SF1* | 5.2448 | 0.11205 | -0.61363 | -0.62875 | 1.58 |
| 5639 | *ZNF777* | 4.2545 | 0.3784 | 0.11011 | -0.13144 | 1.58 |
| 5640 | *ZNF704* | 3.6416 | 1.599 | -0.49902 | -1.035 | 1.58 |
| 5641 | *SNUPN* | 3.7568 | 2.2564 | -1.2724 | -1.3994 | 1.58 |
| 5642 | *OSBPL7* | 3.4033 | 2.2737 | -0.93626 | -1.2952 | 1.58 |
| 5643 | *DACT3* | 4.7478 | 0.077652 | -0.08582 | -0.39235 | 1.58 |
| 5644 | *GABRG1* | 3.2554 | 0.90002 | 0.58385 | -0.19577 | 1.58 |
| 5645 | *PRODH2* | 2.9326 | 2.2739 | -0.46733 | -1.5598 | 1.58 |
| 5646 | *CDHR3* | 3.011 | 2.1076 | -0.37998 | -2.5707 | 1.58 |
| 5647 | *FAM124B* | 5.3778 | -0.2804 | -0.36102 | -1.6992 | 1.58 |
| 5648 | *FRMD6* | 3.4317 | 1.663 | -0.3584 | -0.68133 | 1.58 |
| 5649 | *OTUD1* | 4.9909 | 0.32397 | -0.57857 | -2.3265 | 1.58 |
| 5650 | *YWHAZ* | 3.9052 | 0.62064 | 0.20989 | -0.44353 | 1.58 |
| 5651 | *ITIH5* | 3.7083 | 1.0022 | 0.02461 | -1.912 | 1.58 |

| | | | | | | |
|---|---|---|---|---|---|---|
| 5652 | *C11orf87* | 2.1523 | 2.1213 | 0.46149 | -0.23999 | 1.58 |
| 5653 | *GRB7* | 3.8732 | 0.67452 | 0.18686 | -0.23958 | 1.58 |
| 5654 | *AKT3* | 3.6752 | 0.54931 | 0.51 | -0.10672 | 1.58 |
| 5655 | *PRAMEF19* | 2.9159 | 1.3255 | 0.49305 | -0.36668 | 1.58 |
| 5656 | *NFE4* | 4.1473 | 0.39995 | 0.18664 | -1.3946 | 1.58 |
| 5657 | *ACSM3* | 3.655 | 0.61017 | 0.4687 | -2.8782 | 1.58 |
| 5658 | *SGPP1* | 5.1012 | -0.02503 | -0.34232 | -1.9398 | 1.58 |
| 5659 | *P2RY6* | 3.2 | 1.5927 | -0.0624 | -0.92375 | 1.58 |
| 5660 | *DISP2* | 1.9172 | 1.5777 | 1.2346 | -0.67842 | 1.58 |
| 5661 | *ADCK5* | 3.5787 | 0.88243 | 0.26813 | -0.61989 | 1.58 |
| 5662 | *FAM43B* | 2.9254 | 0.99848 | 0.80493 | -1.0737 | 1.58 |
| 5663 | *IL2RA* | 2.3183 | 2.1325 | 0.27751 | -1.9175 | 1.58 |
| 5664 | *TMEM231* | 3.5432 | 0.74344 | 0.43942 | -2.5065 | 1.58 |
| 5665 | *FAM133A* | 4.2316 | 0.92554 | -0.43276 | -0.49953 | 1.57 |
| 5666 | *SMARCD3* | 3.1974 | 0.88221 | 0.64429 | -1.1825 | 1.57 |
| 5667 | *IGFBP3* | 2.9697 | 1.0724 | 0.68104 | -0.94498 | 1.57 |
| 5668 | *RNF222* | 2.58 | 2.4771 | -0.33428 | -1.2995 | 1.57 |
| 5669 | *SLC12A9* | 3.0693 | 3.0505 | -1.3978 | -2.1185 | 1.57 |
| 5670 | *TPH1* | 3.8783 | 0.46918 | 0.37429 | -0.97381 | 1.57 |
| 5671 | *CNKSR2* | 3.6608 | 0.56758 | 0.49275 | -1.1135 | 1.57 |
| 5672 | *CADM1* | 3.245 | 1.5116 | -0.03588 | -0.86242 | 1.57 |
| 5673 | *PCDHA3* | 4.0288 | 0.37743 | 0.3141 | -1.438 | 1.57 |
| 5674 | *FABP5* | 5.2714 | 0.53813 | -1.0898 | -1.3184 | 1.57 |
| 5675 | *ZFAND3* | 1.8356 | 1.4492 | 1.4337 | -0.66873 | 1.57 |
| 5676 | *NAA60* | 2.3462 | 1.9177 | 0.45389 | -1.4747 | 1.57 |
| 5677 | *LOC339862* | 3.3545 | 1.0341 | 0.32908 | 0.31279 | 1.57 |
| 5678 | *BCKDK* | 5.5419 | -0.17554 | -0.64918 | -0.76606 | 1.57 |
| 5679 | *ZFAND5* | 5.17 | 0.18753 | -0.6408 | -2.0165 | 1.57 |
| 5680 | *C19orf66* | 2.9737 | 0.90081 | 0.84174 | -0.96857 | 1.57 |
| 5681 | *TUSC1* | 4.1924 | 1.8576 | -1.3338 | -1.4767 | 1.57 |
| 5682 | *ACTG2* | 2.4312 | 1.4911 | 0.79352 | -0.61595 | 1.57 |
| 5683 | *CD69* | 2.9909 | 2.2578 | -0.53297 | -0.90576 | 1.57 |
| 5684 | *CSNK2A3* | 4.7985 | 0.13984 | -0.224 | -0.8338 | 1.57 |
| 5685 | *TFAP2D* | 3.6645 | 1.8886 | -0.83891 | -1.8789 | 1.57 |
| 5686 | *BAX* | 2.4996 | 1.6024 | 0.61199 | -1.1385 | 1.57 |
| 5687 | *ENPP5* | 4.6734 | 0.41806 | -0.3787 | -1.2655 | 1.57 |
| 5688 | *OR4N2* | 1.8254 | 1.7669 | 1.1203 | -0.19839 | 1.57 |
| 5689 | *GABRP* | 2.0832 | 2.0188 | 0.61017 | -0.066 | 1.57 |
| 5690 | *RECQL4* | 3.3787 | 0.94489 | 0.38708 | 0.35867 | 1.57 |
| 5691 | *LACTB2* | 3.9625 | 0.8258 | -0.07799 | -0.80498 | 1.57 |
| 5692 | *SLC26A6* | 2.714 | 1.5597 | 0.43635 | -1.4469 | 1.57 |
| 5693 | *CDH24* | 3.5126 | 2.2573 | -1.06 | -2.6275 | 1.57 |
| 5694 | *HOXD3* | 4.5824 | 0.56115 | -0.43473 | -0.78951 | 1.57 |
| 5695 | *RRAS* | 4.5358 | 0.12602 | 0.04683 | -1.2651 | 1.57 |
| 5696 | *TNKS2* | 3.325 | 0.70578 | 0.67665 | -1.5304 | 1.57 |
| 5697 | *CFHR1* | 4.6473 | 0.90791 | -0.84815 | -1.7743 | 1.57 |

| | | | | | | |
|---|---|---|---|---|---|---|
| 5698 | *ALAS1* | 3.9562 | 1.394 | -0.64362 | -1.3166 | 1.57 |
| 5699 | *TCF20* | 4.5312 | 0.30498 | -0.12977 | -2.1151 | 1.57 |
| 5700 | *VASN* | 2.8626 | 1.7558 | 0.08672 | 0.079497 | 1.57 |
| 5701 | *PEX10* | 4.3956 | 0.23325 | 0.07562 | -0.82041 | 1.57 |
| 5702 | *ZZEF1* | 2.406 | 1.7162 | 0.58163 | -1.0519 | 1.57 |
| 5703 | *KIF2A* | 3.2884 | 0.97931 | 0.43594 | -0.64806 | 1.57 |
| 5704 | *HPRT1* | 4.4131 | 1.0775 | -0.78775 | -0.84504 | 1.57 |
| 5705 | *OR5B21* | 2.6235 | 1.1271 | 0.95207 | -0.46711 | 1.57 |
| 5706 | *MYPOP* | 3.3338 | 1.9253 | -0.55655 | -1.9287 | 1.57 |
| 5707 | *UGT1A5* | 2.9974 | 2.6562 | -0.95136 | -1.3071 | 1.57 |
| 5708 | *TRPA1* | 3.405 | 2.2486 | -0.95378 | -1.7365 | 1.57 |
| 5709 | *NAPEPLD* | 3.4002 | 1.4361 | -0.1371 | -0.19183 | 1.57 |
| 5710 | *GPR6* | 3.7772 | 0.68351 | 0.23842 | -1.7459 | 1.57 |
| 5711 | *OGN* | 4.4025 | 0.65697 | -0.36045 | -0.63687 | 1.57 |
| 5712 | *POLDIP3* | 4.8851 | 0.65278 | -0.83912 | -0.94296 | 1.57 |
| 5713 | *CPLX3* | 3.2026 | 1.5814 | -0.08582 | -0.95658 | 1.57 |
| 5714 | *THEM4* | 2.7114 | 2.3568 | -0.3703 | -0.99182 | 1.57 |
| 5715 | *RPP25L* | 2.9015 | 2.5844 | -0.78866 | -0.968 | 1.57 |
| 5716 | *TYW1B* | 5.9922 | -0.13895 | -1.1573 | -1.7862 | 1.57 |
| 5717 | *TRIM3* | 4.6609 | 0.42052 | -0.38553 | -1.1487 | 1.57 |
| 5718 | *SIGLEC11* | 2.3773 | 2.0182 | 0.29909 | -0.42859 | 1.56 |
| 5719 | *ENOX2* | 2.4283 | 2.1606 | 0.10565 | -0.85236 | 1.56 |
| 5720 | *IFNA16* | 3.7557 | 1.0308 | -0.09223 | -1.7507 | 1.56 |
| 5721 | *CLDN19* | 4.225 | 0.35648 | 0.11177 | 0.049512 | 1.56 |
| 5722 | *SLC25A6* | 3.8719 | 0.57734 | 0.24264 | -2.8019 | 1.56 |
| 5723 | *MB* | 4.9504 | 0.073858 | -0.33241 | -1.606 | 1.56 |
| 5724 | *KLK5* | 4.7044 | 0.69767 | -0.71041 | -1.7153 | 1.56 |
| 5725 | *IQCD* | 3.3624 | 3.0088 | -1.6802 | -2.7483 | 1.56 |
| 5726 | *HIST2H2AB* | 4.7725 | 0.93187 | -1.0149 | -1.584 | 1.56 |
| 5727 | *DNAI2* | 3.0665 | 1.4736 | 0.14903 | -1.7994 | 1.56 |
| 5728 | *FAM212B* | 3.041 | 1.3891 | 0.25881 | -0.22469 | 1.56 |
| 5729 | *UBE2D1* | 4.3566 | 0.29986 | 0.03187 | -0.62307 | 1.56 |
| 5730 | *LAP3* | 2.8129 | 1.5677 | 0.30522 | -1.1905 | 1.56 |
| 5731 | *PITPNM1* | 4.4645 | 1.1653 | -0.94399 | -1.7906 | 1.56 |
| 5732 | *PPP2R2B* | 3.5487 | 0.75424 | 0.38261 | -0.29222 | 1.56 |
| 5733 | *POTEI* | 4.115 | 0.46994 | 0.0985 | -1.6649 | 1.56 |
| 5734 | *IL15RA* | 2.737 | 1.6313 | 0.3145 | -1.4916 | 1.56 |
| 5735 | *ALKBH2* | 4.3079 | 0.25115 | 0.1237 | -0.21429 | 1.56 |
| 5736 | *ZNRF3* | 2.5546 | 2.0242 | 0.10131 | -1.1833 | 1.56 |
| 5737 | *C3orf67* | 4.4927 | 0.26829 | -0.08104 | -1.6368 | 1.56 |
| 5738 | *HMBS* | 3.6448 | 1.0432 | -0.00825 | -1.4606 | 1.56 |
| 5739 | *B3GALT6* | 2.9457 | 1.3294 | 0.40444 | -1.6263 | 1.56 |
| 5740 | *CEMP1* | 4.6641 | 0.34615 | -0.33071 | -1.4852 | 1.56 |
| 5741 | *PRDX5* | 3.9247 | 0.61957 | 0.13492 | 0.093787 | 1.56 |
| 5742 | *FXYD6* | 3.5965 | 1.159 | -0.07689 | -0.5047 | 1.56 |
| 5743 | *CRIP1* | 5.1826 | -0.20038 | -0.30376 | -1.1209 | 1.56 |

| | | | | | | |
|---|---|---|---|---|---|---|
| 5744 | *TARM1* | 3.484 | 0.68961 | 0.50476 | -0.02553 | 1.56 |
| 5745 | *PARD6B* | 5.0139 | 0.048059 | -0.38361 | -1.6105 | 1.56 |
| 5746 | *GPATCH8* | 3.04 | 1.0113 | 0.62659 | -1.1882 | 1.56 |
| 5747 | *NBPF12* | 3.539 | 0.76813 | 0.37051 | -1.9573 | 1.56 |
| 5748 | *KLHL12* | 3.9886 | 0.56958 | 0.11931 | -2.2745 | 1.56 |
| 5749 | *ITGB1* | 2.2905 | 1.635 | 0.75188 | -1.5645 | 1.56 |
| 5750 | *CAPN8* | 3.7134 | 1.9345 | -0.97066 | -1.0945 | 1.56 |
| 5751 | *TMEM106B* | 3.8375 | 0.82849 | 0.01096 | -1.5946 | 1.56 |
| 5752 | *ARMT1* | 3.3367 | 1.8461 | -0.5059 | -0.77018 | 1.56 |
| 5753 | *REP15* | 5.1606 | -0.17566 | -0.30804 | -1.0359 | 1.56 |
| 5754 | *OR6N1* | 2.7517 | 1.1461 | 0.77801 | 0.47239 | 1.56 |
| 5755 | *S100PBP* | 3.5311 | 0.74126 | 0.40191 | 0.34241 | 1.56 |
| 5756 | *FRMPD3* | 3.4713 | 0.96682 | 0.23609 | -1.2105 | 1.56 |
| 5757 | *OR2AT4* | 4.4003 | 0.2658 | 0.00756 | -0.72201 | 1.56 |
| 5758 | *DEFB130* | 2.5489 | 2.0915 | 0.03282 | -1.2915 | 1.56 |
| 5759 | *CHST11* | 3.2995 | 1.2335 | 0.13995 | -0.3311 | 1.56 |
| 5760 | *LRP3* | 5.2759 | 0.35512 | -0.95838 | -1.6926 | 1.56 |
| 5761 | *SLC35D3* | 5.071 | 0.86953 | -1.2683 | -2.254 | 1.56 |
| 5762 | *SLC10A1* | 2.2251 | 1.7583 | 0.6884 | -0.65145 | 1.56 |
| 5763 | *BROX* | 2.6415 | 2.4481 | -0.41803 | -1.095 | 1.56 |
| 5764 | *GRM8* | 2.7718 | 2.137 | -0.23846 | -0.49311 | 1.56 |
| 5765 | *KDM4A* | 3.0683 | 1.7353 | -0.13328 | -0.35744 | 1.56 |
| 5766 | *MMGT1* | 4.9248 | 1.7282 | -1.985 | -2.1058 | 1.56 |
| 5767 | *ATF3* | 4.0311 | 0.63571 | -0.00066 | -1.2657 | 1.56 |
| 5768 | *WNT5B* | 3.854 | 0.84309 | -0.03116 | -0.78941 | 1.56 |
| 5769 | *AADAC* | 2.7854 | 1.0559 | 0.8246 | 0.088509 | 1.56 |
| 5770 | *GRID2IP* | 3.0945 | 0.80412 | 0.76597 | 0.2955 | 1.55 |
| 5771 | *UPB1* | 3.6412 | 0.80025 | 0.22188 | -0.39689 | 1.55 |
| 5772 | *MAK16* | 4.001 | 0.56611 | 0.09562 | -1.5956 | 1.55 |
| 5773 | *PIM2* | 2.759 | 1.7913 | 0.11235 | -0.4676 | 1.55 |
| 5774 | *ADH5* | 4.2177 | 0.23314 | 0.21148 | -2.6683 | 1.55 |
| 5775 | *METTL2A* | 4.4535 | 0.11496 | 0.0936 | -1.6271 | 1.55 |
| 5776 | *KATNA1* | 3.6915 | 2.1815 | -1.2114 | -1.4306 | 1.55 |
| 5777 | *ASPG* | 3.7347 | 2.942 | -2.0161 | -2.5419 | 1.55 |
| 5778 | *ADIG* | 5.0002 | 1.239 | -1.5788 | -2.0842 | 1.55 |
| 5779 | *GLS2* | 4.7308 | 0.099866 | -0.17081 | -2.1817 | 1.55 |
| 5780 | *OR4C12* | 2.2931 | 2.2873 | 0.07918 | -1.1021 | 1.55 |
| 5781 | *CSF1* | 3.6208 | 0.77738 | 0.26076 | 0.1939 | 1.55 |
| 5782 | *COX4I2* | 2.9169 | 2.0041 | -0.26245 | -1.8038 | 1.55 |
| 5783 | *TEX14* | 2.6621 | 1.9913 | 0.00305 | -0.3377 | 1.55 |
| 5784 | *TTLL11* | 5.4564 | -0.38503 | -0.41533 | -1.3835 | 1.55 |
| 5785 | *GOLGA8K* | 4.3952 | 0.21295 | 0.04757 | -0.99729 | 1.55 |
| 5786 | *ZNF479* | 2.7889 | 1.3075 | 0.55821 | -0.17622 | 1.55 |
| 5787 | *C19orf81* | 2.2309 | 2.1613 | 0.26207 | -2.5686 | 1.55 |
| 5788 | *C17orf77* | 2.6692 | 2.0087 | -0.02387 | -0.39012 | 1.55 |
| 5789 | *SCHIP1* | 4.3797 | 0.43954 | -0.16569 | -1.7263 | 1.55 |

| | | | | | | |
|---|---|---|---|---|---|---|
| 5790 | *MYO5B* | 4.5304 | 0.48797 | -0.36579 | -1.0383 | 1.55 |
| 5791 | *UPF1* | 3.7447 | 0.47055 | 0.43717 | -2.6024 | 1.55 |
| 5792 | *STK33* | 2.5674 | 2.1822 | -0.09731 | -2.0837 | 1.55 |
| 5793 | *KHDRBS3* | 3.3429 | 1.0577 | 0.25162 | -0.76582 | 1.55 |
| 5794 | *INHBA* | 3.1754 | 1.3542 | 0.12258 | -0.60072 | 1.55 |
| 5795 | *OMA1* | 3.5528 | 1.1051 | -0.0083 | -0.37213 | 1.55 |
| 5796 | *THUMPD2* | 2.6154 | 1.3485 | 0.68562 | -1.8387 | 1.55 |
| 5797 | *TAS2R39* | 2.8021 | 1.066 | 0.78063 | -0.14173 | 1.55 |
| 5798 | *SAMD3* | 2.8317 | 1.2959 | 0.52111 | 0.32176 | 1.55 |
| 5799 | *WDR20* | 2.9752 | 1.7479 | -0.0746 | -0.12945 | 1.55 |
| 5800 | *RHOXF2B* | 2.5622 | 1.513 | 0.57321 | -1.3668 | 1.55 |
| 5801 | *SMG8* | 3.508 | 1.4165 | -0.27612 | -0.56492 | 1.55 |
| 5802 | *HSBP1* | 4.3431 | 0.52376 | -0.22042 | -0.61585 | 1.55 |
| 5803 | *KRTAP10-5* | 4.3174 | 0.30382 | 0.02521 | -0.02965 | 1.55 |
| 5804 | *KIAA2026* | 3.3519 | 0.99875 | 0.29535 | -1.5167 | 1.55 |
| 5805 | *HSD17B14* | 3.9173 | 2.0317 | -1.3036 | -1.5267 | 1.55 |
| 5806 | *GOT1* | 4.1722 | 0.47018 | 0.00283 | -0.91981 | 1.55 |
| 5807 | *PCMTD2* | 4.1158 | 0.94146 | -0.41237 | -1.6044 | 1.55 |
| 5808 | *SCN5A* | 4.747 | 0.51984 | -0.62395 | -1.0359 | 1.55 |
| 5809 | *ENPP4* | 4.9015 | 0.077732 | -0.33682 | -1.5268 | 1.55 |
| 5810 | *PDLIM5* | 3.4033 | 0.92528 | 0.31371 | -0.35449 | 1.55 |
| 5811 | *REXO4* | 3.6211 | 1.5286 | -0.50748 | -2.1984 | 1.55 |
| 5812 | *TMEM55B* | 3.2017 | 0.85669 | 0.58307 | -0.07087 | 1.55 |
| 5813 | *CLK2* | 4.2692 | 0.74226 | -0.3701 | -0.72196 | 1.55 |
| 5814 | *HIST1H1E* | 3.073 | 2.7867 | -1.219 | -1.9533 | 1.55 |
| 5815 | *REG3G* | 3.9486 | 0.48577 | 0.2057 | -1.9919 | 1.55 |
| 5816 | *DACT2* | 4.1789 | 0.25015 | 0.21099 | -1.2693 | 1.55 |
| 5817 | *PIGA* | 2.7464 | 1.7034 | 0.19018 | -1.6065 | 1.55 |
| 5818 | *SYNE4* | 3.4795 | 2.0049 | -0.84452 | -1.6182 | 1.55 |
| 5819 | *ZNF320* | 3.8484 | 1.5405 | -0.74972 | -1.0872 | 1.55 |
| 5820 | *PPM1J* | 3.0961 | 1.3549 | 0.18793 | -0.26332 | 1.55 |
| 5821 | *GRHL3* | 3.4159 | 0.99591 | 0.22709 | -0.48606 | 1.55 |
| 5822 | *SHANK2* | 3.4101 | 0.71987 | 0.50868 | -1.5167 | 1.55 |
| 5823 | *CCDC71L* | 3.498 | 1.7515 | -0.61092 | -1.5544 | 1.55 |
| 5824 | *DCTN2* | 3.5954 | 1.3737 | -0.33109 | -0.47527 | 1.55 |
| 5825 | *STK35* | 4.2845 | 0.819 | -0.46562 | -1.9099 | 1.55 |
| 5826 | *C2orf82* | 4.0043 | 1.1681 | -0.53487 | -1.1399 | 1.55 |
| 5827 | *ALX3* | 2.8949 | 2.6305 | -0.88839 | -1.3118 | 1.55 |
| 5828 | *PACRGL* | 2.1479 | 1.6888 | 0.80023 | -1.2568 | 1.55 |
| 5829 | *ZSWIM2* | 4.049 | 1.7504 | -1.1631 | -2.252 | 1.55 |
| 5830 | *STAM2* | 1.8897 | 1.6793 | 1.0664 | 0.8032 | 1.55 |
| 5831 | *ELOVL5* | 3.3703 | 0.66261 | 0.60198 | -0.74913 | 1.54 |
| 5832 | *OR9I1* | 2.9823 | 2.5142 | -0.86546 | -1.4684 | 1.54 |
| 5833 | *PLEKHF2* | 3.4625 | 0.97837 | 0.19015 | -0.78229 | 1.54 |
| 5834 | *HRASLS5* | 3.5868 | 1.2945 | -0.251 | -0.50133 | 1.54 |
| 5835 | *SLC2A1* | 2.3421 | 2.2401 | 0.04765 | -1.2486 | 1.54 |

| | | | | | | |
|---|---|---|---|---|---|---|
| 5836 | *LCAT* | 3.2138 | 0.98643 | 0.42918 | -1.4946 | 1.54 |
| 5837 | *GCKR* | 5.059 | 0.013874 | -0.44353 | -0.93365 | 1.54 |
| 5838 | *CCDC80* | 3.7881 | 1.6181 | -0.77733 | -0.89957 | 1.54 |
| 5839 | *GDPGP1* | 4.2053 | 0.34192 | 0.08118 | -0.43993 | 1.54 |
| 5840 | *EIF5AL1* | 3.6211 | 0.64483 | 0.36198 | -1.4248 | 1.54 |
| 5841 | *SPATS2L* | 1.8668 | 1.8051 | 0.95576 | 0.23888 | 1.54 |
| 5842 | *TRIM10* | 3.4269 | 1.1176 | 0.08246 | -0.4457 | 1.54 |
| 5843 | *IPCEF1* | 4.9465 | 0.71441 | -1.0343 | -1.122 | 1.54 |
| 5844 | *PKDREJ* | 4.72 | 0.088291 | -0.18187 | -1.6636 | 1.54 |
| 5845 | *OR11H12* | 3.3887 | 1.6617 | -0.42401 | -0.56773 | 1.54 |
| 5846 | *ASCL2* | 3.526 | 0.5631 | 0.53629 | -2.0908 | 1.54 |
| 5847 | *ARHGAP6* | 2.2939 | 2.2112 | 0.11908 | -0.16976 | 1.54 |
| 5848 | *ZNF44* | 2.6102 | 1.9211 | 0.09188 | -0.43772 | 1.54 |
| 5849 | *NOXA1* | 2.6486 | 1.116 | 0.85815 | -0.62629 | 1.54 |
| 5850 | *PMM2* | 3.812 | 0.5922 | 0.21848 | -0.76355 | 1.54 |
| 5851 | *ACTR3B* | 3.6542 | 0.97636 | -0.00822 | -0.9169 | 1.54 |
| 5852 | *EEF1D* | 4.6284 | 0.17514 | -0.18158 | -1.3402 | 1.54 |
| 5853 | *GAGE12H* | 4.2959 | 0.53466 | -0.20985 | -0.32243 | 1.54 |
| 5854 | *NKRF* | 2.9307 | 1.3642 | 0.3254 | -0.47577 | 1.54 |
| 5855 | *CCDC61* | 4.3883 | 0.33301 | -0.1025 | -1.6001 | 1.54 |
| 5856 | *FAM135B* | 1.9278 | 1.5983 | 1.0904 | -0.79332 | 1.54 |
| 5857 | *CCDC185* | 2.5884 | 2.1869 | -0.15884 | -0.30145 | 1.54 |
| 5858 | *SLC6A18* | 4.4989 | 0.22384 | -0.10704 | -1.72 | 1.54 |
| 5859 | *IL12RB2* | 2.4797 | 1.6538 | 0.48175 | -1.0089 | 1.54 |
| 5860 | *MYO9A* | 3.6245 | 0.6062 | 0.38442 | -0.98238 | 1.54 |
| 5861 | *MARK4* | 4.1376 | 0.31778 | 0.15924 | -1.0997 | 1.54 |
| 5862 | *ZBTB7B* | 2.5679 | 1.1423 | 0.90406 | -1.6754 | 1.54 |
| 5863 | *VPS37B* | 3.0452 | 1.4874 | 0.08093 | -1.2411 | 1.54 |
| 5864 | *GDPD5* | 3.7603 | 1.4163 | -0.56329 | -2.5264 | 1.54 |
| 5865 | *PROSER1* | 3.4541 | 0.8257 | 0.33331 | -1.077 | 1.54 |
| 5866 | *ENTPD7* | 3.7824 | 1.2152 | -0.38472 | -0.94113 | 1.54 |
| 5867 | *EXT2* | 3.7938 | 0.68584 | 0.13256 | -0.90256 | 1.54 |
| 5868 | *CERCAM* | 3.6898 | 1.6255 | -0.70417 | -1.1126 | 1.54 |
| 5869 | *MILR1* | 3.2357 | 0.79998 | 0.57509 | -0.49704 | 1.54 |
| 5870 | *C4orf46* | 2.6859 | 1.2691 | 0.6556 | -1.3837 | 1.54 |
| 5871 | *VCY* | 2.9361 | 2.1543 | -0.47993 | -2.2014 | 1.54 |
| 5872 | *ACTBL2* | 3.856 | 0.74161 | 0.01275 | -0.94563 | 1.54 |
| 5873 | *IL19* | 3.7822 | 1.5932 | -0.76671 | -1.9219 | 1.54 |
| 5874 | *LRRD1* | 3.9281 | 1.148 | -0.46743 | -1.7683 | 1.54 |
| 5875 | *SOD3* | 2.5966 | 1.3932 | 0.61884 | -1.9041 | 1.54 |
| 5876 | *KIAA1328* | 4.1038 | 0.35651 | 0.14773 | -1.3542 | 1.54 |
| 5877 | *DDHD1* | 2.8401 | 1.4534 | 0.31365 | -1.5052 | 1.54 |
| 5878 | *MRPL32* | 4.4906 | 0.95507 | -0.83862 | -1.2787 | 1.54 |
| 5879 | *SPEG* | 3.9117 | 1.0884 | -0.39318 | -1.3711 | 1.54 |
| 5880 | *MKL1* | 3.5252 | 2.5836 | -1.502 | -1.9814 | 1.54 |
| 5881 | *CLCN1* | 3.3447 | 2.0772 | -0.81549 | -1.34 | 1.54 |

| | | | | | | |
|---|---|---|---|---|---|---|
| 5882 | *BATF2* | 3.8207 | 1.9017 | -1.1169 | -2.3563 | 1.54 |
| 5883 | *NANOS1* | 3.1332 | 1.2943 | 0.17765 | -1.4434 | 1.54 |
| 5884 | *TYROBP* | 3.7918 | 0.57939 | 0.23385 | -2.4394 | 1.54 |
| 5885 | *RLF* | 2.6494 | 1.5099 | 0.44527 | -0.05733 | 1.53 |
| 5886 | *SYDE2* | 4.11 | 0.61764 | -0.12329 | -1.4067 | 1.53 |
| 5887 | *FZD3* | 3.1039 | 2.0844 | -0.58404 | -2.6675 | 1.53 |
| 5888 | *UBAP1L* | 3.2906 | 1.1866 | 0.12627 | -0.85821 | 1.53 |
| 5889 | *TDRD6* | 2.529 | 1.4304 | 0.64397 | -0.95321 | 1.53 |
| 5890 | *CITED1* | 4.4529 | 0.42261 | -0.27291 | -1.9622 | 1.53 |
| 5891 | *C19orf12* | 4.1202 | 1.0431 | -0.56225 | -1.5297 | 1.53 |
| 5892 | *PAGE1* | 2.9723 | 2.3039 | -0.67536 | -1.3254 | 1.53 |
| 5893 | *CTNNA3* | 3.7452 | 1.4723 | -0.61675 | -1.8157 | 1.53 |
| 5894 | *CCDC153* | 2.0276 | 1.8785 | 0.69402 | 0.22001 | 1.53 |
| 5895 | *PPP3CC* | 5.594 | -0.39082 | -0.60371 | -0.68668 | 1.53 |
| 5896 | *CXCR1* | 3.9888 | 1.0612 | -0.45103 | -2.2664 | 1.53 |
| 5897 | *ZNF471* | 3.1364 | 1.0111 | 0.45096 | -0.62017 | 1.53 |
| 5898 | *FUT2* | 3.9039 | 0.53655 | 0.15769 | -0.90922 | 1.53 |
| 5899 | *AGO2* | 2.4425 | 1.1331 | 1.0212 | 0.063662 | 1.53 |
| 5900 | *SLC16A8* | 3.0295 | 1.5519 | 0.01539 | -1.1122 | 1.53 |
| 5901 | *OSBPL5* | 5.2216 | 0.37684 | -1.0019 | -1.9624 | 1.53 |
| 5902 | *LECT2* | 3.1098 | 0.8546 | 0.63147 | -2.3696 | 1.53 |
| 5903 | *PSTPIP1* | 3.7016 | 0.70846 | 0.18579 | -0.84136 | 1.53 |
| 5904 | *NEUROD4* | 5.1602 | -0.10716 | -0.45754 | -1.2837 | 1.53 |
| 5905 | *APOA2* | 3.6674 | 0.50231 | 0.42443 | -1.4715 | 1.53 |
| 5906 | *CHST6* | 2.3783 | 1.5092 | 0.70661 | -1.95 | 1.53 |
| 5907 | *DAND5* | 4.5555 | 0.074813 | -0.03668 | -0.59622 | 1.53 |
| 5908 | *BMP4* | 3.2461 | 2.7347 | -1.3875 | -1.9622 | 1.53 |
| 5909 | *TXLNG* | 4.8715 | -0.06413 | -0.2143 | -0.62154 | 1.53 |
| 5910 | *DCAF4L1* | 4.5798 | 0.020906 | -0.00832 | -0.46747 | 1.53 |
| 5911 | *ZNF197* | 3.8308 | 0.39905 | 0.36174 | -0.49034 | 1.53 |
| 5912 | *CEACAM20* | 2.4897 | 2.3577 | -0.2562 | -1.3282 | 1.53 |
| 5913 | *LZIC* | 2.9903 | 1.3983 | 0.20104 | -1.2782 | 1.53 |
| 5914 | *TAS2R1* | 3.1445 | 1.9914 | -0.54657 | -1.8996 | 1.53 |
| 5915 | *ARL13A* | 3.8784 | 1.9173 | -1.2079 | -2.2021 | 1.53 |
| 5916 | *PLAC8L1* | 2.9797 | 1.818 | -0.21148 | -1.2442 | 1.53 |
| 5917 | *PRTN3* | 2.5049 | 2.1888 | -0.10769 | -1.1075 | 1.53 |
| 5918 | *EPDR1* | 4.1304 | 0.94623 | -0.49071 | -1.0153 | 1.53 |
| 5919 | *MRVI1* | 3.3643 | 0.72927 | 0.49156 | -1.3416 | 1.53 |
| 5920 | *HIST1H2BH* | 1.6951 | 1.5326 | 1.3569 | 1.0688 | 1.53 |
| 5921 | *PRM2* | 3.5423 | 0.7007 | 0.34108 | -1.2218 | 1.53 |
| 5922 | *FRRS1L* | 2.5485 | 1.5652 | 0.46992 | -0.59259 | 1.53 |
| 5923 | *TRIML1* | 4.1432 | 0.38273 | 0.05725 | -2.8521 | 1.53 |
| 5924 | *N4BP2L2* | 3.6338 | 1.3955 | -0.44661 | -1.6885 | 1.53 |
| 5925 | *EBLN1* | 2.0205 | 1.5253 | 1.0355 | -1.1479 | 1.53 |
| 5926 | *PAX2* | 4.386 | 0.55094 | -0.35604 | -0.93011 | 1.53 |
| 5927 | *OLFML2A* | 3.2199 | 2.5117 | -1.1508 | -1.2277 | 1.53 |

| | | | | | | |
|---|---|---|---|---|---|---|
| 5928 | *PMCH* | 2.6567 | 2.5935 | -0.66948 | -0.88213 | 1.53 |
| 5929 | *ZNF550* | 3.8053 | 1.4855 | -0.71019 | -1.0088 | 1.53 |
| 5930 | *WDR41* | 3.4923 | 0.7883 | 0.29984 | -0.40822 | 1.53 |
| 5931 | *ADAT3* | 1.9487 | 1.424 | 1.2058 | -0.12729 | 1.53 |
| 5932 | *IMPA2* | 4.1061 | 0.53073 | -0.05841 | -1.4684 | 1.53 |
| 5933 | *CBFB* | 5.0406 | -0.06015 | -0.4027 | -1.5031 | 1.53 |
| 5934 | *TXNDC11* | 3.8667 | 0.64291 | 0.0674 | -0.20165 | 1.53 |
| 5935 | *CTAGE8* | 2.6224 | 1.2524 | 0.70097 | -1.661 | 1.53 |
| 5936 | *PREPL* | 3.0065 | 1.0717 | 0.49689 | -0.00832 | 1.53 |
| 5937 | *STX3* | 2.935 | 2.2703 | -0.63092 | -1.9748 | 1.52 |
| 5938 | *C2CD2* | 3.2376 | 0.70899 | 0.62646 | -1.3393 | 1.52 |
| 5939 | *ATG9A* | 3.4146 | 1.826 | -0.6677 | -0.96751 | 1.52 |
| 5940 | *COL4A3* | 2.0194 | 1.721 | 0.83229 | 0.2638 | 1.52 |
| 5941 | *SHCBP1* | 3.3574 | 1.2518 | -0.03705 | -0.20554 | 1.52 |
| 5942 | *HOXB13* | 2.8692 | 1.7362 | -0.03462 | -0.60371 | 1.52 |
| 5943 | *TSNARE1* | 3.1481 | 1.2292 | 0.19313 | -1.3591 | 1.52 |
| 5944 | *SMU1* | 2.574 | 1.9207 | 0.07442 | -1.165 | 1.52 |
| 5945 | *HIVEP3* | 5.2661 | 0.33847 | -1.0364 | -4.9314 | 1.52 |
| 5946 | *TIMM9* | 4.3832 | 0.33837 | -0.15443 | -0.93399 | 1.52 |
| 5947 | *SIRT4* | 3.6235 | 1.1532 | -0.20958 | -1.5598 | 1.52 |
| 5948 | *ZNF415* | 2.3149 | 2.0769 | 0.17276 | -0.64793 | 1.52 |
| 5949 | *PHYHD1* | 3.8558 | 0.89213 | -0.18348 | -1.4185 | 1.52 |
| 5950 | *MCEMP1* | 2.8632 | 1.433 | 0.26822 | -0.96583 | 1.52 |
| 5951 | *OCEL1* | 3.6048 | 0.62865 | 0.33086 | -0.56634 | 1.52 |
| 5952 | *DNAH1* | 2.4965 | 1.9822 | 0.08527 | -0.64857 | 1.52 |
| 5953 | *SCRT1* | 3.4547 | 1.5149 | -0.4057 | -1.0951 | 1.52 |
| 5954 | *FAM65A* | 2.7201 | 1.3358 | 0.50798 | -1.106 | 1.52 |
| 5955 | *ABCG4* | 1.8494 | 1.658 | 1.0558 | 0.53797 | 1.52 |
| 5956 | *BRWD1* | 2.5579 | 1.2436 | 0.76087 | 0.53697 | 1.52 |
| 5957 | *ORMDL2* | 3.9371 | 2.2047 | -1.5805 | -1.6428 | 1.52 |
| 5958 | *ERRFI1* | 4.3803 | 0.15612 | 0.02471 | -0.13006 | 1.52 |
| 5959 | *PKN3* | 2.218 | 1.4518 | 0.89072 | -0.35472 | 1.52 |
| 5960 | *CELA2A* | 3.2579 | 0.69715 | 0.60532 | -2.127 | 1.52 |
| 5961 | *CTSZ* | 3.4639 | 1.1286 | -0.03492 | -1 | 1.52 |
| 5962 | *GUF1* | 3.4191 | 0.84881 | 0.2887 | -1.2746 | 1.52 |
| 5963 | *C17orf78* | 3.3695 | 2.3671 | -1.1808 | -1.2097 | 1.52 |
| 5964 | *SP6* | 3.9007 | 1.0977 | -0.44353 | -0.75351 | 1.52 |
| 5965 | *BHLHE23* | 3.7808 | 1.9684 | -1.1944 | -2.6805 | 1.52 |
| 5966 | *MTL5* | 4.7079 | 0.18841 | -0.34158 | -2.6549 | 1.52 |
| 5967 | *MCIDAS* | 4.9072 | 0.61108 | -0.96367 | -1.0582 | 1.52 |
| 5968 | *ST6GALNAC4* | 4.7679 | 0.053968 | -0.26752 | -1.2872 | 1.52 |
| 5969 | *SFRP1* | 4.0729 | 0.75336 | -0.27244 | -1.3728 | 1.52 |
| 5970 | *SCARF1* | 3.6188 | 0.67328 | 0.2606 | -0.2279 | 1.52 |
| 5971 | *FRMD4B* | 4.6254 | 0.3074 | -0.38059 | -2.0586 | 1.52 |
| 5972 | *PRKAB1* | 2.6134 | 1.6356 | 0.30316 | -0.51929 | 1.52 |
| 5973 | *THOP1* | 4.4139 | 1.0645 | -0.92629 | -2.4018 | 1.52 |

| | | | | | | |
|---|---|---|---|---|---|---|
| 5974 | *LYSMD2* | 2.3279 | 2.2872 | -0.06311 | -0.13638 | 1.52 |
| 5975 | *MTMR3* | 1.7152 | 1.5909 | 1.2457 | -0.36761 | 1.52 |
| 5976 | *OR4K17* | 2.4264 | 2.0995 | 0.0254 | -0.81906 | 1.52 |
| 5977 | *GNRHR* | 4.1587 | 0.22274 | 0.16985 | -1.7177 | 1.52 |
| 5978 | *NAIP* | 2.4662 | 1.146 | 0.93884 | -0.79052 | 1.52 |
| 5979 | *LARP7* | 4.06 | 0.97477 | -0.48538 | -0.74192 | 1.52 |
| 5980 | *ANKRD33* | 3.0466 | 1.0973 | 0.40512 | -0.75395 | 1.52 |
| 5981 | *ATP11A* | 2.8273 | 2.6453 | -0.92375 | -1.0153 | 1.52 |
| 5982 | *KCTD3* | 4.313 | 0.41346 | -0.17766 | -0.43231 | 1.52 |
| 5983 | *RABGAP1* | 4.328 | 0.42665 | -0.20648 | -1.2456 | 1.52 |
| 5984 | *CLN3* | 2.9133 | 2.4063 | -0.77162 | -0.78314 | 1.52 |
| 5985 | *PPARGC1B* | 3.423 | 0.60842 | 0.51629 | -1.4797 | 1.52 |
| 5986 | *LAPTM4B* | 4.7814 | 0.78538 | -1.0198 | -2.2676 | 1.52 |
| 5987 | *KRT7* | 3.9132 | 0.70572 | -0.07219 | -0.78229 | 1.52 |
| 5988 | *ZNF431* | 4.6196 | -0.00232 | -0.07212 | -0.55284 | 1.52 |
| 5989 | *HOMER1* | 4.1393 | 1.8396 | -1.4338 | -2.1369 | 1.52 |
| 5990 | *FAM159B* | 2.3968 | 2.0004 | 0.14783 | -0.28778 | 1.52 |
| 5991 | *ADAM11* | 2.9599 | 2.4095 | -0.82439 | -1.407 | 1.52 |
| 5992 | *ADIPOR1* | 3.4525 | 1.3155 | -0.22313 | -0.92278 | 1.51 |
| 5993 | *ZNF101* | 3.6773 | 0.62206 | 0.24498 | -3.1038 | 1.51 |
| 5994 | *MRGPRE* | 4.0011 | 1.2929 | -0.74979 | -0.8846 | 1.51 |
| 5995 | *CCNK* | 3.6349 | 1.0787 | -0.16951 | -0.42074 | 1.51 |
| 5996 | *PRAMEF9* | 4.9456 | 0.51906 | -0.92086 | -1.3955 | 1.51 |
| 5997 | *ENC1* | 3.5752 | 1.7465 | -0.77828 | -1.3039 | 1.51 |
| 5998 | *KRT80* | 3.9271 | 0.3755 | 0.2405 | 0.080834 | 1.51 |
| 5999 | *SYNE3* | 2.7393 | 1.147 | 0.65655 | 0.35036 | 1.51 |
| 6000 | *PES1* | 2.7594 | 2.2033 | -0.42163 | -1.5291 | 1.51 |
| 6001 | *PRLHR* | 2.6415 | 1.9487 | -0.04919 | -0.74141 | 1.51 |
| 6002 | *RBFA* | 2.5661 | 1.0827 | 0.89141 | -1.3542 | 1.51 |
| 6003 | *STOX1* | 4.7426 | 0.52683 | -0.72985 | -0.83278 | 1.51 |
| 6004 | *OR9A4* | 2.5544 | 2.3891 | -0.40581 | -0.41317 | 1.51 |
| 6005 | *NPHP4* | 3.7646 | 0.59379 | 0.17837 | -0.82326 | 1.51 |
| 6006 | *ZNF708* | 4.4993 | 0.71788 | -0.68153 | -1.5663 | 1.51 |
| 6007 | *LRRC8D* | 2.3883 | 1.3338 | 0.81294 | 0.57088 | 1.51 |
| 6008 | *DECR2* | 4.3226 | 0.33517 | -0.12414 | -1.5563 | 1.51 |
| 6009 | *SPATS2* | 3.4659 | 0.67506 | 0.39196 | -0.38144 | 1.51 |
| 6010 | *PRAMEF14* | 4.8154 | 0.19463 | -0.47739 | -0.56922 | 1.51 |
| 6011 | *GAS2L3* | 3.2888 | 0.63413 | 0.60935 | 0.51748 | 1.51 |
| 6012 | *XPO4* | 3.9914 | 1.7358 | -1.1959 | -2.1213 | 1.51 |
| 6013 | *MPZL3* | 3.5849 | 0.73531 | 0.21027 | -1.4025 | 1.51 |
| 6014 | *OR5D13* | 2.3371 | 1.4199 | 0.77348 | -0.61114 | 1.51 |
| 6015 | *RNF111* | 3.9507 | 0.35451 | 0.22523 | -0.94493 | 1.51 |
| 6016 | *GRASP* | 2.4023 | 1.9392 | 0.18854 | -1.8354 | 1.51 |
| 6017 | *MROH6* | 3.3179 | 1.651 | -0.43886 | -2.2537 | 1.51 |
| 6018 | *HOXD10* | 2.7578 | 2.4709 | -0.69913 | -1.8154 | 1.51 |
| 6019 | *FBXW10* | 2.9465 | 1.5968 | -0.0139 | -0.95322 | 1.51 |

| | | | | | | |
|---|---|---|---|---|---|---|
| 6020 | *XDH* | 2.9171 | 2.1298 | -0.51785 | -0.94472 | 1.51 |
| 6021 | *LOC100133267* | 2.3184 | 1.3359 | 0.87369 | -0.78444 | 1.51 |
| 6022 | *SUV39H1* | 3.2026 | 0.69539 | 0.62925 | -0.00805 | 1.51 |
| 6023 | *ARID5B* | 4.3595 | 0.17547 | -0.00826 | -2.3424 | 1.51 |
| 6024 | *CADM3* | 4.7863 | 0.5069 | -0.76716 | -1.3957 | 1.51 |
| 6025 | *C14orf119* | 4.3286 | 1.096 | -0.90104 | -1.1366 | 1.51 |
| 6026 | *CKAP4* | 4.5037 | 0.1016 | -0.08183 | -0.42958 | 1.51 |
| 6027 | *PTMA* | 3.6637 | 0.75852 | 0.10111 | -1.5428 | 1.51 |
| 6028 | *C16orf46* | 4.2158 | 0.87192 | -0.56492 | -1.1357 | 1.51 |
| 6029 | *PLEKHA7* | 2.6399 | 1.6958 | 0.18653 | -2.3287 | 1.51 |
| 6030 | *SFTPB* | 3.936 | 1.3886 | -0.80315 | -0.94466 | 1.51 |
| 6031 | *ALPL* | 1.9456 | 1.9449 | 0.63028 | 0.41919 | 1.51 |
| 6032 | *GRAMD1C* | 2.9342 | 0.94755 | 0.6389 | -2.0429 | 1.51 |
| 6033 | *GPBAR1* | 2.3314 | 1.1663 | 1.0221 | 0.49035 | 1.51 |
| 6034 | *ARHGEF28* | 3.2531 | 2.5113 | -1.2455 | -1.8627 | 1.51 |
| 6035 | *PRR23B* | 2.2765 | 1.2286 | 1.0136 | -1.0508 | 1.51 |
| 6036 | *ZNF419* | 3.9939 | 0.32154 | 0.20314 | -0.51 | 1.51 |
| 6037 | *GJC3* | 3.826 | 0.94409 | -0.25163 | -2.1113 | 1.51 |
| 6038 | *TSPYL6* | 3.0224 | 1.373 | 0.1224 | -0.3308 | 1.51 |
| 6039 | *MUC21* | 3.1416 | 2.5911 | -1.2149 | -1.4683 | 1.51 |
| 6040 | *SPAM1* | 3.0725 | 1.409 | 0.03609 | -1.7307 | 1.51 |
| 6041 | *CXorf65* | 2.7934 | 1.1518 | 0.57176 | -2.1055 | 1.51 |
| 6042 | *DDX53* | 3.7364 | 0.55612 | 0.22441 | -1.63 | 1.51 |
| 6043 | *LYPD6* | 3.2738 | 1.6245 | -0.38163 | -0.42945 | 1.51 |
| 6044 | *SERPINA7* | 4.1688 | 1.1027 | -0.75495 | -0.87198 | 1.51 |
| 6045 | *SLC30A1* | 4.1504 | 1.6939 | -1.3279 | -1.479 | 1.51 |
| 6046 | *PTGER1* | 3.3381 | 1.3602 | -0.18304 | -0.5728 | 1.51 |
| 6047 | *RXFP4* | 2.9624 | 0.8259 | 0.72657 | -0.8424 | 1.50 |
| 6048 | *RPTOR* | 4.3111 | 0.44004 | -0.23628 | -1.9693 | 1.50 |
| 6049 | *SSH1* | 3.084 | 0.94562 | 0.4833 | -0.64186 | 1.50 |
| 6050 | *SLC22A31* | 3.0526 | 1.6177 | -0.15831 | -1.4104 | 1.50 |
| 6051 | *ADSS* | 4.0261 | 1.3752 | -0.8903 | -1.027 | 1.50 |
| 6052 | *GRN* | 4.1367 | 0.60721 | -0.23497 | -0.54246 | 1.50 |
| 6053 | *IFFO2* | 2.5388 | 1.0251 | 0.94391 | -0.3486 | 1.50 |
| 6054 | *CCL5* | 2.9119 | 2.1257 | -0.53045 | -0.70406 | 1.50 |
| 6055 | *KCNG1* | 3.7007 | 1.4517 | -0.64596 | -1.9746 | 1.50 |
| 6056 | *CCDC47* | 3.6355 | 0.46797 | 0.40295 | -0.84337 | 1.50 |
| 6057 | *GPR161* | 2.3735 | 1.3676 | 0.76501 | -1.4527 | 1.50 |
| 6058 | *MAGEA8* | 4.5555 | 1.0346 | -1.084 | -1.8566 | 1.50 |
| 6059 | *FAM126A* | 2.2056 | 1.1674 | 1.1325 | 1.1166 | 1.50 |
| 6060 | *STK25* | 3.4226 | 1.2938 | -0.21216 | -2.012 | 1.50 |
| 6061 | *DNAH17* | 4.0278 | 1.1267 | -0.65113 | -1.4996 | 1.50 |
| 6062 | *CMAS* | 2.5616 | 2.048 | -0.10664 | -0.17658 | 1.50 |
| 6063 | *KRTAP3-2* | 3.5781 | 0.79423 | 0.12988 | -0.9977 | 1.50 |
| 6064 | *PIP4K2C* | 5.1316 | -0.27291 | -0.35694 | -2.3534 | 1.50 |
| 6065 | *OR10X1* | 3.6514 | 0.45979 | 0.39048 | -0.29227 | 1.50 |

| | | | | | | |
|---|---|---|---|---|---|---|
| 6066 | *LRRIQ4* | 1.9428 | 1.2992 | 1.2594 | -0.03448 | 1.50 |
| 6067 | *TARSL2* | 4.313 | 1.9401 | -1.7526 | -1.7587 | 1.50 |
| 6068 | *METTL15* | 2.5931 | 2.5545 | -0.64806 | -1.9748 | 1.50 |
| 6069 | *PIK3CA* | 5.1511 | 0.91449 | -1.5663 | -2.7705 | 1.50 |
| 6070 | *MNDA* | 4.021 | 0.33537 | 0.14261 | -0.68565 | 1.50 |
| 6071 | *PFKL* | 5.0933 | 0.18976 | -0.7847 | -2.0838 | 1.50 |
| 6072 | *ALDH9A1* | 3.0779 | 1.1166 | 0.30364 | -1.2802 | 1.50 |
| 6073 | *OTP* | 3.0775 | 1.0379 | 0.38094 | -0.20942 | 1.50 |
| 6074 | *MMP23B* | 2.7268 | 2.2093 | -0.44094 | -0.49563 | 1.50 |
| 6075 | *PATL1* | 2.7601 | 2.09 | -0.35585 | -1.3627 | 1.50 |
| 6076 | *BMP6* | 4.1649 | 0.22609 | 0.10206 | -0.54847 | 1.50 |
| 6077 | *SLC4A10* | 5.2543 | 0.50009 | -1.2623 | -2.0553 | 1.50 |
| 6078 | *PTPN5* | 2.8205 | 1.2664 | 0.40464 | -0.25609 | 1.50 |
| 6079 | *DAZAP1* | 2.1484 | 1.9032 | 0.43965 | -1.8112 | 1.50 |
| 6080 | *MPC1* | 5.264 | -0.37343 | -0.39967 | -0.79779 | 1.50 |
| 6081 | *CNPY3* | 4.7688 | 0.015765 | -0.29442 | -1.2272 | 1.50 |
| 6082 | *SNRNP25* | 3.7973 | 0.58593 | 0.10647 | -1.0261 | 1.50 |
| 6083 | *ARMS2* | 3.1266 | 2.0917 | -0.7288 | -0.93696 | 1.50 |
| 6084 | *ALOXE3* | 2.4496 | 1.9466 | 0.08994 | -0.47293 | 1.50 |
| 6085 | *PTPN13* | 3.7827 | 1.184 | -0.48156 | -1.0421 | 1.50 |
| 6086 | *UGT2A2* | 2.8559 | 1.3673 | 0.26163 | -0.45012 | 1.49 |
| 6087 | *SSH3* | 4.1359 | 0.36987 | -0.0211 | -1.7882 | 1.49 |
| 6088 | *TAP2* | 2.6417 | 2.2591 | -0.41621 | -0.47948 | 1.49 |
| 6089 | *VRK3* | 3.2329 | 1.2206 | 0.03043 | -0.28209 | 1.49 |
| 6090 | *PCMTD1* | 2.9014 | 2.7785 | -1.1962 | -1.5077 | 1.49 |
| 6091 | *VWA2* | 4.9662 | 0.42941 | -0.91296 | -1.3627 | 1.49 |
| 6092 | *GPX8* | 3.3379 | 0.62661 | 0.51793 | -0.65839 | 1.49 |
| 6093 | *OR4S1* | 2.3399 | 1.411 | 0.73114 | 0.28875 | 1.49 |
| 6094 | *FABP9* | 5.1415 | 0.18203 | -0.84158 | -0.94954 | 1.49 |
| 6095 | *RNF168* | 2.7381 | 2.2969 | -0.55311 | -0.63312 | 1.49 |
| 6096 | *FOXE3* | 2.4348 | 2.2619 | -0.21499 | -0.92451 | 1.49 |
| 6097 | *NUDT19* | 4.2982 | 0.65015 | -0.4674 | -2.3381 | 1.49 |
| 6098 | *CNTN5* | 3.9499 | 0.7408 | -0.2098 | -1.8537 | 1.49 |
| 6099 | *VDR* | 3.1125 | 0.73902 | 0.62915 | -1.536 | 1.49 |
| 6100 | *COL1A1* | 2.7914 | 1.3797 | 0.30903 | 0.057237 | 1.49 |
| 6101 | *RIN3* | 2.5329 | 1.8584 | 0.08825 | -0.86123 | 1.49 |
| 6102 | *OIT3* | 4.7015 | 0.5272 | -0.75023 | -0.84248 | 1.49 |
| 6103 | *TLR2* | 3.4431 | 1.2468 | -0.21148 | -1.3751 | 1.49 |
| 6104 | *SCGB1A1* | 1.8439 | 1.5147 | 1.1197 | -0.08362 | 1.49 |
| 6105 | *SHISA9* | 3.1246 | 0.91197 | 0.43954 | 0.36586 | 1.49 |
| 6106 | *SYN3* | 4.7186 | -0.04085 | -0.20202 | -1.4688 | 1.49 |
| 6107 | *SLC5A8* | 3.4673 | 0.65594 | 0.35194 | -0.56815 | 1.49 |
| 6108 | *HIST1H2AA* | 2.7916 | 1.6509 | 0.03259 | -0.02811 | 1.49 |
| 6109 | *GBE1* | 2.8806 | 1.2086 | 0.38377 | -0.67943 | 1.49 |
| 6110 | *KLK1* | 4.3843 | 0.14115 | -0.05348 | -0.94493 | 1.49 |
| 6111 | *DDX24* | 1.7368 | 1.673 | 1.0602 | 0.049989 | 1.49 |

| | | | | | | |
|---|---|---|---|---|---|---|
| 6112 | *MMP20* | 3.6632 | 2.1098 | -1.3031 | -1.5886 | 1.49 |
| 6113 | *SPRY3* | 3.0299 | 1.607 | -0.16774 | -1.2015 | 1.49 |
| 6114 | *TMF1* | 4.0798 | 0.52357 | -0.13436 | -0.50419 | 1.49 |
| 6115 | *DCHS1* | 4.2969 | 0.78607 | -0.6141 | -2.4699 | 1.49 |
| 6116 | *TSN* | 4.1635 | 0.99573 | -0.69112 | -1.5821 | 1.49 |
| 6117 | *EBP* | 2.927 | 1.0233 | 0.51758 | 0.050728 | 1.49 |
| 6118 | *ABHD11* | 1.771 | 1.5819 | 1.1137 | 0.58526 | 1.49 |
| 6119 | *TRAPPC12* | 4.5346 | 0.83393 | -0.9034 | -1.3327 | 1.49 |
| 6120 | *SPIN2A* | 3.5462 | 0.70943 | 0.20802 | 0.029689 | 1.49 |
| 6121 | *ZNF518A* | 3.0322 | 1.4607 | -0.0298 | -0.61281 | 1.49 |
| 6122 | *MAGEE1* | 2.5162 | 1.2983 | 0.64807 | -0.86772 | 1.49 |
| 6123 | *PSAPL1* | 2.9356 | 1.7807 | -0.25444 | -0.60554 | 1.49 |
| 6124 | *RGS13* | 3.1604 | 0.82505 | 0.47547 | -2.223 | 1.49 |
| 6125 | *ATP13A5* | 2.2921 | 1.4682 | 0.70052 | -0.35473 | 1.49 |
| 6126 | *CCDC34* | 5.6109 | -0.49902 | -0.65132 | -1.1145 | 1.49 |
| 6127 | *SLC5A4* | 2.5195 | 2.2225 | -0.28162 | -0.64967 | 1.49 |
| 6128 | *FGD6* | 3.7007 | 0.65441 | 0.1049 | -1.5568 | 1.49 |
| 6129 | *ZNF551* | 3.1558 | 1.2925 | 0.01116 | -0.33081 | 1.49 |
| 6130 | *OR4F21* | 4.1003 | 0.80221 | -0.44353 | -1.7493 | 1.49 |
| 6131 | *TXNDC2* | 2.8287 | 1.2108 | 0.41933 | -1.5715 | 1.49 |
| 6132 | *CTAGE5* | 3.575 | 1.6065 | -0.72283 | -0.84816 | 1.49 |
| 6133 | *ATP7B* | 3.9789 | 1.6178 | -1.1383 | -1.748 | 1.49 |
| 6134 | *SLC23A3* | 4.2258 | 0.26461 | -0.03283 | -1.2697 | 1.49 |
| 6135 | *ZIC5* | 2.9845 | 0.97705 | 0.4958 | -0.30608 | 1.49 |
| 6136 | *AHCTF1* | 2.9298 | 1.1257 | 0.40166 | -0.96857 | 1.49 |
| 6137 | *RNF14* | 2.8044 | 1.2124 | 0.43983 | 0.3014 | 1.49 |
| 6138 | *TSPAN2* | 5.4831 | 0.43639 | -1.463 | -2.8398 | 1.49 |
| 6139 | *MLF2* | 3.3245 | 0.68213 | 0.44898 | -0.06794 | 1.49 |
| 6140 | *UMODL1* | 4.8466 | 0.18614 | -0.57742 | -0.67482 | 1.49 |
| 6141 | *CCDC92* | 4.7276 | 0.16555 | -0.43886 | -0.50087 | 1.48 |
| 6142 | *CDK10* | 4.5277 | 1.107 | -1.181 | -1.5707 | 1.48 |
| 6143 | *FAM150B* | 4.2722 | 0.14912 | 0.03238 | -0.91127 | 1.48 |
| 6144 | *GTF2A1* | 3.2611 | 1.6641 | -0.47192 | -1.16 | 1.48 |
| 6145 | *TMCC1* | 4.2116 | 0.31701 | -0.07545 | -2.0436 | 1.48 |
| 6146 | *GSTA5* | 2.6793 | 1.4538 | 0.31936 | -0.03366 | 1.48 |
| 6147 | *SRSF10* | 3.9363 | 0.40972 | 0.1062 | -1.5663 | 1.48 |
| 6148 | *CDCA8* | 2.5173 | 1.6419 | 0.29244 | 0.17431 | 1.48 |
| 6149 | *FAM81A* | 3.632 | 0.68632 | 0.13245 | -0.47227 | 1.48 |
| 6150 | *GRM6* | 4.5239 | 0.96553 | -1.0402 | -1.6158 | 1.48 |
| 6151 | *ACTG1* | 3.543 | 1.3579 | -0.4521 | -0.8934 | 1.48 |
| 6152 | *FAM127C* | 4.0678 | 0.25825 | 0.12253 | -0.35735 | 1.48 |
| 6153 | *CCL14* | 3.0437 | 1.3183 | 0.08633 | -1.135 | 1.48 |
| 6154 | *CCDC28A* | 3.4171 | 1.6207 | -0.58949 | -0.81548 | 1.48 |
| 6155 | *APOBEC3H* | 1.9886 | 1.8635 | 0.59553 | -0.84872 | 1.48 |
| 6156 | *ASTN1* | 2.9474 | 0.80389 | 0.6956 | -0.87504 | 1.48 |
| 6157 | *FGFBP2* | 4.0468 | 1.3472 | -0.94739 | -2.3496 | 1.48 |

| | | | | | | |
|---|---|---|---|---|---|---|
| 6158 | *RFWD3* | 3.6466 | 0.82493 | -0.02638 | -2.5707 | 1.48 |
| 6159 | *TMEM9* | 2.9824 | 0.76114 | 0.7014 | -2.01 | 1.48 |
| 6160 | *LRRC6* | 2.5138 | 1.2294 | 0.70073 | -0.41995 | 1.48 |
| 6161 | *AP1B1* | 3.8933 | 0.29458 | 0.25604 | -0.97477 | 1.48 |
| 6162 | *ZDHHC20* | 3.0736 | 0.80978 | 0.55854 | -0.46415 | 1.48 |
| 6163 | *FAN1* | 1.6681 | 1.4734 | 1.3001 | -1.1175 | 1.48 |
| 6164 | *DHRS13* | 5.556 | -0.22235 | -0.89219 | -2.0952 | 1.48 |
| 6165 | *CT62* | 3.0891 | 1.8581 | -0.50695 | -3.4184 | 1.48 |
| 6166 | *DLEU7* | 4.5702 | 0.72019 | -0.85059 | -1.4312 | 1.48 |
| 6167 | *MZT2A* | 3.3021 | 0.8495 | 0.2868 | -1.2937 | 1.48 |
| 6168 | *TMEM35* | 3.306 | 1.8619 | -0.72973 | -0.85467 | 1.48 |
| 6169 | *HMX2* | 3.3225 | 0.56133 | 0.55401 | -1.2357 | 1.48 |
| 6170 | *TBC1D20* | 2.7968 | 0.91011 | 0.73079 | -1.8645 | 1.48 |
| 6171 | *OR2A1* | 3.1432 | 0.74407 | 0.54869 | -0.52495 | 1.48 |
| 6172 | *SENP7* | 3.6521 | 0.49803 | 0.28553 | 0.10263 | 1.48 |
| 6173 | *RNASE8* | 2.9434 | 2.0513 | -0.55909 | -0.75665 | 1.48 |
| 6174 | *UBL7* | 3.6276 | 0.48211 | 0.32561 | -0.42137 | 1.48 |
| 6175 | *MRPL2* | 3.3424 | 0.60733 | 0.48467 | -1.7164 | 1.48 |
| 6176 | *OPN1SW* | 2.5202 | 1.9901 | -0.07636 | -1.3047 | 1.48 |
| 6177 | *POLI* | 1.9885 | 1.5518 | 0.89214 | -1.5171 | 1.48 |
| 6178 | *IL23R* | 4.9192 | -0.10195 | -0.38522 | -2.3534 | 1.48 |
| 6179 | *CTGF* | 3.8952 | 0.42771 | 0.1083 | -1.7782 | 1.48 |
| 6180 | *ATOH1* | 5.6004 | 0.25219 | -1.4214 | -1.7434 | 1.48 |
| 6181 | *PCDHA13* | 2.9092 | 1.1127 | 0.40872 | -0.65417 | 1.48 |
| 6182 | *PTPRU* | 5.3662 | 0.25861 | -1.1945 | -1.8046 | 1.48 |
| 6183 | *SMC1B* | 4.5417 | 0.025264 | -0.13683 | -0.53058 | 1.48 |
| 6184 | *WHSC1L1* | 4.2659 | 1.0664 | -0.90281 | -1.135 | 1.48 |
| 6185 | *ADGRL3* | 3.1226 | 0.79887 | 0.50528 | -0.19089 | 1.48 |
| 6186 | *LEP* | 1.6156 | 1.5447 | 1.2651 | -2.1391 | 1.48 |
| 6187 | *SPATA17* | 4.0667 | 1.3675 | -1.01 | -1.0604 | 1.47 |
| 6188 | *TESPA1* | 3.9484 | 1.1794 | -0.70361 | -0.96072 | 1.47 |
| 6189 | *IFNA17* | 3.8343 | 0.58647 | 0.00329 | -0.17991 | 1.47 |
| 6190 | *ADAM22* | 2.9902 | 1.6921 | -0.25859 | -1.3033 | 1.47 |
| 6191 | *AVL9* | 4.2644 | 0.60236 | -0.44353 | -0.82366 | 1.47 |
| 6192 | *LRFN1* | 2.5959 | 1.9514 | -0.12456 | -0.83305 | 1.47 |
| 6193 | *RIMS1* | 2.645 | 1.9146 | -0.13688 | -1.2633 | 1.47 |
| 6194 | *COL9A3* | 2.913 | 2.5135 | -1.0066 | -1.128 | 1.47 |
| 6195 | *SLCO1B3* | 3.183 | 1.2618 | -0.02502 | -2.3452 | 1.47 |
| 6196 | *FAM131C* | 3.0734 | 2.9311 | -1.5848 | -2.1357 | 1.47 |
| 6197 | *HIGD1A* | 1.9265 | 1.368 | 1.1246 | -1.6665 | 1.47 |
| 6198 | *ZNF571* | 3.1776 | 0.7079 | 0.53314 | -2.1796 | 1.47 |
| 6199 | *EDN3* | 2.2426 | 2.1601 | 0.01581 | -0.74282 | 1.47 |
| 6200 | *CCL16* | 3.9235 | 0.81334 | -0.31848 | -2.9244 | 1.47 |
| 6201 | *FAM110C* | 4.6365 | -0.08956 | -0.12896 | -1.498 | 1.47 |
| 6202 | *SCRG1* | 2.4073 | 1.9871 | 0.02267 | -0.4221 | 1.47 |
| 6203 | *PTK2* | 3.5364 | 1.5806 | -0.69995 | -2.1576 | 1.47 |

| | | | | | | |
|---|---|---|---|---|---|---|
| 6204 | *ACTRT1* | 2.4394 | 1.25 | 0.72742 | -2.0116 | 1.47 |
| 6205 | *C14orf105* | 4.1527 | 0.93005 | -0.66616 | -0.90906 | 1.47 |
| 6206 | *SIGLEC1* | 4.8491 | -0.20434 | -0.2283 | -0.76055 | 1.47 |
| 6207 | *NSMCE1* | 3.0433 | 0.97867 | 0.39359 | -1.3402 | 1.47 |
| 6208 | *FTHL17* | 3.5064 | 0.55416 | 0.35463 | -0.63673 | 1.47 |
| 6209 | *HID1* | 3.4408 | 2.1416 | -1.1673 | -1.7285 | 1.47 |
| 6210 | *FCER1G* | 2.2126 | 1.457 | 0.74434 | -0.35694 | 1.47 |
| 6211 | *GPR89A* | 3.8211 | 1.0347 | -0.44236 | -1.9511 | 1.47 |
| 6212 | *IFNA14* | 2.3704 | 1.3138 | 0.72916 | -2.5571 | 1.47 |
| 6213 | *HIST1H2BM* | 4.6961 | 0.45493 | -0.73823 | -1.7445 | 1.47 |
| 6214 | *SLC6A1* | 5.485 | -0.3146 | -0.758 | -1.6494 | 1.47 |
| 6215 | *INPP4A* | 4.4979 | 0.70725 | -0.79316 | -1.6848 | 1.47 |
| 6216 | *MAP4K1* | 3.4368 | 2.4013 | -1.4265 | -1.5337 | 1.47 |
| 6217 | *GAS2L1* | 3.9047 | 0.69628 | -0.18996 | -1.3232 | 1.47 |
| 6218 | *TMEM234* | 3.7894 | 0.86308 | -0.24157 | -1.5502 | 1.47 |
| 6219 | *PDZD9* | 3.1194 | 1.7344 | -0.44353 | -0.64748 | 1.47 |
| 6220 | *FHL5* | 2.9149 | 1.0007 | 0.49436 | -1.2935 | 1.47 |
| 6221 | *PDIA2* | 4.3719 | 0.4232 | -0.38541 | -0.75659 | 1.47 |
| 6222 | *KIAA1191* | 4.4552 | 0.17232 | -0.21796 | -0.33844 | 1.47 |
| 6223 | *UGT2B7* | 3.8132 | 1.2428 | -0.64654 | -1.2957 | 1.47 |
| 6224 | *PLP1* | 3.1629 | 1.206 | 0.04036 | -1.3619 | 1.47 |
| 6225 | *CCDC43* | 3.9197 | 0.73245 | -0.24327 | -1.2192 | 1.47 |
| 6226 | *CAMK2N1* | 3.651 | 0.6471 | 0.11031 | -0.33584 | 1.47 |
| 6227 | *C11orf95* | 3.7704 | 0.8689 | -0.23109 | -1.4356 | 1.47 |
| 6228 | *MMP10* | 2.0421 | 1.9844 | 0.38166 | 0.11185 | 1.47 |
| 6229 | *ARHGEF5* | 2.9164 | 1.089 | 0.40274 | -0.35232 | 1.47 |
| 6230 | *FAM170B* | 3.4409 | 0.49118 | 0.47556 | -1.8856 | 1.47 |
| 6231 | *SLC25A19* | 2.2808 | 1.2413 | 0.88399 | -0.45336 | 1.47 |
| 6232 | *ABHD16A* | 3.0143 | 1.8704 | -0.47965 | -0.70448 | 1.47 |
| 6233 | *CCL28* | 4.1655 | 0.27961 | -0.04015 | -1.4333 | 1.47 |
| 6234 | *NPHS2* | 5.3704 | 0.39367 | -1.3593 | -2.4168 | 1.47 |
| 6235 | *GRK7* | 2.6877 | 2.0384 | -0.32162 | -1.5103 | 1.47 |
| 6236 | *TRAPPC9* | 4.5678 | 0.31026 | -0.47412 | -1.0552 | 1.47 |
| 6237 | *DEGS2* | 2.85 | 2.353 | -0.79987 | -1.8908 | 1.47 |
| 6238 | *DCDC2* | 3.6523 | 0.5676 | 0.18274 | -0.41522 | 1.47 |
| 6239 | *MCMDC2* | 4.0908 | 0.25563 | 0.05615 | -1.1912 | 1.47 |
| 6240 | *HAX1* | 2.8148 | 2.2365 | -0.64902 | -1.2697 | 1.47 |
| 6241 | *CBWD6* | 3.2626 | 1.4385 | -0.29885 | -1.1797 | 1.47 |
| 6242 | *HN1* | 5.2309 | 0.2261 | -1.0549 | -1.296 | 1.47 |
| 6243 | *GALR1* | 3.7027 | 0.37308 | 0.32631 | -3.4188 | 1.47 |
| 6244 | *RGMB* | 2.5809 | 2.5118 | -0.69102 | -1.4685 | 1.47 |
| 6245 | *KCNAB3* | 3.2725 | 1.0666 | 0.06244 | -0.60543 | 1.47 |
| 6246 | *JAG2* | 2.1933 | 1.369 | 0.83917 | -0.50087 | 1.47 |
| 6247 | *PRMT1* | 2.5082 | 1.8413 | 0.05143 | -0.58005 | 1.47 |
| 6248 | *MIB2* | 3.0987 | 0.71068 | 0.59146 | -1.191 | 1.47 |
| 6249 | *SHC1* | 2.0428 | 1.6222 | 0.73576 | 0.42378 | 1.47 |

| | | | | | | |
|---|---|---|---|---|---|---|
| 6250 | *FRMD8* | 4.7501 | 0.38184 | -0.73155 | -2.5414 | 1.47 |
| 6251 | *C10orf91* | 1.9743 | 1.2519 | 1.1737 | -0.30939 | 1.47 |
| 6252 | *PFKFB2* | 2.1596 | 1.3582 | 0.88107 | -1.3127 | 1.47 |
| 6253 | *NKAIN1* | 4.2496 | 0.33703 | -0.18961 | -0.34632 | 1.47 |
| 6254 | *PCDHA8* | 2.9446 | 1.1014 | 0.35085 | -2.2136 | 1.47 |
| 6255 | *TTC39B* | 4.2125 | 1.5056 | -1.3218 | -2.1519 | 1.47 |
| 6256 | *NEFL* | 3.5248 | 0.80808 | 0.06312 | -2.7627 | 1.47 |
| 6257 | *TRIM44* | 4.2421 | 1.1014 | -0.94777 | -3.0961 | 1.47 |
| 6258 | *EPAS1* | 2.92 | 0.86165 | 0.61334 | -1.5635 | 1.46 |
| 6259 | *MGAT5* | 3.6265 | 0.86615 | -0.0981 | -2.669 | 1.46 |
| 6260 | *WBSCR28* | 2.8787 | 1.8635 | -0.34777 | -0.95423 | 1.46 |
| 6261 | *NTPCR* | 2.7174 | 1.895 | -0.218 | -1.4034 | 1.46 |
| 6262 | *ITLN1* | 2.3809 | 1.42 | 0.59348 | -0.77346 | 1.46 |
| 6263 | *ZNF705E* | 4.8817 | 0.68351 | -1.1713 | -1.9748 | 1.46 |
| 6264 | *KIAA0195* | 4.5338 | 0.41626 | -0.55741 | -0.73834 | 1.46 |
| 6265 | *OR10H3* | 3.3495 | 1.0212 | 0.02183 | -0.02738 | 1.46 |
| 6266 | *KLF12* | 3.1647 | 0.94627 | 0.2813 | -1.522 | 1.46 |
| 6267 | *WSCD1* | 3.3965 | 0.63053 | 0.36374 | -0.73708 | 1.46 |
| 6268 | *ID3* | 3.4146 | 0.56254 | 0.41311 | 0.047097 | 1.46 |
| 6269 | *ZNF805* | 2.7492 | 0.84092 | 0.80002 | 0.16779 | 1.46 |
| 6270 | *TYW1* | 3.9676 | 0.71917 | -0.29699 | -1.1791 | 1.46 |
| 6271 | *TRIAP1* | 5.6049 | -0.1589 | -1.0567 | -1.1403 | 1.46 |
| 6272 | *PQLC1* | 4.3983 | 0.19404 | -0.20434 | -1.1322 | 1.46 |
| 6273 | *DYDC1* | 3.0189 | 1.5545 | -0.18547 | -0.48413 | 1.46 |
| 6274 | *GMPPA* | 5.2342 | -0.37408 | -0.47303 | -2.4141 | 1.46 |
| 6275 | *IQCF1* | 1.9867 | 1.3632 | 1.0361 | -0.16868 | 1.46 |
| 6276 | *GABRG2* | 4.7341 | 0.50722 | -0.85542 | -1.9408 | 1.46 |
| 6277 | *CALML5* | 3.5457 | 1.0371 | -0.19706 | -0.52949 | 1.46 |
| 6278 | *SMDT1* | 3.4941 | 1.3099 | -0.41831 | -2.9564 | 1.46 |
| 6279 | *SF3B4* | 3.8009 | 0.99599 | -0.41231 | -1.9075 | 1.46 |
| 6280 | *LARP4* | 3.6618 | 0.59723 | 0.12515 | 0.041717 | 1.46 |
| 6281 | *IDO1* | 2.2751 | 2.2624 | -0.15437 | -1.9287 | 1.46 |
| 6282 | *C20orf202* | 3.4063 | 0.77009 | 0.20613 | -1.7691 | 1.46 |
| 6283 | *CAMK2G* | 4.3426 | 0.32946 | -0.28961 | -2.1018 | 1.46 |
| 6284 | *TIMP1* | 3.458 | 1.4318 | -0.50754 | -1.9213 | 1.46 |
| 6285 | *C5orf15* | 3.2181 | 0.6696 | 0.49392 | 0.29764 | 1.46 |
| 6286 | *PRICKLE3* | 3.607 | 0.55103 | 0.22289 | 0.20675 | 1.46 |
| 6287 | *SNAP47* | 4.2032 | 0.89502 | -0.71819 | -0.87395 | 1.46 |
| 6288 | *STEAP1B* | 2.2268 | 1.385 | 0.76817 | -2.0632 | 1.46 |
| 6289 | *CTSA* | 2.9635 | 2.2358 | -0.82081 | -1.5703 | 1.46 |
| 6290 | *NKX2-4* | 4.1511 | 0.37448 | -0.14736 | -0.44353 | 1.46 |
| 6291 | *BCAT1* | 2.9683 | 1.5158 | -0.1062 | -1.3132 | 1.46 |
| 6292 | *LIMS3* | 3.7407 | 0.52529 | 0.11094 | -0.46967 | 1.46 |
| 6293 | *CST11* | 2.9008 | 2.6703 | -1.1945 | -3.0779 | 1.46 |
| 6294 | *MRFAP1* | 4.5277 | 0.31569 | -0.46716 | -1.9727 | 1.46 |
| 6295 | *ASMT* | 3.6269 | 0.61161 | 0.13741 | -0.0748 | 1.46 |

| | | | | | | |
|---|---|---|---|---|---|---|
| 6296 | *YY2* | 4.224 | 0.32725 | -0.17641 | -1.1972 | 1.46 |
| 6297 | *HSD17B1* | 4.0342 | 0.67879 | -0.33895 | -1.0669 | 1.46 |
| 6298 | *TEX13A* | 3.311 | 0.89105 | 0.17146 | -0.72515 | 1.46 |
| 6299 | *PRR30* | 3.9364 | 1.9579 | -1.5212 | -1.5255 | 1.46 |
| 6300 | *HECW1* | 3.0446 | 0.77323 | 0.55522 | -1.5271 | 1.46 |
| 6301 | *MAPRE2* | 2.6235 | 2.0484 | -0.29906 | -1.659 | 1.46 |
| 6302 | *GPR52* | 4.5114 | 0.4092 | -0.54786 | -1.7044 | 1.46 |
| 6303 | *AADACL2* | 3.5087 | 0.53457 | 0.32938 | -1.1836 | 1.46 |
| 6304 | *ZBED2* | 2.3147 | 1.638 | 0.41857 | -0.89974 | 1.46 |
| 6305 | *GJC1* | 4.2514 | 0.446 | -0.32689 | -1.2075 | 1.46 |
| 6306 | *SYNJ1* | 5.2149 | 0.5653 | -1.4104 | -1.5781 | 1.46 |
| 6307 | *FZD2* | 2.6717 | 1.6078 | 0.0903 | -1.632 | 1.46 |
| 6308 | *ATPIF1* | 2.4931 | 1.8217 | 0.05498 | -1.5992 | 1.46 |
| 6309 | *DLG3* | 5.1789 | 0.67817 | -1.4873 | -2.7127 | 1.46 |
| 6310 | *SIRT6* | 3.0031 | 1.0431 | 0.32339 | 0.11495 | 1.46 |
| 6311 | *AKR1D1* | 3.4779 | 1.3365 | -0.44506 | -0.82326 | 1.46 |
| 6312 | *ZNF524* | 2.9252 | 1.5404 | -0.09658 | -0.3644 | 1.46 |
| 6313 | *NSA2* | 4.0303 | 2.3303 | -1.9919 | -2.4312 | 1.46 |
| 6314 | *FAM175B* | 3.2694 | 1.4547 | -0.35641 | -0.46767 | 1.46 |
| 6315 | *TLL1* | 2.9069 | 1.7544 | -0.29571 | -0.48701 | 1.46 |
| 6316 | *ALOX12B* | 3.9621 | 0.2261 | 0.1771 | -0.88189 | 1.46 |
| 6317 | *GPR87* | 3.6607 | 0.49164 | 0.21281 | -1.1907 | 1.46 |
| 6318 | *ARMC4* | 3.3791 | 2.2794 | -1.2935 | -1.747 | 1.46 |
| 6319 | *NGFR* | 2.7533 | 0.97486 | 0.63663 | -1.3773 | 1.45 |
| 6320 | *PGR* | 3.7689 | 0.71281 | -0.11939 | -0.80713 | 1.45 |
| 6321 | *OR52L1* | 3.2205 | 2.187 | -1.0452 | -1.0614 | 1.45 |
| 6322 | *MPP2* | 5.0556 | 0.1106 | -0.8046 | -1.4878 | 1.45 |
| 6323 | *BTN3A2* | 4.2332 | 1.0184 | -0.89059 | -1.0681 | 1.45 |
| 6324 | *SAMD5* | 3.7821 | 1.0033 | -0.42469 | -1.9196 | 1.45 |
| 6325 | *SLC25A47* | 5.1325 | -0.05534 | -0.71717 | -2.7546 | 1.45 |
| 6326 | *SRSF4* | 2.7527 | 1.2231 | 0.38398 | -0.93707 | 1.45 |
| 6327 | *ZNF688* | 4.9996 | -0.03366 | -0.60641 | -1.3016 | 1.45 |
| 6328 | *CDC20* | 3.7783 | 1.0426 | -0.46228 | -0.7647 | 1.45 |
| 6329 | *OR13C8* | 4.1173 | 0.51223 | -0.27102 | -0.61409 | 1.45 |
| 6330 | *LAMC3* | 2.2469 | 1.905 | 0.20556 | -0.17982 | 1.45 |
| 6331 | *ZC3H7B* | 2.1895 | 1.567 | 0.60047 | -0.13255 | 1.45 |
| 6332 | *KLHL11* | 3.2276 | 1.3741 | -0.24474 | -0.8597 | 1.45 |
| 6333 | *WDR19* | 2.9834 | 2.1682 | -0.79525 | -1.9758 | 1.45 |
| 6334 | *ZMYM5* | 3.0992 | 1.1009 | 0.15608 | -0.35805 | 1.45 |
| 6335 | *C5orf38* | 3.5048 | 1.7874 | -0.93621 | -1.7459 | 1.45 |
| 6336 | *PPM1M* | 3.2275 | 0.76799 | 0.36049 | -2.1947 | 1.45 |
| 6337 | *INO80D* | 4.4803 | 0.081586 | -0.20681 | -0.69124 | 1.45 |
| 6338 | *ARFGEF1* | 4.4196 | 0.81748 | -0.88274 | -1.773 | 1.45 |
| 6339 | *NAP1L4* | 3.4438 | 0.47547 | 0.43475 | -1.1734 | 1.45 |
| 6340 | *TCL1A* | 3.5203 | 0.51211 | 0.32125 | -0.70424 | 1.45 |
| 6341 | *PSKH1* | 3.0395 | 0.7993 | 0.51452 | -0.01323 | 1.45 |

| | | | | | | |
|---|---|---|---|---|---|---|
| 6342 | *ADRA2A* | 2.9697 | 1.0048 | 0.37868 | 0.29932 | 1.45 |
| 6343 | *SPATA31D4* | 3.7352 | 0.84707 | -0.22963 | -0.54811 | 1.45 |
| 6344 | *NEK10* | 3.5271 | 1.2851 | -0.45989 | -1.0777 | 1.45 |
| 6345 | *GPHB5* | 4.8361 | -0.24019 | -0.24401 | -1.615 | 1.45 |
| 6346 | *CTRB1* | 3.154 | 2.5088 | -1.3114 | -1.7169 | 1.45 |
| 6347 | *SIK3* | 4.057 | 0.74438 | -0.44998 | -1.0645 | 1.45 |
| 6348 | *FBXO5* | 4.2891 | 0.26733 | -0.20532 | -1.8024 | 1.45 |
| 6349 | *ERMP1* | 2.3039 | 1.7345 | 0.31212 | -0.04465 | 1.45 |
| 6350 | *KIAA0319* | 3.0113 | 0.73122 | 0.6071 | -0.15855 | 1.45 |
| 6351 | *NUTM1* | 2.9706 | 1.1282 | 0.25012 | -0.29396 | 1.45 |
| 6352 | *SEPT11* | 3.4946 | 1.5885 | -0.73483 | -1.257 | 1.45 |
| 6353 | *SMARCD2* | 3.1052 | 2.3698 | -1.127 | -1.8833 | 1.45 |
| 6354 | *STX18* | 4.1847 | 0.58179 | -0.41857 | -0.77624 | 1.45 |
| 6355 | *C4orf32* | 3.2238 | 1.391 | -0.26747 | -1.3125 | 1.45 |
| 6356 | *IZUMO1R* | 2.8259 | 2.1284 | -0.60701 | -1.1559 | 1.45 |
| 6357 | *TMEM211* | 2.7719 | 1.3823 | 0.19282 | -1.3457 | 1.45 |
| 6358 | *LCORL* | 2.2086 | 1.0831 | 1.0549 | -0.76728 | 1.45 |
| 6359 | *TNFRSF6B* | 2.8346 | 2.6257 | -1.1138 | -1.2975 | 1.45 |
| 6360 | *GNAO1* | 4.3104 | 0.51695 | -0.48085 | -0.83621 | 1.45 |
| 6361 | *DESI1* | 3.7324 | 0.68075 | -0.06678 | -0.3361 | 1.45 |
| 6362 | *PRSS16* | 4.0095 | 0.68971 | -0.35445 | -0.51301 | 1.45 |
| 6363 | *CBLB* | 4.0462 | 0.17117 | 0.12714 | -2.1268 | 1.45 |
| 6364 | *UBE2I* | 4.1092 | 0.3755 | -0.14048 | -0.94252 | 1.45 |
| 6365 | *PIRT* | 3.8458 | 0.75122 | -0.25314 | -1.0838 | 1.45 |
| 6366 | *MLH3* | 2.6946 | 0.86859 | 0.78013 | -0.64632 | 1.45 |
| 6367 | *P2RY4* | 2.2787 | 1.667 | 0.39718 | -1.7364 | 1.45 |
| 6368 | *COMP* | 4.0849 | 1.1004 | -0.84261 | -1.4183 | 1.45 |
| 6369 | *KIAA1045* | 4.4397 | 1.0882 | -1.186 | -1.9302 | 1.45 |
| 6370 | *SAA4* | 2.1191 | 1.549 | 0.67352 | -1.7113 | 1.45 |
| 6371 | *PIBF1* | 3.2788 | 0.61791 | 0.44395 | -0.56564 | 1.45 |
| 6372 | *MRPS22* | 4.4107 | 0.34029 | -0.41131 | -0.7873 | 1.45 |
| 6373 | *PKN1* | 2.8422 | 1.5545 | -0.05809 | -0.25339 | 1.45 |
| 6374 | *POLR3GL* | 2.8669 | 0.76148 | 0.70943 | -0.17393 | 1.45 |
| 6375 | *DRD4* | 2.3231 | 1.6024 | 0.41185 | -0.70626 | 1.45 |
| 6376 | *C17orf75* | 3.5481 | 1.1122 | -0.32471 | -0.82174 | 1.45 |
| 6377 | *IQCJ-SCHIP1* | 3.8666 | 0.68028 | -0.21138 | -1.0016 | 1.45 |
| 6378 | *ZKSCAN3* | 3.7126 | 0.97904 | -0.35641 | -1.0323 | 1.45 |
| 6379 | *USHBP1* | 2.5458 | 1.0991 | 0.69008 | -0.24548 | 1.44 |
| 6380 | *CSNK2A1* | 3.8868 | 1.1926 | -0.74471 | -0.87382 | 1.44 |
| 6381 | *ATP2C2* | 2.2265 | 2.0396 | 0.06853 | -1.3847 | 1.44 |
| 6382 | *JKAMP* | 3.3179 | 1.069 | -0.05241 | -0.22114 | 1.44 |
| 6383 | *ARMC1* | 2.398 | 1.0134 | 0.92298 | -1.3988 | 1.44 |
| 6384 | *IER3* | 3.0514 | 1.7127 | -0.42994 | -0.56961 | 1.44 |
| 6385 | *SLC36A1* | 2.7119 | 0.9875 | 0.63384 | -1.3821 | 1.44 |
| 6386 | *DDX50* | 4.2096 | 0.31183 | -0.18898 | -2.1902 | 1.44 |
| 6387 | *ANO3* | 3.0218 | 1.3684 | -0.05776 | -0.95 | 1.44 |

| | | | | | | |
|---|---|---|---|---|---|---|
| 6388 | *INSR* | 2.3975 | 1.9387 | -0.0048 | -1.5956 | 1.44 |
| 6389 | *USP46* | 4.2476 | 0.33116 | -0.2481 | -0.71493 | 1.44 |
| 6390 | *CTNND1* | 3.8178 | 0.54869 | -0.03639 | -2.2517 | 1.44 |
| 6391 | *OR2B2* | 2.8822 | 1.6876 | -0.23983 | -0.46371 | 1.44 |
| 6392 | *FA2H* | 4.296 | 0.33483 | -0.30188 | -1.2805 | 1.44 |
| 6393 | *SBF1* | 2.3646 | 1.074 | 0.88948 | -3.3818 | 1.44 |
| 6394 | *RASGRP2* | 2.5153 | 1.7295 | 0.08307 | -2.0028 | 1.44 |
| 6395 | *TEX38* | 2.6432 | 2.5728 | -0.88877 | -1.5863 | 1.44 |
| 6396 | *GHITM* | 5.2371 | -0.01034 | -0.90001 | -1.1507 | 1.44 |
| 6397 | *TSNAXIP1* | 2.8647 | 1.6748 | -0.21304 | -1.2571 | 1.44 |
| 6398 | *ANKHD1-EIF4EBP*[illegible] | 3.5805 | 0.40753 | 0.3381 | 0.063283 | 1.44 |
| 6399 | *ATG4C* | 4.8072 | 0.040213 | -0.5221 | -1.6452 | 1.44 |
| 6400 | *HDHD3* | 4.5642 | 1.3724 | -1.6115 | -3.1273 | 1.44 |
| 6401 | *ZNF212* | 4.9952 | 0.40466 | -1.0756 | -1.5039 | 1.44 |
| 6402 | *ZNF510* | 2.9171 | 0.79678 | 0.60886 | -1.7232 | 1.44 |
| 6403 | *METTL25* | 3.7212 | 1.9912 | -1.3902 | -2.1334 | 1.44 |
| 6404 | *SLC22A16* | 4.0671 | 0.27872 | -0.02488 | -0.58152 | 1.44 |
| 6405 | *KCNK13* | 2.4621 | 1.4484 | 0.41022 | -0.5591 | 1.44 |
| 6406 | *KLHL20* | 3.9353 | 0.61455 | -0.2293 | -0.46755 | 1.44 |
| 6407 | *ZFP28* | 3.0887 | 1.2589 | -0.0274 | -2.1489 | 1.44 |
| 6408 | *SPR* | 3.9079 | 0.4561 | -0.04469 | -0.16974 | 1.44 |
| 6409 | *TAS2R10* | 2.5184 | 1.5566 | 0.24349 | -1.5888 | 1.44 |
| 6410 | *STARD6* | 3.7096 | 0.86903 | -0.26068 | -2.5354 | 1.44 |
| 6411 | *ACVRL1* | 3.4748 | 1.0393 | -0.19658 | -1.5781 | 1.44 |
| 6412 | *CTTN* | 2.6692 | 2.2901 | -0.64186 | -1.3346 | 1.44 |
| 6413 | *MYO1D* | 3.7466 | 0.53623 | 0.03435 | -0.10838 | 1.44 |
| 6414 | *CLNS1A* | 4.016 | 0.87013 | -0.56924 | -2.1044 | 1.44 |
| 6415 | *ZSCAN20* | 3.7744 | 0.63096 | -0.08868 | -2.1709 | 1.44 |
| 6416 | *CTNNA1* | 5.007 | 1.1996 | -1.8906 | -1.9622 | 1.44 |
| 6417 | *FARSA* | 3.1316 | 1.4419 | -0.25953 | -0.93329 | 1.44 |
| 6418 | *FKBP6* | 2.0323 | 1.7965 | 0.48483 | -0.62596 | 1.44 |
| 6419 | *RBFOX3* | 2.4897 | 0.91386 | 0.90965 | -1.9304 | 1.44 |
| 6420 | *CD82* | 3.3634 | 0.61278 | 0.337 | -0.52458 | 1.44 |
| 6421 | *OSER1* | 5.5545 | -0.39121 | -0.85107 | -1.8154 | 1.44 |
| 6422 | *TRIM49B* | 2.5723 | 1.547 | 0.19251 | -1.0619 | 1.44 |
| 6423 | *DUSP4* | 5.1102 | -0.26208 | -0.53661 | -1.0283 | 1.44 |
| 6424 | *HDAC3* | 3.5431 | 0.98637 | -0.21797 | -1.8575 | 1.44 |
| 6425 | *C1orf198* | 2.5107 | 1.7279 | 0.07189 | -0.28742 | 1.44 |
| 6426 | *IL17RA* | 3.5824 | 1.2017 | -0.47447 | -1.3818 | 1.44 |
| 6427 | *KRTAP19-3* | 3.8705 | 0.22299 | 0.21603 | 0.18322 | 1.44 |
| 6428 | *NIPSNAP3A* | 2.4389 | 1.6008 | 0.26973 | -0.24738 | 1.44 |
| 6429 | *NELL1* | 2.9549 | 1.6887 | -0.33488 | -1.63 | 1.44 |
| 6430 | *C1QL4* | 4.1542 | 0.15426 | 0.00025 | -0.57147 | 1.44 |
| 6431 | *CAMLG* | 2.8384 | 0.76951 | 0.69978 | -0.74572 | 1.44 |
| 6432 | *RDH12* | 1.8312 | 1.3037 | 1.1725 | -2.1848 | 1.44 |
| 6433 | *NCF4* | 4.8094 | 0.88398 | -1.3865 | -1.8642 | 1.44 |

| | | | | | | |
|---|---|---|---|---|---|---|
| 6434 | *NDUFAF7* | 2.3586 | 1.4492 | 0.49894 | -1.3818 | 1.44 |
| 6435 | *RGPD1* | 3.044 | 1.246 | 0.01674 | -0.43292 | 1.44 |
| 6436 | *BDH1* | 2.0299 | 1.7551 | 0.52168 | -1.0528 | 1.44 |
| 6437 | *HSPB3* | 5.1254 | 0.34077 | -1.1595 | -3.2662 | 1.44 |
| 6438 | *ESYT1* | 4.263 | 0.17607 | -0.1331 | -1.8645 | 1.44 |
| 6439 | *DLX5* | 3.1397 | 0.95926 | 0.2061 | -0.71684 | 1.44 |
| 6440 | *ILK* | 5.7 | 0.2124 | -1.6085 | -1.6112 | 1.43 |
| 6441 | *ARMCX3* | 2.3831 | 1.3488 | 0.57086 | 0.45952 | 1.43 |
| 6442 | *CASP1* | 3.7384 | 0.5829 | -0.01883 | -0.47557 | 1.43 |
| 6443 | *GNPAT* | 4.6787 | 0.99391 | -1.371 | -1.4429 | 1.43 |
| 6444 | *SELL* | 3.8172 | 1.2006 | -0.71622 | -2.0384 | 1.43 |
| 6445 | *MACROD2* | 3.6367 | 1.4473 | -0.78325 | -0.8191 | 1.43 |
| 6446 | *SOX6* | 2.4813 | 1.2254 | 0.59281 | -0.07202 | 1.43 |
| 6447 | *PDE9A* | 3.8302 | 0.35573 | 0.1134 | -0.67221 | 1.43 |
| 6448 | *ZNF629* | 2.2278 | 2.0923 | -0.02105 | -1.0003 | 1.43 |
| 6449 | *XPOT* | 3.3823 | 0.6751 | 0.24138 | -0.31191 | 1.43 |
| 6450 | *SOCS4* | 2.7032 | 2.197 | -0.60147 | -1.478 | 1.43 |
| 6451 | *NPC1L1* | 3.5635 | 0.51516 | 0.21977 | -3.1918 | 1.43 |
| 6452 | *RC3H2* | 3.3211 | 0.73927 | 0.23758 | -0.60402 | 1.43 |
| 6453 | *P2RY2* | 4.7844 | 1.0934 | -1.5806 | -2.312 | 1.43 |
| 6454 | *ARHGDIB* | 2.7271 | 1.4338 | 0.13599 | -0.5731 | 1.43 |
| 6455 | *TAF9B* | 1.9063 | 1.4368 | 0.95354 | 0.76184 | 1.43 |
| 6456 | *CACNG8* | 4.743 | -0.00816 | -0.43841 | -2.0351 | 1.43 |
| 6457 | *OSGIN2* | 2.9311 | 1.3587 | 0.00628 | -0.08851 | 1.43 |
| 6458 | *CT47A9* | 4.4617 | 0.066291 | -0.23223 | -0.89429 | 1.43 |
| 6459 | *BCHE* | 2.0659 | 1.623 | 0.60677 | -0.91862 | 1.43 |
| 6460 | *CSAG3* | 4.3679 | 0.56073 | -0.63328 | -1.9891 | 1.43 |
| 6461 | *ELF2* | 2.9966 | 1.5007 | -0.20249 | -1.2577 | 1.43 |
| 6462 | *CLU* | 3.1645 | 0.951 | 0.17837 | -1.5855 | 1.43 |
| 6463 | *BHLHA9* | 2.4865 | 1.4461 | 0.35921 | 0.258 | 1.43 |
| 6464 | *BCOR* | 4.3751 | 0.47286 | -0.55741 | -2.9354 | 1.43 |
| 6465 | *SPOCK2* | 3.0178 | 1.5141 | -0.24164 | -0.28073 | 1.43 |
| 6466 | *ST18* | 3.536 | 3.0486 | -2.2951 | -2.458 | 1.43 |
| 6467 | *RASL11B* | 3.3219 | 0.84494 | 0.12253 | -1.1511 | 1.43 |
| 6468 | *PSG6* | 1.756 | 1.7431 | 0.78918 | -0.47273 | 1.43 |
| 6469 | *RAB8A* | 2.9813 | 0.83525 | 0.4696 | 0.10049 | 1.43 |
| 6470 | *HLCS* | 5.3065 | -0.33116 | -0.68934 | -2.255 | 1.43 |
| 6471 | *CDX1* | 3.1251 | 0.88257 | 0.27826 | -1.6158 | 1.43 |
| 6472 | *SLC26A1* | 3.4066 | 1.0746 | -0.19808 | -1.6711 | 1.43 |
| 6473 | *TRIM68* | 1.9554 | 1.433 | 0.89321 | -1.4747 | 1.43 |
| 6474 | *MOGAT1* | 3.3242 | 2.3313 | -1.3743 | -1.7945 | 1.43 |
| 6475 | *CISD3* | 5.5391 | -0.34533 | -0.91283 | -3.887 | 1.43 |
| 6476 | *HIF1A* | 2.1333 | 1.9632 | 0.18357 | -2.4227 | 1.43 |
| 6477 | *TXLNB* | 5.577 | 0.40726 | -1.7044 | -1.9168 | 1.43 |
| 6478 | *GOLGA4* | 3.084 | 2.0606 | -0.86494 | -0.95545 | 1.43 |
| 6479 | *PROK2* | 4.1176 | 1.289 | -1.127 | -1.3523 | 1.43 |

| | | | | | | |
|---|---|---|---|---|---|---|
| 6480 | *MON1B* | 3.7903 | 0.36827 | 0.12068 | -1.808 | 1.43 |
| 6481 | *ARHGAP18* | 2.5826 | 2.3504 | -0.65378 | -1.0087 | 1.43 |
| 6482 | *ZNF227* | 5.3718 | -0.28704 | -0.80594 | -1.2797 | 1.43 |
| 6483 | *DDX31* | 3.6557 | 0.38148 | 0.2411 | -0.44353 | 1.43 |
| 6484 | *TTC31* | 4.1422 | 0.62538 | -0.48982 | -1.7306 | 1.43 |
| 6485 | *POMZP3* | 3.3189 | 0.53423 | 0.42428 | -1.9567 | 1.43 |
| 6486 | *PUS3* | 4.2089 | 0.18739 | -0.11939 | -1.4483 | 1.43 |
| 6487 | *PYGM* | 3.6437 | 0.50718 | 0.12579 | -1.9945 | 1.43 |
| 6488 | *NUDT9* | 3.6987 | 0.9968 | -0.41987 | -5.3559 | 1.43 |
| 6489 | *CDY2A* | 3.8305 | 0.23536 | 0.20867 | -0.11543 | 1.42 |
| 6490 | *C3orf14* | 3.8952 | 0.21667 | 0.16246 | -0.67874 | 1.42 |
| 6491 | *UGGT1* | 2.6181 | 1.4502 | 0.20594 | -0.73202 | 1.42 |
| 6492 | *MBD3L1* | 2.4086 | 1.7317 | 0.13367 | -0.41307 | 1.42 |
| 6493 | *SPATA31A1* | 4.304 | 0.48005 | -0.5101 | -1.1043 | 1.42 |
| 6494 | *CLCF1* | 2.7969 | 1.1387 | 0.33821 | -1.1872 | 1.42 |
| 6495 | *ATG16L1* | 2.833 | 2.2076 | -0.76857 | -1.1164 | 1.42 |
| 6496 | *BRWD3* | 3.5414 | 0.42209 | 0.30839 | -1.748 | 1.42 |
| 6497 | *SPATA31A3* | 4.6933 | 0.10802 | -0.53086 | -0.76055 | 1.42 |
| 6498 | *NADK2* | 5.121 | -0.30496 | -0.54559 | -2.1463 | 1.42 |
| 6499 | *PLAG1* | 3.3131 | 2.1847 | -1.2277 | -1.6876 | 1.42 |
| 6500 | *MRPL40* | 4.385 | 1.0108 | -1.1263 | -1.8778 | 1.42 |
| 6501 | *HPSE2* | 3.6837 | 0.51031 | 0.07429 | -0.11897 | 1.42 |
| 6502 | *IL13RA2* | 3.4419 | 0.76799 | 0.05813 | 0.023338 | 1.42 |
| 6503 | *AURKAIP1* | 2.4882 | 0.95328 | 0.8264 | -2.3013 | 1.42 |
| 6504 | *MAGEA2B* | 3.0958 | 0.73418 | 0.43717 | -1.613 | 1.42 |
| 6505 | *ATG14* | 4.6431 | 0.4011 | -0.77755 | -0.98951 | 1.42 |
| 6506 | *SATB1* | 3.099 | 0.82348 | 0.34374 | -0.58305 | 1.42 |
| 6507 | *DENND4B* | 2.0659 | 1.9129 | 0.28548 | 0.23049 | 1.42 |
| 6508 | *L3MBTL3* | 4.1878 | 0.16544 | -0.08915 | -0.98719 | 1.42 |
| 6509 | *FBXL21* | 3.737 | 0.89212 | -0.36582 | -1.7802 | 1.42 |
| 6510 | *ZNF221* | 2.328 | 1.5748 | 0.36026 | 0.28019 | 1.42 |
| 6511 | *ANKRD60* | 3.7649 | 1.0616 | -0.56365 | -1.205 | 1.42 |
| 6512 | *LTB4R2* | 3.6395 | 0.40797 | 0.21504 | -0.63002 | 1.42 |
| 6513 | *OR2W3* | 3.2725 | 1.4206 | -0.43077 | -1.3085 | 1.42 |
| 6514 | *ACBD4* | 2.8526 | 1.8665 | -0.45727 | -0.99699 | 1.42 |
| 6515 | *FIBCD1* | 4.4043 | -0.06913 | -0.07367 | -1.5194 | 1.42 |
| 6516 | *FICD* | 3.004 | 0.87216 | 0.3849 | -1.9375 | 1.42 |
| 6517 | *PROL1* | 2.0218 | 1.885 | 0.35392 | -1.2405 | 1.42 |
| 6518 | *LHX3* | 4.2807 | 0.73657 | -0.75896 | -0.83411 | 1.42 |
| 6519 | *CTAGE1* | 4.079 | 1.1775 | -0.99958 | -1.714 | 1.42 |
| 6520 | *CACNG5* | 3.319 | 0.98984 | -0.0525 | -0.96962 | 1.42 |
| 6521 | *DIXDC1* | 1.7727 | 1.296 | 1.1873 | -0.47386 | 1.42 |
| 6522 | *PVRIG* | 2.7029 | 1.3918 | 0.16118 | -1.1226 | 1.42 |
| 6523 | *C19orf18* | 3.3411 | 1.9197 | -1.0051 | -1.9474 | 1.42 |
| 6524 | *NRIP3* | 5.5302 | -0.62306 | -0.65191 | -1.377 | 1.42 |
| 6525 | *UBQLN4* | 4.2545 | 0.019125 | -0.01915 | -1.5304 | 1.42 |

| | | | | | | |
|---|---|---|---|---|---|---|
| 6526 | *SPOCD1* | 4.0573 | 0.67787 | -0.48085 | -1.2629 | 1.42 |
| 6527 | *KLHL9* | 2.3518 | 0.96068 | 0.94137 | -0.49215 | 1.42 |
| 6528 | *ZNF676* | 3.3531 | 1.5137 | -0.61579 | -1.1689 | 1.42 |
| 6529 | *PGBD3* | 1.9967 | 1.4038 | 0.85027 | 0.43945 | 1.42 |
| 6530 | *KIAA1614* | 5.0434 | 0.37485 | -1.1685 | -1.1925 | 1.42 |
| 6531 | *PIGT* | 3.4444 | 0.66709 | 0.13813 | -0.34516 | 1.42 |
| 6532 | *CD93* | 3.4587 | 0.98997 | -0.19915 | -1.4633 | 1.42 |
| 6533 | *IMPA1* | 2.5441 | 2.4377 | -0.73244 | -1.2728 | 1.42 |
| 6534 | *WFS1* | 2.5278 | 2.4373 | -0.71626 | -2.4172 | 1.42 |
| 6535 | *GINS4* | 3.851 | 0.79457 | -0.39711 | -0.53271 | 1.42 |
| 6536 | *FTCD* | 2.3837 | 1.1841 | 0.6799 | -0.87898 | 1.42 |
| 6537 | *CIART* | 4.5341 | -0.1097 | -0.17679 | -0.60083 | 1.42 |
| 6538 | *DLL4* | 3.3087 | 0.93597 | 0.00239 | -1.2716 | 1.42 |
| 6539 | *ZNF705D* | 1.9408 | 1.8021 | 0.50377 | 0.26655 | 1.42 |
| 6540 | *UBE2D4* | 3.3114 | 0.77207 | 0.1628 | 0.068922 | 1.42 |
| 6541 | *KRTAP29-1* | 1.6302 | 1.6052 | 1.0106 | -0.76088 | 1.42 |
| 6542 | *CCNO* | 2.6448 | 1.3984 | 0.20224 | -1.6885 | 1.42 |
| 6543 | *ZNF564* | 4.0614 | 0.11498 | 0.06868 | 0.041724 | 1.42 |
| 6544 | *KCTD21* | 2.7307 | 2.4417 | -0.92769 | -1.4906 | 1.41 |
| 6545 | *RSG1* | 4.0801 | 1.7365 | -1.5719 | -1.9455 | 1.41 |
| 6546 | *ZNF780A* | 2.8808 | 1.4594 | -0.09668 | -1.9284 | 1.41 |
| 6547 | *NACC2* | 2.8233 | 1.2078 | 0.21184 | -2.3154 | 1.41 |
| 6548 | *GPR183* | 2.3865 | 1.9757 | -0.11939 | -0.70137 | 1.41 |
| 6549 | *FAM212A* | 4.6681 | 0.026872 | -0.4533 | -2.8807 | 1.41 |
| 6550 | *PDIA3* | 3.2595 | 1.3077 | -0.32773 | -0.38767 | 1.41 |
| 6551 | *GPX2* | 4.8609 | -0.30378 | -0.31787 | -0.72246 | 1.41 |
| 6552 | *C9orf152* | 4.3124 | 0.068008 | -0.14163 | -0.70506 | 1.41 |
| 6553 | *MTRNR2L1* | 3.0275 | -0.20181 | | | 1.41 |
| 6554 | *LACTBL1* | 2.4874 | 1.2317 | 0.51899 | 0.083394 | 1.41 |
| 6555 | *BFAR* | 2.6168 | 2.0744 | -0.4541 | -2.4633 | 1.41 |
| 6556 | *CD302* | 3.0663 | 1.3494 | -0.17979 | -0.79449 | 1.41 |
| 6557 | *COA5* | 2.9742 | 0.98388 | 0.27774 | -1.338 | 1.41 |
| 6558 | *LRFN2* | 2.7409 | 2.0388 | -0.54417 | -0.93112 | 1.41 |
| 6559 | *TNK1* | 3.737 | 0.31825 | 0.17974 | -1.398 | 1.41 |
| 6560 | *SOWAHD* | 3.769 | 1.5784 | -1.1125 | -1.3033 | 1.41 |
| 6561 | *TMEM150B* | 4.3147 | 0.18955 | -0.27034 | -0.31494 | 1.41 |
| 6562 | *FER1L6* | 2.5308 | 0.85717 | 0.84552 | -0.32728 | 1.41 |
| 6563 | *PGA5* | 3.0644 | 2.2639 | -1.0949 | -3.0564 | 1.41 |
| 6564 | *PAX5* | 3.7314 | 0.77385 | -0.2723 | -1.5294 | 1.41 |
| 6565 | *CPB1* | 3.8572 | 0.38765 | -0.01245 | -0.46228 | 1.41 |
| 6566 | *FBXL16* | 4.228 | 0.062296 | -0.0579 | -0.90991 | 1.41 |
| 6567 | *DPYD* | 3.7682 | 0.90769 | -0.44353 | -1.0153 | 1.41 |
| 6568 | *ZCCHC13* | 2.7739 | 0.76069 | 0.69741 | -0.66767 | 1.41 |
| 6569 | *CAPN11* | 4.463 | 0.31952 | -0.55134 | -1.6289 | 1.41 |
| 6570 | *SULT4A1* | 2.0541 | 1.2815 | 0.89532 | -0.39136 | 1.41 |
| 6571 | *DCAF11* | 2.5035 | 1.4798 | 0.24742 | 0.22935 | 1.41 |

| | | | | | | |
|---|---|---|---|---|---|---|
| 6572 | *SLC36A2* | 3.8006 | 0.42348 | 0.00646 | -0.98938 | 1.41 |
| 6573 | *CLRN3* | 3.3774 | 0.95947 | -0.10633 | -0.9272 | 1.41 |
| 6574 | *DEFB127* | 4.2256 | 0.34738 | -0.34292 | -0.86007 | 1.41 |
| 6575 | *ZDHHC2* | 3.9334 | 0.99324 | -0.69683 | -1.0393 | 1.41 |
| 6576 | *LRP1B* | 3.2726 | 0.60262 | 0.3544 | -1.6131 | 1.41 |
| 6577 | *GH2* | 3.1706 | 1.1915 | -0.13255 | -0.54274 | 1.41 |
| 6578 | *SULT6B1* | 5.182 | 0.10417 | -1.0571 | -1.4485 | 1.41 |
| 6579 | *HIST2H4B* | 2.6521 | 1.2047 | 0.37149 | 0.18586 | 1.41 |
| 6580 | *CLDN6* | 3.679 | 0.49337 | 0.05557 | -1.5369 | 1.41 |
| 6581 | *B2M* | 2.7146 | 0.87211 | 0.64115 | -0.3159 | 1.41 |
| 6582 | *USP15* | 1.7071 | 1.2758 | 1.2447 | 0.11435 | 1.41 |
| 6583 | *ZNF609* | 4.7995 | -0.13528 | -0.43664 | -3.182 | 1.41 |
| 6584 | *FUT1* | 1.8956 | 1.8521 | 0.47987 | -0.00826 | 1.41 |
| 6585 | *SLC22A20* | 2.4221 | 1.6321 | 0.17237 | -1.7018 | 1.41 |
| 6586 | *SLC25A21* | 4.1141 | 0.2034 | -0.09112 | -0.12613 | 1.41 |
| 6587 | *MTF2* | 2.8448 | 0.73877 | 0.64259 | -0.53488 | 1.41 |
| 6588 | *NXPH1* | 2.3126 | 2.2111 | -0.29801 | -0.32699 | 1.41 |
| 6589 | *ERCC3* | 5.0242 | 0.073424 | -0.87256 | -1.202 | 1.41 |
| 6590 | *LRRC28* | 2.1494 | 1.2979 | 0.77774 | -0.79558 | 1.41 |
| 6591 | *TCEA3* | 4.2196 | 0.27524 | -0.26992 | -0.53029 | 1.41 |
| 6592 | *MAP2K1* | 4.1261 | 1.3179 | -1.2198 | -1.6828 | 1.41 |
| 6593 | *MAP7D3* | 2.8304 | 0.87631 | 0.51694 | -0.21491 | 1.41 |
| 6594 | *SRD5A3* | 4.3772 | 0.008092 | -0.16176 | -0.95802 | 1.41 |
| 6595 | *CELF3* | 3.5965 | 1.4578 | -0.83095 | -1.6617 | 1.41 |
| 6596 | *EGR3* | 3.0972 | 1.9731 | -0.84748 | -1.4538 | 1.41 |
| 6597 | *KPNA2* | 4.6021 | 0.14607 | -0.52557 | -1.9295 | 1.41 |
| 6598 | *P2RY10* | 3.6173 | 1.0718 | -0.46698 | -3.2255 | 1.41 |
| 6599 | *DIO2* | 2.3357 | 1.1549 | 0.7309 | -0.20874 | 1.41 |
| 6600 | *SHISA7* | 2.9418 | 1.4075 | -0.12908 | -0.13539 | 1.41 |
| 6601 | *TRIP10* | 2.8578 | 1.4816 | -0.11939 | -1.4824 | 1.41 |
| 6602 | *TMPRSS9* | 2.0645 | 1.2786 | 0.87648 | 0.71892 | 1.41 |
| 6603 | *FAT2* | 2.9445 | 0.99307 | 0.28057 | -0.01923 | 1.41 |
| 6604 | *PTPRB* | 4.7227 | 0.41904 | -0.92375 | -1.5213 | 1.41 |
| 6605 | *CADPS* | 2.17 | 1.7447 | 0.30309 | -1.3978 | 1.41 |
| 6606 | *KHDC3L* | 4.6864 | -0.06132 | -0.4073 | -1.082 | 1.41 |
| 6607 | *AMPD2* | 3.8348 | 0.62779 | -0.24578 | -1.478 | 1.41 |
| 6608 | *ELK4* | 3.2717 | 1.0856 | -0.14323 | -0.17488 | 1.40 |
| 6609 | *OSGEP* | 4.1231 | 0.43117 | -0.34067 | -0.42708 | 1.40 |
| 6610 | *ADAM29* | 3.4225 | 1.1957 | -0.40465 | -0.48156 | 1.40 |
| 6611 | *FAM73A* | 3.6677 | 0.59177 | -0.04593 | -1.5167 | 1.40 |
| 6612 | *WDR53* | 4.4125 | 0.61018 | -0.80946 | -1.3165 | 1.40 |
| 6613 | *UGT2B11* | 2.636 | 1.5469 | 0.03025 | -0.95221 | 1.40 |
| 6614 | *ZGRF1* | 2.6116 | 0.83414 | 0.76682 | -0.85612 | 1.40 |
| 6615 | *UNC5C* | 4.1165 | 0.083386 | 0.01252 | -0.42859 | 1.40 |
| 6616 | *OR10J1* | 3.5685 | 1.7481 | -1.1043 | -2.5403 | 1.40 |
| 6617 | *CYTH3* | 2.3919 | 2.0138 | -0.19357 | -1.9089 | 1.40 |

| | | | | | | |
|---|---|---|---|---|---|---|
| 6618 | *ZCCHC2* | 4.5652 | 0.60331 | -0.95658 | -1.6528 | 1.40 |
| 6619 | *CPA6* | 2.7294 | 2.287 | -0.80517 | -1.864 | 1.40 |
| 6620 | *TXK* | 5.0516 | -0.31508 | -0.52552 | -1.088 | 1.40 |
| 6621 | *AP3B2* | 4.2809 | 0.077878 | -0.14796 | -1.3074 | 1.40 |
| 6622 | *SUSD4* | 3.4664 | 0.52804 | 0.2159 | -1.57 | 1.40 |
| 6623 | *SLC5A12* | 3.2547 | 1.4578 | -0.50265 | -2.3065 | 1.40 |
| 6624 | *GPR153* | 3.4637 | 1.6038 | -0.85771 | -1.8896 | 1.40 |
| 6625 | *PEG10* | 3.4117 | 0.55896 | 0.23877 | -2.217 | 1.40 |
| 6626 | *PDZK1* | 4.6483 | 0.83615 | -1.2755 | -1.8908 | 1.40 |
| 6627 | *RASGRP3* | 2.2335 | 1.0201 | 0.95355 | -1.4501 | 1.40 |
| 6628 | *ACOX1* | 3.5736 | 2.1228 | -1.4894 | -1.8604 | 1.40 |
| 6629 | *SNAI3* | 3.2481 | 2.2123 | -1.2543 | -3.0045 | 1.40 |
| 6630 | *TMEM88* | 4.5121 | 0.85593 | -1.162 | -2.7568 | 1.40 |
| 6631 | *REPIN1* | 2.6885 | 1.1318 | 0.3848 | -0.26018 | 1.40 |
| 6632 | *UGT2A1* | 2.3418 | 1.4898 | 0.37261 | 0.023843 | 1.40 |
| 6633 | *IL2RG* | 3.6888 | 2.0532 | -1.5382 | -1.6564 | 1.40 |
| 6634 | *SLC47A2* | 3.3086 | 0.7297 | 0.16515 | -0.78313 | 1.40 |
| 6635 | *SIGLEC7* | 3.9626 | 0.82252 | -0.58227 | -0.89052 | 1.40 |
| 6636 | *CT83* | 4.6174 | 0.13442 | -0.55002 | -1.6921 | 1.40 |
| 6637 | *SLC25A40* | 3.6138 | 2.018 | -1.4306 | -1.6004 | 1.40 |
| 6638 | *RALGAPA2* | 2.8322 | 1.599 | -0.23042 | -1.6128 | 1.40 |
| 6639 | *RPS4Y1* | 3.3529 | 0.83413 | 0.01369 | -0.83856 | 1.40 |
| 6640 | *PTDSS1* | 3.5973 | 1.1942 | -0.59099 | -1.4775 | 1.40 |
| 6641 | *DPH7* | 3.7368 | 0.61455 | -0.15258 | -2.9706 | 1.40 |
| 6642 | *NWD1* | 3.4337 | 0.89283 | -0.12977 | -1.0413 | 1.40 |
| 6643 | *FAM209A* | 2.7506 | 1.4954 | -0.04931 | -1.1989 | 1.40 |
| 6644 | *ITIH3* | 4.2542 | 0.16594 | -0.22378 | -0.85739 | 1.40 |
| 6645 | *WASF3* | 3.6472 | 0.3453 | 0.20381 | -0.66253 | 1.40 |
| 6646 | *TXNRD2* | 2.9776 | 0.70533 | 0.51104 | -4.3821 | 1.40 |
| 6647 | *BTN2A2* | 2.9323 | 1.3376 | -0.07721 | -0.88507 | 1.40 |
| 6648 | *CST1* | 3.2168 | 1.0314 | -0.05706 | -2.0885 | 1.40 |
| 6649 | *OR5D18* | 2.2872 | 1.5523 | 0.3515 | -0.90084 | 1.40 |
| 6650 | *MAGEA4* | 3.7864 | 0.59057 | -0.18682 | -1.1196 | 1.40 |
| 6651 | *LILRA2* | 3.7587 | 0.39597 | 0.03546 | -0.47964 | 1.40 |
| 6652 | *GALP* | 3.0917 | 1.5656 | -0.46745 | -2.0838 | 1.40 |
| 6653 | *PPP4R1* | 1.9066 | 1.1643 | 1.1182 | -1.1388 | 1.40 |
| 6654 | *DAW1* | 4.4136 | 0.66956 | -0.89424 | -0.92553 | 1.40 |
| 6655 | *STX5* | 3.7728 | 0.79456 | -0.37845 | -1.5671 | 1.40 |
| 6656 | *RPL36A* | 1.4944 | 1.4887 | 1.2058 | -2.017 | 1.40 |
| 6657 | *KIAA1210* | 3.8299 | 0.28562 | 0.073 | -0.24946 | 1.40 |
| 6658 | *PTGER2* | 3.5009 | 0.54097 | 0.1466 | -0.742 | 1.40 |
| 6659 | *IDH3B* | 2.9388 | 1.1471 | 0.10177 | -1.4485 | 1.40 |
| 6660 | *HIST1H3D* | 4.321 | 0.12246 | -0.25589 | -1.5876 | 1.40 |
| 6661 | *TEKT5* | 4.6395 | 1.3199 | -1.7743 | -1.9125 | 1.40 |
| 6662 | *ELANE* | 3.2047 | 0.94323 | 0.03632 | -0.11697 | 1.39 |
| 6663 | *LCE1F* | 3.4249 | 0.47297 | 0.28463 | -1.0699 | 1.39 |

| | | | | | | |
|---|---|---|---|---|---|---|
| 6664 | *STARD3NL* | 1.7301 | 1.2749 | 1.1771 | -0.82656 | 1.39 |
| 6665 | *TMEM246* | 2.5553 | 1.0828 | 0.54388 | -1.9558 | 1.39 |
| 6666 | *BPI* | 4.7843 | 0.10613 | -0.70993 | -1.2426 | 1.39 |
| 6667 | *PPM1F* | 3.1418 | 0.82682 | 0.21136 | -0.9311 | 1.39 |
| 6668 | *FDXR* | 3.5495 | 0.77657 | -0.14689 | -1.7397 | 1.39 |
| 6669 | *PRPS1L1* | 2.9223 | 0.93595 | 0.32086 | -1.3016 | 1.39 |
| 6670 | *RGS17* | 3.1844 | 0.50316 | 0.49141 | -2.1089 | 1.39 |
| 6671 | *DOLPP1* | 3.7691 | 0.62632 | -0.21666 | -0.89401 | 1.39 |
| 6672 | *FBXO36* | 3.7241 | 1.206 | -0.75275 | -0.95577 | 1.39 |
| 6673 | *SLC2A2* | 2.8812 | 1.1879 | 0.1078 | -1.6494 | 1.39 |
| 6674 | *RHEB* | 2.9906 | 2.1603 | -0.97599 | -2.0682 | 1.39 |
| 6675 | *TFE3* | 4.4357 | 0.82008 | -1.0811 | -2.095 | 1.39 |
| 6676 | *XKR6* | 3.4618 | 1.0236 | -0.31166 | -1.4174 | 1.39 |
| 6677 | *PRKACG* | 2.7902 | 1.9788 | -0.59676 | -0.95767 | 1.39 |
| 6678 | *CCDC87* | 3.1529 | 0.59135 | 0.42556 | -0.64956 | 1.39 |
| 6679 | *LRRC47* | 5.1937 | 0.79593 | -1.8202 | -2.0337 | 1.39 |
| 6680 | *SPATA12* | 1.8494 | 1.3526 | 0.96644 | -1.6524 | 1.39 |
| 6681 | *FTSJ3* | 3.7677 | 0.81861 | -0.41812 | -2.1332 | 1.39 |
| 6682 | *HSD17B6* | 3.8063 | 0.63825 | -0.27669 | -1.2262 | 1.39 |
| 6683 | *ISOC2* | 4.0223 | 1.1842 | -1.0388 | -1.2864 | 1.39 |
| 6684 | *APOO* | 3.1962 | 0.57788 | 0.39221 | 0.35272 | 1.39 |
| 6685 | *TSHZ1* | 2.6621 | 0.90894 | 0.59472 | 0.2083 | 1.39 |
| 6686 | *RPS6KB1* | 2.5112 | 0.95708 | 0.69731 | -1.6173 | 1.39 |
| 6687 | *ERLEC1* | 3.3706 | 0.45364 | 0.34102 | -0.45692 | 1.39 |
| 6688 | *SLC27A5* | 2.75 | 1.8325 | -0.41758 | -1.5956 | 1.39 |
| 6689 | *GRAMD1A* | 5.2663 | -0.41285 | -0.68896 | -1.3169 | 1.39 |
| 6690 | *DEF6* | 4.1105 | 0.35673 | -0.30375 | -1.8007 | 1.39 |
| 6691 | *KRTAP4-2* | 3.9765 | 0.64511 | -0.45886 | -0.78048 | 1.39 |
| 6692 | *SNX9* | 3.6913 | 0.87559 | -0.40515 | -1.2567 | 1.39 |
| 6693 | *GAGE12J* | 3.0531 | 1.6812 | -0.57301 | -0.71009 | 1.39 |
| 6694 | *FSCN3* | 2.815 | 2.1577 | -0.81256 | -1.507 | 1.39 |
| 6695 | *MR1* | 4.6778 | 0.057706 | -0.57579 | -1.217 | 1.39 |
| 6696 | *PAN2* | 3.3363 | 0.49068 | 0.33245 | -0.83773 | 1.39 |
| 6697 | *CDK5R1* | 3.5253 | 0.39971 | 0.23428 | -1.691 | 1.39 |
| 6698 | *MROH1* | 2.9241 | 1.1364 | 0.09875 | -0.04564 | 1.39 |
| 6699 | *NEU2* | 3.0322 | 0.60487 | 0.52138 | -0.51689 | 1.39 |
| 6700 | *SERP1* | 1.9041 | 1.4992 | 0.75411 | 0.30481 | 1.39 |
| 6701 | *SLCO4C1* | 3.1088 | 0.77732 | 0.27048 | -1.5715 | 1.39 |
| 6702 | *USP17L28* | 5.6162 | -0.62833 | -0.83138 | -2.9308 | 1.39 |
| 6703 | *PRSS57* | 1.7256 | 1.606 | 0.82447 | -0.89595 | 1.39 |
| 6704 | *UTS2B* | 2.2042 | 1.8991 | 0.05183 | -1.2884 | 1.39 |
| 6705 | *IFNG* | 2.8408 | 2.7438 | -1.4302 | -2.3273 | 1.38 |
| 6706 | *TMEM223* | 4.2979 | 0.095183 | -0.23941 | -0.7423 | 1.38 |
| 6707 | *LCP1* | 3.5813 | 0.51261 | 0.05963 | -0.57542 | 1.38 |
| 6708 | *TPH2* | 2.2094 | 1.5929 | 0.35115 | -2.3602 | 1.38 |
| 6709 | *USP45* | 4.7447 | 1.0215 | -1.6129 | -2.5008 | 1.38 |

| | | | | | | |
|---|---|---|---|---|---|---|
| 6710 | *PANK3* | 3.3561 | 0.69192 | 0.10415 | 0.01524 | 1.38 |
| 6711 | *OR6C65* | 3.9095 | 1.1588 | -0.91647 | -1.4702 | 1.38 |
| 6712 | *MLF1* | 5.4541 | -0.3342 | -0.96857 | -2.1312 | 1.38 |
| 6713 | *D2HGDH* | 3.841 | 0.77621 | -0.46589 | -1.7149 | 1.38 |
| 6714 | *PAK3* | 4.2922 | 0.73444 | -0.87536 | -2.1848 | 1.38 |
| 6715 | *CNGA2* | 3.5061 | 1.7719 | -1.127 | -2.4208 | 1.38 |
| 6716 | *KLRC3* | 1.8217 | 1.4131 | 0.91566 | -1.0142 | 1.38 |
| 6717 | *DIRC1* | 2.2148 | 1.9547 | -0.02137 | -0.88465 | 1.38 |
| 6718 | *TAS2R42* | 3.926 | 0.65032 | -0.42892 | -0.61908 | 1.38 |
| 6719 | *HGSNAT* | 4.1951 | 0.069412 | -0.11762 | -1.0014 | 1.38 |
| 6720 | *PCBP2* | 4.3426 | 0.88958 | -1.0858 | -1.6676 | 1.38 |
| 6721 | *FHAD1* | 4.4564 | 0.056806 | -0.36689 | -0.4978 | 1.38 |
| 6722 | *DDX6* | 3.3524 | 0.66954 | 0.12418 | -0.76253 | 1.38 |
| 6723 | *PQBP1* | 3.4293 | 1.0607 | -0.34416 | -1.0386 | 1.38 |
| 6724 | *ELP5* | 3.8882 | 0.68323 | -0.42585 | -0.79443 | 1.38 |
| 6725 | *HS3ST2* | 4.0281 | 0.69218 | -0.57479 | -1.5552 | 1.38 |
| 6726 | *ZMYND12* | 2.8623 | 1.3678 | -0.08472 | -0.65806 | 1.38 |
| 6727 | *FOXN2* | 4.2073 | 0.38153 | -0.44353 | -1.6469 | 1.38 |
| 6728 | *BABAM1* | 3.6452 | 0.35364 | 0.14603 | -2.3086 | 1.38 |
| 6729 | *NDUFA7* | 2.6636 | 1.2814 | 0.19908 | -1.714 | 1.38 |
| 6730 | *RPS6KA1* | 4.5055 | -0.16749 | -0.19457 | -1.246 | 1.38 |
| 6731 | *FGD5* | 3.7151 | 0.3616 | 0.06576 | -0.44353 | 1.38 |
| 6732 | *FAM57A* | 3.9911 | 0.45577 | -0.30454 | -1.5401 | 1.38 |
| 6733 | *RTKN* | 4.2604 | 0.54286 | -0.66194 | -1.479 | 1.38 |
| 6734 | *ASCL1* | 3.8264 | 0.61569 | -0.30078 | -1.9622 | 1.38 |
| 6735 | *RAMP2* | 2.5007 | 1.222 | 0.41695 | 0.35667 | 1.38 |
| 6736 | *HMBOX1* | 4.2454 | 0.59077 | -0.69655 | -1.5014 | 1.38 |
| 6737 | *LOC100506422* | 2.4528 | 1.1088 | 0.57743 | -2.808 | 1.38 |
| 6738 | *RBMY1A1* | 3.79 | 1.4197 | -1.0707 | -2.7863 | 1.38 |
| 6739 | *KIAA1143* | 2.8469 | 1.1776 | 0.11411 | -1.3285 | 1.38 |
| 6740 | *KLHL4* | 2.6817 | 1.3418 | 0.11502 | -1.7613 | 1.38 |
| 6741 | *NGRN* | 3.4176 | 1.1139 | -0.39313 | -1.2158 | 1.38 |
| 6742 | *TMEM179* | 4.8554 | -0.25927 | -0.45804 | -2.4817 | 1.38 |
| 6743 | *PDLIM7* | 4.4124 | 0.89123 | -1.1659 | -1.3738 | 1.38 |
| 6744 | *SMARCAD1* | 2.4495 | 1.4245 | 0.26293 | -0.81761 | 1.38 |
| 6745 | *COL23A1* | 2.4905 | 1.7183 | -0.07224 | -1.0901 | 1.38 |
| 6746 | *SPANXA2* | 3.2712 | 1.3712 | -0.50589 | -1.2568 | 1.38 |
| 6747 | *CNOT4* | 1.9725 | 1.8286 | 0.33194 | -0.29752 | 1.38 |
| 6748 | *TECR* | 4.9309 | 0.42935 | -1.2284 | -1.4794 | 1.38 |
| 6749 | *RGAG4* | 3.2779 | 1.0364 | -0.18246 | -1.1153 | 1.38 |
| 6750 | *MYH9* | 2.1116 | 1.8469 | 0.17326 | -0.94121 | 1.38 |
| 6751 | *ZC4H2* | 1.9357 | 1.4878 | 0.70814 | -0.98366 | 1.38 |
| 6752 | *PXMP4* | 3.9829 | 0.5581 | -0.40961 | -0.86229 | 1.38 |
| 6753 | *MNT* | 2.3936 | 1.184 | 0.55356 | -0.44353 | 1.38 |
| 6754 | *CNGB3* | 3.0684 | 1.0311 | 0.03091 | -0.28943 | 1.38 |
| 6755 | *SLC15A2* | 3.5466 | 1.1969 | -0.61356 | -0.99585 | 1.38 |

| | | | | | | |
|---|---|---|---|---|---|---|
| 6756 | *LMBRD2* | 3.247 | 0.58375 | 0.29853 | -1.2272 | 1.38 |
| 6757 | *ZMAT3* | 2.566 | 1.3018 | 0.2606 | -0.44687 | 1.38 |
| 6758 | *LYPD8* | 4.6538 | 0.55611 | -1.0819 | -1.161 | 1.38 |
| 6759 | *LBR* | 5.4077 | -0.07056 | -1.2097 | -1.6383 | 1.38 |
| 6760 | *C5* | 5.3263 | -0.17629 | -1.0228 | -1.5152 | 1.38 |
| 6761 | *MEX3C* | 2.7517 | 1.6429 | -0.26743 | -0.64399 | 1.38 |
| 6762 | *ARL4C* | 2.1875 | 1.677 | 0.26174 | -2.1859 | 1.38 |
| 6763 | *TSSK1B* | 3.3228 | 0.75389 | 0.04924 | -0.91913 | 1.38 |
| 6764 | *MICU3* | 4.7149 | 0.001963 | -0.59119 | -2.1706 | 1.38 |
| 6765 | *KLK13* | 3.3421 | 1.0481 | -0.26468 | -0.54179 | 1.38 |
| 6766 | *ALMS1* | 4.8774 | 0.28022 | -1.0328 | -1.8157 | 1.37 |
| 6767 | *MEIS3* | 4.2505 | 0.18624 | -0.31248 | -1.1028 | 1.37 |
| 6768 | *SLC16A5* | 3.1472 | 1.9584 | -0.9815 | -1.2684 | 1.37 |
| 6769 | *PPP1R32* | 4.7171 | 0.30757 | -0.90073 | -2.0609 | 1.37 |
| 6770 | *EFR3A* | 2.8431 | 0.70788 | 0.5727 | 0.46963 | 1.37 |
| 6771 | *AMMECR1L* | 3.6889 | 1.2065 | -0.77236 | -1.3132 | 1.37 |
| 6772 | *RNF114* | 4.4836 | -0.11406 | -0.24665 | -2.1185 | 1.37 |
| 6773 | *ADGRV1* | 4.0587 | 0.25015 | -0.18612 | -2.2379 | 1.37 |
| 6774 | *AIFM2* | 4.3514 | 0.55834 | -0.78718 | -2.4268 | 1.37 |
| 6775 | *CTAGE9* | 5.0133 | -0.39277 | -0.49805 | -2.4933 | 1.37 |
| 6776 | *OR9A2* | 2.6221 | 0.90028 | 0.59986 | -0.29054 | 1.37 |
| 6777 | *SLFN13* | 5.269 | -0.23796 | -0.90922 | -1.0151 | 1.37 |
| 6778 | *TTYH3* | 4.7768 | 0.10845 | -0.76352 | -1.3955 | 1.37 |
| 6779 | *GPSM3* | 2.4971 | 1.3876 | 0.23583 | 0.059093 | 1.37 |
| 6780 | *ATP1B4* | 2.116 | 1.4925 | 0.51024 | -1.0942 | 1.37 |
| 6781 | *LRRN4* | 2.212 | 1.5949 | 0.3109 | -0.45563 | 1.37 |
| 6782 | *ZNF486* | 4.013 | 0.58265 | -0.47821 | -2.8463 | 1.37 |
| 6783 | *SH2D1A* | 1.932 | 1.6426 | 0.54262 | 0.21358 | 1.37 |
| 6784 | *CCNI2* | 4.1384 | 0.65248 | -0.67378 | -0.77409 | 1.37 |
| 6785 | *CCDC82* | 2.669 | 1.0877 | 0.36026 | -1.2246 | 1.37 |
| 6786 | *SPEM1* | 2.4703 | 2.0822 | -0.43572 | -0.45505 | 1.37 |
| 6787 | *TMEM9B* | 3.2873 | 1.2751 | -0.44661 | -0.77418 | 1.37 |
| 6788 | *STXBP1* | 2.9209 | 2.1025 | -0.90797 | -1.2393 | 1.37 |
| 6789 | *PARD3* | 2.5934 | 2.1056 | -0.5836 | -0.73659 | 1.37 |
| 6790 | *OR2T6* | 2.3489 | 1.9779 | -0.21208 | -1.047 | 1.37 |
| 6791 | *ERCC6L2* | 3.3527 | 0.86882 | -0.10743 | -0.71479 | 1.37 |
| 6792 | *ENTHD2* | 5.7288 | -0.77284 | -0.84337 | -1.1758 | 1.37 |
| 6793 | *PLCL1* | 3.3904 | 0.49027 | 0.23168 | -0.59662 | 1.37 |
| 6794 | *NAALAD2* | 4.1051 | 1.937 | -1.9302 | -2.3257 | 1.37 |
| 6795 | *C1orf195* | 4.5766 | -0.1707 | -0.29411 | -1.3702 | 1.37 |
| 6796 | *OTC* | 3.4056 | 1.4073 | -0.70147 | -1.2437 | 1.37 |
| 6797 | *ZNF286A* | 3.9953 | 0.35705 | -0.24148 | -0.26052 | 1.37 |
| 6798 | *RP2* | 3.2285 | 0.73632 | 0.1454 | -1.3249 | 1.37 |
| 6799 | *FAM163B* | 2.9617 | 2.3484 | -1.2007 | -1.7561 | 1.37 |
| 6800 | *C22orf39* | 4.0559 | 1.4597 | -1.4064 | -1.7139 | 1.37 |
| 6801 | *SCAP* | 4.0806 | 0.81451 | -0.78591 | -1.4972 | 1.37 |

| | | | | | | |
|---|---|---|---|---|---|---|
| 6802 | *RNF169* | 1.9286 | 1.8958 | 0.2839 | -1.8181 | 1.37 |
| 6803 | *HIST2H3C* | 2.4714 | 1.0934 | 0.54272 | -1.3805 | 1.37 |
| 6804 | *WFDC11* | 3.661 | 1.1369 | -0.69059 | -1.0232 | 1.37 |
| 6805 | *LPAR2* | 3.5831 | 0.32932 | 0.19435 | -0.92629 | 1.37 |
| 6806 | *FAM3B* | 3.62 | 1.6618 | -1.1759 | -1.3517 | 1.37 |
| 6807 | *SLC22A6* | 1.5758 | 1.536 | 0.99288 | -0.96047 | 1.37 |
| 6808 | *ZRANB2* | 4.498 | -0.19465 | -0.19935 | -1.1717 | 1.37 |
| 6809 | *NLGN4Y* | 3.834 | 1.8066 | -1.5376 | -2.7587 | 1.37 |
| 6810 | *ENTPD5* | 4.635 | -0.25181 | -0.28173 | -0.6981 | 1.37 |
| 6811 | *ASB15* | 3.099 | 0.95209 | 0.04951 | -1.0638 | 1.37 |
| 6812 | *LOC100129083* | 4.5282 | 0.0776 | -0.50548 | -0.61167 | 1.37 |
| 6813 | *WWC1* | 2.1072 | 1.447 | 0.54576 | -0.47739 | 1.37 |
| 6814 | *ANKH* | 2.6715 | 1.6079 | -0.18132 | -0.98459 | 1.37 |
| 6815 | *ALOX15* | 3.6526 | 1.274 | -0.82951 | -0.87719 | 1.37 |
| 6816 | *NDUFS8* | 3.3251 | 1.022 | -0.25018 | -1.72 | 1.37 |
| 6817 | *PSMA4* | 4.1764 | 1.1231 | -1.2027 | -2.8111 | 1.37 |
| 6818 | *TMEM39B* | 2.3377 | 1.285 | 0.47357 | -1.7328 | 1.37 |
| 6819 | *PRDM9* | 3.0288 | 2.404 | -1.3366 | -1.7219 | 1.37 |
| 6820 | *SLC2A12* | 4.6669 | 0.95925 | -1.5309 | -1.9055 | 1.37 |
| 6821 | *LZTS2* | 2.2586 | 1.6872 | 0.14798 | -0.36232 | 1.36 |
| 6822 | *ASTL* | 3.7077 | 1.7269 | -1.3412 | -2.7127 | 1.36 |
| 6823 | *PAEP* | 3.6127 | 0.30283 | 0.17653 | -0.18778 | 1.36 |
| 6824 | *FKBP11* | 2.0848 | 1.2136 | 0.79268 | -1.3513 | 1.36 |
| 6825 | *GRAMD3* | 4.2112 | -0.05026 | -0.07219 | -0.91913 | 1.36 |
| 6826 | *LEPROT* | 3.8843 | 0.19156 | 0.01243 | -0.15684 | 1.36 |
| 6827 | *CDH19* | 3.6067 | 0.83508 | -0.35478 | -0.78229 | 1.36 |
| 6828 | *PPAPDC3* | 3.4941 | 0.37342 | 0.21945 | -2.6806 | 1.36 |
| 6829 | *ZNF521* | 2.7231 | 2.5065 | -1.1427 | -1.3088 | 1.36 |
| 6830 | *SGTB* | 3.0716 | 1.0357 | -0.02062 | -0.83101 | 1.36 |
| 6831 | *KRT222* | 3.033 | 1.8263 | -0.77284 | -2.0423 | 1.36 |
| 6832 | *SEP15* | 3.9385 | 0.63648 | -0.48897 | -2.1496 | 1.36 |
| 6833 | *TM7SF3* | 2.9073 | 0.71886 | 0.45962 | -0.44353 | 1.36 |
| 6834 | *FGL2* | 3.2573 | 2.5518 | -1.7235 | -2.3684 | 1.36 |
| 6835 | *CHN1* | 2.2429 | 1.5572 | 0.28495 | -0.19268 | 1.36 |
| 6836 | *MFI2* | 2.2121 | 1.8308 | 0.04151 | -1.3128 | 1.36 |
| 6837 | *GAS7* | 4.5785 | -0.15891 | -0.33536 | -1.3896 | 1.36 |
| 6838 | *ANAPC2* | 4.8695 | 0.31494 | -1.1006 | -1.4061 | 1.36 |
| 6839 | *TM4SF18* | 3.2716 | 0.56133 | 0.25068 | -1.1303 | 1.36 |
| 6840 | *CD320* | 2.4606 | 1.681 | -0.05837 | -0.86735 | 1.36 |
| 6841 | *NFS1* | 2.816 | 1.525 | -0.25832 | -0.80909 | 1.36 |
| 6842 | *TBC1D10B* | 2.6725 | 2.6271 | -1.2185 | -1.8909 | 1.36 |
| 6843 | *CACNA1A* | 4.7244 | -0.20023 | -0.44353 | -1.6917 | 1.36 |
| 6844 | *ZNF354B* | 4.7169 | 0.68464 | -1.3209 | -1.9734 | 1.36 |
| 6845 | *CENPU* | 3.2722 | 0.58691 | 0.22126 | -1.5375 | 1.36 |
| 6846 | *ARL6IP6* | 4.5851 | 1.5565 | -2.0629 | -2.0954 | 1.36 |
| 6847 | *OST4* | 2.7378 | 1.184 | 0.15653 | -0.13461 | 1.36 |

| | | | | | | |
|---|---|---|---|---|---|---|
| 6848 | *COBL* | 2.535 | 1.8504 | -0.30726 | -0.90003 | 1.36 |
| 6849 | *OR10J3* | 1.7947 | 1.2483 | 1.035 | -0.03157 | 1.36 |
| 6850 | *PXMP2* | 4.0174 | 0.11858 | -0.05851 | -0.17298 | 1.36 |
| 6851 | *TIGD6* | 4.8745 | -0.34816 | -0.44888 | -1.329 | 1.36 |
| 6852 | *PEAR1* | 3.5597 | 0.32001 | 0.19714 | -1.0863 | 1.36 |
| 6853 | *IKZF1* | 3.3666 | 0.36163 | 0.34808 | -3.6506 | 1.36 |
| 6854 | *ARRDC4* | 2.3484 | 1.0177 | 0.7095 | 0.63999 | 1.36 |
| 6855 | *PRKG1* | 4.7415 | 0.37956 | -1.0467 | -1.9678 | 1.36 |
| 6856 | *CD14* | 4.8107 | 0.21086 | -0.94759 | -1.6469 | 1.36 |
| 6857 | *EXTL1* | 2.3661 | 0.87842 | 0.82897 | -1.8591 | 1.36 |
| 6858 | *VNN2* | 3.1593 | 0.81936 | 0.09429 | -2.4832 | 1.36 |
| 6859 | *AKAP14* | 2.5653 | 1.6927 | -0.186 | -0.49925 | 1.36 |
| 6860 | *CD1A* | 2.6607 | 0.81522 | 0.596 | -0.81509 | 1.36 |
| 6861 | *APC* | 3.4671 | 1.0497 | -0.44499 | -0.81793 | 1.36 |
| 6862 | *TMEM229A* | 4.6913 | -0.18175 | -0.43776 | -1.6682 | 1.36 |
| 6863 | *LOC388849* | 4.1392 | 0.71476 | -0.78229 | -1.7378 | 1.36 |
| 6864 | *HIST1H4I* | 3.9222 | 0.50064 | -0.35205 | -1.2304 | 1.36 |
| 6865 | *ANKRD42* | 2.3111 | 1.2808 | 0.47809 | -1.2867 | 1.36 |
| 6866 | *KDM6B* | 3.7509 | 0.28744 | 0.03084 | -1.1212 | 1.36 |
| 6867 | *FPGT-TNNI3K* | 3.763 | 0.79011 | -0.48419 | -0.76934 | 1.36 |
| 6868 | *ADAM10* | 3.3096 | 0.6633 | 0.0951 | -2.0022 | 1.36 |
| 6869 | *FABP1* | 2.5793 | 0.97892 | 0.5094 | 0.12387 | 1.36 |
| 6870 | *FABP6* | 4.0383 | 0.25313 | -0.22451 | -1.5155 | 1.36 |
| 6871 | *STRIP2* | 3.8489 | 0.11518 | 0.10265 | -0.87195 | 1.36 |
| 6872 | *CLDN34* | 2.1949 | 1.3291 | 0.54269 | -2.72 | 1.36 |
| 6873 | *RXRA* | 2.6509 | 1.6964 | -0.28084 | -1.1442 | 1.36 |
| 6874 | *SLC26A3* | 3.1501 | 0.59304 | 0.32211 | -1.4951 | 1.36 |
| 6875 | *SOS1* | 2.8474 | 1.4212 | -0.20601 | -1.729 | 1.35 |
| 6876 | *SYT15* | 4.0229 | 0.39939 | -0.36006 | -1.8462 | 1.35 |
| 6877 | *CLDN10* | 1.8822 | 1.3231 | 0.85686 | -0.54287 | 1.35 |
| 6878 | *ICA1L* | 3.3434 | 0.45747 | 0.26124 | -1.8604 | 1.35 |
| 6879 | *CXorf40B* | 2.4275 | 1.6285 | 0.00488 | -0.86925 | 1.35 |
| 6880 | *TBCEL* | 3.3763 | 0.86685 | -0.18411 | -0.97102 | 1.35 |
| 6881 | *RHCE* | 2.8261 | 1.0307 | 0.20167 | -0.33603 | 1.35 |
| 6882 | *WNT9B* | 1.9185 | 1.8295 | 0.31041 | -0.12128 | 1.35 |
| 6883 | *CHMP4A* | 3.2889 | 1.6194 | -0.85027 | -1.2935 | 1.35 |
| 6884 | *NUDT17* | 2.9366 | 0.87361 | 0.24775 | -1.2497 | 1.35 |
| 6885 | *NCAPG* | 2.8436 | 1.2401 | -0.02598 | -0.20681 | 1.35 |
| 6886 | *C16orf52* | 2.1982 | 1.5227 | 0.33645 | 0.038212 | 1.35 |
| 6887 | *TNFSF12* | 2.9288 | 0.65812 | 0.47012 | -0.17105 | 1.35 |
| 6888 | *C11orf24* | 2.4149 | 1.2329 | 0.40906 | -0.41454 | 1.35 |
| 6889 | *MAGEA1* | 2.5081 | 1.4411 | 0.10729 | -0.04693 | 1.35 |
| 6890 | *MOB3B* | 2.6449 | 0.97036 | 0.44018 | -0.97623 | 1.35 |
| 6891 | *NEU4* | 2.8866 | 0.86502 | 0.3027 | -0.22067 | 1.35 |
| 6892 | *FAS* | 2.1044 | 1.7887 | 0.16109 | -1.611 | 1.35 |
| 6893 | *STXBP5L* | 2.4011 | 2.0534 | -0.40174 | -1.0711 | 1.35 |

| | | | | | | |
|---|---|---|---|---|---|---|
| 6894 | *ZSCAN5B* | 3.5947 | 0.45427 | 0.00369 | -0.58305 | 1.35 |
| 6895 | *MIER1* | 4.1867 | 0.95064 | -1.0853 | -1.7829 | 1.35 |
| 6896 | *ROCK2* | 3.6475 | 0.79669 | -0.39244 | -1.6065 | 1.35 |
| 6897 | *CELF4* | 4.7406 | 0.52243 | -1.2114 | -1.481 | 1.35 |
| 6898 | *ZNF17* | 3.9415 | 0.07586 | 0.03288 | -1.2007 | 1.35 |
| 6899 | *GLTSCR1* | 2.7353 | 2.2392 | -0.92427 | -1.481 | 1.35 |
| 6900 | *IFNA8* | 2.9198 | 1.6174 | -0.48744 | -1.1121 | 1.35 |
| 6901 | *KRTAP2-1* | 1.9562 | 1.9557 | 0.13743 | -1.0436 | 1.35 |
| 6902 | *NR0B1* | 3.9015 | 0.34011 | -0.19331 | -1.856 | 1.35 |
| 6903 | *ZFPL1* | 2.7584 | 2.3416 | -1.0528 | -1.3162 | 1.35 |
| 6904 | *C16orf47* | 1.9675 | 1.1744 | 0.90495 | -0.91793 | 1.35 |
| 6905 | *LMO1* | 3.328 | 0.77097 | -0.05219 | -0.7463 | 1.35 |
| 6906 | *KRTAP13-1* | 3.8858 | 0.46702 | -0.30606 | -2.04 | 1.35 |
| 6907 | *TDO2* | 1.7694 | 1.408 | 0.86936 | -0.59952 | 1.35 |
| 6908 | *CKM* | 4.4956 | -0.18133 | -0.26854 | -0.98864 | 1.35 |
| 6909 | *BTBD6* | 2.4771 | 0.94807 | 0.61992 | -1.8248 | 1.35 |
| 6910 | *KIR3DL1* | 4.9908 | -0.01817 | -0.92764 | -1.6085 | 1.35 |
| 6911 | *EIF1AX* | 3.3572 | 0.40781 | 0.27912 | -1.6656 | 1.35 |
| 6912 | *DCLK1* | 3.267 | 0.45371 | 0.3233 | -1.9378 | 1.35 |
| 6913 | *ATRNL1* | 3.3772 | 0.72184 | -0.05504 | -0.77282 | 1.35 |
| 6914 | *HAO2* | 2.8211 | 1.185 | 0.0373 | -0.9067 | 1.35 |
| 6915 | *CAMKV* | 3.8269 | 0.51973 | -0.30376 | -1.0481 | 1.35 |
| 6916 | *PTCRA* | 2.1025 | 1.0909 | 0.8494 | -0.52726 | 1.35 |
| 6917 | *EP300* | 2.4169 | 0.84925 | 0.77587 | -0.63127 | 1.35 |
| 6918 | *ZNF841* | 3.9569 | 0.73147 | -0.64806 | -0.77018 | 1.35 |
| 6919 | *IFT52* | 1.8902 | 1.8188 | 0.33126 | -0.73695 | 1.35 |
| 6920 | *SMYD3* | 3.8921 | 0.11994 | 0.02759 | -1.25 | 1.35 |
| 6921 | *CTNNA2* | 3.6259 | 0.41678 | -0.00309 | -0.67149 | 1.35 |
| 6922 | *SESN3* | 4.394 | 0.60871 | -0.96407 | -1.6072 | 1.35 |
| 6923 | *AARD* | 3.4306 | 0.37639 | 0.23163 | -1.2573 | 1.35 |
| 6924 | *ARID3C* | 4.8669 | 0.12753 | -0.95708 | -2.2607 | 1.35 |
| 6925 | *PAPPA2* | 3.3689 | 0.66136 | 0.00702 | -0.36654 | 1.35 |
| 6926 | *DEPDC5* | 4.3397 | 0.50617 | -0.81003 | -1.7488 | 1.35 |
| 6927 | *PPP2R1B* | 3.9426 | 0.40532 | -0.31232 | -2.2844 | 1.35 |
| 6928 | *ATE1* | 3.1183 | 0.78356 | 0.13367 | -0.34819 | 1.35 |
| 6929 | *CNNM1* | 4.1469 | 0.12602 | -0.23802 | -1.0112 | 1.34 |
| 6930 | *FFAR4* | 3.9425 | 0.50438 | -0.41207 | -1.183 | 1.34 |
| 6931 | *MYEOV* | 4.151 | 0.54144 | -0.65906 | -1.8297 | 1.34 |
| 6932 | *ZDHHC7* | 3.5363 | 0.41203 | 0.0844 | -1.6239 | 1.34 |
| 6933 | *CTXN3* | 3.3815 | 1.5399 | -0.88876 | -1.9686 | 1.34 |
| 6934 | *FHL2* | 2.6278 | 1.4527 | -0.04847 | -0.93562 | 1.34 |
| 6935 | *MEF2C* | 3.6228 | 0.60429 | -0.19515 | -1.3727 | 1.34 |
| 6936 | *WNT6* | 2.4032 | 2.3762 | -0.748 | -0.78445 | 1.34 |
| 6937 | *RDH8* | 3.7764 | 1.0697 | -0.81528 | -1.3226 | 1.34 |
| 6938 | *B4GALNT4* | 3.143 | 2.5186 | -1.6316 | -2.7705 | 1.34 |
| 6939 | *TMSB15A* | 4.3465 | -0.1313 | -0.18589 | -1.3756 | 1.34 |

| | | | | | | |
|---|---|---|---|---|---|---|
| 6940 | *RRS1* | 3.1529 | 1.5474 | -0.67149 | -2.2254 | 1.34 |
| 6941 | *RASGRP1* | 4.553 | 0.19242 | -0.71669 | -0.81448 | 1.34 |
| 6942 | *IFRD1* | 2.7718 | 0.74863 | 0.5083 | 0.25102 | 1.34 |
| 6943 | *LDOC1* | 2.7192 | 0.69807 | 0.61139 | -0.44353 | 1.34 |
| 6944 | *CIPC* | 4.6642 | 1.0561 | -1.6921 | -2.5795 | 1.34 |
| 6945 | *INVS* | 3.169 | 2.003 | -1.1444 | -1.463 | 1.34 |
| 6946 | *IDO2* | 4.5802 | 0.14208 | -0.6951 | -0.88098 | 1.34 |
| 6947 | *LRCH1* | 2.7204 | 1.5935 | -0.28752 | -0.40577 | 1.34 |
| 6948 | *CASR* | 2.0977 | 1.49 | 0.4386 | -1.2208 | 1.34 |
| 6949 | *PKHD1* | 2.0049 | 1.0765 | 0.94481 | -1.1945 | 1.34 |
| 6950 | *IL18* | 3.5229 | 2.0618 | -1.5588 | -1.7316 | 1.34 |
| 6951 | *PRKAR2B* | 2.3981 | 2.0632 | -0.43551 | -1.5943 | 1.34 |
| 6952 | *ZNF783* | 2.401 | 1.4923 | 0.13052 | -0.73552 | 1.34 |
| 6953 | *PAPD5* | 2.846 | 0.98721 | 0.19051 | -1.0288 | 1.34 |
| 6954 | *DSCR4* | 1.6527 | 1.2751 | 1.0951 | -1.2618 | 1.34 |
| 6955 | *AMER1* | 3.7376 | 2.4815 | -2.197 | -3.0974 | 1.34 |
| 6956 | *WIPI2* | 3.1498 | 1.405 | -0.53346 | -0.85512 | 1.34 |
| 6957 | *C3orf30* | 3.0621 | 0.63937 | 0.31943 | -1.1039 | 1.34 |
| 6958 | *RNF186* | 2.8464 | 1.1781 | -0.00464 | -0.22601 | 1.34 |
| 6959 | *NPVF* | 2.7149 | 1.106 | 0.19776 | -0.64956 | 1.34 |
| 6960 | *MTMR10* | 3.1627 | 1.0368 | -0.18093 | -0.62307 | 1.34 |
| 6961 | *LCE1E* | 1.4134 | 1.4001 | 1.2044 | 0.91707 | 1.34 |
| 6962 | *ETV7* | 2.4662 | 1.574 | -0.0224 | -1.6974 | 1.34 |
| 6963 | *TSPAN10* | 1.814 | 1.7935 | 0.40999 | -0.94184 | 1.34 |
| 6964 | *ANK3* | 2.8418 | 2.3612 | -1.1861 | -1.3419 | 1.34 |
| 6965 | *BTG1* | 2.5809 | 1.4148 | 0.02091 | -0.33119 | 1.34 |
| 6966 | *AK1* | 3.7836 | 0.85028 | -0.61749 | -1.2008 | 1.34 |
| 6967 | *IFNW1* | 2.8231 | 1.6781 | -0.48492 | -1.9168 | 1.34 |
| 6968 | *IRF9* | 3.2557 | 1.0209 | -0.26055 | -1.2684 | 1.34 |
| 6969 | *CNTN2* | 3.0449 | 2.2751 | -1.3044 | -1.4824 | 1.34 |
| 6970 | *NCALD* | 2.1652 | 1.8636 | -0.01587 | -0.16506 | 1.34 |
| 6971 | *CNTD2* | 3.0046 | 1.8181 | -0.80978 | -1.0212 | 1.34 |
| 6972 | *HIST1H4F* | 2.1079 | 0.99453 | 0.91012 | -0.38491 | 1.34 |
| 6973 | *BRCC3* | 3.8004 | 0.44057 | -0.22873 | -1.0033 | 1.34 |
| 6974 | *FURIN* | 2.9692 | 2.2998 | -1.257 | -1.3353 | 1.34 |
| 6975 | *PTGS2* | 4.7756 | 0.21713 | -0.98104 | -1.4075 | 1.34 |
| 6976 | *IDUA* | 3.2146 | 0.95422 | -0.15855 | -1.1001 | 1.34 |
| 6977 | *OR6V1* | 1.586 | 1.2548 | 1.1692 | 0.22976 | 1.34 |
| 6978 | *MTHFSD* | 2.0211 | 1.4459 | 0.54299 | 0.081831 | 1.34 |
| 6979 | *SLC17A4* | 2.1777 | 1.2829 | 0.54921 | 0.094326 | 1.34 |
| 6980 | *SNRPC* | 2.7444 | 1.708 | -0.44261 | -1.0268 | 1.34 |
| 6981 | *PPT2* | 2.919 | 1.8867 | -0.79641 | -1.135 | 1.34 |
| 6982 | *FAM83F* | 2.2462 | 1.1964 | 0.56668 | 0.060946 | 1.34 |
| 6983 | *MDP1* | 1.8014 | 1.3144 | 0.89321 | -2.0187 | 1.34 |
| 6984 | *SLC35A3* | 4.0353 | 0.82493 | -0.85163 | -1.1309 | 1.34 |
| 6985 | *CCNY* | 2.1875 | 1.5472 | 0.27373 | -0.34653 | 1.34 |

| | | | | | | |
|---|---|---|---|---|---|---|
| 6986 | *ROBO4* | 1.68 | 1.1706 | 1.1576 | 0.26156 | 1.34 |
| 6987 | *SPINK7* | 4.3529 | 0.098095 | -0.44353 | -0.51761 | 1.34 |
| 6988 | *SEC14L5* | 3.5696 | 1.1771 | -0.73972 | -0.91076 | 1.34 |
| 6989 | *MOK* | 4.3409 | 0.76489 | -1.1005 | -1.5312 | 1.34 |
| 6990 | *GLRA4* | 4.1076 | 0.80786 | -0.9105 | -1.2457 | 1.33 |
| 6991 | *PCTP* | 2.0268 | 1.7137 | 0.26438 | -1.3622 | 1.33 |
| 6992 | *HTR4* | 2.5229 | 1.1505 | 0.33106 | -1.205 | 1.33 |
| 6993 | *ABL1* | 3.116 | 0.5268 | 0.36116 | -2.2104 | 1.33 |
| 6994 | *ZNF645* | 4.4757 | 0.28106 | -0.75293 | -1.9284 | 1.33 |
| 6995 | *RIT1* | 3.1283 | 1.0509 | -0.17647 | -0.58837 | 1.33 |
| 6996 | *NDRG2* | 2.8753 | 1.5107 | -0.38376 | -1.1958 | 1.33 |
| 6997 | *CCDC124* | 5.0444 | -0.46218 | -0.58088 | -0.96012 | 1.33 |
| 6998 | *RCOR3* | 3.36 | 0.51318 | 0.12772 | -0.4606 | 1.33 |
| 6999 | *C11orf97* | 2.3764 | 1.4189 | 0.2053 | -0.89866 | 1.33 |
| 7000 | *FGF7* | 1.6961 | 1.6173 | 0.68701 | 0.46419 | 1.33 |
| 7001 | *PRSS2* | 3.6538 | 1.2767 | -0.93011 | -1.3285 | 1.33 |
| 7002 | *NID1* | 4.6639 | 0.034566 | -0.69841 | -0.88416 | 1.33 |
| 7003 | *PCDHGA2* | 4.9078 | 0.13729 | -1.0453 | -1.2935 | 1.33 |
| 7004 | *SOX4* | 2.5399 | 0.87689 | 0.58268 | -2.7277 | 1.33 |
| 7005 | *SNRPA1* | 4.827 | 0.66293 | -1.492 | -2.5419 | 1.33 |
| 7006 | *SLC25A48* | 3.6649 | 0.58909 | -0.25608 | -1.7836 | 1.33 |
| 7007 | *ACAT2* | 1.5162 | 1.2603 | 1.2209 | -2.3949 | 1.33 |
| 7008 | *C6orf25* | 3.4866 | 1.0055 | -0.49481 | -1.3961 | 1.33 |
| 7009 | *TMPRSS3* | 2.7549 | 0.95084 | 0.29117 | 0.19992 | 1.33 |
| 7010 | *UBOX5* | 4.1782 | -0.06272 | -0.11882 | -0.75361 | 1.33 |
| 7011 | *ASIC2* | 3.0518 | 0.99326 | -0.04876 | -0.23045 | 1.33 |
| 7012 | *PLCXD3* | 4.6638 | 0.59244 | -1.26 | -1.2827 | 1.33 |
| 7013 | *ZNF330* | 1.8459 | 1.2814 | 0.86883 | -0.23607 | 1.33 |
| 7014 | *KANSL1* | 4.164 | 0.92957 | -1.0986 | -1.7718 | 1.33 |
| 7015 | *FKBP1A* | 5.4207 | -0.58603 | -0.84001 | -2.2142 | 1.33 |
| 7016 | *CA13* | 4.0901 | 0.077839 | -0.17375 | -0.93889 | 1.33 |
| 7017 | *MS4A15* | 2.6942 | 1.216 | 0.08386 | -0.17622 | 1.33 |
| 7018 | *TFAP2B* | 5.0574 | -0.46764 | -0.5959 | -1.8949 | 1.33 |
| 7019 | *SOWAHB* | 2.1287 | 1.0265 | 0.83804 | -0.97944 | 1.33 |
| 7020 | *LDLRAD1* | 2.6942 | 2.5526 | -1.2541 | -1.416 | 1.33 |
| 7021 | *REG4* | 3.8412 | 0.21972 | -0.06857 | -0.12688 | 1.33 |
| 7022 | *CFH* | 2.2751 | 1.7633 | -0.04669 | -1.6759 | 1.33 |
| 7023 | *TRIM22* | 3.6448 | 0.79236 | -0.44573 | -0.68674 | 1.33 |
| 7024 | *OR5AP2* | 2.793 | 1.8462 | -0.64793 | -1.9036 | 1.33 |
| 7025 | *SPACA5* | 3.3156 | 0.96833 | -0.29349 | -0.65848 | 1.33 |
| 7026 | *APOBEC3C* | 2.5102 | 1.0725 | 0.40742 | -0.47028 | 1.33 |
| 7027 | *NEUROD2* | 4.7611 | -0.08348 | -0.68825 | -1.0088 | 1.33 |
| 7028 | *C6orf132* | 1.7089 | 1.1911 | 1.0892 | -1.5866 | 1.33 |
| 7029 | *RNASE10* | 2.1546 | 1.4732 | 0.36133 | -0.27359 | 1.33 |
| 7030 | *ZNF469* | 2.1472 | 1.7579 | 0.084 | -1.5741 | 1.33 |
| 7031 | *OR8H1* | 2.5528 | 2.3268 | -0.89269 | -1.1325 | 1.33 |

| | | | | | | |
|---|---|---|---|---|---|---|
| 7032 | *ALOX15B* | 2.6812 | 2.6286 | -1.3237 | -2.1706 | 1.33 |
| 7033 | *MRPS7* | 2.2127 | 1.0744 | 0.69893 | -0.6114 | 1.33 |
| 7034 | *OR52B6* | 3.0978 | 1.6791 | -0.79147 | -1.3593 | 1.33 |
| 7035 | *KIRREL3* | 2.118 | 2.079 | -0.21169 | -1.4214 | 1.33 |
| 7036 | *GPR156* | 3.1695 | 0.98943 | -0.1741 | -2.4011 | 1.33 |
| 7037 | *RTP3* | 3.1694 | 1.3228 | -0.50754 | -1.1879 | 1.33 |
| 7038 | *ZNF213* | 3.2864 | 0.41473 | 0.28351 | -0.66535 | 1.33 |
| 7039 | *KIF2C* | 3.985 | 0.69988 | -0.70064 | -1.9713 | 1.33 |
| 7040 | *LCOR* | 2.9226 | 1.1445 | -0.0834 | -0.14163 | 1.33 |
| 7041 | *RBP1* | 2.3253 | 2.1401 | -0.4818 | -0.98421 | 1.33 |
| 7042 | *CPNE3* | 4.621 | 0.13558 | -0.77418 | -1.536 | 1.33 |
| 7043 | *JMJD1C* | 3.0947 | 0.84507 | 0.0418 | -0.54826 | 1.33 |
| 7044 | *ATP6V0E2* | 4.4035 | -0.01791 | -0.40501 | -1.8087 | 1.33 |
| 7045 | *LPL* | 1.6219 | 1.2966 | 1.0619 | -1.632 | 1.33 |
| 7046 | *PDE8B* | 2.4049 | 1.3683 | 0.20636 | -1.7363 | 1.33 |
| 7047 | *TMEM45A* | 3.2194 | 3.0931 | -2.3334 | -4.1669 | 1.33 |
| 7048 | *UBE4A* | 3.991 | 0.315 | -0.32717 | -1.6989 | 1.33 |
| 7049 | *HKR1* | 3.9873 | 0.21869 | -0.22726 | -1.5194 | 1.33 |
| 7050 | *CCDC71* | 3.3385 | 0.85499 | -0.21647 | -1.4046 | 1.33 |
| 7051 | *CAMKMT* | 3.4825 | 0.91249 | -0.4187 | -1.1492 | 1.33 |
| 7052 | *SIRT1* | 4.4238 | 0.97399 | -1.4233 | -2.6889 | 1.32 |
| 7053 | *PPP3CB* | 3.1967 | 0.85124 | -0.0741 | -0.09188 | 1.32 |
| 7054 | *CD4* | 2.2534 | 1.0266 | 0.69327 | 0.12362 | 1.32 |
| 7055 | *IL17REL* | 1.5302 | 1.3194 | 1.123 | -1.9041 | 1.32 |
| 7056 | *C16orf54* | 3.6639 | 0.36987 | -0.0614 | -0.33711 | 1.32 |
| 7057 | *TTLL6* | 4.9512 | -0.14289 | -0.83631 | -1.8996 | 1.32 |
| 7058 | *MINA* | 5.5787 | -0.15702 | -1.4507 | -2.4832 | 1.32 |
| 7059 | *MXD1* | 2.9848 | 0.8649 | 0.12112 | -0.58676 | 1.32 |
| 7060 | *ABHD14B* | 2.7783 | 1.0734 | 0.11886 | -1.0414 | 1.32 |
| 7061 | *ZNF556* | 3.1523 | 1.7112 | -0.89401 | -2.6805 | 1.32 |
| 7062 | *RFX7* | 3.2502 | 0.53274 | 0.18642 | -2.0383 | 1.32 |
| 7063 | *TMEM171* | 2.1235 | 1.6448 | 0.19987 | -1.3743 | 1.32 |
| 7064 | *PUS7* | 4.7295 | 0.51379 | -1.2752 | -1.4754 | 1.32 |
| 7065 | *ZNF883* | 3.7153 | 0.23344 | 0.01861 | -0.51805 | 1.32 |
| 7066 | *BLOC1S1* | 3.7423 | 1.1265 | -0.90154 | -0.9653 | 1.32 |
| 7067 | *C1orf194* | 2.6022 | 1.3211 | 0.04392 | -0.76355 | 1.32 |
| 7068 | *AKAP8* | 3.5756 | 0.41049 | -0.01897 | -0.64397 | 1.32 |
| 7069 | *UTRN* | 2.7774 | 0.77622 | 0.41252 | -2.0629 | 1.32 |
| 7070 | *PRDM6* | 3.2703 | 0.53634 | 0.15921 | -0.99079 | 1.32 |
| 7071 | *KRTAP1-5* | 3.0326 | 0.79772 | 0.13475 | -0.62629 | 1.32 |
| 7072 | *ACOXL* | 1.8327 | 1.423 | 0.70864 | -1.6891 | 1.32 |
| 7073 | *GRM7* | 3.4453 | 1.6026 | -1.0841 | -1.2278 | 1.32 |
| 7074 | *HSFY1* | 4.7755 | -0.13288 | -0.67934 | -1.8187 | 1.32 |
| 7075 | *PPRC1* | 2.566 | 0.89321 | 0.50404 | -0.37669 | 1.32 |
| 7076 | *GKAP1* | 4.9622 | 0.18317 | -1.1826 | -2.1342 | 1.32 |
| 7077 | *KIR3DL3* | 3.0451 | 0.53014 | 0.38747 | -0.88277 | 1.32 |

| | | | | | | |
|---|---|---|---|---|---|---|
| 7078 | *CLIP3* | 4.1445 | 0.69134 | -0.87382 | -1.7999 | 1.32 |
| 7079 | *ESM1* | 2.0391 | 1.8217 | 0.10069 | -0.72406 | 1.32 |
| 7080 | *NOC3L* | 2.193 | 0.97778 | 0.79009 | -1.1207 | 1.32 |
| 7081 | *SLC27A2* | 4.5003 | 0.53285 | -1.0726 | -1.62 | 1.32 |
| 7082 | *MAG* | 2.6198 | 2.4333 | -1.093 | -1.244 | 1.32 |
| 7083 | *RSPH14* | 4.0725 | 0.20802 | -0.32087 | -2.5571 | 1.32 |
| 7084 | *OR5P2* | 4.7875 | -0.21291 | -0.61528 | -2.0629 | 1.32 |
| 7085 | *DUSP8* | 1.59 | 1.2572 | 1.1117 | -1.2198 | 1.32 |
| 7086 | *ABCG8* | 2.4523 | 1.4936 | 0.01271 | -1.078 | 1.32 |
| 7087 | *C7orf50* | 3.3902 | 2.1643 | -1.5962 | -5.7134 | 1.32 |
| 7088 | *MRGPRX1* | 3.9297 | 0.48812 | -0.46017 | -3.3281 | 1.32 |
| 7089 | *RICTOR* | 2.0778 | 0.98916 | 0.89019 | -1.6597 | 1.32 |
| 7090 | *WNT7A* | 1.8497 | 1.1133 | 0.99414 | -1.2935 | 1.32 |
| 7091 | *INTS12* | 2.4746 | 0.81241 | 0.66962 | -1.5258 | 1.32 |
| 7092 | *KLHDC7B* | 3.9953 | 1.3206 | -1.3593 | -3.087 | 1.32 |
| 7093 | *DPCR1* | 1.7666 | 1.5589 | 0.63099 | -2.0653 | 1.32 |
| 7094 | *DPH5* | 3.9618 | 0.7594 | -0.76606 | -1.3875 | 1.32 |
| 7095 | *XPO7* | 2.1269 | 1.7571 | 0.07113 | -0.10524 | 1.32 |
| 7096 | *NGFRAP1* | 2.5009 | 1.9294 | -0.47527 | -1.5406 | 1.32 |
| 7097 | *MAML3* | 3.0305 | 0.66824 | 0.25595 | -0.59007 | 1.32 |
| 7098 | *TIMM21* | 3.4952 | 0.71707 | -0.25767 | -1.9062 | 1.32 |
| 7099 | *MN1* | 3.9493 | 0.017858 | -0.0131 | -0.3193 | 1.32 |
| 7100 | *BTN3A3* | 3.523 | 0.55409 | -0.124 | -0.66945 | 1.32 |
| 7101 | *LOC100289561* | 2.3111 | 1.3422 | 0.29979 | -2.6365 | 1.32 |
| 7102 | *ERP27* | 3.5899 | 0.19846 | 0.16442 | -1.4597 | 1.32 |
| 7103 | *RAB42* | 1.8863 | 1.529 | 0.53697 | -2.0559 | 1.32 |
| 7104 | *L2HGDH* | 3.8329 | 1.0564 | -0.93837 | -2.1987 | 1.32 |
| 7105 | *ZNF584* | 3.8888 | 0.50541 | -0.44374 | -1.1303 | 1.32 |
| 7106 | *IKBKG* | 5.1695 | -0.33937 | -0.88008 | -3.2582 | 1.32 |
| 7107 | *WFDC9* | 2.3553 | 2.2813 | -0.68729 | -0.79745 | 1.32 |
| 7108 | *LMF1* | 2.2138 | 1.5151 | 0.22013 | -1.2891 | 1.32 |
| 7109 | *FLT4* | 4.3119 | 0.24399 | -0.60717 | -1.742 | 1.32 |
| 7110 | *GSK3B* | 2.3368 | 1.9769 | -0.36527 | -0.64311 | 1.32 |
| 7111 | *CEP55* | 4.3669 | 0.38844 | -0.80743 | -1.6851 | 1.32 |
| 7112 | *RBP4* | 2.7229 | 1.2113 | 0.01338 | -0.19136 | 1.32 |
| 7113 | *THAP7* | 2.0425 | 1.2337 | 0.66959 | -1.5268 | 1.32 |
| 7114 | *NRL* | 2.3568 | 1.0544 | 0.53445 | -0.19289 | 1.32 |
| 7115 | *SPOPL* | 2.6576 | 2.5561 | -1.2705 | -2.3204 | 1.31 |
| 7116 | *GYG1* | 3.1551 | 0.44405 | 0.34401 | -1.5313 | 1.31 |
| 7117 | *KRTAP7-1* | 2.0191 | 0.97534 | 0.94778 | -0.95 | 1.31 |
| 7118 | *GPR75* | 2.2885 | 0.9747 | 0.67879 | -1.4102 | 1.31 |
| 7119 | *GABRE* | 3.9945 | 0.006056 | -0.05926 | -1.5125 | 1.31 |
| 7120 | *CYP2U1* | 4.9476 | 1.4194 | -2.4268 | -3.0179 | 1.31 |
| 7121 | *DIP2B* | 2.691 | 2.4978 | -1.2493 | -1.4873 | 1.31 |
| 7122 | *HBA2* | 3.2382 | 1.2427 | -0.54188 | -0.71467 | 1.31 |
| 7123 | *NSRP1* | 4.2611 | -0.06433 | -0.25794 | -2.1239 | 1.31 |

| | | | | | | |
|---|---|---|---|---|---|---|
| 7124 | *KIF16B* | 4.7977 | 0.67722 | -1.5382 | -1.6826 | 1.31 |
| 7125 | *PRSS45* | 2.8468 | 2.4142 | -1.325 | -1.9622 | 1.31 |
| 7126 | *OR4D6* | 4.8061 | 0.24672 | -1.1169 | -2.056 | 1.31 |
| 7127 | *PYCR2* | 3.5361 | 0.48977 | -0.0907 | -0.63309 | 1.31 |
| 7128 | *MMD2* | 1.9755 | 1.8166 | 0.14299 | -0.84738 | 1.31 |
| 7129 | *YY1* | 2.7165 | 0.67857 | 0.53966 | 0.44402 | 1.31 |
| 7130 | *OR13C3* | 2.8451 | 0.96541 | 0.12367 | -1.2997 | 1.31 |
| 7131 | *MTMR11* | 3.5541 | 1.3758 | -0.99608 | -2.7871 | 1.31 |
| 7132 | *SIDT2* | 3.0415 | 1.2428 | -0.35064 | -0.93057 | 1.31 |
| 7133 | *ZDHHC22* | 2.6533 | 0.90462 | 0.37546 | -1.6477 | 1.31 |
| 7134 | *CPVL* | 4.1546 | -0.02105 | -0.20044 | -0.58231 | 1.31 |
| 7135 | *C2orf48* | 2.3703 | 1.1355 | 0.42713 | -0.5656 | 1.31 |
| 7136 | *C9orf43* | 3.7773 | 0.85526 | -0.69985 | -1.1095 | 1.31 |
| 7137 | *KDM2B* | 3.0334 | 1.7624 | -0.86358 | -1.4473 | 1.31 |
| 7138 | *FOXO4* | 2.222 | 1.5844 | 0.12471 | -0.30496 | 1.31 |
| 7139 | *KCNK10* | 3.1411 | 0.8332 | -0.04324 | -1.2572 | 1.31 |
| 7140 | *TCF19* | 4.2639 | 0.090632 | -0.42397 | -0.44135 | 1.31 |
| 7141 | *TBX3* | 5.4205 | 0.21634 | -1.7065 | -1.7528 | 1.31 |
| 7142 | *CERK* | 4.3361 | -0.0828 | -0.32321 | -0.39775 | 1.31 |
| 7143 | *OR2K2* | 4.8017 | 0.18292 | -1.0553 | -2.875 | 1.31 |
| 7144 | *SSX3* | 2.5744 | 1.3592 | -0.00492 | -2.0411 | 1.31 |
| 7145 | *TMEM242* | 2.1705 | 2.0964 | -0.33826 | -1.8095 | 1.31 |
| 7146 | *CRABP2* | 2.9061 | 0.57026 | 0.4521 | -1.1344 | 1.31 |
| 7147 | *ZNF653* | 2.7195 | 0.70323 | 0.50565 | -0.51722 | 1.31 |
| 7148 | *PREX2* | 2.631 | 1.177 | 0.11982 | -1.0614 | 1.31 |
| 7149 | *OR4M1* | 4.7924 | -0.35565 | -0.50904 | -1.1558 | 1.31 |
| 7150 | *AGBL3* | 1.8799 | 1.3411 | 0.70642 | -0.47179 | 1.31 |
| 7151 | *TMEM181* | 3.4361 | 0.31906 | 0.1718 | -1.3141 | 1.31 |
| 7152 | *WDR70* | 5.4763 | -0.47183 | -1.0779 | -1.4657 | 1.31 |
| 7153 | *LCE3D* | 3.1191 | 0.58201 | 0.22523 | -0.8792 | 1.31 |
| 7154 | *ATG4A* | 4.0726 | 0.24868 | -0.39533 | -1.2646 | 1.31 |
| 7155 | *DPF1* | 4.2404 | 0.049395 | -0.36454 | -0.86182 | 1.31 |
| 7156 | *INHBE* | 3.6003 | 1.4194 | -1.0949 | -1.4337 | 1.31 |
| 7157 | *ALDH3B2* | 4.1128 | 0.12694 | -0.31504 | -0.53458 | 1.31 |
| 7158 | *FAM153A* | 2.9512 | 1.3562 | -0.38348 | -0.8338 | 1.31 |
| 7159 | *SPATA16* | 2.5006 | 1.3018 | 0.12149 | -0.28806 | 1.31 |
| 7160 | *MTG1* | 2.1895 | 1.6962 | 0.03704 | -0.67203 | 1.31 |
| 7161 | *CBLN2* | 3.8238 | 0.3533 | -0.25501 | -1.1117 | 1.31 |
| 7162 | *OTUD6B* | 2.3485 | 1.5089 | 0.06442 | -2.0392 | 1.31 |
| 7163 | *CLIC4* | 2.7692 | 1.3055 | -0.15424 | -0.58589 | 1.31 |
| 7164 | *TAC4* | 5.3247 | -0.63214 | -0.77241 | -1.3627 | 1.31 |
| 7165 | *ASXL2* | 4.2971 | 0.32236 | -0.69972 | -1.0644 | 1.31 |
| 7166 | *CCDC37* | 2.9937 | 0.92051 | 0.00551 | -1.7438 | 1.31 |
| 7167 | *SLC6A3* | 2.3772 | 1.2172 | 0.32504 | -1.4234 | 1.31 |
| 7168 | *FXR2* | 3.4711 | 0.97836 | -0.53058 | -1.1729 | 1.31 |
| 7169 | *LMLN* | 2.4429 | 0.79471 | 0.68105 | -1.5121 | 1.31 |

| | | | | | | |
|---|---|---|---|---|---|---|
| 7170 | *FAM207A* | 3.2667 | 0.97589 | -0.32397 | -0.80909 | 1.31 |
| 7171 | *C1orf68* | 2.3352 | 1.6784 | -0.09539 | -1.2234 | 1.31 |
| 7172 | *CSNK1A1L* | 2.0626 | 1.7766 | 0.07871 | -0.06657 | 1.31 |
| 7173 | *OR8G5* | 2.0391 | 0.95321 | 0.92534 | -1.7874 | 1.31 |
| 7174 | *MFSD10* | 4.056 | 0.39486 | -0.53361 | -2.1419 | 1.31 |
| 7175 | *SCX* | 3.3862 | 1.0633 | -0.53268 | -0.58787 | 1.31 |
| 7176 | *KIAA1211* | 3.3307 | 1.2519 | -0.66583 | -1.3745 | 1.31 |
| 7177 | *CEL* | 2.5265 | 0.77752 | 0.61125 | -0.03871 | 1.31 |
| 7178 | *PPP1R27* | 3.0273 | 0.5137 | 0.37426 | -1.3294 | 1.31 |
| 7179 | *OR52H1* | 3.6427 | 0.784 | -0.51152 | -0.75974 | 1.31 |
| 7180 | *PAX8* | 4.0787 | 0.68352 | -0.84806 | -0.8504 | 1.30 |
| 7181 | *SMURF1* | 2.5977 | 0.65859 | 0.65733 | -0.30606 | 1.30 |
| 7182 | *TIMP3* | 1.7794 | 1.1567 | 0.97736 | -1.6225 | 1.30 |
| 7183 | *SYNGR1* | 2.9393 | 0.75055 | 0.22282 | -1.48 | 1.30 |
| 7184 | *SEPT3* | 2.973 | 0.86459 | 0.07442 | -1.4171 | 1.30 |
| 7185 | *PAG1* | 2.1723 | 1.2393 | 0.50023 | -0.44353 | 1.30 |
| 7186 | *KLF11* | 4.7048 | -0.28873 | -0.50426 | -2.5956 | 1.30 |
| 7187 | *NPDC1* | 1.8415 | 1.057 | 1.0132 | -0.89592 | 1.30 |
| 7188 | *RPUSD2* | 4.2797 | -0.05721 | -0.31103 | -0.9939 | 1.30 |
| 7189 | *RASGRF1* | 1.8714 | 1.3102 | 0.72978 | -0.4854 | 1.30 |
| 7190 | *MYBPC3* | 3.0569 | 1.4909 | -0.63643 | -2.0758 | 1.30 |
| 7191 | *VN1R1* | 2.8319 | 2.069 | -0.98966 | -1.6176 | 1.30 |
| 7192 | *KIAA0141* | 4.0373 | 0.088541 | -0.21468 | -1.0326 | 1.30 |
| 7193 | *COX8A* | 4.7297 | 0.23807 | -1.0567 | -1.4213 | 1.30 |
| 7194 | *S100A16* | 2.9353 | 0.67615 | 0.29865 | -0.52397 | 1.30 |
| 7195 | *GPC2* | 2.2034 | 0.9061 | 0.79872 | -1.4692 | 1.30 |
| 7196 | *LCLAT1* | 4.5576 | 0.85005 | -1.4997 | -1.5243 | 1.30 |
| 7197 | *OR51M1* | 3.8078 | 0.76316 | -0.66301 | -2.1759 | 1.30 |
| 7198 | *OR4X2* | 3.2538 | 0.87821 | -0.22422 | -0.3146 | 1.30 |
| 7199 | *MURC* | 2.9255 | 2.1189 | -1.137 | -1.2825 | 1.30 |
| 7200 | *DEFA3* | 4.625 | 0.18105 | -0.89947 | -1.2108 | 1.30 |
| 7201 | *TMEM52B* | 4.079 | 0.42931 | -0.602 | -1.3929 | 1.30 |
| 7202 | *GRK4* | 2.1719 | 1.1445 | 0.58984 | -0.72334 | 1.30 |
| 7203 | *HIST1H4L* | 2.1864 | 1.9615 | -0.24262 | -0.31848 | 1.30 |
| 7204 | *GATAD1* | 2.6957 | 1.9364 | -0.72684 | -1.2895 | 1.30 |
| 7205 | *TRPM5* | 2.7355 | 0.62805 | 0.53898 | -1.271 | 1.30 |
| 7206 | *SDSL* | 2.159 | 1.4115 | 0.33085 | -2.4405 | 1.30 |
| 7207 | *HEXIM2* | 3.608 | 1.6615 | -1.3683 | -1.8251 | 1.30 |
| 7208 | *CSRNP3* | 4.7686 | -0.22313 | -0.64467 | -1.2356 | 1.30 |
| 7209 | *AMER3* | 2.6671 | 1.9177 | -0.68565 | -1.1226 | 1.30 |
| 7210 | *SYT7* | 3.0581 | 1.3403 | -0.49926 | -1.246 | 1.30 |
| 7211 | *AIFM1* | 3.2547 | 0.9159 | -0.2715 | -0.55002 | 1.30 |
| 7212 | *PIR* | 2.0995 | 1.886 | -0.08748 | -0.20739 | 1.30 |
| 7213 | *LOC645382* | 4.6219 | -0.31092 | -0.41307 | -2.0267 | 1.30 |
| 7214 | *PLEKHG4B* | 4.9919 | 0.046873 | -1.141 | -1.291 | 1.30 |
| 7215 | *PTH2* | 4.8089 | -0.15953 | -0.75169 | -1.237 | 1.30 |

| | | | | | | |
|---|---|---|---|---|---|---|
| 7216 | *C5orf24* | 3.5332 | 0.9019 | -0.53775 | -1.3218 | 1.30 |
| 7217 | *GULP1* | 1.8217 | 1.1911 | 0.88385 | -2.7379 | 1.30 |
| 7218 | *RCN3* | 2.3009 | 1.0569 | 0.53845 | -1.6128 | 1.30 |
| 7219 | *EFCAB9* | 2.8489 | 0.90742 | 0.13911 | -1.5125 | 1.30 |
| 7220 | *PIF1* | 3.5906 | 0.74765 | -0.44353 | -1.6626 | 1.30 |
| 7221 | *DPYSL5* | 2.6095 | 1.049 | 0.23504 | -0.70231 | 1.30 |
| 7222 | *RBBP9* | 3.4914 | 1.6973 | -1.2954 | -2.5119 | 1.30 |
| 7223 | *MRC2* | 4.1735 | 0.058484 | -0.33933 | -0.34308 | 1.30 |
| 7224 | *ZNF121* | 1.6588 | 1.6054 | 0.62741 | -1.6587 | 1.30 |
| 7225 | *RALGAPA1* | 2.002 | 1.6464 | 0.24309 | -0.16438 | 1.30 |
| 7226 | *GLYATL3* | 4.0865 | -0.06374 | -0.13182 | -0.62596 | 1.30 |
| 7227 | *SCAND1* | 3.4044 | 0.59341 | -0.10716 | -0.35367 | 1.30 |
| 7228 | *OR2C3* | 1.9997 | 1.8217 | 0.06813 | -0.06319 | 1.30 |
| 7229 | *LOC389895* | 4.2916 | 0.15417 | -0.55675 | -0.85345 | 1.30 |
| 7230 | *TBPL1* | 2.6262 | 0.91268 | 0.3501 | -1.0039 | 1.30 |
| 7231 | *GGH* | 3.1589 | 0.89943 | -0.16946 | -1.988 | 1.30 |
| 7232 | *RNASEH2B* | 5.6948 | -0.2715 | -1.5346 | -3.0054 | 1.30 |
| 7233 | *TRAM1* | 2.3853 | 1.4271 | 0.07586 | -1.1004 | 1.30 |
| 7234 | *CPNE4* | 4.323 | 0.031912 | -0.46764 | -0.68781 | 1.30 |
| 7235 | *OR51Q1* | 3.7742 | 0.95686 | -0.84419 | -0.87318 | 1.30 |
| 7236 | *GNPTAB* | 3.7536 | 0.43378 | -0.30056 | -0.67298 | 1.30 |
| 7237 | *OR51A7* | 3.1185 | 1.5112 | -0.7436 | -0.7757 | 1.30 |
| 7238 | *SPATA31E1* | 3.1493 | 0.77005 | -0.03348 | -0.59143 | 1.30 |
| 7239 | *GEMIN2* | 4.6848 | 0.26156 | -1.0614 | -2.1321 | 1.29 |
| 7240 | *REG3A* | 2.9876 | 0.72377 | 0.17348 | -1.1512 | 1.29 |
| 7241 | *XKRY* | 2.8395 | 1.4887 | -0.44353 | -0.55163 | 1.29 |
| 7242 | *SPRR3* | 2.4768 | 0.82135 | 0.58647 | -2.2075 | 1.29 |
| 7243 | *PLET1* | 4.8901 | -0.08846 | -0.91721 | -1.4122 | 1.29 |
| 7244 | *DFNA5* | 3.1008 | 1.1496 | -0.36668 | -0.90462 | 1.29 |
| 7245 | *STOM* | 4.7291 | -0.23831 | -0.60717 | -1.9065 | 1.29 |
| 7246 | *ADAM19* | 4.7643 | -0.39856 | -0.48231 | -0.89974 | 1.29 |
| 7247 | *NEBL* | 3.8583 | 0.17873 | -0.15439 | -1.287 | 1.29 |
| 7248 | *ADAM8* | 5.1858 | 0.002076 | -1.3055 | -1.8157 | 1.29 |
| 7249 | *SMARCC2* | 3.7468 | 0.3881 | -0.25264 | -1.0796 | 1.29 |
| 7250 | *KCNK17* | 4.4524 | 0.51759 | -1.0878 | -1.2353 | 1.29 |
| 7251 | *UGT8* | 3.7198 | 0.13368 | 0.02789 | -1.7018 | 1.29 |
| 7252 | *KCNA5* | 2.4416 | 1.3565 | 0.08309 | -0.75448 | 1.29 |
| 7253 | *IDH3G* | 3.6141 | 0.50127 | -0.23444 | -1.7065 | 1.29 |
| 7254 | *ADAM32* | 2.5947 | 1.2782 | 0.008 | -0.21201 | 1.29 |
| 7255 | *PARP4* | 4.1306 | 0.038291 | -0.28896 | -1.6597 | 1.29 |
| 7256 | *TSC22D4* | 1.6001 | 1.3007 | 0.97805 | 0.7565 | 1.29 |
| 7257 | *HCRTR1* | 1.9177 | 1.3807 | 0.58021 | 0.40441 | 1.29 |
| 7258 | *C14orf93* | 4.4148 | 0.55654 | -1.093 | -1.4061 | 1.29 |
| 7259 | *DCAF12L2* | 3.0528 | 1.0421 | -0.21666 | -0.90542 | 1.29 |
| 7260 | *HAUS1* | 3.1542 | 2.2204 | -1.4969 | -1.6262 | 1.29 |
| 7261 | *DTNB* | 3.5177 | 0.7429 | -0.38361 | -0.62306 | 1.29 |

| | | | | | | |
|---|---|---|---|---|---|---|
| 7262 | *UBTD1* | 2.2221 | 1.2246 | 0.43028 | -0.2146 | 1.29 |
| 7263 | *MYOC* | 4.7644 | 0.24209 | -1.13 | -2.2352 | 1.29 |
| 7264 | *HDDC2* | 2.7241 | 0.90791 | 0.24325 | -0.7602 | 1.29 |
| 7265 | *EPS8L1* | 3.6517 | 1.3372 | -1.1138 | -2.2473 | 1.29 |
| 7266 | *SLC25A32* | 2.3517 | 1.3362 | 0.18614 | 0.12559 | 1.29 |
| 7267 | *NEU3* | 3.0599 | 0.58197 | 0.23139 | -1.1108 | 1.29 |
| 7268 | *KCNA2* | 2.4866 | 1.4731 | -0.08646 | -0.14649 | 1.29 |
| 7269 | *ERVW-1* | 4.5291 | 0.3464 | -1.0024 | -1.4541 | 1.29 |
| 7270 | *HENMT1* | 2.7957 | 1.0817 | -0.005 | -1.3139 | 1.29 |
| 7271 | *PPP1R17* | 3.7007 | 1.8652 | -1.6939 | -1.9431 | 1.29 |
| 7272 | *OR11A1* | 4.6541 | 0.64084 | -1.4233 | -1.6339 | 1.29 |
| 7273 | *ANXA13* | 2.1933 | 1.3103 | 0.36776 | -0.57084 | 1.29 |
| 7274 | *RGS3* | 2.9156 | 0.92011 | 0.03509 | -0.18775 | 1.29 |
| 7275 | *SLC25A24* | 2.4701 | 0.84333 | 0.55656 | -0.35259 | 1.29 |
| 7276 | *PCDH19* | 1.8617 | 1.3492 | 0.65871 | -1.0644 | 1.29 |
| 7277 | *CAPN2* | 3.4293 | 0.26122 | 0.17905 | -1.4467 | 1.29 |
| 7278 | *CD27* | 4.6354 | 1.0162 | -1.7822 | -2.1412 | 1.29 |
| 7279 | *DCLRE1B* | 2.4462 | 0.87831 | 0.54424 | -2.9338 | 1.29 |
| 7280 | *MANSC1* | 4.9355 | -0.44353 | -0.62362 | -1.3235 | 1.29 |
| 7281 | *TMEM125* | 5.0272 | 0.72256 | -1.8816 | -2.7384 | 1.29 |
| 7282 | *AP5Z1* | 4.021 | 0.78129 | -0.93517 | -1.7338 | 1.29 |
| 7283 | *SSR2* | 5.8916 | -0.71939 | -1.3055 | -1.9722 | 1.29 |
| 7284 | *CYP39A1* | 2.5469 | 0.904 | 0.41542 | -0.80828 | 1.29 |
| 7285 | *PSMD13* | 3.5844 | 0.54691 | -0.26518 | -0.82044 | 1.29 |
| 7286 | *FGF17* | 4.6452 | 1.3252 | -2.1055 | -2.2311 | 1.29 |
| 7287 | *FAM229B* | 3.0028 | 0.48398 | 0.37747 | 0.23574 | 1.29 |
| 7288 | *ALX4* | 4.6201 | -0.08194 | -0.67482 | -2.3689 | 1.29 |
| 7289 | *NETO2* | 4.0204 | 0.66344 | -0.82208 | -2.2599 | 1.29 |
| 7290 | *FGFR2* | 3.2723 | 0.67415 | -0.08549 | -1.0579 | 1.29 |
| 7291 | *CAP1* | 5.3058 | -0.11854 | -1.3271 | -1.3464 | 1.29 |
| 7292 | *FRG2C* | 2.8482 | 0.69513 | 0.31649 | -0.45015 | 1.29 |
| 7293 | *ABCB7* | 4.678 | 0.34899 | -1.1673 | -2.0593 | 1.29 |
| 7294 | *FAM120A* | 2.3723 | 0.92356 | 0.56241 | -2.3378 | 1.29 |
| 7295 | *AGMAT* | 3.8084 | 0.56162 | -0.51398 | -1.9558 | 1.29 |
| 7296 | *PCDH12* | 3.9092 | 0.33434 | -0.38862 | -1.3728 | 1.28 |
| 7297 | *IGSF5* | 2.4301 | 2.3703 | -0.94563 | -1.6339 | 1.28 |
| 7298 | *F11* | 3.6649 | 1.7015 | -1.5121 | -1.7462 | 1.28 |
| 7299 | *DEFB108B* | 4.2026 | 0.31468 | -0.66358 | -0.91215 | 1.28 |
| 7300 | *LRRC46* | 1.9448 | 1.0363 | 0.87239 | -1.2963 | 1.28 |
| 7301 | *C19orf71* | 2.903 | 1.3588 | -0.40863 | -1.119 | 1.28 |
| 7302 | *OR52E4* | 3.8908 | 0.63968 | -0.67801 | -1.3287 | 1.28 |
| 7303 | *GIF* | 5.3094 | -0.66948 | -0.78784 | -0.84141 | 1.28 |
| 7304 | *CALN1* | 3.4614 | 0.54241 | -0.15385 | -1.57 | 1.28 |
| 7305 | *TAS2R3* | 2.1445 | 0.88379 | 0.82106 | -0.18009 | 1.28 |
| 7306 | *CHPF* | 3.75 | 0.16085 | -0.0629 | -0.31504 | 1.28 |
| 7307 | *OR52I1* | 3.4751 | 0.5531 | -0.18133 | -1.7585 | 1.28 |

| | | | | | | |
|---|---|---|---|---|---|---|
| 7308 | *OR11H6* | 4.3176 | -0.09312 | -0.37797 | -1.626 | 1.28 |
| 7309 | *ZBTB47* | 3.4923 | 0.37835 | -0.02428 | -1.6152 | 1.28 |
| 7310 | *DRP2* | 3.5466 | 0.30505 | -0.00613 | -0.46153 | 1.28 |
| 7311 | *CFAP43* | 3.4616 | 0.98984 | -0.60604 | -1.4606 | 1.28 |
| 7312 | *SHH* | 4.0234 | 0.80877 | -0.98709 | -1.1043 | 1.28 |
| 7313 | *RBM44* | 3.475 | 0.81091 | -0.44185 | -1.5631 | 1.28 |
| 7314 | *IL10* | 4.4052 | 0.24958 | -0.811 | -1.371 | 1.28 |
| 7315 | *SH3BP1* | 4.6749 | 0.18818 | -1.0197 | -2.5349 | 1.28 |
| 7316 | *HIP1* | 4.1992 | 0.35716 | -0.7131 | -0.79813 | 1.28 |
| 7317 | *C3AR1* | 2.4472 | 1.0512 | 0.34426 | -0.21809 | 1.28 |
| 7318 | *VWDE* | 3.4396 | 0.34603 | 0.0552 | -0.00504 | 1.28 |
| 7319 | *AEN* | 3.6013 | 1.4496 | -1.2104 | -1.5121 | 1.28 |
| 7320 | *MIOX* | 3.1999 | 0.60219 | 0.03815 | -1.4151 | 1.28 |
| 7321 | *TOMM40* | 3.41 | 0.33072 | 0.09878 | -1.6374 | 1.28 |
| 7322 | *TRIM32* | 3.6924 | 0.57397 | -0.42743 | -0.47358 | 1.28 |
| 7323 | *CYP2C19* | 4.2287 | 0.42547 | -0.81535 | -1.36 | 1.28 |
| 7324 | *COQ9* | 1.8848 | 1.8362 | 0.11748 | -3.6696 | 1.28 |
| 7325 | *BIRC7* | 2.6827 | 0.76864 | 0.38682 | -1.8604 | 1.28 |
| 7326 | *OR10Q1* | 3.3924 | 1.171 | -0.72635 | -1.088 | 1.28 |
| 7327 | *FBXO17* | 3.3516 | 1.1966 | -0.71161 | -1.2404 | 1.28 |
| 7328 | *GATS* | 2.9259 | 1.1296 | -0.21897 | -1.8224 | 1.28 |
| 7329 | *MIR1-1HG* | 2.784 | 1.5922 | -0.54089 | -1.162 | 1.28 |
| 7330 | *NTN5* | 3.222 | 0.69164 | -0.07903 | -2.2951 | 1.28 |
| 7331 | *FAM76B* | 2.2638 | 1.3582 | 0.21148 | -0.76344 | 1.28 |
| 7332 | *CD300LD* | 3.1184 | 2.4349 | -1.72 | -2.1965 | 1.28 |
| 7333 | *COMMD3-BMI1* | 2.4367 | 1.9696 | -0.57359 | -1.9431 | 1.28 |
| 7334 | *DMRT3* | 3.8487 | 0.66955 | -0.68598 | -2.3904 | 1.28 |
| 7335 | *BGLAP* | 2.8566 | 1.1018 | -0.12651 | -0.16579 | 1.28 |
| 7336 | *EIF4B* | 3.8434 | 0.19975 | -0.21126 | -1.5031 | 1.28 |
| 7337 | *IL32* | 3.4993 | 0.82828 | -0.4958 | -0.66788 | 1.28 |
| 7338 | *REM1* | 2.4931 | 1.8689 | -0.5305 | -1.9417 | 1.28 |
| 7339 | *MMP7* | 3.1422 | 0.80664 | -0.11781 | -0.2122 | 1.28 |
| 7340 | *C11orf94* | 3.8525 | 0.12724 | -0.14995 | -0.2492 | 1.28 |
| 7341 | *TRPM6* | 2.6304 | 0.63006 | 0.56844 | -1.7399 | 1.28 |
| 7342 | *PPM1N* | 3.288 | 0.85474 | -0.31434 | -0.7819 | 1.28 |
| 7343 | *HTR2B* | 2.3363 | 2.1203 | -0.62833 | -1.6526 | 1.28 |
| 7344 | *PRKCZ* | 2.7798 | 0.80835 | 0.23983 | -1.2145 | 1.28 |
| 7345 | *SAMD15* | 5.3648 | -0.43113 | -1.1067 | -1.3023 | 1.28 |
| 7346 | *THAP9* | 3.9943 | 0.20894 | -0.37686 | -1.987 | 1.28 |
| 7347 | *OPN4* | 4.295 | 0.46124 | -0.93011 | -1.0102 | 1.28 |
| 7348 | *KLLN* | 4.0314 | 1.6389 | -1.8443 | -1.8642 | 1.28 |
| 7349 | *ETHE1* | 2.9182 | 0.53553 | 0.37201 | -0.53481 | 1.28 |
| 7350 | *RBP2* | 3.7934 | 0.45284 | -0.42062 | -1.077 | 1.28 |
| 7351 | *CAST* | 2.2702 | 1.5206 | 0.03442 | -0.65754 | 1.28 |
| 7352 | *EEF2K* | 3.8109 | 0.18332 | -0.16908 | -1.0684 | 1.28 |
| 7353 | *KCTD17* | 2.5167 | 1.1462 | 0.16091 | 0.12463 | 1.27 |

| | | | | | | |
|---|---|---|---|---|---|---|
| 7354 | *USP27X* | 3.3499 | 0.81171 | -0.33826 | -0.64327 | 1.27 |
| 7355 | *PHF7* | 2.7336 | 0.60473 | 0.48364 | -1.0187 | 1.27 |
| 7356 | *BARX1* | 3.4674 | 0.37307 | -0.02108 | -0.0279 | 1.27 |
| 7357 | *ZNF816* | 3.7493 | 0.43589 | -0.36624 | -0.72789 | 1.27 |
| 7358 | *MLXIPL* | 3.3784 | 0.72221 | -0.28198 | -1.8792 | 1.27 |
| 7359 | *ZBTB2* | 3.1493 | 0.37192 | 0.29708 | -1.6664 | 1.27 |
| 7360 | *PRCP* | 3.0273 | 1.0328 | -0.24196 | -1.2951 | 1.27 |
| 7361 | *CYB5R2* | 1.5482 | 1.3267 | 0.94203 | -0.54875 | 1.27 |
| 7362 | *CCDC15* | 3.0311 | 1.7456 | -0.96 | -1.4757 | 1.27 |
| 7363 | *NBPF11* | 3.7448 | 1.7004 | -1.6288 | -1.9305 | 1.27 |
| 7364 | *CDK3* | 2.4333 | 0.9539 | 0.42909 | -2.0126 | 1.27 |
| 7365 | *CADPS2* | 4.0813 | 0.3841 | -0.64943 | -1.6419 | 1.27 |
| 7366 | *OR10A6* | 3.1119 | 0.49966 | 0.20388 | -0.23883 | 1.27 |
| 7367 | *TNFSF10* | 4.5647 | 0.32564 | -1.0761 | -1.9984 | 1.27 |
| 7368 | *WASF2* | 5.5835 | -0.66718 | -1.1021 | -1.1598 | 1.27 |
| 7369 | *EEA1* | 3.9197 | 0.87193 | -0.97776 | -1.2474 | 1.27 |
| 7370 | *TMEM86B* | 1.7583 | 1.6308 | 0.42399 | -1.151 | 1.27 |
| 7371 | *CPA4* | 2.1052 | 1.2525 | 0.45515 | -0.21486 | 1.27 |
| 7372 | *INPP4B* | 2.3287 | 0.78498 | 0.69873 | -2.0205 | 1.27 |
| 7373 | *TEX28* | 1.9586 | 1.4508 | 0.40221 | -1.6443 | 1.27 |
| 7374 | *ACAN* | 3.6241 | 0.37835 | -0.19089 | -0.6908 | 1.27 |
| 7375 | *FABP3* | 3.4995 | 1.4121 | -1.1005 | -1.5243 | 1.27 |
| 7376 | *MYCN* | 2.6619 | 1.1856 | -0.03643 | -1.4195 | 1.27 |
| 7377 | *HEPN1* | 4.2855 | 0.34932 | -0.82402 | -1.0952 | 1.27 |
| 7378 | *HSD17B11* | 2.9203 | 1.5786 | -0.68813 | -0.91253 | 1.27 |
| 7379 | *TIGD1* | 4.7804 | 0.89452 | -1.8642 | -3.0112 | 1.27 |
| 7380 | *TNNC2* | 3.6783 | 0.6253 | -0.49326 | -1.175 | 1.27 |
| 7381 | *MARCH4* | 4.117 | 0.53571 | -0.8424 | -1.1571 | 1.27 |
| 7382 | *PURB* | 1.6894 | 1.5827 | 0.53818 | -1.5901 | 1.27 |
| 7383 | *TET3* | 5.2761 | 0.61232 | -2.0791 | -2.5754 | 1.27 |
| 7384 | *C9orf47* | 2.4637 | 1.6181 | -0.2732 | -1.4454 | 1.27 |
| 7385 | *GBA2* | 4.1265 | 0.82809 | -1.1461 | -1.8836 | 1.27 |
| 7386 | *NUAK1* | 1.7712 | 1.4397 | 0.59722 | -0.58662 | 1.27 |
| 7387 | *LNPEP* | 2.5259 | 1.1759 | 0.106 | -1.5305 | 1.27 |
| 7388 | *CTSL* | 3.9735 | -0.00014 | -0.16607 | -1.0506 | 1.27 |
| 7389 | *SERTAD3* | 3.5193 | 1.0142 | -0.72725 | -1.6033 | 1.27 |
| 7390 | *SPATA4* | 3.0221 | 0.96975 | -0.18588 | -1.067 | 1.27 |
| 7391 | *ICT1* | 2.5136 | 0.78731 | 0.50439 | -2.2825 | 1.27 |
| 7392 | *NDP* | 1.6002 | 1.1997 | 1.0041 | -1.0924 | 1.27 |
| 7393 | *AMBN* | 1.9337 | 1.5244 | 0.34501 | -0.09546 | 1.27 |
| 7394 | *IMPACT* | 1.9654 | 1.6006 | 0.23698 | 0.18324 | 1.27 |
| 7395 | *IL18BP* | 2.5686 | 2.1647 | -0.93114 | -1.251 | 1.27 |
| 7396 | *NUP93* | 3.5835 | 1.1864 | -0.96776 | -1.3946 | 1.27 |
| 7397 | *PLEKHH3* | 2.9026 | 0.86412 | 0.03392 | -0.04156 | 1.27 |
| 7398 | *REG1A* | 4.8279 | -0.31688 | -0.71077 | -2.3487 | 1.27 |
| 7399 | *SLC15A3* | 3.5758 | 0.15401 | 0.06988 | -1.6112 | 1.27 |

| | | | | | | |
|---|---|---|---|---|---|---|
| 7400 | *ZNF429* | 2.7101 | 1.6659 | -0.5765 | -1.2481 | 1.27 |
| 7401 | *LTN1* | 3.5872 | 0.87634 | -0.66462 | -1.8646 | 1.27 |
| 7402 | *DRICH1* | 1.7906 | 1.2067 | 0.80044 | 0.068299 | 1.27 |
| 7403 | *DEFA6* | 2.8426 | 1.4752 | -0.5203 | -1.025 | 1.27 |
| 7404 | *WDR18* | 4.1254 | 0.16776 | -0.49897 | -0.86066 | 1.26 |
| 7405 | *NDUFS4* | 2.3552 | 1.3266 | 0.1119 | -1.6701 | 1.26 |
| 7406 | *PCDHA11* | 3.1636 | 0.97929 | -0.34959 | -0.83145 | 1.26 |
| 7407 | *COPZ2* | 1.9887 | 0.99686 | 0.80755 | -0.93325 | 1.26 |
| 7408 | *LYST* | 3.5968 | 0.54365 | -0.34886 | -0.91981 | 1.26 |
| 7409 | *TNFSF11* | 3.9835 | 1.5433 | -1.7364 | -2.4386 | 1.26 |
| 7410 | *DCC* | 2.1994 | 1.4427 | 0.14574 | -2.2044 | 1.26 |
| 7411 | *ESYT2* | 3.2331 | 0.35833 | 0.19604 | -1.1932 | 1.26 |
| 7412 | *ASB5* | 3.3699 | 0.24401 | 0.17348 | -0.22952 | 1.26 |
| 7413 | *CTAG1B* | 4.7478 | 0.8258 | -1.7865 | -2.3517 | 1.26 |
| 7414 | *AKAP6* | 2.5696 | 1.7034 | -0.48623 | -0.49984 | 1.26 |
| 7415 | *KIF25* | 2.566 | 1.2403 | -0.0205 | -3.9666 | 1.26 |
| 7416 | *ZCWPW1* | 2.17 | 0.94524 | 0.66962 | -0.41669 | 1.26 |
| 7417 | *C17orf89* | 4.7358 | -0.38912 | -0.56205 | -1.4365 | 1.26 |
| 7418 | *DNER* | 4.0871 | 0.9147 | -1.2174 | -1.3004 | 1.26 |
| 7419 | *CFI* | 2.0293 | 1.7785 | -0.02387 | -0.80654 | 1.26 |
| 7420 | *LATS1* | 5.0234 | 0.24777 | -1.4873 | -1.8297 | 1.26 |
| 7421 | *ARHGEF25* | 2.281 | 1.0478 | 0.45426 | -1.4189 | 1.26 |
| 7422 | *PNMA2* | 2.891 | 1.0227 | -0.13144 | -0.64177 | 1.26 |
| 7423 | *TMEM95* | 2.2372 | 1.0171 | 0.52665 | -1.7935 | 1.26 |
| 7424 | *PI4K2B* | 2.9717 | 0.69654 | 0.11229 | -1.0884 | 1.26 |
| 7425 | *FDFT1* | 3.0014 | 0.85237 | -0.07334 | -0.65444 | 1.26 |
| 7426 | *TEKT4* | 3.2736 | 0.79994 | -0.29314 | -3.887 | 1.26 |
| 7427 | *LYG2* | 4.098 | 0.15963 | -0.47757 | -0.48808 | 1.26 |
| 7428 | *OSTN* | 2.7748 | 0.71822 | 0.28645 | -1.2568 | 1.26 |
| 7429 | *ZBTB44* | 2.539 | 1.0229 | 0.21736 | -2.4862 | 1.26 |
| 7430 | *N4BP2L1* | 4.7873 | 0.15282 | -1.162 | -1.2772 | 1.26 |
| 7431 | *NDUFV3* | 3.0943 | 0.56322 | 0.12053 | -0.48131 | 1.26 |
| 7432 | *MAS1L* | 2.1671 | 1.4249 | 0.186 | -1.3141 | 1.26 |
| 7433 | *NSMF* | 4.7535 | 0.020642 | -0.99644 | -1.9984 | 1.26 |
| 7434 | *SRRM5* | 3.0553 | 0.3799 | 0.34244 | -0.53931 | 1.26 |
| 7435 | *ARNTL2* | 4.6861 | 0.70719 | -1.6158 | -1.6867 | 1.26 |
| 7436 | *RPA1* | 3.4039 | 1.4928 | -1.1206 | -1.2466 | 1.26 |
| 7437 | *TRIM6-TRIM34* | 3.3736 | 1.1492 | -0.74671 | -1.1271 | 1.26 |
| 7438 | *TMEM180* | 3.1813 | 1.8678 | -1.2736 | -2.213 | 1.26 |
| 7439 | *ZNF182* | 3.3824 | 0.55728 | -0.16483 | -0.72625 | 1.26 |
| 7440 | *SIX1* | 2.4606 | 1.7652 | -0.45106 | -0.73844 | 1.26 |
| 7441 | *ABCB1* | 3.4826 | 0.74915 | -0.45705 | -1.1316 | 1.26 |
| 7442 | *LRRC58* | 2.836 | 1.1714 | -0.23523 | -1.1815 | 1.26 |
| 7443 | *ODF1* | 3.6692 | 0.53391 | -0.43101 | -1.9867 | 1.26 |
| 7444 | *CSRP3* | 3.4898 | 0.14582 | 0.13631 | 0.024092 | 1.26 |
| 7445 | *APAF1* | 2.9643 | 0.74443 | 0.06152 | -0.68729 | 1.26 |

| | | | | | | |
|---|---|---|---|---|---|---|
| 7446 | *MXI1* | 3.888 | 0.59146 | -0.70952 | -1.0005 | 1.26 |
| 7447 | *HIPK4* | 2.8679 | 1.0903 | -0.18962 | -2.8684 | 1.26 |
| 7448 | *GHRHR* | 3.0952 | 1.5069 | -0.83436 | -1.027 | 1.26 |
| 7449 | *SH2D4A* | 3.6303 | 0.074813 | 0.06259 | -2.4955 | 1.26 |
| 7450 | *NLGN3* | 4.217 | 0.38255 | -0.83222 | -1.5096 | 1.26 |
| 7451 | *CNEP1R1* | 4.3634 | 0.73803 | -1.3343 | -1.5658 | 1.26 |
| 7452 | *PIAS4* | 2.7763 | 2.0204 | -1.0298 | -1.9661 | 1.26 |
| 7453 | *DSG2* | 3.9832 | 0.097096 | -0.31475 | -0.33232 | 1.26 |
| 7454 | *CPS1* | 2.8934 | 1.1887 | -0.31698 | -2.0593 | 1.26 |
| 7455 | *HEG1* | 2.6052 | 0.59473 | 0.56468 | 0.12836 | 1.25 |
| 7456 | *PLA2R1* | 3.679 | 0.27668 | -0.19107 | -0.49657 | 1.25 |
| 7457 | *YJEFN3* | 2.606 | 0.96659 | 0.19033 | -0.83497 | 1.25 |
| 7458 | *P4HA3* | 1.5896 | 1.332 | 0.84117 | -0.93273 | 1.25 |
| 7459 | *ZNF490* | 5.0194 | -0.11759 | -1.1391 | -2.3389 | 1.25 |
| 7460 | *MTRNR2L2* | 4.9107 | -0.12446 | -1.0236 | -1.433 | 1.25 |
| 7461 | *UCHL5* | 2.1662 | 1.102 | 0.49431 | -1.7829 | 1.25 |
| 7462 | *CARD6* | 2.8008 | 1.2974 | -0.33659 | -1.3053 | 1.25 |
| 7463 | *CREB3L2* | 3.9675 | 0.42895 | -0.63486 | -1.1679 | 1.25 |
| 7464 | *AIFM3* | 2.1451 | 1.6968 | -0.08064 | -1.1135 | 1.25 |
| 7465 | *MZT2B* | 4.904 | 0.80776 | -1.9513 | -2.5865 | 1.25 |
| 7466 | *FSCN1* | 3.9889 | 0.41801 | -0.64671 | -1.3218 | 1.25 |
| 7467 | *RSRC1* | 3.4765 | 0.1642 | 0.11931 | -0.85392 | 1.25 |
| 7468 | *FBXL20* | 2.5393 | 0.68934 | 0.53093 | -0.09021 | 1.25 |
| 7469 | *LCE2A* | 4.0752 | 0.71306 | -1.0293 | -1.1395 | 1.25 |
| 7470 | *ADM* | 3.602 | 0.30735 | -0.15058 | -1.3983 | 1.25 |
| 7471 | *CEND1* | 2.3513 | 1.6176 | -0.21126 | -1.9295 | 1.25 |
| 7472 | *TP53AIP1* | 3.058 | 1.3264 | -0.62684 | -1.4841 | 1.25 |
| 7473 | *FSTL1* | 1.5699 | 1.1381 | 1.0487 | -2.2065 | 1.25 |
| 7474 | *DDX28* | 3.7847 | 0.006683 | -0.03623 | -0.51023 | 1.25 |
| 7475 | *MPST* | 3.4612 | 0.80292 | -0.51023 | -1.098 | 1.25 |
| 7476 | *SLC33A1* | 2.6001 | 2.2148 | -1.0611 | -1.6622 | 1.25 |
| 7477 | *CLDN20* | 2.0501 | 1.6151 | 0.08818 | -1.0576 | 1.25 |
| 7478 | *CD244* | 3.3221 | 0.26954 | 0.16167 | -1.2531 | 1.25 |
| 7479 | *HILPDA* | 3.1641 | 0.94374 | -0.35499 | -0.68811 | 1.25 |
| 7480 | *ELMSAN1* | 2.3851 | 1.5862 | -0.21868 | -1.6726 | 1.25 |
| 7481 | *ATP1A1* | 3.0466 | 0.35864 | 0.34715 | -0.68259 | 1.25 |
| 7482 | *WTAP* | 2.3733 | 1.0972 | 0.28182 | -1.4717 | 1.25 |
| 7483 | *RAB3D* | 3.4305 | 1.8553 | -1.5382 | -1.8589 | 1.25 |
| 7484 | *RNF185* | 5.3076 | -0.20784 | -1.3523 | -1.4997 | 1.25 |
| 7485 | *HNRNPH3* | 1.8727 | 1.832 | 0.04272 | 0.028035 | 1.25 |
| 7486 | *IL34* | 3.9402 | 0.70459 | -0.89752 | -1.3246 | 1.25 |
| 7487 | *ACTL6A* | 3.3999 | 0.71489 | -0.36971 | -0.56818 | 1.25 |
| 7488 | *CLDN18* | 5.1547 | -0.67149 | -0.73814 | -1.9534 | 1.25 |
| 7489 | *ZBTB21* | 4.1903 | -0.15399 | -0.29139 | -0.31916 | 1.25 |
| 7490 | *PFDN1* | 3.8079 | 0.058704 | -0.12212 | -2.0066 | 1.25 |
| 7491 | *PM20D2* | 2.2537 | 1.0798 | 0.40995 | 0.075935 | 1.25 |

| | | | | | | |
|---|---|---|---|---|---|---|
| 7492 | *PARM1* | 4.0101 | 0.52402 | -0.79149 | -1.7526 | 1.25 |
| 7493 | *FOXD4L4* | 3.9113 | 0.13896 | -0.3077 | -1.6014 | 1.25 |
| 7494 | *HMGCL* | 2.2539 | 1.0669 | 0.42137 | -0.93415 | 1.25 |
| 7495 | *ARFIP2* | 2.2898 | 1.7296 | -0.27771 | -1.5436 | 1.25 |
| 7496 | *ESRRB* | 2.5347 | 0.95373 | 0.25246 | -0.7443 | 1.25 |
| 7497 | *GBGT1* | 2.7761 | 1.0264 | -0.062 | -0.08586 | 1.25 |
| 7498 | *GHSR* | 3.8415 | 0.0528 | -0.15545 | -0.43919 | 1.25 |
| 7499 | *FAM184B* | 4.7544 | 0.037251 | -1.0529 | -1.1344 | 1.25 |
| 7500 | *NR3C2* | 4.0475 | 0.77323 | -1.0824 | -1.1025 | 1.25 |
| 7501 | *TAS2R8* | 3.0925 | 0.64417 | 0.00067 | -1.2141 | 1.25 |
| 7502 | *PRAMEF20* | 2.6339 | 0.77941 | 0.32373 | -2.4244 | 1.25 |
| 7503 | *STXBP2* | 2.509 | 1.2977 | -0.07205 | -0.18488 | 1.24 |
| 7504 | *DBX2* | 2.9045 | 0.6742 | 0.1557 | -1.9794 | 1.24 |
| 7505 | *MRPS5* | 3.0344 | 0.75616 | -0.05693 | -0.95677 | 1.24 |
| 7506 | *OLIG3* | 1.9833 | 1.0072 | 0.742 | -1.6109 | 1.24 |
| 7507 | *TBCB* | 4.635 | 0.06955 | -0.9728 | -1.5865 | 1.24 |
| 7508 | *P2RX7* | 2.1787 | 1.1636 | 0.38938 | -1.1222 | 1.24 |
| 7509 | *AMTN* | 3.5216 | 0.74907 | -0.54271 | -1.0459 | 1.24 |
| 7510 | *RDH10* | 2.6959 | 0.90329 | 0.12843 | -0.65489 | 1.24 |
| 7511 | *H2BFM* | 2.3492 | 1.7099 | -0.33251 | -0.44088 | 1.24 |
| 7512 | *MBNL1* | 4.25 | 0.55825 | -1.0824 | -1.6401 | 1.24 |
| 7513 | *PARP16* | 3.953 | 0.58649 | -0.81381 | -1.908 | 1.24 |
| 7514 | *C20orf24* | 4.9653 | -0.36705 | -0.87318 | -1.8076 | 1.24 |
| 7515 | *CYP4X1* | 3.3465 | 1.9291 | -1.5508 | -2.252 | 1.24 |
| 7516 | *PPFIA2* | 3.5016 | 1.5631 | -1.341 | -1.9099 | 1.24 |
| 7517 | *RABEP2* | 3.4536 | 1.0813 | -0.81147 | -1.0062 | 1.24 |
| 7518 | *BEX4* | 1.8778 | 1.8291 | 0.01647 | -0.07612 | 1.24 |
| 7519 | *RANBP10* | 2.7933 | 0.74308 | 0.18668 | -0.02674 | 1.24 |
| 7520 | *EREG* | 3.0098 | 0.64728 | 0.06592 | -1.2347 | 1.24 |
| 7521 | *PDHA1* | 4.6651 | -0.29292 | -0.64932 | -0.83059 | 1.24 |
| 7522 | *OR13C4* | 2.4838 | 0.6614 | 0.57711 | -2.5004 | 1.24 |
| 7523 | *SPANXN3* | 3.2452 | 1.6189 | -1.1421 | -1.4856 | 1.24 |
| 7524 | *HOXA3* | 2.4074 | 1.0854 | 0.2289 | -2.1616 | 1.24 |
| 7525 | *TET2* | 3.8937 | 0.43341 | -0.60547 | -0.89392 | 1.24 |
| 7526 | *OCSTAMP* | 2.7318 | 1.1676 | -0.17783 | -1.4448 | 1.24 |
| 7527 | *PDXK* | 1.9806 | 1.0588 | 0.68194 | 0.018783 | 1.24 |
| 7528 | *PHF13* | 4.2037 | 0.46168 | -0.9442 | -1.9023 | 1.24 |
| 7529 | *SAMD7* | 3.9873 | 0.17513 | -0.44236 | -1.987 | 1.24 |
| 7530 | *NLRC5* | 2.9814 | 0.39691 | 0.34146 | -0.3221 | 1.24 |
| 7531 | *CACUL1* | 4.1269 | -0.20194 | -0.20571 | -0.36531 | 1.24 |
| 7532 | *C11orf44* | 2.9554 | 0.5809 | 0.18195 | -0.30192 | 1.24 |
| 7533 | *SLC35G6* | 3.9057 | 0.27966 | -0.46731 | -1.6014 | 1.24 |
| 7534 | *LDAH* | 1.4862 | 1.3324 | 0.89945 | 0.34054 | 1.24 |
| 7535 | *ADORA1* | 1.7826 | 1.7589 | 0.17647 | -0.55585 | 1.24 |
| 7536 | *C8orf74* | 5.2678 | 0.48487 | -2.0363 | -2.3424 | 1.24 |
| 7537 | *ZFX* | 2.551 | 0.91279 | 0.252 | -0.87208 | 1.24 |

| | | | | | | |
|---|---|---|---|---|---|---|
| 7538 | *MSX1* | 1.832 | 1.1229 | 0.76079 | 0.66962 | 1.24 |
| 7539 | *POTEG* | 4.0362 | 0.5484 | -0.86925 | -1.4747 | 1.24 |
| 7540 | *C12orf40* | 3.1739 | 1.9659 | -1.4252 | -1.7364 | 1.24 |
| 7541 | *PYGB* | 2.2592 | 1.6714 | -0.21666 | -0.44353 | 1.24 |
| 7542 | *CCL4* | 2.7556 | 0.51145 | 0.44547 | -0.39081 | 1.24 |
| 7543 | *SPESP1* | 2.9145 | 1.3258 | -0.52853 | -0.8604 | 1.24 |
| 7544 | *OR12D3* | 3.9559 | 0.69701 | -0.94119 | -1.1825 | 1.24 |
| 7545 | *TMEM254* | 3.9084 | 0.34706 | -0.54402 | -2.095 | 1.24 |
| 7546 | *FANCE* | 4.4939 | -0.0426 | -0.74015 | -0.75351 | 1.24 |
| 7547 | *AREG* | 2.1569 | 1.7741 | -0.222 | -1.6128 | 1.24 |
| 7548 | *SQLE* | 3.3342 | 0.30309 | 0.0695 | -0.03675 | 1.24 |
| 7549 | *RBL2* | 3.9329 | 0.78604 | -1.0134 | -1.4023 | 1.24 |
| 7550 | *DEFA4* | 1.8247 | 1.5369 | 0.34331 | -0.01323 | 1.23 |
| 7551 | *SERPINA4* | 2.0336 | 1.6278 | 0.04297 | -0.92312 | 1.23 |
| 7552 | *CXCL5* | 3.3152 | 0.42215 | -0.03304 | -0.50429 | 1.23 |
| 7553 | *C1orf94* | 2.4451 | 1.2567 | 0.00192 | -0.59804 | 1.23 |
| 7554 | *CDH2* | 2.8924 | 2.3304 | -1.5194 | -1.8636 | 1.23 |
| 7555 | *APOC1* | 1.6597 | 1.345 | 0.69837 | -0.81163 | 1.23 |
| 7556 | *VBP1* | 3.0113 | 1.5256 | -0.83435 | -1.9541 | 1.23 |
| 7557 | *RPS3* | 4.234 | 0.36888 | -0.9007 | -1.1865 | 1.23 |
| 7558 | *MRPS6* | 4.1727 | -0.21998 | -0.25124 | -1.1958 | 1.23 |
| 7559 | *DCUN1D4* | 3.6564 | 1.505 | -1.4614 | -2.095 | 1.23 |
| 7560 | *PNMT* | 4.1345 | 0.078186 | -0.51302 | -2.5004 | 1.23 |
| 7561 | *PCDHAC2* | 2.5956 | 1.038 | 0.06587 | -2.1687 | 1.23 |
| 7562 | *RNPEP* | 3.9361 | -0.09946 | -0.13753 | -0.80234 | 1.23 |
| 7563 | *MESDC1* | 3.3935 | 0.3665 | -0.06165 | -0.11145 | 1.23 |
| 7564 | *FAM189B* | 3.5654 | 0.29836 | -0.16574 | -0.61483 | 1.23 |
| 7565 | *CCIN* | 4.2266 | -0.03857 | -0.49071 | -2.1906 | 1.23 |
| 7566 | *KRT6B* | 2.7322 | 0.72574 | 0.23822 | -0.02576 | 1.23 |
| 7567 | *CENPM* | 3.8629 | 0.16852 | -0.33538 | -1.7424 | 1.23 |
| 7568 | *SLC44A3* | 3.2916 | 0.97008 | -0.56564 | -1.9844 | 1.23 |
| 7569 | *COL5A2* | 4.8456 | -0.08609 | -1.0635 | -2.3915 | 1.23 |
| 7570 | *PEX26* | 1.6232 | 1.1041 | 0.96862 | 0.19922 | 1.23 |
| 7571 | *DGAT2L6* | 3.8173 | 0.52584 | -0.64748 | -1.2135 | 1.23 |
| 7572 | *TNFRSF10A* | 2.4264 | 1.4704 | -0.20127 | -1.5395 | 1.23 |
| 7573 | *ZNF658* | 2.9437 | 1.1508 | -0.39907 | -1.5663 | 1.23 |
| 7574 | *NUP54* | 4.0097 | 0.069497 | -0.38428 | -1.9023 | 1.23 |
| 7575 | *PSMA8* | 3.9542 | 0.54044 | -0.80144 | -1.9567 | 1.23 |
| 7576 | *CALM1* | 1.5327 | 1.3695 | 0.7883 | -0.21814 | 1.23 |
| 7577 | *SP3* | 2.8959 | 1.7157 | -0.92131 | -1.1145 | 1.23 |
| 7578 | *C9orf135* | 3.6172 | 1.6316 | -1.5587 | -1.6776 | 1.23 |
| 7579 | *CCNE1* | 1.6988 | 1.3042 | 0.6871 | 0.13805 | 1.23 |
| 7580 | *ALOX5AP* | 1.9167 | 1.5155 | 0.25747 | -0.09554 | 1.23 |
| 7581 | *CCDC74B* | 2.2626 | 2.0812 | -0.6548 | -1.1222 | 1.23 |
| 7582 | *TEX2* | 2.9742 | 0.78382 | -0.07026 | -1.0799 | 1.23 |
| 7583 | *TMEM126A* | 2.4418 | 0.71297 | 0.53261 | -0.14606 | 1.23 |

| | | | | | | |
|---|---|---|---|---|---|---|
| 7584 | *GBP7* | 3.35 | 0.30932 | 0.02789 | -1.7805 | 1.23 |
| 7585 | *HNRNPUL2* | 2.8776 | 2.5877 | -1.7783 | -2.4829 | 1.23 |
| 7586 | *MYOZ2* | 2.8397 | 0.44248 | 0.40476 | -0.15226 | 1.23 |
| 7587 | *SSR4* | 2.3944 | 2.1694 | -0.87706 | -1.6696 | 1.23 |
| 7588 | *C21orf91* | 2.2786 | 1.6277 | -0.21993 | -1.329 | 1.23 |
| 7589 | *PIEZO2* | 3.4273 | 1.2679 | -1.0089 | -1.1848 | 1.23 |
| 7590 | *SYN2* | 1.8587 | 1.5494 | 0.27752 | -1.2421 | 1.23 |
| 7591 | *CT47A8* | 2.4949 | 1.9063 | -0.71588 | -1.7591 | 1.23 |
| 7592 | *RNF34* | 4.5254 | 0.38113 | -1.222 | -3.301 | 1.23 |
| 7593 | *C6orf163* | 1.5792 | 1.1047 | 1.0002 | 0.7564 | 1.23 |
| 7594 | *CDKN3* | 2.8091 | 1.5606 | -0.68606 | -2.2413 | 1.23 |
| 7595 | *KCNA6* | 2.7747 | 1.2149 | -0.30609 | -3.4888 | 1.23 |
| 7596 | *GLTP* | 2.3304 | 0.99354 | 0.35935 | -1.7676 | 1.23 |
| 7597 | *FGF22* | 2.9007 | 0.68001 | 0.10175 | -0.45544 | 1.23 |
| 7598 | *AMY1A* | 3.5669 | 0.48657 | -0.37108 | -0.47557 | 1.23 |
| 7599 | *MB21D1* | 1.5224 | 1.2902 | 0.86898 | -0.77184 | 1.23 |
| 7600 | *SIX5* | 3.9615 | 0.014205 | -0.29442 | -1.4507 | 1.23 |
| 7601 | *C1QTNF4* | 1.3946 | 1.2828 | 1.0037 | 0.050642 | 1.23 |
| 7602 | *KBTBD11* | 4.4156 | 0.13052 | -0.86698 | -1.0846 | 1.23 |
| 7603 | *HOXC5* | 2.6564 | 1.6272 | -0.60563 | -1.4154 | 1.23 |
| 7604 | *TMEFF1* | 2.4269 | 0.81814 | 0.43282 | 0.36064 | 1.23 |
| 7605 | *OR8G1* | 2.8937 | 1.1571 | -0.37334 | -0.81947 | 1.23 |
| 7606 | *THSD1* | 1.6556 | 1.4915 | 0.53014 | 0.40872 | 1.23 |
| 7607 | *SZT2* | 2.4534 | 1.0128 | 0.20959 | -1.0995 | 1.23 |
| 7608 | *MAPK4* | 1.9744 | 1.2586 | 0.44215 | -1.067 | 1.23 |
| 7609 | *FBRS* | 3.9921 | 0.88291 | -1.2001 | -1.3479 | 1.22 |
| 7610 | *LHFPL5* | 3.8969 | 0.35544 | -0.57822 | -0.63399 | 1.22 |
| 7611 | *NAV1* | 4.7387 | 0.2261 | -1.2912 | -1.9623 | 1.22 |
| 7612 | *NIT2* | 2.4717 | 1.741 | -0.53931 | -1.9979 | 1.22 |
| 7613 | *PALM* | 3.7225 | 0.14909 | -0.19842 | -0.27924 | 1.22 |
| 7614 | *PPP2R1A* | 1.5062 | 1.3704 | 0.79651 | -2.0069 | 1.22 |
| 7615 | *PAFAH1B1* | 2.7597 | 0.50952 | 0.40308 | -1.772 | 1.22 |
| 7616 | *LIN37* | 4.1139 | -0.00702 | -0.43466 | -0.78203 | 1.22 |
| 7617 | *ADPRH* | 3.7356 | 0.058977 | -0.1227 | -2.3488 | 1.22 |
| 7618 | *SOCS7* | 4.1499 | 0.74421 | -1.2234 | -1.609 | 1.22 |
| 7619 | *GFOD2* | 3.8748 | 0.12479 | -0.32892 | -0.9858 | 1.22 |
| 7620 | *C1orf226* | 3.1537 | 0.66958 | -0.15385 | -0.22068 | 1.22 |
| 7621 | *MARK1* | 2.9547 | 0.93209 | -0.21803 | -0.64574 | 1.22 |
| 7622 | *ABCA9* | 4.1836 | 1.0157 | -1.5309 | -5.3042 | 1.22 |
| 7623 | *ADRA2C* | 4.2347 | -0.06231 | -0.50473 | -2.2844 | 1.22 |
| 7624 | *FRMD1* | 3.4263 | 0.95819 | -0.71717 | -1.5082 | 1.22 |
| 7625 | *SMARCA4* | 3.1024 | 2.0771 | -1.5138 | -1.7342 | 1.22 |
| 7626 | *OR52N5* | 3.9267 | -0.05599 | -0.20537 | -0.79726 | 1.22 |
| 7627 | *PID1* | 2.8047 | 1.1948 | -0.3342 | -1.2084 | 1.22 |
| 7628 | *ZNF502* | 1.3263 | 1.1168 | | | 1.22 |
| 7629 | *RAB19* | 1.9062 | 1.0011 | 0.75727 | -1.0811 | 1.22 |

| | | | | | | |
|---|---|---|---|---|---|---|
| 7630 | *CNIH1* | 4.5338 | -0.03058 | -0.83938 | -2.3424 | 1.22 |
| 7631 | *MAP3K7* | 1.7654 | 1.7615 | 0.13677 | 0.021444 | 1.22 |
| 7632 | *KRT20* | 2.4976 | 1.6933 | -0.52739 | -0.61779 | 1.22 |
| 7633 | *ROS1* | 4.3711 | 0.64807 | -1.3574 | -2.1749 | 1.22 |
| 7634 | *FBXW9* | 2.1593 | 1.9401 | -0.43875 | -0.70466 | 1.22 |
| 7635 | *TNFRSF21* | 3.5704 | 0.048185 | 0.0405 | -0.75778 | 1.22 |
| 7636 | *JOSD2* | 3.7761 | 0.12559 | -0.24368 | -0.65339 | 1.22 |
| 7637 | *MECR* | 2.285 | 1.5238 | -0.15139 | -0.56076 | 1.22 |
| 7638 | *PRICKLE1* | 3.9997 | 0.96682 | -1.3092 | -2.3857 | 1.22 |
| 7639 | *CCP110* | 2.8679 | 2.2831 | -1.4944 | -2.0535 | 1.22 |
| 7640 | *SLC8A1* | 3.4783 | 0.90579 | -0.72764 | -1.9713 | 1.22 |
| 7641 | *HDGF* | 3.6655 | 0.43401 | -0.44353 | -0.97786 | 1.22 |
| 7642 | *RASAL2* | 3.7054 | 0.2372 | -0.28715 | -0.82402 | 1.22 |
| 7643 | *TTI1* | 4.4497 | -0.17402 | -0.62145 | -1.1709 | 1.22 |
| 7644 | *ECHDC1* | 2.1447 | 0.76987 | 0.73907 | -1.8094 | 1.22 |
| 7645 | *BANF2* | 3.7534 | 0.28631 | -0.3863 | -3.1439 | 1.22 |
| 7646 | *LY6G6F* | 3.0467 | 0.38259 | 0.22286 | -1.3205 | 1.22 |
| 7647 | *HDAC8* | 1.9404 | 1.6784 | 0.03318 | -0.06973 | 1.22 |
| 7648 | *ZNF260* | 4.5284 | -0.43407 | -0.44353 | -1.5722 | 1.22 |
| 7649 | *TSPAN31* | 2.4207 | 1.4757 | -0.24569 | -1.8095 | 1.22 |
| 7650 | *SGMS2* | 4.8302 | -0.29739 | -0.88267 | -2.0629 | 1.22 |
| 7651 | *PRG4* | 4.1289 | -0.03528 | -0.44353 | -1.2171 | 1.22 |
| 7652 | *ABCA6* | 4.5922 | 0.1106 | -1.0528 | -1.3585 | 1.22 |
| 7653 | *HGF* | 4.9994 | -0.52601 | -0.82368 | -1.2004 | 1.22 |
| 7654 | *C7orf66* | 3.7537 | 0.70583 | -0.81001 | -2.1315 | 1.22 |
| 7655 | *CHCHD6* | 2.846 | 0.85708 | -0.05396 | -0.13322 | 1.22 |
| 7656 | *OCIAD1* | 4.9851 | -0.44261 | -0.8934 | -1.3053 | 1.22 |
| 7657 | *OR9Q2* | 3.838 | 0.22342 | -0.41239 | -0.83337 | 1.22 |
| 7658 | *PTAR1* | 5.3454 | -0.78959 | -0.90713 | -1.5095 | 1.22 |
| 7659 | *RUFY2* | 2.8545 | 1.1651 | -0.37108 | -1.8779 | 1.22 |
| 7660 | *GK5* | 3.3209 | 1.4758 | -1.1484 | -1.9693 | 1.22 |
| 7661 | *IQSEC2* | 1.7552 | 1.3233 | 0.56967 | -0.95184 | 1.22 |
| 7662 | *EIF1AY* | 3.8213 | 0.14253 | -0.31596 | -1.7567 | 1.22 |
| 7663 | *ST20* | 3.5606 | 0.26445 | -0.17734 | -0.23315 | 1.22 |
| 7664 | *ABHD17C* | 1.9374 | 1.1403 | 0.56986 | -0.22324 | 1.22 |
| 7665 | *CEP57* | 4.3564 | 0.097322 | -0.80636 | -0.85943 | 1.22 |
| 7666 | *TXNRD3NB* | 3.3921 | 0.74036 | -0.48606 | -1.4619 | 1.22 |
| 7667 | *MTCH1* | 2.6069 | 0.91221 | 0.12611 | -2.9211 | 1.22 |
| 7668 | *XAF1* | 3.56 | 0.28611 | -0.20142 | -1.3729 | 1.21 |
| 7669 | *MMRN1* | 3.7171 | 1.2204 | -1.2935 | -1.3346 | 1.21 |
| 7670 | *PDE1B* | 2.8388 | 1.1617 | -0.35665 | -2.3244 | 1.21 |
| 7671 | *ZNF266* | 3.582 | 1.5081 | -1.4474 | -1.6092 | 1.21 |
| 7672 | *PIK3CB* | 2.8509 | 1.9011 | -1.1094 | -1.9094 | 1.21 |
| 7673 | *SNX30* | 2.3018 | 1.7341 | -0.39342 | -1.213 | 1.21 |
| 7674 | *INTS4* | 2.9298 | 0.66252 | 0.05 | -1.316 | 1.21 |
| 7675 | *RIMKLA* | 2.3333 | 0.80747 | 0.50015 | -1.2163 | 1.21 |

| | | | | | | |
|---|---|---|---|---|---|---|
| 7676 | *ABO* | 2.5311 | 0.56905 | 0.54066 | -1.3279 | 1.21 |
| 7677 | *WNT8A* | 3.0294 | 0.5128 | 0.09836 | -2.229 | 1.21 |
| 7678 | *TMEM41A* | 3.533 | 0.90858 | -0.80129 | -1.3864 | 1.21 |
| 7679 | *DOK5* | 3.0494 | 1.0344 | -0.44353 | -1.855 | 1.21 |
| 7680 | *GAS6* | 3.1314 | 0.43955 | 0.06875 | -0.43966 | 1.21 |
| 7681 | *MAPK9* | 4.1381 | -0.06539 | -0.43335 | -0.43587 | 1.21 |
| 7682 | *NUDT16* | 3.1328 | 0.38449 | 0.12206 | -0.74875 | 1.21 |
| 7683 | *FAM47B* | 2.8953 | 2.2742 | -1.5304 | -1.7165 | 1.21 |
| 7684 | *ZFHX4* | 2.7949 | 0.68476 | 0.15937 | -1.4717 | 1.21 |
| 7685 | *C21orf33* | 3.2171 | 0.89646 | -0.47577 | -0.92718 | 1.21 |
| 7686 | *CLEC2L* | 3.9214 | -0.12896 | -0.15499 | -0.7771 | 1.21 |
| 7687 | *AP4M1* | 4.5009 | 0.91475 | -1.7802 | -2.0218 | 1.21 |
| 7688 | *DPH3P1* | 3.518 | 1.3297 | -1.2132 | -1.3132 | 1.21 |
| 7689 | *HYI* | 3.8621 | 0.45661 | -0.68659 | -1.7782 | 1.21 |
| 7690 | *CLIC1* | 2.5553 | 1.4051 | -0.32863 | -1.707 | 1.21 |
| 7691 | *NAALADL1* | 2.8618 | 0.62168 | 0.14824 | -2.3251 | 1.21 |
| 7692 | *OR6C68* | 2.038 | 1.4719 | 0.12154 | -1.6525 | 1.21 |
| 7693 | *SHROOM4* | 2.4852 | 1.4789 | -0.33266 | -1.1451 | 1.21 |
| 7694 | *KIDINS220* | 2.9525 | 0.55812 | 0.11975 | -0.49438 | 1.21 |
| 7695 | *CALR* | 3.6254 | 0.7937 | -0.78899 | -1.8567 | 1.21 |
| 7696 | *LCE3B* | 2.6006 | 0.74132 | 0.28806 | -1.7049 | 1.21 |
| 7697 | *BLM* | 3.2571 | 0.41919 | -0.04635 | -1.9554 | 1.21 |
| 7698 | *RTEL1* | 5.9886 | -1.1067 | -1.2525 | -1.2964 | 1.21 |
| 7699 | *FAM155A* | 3.4128 | 0.29118 | -0.07613 | -0.45644 | 1.21 |
| 7700 | *GPCPD1* | 4.0571 | -0.00833 | -0.42127 | -0.70752 | 1.21 |
| 7701 | *HDAC9* | 2.0212 | 0.92582 | 0.68011 | -0.92975 | 1.21 |
| 7702 | *RAB11FIP4* | 5.1448 | 0.00711 | -1.525 | -1.8085 | 1.21 |
| 7703 | *PROC* | 3.019 | 0.41861 | 0.18898 | -2.6717 | 1.21 |
| 7704 | *RIOK3* | 4.8841 | -0.42195 | -0.83627 | -1.2192 | 1.21 |
| 7705 | *POLR1A* | 3.9454 | 1.1218 | -1.4416 | -1.6078 | 1.21 |
| 7706 | *SLC16A2* | 3.5535 | 0.98197 | -0.91045 | -2.0343 | 1.21 |
| 7707 | *FAM129A* | 2.0291 | 0.93044 | 0.66404 | -1.7947 | 1.21 |
| 7708 | *KIAA1161* | 3.0525 | 0.89569 | -0.32515 | -2.1103 | 1.21 |
| 7709 | *LCK* | 3.5734 | 1.2133 | -1.1637 | -1.6025 | 1.21 |
| 7710 | *ZNF582* | 2.972 | 0.86377 | -0.21291 | -0.25444 | 1.21 |
| 7711 | *C10orf99* | 4.8862 | -0.01286 | -1.2505 | -2.4633 | 1.21 |
| 7712 | *AP3M2* | 2.0813 | 1.2916 | 0.24988 | -1.3542 | 1.21 |
| 7713 | *ATP8A2* | 2.3439 | 1.7327 | -0.45397 | -1.4689 | 1.21 |
| 7714 | *THEM6* | 3.2286 | 0.34157 | 0.05235 | -0.08768 | 1.21 |
| 7715 | *PCBD1* | 1.8841 | 0.92052 | 0.81777 | -0.68535 | 1.21 |
| 7716 | *PLEKHA5* | 2.2283 | 0.73304 | 0.6604 | -0.48247 | 1.21 |
| 7717 | *FAT4* | 2.9011 | 1.3902 | -0.67019 | -0.78229 | 1.21 |
| 7718 | *TFPI* | 1.726 | 1.0102 | 0.8844 | -0.68869 | 1.21 |
| 7719 | *OR5L2* | 3.8543 | 0.091295 | -0.32524 | -1.2111 | 1.21 |
| 7720 | *MUC15* | 4.0048 | 0.50678 | -0.89155 | -1.1468 | 1.21 |
| 7721 | *PTPRS* | 2.3482 | 1.0339 | 0.23731 | -1.7547 | 1.21 |

| | | | | | | |
|---|---|---|---|---|---|---|
| 7722 | *ART5* | 2.0334 | 1.1781 | 0.40742 | -1.8382 | 1.21 |
| 7723 | *AP1M1* | 2.2705 | 0.73002 | 0.61789 | -0.30737 | 1.21 |
| 7724 | *LRRC30* | 2.6656 | 0.82742 | 0.12492 | -1.8349 | 1.21 |
| 7725 | *HRCT1* | 3.2832 | 2.087 | -1.7526 | -1.7782 | 1.21 |
| 7726 | *STK4* | 3.2624 | 1.9262 | -1.5729 | -1.9105 | 1.21 |
| 7727 | *BRK1* | 4.8795 | -0.17509 | -1.0904 | -1.3371 | 1.20 |
| 7728 | *STXBP6* | 3.9697 | 0.19797 | -0.55441 | -0.60701 | 1.20 |
| 7729 | *KLRF2* | 2.702 | 0.97109 | -0.06021 | -1.343 | 1.20 |
| 7730 | *PLEKHM1* | 2.5008 | 1.173 | -0.06253 | -0.6878 | 1.20 |
| 7731 | *TMEM51* | 1.7991 | 1.393 | 0.41913 | -1.2014 | 1.20 |
| 7732 | *ZBTB33* | 2.9012 | 0.56486 | 0.14465 | -0.95896 | 1.20 |
| 7733 | *BLOC1S5* | 4.5332 | -0.33456 | -0.58797 | -1.224 | 1.20 |
| 7734 | *CUL1* | 3.2402 | 0.33218 | 0.03797 | -2.1599 | 1.20 |
| 7735 | *GOPC* | 3.903 | 0.46788 | -0.76083 | -1.2234 | 1.20 |
| 7736 | *LTA* | 4.5875 | -0.4735 | -0.50405 | -1.9623 | 1.20 |
| 7737 | *ACPT* | 2.4732 | 0.83429 | 0.30228 | -2.2125 | 1.20 |
| 7738 | *SNRPB* | 4.1518 | -0.15232 | -0.3908 | -1.4596 | 1.20 |
| 7739 | *CASS4* | 3.4066 | 0.14037 | 0.06168 | -0.69286 | 1.20 |
| 7740 | *POGK* | 4.1071 | 0.50129 | -1.0003 | -1.8478 | 1.20 |
| 7741 | *IFNAR1* | 1.6438 | 1.3949 | 0.56882 | -2.6072 | 1.20 |
| 7742 | *ACVR2A* | 3.2443 | 0.50239 | -0.13938 | -0.81247 | 1.20 |
| 7743 | *ZFHX2* | 4.0595 | 0.80351 | -1.2572 | -1.7645 | 1.20 |
| 7744 | *PERM1* | 4.8335 | -0.10979 | -1.1189 | -1.2694 | 1.20 |
| 7745 | *ZNF681* | 4.0979 | 0.095292 | -0.58894 | -0.68303 | 1.20 |
| 7746 | *SLC4A1* | 1.7242 | 1.6845 | 0.19472 | -2.0895 | 1.20 |
| 7747 | *ALKBH6* | 2.6325 | 2.532 | -1.5611 | -1.725 | 1.20 |
| 7748 | *OR5K2* | 2.8738 | 1.2477 | -0.51845 | -1.0955 | 1.20 |
| 7749 | *METRNL* | 1.5856 | 1.4735 | 0.54394 | -1.1034 | 1.20 |
| 7750 | *COMTD1* | 2.2192 | 1.593 | -0.20946 | -1.4407 | 1.20 |
| 7751 | *CTTNBP2* | 2.3106 | 1.2797 | 0.01243 | -0.72279 | 1.20 |
| 7752 | *SUPT20HL1* | 4.2274 | 0.000118 | -0.62527 | -0.81863 | 1.20 |
| 7753 | *GIPC1* | 5.4437 | -0.08164 | -1.7602 | -1.8157 | 1.20 |
| 7754 | *CDY2B* | 3.3877 | 0.42034 | -0.20663 | -0.24491 | 1.20 |
| 7755 | *ZNF511* | 3.6329 | 0.64987 | -0.68192 | -1.4185 | 1.20 |
| 7756 | *C1orf127* | 3.7862 | 0.17208 | -0.35757 | -0.52494 | 1.20 |
| 7757 | *OLFM1* | 3.0764 | 0.96863 | -0.44459 | -0.74059 | 1.20 |
| 7758 | *TMED3* | 4.3305 | 0.058664 | -0.78959 | -1.337 | 1.20 |
| 7759 | *ZNF528* | 2.9908 | 0.53903 | 0.06919 | -0.05737 | 1.20 |
| 7760 | *MMP16* | 3.147 | 0.5381 | -0.08622 | -0.93002 | 1.20 |
| 7761 | *KLK8* | 4.1207 | -0.09698 | -0.42503 | -0.8458 | 1.20 |
| 7762 | *OR3A3* | 3.068 | 0.96971 | -0.43928 | -0.92742 | 1.20 |
| 7763 | *MUC3A* | 4.3856 | -0.10619 | -0.68136 | -2.3487 | 1.20 |
| 7764 | *SEPHS2* | 2.058 | 2.0458 | -0.50601 | -1.3742 | 1.20 |
| 7765 | *PDZD2* | 1.542 | 1.3183 | 0.73687 | 0.38767 | 1.20 |
| 7766 | *PGPEP1L* | 3.344 | 0.79798 | -0.54484 | -1.1305 | 1.20 |
| 7767 | *GORASP1* | 3.0699 | 0.79324 | -0.26625 | -1.8908 | 1.20 |

| | | | | | | |
|---|---|---|---|---|---|---|
| 7768 | *SIGLEC10* | 2.408 | 1.2453 | -0.0566 | -2.2276 | 1.20 |
| 7769 | *IGFL3* | 3.6303 | 0.22193 | -0.25622 | -0.88053 | 1.20 |
| 7770 | *KRT15* | 4.1728 | -0.28047 | -0.29652 | -1.525 | 1.20 |
| 7771 | *DLG4* | 3.7208 | -0.04218 | -0.08352 | -0.53239 | 1.20 |
| 7772 | *FGF3* | 2.7368 | 0.88904 | -0.03148 | -0.99638 | 1.20 |
| 7773 | *PTPRZ1* | 4.3738 | 0.48467 | -1.2646 | -1.3966 | 1.20 |
| 7774 | *RPS15* | 4.6371 | 0.056101 | -1.0999 | -1.1925 | 1.20 |
| 7775 | *KRT23* | 3.4335 | 0.1389 | 0.02064 | -0.59743 | 1.20 |
| 7776 | *ERBB4* | 2.2745 | 1.1471 | 0.17035 | -0.84158 | 1.20 |
| 7777 | *GTF2B* | 4.3221 | -0.03304 | -0.69735 | -1.7029 | 1.20 |
| 7778 | *SYTL5* | 1.863 | 0.99426 | 0.73388 | -0.93696 | 1.20 |
| 7779 | *PNPLA6* | 4.4164 | -0.04968 | -0.7757 | -2.5153 | 1.20 |
| 7780 | *NT5C1B* | 2.3811 | 1.6049 | -0.3952 | -1.1008 | 1.20 |
| 7781 | *HBM* | 1.8746 | 1.2645 | 0.45091 | -0.90014 | 1.20 |
| 7782 | *MARVELD2* | 2.7847 | 1.1162 | -0.31244 | -1.6707 | 1.20 |
| 7783 | *CCNF* | 2.5574 | 2.5261 | -1.4954 | -1.6718 | 1.20 |
| 7784 | *AKR1C2* | 3.6513 | 0.51054 | -0.57408 | -1.666 | 1.20 |
| 7785 | *DUSP1* | 1.975 | 0.88343 | 0.72823 | -1.0863 | 1.20 |
| 7786 | *GUCY2C* | 2.2808 | 2.1203 | -0.81524 | -2.9931 | 1.20 |
| 7787 | *MYO6* | 2.4758 | 0.99078 | 0.11922 | -0.96433 | 1.20 |
| 7788 | *SUPT3H* | 2.594 | 1.4708 | -0.47929 | -1.072 | 1.20 |
| 7789 | *SFTPD* | 2.7094 | 0.61412 | 0.26182 | -1.5134 | 1.20 |
| 7790 | *C10orf142* | 3.1871 | 1.0122 | -0.6141 | -1.0954 | 1.20 |
| 7791 | *C5orf49* | 3.9599 | 0.32436 | -0.69972 | -1.3053 | 1.19 |
| 7792 | *CHRNA6* | 2.3552 | 1.439 | -0.20969 | -0.30843 | 1.19 |
| 7793 | *IRF5* | 4.1916 | -0.21059 | -0.39673 | -1.6093 | 1.19 |
| 7794 | *ZNF12* | 2.9298 | 0.64084 | 0.01338 | -0.95224 | 1.19 |
| 7795 | *SPATA3* | 1.8031 | 1.1521 | 0.62871 | -1.5269 | 1.19 |
| 7796 | *MCAT* | 3.7567 | 1.154 | -1.327 | -1.4804 | 1.19 |
| 7797 | *NSMCE2* | 3.5467 | 0.31424 | -0.27841 | -0.69006 | 1.19 |
| 7798 | *PINX1* | 2.9273 | 0.85515 | -0.19995 | -1.2027 | 1.19 |
| 7799 | *LIPE* | 2.7558 | 1.9438 | -1.1173 | -1.7117 | 1.19 |
| 7800 | *TIMP2* | 3.4606 | 0.52501 | -0.4046 | -1.8297 | 1.19 |
| 7801 | *FAM136A* | 3.8997 | 0.52416 | -0.84436 | -1.4156 | 1.19 |
| 7802 | *MRGPRD* | 1.9441 | 0.83454 | 0.80057 | -0.86494 | 1.19 |
| 7803 | *SSTR3* | 4.3585 | 0.97965 | -1.7591 | -2.2888 | 1.19 |
| 7804 | *ATP5G3* | 3.9127 | 0.50345 | -0.83713 | -1.448 | 1.19 |
| 7805 | *SUPT16H* | 4.905 | -0.54155 | -0.78482 | -2.1073 | 1.19 |
| 7806 | *DTWD1* | 2.7817 | 0.80823 | -0.01218 | -0.63512 | 1.19 |
| 7807 | *SHE* | 2.7783 | 0.64171 | 0.15724 | -0.44691 | 1.19 |
| 7808 | *RAB39A* | 1.8788 | 1.1576 | 0.53875 | 0.000993 | 1.19 |
| 7809 | *CNPPD1* | 4.2563 | -0.09866 | -0.58315 | -0.70463 | 1.19 |
| 7810 | *GOLGA7* | 4.2551 | -0.15961 | -0.52182 | -1.5243 | 1.19 |
| 7811 | *ST6GALNAC3* | 5.3648 | -0.70442 | -1.0868 | -1.6356 | 1.19 |
| 7812 | *C6orf89* | 2.4026 | 0.69851 | 0.47241 | -1.4039 | 1.19 |
| 7813 | *UQCC3* | 2.7458 | 1.6926 | -0.86494 | -1.3517 | 1.19 |

| | | | | | | |
|---|---|---|---|---|---|---|
| 7814 | *CKS2* | 3.6217 | 1.6821 | -1.7322 | -2.0682 | 1.19 |
| 7815 | *CYP4F2* | 4.5461 | -0.2347 | -0.74003 | -1.9623 | 1.19 |
| 7816 | *WDR34* | 3.388 | 1.458 | -1.2755 | -1.4168 | 1.19 |
| 7817 | *SLC37A4* | 3.3314 | 0.79492 | -0.55641 | -1.8627 | 1.19 |
| 7818 | *KCNQ5* | 2.8983 | 1.1048 | -0.43473 | -1.2448 | 1.19 |
| 7819 | *GAB1* | 3.086 | 0.96745 | -0.48669 | -1.3007 | 1.19 |
| 7820 | *SLC26A8* | 3.851 | 0.85891 | -1.1432 | -1.316 | 1.19 |
| 7821 | *ADGRG7* | 2.4111 | 1.1494 | 0.00598 | -0.27333 | 1.19 |
| 7822 | *ZNF668* | 5.17 | -0.26844 | -1.3353 | -1.3373 | 1.19 |
| 7823 | *ICMT* | 2.769 | 1.1804 | -0.38349 | -1.4048 | 1.19 |
| 7824 | *CRYL1* | 3.5968 | 1.05 | -1.0811 | -1.3643 | 1.19 |
| 7825 | *CD83* | 3.8169 | 0.52736 | -0.77889 | -0.90387 | 1.19 |
| 7826 | *PIEZO1* | 3.9061 | -0.08544 | -0.25611 | -1.4707 | 1.19 |
| 7827 | *KCNA7* | 2.8489 | 1.4099 | -0.69463 | -1.4206 | 1.19 |
| 7828 | *RSRP1* | 2.879 | 0.67899 | 0.00504 | -1.3645 | 1.19 |
| 7829 | *PNPLA1* | 1.9151 | 1.1325 | 0.51482 | -0.86884 | 1.19 |
| 7830 | *B4GAT1* | 1.683 | 1.6501 | 0.22919 | -1.8157 | 1.19 |
| 7831 | *CAPZA1* | 2.0375 | 0.95152 | 0.57307 | -0.39998 | 1.19 |
| 7832 | *TMEM175* | 4.5119 | 0.4967 | -1.4473 | -2.1685 | 1.19 |
| 7833 | *TRPV3* | 4.7069 | -0.25225 | -0.8934 | -1.4636 | 1.19 |
| 7834 | *ZNF568* | 2.5445 | 1.52 | -0.5038 | -1.2755 | 1.19 |
| 7835 | *DVL2* | 3.7444 | 0.57317 | -0.75721 | -2.7427 | 1.19 |
| 7836 | *FOXO1* | 2.5371 | 0.72283 | 0.30033 | -0.09792 | 1.19 |
| 7837 | *CNPY1* | 1.9983 | 1.7146 | -0.1534 | -1.3694 | 1.19 |
| 7838 | *ACSL6* | 4.2005 | -0.01037 | -0.63112 | -1.4014 | 1.19 |
| 7839 | *MYOM3* | 3.805 | 0.62155 | -0.86775 | -1.4173 | 1.19 |
| 7840 | *SLC35E2B* | 2.7305 | 1.6926 | -0.86479 | -1.5382 | 1.19 |
| 7841 | *CLPX* | 2.4842 | 0.8415 | 0.23161 | -1.1848 | 1.19 |
| 7842 | *CLK3* | 4.6162 | -0.5196 | -0.53963 | -0.88458 | 1.19 |
| 7843 | *SEL1L2* | 4.417 | 1.1015 | -1.962 | -2.3086 | 1.19 |
| 7844 | *HECTD3* | 3.5749 | 0.40139 | -0.42014 | -0.54345 | 1.19 |
| 7845 | *PINLYP* | 5.6103 | -0.36751 | -1.687 | -1.8631 | 1.19 |
| 7846 | *CTNND2* | 5.7283 | -0.99768 | -1.175 | -1.7061 | 1.19 |
| 7847 | *INPP5J* | 2.8375 | 1.1074 | -0.38933 | -0.53947 | 1.19 |
| 7848 | *RPAIN* | 4.8796 | -0.64164 | -0.68303 | -2.1794 | 1.18 |
| 7849 | *ZFP62* | 3.3605 | 1.8761 | -1.6824 | -1.9023 | 1.18 |
| 7850 | *KIF18B* | 4.5331 | 0.19497 | -1.1739 | -1.7072 | 1.18 |
| 7851 | *DAPK3* | 3.0869 | 0.75028 | -0.28371 | -0.72725 | 1.18 |
| 7852 | *SPTSSA* | 3.2027 | 0.95268 | -0.60223 | -2.2381 | 1.18 |
| 7853 | *B4GALT4* | 4.8653 | -0.54329 | -0.76934 | -2.6964 | 1.18 |
| 7854 | *FAM98C* | 2.3687 | 1.097 | 0.08672 | -1.57 | 1.18 |
| 7855 | *BNIPL* | 2.6946 | 1.9988 | -1.141 | -1.1935 | 1.18 |
| 7856 | *SNRPN* | 3.9515 | -0.12926 | -0.26999 | -0.7015 | 1.18 |
| 7857 | *BBS5* | 3.0727 | 0.91787 | -0.43841 | -1.057 | 1.18 |
| 7858 | *OR1A1* | 2.1699 | 1.6729 | -0.29089 | -1.2426 | 1.18 |
| 7859 | *NCF2* | 3.1004 | 0.30209 | 0.14941 | -0.06021 | 1.18 |

| | | | | | | |
|---|---|---|---|---|---|---|
| 7860 | *PLCG1* | 3.4909 | 0.31685 | -0.2562 | -1.2364 | 1.18 |
| 7861 | *IL18RAP* | 2.5625 | 1.6373 | -0.64851 | -1.1665 | 1.18 |
| 7862 | *YBX1* | 2.5944 | 2.2089 | -1.2525 | -1.6542 | 1.18 |
| 7863 | *C16orf97* | 2.0421 | 1.1401 | 0.36806 | -1.6712 | 1.18 |
| 7864 | *NME7* | 1.8375 | 1.3748 | 0.33789 | -0.01628 | 1.18 |
| 7865 | *PEX5L* | 3.2328 | 0.25015 | 0.06683 | 0.045836 | 1.18 |
| 7866 | *STEAP3* | 3.8053 | 0.022271 | -0.27844 | -0.72142 | 1.18 |
| 7867 | *MGC57346-CRHR1* | 4.5951 | -0.44661 | -0.60003 | -2.3552 | 1.18 |
| 7868 | *ZNF572* | 4.2756 | 0.15744 | -0.88465 | -1.448 | 1.18 |
| 7869 | *PDK3* | 3.4386 | 0.37011 | -0.26093 | -1.5954 | 1.18 |
| 7870 | *URGCP* | 3.9964 | 0.05116 | -0.50067 | -1.4509 | 1.18 |
| 7871 | *MED10* | 3.3362 | 0.57164 | -0.36165 | -0.70315 | 1.18 |
| 7872 | *CORO7* | 3.0049 | 1.458 | -0.91676 | -1.0935 | 1.18 |
| 7873 | *DUPD1* | 3.6617 | 0.28741 | -0.40305 | -0.97791 | 1.18 |
| 7874 | *ZNF99* | 2.414 | 0.90689 | 0.22402 | -1.3554 | 1.18 |
| 7875 | *TAF1B* | 3.0567 | 0.74596 | -0.25802 | -0.52105 | 1.18 |
| 7876 | *KHDC1L* | 4.4575 | -0.18092 | -0.73202 | -1.5082 | 1.18 |
| 7877 | *KCNG2* | 3.0095 | 0.73559 | -0.20105 | -1.9107 | 1.18 |
| 7878 | *BCO2* | 3.1467 | 1.1578 | -0.76055 | -1.5022 | 1.18 |
| 7879 | *CHST4* | 1.6541 | 1.2003 | 0.68939 | -0.15375 | 1.18 |
| 7880 | *TBKBP1* | 2.9558 | 0.33012 | 0.25756 | 0.20167 | 1.18 |
| 7881 | *MTSS1* | 2.662 | 1.1939 | -0.31487 | -1.3183 | 1.18 |
| 7882 | *BEST3* | 3.4017 | 0.43825 | -0.30003 | -0.96173 | 1.18 |
| 7883 | *HBZ* | 2.885 | 0.75115 | -0.09681 | -0.54121 | 1.18 |
| 7884 | *LMO4* | 1.4053 | 1.1664 | 0.96756 | 0.55616 | 1.18 |
| 7885 | *SLC39A5* | 2.8813 | 0.85912 | -0.20181 | -2.6921 | 1.18 |
| 7886 | *MS4A6E* | 3.0334 | 1.4498 | -0.94546 | -1.1598 | 1.18 |
| 7887 | *CYB5RL* | 2.4309 | 1.2483 | -0.14166 | -0.78229 | 1.18 |
| 7888 | *ANKRD7* | 2.9901 | 1.7682 | -1.2211 | -2.7629 | 1.18 |
| 7889 | *NAA11* | 3.3344 | 1.1134 | -0.91096 | -1.3294 | 1.18 |
| 7890 | *ABHD17B* | 3.9662 | 0.092673 | -0.52255 | -1.3207 | 1.18 |
| 7891 | *BEND2* | 2.1465 | 1.0699 | 0.31982 | -0.66505 | 1.18 |
| 7892 | *CERKL* | 2.0343 | 1.3282 | 0.17236 | -0.20958 | 1.18 |
| 7893 | *FAM222B* | 4.898 | -0.25295 | -1.1105 | -2.7374 | 1.18 |
| 7894 | *ZNF432* | 2.6355 | 0.49518 | 0.40372 | -1.7802 | 1.18 |
| 7895 | *POFUT2* | 2.3816 | 1.1659 | -0.01388 | -0.9363 | 1.18 |
| 7896 | *PSG7* | 2.9815 | 1.1732 | -0.62154 | -1.2071 | 1.18 |
| 7897 | *LZTFL1* | 2.8355 | 0.64793 | 0.04902 | -2.4435 | 1.18 |
| 7898 | *FAM200A* | 2.5896 | 1.4412 | -0.49902 | -1.135 | 1.18 |
| 7899 | *HLA-E* | 2.6848 | 1.0852 | -0.23916 | -0.94576 | 1.18 |
| 7900 | *CD44* | 2.2946 | 0.6802 | 0.55551 | 0.22548 | 1.18 |
| 7901 | *MASP1* | 4.3936 | 0.047789 | -0.91215 | -1.1642 | 1.18 |
| 7902 | *PRB4* | 3.2906 | 1.4652 | -1.2267 | -1.7134 | 1.18 |
| 7903 | *RPS6KA4* | 1.6585 | 1.2771 | 0.59348 | -1.6597 | 1.18 |
| 7904 | *ASB3* | 3.0242 | 0.34095 | 0.16383 | -0.29114 | 1.18 |
| 7905 | *AP5B1* | 5.3973 | -0.48654 | -1.3818 | -1.4607 | 1.18 |

| | | | | | | |
|---|---|---|---|---|---|---|
| 7906 | *IMMP1L* | 3.1841 | 0.37761 | -0.03288 | -0.95224 | 1.18 |
| 7907 | *P2RY13* | 2.4911 | 2.262 | -1.2244 | -1.2958 | 1.18 |
| 7908 | *UGCG* | 4.1891 | 0.012544 | -0.67313 | -1.1853 | 1.18 |
| 7909 | *GPATCH11* | 4.6211 | 0.15112 | -1.2442 | -1.8157 | 1.18 |
| 7910 | *EPX* | 3.7398 | 0.66159 | -0.87366 | -1.258 | 1.18 |
| 7911 | *APOB* | 2.4046 | 0.57679 | 0.54514 | -1.3481 | 1.18 |
| 7912 | *MEGF8* | 2.1314 | 1.4935 | -0.09869 | -1.2775 | 1.18 |
| 7913 | *TMEM54* | 2.456 | 2.0735 | -1.0036 | -4.2747 | 1.18 |
| 7914 | *UNC5CL* | 3.8173 | 0.29852 | -0.59128 | -2.5056 | 1.17 |
| 7915 | *EFCAB7* | 1.6475 | 1.1684 | 0.70854 | -0.73443 | 1.17 |
| 7916 | *GNB3* | 2.228 | 1.1924 | 0.10343 | -1.1519 | 1.17 |
| 7917 | *CLDND1* | 4.0786 | 0.093218 | -0.64808 | -1.9397 | 1.17 |
| 7918 | *KCNK7* | 3.7892 | 0.14449 | -0.41079 | -1.5696 | 1.17 |
| 7919 | *EYA1* | 3.6903 | 1.0558 | -1.2242 | -1.3255 | 1.17 |
| 7920 | *SLC22A24* | 2.7919 | 0.76967 | -0.04015 | -1.1319 | 1.17 |
| 7921 | *EXT1* | 3.9866 | 0.10415 | -0.56961 | -0.92718 | 1.17 |
| 7922 | *HBD* | 4.2548 | 0.040234 | -0.77398 | -1.4536 | 1.17 |
| 7923 | *HIST1H2BB* | 3.9282 | 0.17732 | -0.58495 | -0.8897 | 1.17 |
| 7924 | *KIR2DL1* | 2.7663 | 0.60896 | 0.14487 | -2.3244 | 1.17 |
| 7925 | *TEX29* | 2.5088 | 0.60803 | 0.40329 | -0.68133 | 1.17 |
| 7926 | *MRPL30* | 3.6515 | 0.27162 | -0.40339 | -1.232 | 1.17 |
| 7927 | *MRPL51* | 4.9331 | 0.048185 | -1.463 | -1.6826 | 1.17 |
| 7928 | *OR4M2* | 3.2959 | 0.27944 | -0.05715 | -2.223 | 1.17 |
| 7929 | *DCPS* | 2.7232 | 1.0098 | -0.21487 | -1.7268 | 1.17 |
| 7930 | *SKIV2L2* | 4.0613 | 1.4077 | -1.9511 | -2.6635 | 1.17 |
| 7931 | *APOL5* | 3.183 | 1.9362 | -1.6023 | -1.9522 | 1.17 |
| 7932 | *ASAH1* | 2.4268 | 1.0614 | 0.02733 | -0.31126 | 1.17 |
| 7933 | *UBE2U* | 3.2339 | 0.30505 | -0.0235 | -0.86725 | 1.17 |
| 7934 | *ZNF770* | 3.232 | 0.1435 | 0.1395 | -1.9559 | 1.17 |
| 7935 | *COX18* | 3.657 | 0.30138 | -0.44353 | -1.5337 | 1.17 |
| 7936 | *MPC2* | 4.6808 | 0.8179 | -1.984 | -2.0007 | 1.17 |
| 7937 | *RAB11FIP1* | 2.2376 | 1.4019 | -0.12485 | -1.4415 | 1.17 |
| 7938 | *KRT85* | 4.3876 | 0.071812 | -0.94511 | -2.0277 | 1.17 |
| 7939 | *UBL4B* | 3.7388 | 0.35683 | -0.58136 | -2.1108 | 1.17 |
| 7940 | *FBXO7* | 3.112 | 0.35566 | 0.04626 | -1.7512 | 1.17 |
| 7941 | *FLJ44635* | 3.5546 | 0.81227 | -0.85296 | -2.3292 | 1.17 |
| 7942 | *COL3A1* | 2.5852 | 1.2116 | -0.28294 | -1.3573 | 1.17 |
| 7943 | *REXO1* | 2.1356 | 1.3166 | 0.06093 | -0.88272 | 1.17 |
| 7944 | *CACNB2* | 3.7682 | 0.43155 | -0.68706 | -0.75665 | 1.17 |
| 7945 | *MROH5* | 1.9667 | 1.2091 | 0.33549 | -2.1816 | 1.17 |
| 7946 | *MAN2A1* | 2.5439 | 0.95306 | 0.01427 | -0.65368 | 1.17 |
| 7947 | *TMUB1* | 3.8827 | 0.002584 | -0.37429 | -1.2024 | 1.17 |
| 7948 | *PHF1* | 4.4652 | 0.36105 | -1.3162 | -1.5225 | 1.17 |
| 7949 | *RASGEF1C* | 4.4783 | -0.15339 | -0.81542 | -2.3577 | 1.17 |
| 7950 | *CHRNB3* | 2.8722 | 1.5431 | -0.90606 | -1.3402 | 1.17 |
| 7951 | *ZPBP2* | 2.5085 | 2.2887 | -1.2884 | -1.6105 | 1.17 |

| | | | | | | |
|---|---|---|---|---|---|---|
| 7952 | *ARNT2* | 2.7145 | 0.78312 | 0.01027 | -1.6935 | 1.17 |
| 7953 | *ANXA8L1* | 3.8224 | 0.51973 | -0.83436 | -2.5125 | 1.17 |
| 7954 | *NPM3* | 2.3679 | 1.2864 | -0.14868 | -1.6663 | 1.17 |
| 7955 | *RFTN1* | 4.2976 | -0.13757 | -0.65453 | -2.9283 | 1.17 |
| 7956 | *LACRT* | 3.1284 | 0.48073 | -0.10397 | -1.2067 | 1.17 |
| 7957 | *XIRP2* | 2.6175 | 1.1079 | -0.22082 | -1.8869 | 1.17 |
| 7958 | *GPC1* | 4.6104 | -0.29271 | -0.81334 | -1.433 | 1.17 |
| 7959 | *ACVR1B* | 4.0561 | -0.09356 | -0.45867 | -1.6664 | 1.17 |
| 7960 | *LRIT2* | 2.4824 | 1.9457 | -0.92485 | -1.2226 | 1.17 |
| 7961 | *AGAP2* | 4.0333 | 0.50033 | -1.0315 | -2.0016 | 1.17 |
| 7962 | *ANKRD27* | 3.2761 | 0.36462 | -0.13947 | -0.29579 | 1.17 |
| 7963 | *CDCA5* | 1.3496 | 1.2433 | 0.90816 | -1.3749 | 1.17 |
| 7964 | *KRT37* | 1.6201 | 1.1862 | 0.69416 | 0.2834 | 1.17 |
| 7965 | *CHMP3* | 3.1337 | 0.18941 | 0.17725 | -0.99182 | 1.17 |
| 7966 | *C20orf144* | 1.638 | 1.0649 | 0.79737 | -0.27548 | 1.17 |
| 7967 | *EXTL2* | 3.565 | 0.44829 | -0.51468 | -1.6688 | 1.17 |
| 7968 | *TEAD3* | 2.7577 | 0.76313 | -0.0224 | -0.73342 | 1.17 |
| 7969 | *PNMA6A* | 3.8416 | 0.16445 | -0.50828 | -1.4151 | 1.17 |
| 7970 | *SIAH1* | 1.6937 | 0.92894 | 0.87499 | -1.5364 | 1.17 |
| 7971 | *NTM* | 3.2406 | 0.66003 | -0.40374 | -2.0682 | 1.17 |
| 7972 | *PKIG* | 3.0787 | 0.58533 | -0.16814 | -1.468 | 1.17 |
| 7973 | *ALDH5A1* | 3.5437 | 0.045209 | -0.09334 | -0.28742 | 1.17 |
| 7974 | *SMAP2* | 2.3269 | 1.0403 | 0.12773 | -1.2226 | 1.16 |
| 7975 | *CNNM4* | 1.9831 | 1.9158 | -0.40406 | -0.7331 | 1.16 |
| 7976 | *RAB18* | 3.5091 | 1.4975 | -1.5121 | -2.0878 | 1.16 |
| 7977 | *PCYT1B* | 4.0377 | 0.24637 | -0.78963 | -1.3198 | 1.16 |
| 7978 | *ZNF540* | 2.3578 | 1.9111 | -0.77475 | -1.0158 | 1.16 |
| 7979 | *CCDC136* | 2.0129 | 0.91268 | 0.56846 | -2.283 | 1.16 |
| 7980 | *NXF1* | 1.7399 | 1.4057 | 0.34841 | -1.3742 | 1.16 |
| 7981 | *FRS3* | 3.2736 | 1.543 | -1.3226 | -1.8157 | 1.16 |
| 7982 | *BRF1* | 1.9682 | 0.82051 | 0.70495 | -1.5568 | 1.16 |
| 7983 | *FGFR4* | 2.7076 | 1.7785 | -0.99277 | -1.9608 | 1.16 |
| 7984 | *CDHR1* | 2.8985 | 1.0383 | -0.44353 | -0.75953 | 1.16 |
| 7985 | *PLSCR1* | 3.3646 | 0.11948 | 0.00876 | -2.4018 | 1.16 |
| 7986 | *ZG16B* | 1.2671 | 1.2623 | 0.96313 | -2.4815 | 1.16 |
| 7987 | *NDUFB5* | 3.4631 | 0.82472 | -0.79548 | -0.79745 | 1.16 |
| 7988 | *ASS1* | 2.4029 | 0.61574 | 0.47368 | -1.1447 | 1.16 |
| 7989 | *HIPK2* | 2.0253 | 1.6855 | -0.21852 | -1.0359 | 1.16 |
| 7990 | *TSHZ2* | 4.9746 | -0.19793 | -1.2846 | -1.8799 | 1.16 |
| 7991 | *C8orf34* | 2.5675 | 0.59519 | 0.32887 | -1.893 | 1.16 |
| 7992 | *PHTF2* | 4.3133 | -0.38495 | -0.43694 | -1.5598 | 1.16 |
| 7993 | *OR9Q1* | 5.1959 | -0.47818 | -1.2264 | -1.7845 | 1.16 |
| 7994 | *ADCK1* | 4.0334 | 0.30562 | -0.8478 | -1.274 | 1.16 |
| 7995 | *OR52K1* | 3.1251 | 0.7875 | -0.42195 | -0.73911 | 1.16 |
| 7996 | *FUNDC2* | 3.1242 | 1.3248 | -0.95893 | -1.9734 | 1.16 |
| 7997 | *MAP3K3* | 3.9514 | 0.82601 | -1.2884 | -1.6711 | 1.16 |

| | | | | | | |
|---|---|---|---|---|---|---|
| 7998 | *HIST1H2BK* | 2.5769 | 1.9847 | -1.0733 | -1.6419 | 1.16 |
| 7999 | *OLFM3* | 4.2513 | 0.41759 | -1.1807 | -1.7512 | 1.16 |
| 8000 | *GNPTG* | 2.472 | 0.64864 | 0.36689 | -1.0033 | 1.16 |
| 8001 | *VSIG10L* | 3.7018 | 0.53353 | -0.74819 | -1.3465 | 1.16 |
| 8002 | *PXDNL* | 3.4351 | 0.68534 | -0.63357 | -0.72095 | 1.16 |
| 8003 | *PBX1* | 2.7387 | 1.5264 | -0.77853 | -1.5426 | 1.16 |
| 8004 | *RRP1B* | 2.365 | 1.0572 | 0.06333 | -1.0795 | 1.16 |
| 8005 | *TRPS1* | 4.5395 | 0.42831 | -1.4824 | -1.7364 | 1.16 |
| 8006 | *PARP6* | 4.209 | 0.44498 | -1.1696 | -1.4524 | 1.16 |
| 8007 | *PDZD4* | 3.2005 | 0.71604 | -0.4323 | -0.72493 | 1.16 |
| 8008 | *PRIMPOL* | 3.7527 | 1.2836 | -1.5522 | -1.8724 | 1.16 |
| 8009 | *LOC388780* | 3.1978 | 0.62119 | -0.33503 | -0.73896 | 1.16 |
| 8010 | *KCTD8* | 2.434 | 0.91407 | 0.1358 | -1.5418 | 1.16 |
| 8011 | *KBTBD12* | 3.8396 | 0.30136 | -0.65768 | -0.7463 | 1.16 |
| 8012 | *UBAC1* | 4.3325 | -0.21563 | -0.63405 | -1.6456 | 1.16 |
| 8013 | *FAM227B* | 4.0901 | 0.031661 | -0.63903 | -0.74568 | 1.16 |
| 8014 | *ZNF280B* | 4.2934 | -0.0509 | -0.7602 | -1.0932 | 1.16 |
| 8015 | *ZZZ3* | 3.5835 | 0.48995 | -0.59217 | -2.0851 | 1.16 |
| 8016 | *SMYD4* | 2.6351 | 1.6453 | -0.79987 | -2.2329 | 1.16 |
| 8017 | *ZNF558* | 3.6574 | 0.1136 | -0.29082 | -1.8405 | 1.16 |
| 8018 | *C1orf54* | 1.7066 | 0.9847 | 0.78807 | -1.3978 | 1.16 |
| 8019 | *CLIC6* | 4.1708 | 0.038212 | -0.73155 | -0.99049 | 1.16 |
| 8020 | *LY6G6C* | 1.8183 | 1.47 | 0.18841 | -0.87312 | 1.16 |
| 8021 | *KLHL3* | 2.6271 | 1.9963 | -1.1468 | -1.9431 | 1.16 |
| 8022 | *SYNDIG1* | 3.512 | 0.51334 | -0.54887 | -1.5647 | 1.16 |
| 8023 | *OR6C1* | 3.4163 | 0.03422 | 0.02538 | -1.3728 | 1.16 |
| 8024 | *TMEM38A* | 2.6231 | 1.1169 | -0.26446 | -2.2578 | 1.16 |
| 8025 | *KLF13* | 3.9723 | -0.12866 | -0.36913 | -0.40962 | 1.16 |
| 8026 | *RGPD5* | 3.7319 | 0.4195 | -0.67707 | -0.78766 | 1.16 |
| 8027 | *KRT5* | 3.4063 | 0.51143 | -0.44353 | -0.60541 | 1.16 |
| 8028 | *TNNI1* | 5.0177 | 0.008772 | -1.5524 | -2.425 | 1.16 |
| 8029 | *LHX8* | 3.26 | 0.47249 | -0.26018 | -0.62489 | 1.16 |
| 8030 | *ARL8B* | 3.3793 | 0.18234 | -0.08993 | -1.8583 | 1.16 |
| 8031 | *C12orf43* | 1.4123 | 1.2007 | 0.85843 | -2.3552 | 1.16 |
| 8032 | *MEI4* | 3.2197 | 0.32373 | -0.07232 | -1.1739 | 1.16 |
| 8033 | *MYO19* | 2.8555 | 0.52622 | 0.08877 | -1.1354 | 1.16 |
| 8034 | *LPPR3* | 4.6794 | -0.18461 | -1.0256 | -1.4682 | 1.16 |
| 8035 | *ATP6V0D2* | 4.8586 | 0.14783 | -1.5375 | -1.5598 | 1.16 |
| 8036 | *EXTL3* | 2.8407 | 0.56773 | 0.06023 | -0.0708 | 1.16 |
| 8037 | *C4orf3* | 3.7833 | 0.29438 | -0.6093 | -1.0889 | 1.16 |
| 8038 | *POU4F3* | 2.2368 | 1.2575 | -0.02594 | -1.0217 | 1.16 |
| 8039 | *C9orf89* | 3.0536 | 1.1324 | -0.71785 | -1.6213 | 1.16 |
| 8040 | *SKOR2* | 2.3554 | 1.4071 | -0.29457 | -0.42279 | 1.16 |
| 8041 | *DCBLD1* | 1.4591 | 1.2843 | 0.72432 | -1.2008 | 1.16 |
| 8042 | *SCGB1D1* | 2.7196 | 0.98327 | -0.23523 | -0.78725 | 1.16 |
| 8043 | *IMPAD1* | 3.968 | -0.04845 | -0.45222 | -0.63187 | 1.16 |

| | | | | | | |
|---|---|---|---|---|---|---|
| 8044 | *H2AFZ* | 2.1443 | 0.73717 | 0.5851 | -0.01839 | 1.16 |
| 8045 | *HSPB1* | 3.44 | 0.73702 | -0.71051 | -1.5235 | 1.16 |
| 8046 | *PPP1R16B* | 3.0853 | 0.89908 | -0.51796 | -2.033 | 1.16 |
| 8047 | *AHSP* | 2.4583 | 0.61099 | 0.39693 | -1.9844 | 1.16 |
| 8048 | *AGT* | 4.8052 | -0.0378 | -1.3026 | -1.4783 | 1.15 |
| 8049 | *CACNA1G* | 4.2035 | 1.0767 | -1.8157 | -2.278 | 1.15 |
| 8050 | *ARHGAP8* | 1.9531 | 0.98367 | 0.52723 | -0.90834 | 1.15 |
| 8051 | *ORAI1* | 2.7074 | 0.69369 | 0.06152 | -2.1523 | 1.15 |
| 8052 | *LYRM2* | 1.4583 | 1.1543 | 0.84982 | 0.66539 | 1.15 |
| 8053 | *MAGEB4* | 3.2428 | 0.22738 | -0.00822 | -0.59136 | 1.15 |
| 8054 | *ZNF740* | 4.0111 | -0.21666 | -0.33268 | -1.0777 | 1.15 |
| 8055 | *MLST8* | 4.082 | -0.07341 | -0.54734 | -2.4858 | 1.15 |
| 8056 | *NRSN1* | 2.8971 | 1.2512 | -0.68811 | -1.3345 | 1.15 |
| 8057 | *ONECUT2* | 4.8326 | -0.48968 | -0.88503 | -2.2888 | 1.15 |
| 8058 | *KRT78* | 2.9034 | 1.1689 | -0.61483 | -2.1143 | 1.15 |
| 8059 | *SSX4* | 3.0154 | 0.81559 | -0.3744 | -1.1126 | 1.15 |
| 8060 | *CBR4* | 2.8019 | 2.1476 | -1.4932 | -3.3669 | 1.15 |
| 8061 | *OTUD7B* | 3.8568 | 0.352 | -0.7532 | -1.8282 | 1.15 |
| 8062 | *ZNF660* | 3.0009 | 0.66964 | -0.21539 | -0.7873 | 1.15 |
| 8063 | *DNM1* | 2.9349 | 0.46691 | 0.05282 | -0.10031 | 1.15 |
| 8064 | *CWH43* | 4.7527 | -0.08123 | -1.217 | -1.6524 | 1.15 |
| 8065 | *TENM2* | 3.5975 | 0.4902 | -0.63348 | -2.1478 | 1.15 |
| 8066 | *LOC100129697* | 2.4002 | 1.8754 | -0.82255 | -0.90922 | 1.15 |
| 8067 | *C9orf41* | 3.0027 | 0.6318 | -0.18229 | -0.20425 | 1.15 |
| 8068 | *NANOS2* | 4.4009 | 0.20541 | -1.1543 | -1.855 | 1.15 |
| 8069 | *SEC61A1* | 3.599 | 0.43086 | -0.57787 | -0.7492 | 1.15 |
| 8070 | *TUBA4A* | 3.1836 | 1.597 | -1.329 | -1.3865 | 1.15 |
| 8071 | *FBXL7* | 4.252 | 0.11283 | -0.91343 | -1.0726 | 1.15 |
| 8072 | *PP2D1* | 2.7785 | 0.98977 | -0.31688 | -2.0476 | 1.15 |
| 8073 | *CAV3* | 2.7209 | 0.80615 | -0.07588 | -0.18786 | 1.15 |
| 8074 | *SLA* | 4.3005 | -0.22469 | -0.62514 | -0.72819 | 1.15 |
| 8075 | *MYD88* | 4.3273 | 0.29629 | -1.1733 | -1.4985 | 1.15 |
| 8076 | *PGGT1B* | 3.3922 | 1.4769 | -1.4192 | -1.7802 | 1.15 |
| 8077 | *PPP1CA* | 1.3753 | 1.0449 | 1.0295 | -0.51886 | 1.15 |
| 8078 | *ADAMTS17* | 4.7623 | -0.19404 | -1.1199 | -1.5526 | 1.15 |
| 8079 | *IRAK2* | 3.2243 | 0.23089 | -0.00686 | -0.14513 | 1.15 |
| 8080 | *OLIG2* | 3.4725 | 0.10278 | -0.12795 | -1.7865 | 1.15 |
| 8081 | *PXT1* | 2.9144 | 0.49825 | 0.03406 | -0.89068 | 1.15 |
| 8082 | *GIPC2* | 2.823 | 1.9433 | -1.3199 | -1.3957 | 1.15 |
| 8083 | *CFAP36* | 4.6832 | 0.046655 | -1.2839 | -1.8751 | 1.15 |
| 8084 | *LETM2* | 3.1507 | 0.28236 | 0.01276 | -0.5584 | 1.15 |
| 8085 | *EVI5L* | 1.8164 | 1.178 | 0.451 | -2.8019 | 1.15 |
| 8086 | *C6* | 2.66 | 0.71412 | 0.07057 | -1.6597 | 1.15 |
| 8087 | *PIGB* | 2.4216 | 2.0386 | -1.0158 | -2.069 | 1.15 |
| 8088 | *PRAC1* | 4.2548 | -0.0391 | -0.77289 | -1.9511 | 1.15 |
| 8089 | *CYTL1* | 2.3983 | 1.7751 | -0.73199 | -0.9858 | 1.15 |

| | | | | | | |
|---|---|---|---|---|---|---|
| 8090 | *KLHL8* | 2.2014 | 0.9997 | 0.24016 | -1.5632 | 1.15 |
| 8091 | *ALPK1* | 2.0876 | 1.0011 | 0.35245 | -1.3996 | 1.15 |
| 8092 | *CGA* | 4.5869 | -0.12108 | -1.0248 | -1.3808 | 1.15 |
| 8093 | *HELZ* | 3.6876 | 0.21634 | -0.46333 | -1.3585 | 1.15 |
| 8094 | *PISD* | 4.1891 | 0.80248 | -1.5524 | -1.9891 | 1.15 |
| 8095 | *SOX18* | 2.7848 | 0.80445 | -0.15141 | -2.9338 | 1.15 |
| 8096 | *WNT7B* | 3.389 | 0.13556 | -0.08748 | -0.2595 | 1.15 |
| 8097 | *TRIM17* | 3.9696 | -0.19524 | -0.33806 | -0.83169 | 1.15 |
| 8098 | *ZDHHC5* | 2.8061 | 1.507 | -0.87725 | -2.746 | 1.15 |
| 8099 | *MTFR1* | 3.1425 | 0.1477 | 0.14546 | -1.3496 | 1.15 |
| 8100 | *PAM16* | 5.2666 | -0.82757 | -1.0041 | -4.1213 | 1.14 |
| 8101 | *SAXO2* | 3.4349 | 0.59593 | -0.59622 | -1.5568 | 1.14 |
| 8102 | *BTBD19* | 2.2559 | 1.0507 | 0.12772 | -0.64707 | 1.14 |
| 8103 | *IL2RB* | 2.7006 | 0.88367 | -0.15085 | -2.2319 | 1.14 |
| 8104 | *SLC25A53* | 3.0913 | 0.47851 | -0.13656 | -1.2019 | 1.14 |
| 8105 | *KRTAP4-8* | 2.3405 | 1.364 | -0.27144 | -1.5315 | 1.14 |
| 8106 | *DTWD2* | 2.7829 | 0.57913 | 0.0708 | 0.052769 | 1.14 |
| 8107 | *ZNF354A* | 2.8512 | 0.38252 | 0.19869 | -1.4206 | 1.14 |
| 8108 | *ZNF470* | 1.6105 | 1.2362 | 0.58504 | -1.808 | 1.14 |
| 8109 | *BOLA1* | 2.9303 | 1.2555 | -0.75504 | -2.6543 | 1.14 |
| 8110 | *PAPL* | 1.7505 | 1.2463 | 0.43325 | -0.53961 | 1.14 |
| 8111 | *ALOX5* | 2.2334 | 0.96268 | 0.2336 | -2.2078 | 1.14 |
| 8112 | *C9orf116* | 4.1878 | -0.04919 | -0.71009 | -1.9619 | 1.14 |
| 8113 | *CCDC85B* | 4.6756 | -0.24946 | -0.99768 | -2.4101 | 1.14 |
| 8114 | *CLRN2* | 4.3759 | -0.22977 | -0.71768 | -1.2426 | 1.14 |
| 8115 | *KIF14* | 2.6527 | 2.2014 | -1.4258 | -2.1155 | 1.14 |
| 8116 | *ZPR1* | 2.743 | 0.42629 | 0.25886 | -0.30608 | 1.14 |
| 8117 | *PCDHB8* | 2.6207 | 0.57608 | 0.23058 | -0.68896 | 1.14 |
| 8118 | *OVOL3* | 2.7189 | 1.1646 | -0.4571 | -1.9116 | 1.14 |
| 8119 | *NAA35* | 2.9074 | 1.5014 | -0.98241 | -1.4789 | 1.14 |
| 8120 | *ANP32B* | 3.9235 | 0.2606 | -0.75778 | -1.3353 | 1.14 |
| 8121 | *MLXIP* | 4.2353 | 0.019024 | -0.82833 | -1.1743 | 1.14 |
| 8122 | *RAX2* | 3.6599 | 0.11094 | -0.34536 | -0.83527 | 1.14 |
| 8123 | *AKR1B10* | 2.2128 | 1.3457 | -0.13369 | -1.0153 | 1.14 |
| 8124 | *MAP4K3* | 3.1743 | 0.66025 | -0.41031 | -1.2438 | 1.14 |
| 8125 | *FBXO44* | 2.7045 | 1.006 | -0.2863 | -0.70618 | 1.14 |
| 8126 | *FLG2* | 3.6416 | 1.468 | -1.6855 | -2.3235 | 1.14 |
| 8127 | *MRPL11* | 3.9111 | 0.90907 | -1.3973 | -2.1261 | 1.14 |
| 8128 | *BUD13* | 4.9948 | 0.1718 | -1.7438 | -2.8085 | 1.14 |
| 8129 | *ZNF131* | 1.6208 | 1.3404 | 0.46135 | -1.5795 | 1.14 |
| 8130 | *ZNF839* | 3.5332 | 1.0354 | -1.1474 | -1.2855 | 1.14 |
| 8131 | *CSAG1* | 1.7738 | 1.5012 | 0.14601 | -0.01918 | 1.14 |
| 8132 | *IRX3* | 3.198 | 0.32903 | -0.10815 | -2.4012 | 1.14 |
| 8133 | *SHISA8* | 2.2197 | 1.2403 | -0.04154 | -0.5893 | 1.14 |
| 8134 | *ACMSD* | 3.2632 | 0.72312 | -0.56963 | -1.4999 | 1.14 |
| 8135 | *USP17L7* | 2.1391 | 1.5739 | -0.2969 | -0.35892 | 1.14 |

| | | | | | | |
|---|---|---|---|---|---|---|
| 8136 | *NOTUM* | 3.1726 | 0.26202 | -0.01938 | -2.2766 | 1.14 |
| 8137 | *APOA1BP* | 3.0399 | 0.32908 | 0.04479 | -1.4082 | 1.14 |
| 8138 | *ZNF526* | 2.9196 | 0.62365 | -0.12998 | -1.3007 | 1.14 |
| 8139 | *PGAP1* | 2.461 | 2.3217 | -1.3702 | -2.1461 | 1.14 |
| 8140 | *BRE* | 3.8963 | 0.70199 | -1.1872 | -1.7658 | 1.14 |
| 8141 | *CHP1* | 3.8538 | 0.021013 | -0.46382 | -1.5 | 1.14 |
| 8142 | *C15orf41* | 2.9883 | 1.1149 | -0.69262 | -1.6386 | 1.14 |
| 8143 | *HOXD8* | 1.8203 | 0.93503 | 0.65508 | -1.0343 | 1.14 |
| 8144 | *C4orf48* | 4.646 | -0.52343 | -0.71238 | -1.1417 | 1.14 |
| 8145 | *ADPRHL2* | 1.8943 | 1.3751 | 0.14058 | -0.99137 | 1.14 |
| 8146 | *WFIKKN1* | 1.6455 | 1.0043 | 0.75948 | 0.17981 | 1.14 |
| 8147 | *TMEM248* | 1.7881 | 1.0263 | 0.59329 | -0.8397 | 1.14 |
| 8148 | *CELF1* | 1.9632 | 1.6195 | -0.17509 | -0.66973 | 1.14 |
| 8149 | *TNS3* | 2.7903 | 2.0731 | -1.4563 | -1.5696 | 1.14 |
| 8150 | *SRRD* | 1.8108 | 1.1106 | 0.48551 | -1.6284 | 1.14 |
| 8151 | *DSTYK* | 3.795 | 0.021801 | -0.41012 | -0.62684 | 1.14 |
| 8152 | *KLF15* | 1.3481 | 1.1016 | 0.95668 | -0.01366 | 1.14 |
| 8153 | *MTRNR2L3* | 3.5298 | 0.40422 | -0.52764 | -1.7251 | 1.14 |
| 8154 | *PLGRKT* | 2.1966 | 0.88742 | 0.32159 | -1.017 | 1.14 |
| 8155 | *C1orf110* | 1.8528 | 1.7029 | -0.15062 | -0.58231 | 1.14 |
| 8156 | *NOTO* | 5.2857 | -0.49724 | -1.3837 | -1.7802 | 1.13 |
| 8157 | *NBEA* | 2.7625 | 0.63882 | 0.00149 | -0.1394 | 1.13 |
| 8158 | *FERMT1* | 2.7251 | 1.0196 | -0.34227 | -1.0494 | 1.13 |
| 8159 | *HAS3* | 3.0566 | 0.26862 | 0.07621 | -1.6007 | 1.13 |
| 8160 | *HIST1H4A* | 3.0266 | 0.87268 | -0.49812 | -1.222 | 1.13 |
| 8161 | *ZSCAN31* | 2.0508 | 0.98974 | 0.36048 | -0.71032 | 1.13 |
| 8162 | *KRTAP5-7* | 2.6924 | 0.38885 | 0.31941 | -0.20946 | 1.13 |
| 8163 | *ADGRL1* | 2.9625 | 0.51107 | -0.07312 | -1.9589 | 1.13 |
| 8164 | *SDCCAG8* | 2.9947 | 0.32118 | 0.08359 | -0.1875 | 1.13 |
| 8165 | *BUB1* | 3.8747 | 1.1091 | -1.5849 | -2.5103 | 1.13 |
| 8166 | *OR6C75* | 4.5992 | -0.08253 | -1.1189 | -1.4064 | 1.13 |
| 8167 | *SPACA7* | 1.5496 | 1.0378 | 0.80961 | -1.122 | 1.13 |
| 8168 | *ARHGAP20* | 2.5109 | 0.74 | 0.14551 | -1.2815 | 1.13 |
| 8169 | *IL6* | 2.0217 | 0.90388 | 0.47012 | -1.9846 | 1.13 |
| 8170 | *NSUN6* | 4.2248 | 0.1475 | -0.97667 | -1.3153 | 1.13 |
| 8171 | *NXPE2* | 2.3158 | 0.56262 | 0.51709 | -0.62862 | 1.13 |
| 8172 | *INAFM1* | 4.204 | 0.42497 | -1.2338 | -1.9338 | 1.13 |
| 8173 | *CHRNB2* | 4.2387 | -0.08672 | -0.75717 | -1.3078 | 1.13 |
| 8174 | *JPH4* | 1.7516 | 1.4127 | 0.22939 | -0.16753 | 1.13 |
| 8175 | *RGSL1* | 2.4804 | 0.46861 | 0.44453 | 0.15487 | 1.13 |
| 8176 | *COX7A1* | 3.3722 | 0.41085 | -0.38975 | -1.0224 | 1.13 |
| 8177 | *FPGT* | 3.09 | 0.8936 | -0.59061 | -1.0364 | 1.13 |
| 8178 | *MMP28* | 2.59 | 0.89123 | -0.08956 | -1.0315 | 1.13 |
| 8179 | *TCHHL1* | 2.6461 | 0.6642 | 0.08106 | -1.5679 | 1.13 |
| 8180 | *RASGEF1B* | 3.8172 | 0.90586 | -1.3322 | -1.505 | 1.13 |
| 8181 | *HOXB2* | 1.3846 | 1.2797 | 0.72654 | -1.6731 | 1.13 |

| | | | | | | |
|---|---|---|---|---|---|---|
| 8182 | *THAP3* | 2.4217 | 1.135 | -0.16614 | -1.0186 | 1.13 |
| 8183 | *ADD3* | 3.6932 | 0.49515 | -0.79821 | -3.1519 | 1.13 |
| 8184 | *PATE3* | 4.8264 | -0.59013 | -0.84687 | -1.5312 | 1.13 |
| 8185 | *CISH* | 3.6429 | 0.31268 | -0.56659 | -0.82663 | 1.13 |
| 8186 | *AKAP7* | 4.3182 | -0.13757 | -0.79282 | -1.6527 | 1.13 |
| 8187 | *ANKRD13D* | 4.9371 | -0.256 | -1.2936 | -1.6119 | 1.13 |
| 8188 | *ERLIN1* | 3.8034 | -0.00829 | -0.40805 | -1.5364 | 1.13 |
| 8189 | *SYNGR4* | 2.5642 | 0.58139 | 0.24131 | 0.077067 | 1.13 |
| 8190 | *DCLK2* | 3.3473 | 0.028984 | 0.01047 | -0.31035 | 1.13 |
| 8191 | *S1PR5* | 2.5726 | 0.84658 | -0.03246 | -1.3554 | 1.13 |
| 8192 | *PLOD2* | 4.4203 | -0.50048 | -0.53406 | -0.81505 | 1.13 |
| 8193 | *CNTF* | 4.9109 | -0.05759 | -1.4685 | -1.7407 | 1.13 |
| 8194 | *OR4C16* | 3.4612 | 0.71758 | -0.79663 | -1.0211 | 1.13 |
| 8195 | *CDH8* | 1.5823 | 1.0849 | 0.71474 | -1.6597 | 1.13 |
| 8196 | *CDA* | 3.223 | 1.293 | -1.135 | -1.2837 | 1.13 |
| 8197 | *TMEM109* | 4.7253 | 0.13174 | -1.4773 | -1.696 | 1.13 |
| 8198 | *CRX* | 2.2558 | 1.3924 | -0.26949 | -1.2412 | 1.13 |
| 8199 | *NICN1* | 3.7807 | 0.065124 | -0.46716 | -0.75619 | 1.13 |
| 8200 | *GPR37L1* | 3.4605 | 0.36129 | -0.44353 | -1.7908 | 1.13 |
| 8201 | *CFAP44* | 2.324 | 0.56974 | 0.4845 | -1.0296 | 1.13 |
| 8202 | *BCL3* | 2.0859 | 0.74962 | 0.54217 | -1.942 | 1.13 |
| 8203 | *SPA17* | 2.6136 | 0.85067 | -0.08719 | -0.18393 | 1.13 |
| 8204 | *PELI1* | 4.2848 | 0.17781 | -1.0858 | -1.6065 | 1.13 |
| 8205 | *LSM12* | 4.0669 | -0.18185 | -0.50837 | -0.78594 | 1.13 |
| 8206 | *CFAP58* | 3.9894 | 0.90538 | -1.5182 | -2.2352 | 1.13 |
| 8207 | *FKRP* | 4.5391 | -0.54851 | -0.61437 | -1.1945 | 1.13 |
| 8208 | *MPO* | 1.9747 | 1.8574 | -0.45628 | -1.8212 | 1.13 |
| 8209 | *TRMT6* | 4.4962 | -0.06464 | -1.0561 | -1.1096 | 1.13 |
| 8210 | *ANGPTL3* | 2.4396 | 1.1459 | -0.21205 | -0.5655 | 1.12 |
| 8211 | *A1BG* | 3.4324 | 0.19996 | -0.25912 | -0.44353 | 1.12 |
| 8212 | *SLC9A3* | 4.4497 | 0.077878 | -1.1547 | -1.5956 | 1.12 |
| 8213 | *CDC23* | 4.3662 | 0.016066 | -1.0094 | -1.9442 | 1.12 |
| 8214 | *PTPN1* | 3.4289 | 1.6428 | -1.6989 | -1.7908 | 1.12 |
| 8215 | *METTL7A* | 4.6248 | -0.57506 | -0.67764 | -0.78482 | 1.12 |
| 8216 | *VPS4A* | 3.1368 | 0.1466 | 0.08735 | -1.7639 | 1.12 |
| 8217 | *CALCRL* | 3.5083 | 0.99725 | -1.1354 | -2.8849 | 1.12 |
| 8218 | *SH3RF2* | 3.4379 | 0.309 | -0.37702 | -1.7646 | 1.12 |
| 8219 | *PKD2* | 1.4277 | 1.1897 | 0.75228 | 0.34745 | 1.12 |
| 8220 | *NRDE2* | 4.8744 | 0.13368 | -1.6386 | -2.0791 | 1.12 |
| 8221 | *MCF2* | 2.9929 | 0.62505 | -0.2486 | -0.61767 | 1.12 |
| 8222 | *ABHD6* | 1.5816 | 1.0477 | 0.73963 | -0.0159 | 1.12 |
| 8223 | *SLC25A27* | 1.8972 | 1.6453 | -0.17383 | -0.57701 | 1.12 |
| 8224 | *RNASE13* | 3.4294 | 0.44696 | -0.50855 | -1.3292 | 1.12 |
| 8225 | *RGR* | 2.1933 | 1.2359 | -0.06182 | -1.9249 | 1.12 |
| 8226 | *DLG2* | 1.6253 | 1.1807 | 0.56093 | -0.7351 | 1.12 |
| 8227 | *PXK* | 3.1026 | 1.7204 | -1.4561 | -1.6913 | 1.12 |

| | | | | | | |
|---|---|---|---|---|---|---|
| 8228 | *ARL14EP* | 1.6741 | 1.6675 | 0.02366 | -0.95708 | 1.12 |
| 8229 | *RPS2* | 3.3803 | -0.00827 | -0.01068 | -2.1347 | 1.12 |
| 8230 | *PTGES3L* | 3.0384 | 0.76604 | -0.44353 | -0.7329 | 1.12 |
| 8231 | *FAM104A* | 3.6122 | 0.67648 | -0.92781 | -2.3036 | 1.12 |
| 8232 | *GAL3ST1* | 2.3903 | 0.72074 | 0.24975 | -0.02864 | 1.12 |
| 8233 | *STX1A* | 3.7678 | -0.01264 | -0.39468 | -1.0126 | 1.12 |
| 8234 | *RNLS* | 3.6332 | -0.0996 | -0.17345 | -0.43674 | 1.12 |
| 8235 | *RPS12* | 2.4757 | 1.8423 | -0.95838 | -1.4809 | 1.12 |
| 8236 | *ADCYAP1* | 2.6516 | 1.4291 | -0.72201 | -2.5878 | 1.12 |
| 8237 | *LAPTM5* | 1.8652 | 1.1042 | 0.38873 | -1.9091 | 1.12 |
| 8238 | *GRIFIN* | 2.7626 | 0.83474 | -0.23931 | -0.32004 | 1.12 |
| 8239 | *CCDC126* | 2.8289 | 1.4298 | -0.9007 | -1.8219 | 1.12 |
| 8240 | *ENTHD1* | 3.3437 | 0.3533 | -0.34014 | -1.7495 | 1.12 |
| 8241 | *RAD54B* | 3.3073 | 1.9234 | -1.8744 | -2.8647 | 1.12 |
| 8242 | *NAA50* | 4.3762 | -0.06876 | -0.9512 | -3.2856 | 1.12 |
| 8243 | *GRAPL* | 2.5253 | 0.72043 | 0.10907 | -0.25484 | 1.12 |
| 8244 | *UTF1* | 5.0169 | -0.74965 | -0.91269 | -4.7884 | 1.12 |
| 8245 | *PI15* | 2.2055 | 1.5761 | -0.42818 | -1.4486 | 1.12 |
| 8246 | *HTN3* | 3.5838 | 0.076911 | -0.3077 | -2.5645 | 1.12 |
| 8247 | *SIRPA* | 4.01 | -0.27376 | -0.38402 | -1.4273 | 1.12 |
| 8248 | *TSPAN33* | 2.4297 | 0.77392 | 0.14852 | -0.227 | 1.12 |
| 8249 | *C7orf25* | 2.6481 | 0.82438 | -0.12153 | -1.3445 | 1.12 |
| 8250 | *ATG10* | 2.0992 | 0.66966 | 0.58006 | 0.27802 | 1.12 |
| 8251 | *OPTC* | 3.217 | 0.70344 | -0.57172 | -0.99574 | 1.12 |
| 8252 | *KRTAP26-1* | 3.2931 | 0.89643 | -0.84158 | -1.2195 | 1.12 |
| 8253 | *FRA10AC1* | 2.335 | 0.67948 | 0.3327 | -0.726 | 1.12 |
| 8254 | *TAF12* | 4.0067 | 0.28371 | -0.94333 | -2.0277 | 1.12 |
| 8255 | *RECK* | 3.1154 | 0.68874 | -0.45727 | -1.8909 | 1.12 |
| 8256 | *XCL1* | 4.9431 | 0.14337 | -1.7397 | -2.5419 | 1.12 |
| 8257 | *KLC1* | 2.6623 | 0.78772 | -0.10329 | -0.53999 | 1.12 |
| 8258 | *MARCH7* | 2.4556 | 1.528 | -0.63691 | -2.588 | 1.12 |
| 8259 | *NCAPH2* | 2.3607 | 0.85271 | 0.13273 | -0.44135 | 1.12 |
| 8260 | *USP17L12* | 2.7914 | 1.1299 | -0.57579 | -0.87208 | 1.12 |
| 8261 | *IQCJ* | 4.8221 | -0.2317 | -1.2451 | -2.4709 | 1.12 |
| 8262 | *PRR11* | 3.6195 | 0.52921 | -0.80395 | -1.2306 | 1.11 |
| 8263 | *NAB2* | 1.6813 | 1.0497 | 0.61354 | 0.49645 | 1.11 |
| 8264 | *RIMBP3C* | 3.4032 | 0.67328 | -0.73219 | -1.0012 | 1.11 |
| 8265 | *GYPC* | 1.6346 | 0.87981 | 0.82888 | -0.61353 | 1.11 |
| 8266 | *CHD3* | 5.0368 | -0.66362 | -1.03 | -1.3473 | 1.11 |
| 8267 | *DAB2* | 3.6681 | 0.31609 | -0.64117 | -2.4151 | 1.11 |
| 8268 | *DPP9-AS1* | 2.1917 | 1.9148 | -0.76366 | -1.4682 | 1.11 |
| 8269 | *AJAP1* | 3.0906 | 0.50002 | -0.24797 | -1.1156 | 1.11 |
| 8270 | *KRTAP12-3* | 2.5106 | 0.7968 | 0.03497 | -0.62372 | 1.11 |
| 8271 | *RAB31* | 2.9204 | 0.85297 | -0.43107 | -1.106 | 1.11 |
| 8272 | *SIGLEC8* | 2.0986 | 1.4923 | -0.24926 | -0.73826 | 1.11 |
| 8273 | *CORO2A* | 2.7446 | 1.2848 | -0.68889 | -1.2475 | 1.11 |

| | | | | | | |
|---|---|---|---|---|---|---|
| 8274 | *EPB41* | 3.4503 | 0.57298 | -0.68303 | -1.8869 | 1.11 |
| 8275 | *PIP* | 2.535 | 1.6504 | -0.84558 | -0.90772 | 1.11 |
| 8276 | *CERS5* | 3.666 | 0.41322 | -0.74086 | -1.8631 | 1.11 |
| 8277 | *TCEB3* | 2.1416 | 0.78541 | 0.41125 | -1.615 | 1.11 |
| 8278 | *EFHD1* | 3.5083 | 0.16218 | -0.33231 | -1.7547 | 1.11 |
| 8279 | *SPATA31C2* | 1.6932 | 1.3427 | 0.30154 | -0.70988 | 1.11 |
| 8280 | *SIX2* | 3.5304 | 1.7306 | -1.9237 | -2.5057 | 1.11 |
| 8281 | *GLUD2* | 2.6745 | 1.14 | -0.47743 | -1.0267 | 1.11 |
| 8282 | *CCDC93* | 2.3806 | 0.57773 | 0.37868 | -1.6984 | 1.11 |
| 8283 | *FIGNL2* | 2.6 | 0.77087 | -0.03476 | -0.91842 | 1.11 |
| 8284 | *FGF4* | 2.0786 | 0.69682 | 0.56035 | -1.2541 | 1.11 |
| 8285 | *CIRH1A* | 5.1245 | -0.85606 | -0.93383 | -1.545 | 1.11 |
| 8286 | *HTR1F* | 3.5514 | -0.06269 | -0.15416 | -1.0564 | 1.11 |
| 8287 | *TRMU* | 3.7969 | 0.67421 | -1.1369 | -1.2511 | 1.11 |
| 8288 | *ALB* | 2.9172 | 1.8698 | -1.4538 | -2.3684 | 1.11 |
| 8289 | *TEDDM1* | 2.6583 | 0.63668 | 0.0369 | -0.90028 | 1.11 |
| 8290 | *GTF2H2C* | 1.9587 | 1.8516 | -0.47864 | -2.6463 | 1.11 |
| 8291 | *IPO7* | 2.7233 | 0.44434 | 0.16383 | -1.0956 | 1.11 |
| 8292 | *EFNB1* | 4.2082 | 0.023261 | -0.90028 | -2.1179 | 1.11 |
| 8293 | *OLFM4* | 3.8505 | 0.38541 | -0.90492 | -1.0277 | 1.11 |
| 8294 | *AMMECR1* | 3.3646 | 0.63264 | -0.66652 | -2.2108 | 1.11 |
| 8295 | *PLEKHG3* | 2.8808 | 0.57025 | -0.12054 | -1.1282 | 1.11 |
| 8296 | *LPP* | 1.5787 | 1.2907 | 0.46058 | -0.03146 | 1.11 |
| 8297 | *CNRIP1* | 3.3743 | 0.003701 | -0.04817 | -0.84844 | 1.11 |
| 8298 | *USP17L22* | 2.3194 | 0.77323 | 0.23632 | -0.07031 | 1.11 |
| 8299 | *RABL6* | 3.1822 | 0.75372 | -0.60701 | -2.4197 | 1.11 |
| 8300 | *TNIK* | 3.9146 | 0.60934 | -1.1953 | -1.7855 | 1.11 |
| 8301 | *FFAR2* | 4.2896 | -0.00355 | -0.95794 | -1.8392 | 1.11 |
| 8302 | *DYSF* | 1.4337 | 1.2124 | 0.6804 | -0.88039 | 1.11 |
| 8303 | *OR2AP1* | 4.0225 | -0.28173 | -0.41561 | -1.2137 | 1.11 |
| 8304 | *CYP4F8* | 2.3637 | 1.331 | -0.36957 | -1.0623 | 1.11 |
| 8305 | *MICALL2* | 3.81 | 0.75411 | -1.2393 | -1.2955 | 1.11 |
| 8306 | *ZNF488* | 4.779 | 0.20146 | -1.6559 | -1.8256 | 1.11 |
| 8307 | *HHAT* | 4.1571 | 0.23996 | -1.0727 | -1.6934 | 1.11 |
| 8308 | *HTRA4* | 2.2167 | 1.8567 | -0.75059 | -1.0769 | 1.11 |
| 8309 | *GSTK1* | 2.0787 | 1.2128 | 0.0303 | -1.5063 | 1.11 |
| 8310 | *TOM1* | 1.7547 | 1.5397 | 0.02597 | -0.2372 | 1.11 |
| 8311 | *RPS5* | 2.9426 | 0.32677 | 0.05003 | -1.1965 | 1.11 |
| 8312 | *DHH* | 3.4075 | 0.003685 | -0.09208 | -1.1977 | 1.11 |
| 8313 | *GLIPR1L1* | 1.5954 | 1.2111 | 0.51192 | -2.7813 | 1.11 |
| 8314 | *BAGE2* | 2.6795 | 1.5038 | -0.86506 | -1.5805 | 1.11 |
| 8315 | *VIMP* | 2.9652 | 1.0838 | -0.73172 | -1.8253 | 1.11 |
| 8316 | *MAN2A2* | 1.8199 | 0.84625 | 0.65096 | -1.175 | 1.11 |
| 8317 | *ZSWIM5* | 2.356 | 1.6588 | -0.69787 | -1.0215 | 1.11 |
| 8318 | *AZIN1* | 3.7466 | 0.70864 | -1.1387 | -3.031 | 1.11 |
| 8319 | *SYNM* | 4.0633 | 0.3674 | -1.1144 | -1.3287 | 1.11 |

| | | | | | | |
|---|---|---|---|---|---|---|
| 8320 | *AFG3L2* | 3.2364 | 0.25096 | -0.17208 | -1.7467 | 1.11 |
| 8321 | *HIST3H2A* | 4.0697 | 1.3872 | -2.1421 | -2.5453 | 1.10 |
| 8322 | *FRK* | 2.8238 | 0.65221 | -0.16176 | -0.73348 | 1.10 |
| 8323 | *HOXC8* | 2.7179 | 0.36628 | 0.2299 | -0.56938 | 1.10 |
| 8324 | *SLC6A17* | 2.4425 | 1.5389 | -0.66773 | -1.0507 | 1.10 |
| 8325 | *KCNMB2* | 3.6917 | 0.032031 | -0.41025 | -0.8903 | 1.10 |
| 8326 | *CACNG1* | 2.5303 | 0.80808 | -0.02502 | -0.02697 | 1.10 |
| 8327 | *IHH* | 4.3186 | 0.061681 | -1.0671 | -1.3264 | 1.10 |
| 8328 | *PDZK1IP1* | 2.2469 | 1.2955 | -0.22972 | -0.41476 | 1.10 |
| 8329 | *DNAJC6* | 4.6637 | -0.198 | -1.154 | -1.6251 | 1.10 |
| 8330 | *WIPF2* | 3.0434 | 0.50218 | -0.23418 | -0.87395 | 1.10 |
| 8331 | *AP3S2* | 3.1248 | 0.14975 | 0.03663 | -2.0116 | 1.10 |
| 8332 | *ERMAP* | 2.7365 | 1.018 | -0.44353 | -1.5128 | 1.10 |
| 8333 | *IQCF3* | 1.392 | 1.1055 | 0.81272 | -0.26161 | 1.10 |
| 8334 | *TRIM60* | 1.6613 | 1.1425 | 0.50615 | -0.7623 | 1.10 |
| 8335 | *CCDC112* | 5.4669 | -0.94052 | -1.2165 | -1.6992 | 1.10 |
| 8336 | *RRH* | 2.4862 | 1.1587 | -0.3351 | -2.1151 | 1.10 |
| 8337 | *PSMB9* | 1.8833 | 0.85162 | 0.57427 | -2.9538 | 1.10 |
| 8338 | *SYCE2* | 1.8857 | 1.8218 | -0.39872 | -0.53971 | 1.10 |
| 8339 | *FGG* | 2.771 | 0.69165 | -0.15424 | -1.882 | 1.10 |
| 8340 | *PADI1* | 1.5377 | 1.2489 | 0.5212 | -1.0168 | 1.10 |
| 8341 | *PEX11A* | 2.9531 | 0.65132 | -0.29729 | -1.515 | 1.10 |
| 8342 | *LILRB5* | 4.1281 | 0.1267 | -0.94859 | -1.358 | 1.10 |
| 8343 | *NOMO3* | 3.5564 | 0.37721 | -0.62747 | -1.5232 | 1.10 |
| 8344 | *GTSE1* | 3.7277 | 0.60138 | -1.0231 | -1.2393 | 1.10 |
| 8345 | *SEPT1* | 4.7208 | -0.64888 | -0.76635 | -2.3251 | 1.10 |
| 8346 | *OR7A5* | 3.8877 | 0.32919 | -0.91165 | -2.2385 | 1.10 |
| 8347 | *CCDC70* | 2.3749 | 0.79265 | 0.13672 | -0.90468 | 1.10 |
| 8348 | *LYN* | 3.0482 | 0.39853 | -0.14252 | -1.5255 | 1.10 |
| 8349 | *SFXN2* | 2.519 | 1.228 | -0.44353 | -1.2655 | 1.10 |
| 8350 | *SLC12A5* | 1.7968 | 1.1444 | 0.36195 | 0.16792 | 1.10 |
| 8351 | *KIAA1755* | 2.5295 | 0.65313 | 0.12019 | -2.9564 | 1.10 |
| 8352 | *CSF3* | 2.6016 | 0.51931 | 0.18166 | -0.25839 | 1.10 |
| 8353 | *ZFP36L2* | 3.8147 | 0.15971 | -0.67199 | -1.3988 | 1.10 |
| 8354 | *FAM173A* | 3.1852 | 0.076377 | 0.04083 | -0.53785 | 1.10 |
| 8355 | *NKX6-3* | 3.1102 | 0.26472 | -0.07341 | -1.3912 | 1.10 |
| 8356 | *LYAR* | 2.1324 | 1.0646 | 0.10419 | -1.5672 | 1.10 |
| 8357 | *CACNG6* | 1.5166 | 1.3145 | 0.46987 | -1.0337 | 1.10 |
| 8358 | *CFAP47* | 3.5544 | 1.0226 | -1.2763 | -1.8112 | 1.10 |
| 8359 | *HIGD1C* | 2.5308 | 1.365 | -0.59511 | -1.2324 | 1.10 |
| 8360 | *PKM* | 3.5985 | 0.86236 | -1.1605 | -1.716 | 1.10 |
| 8361 | *LRRC75B* | 4.1308 | -0.41207 | -0.4187 | -0.93011 | 1.10 |
| 8362 | *GPX1* | 2.9941 | 0.60057 | -0.29546 | -1.1666 | 1.10 |
| 8363 | *MTERF3* | 3.1113 | 0.74287 | -0.55528 | -1.2703 | 1.10 |
| 8364 | *PALM2-AKAP2* | 3.4033 | 0.44829 | -0.55288 | -1.6891 | 1.10 |
| 8365 | *RAB30* | 3.7592 | 1.719 | -2.1798 | -2.201 | 1.10 |

| | | | | | | |
|---|---|---|---|---|---|---|
| 8366 | *C9orf66* | 3.3672 | 1.4751 | -1.5447 | -1.545 | 1.10 |
| 8367 | *HMGB2* | 1.7276 | 1.4971 | 0.07273 | -0.24137 | 1.10 |
| 8368 | *TIMM23B* | 3.3857 | 0.10638 | -0.19494 | -0.9879 | 1.10 |
| 8369 | *DNAH3* | 5.379 | -0.70074 | -1.3814 | -2.0062 | 1.10 |
| 8370 | *MRPL23* | 3.1417 | 1.1899 | -1.0351 | -1.1135 | 1.10 |
| 8371 | *MGST1* | 2.3394 | 1.1475 | -0.19054 | -0.73781 | 1.10 |
| 8372 | *SAPCD1* | 1.5938 | 1.074 | 0.62787 | -0.97237 | 1.10 |
| 8373 | *ODF3* | 4.6633 | -0.07557 | -1.2927 | -1.9625 | 1.10 |
| 8374 | *CD300LF* | 2.3739 | 0.54622 | 0.37489 | -0.4674 | 1.10 |
| 8375 | *PLBD1* | 3.7357 | -0.20695 | -0.23481 | -0.4674 | 1.10 |
| 8376 | *SYNC* | 4.2791 | -0.20361 | -0.78221 | -2.5806 | 1.10 |
| 8377 | *PMM1* | 3.7551 | 0.61838 | -1.0814 | -2.4642 | 1.10 |
| 8378 | *PCDH8* | 1.3778 | 1.3669 | 0.54736 | -2.2993 | 1.10 |
| 8379 | *LSS* | 4.5599 | -0.40353 | -0.86435 | -1.6243 | 1.10 |
| 8380 | *SLC2A7* | 1.3559 | 1.0074 | 0.92779 | 0.049777 | 1.10 |
| 8381 | *ANGPT2* | 1.8272 | 1.3953 | 0.06847 | -0.94841 | 1.10 |
| 8382 | *KIF19* | 1.7327 | 1.153 | 0.40496 | -0.34251 | 1.10 |
| 8383 | *LIFR* | 2.8935 | 0.59474 | -0.198 | -0.51302 | 1.10 |
| 8384 | *LAMA3* | 2.8537 | 0.54764 | -0.1112 | -0.83623 | 1.10 |
| 8385 | *MX1* | 3.542 | 1.3541 | -1.606 | -2.1936 | 1.10 |
| 8386 | *LPPR1* | 2.0624 | 0.74524 | 0.48176 | -2.6289 | 1.10 |
| 8387 | *GYPB* | 2.1683 | 0.58529 | 0.53505 | -2.0895 | 1.10 |
| 8388 | *ARHGDIG* | 2.2692 | 0.64483 | 0.37448 | -1.9622 | 1.10 |
| 8389 | *ZNF691* | 2.3008 | 0.81226 | 0.17505 | -0.32569 | 1.10 |
| 8390 | *AACS* | 1.7943 | 0.99063 | 0.50286 | 0.3576 | 1.10 |
| 8391 | *SNAPC5* | 3.4318 | 0.40936 | -0.55388 | -0.99608 | 1.10 |
| 8392 | *ZNF865* | 3.2778 | 0.044461 | -0.03528 | -0.17663 | 1.10 |
| 8393 | *ACSL3* | 2.7356 | 2.3703 | -1.8191 | -1.9673 | 1.10 |
| 8394 | *ATG2B* | 4.4666 | 1.1942 | -2.374 | -2.6275 | 1.10 |
| 8395 | *UGT2B10* | 3.5209 | 1.3507 | -1.5848 | -1.8503 | 1.10 |
| 8396 | *BNC2* | 3.6084 | -0.09312 | -0.2287 | -1.1421 | 1.10 |
| 8397 | *URGCP-MRPS24* | 4.0681 | -0.36649 | -0.4155 | -1.3211 | 1.10 |
| 8398 | *DNAH9* | 2.5677 | 1.9002 | -1.1822 | -1.5373 | 1.10 |
| 8399 | *EPB41L4A* | 3.6059 | 0.28812 | -0.60845 | -2.0437 | 1.10 |
| 8400 | *MTMR4* | 3.1975 | 0.090585 | -0.00311 | -1.7591 | 1.09 |
| 8401 | *FGF11* | 3.6011 | 0.18574 | -0.50208 | -0.80144 | 1.09 |
| 8402 | *DHX30* | 3.489 | 0.054696 | -0.25904 | -1.7874 | 1.09 |
| 8403 | *GZMB* | 2.6471 | 0.88031 | -0.24337 | -1.4764 | 1.09 |
| 8404 | *LRRTM3* | 4.3643 | -0.45923 | -0.62154 | -1.9758 | 1.09 |
| 8405 | *SRPK1* | 3.4014 | 1.6978 | -1.8157 | -1.9055 | 1.09 |
| 8406 | *RNF139* | 2.4042 | 0.86236 | 0.01636 | -0.88634 | 1.09 |
| 8407 | *OR2M4* | 3.2055 | 1.0354 | -0.95816 | -1.6802 | 1.09 |
| 8408 | *LRRC32* | 3.6642 | 0.033637 | -0.41538 | -0.67138 | 1.09 |
| 8409 | *LHB* | 4.4208 | -0.46747 | -0.6714 | -1.2215 | 1.09 |
| 8410 | *BTBD8* | 4.7803 | -0.31688 | -1.1825 | -1.2867 | 1.09 |
| 8411 | *NCAPG2* | 3.9756 | 0.66956 | -1.3644 | -2.1217 | 1.09 |

| | | | | | | |
|---|---|---|---|---|---|---|
| 8412 | *NPFFR2* | 2.694 | 0.95737 | -0.3713 | -0.92879 | 1.09 |
| 8413 | *RFPL2* | 2.3007 | 0.66577 | 0.31308 | -1.5267 | 1.09 |
| 8414 | *PLCB2* | 2.6388 | 1.1411 | -0.50165 | -1.5128 | 1.09 |
| 8415 | *UBE2Q1* | 3.6686 | -0.13545 | -0.25527 | -1.7107 | 1.09 |
| 8416 | *MARCKSL1* | 4.521 | -0.29114 | -0.95321 | -1.1605 | 1.09 |
| 8417 | *SH2D5* | 3.2501 | 0.12282 | -0.09677 | -0.81541 | 1.09 |
| 8418 | *CA4* | 2.3874 | 0.79764 | 0.09091 | -1.445 | 1.09 |
| 8419 | *ALAD* | 5.3253 | -0.97549 | -1.0741 | -1.6855 | 1.09 |
| 8420 | *HINT3* | 1.8647 | 1.7307 | -0.32009 | -1.2438 | 1.09 |
| 8421 | *TAS1R3* | 3.9004 | 0.71022 | -1.3356 | -1.5304 | 1.09 |
| 8422 | *CDX4* | 2.4931 | 0.80519 | -0.02354 | -0.9142 | 1.09 |
| 8423 | *CSTL1* | 2.0711 | 0.91461 | 0.28831 | -1.177 | 1.09 |
| 8424 | *NFKB2* | 4.0923 | 0.48317 | -1.3016 | -1.4913 | 1.09 |
| 8425 | *OTUB2* | 4.2367 | 0.36231 | -1.3255 | -1.3292 | 1.09 |
| 8426 | *TMEM182* | 4.5511 | -0.49564 | -0.78229 | -1.1034 | 1.09 |
| 8427 | *TGFBRAP1* | 3.8841 | 0.15223 | -0.7635 | -3.3669 | 1.09 |
| 8428 | *MAPRE3* | 2.3793 | 1.0336 | -0.14009 | -2.0383 | 1.09 |
| 8429 | *SUSD1* | 3.8763 | 0.30635 | -0.9105 | -1.124 | 1.09 |
| 8430 | *BTBD1* | 2.7528 | 1.6903 | -1.1713 | -1.5654 | 1.09 |
| 8431 | *PFKP* | 3.7162 | -0.15293 | -0.29241 | -0.82172 | 1.09 |
| 8432 | *LOC100996693* | 3.7053 | 0.40039 | -0.83542 | -1.3148 | 1.09 |
| 8433 | *CCT8L2* | 2.9601 | 0.36641 | -0.05663 | -2.6313 | 1.09 |
| 8434 | *AFM* | 3.0674 | 0.38542 | -0.18358 | -1.2705 | 1.09 |
| 8435 | *CHCHD3* | 2.6967 | 0.35881 | 0.21342 | -1.0095 | 1.09 |
| 8436 | *NLRX1* | 3.5929 | -0.02502 | -0.29995 | -3.914 | 1.09 |
| 8437 | *SEC16A* | 4.6206 | 1.1774 | -2.5302 | -2.5386 | 1.09 |
| 8438 | *CMTR1* | 5.5319 | -1.082 | -1.1825 | -1.9748 | 1.09 |
| 8439 | *GABPA* | 2.4799 | 1.1184 | -0.3316 | -1.0813 | 1.09 |
| 8440 | *HSPA14* | 2.5847 | 0.93943 | -0.25748 | -1.1743 | 1.09 |
| 8441 | *LANCL1* | 1.7789 | 1.7694 | -0.2827 | -1.8193 | 1.09 |
| 8442 | *PIN4* | 2.7067 | 0.70292 | -0.14406 | -0.36513 | 1.09 |
| 8443 | *OPTN* | 2.1864 | 1.1152 | -0.03607 | -1.6649 | 1.09 |
| 8444 | *ERI3* | 2.0971 | 0.84672 | 0.32159 | -1.2744 | 1.09 |
| 8445 | *ARHGAP4* | 5.3019 | -0.73219 | -1.3043 | -1.7561 | 1.09 |
| 8446 | *AURKB* | 3.4793 | 0.36922 | -0.5834 | -1.1846 | 1.09 |
| 8447 | *MUC5B* | 1.6911 | 1.1009 | 0.47288 | -1.917 | 1.09 |
| 8448 | *CLPSL1* | 2.7713 | 1.7383 | -1.2448 | -1.3453 | 1.09 |
| 8449 | *LRRC25* | 4.0213 | -0.11781 | -0.63872 | -1.4682 | 1.09 |
| 8450 | *VAPA* | 1.9004 | 1.4356 | -0.07151 | -1.2304 | 1.09 |
| 8451 | *CSHL1* | 3.1116 | 1.0759 | -0.92339 | -1.0328 | 1.09 |
| 8452 | *C11orf85* | 2.9778 | 0.1507 | 0.13561 | -1.3343 | 1.09 |
| 8453 | *AK5* | 2.5991 | 0.35272 | 0.31206 | -0.98361 | 1.09 |
| 8454 | *FOLH1* | 3.6581 | 0.85874 | -1.2534 | -3.0644 | 1.09 |
| 8455 | *APOE* | 4.0372 | 0.13598 | -0.9101 | -1.3937 | 1.09 |
| 8456 | *TTC39C* | 3.7856 | 1.1477 | -1.6703 | -2.1202 | 1.09 |
| 8457 | *NAT8L* | 3.2101 | 0.89551 | -0.8428 | -1.0031 | 1.09 |

| | | | | | | |
|---|---|---|---|---|---|---|
| 8458 | *WDR62* | 2.8988 | 0.39617 | -0.03222 | -0.35836 | 1.09 |
| 8459 | *COMMD5* | 4.6063 | -0.55291 | -0.79083 | -1.2684 | 1.09 |
| 8460 | *FUBP3* | 3.5687 | 0.70185 | -1.0091 | -2.1282 | 1.09 |
| 8461 | *ADRB2* | 2.0237 | 1.3848 | -0.14749 | -1.9912 | 1.09 |
| 8462 | *CREBL2* | 2.0882 | 0.88078 | 0.29118 | -0.02978 | 1.09 |
| 8463 | *CIC* | 4.6436 | -0.27145 | -1.1122 | -2.4084 | 1.09 |
| 8464 | *ARHGAP12* | 3.5396 | -0.09878 | -0.18133 | -0.77284 | 1.09 |
| 8465 | *DHX58* | 3.7859 | 1.4353 | -1.9622 | -2.3466 | 1.09 |
| 8466 | *C11orf63* | 2.8335 | 0.56823 | -0.14309 | -0.59981 | 1.09 |
| 8467 | *TNFRSF10C* | 1.8882 | 0.87658 | 0.49376 | -0.95469 | 1.09 |
| 8468 | *ZNF596* | 1.9873 | 0.65056 | 0.62056 | -1.8484 | 1.09 |
| 8469 | *RGN* | 1.2559 | 1.1805 | 0.82133 | 0.083113 | 1.09 |
| 8470 | *IGFBP6* | 4.0746 | 1.0057 | -1.8227 | -1.9175 | 1.09 |
| 8471 | *CSH1* | 1.6572 | 1.2945 | 0.30552 | -1.6434 | 1.09 |
| 8472 | *TRIM58* | 1.7912 | 1.197 | 0.26863 | -1.4754 | 1.09 |
| 8473 | *PTGR1* | 3.5967 | 0.23339 | -0.57379 | -0.80771 | 1.09 |
| 8474 | *ABCC1* | 3.4252 | -0.03882 | -0.13022 | -2.0953 | 1.09 |
| 8475 | *PLD4* | 2.6638 | 0.85239 | -0.26009 | -0.96962 | 1.09 |
| 8476 | *SLC6A7* | 3.6224 | 1.8723 | -2.2388 | -2.2947 | 1.09 |
| 8477 | *ZNF706* | 4.5172 | -0.47107 | -0.79079 | -1.203 | 1.09 |
| 8478 | *UBE2QL1* | 2.8677 | 0.56312 | -0.17591 | -0.31116 | 1.08 |
| 8479 | *DOCK6* | 2.1448 | 0.99439 | 0.11544 | -0.14132 | 1.08 |
| 8480 | *KCMF1* | 3.3568 | 0.23696 | -0.33933 | -2.1612 | 1.08 |
| 8481 | *TRAF3IP2* | 2.7539 | 1.674 | -1.1742 | -2.1541 | 1.08 |
| 8482 | *ZBED5* | 1.8108 | 1.1439 | 0.29893 | -0.07475 | 1.08 |
| 8483 | *TUBG2* | 2.7433 | 2.022 | -1.5118 | -1.9559 | 1.08 |
| 8484 | *OR4D2* | 2.1108 | 0.9731 | 0.16938 | -0.65924 | 1.08 |
| 8485 | *C16orf86* | 3.5253 | 0.44472 | -0.71743 | -1.5104 | 1.08 |
| 8486 | *PRRC2A* | 2.8046 | 1.2861 | -0.83869 | -1.0584 | 1.08 |
| 8487 | *POU2F3* | 1.1703 | 1.1657 | 0.91466 | -1.0766 | 1.08 |
| 8488 | *IFITM2* | 1.7592 | 1.6501 | -0.1591 | -1.4902 | 1.08 |
| 8489 | *DKKL1* | 2.5207 | 1.036 | -0.3068 | -1.0576 | 1.08 |
| 8490 | *GOLGA6L4* | 3.3463 | 0.7069 | -0.80351 | -0.83753 | 1.08 |
| 8491 | *RGS10* | 3.7964 | 0.4308 | -0.97768 | -1.1148 | 1.08 |
| 8492 | *MYL1* | 2.3252 | 1.3159 | -0.39171 | -0.87846 | 1.08 |
| 8493 | *KIF1A* | 3.7576 | -0.20239 | -0.30609 | -0.34036 | 1.08 |
| 8494 | *TMEM263* | 2.7964 | 0.48193 | -0.02974 | -2.4873 | 1.08 |
| 8495 | *TFAP4* | 3.1495 | 0.18027 | -0.08293 | -1.4061 | 1.08 |
| 8496 | *TRIM52* | 2.4485 | 0.55552 | 0.24278 | -0.30208 | 1.08 |
| 8497 | *RNF217* | 2.3643 | 0.69777 | 0.18343 | -0.30609 | 1.08 |
| 8498 | *RP1L1* | 2.5769 | 0.56242 | 0.10613 | -2.1197 | 1.08 |
| 8499 | *LCE1D* | 1.5016 | 1.0236 | 0.71966 | -0.79485 | 1.08 |
| 8500 | *BICC1* | 2.8176 | 0.50751 | -0.08039 | -0.10575 | 1.08 |
| 8501 | *CYBB* | 2.5542 | 1.0365 | -0.34635 | -0.44353 | 1.08 |
| 8502 | *GPR37* | 4.5043 | -0.00828 | -1.2517 | -1.9558 | 1.08 |
| 8503 | *EVL* | 1.5501 | 1.468 | 0.22609 | 5.25E-05 | 1.08 |

| | | | | | | |
|---|---|---|---|---|---|---|
| 8504 | *ADIPOR2* | 2.294 | 2.1555 | -1.2056 | -1.6998 | 1.08 |
| 8505 | *S100A7A* | 3.7907 | -0.18381 | -0.36311 | -2.3876 | 1.08 |
| 8506 | *DEFB106B* | 2.7492 | -0.58705 | | | 1.08 |
| 8507 | *LRRC29* | 2.2042 | 0.67494 | 0.36389 | -0.57207 | 1.08 |
| 8508 | *VHL* | 5.1905 | -0.64769 | -1.3004 | -2.2343 | 1.08 |
| 8509 | *KLRG1* | 2.9953 | 1.0231 | -0.77602 | -2.7566 | 1.08 |
| 8510 | *SDHD* | 4.2611 | -0.32009 | -0.69866 | -1.2666 | 1.08 |
| 8511 | *IFITM10* | 2.8539 | 0.43059 | -0.04339 | -0.91192 | 1.08 |
| 8512 | *FOXN1* | 3.7385 | -0.07348 | -0.4247 | -0.49968 | 1.08 |
| 8513 | *KIF21A* | 1.7241 | 1.6993 | -0.18327 | -0.64767 | 1.08 |
| 8514 | *UHRF1BP1* | 3.4727 | -0.06034 | -0.17238 | -1.3994 | 1.08 |
| 8515 | *OR1L8* | 2.515 | 0.7349 | -0.01 | -1.3973 | 1.08 |
| 8516 | *KIAA1462* | 2.693 | 0.77875 | -0.2325 | -2.1541 | 1.08 |
| 8517 | *QKI* | 3.6399 | 0.091453 | -0.49216 | -0.9472 | 1.08 |
| 8518 | *OR4F16* | 3.3439 | 0.30728 | -0.41239 | -1.3694 | 1.08 |
| 8519 | *MIA2* | 3.0898 | 0.75887 | -0.61017 | -1.3409 | 1.08 |
| 8520 | *SIK2* | 3.2964 | 0.39038 | -0.44832 | -1.0923 | 1.08 |
| 8521 | *C20orf173* | 1.7069 | 1.442 | 0.08929 | -1.124 | 1.08 |
| 8522 | *MOGAT3* | 3.829 | -0.25941 | -0.33152 | -0.96661 | 1.08 |
| 8523 | *EHF* | 4.5766 | -0.22726 | -1.1114 | -1.9673 | 1.08 |
| 8524 | *AQP8* | 2.342 | 0.82905 | 0.06658 | -0.50731 | 1.08 |
| 8525 | *OR13A1* | 2.9438 | 1.0617 | -0.76857 | -1.5073 | 1.08 |
| 8526 | *C19orf45* | 2.1694 | 0.99891 | 0.06774 | 0.05242 | 1.08 |
| 8527 | *RINT1* | 3.5922 | 0.36232 | -0.71905 | -2.3389 | 1.08 |
| 8528 | *TSPAN18* | 3.8211 | 0.62581 | -1.2125 | -1.4685 | 1.08 |
| 8529 | *NTN4* | 4.1574 | 0.21938 | -1.1426 | -1.6525 | 1.08 |
| 8530 | *C9orf171* | 4.1326 | -0.42378 | -0.47485 | -1.615 | 1.08 |
| 8531 | *SPAG7* | 2.0171 | 1.8722 | -0.65553 | -1.3769 | 1.08 |
| 8532 | *TMEM17* | 1.4447 | 1.2343 | 0.55401 | -0.51931 | 1.08 |
| 8533 | *TYMS* | 2.3813 | 0.56035 | 0.29118 | -2.0256 | 1.08 |
| 8534 | *EIF2S1* | 3.5989 | 1.3938 | -1.7614 | -2.0452 | 1.08 |
| 8535 | *FILIP1* | 4.5215 | -0.45505 | -0.83527 | -0.9857 | 1.08 |
| 8536 | *RPUSD4* | 2.6709 | 0.38514 | 0.17508 | -0.69006 | 1.08 |
| 8537 | *IPO8* | 2.7193 | 0.47026 | 0.04149 | -1.2324 | 1.08 |
| 8538 | *METTL14* | 3.1621 | 0.72355 | -0.65553 | -1.6672 | 1.08 |
| 8539 | *KIAA0922* | 3.0575 | 0.29641 | -0.12408 | -0.8083 | 1.08 |
| 8540 | *KRTAP20-1* | 3.06 | 2.2243 | -2.0545 | -2.0834 | 1.08 |
| 8541 | *TFCP2* | 2.1001 | 1.9748 | -0.84529 | -1.3465 | 1.08 |
| 8542 | *ESRRA* | 3.8302 | 0.47086 | -1.0716 | -1.6286 | 1.08 |
| 8543 | *SMAP1* | 2.5478 | 0.57218 | 0.10941 | -1.8692 | 1.08 |
| 8544 | *ZSWIM8* | 4.1707 | 0.11251 | -1.054 | -1.626 | 1.08 |
| 8545 | *DDN* | 4.4411 | 0.048425 | -1.2608 | -1.3098 | 1.08 |
| 8546 | *OR2J3* | 2.3266 | 1.2261 | -0.3247 | -0.3658 | 1.08 |
| 8547 | *STK24* | 3.5753 | 0.39078 | -0.73853 | -1.6289 | 1.08 |
| 8548 | *B3GAT1* | 2.8552 | 0.77193 | -0.39998 | -0.85467 | 1.08 |
| 8549 | *SPHKAP* | 4.575 | -0.65035 | -0.69815 | -1.1031 | 1.08 |

| | | | | | | |
|---|---|---|---|---|---|---|
| 8550 | *SETDB1* | 3.3853 | -0.04486 | -0.11452 | -1.3842 | 1.08 |
| 8551 | *APLP2* | 4.2916 | 0.05102 | -1.1169 | -2.3602 | 1.08 |
| 8552 | *CTBS* | 1.6141 | 1.0557 | 0.55553 | -2.2432 | 1.08 |
| 8553 | *PSG2* | 2.4485 | 0.68003 | 0.09665 | -0.91236 | 1.08 |
| 8554 | *FAM181A* | 1.2611 | 1.1849 | 0.77887 | -0.42064 | 1.07 |
| 8555 | *PGLYRP4* | 1.5606 | 1.3536 | 0.31054 | -1.6368 | 1.07 |
| 8556 | *TNNT2* | 2.8666 | 0.80045 | -0.44353 | -0.71077 | 1.07 |
| 8557 | *ZIK1* | 4.8723 | -0.61338 | -1.0355 | -1.2172 | 1.07 |
| 8558 | *VIPR2* | 3.6118 | -0.04513 | -0.34327 | -0.87109 | 1.07 |
| 8559 | *GDA* | 2.8969 | 0.58537 | -0.2594 | -1.1919 | 1.07 |
| 8560 | *CBR3* | 2.7194 | 0.65655 | -0.15337 | -1.8366 | 1.07 |
| 8561 | *LY6H* | 2.7864 | 0.93864 | -0.50265 | -2.5707 | 1.07 |
| 8562 | *FANCA* | 2.5217 | 1.2197 | -0.51912 | -1.0032 | 1.07 |
| 8563 | *RNF215* | 2.8483 | 0.65567 | -0.28176 | -0.30739 | 1.07 |
| 8564 | *HCFC1* | 4.0138 | 0.44884 | -1.2411 | -2.1239 | 1.07 |
| 8565 | *TAS2R30* | 2.4913 | 0.80445 | -0.07427 | -0.4221 | 1.07 |
| 8566 | *SMG1* | 5.2505 | -0.91779 | -1.1114 | -1.9984 | 1.07 |
| 8567 | *KCNIP1* | 4.2138 | -0.44353 | -0.55158 | -0.69754 | 1.07 |
| 8568 | *PTPN11* | 4.1984 | -0.13539 | -0.84523 | -2.9646 | 1.07 |
| 8569 | *PANK4* | 4.3019 | 0.47862 | -1.5635 | -2.6289 | 1.07 |
| 8570 | *RBM10* | 2.6428 | 0.80141 | -0.2283 | -1.2574 | 1.07 |
| 8571 | *TCERG1L* | 3.7968 | -0.13429 | -0.44661 | -1.0576 | 1.07 |
| 8572 | *TFF3* | 2.4234 | 0.42358 | 0.36833 | -1.3117 | 1.07 |
| 8573 | *TAF15* | 2.782 | 0.61845 | -0.18662 | -1.3711 | 1.07 |
| 8574 | *COCH* | 2.1425 | 0.72059 | 0.3488 | -0.3351 | 1.07 |
| 8575 | *COL24A1* | 4.5225 | -0.0304 | -1.2815 | -1.3583 | 1.07 |
| 8576 | *CCT3* | 2.8823 | 0.26025 | 0.06783 | -2.8051 | 1.07 |
| 8577 | *MYLIP* | 1.5725 | 1.3056 | 0.33119 | -2.0107 | 1.07 |
| 8578 | *FLVCR2* | 2.0363 | 0.97658 | 0.19565 | -1.7886 | 1.07 |
| 8579 | *BDKRB2* | 2.3071 | 0.87275 | 0.02849 | -1.1094 | 1.07 |
| 8580 | *EVA1A* | 2.5198 | 0.91317 | -0.22469 | -1.348 | 1.07 |
| 8581 | *NCKAP1L* | 3.6328 | 0.12276 | -0.54818 | -0.59823 | 1.07 |
| 8582 | *ZNF496* | 3.6612 | 0.16511 | -0.61946 | -0.92629 | 1.07 |
| 8583 | *PGM2L1* | 2.6643 | 0.58168 | -0.03924 | -1.2024 | 1.07 |
| 8584 | *USP13* | 2.5264 | 0.42153 | 0.25877 | -1.1331 | 1.07 |
| 8585 | *FGFBP1* | 2.0747 | 1.0155 | 0.11535 | 0.059357 | 1.07 |
| 8586 | *SMCP* | 3.2888 | 0.93563 | -1.0191 | -2.0304 | 1.07 |
| 8587 | *KIF22* | 2.7623 | 0.75738 | -0.31475 | -2.0301 | 1.07 |
| 8588 | *ITGB1BP1* | 3.7988 | -0.2873 | -0.30677 | -0.81217 | 1.07 |
| 8589 | *SLAMF1* | 4.2201 | -0.45928 | -0.55655 | -0.94777 | 1.07 |
| 8590 | *DTX3* | 2.5911 | 1.1776 | -0.56586 | -2.6376 | 1.07 |
| 8591 | *ITGB1BP2* | 1.8341 | 1.6752 | -0.30723 | -0.59716 | 1.07 |
| 8592 | *BCL6B* | 2.2903 | 2.0172 | -1.1063 | -2.2154 | 1.07 |
| 8593 | *OR4F4* | 2.5013 | 1.1856 | -0.48575 | -1.4415 | 1.07 |
| 8594 | *ECE1* | 1.2599 | 1.2455 | 0.6956 | -0.17111 | 1.07 |
| 8595 | *XRCC3* | 5.7698 | -1.2027 | -1.3662 | -1.6007 | 1.07 |

| | | | | | | |
|---|---|---|---|---|---|---|
| 8596 | *GLT8D1* | 4.0124 | -0.34411 | -0.46748 | -1.4206 | 1.07 |
| 8597 | *SLC12A3* | 3.7463 | 0.22275 | -0.76882 | -1.743 | 1.07 |
| 8598 | *GRAMD1B* | 3.6061 | 0.3645 | -0.77042 | -1.5884 | 1.07 |
| 8599 | *RHBDD2* | 3.3444 | -0.00492 | -0.13962 | -0.77848 | 1.07 |
| 8600 | *EIF4A2* | 1.3049 | 1.0362 | 0.85862 | 0.30154 | 1.07 |
| 8601 | *MGLL* | 2.5471 | 0.64819 | 0.00396 | -0.5183 | 1.07 |
| 8602 | *LFNG* | 2.0993 | 2.0167 | -0.91701 | -1.5608 | 1.07 |
| 8603 | *C9orf57* | 3.3284 | 0.9896 | -1.1193 | -2.3496 | 1.07 |
| 8604 | *MBD6* | 3.9239 | 0.078829 | -0.80442 | -1.1212 | 1.07 |
| 8605 | *IFIT2* | 3.0542 | 0.25827 | -0.11468 | -0.82826 | 1.07 |
| 8606 | *PIH1D2* | 1.4906 | 1.1841 | 0.52293 | -0.70976 | 1.07 |
| 8607 | *ANKRD18B* | 2.0043 | 0.71805 | 0.47389 | -0.73143 | 1.07 |
| 8608 | *ZNF311* | 2.8116 | 1.6121 | -1.2277 | -1.371 | 1.07 |
| 8609 | *LAMA4* | 3.4608 | 0.56332 | -0.82833 | -1.6218 | 1.07 |
| 8610 | *HVCN1* | 3.4203 | 0.77017 | -0.99474 | -1.2915 | 1.07 |
| 8611 | *TRIM33* | 2.2064 | 0.88068 | 0.10822 | -1.9008 | 1.07 |
| 8612 | *BEND5* | 2.9842 | 0.5894 | -0.38028 | -2.9834 | 1.06 |
| 8613 | *GABRA4* | 1.5655 | 0.88248 | 0.74438 | -0.53661 | 1.06 |
| 8614 | *TCAP* | 1.9671 | 1.7344 | -0.50935 | -1.5149 | 1.06 |
| 8615 | *DACH1* | 2.2152 | 0.50053 | 0.47612 | -0.42226 | 1.06 |
| 8616 | *TSC22D2* | 1.8617 | 0.82981 | 0.50025 | -0.73787 | 1.06 |
| 8617 | *MYBPC2* | 2.6062 | 1.5082 | -0.92384 | -2.3577 | 1.06 |
| 8618 | *SYT4* | 5.3551 | -0.55815 | -1.6065 | -1.6208 | 1.06 |
| 8619 | *LRIG2* | 3.2571 | -0.02341 | -0.04332 | -0.33752 | 1.06 |
| 8620 | *C1GALT1* | 1.6215 | 1.0507 | 0.51733 | -0.67702 | 1.06 |
| 8621 | *BNIP2* | 1.8878 | 1.6447 | -0.34315 | -0.61017 | 1.06 |
| 8622 | *RDH5* | 2.2649 | 1.0575 | -0.13381 | -0.13622 | 1.06 |
| 8623 | *ZNF595* | 5.1107 | -0.72038 | -1.202 | -1.5082 | 1.06 |
| 8624 | *FZD5* | 4.4162 | -0.15551 | -1.0726 | -1.1053 | 1.06 |
| 8625 | *CDK16* | 2.8619 | 0.65057 | -0.32567 | -1.5246 | 1.06 |
| 8626 | *HLA-DQB1* | 3.357 | 0.24134 | -0.41207 | -1.1461 | 1.06 |
| 8627 | *CT47A3* | 2.2023 | 0.9802 | 0.00363 | -0.04968 | 1.06 |
| 8628 | *EXOSC7* | 2.3206 | 0.62291 | 0.24242 | -1.6469 | 1.06 |
| 8629 | *RSC1A1* | 2.0603 | 0.76148 | 0.36413 | -0.6951 | 1.06 |
| 8630 | *GZMM* | 2.3464 | 1.6831 | -0.84375 | -1.8711 | 1.06 |
| 8631 | *PNCK* | 2.1835 | 0.6876 | 0.31419 | -0.63976 | 1.06 |
| 8632 | *BBS7* | 2.5556 | 1.6221 | -0.9925 | -1.0752 | 1.06 |
| 8633 | *TMEM110* | 4.622 | -0.47696 | -0.9599 | -1.4325 | 1.06 |
| 8634 | *CYP3A7-CYP3A51* | 2.0912 | 0.60261 | 0.49122 | -1.8741 | 1.06 |
| 8635 | *ACOT13* | 2.6962 | 0.68499 | -0.19626 | -2.2866 | 1.06 |
| 8636 | *AIM1L* | 3.7438 | -0.16884 | -0.39081 | -0.68889 | 1.06 |
| 8637 | *NAP1L5* | 2.0351 | 1.6712 | -0.52255 | -1.5952 | 1.06 |
| 8638 | *SLC22A23* | 3.2374 | 0.12678 | -0.1808 | -2.2432 | 1.06 |
| 8639 | *IL12B* | 3.8952 | -0.13563 | -0.5763 | -0.66034 | 1.06 |
| 8640 | *JSRP1* | 3.1471 | 0.55041 | -0.51464 | -1.5744 | 1.06 |
| 8641 | *SPATA6L* | 1.537 | 1.1245 | 0.52057 | -0.30091 | 1.06 |

| | | | | | | |
|---|---|---|---|---|---|---|
| 8642 | *CRTAP* | 3.1867 | 1.028 | -1.0347 | -1.1053 | 1.06 |
| 8643 | *MGAM2* | 4.8801 | -0.45972 | -1.2411 | -1.3591 | 1.06 |
| 8644 | *LCE1A* | 2.125 | 0.81979 | 0.23375 | -0.25241 | 1.06 |
| 8645 | *AP1AR* | 2.6837 | 0.7659 | -0.27146 | -1.4684 | 1.06 |
| 8646 | *FAM160A2* | 4.4538 | 0.088577 | -1.3643 | -1.4306 | 1.06 |
| 8647 | *TPRG1L* | 3.729 | -0.14256 | -0.40933 | -2.0791 | 1.06 |
| 8648 | *MAD2L2* | 2.7523 | 1.6317 | -1.2071 | -1.7766 | 1.06 |
| 8649 | *TOPORS* | 2.3819 | 0.79669 | -0.00314 | -1.3194 | 1.06 |
| 8650 | *UROD* | 3.1996 | 0.36129 | -0.38553 | -0.95842 | 1.06 |
| 8651 | *TMEM132C* | 2.9059 | 0.13862 | 0.13058 | -0.13916 | 1.06 |
| 8652 | *PRKDC* | 3.1158 | 1.3178 | -1.2586 | -1.4272 | 1.06 |
| 8653 | *RGS21* | 3.9945 | -0.14795 | -0.67235 | -1.4204 | 1.06 |
| 8654 | *ZNF232* | 2.8618 | 0.75131 | -0.43946 | -3.8314 | 1.06 |
| 8655 | *TMEM69* | 1.6311 | 1.3467 | 0.1957 | -2.9818 | 1.06 |
| 8656 | *SPPL2C* | 3.6399 | 0.50741 | -0.97433 | -1.3577 | 1.06 |
| 8657 | *LRRC55* | 2.3598 | 0.75405 | 0.05894 | -0.27755 | 1.06 |
| 8658 | *CRLF1* | 2.5485 | 0.41343 | 0.21086 | -2.1143 | 1.06 |
| 8659 | *DDIT4L* | 1.8655 | 1.2163 | 0.09098 | -1.124 | 1.06 |
| 8660 | *C8A* | 2.0817 | 0.75214 | 0.33838 | -0.36391 | 1.06 |
| 8661 | *TPBGL* | 1.7907 | 1.0612 | 0.31678 | -1.7807 | 1.06 |
| 8662 | *FOXP4* | 1.983 | 0.76584 | 0.41968 | -1.0888 | 1.06 |
| 8663 | *CRYBA1* | 3.1456 | 0.21358 | -0.19096 | -0.4735 | 1.06 |
| 8664 | *C11orf45* | 2.0417 | 0.80629 | 0.31999 | -0.38559 | 1.06 |
| 8665 | *HIST1H4C* | 3.9345 | 0.58153 | -1.3481 | -2.7991 | 1.06 |
| 8666 | *IL25* | 3.6579 | 0.40039 | -0.89129 | -2.4328 | 1.06 |
| 8667 | *DHRS1* | 2.6399 | 0.26577 | 0.2606 | -1.9326 | 1.06 |
| 8668 | *RMND1* | 2.0123 | 1.2717 | -0.11773 | -1.7317 | 1.06 |
| 8669 | *FCRL1* | 2.1643 | 0.73466 | 0.2672 | -0.98292 | 1.06 |
| 8670 | *KCNH8* | 2.4032 | 1.5206 | -0.75796 | -1.3668 | 1.06 |
| 8671 | *POU1F1* | 3.6929 | -0.10911 | -0.4187 | -1.0092 | 1.06 |
| 8672 | *DGKG* | 2.4108 | 0.52995 | 0.22384 | -3.1836 | 1.05 |
| 8673 | *FGF23* | 3.6964 | -0.0886 | -0.44353 | -1.7888 | 1.05 |
| 8674 | *PRSS48* | 4.87 | -0.14289 | -1.5635 | -2.1342 | 1.05 |
| 8675 | *ISLR* | 2.4248 | 1.6983 | -0.95964 | -1.0802 | 1.05 |
| 8676 | *COL25A1* | 3.2168 | 0.31543 | -0.36917 | -1.3127 | 1.05 |
| 8677 | *TPP2* | 4.6058 | -0.22943 | -1.2141 | -1.2277 | 1.05 |
| 8678 | *MYBL1* | 4.2591 | 0.14094 | -1.2383 | -2.0337 | 1.05 |
| 8679 | *AICDA* | 4.4909 | -0.56598 | -0.76495 | -1.2867 | 1.05 |
| 8680 | *MAF1* | 4.079 | 0.51745 | -1.4373 | -1.4996 | 1.05 |
| 8681 | *NFKBIA* | 2.3463 | 1.7399 | -0.92823 | -1.6649 | 1.05 |
| 8682 | *CD37* | 2.8448 | 1.257 | -0.94399 | -1.8478 | 1.05 |
| 8683 | *PARP15* | 4.6417 | -0.1549 | -1.329 | -1.7466 | 1.05 |
| 8684 | *NKAIN2* | 2.1127 | 1.2731 | -0.22803 | -0.31244 | 1.05 |
| 8685 | *FLNA* | 3.5534 | 0.30172 | -0.69831 | -1.2783 | 1.05 |
| 8686 | *TMEM232* | 1.292 | 1.0049 | 0.85962 | -1.2815 | 1.05 |
| 8687 | *RNF212B* | 3.9876 | 0.3541 | -1.1858 | -2.0351 | 1.05 |

| | | | | | | |
|---|---|---|---|---|---|---|
| 8688 | *DES* | 3.2008 | 0.22548 | -0.2715 | -1.094 | 1.05 |
| 8689 | *RPLP1* | 5.39 | -0.40121 | -1.8341 | -2.2771 | 1.05 |
| 8690 | *ALYREF* | 2.611 | 0.53812 | 0.00556 | -0.71123 | 1.05 |
| 8691 | *P2RX1* | 3.5285 | -0.01633 | -0.35899 | -2.3534 | 1.05 |
| 8692 | *ATP1B3* | 3.0477 | 1.1877 | -1.0824 | -1.872 | 1.05 |
| 8693 | *ENHO* | 2.4853 | 0.93921 | -0.27198 | -1.1547 | 1.05 |
| 8694 | *KLHL32* | 2.8707 | 1.2006 | -0.91901 | -1.4051 | 1.05 |
| 8695 | *KMT2E* | 3.5755 | 0.025902 | -0.44951 | -1.1957 | 1.05 |
| 8696 | *HERC3* | 2.0936 | 1.3714 | -0.31342 | -1.7695 | 1.05 |
| 8697 | *LCTL* | 3.1054 | 0.50263 | -0.45663 | -0.67253 | 1.05 |
| 8698 | *HES6* | 3.1967 | 0.67976 | -0.72585 | -1.7323 | 1.05 |
| 8699 | *PAGE5* | 3.5647 | -0.03832 | -0.37612 | -2.7442 | 1.05 |
| 8700 | *LEFTY1* | 4.4138 | -0.49423 | -0.76966 | -1.6801 | 1.05 |
| 8701 | *CENPN* | 2.8895 | 1.2143 | -0.95423 | -1.5022 | 1.05 |
| 8702 | *PARD6G* | 3.61 | 0.13046 | -0.59168 | -2.0483 | 1.05 |
| 8703 | *DENND2C* | 2.7668 | 1.5577 | -1.1759 | -1.8248 | 1.05 |
| 8704 | *CGREF1* | 2.7834 | 0.42847 | -0.06333 | -1.9237 | 1.05 |
| 8705 | *LDHAL6B* | 3.0311 | 0.36086 | -0.24387 | -0.41139 | 1.05 |
| 8706 | *PPP1R21* | 4.2507 | -0.35718 | -0.74566 | -1.9435 | 1.05 |
| 8707 | *SRPK2* | 2.5989 | 0.6408 | -0.0923 | -1.3184 | 1.05 |
| 8708 | *HSPB11* | 1.9607 | 1.1581 | 0.02802 | -1.6999 | 1.05 |
| 8709 | *OR11H2* | 3.3269 | 0.2962 | -0.47636 | -1.0904 | 1.05 |
| 8710 | *FBXO4* | 4.5074 | 0.29113 | -1.653 | -2.4022 | 1.05 |
| 8711 | *SCAPER* | 1.6281 | 1.2432 | 0.27398 | -2.0181 | 1.05 |
| 8712 | *DLX3* | 2.164 | 0.98874 | -0.0083 | -1.2301 | 1.05 |
| 8713 | *ACVR2B* | 1.6074 | 1.5151 | 0.02187 | -0.56681 | 1.05 |
| 8714 | *FLII* | 3.6638 | -0.2283 | -0.29114 | -0.96196 | 1.05 |
| 8715 | *ETFA* | 1.5309 | 1.332 | 0.28141 | -0.04288 | 1.05 |
| 8716 | *CD8A* | 2.0656 | 2.0511 | -0.97357 | -1.7316 | 1.05 |
| 8717 | *CPLX4* | 3.4112 | 0.50376 | -0.77255 | -1.6527 | 1.05 |
| 8718 | *RHBG* | 2.2283 | 0.64162 | 0.27221 | -1.0386 | 1.05 |
| 8719 | *ZMYND15* | 2.3134 | 1.5568 | -0.72825 | -1.4536 | 1.05 |
| 8720 | *MAP7D2* | 2.4689 | 1.287 | -0.61436 | -2.9538 | 1.05 |
| 8721 | *SCGB1D4* | 4.8442 | -0.4125 | -1.2902 | -1.3556 | 1.05 |
| 8722 | *LDLRAD2* | 3.2397 | 0.23598 | -0.33426 | -2.3036 | 1.05 |
| 8723 | *AHSG* | 3.7585 | 1.1776 | -1.7947 | -2.537 | 1.05 |
| 8724 | *ZFP37* | 4.238 | 0.15417 | -1.251 | -2.2811 | 1.05 |
| 8725 | *MOCS3* | 2.5197 | 1.4166 | -0.79558 | -2.7689 | 1.05 |
| 8726 | *PCDH11Y* | 1.5185 | 1.2408 | 0.38129 | -0.77095 | 1.05 |
| 8727 | *PRAMEF1* | 2.0881 | 1.2339 | -0.18195 | -0.46586 | 1.05 |
| 8728 | *CFTR* | 2.4426 | 0.83229 | -0.13487 | -1.1385 | 1.05 |
| 8729 | *LEMD3* | 2.4204 | 0.54994 | 0.16959 | -0.0014 | 1.05 |
| 8730 | *COLEC10* | 2.3121 | 1.1219 | -0.29418 | -1.5182 | 1.05 |
| 8731 | *SYNCRIP* | 3.2817 | 0.94319 | -1.0863 | -2.4221 | 1.05 |
| 8732 | *TPX2* | 2.9224 | 1.189 | -0.97381 | -1.7782 | 1.05 |
| 8733 | *CLDN1* | 2.1067 | 0.73695 | 0.29391 | -0.31505 | 1.05 |

| | | | | | | |
|---|---|---|---|---|---|---|
| 8734 | *GK2* | 1.7536 | 0.95032 | 0.43341 | -1.4154 | 1.05 |
| 8735 | *UPP2* | 2.0413 | 0.84243 | 0.25253 | -0.05667 | 1.05 |
| 8736 | *CWC25* | 5.1005 | -0.7409 | -1.2234 | -2.1938 | 1.05 |
| 8737 | *SLC9A2* | 3.6564 | -0.1386 | -0.38361 | -2.0537 | 1.04 |
| 8738 | *XKR9* | 2.8198 | 0.87394 | -0.5603 | -1.0529 | 1.04 |
| 8739 | *LRMP* | 2.582 | 0.49105 | 0.06014 | -0.44353 | 1.04 |
| 8740 | *CHKB* | 3.2547 | 0.59723 | -0.71893 | -2.7849 | 1.04 |
| 8741 | *RHPN1* | 1.7063 | 1.2334 | 0.19304 | -0.87335 | 1.04 |
| 8742 | *NUPL2* | 3.1274 | 0.45962 | -0.45531 | -1.5393 | 1.04 |
| 8743 | *LRRN2* | 3.0409 | 0.12142 | -0.03097 | -1.9673 | 1.04 |
| 8744 | *DIS3L* | 3.7631 | -0.1865 | -0.44774 | -1.94 | 1.04 |
| 8745 | *SRD5A1* | 2.7562 | 0.57985 | -0.20743 | -0.94881 | 1.04 |
| 8746 | *MDM4* | 3.0483 | 0.62398 | -0.54373 | -2.1342 | 1.04 |
| 8747 | *TMEM198* | 1.4055 | 0.96502 | 0.75622 | -2.3235 | 1.04 |
| 8748 | *WDR13* | 1.4323 | 0.95746 | 0.73692 | -1.7646 | 1.04 |
| 8749 | *EIF3H* | 2.202 | 1.2851 | -0.36106 | -1.1322 | 1.04 |
| 8750 | *PRUNE* | 4.0432 | 0.35324 | -1.2705 | -1.2837 | 1.04 |
| 8751 | *PIPOX* | 2.0195 | 0.74601 | 0.3593 | -0.5183 | 1.04 |
| 8752 | *FLYWCH1* | 2.395 | 0.69891 | 0.03066 | -1.2174 | 1.04 |
| 8753 | *SCP2D1* | 2.1131 | 1.9447 | -0.93329 | -1.5125 | 1.04 |
| 8754 | *FES* | 1.0551 | 1.054 | 1.0152 | -0.03588 | 1.04 |
| 8755 | *SPINK9* | 4.6993 | -0.17772 | -1.3982 | -1.4143 | 1.04 |
| 8756 | *NAA38* | 3.5771 | 0.37382 | -0.82796 | -1.1726 | 1.04 |
| 8757 | *TAF3* | 1.3517 | 1.1841 | 0.58686 | -1.697 | 1.04 |
| 8758 | *ABHD13* | 3.9202 | -0.01403 | -0.78364 | -2.3248 | 1.04 |
| 8759 | *NTRK1* | 2.7724 | 0.20593 | 0.14415 | -0.07288 | 1.04 |
| 8760 | *OR8B3* | 3.2511 | 0.027145 | -0.15596 | -0.75786 | 1.04 |
| 8761 | *SHISA4* | 3.1747 | 0.2658 | -0.31838 | -1.4373 | 1.04 |
| 8762 | *GZMA* | 2.1887 | 1.4523 | -0.51929 | -1.8227 | 1.04 |
| 8763 | *ZNF630* | 2.196 | 1.4711 | -0.54544 | -1.0355 | 1.04 |
| 8764 | *PLA1A* | 3.446 | 0.022499 | -0.34751 | -0.46757 | 1.04 |
| 8765 | *PRRC2B* | 1.4581 | 1.2847 | 0.37817 | -0.35462 | 1.04 |
| 8766 | *NIPA2* | 3.1933 | 1.1014 | -1.1739 | -1.505 | 1.04 |
| 8767 | *SCNN1B* | 4.1925 | -0.47206 | -0.60003 | -0.93865 | 1.04 |
| 8768 | *CCNA1* | 3.2372 | 0.71119 | -0.82805 | -0.95629 | 1.04 |
| 8769 | *GORAB* | 2.2595 | 1.6211 | -0.76033 | -1.4684 | 1.04 |
| 8770 | *FNDC3A* | 1.8392 | 1.7221 | -0.44134 | -1.0918 | 1.04 |
| 8771 | *SNRNP70* | 3.4318 | 0.060946 | -0.37317 | -0.83145 | 1.04 |
| 8772 | *CXCL14* | 2.6834 | 0.50977 | -0.07541 | -0.30078 | 1.04 |
| 8773 | *ACOT11* | 3.0281 | 0.46482 | -0.37592 | -1.5631 | 1.04 |
| 8774 | *TNNT1* | 1.7759 | 1.6155 | -0.275 | -1.359 | 1.04 |
| 8775 | *CCDC42* | 2.0238 | 0.8693 | 0.22311 | -0.03866 | 1.04 |
| 8776 | *MSANTD4* | 2.8492 | 0.20352 | 0.06291 | -0.25317 | 1.04 |
| 8777 | *F10* | 4.7354 | -0.41681 | -1.2032 | -2.4394 | 1.04 |
| 8778 | *CARNS1* | 2.7162 | 0.9243 | -0.52521 | -0.73938 | 1.04 |
| 8779 | *LOC100130357* | 3.9351 | -0.03767 | -0.78229 | -0.85389 | 1.04 |

| | | | | | | |
|---|---|---|---|---|---|---|
| 8780 | *STMN4* | 1.3491 | 1.0022 | 0.76255 | -0.1307 | 1.04 |
| 8781 | *SLC39A14* | 1.1151 | 1.0863 | 0.912 | 0.46625 | 1.04 |
| 8782 | *FABP4* | 2.1463 | 1.8159 | -0.85027 | -2.203 | 1.04 |
| 8783 | *ZNF256* | 2.9758 | 0.57062 | -0.43507 | -0.81509 | 1.04 |
| 8784 | *FAM118B* | 3.9077 | -0.23523 | -0.56145 | -1.3762 | 1.04 |
| 8785 | *SLC25A42* | 2.2312 | 1.1566 | -0.27844 | -1.7232 | 1.04 |
| 8786 | *FOXF2* | 2.2831 | 2.1284 | -1.3031 | -1.5658 | 1.04 |
| 8787 | *STK19* | 2.6487 | 0.36866 | 0.08979 | -0.89741 | 1.04 |
| 8788 | *MGAM* | 4.3202 | -0.33104 | -0.88213 | -1.0385 | 1.04 |
| 8789 | *MTHFS* | 1.7933 | 0.99901 | 0.31424 | -2.2844 | 1.04 |
| 8790 | *WDR92* | 2.1184 | 0.71004 | 0.27765 | -1.0444 | 1.04 |
| 8791 | *CDC42BPA* | 4.012 | 1.0695 | -1.9758 | -2.0258 | 1.04 |
| 8792 | *TTC8* | 3.1253 | 1.3738 | -1.3935 | -1.4405 | 1.04 |
| 8793 | *TMEM209* | 3.7855 | -0.30094 | -0.37913 | -1.707 | 1.04 |
| 8794 | *C19orf26* | 4.1717 | -0.30777 | -0.75876 | -0.84858 | 1.04 |
| 8795 | *C9orf91* | 2.6478 | 0.96477 | -0.50793 | -1.5171 | 1.03 |
| 8796 | *KLK4* | 3.3485 | 0.48942 | -0.73333 | -1.8366 | 1.03 |
| 8797 | *VSX2* | 2.6519 | 2.268 | -1.8157 | -2.5198 | 1.03 |
| 8798 | *RADIL* | 3.829 | -0.33788 | -0.38754 | -1.7505 | 1.03 |
| 8799 | *NLRP1* | 3.7585 | -0.01099 | -0.64496 | -2.4331 | 1.03 |
| 8800 | *HDHD2* | 3.0912 | 0.21428 | -0.20338 | -1.1685 | 1.03 |
| 8801 | *SAMD11* | 2.3689 | 0.51562 | 0.21727 | -1.2936 | 1.03 |
| 8802 | *RXRB* | 3.7913 | 0.82957 | -1.5194 | -1.8929 | 1.03 |
| 8803 | *PTPRR* | 3.5207 | 0.58984 | -1.0092 | -1.3416 | 1.03 |
| 8804 | *NQO1* | 3.1386 | 0.10416 | -0.14252 | -0.52034 | 1.03 |
| 8805 | *WBP1L* | 4.2137 | -0.28391 | -0.82989 | -1.696 | 1.03 |
| 8806 | *ABI1* | 2.6616 | 0.52823 | -0.09238 | -1.1444 | 1.03 |
| 8807 | *TMEM190* | 2.31 | 1.817 | -1.0297 | -1.146 | 1.03 |
| 8808 | *HSF5* | 2.69 | 0.82961 | -0.42248 | -0.63826 | 1.03 |
| 8809 | *TUBD1* | 3.0061 | 1.0594 | -0.96938 | -1.1742 | 1.03 |
| 8810 | *C10orf131* | 3.0051 | 0.10931 | -0.01845 | -2.3154 | 1.03 |
| 8811 | *MINK1* | 3.5458 | 1.2422 | -1.6926 | -2.6973 | 1.03 |
| 8812 | *DGCR6* | 2.1939 | 0.63881 | 0.26193 | -0.60095 | 1.03 |
| 8813 | *HPCAL1* | 2.171 | 1.1589 | -0.23607 | -0.58929 | 1.03 |
| 8814 | *ASIC5* | 2.2407 | 1.0165 | -0.16343 | -2.0895 | 1.03 |
| 8815 | *SNRPB2* | 2.0766 | 1.0578 | -0.04085 | -0.91307 | 1.03 |
| 8816 | *LRRK1* | 2.4223 | 0.85084 | -0.17986 | -0.30525 | 1.03 |
| 8817 | *FBXO8* | 4.4808 | -0.61358 | -0.77462 | -0.81531 | 1.03 |
| 8818 | *CACNA1E* | 1.8765 | 0.98304 | 0.23292 | -0.72142 | 1.03 |
| 8819 | *RPE65* | 4.2183 | -0.17129 | -0.95461 | -1.7591 | 1.03 |
| 8820 | *RCAN2* | 2.1611 | 0.82695 | 0.10416 | 0.10106 | 1.03 |
| 8821 | *PPAP2B* | 1.1869 | 1.1027 | 0.8022 | -1.5997 | 1.03 |
| 8822 | *CSMD3* | 3.2009 | 0.26809 | -0.378 | -0.402 | 1.03 |
| 8823 | *WDHD1* | 5.17 | -0.63348 | -1.4467 | -2.2311 | 1.03 |
| 8824 | *C1GALT1C1* | 2.3758 | 0.44439 | 0.26891 | -1.9936 | 1.03 |
| 8825 | *POP1* | 2.8908 | 0.53063 | -0.33357 | -1.2104 | 1.03 |

| | | | | | | |
|---|---|---|---|---|---|---|
| 8826 | *BOLL* | 1.6322 | 1.1975 | 0.2579 | -0.55387 | 1.03 |
| 8827 | *ITPK1* | 2.2417 | 1.32 | -0.47438 | -1.6003 | 1.03 |
| 8828 | *INSM1* | 2.362 | 1.329 | -0.60439 | -2.4858 | 1.03 |
| 8829 | *TMEM44* | 2.6809 | 0.29032 | 0.11441 | -0.27925 | 1.03 |
| 8830 | *CYP1A1* | 4.7297 | -0.41447 | -1.2301 | -1.3686 | 1.03 |
| 8831 | *DTHD1* | 2.6999 | 0.9226 | -0.5377 | -1.0212 | 1.03 |
| 8832 | *UVRAG* | 3.4536 | 0.2124 | -0.58173 | -0.65924 | 1.03 |
| 8833 | *CCDC173* | 2.4036 | 1.358 | -0.67764 | -2.6485 | 1.03 |
| 8834 | *OR52E6* | 3.0881 | 0.48987 | -0.49483 | -0.80442 | 1.03 |
| 8835 | *IPP* | 3.9672 | -0.44133 | -0.44353 | -1.2273 | 1.03 |
| 8836 | *ENPP1* | 3.4065 | 0.06324 | -0.38767 | -1.9988 | 1.03 |
| 8837 | *RPTN* | 2.8728 | 0.14678 | 0.06247 | -0.39944 | 1.03 |
| 8838 | *TMEM177* | 1.5782 | 1.3279 | 0.17503 | -0.34964 | 1.03 |
| 8839 | *OR10G8* | 4.1191 | -0.11939 | -0.91886 | -2.9198 | 1.03 |
| 8840 | *PSMD4* | 2.398 | 0.3618 | 0.32041 | -2.1068 | 1.03 |
| 8841 | *CHST5* | 1.6245 | 1.2076 | 0.24723 | -0.2765 | 1.03 |
| 8842 | *HINT1* | 1.4037 | 1.1895 | 0.48548 | 0.15563 | 1.03 |
| 8843 | *SMN1* | 3.3708 | -0.07288 | -0.22006 | -1.8049 | 1.03 |
| 8844 | *PPP6R3* | 3.2713 | 1.5751 | -1.7688 | -1.9341 | 1.03 |
| 8845 | *SPTB* | 4.9707 | -0.43077 | -1.4625 | -2.2844 | 1.03 |
| 8846 | *SSMEM1* | 3.5251 | 0.019024 | -0.4674 | -1.474 | 1.03 |
| 8847 | *CCL2* | 2.6802 | 0.5365 | -0.14068 | -0.91357 | 1.03 |
| 8848 | *FAM156A* | 4.0595 | -0.25875 | -0.72612 | -0.86267 | 1.02 |
| 8849 | *PAPOLB* | 1.8835 | 0.74429 | 0.44594 | -1.2219 | 1.02 |
| 8850 | *EGLN2* | 4.1999 | -0.44572 | -0.68122 | -1.4469 | 1.02 |
| 8851 | *CASP5* | 2.03 | 1.0387 | 0.00408 | -0.48021 | 1.02 |
| 8852 | *FAT1* | 2.8801 | 0.41169 | -0.22081 | -0.77217 | 1.02 |
| 8853 | *FXYD3* | 3.7391 | -0.26798 | -0.40118 | -1.9326 | 1.02 |
| 8854 | *SGCE* | 1.5635 | 1.5385 | -0.03257 | -0.84542 | 1.02 |
| 8855 | *CRELD1* | 1.75 | 0.80329 | 0.51607 | -1.3599 | 1.02 |
| 8856 | *TBC1D31* | 3.5936 | -0.04968 | -0.47499 | -0.60054 | 1.02 |
| 8857 | *AGBL4* | 2.4898 | 0.53176 | 0.04723 | -1.8707 | 1.02 |
| 8858 | *NPL* | 3.5289 | -0.16814 | -0.29292 | -1.0529 | 1.02 |
| 8859 | *SLC22A14* | 3.1235 | 0.93401 | -0.99037 | -1.2906 | 1.02 |
| 8860 | *IFNL2* | 3.3238 | 1.204 | -1.4607 | -2.1018 | 1.02 |
| 8861 | *ADGRA3* | 3.9895 | 0.08603 | -1.0091 | -1.6559 | 1.02 |
| 8862 | *C19orf80* | 3.4111 | 0.78316 | -1.1282 | -1.8076 | 1.02 |
| 8863 | *TMEM243* | 2.4403 | 0.49111 | 0.13447 | -2.3804 | 1.02 |
| 8864 | *PSG5* | 4.4827 | -0.51845 | -0.89866 | -2.0267 | 1.02 |
| 8865 | *PHYH* | 2.31 | 0.69252 | 0.06301 | -0.78229 | 1.02 |
| 8866 | *ANKRD20A4* | 1.599 | 1.092 | 0.37379 | -0.37704 | 1.02 |
| 8867 | *MMADHC* | 3.5018 | 0.61867 | -1.0561 | -2.6973 | 1.02 |
| 8868 | *PARL* | 3.4043 | 0.50725 | -0.8475 | -1.6907 | 1.02 |
| 8869 | *RND3* | 2.8164 | 0.38155 | -0.13436 | -0.69128 | 1.02 |
| 8870 | *FZD10* | 3.6345 | 0.21342 | -0.78444 | -0.87673 | 1.02 |
| 8871 | *SPOCK1* | 3.0177 | 0.50672 | -0.46155 | -0.95933 | 1.02 |

| | | | | | | |
|---|---|---|---|---|---|---|
| 8872 | *ANGPTL7* | 3.7351 | 0.18707 | -0.85943 | -1.4567 | 1.02 |
| 8873 | *NRM* | 3.3255 | 1.0421 | -1.305 | -1.4299 | 1.02 |
| 8874 | *AFTPH* | 3.4413 | 0.81109 | -1.1898 | -1.536 | 1.02 |
| 8875 | *GDAP1L1* | 2.8394 | 0.67439 | -0.45191 | -1.0265 | 1.02 |
| 8876 | *BCAR3* | 2.3891 | 0.39999 | 0.27273 | -0.24269 | 1.02 |
| 8877 | *XAGE3* | 3.934 | 0.66498 | -1.5375 | -1.6586 | 1.02 |
| 8878 | *UNC119B* | 2.7795 | 0.35184 | -0.0702 | -0.27672 | 1.02 |
| 8879 | *CLDN16* | 3.9877 | -0.14198 | -0.78472 | -1.6524 | 1.02 |
| 8880 | *BOC* | 2.1746 | 0.52169 | 0.36413 | -0.76361 | 1.02 |
| 8881 | *ZBED9* | 3.4089 | 0.53353 | -0.88203 | -1.8642 | 1.02 |
| 8882 | *AIRE* | 4.4743 | -0.3576 | -1.0564 | -1.4151 | 1.02 |
| 8883 | *GLIPR1* | 3.8823 | -0.06824 | -0.75408 | -1.1617 | 1.02 |
| 8884 | *SNCG* | 5.0882 | -0.06618 | -1.9622 | -2.1947 | 1.02 |
| 8885 | *RAB17* | 3.0599 | 0.19327 | -0.19354 | -0.84426 | 1.02 |
| 8886 | *AQP7* | 2.7136 | 0.50617 | -0.16038 | -0.95527 | 1.02 |
| 8887 | *ACTN2* | 3.0796 | 0.53281 | -0.55311 | -1.6847 | 1.02 |
| 8888 | *CHUK* | 3.0686 | 0.016271 | -0.02658 | -0.40738 | 1.02 |
| 8889 | *STAMBP* | 3.1971 | 0.7294 | -0.86821 | -2.4472 | 1.02 |
| 8890 | *OR4Q3* | 3.3657 | 0.71832 | -1.0267 | -1.8077 | 1.02 |
| 8891 | *FAM173B* | 5.9432 | -0.92427 | -1.962 | -2.0851 | 1.02 |
| 8892 | *GUCA1C* | 4.1886 | 0.28749 | -1.4197 | -1.5781 | 1.02 |
| 8893 | *MED31* | 2.315 | 0.64847 | 0.09277 | -2.1287 | 1.02 |
| 8894 | *SLURP1* | 2.1736 | 1.4296 | -0.54708 | -1.0481 | 1.02 |
| 8895 | *ZNF235* | 4.1989 | 0.3057 | -1.4485 | -1.8725 | 1.02 |
| 8896 | *TMOD3* | 1.9611 | 1.0137 | 0.07872 | -2.6635 | 1.02 |
| 8897 | *SLC40A1* | 3.7128 | 0.88608 | -1.5454 | -2.2276 | 1.02 |
| 8898 | *PPBP* | 2.5422 | 1.1447 | -0.63357 | -1.4465 | 1.02 |
| 8899 | *IGFLR1* | 4.6865 | -0.70391 | -0.92956 | -1.344 | 1.02 |
| 8900 | *OR2A12* | 3.4939 | 0.57693 | -1.0178 | -1.4935 | 1.02 |
| 8901 | *LENG1* | 3.6812 | 0.11798 | -0.74815 | -1.035 | 1.02 |
| 8902 | *DBT* | 2.1953 | 1.4108 | -0.55531 | -1.1706 | 1.02 |
| 8903 | *XKR3* | 4.1088 | -0.34014 | -0.71819 | -1.9619 | 1.02 |
| 8904 | *C8orf48* | 3.461 | 0.46602 | -0.87694 | -1.0104 | 1.02 |
| 8905 | *KCTD2* | 2.3475 | 0.3669 | 0.33545 | -1.4487 | 1.02 |
| 8906 | *INHA* | 2.5353 | 0.72132 | -0.20681 | -1.8909 | 1.02 |
| 8907 | *NMU* | 4.4242 | -0.01876 | -1.3559 | -1.9168 | 1.02 |
| 8908 | *NLRP4* | 3.1711 | 0.282 | -0.40374 | -1.5462 | 1.02 |
| 8909 | *HSD17B12* | 3.5036 | 0.13485 | -0.59008 | -1.9705 | 1.02 |
| 8910 | *IL17F* | 2.0317 | 1.1631 | -0.14649 | -1.4359 | 1.02 |
| 8911 | *GPRC6A* | 1.064 | 0.99401 | 0.98991 | 0.60444 | 1.02 |
| 8912 | *MIA* | 3.6188 | 0.2269 | -0.79798 | -1.4364 | 1.02 |
| 8913 | *ADRA2B* | 3.9441 | -0.07649 | -0.82 | -1.7553 | 1.02 |
| 8914 | *MRPS27* | 4.9951 | -0.74815 | -1.1998 | -1.4299 | 1.02 |
| 8915 | *CLPS* | 4.0913 | -0.02692 | -1.0175 | -1.479 | 1.02 |
| 8916 | *MTRNR2L5* | 1.9642 | 0.74969 | 0.33271 | -1.4364 | 1.02 |
| 8917 | *CYTH4* | 1.693 | 0.83303 | 0.51984 | -1.3783 | 1.02 |

| | | | | | | |
|---|---|---|---|---|---|---|
| 8918 | *DTD1* | 1.3342 | 1.2033 | 0.50771 | -1.2867 | 1.02 |
| 8919 | *SYCP2* | 2.1411 | 0.80418 | 0.09972 | -0.60841 | 1.02 |
| 8920 | *ACTN1* | 3.6628 | 1.1692 | -1.7874 | -2.2277 | 1.01 |
| 8921 | *IL5* | 3.7613 | 0.1111 | -0.82811 | -1.505 | 1.01 |
| 8922 | *MICALL1* | 2.1181 | 0.48969 | 0.43614 | -0.77185 | 1.01 |
| 8923 | *KHSRP* | 6.0156 | -1.0718 | -1.8999 | -2.0837 | 1.01 |
| 8924 | *USP17L30* | 2.22 | 1.3373 | -0.51363 | -0.55492 | 1.01 |
| 8925 | *AATK* | 3.3181 | 0.53611 | -0.81072 | -2.1111 | 1.01 |
| 8926 | *DCAF12* | 2.8984 | 0.54977 | -0.40642 | -1.0884 | 1.01 |
| 8927 | *OR2A5* | 2.5323 | 0.94018 | -0.43077 | -1.8478 | 1.01 |
| 8928 | *COL1A2* | 3.0392 | 0.17599 | -0.17403 | -0.30038 | 1.01 |
| 8929 | *RASSF8* | 3.1683 | 0.89685 | -1.0243 | -1.6463 | 1.01 |
| 8930 | *B3GALT5* | 2.9836 | 0.12081 | -0.06364 | -2.5386 | 1.01 |
| 8931 | *NARR* | 4.0094 | -0.00439 | -0.96471 | -1.3209 | 1.01 |
| 8932 | *CYP2C8* | 4.8738 | -0.39389 | -1.4402 | -2.0303 | 1.01 |
| 8933 | *CRHR1* | 4.3539 | -0.63826 | -0.6765 | -0.72326 | 1.01 |
| 8934 | *CRB2* | 2.9179 | 1.0612 | -0.94027 | -1.3266 | 1.01 |
| 8935 | *NXPH4* | 4.4128 | -0.10845 | -1.2665 | -2.2692 | 1.01 |
| 8936 | *NDUFS6* | 2.3808 | 1.5575 | -0.90154 | -1.5984 | 1.01 |
| 8937 | *REM2* | 4.0353 | -0.13096 | -0.86793 | -2.4784 | 1.01 |
| 8938 | *FAM186A* | 2.4863 | 0.81532 | -0.26617 | -1.1289 | 1.01 |
| 8939 | *PPIP5K1* | 1.975 | 0.75422 | 0.30583 | -0.61852 | 1.01 |
| 8940 | *FABP12* | 2.4867 | 0.33905 | 0.20899 | -2.5334 | 1.01 |
| 8941 | *RPGR* | 3.5203 | 0.44492 | -0.93112 | -1.2086 | 1.01 |
| 8942 | *CT47A11* | 3.1799 | 0.78283 | -0.92911 | -1.9446 | 1.01 |
| 8943 | *AP2A1* | 1.5899 | 1.0944 | 0.34843 | 0.12788 | 1.01 |
| 8944 | *VPS51* | 3.5352 | 0.63571 | -1.1385 | -1.407 | 1.01 |
| 8945 | *BIN2* | 3.5072 | 0.72975 | -1.2051 | -1.4141 | 1.01 |
| 8946 | *MAMSTR* | 4.2981 | 0.008409 | -1.2753 | -1.9513 | 1.01 |
| 8947 | *ZPLD1* | 2.1163 | 1.0715 | -0.15678 | -1.6755 | 1.01 |
| 8948 | *APOLD1* | 2.8419 | 0.9954 | -0.80641 | -2.0198 | 1.01 |
| 8949 | *HLA-G* | 3.0675 | 1.7164 | -1.7541 | -2.5477 | 1.01 |
| 8950 | *MED26* | 2.5496 | 1.7192 | -1.2391 | -1.5644 | 1.01 |
| 8951 | *PIGQ* | 2.4363 | 0.52691 | 0.06617 | -0.23536 | 1.01 |
| 8952 | *DIRAS3* | 1.2571 | 1.1341 | 0.63785 | 0.39417 | 1.01 |
| 8953 | *ARHGAP21* | 3.0388 | 0.68386 | -0.69497 | -1.9935 | 1.01 |
| 8954 | *CCL22* | 3.2805 | 0.26885 | -0.52167 | -0.79934 | 1.01 |
| 8955 | *C2orf70* | 4.4012 | -0.21964 | -1.1545 | -1.4325 | 1.01 |
| 8956 | *KRCC1* | 3.4469 | 0.58818 | -1.0095 | -1.577 | 1.01 |
| 8957 | *SLC6A2* | 3.5719 | -0.15153 | -0.39494 | -0.96173 | 1.01 |
| 8958 | *PRDM16* | 4.6127 | -0.61515 | -0.97252 | -2.0488 | 1.01 |
| 8959 | *PDE4C* | 2.1391 | 1.6496 | -0.76379 | -0.93525 | 1.01 |
| 8960 | *CMKLR1* | 3.4725 | 0.11075 | -0.56003 | -0.97549 | 1.01 |
| 8961 | *LOC730159* | 2.3141 | 0.72758 | -0.01854 | -0.41642 | 1.01 |
| 8962 | *METTL11B* | 3.6448 | -0.14906 | -0.4744 | -2.9632 | 1.01 |
| 8963 | *MED13L* | 2.6925 | 0.47245 | -0.14563 | -0.17323 | 1.01 |

| | | | | | | |
|---|---|---|---|---|---|---|
| 8964 | *SRY* | 4.6778 | -0.37125 | -1.2876 | -2.1696 | 1.01 |
| 8965 | *S100A8* | 3.9913 | -0.00829 | -0.96497 | -2.4112 | 1.01 |
| 8966 | *KLHL41* | 3.3922 | 0.06609 | -0.44136 | -1.0212 | 1.01 |
| 8967 | *PLA2G15* | 2.7634 | 0.64093 | -0.38754 | -0.84395 | 1.01 |
| 8968 | *TMEM260* | 2.8475 | 0.34671 | -0.17752 | -0.53029 | 1.01 |
| 8969 | *LSP1* | 1.6511 | 0.90252 | 0.46288 | -1.1713 | 1.01 |
| 8970 | *MMP25* | 4.0456 | 0.059245 | -1.0884 | -1.8341 | 1.01 |
| 8971 | *TENM4* | 4.0392 | 0.44389 | -1.468 | -2.6939 | 1.01 |
| 8972 | *PRAC2* | 3.4218 | 0.64939 | -1.0564 | -1.6763 | 1.00 |
| 8973 | *FARP1* | 3.7033 | -0.22943 | -0.46017 | -2.0908 | 1.00 |
| 8974 | *CDC73* | 2.9981 | 0.26556 | -0.25022 | -2.2644 | 1.00 |
| 8975 | *SESN1* | 4.0759 | -0.11897 | -0.94536 | -0.96395 | 1.00 |
| 8976 | *DUSP5* | 2.87 | 1.0578 | -0.91701 | -2.0675 | 1.00 |
| 8977 | *SERTM1* | 3.9686 | 0.38081 | -1.3387 | -1.7407 | 1.00 |
| 8978 | *ARHGEF9* | 3.1592 | 1.0898 | -1.2386 | -1.8293 | 1.00 |
| 8979 | *ZNF436* | 2.2793 | 1.2514 | -0.5206 | -0.73823 | 1.00 |
| 8980 | *HUS1B* | 1.8952 | 1.2392 | -0.12433 | -1.9338 | 1.00 |
| 8981 | *TSC22D1* | 1.2851 | 1.1512 | 0.57229 | -0.71317 | 1.00 |
| 8982 | *PROM2* | 2.5624 | 1.5177 | -1.072 | -1.5671 | 1.00 |
| 8983 | *KRTAP9-2* | 2.2981 | 0.55603 | 0.15285 | -1.6143 | 1.00 |
| 8984 | *WFDC13* | 2.5899 | 0.5712 | -0.15567 | -1.268 | 1.00 |
| 8985 | *FXYD4* | 3.4824 | -0.02342 | -0.45443 | -0.99418 | 1.00 |
| 8986 | *GNG3* | 4.1056 | -0.24591 | -0.85552 | -1.1756 | 1.00 |
| 8987 | *ZNF333* | 1.3421 | 0.89593 | 0.76588 | -0.73728 | 1.00 |
| 8988 | *C10orf25* | 3.2349 | 0.074008 | -0.30607 | -1.2884 | 1.00 |
| 8989 | *MOBP* | 2.3206 | 0.80359 | -0.12213 | -0.53361 | 1.00 |
| 8990 | *FIZ1* | 2.3968 | 0.60687 | -0.00164 | -0.48101 | 1.00 |
| 8991 | *CD1D* | 3.5188 | 0.54668 | -1.0635 | -1.257 | 1.00 |
| 8992 | *PTPRN* | 2.7879 | 1.0757 | -0.86182 | -2.3145 | 1.00 |
| 8993 | *ANKRD13B* | 1.3331 | 0.93177 | 0.73674 | -1.4234 | 1.00 |
| 8994 | *FNDC8* | 3.0985 | 0.30051 | -0.39779 | -0.67731 | 1.00 |
| 8995 | *KRT82* | 1.7948 | 0.66293 | 0.54271 | -1.9659 | 1.00 |
| 8996 | *IFI16* | 3.1216 | 0.15514 | -0.27676 | -0.81509 | 1.00 |
| 8997 | *MANSC4* | 3.6962 | -0.01874 | -0.67801 | -1.6356 | 1.00 |
| 8998 | *DCLRE1A* | 3.4129 | 0.13535 | -0.54889 | -1.3419 | 1.00 |
| 8999 | *LRP11* | 2.7187 | 0.16358 | 0.11566 | -0.14427 | 1.00 |
| 9000 | *AK9* | 1.4294 | 1.0145 | 0.55379 | 0.19481 | 1.00 |
| 9001 | *ADAMTSL4* | 3.2182 | -0.06587 | -0.15464 | -0.58005 | 1.00 |
| 9002 | *CLEC3B* | 2.6881 | 0.60138 | -0.29186 | -1.2474 | 1.00 |
| 9003 | *ZNF35* | 3.4853 | -0.0265 | -0.46124 | -1.4062 | 1.00 |
| 9004 | *MYL3* | 2.5154 | 1.0341 | -0.55207 | -1.4414 | 1.00 |
| 9005 | *GBA* | 3.6178 | 0.30722 | -0.9279 | -1.0948 | 1.00 |
| 9006 | *TEC* | 2.8036 | 0.17295 | 0.02003 | -2.0593 | 1.00 |
| 9007 | *MMD* | 2.479 | 0.50113 | 0.01511 | -0.20532 | 1.00 |
| 9008 | *INPP1* | 2.2553 | 0.56457 | 0.17526 | -0.51182 | 1.00 |
| 9009 | *PABPC5* | 2.225 | 0.7435 | 0.02648 | -4.1136 | 1.00 |

| | | | | | | |
|---|---|---|---|---|---|---|
| 9010 | *ANKRD33B* | 4.3933 | 0.3701 | -1.7688 | -2.1357 | 1.00 |
| 9011 | *TRIM28* | 2.8247 | 0.2669 | -0.09731 | -0.41642 | 1.00 |
| 9012 | *ALS2* | 4.3147 | -0.03091 | -1.2896 | -2.298 | 1.00 |
| 9013 | *CCDC67* | 4.3459 | -0.30877 | -1.043 | -2.7221 | 1.00 |
| 9014 | *MTURN* | 2.3177 | 0.78549 | -0.10911 | -2.2075 | 1.00 |
| 9015 | *ASIP* | 2.4221 | 1.3946 | -0.82345 | -2.3101 | 1.00 |
| 9016 | *LMO3* | 2.2632 | 1.2839 | -0.55424 | -1.5544 | 1.00 |
| 9017 | *SPACA3* | 3.7671 | 0.67638 | -1.4507 | -1.5912 | 1.00 |
| 9018 | *SHANK3* | 3.0007 | 0.48128 | -0.49016 | -1.1685 | 1.00 |
| 9019 | *RMI2* | 2.6024 | 0.66261 | -0.27548 | -1.6323 | 1.00 |
| 9020 | *HIST1H3E* | 5.1982 | -0.15148 | -2.0574 | -2.278 | 1.00 |
| 9021 | *CEP76* | 1.8612 | 1.7298 | -0.60262 | -1.1199 | 1.00 |
| 9022 | *ATG2A* | 1.6646 | 0.85125 | 0.47215 | 0.11075 | 1.00 |
| 9023 | *FBXL19* | 3.7955 | -0.38428 | -0.42356 | -1.578 | 1.00 |
| 9024 | *VDAC2* | 2.7654 | 0.21699 | 0.00499 | -1.4615 | 1.00 |
| 9025 | *C6orf47* | 3.9191 | -0.31508 | -0.61829 | -1.4932 | 1.00 |
| 9026 | *SMCO3* | 1.2847 | 0.89979 | 0.80093 | -0.66488 | 1.00 |
| 9027 | *IPO13* | 3.2947 | 0.57564 | -0.88507 | -1.7807 | 1.00 |
| 9028 | *NLRP14* | 5.5107 | -0.71009 | -1.8157 | -2.3032 | 0.99 |
| 9029 | *CTSE* | 2.6504 | 0.49173 | -0.15746 | -1.3184 | 0.99 |
| 9030 | *LGR6* | 3.1539 | 0.090093 | -0.25941 | -0.44866 | 0.99 |
| 9031 | *LHFPL3* | 1.9365 | 0.66302 | 0.3849 | -2.5934 | 0.99 |
| 9032 | *CHRNA9* | 3.6242 | -0.28994 | -0.35044 | -1.2464 | 0.99 |
| 9033 | *BAZ2B* | 3.6769 | -0.02287 | -0.67138 | -2.3096 | 0.99 |
| 9034 | *SIGLECL1* | 2.8831 | 1.219 | -1.12 | -1.4994 | 0.99 |
| 9035 | *RSPO3* | 4.007 | 0.74959 | -1.7745 | -3.5428 | 0.99 |
| 9036 | *KRT12* | 1.5256 | 1.0352 | 0.42116 | -1.5962 | 0.99 |
| 9037 | *ATP9B* | 3.0833 | 1.4025 | -1.5039 | -1.6636 | 0.99 |
| 9038 | *CWC22* | 3.8166 | 0.65738 | -1.4924 | -1.6243 | 0.99 |
| 9039 | *LGALS9* | 3.392 | 0.50675 | -0.91874 | -1.0688 | 0.99 |
| 9040 | *OR2AG1* | 1.8051 | 1.4952 | -0.3205 | -0.37417 | 0.99 |
| 9041 | *GAGE12I* | 2.169 | 2.0929 | -1.2827 | -1.8102 | 0.99 |
| 9042 | *RNF2* | 3.0988 | 0.054625 | -0.17432 | -1.278 | 0.99 |
| 9043 | *ANKRD55* | 2.7454 | 0.59967 | -0.36689 | -1.6085 | 0.99 |
| 9044 | *ZRSR2* | 3.6445 | 0.19726 | -0.86358 | -1.3557 | 0.99 |
| 9045 | *IRF1* | 4.8308 | -0.52426 | -1.3285 | -1.6597 | 0.99 |
| 9046 | *CMTM8* | 3.485 | 0.5883 | -1.0953 | -1.161 | 0.99 |
| 9047 | *ZNF180* | 2.165 | 1.0832 | -0.2705 | -0.45152 | 0.99 |
| 9048 | *SEMA6B* | 3.6883 | -0.17054 | -0.54008 | -1.2792 | 0.99 |
| 9049 | *FARS2* | 3.0505 | 0.26985 | -0.34275 | -0.80061 | 0.99 |
| 9050 | *KIAA1024* | 5.0758 | -0.25597 | -1.8427 | -2.8628 | 0.99 |
| 9051 | *PRAMEF15* | 2.0277 | 1.7712 | -0.82208 | -1.6764 | 0.99 |
| 9052 | *LPCAT3* | 1.35 | 1.0769 | 0.54939 | -0.18229 | 0.99 |
| 9053 | *LSM11* | 3.324 | 0.54891 | -0.89667 | -1.803 | 0.99 |
| 9054 | *HFM1* | 2.2777 | 1.125 | -0.42658 | -1.8598 | 0.99 |
| 9055 | *FAM50A* | 3.7177 | 0.6833 | -1.4256 | -2.0553 | 0.99 |

| | | | | | | |
|---|---|---|---|---|---|---|
| 9056 | *ESPN* | 5.0614 | -0.47327 | -1.6129 | -1.7743 | 0.99 |
| 9057 | *FN3KRP* | 4.5944 | -0.35652 | -1.2628 | -1.3574 | 0.99 |
| 9058 | *ERICH1* | 1.4648 | 1.1674 | 0.34276 | -0.78229 | 0.99 |
| 9059 | *PECAM1* | 3.3543 | -0.17238 | -0.2072 | -0.51469 | 0.99 |
| 9060 | *CHRFAM7A* | 1.4106 | 0.84854 | 0.71536 | 0.062078 | 0.99 |
| 9061 | *MUM1* | 4.0893 | -0.2505 | -0.86479 | -2.555 | 0.99 |
| 9062 | *ALG10B* | 2.9177 | 0.16038 | -0.1049 | -2.1576 | 0.99 |
| 9063 | *NEUROD6* | 3.234 | 0.82344 | -1.0843 | -1.3016 | 0.99 |
| 9064 | *SNX32* | 3.3575 | 0.48331 | -0.86861 | -2.2171 | 0.99 |
| 9065 | *ARIH2* | 3.5509 | -0.23428 | -0.34535 | -0.72038 | 0.99 |
| 9066 | *RELA* | 3.3522 | -0.1666 | -0.21563 | -2.224 | 0.99 |
| 9067 | *EPS8L2* | 3.7651 | -0.26098 | -0.53437 | -1.8591 | 0.99 |
| 9068 | *ING1* | 3.27 | 0.004671 | -0.30535 | -2.7736 | 0.99 |
| 9069 | *ANP32D* | 2.8828 | 0.20422 | -0.11788 | -0.41359 | 0.99 |
| 9070 | *ATP1A2* | 1.4164 | 0.81004 | 0.74243 | 0.44887 | 0.99 |
| 9071 | *CNOT1* | 2.5459 | 0.32329 | 0.09956 | -1.8223 | 0.99 |
| 9072 | *ZBED6CL* | 3.5713 | -0.06413 | -0.53845 | -0.92214 | 0.99 |
| 9073 | *MARC2* | 3.5304 | 1.0298 | -1.5916 | -1.9404 | 0.99 |
| 9074 | *PAICS* | 4.2383 | -0.60655 | -0.66362 | -1.0576 | 0.99 |
| 9075 | *RAB40B* | 2.196 | 1.1243 | -0.35447 | -1.7538 | 0.99 |
| 9076 | *MYH10* | 1.6035 | 0.76737 | 0.59474 | -2.0214 | 0.99 |
| 9077 | *AGPS* | 3.024 | 0.095844 | -0.15499 | -0.39017 | 0.99 |
| 9078 | *CRISP2* | 3.2864 | 0.002484 | -0.32495 | -2.7221 | 0.99 |
| 9079 | *ARHGAP31* | 3.6555 | 0.006708 | -0.70051 | -0.7757 | 0.99 |
| 9080 | *WNK2* | 2.6418 | 0.45325 | -0.13353 | -0.80249 | 0.99 |
| 9081 | *DDX54* | 2.1083 | 0.89187 | -0.03962 | -0.57078 | 0.99 |
| 9082 | *RABIF* | 3.4464 | -0.03492 | -0.45254 | -1.6495 | 0.99 |
| 9083 | *CCDC116* | 3.1361 | 0.94342 | -1.1207 | -1.9047 | 0.99 |
| 9084 | *EXOC3L4* | 2.7964 | 0.67915 | -0.51722 | -1.3954 | 0.99 |
| 9085 | *CSNK1G3* | 1.4257 | 1.0702 | 0.46191 | -1.4622 | 0.99 |
| 9086 | *TEX264* | 2.7436 | 1.6993 | -1.4852 | -1.7709 | 0.99 |
| 9087 | *RNF224* | 3.0568 | 0.24928 | -0.34927 | -1.448 | 0.99 |
| 9088 | *NRCAM* | 2.8461 | 0.24289 | -0.13244 | -0.51302 | 0.99 |
| 9089 | *LRP6* | 3.5073 | 0.30398 | -0.855 | -2.179 | 0.99 |
| 9090 | *ZNF329* | 4.2139 | 0.32898 | -1.5866 | -1.7153 | 0.99 |
| 9091 | *RAB3IL1* | 3.7902 | -0.18039 | -0.6536 | -1.6251 | 0.99 |
| 9092 | *HEPACAM* | 3.0032 | 0.36038 | -0.40761 | -1.0826 | 0.99 |
| 9093 | *TPSB2* | 2.5098 | 0.23771 | 0.20794 | -0.42074 | 0.99 |
| 9094 | *TTC14* | 3.2298 | 0.071102 | -0.34558 | -2.7008 | 0.99 |
| 9095 | *H1FNT* | 2.9934 | 1.3432 | -1.3813 | -2.1143 | 0.99 |
| 9096 | *ZFP30* | 1.1865 | 1.0724 | 0.69576 | -1.1995 | 0.98 |
| 9097 | *CALR3* | 2.8546 | 0.71901 | -0.61958 | -1.9794 | 0.98 |
| 9098 | *NGB* | 3.7967 | -0.18717 | -0.65581 | -1.0908 | 0.98 |
| 9099 | *PBX2* | 2.9458 | 0.11388 | -0.10615 | -1.3827 | 0.98 |
| 9100 | *ATP5H* | 1.2805 | 1.0105 | 0.66253 | -1.4282 | 0.98 |
| 9101 | *RAB32* | 2.1478 | 1.4529 | -0.64748 | -1.25 | 0.98 |

| | | | | | | |
|---|---|---|---|---|---|---|
| 9102 | *SCGB1C1* | 2.3894 | 0.94458 | -0.38112 | -1.6654 | 0.98 |
| 9103 | *NOC2L* | 4.181 | 0.33766 | -1.5658 | -1.8405 | 0.98 |
| 9104 | *ABHD12* | 2.6646 | 0.16344 | 0.12402 | -1.4997 | 0.98 |
| 9105 | *EDAR* | 1.9337 | 1.7062 | -0.68818 | -1.3298 | 0.98 |
| 9106 | *ADAMTS16* | 4.05 | 0.2003 | -1.2987 | -3.1335 | 0.98 |
| 9107 | *KIAA1644* | 1.9024 | 0.84645 | 0.20268 | -2.312 | 0.98 |
| 9108 | *TBATA* | 1.42 | 1.0564 | 0.47468 | -1.4754 | 0.98 |
| 9109 | *SEMA4F* | 2.9763 | 0.17739 | -0.20284 | -0.20338 | 0.98 |
| 9110 | *NAGS* | 1.8634 | 1.6843 | -0.59696 | -1.0342 | 0.98 |
| 9111 | *CCDC12* | 2.2898 | 0.4413 | 0.21879 | -0.69615 | 0.98 |
| 9112 | *PRSS55* | 3.3475 | 0.045365 | -0.44299 | -1.2656 | 0.98 |
| 9113 | *CFAP46* | 2.6354 | 1.0513 | -0.73708 | -0.84659 | 0.98 |
| 9114 | *C15orf56* | 2.3026 | 0.82598 | -0.17928 | -0.27631 | 0.98 |
| 9115 | *BAHCC1* | 4.9089 | -0.41738 | -1.544 | -2.1412 | 0.98 |
| 9116 | *ACSM2A* | 5.0652 | -0.8606 | -1.2574 | -1.4467 | 0.98 |
| 9117 | *LRRIQ1* | 1.8591 | 1.276 | -0.18822 | -1.0271 | 0.98 |
| 9118 | *SNRPE* | 1.7132 | 1.0795 | 0.15417 | -1.5658 | 0.98 |
| 9119 | *C22orf15* | 2.5758 | 0.39754 | -0.02667 | -0.13281 | 0.98 |
| 9120 | *SORD* | 1.5983 | 1.2992 | 0.04902 | -1.1564 | 0.98 |
| 9121 | *C5orf47* | 3.304 | 0.055982 | -0.41383 | -1.6695 | 0.98 |
| 9122 | *CMTM6* | 3.4386 | 0.47673 | -0.96962 | -1.049 | 0.98 |
| 9123 | *EPC1* | 4.3165 | -0.05477 | -1.3165 | -2.2955 | 0.98 |
| 9124 | *NAT8B* | 3.4469 | 0.17441 | -0.67625 | -0.827 | 0.98 |
| 9125 | *PTS* | 4.9358 | -0.35735 | -1.6339 | -1.9237 | 0.98 |
| 9126 | *ELMOD2* | 3.1789 | 0.35655 | -0.59215 | -0.72053 | 0.98 |
| 9127 | *PRNP* | 3.399 | 0.32829 | -0.78444 | -1.9891 | 0.98 |
| 9128 | *ANHX* | 3.0159 | 0.96387 | -1.0372 | -1.7266 | 0.98 |
| 9129 | *MLNR* | 5.2551 | -0.61119 | -1.7018 | -2.4112 | 0.98 |
| 9130 | *ZNF75D* | 3.0332 | 0.44884 | -0.54101 | -2.1599 | 0.98 |
| 9131 | *NR2F2* | 2.4042 | 0.58598 | -0.04919 | -0.14793 | 0.98 |
| 9132 | *DEPTOR* | 3.311 | 0.18336 | -0.55359 | -1.4783 | 0.98 |
| 9133 | *PRAMEF8* | 2.9873 | 0.052475 | -0.09982 | -0.14858 | 0.98 |
| 9134 | *SLC43A3* | 4.0196 | -0.48486 | -0.59521 | -0.68222 | 0.98 |
| 9135 | *AKT1* | 2.8587 | 0.30689 | -0.22617 | -0.5204 | 0.98 |
| 9136 | *LRBA* | 4.1914 | -0.44198 | -0.81034 | -0.86698 | 0.98 |
| 9137 | *IFNGR1* | 4.497 | -0.74011 | -0.81798 | -1.2955 | 0.98 |
| 9138 | *TP53BP2* | 1.4071 | 1.1234 | 0.40836 | -0.10044 | 0.98 |
| 9139 | *ANKRD20A2* | 2.3161 | 1.8075 | -1.1848 | -1.6636 | 0.98 |
| 9140 | *DDT* | 3.0399 | 0.30927 | -0.41087 | -2.6485 | 0.98 |
| 9141 | *DEFB116* | 2.2408 | 0.49507 | 0.20228 | -0.94848 | 0.98 |
| 9142 | *CYP2A13* | 2.107 | 0.86544 | -0.03586 | -0.27903 | 0.98 |
| 9143 | *ZNF154* | 1.2419 | 0.95004 | 0.74421 | -0.75192 | 0.98 |
| 9144 | *MTPN* | 3.771 | 0.31864 | -1.1537 | -2.056 | 0.98 |
| 9145 | *SP9* | 1.3408 | 0.84242 | 0.75259 | 0.49538 | 0.98 |
| 9146 | *TBCE* | 2.7864 | 1.3877 | -1.2386 | -1.2935 | 0.98 |
| 9147 | *HRH1* | 2.0518 | 0.66151 | 0.22209 | -0.15385 | 0.98 |

| | | | | | | |
|---|---|---|---|---|---|---|
| 9148 | *ECH1* | 1.7263 | 1.6641 | -0.45553 | -2.0347 | 0.98 |
| 9149 | *METTL8* | 3.7406 | 0.15092 | -0.95682 | -1.5458 | 0.98 |
| 9150 | *MRPL37* | 4.3924 | -0.1342 | -1.3237 | -1.7493 | 0.98 |
| 9151 | *PTPN18* | 4.388 | -0.31869 | -1.1351 | -1.5396 | 0.98 |
| 9152 | *UVSSA* | 2.346 | 0.97686 | -0.38901 | -1.0224 | 0.98 |
| 9153 | *FAM105A* | 3.9472 | 0.71324 | -1.7266 | -3.064 | 0.98 |
| 9154 | *BTNL2* | 3.3373 | -0.14247 | -0.26159 | -0.80581 | 0.98 |
| 9155 | *PNPO* | 2.0429 | 1.5579 | -0.66897 | -2.3719 | 0.98 |
| 9156 | *KCTD7* | 2.1651 | 0.46439 | 0.30209 | -1.5115 | 0.98 |
| 9157 | *ZC2HC1A* | 2.0922 | 0.76694 | 0.07197 | -1.5544 | 0.98 |
| 9158 | *RAB11A* | 3.8976 | -0.44353 | -0.52392 | -1.5722 | 0.98 |
| 9159 | *HIST1H2BG* | 1.6003 | 1.4728 | -0.14304 | -0.73668 | 0.98 |
| 9160 | *IBTK* | 1.5841 | 0.87244 | 0.47122 | -1.2647 | 0.98 |
| 9161 | *FGL1* | 2.9792 | 0.29492 | -0.34675 | -1.1709 | 0.98 |
| 9162 | *GPX6* | 2.1492 | 0.46212 | 0.31588 | 0.14394 | 0.98 |
| 9163 | *TCP10L2* | 2.9045 | 0.10118 | -0.07893 | -0.15499 | 0.98 |
| 9164 | *EML3* | 2.0357 | 0.55567 | 0.33523 | 0.038507 | 0.98 |
| 9165 | *STK32B* | 2.9118 | 1.2013 | -1.1866 | -2.3795 | 0.98 |
| 9166 | *FMNL2* | 4.2655 | -0.52447 | -0.81505 | -1.2261 | 0.98 |
| 9167 | *LOC100134391* | 2.2261 | 0.38204 | 0.31742 | -0.30822 | 0.98 |
| 9168 | *PCSK5* | 1.3828 | 0.97 | 0.57247 | -0.49481 | 0.98 |
| 9169 | *NDUFA2* | 2.7021 | 0.94515 | -0.72294 | -0.93525 | 0.97 |
| 9170 | *VASP* | 4.5156 | -0.19249 | -1.3988 | -2.8823 | 0.97 |
| 9171 | *USP19* | 3.2289 | 0.84912 | -1.154 | -1.802 | 0.97 |
| 9172 | *KRTAP3-3* | 2.6784 | 1.8435 | -1.5984 | -2.0791 | 0.97 |
| 9173 | *C9orf40* | 2.1054 | 0.51917 | 0.29864 | -0.11527 | 0.97 |
| 9174 | *NAP1L1* | 3.272 | 0.83922 | -1.1881 | -2.6716 | 0.97 |
| 9175 | *DNAI1* | 3.193 | 0.58934 | -0.85931 | -2.1502 | 0.97 |
| 9176 | *OR5D16* | 2.6007 | 0.58937 | -0.26712 | -0.70532 | 0.97 |
| 9177 | *PRR5L* | 3.5071 | 0.041147 | -0.62649 | -1.9172 | 0.97 |
| 9178 | *CKMT2* | 3.8165 | -0.34199 | -0.55302 | -1.2426 | 0.97 |
| 9179 | *PACSIN1* | 1.5272 | 0.75428 | 0.63967 | -1.0949 | 0.97 |
| 9180 | *PLCD4* | 3.4011 | 0.13771 | -0.61779 | -2.29 | 0.97 |
| 9181 | *ADIRF* | 2.9607 | 0.96998 | -1.0097 | -1.6731 | 0.97 |
| 9182 | *POF1B* | 2.0921 | 0.70247 | 0.12494 | -2.0433 | 0.97 |
| 9183 | *BMP10* | 2.2945 | 0.53438 | 0.09061 | -1.3694 | 0.97 |
| 9184 | *PRR20E* | 2.7877 | 0.71607 | -0.5845 | -1.6755 | 0.97 |
| 9185 | *ZNF827* | 1.1772 | 0.93576 | 0.80518 | -1.1875 | 0.97 |
| 9186 | *RPRM* | 4.2649 | 0.39699 | -1.7438 | -2.3174 | 0.97 |
| 9187 | *SNAP29* | 3.336 | 0.42879 | -0.84674 | -1.8157 | 0.97 |
| 9188 | *PRKAG3* | 2.4377 | 0.99692 | -0.51745 | -0.65145 | 0.97 |
| 9189 | *TAC3* | 2.2436 | 0.61237 | 0.06116 | -0.24191 | 0.97 |
| 9190 | *PCDHB2* | 4.1791 | -0.41383 | -0.84816 | -3.1921 | 0.97 |
| 9191 | *OR5AR1* | 1.3882 | 1.3491 | 0.1795 | -1.4354 | 0.97 |
| 9192 | *OR7G1* | 3.8427 | 0.16896 | -1.0953 | -1.8995 | 0.97 |
| 9193 | *MT1E* | 3.4695 | 0.021388 | -0.57503 | -1.268 | 0.97 |

| | | | | | | |
|---|---|---|---|---|---|---|
| 9194 | *SLC26A11* | 2.7646 | 0.53121 | -0.38044 | -0.88143 | 0.97 |
| 9195 | *VPS11* | 3.6193 | 0.12446 | -0.8285 | -1.2472 | 0.97 |
| 9196 | *ZC3H6* | 1.7367 | 0.62912 | 0.54939 | -0.91028 | 0.97 |
| 9197 | *KRTAP5-8* | 1.3424 | 0.80548 | 0.76656 | 0.18083 | 0.97 |
| 9198 | *OR2M2* | 1.4823 | 0.9173 | 0.51379 | -1.94 | 0.97 |
| 9199 | *SLC27A4* | 2.2914 | 0.47748 | 0.14437 | -1.5448 | 0.97 |
| 9200 | *OTX1* | 3.6906 | -0.27333 | -0.50436 | -1.9943 | 0.97 |
| 9201 | *NCEH1* | 3.243 | 0.30923 | -0.63935 | -0.65393 | 0.97 |
| 9202 | *PAQR7* | 3.81 | -0.23297 | -0.66419 | -1.3266 | 0.97 |
| 9203 | *TMEM14A* | 3.2098 | 0.002272 | -0.29982 | -0.33191 | 0.97 |
| 9204 | *ADD2* | 2.6955 | 0.495 | -0.2783 | -0.37686 | 0.97 |
| 9205 | *GLYATL2* | 4.451 | -0.14828 | -1.3907 | -1.6276 | 0.97 |
| 9206 | *SOX11* | 2.6892 | 0.71252 | -0.48982 | -1.0323 | 0.97 |
| 9207 | *AMPD3* | 3.615 | 0.006369 | -0.70957 | -0.99448 | 0.97 |
| 9208 | *AP3D1* | 3.2042 | 0.24472 | -0.53766 | -0.54744 | 0.97 |
| 9209 | *GTF3C2* | 2.4831 | 1.6372 | -1.2094 | -1.3457 | 0.97 |
| 9210 | *CCL17* | 2.3897 | 1.0766 | -0.55601 | -0.86123 | 0.97 |
| 9211 | *TWIST1* | 3.4734 | 0.24772 | -0.81136 | -1.5687 | 0.97 |
| 9212 | *MYH4* | 3.1562 | 0.045209 | -0.29305 | -0.99526 | 0.97 |
| 9213 | *ATG13* | 3.5981 | -0.27707 | -0.41308 | -0.50474 | 0.97 |
| 9214 | *MIB1* | 4.4873 | -0.15943 | -1.4204 | -3.2419 | 0.97 |
| 9215 | *BPIFA3* | 3.7406 | 0.021312 | -0.85456 | -2.0277 | 0.97 |
| 9216 | *OARD1* | 4.0893 | -0.50419 | -0.67888 | -2.0927 | 0.97 |
| 9217 | *FPGS* | 4.0759 | 0.49846 | -1.6682 | -3.1229 | 0.97 |
| 9218 | *ADGRG6* | 1.7931 | 0.92282 | 0.19018 | -1.8046 | 0.97 |
| 9219 | *FAM160B2* | 1.5882 | 0.71542 | 0.60228 | -0.32466 | 0.97 |
| 9220 | *NSDHL* | 2.2471 | 0.4371 | 0.22117 | -0.43859 | 0.97 |
| 9221 | *SLC24A5* | 3.5726 | -0.05505 | -0.61329 | -0.66577 | 0.97 |
| 9222 | *TDRD9* | 4.1374 | -0.07224 | -1.1619 | -1.2574 | 0.97 |
| 9223 | *FBXO28* | 2.4771 | 1.0303 | -0.60628 | -1.0442 | 0.97 |
| 9224 | *SFTPA2* | 1.3733 | 0.78594 | 0.74098 | -0.95258 | 0.97 |
| 9225 | *GLT6D1* | 2.371 | 0.7515 | -0.22313 | -2.3629 | 0.97 |
| 9226 | *RAPGEF6* | 1.3316 | 0.7876 | 0.77938 | -2.3949 | 0.97 |
| 9227 | *OPRM1* | 1.909 | 0.58665 | 0.40281 | 0.16544 | 0.97 |
| 9228 | *RSAD2* | 4.5035 | -0.40427 | -1.2008 | -1.2097 | 0.97 |
| 9229 | *NUP88* | 1.5552 | 1.4705 | -0.12896 | -1.8995 | 0.97 |
| 9230 | *MBP* | 2.2984 | 1.9791 | -1.3808 | -1.6601 | 0.97 |
| 9231 | *RGS1* | 4.4869 | -0.64851 | -0.94171 | -1.8714 | 0.97 |
| 9232 | *NR4A1* | 1.5219 | 1.0386 | 0.33583 | -0.14762 | 0.97 |
| 9233 | *SLC34A2* | 1.7156 | 0.742 | 0.43844 | -1.4747 | 0.97 |
| 9234 | *ERCC6* | 4.3141 | 0.17607 | -1.5949 | -2.1108 | 0.97 |
| 9235 | *DMBX1* | 3.1248 | 0.82516 | -1.0564 | -2.0992 | 0.96 |
| 9236 | *CD163L1* | 2.4991 | 0.39392 | 0.00046 | -0.96878 | 0.96 |
| 9237 | *SSUH2* | 3.2241 | -0.05841 | -0.27244 | -1.1199 | 0.96 |
| 9238 | *STMND1* | 1.8417 | 0.8072 | 0.24339 | -1.5373 | 0.96 |
| 9239 | *HEPHL1* | 2.5618 | 0.71832 | -0.38815 | -1.8157 | 0.96 |

| | | | | | | |
|---|---|---|---|---|---|---|
| 9240 | *EXOSC5* | 2.9287 | 0.12559 | -0.16234 | -2.3452 | 0.96 |
| 9241 | *PLAUR* | 3.0465 | -0.07748 | -0.07903 | -1.1904 | 0.96 |
| 9242 | *LGALS7B* | 3.8126 | -0.11865 | -0.80675 | -2.8553 | 0.96 |
| 9243 | *ORMDL3* | 2.5128 | 0.22286 | 0.15053 | -0.77038 | 0.96 |
| 9244 | *RNF157* | 2.2569 | 0.4966 | 0.13265 | -1.9367 | 0.96 |
| 9245 | *MFSD11* | 3.2345 | -0.1496 | -0.19915 | -1.01 | 0.96 |
| 9246 | *KBTBD4* | 1.3941 | 0.93201 | 0.55956 | -1.765 | 0.96 |
| 9247 | *KDM1B* | 3.127 | 0.26849 | -0.50989 | -1.4019 | 0.96 |
| 9248 | *IL7R* | 1.9017 | 1.7869 | -0.80341 | -2.0134 | 0.96 |
| 9249 | *SULF1* | 3.8542 | -0.19043 | -0.7789 | -1.1529 | 0.96 |
| 9250 | *B3GNTL1* | 2.3889 | 0.5969 | -0.10116 | -0.35328 | 0.96 |
| 9251 | *ERGIC1* | 1.6161 | 1.2633 | 0.00452 | -1.3769 | 0.96 |
| 9252 | *DLX4* | 2.6855 | 1.2253 | -1.0273 | -1.6246 | 0.96 |
| 9253 | *INTS8* | 3.4365 | 0.29458 | -0.8478 | -2.5561 | 0.96 |
| 9254 | *ATP5L* | 3.3143 | 0.39047 | -0.82166 | -1.8572 | 0.96 |
| 9255 | *HMCN1* | 4.7491 | -0.25597 | -1.6105 | -2.381 | 0.96 |
| 9256 | *TXNDC16* | 3.6241 | -0.35899 | -0.38248 | -0.57513 | 0.96 |
| 9257 | *GTPBP2* | 2.8305 | 0.68209 | -0.63014 | -1.3542 | 0.96 |
| 9258 | *SLC16A3* | 3.1874 | 0.53862 | -0.84457 | -1.1338 | 0.96 |
| 9259 | *C17orf104* | 2.4015 | 0.4429 | 0.03663 | -1.5901 | 0.96 |
| 9260 | *CDY1* | 2.0628 | 0.97988 | -0.16266 | -1.5544 | 0.96 |
| 9261 | *TRMT2A* | 2.241 | 1.4002 | -0.76124 | -1.0501 | 0.96 |
| 9262 | *BTD* | 2.7196 | 0.42427 | -0.26466 | -1.1904 | 0.96 |
| 9263 | *KIAA0101* | 2.8833 | 0.68986 | -0.69411 | -0.70463 | 0.96 |
| 9264 | *PRDM12* | 4.0072 | 0.16975 | -1.2985 | -1.6033 | 0.96 |
| 9265 | *NOS3* | 3.9994 | 0.26037 | -1.3814 | -1.4636 | 0.96 |
| 9266 | *ATP10B* | 1.6737 | 1.2651 | -0.06044 | -1.7571 | 0.96 |
| 9267 | *ZNF639* | 2.0545 | 0.49193 | 0.33119 | -1.151 | 0.96 |
| 9268 | *CUX2* | 2.0644 | 0.69779 | 0.1148 | -1.6339 | 0.96 |
| 9269 | *TM2D3* | 2.84 | 1.0813 | -1.0446 | -1.9179 | 0.96 |
| 9270 | *WDFY2* | 3.0273 | -0.018 | -0.13273 | -1.4856 | 0.96 |
| 9271 | *CPSF6* | 3.0341 | -0.01264 | -0.14517 | -1.1453 | 0.96 |
| 9272 | *DNAAF2* | 3.6141 | -0.21938 | -0.51929 | -0.97776 | 0.96 |
| 9273 | *VPREB1* | 1.7377 | 1.1056 | 0.03207 | -0.04323 | 0.96 |
| 9274 | *CHIT1* | 1.3122 | 1.092 | 0.47102 | -0.39433 | 0.96 |
| 9275 | *IL27RA* | 1.1476 | 0.96542 | 0.76167 | -1.1676 | 0.96 |
| 9276 | *CBFA2T3* | 2.3895 | 0.47483 | 0.01 | -1.7495 | 0.96 |
| 9277 | *LAMB3* | 2.5815 | 0.45714 | -0.16543 | -0.24551 | 0.96 |
| 9278 | *SLC22A18AS* | 3.6098 | -0.34364 | -0.3931 | -1.0621 | 0.96 |
| 9279 | *PDDC1* | 1.7874 | 0.84999 | 0.2352 | -0.03676 | 0.96 |
| 9280 | *C18orf32* | 2.3719 | 0.36198 | 0.13733 | -0.20663 | 0.96 |
| 9281 | *CSDC2* | 2.0309 | 1.0682 | -0.22813 | -0.82398 | 0.96 |
| 9282 | *CRB3* | 3.7945 | 0.53127 | -1.4555 | -1.5952 | 0.96 |
| 9283 | *ENPP6* | 3.6968 | -0.25206 | -0.57484 | -1.2935 | 0.96 |
| 9284 | *CEACAM1* | 2.4943 | 0.21771 | 0.1572 | -1.0948 | 0.96 |
| 9285 | *TRADD* | 3.6599 | 0.94688 | -1.7392 | -1.8184 | 0.96 |

| | | | | | | |
|---|---|---|---|---|---|---|
| 9286 | *SGK223* | 5.6991 | -0.88272 | -1.9502 | -2.8849 | 0.96 |
| 9287 | *KDM7A* | 3.0787 | -0.06913 | -0.14391 | -0.92046 | 0.96 |
| 9288 | *DMRTA1* | 2.9139 | 0.27812 | -0.32728 | -2.1321 | 0.95 |
| 9289 | *LRRC4C* | 3.5319 | -0.3128 | -0.35441 | -0.49953 | 0.95 |
| 9290 | *MRGPRF* | 3.0866 | 0.88791 | -1.1105 | -1.2935 | 0.95 |
| 9291 | *PIK3C2A* | 2.655 | 0.62398 | -0.41502 | -2.1419 | 0.95 |
| 9292 | *DLGAP3* | 1.8324 | 1.5233 | -0.49322 | -1.9622 | 0.95 |
| 9293 | *EFHC2* | 1.5408 | 0.67206 | 0.64951 | -0.19342 | 0.95 |
| 9294 | *CHGB* | 4.2167 | -0.51642 | -0.83808 | -2.8628 | 0.95 |
| 9295 | *CLDN3* | 3.5378 | 0.81333 | -1.4894 | -1.6281 | 0.95 |
| 9296 | *RNF128* | 2.8861 | 0.13021 | -0.15509 | -1.0458 | 0.95 |
| 9297 | *PDGFC* | 3.1856 | -0.11363 | -0.21267 | -0.29089 | 0.95 |
| 9298 | *BAHD1* | 1.8661 | 1.1693 | -0.17647 | -0.85236 | 0.95 |
| 9299 | *CCL26* | 2.4241 | 0.79277 | -0.35816 | -1.0315 | 0.95 |
| 9300 | *KEL* | 4.5804 | -0.59174 | -1.1305 | -2.8327 | 0.95 |
| 9301 | *HSPB9* | 4.4311 | 0.19946 | -1.7726 | -2.1572 | 0.95 |
| 9302 | *FGF14* | 4.3021 | -0.23957 | -1.2055 | -1.8554 | 0.95 |
| 9303 | *ZBTB8A* | 3.1418 | 1.6772 | -1.9621 | -2.5707 | 0.95 |
| 9304 | *RASD2* | 2.91 | 0.41436 | -0.46757 | -2.1108 | 0.95 |
| 9305 | *LRIG1* | 4.1883 | -0.51666 | -0.81517 | -0.84523 | 0.95 |
| 9306 | *SP2* | 4.6075 | -0.80256 | -0.94965 | -2.3751 | 0.95 |
| 9307 | *PTK6* | 1.3074 | 1.1777 | 0.36943 | -2.1669 | 0.95 |
| 9308 | *SP4* | 3.0703 | 0.99568 | -1.2118 | -2.0697 | 0.95 |
| 9309 | *EPHA4* | 2.7049 | 0.72608 | -0.57742 | -2.2528 | 0.95 |
| 9310 | *CNBD1* | 3.04 | 1.3207 | -1.5073 | -2.2951 | 0.95 |
| 9311 | *HIST2H2AA4* | 4.4314 | -0.5231 | -1.0564 | -1.8382 | 0.95 |
| 9312 | *HIST1H4G* | 3.3448 | 0.52919 | -1.0236 | -1.6313 | 0.95 |
| 9313 | *DYNLT3* | 3.2556 | -0.10364 | -0.30215 | -0.4492 | 0.95 |
| 9314 | *CMBL* | 2.0414 | 0.99411 | -0.18589 | -0.98026 | 0.95 |
| 9315 | *C21orf2* | 2.4683 | 0.21218 | 0.16903 | -0.25605 | 0.95 |
| 9316 | *C14orf178* | 4.8173 | 0.26323 | -2.2311 | -3.1921 | 0.95 |
| 9317 | *IGSF22* | 1.8366 | 0.94913 | 0.06301 | -1.6575 | 0.95 |
| 9318 | *SRPX* | 2.0686 | 0.53345 | 0.24615 | -0.40104 | 0.95 |
| 9319 | *ZSCAN5A* | 4.9026 | -0.64769 | -1.4076 | -2.0782 | 0.95 |
| 9320 | *ZNF57* | 2.3731 | 0.40051 | 0.07356 | -1.3113 | 0.95 |
| 9321 | *SOX1* | 2.297 | 0.47342 | 0.07632 | -1.4893 | 0.95 |
| 9322 | *PROSC* | 2.1766 | 0.63011 | 0.03998 | 0.011761 | 0.95 |
| 9323 | *SENP3* | 4.5589 | -0.50133 | -1.2109 | -2.4595 | 0.95 |
| 9324 | *C12orf10* | 1.6188 | 0.9699 | 0.25747 | -1.359 | 0.95 |
| 9325 | *SLC9A4* | 2.5869 | 1.0304 | -0.77114 | -1.6488 | 0.95 |
| 9326 | *PLD3* | 3.0505 | 0.7254 | -0.92987 | -0.9327 | 0.95 |
| 9327 | *BRINP1* | 4.4413 | -0.2715 | -1.3247 | -1.4998 | 0.95 |
| 9328 | *CD1E* | 5.054 | -1.0283 | -1.1812 | -1.3023 | 0.95 |
| 9329 | *CMTR2* | 1.9029 | 0.57888 | 0.36202 | -1.2834 | 0.95 |
| 9330 | *GLMP* | 2.2692 | 0.44355 | 0.13096 | -1.478 | 0.95 |
| 9331 | *TBC1D30* | 2.2726 | 0.55491 | 0.0157 | -0.72157 | 0.95 |

| | | | | | | |
|---|---|---|---|---|---|---|
| 9332 | *CEACAM3* | 1.9184 | 0.56676 | 0.35793 | -0.74604 | 0.95 |
| 9333 | *CCDC177* | 2.3437 | 1.084 | -0.58534 | -1.7782 | 0.95 |
| 9334 | *ULBP1* | 2.4394 | 0.82505 | -0.4221 | -1.4182 | 0.95 |
| 9335 | *CDKN2A* | 1.9514 | 0.67735 | 0.21332 | -0.78096 | 0.95 |
| 9336 | *DDX3Y* | 3.4573 | 0.039168 | -0.6548 | -2.3623 | 0.95 |
| 9337 | *C22orf24* | 3.3539 | -0.15561 | -0.35674 | -1.6125 | 0.95 |
| 9338 | *RASA2* | 1.9364 | 0.83652 | 0.06861 | -0.43964 | 0.95 |
| 9339 | *IL17B* | 1.4896 | 1.2595 | 0.09207 | -0.74576 | 0.95 |
| 9340 | *ZNF16* | 3.101 | -0.07692 | -0.18381 | -1.2049 | 0.95 |
| 9341 | *C10orf90* | 2.6174 | 0.92252 | -0.70132 | -0.84278 | 0.95 |
| 9342 | *MIEN1* | 3.0649 | 0.39779 | -0.62489 | -1.9308 | 0.95 |
| 9343 | *CEBPA* | 3.0397 | 0.26983 | -0.47206 | -1.4033 | 0.95 |
| 9344 | *SLC44A2* | 4.7917 | -0.38059 | -1.5744 | -1.7577 | 0.95 |
| 9345 | *ACTRT3* | 2.1076 | 0.48816 | 0.24082 | -0.68813 | 0.95 |
| 9346 | *IFT122* | 3.4517 | 0.73607 | -1.3523 | -2.2169 | 0.95 |
| 9347 | *HIST1H3H* | 2.9244 | 0.16197 | -0.25095 | -0.8397 | 0.95 |
| 9348 | *VWA1* | 4.2712 | 0.15938 | -1.5962 | -2.1605 | 0.94 |
| 9349 | *SLC41A3* | 2.4008 | 1.622 | -1.1888 | -1.3664 | 0.94 |
| 9350 | *TBX4* | 2.5692 | 0.55068 | -0.28588 | -1.4914 | 0.94 |
| 9351 | *EID2* | 3.9228 | 0.39016 | -1.479 | -2.1412 | 0.94 |
| 9352 | *SLC25A12* | 4.383 | -0.59183 | -0.95753 | -1.6419 | 0.94 |
| 9353 | *KLB* | 2.3737 | 0.39468 | 0.06461 | -1.2324 | 0.94 |
| 9354 | *MUTYH* | 3.9397 | -0.36786 | -0.73911 | -0.97369 | 0.94 |
| 9355 | *CDK11A* | 2.7265 | 0.15254 | -0.04703 | -0.48416 | 0.94 |
| 9356 | *RAB41* | 1.4741 | 1.1154 | 0.24234 | -0.55675 | 0.94 |
| 9357 | *ARID4B* | 4.3895 | -0.26423 | -1.2936 | -1.5724 | 0.94 |
| 9358 | *DBNL* | 3.7336 | 0.47677 | -1.379 | -1.8102 | 0.94 |
| 9359 | *HBA1* | 3.1367 | 0.46651 | -0.77196 | -1.0181 | 0.94 |
| 9360 | *COMMD8* | 3.1786 | 0.038005 | -0.3859 | -0.9109 | 0.94 |
| 9361 | *TMEM143* | 2.0862 | 1.1336 | -0.38998 | -0.62047 | 0.94 |
| 9362 | *SEPT7* | 1.7953 | 0.52306 | 0.51143 | -0.60441 | 0.94 |
| 9363 | *GALNTL5* | 5.6911 | -1.2623 | -1.5992 | -2.0343 | 0.94 |
| 9364 | *R3HDM4* | 1.9226 | 0.49003 | 0.4166 | -0.72095 | 0.94 |
| 9365 | *KRTAP23-1* | 3.4964 | -0.05428 | -0.61297 | -1.5131 | 0.94 |
| 9366 | *NHLH1* | 2.0974 | 0.53893 | 0.19272 | -0.24741 | 0.94 |
| 9367 | *OR2L5* | 3.4078 | 0.13081 | -0.70998 | -2.7221 | 0.94 |
| 9368 | *TBC1D2* | 3.3866 | 0.76489 | -1.3237 | -1.5052 | 0.94 |
| 9369 | *DGKQ* | 3.8656 | 0.1573 | -1.1958 | -1.6836 | 0.94 |
| 9370 | *MRPL55* | 3.1901 | -0.03784 | -0.32546 | -2.1368 | 0.94 |
| 9371 | *KMO* | 1.5552 | 0.67432 | 0.59648 | -0.95802 | 0.94 |
| 9372 | *MORC4* | 1.5739 | 0.83166 | 0.41996 | -0.80248 | 0.94 |
| 9373 | *GRHL2* | 2.6201 | 0.95101 | -0.7458 | -1.095 | 0.94 |
| 9374 | *PUSL1* | 3.3628 | 0.15963 | -0.69712 | -1.3627 | 0.94 |
| 9375 | *C5orf34* | 2.621 | 1.146 | -0.94171 | -1.691 | 0.94 |
| 9376 | *HYPK* | 2.5631 | 0.25289 | 0.00884 | -2.6217 | 0.94 |
| 9377 | *NXF2* | 1.4083 | 0.84496 | 0.57155 | -0.85059 | 0.94 |

| | | | | | | |
|---|---|---|---|---|---|---|
| 9378 | *TRAPPC10* | 2.3892 | 0.59437 | -0.15953 | -0.18876 | 0.94 |
| 9379 | *USO1* | 2.0404 | 0.76144 | 0.0217 | -1.8341 | 0.94 |
| 9380 | *FAM209B* | 5.2574 | -1.0005 | -1.4337 | -1.8856 | 0.94 |
| 9381 | *RPS16* | 2.2126 | 0.89422 | -0.28374 | -0.94956 | 0.94 |
| 9382 | *OR13H1* | 3.2809 | 0.24924 | -0.70708 | -0.73708 | 0.94 |
| 9383 | *TAS2R38* | 2.01 | 0.87194 | -0.0597 | -0.15447 | 0.94 |
| 9384 | *CCL13* | 1.7462 | 1.585 | -0.50953 | -0.77159 | 0.94 |
| 9385 | *LLPH* | 3.5694 | 0.83015 | -1.578 | -1.6796 | 0.94 |
| 9386 | *CPED1* | 3.2958 | -0.07644 | -0.39817 | -1.3039 | 0.94 |
| 9387 | *NCOA7* | 3.6548 | 0.046816 | -0.88062 | -1.5458 | 0.94 |
| 9388 | *TBC1D23* | 4.0113 | 0.34839 | -1.539 | -1.83 | 0.94 |
| 9389 | *PIK3R1* | 2.8474 | 0.83693 | -0.86399 | -1.0453 | 0.94 |
| 9390 | *SDCBP* | 2.8911 | 0.44675 | -0.51799 | -1.6759 | 0.94 |
| 9391 | *KCNMB1* | 1.9318 | 0.79459 | 0.09276 | -0.44843 | 0.94 |
| 9392 | *HBG1* | 1.7886 | 0.5743 | 0.45582 | -0.54252 | 0.94 |
| 9393 | *NFYC* | 1.8745 | 1.5695 | -0.62644 | -1.9404 | 0.94 |
| 9394 | *CAMK2D* | 2.9476 | 1.0379 | -1.1685 | -1.3542 | 0.94 |
| 9395 | *EBLN2* | 2.438 | 0.81815 | -0.43966 | -1.4302 | 0.94 |
| 9396 | *RSPH3* | 2.3738 | 0.65244 | -0.20988 | -2.4633 | 0.94 |
| 9397 | *PYCR1* | 2.9595 | 0.20565 | -0.34887 | -0.56098 | 0.94 |
| 9398 | *SYT2* | 2.3481 | 0.94455 | -0.47642 | -0.83278 | 0.94 |
| 9399 | *C12orf54* | 4.0805 | -0.22469 | -1.0399 | -1.2167 | 0.94 |
| 9400 | *VPS18* | 2.5281 | 1.0166 | -0.72881 | -1.5787 | 0.94 |
| 9401 | *MIIP* | 2.1451 | 0.72634 | -0.05599 | -1.1592 | 0.94 |
| 9402 | *HAP1* | 2.2415 | 0.36658 | 0.20721 | 0.011291 | 0.94 |
| 9403 | *KIAA1804* | 2.6638 | 0.63413 | -0.48272 | -3.4957 | 0.94 |
| 9404 | *SMPD3* | 3.8821 | 0.41678 | -1.4841 | -2.0147 | 0.94 |
| 9405 | *ARHGDIA* | 3.1577 | 0.39153 | -0.73465 | -2.8398 | 0.94 |
| 9406 | *FCGR3A* | 2.8159 | 0.27065 | -0.2735 | -1.6802 | 0.94 |
| 9407 | *SPCS2* | 3.386 | 0.14048 | -0.71363 | -1.0093 | 0.94 |
| 9408 | *PLSCR2* | 3.3652 | 1.1184 | -1.6709 | -1.8751 | 0.94 |
| 9409 | *TMEM52* | 1.2379 | 0.83583 | 0.73815 | 0.5381 | 0.94 |
| 9410 | *PARK2* | 3.4318 | -0.13189 | -0.48829 | -1.3818 | 0.94 |
| 9411 | *ZNF557* | 5.0886 | -0.91269 | -1.3643 | -1.6065 | 0.94 |
| 9412 | *EMP1* | 2.9648 | 0.58987 | -0.74389 | -2.1187 | 0.94 |
| 9413 | *PLAC9* | 1.9743 | 0.92295 | -0.08675 | -1.2719 | 0.94 |
| 9414 | *PCDHA10* | 2.5088 | 0.84114 | -0.54219 | -1.4364 | 0.94 |
| 9415 | *COX6B2* | 3.424 | -0.01745 | -0.59981 | -1.6567 | 0.94 |
| 9416 | *NEK1* | 2.946 | 0.000336 | -0.13961 | -1.2603 | 0.94 |
| 9417 | *TAS2R31* | 2.1367 | 1.1372 | -0.46724 | -0.82172 | 0.94 |
| 9418 | *CIB3* | 2.9718 | 0.7152 | -0.88062 | -1.3279 | 0.94 |
| 9419 | *OR6P1* | 4.3523 | 0.10613 | -1.6524 | -1.9727 | 0.94 |
| 9420 | *ZNF384* | 1.0873 | 1.0385 | 0.68013 | -1.5183 | 0.94 |
| 9421 | *SALL2* | 3.6538 | -0.4046 | -0.4433 | -1.1096 | 0.94 |
| 9422 | *KRTAP10-6* | 1.4912 | 0.96328 | 0.3501 | -1.3633 | 0.93 |
| 9423 | *BLZF1* | 3.0731 | 0.64348 | -0.91234 | -1.4168 | 0.93 |

| | | | | | | |
|---|---|---|---|---|---|---|
| 9424 | *NRBP2* | 3.459 | -0.02912 | -0.62672 | -0.82921 | 0.93 |
| 9425 | *MS4A13* | 3.6239 | 0.25077 | -1.0723 | -1.2651 | 0.93 |
| 9426 | *PRDM8* | 1.5954 | 1.4433 | -0.23638 | -1.1317 | 0.93 |
| 9427 | *TSPYL2* | 2.3061 | 0.67237 | -0.17629 | -0.9612 | 0.93 |
| 9428 | *GIP* | 2.5726 | 0.44348 | -0.2146 | -0.85931 | 0.93 |
| 9429 | *C4orf47* | 1.4317 | 0.81801 | 0.55092 | 0.12097 | 0.93 |
| 9430 | *MAVS* | 1.8747 | 1.5241 | -0.5986 | -1.438 | 0.93 |
| 9431 | *ZNF474* | 3.0447 | 0.50855 | -0.75354 | -3.3966 | 0.93 |
| 9432 | *CHMP1B* | 3.2781 | 0.56038 | -1.0388 | -2.5743 | 0.93 |
| 9433 | *OVCH2* | 4.0698 | -0.40961 | -0.86089 | -1.802 | 0.93 |
| 9434 | *EVPLL* | 1.1244 | 0.85331 | 0.82137 | 0.09768 | 0.93 |
| 9435 | *FOSL1* | 2.6438 | 0.12044 | 0.03435 | -1.7134 | 0.93 |
| 9436 | *LARS2* | 4.5952 | -0.02993 | -1.767 | -1.9819 | 0.93 |
| 9437 | *SNTG2* | 3.5292 | 0.54452 | -1.2755 | -1.3358 | 0.93 |
| 9438 | *L3MBTL4* | 3.4007 | -0.01938 | -0.58353 | -0.68133 | 0.93 |
| 9439 | *SLC19A1* | 2.5411 | 0.19604 | 0.0606 | -0.05573 | 0.93 |
| 9440 | *ATG16L2* | 3.1649 | 0.62059 | -0.98803 | -1.6085 | 0.93 |
| 9441 | *FOXP3* | 3.0095 | 0.37301 | -0.58554 | -0.61169 | 0.93 |
| 9442 | *HCFC2* | 4.0533 | 0.32994 | -1.5865 | -1.7068 | 0.93 |
| 9443 | *SOBP* | 1.9913 | 1.4652 | -0.66008 | -1.666 | 0.93 |
| 9444 | *SMPX* | 3.0412 | 0.6846 | -0.92948 | -1.5167 | 0.93 |
| 9445 | *SIX6* | 2.5174 | 0.65788 | -0.37913 | -0.61483 | 0.93 |
| 9446 | *TNFRSF4* | 3.7578 | 0.1059 | -1.0691 | -1.3479 | 0.93 |
| 9447 | *TCF3* | 2.1748 | 0.66948 | -0.05 | -1.536 | 0.93 |
| 9448 | *TRIML2* | 1.8651 | 0.56815 | 0.36017 | 0.28429 | 0.93 |
| 9449 | *CPSF4L* | 2.0108 | 0.44573 | 0.3359 | -2.0545 | 0.93 |
| 9450 | *ST3GAL4* | 1.3246 | 1.0754 | 0.39221 | -0.80675 | 0.93 |
| 9451 | *NCAM1* | 3.4627 | 0.19462 | -0.86556 | -1.683 | 0.93 |
| 9452 | *SLC43A2* | 2.8326 | 0.22545 | -0.26652 | -1.203 | 0.93 |
| 9453 | *OR51F2* | 2.2412 | 0.27944 | 0.27053 | -1.3886 | 0.93 |
| 9454 | *TRIM6* | 3.6092 | 0.1385 | -0.95749 | -1.4994 | 0.93 |
| 9455 | *SLC7A13* | 3.1237 | 0.39289 | -0.72656 | -0.74943 | 0.93 |
| 9456 | *HARS2* | 2.3626 | 1.0547 | -0.62762 | -1.1034 | 0.93 |
| 9457 | *RBM26* | 2.821 | 0.55802 | -0.59045 | -0.72654 | 0.93 |
| 9458 | *CKB* | 4.1498 | -0.42038 | -0.9414 | -2.2856 | 0.93 |
| 9459 | *DEFB135* | 4.1227 | 0.46402 | -1.7991 | -5.5895 | 0.93 |
| 9460 | *ZNF304* | 1.3858 | 0.92518 | 0.47636 | -3.2419 | 0.93 |
| 9461 | *SULT1A3* | 2.2727 | 0.7429 | -0.22915 | -0.8756 | 0.93 |
| 9462 | *AMDHD2* | 3.4699 | 0.25706 | -0.94152 | -1.4452 | 0.93 |
| 9463 | *CDHR2* | 3.566 | 0.00147 | -0.78229 | -0.83081 | 0.93 |
| 9464 | *ZNF541* | 3.9516 | -0.57921 | -0.58771 | -1.879 | 0.93 |
| 9465 | *SARDH* | 3.9953 | 0.61452 | -1.8253 | -2.565 | 0.93 |
| 9466 | *POLR3G* | 4.7638 | -0.68606 | -1.2934 | -1.988 | 0.93 |
| 9467 | *NCR2* | 2.0102 | 1.0184 | -0.24578 | -2.8849 | 0.93 |
| 9468 | *SPG21* | 3.1702 | 0.46938 | -0.85697 | -2.278 | 0.93 |
| 9469 | *DDX19A* | 2.4808 | 1.0335 | -0.73185 | -1.1986 | 0.93 |

| | | | | | | |
|---|---|---|---|---|---|---|
| 9470 | *PPAPDC1A* | 2.2368 | 0.35673 | 0.18761 | -0.27811 | 0.93 |
| 9471 | *FNDC9* | 1.6845 | 0.97526 | 0.12081 | -1.304 | 0.93 |
| 9472 | *EML5* | 2.6725 | 0.35935 | -0.25163 | -1.5672 | 0.93 |
| 9473 | *COL15A1* | 2.6288 | 0.40183 | -0.2505 | -0.54142 | 0.93 |
| 9474 | *VWA5B1* | 4.2138 | -0.48659 | -0.94759 | -2.7384 | 0.93 |
| 9475 | *SLC35A2* | 4.4 | -0.13539 | -1.4856 | -2.2154 | 0.93 |
| 9476 | *SHROOM1* | 1.4619 | 0.73309 | 0.58354 | -1.7312 | 0.93 |
| 9477 | *ARG2* | 2.7667 | 0.23572 | -0.224 | -0.40319 | 0.93 |
| 9478 | *ECI1* | 2.8841 | 0.62385 | -0.72969 | -1.4584 | 0.93 |
| 9479 | *LACC1* | 2.3557 | 0.75415 | -0.33253 | -0.93892 | 0.93 |
| 9480 | *CCDC58* | 2.4194 | 2.4124 | -2.0545 | -2.2154 | 0.93 |
| 9481 | *LOC113230* | 2.4322 | 1.2853 | -0.94119 | -1.8975 | 0.93 |
| 9482 | *VWC2L* | 2.9426 | 0.34843 | -0.51585 | -1.6934 | 0.93 |
| 9483 | *BARHL2* | 2.6242 | 0.26086 | -0.11023 | -2.3811 | 0.92 |
| 9484 | *CYB5R1* | 3.6774 | -0.2293 | -0.67342 | -1.5099 | 0.92 |
| 9485 | *ADM2* | 2.4988 | 0.23694 | 0.03866 | -0.4311 | 0.92 |
| 9486 | *TMEM151A* | 3.9742 | -0.10473 | -1.0951 | -2.6889 | 0.92 |
| 9487 | *MPP6* | 3.265 | -0.18269 | -0.30801 | -1.1674 | 0.92 |
| 9488 | *PCGF5* | 1.9054 | 0.81614 | 0.05273 | -0.30883 | 0.92 |
| 9489 | *MC5R* | 2.6315 | 0.10737 | 0.03508 | -1.0799 | 0.92 |
| 9490 | *BPIFB2* | 1.7472 | 0.99021 | 0.03515 | -1.2421 | 0.92 |
| 9491 | *HEBP2* | 2.6945 | 1.1896 | -1.1138 | -3.0595 | 0.92 |
| 9492 | *WDR43* | 4.4837 | -0.59507 | -1.1184 | -1.2295 | 0.92 |
| 9493 | *DOC2B* | 4.1417 | -0.4449 | -0.92828 | -1.9802 | 0.92 |
| 9494 | *BCAP31* | 4.3016 | -0.04002 | -1.4936 | -1.5979 | 0.92 |
| 9495 | *BLOC1S3* | 4.2718 | -0.12489 | -1.379 | -2.9234 | 0.92 |
| 9496 | *KCNH5* | 1.6393 | 0.8389 | 0.28866 | 0.037225 | 0.92 |
| 9497 | *TMEM179B* | 2.4139 | 0.87217 | -0.51928 | -1.3019 | 0.92 |
| 9498 | *C3orf20* | 2.1129 | 1.2035 | -0.55011 | -1.9763 | 0.92 |
| 9499 | *PROKR1* | 1.8085 | 0.64943 | 0.30814 | 0.18128 | 0.92 |
| 9500 | *MT1HL1* | 3.1737 | 0.63025 | -1.0385 | -1.9885 | 0.92 |
| 9501 | *HTR3B* | 1.4489 | 0.73632 | 0.57896 | -1.9372 | 0.92 |
| 9502 | *FAIM2* | 2.0716 | 0.6696 | 0.02237 | -1.4028 | 0.92 |
| 9503 | *GATA3* | 2.8832 | -0.02494 | -0.09541 | -0.49999 | 0.92 |
| 9504 | *SLC5A6* | 4.4423 | -0.60138 | -1.0785 | -1.2628 | 0.92 |
| 9505 | *LITAF* | 2.8154 | 0.12969 | -0.18282 | -1.5312 | 0.92 |
| 9506 | *TOB1* | 1.8 | 1.7174 | -0.75603 | -1.3162 | 0.92 |
| 9507 | *OSTF1* | 1.0972 | 1.0013 | 0.6624 | -0.42777 | 0.92 |
| 9508 | *H3F3B* | 2.3259 | 1.5568 | -1.122 | -2.6593 | 0.92 |
| 9509 | *TMEM57* | 1.4994 | 1.1142 | 0.14641 | -1.8157 | 0.92 |
| 9510 | *ZFP91* | 1.3777 | 0.89978 | 0.48216 | -1.4639 | 0.92 |
| 9511 | *CSF1R* | 2.3794 | 1.7006 | -1.3205 | -1.5671 | 0.92 |
| 9512 | *CAPS2* | 3.1442 | -0.06466 | -0.32015 | -0.95044 | 0.92 |
| 9513 | *BZW1* | 2.7867 | 0.090471 | -0.11811 | -1.8435 | 0.92 |
| 9514 | *ZC2HC1C* | 2.7501 | 0.32836 | -0.31968 | -0.9745 | 0.92 |
| 9515 | *RHOV* | 2.6635 | 1.0716 | -0.97648 | -1.2884 | 0.92 |

| | | | | | | |
|---|---|---|---|---|---|---|
| 9516 | *COL28A1* | 2.8282 | 0.68455 | -0.75414 | -0.81496 | 0.92 |
| 9517 | *ABCC12* | 1.9301 | 1.3644 | -0.53613 | -0.56145 | 0.92 |
| 9518 | *ARHGAP28* | 3.0147 | 0.34019 | -0.59847 | -0.78221 | 0.92 |
| 9519 | *CXXC4* | 1.6417 | 0.80145 | 0.313 | -1.6243 | 0.92 |
| 9520 | *SH3D21* | 3.3083 | 0.54097 | -1.0942 | -2.278 | 0.92 |
| 9521 | *TIAM2* | 4.133 | -0.13094 | -1.2471 | -2.1896 | 0.92 |
| 9522 | *ESCO2* | 1.9398 | 0.55092 | 0.26289 | -0.12795 | 0.92 |
| 9523 | *SLC12A8* | 3.7222 | -0.31414 | -0.6548 | -1.658 | 0.92 |
| 9524 | *TSSK3* | 2.2837 | 0.31015 | 0.15931 | -0.0211 | 0.92 |
| 9525 | *ABLIM2* | 1.101 | 0.91654 | 0.73478 | -1.5073 | 0.92 |
| 9526 | *GALNT6* | 2.8636 | 0.14215 | -0.25415 | -0.51363 | 0.92 |
| 9527 | *HYKK* | 1.3067 | 1.2013 | 0.24312 | -0.30933 | 0.92 |
| 9528 | *CDKL4* | 2.177 | 0.38209 | 0.19045 | -1.5039 | 0.92 |
| 9529 | *HEMK1* | 2.1984 | 0.61278 | -0.06264 | -0.49138 | 0.92 |
| 9530 | *TRIM41* | 2.3773 | 0.27695 | 0.09356 | -0.52255 | 0.92 |
| 9531 | *SEC61B* | 2.5975 | 1.2007 | -1.0506 | -1.3689 | 0.92 |
| 9532 | *M1AP* | 3.8062 | 0.46677 | -1.5271 | -1.7766 | 0.92 |
| 9533 | *CHST3* | 2.5891 | 0.46379 | -0.30715 | -1.0704 | 0.92 |
| 9534 | *NHLRC1* | 2.926 | 0.8055 | -0.9858 | -1.6371 | 0.92 |
| 9535 | *SLC22A10* | 1.1387 | 0.87806 | 0.72892 | -1.0863 | 0.92 |
| 9536 | *C15orf61* | 2.3055 | 1.2767 | -0.83676 | -1.26 | 0.92 |
| 9537 | *ZBTB37* | 2.738 | 1.3069 | -1.3 | -1.5153 | 0.91 |
| 9538 | *E2F3* | 3.5402 | -0.1095 | -0.68727 | -1.578 | 0.91 |
| 9539 | *PDK4* | 1.3388 | 0.96396 | 0.44041 | -1.1414 | 0.91 |
| 9540 | *KRTAP20-2* | 4.0101 | 0.088405 | -1.3556 | -1.6731 | 0.91 |
| 9541 | *ALPK3* | 1.6861 | 1.3632 | -0.30653 | -0.36236 | 0.91 |
| 9542 | *TFB2M* | 2.3353 | 0.92907 | -0.52197 | -1.6571 | 0.91 |
| 9543 | *LDLRAD4* | 1.9211 | 0.51292 | 0.30732 | -0.69754 | 0.91 |
| 9544 | *EPHX3* | 2.8746 | 0.20971 | -0.34304 | -0.67458 | 0.91 |
| 9545 | *SNAP25* | 1.457 | 0.84267 | 0.44136 | -0.15323 | 0.91 |
| 9546 | *ANO4* | 2.0063 | 0.68585 | 0.04818 | -0.38428 | 0.91 |
| 9547 | *KAZN* | 3.47 | 0.033716 | -0.76389 | -1.5373 | 0.91 |
| 9548 | *PNRC1* | 2.6212 | 0.18656 | -0.06824 | -2.3449 | 0.91 |
| 9549 | *GABRR2* | 3.9966 | 0.39547 | -1.6526 | -2.0301 | 0.91 |
| 9550 | *GLDC* | 2.5725 | 0.24598 | -0.07903 | -0.16277 | 0.91 |
| 9551 | *SP110* | 5.1005 | -0.94219 | -1.4197 | -1.8775 | 0.91 |
| 9552 | *DNAL1* | 3.4663 | -0.06938 | -0.65839 | -1.4557 | 0.91 |
| 9553 | *HTATSF1* | 4.1741 | -0.1386 | -1.2975 | -1.2983 | 0.91 |
| 9554 | *TOMM20* | 3.1527 | -0.03366 | -0.38195 | -1.2027 | 0.91 |
| 9555 | *COG7* | 4.2669 | -0.5038 | -1.0261 | -1.6157 | 0.91 |
| 9556 | *KIAA1147* | 3.249 | 1.2905 | -1.8038 | -1.9075 | 0.91 |
| 9557 | *KLK12* | 3.5859 | 0.025101 | -0.8756 | -2.3338 | 0.91 |
| 9558 | *SAP30L* | 3.7334 | -0.28508 | -0.71305 | -0.76866 | 0.91 |
| 9559 | *ERG* | 1.8317 | 0.62448 | 0.27907 | -0.54354 | 0.91 |
| 9560 | *UBR3* | 2.3346 | 0.87263 | -0.47246 | -1.061 | 0.91 |
| 9561 | *EIF3B* | 3.3727 | 0.23298 | -0.87129 | -1.1267 | 0.91 |

| | | | | | | |
|---|---|---|---|---|---|---|
| 9562 | *TNFSF15* | 1.6353 | 1.4303 | -0.33191 | -0.62306 | 0.91 |
| 9563 | *SLMAP* | 3.1225 | 0.89541 | -1.2845 | -1.5817 | 0.91 |
| 9564 | *UTY* | 2.1672 | 0.47158 | 0.09435 | -0.14878 | 0.91 |
| 9565 | *POU6F1* | 4.1259 | 0.33313 | -1.7266 | -2.5005 | 0.91 |
| 9566 | *MC2R* | 1.7503 | 0.84114 | 0.14058 | -1.7496 | 0.91 |
| 9567 | *LOC100129924* | 3.3086 | 0.49297 | -1.0705 | -2.2117 | 0.91 |
| 9568 | *PLA2G6* | 2.7606 | 0.96402 | -0.99448 | -1.2064 | 0.91 |
| 9569 | *C5orf28* | 3.5551 | 0.13475 | -0.96031 | -3.8624 | 0.91 |
| 9570 | *NADSYN1* | 4.5737 | -0.17566 | -1.6689 | -1.8098 | 0.91 |
| 9571 | *ADRA1B* | 2.5884 | 0.94 | -0.79932 | -1.4683 | 0.91 |
| 9572 | *GCA* | 3.0505 | 0.8264 | -1.1487 | -2.428 | 0.91 |
| 9573 | *CLPSL2* | 1.6121 | 0.79624 | 0.31975 | -0.42859 | 0.91 |
| 9574 | *PRIMA1* | 2.2834 | 0.76588 | -0.32123 | -0.48119 | 0.91 |
| 9575 | *DLC1* | 1.1956 | 1.1645 | 0.36744 | -0.74618 | 0.91 |
| 9576 | *TBX15* | 2.0155 | 0.43185 | 0.28018 | -1.7602 | 0.91 |
| 9577 | *YKT6* | 2.4374 | 0.83128 | -0.54126 | -1.2806 | 0.91 |
| 9578 | *ELAC2* | 2.9405 | 0.84196 | -1.0551 | -1.3532 | 0.91 |
| 9579 | *IGFBP2* | 3.5383 | -0.08187 | -0.72907 | -1.3942 | 0.91 |
| 9580 | *NHLRC3* | 1.821 | 0.67689 | 0.22922 | -2.4015 | 0.91 |
| 9581 | *FOLR2* | 2.2408 | 0.74076 | -0.25493 | -1.2973 | 0.91 |
| 9582 | *RFNG* | 2.8939 | 0.18854 | -0.35606 | -1.5068 | 0.91 |
| 9583 | *IL20RB* | 3.5747 | -0.34738 | -0.50138 | -1.1524 | 0.91 |
| 9584 | *FCN2* | 3.2784 | 0.30856 | -0.86147 | -1.0104 | 0.91 |
| 9585 | *SLC23A1* | 3.3884 | -0.23777 | -0.42524 | -1.2163 | 0.91 |
| 9586 | *NUP155* | 3.0717 | 0.19791 | -0.54422 | -1.1634 | 0.91 |
| 9587 | *OR4C3* | 2.0548 | 1.4528 | -0.78229 | -1.602 | 0.91 |
| 9588 | *ACVR1C* | 2.9642 | 0.30269 | -0.54219 | -1.877 | 0.91 |
| 9589 | *TSEN2* | 3.4481 | -0.20557 | -0.51817 | -1.2253 | 0.91 |
| 9590 | *RPS10-NUDT3* | 3.5517 | 0.81691 | -1.6447 | -2.4557 | 0.91 |
| 9591 | *FANCI* | 1.9946 | 1.3275 | -0.59823 | -0.71834 | 0.91 |
| 9592 | *ACAD11* | 2.9239 | 1.2526 | -1.4527 | -1.5956 | 0.91 |
| 9593 | *SLC22A15* | 1.5079 | 1.3315 | -0.11567 | -1.4534 | 0.91 |
| 9594 | *ZCCHC14* | 3.1232 | 0.45733 | -0.85697 | -1.0254 | 0.91 |
| 9595 | *NSUN3* | 2.3656 | 0.47116 | -0.1138 | -0.7391 | 0.91 |
| 9596 | *CGRRF1* | 2.5767 | 0.48864 | -0.34255 | -2.254 | 0.91 |
| 9597 | *ALDH4A1* | 1.7867 | 0.47172 | 0.46356 | -0.59762 | 0.91 |
| 9598 | *UNKL* | 2.3262 | 0.87414 | -0.47911 | -2.005 | 0.91 |
| 9599 | *JAKMIP1* | 3.6855 | -0.15504 | -0.81007 | -0.84542 | 0.91 |
| 9600 | *RAB3A* | 2.4736 | 0.15938 | 0.08699 | -2.2381 | 0.91 |
| 9601 | *ACKR2* | 1.1601 | 0.79894 | 0.76079 | -0.52896 | 0.91 |
| 9602 | *SLC39A6* | 3.0702 | 0.93679 | -1.2872 | -1.8591 | 0.91 |
| 9603 | *PWWP2B* | 2.1063 | 0.36161 | 0.25156 | -1.1555 | 0.91 |
| 9604 | *AVP* | 4.2803 | -0.19525 | -1.3666 | -1.9251 | 0.91 |
| 9605 | *RAD54L* | 5.1771 | -1.1385 | -1.3209 | -1.8219 | 0.91 |
| 9606 | *MRPS24* | 2.2514 | 0.90315 | -0.43808 | -1.7783 | 0.91 |
| 9607 | *BEGAIN* | 1.9764 | 1.8975 | -1.1583 | -1.4545 | 0.91 |

| | | | | | | |
|---|---|---|---|---|---|---|
| 9608 | *CCNB2* | 1.6471 | 1.4649 | -0.39804 | -2.0277 | 0.90 |
| 9609 | *KIAA0895L* | 3.9357 | 0.13885 | -1.3619 | -2.3204 | 0.90 |
| 9610 | *ADAM18* | 4.782 | -0.95154 | -1.1202 | -1.6216 | 0.90 |
| 9611 | *FBXL4* | 4.0098 | -0.36987 | -0.92975 | -1.2411 | 0.90 |
| 9612 | *DOCK7* | 2.9729 | -0.08163 | -0.18116 | -1.714 | 0.90 |
| 9613 | *CCDC105* | 3.3513 | 0.36392 | -1.0055 | -1.2647 | 0.90 |
| 9614 | *SUCLG1* | 1.9473 | 0.72975 | 0.03151 | -1.2666 | 0.90 |
| 9615 | *STAG2* | 2.7656 | 1.3416 | -1.3987 | -1.8822 | 0.90 |
| 9616 | *VWA7* | 3.3248 | -0.10704 | -0.51002 | -1.4168 | 0.90 |
| 9617 | *VIT* | 1.9805 | 0.59361 | 0.13267 | -1.8575 | 0.90 |
| 9618 | *NR1I3* | 2.6513 | 0.67399 | -0.61892 | -1.5959 | 0.90 |
| 9619 | *IL3RA* | 1.367 | 1.3455 | -0.0062 | -0.28677 | 0.90 |
| 9620 | *TTC4* | 3.1644 | 0.5799 | -1.0388 | -1.5448 | 0.90 |
| 9621 | *CCDC113* | 1.4594 | 1.3327 | -0.08675 | -0.84457 | 0.90 |
| 9622 | *DEFB123* | 1.8754 | 0.51648 | 0.31323 | -0.30045 | 0.90 |
| 9623 | *REG1B* | 2.1666 | 1.5901 | -1.0516 | -1.078 | 0.90 |
| 9624 | *CCER2* | 2.6548 | 0.55698 | -0.50689 | -0.70442 | 0.90 |
| 9625 | *LYPD1* | 1.3697 | 0.80123 | 0.53367 | -0.40742 | 0.90 |
| 9626 | *HOXC10* | 3.1253 | 0.25015 | -0.6714 | -0.78229 | 0.90 |
| 9627 | *RSPH10B2* | 1.9241 | 0.65427 | 0.12494 | -1.0528 | 0.90 |
| 9628 | *ARHGEF26* | 1.5436 | 0.78732 | 0.37229 | -0.42708 | 0.90 |
| 9629 | *MAEL* | 1.3683 | 0.73915 | 0.59519 | -1.339 | 0.90 |
| 9630 | *BIN3* | 3.3333 | -0.0248 | -0.60604 | -1.5268 | 0.90 |
| 9631 | *PSMG4* | 4.0744 | -0.5967 | -0.77562 | -1.7577 | 0.90 |
| 9632 | *BAALC* | 3.1463 | -0.17129 | -0.27423 | -1.3364 | 0.90 |
| 9633 | *ZNF593* | 1.3233 | 0.7057 | 0.67131 | -0.1288 | 0.90 |
| 9634 | *TM9SF1* | 2.4594 | 1.2633 | -1.0231 | -2.382 | 0.90 |
| 9635 | *EFNA1* | 2.16 | 0.27747 | 0.26202 | 0.043375 | 0.90 |
| 9636 | *FMNL1* | 4.1688 | -0.21944 | -1.25 | -1.4214 | 0.90 |
| 9637 | *CPEB3* | 2.9103 | 0.3323 | -0.5442 | -1.4924 | 0.90 |
| 9638 | *SPATA22* | 1.8732 | 1.0774 | -0.25225 | -2.6072 | 0.90 |
| 9639 | *OR4E2* | 2.3769 | 0.19975 | 0.12166 | -1.1964 | 0.90 |
| 9640 | *MSH4* | 1.9458 | 0.38375 | 0.36873 | -0.75889 | 0.90 |
| 9641 | *CHRM3* | 1.2977 | 0.76001 | 0.64033 | -0.12598 | 0.90 |
| 9642 | *KRTAP24-1* | 1.4741 | 1.2485 | -0.02475 | -0.42354 | 0.90 |
| 9643 | *NPR3* | 3.41 | -0.30232 | -0.4102 | -0.7532 | 0.90 |
| 9644 | *ASUN* | 1.9264 | 0.70807 | 0.06281 | -0.43628 | 0.90 |
| 9645 | *HECTD2* | 3.1029 | 0.21507 | -0.62099 | -1.1754 | 0.90 |
| 9646 | *RARRES1* | 2.002 | 0.78671 | -0.0918 | -0.87019 | 0.90 |
| 9647 | *KLF2* | 2.483 | 0.50376 | -0.28994 | -1.7177 | 0.90 |
| 9648 | *LSM10* | 5.0573 | -0.84291 | -1.5176 | -2.907 | 0.90 |
| 9649 | *ARGLU1* | 1.7518 | 0.56676 | 0.37761 | -1.4597 | 0.90 |
| 9650 | *GYLTL1B* | 3.2046 | 0.14183 | -0.65109 | -0.79225 | 0.90 |
| 9651 | *OR2G2* | 2.7622 | 0.085857 | -0.15357 | -0.15385 | 0.90 |
| 9652 | *C15orf53* | 2.5865 | 0.40216 | -0.29484 | -2.641 | 0.90 |
| 9653 | *SCAF1* | 3.5455 | 0.83686 | -1.6891 | -2.5016 | 0.90 |

| | | | | | | |
|---|---|---|---|---|---|---|
| 9654 | *TIMM17A* | 2.0461 | 0.46952 | 0.17644 | 0.09986 | 0.90 |
| 9655 | *EMCN* | 2.2353 | 0.9419 | -0.48524 | -1.589 | 0.90 |
| 9656 | *ARHGAP24* | 2.2194 | 0.6295 | -0.15699 | -1.0426 | 0.90 |
| 9657 | *PRDM2* | 3.8366 | -0.34927 | -0.79573 | -1.5165 | 0.90 |
| 9658 | *RIC3* | 1.3613 | 0.96187 | 0.36623 | -0.19136 | 0.90 |
| 9659 | *H2AFJ* | 3.985 | -0.12176 | -1.1754 | -1.8646 | 0.90 |
| 9660 | *MIR205HG* | 3.3323 | -0.20105 | -0.44353 | -0.77159 | 0.90 |
| 9661 | *APOBEC3D* | 1.9983 | 0.36925 | 0.31999 | -1.2756 | 0.90 |
| 9662 | *ACVR1* | 2.0941 | 0.39435 | 0.19892 | -0.28712 | 0.90 |
| 9663 | *ZNF446* | 2.6474 | 0.068356 | -0.02864 | -0.74774 | 0.90 |
| 9664 | *KDM5C* | 2.9345 | 1.0342 | -1.2816 | -2.4815 | 0.90 |
| 9665 | *KIAA1919* | 2.3799 | 0.21235 | 0.09433 | -0.00092 | 0.90 |
| 9666 | *PRMT8* | 3.2325 | 0.61905 | -1.1651 | -1.3785 | 0.90 |
| 9667 | *RPL37A* | 3.0525 | 0.060454 | -0.42658 | -1.5195 | 0.90 |
| 9668 | *HGH1* | 3.8216 | 0.027969 | -1.1637 | -1.4204 | 0.90 |
| 9669 | *ZMYND8* | 4.4284 | -0.05504 | -1.6885 | -1.7766 | 0.89 |
| 9670 | *CXCL1* | 4.0697 | 0.05242 | -1.4375 | -2.3154 | 0.89 |
| 9671 | *TBX20* | 2.3788 | 0.48588 | -0.18006 | -1.9168 | 0.89 |
| 9672 | *ARHGEF39* | 4.7226 | 0.61036 | -2.6495 | -3.2745 | 0.89 |
| 9673 | *SPANXN5* | 1.9071 | 1.2073 | -0.43114 | -1.554 | 0.89 |
| 9674 | *TSC22D3* | 1.5916 | 0.8577 | 0.23385 | -2.0383 | 0.89 |
| 9675 | *USP4* | 1.7128 | 0.5995 | 0.37073 | -0.11745 | 0.89 |
| 9676 | *SPATA19* | 1.8514 | 0.68622 | 0.14537 | -1.277 | 0.89 |
| 9677 | *SLC6A15* | 1.7421 | 1.0848 | -0.14427 | -1.2628 | 0.89 |
| 9678 | *TRMT44* | 2.7297 | 0.57812 | -0.62531 | -0.81448 | 0.89 |
| 9679 | *ANKRD13C* | 3.3854 | -0.13849 | -0.56507 | -1.4757 | 0.89 |
| 9680 | *FAM216A* | 5.097 | -0.33156 | -2.0837 | -2.3876 | 0.89 |
| 9681 | *LYZ* | 3.0614 | 0.4506 | -0.83027 | -1.3851 | 0.89 |
| 9682 | *ACACB* | 3.1576 | 0.52808 | -1.0041 | -2.101 | 0.89 |
| 9683 | *AAMP* | 2.0526 | 0.73484 | -0.10647 | -1.0769 | 0.89 |
| 9684 | *GABPB1* | 3.8563 | -0.18934 | -0.98634 | -1.9866 | 0.89 |
| 9685 | *SEMA3F* | 4.807 | -0.51363 | -1.6137 | -2.0466 | 0.89 |
| 9686 | *ZNF414* | 2.5777 | 0.50912 | -0.40765 | -0.91632 | 0.89 |
| 9687 | *RBM42* | 4.6004 | -0.14513 | -1.7766 | -1.984 | 0.89 |
| 9688 | *ITPR3* | 1.6531 | 0.99657 | 0.02895 | -0.50828 | 0.89 |
| 9689 | *IFRD2* | 1.1841 | 0.91331 | 0.58115 | -0.97784 | 0.89 |
| 9690 | *TADA2A* | 1.3559 | 0.92367 | 0.39727 | -1.9175 | 0.89 |
| 9691 | *SVEP1* | 2.8294 | 0.013685 | -0.16644 | -0.50207 | 0.89 |
| 9692 | *SART1* | 3.2886 | -0.15787 | -0.45454 | -0.78177 | 0.89 |
| 9693 | *STARD9* | 4.0167 | -0.28822 | -1.0529 | -1.259 | 0.89 |
| 9694 | *FOXR1* | 2.3661 | 0.45259 | -0.14362 | -0.85564 | 0.89 |
| 9695 | *TNKS1BP1* | 4.6696 | -0.74392 | -1.2517 | -5.8457 | 0.89 |
| 9696 | *PIM3* | 2.912 | 0.29119 | -0.52944 | -2.092 | 0.89 |
| 9697 | *HIST1H2AD* | 4.2802 | 0.050441 | -1.6587 | -2.1347 | 0.89 |
| 9698 | *STAP2* | 2.571 | 0.84298 | -0.74308 | -1.8272 | 0.89 |
| 9699 | *SDC2* | 2.6917 | 0.45188 | -0.47304 | -1.6525 | 0.89 |

| | | | | | | |
|---|---|---|---|---|---|---|
| 9700 | *COMMD10* | 3.6069 | 0.39778 | -1.3358 | -2.5004 | 0.89 |
| 9701 | *CPXM2* | 2.6442 | 0.030467 | -0.00586 | -1.6827 | 0.89 |
| 9702 | *DDX60* | 1.0506 | 1.0069 | 0.61041 | -1.5494 | 0.89 |
| 9703 | *SMCR8* | 3.7247 | -0.3548 | -0.70202 | -1.7517 | 0.89 |
| 9704 | *CD99* | 2.0377 | 0.8262 | -0.19658 | -0.76337 | 0.89 |
| 9705 | *PDS5A* | 3.0893 | 0.62099 | -1.043 | -2.8928 | 0.89 |
| 9706 | *CNTRL* | 3.6166 | -0.34327 | -0.60692 | -1.5913 | 0.89 |
| 9707 | *ZNF25* | 4.0658 | -0.60845 | -0.79147 | -1.9065 | 0.89 |
| 9708 | *FAM170A* | 2.1972 | 0.32443 | 0.14408 | -1.2111 | 0.89 |
| 9709 | *RASSF2* | 2.5126 | 2.0209 | -1.8679 | -2.6003 | 0.89 |
| 9710 | *ZNF385A* | 1.9682 | 0.56191 | 0.13547 | -0.15718 | 0.89 |
| 9711 | *WISP3* | 1.3031 | 1.0361 | 0.3263 | -0.12728 | 0.89 |
| 9712 | *PRKRIP1* | 2.136 | 0.42233 | 0.10696 | -1.466 | 0.89 |
| 9713 | *ABCA12* | 3.1933 | 0.19686 | -0.72488 | -2.2766 | 0.89 |
| 9714 | *DOLK* | 3.3967 | -0.21348 | -0.51805 | -1.257 | 0.89 |
| 9715 | *STARD10* | 4.4022 | -0.16927 | -1.5684 | -1.677 | 0.89 |
| 9716 | *SNTN* | 3.5899 | 0.68912 | -1.6146 | -2.012 | 0.89 |
| 9717 | *ZASP* | 1.4659 | 1.0104 | 0.18807 | -0.63866 | 0.89 |
| 9718 | *SORBS3* | 2.4081 | 0.7712 | -0.51496 | -0.89387 | 0.89 |
| 9719 | *VEZF1* | 1.9293 | 1.4314 | -0.6968 | -1.7567 | 0.89 |
| 9720 | *PLAC4* | 1.9248 | 0.55632 | 0.18216 | -0.2302 | 0.89 |
| 9721 | *CCDC89* | 2.3525 | 0.22074 | 0.08999 | -1.5095 | 0.89 |
| 9722 | *GFAP* | 2.6825 | 0.71641 | -0.73604 | -3.9446 | 0.89 |
| 9723 | *FAXDC2* | 4.557 | -0.44353 | -1.4507 | -2.8402 | 0.89 |
| 9724 | *RACGAP1* | 2.055 | 1.0201 | -0.41239 | -1.1532 | 0.89 |
| 9725 | *ZNF215* | 3.0894 | 0.53844 | -0.96632 | -1.0528 | 0.89 |
| 9726 | *INS* | 1.5551 | 0.99547 | 0.11075 | -0.66604 | 0.89 |
| 9727 | *MYO18A* | 2.1165 | 1.1798 | -0.63554 | -2.0538 | 0.89 |
| 9728 | *PAFAH2* | 2.1796 | 1.7229 | -1.2427 | -1.8817 | 0.89 |
| 9729 | *PLA2G7* | 1.7246 | 0.60331 | 0.33119 | -2.5153 | 0.89 |
| 9730 | *OR2S2* | 1.4951 | 0.78316 | 0.38056 | -0.59263 | 0.89 |
| 9731 | *NSMAF* | 3.6145 | -0.21666 | -0.73911 | -1.1193 | 0.89 |
| 9732 | *PRKG2* | 1.7003 | 1.2223 | -0.26399 | -1.8631 | 0.89 |
| 9733 | *PHLPP1* | 2.1009 | 0.51115 | 0.04617 | -0.66587 | 0.89 |
| 9734 | *HSD17B4* | 3.0977 | 0.72497 | -1.165 | -1.9624 | 0.89 |
| 9735 | *FAM69C* | 3.0014 | -0.04759 | -0.29647 | -0.60321 | 0.89 |
| 9736 | *AHDC1* | 1.3257 | 0.69063 | 0.64095 | -1.4684 | 0.89 |
| 9737 | *KRBOX1* | 4.543 | 0.076234 | -1.9622 | -2.1334 | 0.89 |
| 9738 | *DLG5* | 2.4685 | 1.601 | -1.4132 | -2.2255 | 0.89 |
| 9739 | *EIF4G3* | 1.8164 | 0.74847 | 0.09132 | -0.19311 | 0.89 |
| 9740 | *NHSL2* | 3.7278 | -0.22492 | -0.84718 | -1.5383 | 0.89 |
| 9741 | *RPS10* | 2.2249 | 0.81638 | -0.3866 | -1.2371 | 0.88 |
| 9742 | *C9orf129* | 3.5135 | -0.25493 | -0.60423 | -1.1503 | 0.88 |
| 9743 | *PLA2G4B* | 2.5342 | 0.80718 | -0.68706 | -1.4335 | 0.88 |
| 9744 | *KISS1R* | 3.9989 | -0.55833 | -0.78651 | -1.1553 | 0.88 |
| 9745 | *MLIP* | 3.0122 | 1.0961 | -1.4553 | -1.5212 | 0.88 |

| | | | | | | |
|---|---|---|---|---|---|---|
| 9746 | *CTAGE15* | 1.6051 | 1.4904 | -0.44353 | -1.1002 | 0.88 |
| 9747 | *GSTM5* | 3.2255 | -0.17285 | -0.40109 | -0.86744 | 0.88 |
| 9748 | *OXLD1* | 4.9087 | -0.78959 | -1.4683 | -4.5653 | 0.88 |
| 9749 | *CSF2RA* | 2.7532 | 0.97997 | -1.0824 | -1.3198 | 0.88 |
| 9750 | *GRK5* | 2.5244 | 0.097241 | 0.02503 | -1.2085 | 0.88 |
| 9751 | *CHRM4* | 2.8355 | -0.03116 | -0.15831 | -0.27982 | 0.88 |
| 9752 | *HMHA1* | 4.7282 | -0.40692 | -1.6753 | -1.7322 | 0.88 |
| 9753 | *TBC1D3B* | 2.6881 | 0.47762 | -0.51994 | -1.6976 | 0.88 |
| 9754 | *SNIP1* | 2.7642 | 0.04324 | -0.16242 | -0.333 | 0.88 |
| 9755 | *OR10C1* | 3.0758 | -0.12659 | -0.3048 | -1.197 | 0.88 |
| 9756 | *GNG5* | 3.9235 | -0.27016 | -1.0092 | -1.7507 | 0.88 |
| 9757 | *GABRA3* | 4.1405 | -0.02008 | -1.4767 | -2.8316 | 0.88 |
| 9758 | *IQCG* | 1.4386 | 1.1146 | 0.09045 | -0.64278 | 0.88 |
| 9759 | *CDH13* | 3.5217 | 0.65513 | -1.5354 | -1.8219 | 0.88 |
| 9760 | *ACTL7A* | 2.2859 | 0.39559 | -0.04064 | -0.5138 | 0.88 |
| 9761 | *RTN4RL2* | 1.8826 | 0.84679 | -0.09043 | -1.043 | 0.88 |
| 9762 | *SLC10A4* | 3.6772 | -0.30237 | -0.73591 | -1.8804 | 0.88 |
| 9763 | *EVX2* | 2.5685 | 0.27344 | -0.20338 | -1.927 | 0.88 |
| 9764 | *MOB3C* | 2.4202 | 0.74607 | -0.52795 | -0.54155 | 0.88 |
| 9765 | *MAPKBP1* | 1.686 | 0.54868 | 0.40342 | -1.2349 | 0.88 |
| 9766 | *SSR3* | 2.7724 | 0.28482 | -0.41917 | -0.47304 | 0.88 |
| 9767 | *EIF4E2* | 1.9248 | 0.77026 | -0.05738 | -5.0141 | 0.88 |
| 9768 | *BOD1* | 1.3239 | 0.95622 | 0.35683 | -0.06044 | 0.88 |
| 9769 | *SH2D7* | 4.1111 | -0.68386 | -0.79065 | -1.5956 | 0.88 |
| 9770 | *ARL8A* | 4.5501 | -0.36792 | -1.5458 | -2.1404 | 0.88 |
| 9771 | *ATP5J2-PTCD1* | 2.7837 | 0.72665 | -0.87418 | -1.9423 | 0.88 |
| 9772 | *TMEM207* | 3.164 | 0.29116 | -0.81932 | -1.1357 | 0.88 |
| 9773 | *PAF1* | 4.1906 | -0.13627 | -1.4189 | -1.6597 | 0.88 |
| 9774 | *CMTM5* | 3.5066 | 0.063013 | -0.93453 | -1.3851 | 0.88 |
| 9775 | *PAIP2B* | 1.8258 | 0.68725 | 0.12097 | -0.41073 | 0.88 |
| 9776 | *RAD21L1* | 0.93194 | 0.85789 | 0.84366 | -0.34187 | 0.88 |
| 9777 | *TGFBR3L* | 2.2351 | 0.31766 | 0.08015 | -0.83784 | 0.88 |
| 9778 | *OR10G4* | 2.7185 | 0.50465 | -0.59028 | -0.94733 | 0.88 |
| 9779 | *HIST1H2AG* | 2.713 | 0.071886 | -0.15203 | -0.60368 | 0.88 |
| 9780 | *DNASE1L2* | 4.191 | -0.68193 | -0.87768 | -0.96379 | 0.88 |
| 9781 | *CT47A10* | 3.0878 | 0.035844 | -0.49322 | -1.4299 | 0.88 |
| 9782 | *RNF31* | 3.6379 | -0.04858 | -0.95933 | -1.1858 | 0.88 |
| 9783 | *FBXO34* | 1.7701 | 0.86996 | -0.01116 | -0.16415 | 0.88 |
| 9784 | *FAM129C* | 1.4509 | 1.2188 | -0.04189 | -0.39698 | 0.88 |
| 9785 | *TESK1* | 3.4275 | -0.17663 | -0.62307 | -0.69961 | 0.88 |
| 9786 | *ZNF507* | 2.8953 | 0.00844 | -0.27596 | -2.0602 | 0.88 |
| 9787 | *GORASP2* | 2.6035 | 0.47351 | -0.44927 | -0.55678 | 0.88 |
| 9788 | *TMEM249* | 4.2199 | -0.28605 | -1.307 | -2.5754 | 0.88 |
| 9789 | *C2orf76* | 1.4428 | 1.3878 | -0.20378 | -1.1263 | 0.88 |
| 9790 | *C17orf99* | 2.5425 | 0.52708 | -0.44353 | -0.45847 | 0.88 |
| 9791 | *INSIG1* | 5.2149 | -0.27719 | -2.3118 | -3.9349 | 0.88 |

| | | | | | | |
|---|---|---|---|---|---|---|
| 9792 | *SPATA21* | 3.6234 | 0.20209 | -1.2001 | -1.8341 | 0.88 |
| 9793 | *LYRM7* | 3.6227 | 0.46266 | -1.4601 | -1.4878 | 0.88 |
| 9794 | *ATP2C1* | 2.2182 | 0.77718 | -0.37139 | -1.5155 | 0.87 |
| 9795 | *BCR* | 5.3145 | 0.66019 | -3.3519 | -3.729 | 0.87 |
| 9796 | *HSD11B2* | 3.2931 | -0.29942 | -0.37103 | -0.63572 | 0.87 |
| 9797 | *FAM175A* | 3.0641 | 0.17706 | -0.61899 | -1.3663 | 0.87 |
| 9798 | *GPLD1* | 1.1548 | 0.91929 | 0.54787 | 0.47483 | 0.87 |
| 9799 | *NOL12* | 3.2849 | -0.2804 | -0.38349 | -1.5022 | 0.87 |
| 9800 | *NARF* | 1.3501 | 1.0537 | 0.21702 | -1.9722 | 0.87 |
| 9801 | *ENTPD1* | 1.5899 | 0.6071 | 0.42375 | -1.2437 | 0.87 |
| 9802 | *DOCK8* | 4.0073 | 0.010651 | -1.3976 | -2.0895 | 0.87 |
| 9803 | *CXorf21* | 2.3134 | 0.27655 | 0.02949 | -0.52712 | 0.87 |
| 9804 | *CXorf36* | 1.9014 | 0.54576 | 0.1719 | 0.006927 | 0.87 |
| 9805 | *ZNF225* | 2.4824 | 0.6576 | -0.52149 | -0.80946 | 0.87 |
| 9806 | *HIST1H2BE* | 2.0286 | 1.4331 | -0.84365 | -1.6158 | 0.87 |
| 9807 | *FOPNL* | 2.4363 | 0.4001 | -0.21964 | -1.1703 | 0.87 |
| 9808 | *NEXN* | 2.6489 | 0.033178 | -0.0658 | -0.26592 | 0.87 |
| 9809 | *GOSR2* | 1.2914 | 1.0718 | 0.25236 | -0.61356 | 0.87 |
| 9810 | *SDK1* | 1.7173 | 0.58659 | 0.31145 | 0.032177 | 0.87 |
| 9811 | *FN1* | 2.7382 | 0.16693 | -0.28986 | -1.1968 | 0.87 |
| 9812 | *TNFRSF8* | 1.6849 | 0.84771 | 0.08154 | -0.27609 | 0.87 |
| 9813 | *RPS6KL1* | 2.7247 | 0.052148 | -0.16282 | -1.2008 | 0.87 |
| 9814 | *MAFB* | 2.8824 | 0.32298 | -0.59244 | -0.62174 | 0.87 |
| 9815 | *RPL21* | 3.5281 | 0.15861 | -1.0741 | -1.6213 | 0.87 |
| 9816 | *ILVBL* | 2.4234 | 0.19737 | -0.00825 | -2.6675 | 0.87 |
| 9817 | *ITIH6* | 2.3695 | 0.84709 | -0.60445 | -1.6383 | 0.87 |
| 9818 | *FAM166A* | 1.3499 | 1.1074 | 0.1539 | -0.70558 | 0.87 |
| 9819 | *ART3* | 3.8323 | -0.02511 | -1.1964 | -1.2936 | 0.87 |
| 9820 | *ZNF76* | 4.9502 | -1.083 | -1.2571 | -1.5956 | 0.87 |
| 9821 | *CENPP* | 2.1388 | 0.91449 | -0.44353 | -2.2432 | 0.87 |
| 9822 | *HEATR5A* | 2.5031 | 0.42059 | -0.3146 | -1.6803 | 0.87 |
| 9823 | *MEI1* | 3.9288 | -0.40584 | -0.91396 | -1.1886 | 0.87 |
| 9824 | *PLEKHO1* | 3.8677 | -0.44445 | -0.81433 | -1.3479 | 0.87 |
| 9825 | *DPM1* | 3.2133 | -0.21484 | -0.39043 | -1.2388 | 0.87 |
| 9826 | *CDH1* | 2.5815 | 0.25825 | -0.23215 | -1.0841 | 0.87 |
| 9827 | *MKKS* | 2.2386 | 0.25294 | 0.1148 | -1.2616 | 0.87 |
| 9828 | *LRRN3* | 3.3167 | -0.01068 | -0.70175 | -3.6928 | 0.87 |
| 9829 | *CYCS* | 3.2375 | 0.74883 | -1.3829 | -1.5346 | 0.87 |
| 9830 | *HARBI1* | 1.2631 | 1.0805 | 0.25931 | 0.15924 | 0.87 |
| 9831 | *OCM* | 3.9387 | -0.0426 | -1.2935 | -2.2165 | 0.87 |
| 9832 | *NCMAP* | 2.284 | 0.18114 | 0.13714 | 0.098257 | 0.87 |
| 9833 | *PRKAA2* | 2.1685 | 1.2567 | -0.82326 | -1.5073 | 0.87 |
| 9834 | *GPR83* | 2.3667 | 0.44838 | -0.21317 | -0.5183 | 0.87 |
| 9835 | *PRODH* | 2.3573 | 0.66202 | -0.41835 | -1.5837 | 0.87 |
| 9836 | *SERPINA10* | 3.6313 | -0.37106 | -0.65949 | -0.84743 | 0.87 |
| 9837 | *SCNN1G* | 4.6133 | -0.96222 | -1.0508 | -1.2365 | 0.87 |

| | | | | | | |
|---|---|---|---|---|---|---|
| 9838 | *SLC38A5* | 1.3729 | 0.70642 | 0.52025 | -0.16695 | 0.87 |
| 9839 | *TMEM173* | 1.3663 | 1.1361 | 0.09713 | -0.043 | 0.87 |
| 9840 | *MFSD2A* | 2.8331 | 0.86742 | -1.1012 | -2.1628 | 0.87 |
| 9841 | *TMEM218* | 2.6483 | 1.3428 | -1.3918 | -1.7074 | 0.87 |
| 9842 | *CDC42SE1* | 3.7799 | -0.36035 | -0.82044 | -0.93885 | 0.87 |
| 9843 | *LRRC38* | 1.7635 | 0.64291 | 0.19223 | -1.4252 | 0.87 |
| 9844 | *RGS4* | 3.9737 | -0.3783 | -0.99768 | -1.3478 | 0.87 |
| 9845 | *KBTBD13* | 4.5697 | -0.79149 | -1.1807 | -2.5839 | 0.87 |
| 9846 | *SH2D1B* | 2.1423 | 0.71068 | -0.25619 | -0.98047 | 0.87 |
| 9847 | *SYT3* | 2.8831 | 0.29117 | -0.57824 | -4.3949 | 0.87 |
| 9848 | *PLLP* | 3.493 | -0.20434 | -0.69333 | -1.348 | 0.87 |
| 9849 | *CNIH4* | 2.7053 | 0.81803 | -0.92872 | -1.0765 | 0.86 |
| 9850 | *SMIM2* | 3.6726 | -0.03406 | -1.0443 | -1.1797 | 0.86 |
| 9851 | *KCNIP3* | 2.578 | 0.3267 | -0.31051 | -1.4764 | 0.86 |
| 9852 | *MSGN1* | 3.2892 | 0.095647 | -0.79108 | -0.92078 | 0.86 |
| 9853 | *ATXN7L1* | 2.9706 | 0.30598 | -0.68388 | -0.69604 | 0.86 |
| 9854 | *GSTP1* | 3.6354 | -0.30497 | -0.73835 | -1.4015 | 0.86 |
| 9855 | *TFEB* | 2.7062 | 0.42375 | -0.53856 | -1.6288 | 0.86 |
| 9856 | *DUOX1* | 2.5654 | 0.401 | -0.37592 | -1.5883 | 0.86 |
| 9857 | *FAM84B* | 3.7305 | -0.50221 | -0.63826 | -1.6208 | 0.86 |
| 9858 | *ZBTB11* | 3.2475 | 0.02563 | -0.68335 | -0.79202 | 0.86 |
| 9859 | *NOSTRIN* | 3.285 | -0.21979 | -0.47567 | -0.57045 | 0.86 |
| 9860 | *PAOX* | 2.8796 | 0.59868 | -0.8895 | -2.1566 | 0.86 |
| 9861 | *TAS2R4* | 1.9359 | 0.3515 | 0.30081 | -0.14778 | 0.86 |
| 9862 | *ABCA10* | 3.3605 | 1.0803 | -1.8529 | -2.1947 | 0.86 |
| 9863 | *CDNF* | 2.534 | 0.49741 | -0.44353 | -0.46763 | 0.86 |
| 9864 | *MYL6B* | 1.3709 | 0.67274 | 0.5439 | -0.69891 | 0.86 |
| 9865 | *SMARCA5* | 2.379 | 0.95694 | -0.74913 | -1.5767 | 0.86 |
| 9866 | *PODXL2* | 3.6666 | -0.47696 | -0.60307 | -1.1095 | 0.86 |
| 9867 | *CCR6* | 1.9755 | 0.37588 | 0.23508 | -1.12 | 0.86 |
| 9868 | *EPS8L3* | 2.2006 | 1.2407 | -0.85564 | -2.0383 | 0.86 |
| 9869 | *CTPS2* | 1.6998 | 0.57698 | 0.30863 | 0.008554 | 0.86 |
| 9870 | *SMC6* | 1.2851 | 0.75598 | 0.54429 | -1.175 | 0.86 |
| 9871 | *BLOC1S4* | 2.2466 | 0.24738 | 0.091 | -0.0466 | 0.86 |
| 9872 | *KIF6* | 1.0183 | 0.96903 | 0.59762 | -0.20621 | 0.86 |
| 9873 | *RBM46* | 2.2795 | 0.742 | -0.43702 | -1.4682 | 0.86 |
| 9874 | *ZNF687* | 2.9521 | 0.81498 | -1.1826 | -1.2797 | 0.86 |
| 9875 | *SLCO1C1* | 3.2497 | 1.4558 | -2.1217 | -3.1051 | 0.86 |
| 9876 | *HHLA1* | 4.4461 | -0.76569 | -1.0975 | -1.7266 | 0.86 |
| 9877 | *BCL2L15* | 3.3169 | -0.17556 | -0.55849 | -1.6116 | 0.86 |
| 9878 | *MRPS30* | 2.5366 | 0.26145 | -0.21532 | -1.2146 | 0.86 |
| 9879 | *PPM1L* | 4.0334 | -0.54786 | -0.90303 | -1.0367 | 0.86 |
| 9880 | *LIMS2* | 2.761 | 0.2274 | -0.40605 | -5.6424 | 0.86 |
| 9881 | *TMEM206* | 3.1295 | 0.47755 | -1.0248 | -1.204 | 0.86 |
| 9882 | *ZNF318* | 3.1201 | 0.76713 | -1.305 | -2.3065 | 0.86 |
| 9883 | *TRIO* | 2.4125 | 0.73762 | -0.56841 | -1.0925 | 0.86 |

| | | | | | | |
|---|---|---|---|---|---|---|
| 9884 | *MXRA5* | 3.2858 | -0.23176 | -0.47244 | -1.5552 | 0.86 |
| 9885 | *NXF3* | 1.9677 | 0.81679 | -0.20324 | -0.27892 | 0.86 |
| 9886 | *CEACAM7* | 4.6922 | -0.79319 | -1.3178 | -1.3465 | 0.86 |
| 9887 | *LTB* | 2.9157 | 1.3866 | -1.7215 | -2.7042 | 0.86 |
| 9888 | *EVA1B* | 1.1674 | 1.0209 | 0.39248 | 0.15417 | 0.86 |
| 9889 | *AURKC* | 2.2398 | 0.21026 | 0.13043 | -1.7378 | 0.86 |
| 9890 | *PLEKHG5* | 4.0072 | -0.67461 | -0.75213 | -1.1989 | 0.86 |
| 9891 | *USP40* | 1.7433 | 0.49398 | 0.34277 | -1.2411 | 0.86 |
| 9892 | *HNRNPCL1* | 4.0154 | 0.063498 | -1.4996 | -1.7782 | 0.86 |
| 9893 | *ADAM23* | 2.9232 | 0.25253 | -0.59662 | -1.3287 | 0.86 |
| 9894 | *LRRC8B* | 2.9005 | 0.10349 | -0.42524 | -1.7616 | 0.86 |
| 9895 | *PRSS54* | 1.4071 | 1.1621 | 0.00943 | -1.6112 | 0.86 |
| 9896 | *SLC31A1* | 3.8025 | -0.48833 | -0.73585 | -1.7507 | 0.86 |
| 9897 | *PIGP* | 2.2891 | 0.31701 | -0.02943 | -0.70443 | 0.86 |
| 9898 | *VPS13B* | 2.485 | 0.55162 | -0.46064 | -1.7935 | 0.86 |
| 9899 | *OR6C70* | 2.8888 | -0.04416 | -0.26898 | -0.28308 | 0.86 |
| 9900 | *KRT16* | 2.7744 | 0.042576 | -0.24142 | -0.64458 | 0.86 |
| 9901 | *LHFP* | 1.4147 | 1.283 | -0.1224 | -1.3694 | 0.86 |
| 9902 | *ZMAT1* | 1.4909 | 0.64084 | 0.44331 | 0.31041 | 0.86 |
| 9903 | *REPS2* | 1.1829 | 1.1175 | 0.27293 | -2.3145 | 0.86 |
| 9904 | *NDEL1* | 4.6205 | -0.42593 | -1.6213 | -1.9734 | 0.86 |
| 9905 | *STK38L* | 2.6739 | 0.15359 | -0.25452 | -1.4946 | 0.86 |
| 9906 | *ZNF385B* | 1.6999 | 0.55449 | 0.31824 | -0.30045 | 0.86 |
| 9907 | *HERC5* | 1.2969 | 1.1068 | 0.16852 | -0.37417 | 0.86 |
| 9908 | *TMSB4X* | 3.6734 | 0.4482 | -1.5499 | -1.8761 | 0.86 |
| 9909 | *KIAA1467* | 2.4629 | 1.1032 | -0.99508 | -1.9227 | 0.86 |
| 9910 | *PAWR* | 1.5895 | 0.54154 | 0.43912 | 0.088767 | 0.86 |
| 9911 | *ARHGAP10* | 3.2248 | 0.17199 | -0.82833 | -1.7911 | 0.86 |
| 9912 | *MAFK* | 3.5131 | -0.07094 | -0.87395 | -2.2676 | 0.86 |
| 9913 | *DQX1* | 2.1166 | 0.76393 | -0.31244 | -0.37791 | 0.86 |
| 9914 | *APEH* | 1.8701 | 0.50306 | 0.19474 | -0.60258 | 0.86 |
| 9915 | *SPNS1* | 2.0238 | 0.8659 | -0.32193 | -1.5556 | 0.86 |
| 9916 | *CDRT4* | 1.2563 | 0.66736 | 0.64372 | 0.148 | 0.86 |
| 9917 | *MSRB1* | 2.4654 | 0.71461 | -0.61264 | -0.97252 | 0.86 |
| 9918 | *ERCC4* | 3.5164 | -0.15394 | -0.79535 | -1.2399 | 0.86 |
| 9919 | *FAM26F* | 4.5457 | -0.80581 | -1.173 | -1.1964 | 0.86 |
| 9920 | *BACE2* | 1.3339 | 0.79061 | 0.4418 | -2.0752 | 0.86 |
| 9921 | *ORAI2* | 1.6237 | 0.83755 | 0.10504 | 0.021712 | 0.86 |
| 9922 | *PARP12* | 4.4271 | -0.45615 | -1.4055 | -1.6447 | 0.86 |
| 9923 | *GRIA2* | 1.9187 | 1.5648 | -0.91842 | -1.5297 | 0.86 |
| 9924 | *SEPT8* | 2.0823 | 1.0832 | -0.60174 | -0.70732 | 0.85 |
| 9925 | *ESD* | 2.2809 | 0.60652 | -0.32428 | -0.85799 | 0.85 |
| 9926 | *FAM161A* | 1.894 | 0.89296 | -0.22455 | -1.3644 | 0.85 |
| 9927 | *ATP2B4* | 3.3216 | 0.4726 | -1.2329 | -1.3496 | 0.85 |
| 9928 | *SLC6A16* | 2.324 | 1.3809 | -1.1442 | -1.7247 | 0.85 |
| 9929 | *MSANTD2* | 1.702 | 0.93187 | -0.07341 | -1.7068 | 0.85 |

| | | | | | | |
|---|---|---|---|---|---|---|
| 9930 | *HBS1L* | 3.1096 | 0.088179 | -0.63753 | -0.77616 | 0.85 |
| 9931 | *ATP6V1E2* | 2.5748 | 0.13558 | -0.15145 | -1.4123 | 0.85 |
| 9932 | *AGAP6* | 1.7609 | 1.6976 | -0.89988 | -2.9678 | 0.85 |
| 9933 | *FITM1* | 3.2916 | -0.15596 | -0.57704 | -1.2867 | 0.85 |
| 9934 | *CREBRF* | 3.5455 | 0.41089 | -1.3978 | -1.9709 | 0.85 |
| 9935 | *OR1Q1* | 2.4986 | 0.30139 | -0.24142 | -0.98486 | 0.85 |
| 9936 | *ASAH2* | 2.9102 | 0.031866 | -0.38472 | -1.0259 | 0.85 |
| 9937 | *PRR23C* | 1.0161 | 0.98392 | 0.55641 | -1.0475 | 0.85 |
| 9938 | *ATP6V1E1* | 2.6758 | 0.84514 | -0.96492 | -1.9065 | 0.85 |
| 9939 | *TMSB10* | 2.4503 | 1.3902 | -1.2852 | -1.9727 | 0.85 |
| 9940 | *NRIP2* | 1.372 | 0.61455 | 0.56796 | 0.25771 | 0.85 |
| 9941 | *ZNF547* | 2.3073 | 0.67866 | -0.43202 | -1.5611 | 0.85 |
| 9942 | *TRMT10C* | 3.5001 | 0.35283 | -1.2995 | -2.7191 | 0.85 |
| 9943 | *C16orf89* | 1.8282 | 0.42359 | 0.30019 | -1.8528 | 0.85 |
| 9944 | *NMRK2* | 1.405 | 1.2706 | -0.12397 | -0.78345 | 0.85 |
| 9945 | *SCRN1* | 3.4312 | 0.17562 | -1.0553 | -1.3857 | 0.85 |
| 9946 | *MYH1* | 4.308 | 0.25771 | -2.0146 | -2.2712 | 0.85 |
| 9947 | *EPHB2* | 1.9247 | 0.93092 | -0.30487 | -1.3495 | 0.85 |
| 9948 | *CXCL3* | 4.1159 | -0.52458 | -1.0413 | -1.1991 | 0.85 |
| 9949 | *COQ2* | 2.5174 | 0.9242 | -0.89167 | -1.0231 | 0.85 |
| 9950 | *CACNB4* | 4.0801 | 0.047718 | -1.578 | -2.6376 | 0.85 |
| 9951 | *GARNL3* | 2.4939 | 0.092509 | -0.0378 | -0.84158 | 0.85 |
| 9952 | *LARP4B* | 2.8387 | 0.052562 | -0.34356 | -0.42609 | 0.85 |
| 9953 | *C3orf35* | 1.7584 | 1.4637 | -0.67472 | -1.1507 | 0.85 |
| 9954 | *ABCD4* | 3.7877 | 0.44439 | -1.6848 | -1.7364 | 0.85 |
| 9955 | *HELB* | 4.3402 | -0.47353 | -1.3198 | -1.8002 | 0.85 |
| 9956 | *GLI1* | 2.287 | 0.17614 | 0.08342 | -0.0907 | 0.85 |
| 9957 | *EFNA5* | 4.1573 | 0.043375 | -1.6561 | -2.3046 | 0.85 |
| 9958 | *EML4* | 3.9111 | 0.039836 | -1.4064 | -1.5651 | 0.85 |
| 9959 | *MPPED1* | 2.3015 | 0.50578 | -0.26429 | -0.59455 | 0.85 |
| 9960 | *NECAP2* | 4.1926 | -0.57448 | -1.0775 | -1.8909 | 0.85 |
| 9961 | *CLTCL1* | 3.7542 | -0.12995 | -1.084 | -2.7941 | 0.85 |
| 9962 | *HOXD12* | 3.0285 | -0.01188 | -0.47665 | -1.0613 | 0.85 |
| 9963 | *ZBTB3* | 2.8375 | 0.81004 | -1.1085 | -2.058 | 0.85 |
| 9964 | *PTK2B* | 2.0127 | 0.61518 | -0.08915 | -1.0551 | 0.85 |
| 9965 | *NPR1* | 2.0479 | 0.71182 | -0.22106 | -0.7331 | 0.85 |
| 9966 | *DEPDC1B* | 2.436 | 0.17071 | -0.06816 | -1.584 | 0.85 |
| 9967 | *UBL5* | 2.5948 | 0.59529 | -0.65184 | -2.2613 | 0.85 |
| 9968 | *ITPRIP* | 3.4011 | 0.068986 | -0.93294 | -1.4878 | 0.85 |
| 9969 | *ZNF280C* | 3.5179 | -0.25608 | -0.72473 | -1.7073 | 0.85 |
| 9970 | *NHS* | 1.5801 | 0.62998 | 0.32673 | -2.8398 | 0.85 |
| 9971 | *RAPGEF5* | 2.3335 | 0.11143 | 0.09132 | -1.1387 | 0.85 |
| 9972 | *ZNF440* | 1.7945 | 1.0942 | -0.35292 | -0.40749 | 0.85 |
| 9973 | *PRKD1* | 3.1886 | 0.070081 | -0.72324 | -1.1768 | 0.85 |
| 9974 | *MCTP2* | 2.938 | 0.5355 | -0.94052 | -1.3729 | 0.84 |
| 9975 | *SIGIRR* | 2.432 | 0.40537 | -0.30564 | -0.57001 | 0.84 |

| | | | | | | |
|---|---|---|---|---|---|---|
| 9976 | *NOTCH2* | 3.0952 | -0.07692 | -0.48707 | -1.6049 | 0.84 |
| 9977 | *SERPINB12* | 1.9066 | 0.49406 | 0.1304 | -1.2951 | 0.84 |
| 9978 | *BMP2K* | 1.2865 | 0.7839 | 0.46044 | -0.34451 | 0.84 |
| 9979 | *BBIP1* | 3.8575 | -0.53069 | -0.79641 | -1.0951 | 0.84 |
| 9980 | *TFEC* | 2.133 | 0.3849 | 0.01243 | -0.2704 | 0.84 |
| 9981 | *SLC9B1* | 2.649 | 0.56059 | -0.67934 | -1.3526 | 0.84 |
| 9982 | *TULP3* | 2.7701 | 1.6686 | -1.9086 | -2.4481 | 0.84 |
| 9983 | *ZNF562* | 1.7724 | 0.39643 | 0.36048 | -0.34886 | 0.84 |
| 9984 | *GATA2* | 3.1228 | 0.106 | -0.69955 | -2.0949 | 0.84 |
| 9985 | *DLG1* | 2.1036 | 0.55641 | -0.1313 | -0.95933 | 0.84 |
| 9986 | *MRPL10* | 2.801 | 0.14671 | -0.41967 | -2.0548 | 0.84 |
| 9987 | *PLTP* | 3.7775 | -0.12428 | -1.1263 | -1.7979 | 0.84 |
| 9988 | *PTPRG* | 4.0858 | -0.37888 | -1.181 | -1.2666 | 0.84 |
| 9989 | *C8B* | 2.6195 | 0.12739 | -0.22181 | -0.68095 | 0.84 |
| 9990 | *SLC38A11* | 2.5987 | 0.36093 | -0.43488 | -0.83459 | 0.84 |
| 9991 | *GABRA1* | 1.9245 | 0.64799 | -0.04796 | -1.3078 | 0.84 |
| 9992 | *ZNF619* | 2.4797 | 0.62365 | -0.5795 | -0.90757 | 0.84 |
| 9993 | *HSPH1* | 3.8595 | 0.19374 | -1.5297 | -2.1445 | 0.84 |
| 9994 | *SEPT10* | 3.8595 | -0.29445 | -1.043 | -1.3353 | 0.84 |
| 9995 | *HLA-DPA1* | 3.4066 | 0.32753 | -1.2125 | -1.2876 | 0.84 |
| 9996 | *B4GALT7* | 2.0334 | 0.71104 | -0.22302 | -1.7577 | 0.84 |
| 9997 | *HAVCR1* | 2.28 | 0.3605 | -0.1198 | -0.21407 | 0.84 |
| 9998 | *ZNF503* | 3.7196 | -0.0164 | -1.1834 | -1.8382 | 0.84 |
| 9999 | *OR2T3* | 2.7909 | 0.12142 | -0.3931 | -0.69961 | 0.84 |
| 10000 | *CAMP* | 2.9027 | -0.0083 | -0.37612 | -0.45185 | 0.84 |
| 10001 | *GPAA1* | 3.0957 | -0.15139 | -0.42652 | -1.1017 | 0.84 |
| 10002 | *GOLT1A* | 3.1278 | 0.44901 | -1.0598 | -2.0095 | 0.84 |
| 10003 | *RBPMS2* | 3.3547 | -0.39433 | -0.44353 | -0.49824 | 0.84 |
| 10004 | *GYG2* | 4.696 | -0.95321 | -1.2261 | -2.1004 | 0.84 |
| 10005 | *UBE2Q2* | 2.0449 | 1.1893 | -0.71753 | -1.1726 | 0.84 |
| 10006 | *ST8SIA3* | 3.2727 | 0.025891 | -0.78295 | -2.5127 | 0.84 |
| 10007 | *ANKRD26* | 3.1008 | 0.33778 | -0.92326 | -1.7618 | 0.84 |
| 10008 | *RRP7A* | 3.5835 | -0.35891 | -0.71009 | -2.3023 | 0.84 |
| 10009 | *OR5M3* | 4.4317 | 0.13096 | -2.0483 | -3.4283 | 0.84 |
| 10010 | *BARD1* | 1.0483 | 0.96611 | 0.4979 | 0.22092 | 0.84 |
| 10011 | *RXRG* | 3.1663 | 0.84519 | -1.4997 | -1.8323 | 0.84 |
| 10012 | *PAPD4* | 2.5791 | -0.00357 | -0.06407 | -0.66107 | 0.84 |
| 10013 | *SMARCB1* | 3.3989 | -0.10473 | -0.78279 | -2.3235 | 0.84 |
| 10014 | *CYP2F1* | 3.2029 | -0.32666 | -0.36583 | -1.4856 | 0.84 |
| 10015 | *DGCR14* | 3.6348 | 0.26867 | -1.3945 | -1.8291 | 0.84 |
| 10016 | *C5AR2* | 1.6331 | 1.1589 | -0.28335 | -0.55284 | 0.84 |
| 10017 | *SYTL4* | 4.2594 | -0.45529 | -1.296 | -2.1648 | 0.84 |
| 10018 | *MFSD12* | 4.011 | -0.69915 | -0.8038 | -1.4998 | 0.84 |
| 10019 | *ZNF772* | 1.2278 | 0.91479 | 0.36479 | -0.41359 | 0.84 |
| 10020 | *UBE2L6* | 1.7726 | 1.0961 | -0.36266 | -0.96962 | 0.84 |
| 10021 | *TAS2R46* | 3.9737 | -0.17663 | -1.2912 | -1.4969 | 0.84 |

| | | | | | | |
|---|---|---|---|---|---|---|
| 10022 | *YIPF6* | 3.2023 | 0.32065 | -1.0178 | -1.6251 | 0.84 |
| 10023 | *IQCH* | 1.2658 | 0.82411 | 0.41519 | -0.45707 | 0.84 |
| 10024 | *NOD1* | 3.3645 | 0.88961 | -1.7493 | -1.8046 | 0.83 |
| 10025 | *HSPA9* | 3.3649 | 0.089583 | -0.94989 | -1.6789 | 0.83 |
| 10026 | *PIGK* | 4.157 | -0.03224 | -1.6208 | -2.0351 | 0.83 |
| 10027 | *CACNA1C* | 2.4287 | 0.59865 | -0.52359 | -1.7438 | 0.83 |
| 10028 | *SCAMP4* | 1.5741 | 1.1166 | -0.18707 | -2.4405 | 0.83 |
| 10029 | *ADAM9* | 1.6095 | 1.2579 | -0.36401 | -1.9284 | 0.83 |
| 10030 | *CD207* | 1.8627 | 0.57942 | 0.05962 | -1.3371 | 0.83 |
| 10031 | *RARS* | 2.7876 | 0.53256 | -0.81932 | -1.1685 | 0.83 |
| 10032 | *CABP2* | 2.1037 | 1.4207 | -1.0248 | -1.8636 | 0.83 |
| 10033 | *SIPA1L3* | 4.9852 | -1.0054 | -1.4809 | -1.5965 | 0.83 |
| 10034 | *OR8D4* | 3.0847 | 0.092413 | -0.67868 | -1.9621 | 0.83 |
| 10035 | *TDG* | 3.892 | 0.064489 | -1.4584 | -2.7784 | 0.83 |
| 10036 | *C8orf76* | 2.0447 | 0.31401 | 0.13862 | 0.12728 | 0.83 |
| 10037 | *DPY19L1* | 1.5727 | 0.90263 | 0.0205 | -2.1956 | 0.83 |
| 10038 | *RHBDD1* | 3.2352 | -0.08265 | -0.65758 | -1.2618 | 0.83 |
| 10039 | *VGLL2* | 2.6159 | 0.76772 | -0.8897 | -1.6469 | 0.83 |
| 10040 | *ADAMTS7* | 1.3991 | 0.80653 | 0.28742 | -0.3271 | 0.83 |
| 10041 | *TNFRSF19* | 1.492 | 0.76676 | 0.23339 | -1.1511 | 0.83 |
| 10042 | *MARCH11* | 0.96981 | 0.77718 | 0.74484 | 0.54726 | 0.83 |
| 10043 | *FXYD1* | 1.2101 | 0.85322 | 0.42813 | -0.44353 | 0.83 |
| 10044 | *FAM13B* | 2.5105 | 0.46439 | -0.48368 | -2.2866 | 0.83 |
| 10045 | *GALNT16* | 3.5257 | 0.0077 | -1.0426 | -1.507 | 0.83 |
| 10046 | *GDF10* | 1.7193 | 1.5835 | -0.81264 | -1.4852 | 0.83 |
| 10047 | *RABGAP1L* | 2.6345 | 0.15379 | -0.29862 | -1.3426 | 0.83 |
| 10048 | *OR11H4* | 3.1758 | 0.17593 | -0.86229 | -1.371 | 0.83 |
| 10049 | *DUSP23* | 1.9567 | 0.39128 | 0.14083 | -0.14716 | 0.83 |
| 10050 | *DENND3* | 2.3873 | 1.157 | -1.0563 | -1.5962 | 0.83 |
| 10051 | *ISOC1* | 2.6303 | 0.13279 | -0.27522 | -1.5273 | 0.83 |
| 10052 | *LACE1* | 1.9758 | 0.30138 | 0.21066 | -0.85617 | 0.83 |
| 10053 | *STON1* | 2.4011 | 0.082463 | 0.00424 | -0.31299 | 0.83 |
| 10054 | *CD46* | 1.3128 | 1.139 | 0.036 | -1.12 | 0.83 |
| 10055 | *ZNF8* | 1.7849 | 0.49953 | 0.20314 | 0.006651 | 0.83 |
| 10056 | *ITLN2* | 3.8768 | -0.2456 | -1.1437 | -2.555 | 0.83 |
| 10057 | *SESN2* | 2.054 | 1.1527 | -0.71935 | -0.96791 | 0.83 |
| 10058 | *SCN2B* | 2.6625 | 0.77231 | -0.94747 | -4.9483 | 0.83 |
| 10059 | *ZNF226* | 2.101 | 0.7244 | -0.33844 | -3.3058 | 0.83 |
| 10060 | *OR2T4* | 2.7921 | 0.80432 | -1.1098 | -2.3461 | 0.83 |
| 10061 | *NEDD9* | 3.4283 | -0.42671 | -0.51549 | -2.0629 | 0.83 |
| 10062 | *SCP2* | 1.8141 | 0.71125 | -0.03944 | -0.56507 | 0.83 |
| 10063 | *CAMSAP2* | 3.9425 | -0.54427 | -0.91234 | -1.8443 | 0.83 |
| 10064 | *TMC8* | 4.3549 | -0.80823 | -1.0613 | -1.8751 | 0.83 |
| 10065 | *ANGEL2* | 1.5773 | 0.54151 | 0.36641 | -1.4824 | 0.83 |
| 10066 | *TCTN1* | 5.0108 | -1.208 | -1.3183 | -2.5125 | 0.83 |
| 10067 | *OPRD1* | 3.7579 | -0.25794 | -1.016 | -1.94 | 0.83 |

| | | | | | | |
|---|---|---|---|---|---|---|
| 10068 | *ADAT1* | 0.87679 | 0.82264 | 0.78393 | -1.7061 | 0.83 |
| 10069 | *RNF10* | 4.3298 | -0.44353 | -1.403 | -2.26 | 0.83 |
| 10070 | *SMN2* | 3.7169 | -0.30753 | -0.92781 | -1.2743 | 0.83 |
| 10071 | *MRPL14* | 4.7705 | -0.92486 | -1.3643 | -2.012 | 0.83 |
| 10072 | *KCNS1* | 3.4914 | 0.31994 | -1.33 | -2.5754 | 0.83 |
| 10073 | *MYL5* | 2.8572 | 0.12226 | -0.49818 | -0.70057 | 0.83 |
| 10074 | *COL6A5* | 2.9161 | -0.15439 | -0.28058 | -0.48118 | 0.83 |
| 10075 | *OGFOD3* | 1.3533 | 1.1495 | -0.02187 | -0.08446 | 0.83 |
| 10076 | *UBFD1* | 4.5715 | -0.46205 | -1.6288 | -2.2343 | 0.83 |
| 10077 | *ACADM* | 1.6998 | 0.6201 | 0.16025 | -1.765 | 0.83 |
| 10078 | *IRF2BPL* | 2.3154 | 0.097924 | 0.06487 | -0.07436 | 0.83 |
| 10079 | *CNTD1* | 3.771 | -0.53009 | -0.76371 | -1.2908 | 0.83 |
| 10080 | *FGFR3* | 1.5139 | 1.2629 | -0.30038 | -0.59643 | 0.83 |
| 10081 | *UBAP2* | 1.3709 | 0.58003 | 0.52482 | -1.3407 | 0.83 |
| 10082 | *SEMA6A* | 2.114 | 0.241 | 0.12019 | -0.2765 | 0.83 |
| 10083 | *SLC29A4* | 1.6982 | 0.45311 | 0.32349 | 0.10037 | 0.82 |
| 10084 | *PLA2G2E* | 1.9677 | 1.9302 | -1.4234 | -2.586 | 0.82 |
| 10085 | *DUOXA2* | 4.3754 | -0.88873 | -1.0129 | -1.0796 | 0.82 |
| 10086 | *TTLL10* | 1.1534 | 1.0462 | 0.27327 | -0.33704 | 0.82 |
| 10087 | *C1QTNF2* | 3.1353 | -0.19532 | -0.46731 | -1.9442 | 0.82 |
| 10088 | *C1orf61* | 1.1894 | 1.1018 | 0.18144 | -0.37557 | 0.82 |
| 10089 | *C7orf62* | 2.4753 | 0.21608 | -0.21887 | -0.40714 | 0.82 |
| 10090 | *IMPG1* | 3.0397 | 0.015765 | -0.58334 | -1.2198 | 0.82 |
| 10091 | *ZFPM2* | 2.3111 | 0.13801 | 0.02287 | -1.7665 | 0.82 |
| 10092 | *ISL2* | 2.4901 | 0.42035 | -0.43946 | -0.5718 | 0.82 |
| 10093 | *TLR6* | 2.7556 | 0.93531 | -1.2201 | -2.6567 | 0.82 |
| 10094 | *ZC3H12B* | 1.4572 | 0.76694 | 0.24637 | -2.1219 | 0.82 |
| 10095 | *BRIX1* | 3.6212 | -0.03006 | -1.1207 | -2.2167 | 0.82 |
| 10096 | *ZNF791* | 1.373 | 1.0167 | 0.08041 | -1.2926 | 0.82 |
| 10097 | *CLECL1* | 3.1408 | -0.28559 | -0.38529 | -1.8816 | 0.82 |
| 10098 | *KHDRBS2* | 1.8803 | 0.40192 | 0.18761 | -1.7911 | 0.82 |
| 10099 | *DAZ2* | 2.3901 | 0.87663 | -0.79798 | -1.7061 | 0.82 |
| 10100 | *SLC35E3* | 2.625 | 1.1303 | -1.2867 | -1.6618 | 0.82 |
| 10101 | *OR8D1* | 1.1824 | 0.69377 | 0.59214 | -0.0937 | 0.82 |
| 10102 | *ROR1* | 3.0823 | 0.7361 | -1.3501 | -1.9341 | 0.82 |
| 10103 | *DOCK10* | 4.6049 | -0.78229 | -1.3567 | -1.8246 | 0.82 |
| 10104 | *OR5B12* | 3.5923 | -0.22653 | -0.90003 | -1.0425 | 0.82 |
| 10105 | *CSNK1D* | 3.6193 | -0.26714 | -0.88661 | -1.2517 | 0.82 |
| 10106 | *GPR50* | 2.7459 | 0.64483 | -0.92613 | -1.0273 | 0.82 |
| 10107 | *NXT1* | 4.0477 | -0.63406 | -0.94912 | -1.4255 | 0.82 |
| 10108 | *TTLL4* | 3.3122 | -0.03081 | -0.817 | -1.4566 | 0.82 |
| 10109 | *WRN* | 3.4161 | 0.014271 | -0.96632 | -1.7757 | 0.82 |
| 10110 | *ZNF343* | 2.6403 | 0.26595 | -0.44353 | -0.63452 | 0.82 |
| 10111 | *ZNF674* | 1.6604 | 0.7476 | 0.05256 | -0.06618 | 0.82 |
| 10112 | *RPL35* | 1.9076 | 0.40349 | 0.1486 | -1.384 | 0.82 |
| 10113 | *ZNF768* | 3.6385 | -0.41139 | -0.76755 | -1.4811 | 0.82 |

| | | | | | | |
|---|---|---|---|---|---|---|
| 10114 | *BMPER* | 2.3231 | 1.1987 | -1.0623 | -1.6648 | 0.82 |
| 10115 | *MAP3K13* | 3.0764 | 0.16733 | -0.78439 | -0.93107 | 0.82 |
| 10116 | *EHD1* | 4.7912 | -0.93592 | -1.3961 | -1.8645 | 0.82 |
| 10117 | *PLGLB2* | 1.1261 | 0.71129 | 0.62128 | -1.72 | 0.82 |
| 10118 | *PCBP1* | 2.6168 | 0.2171 | -0.37602 | -1.1587 | 0.82 |
| 10119 | *SYAP1* | 2.1559 | 1.1167 | -0.81541 | -1.1812 | 0.82 |
| 10120 | *SPATC1* | 2.6064 | 0.46514 | -0.61467 | -0.67846 | 0.82 |
| 10121 | *KATNAL1* | 2.2077 | 0.64807 | -0.39997 | -0.40398 | 0.82 |
| 10122 | *MC4R* | 3.548 | -0.43674 | -0.65553 | -2.5103 | 0.82 |
| 10123 | *FAM149B1* | 4.9997 | -1.0372 | -1.5077 | -1.8966 | 0.82 |
| 10124 | *RABEP1* | 2.9816 | 0.52282 | -1.0497 | -1.455 | 0.82 |
| 10125 | *C3orf70* | 4.7884 | -0.93287 | -1.4031 | -2.4955 | 0.82 |
| 10126 | *PAQR5* | 4.0369 | 0.098257 | -1.6836 | -2.6463 | 0.82 |
| 10127 | *DDIT3* | 1.6964 | 0.97771 | -0.2227 | -0.69815 | 0.82 |
| 10128 | *MCCC1* | 1.3955 | 0.99784 | 0.05634 | -0.43478 | 0.82 |
| 10129 | *SH3PXD2A* | 2.704 | 0.78829 | -1.0435 | -1.4402 | 0.82 |
| 10130 | *MXD4* | 2.7435 | 0.45984 | -0.75472 | -1.6913 | 0.82 |
| 10131 | *CRYGS* | 2.9503 | 0.72805 | -1.2302 | -2.0818 | 0.82 |
| 10132 | *ASPDH* | 1.7897 | 0.75012 | -0.0925 | -0.5898 | 0.82 |
| 10133 | *FSBP* | 1.7212 | 0.43453 | 0.29115 | 0.076188 | 0.82 |
| 10134 | *GABRG3* | 4.8881 | -1.1974 | -1.244 | -1.6597 | 0.82 |
| 10135 | *KPRP* | 1.5488 | 0.58812 | 0.30864 | -0.53911 | 0.82 |
| 10136 | *RRAD* | 1.99 | 1.0721 | -0.61737 | -1.5634 | 0.81 |
| 10137 | *DERL3* | 1.748 | 1.3235 | -0.62814 | -0.81101 | 0.81 |
| 10138 | *GPATCH2L* | 2.4009 | 0.63732 | -0.5961 | -1.1729 | 0.81 |
| 10139 | *PPP1R42* | 2.1358 | 0.35781 | -0.05252 | -2.3857 | 0.81 |
| 10140 | *TCERG1* | 2.4719 | 0.78652 | -0.81757 | -2.0126 | 0.81 |
| 10141 | *GLB1* | 1.6773 | 0.69338 | 0.0693 | -0.17549 | 0.81 |
| 10142 | *SEL1L3* | 2.6309 | 0.49202 | -0.68303 | -1.9827 | 0.81 |
| 10143 | *SLCO1B1* | 3.8939 | -0.5656 | -0.88843 | -1.0724 | 0.81 |
| 10144 | *STX1B* | 2.8858 | 0.46392 | -0.90991 | -1.6817 | 0.81 |
| 10145 | *GRXCR1* | 1.9572 | 0.36875 | 0.1134 | -1.677 | 0.81 |
| 10146 | *DHRS2* | 4.4118 | 0.19804 | -2.1714 | -2.3461 | 0.81 |
| 10147 | *ADGRF4* | 1.3873 | 0.62252 | 0.42783 | -0.48085 | 0.81 |
| 10148 | *PTX4* | 2.1314 | 1.227 | -0.92097 | -1.213 | 0.81 |
| 10149 | *ATP5J* | 5.5467 | -1.0145 | -2.0953 | -2.1217 | 0.81 |
| 10150 | *MAPK11* | 3.9487 | 0.33438 | -1.8462 | -2.1151 | 0.81 |
| 10151 | *C11orf52* | 1.4165 | 1.3186 | -0.29861 | -0.83749 | 0.81 |
| 10152 | *DYX1C1* | 3.0314 | 0.04945 | -0.64441 | -0.67629 | 0.81 |
| 10153 | *SGK494* | 4.5489 | -0.87673 | -1.236 | -1.2729 | 0.81 |
| 10154 | *TBC1D7* | 2.3752 | 0.88995 | -0.82921 | -2.2537 | 0.81 |
| 10155 | *KTI12* | 1.6313 | 1.1136 | -0.30975 | -1.5722 | 0.81 |
| 10156 | *IGFALS* | 2.4121 | 0.37386 | -0.35098 | -1.3542 | 0.81 |
| 10157 | *LCN9* | 3.2796 | 0.69659 | -1.5425 | -1.9112 | 0.81 |
| 10158 | *CYB5R4* | 3.1944 | -0.27076 | -0.49095 | -2.5878 | 0.81 |
| 10159 | *EDNRA* | 1.2108 | 1.0111 | 0.20954 | -0.96572 | 0.81 |

| | | | | | | |
|---|---|---|---|---|---|---|
| 10160 | *TCF7L2* | 2.9889 | -0.1376 | -0.41994 | -0.70705 | 0.81 |
| 10161 | *LMCD1* | 3.3554 | -0.41643 | -0.50804 | -2.005 | 0.81 |
| 10162 | *PTPRA* | 1.2039 | 0.80114 | 0.42522 | -1.5911 | 0.81 |
| 10163 | *ZNF236* | 3.5884 | -0.31164 | -0.84652 | -2.2508 | 0.81 |
| 10164 | *DPYSL3* | 1.0441 | 0.90552 | 0.47937 | -1.5212 | 0.81 |
| 10165 | *WDR54* | 2.0011 | 0.55139 | -0.12414 | -0.14312 | 0.81 |
| 10166 | *C1orf233* | 2.2331 | 1.0457 | -0.85194 | -1.117 | 0.81 |
| 10167 | *SSTR4* | 1.0047 | 0.79948 | 0.6225 | -1.2934 | 0.81 |
| 10168 | *ACTR1B* | 2.3007 | 1.988 | -1.8623 | -2.0181 | 0.81 |
| 10169 | *FOSB* | 3.8846 | 0.063662 | -1.5225 | -1.8908 | 0.81 |
| 10170 | *XRCC6BP1* | 2.6581 | -0.04064 | -0.19179 | -1.3579 | 0.81 |
| 10171 | *FZD7* | 1.7185 | 0.69837 | 0.00877 | -1.7068 | 0.81 |
| 10172 | *ROPN1* | 3.9473 | 0.63392 | -2.1566 | -2.7063 | 0.81 |
| 10173 | *TDRP* | 3.2093 | 0.32948 | -1.1145 | -1.3111 | 0.81 |
| 10174 | *KAZALD1* | 1.8671 | 0.77261 | -0.21587 | -1.7044 | 0.81 |
| 10175 | *ARHGEF37* | 4.6497 | -0.46721 | -1.7591 | -2.2214 | 0.81 |
| 10176 | *SERINC2* | 2.7872 | -0.03697 | -0.3271 | -2.4785 | 0.81 |
| 10177 | *NDUFAF2* | 2.3166 | 0.59545 | -0.48916 | -0.91343 | 0.81 |
| 10178 | *MYL2* | 2.2955 | 0.44138 | -0.31411 | -0.35896 | 0.81 |
| 10179 | *DSE* | 4.43 | -0.79711 | -1.2109 | -1.7956 | 0.81 |
| 10180 | *PARVG* | 2.6276 | 0.41381 | -0.61967 | -1.6547 | 0.81 |
| 10181 | *FILIP1L* | 2.764 | -0.09774 | -0.24493 | -0.93696 | 0.81 |
| 10182 | *SNX3* | 3.0324 | -0.27219 | -0.33922 | -0.45223 | 0.81 |
| 10183 | *UROC1* | 1.9702 | 0.28838 | 0.16177 | -1.7512 | 0.81 |
| 10184 | *RPAP3* | 4.9736 | -1.1558 | -1.3985 | -1.6891 | 0.81 |
| 10185 | *GALNT12* | 2.3242 | 0.24021 | -0.14558 | -0.7819 | 0.81 |
| 10186 | *C12orf80* | 3.5988 | 0.090452 | -1.2705 | -1.6072 | 0.81 |
| 10187 | *MTRF1L* | 3.1189 | -0.00826 | -0.69199 | -1.5956 | 0.81 |
| 10188 | *CACNG4* | 1.8754 | 1.0862 | -0.54329 | -1.5959 | 0.81 |
| 10189 | *CCDC84* | 3.1957 | 0.34155 | -1.1196 | -1.3686 | 0.81 |
| 10190 | *AKAP1* | 4.0021 | 0.23099 | -1.8157 | -1.8872 | 0.81 |
| 10191 | *SMOC1* | 2.1973 | 0.2159 | 0.00418 | -0.11018 | 0.81 |
| 10192 | *FAR1* | 2.9742 | 0.78246 | -1.3403 | -1.3542 | 0.81 |
| 10193 | *HIPK3* | 4.4867 | -0.96878 | -1.1016 | -1.8938 | 0.81 |
| 10194 | *CELF5* | 2.4087 | 0.40424 | -0.39731 | -1.095 | 0.81 |
| 10195 | *DGCR2* | 1.7535 | 1.5061 | -0.84411 | -1.5724 | 0.81 |
| 10196 | *OR10G7* | 3.4273 | -0.04795 | -0.9644 | -1.3836 | 0.80 |
| 10197 | *PARP9* | 1.7971 | 0.90587 | -0.28915 | -2.0952 | 0.80 |
| 10198 | *MCM10* | 1.6991 | 0.4543 | 0.26031 | -2.3243 | 0.80 |
| 10199 | *PRSS58* | 1.8904 | 0.954 | -0.43077 | -2.312 | 0.80 |
| 10200 | *PROM1* | 3.938 | -0.15715 | -1.3686 | -1.8462 | 0.80 |
| 10201 | *IL36A* | 2.7109 | 0.88918 | -1.1879 | -2.072 | 0.80 |
| 10202 | *EXOC3L2* | 2.7345 | 0.26387 | -0.58691 | -0.77848 | 0.80 |
| 10203 | *ITGA7* | 2.1859 | 0.50484 | -0.27941 | -1.8859 | 0.80 |
| 10204 | *PNP* | 2.4407 | 0.36257 | -0.39214 | -1.0914 | 0.80 |
| 10205 | *PLA2G1B* | 2.1633 | 1.0756 | -0.82921 | -1.7076 | 0.80 |

| | | | | | | |
|---|---|---|---|---|---|---|
| 10206 | *SERTAD2* | 0.92033 | 0.79291 | 0.69449 | -1.2985 | 0.80 |
| 10207 | *TRIM49C* | 1.6213 | 0.63529 | 0.15108 | -0.79057 | 0.80 |
| 10208 | *DMKN* | 3.0016 | -0.24442 | -0.35015 | -1.1836 | 0.80 |
| 10209 | *TRA2A* | 3.8633 | -0.67458 | -0.78229 | -1.0564 | 0.80 |
| 10210 | *ECD* | 3.052 | -0.07232 | -0.57397 | -1.3903 | 0.80 |
| 10211 | *BCL2L10* | 3.3547 | -0.1309 | -0.81943 | -1.122 | 0.80 |
| 10212 | *C2orf78* | 3.1157 | -0.33821 | -0.37335 | -0.49056 | 0.80 |
| 10213 | *NWD2* | 3.0626 | -0.20771 | -0.45122 | -2.4901 | 0.80 |
| 10214 | *UCKL1* | 0.98949 | 0.89401 | 0.51997 | -0.26267 | 0.80 |
| 10215 | *NNMT* | 3.9111 | -0.70002 | -0.80847 | -1.7279 | 0.80 |
| 10216 | *PXDC1* | 3.457 | 0.11845 | -1.173 | -1.2703 | 0.80 |
| 10217 | *RNF180* | 2.3375 | 0.50115 | -0.43706 | -1.6684 | 0.80 |
| 10218 | *HIVEP2* | 4.2542 | -0.20023 | -1.6525 | -2.1357 | 0.80 |
| 10219 | *DPP9* | 3.7134 | 0.24439 | -1.5568 | -1.5724 | 0.80 |
| 10220 | *CDKN2AIPNL* | 2.7322 | 0.26787 | -0.59987 | -0.80599 | 0.80 |
| 10221 | *ZCCHC4* | 2.4241 | 0.040889 | -0.0648 | -1.0059 | 0.80 |
| 10222 | *YTHDC1* | 1.8067 | 0.5365 | 0.0566 | -1.3593 | 0.80 |
| 10223 | *SGSM2* | 2.5716 | 0.15656 | -0.32842 | -1.3572 | 0.80 |
| 10224 | *HEYL* | 1.1293 | 1.0951 | 0.17443 | 0.11456 | 0.80 |
| 10225 | *GPR27* | 2.9498 | 0.45695 | -1.011 | -1.1043 | 0.80 |
| 10226 | *JAM2* | 2.6734 | 0.1087 | -0.3866 | -0.44313 | 0.80 |
| 10227 | *FADS1* | 1.764 | 0.37182 | 0.25944 | -1.9259 | 0.80 |
| 10228 | *FAIM3* | 3.4429 | -0.1931 | -0.85508 | -1.7911 | 0.80 |
| 10229 | *HSD3B1* | 3.591 | -0.18235 | -1.0142 | -2.7171 | 0.80 |
| 10230 | *KLHDC3* | 3.3483 | 0.10131 | -1.0553 | -1.9567 | 0.80 |
| 10231 | *OGFOD2* | 2.9211 | 0.42976 | -0.95749 | -1.1305 | 0.80 |
| 10232 | *MYO5C* | 1.4875 | 0.55717 | 0.34868 | -1.0528 | 0.80 |
| 10233 | *C4orf27* | 1.2598 | 0.59737 | 0.53514 | -0.59711 | 0.80 |
| 10234 | *SETX* | 1.3645 | 1.3359 | -0.30811 | -2.4338 | 0.80 |
| 10235 | *CAPN1* | 2.4362 | 0.82342 | -0.86756 | -2.1921 | 0.80 |
| 10236 | *FBXO15* | 2.0406 | 0.64946 | -0.2982 | -1.0765 | 0.80 |
| 10237 | *SOX30* | 2.5254 | 0.33086 | -0.46441 | -1.7718 | 0.80 |
| 10238 | *TMEM170A* | 3.4879 | -0.18864 | -0.90791 | -1.7112 | 0.80 |
| 10239 | *LDB2* | 1.9085 | 0.53479 | -0.0534 | -1.0558 | 0.80 |
| 10240 | *TOM1L2* | 2.498 | 0.072685 | -0.1815 | -1.5862 | 0.80 |
| 10241 | *FOXJ2* | 2.8448 | -0.12456 | -0.33191 | -1.2867 | 0.80 |
| 10242 | *F7* | 2.8334 | -0.02163 | -0.42369 | -0.87128 | 0.80 |
| 10243 | *C14orf142* | 2.4676 | 0.9346 | -1.0158 | -1.5568 | 0.80 |
| 10244 | *SH3RF1* | 3.4191 | -0.30692 | -0.72666 | -1.4299 | 0.80 |
| 10245 | *UBE2A* | 1.4854 | 1.248 | -0.34814 | -0.44353 | 0.80 |
| 10246 | *SCUBE1* | 3.8755 | -0.70945 | -0.78236 | -1.7585 | 0.79 |
| 10247 | *RTTN* | 2.9798 | 0.07636 | -0.67292 | -0.86412 | 0.79 |
| 10248 | *AIF1* | 2.6678 | 0.4863 | -0.77144 | -1.1805 | 0.79 |
| 10249 | *CTDSPL* | 4.6434 | -0.34882 | -1.9123 | -2.0196 | 0.79 |
| 10250 | *OR51B6* | 3.0722 | -0.20283 | -0.48725 | -0.54741 | 0.79 |
| 10251 | *RIMS2* | 5.6746 | -0.93107 | -2.3615 | -2.5006 | 0.79 |

| | | | | | | |
|---|---|---|---|---|---|---|
| 10252 | *HRC* | 2.3755 | 0.37028 | -0.36451 | -1.2662 | 0.79 |
| 10253 | *BPTF* | 1.7327 | 0.95951 | -0.31109 | -0.68416 | 0.79 |
| 10254 | *DCP1B* | 2.4019 | 0.88385 | -0.90606 | -2.1151 | 0.79 |
| 10255 | *RASIP1* | 2.1713 | 0.16197 | 0.04624 | -2.0326 | 0.79 |
| 10256 | *USP38* | 3.0617 | -0.31535 | -0.36724 | -0.44353 | 0.79 |
| 10257 | *CD40LG* | 4.1433 | -0.47929 | -1.2852 | -3.4111 | 0.79 |
| 10258 | *DHX32* | 2.7103 | 0.57743 | -0.90933 | -1.8352 | 0.79 |
| 10259 | *MCRS1* | 1.2928 | 0.95876 | 0.12623 | -0.65489 | 0.79 |
| 10260 | *PGM5* | 3.5085 | 0.014683 | -1.1456 | -1.8238 | 0.79 |
| 10261 | *SLC37A1* | 0.94095 | 0.87752 | 0.55809 | -0.28633 | 0.79 |
| 10262 | *CBWD5* | 2.233 | 1.1794 | -1.0359 | -1.1479 | 0.79 |
| 10263 | *ANKRD36* | 3.0343 | -0.3129 | -0.34496 | -0.50283 | 0.79 |
| 10264 | *PLAC8* | 1.3329 | 0.74716 | 0.29622 | -0.7167 | 0.79 |
| 10265 | *LHFPL1* | 3.7187 | 0.30874 | -1.6524 | -2.5754 | 0.79 |
| 10266 | *PROP1* | 1.4879 | 0.9652 | -0.07947 | -0.15667 | 0.79 |
| 10267 | *SLC29A3* | 2.3327 | 0.58116 | -0.54402 | -0.82472 | 0.79 |
| 10268 | *TSPAN12* | 3.0537 | -0.3077 | -0.37641 | -0.92427 | 0.79 |
| 10269 | *HSDL1* | 3.2658 | -0.00278 | -0.89392 | -1.0589 | 0.79 |
| 10270 | *PUS7L* | 2.219 | 0.35402 | -0.20552 | -1.9054 | 0.79 |
| 10271 | *ANKRD22* | 1.6598 | 0.85048 | -0.14309 | -1.7526 | 0.79 |
| 10272 | *NFU1* | 2.6176 | 0.21699 | -0.46759 | -1.1401 | 0.79 |
| 10273 | *TOMM34* | 1.2201 | 0.79518 | 0.35163 | -1.9099 | 0.79 |
| 10274 | *FCGR1A* | 3.0456 | 0.23395 | -0.91283 | -2.1947 | 0.79 |
| 10275 | *OR7C1* | 1.2928 | 1.2494 | -0.17577 | -0.54274 | 0.79 |
| 10276 | *ZKSCAN8* | 1.6238 | 1.394 | -0.65178 | -1.3382 | 0.79 |
| 10277 | *CS* | 3.9581 | -0.76973 | -0.82311 | -1.1205 | 0.79 |
| 10278 | *RAP1B* | 4.989 | -1.1856 | -1.4388 | -2.3294 | 0.79 |
| 10279 | *PDE7B* | 1.8563 | 0.91726 | -0.40938 | -0.58894 | 0.79 |
| 10280 | *SLC35G5* | 3.4101 | -0.25892 | -0.78812 | -1.785 | 0.79 |
| 10281 | *KRT3* | 2.1603 | 0.28586 | -0.08354 | -0.25981 | 0.79 |
| 10282 | *WISP1* | 3.9115 | -0.21146 | -1.3379 | -1.6325 | 0.79 |
| 10283 | *OR6S1* | 3.0853 | 0.42056 | -1.1438 | -2.0099 | 0.79 |
| 10284 | *COX5B* | 2.6825 | 0.25176 | -0.57241 | -1.1487 | 0.79 |
| 10285 | *STK16* | 2.0771 | 0.49312 | -0.20848 | -1.8538 | 0.79 |
| 10286 | *SLC50A1* | 3.3863 | -0.4636 | -0.56098 | -1.2662 | 0.79 |
| 10287 | *VWC2* | 1.3414 | 1.2012 | -0.18093 | -0.50246 | 0.79 |
| 10288 | *DTYMK* | 1.5363 | 1.5131 | -0.68804 | -0.69008 | 0.79 |
| 10289 | *RPGRIP1L* | 1.5775 | 0.53279 | 0.24924 | -0.14685 | 0.79 |
| 10290 | *TAS2R7* | 1.1553 | 0.89213 | 0.31154 | -1.6635 | 0.79 |
| 10291 | *KRTAP10-11* | 2.3198 | 0.59321 | -0.55437 | -0.97658 | 0.79 |
| 10292 | *MBTD1* | 2.3235 | 0.14713 | -0.11306 | -2.1318 | 0.79 |
| 10293 | *METTL18* | 1.7283 | 1.4075 | -0.77908 | -0.82514 | 0.79 |
| 10294 | *LYL1* | 2.4638 | 0.5491 | -0.65714 | -2.003 | 0.79 |
| 10295 | *LRRC57* | 2.1731 | 0.8289 | -0.64683 | -0.88277 | 0.79 |
| 10296 | *MPV17* | 2.1416 | 0.74753 | -0.53437 | -0.58485 | 0.78 |
| 10297 | *TCEB3CL2* | 1.8116 | 1.3373 | -0.79453 | -1.0111 | 0.78 |

| | | | | | | |
|---|---|---|---|---|---|---|
| 10298 | *SGK2* | 3.2979 | 0.4078 | -1.3517 | -1.6612 | 0.78 |
| 10299 | *MANEAL* | 2.8958 | 1.3201 | -1.862 | -2.312 | 0.78 |
| 10300 | *TMX1* | 2.5516 | 0.91611 | -1.1145 | -1.5671 | 0.78 |
| 10301 | *ASIC1* | 1.5611 | 0.42459 | 0.36727 | -0.97794 | 0.78 |
| 10302 | *ZBTB40* | 2.3038 | 0.75877 | -0.70998 | -1.677 | 0.78 |
| 10303 | *SIGLEC15* | 3.6398 | 0.34346 | -1.631 | -1.6663 | 0.78 |
| 10304 | *BARHL1* | 3.5575 | -0.11162 | -1.0942 | -1.8095 | 0.78 |
| 10305 | *NFIA* | 4.4105 | -0.44445 | -1.6146 | -1.8792 | 0.78 |
| 10306 | *C2orf81* | 2.2473 | 0.58543 | -0.48156 | -3.1229 | 0.78 |
| 10307 | *HHATL* | 2.6713 | -0.05456 | -0.26575 | -1.8869 | 0.78 |
| 10308 | *RAG1* | 3.9321 | -0.6176 | -0.96422 | -1.0231 | 0.78 |
| 10309 | *CDK17* | 2.453 | 0.74731 | -0.85194 | -2.3573 | 0.78 |
| 10310 | *KRT77* | 1.5119 | 0.74882 | 0.08715 | -1.0753 | 0.78 |
| 10311 | *C16orf62* | 4.4253 | -0.98653 | -1.0912 | -1.172 | 0.78 |
| 10312 | *UGT3A2* | 2.9383 | 0.016992 | -0.60837 | -1.7954 | 0.78 |
| 10313 | *SCGB1D2* | 1.7666 | 0.43928 | 0.14042 | -1.1805 | 0.78 |
| 10314 | *BOD1L1* | 2.6078 | 0.19521 | -0.45672 | -1.1317 | 0.78 |
| 10315 | *FAM71B* | 1.4012 | 1.259 | -0.31396 | -1.4247 | 0.78 |
| 10316 | *PIKFYVE* | 3.0749 | 1.5886 | -2.3174 | -2.3885 | 0.78 |
| 10317 | *ATL1* | 3.4562 | -0.46123 | -0.64888 | -0.99368 | 0.78 |
| 10318 | *DACT1* | 3.6879 | -0.1037 | -1.2395 | -1.5603 | 0.78 |
| 10319 | *NFATC2* | 2.5501 | 0.52293 | -0.72916 | -1.3574 | 0.78 |
| 10320 | *ZFYVE16* | 1.7454 | 0.54063 | 0.05771 | -1.3496 | 0.78 |
| 10321 | *GPR88* | 3.4982 | -0.34767 | -0.8071 | -1.2954 | 0.78 |
| 10322 | *SUZ12* | 2.8185 | -0.07138 | -0.40374 | -2.207 | 0.78 |
| 10323 | *ONECUT3* | 1.4024 | 0.834 | 0.10662 | -1.301 | 0.78 |
| 10324 | *HBB* | 2.2357 | 1.0171 | -0.9101 | -4.5174 | 0.78 |
| 10325 | *PIK3R5* | 3.0976 | -0.19793 | -0.55741 | -1.1337 | 0.78 |
| 10326 | *GPR149* | 3.3876 | -0.11049 | -0.93525 | -1.3982 | 0.78 |
| 10327 | *PLK5* | 2.9704 | -0.14637 | -0.48231 | -1.3356 | 0.78 |
| 10328 | *PNMAL2* | 2.3689 | 0.11868 | -0.14622 | -0.54973 | 0.78 |
| 10329 | *RAB36* | 1.5351 | 0.97281 | -0.16687 | -1.4773 | 0.78 |
| 10330 | *HTR5A* | 2.3905 | 0.41593 | -0.46647 | -1.4584 | 0.78 |
| 10331 | *TPD52L1* | 3.9585 | -0.09964 | -1.5193 | -1.6274 | 0.78 |
| 10332 | *MTRF1* | 1.3708 | 0.78469 | 0.18352 | -0.67253 | 0.78 |
| 10333 | *MBIP* | 1.4205 | 0.78556 | 0.1311 | -1.3946 | 0.78 |
| 10334 | *OR6T1* | 2.2119 | 0.21342 | -0.0887 | -1.695 | 0.78 |
| 10335 | *RBP3* | 1.8972 | 1.1339 | -0.6951 | -2.0505 | 0.78 |
| 10336 | *ZFHX3* | 3.6934 | -0.17054 | -1.1879 | -1.9013 | 0.78 |
| 10337 | *GARS* | 1.6059 | 0.70459 | 0.02439 | -1.2169 | 0.78 |
| 10338 | *CD300E* | 3.2432 | 0.31562 | -1.2246 | -2.3517 | 0.78 |
| 10339 | *PLCXD2* | 1.9618 | 0.50452 | -0.13218 | -1.7338 | 0.78 |
| 10340 | *IL1F10* | 1.848 | 1.1674 | -0.68222 | -1.8577 | 0.78 |
| 10341 | *C17orf64* | 1.9436 | 0.33355 | 0.05556 | -1.1177 | 0.78 |
| 10342 | *PRKCSH* | 2.3979 | 0.1829 | -0.24887 | -1.388 | 0.78 |
| 10343 | *FAM210B* | 2.706 | 0.25769 | -0.63207 | -1.6194 | 0.78 |

| | | | | | | |
|---|---|---|---|---|---|---|
| 10344 | *TNNI3* | 3.3665 | -0.39944 | -0.63554 | -0.85931 | 0.78 |
| 10345 | *OTOG* | 2.6155 | 0.51683 | -0.8012 | -1.8205 | 0.78 |
| 10346 | *PPEF1* | 1.0245 | 1.0122 | 0.29337 | -0.66353 | 0.78 |
| 10347 | *CD9* | 1.6089 | 0.50517 | 0.21583 | -1.4185 | 0.78 |
| 10348 | *TPSAB1* | 1.7904 | 0.39161 | 0.1469 | -0.95461 | 0.78 |
| 10349 | *ALDH1L1* | 3.2387 | -0.41748 | -0.49241 | -2.2385 | 0.78 |
| 10350 | *ZDHHC23* | 3.2859 | -0.18547 | -0.77185 | -1.0564 | 0.78 |
| 10351 | *BPIFA1* | 3.1265 | 0.35466 | -1.1529 | -1.3336 | 0.78 |
| 10352 | *C17orf74* | 3.5832 | 0.056101 | -1.3114 | -2.5764 | 0.78 |
| 10353 | *ISG15* | 1.5932 | 1.3654 | -0.63125 | -1.8076 | 0.78 |
| 10354 | *CACTIN* | 4.2852 | -0.92339 | -1.0354 | -2.0134 | 0.78 |
| 10355 | *TSPAN3* | 2.5941 | 0.27663 | -0.54465 | -1.5382 | 0.78 |
| 10356 | *CENPW* | 4.7051 | -0.64944 | -1.7299 | -2.112 | 0.78 |
| 10357 | *PNMA5* | 1.4454 | 0.60772 | 0.27244 | -0.52917 | 0.78 |
| 10358 | *GPR146* | 3.6666 | -0.26093 | -1.0803 | -2.7416 | 0.78 |
| 10359 | *NUDT2* | 1.7789 | 0.41266 | 0.13367 | -1.7373 | 0.78 |
| 10360 | *ELK3* | 1.9592 | 0.28077 | 0.08526 | -0.49347 | 0.78 |
| 10361 | *RABAC1* | 1.9404 | 0.51563 | -0.13218 | -2.1134 | 0.77 |
| 10362 | *ATP2A1* | 2.9209 | 0.45067 | -1.0485 | -1.368 | 0.77 |
| 10363 | *IPO11* | 5.4226 | -1.4112 | -1.6885 | -2.4456 | 0.77 |
| 10364 | *CTNNBIP1* | 3.4427 | 0.008243 | -1.1282 | -1.4541 | 0.77 |
| 10365 | *ANKRD54* | 0.94214 | 0.85405 | 0.5251 | -1.658 | 0.77 |
| 10366 | *PRKRA* | 1.234 | 0.75099 | 0.3359 | 0.079335 | 0.77 |
| 10367 | *CUL7* | 3.2959 | -0.16245 | -0.81272 | -1.0581 | 0.77 |
| 10368 | *ZSCAN25* | 2.2769 | 0.32666 | -0.28348 | -0.31252 | 0.77 |
| 10369 | *SYNE2* | 0.87217 | 0.82828 | 0.61886 | -0.4573 | 0.77 |
| 10370 | *CASP7* | 3.9051 | 0.15433 | -1.7407 | -1.9495 | 0.77 |
| 10371 | *TTC17* | 1.7117 | 0.49843 | 0.10845 | -0.44353 | 0.77 |
| 10372 | *SETD7* | 2.8948 | 0.089763 | -0.66696 | -1.4388 | 0.77 |
| 10373 | *MIF4GD* | 1.0762 | 1.0349 | 0.20614 | -1.1113 | 0.77 |
| 10374 | *CCDC51* | 3.1432 | -0.39872 | -0.4273 | -2.7509 | 0.77 |
| 10375 | *MASTL* | 3.9421 | -0.47672 | -1.1492 | -1.4185 | 0.77 |
| 10376 | *ECE2* | 2.4829 | 0.021801 | -0.18978 | -0.99699 | 0.77 |
| 10377 | *ST8SIA4* | 3.242 | 0.61104 | -1.5383 | -1.6158 | 0.77 |
| 10378 | *ACY3* | 4.8338 | -1.2008 | -1.3184 | -1.5493 | 0.77 |
| 10379 | *ZNF483* | 2.8293 | 0.6123 | -1.1275 | -2.3121 | 0.77 |
| 10380 | *DDI2* | 2.6504 | -0.05445 | -0.28335 | -1.1488 | 0.77 |
| 10381 | *FAF2* | 3.8488 | -0.6202 | -0.91807 | -1.4681 | 0.77 |
| 10382 | *TRIM14* | 2.5264 | 0.46572 | -0.68192 | -1.9321 | 0.77 |
| 10383 | *COTL1* | 3.3776 | 0.34142 | -1.4089 | -2.0645 | 0.77 |
| 10384 | *CKAP2* | 2.6439 | 0.29378 | -0.62792 | -1.0567 | 0.77 |
| 10385 | *KLK10* | 1.3456 | 1.0208 | -0.05715 | -1.4016 | 0.77 |
| 10386 | *MDFI* | 1.9393 | 0.32059 | 0.04779 | -1.3203 | 0.77 |
| 10387 | *RIOK2* | 3.1839 | 0.22609 | -1.1034 | -1.6306 | 0.77 |
| 10388 | *GOLGA6C* | 3.8896 | -0.77396 | -0.8098 | -1.4584 | 0.77 |
| 10389 | *LPO* | 1.8337 | 0.23758 | 0.23385 | -3.2856 | 0.77 |

| | | | | | | |
|---|---|---|---|---|---|---|
| 10390 | *INCA1* | 1.3268 | 1.0566 | -0.07831 | -1.6071 | 0.77 |
| 10391 | *STRBP* | 2.2306 | 0.30193 | -0.22784 | -1.3034 | 0.77 |
| 10392 | *LAMB1* | 2.7461 | 0.44134 | -0.88278 | -0.97549 | 0.77 |
| 10393 | *USP29* | 2.6844 | 0.045563 | -0.42545 | -0.7513 | 0.77 |
| 10394 | *FOXC1* | 1.4373 | 1.1206 | -0.25387 | -0.51851 | 0.77 |
| 10395 | *STOML1* | 4.6545 | -0.90296 | -1.448 | -2.1342 | 0.77 |
| 10396 | *SMIM4* | 3.3709 | 0.14654 | -1.2141 | -2.0895 | 0.77 |
| 10397 | *POFUT1* | 2.7014 | 0.28917 | -0.6878 | -1.3931 | 0.77 |
| 10398 | *KRT33A* | 1.7872 | 1.2934 | -0.77853 | -1.7014 | 0.77 |
| 10399 | *GAPT* | 2.0807 | 0.6854 | -0.46404 | -1.8031 | 0.77 |
| 10400 | *SMIM22* | 2.0969 | 0.2171 | -0.01202 | -1.7125 | 0.77 |
| 10401 | *DGCR8* | 3.8845 | -0.51454 | -1.0682 | -2.508 | 0.77 |
| 10402 | *LRRC49* | 3.0962 | 1.1802 | -1.9748 | -2.3204 | 0.77 |
| 10403 | *MTFR2* | 2.0871 | 0.13367 | 0.08063 | -0.94912 | 0.77 |
| 10404 | *DUOX2* | 3.0691 | 0.55428 | -1.3221 | -2.1574 | 0.77 |
| 10405 | *ZNF654* | 4.3949 | -0.01938 | -2.0752 | -4.0172 | 0.77 |
| 10406 | *QDPR* | 3.1555 | -0.38393 | -0.47147 | -2.9638 | 0.77 |
| 10407 | *NKX2-5* | 2.2128 | 1.9665 | -1.8792 | -2.382 | 0.77 |
| 10408 | *PCDHA7* | 4.9612 | -1.2444 | -1.4168 | -2.3689 | 0.77 |
| 10409 | *GLYATL1* | 1.124 | 0.71441 | 0.46116 | -0.78592 | 0.77 |
| 10410 | *TOR1AIP1* | 4.7279 | -0.60083 | -1.8282 | -1.8759 | 0.77 |
| 10411 | *WWP1* | 3.5019 | -0.59763 | -0.60543 | -1.2465 | 0.77 |
| 10412 | *NDUFS7* | 3.227 | -0.13962 | -0.78963 | -1.4123 | 0.77 |
| 10413 | *MFSD6L* | 1.5384 | 0.7873 | -0.02799 | -0.5775 | 0.77 |
| 10414 | *C10orf120* | 3.1737 | 0.1136 | -0.98976 | -1.1826 | 0.77 |
| 10415 | *REC8* | 3.0241 | 0.53052 | -1.2573 | -2.3936 | 0.77 |
| 10416 | *TASP1* | 4.6 | -0.93489 | -1.3686 | -1.4234 | 0.77 |
| 10417 | *ARPC1A* | 3.9292 | -0.41967 | -1.2141 | -1.3199 | 0.77 |
| 10418 | *RNGTT* | 2.1735 | 0.48118 | -0.36023 | -1.2193 | 0.76 |
| 10419 | *GABRA2* | 3.7967 | -0.09975 | -1.4034 | -1.7945 | 0.76 |
| 10420 | *PHLDB2* | 4.5413 | -1.0039 | -1.2448 | -1.4919 | 0.76 |
| 10421 | *MAGEA9* | 2.9546 | 0.5894 | -1.2515 | -2.0952 | 0.76 |
| 10422 | *EPHA2* | 3.1766 | 0.092716 | -0.97716 | -1.197 | 0.76 |
| 10423 | *HLA-DOA* | 3.3724 | -0.22166 | -0.85931 | -1.0852 | 0.76 |
| 10424 | *RNF216* | 2.5263 | 0.23188 | -0.46748 | -0.7763 | 0.76 |
| 10425 | *MAPRE1* | 2.9515 | -0.18996 | -0.47116 | -1.8248 | 0.76 |
| 10426 | *THTPA* | 2.1898 | 1.184 | -1.0849 | -2.3452 | 0.76 |
| 10427 | *RPL17-C18orf32* | 3.8077 | -0.09142 | -1.4277 | -1.8109 | 0.76 |
| 10428 | *MAGEL2* | 2.5008 | -0.01099 | -0.20264 | -0.36394 | 0.76 |
| 10429 | *OGDHL* | 4.4222 | -0.60425 | -1.5308 | -1.671 | 0.76 |
| 10430 | *ZNF98* | 0.9713 | 0.86127 | 0.45457 | -1.1014 | 0.76 |
| 10431 | *KCNJ4* | 3.6205 | -0.49212 | -0.84195 | -1.2846 | 0.76 |
| 10432 | *PRR23D1* | 2.1571 | 1.4095 | -1.2802 | -2.2311 | 0.76 |
| 10433 | *IFI27L1* | 1.1925 | 1.1617 | -0.06838 | -0.94653 | 0.76 |
| 10434 | *ANKRD65* | 3.0723 | 0.16666 | -0.9532 | -1.7322 | 0.76 |
| 10435 | *USP54* | 3.9011 | -0.6122 | -1.0033 | -1.2792 | 0.76 |

| | | | | | | |
|---|---|---|---|---|---|---|
| 10436 | *ZNF709* | 2.5472 | -0.00422 | -0.25762 | -2.3877 | 0.76 |
| 10437 | *CLCA2* | 2.3211 | 0.77231 | -0.80812 | -1.1028 | 0.76 |
| 10438 | *CTSH* | 2.7602 | -0.06576 | -0.40976 | -2.2694 | 0.76 |
| 10439 | *GRIP1* | 4.2359 | -0.55773 | -1.3935 | -1.5235 | 0.76 |
| 10440 | *ST8SIA5* | 3.3157 | 0.56201 | -1.5941 | -1.7503 | 0.76 |
| 10441 | *LOC100505478* | 1.2296 | 0.7448 | 0.30903 | -0.00542 | 0.76 |
| 10442 | *CYP27C1* | 3.1058 | -0.18908 | -0.63355 | -0.78199 | 0.76 |
| 10443 | *LSM7* | 4.5996 | -0.60976 | -1.7074 | -2.2721 | 0.76 |
| 10444 | *DNAJC19* | 3.0347 | 0.61924 | -1.372 | -2.1187 | 0.76 |
| 10445 | *MARCO* | 2.0644 | 0.97641 | -0.75944 | -2.3557 | 0.76 |
| 10446 | *TBXA2R* | 2.3577 | 0.017123 | -0.09431 | -0.2424 | 0.76 |
| 10447 | *MACROD1* | 1.3549 | 0.97787 | -0.05226 | -0.81793 | 0.76 |
| 10448 | *PPARGC1A* | 3.5336 | 0.57784 | -1.8318 | -2.1541 | 0.76 |
| 10449 | *EXOC7* | 1.2693 | 1.0277 | -0.01774 | -1.8157 | 0.76 |
| 10450 | *TPRKB* | 2.9052 | -0.15922 | -0.46741 | -1.4616 | 0.76 |
| 10451 | *RTCA* | 2.6384 | 0.33037 | -0.69045 | -1.1251 | 0.76 |
| 10452 | *SERPINB11* | 1.2158 | 0.58065 | 0.48128 | -0.26779 | 0.76 |
| 10453 | *ACRC* | 2.6919 | 0.52555 | -0.93999 | -0.95184 | 0.76 |
| 10454 | *HSPB6* | 3.9257 | -0.5911 | -1.0576 | -1.4299 | 0.76 |
| 10455 | *HSDL2* | 4.2658 | 0.12307 | -2.112 | -2.5844 | 0.76 |
| 10456 | *FBXL8* | 3.5556 | -0.53114 | -0.74875 | -1.5548 | 0.76 |
| 10457 | *PCDHB7* | 2.8431 | 1.01 | -1.5789 | -1.6352 | 0.76 |
| 10458 | *ZNF189* | 1.6338 | 0.51107 | 0.12927 | -1.4423 | 0.76 |
| 10459 | *FAM71C* | 2.6682 | 0.33523 | -0.72936 | -1.0753 | 0.76 |
| 10460 | *SLC16A1* | 2.3653 | 0.042209 | -0.13357 | -1.6564 | 0.76 |
| 10461 | *SUSD2* | 2.3985 | 1.077 | -1.202 | -1.2302 | 0.76 |
| 10462 | *PRR36* | 1.6408 | 0.80931 | -0.17663 | -0.33133 | 0.76 |
| 10463 | *RASA4B* | 1.1529 | 1.0642 | 0.05627 | -1.1395 | 0.76 |
| 10464 | *KRTAP5-9* | 2.0273 | 1.2836 | -1.0376 | -4.134 | 0.76 |
| 10465 | *AIG1* | 3.8335 | -0.72725 | -0.83327 | -1.8631 | 0.76 |
| 10466 | *ZGPAT* | 1.4667 | 1.0639 | -0.25773 | -1.6455 | 0.76 |
| 10467 | *EMX1* | 2.5562 | -0.06903 | -0.21452 | -1.9133 | 0.76 |
| 10468 | *SEPT9* | 3.9003 | -0.58499 | -1.0433 | -2.2947 | 0.76 |
| 10469 | *CR1L* | 1.6363 | 0.47547 | 0.15986 | -0.06441 | 0.76 |
| 10470 | *VAMP2* | 1.4738 | 0.46987 | 0.32795 | -0.07273 | 0.76 |
| 10471 | *CBLN1* | 3.2949 | 0.15245 | -1.1759 | -1.9624 | 0.76 |
| 10472 | *ATP11B* | 2.2638 | 0.12189 | -0.11452 | -1.8615 | 0.76 |
| 10473 | *OR10V1* | 1.7072 | 1.0064 | -0.44353 | -2.0629 | 0.76 |
| 10474 | *KRTAP10-9* | 3.2704 | -0.25286 | -0.74755 | -1.0492 | 0.76 |
| 10475 | *CAPSL* | 3.521 | -0.25821 | -0.9929 | -3.3367 | 0.76 |
| 10476 | *PAQR4* | 1.3935 | 0.56342 | 0.31269 | 0.044791 | 0.76 |
| 10477 | *NVL* | 3.0473 | 0.14191 | -0.91968 | -1.2264 | 0.76 |
| 10478 | *SYT14* | 2.6364 | 0.30572 | -0.67265 | -1.0212 | 0.76 |
| 10479 | *BEAN1* | 2.3348 | 0.49557 | -0.56218 | -1.1256 | 0.76 |
| 10480 | *IL17RE* | 4.6003 | -0.8966 | -1.4356 | -2.1282 | 0.76 |
| 10481 | *PCDHGB6* | 2.9764 | 0.004801 | -0.71382 | -1.0677 | 0.76 |

| | | | | | | |
|---|---|---|---|---|---|---|
| 10482 | *ZCCHC8* | 2.0206 | 0.20314 | 0.04359 | -1.207 | 0.76 |
| 10483 | *ENKUR* | 2.741 | -0.14289 | -0.3328 | -1.9242 | 0.76 |
| 10484 | *HOXC12* | 2.4482 | 0.10825 | -0.29194 | -2.0102 | 0.75 |
| 10485 | *OPLAH* | 3.0453 | 0.14068 | -0.9215 | -1.205 | 0.75 |
| 10486 | *KRTAP12-2* | 2.7058 | 0.094786 | -0.53704 | -1.4823 | 0.75 |
| 10487 | *C11orf84* | 1.4269 | 0.96977 | -0.13392 | -1.257 | 0.75 |
| 10488 | *GTF3C3* | 3.3242 | 0.29391 | -1.3556 | -1.4736 | 0.75 |
| 10489 | *XRN1* | 3.5321 | -0.11437 | -1.1569 | -1.5337 | 0.75 |
| 10490 | *ABHD16B* | 2.9215 | 0.1106 | -0.77241 | -1.4299 | 0.75 |
| 10491 | *UPF2* | 2.439 | 0.019779 | -0.19969 | -0.56681 | 0.75 |
| 10492 | *PXN* | 3.7188 | -0.58305 | -0.87681 | -1.1017 | 0.75 |
| 10493 | *TMEM14E* | 1.4726 | 1.3784 | -0.59217 | -0.6683 | 0.75 |
| 10494 | *ZFAND1* | 2.3348 | 0.99791 | -1.0739 | -1.403 | 0.75 |
| 10495 | *MOS* | 4.6142 | -1.073 | -1.2827 | -2.5753 | 0.75 |
| 10496 | *LRRC10* | 3.0993 | 0.45939 | -1.3007 | -1.3604 | 0.75 |
| 10497 | *SERPINI2* | 1.2443 | 0.92761 | 0.0859 | -0.23531 | 0.75 |
| 10498 | *CHSY1* | 2.0827 | 1.1608 | -0.98572 | -2.4333 | 0.75 |
| 10499 | *DNAJC5* | 5.5677 | -1.3836 | -1.9269 | -5.6631 | 0.75 |
| 10500 | *GABRB3* | 1.599 | 0.68647 | -0.02943 | -1.7263 | 0.75 |
| 10501 | *SLC6A19* | 2.3458 | 0.26904 | -0.3589 | -1.5314 | 0.75 |
| 10502 | *SAMD12* | 3.427 | -0.57564 | -0.59581 | -1.962 | 0.75 |
| 10503 | *COA4* | 1.9336 | 0.57958 | -0.25767 | -1.3823 | 0.75 |
| 10504 | *OCIAD2* | 1.2361 | 0.81182 | 0.20724 | -0.27315 | 0.75 |
| 10505 | *RRM1* | 2.6745 | 0.058631 | -0.47862 | -0.63584 | 0.75 |
| 10506 | *TAF1A* | 1.0734 | 1.0671 | 0.11383 | -1.5138 | 0.75 |
| 10507 | *SERPINF2* | 1.9636 | 1.0611 | -0.77117 | -1.1075 | 0.75 |
| 10508 | *CUL9* | 4.1801 | -0.4628 | -1.4639 | -1.9308 | 0.75 |
| 10509 | *MGRN1* | 2.98 | -0.2562 | -0.47125 | -1.5611 | 0.75 |
| 10510 | *ABCC5* | 1.4967 | 1.2286 | -0.47304 | -1.6836 | 0.75 |
| 10511 | *PLEKHN1* | 4.9611 | -1.1665 | -1.5429 | -2.5655 | 0.75 |
| 10512 | *CHCHD2* | 2.7683 | 0.32334 | -0.84015 | -1.7122 | 0.75 |
| 10513 | *YPEL2* | 2.7027 | 1.067 | -1.5186 | -1.7954 | 0.75 |
| 10514 | *PEBP4* | 2.2536 | 1.6609 | -1.6635 | -1.8107 | 0.75 |
| 10515 | *HNRNPCL3* | 1.7744 | 0.37511 | 0.10111 | -0.53661 | 0.75 |
| 10516 | *USP53* | 2.849 | 0.004468 | -0.60289 | -1.25 | 0.75 |
| 10517 | *TAF5L* | 1.6873 | 0.35719 | 0.2048 | -0.49598 | 0.75 |
| 10518 | *CATSPER3* | 2.0556 | 0.15571 | 0.03686 | -0.31976 | 0.75 |
| 10519 | *C16orf90* | 1.3453 | 0.56824 | 0.33441 | -2.2075 | 0.75 |
| 10520 | *DEFB132* | 2.3372 | 0.58173 | -0.67149 | -0.69748 | 0.75 |
| 10521 | *PHGDH* | 1.8263 | 1.4315 | -1.0104 | -1.5913 | 0.75 |
| 10522 | *TRPC3* | 0.94689 | 0.90031 | 0.39967 | -1.6542 | 0.75 |
| 10523 | *SHB* | 2.8776 | 0.39634 | -1.0272 | -1.3162 | 0.75 |
| 10524 | *ITGAV* | 3.2762 | 0.18588 | -1.2159 | -1.9234 | 0.75 |
| 10525 | *GTPBP3* | 1.1573 | 0.56379 | 0.52455 | 0.047591 | 0.75 |
| 10526 | *NFASC* | 3.4622 | 0.060357 | -1.2774 | -1.6956 | 0.75 |
| 10527 | *IDS* | 0.83163 | 0.81238 | 0.60109 | -0.58554 | 0.75 |

| | | | | | | |
|---|---|---|---|---|---|---|
| 10528 | *LBX1* | 2.3495 | 0.26856 | -0.37343 | -0.54515 | 0.75 |
| 10529 | *KCNV2* | 1.4998 | 0.66554 | 0.07918 | 0.018683 | 0.75 |
| 10530 | *IKZF4* | 2.9339 | -0.20849 | -0.48169 | -1.0733 | 0.75 |
| 10531 | *LAMTOR4* | 3.3623 | 0.8165 | -1.9351 | -2.0661 | 0.75 |
| 10532 | *LIG4* | 3.7754 | 0.046811 | -1.5788 | -1.6617 | 0.75 |
| 10533 | *PRR19* | 4.1453 | -0.00831 | -1.8938 | -2.0062 | 0.75 |
| 10534 | *S100A10* | 3.5434 | 0.32485 | -1.626 | -2.1648 | 0.75 |
| 10535 | *SDHAF3* | 1.9468 | 0.3523 | -0.05738 | -0.67231 | 0.75 |
| 10536 | *GGT5* | 4.6395 | -0.44353 | -1.9548 | -2.2743 | 0.75 |
| 10537 | *POU2F1* | 2.5406 | 0.76167 | -1.0629 | -1.0989 | 0.75 |
| 10538 | *ZEB1* | 2.188 | 0.095742 | -0.04457 | -2.0725 | 0.75 |
| 10539 | *FLT3* | 2.0567 | 0.91797 | -0.73673 | -1.3089 | 0.75 |
| 10540 | *LRRC61* | 1.3456 | 1.1752 | -0.28384 | -1.5568 | 0.75 |
| 10541 | *ADCY4* | 2.7217 | -0.23929 | -0.24596 | -1.2135 | 0.75 |
| 10542 | *NIFK* | 2.328 | 0.23041 | -0.32237 | -1.8127 | 0.75 |
| 10543 | *HDGFRP3* | 2.3809 | 1.8729 | -2.0181 | -2.2044 | 0.75 |
| 10544 | *MSI2* | 1.7822 | 0.45832 | -0.00503 | -0.43756 | 0.75 |
| 10545 | *PAPSS1* | 1.4646 | 1.214 | -0.44353 | -1.0275 | 0.75 |
| 10546 | *ZRANB1* | 0.9894 | 0.81238 | 0.43322 | -0.49127 | 0.75 |
| 10547 | *FUT11* | 1.7917 | 0.7941 | -0.35134 | -2.8922 | 0.74 |
| 10548 | *NDST1* | 1.5351 | 1.1424 | -0.44353 | -0.61783 | 0.74 |
| 10549 | *CDCA3* | 3.2398 | 0.94595 | -1.9523 | -2.4215 | 0.74 |
| 10550 | *MSMB* | 1.2395 | 0.85245 | 0.14129 | -1.8714 | 0.74 |
| 10551 | *UAP1L1* | 3.9731 | -0.44498 | -1.2952 | -1.9417 | 0.74 |
| 10552 | *HOXB5* | 1.4487 | 1.4433 | -0.65919 | -2.1798 | 0.74 |
| 10553 | *OR2L13* | 4.1826 | -0.59045 | -1.3604 | -1.5153 | 0.74 |
| 10554 | *RPS6KB2* | 1.6854 | 0.3323 | 0.21388 | -0.06364 | 0.74 |
| 10555 | *MYCBP* | 3.3566 | 0.040889 | -1.1665 | -2.8007 | 0.74 |
| 10556 | *YPEL4* | 3.6255 | -0.30965 | -1.0858 | -2.0138 | 0.74 |
| 10557 | *FAM86B1* | 4.558 | -1.0467 | -1.2816 | -1.3399 | 0.74 |
| 10558 | *RPL22* | 1.2125 | 0.72339 | 0.29361 | -1.1814 | 0.74 |
| 10559 | *PSIP1* | 1.904 | 0.95444 | -0.62947 | -1.6796 | 0.74 |
| 10560 | *WNT11* | 1.1254 | 0.55491 | 0.5484 | 0.15515 | 0.74 |
| 10561 | *TPM4* | 3.0065 | 0.24242 | -1.0204 | -2.4312 | 0.74 |
| 10562 | *TIMM50* | 3.1032 | 0.70647 | -1.5817 | -3.1149 | 0.74 |
| 10563 | *PPP3R2* | 1.9976 | 0.12243 | 0.10724 | -0.58602 | 0.74 |
| 10564 | *XPNPEP2* | 1.5796 | 0.84092 | -0.1934 | -1.3451 | 0.74 |
| 10565 | *GAGE2D* | 3.4323 | -0.15039 | -1.0551 | -1.3457 | 0.74 |
| 10566 | *APOC3* | 2.8716 | 0.059357 | -0.70503 | -1.0936 | 0.74 |
| 10567 | *CAMSAP1* | 1.5442 | 0.40718 | 0.27427 | -0.10341 | 0.74 |
| 10568 | *C19orf67* | 2.3847 | -0.03335 | -0.1258 | -0.69961 | 0.74 |
| 10569 | *SMYD5* | 1.8978 | 1.5711 | -1.2448 | -2.3244 | 0.74 |
| 10570 | *TXNDC17* | 1.1318 | 0.91214 | 0.18015 | 0.03574 | 0.74 |
| 10571 | *PPP3CA* | 3.7019 | -0.69463 | -0.78345 | -1.5302 | 0.74 |
| 10572 | *PI4KB* | 2.0869 | 0.19793 | -0.0616 | -1.3437 | 0.74 |
| 10573 | *SFI1* | 2.13 | 0.61705 | -0.52447 | -1.711 | 0.74 |

| | | | | | | |
|---|---|---|---|---|---|---|
| 10574 | *PKP1* | 3.1271 | -0.20857 | -0.696 | -0.90387 | 0.74 |
| 10575 | *DCTPP1* | 3.0675 | 0.15514 | -1.0016 | -1.142 | 0.74 |
| 10576 | *CXorf51B* | 0.95551 | 0.70299 | 0.56243 | -0.73073 | 0.74 |
| 10577 | *CHN2* | 3.876 | -0.64398 | -1.0111 | -1.1076 | 0.74 |
| 10578 | *TRAPPC6B* | 2.7485 | 0.58786 | -1.1155 | -1.1912 | 0.74 |
| 10579 | *SMG6* | 2.097 | 0.19646 | -0.07297 | -1.8283 | 0.74 |
| 10580 | *PSMG3* | 3.4453 | -0.40785 | -0.8202 | -1.6993 | 0.74 |
| 10581 | *LOXHD1* | 1.8786 | 0.68258 | -0.34402 | -2.4333 | 0.74 |
| 10582 | *CYP2J2* | 3.5391 | -0.31807 | -1.0041 | -1.4043 | 0.74 |
| 10583 | *TTC5* | 1.3708 | 1.0236 | -0.17749 | -1.1944 | 0.74 |
| 10584 | *LILRA4* | 1.0324 | 0.71163 | 0.47269 | -2.1791 | 0.74 |
| 10585 | *SDF2* | 2.7299 | 0.078277 | -0.5915 | -1.9362 | 0.74 |
| 10586 | *SLC2A8* | 3.0398 | 0.73702 | -1.5604 | -1.7306 | 0.74 |
| 10587 | *CDK2AP2* | 3.0782 | -0.35746 | -0.50463 | -0.59791 | 0.74 |
| 10588 | *AK2* | 2.2904 | 1.7971 | -1.872 | -2.3488 | 0.74 |
| 10589 | *LOC101929983* | 3.0205 | -0.20874 | -0.59716 | -1.2234 | 0.74 |
| 10590 | *OPHN1* | 1.3451 | 0.66293 | 0.20649 | -0.09423 | 0.74 |
| 10591 | *TIMM8A* | 1.9066 | 0.27266 | 0.03497 | -0.91842 | 0.74 |
| 10592 | *FANCM* | 1.6392 | 0.57241 | 0.00226 | -1.3701 | 0.74 |
| 10593 | *CLP1* | 1.8326 | 0.86339 | -0.48231 | -1.3804 | 0.74 |
| 10594 | *ST8SIA1* | 2.6154 | 1.3089 | -1.7107 | -2.7849 | 0.74 |
| 10595 | *NACA2* | 3.2674 | -0.18229 | -0.87257 | -1.6571 | 0.74 |
| 10596 | *PKLR* | 1.5706 | 1.1761 | -0.53554 | -1.5719 | 0.74 |
| 10597 | *CRYAA* | 2.1228 | 0.73736 | -0.64982 | -1.3903 | 0.74 |
| 10598 | *WHSC1* | 3.2188 | 0.13705 | -1.146 | -2.5103 | 0.74 |
| 10599 | *WDR73* | 1.0584 | 0.90972 | 0.24167 | -1.8813 | 0.74 |
| 10600 | *TRIM77* | 3.6279 | -0.27421 | -1.1442 | -1.8267 | 0.74 |
| 10601 | *NKD2* | 2.9405 | -0.27631 | -0.45475 | -0.46752 | 0.74 |
| 10602 | *WNT8B* | 2.498 | 0.060229 | -0.34926 | -2.0351 | 0.74 |
| 10603 | *TMEM221* | 1.318 | 0.56804 | 0.32277 | -1.358 | 0.74 |
| 10604 | *ATP9A* | 3.1997 | -0.19915 | -0.79319 | -1.5633 | 0.74 |
| 10605 | *CDC42BPB* | 5.1189 | -1.0737 | -1.8382 | -2.0368 | 0.74 |
| 10606 | *APLP1* | 1.3795 | 0.61645 | 0.21028 | -0.49016 | 0.74 |
| 10607 | *DDX23* | 1.0246 | 0.91369 | 0.2678 | -0.61818 | 0.74 |
| 10608 | *RRP12* | 1.1602 | 1.046 | -0.00081 | -0.88561 | 0.74 |
| 10609 | *GLRA2* | 1.8761 | 1.1164 | -0.78728 | -1.5195 | 0.74 |
| 10610 | *ENOX1* | 3.4726 | -0.55676 | -0.71077 | -1.0796 | 0.74 |
| 10611 | *PPP2R5B* | 1.8479 | 1.093 | -0.73599 | -1.8908 | 0.73 |
| 10612 | *UBAP2L* | 2.5273 | 0.36907 | -0.69215 | -1.3517 | 0.73 |
| 10613 | *ELAC1* | 2.4024 | -0.05841 | -0.14048 | -0.77404 | 0.73 |
| 10614 | *CARD14* | 3.7104 | -0.48179 | -1.0258 | -1.95 | 0.73 |
| 10615 | *FGR* | 2.8115 | -0.03371 | -0.575 | -1.6599 | 0.73 |
| 10616 | *FBXO16* | 1.5232 | 1.1982 | -0.51886 | -1.4164 | 0.73 |
| 10617 | *DNAH6* | 2.0996 | 1.3594 | -1.2566 | -1.6524 | 0.73 |
| 10618 | *EHD3* | 3.413 | -0.30863 | -0.9027 | -1.2815 | 0.73 |
| 10619 | *EFHD2* | 2.0644 | 0.92295 | -0.78591 | -0.79392 | 0.73 |

| | | | | | | |
|---|---|---|---|---|---|---|
| 10620 | *ZNF324* | 2.0581 | 0.086332 | 0.05664 | -0.56598 | 0.73 |
| 10621 | *DMRTC1* | 1.3 | 0.57159 | 0.32946 | -1.5155 | 0.73 |
| 10622 | *ALPP* | 1.4167 | 0.96716 | -0.18283 | -1.3549 | 0.73 |
| 10623 | *C10orf32* | 1.9352 | 0.48891 | -0.22327 | -1.3231 | 0.73 |
| 10624 | *LOC158434* | 1.8914 | 1.4647 | -1.1553 | -2.9028 | 0.73 |
| 10625 | *POLB* | 3.7864 | -0.45012 | -1.1367 | -1.3875 | 0.73 |
| 10626 | *GUCY1A2* | 1.6915 | 0.8482 | -0.34037 | -0.36731 | 0.73 |
| 10627 | *NPHS1* | 3.9841 | -0.78591 | -0.99906 | -2.0538 | 0.73 |
| 10628 | *DRD3* | 1.7342 | 0.75036 | -0.28742 | -1.0889 | 0.73 |
| 10629 | *RBM18* | 3.2318 | 0.15901 | -1.1944 | -3.1836 | 0.73 |
| 10630 | *ZMYM3* | 2.3014 | 0.83796 | -0.94341 | -1.4448 | 0.73 |
| 10631 | *VAV2* | 1.3477 | 0.75766 | 0.09044 | -0.49095 | 0.73 |
| 10632 | *TCP11L2* | 2.4487 | 0.097322 | -0.35023 | -0.37629 | 0.73 |
| 10633 | *SLC52A3* | 2.2088 | 0.56958 | -0.58495 | -1.4841 | 0.73 |
| 10634 | *HOXA2* | 3.0306 | 0.47742 | -1.3148 | -1.5663 | 0.73 |
| 10635 | *ZNF764* | 5.2834 | -0.89186 | -2.1984 | -2.2879 | 0.73 |
| 10636 | *NCL* | 2.134 | 0.56844 | -0.50961 | -0.781 | 0.73 |
| 10637 | *GOLGA8J* | 1.7815 | 0.83556 | -0.42545 | -0.80594 | 0.73 |
| 10638 | *GDPD3* | 2.5946 | 1.0199 | -1.4234 | -1.8348 | 0.73 |
| 10639 | *SLC25A18* | 3.6215 | -0.6408 | -0.78966 | -1.739 | 0.73 |
| 10640 | *SLC16A12* | 2.0456 | 0.53271 | -0.38744 | -1.7595 | 0.73 |
| 10641 | *GGT6* | 2 | 0.75545 | -0.56461 | -0.87748 | 0.73 |
| 10642 | *ZFP69B* | 2.9932 | 0.51453 | -1.3199 | -1.3966 | 0.73 |
| 10643 | *C11orf73* | 2.8032 | 1.0756 | -1.6917 | -1.8792 | 0.73 |
| 10644 | *PRRT4* | 1.1402 | 0.74863 | 0.2981 | -1.7364 | 0.73 |
| 10645 | *FAM72A* | 2.1011 | 0.60353 | -0.5183 | -1.0954 | 0.73 |
| 10646 | *SPAG8* | 2.1774 | 0.075908 | -0.06698 | -0.78965 | 0.73 |
| 10647 | *BMI1* | 4.4632 | -0.99175 | -1.2852 | -1.6288 | 0.73 |
| 10648 | *PRPF6* | 1.688 | 0.89755 | -0.39932 | -1.0948 | 0.73 |
| 10649 | *TGFBR2* | 3.4332 | -0.07137 | -1.1758 | -1.6164 | 0.73 |
| 10650 | *WSCD2* | 2.4353 | -0.05219 | -0.19778 | -2.0488 | 0.73 |
| 10651 | *SMS* | 3.1745 | -0.3433 | -0.64639 | -2.1328 | 0.73 |
| 10652 | *WDR83OS* | 1.2223 | 0.57596 | 0.3861 | -1.3676 | 0.73 |
| 10653 | *KCNV1* | 1.8131 | 0.35814 | 0.01234 | -0.88279 | 0.73 |
| 10654 | *OR10R2* | 1.1235 | 1.0338 | 0.02626 | -0.70391 | 0.73 |
| 10655 | *KISS1* | 3.938 | -0.30339 | -1.4527 | -3.0804 | 0.73 |
| 10656 | *UNC13D* | 2.8828 | -0.23196 | -0.46973 | -2.0858 | 0.73 |
| 10657 | *HIST1H3C* | 2.7657 | 0.51931 | -1.1043 | -1.3721 | 0.73 |
| 10658 | *TTC24* | 4.9931 | -0.47211 | -2.3405 | -2.4961 | 0.73 |
| 10659 | *NUBPL* | 2.314 | 0.32643 | -0.46017 | -1.7407 | 0.73 |
| 10660 | *SLC38A2* | 2.6484 | 1.2379 | -1.7061 | -1.9075 | 0.73 |
| 10661 | *OCLN* | 2.2509 | -0.00437 | -0.0665 | -1.3666 | 0.73 |
| 10662 | *SPATA13* | 2.3113 | 0.34215 | -0.47381 | -2.9772 | 0.73 |
| 10663 | *DLGAP2* | 2.8046 | -0.00826 | -0.61682 | -1.1169 | 0.73 |
| 10664 | *PA2G4* | 1.2197 | 0.616 | 0.34374 | -1.7513 | 0.73 |
| 10665 | *CST6* | 2.3056 | 0.13558 | -0.26177 | -1.4829 | 0.73 |

| | | | | | | |
|---|---|---|---|---|---|---|
| 10666 | *ALPPL2* | 2.0211 | 0.23258 | -0.07439 | -0.56922 | 0.73 |
| 10667 | *NRGN* | 2.303 | 0.14733 | -0.27244 | -1.4276 | 0.73 |
| 10668 | *RFFL* | 1.6287 | 0.66682 | -0.11771 | -0.54058 | 0.73 |
| 10669 | *ATF4* | 2.377 | 0.24339 | -0.44277 | -0.44353 | 0.73 |
| 10670 | *EDN2* | 2.9472 | 0.23006 | -1.0005 | -1.8702 | 0.73 |
| 10671 | *CSAD* | 1.4586 | 0.46327 | 0.25333 | -0.21499 | 0.73 |
| 10672 | *OR6K3* | 2.5893 | 0.44838 | -0.86298 | -1.8227 | 0.72 |
| 10673 | *STK40* | 1.9869 | 0.43201 | -0.24519 | -0.70517 | 0.72 |
| 10674 | *COL6A2* | 1.8005 | 0.39495 | -0.02184 | -1.5467 | 0.72 |
| 10675 | *PTMS* | 1.36 | 0.84302 | -0.02943 | -0.1449 | 0.72 |
| 10676 | *ADAMTSL1* | 1.3052 | 0.87644 | -0.00826 | -0.31976 | 0.72 |
| 10677 | *EPT1* | 1.2849 | 0.62916 | 0.25904 | -0.91396 | 0.72 |
| 10678 | *ERAS* | 1.1894 | 0.58811 | 0.3946 | -0.26083 | 0.72 |
| 10679 | *GPR182* | 2.1368 | 0.026786 | 0.0073 | -1.1747 | 0.72 |
| 10680 | *SLC35F1* | 1.574 | 1.4306 | -0.83459 | -1.0475 | 0.72 |
| 10681 | *HCN3* | 1.9657 | 1.7647 | -1.5604 | -2.0952 | 0.72 |
| 10682 | *ANXA3* | 1.2378 | 0.63583 | 0.29569 | -2.2167 | 0.72 |
| 10683 | *ANK2* | 1.399 | 0.88455 | -0.11452 | -1.8068 | 0.72 |
| 10684 | *FEM1C* | 3.9448 | -0.82439 | -0.95184 | -1.049 | 0.72 |
| 10685 | *FGFRL1* | 1.8587 | 0.36774 | -0.05814 | -2.7245 | 0.72 |
| 10686 | *SEPN1* | 3.1024 | -0.35892 | -0.57564 | -2.1145 | 0.72 |
| 10687 | *RAB2A* | 2.8452 | 0.27879 | -0.95629 | -1.1325 | 0.72 |
| 10688 | *MED24* | 1.9693 | 0.28022 | -0.08248 | -0.80069 | 0.72 |
| 10689 | *NELL2* | 3.089 | 0.5897 | -1.5137 | -2.0146 | 0.72 |
| 10690 | *DEC1* | 1.6604 | 0.3623 | 0.14219 | -1.0323 | 0.72 |
| 10691 | *SMCHD1* | 1.2997 | 0.61432 | 0.2485 | -1.7044 | 0.72 |
| 10692 | *RBM48* | 2.4038 | 0.005991 | -0.24741 | -1.8507 | 0.72 |
| 10693 | *TOX4* | 3.1759 | -0.33404 | -0.68027 | -1.2061 | 0.72 |
| 10694 | *CTSS* | 1.8928 | 0.26025 | 0.00845 | -0.78588 | 0.72 |
| 10695 | *CDC34* | 2.0993 | 0.9191 | -0.85712 | -1.5775 | 0.72 |
| 10696 | *MGAT4D* | 3.1025 | -0.28431 | -0.65695 | -1.8038 | 0.72 |
| 10697 | *NHSL1* | 1.8381 | 1.2205 | -0.89741 | -1.677 | 0.72 |
| 10698 | *FASTKD5* | 2.8034 | 0.10266 | -0.745 | -1.5883 | 0.72 |
| 10699 | *SLC30A4* | 3.0494 | -0.01623 | -0.87328 | -1.3814 | 0.72 |
| 10700 | *ADARB2* | 1.0231 | 0.93431 | 0.20188 | -0.1207 | 0.72 |
| 10701 | *USP33* | 2.5795 | 0.50701 | -0.92905 | -1.5696 | 0.72 |
| 10702 | *PIGG* | 1.3661 | 0.95719 | -0.16624 | -2.4815 | 0.72 |
| 10703 | *TMEM67* | 2.6095 | 0.36252 | -0.8152 | -2.5571 | 0.72 |
| 10704 | *PHF14* | 3.5516 | -0.2671 | -1.1277 | -1.2195 | 0.72 |
| 10705 | *CANT1* | 3.1815 | 0.30181 | -1.3268 | -2.2593 | 0.72 |
| 10706 | *CRYGN* | 3.4959 | 0.049791 | -1.3903 | -1.8996 | 0.72 |
| 10707 | *SAP130* | 1.9436 | 1.244 | -1.0342 | -2.4576 | 0.72 |
| 10708 | *RARB* | 5.0722 | -0.91317 | -2.0062 | -2.0871 | 0.72 |
| 10709 | *ABT1* | 1.1969 | 1.1314 | -0.17597 | -1.4636 | 0.72 |
| 10710 | *EIF2B3* | 0.73531 | 0.73146 | 0.68528 | -1.2371 | 0.72 |
| 10711 | *DNAJC21* | 4.2711 | -0.78313 | -1.3364 | -1.9622 | 0.72 |

| | | | | | | |
|---|---|---|---|---|---|---|
| 10712 | *ZDHHC18* | 3.2022 | 0.15856 | -1.2094 | -2.1298 | 0.72 |
| 10713 | *OR8H2* | 1.4171 | 0.52142 | 0.21281 | -0.27159 | 0.72 |
| 10714 | *EIF3E* | 1.3986 | 0.69035 | 0.06199 | -0.2313 | 0.72 |
| 10715 | *PRPH* | 2.3608 | 0.78265 | -0.9929 | -1.0436 | 0.72 |
| 10716 | *RARA* | 1.6387 | 0.83056 | -0.31878 | -1.6095 | 0.72 |
| 10717 | *NOSIP* | 1.9326 | 0.30864 | -0.09119 | -1.3288 | 0.72 |
| 10718 | *TNFRSF18* | 1.1746 | 0.82307 | 0.1517 | -1.0013 | 0.72 |
| 10719 | *RHOT1* | 2.4573 | 0.20894 | -0.51764 | -3.1751 | 0.72 |
| 10720 | *ASB18* | 1.374 | 0.91767 | -0.1435 | -2.6313 | 0.72 |
| 10721 | *OR10G2* | 4.6463 | -0.78579 | -1.7129 | -2.462 | 0.72 |
| 10722 | *CILP2* | 3.2712 | 0.25132 | -1.3753 | -1.5176 | 0.72 |
| 10723 | *TCEANC2* | 2.6296 | 0.6556 | -1.1383 | -1.9622 | 0.72 |
| 10724 | *MAT2B* | 2.4373 | 0.48151 | -0.77241 | -1.9372 | 0.72 |
| 10725 | *COBLL1* | 3.5375 | -0.52469 | -0.86756 | -1.2141 | 0.72 |
| 10726 | *SMCO1* | 1.4666 | 0.51053 | 0.16756 | -1.3551 | 0.71 |
| 10727 | *C16orf95* | 2.1558 | 0.63221 | -0.6445 | -1.8581 | 0.71 |
| 10728 | *VWA3A* | 2.8419 | 0.26262 | -0.96206 | -2.1859 | 0.71 |
| 10729 | *STPG2* | 1.9492 | 0.32601 | -0.13354 | -0.52649 | 0.71 |
| 10730 | *UQCC2* | 2.036 | 0.39273 | -0.28734 | -0.75287 | 0.71 |
| 10731 | *TTC34* | 3.4692 | -0.15153 | -1.1765 | -1.2867 | 0.71 |
| 10732 | *FAM53A* | 2.8677 | -0.24896 | -0.4783 | -0.87681 | 0.71 |
| 10733 | *OR52E2* | 1.5657 | 0.67663 | -0.10288 | -2.8636 | 0.71 |
| 10734 | *ZCCHC18* | 1.429 | 0.53063 | 0.17824 | -0.87332 | 0.71 |
| 10735 | *SPG11* | 2.4136 | 0.4304 | -0.70685 | -0.77399 | 0.71 |
| 10736 | *F2RL1* | 3.2627 | -0.51363 | -0.61264 | -0.96492 | 0.71 |
| 10737 | *MAGEA11* | 3.1644 | 0.034674 | -1.063 | -1.5598 | 0.71 |
| 10738 | *NOP9* | 2.9647 | 0.05102 | -0.88008 | -2.4141 | 0.71 |
| 10739 | *KIFC1* | 2.1759 | 0.5224 | -0.56324 | -2.485 | 0.71 |
| 10740 | *GJB6* | 4.7014 | -0.92781 | -1.6386 | -1.7928 | 0.71 |
| 10741 | *LRRC4B* | 2.877 | -0.29546 | -0.44675 | -0.91335 | 0.71 |
| 10742 | *ILDR1* | 0.96695 | 0.66549 | 0.50216 | 0.46644 | 0.71 |
| 10743 | *CCND1* | 1.709 | 0.56162 | -0.13651 | -2.2167 | 0.71 |
| 10744 | *LYPD6B* | 3.3653 | -0.59028 | -0.64164 | -1.2864 | 0.71 |
| 10745 | *FAM98B* | 2.6602 | 0.86884 | -1.3961 | -2.2809 | 0.71 |
| 10746 | *BCL7B* | 1.0595 | 0.5928 | 0.48014 | -1.5809 | 0.71 |
| 10747 | *DCDC2C* | 2.4799 | 0.42265 | -0.77019 | -1.6979 | 0.71 |
| 10748 | *SURF6* | 3.3483 | -0.47476 | -0.74184 | -1.6796 | 0.71 |
| 10749 | *HKDC1* | 1.1253 | 0.58403 | 0.42221 | -0.51467 | 0.71 |
| 10750 | *CHST2* | 4.415 | -0.89941 | -1.3842 | -2.0452 | 0.71 |
| 10751 | *ROM1* | 3.3104 | -0.43292 | -0.74611 | -0.90149 | 0.71 |
| 10752 | *NTHL1* | 2.2265 | 0.031866 | -0.12725 | -0.55815 | 0.71 |
| 10753 | *LEMD2* | 3.0054 | -0.14406 | -0.73077 | -1.0585 | 0.71 |
| 10754 | *BIVM-ERCC5* | 1.5848 | 0.40132 | 0.14433 | 0.011164 | 0.71 |
| 10755 | *SPSB4* | 2.2114 | 0.10585 | -0.18699 | -0.51931 | 0.71 |
| 10756 | *FGF5* | 2.4902 | -0.0175 | -0.34308 | -0.98194 | 0.71 |
| 10757 | *KRTAP16-1* | 2.0071 | 0.69902 | -0.57681 | -0.71896 | 0.71 |

| | | | | | | |
|---|---|---|---|---|---|---|
| 10758 | *SCNN1D* | 3.6731 | -0.73517 | -0.80907 | -0.91982 | 0.71 |
| 10759 | *NKAIN4* | 1.1966 | 0.66293 | 0.26888 | -0.69325 | 0.71 |
| 10760 | *ERN1* | 1.9628 | 0.49689 | -0.33148 | -0.44353 | 0.71 |
| 10761 | *FEZ1* | 2.7238 | 0.30989 | -0.90652 | -1.2667 | 0.71 |
| 10762 | *GGT7* | 2.9339 | -0.40312 | -0.40398 | -2.2456 | 0.71 |
| 10763 | *FAM71A* | 1.3187 | 0.7847 | 0.02334 | -0.04564 | 0.71 |
| 10764 | *ABCC9* | 1.4717 | 0.57601 | 0.07828 | -0.23796 | 0.71 |
| 10765 | *TNPO1* | 3.9758 | -0.34227 | -1.5099 | -1.8952 | 0.71 |
| 10766 | *CLDN11* | 2.6681 | -0.271 | -0.2735 | -1.4584 | 0.71 |
| 10767 | *LRRC37A2* | 0.91814 | 0.77708 | 0.42745 | -0.19975 | 0.71 |
| 10768 | *PTGER3* | 1.6557 | 0.5484 | -0.08217 | -1.9397 | 0.71 |
| 10769 | *OXSM* | 2.7274 | 0.30691 | -0.91429 | -1.8338 | 0.71 |
| 10770 | *SCUBE2* | 2.3678 | 0.30747 | -0.55541 | -1.1637 | 0.71 |
| 10771 | *TROVE2* | 2.402 | -0.10835 | -0.17383 | -0.78965 | 0.71 |
| 10772 | *TIMM22* | 4.6784 | -1.2079 | -1.3507 | -2.3204 | 0.71 |
| 10773 | *CXorf51A* | 2.3881 | 0.18947 | -0.45847 | -0.82974 | 0.71 |
| 10774 | *RNF40* | 3.1367 | 0.056625 | -1.0757 | -2.8019 | 0.71 |
| 10775 | *LENG9* | 1.6846 | 0.42831 | 0.00368 | -0.74331 | 0.71 |
| 10776 | *RFXAP* | 2.2698 | 0.30207 | -0.45544 | -1.2411 | 0.71 |
| 10777 | *OR13J1* | 3.4841 | -0.44353 | -0.92415 | -2.3707 | 0.71 |
| 10778 | *BIRC8* | 1.2643 | 0.75092 | 0.10118 | -0.33985 | 0.71 |
| 10779 | *FRMPD4* | 2.5368 | -0.06121 | -0.35946 | -0.80594 | 0.71 |
| 10780 | *SULT1C2* | 2.7831 | 0.35036 | -1.018 | -1.7591 | 0.71 |
| 10781 | *SGSM3* | 1.9665 | 1.0225 | -0.87429 | -2.5901 | 0.70 |
| 10782 | *SELENBP1* | 2.866 | 0.093836 | -0.84595 | -1.0884 | 0.70 |
| 10783 | *DUS1L* | 4.0338 | -0.44099 | -1.4803 | -1.8441 | 0.70 |
| 10784 | *TMEM100* | 4.3308 | -0.51995 | -1.6984 | -2.3557 | 0.70 |
| 10785 | *CYSLTR1* | 1.5529 | 0.48201 | 0.0774 | -0.61682 | 0.70 |
| 10786 | *TMED2* | 2.4347 | 0.7361 | -1.0586 | -1.134 | 0.70 |
| 10787 | *UGT1A8* | 2.2118 | 0.009703 | -0.11119 | -1.1977 | 0.70 |
| 10788 | *SLC2A14* | 1.5007 | 0.83896 | -0.23034 | -0.99871 | 0.70 |
| 10789 | *IFI44* | 1.9312 | 0.25525 | -0.07822 | -1.2684 | 0.70 |
| 10790 | *EPB41L3* | 1.0123 | 0.82489 | 0.27022 | -0.35718 | 0.70 |
| 10791 | *IL20RA* | 3.3019 | -0.04172 | -1.1529 | -1.333 | 0.70 |
| 10792 | *BEST4* | 1.8928 | 0.36404 | -0.14994 | -0.87128 | 0.70 |
| 10793 | *LMTK2* | 1.9691 | 0.22358 | -0.08618 | -1.2852 | 0.70 |
| 10794 | *TYRO3* | 2.3478 | 0.44266 | -0.68453 | -1.9099 | 0.70 |
| 10795 | *RPL26* | 3.0495 | -0.36877 | -0.57546 | -2.1117 | 0.70 |
| 10796 | *ZNF467* | 0.81473 | 0.66112 | 0.62883 | -0.19183 | 0.70 |
| 10797 | *GRPEL2* | 3.8919 | -0.79108 | -0.99694 | -1.0654 | 0.70 |
| 10798 | *PPP1R1C* | 2.7596 | 0.53774 | -1.1944 | -1.9068 | 0.70 |
| 10799 | *GET4* | 2.8258 | -0.08975 | -0.63348 | -2.7016 | 0.70 |
| 10800 | *PHF3* | 3.4143 | 0.050668 | -1.3627 | -1.8033 | 0.70 |
| 10801 | *ERGIC2* | 2.1204 | 0.42502 | -0.44353 | -0.81329 | 0.70 |
| 10802 | *RAB22A* | 1.5667 | 0.29221 | 0.24249 | -0.29798 | 0.70 |
| 10803 | *DEFB131* | 2.3459 | 0.18009 | -0.42545 | -1.1081 | 0.70 |

| | | | | | | |
|---|---|---|---|---|---|---|
| 10804 | *AZGP1* | 3.4506 | 0.047153 | -1.3973 | -1.5831 | 0.70 |
| 10805 | *TIAF1* | 1.5675 | 0.41298 | 0.11874 | -0.41642 | 0.70 |
| 10806 | *FCGR2A* | 1.4002 | 0.71902 | -0.02022 | -0.97123 | 0.70 |
| 10807 | *PRX* | 2.8935 | -0.2408 | -0.55371 | -1.1316 | 0.70 |
| 10808 | *SPINT3* | 4.1234 | -0.59716 | -1.4273 | -1.5672 | 0.70 |
| 10809 | *MPP1* | 2.2779 | 0.84201 | -1.021 | -1.4994 | 0.70 |
| 10810 | *HTR3E* | 1.367 | 0.89372 | -0.16249 | -0.29713 | 0.70 |
| 10811 | *C17orf67* | 3.621 | -0.58451 | -0.93837 | -0.93878 | 0.70 |
| 10812 | *PTP4A2* | 2.093 | 0.47131 | -0.46623 | -0.55927 | 0.70 |
| 10813 | *HOMEZ* | 0.8032 | 0.73902 | 0.55522 | -1.4688 | 0.70 |
| 10814 | *MCFD2* | 1.2626 | 0.97328 | -0.13903 | -0.29232 | 0.70 |
| 10815 | *CPEB1* | 3.2785 | 0.41241 | -1.5949 | -2.0995 | 0.70 |
| 10816 | *FMO4* | 1.5934 | 0.37149 | 0.13016 | -0.24829 | 0.70 |
| 10817 | *NBPF3* | 2.9762 | -0.11759 | -0.76389 | -1.4789 | 0.70 |
| 10818 | *YWHAB* | 3.4907 | -0.66731 | -0.72877 | -1.7702 | 0.70 |
| 10819 | *MKS1* | 2.2153 | 0.93545 | -1.0564 | -1.5201 | 0.70 |
| 10820 | *TMEM150C* | 2.3551 | -0.09142 | -0.16944 | -1.4415 | 0.70 |
| 10821 | *PCP2* | 0.89844 | 0.66958 | 0.52466 | 0.21805 | 0.70 |
| 10822 | *MYOZ1* | 1.6782 | 0.48958 | -0.07561 | -0.84337 | 0.70 |
| 10823 | *CT47A4* | 2.8987 | 0.12297 | -0.92956 | -1.8297 | 0.70 |
| 10824 | *FAM151A* | 3.3447 | -0.4046 | -0.84841 | -1.8856 | 0.70 |
| 10825 | *DHDDS* | 1.4748 | 1.195 | -0.5788 | -0.88272 | 0.70 |
| 10826 | *R3HDML* | 4.5451 | -0.95321 | -1.5022 | -1.7322 | 0.70 |
| 10827 | *ADCY5* | 0.93097 | 0.75357 | 0.40349 | -0.63399 | 0.70 |
| 10828 | *ARHGAP44* | 1.8907 | 0.78149 | -0.58611 | -1.456 | 0.70 |
| 10829 | *GIN1* | 1.0749 | 0.84029 | 0.169 | -0.01111 | 0.69 |
| 10830 | *ZNF710* | 1.4751 | 0.78283 | -0.17383 | -0.49635 | 0.69 |
| 10831 | *ZNF493* | 0.73626 | 0.73253 | 0.61455 | -1.0134 | 0.69 |
| 10832 | *POLD4* | 2.8192 | -0.17635 | -0.55958 | -1.4725 | 0.69 |
| 10833 | *PEX11G* | 5.3862 | -1.4234 | -1.8804 | -2.3063 | 0.69 |
| 10834 | *KATNAL2* | 2.2656 | 0.73478 | -0.91913 | -1.0097 | 0.69 |
| 10835 | *MMP8* | 4.3948 | -0.54523 | -1.7709 | -2.2046 | 0.69 |
| 10836 | *WDR81* | 1.3187 | 0.39822 | 0.36147 | -1.4284 | 0.69 |
| 10837 | *ACO1* | 1.9945 | 0.14345 | -0.05985 | -2.0697 | 0.69 |
| 10838 | *FAM220A* | 3.3409 | 0.8408 | -2.1036 | -2.1347 | 0.69 |
| 10839 | *ADGRA1* | 2.9072 | -0.20318 | -0.62596 | -2.1648 | 0.69 |
| 10840 | *SAP25* | 3.0232 | -0.07202 | -0.87318 | -1.6525 | 0.69 |
| 10841 | *SHKBP1* | 2.6126 | -0.0749 | -0.46213 | -0.55015 | 0.69 |
| 10842 | *ADPRHL1* | 2.8187 | 0.2138 | -0.95708 | -0.95775 | 0.69 |
| 10843 | *MS4A7* | 2.3252 | 0.5013 | -0.75169 | -1.3955 | 0.69 |
| 10844 | *PPP1R15B* | 3.3141 | -0.60083 | -0.63939 | -0.95105 | 0.69 |
| 10845 | *RPRD2* | 2.7643 | -0.32677 | -0.36443 | -1.8656 | 0.69 |
| 10846 | *MED7* | 2.791 | 0.88044 | -1.5984 | -1.855 | 0.69 |
| 10847 | *ECI2* | 1.6426 | 0.41919 | 0.01102 | -0.02486 | 0.69 |
| 10848 | *AMT* | 3.3401 | 0.34728 | -1.615 | -1.9107 | 0.69 |
| 10849 | *ZNF862* | 1.7694 | 0.35153 | -0.04898 | -0.52106 | 0.69 |

| | | | | | | |
|---|---|---|---|---|---|---|
| 10850 | *UBE2D2* | 2.9323 | 0.13442 | -0.99508 | -1.251 | 0.69 |
| 10851 | *MMP3* | 3.7466 | -0.77018 | -0.90501 | -2.0277 | 0.69 |
| 10852 | *HLX* | 0.76394 | 0.69539 | 0.61199 | -2.1669 | 0.69 |
| 10853 | *NOBOX* | 0.78024 | 0.70154 | 0.58939 | -1.2018 | 0.69 |
| 10854 | *FOXD4L6* | 3.1668 | 0.68881 | -1.7845 | -2.5981 | 0.69 |
| 10855 | *KDM5D* | 1.1682 | 0.89872 | 0.00408 | -0.01352 | 0.69 |
| 10856 | *MCTS1* | 4.1778 | -1.0514 | -1.0561 | -1.2547 | 0.69 |
| 10857 | *TRMT1* | 1.9385 | 1.792 | -1.6604 | -2.2794 | 0.69 |
| 10858 | *MZF1* | 1.3913 | 0.54343 | 0.13481 | 0.028062 | 0.69 |
| 10859 | *CELA3A* | 2.3392 | 0.58419 | -0.85427 | -1.7577 | 0.69 |
| 10860 | *USP5* | 2.3304 | -0.04198 | -0.22069 | -1.0528 | 0.69 |
| 10861 | *LRP1* | 3.976 | -0.43998 | -1.4684 | -1.9567 | 0.69 |
| 10862 | *GPI* | 3.4563 | -0.12158 | -1.2694 | -1.6213 | 0.69 |
| 10863 | *UBALD1* | 1.8413 | 0.24605 | -0.02223 | -1.6907 | 0.69 |
| 10864 | *DLGAP4* | 1.6774 | 1.1515 | -0.76505 | -0.77284 | 0.69 |
| 10865 | *TNNC1* | 0.87135 | 0.65572 | 0.53623 | -0.47527 | 0.69 |
| 10866 | *DDHD2* | 1.8264 | 1.1126 | -0.8759 | -1.0491 | 0.69 |
| 10867 | *FCRLB* | 4.0831 | 0.001156 | -2.0218 | -2.2809 | 0.69 |
| 10868 | *STRA13* | 1.4676 | 1.1152 | -0.52055 | -0.90268 | 0.69 |
| 10869 | *RFX1* | 1.8364 | 0.48166 | -0.2561 | -1.794 | 0.69 |
| 10870 | *RGPD8* | 2.261 | 0.24325 | -0.44444 | -2.2948 | 0.69 |
| 10871 | *VGF* | 1.2105 | 1.1254 | -0.27755 | -0.27933 | 0.69 |
| 10872 | *GAGE1* | 1.5209 | 0.71498 | -0.17764 | -0.53036 | 0.69 |
| 10873 | *TSPAN17* | 3.0709 | 0.31054 | -1.3232 | -1.5212 | 0.69 |
| 10874 | *C1orf167* | 3.3471 | -0.16631 | -1.1226 | -1.4402 | 0.69 |
| 10875 | *NAGK* | 3.0571 | -0.42399 | -0.57513 | -1.8714 | 0.69 |
| 10876 | *TMSB4Y* | 2.2184 | 0.37005 | -0.53069 | -1.803 | 0.69 |
| 10877 | *FSIP1* | 3.2828 | 0.083089 | -1.3085 | -2.1072 | 0.69 |
| 10878 | *MORF4L2* | 3.8671 | -0.27453 | -1.5356 | -1.8751 | 0.69 |
| 10879 | *RANBP6* | 1.4486 | 1.1144 | -0.50668 | -2.0953 | 0.69 |
| 10880 | *NUP50* | 1.5952 | 1.0336 | -0.57296 | -0.75169 | 0.69 |
| 10881 | *PKHD1L1* | 1.2833 | 0.52691 | 0.2448 | -2.217 | 0.69 |
| 10882 | *BMS1* | 4.9779 | -0.93878 | -1.9846 | -2.1947 | 0.68 |
| 10883 | *SMOX* | 4.725 | -1.3111 | -1.3602 | -1.7662 | 0.68 |
| 10884 | *SCOC* | 1.7704 | 1.1617 | -0.8789 | -2.5655 | 0.68 |
| 10885 | *CHST15* | 2.8622 | 0.98804 | -1.7971 | -1.9479 | 0.68 |
| 10886 | *LYRM5* | 1.6596 | 0.87135 | -0.47911 | -0.87898 | 0.68 |
| 10887 | *GGCT* | 4.6474 | -1.2264 | -1.3694 | -2.1802 | 0.68 |
| 10888 | *EMX2* | 1.1465 | 1.04 | -0.13748 | -0.50978 | 0.68 |
| 10889 | *ROR2* | 3.4538 | -0.23223 | -1.1727 | -1.9919 | 0.68 |
| 10890 | *NEUROG1* | 1.6305 | 0.87812 | -0.46017 | -3.4924 | 0.68 |
| 10891 | *TSR1* | 2.3906 | 0.88146 | -1.2241 | -1.4221 | 0.68 |
| 10892 | *IQGAP3* | 1.0588 | 0.93435 | 0.05456 | -0.99888 | 0.68 |
| 10893 | *SNX11* | 1.3991 | 1.0071 | -0.35889 | -0.40467 | 0.68 |
| 10894 | *NUGGC* | 4.6713 | -1.2145 | -1.4102 | -2.1187 | 0.68 |
| 10895 | *HERC6* | 2.422 | 0.26252 | -0.63956 | -1.2051 | 0.68 |

| | | | | | | |
|---|---|---|---|---|---|---|
| 10896 | *RNF126* | 3.4066 | -0.4433 | -0.91862 | -1.1989 | 0.68 |
| 10897 | *OMD* | 4.894 | -1.3582 | -1.492 | -1.7029 | 0.68 |
| 10898 | *HHIPL2* | 1.0559 | 0.53068 | 0.45695 | 0.38259 | 0.68 |
| 10899 | *GLUL* | 1.6165 | 0.27912 | 0.14652 | -0.43543 | 0.68 |
| 10900 | *YBEY* | 1.6013 | 0.31697 | 0.12189 | -0.17554 | 0.68 |
| 10901 | *PSMC3IP* | 1.4999 | 0.83258 | -0.29327 | -0.64665 | 0.68 |
| 10902 | *PLOD1* | 3.1882 | -0.20166 | -0.94739 | -1.329 | 0.68 |
| 10903 | *GDF11* | 2.3851 | -0.16966 | -0.17643 | -1.4622 | 0.68 |
| 10904 | *SENP8* | 3.3178 | -0.62307 | -0.65695 | -0.92308 | 0.68 |
| 10905 | *BTBD10* | 2.6278 | -0.12995 | -0.46011 | -1.3617 | 0.68 |
| 10906 | *MCF2L2* | 0.97445 | 0.57144 | 0.49105 | -1.1977 | 0.68 |
| 10907 | *ABHD1* | 2.5883 | 0.35573 | -0.90713 | -1.1747 | 0.68 |
| 10908 | *KY* | 3.2097 | 0.44703 | -1.62 | -2.1796 | 0.68 |
| 10909 | *AMICA1* | 3.1958 | -0.24304 | -0.91612 | -0.95414 | 0.68 |
| 10910 | *C1orf111* | 2.0466 | 1.0358 | -1.0466 | -2.1224 | 0.68 |
| 10911 | *DNAJC8* | 3.9959 | -0.54094 | -1.4192 | -2.2088 | 0.68 |
| 10912 | *AHCYL1* | 3.8334 | -0.67137 | -1.1263 | -2.0629 | 0.68 |
| 10913 | *PIGR* | 2.1788 | 1.6705 | -1.8157 | -1.8645 | 0.68 |
| 10914 | *PDE4D* | 3.3015 | -0.43841 | -0.83005 | -1.987 | 0.68 |
| 10915 | *ADGRL2* | 0.85373 | 0.84271 | 0.33605 | -1.5865 | 0.68 |
| 10916 | *SALL3* | 2.6068 | 0.89315 | -1.4683 | -1.9727 | 0.68 |
| 10917 | *LCE1C* | 2.8802 | 0.054208 | -0.90518 | -1.4375 | 0.68 |
| 10918 | *MSX2* | 2.3226 | -0.12189 | -0.17172 | -1.9953 | 0.68 |
| 10919 | *ITGB5* | 1.5346 | 0.67147 | -0.17781 | -1.6565 | 0.68 |
| 10920 | *NFE2L3* | 1.6416 | 0.96132 | -0.57573 | -2.1445 | 0.68 |
| 10921 | *PPP1R1A* | 1.5071 | 0.69628 | -0.17622 | -1.0344 | 0.68 |
| 10922 | *CCNB1* | 5.2913 | -1.2845 | -1.9802 | -2.0687 | 0.68 |
| 10923 | *IFNGR2* | 1.9223 | 0.098389 | 0.00589 | -1.3198 | 0.68 |
| 10924 | *VKORC1* | 1.2329 | 0.72973 | 0.06374 | -1.6525 | 0.68 |
| 10925 | *CES4A* | 2.6632 | -0.04621 | -0.59168 | -1.0151 | 0.68 |
| 10926 | *PRPH2* | 1.4271 | 1.2777 | -0.68115 | -1.2109 | 0.67 |
| 10927 | *ADAM15* | 1.3325 | 0.86771 | -0.17721 | -0.81883 | 0.67 |
| 10928 | *PPP5C* | 4.1671 | -0.73483 | -1.4094 | -2.2948 | 0.67 |
| 10929 | *TTLL12* | 3.1261 | -0.0484 | -1.0557 | -1.8157 | 0.67 |
| 10930 | *SFT2D1* | 2.0908 | 0.50298 | -0.5718 | -1.2386 | 0.67 |
| 10931 | *JUND* | 1.8488 | 0.85619 | -0.68387 | -0.9722 | 0.67 |
| 10932 | *MPLKIP* | 1.0537 | 0.84427 | 0.12307 | -0.0236 | 0.67 |
| 10933 | *C8orf89* | 1.0091 | 0.71452 | 0.29731 | 0.24772 | 0.67 |
| 10934 | *NEUROG2* | 2.4266 | 0.59007 | -0.99574 | -1.3864 | 0.67 |
| 10935 | *LOC400736* | 1.7634 | 0.25978 | -0.00264 | -0.54121 | 0.67 |
| 10936 | *RNF122* | 2.9854 | 0.40145 | -1.3666 | -2.0146 | 0.67 |
| 10937 | *DISC1* | 2.1493 | 0.30113 | -0.43022 | -1.1531 | 0.67 |
| 10938 | *DCXR* | 1.3543 | 0.63608 | 0.02895 | -0.53775 | 0.67 |
| 10939 | *MTCH2* | 1.1094 | 0.74739 | 0.16192 | -0.46298 | 0.67 |
| 10940 | *LIG1* | 3.7741 | -0.48659 | -1.2697 | -1.5167 | 0.67 |
| 10941 | *MYCBPAP* | 4.8641 | -0.99146 | -1.8557 | -2.5292 | 0.67 |

| | | | | | | |
|---|---|---|---|---|---|---|
| 10942 | *CDS1* | 1.5188 | 0.48991 | 0.00711 | -1.7882 | 0.67 |
| 10943 | *ALDH16A1* | 3.2961 | -0.17601 | -1.1043 | -1.1951 | 0.67 |
| 10944 | *MTMR7* | 1.5051 | 0.5186 | -0.00832 | -2.5461 | 0.67 |
| 10945 | *PSORS1C1* | 1.4856 | 0.86236 | -0.33321 | -0.50441 | 0.67 |
| 10946 | *TSSK2* | 2.6277 | 0.092625 | -0.70558 | -0.86267 | 0.67 |
| 10947 | *LMBRD1* | 1.2852 | 0.63036 | 0.09894 | -0.92086 | 0.67 |
| 10948 | *CFAP57* | 1.6153 | 0.79326 | -0.39481 | -1.1805 | 0.67 |
| 10949 | *ZNF286B* | 2.203 | 0.35709 | -0.54665 | -0.90279 | 0.67 |
| 10950 | *MAPKAPK2* | 2.405 | 0.11036 | -0.50246 | -0.51679 | 0.67 |
| 10951 | *LGI3* | 1.3368 | 0.92762 | -0.25227 | -2.3629 | 0.67 |
| 10952 | *GAD2* | 3.6253 | -0.57042 | -1.0444 | -1.4331 | 0.67 |
| 10953 | *SLC17A2* | 1.6179 | 0.2397 | 0.15281 | -1.4303 | 0.67 |
| 10954 | *RLIM* | 2.514 | 0.98748 | -1.4913 | -1.5754 | 0.67 |
| 10955 | *FAM117A* | 1.2228 | 0.44215 | 0.34522 | -1.2318 | 0.67 |
| 10956 | *MCF2L* | 1.1816 | 0.47688 | 0.35163 | -1.3633 | 0.67 |
| 10957 | *ADGRG5* | 1.139 | 0.7937 | 0.07702 | -2.4015 | 0.67 |
| 10958 | *DPYS* | 1.2718 | 0.72139 | 0.01577 | -2.1888 | 0.67 |
| 10959 | *TNFRSF1A* | 2.9001 | -0.26314 | -0.62807 | -1.2915 | 0.67 |
| 10960 | *PRSS35* | 4.6194 | -1.0796 | -1.5312 | -3.1458 | 0.67 |
| 10961 | *KHK* | 3.6231 | 0.67406 | -2.2888 | -4.3274 | 0.67 |
| 10962 | *FAM20A* | 0.97824 | 0.92294 | 0.10682 | -1.12 | 0.67 |
| 10963 | *NCDN* | 2.8459 | -0.17525 | -0.66274 | -1.7652 | 0.67 |
| 10964 | *ARL14* | 1.6521 | 0.97726 | -0.62154 | -2.1478 | 0.67 |
| 10965 | *PDZD7* | 1.8899 | 0.62398 | -0.50623 | -0.78439 | 0.67 |
| 10966 | *CEBPD* | 1.1911 | 0.52712 | 0.28939 | -1.1704 | 0.67 |
| 10967 | *PRSS41* | 1.3563 | 1.2836 | -0.63248 | -0.95445 | 0.67 |
| 10968 | *SLC1A3* | 1.0137 | 0.66215 | 0.33123 | -1.6932 | 0.67 |
| 10969 | *LOC102723859* | 2.7311 | 0.003835 | -0.72803 | -1.7528 | 0.67 |
| 10970 | *SKA3* | 1.5874 | 0.25924 | 0.15946 | -0.14145 | 0.67 |
| 10971 | *PCYT2* | 1.5275 | 0.69249 | -0.21392 | -0.73455 | 0.67 |
| 10972 | *DSTN* | 1.4704 | 0.32956 | 0.2058 | -0.78444 | 0.67 |
| 10973 | *PLEKHA6* | 1.8802 | 0.83564 | -0.71077 | -1.0614 | 0.67 |
| 10974 | *CINP* | 1.3901 | 0.82759 | -0.21291 | -1.3627 | 0.67 |
| 10975 | *GLS* | 3.6119 | -0.2227 | -1.3851 | -1.8816 | 0.67 |
| 10976 | *PTPN6* | 1.8217 | 0.82297 | -0.64096 | -2.9931 | 0.67 |
| 10977 | *TRPT1* | 2.2299 | -0.08813 | -0.13963 | -0.72941 | 0.67 |
| 10978 | *NOX4* | 1.0497 | 0.85887 | 0.09355 | -1.8568 | 0.67 |
| 10979 | *ANGPT1* | 2.6558 | 0.48551 | -1.1402 | -1.4127 | 0.67 |
| 10980 | *ITGB6* | 3.3416 | -0.64654 | -0.69399 | -1.28 | 0.67 |
| 10981 | *CDHR4* | 1.5618 | 0.54202 | -0.10315 | -0.27407 | 0.67 |
| 10982 | *CD160* | 1.9662 | 0.05644 | -0.02351 | -0.6408 | 0.67 |
| 10983 | *SOCS6* | 1.172 | 0.76704 | 0.05866 | -1.5393 | 0.67 |
| 10984 | *KCNK2* | 2.0848 | 1.0526 | -1.1402 | -1.3976 | 0.67 |
| 10985 | *RSBN1* | 2.4098 | 0.38242 | -0.79596 | -1.5635 | 0.67 |
| 10986 | *KRTAP4-11* | 1.0273 | 0.62858 | 0.34019 | -1.0091 | 0.67 |
| 10987 | *MAGEA10* | 0.96453 | 0.68993 | 0.34021 | -0.0881 | 0.66 |

| | | | | | | |
|---|---|---|---|---|---|---|
| 10988 | *LCMT1* | 0.7618 | 0.75688 | 0.47594 | -1.3689 | 0.66 |
| 10989 | *DOHH* | 1.8396 | 1.0106 | -0.85586 | -1.2833 | 0.66 |
| 10990 | *PRKAB2* | 2.8024 | 0.44187 | -1.251 | -1.2572 | 0.66 |
| 10991 | *HSD17B10* | 2.4118 | 0.20193 | -0.62047 | -2.5057 | 0.66 |
| 10992 | *C5orf64* | 2.8843 | -0.20512 | -0.68684 | -1.2352 | 0.66 |
| 10993 | *EMG1* | 3.3194 | 0.21608 | -1.5441 | -2.0882 | 0.66 |
| 10994 | *ESR2* | 2.6255 | 0.16047 | -0.79488 | -0.99249 | 0.66 |
| 10995 | *SPDYE5* | 3.0109 | -0.17719 | -0.84326 | -2.0414 | 0.66 |
| 10996 | *C9orf170* | 3.1502 | -0.38473 | -0.77597 | -1.8305 | 0.66 |
| 10997 | *RDH13* | 3.9384 | -0.90553 | -1.0442 | -1.5265 | 0.66 |
| 10998 | *CDC123* | 2.9984 | 0.18128 | -1.1912 | -1.5276 | 0.66 |
| 10999 | *OR10A5* | 2.7438 | -0.08107 | -0.67441 | -2.1357 | 0.66 |
| 11000 | *NOL3* | 0.95207 | 0.85505 | 0.18069 | -1.0318 | 0.66 |
| 11001 | *BRD7* | 0.79384 | 0.66953 | 0.52359 | -0.14722 | 0.66 |
| 11002 | *ZSCAN29* | 1.1161 | 0.73926 | 0.13144 | -1.3294 | 0.66 |
| 11003 | *SGIP1* | 1.8297 | 1.0774 | -0.92087 | -1.6105 | 0.66 |
| 11004 | *PALM3* | 1.8396 | 0.36895 | -0.22235 | -0.78229 | 0.66 |
| 11005 | *KRBA2* | 3.061 | 0.64372 | -1.7188 | -2.8174 | 0.66 |
| 11006 | *PLXNA3* | 1.7969 | 1.4897 | -1.3007 | -1.589 | 0.66 |
| 11007 | *HHIPL1* | 1.2004 | 0.93248 | -0.14701 | -0.55002 | 0.66 |
| 11008 | *FOXE1* | 4.0081 | -0.55492 | -1.4684 | -1.9938 | 0.66 |
| 11009 | *SLC6A12* | 2.5975 | 0.023861 | -0.6373 | -2.9068 | 0.66 |
| 11010 | *SFTPC* | 3.8616 | -0.8462 | -1.0328 | -1.3165 | 0.66 |
| 11011 | *ARPP21* | 2.4483 | -0.14396 | -0.32231 | -0.37494 | 0.66 |
| 11012 | *SMIM20* | 1.4344 | 1.219 | -0.67137 | -1.2027 | 0.66 |
| 11013 | *EIF2B4* | 3.6122 | -0.52061 | -1.1103 | -1.2312 | 0.66 |
| 11014 | *KRTAP10-2* | 1.4655 | 0.83745 | -0.32191 | -1.6564 | 0.66 |
| 11015 | *FAM206A* | 2.9682 | 0.49083 | -1.478 | -5.3559 | 0.66 |
| 11016 | *TELO2* | 3.2882 | -0.59772 | -0.7095 | -1.4624 | 0.66 |
| 11017 | *STAT4* | 4.0024 | -0.10524 | -1.9179 | -2.0904 | 0.66 |
| 11018 | *LYZL2* | 2.4993 | -0.02722 | -0.49564 | -2.0863 | 0.66 |
| 11019 | *LCN10* | 2.0178 | 0.38763 | -0.42923 | -0.85389 | 0.66 |
| 11020 | *SALL4* | 2.7624 | -0.20302 | -0.58349 | -0.72157 | 0.66 |
| 11021 | *EML1* | 1.4665 | 0.94074 | -0.4317 | -2.1834 | 0.66 |
| 11022 | *HSP90B1* | 2.6724 | -0.29255 | -0.40539 | -1.6597 | 0.66 |
| 11023 | *MSLN* | 2.4391 | -0.21258 | -0.25225 | -0.58336 | 0.66 |
| 11024 | *ZNF513* | 2.2057 | 1.1225 | -1.3547 | -2.0447 | 0.66 |
| 11025 | *S1PR3* | 3.6088 | -0.51893 | -1.1169 | -1.473 | 0.66 |
| 11026 | *PCNXL2* | 3.9201 | -0.79548 | -1.1521 | -1.4116 | 0.66 |
| 11027 | *CANX* | 1.3376 | 0.64786 | -0.01323 | -2.0653 | 0.66 |
| 11028 | *NANOS3* | 1.3665 | 0.34273 | 0.26279 | -1.7078 | 0.66 |
| 11029 | *INS-IGF2* | 3.5268 | -0.48911 | -1.0669 | -2.9354 | 0.66 |
| 11030 | *PMEPA1* | 1.8761 | 0.061507 | 0.03203 | -1.9935 | 0.66 |
| 11031 | *FOXN3* | 2.0952 | 0.59853 | -0.7241 | -1.3259 | 0.66 |
| 11032 | *CRELD2* | 2.4441 | 0.17873 | -0.65324 | -2.069 | 0.66 |
| 11033 | *C1QTNF9B-AS1* | 1.6659 | 0.4209 | -0.1178 | -1.5282 | 0.66 |

| | | | | | | |
|---|---|---|---|---|---|---|
| 11034 | *CAMKK1* | 1.2166 | 0.71224 | 0.04013 | -0.98983 | 0.66 |
| 11035 | *LRRC27* | 0.80421 | 0.59287 | 0.57091 | -0.61254 | 0.66 |
| 11036 | *NPHP1* | 2.8616 | -0.023 | -0.87096 | -3.4957 | 0.66 |
| 11037 | *MSANTD3-TMEFF1* | 0.91989 | 0.734 | 0.31235 | -0.49505 | 0.66 |
| 11038 | *OR8B8* | 1.3291 | 0.62527 | 0.01175 | -0.60388 | 0.66 |
| 11039 | *LOC101928436* | 1.1484 | 0.96942 | -0.15226 | -0.72059 | 0.66 |
| 11040 | *YOD1* | 2.9427 | 0.064978 | -1.0422 | -1.0424 | 0.66 |
| 11041 | *ELMO1* | 1.1107 | 0.60513 | 0.24963 | -0.64671 | 0.66 |
| 11042 | *IBSP* | 0.84181 | 0.60152 | 0.52169 | -1.18 | 0.66 |
| 11043 | *FLAD1* | 1.214 | 0.48689 | 0.264 | -1.7566 | 0.65 |
| 11044 | *CD247* | 2.8801 | 0.3663 | -1.2823 | -1.4926 | 0.65 |
| 11045 | *APRT* | 1.9362 | 0.23415 | -0.20626 | -0.86493 | 0.65 |
| 11046 | *PRR20A* | 1.0341 | 0.56483 | 0.36502 | -1.5663 | 0.65 |
| 11047 | *GPKOW* | 4.6663 | -1.0498 | -1.6528 | -2.1532 | 0.65 |
| 11048 | *DCAKD* | 2.8217 | 0.51644 | -1.3762 | -1.5826 | 0.65 |
| 11049 | *ZNF143* | 0.85686 | 0.73966 | 0.36468 | -1.9367 | 0.65 |
| 11050 | *SP8* | 1.7247 | 0.38315 | -0.14672 | -1.0271 | 0.65 |
| 11051 | *TMEM163* | 1.0963 | 0.54299 | 0.3214 | -0.30715 | 0.65 |
| 11052 | *KRT27* | 2.3902 | -0.15596 | -0.27379 | -1.0033 | 0.65 |
| 11053 | *HDAC6* | 1.5067 | 0.70146 | -0.24801 | -1.5687 | 0.65 |
| 11054 | *ANKRD2* | 1.8578 | 0.4005 | -0.29905 | -1.3751 | 0.65 |
| 11055 | *C2orf49* | 3.2961 | -0.24966 | -1.0872 | -1.3766 | 0.65 |
| 11056 | *NIPAL4* | 1.4963 | 0.58191 | -0.11939 | -2.103 | 0.65 |
| 11057 | *SERPINA12* | 3.5548 | -0.45016 | -1.1463 | -1.2444 | 0.65 |
| 11058 | *BLOC1S6* | 1.3988 | 0.32954 | 0.22978 | -0.80958 | 0.65 |
| 11059 | *CCER1* | 2.8738 | 0.18854 | -1.1043 | -1.3226 | 0.65 |
| 11060 | *NUP107* | 2.481 | 0.031866 | -0.55492 | -0.87328 | 0.65 |
| 11061 | *ADCY6* | 3.1408 | 0.087344 | -1.2705 | -1.4619 | 0.65 |
| 11062 | *PCDHB10* | 1.1023 | 0.80071 | 0.05429 | -1.1977 | 0.65 |
| 11063 | *NKIRAS1* | 2.4507 | -0.02041 | -0.4735 | -1.8066 | 0.65 |
| 11064 | *CLK4* | 1.7287 | 1.0991 | -0.87128 | -0.994 | 0.65 |
| 11065 | *IL1RL1* | 1.2622 | 1.1899 | -0.49563 | -0.54027 | 0.65 |
| 11066 | *DNAJC5B* | 3.3617 | -0.62881 | -0.77667 | -3.3058 | 0.65 |
| 11067 | *MTA2* | 1.371 | 0.38425 | 0.20086 | -1.8157 | 0.65 |
| 11068 | *MT2A* | 2.236 | 1.1931 | -1.4732 | -1.6243 | 0.65 |
| 11069 | *MAFF* | 3.6688 | -0.44353 | -1.2699 | -2.8111 | 0.65 |
| 11070 | *HLA-DQA1* | 1.6894 | 0.52817 | -0.26252 | -0.3589 | 0.65 |
| 11071 | *BRCA1* | 1.5189 | 0.96531 | -0.52944 | -1.2797 | 0.65 |
| 11072 | *PLA2G2D* | 1.8043 | 0.38309 | -0.23289 | -0.66434 | 0.65 |
| 11073 | *TLK2* | 2.5209 | 0.49769 | -1.0641 | -1.6664 | 0.65 |
| 11074 | *MESP2* | 1.3413 | 0.86176 | -0.24874 | -0.97167 | 0.65 |
| 11075 | *OR51B4* | 3.1051 | -0.14239 | -1.0092 | -2.0989 | 0.65 |
| 11076 | *C6orf7* | 2.6397 | 0.48764 | -1.1743 | -1.8642 | 0.65 |
| 11077 | *HSD17B3* | 3.0313 | -0.1262 | -0.95321 | -2.5153 | 0.65 |
| 11078 | *TPM1* | 3.1584 | 0.18055 | -1.3878 | -1.9801 | 0.65 |
| 11079 | *PRKCE* | 2.468 | 0.42253 | -0.93991 | -1.0502 | 0.65 |

| | | | | | | |
|---|---|---|---|---|---|---|
| 11080 | *USP32* | 3.2544 | -0.23015 | -1.0737 | -1.2568 | 0.65 |
| 11081 | *UQCRB* | 3.4362 | 0.069884 | -1.5558 | -2.5103 | 0.65 |
| 11082 | *KRTAP9-1* | 2.8626 | 0.34469 | -1.2572 | -1.4136 | 0.65 |
| 11083 | *SULT1C3* | 1.0073 | 0.96104 | -0.0185 | -0.19675 | 0.65 |
| 11084 | *APCDD1* | 1.291 | 0.3539 | 0.30457 | 0.25927 | 0.65 |
| 11085 | *CEP170* | 2.7232 | 0.29504 | -1.0688 | -1.2246 | 0.65 |
| 11086 | *IFNA13* | 3.4714 | -0.74015 | -0.78314 | -1.5264 | 0.65 |
| 11087 | *RBM17* | 4.1221 | -0.9105 | -1.2635 | -1.5404 | 0.65 |
| 11088 | *CDH18* | 1.8569 | 0.94904 | -0.858 | -1.0584 | 0.65 |
| 11089 | *TAB3* | 1.6342 | 0.19879 | 0.11423 | -0.52255 | 0.65 |
| 11090 | *CPA3* | 3.8913 | -0.69325 | -1.2509 | -1.7793 | 0.65 |
| 11091 | *LRTM1* | 2.5121 | 0.66217 | -1.2277 | -1.9179 | 0.65 |
| 11092 | *USE1* | 2.3563 | -0.01683 | -0.39331 | -1.2908 | 0.65 |
| 11093 | *SNX18* | 1.8654 | 0.042086 | 0.03839 | -0.28228 | 0.65 |
| 11094 | *TTC13* | 5.3573 | -1.2021 | -2.2094 | -2.9931 | 0.65 |
| 11095 | *ZNF322* | 2.0154 | 0.12367 | -0.19364 | -0.55764 | 0.65 |
| 11096 | *TRIM8* | 4.6276 | -1.04 | -1.6435 | -1.7658 | 0.65 |
| 11097 | *ZNF699* | 1.5342 | 0.45269 | -0.04308 | -1.2895 | 0.65 |
| 11098 | *GPER1* | 2.5906 | -0.27877 | -0.36821 | -1.3101 | 0.65 |
| 11099 | *OS9* | 2.5728 | 1.1637 | -1.7935 | -1.8364 | 0.65 |
| 11100 | *SNRNP48* | 2.8491 | 0.32631 | -1.2329 | -1.5795 | 0.65 |
| 11101 | *GRXCR2* | 4.0516 | -0.54501 | -1.5644 | -1.6368 | 0.65 |
| 11102 | *STT3B* | 3.8596 | -0.81515 | -1.1027 | -1.9321 | 0.65 |
| 11103 | *ASB9* | 3.2589 | -0.06657 | -1.2507 | -1.6731 | 0.65 |
| 11104 | *COL8A1* | 3.2163 | -0.57429 | -0.70244 | -1.8306 | 0.65 |
| 11105 | *CASC10* | 3.3999 | -0.09423 | -1.367 | -1.4289 | 0.65 |
| 11106 | *CD248* | 1.4535 | 0.27393 | 0.2099 | -0.50752 | 0.65 |
| 11107 | *RRAS2* | 3.5743 | -0.77534 | -0.86181 | -2.01 | 0.65 |
| 11108 | *MAGED1* | 1.0885 | 0.45861 | 0.38996 | -0.24055 | 0.65 |
| 11109 | *PLXNA1* | 2.0633 | 0.3822 | -0.50924 | -1.7845 | 0.65 |
| 11110 | *GLCE* | 2.6609 | 0.10103 | -0.82582 | -1.7466 | 0.65 |
| 11111 | *KIF21B* | 1.278 | 0.63205 | 0.02589 | -1.1907 | 0.65 |
| 11112 | *MISP* | 1.8791 | 0.7166 | -0.66008 | -0.71093 | 0.65 |
| 11113 | *CNTN1* | 2.4931 | 0.11577 | -0.67378 | -1.1974 | 0.65 |
| 11114 | *LLGL2* | 5.0216 | -0.74141 | -2.3452 | -2.6254 | 0.64 |
| 11115 | *DBNDD2* | 1.0044 | 0.53966 | 0.39053 | -1.1463 | 0.64 |
| 11116 | *ETAA1* | 1.1657 | 0.39713 | 0.37148 | -0.80771 | 0.64 |
| 11117 | *ZNF484* | 1.0967 | 0.60551 | 0.23201 | -1.6844 | 0.64 |
| 11118 | *SLC25A33* | 1.3533 | 0.55443 | 0.02541 | -2.5934 | 0.64 |
| 11119 | *SLC22A17* | 1.3658 | 1.1744 | -0.60959 | -2.0955 | 0.64 |
| 11120 | *PAXIP1* | 2.9542 | -0.43702 | -0.58822 | -0.97565 | 0.64 |
| 11121 | *UQCRH* | 2.0902 | 0.75889 | -0.92174 | -1.0908 | 0.64 |
| 11122 | *ARMCX6* | 2.025 | 0.2658 | -0.36412 | -1.06 | 0.64 |
| 11123 | *EFEMP2* | 3.4079 | -0.47282 | -1.0086 | -1.0191 | 0.64 |
| 11124 | *TP53BP1* | 1.7049 | 0.1233 | 0.09805 | -0.17597 | 0.64 |
| 11125 | *CT55* | 1.3969 | 0.95444 | -0.42524 | -0.52643 | 0.64 |

| | | | | | | |
|---|---|---|---|---|---|---|
| 11126 | *TANGO2* | 1.737 | 0.87673 | -0.68766 | -0.74768 | 0.64 |
| 11127 | *EFCAB3* | 3.3168 | 0.81982 | -2.2108 | -2.2456 | 0.64 |
| 11128 | *ARPC2* | 1.1498 | 0.46726 | 0.30819 | -0.77997 | 0.64 |
| 11129 | *ERP44* | 1.9726 | 1.0437 | -1.0912 | -1.3573 | 0.64 |
| 11130 | *TRAF3* | 2.5213 | 0.42168 | -1.0187 | -1.1084 | 0.64 |
| 11131 | *GOLGA8O* | 2.2015 | 0.30668 | -0.58403 | -1.9919 | 0.64 |
| 11132 | *RALA* | 2.575 | -0.00268 | -0.64851 | -1.6208 | 0.64 |
| 11133 | *ZSWIM6* | 1.1399 | 0.60184 | 0.18203 | -3.7736 | 0.64 |
| 11134 | *OXCT1* | 1.383 | 0.43228 | 0.10748 | -0.45572 | 0.64 |
| 11135 | *LRIT1* | 1.0849 | 0.5982 | 0.23887 | 0.10206 | 0.64 |
| 11136 | *C6orf203* | 2.2672 | 0.49542 | -0.84069 | -1.6435 | 0.64 |
| 11137 | *CD2* | 1.1739 | 0.88085 | -0.13339 | -1.8321 | 0.64 |
| 11138 | *PWP1* | 1.485 | 0.75512 | -0.31884 | -1.4385 | 0.64 |
| 11139 | *DCAF16* | 1.3021 | 0.32886 | 0.28896 | -1.0054 | 0.64 |
| 11140 | *MYSM1* | 2.3371 | 0.16551 | -0.58305 | -3.2419 | 0.64 |
| 11141 | *ARTN* | 3.6806 | -0.79788 | -0.96572 | -2.4112 | 0.64 |
| 11142 | *ZNF91* | 1.7282 | 1.0742 | -0.88547 | -1.42 | 0.64 |
| 11143 | *RGPD4* | 1.369 | 0.49817 | 0.04928 | -0.17013 | 0.64 |
| 11144 | *ATL3* | 3.4915 | -0.31808 | -1.2573 | -1.6769 | 0.64 |
| 11145 | *ATG4D* | 3.9089 | -0.17714 | -1.8157 | -2.072 | 0.64 |
| 11146 | *TSPY10* | 3.0744 | -0.39423 | -0.76457 | -1.7566 | 0.64 |
| 11147 | *FBXW5* | 2.3337 | 0.28617 | -0.70618 | -2.1523 | 0.64 |
| 11148 | *KAT6B* | 3.7577 | -0.37544 | -1.4686 | -1.8885 | 0.64 |
| 11149 | *PEG3* | 2.0321 | 0.99015 | -1.1096 | -1.6001 | 0.64 |
| 11150 | *MAGEB5* | 2.5052 | 0.53673 | -1.1298 | -1.522 | 0.64 |
| 11151 | *FABP7* | 2.4793 | 0.4888 | -1.0564 | -1.8033 | 0.64 |
| 11152 | *LY6E* | 1.2195 | 0.42698 | 0.26472 | -1.0986 | 0.64 |
| 11153 | *ATF7* | 3.0096 | -0.53947 | -0.55993 | -0.81943 | 0.64 |
| 11154 | *MAGOH* | 1.7762 | 1.4073 | -1.2737 | -1.4474 | 0.64 |
| 11155 | *MYADM* | 4.5038 | -0.76049 | -1.8341 | -2.0574 | 0.64 |
| 11156 | *HPS3* | 1.9287 | 0.21027 | -0.22977 | -0.94583 | 0.64 |
| 11157 | *OR1N1* | 1.2348 | 0.85468 | -0.18043 | -1.6143 | 0.64 |
| 11158 | *NCR1* | 3.6737 | -0.85296 | -0.91221 | -1.3542 | 0.64 |
| 11159 | *F2RL3* | 2.6218 | 1.0925 | -1.806 | -1.8999 | 0.64 |
| 11160 | *IFNA5* | 2.9411 | -0.16888 | -0.86403 | -1.8792 | 0.64 |
| 11161 | *CRYZ* | 1.7463 | 0.27022 | -0.10861 | -2.0476 | 0.64 |
| 11162 | *TK2* | 2.9962 | -0.1372 | -0.95119 | -1.9125 | 0.64 |
| 11163 | *PCDHGA10* | 1.2094 | 0.40298 | 0.29519 | -1.8523 | 0.64 |
| 11164 | *RBBP6* | 2.9965 | 0.16057 | -1.25 | -2.1068 | 0.64 |
| 11165 | *NUP210L* | 1.9951 | 0.94688 | -1.0358 | -1.4329 | 0.64 |
| 11166 | *C3orf38* | 2.3391 | 0.49378 | -0.92823 | -1.9953 | 0.63 |
| 11167 | *XIRP1* | 1.5726 | 0.32967 | 0.00158 | -1.4455 | 0.63 |
| 11168 | *OAS1* | 4.4055 | -1.2226 | -1.2792 | -1.4081 | 0.63 |
| 11169 | *KANK4* | 1.4199 | 0.69407 | -0.21079 | -0.81034 | 0.63 |
| 11170 | *A4GALT* | 3.6443 | -0.82897 | -0.91236 | -4.2477 | 0.63 |
| 11171 | *WDR55* | 0.99996 | 0.49335 | 0.4092 | -0.95842 | 0.63 |

| | | | | | | |
|---|---|---|---|---|---|---|
| 11172 | *TTC9* | 1.3308 | 1.2816 | -0.71009 | -0.83497 | 0.63 |
| 11173 | *FTMT* | 1.0703 | 0.79764 | 0.03435 | -0.03422 | 0.63 |
| 11174 | *PLAGL1* | 3.4298 | -0.5639 | -0.96463 | -1.0933 | 0.63 |
| 11175 | *RGS16* | 4.1995 | -0.70942 | -1.5901 | -1.9326 | 0.63 |
| 11176 | *YBX3* | 2.1931 | 1.2476 | -1.5408 | -1.7158 | 0.63 |
| 11177 | *TMC2* | 1.4328 | 1.0457 | -0.57899 | -1.5568 | 0.63 |
| 11178 | *CDH12* | 1.267 | 0.99902 | -0.36705 | -1.451 | 0.63 |
| 11179 | *CXCR4* | 1.3913 | 0.52992 | -0.02242 | -1.8877 | 0.63 |
| 11180 | *ZNF735* | 3.6384 | -0.53589 | -1.2051 | -1.6474 | 0.63 |
| 11181 | *C12orf45* | 4.4433 | -0.99604 | -1.55 | -1.8324 | 0.63 |
| 11182 | *ATP8B2* | 0.92266 | 0.72637 | 0.24793 | -0.94777 | 0.63 |
| 11183 | *MKX* | 2.8841 | 0.098319 | -1.0863 | -1.5073 | 0.63 |
| 11184 | *FAM19A5* | 4.2843 | -0.64632 | -1.742 | -2.3795 | 0.63 |
| 11185 | *OR56A4* | 1.4017 | 0.85667 | -0.36467 | -2.3178 | 0.63 |
| 11186 | *CAND2* | 2.4534 | 0.44249 | -1.0028 | -1.337 | 0.63 |
| 11187 | *PABPC1L2B* | 2.4747 | 0.48959 | -1.0718 | -1.2139 | 0.63 |
| 11188 | *C6orf10* | 0.95467 | 0.68352 | 0.25425 | -0.04036 | 0.63 |
| 11189 | *TMEM203* | 3.6932 | -0.60183 | -1.1995 | -1.2635 | 0.63 |
| 11190 | *AMER2* | 1.9725 | 0.39392 | -0.47485 | -1.072 | 0.63 |
| 11191 | *KIF24* | 3.4734 | -0.44353 | -1.1391 | -1.9844 | 0.63 |
| 11192 | *NECAB2* | 1.8029 | 0.06743 | 0.02021 | -0.10273 | 0.63 |
| 11193 | *INTS6* | 2.0989 | 0.35226 | -0.56079 | -1.5712 | 0.63 |
| 11194 | *GPR4* | 2.905 | 0.23041 | -1.2457 | -1.4181 | 0.63 |
| 11195 | *CYP51A1* | 2.9056 | -0.33657 | -0.68021 | -1.3096 | 0.63 |
| 11196 | *ZNF804A* | 4.1854 | -0.59662 | -1.7001 | -2.0791 | 0.63 |
| 11197 | *PDPK1* | 4.3047 | -1.1968 | -1.2195 | -1.7316 | 0.63 |
| 11198 | *C11orf58* | 3.3287 | -0.53254 | -0.90824 | -1.2159 | 0.63 |
| 11199 | *TFAP2E* | 0.9693 | 0.47553 | 0.44254 | -0.67546 | 0.63 |
| 11200 | *CMC4* | 3.861 | -0.58349 | -1.3905 | -2.0197 | 0.63 |
| 11201 | *TMEM5* | 1.9715 | 0.3605 | -0.44547 | -1.0068 | 0.63 |
| 11202 | *DDX59* | 1.1667 | 1.1422 | -0.42248 | -2.3435 | 0.63 |
| 11203 | *SLC7A11* | 1.7486 | 0.19445 | -0.05663 | -0.3744 | 0.63 |
| 11204 | *PCSK9* | 1.3598 | 0.62749 | -0.10176 | -0.1339 | 0.63 |
| 11205 | *SNX33* | 2.5597 | -0.02542 | -0.64888 | -1.6587 | 0.63 |
| 11206 | *NMI* | 2.0459 | 0.25529 | -0.41596 | -0.42369 | 0.63 |
| 11207 | *ITGB2* | 3.94 | -0.6764 | -1.3787 | -1.4206 | 0.63 |
| 11208 | *TMEM253* | 2.5835 | 0.2881 | -0.98749 | -1.0633 | 0.63 |
| 11209 | *FBXO24* | 3.2028 | -0.55278 | -0.76606 | -2.103 | 0.63 |
| 11210 | *ATRIP* | 1.5857 | 1.0157 | -0.71746 | -1.4501 | 0.63 |
| 11211 | *RABGGTB* | 2.7312 | -0.09828 | -0.75049 | -1.989 | 0.63 |
| 11212 | *PDLIM3* | 1.0691 | 0.87312 | -0.06044 | -1.1631 | 0.63 |
| 11213 | *PCDHB1* | 1.9243 | 0.55379 | -0.59703 | -1.1071 | 0.63 |
| 11214 | *LAMP3* | 1.4039 | 0.71221 | -0.23523 | -3.2089 | 0.63 |
| 11215 | *ZNF7* | 1.7542 | 0.25601 | -0.12957 | -0.46614 | 0.63 |
| 11216 | *CD36* | 2.0102 | 0.32484 | -0.45449 | -1.4385 | 0.63 |
| 11217 | *PARVB* | 2.6815 | 0.29391 | -1.0951 | -1.1114 | 0.63 |

| | | | | | | |
|---|---|---|---|---|---|---|
| 11218 | *ARHGEF17* | 2.8126 | -0.15013 | -0.78229 | -1.7585 | 0.63 |
| 11219 | *TAS2R43* | 2.613 | 0.15359 | -0.88702 | -1.2169 | 0.63 |
| 11220 | *MYPN* | 1.5658 | 0.96819 | -0.65503 | -1.2743 | 0.63 |
| 11221 | *TRIM43* | 1.6247 | 0.21814 | 0.03541 | -0.15476 | 0.63 |
| 11222 | *XK* | 1.3776 | 0.70759 | -0.2073 | -2.221 | 0.63 |
| 11223 | *ITGAM* | 2.2041 | 0.18203 | -0.50911 | -1.0947 | 0.63 |
| 11224 | *UQCRHL* | 2.7463 | -0.42564 | -0.44417 | -1.3366 | 0.63 |
| 11225 | *NUDT15* | 1.5169 | 0.74008 | -0.38161 | -1.4951 | 0.63 |
| 11226 | *CARS* | 1.7802 | 0.72606 | -0.63157 | -1.1075 | 0.62 |
| 11227 | *C11orf68* | 1.1929 | 1.0541 | -0.37275 | -1.0728 | 0.62 |
| 11228 | *PALMD* | 2.1438 | 0.82008 | -1.0898 | -1.1126 | 0.62 |
| 11229 | *FOXB2* | 1.0487 | 0.66663 | 0.15852 | -0.17745 | 0.62 |
| 11230 | *MAP2K5* | 2.3079 | -0.0706 | -0.36427 | -0.9879 | 0.62 |
| 11231 | *TMEM101* | 0.76637 | 0.57476 | 0.5309 | -0.96788 | 0.62 |
| 11232 | *POLA1* | 4.0439 | 0.01398 | -2.1859 | -2.6635 | 0.62 |
| 11233 | *TMPRSS11E* | 1.8712 | 0.3341 | -0.33336 | -0.43005 | 0.62 |
| 11234 | *GPATCH1* | 2.5413 | -0.24665 | -0.42373 | -2.8019 | 0.62 |
| 11235 | *C9orf85* | 1.1614 | 0.70917 | -4.4E-05 | -0.10184 | 0.62 |
| 11236 | *SPEF2* | 1.7297 | 0.082695 | 0.05813 | -0.33397 | 0.62 |
| 11237 | *URAD* | 2.5281 | 0.18586 | -0.84348 | -0.95658 | 0.62 |
| 11238 | *PDE12* | 1.3783 | 0.34219 | 0.14984 | -0.5687 | 0.62 |
| 11239 | *MPHOSPH8* | 2.3584 | -0.02915 | -0.45906 | -0.88274 | 0.62 |
| 11240 | *GREB1* | 1.998 | 0.40461 | -0.53254 | -1.2362 | 0.62 |
| 11241 | *MAGEB6* | 3.9122 | -0.2367 | -1.8057 | -2.2004 | 0.62 |
| 11242 | *PRSS23* | 2.4817 | 0.19661 | -0.80909 | -1.6403 | 0.62 |
| 11243 | *C19orf70* | 1.0136 | 0.86372 | -0.00826 | -1.5167 | 0.62 |
| 11244 | *ZGLP1* | 2.6967 | 0.2105 | -1.0393 | -1.0399 | 0.62 |
| 11245 | *MT4* | 2.5834 | -0.13072 | -0.58485 | -1.6289 | 0.62 |
| 11246 | *LRRC39* | 2.1375 | 0.61068 | -0.88098 | -2.1984 | 0.62 |
| 11247 | *FASLG* | 3.198 | -0.63753 | -0.69365 | -1.1536 | 0.62 |
| 11248 | *GOLIM4* | 1.6668 | 0.10682 | 0.09241 | -1.6763 | 0.62 |
| 11249 | *C1S* | 1.0246 | 0.45188 | 0.38954 | -0.49322 | 0.62 |
| 11250 | *TMEM135* | 1.5809 | 0.56676 | -0.28197 | -0.55492 | 0.62 |
| 11251 | *SSBP2* | 1.7844 | 0.91836 | -0.83718 | -2.4527 | 0.62 |
| 11252 | *SBK1* | 3.3208 | 0.13715 | -1.5949 | -2.7838 | 0.62 |
| 11253 | *ARL17B* | 3.9853 | -0.93474 | -1.1887 | -3.2616 | 0.62 |
| 11254 | *TM9SF3* | 1.173 | 0.70104 | -0.01292 | -0.59739 | 0.62 |
| 11255 | *FHL3* | 1.7782 | 0.079734 | 0.00312 | -0.31052 | 0.62 |
| 11256 | *SMNDC1* | 2.0842 | 0.91674 | -1.1407 | -2.1811 | 0.62 |
| 11257 | *CSN2* | 4.0184 | -1.0434 | -1.1152 | -1.5458 | 0.62 |
| 11258 | *VSIG1* | 1.9272 | 0.75503 | -0.82447 | -1.6641 | 0.62 |
| 11259 | *ARHGEF10* | 1.7925 | 0.73512 | -0.67029 | -1.1449 | 0.62 |
| 11260 | *NKG7* | 1.3614 | 0.35399 | 0.14194 | -0.40605 | 0.62 |
| 11261 | *NFX1* | 2.0501 | 0.25796 | -0.45119 | -1.6867 | 0.62 |
| 11262 | *HIST1H2BJ* | 4.1269 | -0.79611 | -1.4747 | -1.5054 | 0.62 |
| 11263 | *THEMIS2* | 1.6593 | 0.14294 | 0.05361 | -0.04952 | 0.62 |

| | | | | | | |
|---|---|---|---|---|---|---|
| 11264 | *MRPS31* | 3.088 | -0.53987 | -0.69287 | -0.78443 | 0.62 |
| 11265 | *CSTF2* | 1.2619 | 0.4209 | 0.17242 | -1.57 | 0.62 |
| 11266 | *PLCXD1* | 2.9019 | 0.62388 | -1.6711 | -2.672 | 0.62 |
| 11267 | *TMEM80* | 0.89643 | 0.6201 | 0.33774 | -0.84897 | 0.62 |
| 11268 | *MYL6* | 1.5776 | 0.16383 | 0.11254 | -1.2476 | 0.62 |
| 11269 | *RMDN3* | 3.5222 | -0.40398 | -1.2646 | -1.5671 | 0.62 |
| 11270 | *CTU1* | 2.7606 | 0.30735 | -1.2145 | -1.606 | 0.62 |
| 11271 | *PRSS1* | 2.7407 | -0.22589 | -0.66194 | -1.8443 | 0.62 |
| 11272 | *ELK1* | 3.6535 | -0.75974 | -1.0424 | -1.2936 | 0.62 |
| 11273 | *ANAPC11* | 3.2404 | -0.64961 | -0.73984 | -1.7228 | 0.62 |
| 11274 | *VPS26B* | 1.1764 | 0.56009 | 0.11435 | -1.6975 | 0.62 |
| 11275 | *SLC22A13* | 2.8792 | -0.24036 | -0.78823 | -2.0483 | 0.62 |
| 11276 | *PPM1D* | 1.9913 | 0.29421 | -0.435 | -1.1049 | 0.62 |
| 11277 | *PON1* | 1.1879 | 1.1061 | -0.44353 | -1.5138 | 0.62 |
| 11278 | *SFMBT1* | 2.23 | -0.17311 | -0.20771 | -1.2867 | 0.62 |
| 11279 | *ZKSCAN2* | 2.0265 | -0.07997 | -0.09757 | -1.77 | 0.62 |
| 11280 | *LRRFIP1* | 3.019 | 0.21771 | -1.3887 | -1.6982 | 0.62 |
| 11281 | *OR4K1* | 1.497 | 0.29421 | 0.0566 | -0.87318 | 0.62 |
| 11282 | *CLDN25* | 0.9357 | 0.93071 | -0.01938 | -2.075 | 0.62 |
| 11283 | *ZFP2* | 1.5732 | 1.1359 | -0.86223 | -1.2277 | 0.62 |
| 11284 | *RBM41* | 0.64116 | 0.62108 | 0.58461 | -1.1448 | 0.62 |
| 11285 | *STEAP4* | 0.90512 | 0.58434 | 0.3573 | -1.1962 | 0.62 |
| 11286 | *SLC27A3* | 2.112 | 0.37416 | -0.64033 | -1.1046 | 0.62 |
| 11287 | *GPR65* | 1.7252 | 0.36032 | -0.24036 | -1.2719 | 0.62 |
| 11288 | *RTKN2* | 1.9774 | -0.01352 | -0.11897 | -0.91913 | 0.61 |
| 11289 | *PARP8* | 1.4079 | 0.22239 | 0.21452 | -1.0985 | 0.61 |
| 11290 | *SCN3B* | 2.0778 | 0.21027 | -0.44353 | -1.0669 | 0.61 |
| 11291 | *NLRP2* | 3.3636 | -0.52811 | -0.99137 | -1.3125 | 0.61 |
| 11292 | *TTC23L* | 3.2732 | 0.15831 | -1.5884 | -2.8402 | 0.61 |
| 11293 | *FDX1* | 3.5636 | -0.2868 | -1.4337 | -1.479 | 0.61 |
| 11294 | *C16orf82* | 1.2292 | 1.0569 | -0.44353 | -1.6526 | 0.61 |
| 11295 | *TAS2R60* | 1.9048 | 1.0134 | -1.078 | -1.8963 | 0.61 |
| 11296 | *PSMA7* | 2.499 | -0.18074 | -0.47862 | -1.5171 | 0.61 |
| 11297 | *TMEM169* | 1.9219 | 0.62281 | -0.7061 | -1.7247 | 0.61 |
| 11298 | *FAM120B* | 1.0441 | 0.66344 | 0.13093 | -0.14548 | 0.61 |
| 11299 | *FAM24B* | 1.4105 | 0.86615 | -0.43845 | -0.62306 | 0.61 |
| 11300 | *DNALI1* | 2.2585 | 0.098877 | -0.5193 | -0.56403 | 0.61 |
| 11301 | *METTL20* | 2.1259 | 0.001801 | -0.28972 | -0.44198 | 0.61 |
| 11302 | *EWSR1* | 3.716 | -0.88905 | -0.98915 | -1.6524 | 0.61 |
| 11303 | *TNFAIP8L3* | 3.1096 | 0.50816 | -1.7813 | -1.916 | 0.61 |
| 11304 | *SERPING1* | 2.5592 | -0.10044 | -0.62306 | -0.94422 | 0.61 |
| 11305 | *KNCN* | 2.5057 | -0.27843 | -0.3921 | -0.60845 | 0.61 |
| 11306 | *P2RY11* | 1.323 | 0.54063 | -0.02974 | -1.144 | 0.61 |
| 11307 | *CABP5* | 1.7122 | 0.33657 | -0.21526 | -1.1641 | 0.61 |
| 11308 | *ZMYM6* | 0.87899 | 0.65764 | 0.29629 | -1.729 | 0.61 |
| 11309 | *PTGDR2* | 2.1938 | 0.48872 | -0.8504 | -1.0945 | 0.61 |

| | | | | | | |
|---|---|---|---|---|---|---|
| 11310 | *DAO* | 1.9338 | 0.4358 | -0.53775 | -1.3173 | 0.61 |
| 11311 | *TNP1* | 3.4568 | -0.78956 | -0.83627 | -1.8698 | 0.61 |
| 11312 | *DNAJC22* | 2.7662 | -0.08018 | -0.85531 | -1.6799 | 0.61 |
| 11313 | *MARVELD3* | 3.3909 | -0.301 | -1.2595 | -1.8398 | 0.61 |
| 11314 | *GPAT2* | 3.3098 | -0.16885 | -1.3112 | -1.3218 | 0.61 |
| 11315 | *PATZ1* | 1.1451 | 1.0922 | -0.4082 | -1.6626 | 0.61 |
| 11316 | *PRR14* | 2.3577 | 0.46888 | -0.99783 | -1.2076 | 0.61 |
| 11317 | *OR4D5* | 1.3485 | 0.2735 | 0.20651 | -0.31052 | 0.61 |
| 11318 | *UBXN8* | 3.1715 | -0.44353 | -0.89957 | -1.7395 | 0.61 |
| 11319 | *UCN3* | 1.1258 | 0.45741 | 0.24469 | -0.18432 | 0.61 |
| 11320 | *PGAP3* | 1.8759 | 0.12888 | -0.17746 | -0.96632 | 0.61 |
| 11321 | *PCDHB5* | 1.9869 | 0.20227 | -0.36295 | -0.67482 | 0.61 |
| 11322 | *TMPRSS4* | 2.0637 | 0.80675 | -1.0443 | -1.3377 | 0.61 |
| 11323 | *P4HTM* | 3.1883 | -0.10479 | -1.2576 | -2.5198 | 0.61 |
| 11324 | *AADAT* | 2.5362 | 0.086847 | -0.79875 | -1.0529 | 0.61 |
| 11325 | *MTFMT* | 1.1645 | 0.43614 | 0.22329 | -2.1921 | 0.61 |
| 11326 | *ZC3H18* | 1.7893 | 0.78918 | -0.75491 | -1.4563 | 0.61 |
| 11327 | *KRT86* | 1.8642 | 0.76409 | -0.80498 | -1.856 | 0.61 |
| 11328 | *BMPR2* | 2.1461 | 0.25432 | -0.57763 | -0.58054 | 0.61 |
| 11329 | *MRPL9* | 1.9183 | -0.03405 | -0.06163 | -0.08357 | 0.61 |
| 11330 | *NUDT10* | 1.4194 | 0.54633 | -0.14312 | -1.0373 | 0.61 |
| 11331 | *AGRP* | 1.7676 | 0.037191 | 0.01713 | -0.23678 | 0.61 |
| 11332 | *TNFAIP8L2-SCNM1* | 0.89994 | 0.62785 | 0.29405 | -0.11049 | 0.61 |
| 11333 | *ARHGAP23* | 1.7132 | 1.6375 | -1.5294 | -2.5806 | 0.61 |
| 11334 | *COQ10A* | 2.9236 | -0.44353 | -0.65906 | -2.4141 | 0.61 |
| 11335 | *USP17L3* | 3.6002 | -0.57558 | -1.2037 | -1.2352 | 0.61 |
| 11336 | *METTL22* | 1.9718 | 0.19156 | -0.34259 | -0.61941 | 0.61 |
| 11337 | *CNIH2* | 2.7359 | 0.11643 | -1.0331 | -1.1504 | 0.61 |
| 11338 | *FASTK* | 1.613 | 0.27818 | -0.07199 | -0.31885 | 0.61 |
| 11339 | *PDE3A* | 1.7964 | 1.5896 | -1.5671 | -3.3136 | 0.61 |
| 11340 | *UFSP2* | 4.3284 | -1.2356 | -1.2745 | -1.8646 | 0.61 |
| 11341 | *C5orf63* | 2.5046 | 0.52671 | -1.2142 | -2.6072 | 0.61 |
| 11342 | *ZBTB9* | 2.2552 | -0.21111 | -0.22803 | -2.4221 | 0.61 |
| 11343 | *ZNF28* | 2.5652 | -0.19851 | -0.55068 | -0.74572 | 0.61 |
| 11344 | *CHST12* | 2.1382 | 0.69694 | -1.0197 | -1.7474 | 0.61 |
| 11345 | *PCSK1N* | 1.5205 | 0.77104 | -0.47692 | -2.1324 | 0.60 |
| 11346 | *IKZF2* | 3.8879 | -0.68565 | -1.3892 | -1.5375 | 0.60 |
| 11347 | *LEKR1* | 2.1353 | 0.21571 | -0.53815 | -1.2801 | 0.60 |
| 11348 | *TTLL1* | 4.9397 | -1.2756 | -1.8527 | -2.3357 | 0.60 |
| 11349 | *SGTA* | 3.3991 | -0.44353 | -1.1442 | -1.7322 | 0.60 |
| 11350 | *HS6ST1* | 2.1613 | 0.95397 | -1.3044 | -1.7435 | 0.60 |
| 11351 | *GRIN3A* | 0.71031 | 0.56364 | 0.53647 | -1.808 | 0.60 |
| 11352 | *TMPRSS12* | 3.0641 | -0.40564 | -0.85002 | -1.3961 | 0.60 |
| 11353 | *KCNG4* | 2.8048 | 0.20881 | -1.2058 | -1.2127 | 0.60 |
| 11354 | *SPAG1* | 1.5 | 1.219 | -0.91134 | -0.9599 | 0.60 |
| 11355 | *ZNF765* | 2.6222 | -0.34443 | -0.47018 | -1.5116 | 0.60 |

| | | | | | | |
|---|---|---|---|---|---|---|
| 11356 | *MGAT4A* | 4.5395 | -1.3554 | -1.3767 | -1.5086 | 0.60 |
| 11357 | *CLEC4C* | 2.0862 | -0.13743 | -0.14198 | -1.2226 | 0.60 |
| 11358 | *ACOT6* | 3.7662 | -0.59239 | -1.3674 | -2.2508 | 0.60 |
| 11359 | *ZSCAN9* | 4.0179 | -0.22769 | -1.984 | -2.0458 | 0.60 |
| 11360 | *DCTN1* | 3.7324 | -0.78221 | -1.144 | -2.5255 | 0.60 |
| 11361 | *DDX19B* | 2.5708 | 0.59245 | -1.3585 | -1.3963 | 0.60 |
| 11362 | *CARD16* | 2.5857 | 0.27104 | -1.0528 | -2.0407 | 0.60 |
| 11363 | *CDH23* | 4.196 | -1.0824 | -1.3098 | -1.7443 | 0.60 |
| 11364 | *PIGZ* | 0.97252 | 0.93061 | -0.09956 | -0.47685 | 0.60 |
| 11365 | *IRS2* | 1.1312 | 0.74124 | -0.06902 | -1.6368 | 0.60 |
| 11366 | *ENGASE* | 2.197 | -0.03288 | -0.36086 | -0.98158 | 0.60 |
| 11367 | *GPATCH4* | 2.3432 | 0.12678 | -0.66989 | -1.9022 | 0.60 |
| 11368 | *OBFC1* | 1.4554 | 0.36854 | -0.02406 | -0.06876 | 0.60 |
| 11369 | *ISCU* | 1.1418 | 0.81513 | -0.15771 | -0.77667 | 0.60 |
| 11370 | *FAM72B* | 5.4301 | -1.0654 | -2.5655 | -2.8928 | 0.60 |
| 11371 | *MAGEB3* | 1.3562 | 0.35324 | 0.08942 | -2.9538 | 0.60 |
| 11372 | *AQP2* | 2.9082 | 0.18797 | -1.2975 | -2.0838 | 0.60 |
| 11373 | *OR1J2* | 3.3964 | -0.44287 | -1.155 | -1.632 | 0.60 |
| 11374 | *RAD51B* | 3.0213 | -0.22492 | -0.99888 | -1.6494 | 0.60 |
| 11375 | *S100A12* | 0.9712 | 0.48113 | 0.34516 | -0.00564 | 0.60 |
| 11376 | *RNASE2* | 1.438 | 0.3037 | 0.05569 | -0.29673 | 0.60 |
| 11377 | *LYVE1* | 3.3176 | -0.33958 | -1.1807 | -3.1397 | 0.60 |
| 11378 | *SLC13A4* | 1.4574 | 0.65496 | -0.31544 | -0.97356 | 0.60 |
| 11379 | *GOLGA6L9* | 2.8082 | 0.86983 | -1.882 | -2.3557 | 0.60 |
| 11380 | *LRRC72* | 2.55 | 0.29818 | -1.0528 | -1.2643 | 0.60 |
| 11381 | *ZNF816-ZNF321P* | 3.5525 | -0.37045 | -1.3878 | -3.0961 | 0.60 |
| 11382 | *RBMS2* | 1.7261 | 0.45012 | -0.38219 | -0.83808 | 0.60 |
| 11383 | *FBXO10* | 2.6889 | -0.16485 | -0.73061 | -1.6691 | 0.60 |
| 11384 | *ZNF608* | 4.0523 | -0.69678 | -1.5634 | -1.9397 | 0.60 |
| 11385 | *PEX12* | 3.8993 | 0.28628 | -2.3936 | -4.8617 | 0.60 |
| 11386 | *TMEM215* | 3.3631 | -0.2192 | -1.3523 | -1.5898 | 0.60 |
| 11387 | *LOC728485* | 1.1222 | 0.76 | -0.09104 | -0.79225 | 0.60 |
| 11388 | *LEAP2* | 1.7569 | 0.36498 | -0.33205 | -3.5362 | 0.60 |
| 11389 | *CCSER2* | 2.1583 | 0.25237 | -0.62125 | -3.1806 | 0.60 |
| 11390 | *NR5A1* | 3.1113 | -0.56773 | -0.75433 | -1.8575 | 0.60 |
| 11391 | *MRPS14* | 2.1769 | -0.12108 | -0.26893 | -2.1791 | 0.60 |
| 11392 | *PAAF1* | 0.83715 | 0.8086 | 0.14066 | -0.7602 | 0.60 |
| 11393 | *UBE2M* | 3.3411 | -0.77241 | -0.78229 | -2.2712 | 0.60 |
| 11394 | *SMARCC1* | 1.8051 | 0.098994 | -0.11826 | -2.4889 | 0.60 |
| 11395 | *TAAR2* | 4.4817 | 0.066478 | -2.7626 | -5.632 | 0.60 |
| 11396 | *CDC5L* | 2.1828 | 0.53982 | -0.93898 | -1.8692 | 0.59 |
| 11397 | *SFTPA1* | 1.3912 | 0.35112 | 0.04102 | -1.0102 | 0.59 |
| 11398 | *CCDC180* | 1.8942 | 0.17683 | -0.28825 | -1.3377 | 0.59 |
| 11399 | *RPP25* | 2.4409 | -0.19176 | -0.46721 | -1.1752 | 0.59 |
| 11400 | *RBM47* | 2.328 | 0.14603 | -0.69223 | -0.7443 | 0.59 |
| 11401 | *NDST4* | 1.7952 | 0.099701 | -0.11448 | -1.7766 | 0.59 |

| | | | | | | |
|---|---|---|---|---|---|---|
| 11402 | *NDUFB2* | 4.1012 | -0.2159 | -2.1055 | -3.4314 | 0.59 |
| 11403 | *SNX31* | 2.2124 | -0.06253 | -0.3703 | -0.75511 | 0.59 |
| 11404 | *ENAM* | 1.275 | 0.28082 | 0.22369 | -1.1512 | 0.59 |
| 11405 | *FAM179B* | 1.2908 | 0.47234 | 0.0155 | -1.2673 | 0.59 |
| 11406 | *EFCAB11* | 3.5638 | -0.40427 | -1.3814 | -2.0055 | 0.59 |
| 11407 | *SOHLH1* | 3.5424 | -0.08417 | -1.6802 | -3.0179 | 0.59 |
| 11408 | *GPHA2* | 2.4456 | -0.00464 | -0.66309 | -1.2293 | 0.59 |
| 11409 | *TTC1* | 1.6207 | 0.23635 | -0.08183 | -1.1326 | 0.59 |
| 11410 | *PRKACA* | 4.0714 | -0.8892 | -1.4076 | -1.7595 | 0.59 |
| 11411 | *PCDHB13* | 2.9067 | 0.25958 | -1.3932 | -1.5348 | 0.59 |
| 11412 | *OCM2* | 1.2068 | 0.35535 | 0.21077 | -1.4442 | 0.59 |
| 11413 | *RPL29* | 2.5557 | -0.07036 | -0.71312 | -1.9758 | 0.59 |
| 11414 | *ALS2CL* | 3.9312 | -0.41373 | -1.7457 | -1.9559 | 0.59 |
| 11415 | *ECT2* | 2.0385 | 0.58978 | -0.85871 | -2.397 | 0.59 |
| 11416 | *PRF1* | 4.1276 | -1.065 | -1.2935 | -2.1798 | 0.59 |
| 11417 | *AATF* | 3.4872 | -0.38348 | -1.3349 | -1.4433 | 0.59 |
| 11418 | *DEFB125* | 1.3964 | 1.26 | -0.88784 | -1.5901 | 0.59 |
| 11419 | *CCDC13* | 1.2893 | 0.55041 | -0.07257 | -0.78337 | 0.59 |
| 11420 | *DDR1* | 1.7814 | 0.087849 | -0.10247 | -1.1552 | 0.59 |
| 11421 | *ZNF628* | 2.1957 | 0.021939 | -0.45122 | -1.2903 | 0.59 |
| 11422 | *DENND1B* | 2.8727 | 0.31877 | -1.4256 | -2.0706 | 0.59 |
| 11423 | *CASC3* | 2.8008 | 0.11584 | -1.151 | -1.2784 | 0.59 |
| 11424 | *RAF1* | 1.985 | 0.84128 | -1.0611 | -1.6208 | 0.59 |
| 11425 | *SLC37A3* | 2.7316 | -0.47383 | -0.49266 | -0.70952 | 0.59 |
| 11426 | *ADCK2* | 1.4572 | 0.20314 | 0.10413 | -1.2321 | 0.59 |
| 11427 | *C10orf82* | 1.7241 | 0.27119 | -0.23297 | -0.3794 | 0.59 |
| 11428 | *BCLAF1* | 3.6451 | -0.46745 | -1.4154 | -2.6778 | 0.59 |
| 11429 | *NALCN* | 2.2345 | -0.02895 | -0.44353 | -0.78842 | 0.59 |
| 11430 | *AQP12B* | 1.3852 | 0.35071 | 0.02416 | 0.015107 | 0.59 |
| 11431 | *ASB13* | 2.813 | -0.11049 | -0.94255 | -1.3801 | 0.59 |
| 11432 | *CSNK2B* | 3.2166 | -0.7057 | -0.75128 | -1.1108 | 0.59 |
| 11433 | *ERCC2* | 3.8633 | -0.60003 | -1.5038 | -1.9125 | 0.59 |
| 11434 | *UCN2* | 2.9486 | -0.43385 | -0.75529 | -0.84151 | 0.59 |
| 11435 | *SLC20A1* | 1.071 | 0.39803 | 0.28926 | -0.41266 | 0.59 |
| 11436 | *CLTC* | 1.1451 | 0.51436 | 0.09832 | -0.49941 | 0.59 |
| 11437 | *ELOVL3* | 1.118 | 0.33851 | 0.29991 | 0.065984 | 0.59 |
| 11438 | *FAM208A* | 2.2759 | -0.03135 | -0.48862 | -0.70327 | 0.59 |
| 11439 | *PHTF1* | 3.2855 | 0.037238 | -1.5671 | -1.8822 | 0.59 |
| 11440 | *ABI3BP* | 3.9434 | -1.0315 | -1.1566 | -2.5008 | 0.59 |
| 11441 | *KRTAP9-7* | 1.7323 | 0.71166 | -0.68869 | -1.6413 | 0.59 |
| 11442 | *FAM3A* | 2.5059 | -0.22444 | -0.52637 | -0.8071 | 0.59 |
| 11443 | *LAT2* | 1.944 | 0.41368 | -0.60315 | -2.033 | 0.58 |
| 11444 | *PRR9* | 1.9239 | 1.1669 | -1.3382 | -1.615 | 0.58 |
| 11445 | *FRMD5* | 0.6911 | 0.56421 | 0.49693 | 0.061693 | 0.58 |
| 11446 | *TMEM126B* | 1.7679 | 0.22609 | -0.2429 | -0.72666 | 0.58 |
| 11447 | *SCPEP1* | 2.889 | -0.03675 | -1.1012 | -1.2411 | 0.58 |

| | | | | | | |
|---|---|---|---|---|---|---|
| 11448 | *FCGRT* | 4.1822 | -0.60845 | -1.8227 | -2.101 | 0.58 |
| 11449 | *WDR90* | 2.3469 | -0.0351 | -0.56132 | -1.4954 | 0.58 |
| 11450 | *PCDHGC5* | 3.2782 | -0.38077 | -1.147 | -2.684 | 0.58 |
| 11451 | *LGALS8* | 1.1346 | 0.52061 | 0.09518 | -0.29314 | 0.58 |
| 11452 | *CLEC1A* | 0.70679 | 0.62558 | 0.41801 | -1.6221 | 0.58 |
| 11453 | *ACSS1* | 1.2654 | 0.28864 | 0.19604 | -0.25941 | 0.58 |
| 11454 | *CCDC17* | 0.76144 | 0.5341 | 0.45442 | -0.52343 | 0.58 |
| 11455 | *CD1C* | 1.3721 | 0.67787 | -0.30029 | -0.48916 | 0.58 |
| 11456 | *OXGR1* | 1.1011 | 0.57714 | 0.07062 | -6.2991 | 0.58 |
| 11457 | *CDX2* | 1.5502 | 0.13826 | 0.0603 | -1.061 | 0.58 |
| 11458 | *TAL2* | 0.6052 | 0.60333 | 0.53974 | -0.49564 | 0.58 |
| 11459 | *LAIR2* | 2.725 | 0.75545 | -1.7322 | -2.0351 | 0.58 |
| 11460 | *PRKX* | 1.7134 | 0.50036 | -0.46749 | -0.92486 | 0.58 |
| 11461 | *LOC389602* | 4.3648 | -1.2324 | -1.3865 | -1.4294 | 0.58 |
| 11462 | *WDFY1* | 3.231 | -0.36549 | -1.1207 | -2.7705 | 0.58 |
| 11463 | *ENPP2* | 0.86459 | 0.49513 | 0.38487 | -0.30457 | 0.58 |
| 11464 | *BANP* | 3.3334 | -0.36047 | -1.2284 | -1.5305 | 0.58 |
| 11465 | *AKR1C4* | 2.4331 | 0.10535 | -0.79541 | -1.226 | 0.58 |
| 11466 | *ANGEL1* | 1.6879 | 1.0031 | -0.94859 | -1.7247 | 0.58 |
| 11467 | *CCDC88B* | 1.9559 | 0.004523 | -0.21821 | -0.87768 | 0.58 |
| 11468 | *ARHGAP32* | 3.461 | -0.43841 | -1.2815 | -1.9727 | 0.58 |
| 11469 | *WFIKKN2* | 1.0008 | 0.92231 | -0.18214 | -1.2379 | 0.58 |
| 11470 | *SLC52A2* | 1.3212 | 0.86765 | -0.44814 | -1.4061 | 0.58 |
| 11471 | *KRTAP5-10* | 2.0155 | 0.048185 | -0.32355 | -0.87109 | 0.58 |
| 11472 | *LGALS3* | 1.5695 | 0.39642 | -0.22593 | -0.59557 | 0.58 |
| 11473 | *OR4F5* | 1.6987 | 0.81253 | -0.77196 | -2.2996 | 0.58 |
| 11474 | *TRMT12* | 3.6209 | -0.31976 | -1.5631 | -1.8849 | 0.58 |
| 11475 | *ZFAND6* | 3.8379 | -0.66215 | -1.4392 | -3.1939 | 0.58 |
| 11476 | *ABCC4* | 3.3009 | -0.66761 | -0.89699 | -1.5415 | 0.58 |
| 11477 | *SPATA31D3* | 1.0905 | 0.72896 | -0.08529 | -1.072 | 0.58 |
| 11478 | *ALG1* | 3.8166 | -0.70394 | -1.3785 | -1.6206 | 0.58 |
| 11479 | *PGAM4* | 4.2053 | -1.1866 | -1.285 | -1.6921 | 0.58 |
| 11480 | *SLC39A3* | 1.0121 | 0.66519 | 0.05607 | -0.52426 | 0.58 |
| 11481 | *SLC1A6* | 2.6735 | 0.18287 | -1.1241 | -1.6526 | 0.58 |
| 11482 | *KANK2* | 1.1487 | 1.0925 | -0.5101 | -1.7512 | 0.58 |
| 11483 | *DIS3L2* | 0.74645 | 0.62542 | 0.35805 | 0.08651 | 0.58 |
| 11484 | *POMT1* | 1.4469 | 1.107 | -0.82415 | -1.2532 | 0.58 |
| 11485 | *FSCN2* | 1.2011 | 0.76053 | -0.23203 | -2.8995 | 0.58 |
| 11486 | *PEX14* | 1.4816 | 0.70038 | -0.45296 | -0.79169 | 0.58 |
| 11487 | *ARSF* | 2.3821 | -0.10831 | -0.5451 | -1.0632 | 0.58 |
| 11488 | *KMT2A* | 1.7004 | 1.1449 | -1.1169 | -1.4428 | 0.58 |
| 11489 | *ZNF581* | 2.2989 | 0.21297 | -0.78362 | -1.3837 | 0.58 |
| 11490 | *LHPP* | 2.1587 | 0.78047 | -1.211 | -1.5901 | 0.58 |
| 11491 | *CD40* | 3.4662 | -0.31469 | -1.4234 | -1.5824 | 0.58 |
| 11492 | *S100A9* | 1.6095 | 0.32681 | -0.20892 | -0.94536 | 0.58 |
| 11493 | *MYBL2* | 2.4546 | -0.28819 | -0.43931 | -2.2732 | 0.58 |

| | | | | | | |
|---|---|---|---|---|---|---|
| 11494 | *SLC2A10* | 2.5838 | 0.38754 | -1.2444 | -1.4713 | 0.58 |
| 11495 | *HNRNPF* | 3.2058 | -0.11232 | -1.3666 | -2.7056 | 0.58 |
| 11496 | *OPA1* | 1.2783 | 0.98148 | -0.53304 | -2.9494 | 0.58 |
| 11497 | *BEND4* | 1.4767 | 0.31276 | -0.06276 | -0.09139 | 0.58 |
| 11498 | *MFSD2B* | 3.4005 | -0.3098 | -1.3644 | -1.7806 | 0.58 |
| 11499 | *POMT2* | 0.86661 | 0.6755 | 0.1825 | -1.7577 | 0.57 |
| 11500 | *TRIM64C* | 1.7336 | 0.70297 | -0.7122 | -0.90296 | 0.57 |
| 11501 | *MINOS1* | 3.453 | -0.80213 | -0.92664 | -1.2422 | 0.57 |
| 11502 | *STARD8* | 1.8326 | 0.051282 | -0.1604 | -1.8066 | 0.57 |
| 11503 | *LYZL1* | 2.6708 | -0.12998 | -0.81765 | -1.197 | 0.57 |
| 11504 | *TNXB* | 3.4487 | -0.50961 | -1.2165 | -1.5965 | 0.57 |
| 11505 | *IQSEC1* | 1.8264 | 0.17543 | -0.27961 | -1.658 | 0.57 |
| 11506 | *AASDHPPT* | 3.308 | -0.41319 | -1.1726 | -1.3627 | 0.57 |
| 11507 | *KRT33B* | 0.96075 | 0.63684 | 0.12445 | -0.87898 | 0.57 |
| 11508 | *BRI3* | 2.0783 | 0.36404 | -0.72128 | -1.4572 | 0.57 |
| 11509 | *RCN1* | 2.7454 | -0.11033 | -0.91429 | -2.4564 | 0.57 |
| 11510 | *SSTR2* | 1.0384 | 0.54045 | 0.14176 | -0.94085 | 0.57 |
| 11511 | *MYH14* | 2.3741 | -0.13429 | -0.51929 | -1.9727 | 0.57 |
| 11512 | *ZUFSP* | 1.1409 | 0.29605 | 0.28305 | -2.5754 | 0.57 |
| 11513 | *TBC1D3K* | 3.392 | -0.76355 | -0.90854 | -1.6876 | 0.57 |
| 11514 | *ZNF263* | 1.3498 | 0.42059 | -0.05081 | -2.1766 | 0.57 |
| 11515 | *CHRNA2* | 3.733 | -0.45069 | -1.5633 | -2.3065 | 0.57 |
| 11516 | *ATXN1L* | 2.8426 | -0.30056 | -0.82307 | -0.90303 | 0.57 |
| 11517 | *TES* | 1.7067 | 0.71653 | -0.70647 | -0.9285 | 0.57 |
| 11518 | *SLC35E1* | 4.1556 | -0.726 | -1.714 | -2.163 | 0.57 |
| 11519 | *UTP20* | 1.9367 | 0.12616 | -0.34846 | -1.0553 | 0.57 |
| 11520 | *CNDP1* | 2.4155 | 0.22886 | -0.93057 | -1.3561 | 0.57 |
| 11521 | *RCAN3* | 0.94071 | 0.77423 | -0.00153 | -1.2474 | 0.57 |
| 11522 | *BMP2* | 2.9994 | 0.26209 | -1.5491 | -1.9442 | 0.57 |
| 11523 | *IFT81* | 1.7957 | 0.6425 | -0.72825 | -0.74586 | 0.57 |
| 11524 | *SCGB2A1* | 1.3514 | 0.32745 | 0.03061 | -0.1624 | 0.57 |
| 11525 | *CCDC54* | 3.3334 | -0.80879 | -0.81542 | -2.42 | 0.57 |
| 11526 | *EFS* | 2.0954 | -0.10888 | -0.27744 | -1.2064 | 0.57 |
| 11527 | *UPK2* | 2.7052 | 0.13547 | -1.1332 | -1.5996 | 0.57 |
| 11528 | *SBK3* | 0.60799 | 0.60149 | 0.49793 | -1.0997 | 0.57 |
| 11529 | *DAPK1* | 1.4841 | 1.1289 | -0.90588 | -1.3496 | 0.57 |
| 11530 | *TREML1* | 1.7783 | 0.32653 | -0.39791 | -0.60109 | 0.57 |
| 11531 | *ZNF135* | 1.1557 | 0.65662 | -0.10547 | -1.0232 | 0.57 |
| 11532 | *ADH4* | 1.1391 | 0.82808 | -0.26125 | -1.0949 | 0.57 |
| 11533 | *ATIC* | 3.1901 | -0.00827 | -1.4764 | -2.537 | 0.57 |
| 11534 | *PPP1R35* | 1.7124 | 1.5305 | -1.5376 | -1.7072 | 0.57 |
| 11535 | *TPRN* | 2.4938 | 0.27856 | -1.0697 | -1.8038 | 0.57 |
| 11536 | *PATE1* | 1.1972 | 0.58517 | -0.07997 | -0.86861 | 0.57 |
| 11537 | *NRTN* | 2.9033 | 0.085787 | -1.2867 | -1.6157 | 0.57 |
| 11538 | *PNN* | 1.4528 | 0.56585 | -0.31644 | -0.41955 | 0.57 |
| 11539 | *DHX57* | 0.91411 | 0.42876 | 0.3593 | 0.039168 | 0.57 |

| | | | | | | |
|---|---|---|---|---|---|---|
| 11540 | *PDE6D* | 1.0849 | 0.51428 | 0.10177 | -2.0831 | 0.57 |
| 11541 | *SULT2B1* | 3.104 | -0.19835 | -1.2049 | -1.7714 | 0.57 |
| 11542 | *RIMBP3* | 2.081 | 0.86464 | -1.245 | -2.1829 | 0.57 |
| 11543 | *LPCAT2* | 2.194 | 0.54825 | -1.0421 | -1.2127 | 0.57 |
| 11544 | *C1GALT1C1L* | 2.8567 | -0.31916 | -0.8397 | -1.8272 | 0.57 |
| 11545 | *MAGEC2* | 1.4761 | 0.35188 | -0.13072 | -1.2534 | 0.57 |
| 11546 | *RGS22* | 1.5574 | 0.36493 | -0.22611 | -0.54402 | 0.57 |
| 11547 | *SRP19* | 3.8298 | -1.058 | -1.0756 | -1.3194 | 0.57 |
| 11548 | *UMPS* | 3.9992 | -1.0585 | -1.2448 | -1.3246 | 0.57 |
| 11549 | *CPN2* | 2.1139 | 0.19616 | -0.61515 | -1.4261 | 0.56 |
| 11550 | *KIAA0907* | 1.4161 | 0.2426 | 0.03565 | -0.20142 | 0.56 |
| 11551 | *MED17* | 2.825 | 0.2536 | -1.3851 | -2.6675 | 0.56 |
| 11552 | *SLC12A1* | 3.1826 | -0.46181 | -1.0273 | -1.6066 | 0.56 |
| 11553 | *ZNF367* | 4.5065 | -1.3038 | -1.5094 | -2.0165 | 0.56 |
| 11554 | *TMPRSS2* | 2.6963 | 0.19908 | -1.2027 | -2.6062 | 0.56 |
| 11555 | *TUBB6* | 3.5236 | -0.75053 | -1.083 | -1.4864 | 0.56 |
| 11556 | *NRP2* | 1.0355 | 0.35068 | 0.30301 | -2.2599 | 0.56 |
| 11557 | *AIMP2* | 1.2719 | 0.77039 | -0.35315 | -1.3718 | 0.56 |
| 11558 | *CLDN14* | 1.3311 | 0.28408 | 0.07384 | -0.96718 | 0.56 |
| 11559 | *SRGAP2C* | 3.0951 | -0.53025 | -0.87594 | -2.5276 | 0.56 |
| 11560 | *CENPF* | 2.5584 | 0.54439 | -1.4143 | -1.8849 | 0.56 |
| 11561 | *SLC19A2* | 0.70889 | 0.52529 | 0.45306 | -0.52786 | 0.56 |
| 11562 | *NDUFC2-KCTD14* | 3.8311 | -0.95321 | -1.1912 | -1.9734 | 0.56 |
| 11563 | *RNPEPL1* | 1.9949 | 0.1473 | -0.45604 | -1.1169 | 0.56 |
| 11564 | *PTP4A1* | 3.8156 | -0.84816 | -1.2829 | -1.545 | 0.56 |
| 11565 | *FAM150A* | 2.3144 | -0.3051 | -0.32581 | -0.97329 | 0.56 |
| 11566 | *SAMD10* | 2.078 | 0.054707 | -0.4497 | -0.62174 | 0.56 |
| 11567 | *ATP6V0E1* | 2.9688 | 0.38711 | -1.6734 | -1.7507 | 0.56 |
| 11568 | *XPNPEP3* | 4.0741 | -1.0838 | -1.309 | -1.3873 | 0.56 |
| 11569 | *NAMPT* | 0.88791 | 0.84999 | -0.05914 | -1.7397 | 0.56 |
| 11570 | *TRAF3IP3* | 2.205 | 0.60213 | -1.1287 | -1.4683 | 0.56 |
| 11571 | *ZBED1* | 1.2758 | 0.35893 | 0.04338 | -0.20808 | 0.56 |
| 11572 | *BCL7C* | 1.4674 | 0.14438 | 0.06595 | -0.69059 | 0.56 |
| 11573 | *FAM122A* | 1.3698 | 0.36987 | -0.06224 | -1.048 | 0.56 |
| 11574 | *IGFBP4* | 2.0445 | 0.18891 | -0.55601 | -1.027 | 0.56 |
| 11575 | *IMPDH1* | 3.0407 | -0.44353 | -0.91981 | -1.0041 | 0.56 |
| 11576 | *TBXAS1* | 1.4444 | 0.7921 | -0.55992 | -1.0313 | 0.56 |
| 11577 | *TNN* | 1.08 | 0.80389 | -0.20805 | -0.30326 | 0.56 |
| 11578 | *ZNF354C* | 3.2088 | 0.27747 | -1.8107 | -2.2413 | 0.56 |
| 11579 | *OR6N2* | 2.3135 | -0.29697 | -0.34122 | -0.75603 | 0.56 |
| 11580 | *SH3BGRL3* | 1.4014 | 0.24174 | 0.03203 | -1.6339 | 0.56 |
| 11581 | *ALG14* | 2.1033 | 0.51745 | -0.94576 | -1.1653 | 0.56 |
| 11582 | *AGBL1* | 0.66959 | 0.56655 | 0.43882 | -1.1144 | 0.56 |
| 11583 | *HIST2H2BF* | 3.0898 | -0.56926 | -0.8462 | -2.2495 | 0.56 |
| 11584 | *RFESD* | 1.4816 | 0.37385 | -0.18158 | -0.20681 | 0.56 |
| 11585 | *DTNBP1* | 2.8361 | -0.34434 | -0.82 | -1.1743 | 0.56 |

| | | | | | | |
|---|---|---|---|---|---|---|
| 11586 | *OR2W5* | 0.85388 | 0.48225 | 0.33495 | -1.9326 | 0.56 |
| 11587 | *RCC2* | 1.9136 | 0.19441 | -0.43704 | -1.5304 | 0.56 |
| 11588 | *PLA2G10* | 2.9742 | -0.56241 | -0.74086 | -1.1479 | 0.56 |
| 11589 | *SLC7A8* | 2.8419 | -0.42177 | -0.74932 | -1.1263 | 0.56 |
| 11590 | *C20orf197* | 2.2032 | 0.57987 | -1.1138 | -1.2736 | 0.56 |
| 11591 | *FUT7* | 2.5276 | -0.16903 | -0.68977 | -1.8076 | 0.56 |
| 11592 | *RNASEH2A* | 3.4913 | 0.23881 | -2.0622 | -2.201 | 0.56 |
| 11593 | *LYG1* | 1.6162 | 0.0628 | -0.0112 | -1.1464 | 0.56 |
| 11594 | *PLEKHB1* | 1.9655 | 0.53727 | -0.83527 | -2.1605 | 0.56 |
| 11595 | *TLR4* | 2.386 | 0.080148 | -0.79904 | -3.0408 | 0.56 |
| 11596 | *BPGM* | 2.8021 | 0.020275 | -1.1553 | -2.1496 | 0.56 |
| 11597 | *ZFYVE28* | 2.5245 | -0.10628 | -0.75161 | -1.392 | 0.56 |
| 11598 | *MOB1A* | 0.91327 | 0.60879 | 0.14319 | -0.34927 | 0.56 |
| 11599 | *GATAD2A* | 1.7382 | 0.3254 | -0.39963 | -2.5043 | 0.55 |
| 11600 | *THOC5* | 4.518 | -1.3056 | -1.5493 | -1.6141 | 0.55 |
| 11601 | *CD48* | 2.0974 | 0.16132 | -0.59653 | -2.0923 | 0.55 |
| 11602 | *COX17* | 1.8137 | 0.49425 | -0.64687 | -1.5508 | 0.55 |
| 11603 | *CDON* | 2.0675 | 0.44707 | -0.85388 | -1.7865 | 0.55 |
| 11604 | *OR51A2* | 2.295 | 0.23339 | -0.86951 | -1.4055 | 0.55 |
| 11605 | *JAK3* | 3.5605 | -0.70558 | -1.1977 | -1.5171 | 0.55 |
| 11606 | *KLF17* | 1.4735 | 0.58315 | -0.39959 | -0.95405 | 0.55 |
| 11607 | *NMB* | 1.4015 | 0.17427 | 0.08072 | -0.01323 | 0.55 |
| 11608 | *RABGGTA* | 1.6807 | 0.86392 | -0.88839 | -1.4994 | 0.55 |
| 11609 | *KLHL2* | 1.612 | 0.028786 | 0.01531 | -0.95136 | 0.55 |
| 11610 | *SC5D* | 1.3848 | 0.57318 | -0.30192 | -0.47304 | 0.55 |
| 11611 | *SKP2* | 2.9037 | -0.33583 | -0.91317 | -2.0181 | 0.55 |
| 11612 | *CHRND* | 0.81498 | 0.78631 | 0.05282 | -0.69815 | 0.55 |
| 11613 | *LRRC24* | 0.80955 | 0.64874 | 0.19573 | -1.7377 | 0.55 |
| 11614 | *KLHL24* | 1.8194 | 0.34848 | -0.51405 | -1.227 | 0.55 |
| 11615 | *COL4A2* | 2.0248 | -0.06779 | -0.30358 | -1.6526 | 0.55 |
| 11616 | *CA14* | 1.5653 | 0.33739 | -0.24951 | -2.5007 | 0.55 |
| 11617 | *IRS1* | 3.2395 | -0.66434 | -0.92255 | -1.6469 | 0.55 |
| 11618 | *STK17A* | 1.9486 | 0.5355 | -0.83276 | -1.6527 | 0.55 |
| 11619 | *PDHX* | 1.491 | 0.27023 | -0.11003 | -0.64538 | 0.55 |
| 11620 | *SLC35B4* | 2.9104 | 0.14254 | -1.4018 | -1.8269 | 0.55 |
| 11621 | *KNTC1* | 2.6501 | 0.041736 | -1.041 | -1.3292 | 0.55 |
| 11622 | *KREMEN1* | 2.5802 | 0.20466 | -1.1343 | -1.202 | 0.55 |
| 11623 | *ALCAM* | 0.83658 | 0.45012 | 0.36321 | -0.17016 | 0.55 |
| 11624 | *MAP3K8* | 1.7485 | 0.098319 | -0.19715 | -1.1432 | 0.55 |
| 11625 | *SLC5A5* | 1.6589 | 0.66019 | -0.66973 | -2.0304 | 0.55 |
| 11626 | *IGSF9B* | 1.7952 | 0.42378 | -0.57018 | -2.0704 | 0.55 |
| 11627 | *LOC100144595* | 3.9554 | -1.0791 | -1.2277 | -1.6306 | 0.55 |
| 11628 | *EDDM3B* | 1.1359 | 0.59052 | -0.07787 | -1.0476 | 0.55 |
| 11629 | *OR51F1* | 1.5292 | 0.51554 | -0.39719 | -2.0116 | 0.55 |
| 11630 | *SPATA31A7* | 1.6075 | 0.12602 | -0.08604 | -1.7658 | 0.55 |
| 11631 | *VSTM2A* | 4.6632 | -1.0695 | -1.9474 | -3.4111 | 0.55 |

| | | | | | | |
|---|---|---|---|---|---|---|
| 11632 | *APOL1* | 1.3359 | 0.87933 | -0.56929 | -0.84204 | 0.55 |
| 11633 | *LGALS4* | 3.2345 | -0.63826 | -0.95221 | -1.5337 | 0.55 |
| 11634 | *OR2A4* | 0.97611 | 0.82709 | -0.16 | -0.80675 | 0.55 |
| 11635 | *IRX1* | 1.743 | 0.016379 | -0.117 | -1.9357 | 0.55 |
| 11636 | *NOD2* | 1.1922 | 0.25432 | 0.19503 | -0.19166 | 0.55 |
| 11637 | *STON1-GTF2A1L* | 0.60905 | 0.59683 | 0.43304 | -1.2421 | 0.55 |
| 11638 | *DNAJB5* | 3.2398 | -0.42506 | -1.1761 | -1.2267 | 0.55 |
| 11639 | *GSG1* | 3.2992 | -0.40961 | -1.251 | -1.6731 | 0.55 |
| 11640 | *LOR* | 0.97216 | 0.41913 | 0.24723 | -2.0995 | 0.55 |
| 11641 | *UQCR10* | 1.8263 | 0.00068 | -0.18847 | -1.247 | 0.55 |
| 11642 | *PDIK1L* | 2.6029 | -0.0886 | -0.8759 | -1.1308 | 0.55 |
| 11643 | *NUDT6* | 2.2277 | 0.29438 | -0.88438 | -1.5634 | 0.55 |
| 11644 | *CRHR2* | 2.5254 | -0.0298 | -0.85943 | -1.2048 | 0.55 |
| 11645 | *ANGPT4* | 1.3321 | 0.57385 | -0.26989 | -0.83059 | 0.55 |
| 11646 | *KRTAP1-3* | 0.87086 | 0.56217 | 0.20255 | -4.6085 | 0.55 |
| 11647 | *SOHLH2* | 1.3385 | 0.32067 | -0.02394 | -0.18143 | 0.55 |
| 11648 | *ZADH2* | 1.591 | 1.2433 | -1.1991 | -1.8275 | 0.55 |
| 11649 | *GIMAP2* | 0.98709 | 0.86957 | -0.22197 | -1.5271 | 0.54 |
| 11650 | *ZFPM1* | 2.5345 | -0.29633 | -0.60362 | -2.1238 | 0.54 |
| 11651 | *SPANXN4* | 1.2713 | 0.24806 | 0.11518 | -1.0557 | 0.54 |
| 11652 | *RBBP5* | 2.8102 | -0.03456 | -1.1417 | -2.4018 | 0.54 |
| 11653 | *NUSAP1* | 1.7543 | -0.0175 | -0.10381 | -1.507 | 0.54 |
| 11654 | *ALOX12* | 2.9772 | -0.32644 | -1.0184 | -1.5001 | 0.54 |
| 11655 | *GRPR* | 4.0043 | -0.55192 | -1.8202 | -1.8596 | 0.54 |
| 11656 | *KIF2B* | 2.3058 | -0.25181 | -0.4221 | -2.0613 | 0.54 |
| 11657 | *KLHL10* | 1.6278 | 0.10111 | -0.0975 | -0.6344 | 0.54 |
| 11658 | *CST8* | 1.5037 | 0.95705 | -0.83005 | -1.0476 | 0.54 |
| 11659 | *GPR25* | 1.77 | 0.031869 | -0.17129 | -0.461 | 0.54 |
| 11660 | *FAM90A1* | 2.4226 | -0.36451 | -0.42771 | -0.8897 | 0.54 |
| 11661 | *MARCH9* | 3.1476 | -0.09208 | -1.4265 | -1.591 | 0.54 |
| 11662 | *SCAF11* | 1.0324 | 0.39738 | 0.19837 | -0.37417 | 0.54 |
| 11663 | *FAM193A* | 0.77457 | 0.68367 | 0.16914 | -0.85617 | 0.54 |
| 11664 | *SERPINB4* | 1.5052 | 0.28617 | -0.16518 | -0.2347 | 0.54 |
| 11665 | *SELM* | 2.0321 | 0.5982 | -1.0051 | -2.4784 | 0.54 |
| 11666 | *HLA-DPB1* | 3.3356 | -0.55196 | -1.1585 | -1.468 | 0.54 |
| 11667 | *SLC6A4* | 2.8119 | -0.36641 | -0.82081 | -1.6051 | 0.54 |
| 11668 | *PPP1R10* | 2.586 | -0.41786 | -0.54374 | -1.5525 | 0.54 |
| 11669 | *ZDHHC6* | 2.3148 | 0.049196 | -0.7412 | -1.4603 | 0.54 |
| 11670 | *LRRC3* | 2.2429 | 0.59641 | -1.2167 | -3.087 | 0.54 |
| 11671 | *DNLZ* | 3.8528 | -0.90606 | -1.3249 | -1.356 | 0.54 |
| 11672 | *C7orf34* | 3.9225 | -1.0614 | -1.2393 | -1.6796 | 0.54 |
| 11673 | *CNTNAP3B* | 3.9911 | -1.1762 | -1.1932 | -3.3213 | 0.54 |
| 11674 | *PJA2* | 2.28 | -0.30777 | -0.35193 | -0.79483 | 0.54 |
| 11675 | *PHOX2B* | 1.1984 | 0.61905 | -0.19718 | -1.0663 | 0.54 |
| 11676 | *SV2B* | 3.9885 | -0.41476 | -1.9559 | -2.3557 | 0.54 |
| 11677 | *COL4A3BP* | 1.7032 | 0.65903 | -0.74489 | -0.9191 | 0.54 |

| | | | | | | |
|---|---|---|---|---|---|---|
| 11678 | *TRPV1* | 2.1076 | 0.88398 | -1.3743 | -1.987 | 0.54 |
| 11679 | *KLHDC10* | 4.0356 | -1.1282 | -1.2915 | -1.4683 | 0.54 |
| 11680 | *LZTS3* | 3.5361 | -0.56047 | -1.3606 | -2.2167 | 0.54 |
| 11681 | *DNA2* | 2.5636 | 0.095684 | -1.0455 | -1.481 | 0.54 |
| 11682 | *TMEM174* | 1.0232 | 0.59813 | -0.00833 | -1.5235 | 0.54 |
| 11683 | *MLYCD* | 2.1201 | 0.76623 | -1.274 | -1.8364 | 0.54 |
| 11684 | *SSNA1* | 2.0061 | 0.33119 | -0.726 | -1.7073 | 0.54 |
| 11685 | *ARX* | 1.4363 | 0.2439 | -0.06926 | -1.6448 | 0.54 |
| 11686 | *CSTF2T* | 1.1267 | 0.39424 | 0.08868 | -1.3615 | 0.54 |
| 11687 | *SUPT20HL2* | 1.5223 | 1.2322 | -1.146 | -2.4454 | 0.54 |
| 11688 | *YIPF3* | 1.1084 | 0.51575 | -0.01568 | -1.6921 | 0.54 |
| 11689 | *OR5AU1* | 3.665 | -0.17931 | -1.8777 | -1.8976 | 0.54 |
| 11690 | *HABP4* | 1.3405 | 0.25344 | 0.01369 | -2.4647 | 0.54 |
| 11691 | *ACBD5* | 1.0826 | 0.53127 | -0.00832 | -0.52021 | 0.54 |
| 11692 | *OR2F1* | 1.9887 | 0.047726 | -0.4312 | -0.86889 | 0.54 |
| 11693 | *NEK4* | 1.397 | 0.40747 | -0.19977 | -1.2763 | 0.53 |
| 11694 | *DTX3L* | 2.154 | 0.20381 | -0.75351 | -1.3007 | 0.53 |
| 11695 | *LATS2* | 1.4855 | 1.4121 | -1.2935 | -2.1574 | 0.53 |
| 11696 | *L1CAM* | 4.9072 | -1.632 | -1.6724 | -1.8636 | 0.53 |
| 11697 | *NDUFA8* | 2.2242 | 0.60177 | -1.2234 | -1.3542 | 0.53 |
| 11698 | *HOXC9* | 2.597 | 0.56009 | -1.5552 | -2.2485 | 0.53 |
| 11699 | *EZR* | 1.7094 | 0.53875 | -0.64632 | -1.794 | 0.53 |
| 11700 | *ADAMTS6* | 1.1588 | 0.42899 | 0.0136 | -1.4143 | 0.53 |
| 11701 | *BAIAP2* | 1.1549 | 0.51621 | -0.07005 | -2.0036 | 0.53 |
| 11702 | *PRRG2* | 0.91932 | 0.39355 | 0.28812 | -1.7235 | 0.53 |
| 11703 | *PROSER3* | 1.2522 | 0.20243 | 0.14582 | -1.4683 | 0.53 |
| 11704 | *NF2* | 0.84302 | 0.80598 | -0.04919 | -0.71553 | 0.53 |
| 11705 | *PPY* | 2.5507 | -0.10934 | -0.84238 | -1.605 | 0.53 |
| 11706 | *TMEM165* | 1.5541 | 0.097548 | -0.05357 | -0.65606 | 0.53 |
| 11707 | *ZNF444* | 1.0425 | 0.41717 | 0.13807 | -0.6251 | 0.53 |
| 11708 | *SMG5* | 1.8411 | 1.193 | -1.4364 | -2.0593 | 0.53 |
| 11709 | *PODXL* | 1.3722 | 0.51748 | -0.29232 | -1.8283 | 0.53 |
| 11710 | *TRIM46* | 0.84518 | 0.64155 | 0.11033 | -0.51549 | 0.53 |
| 11711 | *ZNF280A* | 0.74242 | 0.46706 | 0.38728 | -0.67688 | 0.53 |
| 11712 | *TLK1* | 2.1302 | 0.82334 | -1.3572 | -1.6523 | 0.53 |
| 11713 | *TST* | 3.4196 | -0.85829 | -0.96529 | -1.7611 | 0.53 |
| 11714 | *MT1B* | 1.8093 | 0.12865 | -0.34252 | -1.5806 | 0.53 |
| 11715 | *PRPS1* | 1.863 | 0.027839 | -0.29652 | -1.2646 | 0.53 |
| 11716 | *LRRN1* | 0.6143 | 0.5758 | 0.40336 | -0.80978 | 0.53 |
| 11717 | *BBS10* | 3.1659 | -0.76237 | -0.81031 | -1.2776 | 0.53 |
| 11718 | *PHGR1* | 1.0874 | 0.44439 | 0.05866 | -2.3449 | 0.53 |
| 11719 | *SEPT6* | 3.0625 | 0.83963 | -2.3121 | -2.3189 | 0.53 |
| 11720 | *ZIM2* | 0.92313 | 0.53037 | 0.13459 | -2.2139 | 0.53 |
| 11721 | *METTL10* | 3.1551 | -0.31035 | -1.2571 | -2.5571 | 0.53 |
| 11722 | *GALNT7* | 0.90034 | 0.6528 | 0.03435 | -0.65035 | 0.53 |
| 11723 | *NUDCD2* | 2.9975 | -0.61958 | -0.79083 | -2.3265 | 0.53 |

| | | | | | | |
|---|---|---|---|---|---|---|
| 11724 | *C7orf49* | 3.4003 | -0.48659 | -1.3268 | -1.7994 | 0.53 |
| 11725 | *MSRA* | 1.1699 | 0.59211 | -0.17683 | -1.5739 | 0.53 |
| 11726 | *STUB1* | 1.8494 | -0.0082 | -0.25606 | -1.4935 | 0.53 |
| 11727 | *CLRN1* | 2.0972 | 0.64116 | -1.1543 | -1.3353 | 0.53 |
| 11728 | *UBE2E3* | 2.0447 | 0.071115 | -0.53321 | -0.73944 | 0.53 |
| 11729 | *PITPNB* | 1.5337 | 0.94819 | -0.89974 | -1.0564 | 0.53 |
| 11730 | *HIST1H4B* | 2.2024 | -0.16041 | -0.46092 | -1.4754 | 0.53 |
| 11731 | *C17orf47* | 3.2547 | -0.28303 | -1.3912 | -1.7572 | 0.53 |
| 11732 | *GNA11* | 0.9309 | 0.35764 | 0.29118 | -1.6893 | 0.53 |
| 11733 | *ACTR5* | 1.5292 | 0.27195 | -0.22166 | -1.5954 | 0.53 |
| 11734 | *LAMB2* | 2.08 | -0.19357 | -0.30743 | -0.32858 | 0.53 |
| 11735 | *BEND3* | 1.4579 | 0.64417 | -0.52349 | -0.81496 | 0.53 |
| 11736 | *TLR3* | 1.5478 | 0.63025 | -0.59959 | -1.9417 | 0.53 |
| 11737 | *ANLN* | 3.1073 | -0.63971 | -0.8897 | -1.4365 | 0.53 |
| 11738 | *ATP1B1* | 2.5484 | -0.11664 | -0.855 | -1.2947 | 0.53 |
| 11739 | *KRTAP22-1* | 4.6247 | -1.4597 | -1.5884 | -1.7266 | 0.53 |
| 11740 | *LRSAM1* | 1.2488 | 0.17905 | 0.14867 | -0.45449 | 0.53 |
| 11741 | *PPCS* | 2.5698 | 0.081535 | -1.0751 | -1.1921 | 0.53 |
| 11742 | *GPT* | 0.87458 | 0.79521 | -0.09362 | -0.53666 | 0.53 |
| 11743 | *RILP* | 1.2106 | 0.1979 | 0.16758 | -1.8714 | 0.53 |
| 11744 | *RELL1* | 0.75233 | 0.41263 | 0.41083 | -1.7078 | 0.53 |
| 11745 | *SDCCAG3* | 0.98237 | 0.54647 | 0.04556 | -1.6339 | 0.52 |
| 11746 | *HIST1H3J* | 2.0458 | 0.83804 | -1.311 | -2.3557 | 0.52 |
| 11747 | *SLC35B1* | 3.4536 | -0.6951 | -1.186 | -1.6709 | 0.52 |
| 11748 | *ZC3H12A* | 1.9355 | 0.60262 | -0.96572 | -1.9727 | 0.52 |
| 11749 | *GDPD4* | 1.4128 | 0.42143 | -0.26207 | -0.51398 | 0.52 |
| 11750 | *DPAGT1* | 3.4182 | -0.77924 | -1.067 | -1.8352 | 0.52 |
| 11751 | *TMEM104* | 1.8262 | 0.86048 | -1.1148 | -2.305 | 0.52 |
| 11752 | *NHLRC4* | 1.2872 | 0.55945 | -0.27486 | -0.96671 | 0.52 |
| 11753 | *BTF3L4* | 0.67451 | 0.62835 | 0.26852 | 0.267 | 0.52 |
| 11754 | *TLE3* | 1.0863 | 0.53073 | -0.04606 | -2.0949 | 0.52 |
| 11755 | *HSF4* | 4.4644 | -0.67141 | -2.2223 | -5.8184 | 0.52 |
| 11756 | *PSMB5* | 3.6299 | -0.55196 | -1.5073 | -1.7822 | 0.52 |
| 11757 | *TMEM63B* | 3.4416 | -0.50877 | -1.3625 | -1.7758 | 0.52 |
| 11758 | *OCLM* | 1.0178 | 0.48465 | 0.06767 | -1.2572 | 0.52 |
| 11759 | *GSN* | 2.5538 | -0.31411 | -0.6706 | -1.7129 | 0.52 |
| 11760 | *ZNF622* | 1.8651 | 0.36012 | -0.65679 | -0.74249 | 0.52 |
| 11761 | *DOC2A* | 2.5829 | -0.26584 | -0.74864 | -2.672 | 0.52 |
| 11762 | *ERV3-1* | 2.2561 | -0.19627 | -0.49216 | -0.93999 | 0.52 |
| 11763 | *STK10* | 3.3591 | -0.75053 | -1.0415 | -2.0064 | 0.52 |
| 11764 | *KLKB1* | 0.7579 | 0.68028 | 0.12606 | -1.6176 | 0.52 |
| 11765 | *TRIM65* | 1.8671 | 0.30414 | -0.60744 | -1.5604 | 0.52 |
| 11766 | *ZFP3* | 1.1831 | 0.78605 | -0.40556 | -0.70244 | 0.52 |
| 11767 | *ORC4* | 1.4847 | 1.4812 | -1.4028 | -1.6527 | 0.52 |
| 11768 | *EIF4EBP2* | 2.6481 | 0.076792 | -1.1619 | -1.6124 | 0.52 |
| 11769 | *BCAS3* | 0.75464 | 0.64586 | 0.16232 | -0.95753 | 0.52 |

| | | | | | | |
|---|---|---|---|---|---|---|
| 11770 | *ARIH1* | 3.1055 | 0.71021 | -2.2533 | -2.2593 | 0.52 |
| 11771 | *AP1M2* | 0.95967 | 0.37249 | 0.23008 | -0.74069 | 0.52 |
| 11772 | *C18orf63* | 1.1004 | 0.50722 | -0.04564 | -0.07138 | 0.52 |
| 11773 | *PSG4* | 1.1394 | 0.23161 | 0.19074 | -1.6525 | 0.52 |
| 11774 | *OR5D14* | 5.5666 | -1.5235 | -2.4815 | -5.3303 | 0.52 |
| 11775 | *ITPR2* | 1.1599 | 0.44961 | -0.04799 | -0.70202 | 0.52 |
| 11776 | *DNAAF3* | 2.6912 | -0.18791 | -0.94255 | -1.1974 | 0.52 |
| 11777 | *TMED7* | 1.9492 | 0.054869 | -0.44353 | -0.67691 | 0.52 |
| 11778 | *ZMYM1* | 0.83524 | 0.72607 | -0.00081 | -0.17138 | 0.52 |
| 11779 | *VRK1* | 1.0894 | 0.24511 | 0.2256 | -0.20023 | 0.52 |
| 11780 | *PGAP2* | 1.8275 | -0.12457 | -0.14293 | -0.81001 | 0.52 |
| 11781 | *OR2D2* | 1.6097 | 0.011164 | -0.06142 | -0.63542 | 0.52 |
| 11782 | *DEFB136* | 2.9924 | -0.62782 | -0.80577 | -1.0999 | 0.52 |
| 11783 | *TCEANC* | 1.3049 | 0.59553 | -0.34275 | -0.5231 | 0.52 |
| 11784 | *RND2* | 1.6556 | 0.030874 | -0.12891 | -0.31698 | 0.52 |
| 11785 | *FGF8* | 1.2621 | 0.64541 | -0.35023 | -1.0626 | 0.52 |
| 11786 | *KCTD12* | 1.1563 | 0.67888 | -0.27844 | -0.75173 | 0.52 |
| 11787 | *NDUFC1* | 4.0294 | -1.1804 | -1.2936 | -1.631 | 0.52 |
| 11788 | *SLC17A9* | 2.4315 | 0.88177 | -1.7585 | -2.1796 | 0.52 |
| 11789 | *RENBP* | 3.2176 | -0.70132 | -0.96161 | -1.5569 | 0.52 |
| 11790 | *NDUFA10* | 2.939 | 0.19377 | -1.5781 | -2.1176 | 0.52 |
| 11791 | *C16orf96* | 1.4276 | 0.53936 | -0.41237 | -2.3292 | 0.52 |
| 11792 | *ZFP42* | 1.8442 | 0.18879 | -0.47843 | -0.7423 | 0.52 |
| 11793 | *OR2A25* | 1.8512 | -0.09709 | -0.19995 | -1.6164 | 0.52 |
| 11794 | *NKAPL* | 2.0035 | 0.37687 | -0.827 | -1.5724 | 0.52 |
| 11795 | *NEURL4* | 1.1292 | 0.94222 | -0.51811 | -1.4234 | 0.52 |
| 11796 | *ACOT9* | 1.4649 | 0.86992 | -0.78183 | -1.0712 | 0.52 |
| 11797 | *HYLS1* | 2.111 | -0.12642 | -0.43167 | -0.49138 | 0.52 |
| 11798 | *YLPM1* | 1.0106 | 0.9931 | -0.45119 | -1.9591 | 0.52 |
| 11799 | *ANKRD35* | 0.8554 | 0.67347 | 0.02334 | -1.6696 | 0.52 |
| 11800 | *DUSP14* | 2.4784 | -0.18633 | -0.7412 | -2.1429 | 0.52 |
| 11801 | *TOP3A* | 2.0559 | 0.90323 | -1.4097 | -1.4787 | 0.52 |
| 11802 | *NPS* | 1.4127 | 0.21632 | -0.08118 | -1.9623 | 0.52 |
| 11803 | *MYH15* | 2.9509 | -0.09652 | -1.3068 | -2.4946 | 0.52 |
| 11804 | *IER5* | 3.4375 | 0.44681 | -2.337 | -2.3426 | 0.52 |
| 11805 | *HR* | 1.6891 | 0.22358 | -0.36569 | -1.8908 | 0.52 |
| 11806 | *ZDHHC3* | 3.4715 | -0.6761 | -1.2485 | -1.6186 | 0.52 |
| 11807 | *ABCC10* | 3.9908 | -1.1756 | -1.2684 | -1.9624 | 0.52 |
| 11808 | *LMOD1* | 1.0094 | 0.87883 | -0.34247 | -0.54744 | 0.52 |
| 11809 | *DNAJA2* | 2.1354 | 0.31287 | -0.90281 | -1.5635 | 0.52 |
| 11810 | *PTCH2* | 1.0379 | 0.55577 | -0.04927 | -0.71347 | 0.51 |
| 11811 | *AMZ1* | 1.458 | 0.082627 | 0.00367 | -2.5935 | 0.51 |
| 11812 | *DPPA3* | 2.8302 | 0.12149 | -1.4076 | -1.526 | 0.51 |
| 11813 | *KIAA1211L* | 0.98126 | 0.46635 | 0.09472 | -0.38248 | 0.51 |
| 11814 | *SHISA6* | 3.4122 | -0.77534 | -1.0948 | -1.7407 | 0.51 |
| 11815 | *RBM34* | 3.5029 | -0.73552 | -1.2258 | -1.2306 | 0.51 |

| | | | | | | |
|---|---|---|---|---|---|---|
| 11816 | *DNAJB2* | 1.3122 | 0.32059 | -0.0925 | -0.57827 | 0.51 |
| 11817 | *LCE4A* | 1.7204 | 1.1937 | -1.3741 | -2.1036 | 0.51 |
| 11818 | *FAM228B* | 2.3505 | -0.16574 | -0.64496 | -1.7024 | 0.51 |
| 11819 | *HPS5* | 1.2437 | 0.30432 | -0.00823 | -0.10101 | 0.51 |
| 11820 | *SLC35E2* | 0.80647 | 0.58201 | 0.15123 | 0.019201 | 0.51 |
| 11821 | *GTF3C4* | 3.2604 | -0.60852 | -1.1126 | -2.1364 | 0.51 |
| 11822 | *MEIG1* | 2.5393 | 0.57941 | -1.5801 | -1.7655 | 0.51 |
| 11823 | *PPP2R5E* | 0.71478 | 0.61639 | 0.20724 | -0.66652 | 0.51 |
| 11824 | *KRTAP2-4* | 2.1809 | 0.72921 | -1.3721 | -1.6989 | 0.51 |
| 11825 | *ANKRD66* | 3.2424 | -0.74638 | -0.95835 | -1.1791 | 0.51 |
| 11826 | *LRP2* | 1.8309 | 0.25143 | -0.54467 | -0.9728 | 0.51 |
| 11827 | *SH3PXD2B* | 1.3806 | 0.67459 | -0.51826 | -1.3617 | 0.51 |
| 11828 | *PTGIR* | 1.2555 | 0.15336 | 0.12753 | -0.91701 | 0.51 |
| 11829 | *LAMB4* | 2.5839 | 0.17894 | -1.227 | -1.8946 | 0.51 |
| 11830 | *PDE1C* | 3.3516 | -0.46763 | -1.3486 | -2.0016 | 0.51 |
| 11831 | *HYAL2* | 4.5299 | -0.76371 | -2.2313 | -3.4957 | 0.51 |
| 11832 | *UBR5* | 0.74519 | 0.42316 | 0.36628 | -1.154 | 0.51 |
| 11833 | *CUTA* | 3.5869 | -0.67313 | -1.3795 | -1.9302 | 0.51 |
| 11834 | *NR1H4* | 0.85111 | 0.84967 | -0.1672 | -2.0466 | 0.51 |
| 11835 | *ILDR2* | 1.2811 | 0.65365 | -0.40171 | -1.0177 | 0.51 |
| 11836 | *KIF1C* | 0.80124 | 0.41519 | 0.31649 | 0.012112 | 0.51 |
| 11837 | *APOPT1* | 0.57432 | 0.54151 | 0.41696 | -1.8946 | 0.51 |
| 11838 | *DNAJA1* | 2.0317 | -0.05801 | -0.44134 | -1.0335 | 0.51 |
| 11839 | *EIF3L* | 2.9267 | -0.19995 | -1.1945 | -2.1214 | 0.51 |
| 11840 | *CX3CR1* | 0.75683 | 0.60832 | 0.167 | -1.5649 | 0.51 |
| 11841 | *BID* | 2.0964 | -0.01656 | -0.54786 | -2.6212 | 0.51 |
| 11842 | *UBASH3A* | 2.1651 | -0.2804 | -0.35306 | -1.0706 | 0.51 |
| 11843 | *DHCR24* | 1.5901 | 0.23258 | -0.29123 | -0.74291 | 0.51 |
| 11844 | *ALG10* | 1.2806 | 1.2301 | -0.97956 | -1.2488 | 0.51 |
| 11845 | *DPY19L3* | 1.9993 | 0.94668 | -1.4149 | -1.547 | 0.51 |
| 11846 | *LRRC16B* | 3.8518 | -0.9728 | -1.3481 | -2.0466 | 0.51 |
| 11847 | *ZNF679* | 2.289 | -0.16581 | -0.59231 | -0.94422 | 0.51 |
| 11848 | *RPL35A* | 2.4919 | -0.3576 | -0.60343 | -2.3828 | 0.51 |
| 11849 | *TBC1D4* | 3.5539 | -0.56405 | -1.4596 | -1.8689 | 0.51 |
| 11850 | *ANKRD34B* | 4.0852 | -0.96997 | -1.5852 | -2.0687 | 0.51 |
| 11851 | *KRTAP2-2* | 3.0932 | -0.38044 | -1.183 | -1.4064 | 0.51 |
| 11852 | *AP3S1* | 1.4934 | 0.033126 | 0.00292 | -2.4094 | 0.51 |
| 11853 | *PRSS22* | 0.89724 | 0.80758 | -0.17554 | -0.18955 | 0.51 |
| 11854 | *LCN15* | 2.7024 | -0.30375 | -0.86996 | -1.9302 | 0.51 |
| 11855 | *HTR7* | 2.0144 | 0.28128 | -0.76725 | -1.4368 | 0.51 |
| 11856 | *SNTB2* | 2.7758 | -0.17952 | -1.068 | -1.2426 | 0.51 |
| 11857 | *TCEB3C* | 0.68036 | 0.48287 | 0.36397 | 0.16344 | 0.51 |
| 11858 | *POLE2* | 1.8978 | -0.10074 | -0.26992 | -1.7561 | 0.51 |
| 11859 | *WFDC1* | 1.0236 | 0.59627 | -0.09312 | -0.19169 | 0.51 |
| 11860 | *NFKBIZ* | 1.3603 | 0.45447 | -0.28836 | -1.1035 | 0.51 |
| 11861 | *RABL2A* | 3.1202 | -0.77475 | -0.81932 | -1.1974 | 0.51 |

| | | | | | | |
|---|---|---|---|---|---|---|
| 11862 | *TSGA10* | 1.6734 | 0.45712 | -0.60604 | -1.25 | 0.51 |
| 11863 | *MGAT3* | 3.0852 | 0.57416 | -2.1351 | -2.796 | 0.51 |
| 11864 | *BRINP3* | 4.1754 | -0.94777 | -1.7034 | -2.4197 | 0.51 |
| 11865 | *SAMD14* | 1.9601 | -0.00164 | -0.43472 | -1.3784 | 0.51 |
| 11866 | *MORC3* | 1.3788 | 0.63157 | -0.48669 | -0.77733 | 0.51 |
| 11867 | *LMBR1* | 0.70898 | 0.58147 | 0.23189 | -0.82921 | 0.51 |
| 11868 | *ARHGEF2* | 1.3101 | 0.45486 | -0.24327 | -2.2277 | 0.51 |
| 11869 | *C3* | 2.7479 | 0.15456 | -1.3818 | -4.4937 | 0.51 |
| 11870 | *HLA-DRB1* | 1.0365 | 0.72734 | -0.24578 | -2.0178 | 0.51 |
| 11871 | *CORT* | 1.2577 | 0.27415 | -0.0146 | -1.6281 | 0.51 |
| 11872 | *NCBP2* | 1.0419 | 0.29943 | 0.17578 | -1.0399 | 0.51 |
| 11873 | *AP1S3* | 0.69404 | 0.50311 | 0.31908 | -0.62185 | 0.51 |
| 11874 | *MCM2* | 2.3113 | -0.30585 | -0.48956 | -1.821 | 0.51 |
| 11875 | *KRTAP4-7* | 1.0697 | 0.50065 | -0.05456 | -0.81798 | 0.51 |
| 11876 | *NMS* | 2.069 | -0.18019 | -0.37466 | -1.4147 | 0.50 |
| 11877 | *STRN3* | 1.9574 | 0.85552 | -1.2988 | -2.1873 | 0.50 |
| 11878 | *LAMC2* | 1.1811 | 0.89846 | -0.56545 | -0.76821 | 0.50 |
| 11879 | *KLF8* | 1.6718 | 0.31802 | -0.47577 | -1.367 | 0.50 |
| 11880 | *ATP5G2* | 2.0977 | -0.05066 | -0.53346 | -0.73673 | 0.50 |
| 11881 | *BLID* | 1.3605 | 0.28161 | -0.12866 | -0.30697 | 0.50 |
| 11882 | *FAR2* | 2.0695 | 0.33669 | -0.8929 | -1.8033 | 0.50 |
| 11883 | *PRG2* | 0.8577 | 0.35803 | 0.29693 | 0.26121 | 0.50 |
| 11884 | *GJA9* | 2.9101 | -0.66107 | -0.73686 | -5.7811 | 0.50 |
| 11885 | *HSPA1L* | 1.8362 | 1.0581 | -1.3823 | -1.6691 | 0.50 |
| 11886 | *POTEM* | 2.5249 | -0.28994 | -0.72304 | -2.9578 | 0.50 |
| 11887 | *TUBA3D* | 1.6197 | 0.092873 | -0.20112 | -1.7188 | 0.50 |
| 11888 | *SEC14L3* | 1.2876 | 0.69094 | -0.4675 | -0.7484 | 0.50 |
| 11889 | *SCIN* | 1.6115 | 0.61323 | -0.71403 | -0.79453 | 0.50 |
| 11890 | *NEIL3* | 1.7434 | 0.70459 | -0.93764 | -0.96846 | 0.50 |
| 11891 | *SLC17A8* | 1.0945 | 1.0741 | -0.65906 | -0.78784 | 0.50 |
| 11892 | *PRRT1* | 0.60155 | 0.52501 | 0.38297 | 0.047056 | 0.50 |
| 11893 | *HADHB* | 1.9215 | 0.61495 | -1.027 | -1.0576 | 0.50 |
| 11894 | *SEC23IP* | 2.4565 | -0.14309 | -0.80594 | -2.8405 | 0.50 |
| 11895 | *ESRRG* | 2.1929 | -0.00303 | -0.68252 | -1.6365 | 0.50 |
| 11896 | *TRMT2B* | 1.8866 | 0.32631 | -0.7057 | -1.8869 | 0.50 |
| 11897 | *NDUFA11* | 2.7852 | -0.61796 | -0.66008 | -0.96878 | 0.50 |
| 11898 | *ANKRD52* | 1.3013 | 0.58021 | -0.37461 | -0.65971 | 0.50 |
| 11899 | *KCND2* | 2.8039 | -0.58822 | -0.71032 | -0.75128 | 0.50 |
| 11900 | *CSRP1* | 1.8487 | 0.45573 | -0.79909 | -1.6691 | 0.50 |
| 11901 | *BPY2B* | 1.4581 | 0.055343 | -0.00832 | -1.2239 | 0.50 |
| 11902 | *C19orf73* | 1.6377 | 0.31206 | -0.44575 | -2.0242 | 0.50 |
| 11903 | *MIER2* | 2.6225 | -0.50324 | -0.61549 | -1.7113 | 0.50 |
| 11904 | *COL26A1* | 1.1665 | 0.72762 | -0.39048 | -1.6636 | 0.50 |
| 11905 | *ACCSL* | 0.95681 | 0.33323 | 0.21343 | -0.57573 | 0.50 |
| 11906 | *C3orf18* | 1.8529 | 0.38805 | -0.73812 | -1.6306 | 0.50 |
| 11907 | *NECAP1* | 3.8524 | -1.0224 | -1.3275 | -1.85 | 0.50 |

| | | | | | | |
|---|---|---|---|---|---|---|
| 11908 | *STAT2* | 2.1016 | -0.14506 | -0.45515 | -2.0368 | 0.50 |
| 11909 | *ACBD7* | 0.98421 | 0.62097 | -0.10405 | -0.80475 | 0.50 |
| 11910 | *CTH* | 1.1904 | 1.1002 | -0.78956 | -1.4197 | 0.50 |
| 11911 | *GNB1* | 2.7828 | -0.60424 | -0.67801 | -1.8842 | 0.50 |
| 11912 | *KIAA0825* | 1.7504 | 0.080802 | -0.33085 | -2.004 | 0.50 |
| 11913 | *RGS9* | 1.1709 | 0.23456 | 0.09354 | 0.082543 | 0.50 |
| 11914 | *TMEM191C* | 3.4825 | -0.64496 | -1.3393 | -2.0478 | 0.50 |
| 11915 | *OR4D9* | 2.7288 | -0.10661 | -1.1244 | -1.9534 | 0.50 |
| 11916 | *SCRN3* | 2.2746 | -0.3185 | -0.45867 | -1.0942 | 0.50 |
| 11917 | *PCDHA5* | 2.3813 | 0.69864 | -1.583 | -2.3602 | 0.50 |
| 11918 | *DYNLL2* | 2.6267 | 0.19074 | -1.3218 | -2.0483 | 0.50 |
| 11919 | *NFRKB* | 2.357 | -0.41786 | -0.44353 | -1.1409 | 0.50 |
| 11920 | *ZNF853* | 1.9531 | 0.24191 | -0.69958 | -1.714 | 0.50 |
| 11921 | *TMEM99* | 0.79809 | 0.50078 | 0.19645 | -1.8085 | 0.50 |
| 11922 | *CCDC102B* | 1.8798 | -0.05408 | -0.33139 | -0.57397 | 0.50 |
| 11923 | *PARS2* | 2.5687 | 0.20565 | -1.2802 | -1.4218 | 0.50 |
| 11924 | *MAGEA2* | 1.8431 | -0.1637 | -0.18612 | -1.0557 | 0.50 |
| 11925 | *RANBP9* | 2.3114 | -0.26895 | -0.5497 | -2.4078 | 0.50 |
| 11926 | *HSF2* | 3.6565 | -0.70945 | -1.4544 | -2.4496 | 0.50 |
| 11927 | *NQO2* | 3.1239 | -0.77404 | -0.85794 | -2.4853 | 0.50 |
| 11928 | *ROPN1L* | 1.1298 | 0.31394 | 0.04796 | -0.41178 | 0.50 |
| 11929 | *DIRAS2* | 3.8378 | -1.0889 | -1.2577 | -1.4919 | 0.50 |
| 11930 | *ACE* | 2.0973 | -0.16122 | -0.4449 | -0.89392 | 0.50 |
| 11931 | *TMEM120B* | 2.6739 | -0.34173 | -0.84147 | -1.4883 | 0.50 |
| 11932 | *PVALB* | 2.3931 | -0.05905 | -0.84345 | -1.3648 | 0.50 |
| 11933 | *MAGEE2* | 1.4298 | 1.1347 | -1.0739 | -1.4853 | 0.50 |
| 11934 | *ANOS1* | 1.4962 | 0.002242 | -0.00823 | -1.2399 | 0.50 |
| 11935 | *NPIPA8* | 1.2935 | 0.84041 | -0.64616 | -1.2616 | 0.50 |
| 11936 | *CGGBP1* | 2.4956 | -0.28076 | -0.72725 | -0.93099 | 0.50 |
| 11937 | *SYCN* | 2.9306 | -0.54469 | -0.89832 | -5.4539 | 0.50 |
| 11938 | *MRPL52* | 3.0775 | -0.0975 | -1.4926 | -3.3502 | 0.50 |
| 11939 | *NOXRED1* | 0.99101 | 0.252 | 0.24358 | -0.32866 | 0.50 |
| 11940 | *TMEM257* | 0.81921 | 0.44223 | 0.22431 | -1.4284 | 0.50 |
| 11941 | *GAREML* | 2.1303 | 0.55963 | -1.2052 | -2.6463 | 0.49 |
| 11942 | *C1orf168* | 0.70424 | 0.41605 | 0.36403 | -1.1407 | 0.49 |
| 11943 | *LANCL3* | 5.2444 | -1.2782 | -2.4825 | -2.8823 | 0.49 |
| 11944 | *URM1* | 1.0713 | 0.85562 | -0.44353 | -1.5552 | 0.49 |
| 11945 | *DCK* | 0.68374 | 0.49003 | 0.30954 | -0.67382 | 0.49 |
| 11946 | *TPD52L2* | 3.257 | -0.68169 | -1.0932 | -3.0757 | 0.49 |
| 11947 | *HEBP1* | 2.8001 | -0.24569 | -1.0728 | -1.1576 | 0.49 |
| 11948 | *ZNF155* | 3.1078 | -0.1375 | -1.4894 | -1.7422 | 0.49 |
| 11949 | *PSME2* | 2.0711 | 0.32892 | -0.92065 | -1.5814 | 0.49 |
| 11950 | *PAM* | 0.78676 | 0.73157 | -0.03951 | -1.8102 | 0.49 |
| 11951 | *G2E3* | 0.61696 | 0.56728 | 0.29457 | 0.011164 | 0.49 |
| 11952 | *OR6C76* | 4.3673 | -1.0944 | -1.7945 | -1.9251 | 0.49 |
| 11953 | *LAG3* | 1.445 | 0.36422 | -0.33109 | -0.66475 | 0.49 |

| | | | | | | |
|---|---|---|---|---|---|---|
| 11954 | *CTSF* | 1.6645 | 0.004855 | -0.19144 | -1.1286 | 0.49 |
| 11955 | *KCNH7* | 1.9837 | 0.10168 | -0.60869 | -1.0168 | 0.49 |
| 11956 | *OR4K2* | 2.5611 | 0.85182 | -1.9366 | -2.0002 | 0.49 |
| 11957 | *OR1M1* | 1.0928 | 0.34843 | 0.03435 | -1.0359 | 0.49 |
| 11958 | *KRTAP9-4* | 1.4493 | 0.11434 | -0.09006 | -0.90588 | 0.49 |
| 11959 | *SGK3* | 2.4835 | -0.1856 | -0.82547 | -0.91292 | 0.49 |
| 11960 | *ANKMY1* | 2.4147 | 0.25651 | -1.1989 | -1.2592 | 0.49 |
| 11961 | *GUCD1* | 0.77129 | 0.57061 | 0.12949 | -0.35343 | 0.49 |
| 11962 | *SEPP1* | 2.9731 | -0.54302 | -0.95895 | -1.8102 | 0.49 |
| 11963 | *GAR1* | 1.8938 | -0.01417 | -0.4093 | -0.58864 | 0.49 |
| 11964 | *LIN28B* | 1.4639 | 0.29777 | -0.29199 | -0.63791 | 0.49 |
| 11965 | *NUTM2F* | 1.6966 | 0.32175 | -0.54875 | -1.067 | 0.49 |
| 11966 | *ABI3* | 1.6078 | 0.7295 | -0.86841 | -1.4352 | 0.49 |
| 11967 | *TBC1D12* | 2.7684 | 0.25614 | -1.5558 | -1.6618 | 0.49 |
| 11968 | *CRAT* | 2.6709 | -0.5223 | -0.68236 | -1.4684 | 0.49 |
| 11969 | *TNFRSF13B* | 1.7264 | 0.71252 | -0.9733 | -1.2615 | 0.49 |
| 11970 | *RTP1* | 1.5899 | 0.70589 | -0.83022 | -1.6796 | 0.49 |
| 11971 | *DNAJC13* | 2.5312 | -0.33152 | -0.73443 | -0.93465 | 0.49 |
| 11972 | *BNIP3L* | 1.201 | 0.61617 | -0.35205 | -1.5502 | 0.49 |
| 11973 | *C6orf226* | 1.205 | 0.68556 | -0.42634 | -1.6567 | 0.49 |
| 11974 | *CSPG4* | 2.7453 | 0.67464 | -1.9559 | -2.4172 | 0.49 |
| 11975 | *TAF8* | 2.3383 | -0.03686 | -0.83749 | -0.85712 | 0.49 |
| 11976 | *C2orf88* | 1.0257 | 0.23725 | 0.20075 | 0.19377 | 0.49 |
| 11977 | *NT5DC3* | 0.82089 | 0.74017 | -0.09789 | -0.24348 | 0.49 |
| 11978 | *RXFP2* | 2.248 | -0.37461 | -0.41025 | -1.2034 | 0.49 |
| 11979 | *RBCK1* | 2.1339 | 1.2014 | -1.8724 | -2.796 | 0.49 |
| 11980 | *KLHL15* | 2.0294 | 1.5741 | -2.1419 | -2.508 | 0.49 |
| 11981 | *CPQ* | 3.6707 | -1.0723 | -1.137 | -1.8016 | 0.49 |
| 11982 | *ADAD2* | 1.0871 | 0.26195 | 0.11164 | -1.9727 | 0.49 |
| 11983 | *SLC3A2* | 1.5314 | 0.39341 | -0.46424 | -1.1605 | 0.49 |
| 11984 | *ICAM3* | 2.55 | 0.52271 | -1.6128 | -1.7428 | 0.49 |
| 11985 | *MYOD1* | 2.9463 | -0.34048 | -1.1461 | -1.9172 | 0.49 |
| 11986 | *ZNF846* | 1.0688 | 0.77299 | -0.38248 | -1.5762 | 0.49 |
| 11987 | *TCEAL8* | 1.5062 | 0.80745 | -0.85471 | -3.1921 | 0.49 |
| 11988 | *DUT* | 2.3629 | -0.42697 | -0.4771 | -0.93838 | 0.49 |
| 11989 | *CNOT6L* | 1.2554 | 0.64076 | -0.43782 | -2.0771 | 0.49 |
| 11990 | *C3orf79* | 1.0049 | 0.62059 | -0.16749 | -0.25757 | 0.49 |
| 11991 | *LSM14B* | 2.4329 | 0.37263 | -1.3476 | -1.6037 | 0.49 |
| 11992 | *NMNAT2* | 2.0012 | 0.023346 | -0.56737 | -1.5355 | 0.49 |
| 11993 | *ACADL* | 2.6974 | -0.56586 | -0.67441 | -1.0942 | 0.49 |
| 11994 | *WNT2* | 2.4031 | -0.1097 | -0.83631 | -1.365 | 0.49 |
| 11995 | *SLC7A2* | 1.1609 | 1.04 | -0.74392 | -2.0871 | 0.49 |
| 11996 | *TPI1* | 1.9303 | -0.22521 | -0.24829 | -0.93726 | 0.49 |
| 11997 | *JAM3* | 0.75524 | 0.38592 | 0.31368 | -0.3513 | 0.48 |
| 11998 | *SLITRK4* | 0.91467 | 0.42618 | 0.11365 | -0.11965 | 0.48 |
| 11999 | *ACE2* | 1.7345 | -0.09482 | -0.18547 | -1.1356 | 0.48 |

| | | | | | | |
|---|---|---|---|---|---|---|
| 12000 | *NTN1* | 1.9171 | 0.2522 | -0.71553 | -1.4951 | 0.48 |
| 12001 | *TYMP* | 2.6563 | -0.44353 | -0.75913 | -1.6848 | 0.48 |
| 12002 | *PHRF1* | 1.304 | 0.46614 | -0.31724 | -1.4931 | 0.48 |
| 12003 | *SNX12* | 3.8111 | -0.76036 | -1.5988 | -2.6652 | 0.48 |
| 12004 | *TRMT13* | 2.5096 | 0.24772 | -1.3057 | -3.1086 | 0.48 |
| 12005 | *L3HYPDH* | 1.3933 | 0.22925 | -0.17105 | -0.28457 | 0.48 |
| 12006 | *UBE2N* | 4.9861 | -1.1817 | -2.3534 | -2.9772 | 0.48 |
| 12007 | *BRCA2* | 2.5997 | -0.28058 | -0.86853 | -1.1125 | 0.48 |
| 12008 | *HP* | 2.0987 | -0.12005 | -0.52826 | -1.6759 | 0.48 |
| 12009 | *PDCD6IP* | 1.2244 | 0.49944 | -0.27379 | -0.82846 | 0.48 |
| 12010 | *OR1L1* | 1.3476 | 0.22988 | -0.12795 | -0.37537 | 0.48 |
| 12011 | *CCDC90B* | 1.1528 | 0.83903 | -0.54264 | -1.3599 | 0.48 |
| 12012 | *MUC6* | 0.94066 | 0.83381 | -0.32545 | -2.4067 | 0.48 |
| 12013 | *SATL1* | 2.643 | -0.45669 | -0.7376 | -0.79573 | 0.48 |
| 12014 | *TMEM199* | 2.5477 | -0.33231 | -0.76716 | -1.8253 | 0.48 |
| 12015 | *UBXN1* | 2.6416 | 0.033945 | -1.2276 | -1.6789 | 0.48 |
| 12016 | *TRIP13* | 1.3146 | 0.33245 | -0.19924 | -0.28484 | 0.48 |
| 12017 | *LRCOL1* | 1.3594 | 0.61421 | -0.52585 | -0.97346 | 0.48 |
| 12018 | *POU4F2* | 0.89385 | 0.63328 | -0.08023 | -0.6794 | 0.48 |
| 12019 | *TCP10L* | 0.54526 | 0.48466 | 0.41678 | -0.21079 | 0.48 |
| 12020 | *TMEM63A* | 2.9562 | -0.67695 | -0.83278 | -1.18 | 0.48 |
| 12021 | *FIG4* | 2.4861 | -0.27911 | -0.76108 | -1.8611 | 0.48 |
| 12022 | *EFNB3* | 3.4828 | -0.8383 | -1.1989 | -1.8487 | 0.48 |
| 12023 | *LYPLA1* | 0.68756 | 0.49511 | 0.26203 | -1.8398 | 0.48 |
| 12024 | *TDRKH* | 0.69509 | 0.54897 | 0.20038 | -0.40359 | 0.48 |
| 12025 | *TMEM176A* | 2.7171 | -0.40646 | -0.86698 | -1.5425 | 0.48 |
| 12026 | *SUV420H2* | 1.0724 | 0.82342 | -0.45219 | -2.3629 | 0.48 |
| 12027 | *STC2* | 1.0594 | 0.23583 | 0.14783 | -1.2815 | 0.48 |
| 12028 | *GPR12* | 1.0333 | 0.21956 | 0.19019 | -1.8376 | 0.48 |
| 12029 | *RGS7* | 2.6096 | 0.52793 | -1.6951 | -1.8341 | 0.48 |
| 12030 | *CARD8* | 1.1804 | 0.91919 | -0.65724 | -1.4765 | 0.48 |
| 12031 | *MUC20* | 1.6336 | 0.070305 | -0.26159 | -0.44417 | 0.48 |
| 12032 | *PRRG1* | 1.6194 | 1.0805 | -1.2576 | -1.7428 | 0.48 |
| 12033 | *ZNF296* | 1.0647 | 0.32273 | 0.05463 | -0.44043 | 0.48 |
| 12034 | *PROKR2* | 3.0876 | -0.58028 | -1.067 | -1.3416 | 0.48 |
| 12035 | *MEAF6* | 1.3334 | 0.16938 | -0.06308 | -0.62444 | 0.48 |
| 12036 | *ABCA1* | 1.9912 | 0.6504 | -1.2021 | -2.9638 | 0.48 |
| 12037 | *PLSCR3* | 1.517 | 0.39788 | -0.47577 | -1.3299 | 0.48 |
| 12038 | *HCN2* | 0.71037 | 0.50701 | 0.22171 | -1.7507 | 0.48 |
| 12039 | *LUZP2* | 1.5454 | 0.36606 | -0.47263 | -1.7845 | 0.48 |
| 12040 | *RGL4* | 3.834 | -0.97381 | -1.4214 | -1.4893 | 0.48 |
| 12041 | *NDUFS3* | 1.6114 | 0.60061 | -0.77346 | -1.2111 | 0.48 |
| 12042 | *UGT2B15* | 1.4536 | 0.17905 | -0.19422 | -0.33142 | 0.48 |
| 12043 | *ATP5G1* | 1.2184 | 0.90241 | -0.68303 | -2.217 | 0.48 |
| 12044 | *TMEM158* | 1.6538 | 0.5756 | -0.79167 | -1.4342 | 0.48 |
| 12045 | *PCP4L1* | 1.0994 | 0.63592 | -0.29759 | -1.3353 | 0.48 |

| | | | | | | |
|---|---|---|---|---|---|---|
| 12046 | *TAGAP* | 4.7119 | -1.285 | -1.9894 | -2.1947 | 0.48 |
| 12047 | *VCX2* | 1.5576 | 1.1187 | -1.2393 | -3.114 | 0.48 |
| 12048 | *RHOC* | 1.2213 | 0.82239 | -0.60687 | -1.2705 | 0.48 |
| 12049 | *ACOT8* | 1.0871 | 0.55181 | -0.20227 | -0.90296 | 0.48 |
| 12050 | *PIP4K2B* | 0.65469 | 0.43125 | 0.35049 | 0.28913 | 0.48 |
| 12051 | *FFAR1* | 1.5926 | 0.76464 | -0.92129 | -2.255 | 0.48 |
| 12052 | *IL27* | 2.2687 | -0.37266 | -0.46032 | -0.62947 | 0.48 |
| 12053 | *SLC2A4RG* | 0.56834 | 0.5573 | 0.30962 | -1.9295 | 0.48 |
| 12054 | *TNRC6C* | 1.0708 | 0.39376 | -0.03039 | -0.06364 | 0.48 |
| 12055 | *DOCK3* | 1.0046 | 0.80511 | -0.37602 | -1.4364 | 0.48 |
| 12056 | *HARS* | 3.4824 | -0.117 | -1.9326 | -1.9342 | 0.48 |
| 12057 | *MLLT1* | 4.0914 | -0.86445 | -1.7945 | -2.0407 | 0.48 |
| 12058 | *COASY* | 2.5863 | -0.44354 | -0.71196 | -1.268 | 0.48 |
| 12059 | *SGCZ* | 1.7462 | 0.49143 | -0.80853 | -1.0254 | 0.48 |
| 12060 | *MTRNR2L10* | 1.1073 | 0.18309 | 0.13831 | -1.1718 | 0.48 |
| 12061 | *MAL2* | 3.4772 | -0.12915 | -1.9195 | -2.0384 | 0.48 |
| 12062 | *LRRC41* | 3.0934 | -0.31126 | -1.3544 | -1.3759 | 0.48 |
| 12063 | *CDV3* | 0.83748 | 0.43539 | 0.15398 | -1.6951 | 0.48 |
| 12064 | *C1orf141* | 0.84303 | 0.67326 | -0.08948 | -0.27841 | 0.48 |
| 12065 | *HIST1H2BA* | 3.0202 | -0.53058 | -1.0635 | -1.398 | 0.48 |
| 12066 | *TCN1* | 2.9096 | -0.25048 | -1.2331 | -1.507 | 0.48 |
| 12067 | *MDN1* | 3.4545 | -0.38219 | -1.6463 | -2.0682 | 0.48 |
| 12068 | *TBC1D14* | 1.3163 | 0.41716 | -0.3077 | -0.45279 | 0.48 |
| 12069 | *ELAVL1* | 2.1798 | 0.23694 | -0.99137 | -1.2303 | 0.48 |
| 12070 | *UTP18* | 3.2162 | -0.80129 | -0.98955 | -2.2743 | 0.48 |
| 12071 | *HNF1A* | 2.563 | -0.52655 | -0.61136 | -1.6285 | 0.48 |
| 12072 | *AP3M1* | 2.7931 | 0.28077 | -1.649 | -1.7989 | 0.47 |
| 12073 | *KRT17* | 3.2045 | -0.51868 | -1.2619 | -1.2915 | 0.47 |
| 12074 | *CTDNEP1* | 2.2589 | -0.35272 | -0.4829 | -1.1823 | 0.47 |
| 12075 | *MS4A10* | 1.2674 | 0.68961 | -0.53437 | -1.1278 | 0.47 |
| 12076 | *MRPL45* | 2.7371 | -0.02083 | -1.2937 | -1.4544 | 0.47 |
| 12077 | *FCN3* | 1.5929 | 0.35194 | -0.52426 | -1.4683 | 0.47 |
| 12078 | *VSX1* | 0.79008 | 0.4525 | 0.17756 | -0.18229 | 0.47 |
| 12079 | *GCGR* | 1.8582 | 0.78817 | -1.227 | -1.8157 | 0.47 |
| 12080 | *HLA-DQA2* | 3.0648 | -0.71772 | -0.93115 | -1.5149 | 0.47 |
| 12081 | *C11orf98* | 1.448 | 1.0632 | -1.0953 | -2.2282 | 0.47 |
| 12082 | *PRB3* | 2.8249 | -0.25832 | -1.151 | -2.3804 | 0.47 |
| 12083 | *LETM1* | 2.4366 | -0.21664 | -0.80641 | -1.8366 | 0.47 |
| 12084 | *IRGQ* | 1.0313 | 0.70405 | -0.32219 | -1.1576 | 0.47 |
| 12085 | *LMNB1* | 2.1954 | 1.0209 | -1.8033 | -1.9559 | 0.47 |
| 12086 | *ZNF433* | 1.3815 | 0.22203 | -0.19136 | -2.08 | 0.47 |
| 12087 | *ZSCAN23* | 1.0776 | 0.17043 | 0.16383 | -0.95893 | 0.47 |
| 12088 | *RPL31* | 2.9126 | -0.31727 | -1.1836 | -1.4572 | 0.47 |
| 12089 | *TM4SF5* | 2.2819 | -0.35172 | -0.51912 | -1.548 | 0.47 |
| 12090 | *CECR5* | 2.8059 | -0.13105 | -1.2647 | -1.6066 | 0.47 |
| 12091 | *FRRS1* | 1.5959 | 0.08651 | -0.2723 | -1.1718 | 0.47 |

| | | | | | | |
|---|---|---|---|---|---|---|
| 12092 | *MSS51* | 2.7206 | -0.47607 | -0.83545 | -1.5815 | 0.47 |
| 12093 | *METTL12* | 0.83337 | 0.38206 | 0.19351 | -0.43694 | 0.47 |
| 12094 | *YY1AP1* | 1.3806 | 0.077177 | -0.04986 | -2.8051 | 0.47 |
| 12095 | *KRTAP10-3* | 2.53 | -0.46923 | -0.65291 | -1.0921 | 0.47 |
| 12096 | *RBP7* | 0.70137 | 0.37869 | 0.32713 | -1.0178 | 0.47 |
| 12097 | *LIMS3L* | 0.7766 | 0.44248 | 0.18809 | -0.42359 | 0.47 |
| 12098 | *SFXN4* | 0.88034 | 0.58987 | -0.0633 | -1.6058 | 0.47 |
| 12099 | *CCNJ* | 2.1956 | -0.15919 | -0.63009 | -1.882 | 0.47 |
| 12100 | *NTSR2* | 3.2184 | -0.41323 | -1.3991 | -1.5424 | 0.47 |
| 12101 | *KIF17* | 3.4979 | -0.56251 | -1.5294 | -2.217 | 0.47 |
| 12102 | *INTU* | 1.0948 | 0.26835 | 0.04252 | -0.05408 | 0.47 |
| 12103 | *MPP7* | 2.3129 | -0.10307 | -0.80436 | -1.3294 | 0.47 |
| 12104 | *C9orf16* | 2.4771 | -0.49564 | -0.57637 | -1.3319 | 0.47 |
| 12105 | *SWI5* | 1.9744 | -0.26055 | -0.30936 | -0.76606 | 0.47 |
| 12106 | *MAK* | 1.3541 | 0.51053 | -0.46268 | -1.6129 | 0.47 |
| 12107 | *MRPL27* | 1.3756 | 0.78334 | -0.758 | -2.2254 | 0.47 |
| 12108 | *SPIDR* | 0.92815 | 0.49737 | -0.02484 | -1.4894 | 0.47 |
| 12109 | *TNIP2* | 0.71918 | 0.61857 | 0.06291 | -1.344 | 0.47 |
| 12110 | *CHRDL2* | 0.72666 | 0.37293 | 0.30093 | -0.66513 | 0.47 |
| 12111 | *KIAA0368* | 1.0529 | 0.40862 | -0.06143 | -1.2692 | 0.47 |
| 12112 | *METAP1D* | 1.0045 | 0.39219 | 0.00337 | -0.69245 | 0.47 |
| 12113 | *CBX6* | 3.0439 | -0.16574 | -1.479 | -2.3491 | 0.47 |
| 12114 | *FRG1* | 2.9838 | 0.40731 | -1.9922 | -3.9836 | 0.47 |
| 12115 | *ANKRD63* | 0.54523 | 0.45584 | 0.39779 | -0.26206 | 0.47 |
| 12116 | *TCAF1* | 1.3986 | 0.016531 | -0.01665 | -2.8216 | 0.47 |
| 12117 | *KMT2D* | 1.0858 | 0.37868 | -0.066 | -1.5871 | 0.47 |
| 12118 | *ADGRE1* | 3.0091 | -0.34496 | -1.2666 | -1.2864 | 0.47 |
| 12119 | *IDNK* | 4.1665 | -0.59008 | -2.1798 | -2.9438 | 0.47 |
| 12120 | *RDM1* | 1.2362 | 1.0199 | -0.86047 | -1.9623 | 0.47 |
| 12121 | *MPPE1* | 1.48 | 0.73494 | -0.81932 | -0.99526 | 0.47 |
| 12122 | *ATP2B2* | 3.1215 | 0.24143 | -1.9711 | -3.0779 | 0.46 |
| 12123 | *TFDP2* | 1.8391 | -0.15649 | -0.29186 | -0.66146 | 0.46 |
| 12124 | *TP63* | 3.1439 | -0.59503 | -1.159 | -1.9797 | 0.46 |
| 12125 | *IGSF3* | 1.9956 | 1.0334 | -1.6394 | -2.3689 | 0.46 |
| 12126 | *RPL38* | 0.75087 | 0.33153 | 0.30635 | -1.8477 | 0.46 |
| 12127 | *CD68* | 2.9548 | -0.23832 | -1.3282 | -1.7407 | 0.46 |
| 12128 | *CD5* | 1.8319 | 0.017998 | -0.46314 | -1.7501 | 0.46 |
| 12129 | *KRTAP4-12* | 3.7156 | -0.96583 | -1.3637 | -1.8366 | 0.46 |
| 12130 | *OR51A4* | 0.68351 | 0.45434 | 0.2481 | -1.7575 | 0.46 |
| 12131 | *KHDC1* | 0.91478 | 0.79312 | -0.32254 | -1.4841 | 0.46 |
| 12132 | *ARPIN* | 0.75407 | 0.5505 | 0.08041 | 0.015519 | 0.46 |
| 12133 | *IL1RN* | 3.2349 | -0.79711 | -1.0529 | -1.9558 | 0.46 |
| 12134 | *GPBP1* | 1.1088 | 1.027 | -0.75099 | -1.7049 | 0.46 |
| 12135 | *TBCCD1* | 2.6297 | -0.00394 | -1.2411 | -2.2886 | 0.46 |
| 12136 | *NLRP9* | 3.9882 | -1.1918 | -1.4123 | -2.0837 | 0.46 |
| 12137 | *PILRA* | 0.92056 | 0.51997 | -0.05701 | -0.36476 | 0.46 |

| | | | | | | |
|---|---|---|---|---|---|---|
| 12138 | *SLC25A38* | 3.6694 | -0.74013 | -1.547 | -2.4858 | 0.46 |
| 12139 | *COQ6* | 1.3548 | 0.67522 | -0.64801 | -0.67722 | 0.46 |
| 12140 | *OSR1* | 3.7155 | -1.0313 | -1.3023 | -2.1984 | 0.46 |
| 12141 | *KCNJ15* | 2.0228 | -0.06833 | -0.57301 | -2.0488 | 0.46 |
| 12142 | *ANKMY2* | 1.1351 | 0.32753 | -0.08177 | -0.82833 | 0.46 |
| 12143 | *GTSF1L* | 1.3545 | 0.7844 | -0.75824 | -1.316 | 0.46 |
| 12144 | *CENPT* | 4.0169 | -1.2635 | -1.3729 | -1.8441 | 0.46 |
| 12145 | *RHOA* | 1.4695 | 1.1341 | -1.2234 | -1.8908 | 0.46 |
| 12146 | *PATL2* | 3.3106 | -0.83184 | -1.0993 | -1.3966 | 0.46 |
| 12147 | *ZBTB7A* | 3.3124 | -0.10427 | -1.829 | -2.112 | 0.46 |
| 12148 | *DEFB105B* | 2.4695 | -0.16038 | -0.93005 | -0.97681 | 0.46 |
| 12149 | *NINL* | 1.469 | 0.012762 | -0.10392 | -2.674 | 0.46 |
| 12150 | *PCDHGB7* | 1.8541 | 1.4692 | -1.9455 | -2.1161 | 0.46 |
| 12151 | *ZNF223* | 1.1639 | 0.93159 | -0.71772 | -1.3393 | 0.46 |
| 12152 | *THEMIS* | 1.5845 | 0.70479 | -0.91192 | -1.0297 | 0.46 |
| 12153 | *NPC1* | 1.6701 | -0.03046 | -0.26252 | -1.2135 | 0.46 |
| 12154 | *DUSP22* | 0.62919 | 0.42022 | 0.32673 | -0.17521 | 0.46 |
| 12155 | *MRPL50* | 1.1406 | 1.0095 | -0.77409 | -1.5201 | 0.46 |
| 12156 | *HOXB1* | 1.2249 | 0.19804 | -0.04713 | -0.94636 | 0.46 |
| 12157 | *PPIA* | 0.73591 | 0.6378 | 0.00068 | -0.84786 | 0.46 |
| 12158 | *ADAMTS12* | 0.97281 | 0.64991 | -0.24887 | -2.0008 | 0.46 |
| 12159 | *PPM1G* | 0.95586 | 0.63872 | -0.22097 | -1.5167 | 0.46 |
| 12160 | *KCNH4* | 1.7943 | -0.08724 | -0.33346 | -1.1135 | 0.46 |
| 12161 | *WDR36* | 1.1959 | 0.96881 | -0.79138 | -2.4363 | 0.46 |
| 12162 | *HINT2* | 4.3183 | -1.4098 | -1.5375 | -1.7647 | 0.46 |
| 12163 | *MRAS* | 1.0299 | 0.9556 | -0.61483 | -2.7254 | 0.46 |
| 12164 | *TWISTNB* | 3.8717 | -1.1244 | -1.377 | -1.8879 | 0.46 |
| 12165 | *TXNDC12* | 1.8871 | 0.58923 | -1.1061 | -2.2397 | 0.46 |
| 12166 | *RAP1GAP2* | 0.89686 | 0.34933 | 0.12305 | -0.28806 | 0.46 |
| 12167 | *TTC7B* | 2.3504 | -0.03316 | -0.94827 | -1.7377 | 0.46 |
| 12168 | *RAB11FIP2* | 0.91607 | 0.725 | -0.27291 | -1.448 | 0.46 |
| 12169 | *PRR34* | 3.1433 | -0.64078 | -1.1349 | -1.28 | 0.46 |
| 12170 | *SNX21* | 1.929 | 0.34832 | -0.91138 | -0.94027 | 0.46 |
| 12171 | *GJB3* | 2.4282 | 0.35573 | -1.4182 | -3.4131 | 0.46 |
| 12172 | *ING3* | 0.90144 | 0.89213 | -0.42817 | -0.44353 | 0.46 |
| 12173 | *ARHGEF19* | 0.94574 | 0.85733 | -0.43776 | -0.84001 | 0.46 |
| 12174 | *MAP1LC3A* | 1.4865 | 0.85764 | -0.98026 | -1.8961 | 0.45 |
| 12175 | *CCDC150* | 2.5235 | -0.41239 | -0.74743 | -1.8087 | 0.45 |
| 12176 | *CAV2* | 0.54334 | 0.41716 | 0.40281 | -1.8154 | 0.45 |
| 12177 | *TMEM261* | 0.71037 | 0.53898 | 0.11383 | -2.2392 | 0.45 |
| 12178 | *CCDC140* | 2.652 | 0.090452 | -1.3795 | -2.4094 | 0.45 |
| 12179 | *FLRT1* | 3.5105 | -0.73577 | -1.4122 | -2.3452 | 0.45 |
| 12180 | *CORO7-PAM16* | 0.60784 | 0.45211 | 0.30223 | -2.4741 | 0.45 |
| 12181 | *POLR2J3* | 2.9007 | -0.12191 | -1.4168 | -1.549 | 0.45 |
| 12182 | *LRRC74A* | 0.948 | 0.30826 | 0.1054 | -1.2505 | 0.45 |
| 12183 | *CELA2B* | 0.85669 | 0.67291 | -0.16836 | -0.20848 | 0.45 |

| | | | | | | |
|---|---|---|---|---|---|---|
| 12184 | *PYGO1* | 2.0049 | -0.26409 | -0.38059 | -1.2302 | 0.45 |
| 12185 | *STAT6* | 0.81944 | 0.36726 | 0.17331 | 0.13894 | 0.45 |
| 12186 | *ACAT1* | 2.7667 | -0.49789 | -0.9092 | -1.5448 | 0.45 |
| 12187 | *SYVN1* | 3.7634 | -0.69155 | -1.7125 | -1.9802 | 0.45 |
| 12188 | *DOK2* | 1.9382 | 0.68459 | -1.2646 | -1.2917 | 0.45 |
| 12189 | *CD79B* | 1.6278 | -0.00829 | -0.26161 | -0.63332 | 0.45 |
| 12190 | *TEX22* | 0.80612 | 0.33652 | 0.21478 | -1.5011 | 0.45 |
| 12191 | *CYP4F3* | 0.60282 | 0.56529 | 0.18914 | -0.54097 | 0.45 |
| 12192 | *ADAMTSL5* | 3.4576 | -0.90606 | -1.195 | -1.4073 | 0.45 |
| 12193 | *WDR82* | 1.9977 | -0.26892 | -0.3723 | -1.4852 | 0.45 |
| 12194 | *PCGF6* | 3.9888 | -1.2132 | -1.4197 | -1.9334 | 0.45 |
| 12195 | *RAMP1* | 0.60604 | 0.44659 | 0.30301 | -2.2998 | 0.45 |
| 12196 | *MNAT1* | 0.66539 | 0.53287 | 0.15715 | -1.5383 | 0.45 |
| 12197 | *EXOC8* | 2.5563 | -0.33723 | -0.86479 | -2.0613 | 0.45 |
| 12198 | *VMAC* | 4.0273 | -1.0942 | -1.5795 | -2.1429 | 0.45 |
| 12199 | *LACTB* | 3.3598 | -0.78229 | -1.2259 | -1.2485 | 0.45 |
| 12200 | *CAAP1* | 1.4284 | 0.761 | -0.83784 | -2.2866 | 0.45 |
| 12201 | *C12orf76* | 2.6836 | -0.55956 | -0.77333 | -0.92868 | 0.45 |
| 12202 | *CHID1* | 3.5099 | -0.3866 | -1.7735 | -2.7191 | 0.45 |
| 12203 | *XPO6* | 3.005 | -0.05908 | -1.5962 | -2.3349 | 0.45 |
| 12204 | *CYP20A1* | 1.68 | 0.38492 | -0.71522 | -0.91913 | 0.45 |
| 12205 | *TFDP1* | 4.3355 | -0.92144 | -2.0653 | -2.716 | 0.45 |
| 12206 | *SFTA3* | 2.235 | -0.12259 | -0.76366 | -1.1704 | 0.45 |
| 12207 | *TNNT3* | 1.6402 | -0.11497 | -0.17679 | -0.48572 | 0.45 |
| 12208 | *ZNF835* | 0.83528 | 0.37727 | 0.13558 | -2.1239 | 0.45 |
| 12209 | *CSNK1E* | 2.3996 | -0.50246 | -0.54934 | -1.7307 | 0.45 |
| 12210 | *NOP2* | 2.6667 | -0.5866 | -0.73362 | -1.2019 | 0.45 |
| 12211 | *RRAGA* | 0.8286 | 0.82664 | -0.30911 | -1.5992 | 0.45 |
| 12212 | *NTF4* | 1.2736 | 0.46396 | -0.39149 | -0.61496 | 0.45 |
| 12213 | *FAM193B* | 2.4773 | 0.15516 | -1.2876 | -2.1319 | 0.45 |
| 12214 | *CDKL1* | 1.2129 | 0.77163 | -0.64011 | -0.77891 | 0.45 |
| 12215 | *PTPRE* | 2.1295 | -0.24516 | -0.54207 | -1.8318 | 0.45 |
| 12216 | *DCN* | 1.9399 | -0.22235 | -0.37656 | -2.0223 | 0.45 |
| 12217 | *PCDHGA7* | 1.0599 | 0.40826 | -0.12803 | -1.3331 | 0.45 |
| 12218 | *ARHGAP9* | 3.2767 | -0.25225 | -1.6847 | -2.6635 | 0.45 |
| 12219 | *SMC3* | 4.2361 | -1.3071 | -1.5894 | -2.0613 | 0.45 |
| 12220 | *CARKD* | 0.71911 | 0.39352 | 0.22608 | -0.36847 | 0.45 |
| 12221 | *SLC24A1* | 2.9622 | -0.4978 | -1.1258 | -2.5865 | 0.45 |
| 12222 | *VTI1A* | 0.97533 | 0.47624 | -0.11355 | -1.0502 | 0.45 |
| 12223 | *SCO2* | 1.3536 | 0.042209 | -0.05841 | -0.74594 | 0.45 |
| 12224 | *FBXW4* | 3.4724 | -0.8929 | -1.2427 | -1.2457 | 0.45 |
| 12225 | *ADRB1* | 3.7835 | -1.0741 | -1.3727 | -2.3602 | 0.45 |
| 12226 | *MVB12B* | 2.3544 | -0.09789 | -0.91981 | -1.3334 | 0.45 |
| 12227 | *NDUFAF4* | 1.6324 | 0.34957 | -0.64632 | -2.2951 | 0.45 |
| 12228 | *POLR3E* | 3.3852 | -0.99298 | -1.0573 | -1.186 | 0.44 |
| 12229 | *CCL3* | 1.594 | 1.3201 | -1.5806 | -2.614 | 0.44 |

| | | | | | | |
|---|---|---|---|---|---|---|
| 12230 | *PCBP4* | 4.0792 | -0.56251 | -2.1834 | -2.329 | 0.44 |
| 12231 | *TMEM186* | 4.102 | -1.18 | -1.589 | -2.0466 | 0.44 |
| 12232 | *MAGI2* | 0.70898 | 0.33655 | 0.28742 | -0.87036 | 0.44 |
| 12233 | *ARL4D* | 1.8507 | -0.10146 | -0.41678 | -0.46155 | 0.44 |
| 12234 | *VPS33A* | 2.2285 | -0.12788 | -0.76965 | -2.2207 | 0.44 |
| 12235 | *WDR93* | 0.86552 | 0.54218 | -0.077 | -2.5725 | 0.44 |
| 12236 | *NFXL1* | 2.1551 | 0.26994 | -1.0946 | -1.9919 | 0.44 |
| 12237 | *ATP6V1A* | 0.82103 | 0.32228 | 0.18664 | 0.17094 | 0.44 |
| 12238 | *TAF10* | 0.7399 | 0.49478 | 0.09453 | -1.9727 | 0.44 |
| 12239 | *CDK2AP1* | 3.2659 | -0.10861 | -1.8291 | -2.6416 | 0.44 |
| 12240 | *TMEM194B* | 1.2412 | 0.11742 | -0.03046 | -1.4433 | 0.44 |
| 12241 | *C2orf57* | 3.5946 | -1.0252 | -1.2417 | -2.3244 | 0.44 |
| 12242 | *UBXN10* | 1.7681 | -0.18053 | -0.25999 | -0.44353 | 0.44 |
| 12243 | *C2orf73* | 1.3302 | 0.45389 | -0.45663 | -1.583 | 0.44 |
| 12244 | *CEP131* | 2.2629 | -0.19609 | -0.73938 | -1.0148 | 0.44 |
| 12245 | *MMP19* | 1.0599 | 0.93868 | -0.67144 | -2.5795 | 0.44 |
| 12246 | *NLRP11* | 0.91376 | 0.2124 | 0.20008 | -1.6891 | 0.44 |
| 12247 | *UBD* | 1.0437 | 0.81522 | -0.5328 | -1.4544 | 0.44 |
| 12248 | *KERA* | 0.80449 | 0.71578 | -0.19422 | -0.98653 | 0.44 |
| 12249 | *MANEA* | 1.695 | -0.02131 | -0.34767 | -1.9811 | 0.44 |
| 12250 | *SPRR2A* | 1.114 | 0.67755 | -0.46565 | -1.2929 | 0.44 |
| 12251 | *MS4A1* | 1.2934 | 0.37684 | -0.34496 | -1.2666 | 0.44 |
| 12252 | *SSPO* | 2.3082 | -0.3586 | -0.62456 | -1.6476 | 0.44 |
| 12253 | *FBXO30* | 1.5884 | 0.47311 | -0.73708 | -2.9032 | 0.44 |
| 12254 | *SI* | 0.82519 | 0.47351 | 0.0251 | -1.8523 | 0.44 |
| 12255 | *INSL4* | 0.56519 | 0.44924 | 0.30727 | -0.3931 | 0.44 |
| 12256 | *ENAH* | 0.86519 | 0.275 | 0.18074 | -0.38618 | 0.44 |
| 12257 | *GSC* | 0.90619 | 0.29909 | 0.11502 | -2.6416 | 0.44 |
| 12258 | *C10orf71* | 2.2278 | -0.33565 | -0.57241 | -0.63035 | 0.44 |
| 12259 | *LGSN* | 1.4375 | 0.258 | -0.37797 | -0.46052 | 0.44 |
| 12260 | *ZBED6* | 3.5621 | -1.0369 | -1.209 | -1.5592 | 0.44 |
| 12261 | *MRPL35* | 3.3942 | -0.78956 | -1.2908 | -1.984 | 0.44 |
| 12262 | *MEP1B* | 2.9532 | 0.083863 | -1.7235 | -1.9559 | 0.44 |
| 12263 | *PAX3* | 3.8667 | -0.92916 | -1.6246 | -1.7149 | 0.44 |
| 12264 | *KCNH3* | 1.6898 | 0.31558 | -0.69362 | -1.0232 | 0.44 |
| 12265 | *AVEN* | 1.3749 | 0.82811 | -0.89155 | -1.1025 | 0.44 |
| 12266 | *NCLN* | 3.0871 | -0.66363 | -1.1122 | -2.1197 | 0.44 |
| 12267 | *DDX52* | 3.6708 | -0.77908 | -1.5805 | -1.6192 | 0.44 |
| 12268 | *KLHL36* | 0.98771 | 0.26017 | 0.06281 | -1.9302 | 0.44 |
| 12269 | *CBY1* | 2.496 | 0.042086 | -1.2276 | -1.6826 | 0.44 |
| 12270 | *TBCC* | 1.4519 | 0.34019 | -0.482 | -2.213 | 0.44 |
| 12271 | *CXCL8* | 3.1787 | -0.81496 | -1.0542 | -2.5006 | 0.44 |
| 12272 | *SERPINB10* | 1.9259 | 0.4676 | -1.084 | -1.0955 | 0.44 |
| 12273 | *SND1* | 3.1901 | -0.73398 | -1.1468 | -1.3651 | 0.44 |
| 12274 | *ETF1* | 1.2853 | 0.55925 | -0.53619 | -2.5153 | 0.44 |
| 12275 | *MARS2* | 2.917 | -0.53766 | -1.0711 | -1.4125 | 0.44 |

| | | | | | | |
|---|---|---|---|---|---|---|
| 12276 | *CAND1* | 1.6326 | 1.1943 | -1.5189 | -1.686 | 0.44 |
| 12277 | *TMOD2* | 1.2891 | 0.016479 | 0.00226 | -0.5795 | 0.44 |
| 12278 | *LTB4R* | 1.5375 | -0.05094 | -0.18068 | -1.4156 | 0.44 |
| 12279 | *LMAN1* | 0.58548 | 0.53298 | 0.18677 | -1.9591 | 0.44 |
| 12280 | *WSB2* | 3.0778 | -0.80641 | -0.96647 | -1.0639 | 0.43 |
| 12281 | *SLC36A4* | 2.3216 | 0.013259 | -1.0302 | -2.5007 | 0.43 |
| 12282 | *TRIM49D1* | 0.98916 | 0.55696 | -0.24157 | -1.1287 | 0.43 |
| 12283 | *ANXA7* | 3.8285 | -1.1692 | -1.3549 | -1.6559 | 0.43 |
| 12284 | *TRAPPC3L* | 2.1574 | -0.23777 | -0.61528 | -1.7585 | 0.43 |
| 12285 | *GABARAPL2* | 2.2982 | -0.33705 | -0.65733 | -2.2653 | 0.43 |
| 12286 | *COL10A1* | 1.3781 | 0.22737 | -0.30181 | -1.2224 | 0.43 |
| 12287 | *GRAP2* | 1.0117 | 0.51997 | -0.22852 | -0.9968 | 0.43 |
| 12288 | *KIAA1107* | 1.5078 | 0.77485 | -0.97982 | -1.2679 | 0.43 |
| 12289 | *MTRNR2L4* | 2.3813 | 0.12888 | -1.2075 | -1.4361 | 0.43 |
| 12290 | *CTR9* | 1.9989 | 0.041704 | -0.7385 | -1.6835 | 0.43 |
| 12291 | *GNE* | 2.0708 | 0.23504 | -1.0041 | -1.0741 | 0.43 |
| 12292 | *SHISA2* | 3.463 | -0.43471 | -1.7266 | -1.7577 | 0.43 |
| 12293 | *SIRT5* | 3.1108 | -0.44353 | -1.3668 | -1.4864 | 0.43 |
| 12294 | *ZNF141* | 3.2152 | 0.051769 | -1.9673 | -2.5934 | 0.43 |
| 12295 | *TREML2* | 3.5391 | -0.86298 | -1.377 | -1.864 | 0.43 |
| 12296 | *KIFC2* | 1.3236 | 0.19646 | -0.22133 | -2.0383 | 0.43 |
| 12297 | *UBQLN1* | 2.5295 | 0.13942 | -1.3711 | -2.1648 | 0.43 |
| 12298 | *ZCCHC7* | 2.1331 | -0.28807 | -0.54791 | -0.58892 | 0.43 |
| 12299 | *PM20D1* | 2.2163 | -0.36401 | -0.55554 | -1.6756 | 0.43 |
| 12300 | *ZCRB1* | 2.7744 | -0.29414 | -1.1848 | -2.3222 | 0.43 |
| 12301 | *SEMA3E* | 2.6069 | -0.51467 | -0.79711 | -1.6682 | 0.43 |
| 12302 | *GOLT1B* | 0.69366 | 0.36994 | 0.23091 | -1.3959 | 0.43 |
| 12303 | *SLMO1* | 0.56109 | 0.48778 | 0.24494 | -0.14701 | 0.43 |
| 12304 | *PPARA* | 2.1475 | -0.38667 | -0.46761 | -0.55844 | 0.43 |
| 12305 | *CCNC* | 0.74926 | 0.42975 | 0.11383 | -0.54373 | 0.43 |
| 12306 | *PLA2G4E* | 1.0717 | 0.41875 | -0.19869 | -0.19996 | 0.43 |
| 12307 | *KRTAP17-1* | 4.2222 | -1.1898 | -1.7407 | -2.1354 | 0.43 |
| 12308 | *ENTPD6* | 1.6019 | -0.06967 | -0.24066 | -0.42884 | 0.43 |
| 12309 | *CSNK1G1* | 1.2605 | 1.21 | -1.179 | -1.7037 | 0.43 |
| 12310 | *KLF16* | 3.4714 | -0.96661 | -1.2135 | -2.9298 | 0.43 |
| 12311 | *POLR2M* | 1.7236 | 0.05102 | -0.48336 | -1.8961 | 0.43 |
| 12312 | *PYCARD* | 3.0603 | -0.11824 | -1.6525 | -1.7936 | 0.43 |
| 12313 | *POGLUT1* | 3.7317 | -1.03 | -1.4132 | -1.4864 | 0.43 |
| 12314 | *ACER1* | 4.1174 | -0.79243 | -2.0387 | -2.1732 | 0.43 |
| 12315 | *HRASLS* | 0.58555 | 0.39867 | 0.30155 | -0.58888 | 0.43 |
| 12316 | *MYLK3* | 3.3958 | -0.88507 | -1.2251 | -1.2318 | 0.43 |
| 12317 | *PCNXL3* | 1.5524 | -0.04556 | -0.22209 | -1.3948 | 0.43 |
| 12318 | *ZNF517* | 2.1909 | 0.42299 | -1.3292 | -1.6741 | 0.43 |
| 12319 | *LOC650293* | 4.1391 | -0.96375 | -1.8909 | -2.069 | 0.43 |
| 12320 | *ZNF358* | 2.3645 | -0.16681 | -0.91456 | -1.4108 | 0.43 |
| 12321 | *GNG13* | 1.8285 | -0.1421 | -0.40338 | -0.61112 | 0.43 |

| | | | | | | |
|---|---|---|---|---|---|---|
| 12322 | *C1D* | 1.1117 | 0.85238 | -0.68122 | -1.5167 | 0.43 |
| 12323 | *USP17L17* | 3.713 | -0.24946 | -2.1807 | -2.829 | 0.43 |
| 12324 | *PRRC2C* | 0.58665 | 0.40273 | 0.29267 | 0.068606 | 0.43 |
| 12325 | *PBX3* | 0.77641 | 0.4888 | 0.01653 | 0.000252 | 0.43 |
| 12326 | *FAM92B* | 2.0952 | -0.16294 | -0.65092 | -1.6904 | 0.43 |
| 12327 | *HPS1* | 1.1576 | 0.42876 | -0.30609 | -0.49266 | 0.43 |
| 12328 | *CNBP* | 1.3146 | 0.20971 | -0.24428 | -1.6322 | 0.43 |
| 12329 | *ATP12A* | 1.3455 | 0.70081 | -0.76702 | -1.2034 | 0.43 |
| 12330 | *GJC2* | 4.2696 | -1.4388 | -1.5522 | -1.8462 | 0.43 |
| 12331 | *PDZD11* | 2.718 | -0.09164 | -1.3478 | -1.9404 | 0.43 |
| 12332 | *CYP4V2* | 0.6088 | 0.39713 | 0.26891 | -0.46024 | 0.42 |
| 12333 | *EVPL* | 3.5183 | -0.87694 | -1.3666 | -1.6193 | 0.42 |
| 12334 | *DNAJA4* | 3.0408 | -0.41087 | -1.3554 | -2.0953 | 0.42 |
| 12335 | *CD200R1* | 1.4517 | -0.0813 | -0.09703 | -0.82452 | 0.42 |
| 12336 | *HELLS* | 1.8296 | -0.0236 | -0.53285 | -0.79798 | 0.42 |
| 12337 | *LRP10* | 1.5809 | 0.88098 | -1.1897 | -1.4758 | 0.42 |
| 12338 | *CBX1* | 2.7317 | -0.66174 | -0.79983 | -0.86772 | 0.42 |
| 12339 | *PDK2* | 2.3356 | 0.50465 | -1.5707 | -1.635 | 0.42 |
| 12340 | *FAM169B* | 1.1482 | 0.45281 | -0.33152 | -0.98574 | 0.42 |
| 12341 | *RFC4* | 0.84092 | 0.67435 | -0.24715 | -1.7065 | 0.42 |
| 12342 | *ODF2L* | 1.3725 | 1.3575 | -1.4625 | -1.5848 | 0.42 |
| 12343 | *APOA1* | 0.62089 | 0.61252 | 0.034 | -0.28228 | 0.42 |
| 12344 | *HIST1H1A* | 3.2505 | 0.038212 | -2.0226 | -2.4094 | 0.42 |
| 12345 | *GCK* | 0.92534 | 0.45421 | -0.11492 | -0.35352 | 0.42 |
| 12346 | *SCFD2* | 2.6985 | -0.62792 | -0.80613 | -1.1547 | 0.42 |
| 12347 | *STARD13* | 1.386 | 0.54697 | -0.66992 | -0.72825 | 0.42 |
| 12348 | *TMC1* | 3.7512 | -0.73247 | -1.7567 | -2.0197 | 0.42 |
| 12349 | *NOVA2* | 1.2202 | 0.40338 | -0.3617 | -1.8371 | 0.42 |
| 12350 | *ANKRD44* | 0.50283 | 0.42627 | 0.33245 | -2.6354 | 0.42 |
| 12351 | *AHCYL2* | 1.6628 | 0.28061 | -0.682 | -1.3089 | 0.42 |
| 12352 | *RBSN* | 1.1566 | 0.9493 | -0.84457 | -1.4469 | 0.42 |
| 12353 | *FBXL17* | 2.3585 | 0.048765 | -1.146 | -1.8207 | 0.42 |
| 12354 | *PRSS38* | 1.9621 | 0.1469 | -0.8478 | -2.3517 | 0.42 |
| 12355 | *KRT1* | 1.3332 | 0.30066 | -0.37282 | -1.1344 | 0.42 |
| 12356 | *SYNPR* | 1.2537 | 0.14818 | -0.14104 | -1.5524 | 0.42 |
| 12357 | *ZNF124* | 1.1996 | 0.15759 | -0.09647 | -0.12613 | 0.42 |
| 12358 | *ASH1L* | 2.5985 | -0.52329 | -0.81536 | -0.94171 | 0.42 |
| 12359 | *XAB2* | 1.8609 | 0.10845 | -0.71009 | -1.1242 | 0.42 |
| 12360 | *PGA4* | 1.2977 | 0.37869 | -0.41716 | -1.5431 | 0.42 |
| 12361 | *PHEX* | 2.7428 | -0.43376 | -1.0503 | -1.3132 | 0.42 |
| 12362 | *CHGA* | 1.6102 | 0.61348 | -0.966 | -2.0946 | 0.42 |
| 12363 | *SKOR1* | 1.4199 | 1.2327 | -1.3951 | -1.6895 | 0.42 |
| 12364 | *ADAP2* | 2.5678 | -0.15626 | -1.1543 | -1.4248 | 0.42 |
| 12365 | *RHBDF1* | 2.9861 | -0.26018 | -1.4688 | -2.0267 | 0.42 |
| 12366 | *ZC2HC1B* | 0.48494 | 0.42625 | 0.34559 | -0.19704 | 0.42 |
| 12367 | *C14orf166* | 1.3827 | 0.41626 | -0.54274 | -0.57379 | 0.42 |

| | | | | | | |
|---|---|---|---|---|---|---|
| 12368 | *CAPRIN2* | 2.9711 | -0.17566 | -1.5395 | -2.1463 | 0.42 |
| 12369 | *MED16* | 2.1349 | -0.2607 | -0.61827 | -1.4031 | 0.42 |
| 12370 | *BCCIP* | 0.97532 | 0.26121 | 0.0193 | -3.6612 | 0.42 |
| 12371 | *COPE* | 1.6135 | 0.31895 | -0.67702 | -0.90151 | 0.42 |
| 12372 | *SEMA6C* | 2.4037 | -0.01458 | -1.1344 | -1.6041 | 0.42 |
| 12373 | *CEP295NL* | 0.99061 | 0.22855 | 0.03549 | -1.2074 | 0.42 |
| 12374 | *NUDT1* | 1.2595 | 0.13369 | -0.13871 | -0.49724 | 0.42 |
| 12375 | *REST* | 1.4296 | 0.50751 | -0.683 | -0.97776 | 0.42 |
| 12376 | *AKTIP* | 2.3749 | -0.43219 | -0.68869 | -1.6974 | 0.42 |
| 12377 | *TRPC5OS* | 1.1176 | 0.54526 | -0.40938 | -0.46724 | 0.42 |
| 12378 | *FOXO6* | 4.1202 | -0.78503 | -2.0817 | -2.3811 | 0.42 |
| 12379 | *CEBPG* | 3.4739 | -0.67702 | -1.5436 | -2.458 | 0.42 |
| 12380 | *HSPA2* | 1.4993 | -0.07816 | -0.16862 | -1.5738 | 0.42 |
| 12381 | *MOGAT2* | 1.8438 | 0.86502 | -1.4566 | -1.7874 | 0.42 |
| 12382 | *CCDC137* | 2.3439 | -0.1834 | -0.90872 | -1.1955 | 0.42 |
| 12383 | *MED15* | 0.97387 | 0.88499 | -0.60719 | -1.5424 | 0.42 |
| 12384 | *C1QL1* | 2.7299 | -0.17525 | -1.3033 | -2.0629 | 0.42 |
| 12385 | *IKBKE* | 2.6466 | -0.37982 | -1.0156 | -1.4087 | 0.42 |
| 12386 | *RPP40* | 4.216 | -1.2075 | -1.7578 | -2.1484 | 0.42 |
| 12387 | *RGS11* | 0.58978 | 0.4775 | 0.18281 | -0.67791 | 0.42 |
| 12388 | *RSPH10B* | 1.5786 | -0.10495 | -0.22525 | -1.3574 | 0.42 |
| 12389 | *CCDC79* | 0.97316 | 0.41564 | -0.14151 | -1.138 | 0.42 |
| 12390 | *GPR150* | 1.1524 | 0.89613 | -0.80146 | -0.92389 | 0.42 |
| 12391 | *KCTD10* | 0.81921 | 0.72204 | -0.29427 | -0.627 | 0.42 |
| 12392 | *PRR20D* | 0.92408 | 0.73219 | -0.41178 | -0.93621 | 0.41 |
| 12393 | *SH3D19* | 1.9194 | -0.2492 | -0.42653 | -1.4576 | 0.41 |
| 12394 | *TNKS* | 2.0574 | 0.036582 | -0.8504 | -2.397 | 0.41 |
| 12395 | *ZNF443* | 0.56964 | 0.45188 | 0.22171 | -1.3795 | 0.41 |
| 12396 | *TSPAN16* | 2.2847 | 0.63242 | -1.6739 | -2.2248 | 0.41 |
| 12397 | *ARMCX1* | 2.5255 | 0.004082 | -1.2867 | -1.8392 | 0.41 |
| 12398 | *DEFA5* | 1.2966 | 0.015107 | -0.06937 | -1.6382 | 0.41 |
| 12399 | *RXFP3* | 1.6509 | -0.08868 | -0.32001 | -0.6908 | 0.41 |
| 12400 | *PCDH18* | 2.1615 | -0.29647 | -0.62307 | -0.86261 | 0.41 |
| 12401 | *FAM162A* | 0.86161 | 0.85134 | -0.47107 | -1.4801 | 0.41 |
| 12402 | *P3H3* | 3.1448 | -0.78951 | -1.1135 | -1.1357 | 0.41 |
| 12403 | *PLXNC1* | 2.8136 | -0.17712 | -1.3949 | -1.4644 | 0.41 |
| 12404 | *PRMT3* | 1.0593 | 0.54744 | -0.36518 | -1.1584 | 0.41 |
| 12405 | *BAMBI* | 0.98874 | 0.82446 | -0.57224 | -1.4368 | 0.41 |
| 12406 | *CACNA2D3* | 3.1434 | -0.37367 | -1.5294 | -1.8463 | 0.41 |
| 12407 | *MALRD1* | 3.4233 | -0.89794 | -1.2852 | -1.3296 | 0.41 |
| 12408 | *RSL1D1* | 1.955 | 0.3437 | -1.0586 | -1.2109 | 0.41 |
| 12409 | *BPIFA2* | 3.3447 | -0.89204 | -1.213 | -1.594 | 0.41 |
| 12410 | *PLGLB1* | 2.846 | -0.48968 | -1.1169 | -2.6485 | 0.41 |
| 12411 | *JCHAIN* | 0.94889 | 0.18972 | 0.10061 | -0.19494 | 0.41 |
| 12412 | *KRT83* | 0.8907 | 0.79361 | -0.44533 | -1.6289 | 0.41 |
| 12413 | *TMX3* | 1.7643 | 0.094216 | -0.62114 | -1.5632 | 0.41 |

| | | | | | | |
|---|---|---|---|---|---|---|
| 12414 | *MCL1* | 3.2326 | -0.78221 | -1.2132 | -1.3884 | 0.41 |
| 12415 | *RAD21* | 0.48216 | 0.47261 | 0.28124 | -0.15139 | 0.41 |
| 12416 | *CHRNG* | 1.3421 | 0.45273 | -0.55958 | -0.78229 | 0.41 |
| 12417 | *AVPR1A* | 1.5693 | 0.19561 | -0.53025 | -1.267 | 0.41 |
| 12418 | *CSN3* | 2.8255 | -0.57531 | -1.0164 | -1.6866 | 0.41 |
| 12419 | *UHRF1BP1L* | 0.79112 | 0.26574 | 0.17659 | -1.72 | 0.41 |
| 12420 | *ZNF445* | 3.3905 | -0.75944 | -1.3978 | -1.4926 | 0.41 |
| 12421 | *LRR1* | 3.9188 | -1.3399 | -1.3457 | -2.1947 | 0.41 |
| 12422 | *IFI35* | 1.3216 | 0.13633 | -0.22544 | -0.93112 | 0.41 |
| 12423 | *ZNF22* | 3.2268 | -0.58985 | -1.4048 | -1.7804 | 0.41 |
| 12424 | *ACADVL* | 1.921 | 0.76167 | -1.4507 | -1.9338 | 0.41 |
| 12425 | *NR1H2* | 2.6785 | -0.50595 | -0.94113 | -1.6524 | 0.41 |
| 12426 | *SLN* | 3.1098 | -0.74972 | -1.1287 | -1.3689 | 0.41 |
| 12427 | *ZFAND4* | 2.6523 | -0.69333 | -0.72881 | -0.78672 | 0.41 |
| 12428 | *WDR33* | 1.2901 | -0.01774 | -0.04233 | -0.36579 | 0.41 |
| 12429 | *INPP5D* | 1.647 | 1.2989 | -1.7177 | -1.9242 | 0.41 |
| 12430 | *CCL21* | 0.90155 | 0.25382 | 0.07237 | 0.061247 | 0.41 |
| 12431 | *PLA2G4C* | 3.6745 | -0.49941 | -1.9479 | -1.995 | 0.41 |
| 12432 | *ZCCHC16* | 1.704 | -0.23796 | -0.23892 | -1.4574 | 0.41 |
| 12433 | *SLC43A1* | 0.79649 | 0.3146 | 0.11572 | -1.8102 | 0.41 |
| 12434 | *B4GALNT2* | 1.4314 | 0.49801 | -0.70309 | -1.0696 | 0.41 |
| 12435 | *TMC6* | 2.7436 | -0.24442 | -1.2733 | -2.2543 | 0.41 |
| 12436 | *OR2AE1* | 1.0279 | 0.15053 | 0.04735 | -0.14944 | 0.41 |
| 12437 | *CXorf56* | 1.7045 | 0.56217 | -1.0423 | -1.5801 | 0.41 |
| 12438 | *MRLN* | 2.0426 | 0.49731 | -1.3165 | -1.4946 | 0.41 |
| 12439 | *MYH6* | 2.3757 | 0.28648 | -1.4392 | -2.3065 | 0.41 |
| 12440 | *RAB6C* | 1.128 | 0.5599 | -0.46583 | -1.077 | 0.41 |
| 12441 | *ZMAT4* | 3.5507 | -1.067 | -1.2619 | -1.3637 | 0.41 |
| 12442 | *EIF5B* | 2.0513 | -0.15561 | -0.67435 | -1.2447 | 0.41 |
| 12443 | *FBXO2* | 1.3358 | 0.70635 | -0.8213 | -1.5485 | 0.41 |
| 12444 | *CALHM2* | 2.7499 | -0.37592 | -1.1533 | -1.4076 | 0.41 |
| 12445 | *B4GALNT3* | 1.8387 | 0.22286 | -0.84151 | -2.3689 | 0.41 |
| 12446 | *PDC* | 0.57427 | 0.34469 | 0.30098 | -0.72157 | 0.41 |
| 12447 | *SERPINI1* | 1.7397 | -0.1305 | -0.38937 | -3.3744 | 0.41 |
| 12448 | *AMIGO2* | 3.1173 | -0.5141 | -1.3835 | -1.5482 | 0.41 |
| 12449 | *COL6A1* | 0.98217 | 0.41953 | -0.182 | -2.1018 | 0.41 |
| 12450 | *C3orf17* | 0.99951 | 0.39299 | -0.17534 | -0.54244 | 0.41 |
| 12451 | *MRGPRX2* | 0.50483 | 0.47914 | 0.23281 | 0.055477 | 0.41 |
| 12452 | *RNF125* | 1.6787 | -0.16531 | -0.2969 | -0.47528 | 0.41 |
| 12453 | *ATXN2* | 2.1535 | -0.17241 | -0.76606 | -2.005 | 0.41 |
| 12454 | *AKAP11* | 2.7151 | -0.29739 | -1.2027 | -1.545 | 0.41 |
| 12455 | *AGRN* | 0.74396 | 0.63535 | -0.16549 | -0.77847 | 0.40 |
| 12456 | *SH2B1* | 1.6701 | -0.21833 | -0.23931 | -1.348 | 0.40 |
| 12457 | *C1orf177* | 1.1593 | 0.77524 | -0.72294 | -1.7414 | 0.40 |
| 12458 | *CDCP2* | 1.0023 | 0.74269 | -0.53362 | -1.7397 | 0.40 |
| 12459 | *FXR1* | 1.0399 | 0.98093 | -0.80954 | -1.364 | 0.40 |

| | | | | | | |
|---|---|---|---|---|---|---|
| 12460 | *XBP1* | 3.1054 | -0.83557 | -1.0589 | -2.9525 | 0.40 |
| 12461 | *CELSR2* | 0.87656 | 0.50541 | -0.17188 | -0.30355 | 0.40 |
| 12462 | *OPN1LW* | 0.93712 | 0.57856 | -0.30692 | -0.57241 | 0.40 |
| 12463 | *THAP6* | 1.8768 | -0.0359 | -0.63405 | -1.4316 | 0.40 |
| 12464 | *DEPDC4* | 1.4676 | 0.046064 | -0.30706 | -0.99037 | 0.40 |
| 12465 | *AGTRAP* | 2.3553 | -0.05456 | -1.095 | -1.4919 | 0.40 |
| 12466 | *SLC35E4* | 2.9184 | -0.78579 | -0.92861 | -1.4597 | 0.40 |
| 12467 | *RNF135* | 2.0479 | -0.12884 | -0.7167 | -1.5115 | 0.40 |
| 12468 | *ZNF648* | 0.70117 | 0.66077 | -0.1604 | -1.8503 | 0.40 |
| 12469 | *MYF5* | 1.8224 | 0.44676 | -1.0679 | -5.3431 | 0.40 |
| 12470 | *MED9* | 1.1523 | 0.10954 | -0.06126 | -0.36196 | 0.40 |
| 12471 | *PQLC2* | 3.9858 | -1.0589 | -1.7268 | -1.8348 | 0.40 |
| 12472 | *ZFP92* | 2.5611 | -0.13185 | -1.2292 | -1.7386 | 0.40 |
| 12473 | *RBMY1F* | 0.55325 | 0.49275 | 0.15316 | 0.044397 | 0.40 |
| 12474 | *CASKIN1* | 0.68774 | 0.39742 | 0.11388 | -0.24079 | 0.40 |
| 12475 | *NRXN3* | 1.4293 | -0.07204 | -0.15842 | -0.52607 | 0.40 |
| 12476 | *WFDC6* | 2.3351 | -0.31342 | -0.82311 | -0.9512 | 0.40 |
| 12477 | *SH3GLB2* | 0.64697 | 0.43672 | 0.11411 | -2.6101 | 0.40 |
| 12478 | *BRSK1* | 1.9993 | -0.35805 | -0.44353 | -1.0922 | 0.40 |
| 12479 | *ANXA9* | 1.6507 | 1.3205 | -1.7745 | -2.2955 | 0.40 |
| 12480 | *GLI3* | 3.0726 | -0.8147 | -1.0614 | -2.4328 | 0.40 |
| 12481 | *SLC22A12* | 4.0513 | -1.2208 | -1.6343 | -2.1407 | 0.40 |
| 12482 | *SCEL* | 2.2425 | -0.44353 | -0.60323 | -1.0146 | 0.40 |
| 12483 | *ZHX1-C8orf76* | 2.139 | -0.31772 | -0.62556 | -1.6682 | 0.40 |
| 12484 | *C1orf210* | 1.8282 | -0.13178 | -0.50075 | -2.2732 | 0.40 |
| 12485 | *ENO1* | 0.67486 | 0.66085 | -0.14009 | -1.7443 | 0.40 |
| 12486 | *WDR5* | 1.3544 | 0.37368 | -0.53304 | -0.95527 | 0.40 |
| 12487 | *STK3* | 0.68041 | 0.34241 | 0.17215 | -1.5467 | 0.40 |
| 12488 | *SLC25A13* | 1.3603 | 1.1396 | -1.305 | -1.9284 | 0.40 |
| 12489 | *C20orf78* | 3.351 | -0.94882 | -1.2079 | -1.4686 | 0.40 |
| 12490 | *IFNB1* | 0.52402 | 0.49202 | 0.17824 | -0.3579 | 0.40 |
| 12491 | *CRYBA2* | 2.1909 | -0.28169 | -0.71723 | -1.4234 | 0.40 |
| 12492 | *MEN1* | 3.1672 | -0.38103 | -1.5959 | -2.0409 | 0.40 |
| 12493 | *TSEN54* | 0.94568 | 0.53106 | -0.28668 | -0.66326 | 0.40 |
| 12494 | *CNTLN* | 1.2598 | 0.1167 | -0.18718 | -1.7908 | 0.40 |
| 12495 | *EXD2* | 2.1889 | 0.52568 | -1.5253 | -1.8885 | 0.40 |
| 12496 | *NPTXR* | 1.0194 | 0.67288 | -0.5059 | -2.3937 | 0.40 |
| 12497 | *MYO7B* | 2.5093 | -0.15513 | -1.1698 | -1.3987 | 0.39 |
| 12498 | *C7orf60* | 2.9673 | -0.70973 | -1.0739 | -1.3496 | 0.39 |
| 12499 | *GMEB2* | 2.2103 | -0.05784 | -0.96962 | -1.1184 | 0.39 |
| 12500 | *HSFY2* | 0.66019 | 0.57321 | -0.0509 | -2.0818 | 0.39 |
| 12501 | *MRO* | 1.1617 | 0.39939 | -0.3787 | -1.6885 | 0.39 |
| 12502 | *LOC100129940* | 1.1918 | 0.71993 | -0.72969 | -2.0807 | 0.39 |
| 12503 | *CCL24* | 0.61272 | 0.42443 | 0.14451 | -1.4511 | 0.39 |
| 12504 | *C12orf60* | 3.4148 | -0.89009 | -1.3433 | -2.0197 | 0.39 |
| 12505 | *OR2M5* | 4.4155 | -0.92255 | -2.312 | -3.2688 | 0.39 |

| | | | | | | |
|---|---|---|---|---|---|---|
| 12506 | *DGKK* | 2.7556 | -0.69287 | -0.88181 | -0.99608 | 0.39 |
| 12507 | *PRPF38A* | 3.5543 | -1.0304 | -1.344 | -2.7705 | 0.39 |
| 12508 | *NSUN5* | 0.70606 | 0.49925 | -0.02563 | -0.31508 | 0.39 |
| 12509 | *HGFAC* | 1.3746 | 0.61493 | -0.81068 | -1.697 | 0.39 |
| 12510 | *TSKS* | 0.78385 | 0.75185 | -0.35685 | -1.0441 | 0.39 |
| 12511 | *RBM15B* | 1.5555 | -0.11349 | -0.26369 | -0.5489 | 0.39 |
| 12512 | *MLLT6* | 3.9119 | -1.3597 | -1.3743 | -1.4507 | 0.39 |
| 12513 | *NLK* | 0.89921 | 0.18918 | 0.08947 | -0.31359 | 0.39 |
| 12514 | *FRMPD2* | 1.516 | -0.05369 | -0.28631 | -2.1984 | 0.39 |
| 12515 | *MAP3K10* | 2.2665 | -0.47781 | -0.61271 | -0.94172 | 0.39 |
| 12516 | *HPX* | 2.293 | -0.46405 | -0.65402 | -1.2393 | 0.39 |
| 12517 | *ELFN1* | 2.1914 | 0.31212 | -1.329 | -1.62 | 0.39 |
| 12518 | *SLC29A1* | 2.8275 | -0.2069 | -1.4467 | -2.1046 | 0.39 |
| 12519 | *ARCN1* | 1.843 | -0.30007 | -0.36913 | -1.1491 | 0.39 |
| 12520 | *RNF113B* | 1.8934 | 0.96509 | -1.6866 | -1.7235 | 0.39 |
| 12521 | *ERICH5* | 1.0502 | 0.073137 | 0.04779 | -1.8002 | 0.39 |
| 12522 | *SH3GL3* | 0.90307 | 0.21188 | 0.05538 | -1.6866 | 0.39 |
| 12523 | *ADNP* | 2.0869 | 0.3488 | -1.2656 | -1.3607 | 0.39 |
| 12524 | *KLK11* | 0.69597 | 0.4323 | 0.04174 | -2.2117 | 0.39 |
| 12525 | *TMEM120A* | 2.1326 | 0.083746 | -1.0466 | -1.5663 | 0.39 |
| 12526 | *HCST* | 2.4 | 0.071123 | -1.3023 | -1.707 | 0.39 |
| 12527 | *TP53TG3B* | 1.0101 | 0.37603 | -0.21797 | -1.05 | 0.39 |
| 12528 | *MON2* | 2.0013 | 0.52005 | -1.3542 | -1.6843 | 0.39 |
| 12529 | *PARVA* | 3.6073 | -0.46749 | -1.9727 | -2.184 | 0.39 |
| 12530 | *GOLGA8N* | 3.676 | -1.0229 | -1.4864 | -2.0476 | 0.39 |
| 12531 | *STX16* | 0.68144 | 0.34241 | 0.14234 | -0.09344 | 0.39 |
| 12532 | *TWF1* | 3.1933 | 0.51644 | -2.5442 | -2.9987 | 0.39 |
| 12533 | *OLR1* | 0.83335 | 0.52902 | -0.19704 | -1.3523 | 0.39 |
| 12534 | *POLR3K* | 1.297 | 0.21436 | -0.34796 | -0.40605 | 0.39 |
| 12535 | *OR9G4* | 0.62834 | 0.46664 | 0.06821 | -0.91343 | 0.39 |
| 12536 | *PCDH7* | 3.3359 | -0.3512 | -1.8219 | -1.9295 | 0.39 |
| 12537 | *GMFB* | 0.62056 | 0.29607 | 0.24612 | -1.3635 | 0.39 |
| 12538 | *WDR7* | 1.8496 | -0.11126 | -0.57586 | -0.89494 | 0.39 |
| 12539 | *SUV420H1* | 0.93804 | 0.38688 | -0.16263 | -0.46056 | 0.39 |
| 12540 | *MRPL1* | 0.82379 | 0.35012 | -0.01168 | -1.4567 | 0.39 |
| 12541 | *UBA6* | 1.1496 | 0.9114 | -0.89892 | -2.0105 | 0.39 |
| 12542 | *C1QB* | 3.2156 | -0.994 | -1.0596 | -1.9833 | 0.39 |
| 12543 | *LMNTD1* | 2.1087 | 0.99561 | -1.9431 | -1.9484 | 0.39 |
| 12544 | *SMC1A* | 1.5674 | 0.028439 | -0.43468 | -0.81761 | 0.39 |
| 12545 | *VPS29* | 3.0671 | -0.82833 | -1.078 | -1.4859 | 0.39 |
| 12546 | *DTD2* | 1.75 | 0.14463 | -0.73389 | -1.4564 | 0.39 |
| 12547 | *NSF* | 1.5296 | 0.57405 | -0.94345 | -1.4894 | 0.39 |
| 12548 | *TPD52* | 1.1386 | 0.078531 | -0.0576 | -1.204 | 0.39 |
| 12549 | *TMEM53* | 2.1636 | -0.17323 | -0.83101 | -1.4819 | 0.39 |
| 12550 | *DAAM2* | 1.0148 | 0.12189 | 0.02109 | -0.48204 | 0.39 |
| 12551 | *PPARG* | 1.7111 | 0.19492 | -0.74868 | -1.4154 | 0.39 |

| | | | | | | |
|---|---|---|---|---|---|---|
| 12552 | *AXIN1* | 0.91802 | 0.60926 | -0.37001 | -0.47447 | 0.39 |
| 12553 | *SOX3* | 2.622 | 0.10226 | -1.5671 | -1.8248 | 0.39 |
| 12554 | *CCDC57* | 0.5598 | 0.49216 | 0.1049 | -3.0219 | 0.39 |
| 12555 | *RAP1GAP* | 1.3651 | 0.653 | -0.86157 | -1.2493 | 0.39 |
| 12556 | *WDR17* | 1.0924 | 0.042547 | 0.02099 | -1.088 | 0.39 |
| 12557 | *IRX2* | 1.167 | 0.48815 | -0.49977 | -1.1533 | 0.39 |
| 12558 | *CCDC18* | 3.312 | -0.80301 | -1.3544 | -2.1902 | 0.38 |
| 12559 | *ZNF561* | 1.7273 | 0.42709 | -1.0005 | -2.1648 | 0.38 |
| 12560 | *CD3D* | 2.9424 | -0.6728 | -1.1169 | -2.1919 | 0.38 |
| 12561 | *GLA* | 1.3152 | 0.65146 | -0.81505 | -1.7428 | 0.38 |
| 12562 | *TESK2* | 3.5263 | -1.0204 | -1.3543 | -1.9748 | 0.38 |
| 12563 | *SLC30A6* | 1.5994 | 0.25501 | -0.70379 | -1.208 | 0.38 |
| 12564 | *DRD5* | 2.0082 | -0.38472 | -0.47395 | -1.5073 | 0.38 |
| 12565 | *SORBS1* | 1.7904 | 0.84999 | -1.4913 | -2.0539 | 0.38 |
| 12566 | *SERPINB5* | 1.9949 | 0.16475 | -1.0113 | -2.3452 | 0.38 |
| 12567 | *CADM4* | 1.2301 | 0.55468 | -0.63659 | -2.6289 | 0.38 |
| 12568 | *F2* | 1.6776 | -0.198 | -0.33205 | -1.1897 | 0.38 |
| 12569 | *TP53TG3C* | 2.8374 | -0.08047 | -1.6105 | -2.4615 | 0.38 |
| 12570 | *SPICE1* | 2.3623 | -0.08915 | -1.1271 | -2.5198 | 0.38 |
| 12571 | *EIF3M* | 4.1389 | -1.1739 | -1.8191 | -2.581 | 0.38 |
| 12572 | *DDX39B* | 0.50009 | 0.4261 | 0.21956 | -1.9341 | 0.38 |
| 12573 | *SPATA9* | 0.9219 | 0.80358 | -0.57974 | -1.2214 | 0.38 |
| 12574 | *SPATC1L* | 1.5994 | 0.39005 | -0.84542 | -1.0324 | 0.38 |
| 12575 | *GHRH* | 1.4558 | -0.10126 | -0.21054 | -1.7301 | 0.38 |
| 12576 | *FOXO3* | 1.1796 | 0.2641 | -0.30018 | -0.96946 | 0.38 |
| 12577 | *PCM1* | 3.2692 | -0.49961 | -1.6263 | -1.9251 | 0.38 |
| 12578 | *SYT11* | 1.495 | 0.60069 | -0.95244 | -1.3407 | 0.38 |
| 12579 | *ANKRD49* | 2.0232 | 0.22286 | -1.1035 | -1.6464 | 0.38 |
| 12580 | *CGB8* | 0.48296 | 0.37081 | 0.28866 | -1.3159 | 0.38 |
| 12581 | *PYCRL* | 2.3279 | -0.03667 | -1.1494 | -1.39 | 0.38 |
| 12582 | *INHBC* | 2.3811 | -0.45777 | -0.78229 | -1.0421 | 0.38 |
| 12583 | *EPS15* | 1.8058 | 0.62639 | -1.2912 | -1.301 | 0.38 |
| 12584 | *ANKRD53* | 4.3206 | -1.4024 | -1.7797 | -2.5663 | 0.38 |
| 12585 | *SIPA1L1* | 1.4119 | -0.13518 | -0.13885 | -0.66358 | 0.38 |
| 12586 | *FAP* | 2.3288 | -0.04968 | -1.1415 | -1.7338 | 0.38 |
| 12587 | *BAP1* | 1.9133 | -0.24569 | -0.53009 | -1.4835 | 0.38 |
| 12588 | *OR2G3* | 0.98954 | 0.5661 | -0.4187 | -2.0953 | 0.38 |
| 12589 | *KIR2DL4* | 1.0557 | 0.52472 | -0.44353 | -0.48535 | 0.38 |
| 12590 | *BBS9* | 1.3297 | 0.15055 | -0.34431 | -1.6842 | 0.38 |
| 12591 | *SNX15* | 1.4758 | 0.021592 | -0.36189 | -1.0576 | 0.38 |
| 12592 | *PLA2G16* | 1.7991 | 0.6944 | -1.358 | -2.7442 | 0.38 |
| 12593 | *COG6* | 0.52748 | 0.49817 | 0.1098 | -0.42403 | 0.38 |
| 12594 | *TMEM37* | 0.84595 | 0.1561 | 0.13308 | -0.88418 | 0.38 |
| 12595 | *SCG5* | 1.9338 | 0.06603 | -0.86479 | -2.7223 | 0.38 |
| 12596 | *AMOTL1* | 2.9443 | -0.23447 | -1.5752 | -2.0404 | 0.38 |
| 12597 | *NFKBIE* | 1.5783 | -0.12074 | -0.32355 | -0.80897 | 0.38 |

| | | | | | | |
|---|---|---|---|---|---|---|
| 12598 | *PCDHB15* | 1.2521 | 0.010651 | -0.12891 | -2.0423 | 0.38 |
| 12599 | *BTG4* | 2.6761 | 0.13956 | -1.6819 | -2.1709 | 0.38 |
| 12600 | *MYZAP* | 2.3724 | -0.45449 | -0.78444 | -1.0824 | 0.38 |
| 12601 | *B3GNT4* | 1.0237 | 0.59615 | -0.48648 | -0.84345 | 0.38 |
| 12602 | *NUDT18* | 1.7341 | -0.11133 | -0.48968 | -0.50355 | 0.38 |
| 12603 | *ZNF778* | 1.5988 | 0.21523 | -0.681 | -1.6098 | 0.38 |
| 12604 | *OR2L3* | 1.638 | 0.15436 | -0.65971 | -0.85329 | 0.38 |
| 12605 | *ESYT3* | 1.4442 | 0.267 | -0.57974 | -2.4084 | 0.38 |
| 12606 | *FKBP10* | 1.9618 | 0.36505 | -1.1955 | -1.4913 | 0.38 |
| 12607 | *EFCAB14* | 1.0103 | 0.15515 | -0.03422 | -0.40772 | 0.38 |
| 12608 | *RPL7L1* | 0.90753 | 0.16852 | 0.0547 | -1.209 | 0.38 |
| 12609 | *PAFAH1B2* | 1.6945 | -0.12039 | -0.44353 | -0.84419 | 0.38 |
| 12610 | *EBF4* | 0.82853 | 0.31938 | -0.01805 | -1.3852 | 0.38 |
| 12611 | *CUL3* | 2.6626 | -0.00698 | -1.5274 | -1.5673 | 0.38 |
| 12612 | *RMDN1* | 3.2094 | -1.0388 | -1.043 | -2.1334 | 0.38 |
| 12613 | *UBE2K* | 3.8734 | -1.1125 | -1.6339 | -2.116 | 0.38 |
| 12614 | *KRTAP22-2* | 2.5537 | -0.36233 | -1.0653 | -1.067 | 0.38 |
| 12615 | *MARS* | 1.1121 | 0.6425 | -0.62881 | -2.0337 | 0.38 |
| 12616 | *PZP* | 3.3298 | -0.90554 | -1.2985 | -2.0164 | 0.38 |
| 12617 | *HK3* | 2.2358 | -0.50569 | -0.60582 | -0.89866 | 0.37 |
| 12618 | *GPR39* | 1.2035 | 0.38825 | -0.46749 | -1.4017 | 0.37 |
| 12619 | *POPDC2* | 2.2026 | -0.12852 | -0.95224 | -2.1849 | 0.37 |
| 12620 | *GRIN2A* | 1.9915 | 0.37011 | -1.2437 | -1.9705 | 0.37 |
| 12621 | *CCDC120* | 1.0853 | 0.12666 | -0.09418 | -0.71768 | 0.37 |
| 12622 | *SLC4A9* | 3.0899 | -0.80004 | -1.1723 | -1.7061 | 0.37 |
| 12623 | *C11orf42* | 0.98977 | 0.9548 | -0.82789 | -1.3282 | 0.37 |
| 12624 | *OSBPL3* | 0.89475 | 0.80088 | -0.57899 | -0.71381 | 0.37 |
| 12625 | *PMVK* | 2.4701 | -0.38472 | -0.96878 | -1.257 | 0.37 |
| 12626 | *KDSR* | 3.5736 | -0.83631 | -1.6213 | -1.7466 | 0.37 |
| 12627 | *SLC1A5* | 0.71094 | 0.27348 | 0.13096 | -1.2543 | 0.37 |
| 12628 | *NPAS1* | 1.9184 | -0.0014 | -0.80168 | -1.6676 | 0.37 |
| 12629 | *KLF9* | 0.92427 | 0.67866 | -0.48801 | -0.64514 | 0.37 |
| 12630 | *SLC24A4* | 0.53037 | 0.53031 | 0.05419 | -0.71381 | 0.37 |
| 12631 | *GSG2* | 2.7351 | -0.34036 | -1.2815 | -1.8938 | 0.37 |
| 12632 | *JMJD7-PLA2G4B* | 2.2336 | -0.09964 | -1.0208 | -1.2444 | 0.37 |
| 12633 | *CCDC7* | 3.1345 | -0.87601 | -1.1456 | -1.1739 | 0.37 |
| 12634 | *AARS* | 2.3376 | -0.28336 | -0.94184 | -1.0884 | 0.37 |
| 12635 | *HTT* | 2.137 | -0.39731 | -0.62747 | -1.2281 | 0.37 |
| 12636 | *TAPBPL* | 0.67758 | 0.57053 | -0.13746 | -1.1637 | 0.37 |
| 12637 | *PANX3* | 0.61736 | 0.35453 | 0.1385 | -1.677 | 0.37 |
| 12638 | *TCN2* | 1.2702 | 0.63945 | -0.79932 | -1.0933 | 0.37 |
| 12639 | *PHF2* | 1.8875 | -0.12971 | -0.64793 | -2.0017 | 0.37 |
| 12640 | *OR51E2* | 0.67857 | 0.46392 | -0.03356 | -1.653 | 0.37 |
| 12641 | *KIAA2022* | 1.8687 | 0.86367 | -1.6237 | -1.9541 | 0.37 |
| 12642 | *CROT* | 2.4427 | -0.15766 | -1.1764 | -2.1419 | 0.37 |
| 12643 | *RPS13* | 3.4723 | -0.51779 | -1.8461 | -2.2248 | 0.37 |

| | | | | | | |
|---|---|---|---|---|---|---|
| 12644 | *AHRR* | 1.3743 | 0.10874 | -0.37522 | -1.2815 | 0.37 |
| 12645 | *PTGFR* | 1.4609 | 0.07789 | -0.43101 | -2.4409 | 0.37 |
| 12646 | *PTPRK* | 1.3203 | 0.054536 | -0.26712 | -0.48659 | 0.37 |
| 12647 | *DCAF8* | 1.3658 | 0.55536 | -0.81348 | -2.2866 | 0.37 |
| 12648 | *PRMT7* | 0.99465 | 0.20646 | -0.09459 | -1.0611 | 0.37 |
| 12649 | *ADGRE2* | 1.6722 | -0.16614 | -0.39971 | -0.51679 | 0.37 |
| 12650 | *ACTL10* | 1.1003 | 0.61328 | -0.60744 | -2.0326 | 0.37 |
| 12651 | *GPD1L* | 1.2055 | 0.26936 | -0.36896 | -1.8227 | 0.37 |
| 12652 | *BCL7A* | 0.76171 | 0.25687 | 0.08711 | -1.6785 | 0.37 |
| 12653 | *CNOT11* | 3.1838 | -1.0346 | -1.0454 | -1.5219 | 0.37 |
| 12654 | *FAM114A2* | 3.7196 | -1.143 | -1.4732 | -1.7956 | 0.37 |
| 12655 | *PLEKHG4* | 1.0034 | 0.28222 | -0.18301 | -1.13 | 0.37 |
| 12656 | *NFIB* | 2.2605 | -0.27861 | -0.8803 | -1.9493 | 0.37 |
| 12657 | *APOC4* | 1.3564 | 0.053608 | -0.30863 | -0.63511 | 0.37 |
| 12658 | *PLB1* | 1.8987 | -0.06618 | -0.73143 | -2.0287 | 0.37 |
| 12659 | *TEAD4* | 3.3841 | -0.65714 | -1.626 | -2.3573 | 0.37 |
| 12660 | *CER1* | 0.93165 | 0.28701 | -0.11781 | -0.72728 | 0.37 |
| 12661 | *FGD2* | 1.6213 | 0.36498 | -0.88566 | -1.035 | 0.37 |
| 12662 | *ELL3* | 2.3582 | 0.21879 | -1.4764 | -1.908 | 0.37 |
| 12663 | *MEOX1* | 2.9793 | -0.92781 | -0.95105 | -1.9997 | 0.37 |
| 12664 | *ZHX2* | 1.6583 | 0.76302 | -1.3209 | -1.3218 | 0.37 |
| 12665 | *OR4K14* | 2.2171 | -0.45486 | -0.66194 | -2.098 | 0.37 |
| 12666 | *HOXC4* | 1.1088 | 0.15662 | -0.16642 | -0.46089 | 0.37 |
| 12667 | *SLC35C1* | 1.6101 | 0.61766 | -1.1288 | -2.1282 | 0.37 |
| 12668 | *CD274* | 0.49817 | 0.46215 | 0.13862 | -0.72612 | 0.37 |
| 12669 | *GNB1L* | 1.55 | 0.89957 | -1.3517 | -1.7753 | 0.37 |
| 12670 | *MAP3K19* | 2.633 | -0.26949 | -1.2662 | -1.739 | 0.37 |
| 12671 | *PFDN5* | 1.2237 | 0.080097 | -0.20663 | -2.5844 | 0.37 |
| 12672 | *CLEC12B* | 0.87871 | 0.20328 | 0.01474 | -1.9751 | 0.37 |
| 12673 | *SYPL1* | 1.0952 | 0.73364 | -0.73247 | -0.8988 | 0.37 |
| 12674 | *HSD17B8* | 1.4777 | 0.1144 | -0.49601 | -1.1169 | 0.37 |
| 12675 | *EYS* | 1.3594 | 0.62816 | -0.89155 | -1.1021 | 0.37 |
| 12676 | *SCT* | 4.4839 | -1.631 | -1.7585 | -2.5943 | 0.36 |
| 12677 | *ZDHHC11* | 2.7244 | -0.52075 | -1.1095 | -2.4652 | 0.36 |
| 12678 | *LIN52* | 3.12 | -0.69754 | -1.3285 | -1.7577 | 0.36 |
| 12679 | *OR4P4* | 1.709 | -0.02738 | -0.58822 | -1.4767 | 0.36 |
| 12680 | *PRPF40B* | 0.94025 | 0.76885 | -0.61591 | -1.2393 | 0.36 |
| 12681 | *GALNT5* | 1.4317 | 0.28284 | -0.62202 | -1.124 | 0.36 |
| 12682 | *SAXO1* | 1.7074 | -0.04763 | -0.56753 | -2.2021 | 0.36 |
| 12683 | *CPB2* | 1.413 | 0.10413 | -0.42536 | -1.4154 | 0.36 |
| 12684 | *PREP* | 0.83862 | 0.20845 | 0.04356 | -1.452 | 0.36 |
| 12685 | *KCNE2* | 2.0604 | -0.32419 | -0.6469 | -1.2867 | 0.36 |
| 12686 | *ARHGAP27* | 0.84876 | 0.62094 | -0.38044 | -2.1616 | 0.36 |
| 12687 | *AAED1* | 4.2956 | -1.4467 | -1.7602 | -2.892 | 0.36 |
| 12688 | *POTEA* | 1.6453 | 0.056297 | -0.61436 | -1.8057 | 0.36 |
| 12689 | *CCL3L3* | 0.81654 | 0.49695 | -0.22645 | -0.66654 | 0.36 |

| | | | | | | |
|---|---|---|---|---|---|---|
| 12690 | *PITRM1* | 1.6565 | 0.61894 | -1.1886 | -1.6497 | 0.36 |
| 12691 | *CCDC149* | 2.071 | -0.31981 | -0.66445 | -0.73707 | 0.36 |
| 12692 | *H2AFB1* | 2.8446 | 0.12335 | -1.8816 | -2.7042 | 0.36 |
| 12693 | *MPDU1* | 1.5424 | -0.19357 | -0.26252 | -1.8463 | 0.36 |
| 12694 | *MCHR1* | 1.9123 | -0.08716 | -0.73896 | -1.9054 | 0.36 |
| 12695 | *C11orf49* | 1.1314 | 0.64303 | -0.68889 | -1.3718 | 0.36 |
| 12696 | *STK11* | 0.75145 | 0.62682 | -0.29292 | -0.76955 | 0.36 |
| 12697 | *PHF5A* | 1.1476 | 0.1018 | -0.16549 | -1.4223 | 0.36 |
| 12698 | *ENTPD8* | 1.5496 | 0.56252 | -1.0283 | -1.6636 | 0.36 |
| 12699 | *HPGD* | 1.3354 | 0.78302 | -1.0347 | -1.3932 | 0.36 |
| 12700 | *APLF* | 0.98283 | 0.96132 | -0.86047 | -1.041 | 0.36 |
| 12701 | *KLF7* | 2.7725 | -0.4119 | -1.277 | -1.3607 | 0.36 |
| 12702 | *PLXNA4* | 0.46167 | 0.43311 | 0.18868 | -1.2864 | 0.36 |
| 12703 | *B3GALNT1* | 1.2618 | 0.83314 | -1.0129 | -1.6176 | 0.36 |
| 12704 | *RBM33* | 1.2327 | 0.60136 | -0.75205 | -1.95 | 0.36 |
| 12705 | *NT5E* | 1.24 | 0.73555 | -0.89382 | -1.2666 | 0.36 |
| 12706 | *NOL4L* | 3.0235 | -0.71987 | -1.222 | -1.5336 | 0.36 |
| 12707 | *SSR1* | 3.1489 | -0.67889 | -1.3884 | -1.4557 | 0.36 |
| 12708 | *KIAA1715* | 0.91503 | 0.64788 | -0.48131 | -1.8689 | 0.36 |
| 12709 | *SELV* | 2.6106 | -0.31109 | -1.2187 | -1.3389 | 0.36 |
| 12710 | *DNAJB12* | 1.1925 | 0.40361 | -0.51539 | -1.084 | 0.36 |
| 12711 | *MYO18B* | 3.4634 | -0.89649 | -1.4864 | -2.075 | 0.36 |
| 12712 | *OSBP2* | 2.1242 | -0.38157 | -0.66232 | -0.75639 | 0.36 |
| 12713 | *RAB4B* | 0.68447 | 0.25125 | 0.14449 | -1.0796 | 0.36 |
| 12714 | *TMEM154* | 2.5931 | -0.70212 | -0.8108 | -2.0572 | 0.36 |
| 12715 | *SLC25A3* | 2.6961 | -0.05653 | -1.5603 | -1.7213 | 0.36 |
| 12716 | *GDF15* | 1.7388 | -0.03348 | -0.6262 | -1.5302 | 0.36 |
| 12717 | *CPA1* | 1.1516 | 0.63805 | -0.71063 | -1.9495 | 0.36 |
| 12718 | *CCDC125* | 1.1277 | 0.4332 | -0.48232 | -1.5467 | 0.36 |
| 12719 | *CLDN12* | 0.75516 | 0.38432 | -0.06111 | -0.48893 | 0.36 |
| 12720 | *LRCH4* | 2.205 | -0.56036 | -0.56634 | -1.6663 | 0.36 |
| 12721 | *F8A2* | 3.6594 | -0.49423 | -2.0871 | -2.2473 | 0.36 |
| 12722 | *PRY* | 1.0289 | 0.49258 | -0.44353 | -0.7431 | 0.36 |
| 12723 | *ENO2* | 3.6901 | -0.89756 | -1.7149 | -2.4381 | 0.36 |
| 12724 | *LRRC19* | 2.0431 | 0.75207 | -1.7179 | -2.2485 | 0.36 |
| 12725 | *C1orf105* | 1.2145 | 1.0868 | -1.2241 | -2.0773 | 0.36 |
| 12726 | *PTPN12* | 0.53526 | 0.37901 | 0.16249 | -0.38553 | 0.36 |
| 12727 | *RPL28* | 3.6355 | -1.1316 | -1.4273 | -2.0146 | 0.36 |
| 12728 | *TIGD2* | 1.0013 | 0.7069 | -0.63336 | -0.94583 | 0.36 |
| 12729 | *OOEP* | 1.2678 | 0.47724 | -0.67064 | -1.3058 | 0.36 |
| 12730 | *SUMO2* | 0.5757 | 0.40177 | 0.09659 | -2.0283 | 0.36 |
| 12731 | *NAT8* | 3.4586 | -0.26713 | -2.1179 | -2.252 | 0.36 |
| 12732 | *CST5* | 2.7874 | -0.39035 | -1.3235 | -1.7667 | 0.36 |
| 12733 | *NNAT* | 3.6634 | -1.0239 | -1.5671 | -2.4858 | 0.36 |
| 12734 | *RNF166* | 1.7542 | 0.85457 | -1.5366 | -1.9938 | 0.36 |
| 12735 | *MATN4* | 3.8604 | -0.97346 | -1.8157 | -2.0113 | 0.36 |

| | | | | | | |
|---|---|---|---|---|---|---|
| 12736 | *BRPF3* | 1.2321 | -0.07537 | -0.08624 | -0.90576 | 0.36 |
| 12737 | *C1orf162* | 1.4046 | 0.35446 | -0.68944 | -1.9785 | 0.36 |
| 12738 | *TGM5* | 3.8588 | -1.3209 | -1.4688 | -1.6193 | 0.36 |
| 12739 | *RAB40A* | 0.52217 | 0.41603 | 0.13046 | -1.3593 | 0.36 |
| 12740 | *TKFC* | 1.4595 | 0.24364 | -0.63554 | -0.82921 | 0.36 |
| 12741 | *HNRNPA1L2* | 0.6695 | 0.39105 | 0.00518 | -0.50283 | 0.36 |
| 12742 | *UBTF* | 1.4784 | 0.01731 | -0.42999 | -1.9269 | 0.36 |
| 12743 | *ADM5* | 1.1349 | 0.27932 | -0.3495 | -0.44353 | 0.35 |
| 12744 | *PRAP1* | 0.93502 | 0.70866 | -0.57967 | -1.2628 | 0.35 |
| 12745 | *NAT6* | 0.76482 | 0.23542 | 0.06366 | -0.35059 | 0.35 |
| 12746 | *SERF2* | 3.5671 | -0.13486 | -2.3696 | -2.4784 | 0.35 |
| 12747 | *SPTAN1* | 1.5532 | 0.26655 | -0.75778 | -1.9455 | 0.35 |
| 12748 | *BLMH* | 2.5896 | -0.53487 | -0.99277 | -1.3235 | 0.35 |
| 12749 | *AMIGO1* | 0.7398 | 0.20384 | 0.11746 | -1.6893 | 0.35 |
| 12750 | *TTF1* | 2.5042 | -0.34292 | -1.1002 | -2.1438 | 0.35 |
| 12751 | *CPN1* | 0.7806 | 0.65953 | -0.37942 | -1.8557 | 0.35 |
| 12752 | *CCDC186* | 1.668 | 0.34895 | -0.95677 | -1.1289 | 0.35 |
| 12753 | *KLF4* | 1.292 | 0.091413 | -0.32332 | -1.4076 | 0.35 |
| 12754 | *VASH1* | 1.2198 | -0.02339 | -0.13843 | -0.60343 | 0.35 |
| 12755 | *PKNOX2* | 2.4154 | -0.03972 | -1.3184 | -1.7782 | 0.35 |
| 12756 | *SPATA2L* | 2.9895 | -0.87955 | -1.0529 | -2.4853 | 0.35 |
| 12757 | *SOGA1* | 1.7806 | -0.16018 | -0.56337 | -1.0564 | 0.35 |
| 12758 | *ALKBH5* | 0.99059 | 0.58984 | -0.52343 | -1.6759 | 0.35 |
| 12759 | *PSG11* | 2.0862 | 0.009609 | -1.039 | -1.9484 | 0.35 |
| 12760 | *ABCF2* | 2.5549 | -0.31727 | -1.181 | -1.6116 | 0.35 |
| 12761 | *THBS2* | 0.69815 | 0.63278 | -0.27443 | -0.53285 | 0.35 |
| 12762 | *NENF* | 1.697 | 0.65971 | -1.3019 | -2.462 | 0.35 |
| 12763 | *KATNBL1* | 0.81875 | 0.42847 | -0.19332 | -1.5865 | 0.35 |
| 12764 | *EEF1A2* | 0.70302 | 0.37588 | -0.02662 | -1.0092 | 0.35 |
| 12765 | *DDX20* | 0.56821 | 0.36513 | 0.11841 | -0.52372 | 0.35 |
| 12766 | *GDI1* | 1.4276 | -0.11212 | -0.2638 | -0.59028 | 0.35 |
| 12767 | *PPP1R3E* | 1.1519 | 0.72222 | -0.82255 | -1.5682 | 0.35 |
| 12768 | *CEPT1* | 0.52289 | 0.30868 | 0.21995 | -0.95895 | 0.35 |
| 12769 | *RDH16* | 1.0448 | 0.14247 | -0.13758 | -0.33091 | 0.35 |
| 12770 | *ZNF780B* | 1.2667 | 0.38033 | -0.59743 | -1.2967 | 0.35 |
| 12771 | *PAK4* | 1.1812 | 0.40829 | -0.5404 | -0.90588 | 0.35 |
| 12772 | *LHFPL4* | 4.6945 | -1.3338 | -2.312 | -2.7465 | 0.35 |
| 12773 | *ZBTB38* | 2.6089 | -0.37495 | -1.1856 | -1.877 | 0.35 |
| 12774 | *GTF2H1* | 2.7433 | -0.74367 | -0.95184 | -1.4914 | 0.35 |
| 12775 | *TMBIM4* | 0.94123 | 0.29744 | -0.1911 | -1.5225 | 0.35 |
| 12776 | *LOC100130880* | 0.81884 | 0.21512 | 0.01282 | -0.5313 | 0.35 |
| 12777 | *CCDC33* | 1.8612 | -0.3196 | -0.49565 | -1.2393 | 0.35 |
| 12778 | *HNMT* | 0.94269 | 0.43923 | -0.33689 | -1.1521 | 0.35 |
| 12779 | *SPIB* | 1.3489 | 0.49055 | -0.79453 | -2.2712 | 0.35 |
| 12780 | *ADH1C* | 0.74964 | 0.24134 | 0.05263 | -1.392 | 0.35 |
| 12781 | *EPO* | 1.2066 | 0.37252 | -0.5357 | -1.5906 | 0.35 |

| | | | | | | |
|---|---|---|---|---|---|---|
| 12782 | *LGALS16* | 3.4113 | -0.92975 | -1.4385 | -2.1342 | 0.35 |
| 12783 | *PPP1R18* | 3.9849 | -0.69865 | -2.2432 | -2.7728 | 0.35 |
| 12784 | *RIF1* | 1.4206 | 0.063606 | -0.44134 | -1.7306 | 0.35 |
| 12785 | *ATOX1* | 1.5499 | -0.06364 | -0.44353 | -0.53069 | 0.35 |
| 12786 | *SLCO1B7* | 0.47277 | 0.38592 | 0.18263 | 0.13578 | 0.35 |
| 12787 | *RAB11B* | 3.4807 | -0.78229 | -1.6571 | -2.1809 | 0.35 |
| 12788 | *KLK3* | 0.75845 | 0.33595 | -0.05495 | -0.44353 | 0.35 |
| 12789 | *COPS4* | 2.4659 | -0.55929 | -0.86744 | -1.5577 | 0.35 |
| 12790 | *ANXA5* | 3.3983 | -0.99275 | -1.367 | -1.5703 | 0.35 |
| 12791 | *POU5F1* | 1.6977 | 0.000252 | -0.65949 | -1.7595 | 0.35 |
| 12792 | *TAAR5* | 0.47011 | 0.46058 | 0.10737 | -1.3294 | 0.35 |
| 12793 | *NEDD8-MDP1* | 1.9249 | 0.35566 | -1.2438 | -1.7915 | 0.35 |
| 12794 | *CABP7* | 2.5286 | -0.0409 | -1.4523 | -1.8816 | 0.35 |
| 12795 | *PPIH* | 0.97026 | 0.11609 | -0.0515 | -1.4597 | 0.34 |
| 12796 | *CALCA* | 2.2054 | 0.011667 | -1.1825 | -1.6847 | 0.34 |
| 12797 | *BTBD3* | 1.4964 | 0.6285 | -1.091 | -1.8761 | 0.34 |
| 12798 | *LRRC37A* | 1.9782 | -0.17026 | -0.77409 | -1.2055 | 0.34 |
| 12799 | *JARID2* | 0.93333 | 0.13547 | -0.0354 | -1.492 | 0.34 |
| 12800 | *ABCG5* | 1.5898 | -0.15509 | -0.40292 | -1.3794 | 0.34 |
| 12801 | *DNAJC3* | 2.3586 | 0.14048 | -1.4683 | -2.1046 | 0.34 |
| 12802 | *LY9* | 1.5719 | 0.65411 | -1.1953 | -1.4864 | 0.34 |
| 12803 | *CBX3* | 1.2969 | 0.53779 | -0.8046 | -1.6276 | 0.34 |
| 12804 | *SLC25A2* | 1.3256 | 0.87789 | -1.1739 | -1.5525 | 0.34 |
| 12805 | *PCK1* | 1.109 | 0.41823 | -0.49778 | -0.93579 | 0.34 |
| 12806 | *DEK* | 0.80898 | 0.12633 | 0.09402 | -0.73502 | 0.34 |
| 12807 | *RAB2B* | 0.90012 | 0.18267 | -0.05348 | -0.52329 | 0.34 |
| 12808 | *DENND6A* | 3.2874 | -0.82366 | -1.4359 | -1.8297 | 0.34 |
| 12809 | *GOLGA8B* | 3.1996 | -0.93011 | -1.2417 | -2.1496 | 0.34 |
| 12810 | *UNC5D* | 1.1826 | 0.17101 | -0.32828 | -1.0206 | 0.34 |
| 12811 | *CAMK4* | 0.99363 | 0.30735 | -0.2765 | -0.72666 | 0.34 |
| 12812 | *PPP3R1* | 1.3114 | 0.10209 | -0.3891 | -1.5992 | 0.34 |
| 12813 | *TEK* | 0.8867 | 0.15992 | -0.0235 | -0.98754 | 0.34 |
| 12814 | *NRN1* | 1.4293 | 0.1293 | -0.53578 | -0.57285 | 0.34 |
| 12815 | *RBMY1D* | 0.47677 | 0.39778 | 0.148 | -1.3165 | 0.34 |
| 12816 | *ZNF408* | 0.83556 | 0.57453 | -0.38767 | -1.6525 | 0.34 |
| 12817 | *TMEM251* | 1.1578 | 0.32387 | -0.46032 | -0.53041 | 0.34 |
| 12818 | *KIAA2013* | 2.7109 | 0.056044 | -1.7458 | -2.1626 | 0.34 |
| 12819 | *MEGF11* | 0.8311 | 0.11435 | 0.0752 | -1.9967 | 0.34 |
| 12820 | *ZNF90* | 1.6769 | 0.50206 | -1.1587 | -1.8646 | 0.34 |
| 12821 | *C9orf139* | 0.64529 | 0.21262 | 0.16214 | -0.53911 | 0.34 |
| 12822 | *SERF1A* | 1.5122 | -0.20806 | -0.28472 | -1.7781 | 0.34 |
| 12823 | *BCAS1* | 0.44955 | 0.39529 | 0.17417 | -1.1807 | 0.34 |
| 12824 | *SDK2* | 1.5789 | 0.097822 | -0.65811 | -0.75491 | 0.34 |
| 12825 | *TENM1* | 0.87423 | 0.099278 | 0.04456 | 0.009954 | 0.34 |
| 12826 | *KCNE5* | 0.76777 | 0.68772 | -0.43793 | -0.86718 | 0.34 |
| 12827 | *FAM20C* | 1.5061 | 0.45617 | -0.94517 | -1.6553 | 0.34 |

| | | | | | | |
|---|---|---|---|---|---|---|
| 12828 | *ATP5J2* | 0.97501 | 0.062587 | -0.02228 | -1.0626 | 0.34 |
| 12829 | *DENR* | 2.5681 | -0.71856 | -0.83436 | -1.8157 | 0.34 |
| 12830 | *CETN3* | 0.93447 | 0.52567 | -0.44575 | -2.2825 | 0.34 |
| 12831 | *CAPG* | 1.1342 | 0.71227 | -0.83276 | -1.7802 | 0.34 |
| 12832 | *ASB12* | 0.47554 | 0.28798 | 0.24891 | -0.53529 | 0.34 |
| 12833 | *NUPR1L* | 3.2638 | -0.75304 | -1.4994 | -1.8478 | 0.34 |
| 12834 | *ACTR1A* | 0.6088 | 0.48551 | -0.08298 | -0.21262 | 0.34 |
| 12835 | *CSK* | 0.83475 | 0.41241 | -0.23607 | -2.7016 | 0.34 |
| 12836 | *SLC25A25* | 1.719 | -0.19422 | -0.51373 | -1.5393 | 0.34 |
| 12837 | *CCDC3* | 1.2282 | 0.95063 | -1.1685 | -2.1732 | 0.34 |
| 12838 | *LHX4* | 3.8991 | -0.4073 | -2.4815 | -3.2901 | 0.34 |
| 12839 | *PPME1* | 1.2824 | 0.033922 | -0.30611 | -0.34533 | 0.34 |
| 12840 | *UBE2Q2L* | 1.2694 | 0.32682 | -0.58603 | -0.94341 | 0.34 |
| 12841 | *ERCC1* | 1.638 | 0.28933 | -0.91767 | -2.1669 | 0.34 |
| 12842 | *TGFBR1* | 0.76478 | 0.4002 | -0.1564 | -2.4518 | 0.34 |
| 12843 | *CD53* | 3.0869 | -0.79225 | -1.2864 | -1.4819 | 0.34 |
| 12844 | *FAM71E2* | 1.0366 | 0.116 | -0.1449 | -1.6213 | 0.34 |
| 12845 | *ERN2* | 1.2424 | 0.25837 | -0.49396 | -0.62307 | 0.34 |
| 12846 | *RRAGD* | 2.3334 | -0.0429 | -1.2837 | -3.0124 | 0.34 |
| 12847 | *CCDC73* | 0.57736 | 0.3849 | 0.04383 | -1.7918 | 0.34 |
| 12848 | *CIB1* | 1.7136 | 0.3073 | -1.0153 | -1.7129 | 0.34 |
| 12849 | *ABL2* | 1.0093 | 0.035623 | -0.04002 | -0.23777 | 0.33 |
| 12850 | *SLC7A4* | 1.1794 | 0.73711 | -0.91253 | -0.94019 | 0.33 |
| 12851 | *CYP4F11* | 3.4966 | -1.0904 | -1.403 | -1.4894 | 0.33 |
| 12852 | *SEPT2* | 1.2831 | 0.20048 | -0.48085 | -1.3991 | 0.33 |
| 12853 | *PCDHGA12* | 1.0896 | 0.84592 | -0.93287 | -2.0452 | 0.33 |
| 12854 | *MB21D2* | 0.90285 | 0.69078 | -0.59119 | -0.97891 | 0.33 |
| 12855 | *EIF4E3* | 1.7355 | 0.57904 | -1.3127 | -1.6951 | 0.33 |
| 12856 | *CCDC42B* | 2.6125 | -0.4939 | -1.1169 | -1.6494 | 0.33 |
| 12857 | *DMTF1* | 2.1402 | -0.49295 | -0.64599 | -2.4633 | 0.33 |
| 12858 | *SSU72* | 0.91161 | 0.74669 | -0.65758 | -1.6635 | 0.33 |
| 12859 | *PROSER2* | 0.94374 | 0.078277 | -0.02138 | -1.4325 | 0.33 |
| 12860 | *CDRT15* | 0.69871 | 0.33634 | -0.03456 | -0.41703 | 0.33 |
| 12861 | *VAT1* | 1.73 | 0.93717 | -1.6672 | -2.3537 | 0.33 |
| 12862 | *CIR1* | 1.4525 | 0.49511 | -0.94777 | -1.9833 | 0.33 |
| 12863 | *KAT7* | 1.5061 | 0.19482 | -0.70136 | -2.565 | 0.33 |
| 12864 | *SAMHD1* | 2.5497 | -0.4705 | -1.0797 | -1.6216 | 0.33 |
| 12865 | *LDHA* | 2.6726 | -0.62886 | -1.0444 | -1.2189 | 0.33 |
| 12866 | *HIST1H2BD* | 0.62614 | 0.37073 | 0.00087 | -1.2525 | 0.33 |
| 12867 | *LCE2D* | 0.85771 | 0.29629 | -0.15766 | -1.2097 | 0.33 |
| 12868 | *GOLGA6L10* | 2.564 | -0.29513 | -1.2728 | -2.5361 | 0.33 |
| 12869 | *NUB1* | 1.1261 | 0.36293 | -0.49311 | -1.1387 | 0.33 |
| 12870 | *CUL5* | 1.4478 | 0.31165 | -0.76371 | -1.2736 | 0.33 |
| 12871 | *TMEM88B* | 2.2254 | -0.13461 | -1.0953 | -2.2167 | 0.33 |
| 12872 | *LINGO1* | 2.76 | -0.03338 | -1.7313 | -2.0548 | 0.33 |
| 12873 | *MEF2B* | 1.9684 | -0.47033 | -0.50339 | -1.0987 | 0.33 |

| | | | | | | |
|---|---|---|---|---|---|---|
| 12874 | *ZNF345* | 0.99926 | 0.20031 | -0.20512 | -0.94472 | 0.33 |
| 12875 | *RGL2* | 1.2872 | 0.74883 | -1.0416 | -1.4234 | 0.33 |
| 12876 | *HNRNPH1* | 1.6498 | -0.17606 | -0.47952 | -0.5328 | 0.33 |
| 12877 | *CDKAL1* | 1.8011 | 0.18336 | -0.99043 | -1.7495 | 0.33 |
| 12878 | *CA3* | 0.49798 | 0.33954 | 0.15608 | 0.13042 | 0.33 |
| 12879 | *ZSCAN10* | 1.3855 | -0.05429 | -0.33799 | -1.0863 | 0.33 |
| 12880 | *ZNF385C* | 1.9626 | -0.38198 | -0.58743 | -2.308 | 0.33 |
| 12881 | *COLGALT1* | 2.0975 | 0.22464 | -1.3292 | -1.9166 | 0.33 |
| 12882 | *TAC1* | 0.64941 | 0.32777 | 0.01511 | -1.0315 | 0.33 |
| 12883 | *MYDGF* | 0.76694 | 0.45479 | -0.23166 | -2.6445 | 0.33 |
| 12884 | *EDIL3* | 4.4831 | -1.2766 | -2.217 | -3.8314 | 0.33 |
| 12885 | *MAP4K5* | 0.99409 | 0.01805 | -0.02345 | -1.2118 | 0.33 |
| 12886 | *SYNDIG1L* | 1.7561 | 0.69943 | -1.4683 | -1.5579 | 0.33 |
| 12887 | *MCM3* | 2.8436 | -0.4933 | -1.3633 | -2.1105 | 0.33 |
| 12888 | *GPR34* | 1.1183 | 0.14582 | -0.27742 | -0.5328 | 0.33 |
| 12889 | *DPH6* | 3.6251 | -0.9027 | -1.7364 | -2.2292 | 0.33 |
| 12890 | *ABR* | 1.1225 | 0.025575 | -0.16242 | -1.1666 | 0.33 |
| 12891 | *DPY30* | 1.2807 | 0.26985 | -0.56518 | -2.2485 | 0.33 |
| 12892 | *C15orf38-AP3S2* | 2.8093 | -0.69146 | -1.1333 | -1.219 | 0.33 |
| 12893 | *SEZ6L2* | 3.2547 | -1.0999 | -1.1706 | -2.7374 | 0.33 |
| 12894 | *LMO7DN* | 0.81768 | 0.19631 | -0.03006 | -0.95753 | 0.33 |
| 12895 | *TOB2* | 1.0159 | 0.70864 | -0.74086 | -1.7134 | 0.33 |
| 12896 | *FMO5* | 1.9313 | -0.02606 | -0.92157 | -1.7846 | 0.33 |
| 12897 | *CGB1* | 0.49383 | 0.26853 | 0.22055 | -1.1617 | 0.33 |
| 12898 | *HCN4* | 1.7489 | 0.56594 | -1.3322 | -2.1829 | 0.33 |
| 12899 | *TMEM133* | 0.93403 | 0.48157 | -0.43305 | -0.44353 | 0.33 |
| 12900 | *MAP2* | 1.8656 | 0.25192 | -1.1357 | -2.2218 | 0.33 |
| 12901 | *TAOK3* | 0.69837 | 0.36116 | -0.07891 | -0.61807 | 0.33 |
| 12902 | *FUCA1* | 1.5604 | 0.24743 | -0.82729 | -1.4207 | 0.33 |
| 12903 | *ZNF239* | 0.67516 | 0.25848 | 0.04685 | -1.6422 | 0.33 |
| 12904 | *GOLM1* | 1.1608 | 0.12623 | -0.30743 | -0.35279 | 0.33 |
| 12905 | *NRG3* | 2.3178 | -0.53506 | -0.80341 | -1.3531 | 0.33 |
| 12906 | *LNX1* | 1.202 | 0.086766 | -0.3099 | -0.37196 | 0.33 |
| 12907 | *IPO9* | 2.5467 | 0.17147 | -1.7412 | -1.743 | 0.33 |
| 12908 | *ADCY9* | 0.63178 | 0.58744 | -0.24287 | -1.1317 | 0.33 |
| 12909 | *MPDZ* | 1.3681 | -0.15181 | -0.24003 | -0.8108 | 0.33 |
| 12910 | *PTTG1IP* | 0.85179 | 0.45202 | -0.32773 | -0.80548 | 0.33 |
| 12911 | *HSPA1A* | 0.75885 | 0.56457 | -0.34814 | -1.6984 | 0.33 |
| 12912 | *ALDH7A1* | 1.0849 | 0.10414 | -0.21405 | -1.4361 | 0.32 |
| 12913 | *C6orf120* | 1.1301 | -0.05863 | -0.09671 | -1.4538 | 0.32 |
| 12914 | *ZNF420* | 1.7312 | -0.32162 | -0.43488 | -0.69157 | 0.32 |
| 12915 | *AKNA* | 1.6721 | 0.40382 | -1.1021 | -1.2975 | 0.32 |
| 12916 | *DUXA* | 2.3561 | -0.36971 | -1.0126 | -2.7317 | 0.32 |
| 12917 | *PDHB* | 3.332 | -1.1595 | -1.1988 | -1.2441 | 0.32 |
| 12918 | *PCDH17* | 1.4266 | -0.15177 | -0.30209 | -0.94364 | 0.32 |
| 12919 | *GLT1D1* | 2.1387 | 0.63518 | -1.8013 | -2.0446 | 0.32 |

| | | | | | | |
|---|---|---|---|---|---|---|
| 12920 | *RFTN2* | 2.2194 | 0.045762 | -1.2935 | -1.7364 | 0.32 |
| 12921 | *ZMYM4* | 2.7648 | -0.25912 | -1.535 | -2.5725 | 0.32 |
| 12922 | *AUNIP* | 3.1188 | -1.0318 | -1.1169 | -2.6672 | 0.32 |
| 12923 | *PMF1* | 1.3071 | 0.10618 | -0.44353 | -2.0791 | 0.32 |
| 12924 | *CCDC176* | 1.0001 | 0.75181 | -0.78229 | -3.0045 | 0.32 |
| 12925 | *CD5L* | 1.0183 | 0.17256 | -0.22287 | -1.0682 | 0.32 |
| 12926 | *AGGF1* | 1.288 | 0.8237 | -1.1439 | -1.9833 | 0.32 |
| 12927 | *ZNF736* | 0.82263 | 0.69988 | -0.55531 | -1.1642 | 0.32 |
| 12928 | *TBC1D22B* | 0.61063 | 0.45104 | -0.09459 | -1.9318 | 0.32 |
| 12929 | *PLEKHG6* | 1.1295 | 0.41902 | -0.58221 | -1.6528 | 0.32 |
| 12930 | *SLC25A39* | 0.51455 | 0.38421 | 0.06361 | 0.014048 | 0.32 |
| 12931 | *OR5M8* | 1.022 | 0.36609 | -0.42708 | -1.9086 | 0.32 |
| 12932 | *CPEB2* | 0.83829 | 0.76876 | -0.64742 | -1.0857 | 0.32 |
| 12933 | *KCTD20* | 0.67581 | 0.32273 | -0.04099 | -1.089 | 0.32 |
| 12934 | *STX4* | 3.2295 | -0.68167 | -1.5913 | -2.0116 | 0.32 |
| 12935 | *TBC1D10A* | 1.0861 | 1.0455 | -1.1759 | -1.9623 | 0.32 |
| 12936 | *HIST1H2BC* | 1.5255 | -0.24569 | -0.32466 | -0.73911 | 0.32 |
| 12937 | *NOV* | 0.78974 | 0.15819 | 0.00684 | -1.2225 | 0.32 |
| 12938 | *TTC23* | 0.46167 | 0.34102 | 0.15122 | -1.186 | 0.32 |
| 12939 | *S100Z* | 2.5203 | -0.61746 | -0.94897 | -1.6952 | 0.32 |
| 12940 | *GNGT2* | 1.7149 | 0.69919 | -1.4607 | -1.7846 | 0.32 |
| 12941 | *ANKDD1A* | 0.90769 | 0.37779 | -0.33315 | -0.89316 | 0.32 |
| 12942 | *TIMMDC1* | 0.66623 | 0.5016 | -0.2157 | -1.7577 | 0.32 |
| 12943 | *FDX1L* | 3.3224 | -0.8071 | -1.5632 | -1.5757 | 0.32 |
| 12944 | *PLEKHA3* | 1.182 | 0.17427 | -0.4046 | -1.3165 | 0.32 |
| 12945 | *PURA* | 2.8444 | -0.90281 | -0.99049 | -1.5406 | 0.32 |
| 12946 | *XRCC6* | 3.8832 | -1.0671 | -1.8651 | -2.0789 | 0.32 |
| 12947 | *PSME4* | 1.413 | 0.080058 | -0.54302 | -0.78775 | 0.32 |
| 12948 | *RNASET2* | 3.9031 | -1.3878 | -1.5658 | -1.6401 | 0.32 |
| 12949 | *LUC7L* | 2.4864 | -0.54054 | -0.99676 | -1.4607 | 0.32 |
| 12950 | *LAMP2* | 0.87891 | 0.87638 | -0.80633 | -2.39 | 0.32 |
| 12951 | *SECTM1* | 0.72393 | 0.46466 | -0.24016 | -1.4498 | 0.32 |
| 12952 | *MSC* | 3.3033 | -1.1739 | -1.183 | -1.6659 | 0.32 |
| 12953 | *FAM188A* | 1.0655 | 0.33295 | -0.45246 | -0.96031 | 0.32 |
| 12954 | *OR5H1* | 0.68288 | 0.46721 | -0.20445 | -1.6555 | 0.32 |
| 12955 | *SEZ6L* | 1.7322 | -0.17647 | -0.61017 | -2.1648 | 0.32 |
| 12956 | *UBE2E1* | 2.2535 | -0.3555 | -0.95322 | -1.4511 | 0.31 |
| 12957 | *APH1B* | 1.5549 | -0.18118 | -0.42934 | -1.2937 | 0.31 |
| 12958 | *KDELC2* | 1.5905 | 0.17568 | -0.82208 | -1.1912 | 0.31 |
| 12959 | *TNFAIP8L1* | 1.6303 | 0.72478 | -1.4116 | -2.7379 | 0.31 |
| 12960 | *C7orf77* | 0.80918 | 0.75046 | -0.61641 | -1.0313 | 0.31 |
| 12961 | *DGKE* | 1.3914 | -0.07282 | -0.37563 | -0.46268 | 0.31 |
| 12962 | *PLAGL2* | 2.5981 | -0.81522 | -0.84001 | -1.9278 | 0.31 |
| 12963 | *TMEM141* | 1.9419 | -0.49095 | -0.50815 | -2.1404 | 0.31 |
| 12964 | *FASTKD3* | 1.4284 | -0.14207 | -0.34363 | -1.631 | 0.31 |
| 12965 | *DNAJC15* | 1.1809 | 0.56796 | -0.80675 | -1.338 | 0.31 |

| | | | | | | |
|---|---|---|---|---|---|---|
| 12966 | *RUVBL1* | 1.3404 | 0.45096 | -0.8504 | -1.0824 | 0.31 |
| 12967 | *CCDC25* | 1.1372 | 0.27529 | -0.47183 | -0.94897 | 0.31 |
| 12968 | *AGAP1* | 1.0901 | 0.12588 | -0.27541 | -1.7338 | 0.31 |
| 12969 | *RNF20* | 2.3173 | 0.056768 | -1.4335 | -2.7623 | 0.31 |
| 12970 | *VEGFC* | 0.55033 | 0.2175 | 0.17217 | -0.58691 | 0.31 |
| 12971 | *CHAD* | 3.6695 | -0.99699 | -1.733 | -1.7995 | 0.31 |
| 12972 | *TBR1* | 0.76489 | 0.5237 | -0.34927 | -2.1251 | 0.31 |
| 12973 | *ERICH3* | 1.1417 | 0.92367 | -1.1263 | -1.6836 | 0.31 |
| 12974 | *SNCB* | 2.1298 | -0.25295 | -0.93868 | -1.8376 | 0.31 |
| 12975 | *MTIF3* | 2.0881 | -0.00995 | -1.141 | -1.6105 | 0.31 |
| 12976 | *TFCP2L1* | 0.84537 | 0.23246 | -0.14117 | -1.8435 | 0.31 |
| 12977 | *EXO5* | 2.9241 | -0.99137 | -0.99608 | -1.9459 | 0.31 |
| 12978 | *ACTL6B* | 2.4473 | 0.020029 | -1.5311 | -2.4244 | 0.31 |
| 12979 | *MIOS* | 2.1233 | -0.29121 | -0.89593 | -0.98574 | 0.31 |
| 12980 | *FLOT1* | 0.95434 | 0.44297 | -0.46155 | -1.2346 | 0.31 |
| 12981 | *GBP2* | 2.2504 | 0.30909 | -1.6243 | -1.7911 | 0.31 |
| 12982 | *PLEKHA1* | 1.6583 | 0.23544 | -0.95868 | -1.3694 | 0.31 |
| 12983 | *TMEM178B* | 1.3228 | 0.43735 | -0.82696 | -1.1852 | 0.31 |
| 12984 | *ASB10* | 1.3815 | 0.10365 | -0.55275 | -1.658 | 0.31 |
| 12985 | *HIATL1* | 1.153 | 0.10354 | -0.32466 | -1.4055 | 0.31 |
| 12986 | *EGFL7* | 2.2986 | -0.01883 | -1.3481 | -1.772 | 0.31 |
| 12987 | *RAB6B* | 1.2885 | 0.56755 | -0.92444 | -1.94 | 0.31 |
| 12988 | *MGEA5* | 1.5611 | 0.71885 | -1.3496 | -1.9997 | 0.31 |
| 12989 | *PSMC5* | 1.9142 | 0.44422 | -1.4299 | -1.6885 | 0.31 |
| 12990 | *MOCS2* | 0.91321 | 0.29904 | -0.28391 | -2.2528 | 0.31 |
| 12991 | *DHX40* | 3.608 | -1.0212 | -1.6587 | -2.2075 | 0.31 |
| 12992 | *KIF11* | 3.168 | -0.79165 | -1.4485 | -1.7065 | 0.31 |
| 12993 | *ARRDC2* | 2.4268 | -0.4155 | -1.0846 | -1.1053 | 0.31 |
| 12994 | *RTF1* | 2.5054 | -0.17705 | -1.4022 | -1.6949 | 0.31 |
| 12995 | *VKORC1L1* | 2.1033 | -0.20423 | -0.97333 | -1.8443 | 0.31 |
| 12996 | *FRY* | 0.6994 | 0.21745 | 0.00806 | -0.44198 | 0.31 |
| 12997 | *DAAM1* | 0.8516 | 0.050931 | 0.02185 | -0.33741 | 0.31 |
| 12998 | *PRKCG* | 0.49899 | 0.48679 | -0.06177 | -1.7591 | 0.31 |
| 12999 | *MAL* | 1.0967 | 0.15078 | -0.32677 | -1.7009 | 0.31 |
| 13000 | *HSD11B1L* | 4.6868 | -1.8366 | -1.9295 | -2.3537 | 0.31 |
| 13001 | *ERO1LB* | 3.7701 | -0.99226 | -1.8572 | -2.3525 | 0.31 |
| 13002 | *KCNRG* | 2.5431 | -0.50472 | -1.1189 | -1.1856 | 0.31 |
| 13003 | *IRX5* | 2.4285 | -0.59921 | -0.91045 | -1.1357 | 0.31 |
| 13004 | *CENPH* | 1.1514 | 0.33222 | -0.56492 | -1.3591 | 0.31 |
| 13005 | *ULBP2* | 0.99062 | 0.53719 | -0.60953 | -0.95504 | 0.31 |
| 13006 | *CLCN4* | 0.77215 | 0.43717 | -0.29114 | -1.2977 | 0.31 |
| 13007 | *RERG* | 2.4874 | -0.35899 | -1.2125 | -1.4019 | 0.31 |
| 13008 | *CCDC30* | 0.57799 | 0.28275 | 0.05503 | -1.317 | 0.31 |
| 13009 | *CHST9* | 1.7734 | 0.72823 | -1.587 | -1.6648 | 0.30 |
| 13010 | *SGSM1* | 2.578 | -0.5946 | -1.0688 | -1.4498 | 0.30 |
| 13011 | *GTPBP6* | 0.83111 | 0.38135 | -0.29792 | -2.1947 | 0.30 |

| | | | | | | |
|---|---|---|---|---|---|---|
| 13012 | *OR10H5* | 1.9062 | 0.16733 | -1.1595 | -1.7533 | 0.30 |
| 13013 | *ASPSCR1* | 0.85138 | 0.17884 | -0.11729 | -0.49684 | 0.30 |
| 13014 | *ANTXR1* | 0.43191 | 0.34151 | 0.1395 | -0.42046 | 0.30 |
| 13015 | *TGS1* | 0.50604 | 0.24958 | 0.1573 | -1.8604 | 0.30 |
| 13016 | *CAMK2N2* | 2.6701 | -0.22742 | -1.5312 | -2.0218 | 0.30 |
| 13017 | *OR56B1* | 1.0411 | 0.26286 | -0.39248 | -0.82768 | 0.30 |
| 13018 | *CDK11B* | 0.60384 | 0.42522 | -0.11781 | -1.5684 | 0.30 |
| 13019 | *USP17L8* | 3.2819 | -0.6568 | -1.714 | -2.0664 | 0.30 |
| 13020 | *KLHL29* | 1.8526 | 0.20767 | -1.1492 | -1.8112 | 0.30 |
| 13021 | *SPANXC* | 0.62844 | 0.19503 | 0.08753 | -0.81155 | 0.30 |
| 13022 | *RSF1* | 2.3497 | 0.53344 | -1.9727 | -2.5007 | 0.30 |
| 13023 | *OR4A15* | 3.3344 | -1.1529 | -1.2719 | -2.6275 | 0.30 |
| 13024 | *KCNS2* | 0.62915 | 0.53561 | -0.25597 | -0.3866 | 0.30 |
| 13025 | *FOXK2* | 1.8886 | 0.29982 | -1.2803 | -3.0219 | 0.30 |
| 13026 | *ZIM3* | 0.55312 | 0.23974 | 0.11525 | -0.1435 | 0.30 |
| 13027 | *TMEM92* | 0.79659 | 0.53569 | -0.4247 | -2.22 | 0.30 |
| 13028 | *IL13* | 1.3078 | 0.9004 | -1.3007 | -1.6508 | 0.30 |
| 13029 | *TARBP1* | 0.93664 | 0.2954 | -0.32665 | -1.027 | 0.30 |
| 13030 | *SLC25A45* | 1.2424 | -0.05096 | -0.28609 | -0.90192 | 0.30 |
| 13031 | *YIPF1* | 0.64766 | 0.32361 | -0.06675 | -1.3718 | 0.30 |
| 13032 | *PEMT* | 0.54202 | 0.39341 | -0.03164 | -2.672 | 0.30 |
| 13033 | *MYO5A* | 0.75598 | 0.5179 | -0.3703 | -1.1492 | 0.30 |
| 13034 | *NARS* | 0.95433 | 0.046036 | -0.09738 | -1.4541 | 0.30 |
| 13035 | *SSFA2* | 3.2399 | -1.1322 | -1.205 | -1.7675 | 0.30 |
| 13036 | *NFYA* | 1.4406 | 0.17112 | -0.71078 | -1.2867 | 0.30 |
| 13037 | *COMMD3* | 1.3577 | -0.19331 | -0.26347 | -0.73202 | 0.30 |
| 13038 | *MRPS17* | 3.1085 | -0.60221 | -1.6069 | -1.6139 | 0.30 |
| 13039 | *TMEM25* | 1.5396 | -0.06631 | -0.57429 | -1.8714 | 0.30 |
| 13040 | *SPINK5* | 0.92109 | 0.30279 | -0.32495 | -0.9968 | 0.30 |
| 13041 | *NT5DC1* | 1.2873 | 0.20251 | -0.5911 | -1.6469 | 0.30 |
| 13042 | *FRG2B* | 1.7986 | -0.44887 | -0.45124 | -0.94493 | 0.30 |
| 13043 | *STATH* | 2.1865 | 0.36457 | -1.6526 | -1.9054 | 0.30 |
| 13044 | *VPS53* | 1.9015 | 0.32523 | -1.3294 | -2.2763 | 0.30 |
| 13045 | *ATXN7L3B* | 2.9559 | -0.45373 | -1.6049 | -2.1732 | 0.30 |
| 13046 | *FLI1* | 0.54479 | 0.4616 | -0.11145 | -0.34111 | 0.30 |
| 13047 | *MORN4* | 2.4335 | -0.14795 | -1.3911 | -1.6691 | 0.30 |
| 13048 | *CASD1* | 1.0623 | 0.66948 | -0.83739 | -1.3665 | 0.30 |
| 13049 | *LDHAL6A* | 0.87112 | 0.24958 | -0.22673 | -0.78229 | 0.30 |
| 13050 | *RTN1* | 0.9072 | 0.5381 | -0.55134 | -2.0384 | 0.30 |
| 13051 | *OR5T1* | 1.7742 | 0.42647 | -1.3068 | -2.6456 | 0.30 |
| 13052 | *OGFOD1* | 0.68585 | 0.30586 | -0.09789 | -1.2099 | 0.30 |
| 13053 | *EIF3J* | 2.7211 | -0.35891 | -1.4689 | -1.5524 | 0.30 |
| 13054 | *CENPJ* | 1.4245 | -0.12333 | -0.40793 | -0.75817 | 0.30 |
| 13055 | *NCOA5* | 1.2528 | 0.8717 | -1.232 | -1.6162 | 0.30 |
| 13056 | *ALDH2* | 1.2301 | -0.04536 | -0.29267 | -1.3617 | 0.30 |
| 13057 | *PEX1* | 2.6256 | -0.66914 | -1.0644 | -1.8792 | 0.30 |

| | | | | | | |
|---|---|---|---|---|---|---|
| 13058 | *KANK3* | 1.2802 | -0.14701 | -0.24137 | -0.6251 | 0.30 |
| 13059 | *ANKRD28* | 1.3502 | 0.11542 | -0.57429 | -1.4597 | 0.30 |
| 13060 | *BPIFB3* | 0.35456 | 0.27765 | 0.25899 | -0.61555 | 0.30 |
| 13061 | *ARNTL* | 1.5436 | 0.56698 | -1.2202 | -1.8682 | 0.30 |
| 13062 | *ZNF569* | 1.002 | 0.59361 | -0.70558 | -0.93162 | 0.30 |
| 13063 | *SMUG1* | 2.9908 | -0.80727 | -1.2936 | -1.7364 | 0.30 |
| 13064 | *TRIM71* | 0.83762 | 0.088247 | -0.03623 | -0.59647 | 0.30 |
| 13065 | *OR5A2* | 2.3062 | 0.52879 | -1.9455 | -2.6072 | 0.30 |
| 13066 | *AGFG2* | 1.0058 | 0.36145 | -0.4783 | -2.011 | 0.30 |
| 13067 | *COPA* | 2.3106 | 0.079723 | -1.5016 | -2.016 | 0.30 |
| 13068 | *IGSF6* | 2.966 | 0.45892 | -2.537 | -3.2595 | 0.30 |
| 13069 | *SLC10A2* | 1.9184 | 0.70584 | -1.7364 | -2.0946 | 0.30 |
| 13070 | *TKTL1* | 0.66358 | 0.21311 | 0.00998 | -0.5203 | 0.30 |
| 13071 | *IRF6* | 2.3461 | -0.27911 | -1.1815 | -2.4172 | 0.30 |
| 13072 | *HNRNPD* | 1.278 | -0.08654 | -0.30609 | -0.91283 | 0.30 |
| 13073 | *TAS2R5* | 0.50885 | 0.26163 | 0.11435 | -1.0751 | 0.29 |
| 13074 | *KTN1* | 2.051 | -0.07227 | -1.0942 | -1.2246 | 0.29 |
| 13075 | *LARP1* | 1.9145 | -0.14211 | -0.88804 | -0.94312 | 0.29 |
| 13076 | *CPNE7* | 2.9284 | -0.62928 | -1.4151 | -1.5611 | 0.29 |
| 13077 | *MFSD1* | 0.73954 | 0.58119 | -0.43694 | -1.0095 | 0.29 |
| 13078 | *TGFBI* | 2.494 | -0.45572 | -1.1545 | -2.5346 | 0.29 |
| 13079 | *LRGUK* | 0.56598 | 0.31369 | 0.00277 | -0.30326 | 0.29 |
| 13080 | *PLRG1* | 2.7792 | -0.8462 | -1.0509 | -1.7649 | 0.29 |
| 13081 | *KIAA0020* | 1.4202 | -0.02854 | -0.50961 | -1.2995 | 0.29 |
| 13082 | *CAPZA3* | 1.2312 | 0.44554 | -0.79474 | -1.6524 | 0.29 |
| 13083 | *PGLYRP3* | 0.7006 | 0.45098 | -0.26996 | -1.0283 | 0.29 |
| 13084 | *TRPV5* | 1.3328 | 0.33838 | -0.78959 | -1.1429 | 0.29 |
| 13085 | *NBR1* | 3.9315 | -1.0761 | -1.9748 | -2.29 | 0.29 |
| 13086 | *OTOP2* | 0.62822 | 0.15387 | 0.09832 | -2.9772 | 0.29 |
| 13087 | *EFCAB2* | 0.76572 | 0.42745 | -0.31294 | -1.9172 | 0.29 |
| 13088 | *LOC100506388* | 1.5721 | -0.10216 | -0.58985 | -1.5039 | 0.29 |
| 13089 | *ISYNA1* | 1.0127 | 0.19141 | -0.32466 | -1.5225 | 0.29 |
| 13090 | *HAO1* | 1.3415 | -0.02092 | -0.44135 | -1.2576 | 0.29 |
| 13091 | *FANCD2OS* | 1.938 | -0.20822 | -0.85199 | -1.3132 | 0.29 |
| 13092 | *DIEXF* | 0.74021 | 0.43498 | -0.29752 | -0.97393 | 0.29 |
| 13093 | *RARS2* | 3.4214 | -0.66731 | -1.877 | -1.8996 | 0.29 |
| 13094 | *GUCY2D* | 0.65784 | 0.64897 | -0.43043 | -2.907 | 0.29 |
| 13095 | *ZNF366* | 1.1717 | -0.06833 | -0.22763 | -0.83277 | 0.29 |
| 13096 | *TMEM43* | 0.9887 | 0.02792 | -0.14117 | -1.0907 | 0.29 |
| 13097 | *OR4C13* | 2.2529 | 0.034936 | -1.4128 | -1.8404 | 0.29 |
| 13098 | *C2CD4C* | 1.5024 | 0.15485 | -0.78229 | -1.4776 | 0.29 |
| 13099 | *USP17L11* | 1.1597 | 0.26433 | -0.54921 | -1.3216 | 0.29 |
| 13100 | *SLC36A3* | 0.99785 | 0.010955 | -0.13487 | -0.36821 | 0.29 |
| 13101 | *ZNF721* | 1.0198 | 0.16511 | -0.31132 | -0.95044 | 0.29 |
| 13102 | *DESI2* | 2.6248 | -0.38025 | -1.3718 | -2.0541 | 0.29 |
| 13103 | *OR51V1* | 1.7734 | -0.44353 | -0.45786 | -1.1395 | 0.29 |

| | | | | | | |
|---|---|---|---|---|---|---|
| 13104 | *QRICH2* | 2.9654 | -0.9619 | -1.1316 | -1.1651 | 0.29 |
| 13105 | *FAU* | 1.6361 | 0.14492 | -0.91013 | -1.6012 | 0.29 |
| 13106 | *SCGB1C2* | 0.70671 | 0.49407 | -0.33089 | -1.9255 | 0.29 |
| 13107 | *EIF2AK2* | 1.7968 | -0.45604 | -0.47136 | -1.6116 | 0.29 |
| 13108 | *FZD8* | 1.6382 | -0.18859 | -0.58049 | -1.3208 | 0.29 |
| 13109 | *PPP2R2C* | 0.88608 | 0.72504 | -0.74219 | -2.1502 | 0.29 |
| 13110 | *POLE* | 3.0951 | -0.6386 | -1.5884 | -2.224 | 0.29 |
| 13111 | *FAM163A* | 1.9951 | -0.00456 | -1.124 | -1.2566 | 0.29 |
| 13112 | *TTC19* | 1.3434 | 0.59503 | -1.0726 | -1.172 | 0.29 |
| 13113 | *ACSL5* | 2.2811 | -0.31885 | -1.098 | -1.6525 | 0.29 |
| 13114 | *PPIL4* | 1.2188 | 0.98977 | -1.3451 | -1.5194 | 0.29 |
| 13115 | *DHRS4L2* | 2.7064 | 0.060969 | -1.9047 | -2.0466 | 0.29 |
| 13116 | *ZNF717* | 0.77656 | 0.59128 | -0.50589 | -0.94827 | 0.29 |
| 13117 | *CLNK* | 2.2781 | -0.19053 | -1.2261 | -1.3346 | 0.29 |
| 13118 | *CDK15* | 1.3864 | 0.78479 | -1.3104 | -2.0999 | 0.29 |
| 13119 | *PLEKHG2* | 2.1257 | 0.54076 | -1.806 | -2.223 | 0.29 |
| 13120 | *SNTB1* | 0.96949 | -0.00833 | -0.10088 | -1.1553 | 0.29 |
| 13121 | *LIN7B* | 1.5701 | -0.02072 | -0.68911 | -1.7649 | 0.29 |
| 13122 | *ZMAT2* | 0.77419 | 0.37868 | -0.2928 | -0.78754 | 0.29 |
| 13123 | *ARHGAP22* | 0.96347 | 0.4644 | -0.56812 | -1.748 | 0.29 |
| 13124 | *STK32A* | 0.99547 | 0.56981 | -0.70581 | -1.3085 | 0.29 |
| 13125 | *RNF212* | 2.3765 | -0.4082 | -1.1096 | -4.7884 | 0.29 |
| 13126 | *INSL3* | 0.56732 | 0.24322 | 0.04806 | -0.17509 | 0.29 |
| 13127 | *SMIM17* | 1.7097 | -0.11793 | -0.73419 | -1.6158 | 0.29 |
| 13128 | *RBM19* | 2.436 | -0.58633 | -0.99217 | -1.3298 | 0.29 |
| 13129 | *MED8* | 0.54972 | 0.16453 | 0.14261 | -1.5348 | 0.29 |
| 13130 | *USP42* | 0.56568 | 0.18851 | 0.10242 | -1.0564 | 0.29 |
| 13131 | *ACYP2* | 2.6787 | -0.89258 | -0.9304 | -1.8293 | 0.29 |
| 13132 | *CCL18* | 2.7361 | -0.66164 | -1.2192 | -1.2485 | 0.29 |
| 13133 | *LGI1* | 1.1162 | -0.06413 | -0.19735 | -1.0178 | 0.28 |
| 13134 | *CREBZF* | 0.52019 | 0.31774 | 0.01649 | -1.8611 | 0.28 |
| 13135 | *UGGT2* | 1.7107 | -0.39664 | -0.46017 | -1.0003 | 0.28 |
| 13136 | *RRP36* | 1.4767 | 0.32911 | -0.95315 | -1.5707 | 0.28 |
| 13137 | *SYTL2* | 2.3223 | -0.66731 | -0.80234 | -1.1676 | 0.28 |
| 13138 | *TGM7* | 0.55195 | 0.2836 | 0.01621 | -1.1466 | 0.28 |
| 13139 | *BAAT* | 0.68631 | 0.53898 | -0.37358 | -2.4328 | 0.28 |
| 13140 | *ZNF701* | 2.0619 | 0.18341 | -1.3941 | -1.5906 | 0.28 |
| 13141 | *BZRAP1* | 2.439 | -0.55029 | -1.0385 | -1.6885 | 0.28 |
| 13142 | *OR1E1* | 1.9571 | -0.168 | -0.93895 | -1.1102 | 0.28 |
| 13143 | *OR2B6* | 1.0245 | 0.91864 | -1.0947 | -1.5104 | 0.28 |
| 13144 | *HIST1H2AI* | 1.0082 | 0.25931 | -0.42004 | -1.8851 | 0.28 |
| 13145 | *U2AF1L4* | 2.7584 | -0.1747 | -1.7365 | -2.4094 | 0.28 |
| 13146 | *OR5M9* | 1.917 | 0.58233 | -1.6524 | -1.773 | 0.28 |
| 13147 | *CYP3A5* | 1.0565 | 0.17295 | -0.38428 | -1.5801 | 0.28 |
| 13148 | *RBBP8NL* | 1.2735 | -0.05456 | -0.37417 | -1.631 | 0.28 |
| 13149 | *TECRL* | 0.98259 | 0.43947 | -0.57825 | -0.80146 | 0.28 |

| | | | | | | |
|---|---|---|---|---|---|---|
| 13150 | *ANXA6* | 1.0343 | 0.012414 | -0.20292 | -2.0059 | 0.28 |
| 13151 | *PRKCA* | 1.6395 | 0.44773 | -1.2437 | -1.9302 | 0.28 |
| 13152 | *TYW3* | 1.3076 | 0.099372 | -0.56444 | -1.843 | 0.28 |
| 13153 | *COL19A1* | 2.7664 | -0.77789 | -1.1461 | -1.1758 | 0.28 |
| 13154 | *DENND4A* | 1.1735 | -0.02163 | -0.30965 | -1.8038 | 0.28 |
| 13155 | *MAP6D1* | 1.2292 | 0.068993 | -0.45604 | -1.8838 | 0.28 |
| 13156 | *ANKRD20A3* | 1.4515 | -0.25762 | -0.35211 | -1.5331 | 0.28 |
| 13157 | *TBL1X* | 1.7042 | -0.17298 | -0.68999 | -1.6306 | 0.28 |
| 13158 | *TMEM26* | 1.5352 | -0.31414 | -0.37995 | -1.0691 | 0.28 |
| 13159 | *RBMX* | 1.8744 | -0.00833 | -1.025 | -1.3632 | 0.28 |
| 13160 | *COPS6* | 0.51639 | 0.45083 | -0.12725 | -0.70976 | 0.28 |
| 13161 | *SEC23B* | 1.5089 | 0.18147 | -0.8504 | -0.95527 | 0.28 |
| 13162 | *IL1R2* | 3.6038 | -1.2705 | -1.4935 | -2.2094 | 0.28 |
| 13163 | *PIP5KL1* | 3.4378 | -1.1944 | -1.4036 | -1.748 | 0.28 |
| 13164 | *ACYP1* | 0.44793 | 0.34532 | 0.04584 | -0.12189 | 0.28 |
| 13165 | *CD7* | 3.8417 | -1.1287 | -1.8742 | -2.9143 | 0.28 |
| 13166 | *C1orf229* | 0.62565 | 0.11317 | 0.09965 | -1.9965 | 0.28 |
| 13167 | *CRISPLD2* | 0.4834 | 0.45449 | -0.09984 | -2.1066 | 0.28 |
| 13168 | *NDUFA4* | 1.1524 | 0.29744 | -0.61436 | -1.3694 | 0.28 |
| 13169 | *SLC7A5* | 0.72742 | 0.24671 | -0.14122 | -1.3849 | 0.28 |
| 13170 | *CHD6* | 1.0251 | 0.10965 | -0.30279 | -0.46665 | 0.28 |
| 13171 | *ASB8* | 1.2781 | 0.083593 | -0.53009 | -1.8996 | 0.28 |
| 13172 | *ESCO1* | 0.74858 | 0.52571 | -0.44353 | -0.56489 | 0.28 |
| 13173 | *CD2AP* | 0.58729 | 0.18898 | 0.05386 | -0.94563 | 0.28 |
| 13174 | *TMED7-TICAM2* | 1.2694 | 0.31229 | -0.75205 | -1.8609 | 0.28 |
| 13175 | *IGF2BP1* | 1.5293 | -0.15679 | -0.54345 | -0.88279 | 0.28 |
| 13176 | *FADS2* | 0.91459 | 0.60736 | -0.69312 | -0.7229 | 0.28 |
| 13177 | *FAM178B* | 0.90985 | 0.039496 | -0.12096 | -0.69073 | 0.28 |
| 13178 | *ANKRD39* | 1.4883 | 0.45681 | -1.1169 | -1.6836 | 0.28 |
| 13179 | *RPL10L* | 0.8797 | -0.01508 | -0.03675 | -0.20848 | 0.28 |
| 13180 | *RNASEK* | 0.61627 | 0.50175 | -0.29027 | -0.63147 | 0.28 |
| 13181 | *USMG5* | 1.981 | -0.53969 | -0.61394 | -0.88465 | 0.28 |
| 13182 | *ZNF157* | 0.59461 | 0.23846 | -0.0062 | -0.84134 | 0.28 |
| 13183 | *EDARADD* | 2.4591 | 0.10421 | -1.7365 | -2.0682 | 0.28 |
| 13184 | *SLC22A11* | 1.5832 | -0.31842 | -0.43845 | -0.67395 | 0.28 |
| 13185 | *RIMBP3B* | 2.2822 | -0.21821 | -1.238 | -1.3293 | 0.28 |
| 13186 | *APOBEC3A* | 2.8935 | 0.16266 | -2.2313 | -3.5756 | 0.27 |
| 13187 | *TSTA3* | 0.63433 | 0.21886 | -0.02908 | -1.7855 | 0.27 |
| 13188 | *SLC46A2* | 0.66345 | 0.61421 | -0.45355 | -0.71063 | 0.27 |
| 13189 | *DDX60L* | 1.3641 | -0.02658 | -0.51396 | -0.55816 | 0.27 |
| 13190 | *TSPYL5* | 1.6153 | 0.36143 | -1.1532 | -2.1364 | 0.27 |
| 13191 | *SCN11A* | 1.9841 | -0.48767 | -0.67298 | -0.79195 | 0.27 |
| 13192 | *VPREB3* | 0.75846 | 0.50819 | -0.44353 | -1.3729 | 0.27 |
| 13193 | *N6AMT2* | 1.0187 | -0.07031 | -0.12552 | -1.4287 | 0.27 |
| 13194 | *CLCN3* | 0.72975 | 0.58279 | -0.49107 | -0.76582 | 0.27 |
| 13195 | *LRRC26* | 1.3025 | 0.087154 | -0.56963 | -1.0824 | 0.27 |

| | | | | | | |
|---|---|---|---|---|---|---|
| 13196 | *MADD* | 0.5014 | 0.1857 | 0.13228 | -1.5653 | 0.27 |
| 13197 | *ZNF552* | 2.1543 | -0.66146 | -0.67395 | -1.9041 | 0.27 |
| 13198 | *CCNT1* | 0.75322 | 0.61081 | -0.5458 | -0.58553 | 0.27 |
| 13199 | *RTL1* | 0.80282 | 0.18187 | -0.16651 | -1.1021 | 0.27 |
| 13200 | *NRSN2* | 3.2998 | -0.32466 | -2.1574 | -2.5647 | 0.27 |
| 13201 | *PRDX2* | 1.5021 | 0.29711 | -0.98314 | -1.7364 | 0.27 |
| 13202 | *IFNK* | 2.2345 | -0.7061 | -0.71268 | -1.613 | 0.27 |
| 13203 | *MYL12B* | 1.0779 | 0.8964 | -1.1598 | -1.6246 | 0.27 |
| 13204 | *NOP16* | 2.5428 | -0.50588 | -1.2229 | -1.2704 | 0.27 |
| 13205 | *MNS1* | 1.6375 | -0.01861 | -0.80607 | -1.1598 | 0.27 |
| 13206 | *PSEN1* | 0.80687 | 0.33157 | -0.32589 | -0.68303 | 0.27 |
| 13207 | *OGT* | 1.6244 | 0.40853 | -1.2208 | -1.6555 | 0.27 |
| 13208 | *ARFGAP3* | 2.2607 | -0.65068 | -0.79813 | -1.6266 | 0.27 |
| 13209 | *NOXO1* | 3.5892 | -1.202 | -1.577 | -2.6939 | 0.27 |
| 13210 | *ST7L* | 2.7256 | -0.95423 | -0.96206 | -2.3509 | 0.27 |
| 13211 | *TRPV4* | 2.2867 | 0.17475 | -1.6524 | -2.1956 | 0.27 |
| 13212 | *LAMC1* | 0.49605 | 0.29665 | 0.01621 | -0.89269 | 0.27 |
| 13213 | *SFXN3* | 1.6173 | 0.8979 | -1.7068 | -2.5008 | 0.27 |
| 13214 | *DAP3* | 1.2193 | -0.16485 | -0.2465 | -1.3433 | 0.27 |
| 13215 | *PITPNC1* | 1.179 | 0.21219 | -0.58349 | -2.1477 | 0.27 |
| 13216 | *ATF5* | 1.8888 | -0.12456 | -0.95682 | -1.4325 | 0.27 |
| 13217 | *CACNA1B* | 0.55932 | 0.15515 | 0.09251 | -0.78651 | 0.27 |
| 13218 | *GCFC2* | 1.0967 | 0.34471 | -0.63511 | -0.65924 | 0.27 |
| 13219 | *IGLON5* | 0.98197 | -0.03697 | -0.13928 | -0.74013 | 0.27 |
| 13220 | *MAT1A* | 0.92643 | 0.38415 | -0.50555 | -1.3517 | 0.27 |
| 13221 | *USP39* | 0.77008 | 0.052299 | -0.0175 | -1.0716 | 0.27 |
| 13222 | *PPP1R3B* | 0.62844 | 0.30144 | -0.12642 | -0.80686 | 0.27 |
| 13223 | *RBM27* | 1.4726 | -0.29042 | -0.37913 | -0.72825 | 0.27 |
| 13224 | *DYTN* | 1.164 | 0.1824 | -0.54373 | -1.4474 | 0.27 |
| 13225 | *STX12* | 1.0582 | 0.21171 | -0.46747 | -1.616 | 0.27 |
| 13226 | *GYPE* | 1.5377 | -0.36747 | -0.36913 | -0.83469 | 0.27 |
| 13227 | *RNFT2* | 1.1825 | 0.25883 | -0.64166 | -0.76982 | 0.27 |
| 13228 | *IL31* | 1.8805 | -0.31145 | -0.76955 | -1.9912 | 0.27 |
| 13229 | *CLCC1* | 1.6782 | 0.23807 | -1.1169 | -1.4591 | 0.27 |
| 13230 | *ARHGEF33* | 1.8217 | 0.27676 | -1.3002 | -1.3148 | 0.27 |
| 13231 | *VTN* | 2.7855 | -0.45705 | -1.5306 | -2.5584 | 0.27 |
| 13232 | *NIPAL3* | 0.82378 | 0.27244 | -0.29905 | -1.4684 | 0.27 |
| 13233 | *SLC38A8* | 3.0906 | -0.89155 | -1.4027 | -1.8212 | 0.27 |
| 13234 | *SLCO6A1* | 0.90604 | 0.47089 | -0.58156 | -1.9019 | 0.27 |
| 13235 | *GRIN2C* | 1.845 | -0.48021 | -0.56947 | -1.7365 | 0.27 |
| 13236 | *VSTM5* | 0.53081 | 0.34989 | -0.08572 | -2.4933 | 0.26 |
| 13237 | *ODF4* | 2.6496 | 0.60333 | -2.458 | -2.825 | 0.26 |
| 13238 | *DGAT1* | 1.3274 | 1.0525 | -1.5855 | -2.2461 | 0.26 |
| 13239 | *DSEL* | 1.1552 | 0.17236 | -0.53361 | -1.3505 | 0.26 |
| 13240 | *FGFR1* | 1.9974 | 0.11075 | -1.3151 | -1.9559 | 0.26 |
| 13241 | *LPPR4* | 1.1912 | 0.77628 | -1.175 | -2.2948 | 0.26 |

| | | | | | | |
|---|---|---|---|---|---|---|
| 13242 | *LTA4H* | 1.3797 | 0.21587 | -0.80349 | -1.6676 | 0.26 |
| 13243 | *PEX6* | 0.81109 | 0.78918 | -0.8083 | -1.333 | 0.26 |
| 13244 | *P2RX5* | 3.246 | -0.38655 | -2.0682 | -2.8316 | 0.26 |
| 13245 | *GJB5* | 0.55758 | 0.4252 | -0.19169 | -0.30349 | 0.26 |
| 13246 | *LRRC52* | 0.99501 | 0.068606 | -0.27291 | -2.2485 | 0.26 |
| 13247 | *C2orf68* | 3.0519 | -0.73071 | -1.5309 | -1.7068 | 0.26 |
| 13248 | *MTM1* | 0.73858 | 0.16089 | -0.1095 | -1.2383 | 0.26 |
| 13249 | *SHPRH* | 2.2139 | -0.51703 | -0.90743 | -1.397 | 0.26 |
| 13250 | *ALK* | 1.3523 | 0.11832 | -0.68133 | -1.1888 | 0.26 |
| 13251 | *MT1X* | 1.9937 | -0.40467 | -0.8003 | -1.6047 | 0.26 |
| 13252 | *GGTLC2* | 0.61199 | 0.33869 | -0.16205 | -0.67321 | 0.26 |
| 13253 | *CD33* | 1.2558 | -0.05698 | -0.41025 | -2.0177 | 0.26 |
| 13254 | *PLXDC2* | 2.8204 | -0.29305 | -1.739 | -2.7317 | 0.26 |
| 13255 | *KLHDC8B* | 0.81699 | 0.65417 | -0.68303 | -0.83527 | 0.26 |
| 13256 | *OSTC* | 0.51793 | 0.25552 | 0.01326 | -0.89059 | 0.26 |
| 13257 | *SEMA4A* | 2.3097 | -0.10243 | -1.4206 | -2.3287 | 0.26 |
| 13258 | *GP5* | 2.5144 | -0.35805 | -1.3701 | -1.8999 | 0.26 |
| 13259 | *UPF3A* | 2.6152 | -0.03587 | -1.7935 | -3.2023 | 0.26 |
| 13260 | *INPP5E* | 0.57318 | 0.39716 | -0.1851 | -0.2694 | 0.26 |
| 13261 | *KRT71* | 0.53795 | 0.23846 | 0.00882 | -0.41703 | 0.26 |
| 13262 | *HBEGF* | 0.86634 | 0.067594 | -0.15041 | -0.93696 | 0.26 |
| 13263 | *GLUD1* | 3.4735 | -0.69961 | -1.9912 | -2.4642 | 0.26 |
| 13264 | *APOL4* | 1.8217 | -0.31309 | -0.72635 | -0.98983 | 0.26 |
| 13265 | *UPK1A* | 1.6714 | -0.30419 | -0.58589 | -0.8597 | 0.26 |
| 13266 | *KRT24* | 0.699 | 0.48025 | -0.39897 | -2.0363 | 0.26 |
| 13267 | *SOX9* | 0.75031 | 0.73596 | -0.70659 | -3.9921 | 0.26 |
| 13268 | *ATP10A* | 2.9983 | -0.18858 | -2.0301 | -2.2108 | 0.26 |
| 13269 | *PWWP2A* | 0.86909 | 0.67348 | -0.76352 | -0.89204 | 0.26 |
| 13270 | *CTNS* | 3.0369 | -1.0528 | -1.2052 | -1.3821 | 0.26 |
| 13271 | *TSEN15* | 2.6108 | -0.19042 | -1.6417 | -1.7887 | 0.26 |
| 13272 | *SKIL* | 1.3252 | 0.37477 | -0.92131 | -1.9622 | 0.26 |
| 13273 | *CACNG3* | 2.6576 | -0.52473 | -1.3543 | -2.5728 | 0.26 |
| 13274 | *UBQLN3* | 2.7407 | 0.01805 | -1.9811 | -2.6257 | 0.26 |
| 13275 | *POLR1E* | 0.60959 | 0.41125 | -0.24327 | -0.2868 | 0.26 |
| 13276 | *C17orf105* | 0.87211 | 0.4154 | -0.51002 | -1.5132 | 0.26 |
| 13277 | *CCR3* | 3.7494 | -1.3199 | -1.6525 | -2.6652 | 0.26 |
| 13278 | *METTL9* | 0.66946 | 0.55033 | -0.44374 | -0.48538 | 0.26 |
| 13279 | *GPR108* | 0.97037 | 0.52156 | -0.7167 | -1.4909 | 0.26 |
| 13280 | *PTTG1* | 3.0774 | -0.36582 | -1.9368 | -2.2473 | 0.26 |
| 13281 | *ATG3* | 0.71308 | 0.3868 | -0.32516 | -0.76942 | 0.26 |
| 13282 | *POLR2E* | 0.96431 | 0.37008 | -0.56027 | -1.4281 | 0.26 |
| 13283 | *TTN* | 2.1231 | -0.26055 | -1.0889 | -1.3773 | 0.26 |
| 13284 | *RPL18A* | 2.5782 | -0.74141 | -1.0633 | -1.4258 | 0.26 |
| 13285 | *C15orf62* | 2.3849 | -0.53615 | -1.0756 | -1.305 | 0.26 |
| 13286 | *ZFP1* | 0.74748 | 0.29981 | -0.27498 | -2.105 | 0.26 |
| 13287 | *SSX2* | 2.2631 | -0.15276 | -1.3382 | -1.4856 | 0.26 |

| | | | | | | |
|---|---|---|---|---|---|---|
| 13288 | *OIP5* | 1.3179 | 0.17128 | -0.71717 | -0.73202 | 0.26 |
| 13289 | *TAS2R50* | 2.1173 | 0.10131 | -1.4473 | -1.7247 | 0.26 |
| 13290 | *FADS6* | 0.80248 | 0.53269 | -0.56444 | -0.77521 | 0.26 |
| 13291 | *ECM1* | 1.1933 | -0.14306 | -0.28003 | -0.95105 | 0.26 |
| 13292 | *AOC3* | 0.81508 | 0.16736 | -0.21291 | -1.0095 | 0.26 |
| 13293 | *HNRNPL* | 1.4919 | 0.25617 | -0.97865 | -2.3154 | 0.26 |
| 13294 | *MMP2* | 1.2626 | 0.54342 | -1.0367 | -2.0301 | 0.26 |
| 13295 | *SMAD5* | 1.2468 | -0.01705 | -0.46056 | -1.2934 | 0.26 |
| 13296 | *ACADSB* | 2.9006 | -0.46738 | -1.6644 | -2.674 | 0.26 |
| 13297 | *FAH* | 1.2823 | 0.3781 | -0.89186 | -1.9722 | 0.26 |
| 13298 | *PIK3AP1* | 0.88845 | 0.32362 | -0.44353 | -1.1727 | 0.26 |
| 13299 | *PIH1D1* | 1.5496 | -0.22708 | -0.55464 | -1.3478 | 0.26 |
| 13300 | *NIT1* | 1.3511 | -0.09762 | -0.48561 | -0.85095 | 0.26 |
| 13301 | *MTX1* | 3.1284 | -0.86941 | -1.4916 | -1.5544 | 0.26 |
| 13302 | *KAT5* | 1.3251 | 0.99971 | -1.5588 | -1.7022 | 0.26 |
| 13303 | *ZDHHC12* | 1.278 | 0.51028 | -1.0224 | -1.485 | 0.26 |
| 13304 | *TNFSF12-TNFSF13* | 0.73844 | 0.55385 | -0.52711 | -2.1926 | 0.26 |
| 13305 | *RAB21* | 3.1968 | -0.9968 | -1.4359 | -1.4747 | 0.25 |
| 13306 | *ACTR10* | 2.9173 | -0.50781 | -1.6463 | -2.4371 | 0.25 |
| 13307 | *ITPRIPL2* | 0.73659 | 0.064489 | -0.03836 | -0.402 | 0.25 |
| 13308 | *ZFP57* | 0.44684 | 0.25904 | 0.05677 | -1.5978 | 0.25 |
| 13309 | *ANKRD11* | 0.87347 | 0.096106 | -0.2073 | -1.8678 | 0.25 |
| 13310 | *PIGC* | 0.75354 | 0.40756 | -0.39924 | -0.95495 | 0.25 |
| 13311 | *ERC2* | 1.2182 | 0.71334 | -1.1698 | -2.1921 | 0.25 |
| 13312 | *ADGB* | 1.3894 | 1.1155 | -1.7438 | -1.8212 | 0.25 |
| 13313 | *TGFB1* | 0.64948 | 0.15662 | -0.04502 | -1.3153 | 0.25 |
| 13314 | *RNF133* | 1.4413 | -0.17744 | -0.503 | -1.4137 | 0.25 |
| 13315 | *TLE6* | 1.9383 | -0.2251 | -0.95321 | -1.0528 | 0.25 |
| 13316 | *COL17A1* | 1.9646 | 0.66956 | -1.8751 | -2.3033 | 0.25 |
| 13317 | *ASAH2B* | 0.6879 | 0.093005 | -0.02187 | -0.80869 | 0.25 |
| 13318 | *LCN12* | 3.9412 | -1.5874 | -1.5959 | -1.773 | 0.25 |
| 13319 | *PPP2R5D* | 0.30678 | 0.29956 | 0.15092 | -0.79319 | 0.25 |
| 13320 | *ZNF385D* | 0.52027 | 0.20646 | 0.03043 | -0.11314 | 0.25 |
| 13321 | *LRRC34* | 0.71993 | 0.66286 | -0.62596 | -1.1872 | 0.25 |
| 13322 | *SLC12A2* | 2.5968 | -0.67141 | -1.1698 | -2.4853 | 0.25 |
| 13323 | *PPP5D1* | 1.5449 | 0.021236 | -0.81058 | -2.2599 | 0.25 |
| 13324 | *SPINK8* | 1.9554 | -0.59959 | -0.60058 | -0.64678 | 0.25 |
| 13325 | *TICAM1* | 1.1025 | -0.10946 | -0.23825 | -1.0014 | 0.25 |
| 13326 | *DR1* | 1.1969 | -0.16069 | -0.28164 | -1.3476 | 0.25 |
| 13327 | *ZNF665* | 0.74341 | 0.21148 | -0.20287 | -1.0088 | 0.25 |
| 13328 | *NUTM2A* | 3.2332 | -0.03356 | -2.4481 | -2.5764 | 0.25 |
| 13329 | *CA6* | 3.2923 | -1.0901 | -1.4509 | -2.0476 | 0.25 |
| 13330 | *FAM189A1* | 1.9656 | 0.45421 | -1.6689 | -2.011 | 0.25 |
| 13331 | *MOSPD2* | 2.1486 | -0.32748 | -1.071 | -3.1297 | 0.25 |
| 13332 | *KRTAP4-6* | 3.9246 | -1.2352 | -1.9397 | -2.3251 | 0.25 |
| 13333 | *NPBWR1* | 1.8463 | -0.03909 | -1.0589 | -1.8646 | 0.25 |

| | | | | | | |
|---|---|---|---|---|---|---|
| 13334 | *UQCRQ* | 1.4283 | 0.86236 | -1.5426 | -1.8095 | 0.25 |
| 13335 | *RXFP1* | 0.73987 | 0.15419 | -0.14639 | -0.3828 | 0.25 |
| 13336 | *ZNHIT3* | 2.3491 | -0.76165 | -0.84069 | -2.0989 | 0.25 |
| 13337 | *TMED6* | 2.1887 | -0.5986 | -0.84337 | -2.2449 | 0.25 |
| 13338 | *CHFR* | 1.4599 | 0.36473 | -1.0782 | -1.9472 | 0.25 |
| 13339 | *SPATA25* | 1.0483 | 0.53134 | -0.83411 | -0.8528 | 0.25 |
| 13340 | *SNRPD1* | 0.31317 | 0.21588 | 0.2158 | -1.6218 | 0.25 |
| 13341 | *PTCD3* | 3.1373 | -1.031 | -1.3619 | -2.444 | 0.25 |
| 13342 | *ZNF468* | 1.8722 | -0.29076 | -0.83739 | -1.2097 | 0.25 |
| 13343 | *PDE4A* | 2.22 | -0.33174 | -1.1451 | -1.9455 | 0.25 |
| 13344 | *RABGEF1* | 2.387 | -0.58495 | -1.0589 | -1.4234 | 0.25 |
| 13345 | *RAB12* | 1.3985 | 0.25633 | -0.91296 | -1.1421 | 0.25 |
| 13346 | *RBM15* | 1.0652 | 0.31825 | -0.64164 | -2.4151 | 0.25 |
| 13347 | *OVOL2* | 0.69701 | 0.32713 | -0.28242 | -1.2098 | 0.25 |
| 13348 | *TRIM49D2* | 0.96786 | 0.057442 | -0.28393 | -1.6788 | 0.25 |
| 13349 | *STARD7* | 2.8125 | -0.74184 | -1.3294 | -2.5189 | 0.25 |
| 13350 | *ABCB11* | 1.4178 | 0.21618 | -0.8936 | -1.8679 | 0.25 |
| 13351 | *CNTNAP4* | 0.77909 | 0.15516 | -0.19422 | -0.82833 | 0.25 |
| 13352 | *LMBR1L* | 1.4782 | 0.21632 | -0.95545 | -2.2473 | 0.25 |
| 13353 | *UTS2* | 2.7028 | -0.00819 | -1.9558 | -2.4785 | 0.25 |
| 13354 | *TRIM69* | 0.42524 | 0.3822 | -0.06892 | -0.61436 | 0.25 |
| 13355 | *TIE1* | 0.65117 | 0.4185 | -0.33191 | -1.0118 | 0.25 |
| 13356 | *ZCCHC24* | 0.84203 | 0.014586 | -0.1201 | -1.992 | 0.25 |
| 13357 | *TFAM* | 1.1842 | 0.46277 | -0.91192 | -1.5526 | 0.25 |
| 13358 | *NFE2L2* | 1.9881 | -0.37417 | -0.8789 | -1.0321 | 0.25 |
| 13359 | *UCN* | 1.0749 | 0.46124 | -0.80191 | -2.0363 | 0.24 |
| 13360 | *PITPNA* | 1.2088 | 0.20909 | -0.68383 | -2.2461 | 0.24 |
| 13361 | *LTV1* | 2.5357 | 0.20215 | -2.0042 | -2.5354 | 0.24 |
| 13362 | *RPS19BP1* | 0.41025 | 0.34397 | -0.02105 | -2.1224 | 0.24 |
| 13363 | *ZNF208* | 1.5239 | -0.37642 | -0.41476 | -1.1034 | 0.24 |
| 13364 | *MOV10L1* | 0.41247 | 0.19462 | 0.12522 | -1.6316 | 0.24 |
| 13365 | *FAM160B1* | 1.6579 | 0.42215 | -1.3478 | -1.4519 | 0.24 |
| 13366 | *C19orf35* | 0.61036 | 0.58268 | -0.46092 | -2.0242 | 0.24 |
| 13367 | *GPR62* | 2.2375 | -0.41127 | -1.0942 | -1.7683 | 0.24 |
| 13368 | *CNTNAP5* | 2.664 | -0.50394 | -1.4287 | -2.2203 | 0.24 |
| 13369 | *OR10H4* | 1.1694 | 0.1957 | -0.63436 | -0.68259 | 0.24 |
| 13370 | *ZNF549* | 2.4161 | 0.16183 | -1.8478 | -2.1036 | 0.24 |
| 13371 | *CR2* | 1.0656 | 0.2336 | -0.56947 | -1.2766 | 0.24 |
| 13372 | *AK8* | 1.0265 | -0.02226 | -0.27453 | -1.5212 | 0.24 |
| 13373 | *LKAAEAR1* | 3.4079 | -1.0584 | -1.6199 | -2.3063 | 0.24 |
| 13374 | *SETD1B* | 1.5048 | 0.49602 | -1.2715 | -1.7192 | 0.24 |
| 13375 | *C9orf69* | 2.2823 | -0.43478 | -1.1184 | -2.1723 | 0.24 |
| 13376 | *ARL6IP5* | 2.482 | -0.73029 | -1.023 | -1.6278 | 0.24 |
| 13377 | *IFNL3* | 0.98458 | 0.72684 | -0.98292 | -1.4258 | 0.24 |
| 13378 | *TTC25* | 3.201 | -0.58479 | -1.8879 | -1.9404 | 0.24 |
| 13379 | *TMEM222* | 3.5197 | -1.3633 | -1.4282 | -1.5393 | 0.24 |

| | | | | | | |
|---|---|---|---|---|---|---|
| 13380 | *NUDT13* | 1.9105 | 0.92337 | -2.1068 | -2.2879 | 0.24 |
| 13381 | *C5orf42* | 0.6021 | 0.33338 | -0.20849 | -0.97103 | 0.24 |
| 13382 | *CYSRT1* | 1.9376 | -0.32611 | -0.88495 | -1.8112 | 0.24 |
| 13383 | *SPATA24* | 1.114 | -0.02658 | -0.36242 | -2.2533 | 0.24 |
| 13384 | *SHC2* | 0.67368 | 0.085982 | -0.0354 | -0.74875 | 0.24 |
| 13385 | *TPST2* | 1.5913 | -0.1875 | -0.6799 | -1.1388 | 0.24 |
| 13386 | *ZDHHC9* | 1.816 | 0.3001 | -1.3946 | -1.6939 | 0.24 |
| 13387 | *CLLU1* | 0.65403 | 0.19443 | -0.12709 | -0.16242 | 0.24 |
| 13388 | *CD55* | 1.5364 | 0.38114 | -1.1962 | -2.3826 | 0.24 |
| 13389 | *CRISP1* | 2.4311 | 0.027886 | -1.739 | -3.4957 | 0.24 |
| 13390 | *GRHPR* | 0.96157 | 0.47648 | -0.71905 | -1.9213 | 0.24 |
| 13391 | *UCP2* | 1.1802 | 0.026897 | -0.48925 | -0.90001 | 0.24 |
| 13392 | *OTX2* | 2.7243 | -0.18167 | -1.8253 | -2.262 | 0.24 |
| 13393 | *THEGL* | 3.389 | -1.146 | -1.526 | -1.5415 | 0.24 |
| 13394 | *AZI2* | 0.69364 | 0.20424 | -0.18175 | -0.2408 | 0.24 |
| 13395 | *MYLK* | 0.75321 | 0.23932 | -0.27672 | -1.589 | 0.24 |
| 13396 | *SLC3A1* | 2.7277 | -0.82226 | -1.1897 | -2.1219 | 0.24 |
| 13397 | *OR7G3* | 0.75178 | 0.37143 | -0.40824 | -2.1014 | 0.24 |
| 13398 | *PBK* | 0.3326 | 0.26202 | 0.11968 | -0.71896 | 0.24 |
| 13399 | *CSN1S1* | 1.1185 | 0.28685 | -0.69155 | -2.5198 | 0.24 |
| 13400 | *WTH3DI* | 3.8492 | -1.2717 | -1.8642 | -2.0267 | 0.24 |
| 13401 | *TCIRG1* | 1.8657 | -0.48086 | -0.6716 | -1.5898 | 0.24 |
| 13402 | *ADGRG3* | 1.0105 | 0.38053 | -0.67812 | -2.2748 | 0.24 |
| 13403 | *C2orf74* | 3.7128 | -1.1383 | -1.8636 | -1.8938 | 0.24 |
| 13404 | *POLR3F* | 1.7991 | -0.30192 | -0.78639 | -1.3814 | 0.24 |
| 13405 | *FANCL* | 1.6115 | 0.59114 | -1.4919 | -2.9308 | 0.24 |
| 13406 | *ANKRD17* | 1.4996 | -0.30241 | -0.48669 | -1.7959 | 0.24 |
| 13407 | *IQCB1* | 0.56929 | 0.55315 | -0.41207 | -0.74875 | 0.24 |
| 13408 | *FRS2* | 0.74631 | 0.4075 | -0.44353 | -1.7034 | 0.24 |
| 13409 | *FIGN* | 1.0721 | 0.60896 | -0.97149 | -1.6456 | 0.24 |
| 13410 | *STAC2* | 0.33158 | 0.25162 | 0.12509 | -0.03046 | 0.24 |
| 13411 | *ZBTB4* | 2.3523 | -0.24856 | -1.397 | -1.4087 | 0.24 |
| 13412 | *MAMDC2* | 0.50708 | 0.40046 | -0.20107 | -1.1912 | 0.24 |
| 13413 | *DNAJC28* | 1.3496 | -0.06253 | -0.58143 | -0.60113 | 0.24 |
| 13414 | *BCL2L11* | 2.9451 | -0.78951 | -1.4501 | -1.9474 | 0.24 |
| 13415 | *UPP1* | 1.1851 | -0.11968 | -0.36066 | -0.98239 | 0.23 |
| 13416 | *SNW1* | 2.5408 | -0.87078 | -0.96529 | -1.1768 | 0.23 |
| 13417 | *PRLR* | 3.0378 | -0.86506 | -1.4682 | -1.7383 | 0.23 |
| 13418 | *OAZ3* | 2.3221 | 0.53001 | -2.1478 | -2.5934 | 0.23 |
| 13419 | *C11orf86* | 1.5238 | 0.049686 | -0.86923 | -1.0178 | 0.23 |
| 13420 | *SEC31B* | 2.4049 | -0.5409 | -1.1605 | -1.7072 | 0.23 |
| 13421 | *EML6* | 2.9174 | -0.00317 | -2.2108 | -3.0959 | 0.23 |
| 13422 | *TRUB1* | 0.60091 | 0.14451 | -0.04218 | -0.65503 | 0.23 |
| 13423 | *TBX2* | 2.0545 | 0.44943 | -1.8007 | -3.0378 | 0.23 |
| 13424 | *GLOD5* | 0.70504 | 0.4209 | -0.42352 | -0.57054 | 0.23 |
| 13425 | *TTLL5* | 1.1759 | 0.10748 | -0.58143 | -1.2431 | 0.23 |

| | | | | | | |
|---|---|---|---|---|---|---|
| 13426 | *SKI* | 1.0392 | 0.48824 | -0.82573 | -1.4509 | 0.23 |
| 13427 | *SGOL2* | 2.5866 | -0.82789 | -1.058 | -1.8335 | 0.23 |
| 13428 | *SGK1* | 0.31766 | 0.21312 | 0.16903 | -1.1738 | 0.23 |
| 13429 | *MYO1E* | 0.59299 | 0.34589 | -0.2391 | -0.97381 | 0.23 |
| 13430 | *CDHR5* | 0.75731 | 0.19282 | -0.25047 | -2.0146 | 0.23 |
| 13431 | *ABHD4* | 1.272 | 0.23519 | -0.80847 | -1.5552 | 0.23 |
| 13432 | *WDR60* | 2.8659 | 0.057397 | -2.2254 | -2.4174 | 0.23 |
| 13433 | *AP2A2* | 0.86619 | 0.003671 | -0.17261 | -1.2284 | 0.23 |
| 13434 | *DCP2* | 1.6363 | -0.19587 | -0.74459 | -2.6805 | 0.23 |
| 13435 | *INTS9* | 0.72339 | 0.5894 | -0.61698 | -1.0944 | 0.23 |
| 13436 | *IGF2BP2* | 0.85062 | 0.64689 | -0.80315 | -1.0528 | 0.23 |
| 13437 | *NDN* | 0.47149 | 0.40964 | -0.18718 | -0.69345 | 0.23 |
| 13438 | *STARD4* | 3.09 | -1.1733 | -1.2235 | -2.1517 | 0.23 |
| 13439 | *SLC26A2* | 0.63127 | 0.54286 | -0.48104 | -1.6832 | 0.23 |
| 13440 | *POLG* | 0.97812 | 0.12402 | -0.41025 | -0.58485 | 0.23 |
| 13441 | *RAD52* | 0.81849 | 0.053516 | -0.18026 | -2.4797 | 0.23 |
| 13442 | *ADGRG1* | 1.7875 | 0.99921 | -2.095 | -2.7931 | 0.23 |
| 13443 | *EEF1B2* | 1.2001 | 0.77063 | -1.2792 | -1.8202 | 0.23 |
| 13444 | *ERO1L* | 1.009 | 0.39196 | -0.70973 | -2.0384 | 0.23 |
| 13445 | *RFT1* | 1.8017 | -0.37949 | -0.73155 | -1.9541 | 0.23 |
| 13446 | *PCDHAC1* | 1.1732 | 0.5811 | -1.0645 | -1.1872 | 0.23 |
| 13447 | *ATP8B3* | 3.7047 | -0.60604 | -2.4094 | -2.7705 | 0.23 |
| 13448 | *CASC1* | 1.6426 | 0.30293 | -1.2569 | -1.7366 | 0.23 |
| 13449 | *SRP54* | 0.79458 | 0.29117 | -0.39722 | -1.3298 | 0.23 |
| 13450 | *ZNF512* | 0.84938 | 0.23402 | -0.39636 | -0.66364 | 0.23 |
| 13451 | *GJA5* | 1.5005 | 0.5661 | -1.3818 | -1.6699 | 0.23 |
| 13452 | *TNFSF13B* | 1.0209 | 0.081493 | -0.41791 | -0.83801 | 0.23 |
| 13453 | *C8orf86* | 0.45389 | 0.15078 | 0.07973 | -0.31309 | 0.23 |
| 13454 | *COL21A1* | 1.1778 | 0.34238 | -0.83627 | -1.5302 | 0.23 |
| 13455 | *FMO2* | 0.43851 | 0.15379 | 0.09154 | -0.06809 | 0.23 |
| 13456 | *HOXD1* | 0.39018 | 0.15845 | 0.13483 | 0.068131 | 0.23 |
| 13457 | *TTC16* | 1.2676 | 0.030628 | -0.61491 | -0.68389 | 0.23 |
| 13458 | *PPP6R2* | 0.70446 | 0.40077 | -0.42358 | -1.3409 | 0.23 |
| 13459 | *ZNF527* | 0.70317 | 0.64679 | -0.67007 | -2.8327 | 0.23 |
| 13460 | *PLCD1* | 1.3418 | 0.068529 | -0.73066 | -1.6128 | 0.23 |
| 13461 | *MET* | 1.3415 | 0.2179 | -0.88055 | -0.90281 | 0.23 |
| 13462 | *LOXL3* | 0.62779 | 0.52933 | -0.47862 | -1.8311 | 0.23 |
| 13463 | *CRABP1* | 2.3734 | 0.18155 | -1.879 | -2.0323 | 0.23 |
| 13464 | *DFNB59* | 1.4269 | 0.45179 | -1.2032 | -2.0999 | 0.23 |
| 13465 | *ACSBG2* | 0.38496 | 0.35826 | -0.06787 | -0.69345 | 0.23 |
| 13466 | *WDR5B* | 1.3123 | 0.33 | -0.96703 | -1.073 | 0.23 |
| 13467 | *TOMM6* | 1.9337 | 0.038871 | -1.298 | -1.6199 | 0.22 |
| 13468 | *F8A1* | 1.7083 | 0.14548 | -1.1793 | -1.3993 | 0.22 |
| 13469 | *RNF103-CHMP3* | 0.99463 | 0.81034 | -1.1316 | -2.2631 | 0.22 |
| 13470 | *ANKRD50* | 1.4545 | -0.05706 | -0.7241 | -1.7004 | 0.22 |
| 13471 | *CYYR1* | 3.1232 | -1.0529 | -1.3973 | -1.8792 | 0.22 |

| | | | | | | |
|---|---|---|---|---|---|---|
| 13472 | *GP9* | 2.976 | -0.63691 | -1.6664 | -1.8941 | 0.22 |
| 13473 | *OR6C74* | 1.0008 | 0.82277 | -1.151 | -1.3336 | 0.22 |
| 13474 | *HECW2* | 1.0594 | 0.49134 | -0.87833 | -0.97499 | 0.22 |
| 13475 | *BDH2* | 0.85877 | 0.057862 | -0.24512 | -1.4326 | 0.22 |
| 13476 | *ZNF852* | 1.0086 | 0.95631 | -1.2935 | -1.6469 | 0.22 |
| 13477 | *OR7A17* | 1.6249 | -0.45803 | -0.49565 | -0.51525 | 0.22 |
| 13478 | *DZIP1L* | 0.59503 | 0.084189 | -0.00816 | -2.0016 | 0.22 |
| 13479 | *MAD2L1* | 0.94392 | 0.32443 | -0.59743 | -1.2151 | 0.22 |
| 13480 | *CRYBA4* | 1.9944 | -0.61338 | -0.71051 | -1.5817 | 0.22 |
| 13481 | *ATP5D* | 1.9731 | -0.44517 | -0.85779 | -1.6632 | 0.22 |
| 13482 | *SFT2D3* | 0.54569 | 0.35573 | -0.23158 | -2.1947 | 0.22 |
| 13483 | *ABHD5* | 0.66949 | 0.16939 | -0.16962 | -0.63511 | 0.22 |
| 13484 | *TMEM151B* | 1.5161 | 0.08533 | -0.93307 | -1.8562 | 0.22 |
| 13485 | *CBX4* | 3.8612 | -1.1934 | -1.9996 | -2.9804 | 0.22 |
| 13486 | *LRP5L* | 1.2144 | 0.078277 | -0.62476 | -0.97952 | 0.22 |
| 13487 | *CDC14A* | 1.3016 | 0.27856 | -0.9124 | -3.6612 | 0.22 |
| 13488 | *GRIN3B* | 1.5634 | -0.37358 | -0.52269 | -1.5707 | 0.22 |
| 13489 | *SLITRK5* | 0.74341 | 0.18925 | -0.26653 | -1.691 | 0.22 |
| 13490 | *FKBP9* | 2.4251 | -0.47447 | -1.2846 | -2.1873 | 0.22 |
| 13491 | *CC2D1B* | 0.86364 | 0.71572 | -0.9135 | -1.9023 | 0.22 |
| 13492 | *ADGRF2* | 1.1249 | 0.056342 | -0.51571 | -1.9112 | 0.22 |
| 13493 | *FNDC7* | 0.93074 | -0.05133 | -0.21407 | -1.8157 | 0.22 |
| 13494 | *MAPK12* | 2.3363 | -0.05575 | -1.6176 | -1.6934 | 0.22 |
| 13495 | *MAP2K2* | 1.7695 | 0.40857 | -1.5165 | -1.6098 | 0.22 |
| 13496 | *MAST4* | 1.1737 | -0.15049 | -0.36195 | -1.9934 | 0.22 |
| 13497 | *MBD3L5* | 1.3596 | 0.6852 | -1.3852 | -1.4442 | 0.22 |
| 13498 | *PHYHIPL* | 0.77685 | 0.39872 | -0.51599 | -0.67137 | 0.22 |
| 13499 | *NR4A3* | 0.87087 | 0.30181 | -0.51321 | -0.88666 | 0.22 |
| 13500 | *KIN* | 1.357 | 0.67524 | -1.3729 | -1.9758 | 0.22 |
| 13501 | *OR6C2* | 3.0566 | -0.62017 | -1.7782 | -2.4015 | 0.22 |
| 13502 | *LSM14A* | 0.37751 | 0.23983 | 0.04075 | -0.26537 | 0.22 |
| 13503 | *C15orf65* | 0.79818 | 0.30088 | -0.443 | -1.7585 | 0.22 |
| 13504 | *AP2B1* | 1.401 | 0.062155 | -0.80713 | -0.92086 | 0.22 |
| 13505 | *STXBP4* | 1.0903 | 0.79324 | -1.2279 | -1.2711 | 0.22 |
| 13506 | *PEPD* | 0.6366 | 0.39867 | -0.37976 | -0.90813 | 0.22 |
| 13507 | *WDR83* | 2.3484 | -0.72095 | -0.97237 | -1.363 | 0.22 |
| 13508 | *TMEM185B* | 3.3601 | -1.2736 | -1.4316 | -2.3811 | 0.22 |
| 13509 | *MFNG* | 3.2808 | -0.93448 | -1.6917 | -1.772 | 0.22 |
| 13510 | *PSG3* | 1.2989 | -0.02908 | -0.61535 | -1.3564 | 0.22 |
| 13511 | *CLASP2* | 0.79312 | 0.46439 | -0.60315 | -1.0564 | 0.22 |
| 13512 | *MRPS18B* | 1.3574 | 0.1059 | -0.80909 | -1.7221 | 0.22 |
| 13513 | *FUT4* | 2.1871 | -0.2083 | -1.3253 | -2.1544 | 0.22 |
| 13514 | *TM2D1* | 0.87942 | -0.10466 | -0.12213 | -1.4076 | 0.22 |
| 13515 | *OR4D11* | 0.83612 | 0.064371 | -0.24794 | -0.43906 | 0.22 |
| 13516 | *PDLIM4* | 4.3509 | -1.7364 | -1.962 | -2.1444 | 0.22 |
| 13517 | *DOK7* | 0.59974 | 0.16736 | -0.11527 | -0.84042 | 0.22 |

| | | | | | | |
|---|---|---|---|---|---|---|
| 13518 | *FAM83A* | 1.4926 | 0.80025 | -1.6411 | -2.2895 | 0.22 |
| 13519 | *HPS4* | 0.82454 | 0.58476 | -0.75778 | -1.696 | 0.22 |
| 13520 | *CALM3* | 1.2434 | -0.0566 | -0.53619 | -1.0019 | 0.22 |
| 13521 | *CRACR2A* | 3.2131 | -0.76405 | -1.7994 | -2.4885 | 0.22 |
| 13522 | *ECSCR* | 1.5214 | -0.29647 | -0.57616 | -1.3605 | 0.22 |
| 13523 | *NDUFB11* | 3.859 | -1.4811 | -1.7292 | -2.4405 | 0.22 |
| 13524 | *CWC15* | 0.42385 | 0.39086 | -0.16624 | -0.43628 | 0.22 |
| 13525 | *CAMK1G* | 2.9499 | -0.76015 | -1.5418 | -2.4961 | 0.22 |
| 13526 | *WNK3* | 0.68969 | 0.48761 | -0.53012 | -1.2014 | 0.22 |
| 13527 | *KDM4C* | 1.0383 | 0.22723 | -0.61836 | -0.75369 | 0.22 |
| 13528 | *ACSF2* | 1.2142 | -0.0083 | -0.55909 | -1.803 | 0.22 |
| 13529 | *PIGW* | 1.6938 | -0.42373 | -0.62423 | -1.2353 | 0.22 |
| 13530 | *MYOT* | 1.1968 | 0.52179 | -1.0733 | -1.2339 | 0.22 |
| 13531 | *THOC1* | 0.68939 | 0.48856 | -0.53312 | -1.6885 | 0.21 |
| 13532 | *MRRF* | 0.79592 | 0.55436 | -0.70558 | -1.7322 | 0.21 |
| 13533 | *ZNF646* | 0.89379 | 0.59291 | -0.84261 | -0.87418 | 0.21 |
| 13534 | *SNAPIN* | 0.91708 | 0.083194 | -0.35638 | -1.0356 | 0.21 |
| 13535 | *NCSTN* | 1.0196 | 0.43794 | -0.8146 | -2.5414 | 0.21 |
| 13536 | *ZNF644* | 2.2859 | -0.46228 | -1.1815 | -1.4932 | 0.21 |
| 13537 | *NKX1-1* | 1.3273 | 0.36346 | -1.0493 | -1.3612 | 0.21 |
| 13538 | *ACSS2* | 1.3275 | -0.26639 | -0.42005 | -0.6937 | 0.21 |
| 13539 | *DDA1* | 1.7905 | -0.54219 | -0.60754 | -1.1401 | 0.21 |
| 13540 | *USH1C* | 0.56882 | 0.17386 | -0.10196 | -1.1607 | 0.21 |
| 13541 | *FGF21* | 1.3035 | 0.28281 | -0.94597 | -1.4841 | 0.21 |
| 13542 | *TRNP1* | 1.4392 | -0.01883 | -0.78096 | -1.0433 | 0.21 |
| 13543 | *ZNF142* | 0.90402 | 0.56025 | -0.82488 | -3.2116 | 0.21 |
| 13544 | *FXN* | 0.90917 | -0.01505 | -0.25612 | -0.46733 | 0.21 |
| 13545 | *MIEF1* | 3.3512 | -1.3098 | -1.4039 | -2.1334 | 0.21 |
| 13546 | *NBN* | 2.3998 | -0.47882 | -1.2837 | -1.3903 | 0.21 |
| 13547 | *ADORA2A* | 0.89512 | 0.012516 | -0.2704 | -0.97237 | 0.21 |
| 13548 | *C17orf53* | 2.3638 | 0.015704 | -1.7438 | -2.6589 | 0.21 |
| 13549 | *PAK7* | 0.99928 | -0.0918 | -0.27178 | -0.5171 | 0.21 |
| 13550 | *DNMBP* | 2.5786 | -0.4187 | -1.5243 | -1.8227 | 0.21 |
| 13551 | *SERTAD1* | 3.4636 | -0.44146 | -2.3871 | -4.0736 | 0.21 |
| 13552 | *TTC30A* | 1.1428 | -0.01665 | -0.49177 | -0.59183 | 0.21 |
| 13553 | *STMN3* | 3.131 | -1.1508 | -1.346 | -1.7947 | 0.21 |
| 13554 | *POLR2G* | 1.7485 | -0.33114 | -0.7835 | -1.1247 | 0.21 |
| 13555 | *FNBP1* | 1.1508 | -0.02433 | -0.49275 | -1.0863 | 0.21 |
| 13556 | *OR8K5* | 1.7575 | -0.04249 | -1.0823 | -1.6841 | 0.21 |
| 13557 | *FTL* | 0.49008 | 0.41457 | -0.27226 | -0.64497 | 0.21 |
| 13558 | *EIF4E1B* | 1.9999 | -0.43422 | -0.93343 | -1.3173 | 0.21 |
| 13559 | *CADM2* | 0.66185 | 0.27562 | -0.30611 | -0.99079 | 0.21 |
| 13560 | *FAM161B* | 1.3098 | -0.05851 | -0.62017 | -0.84801 | 0.21 |
| 13561 | *CBWD2* | 2.1736 | 0.20646 | -1.7495 | -1.9938 | 0.21 |
| 13562 | *BBS4* | 0.95253 | 0.62527 | -0.94777 | -1.8572 | 0.21 |
| 13563 | *C9orf3* | 1.2081 | 0.13115 | -0.70954 | -1.1422 | 0.21 |

| | | | | | | |
|---|---|---|---|---|---|---|
| 13564 | *MDC1* | 1.0339 | 0.40146 | -0.80577 | -0.91165 | 0.21 |
| 13565 | *ELN* | 3.5805 | -1.3878 | -1.5635 | -1.8212 | 0.21 |
| 13566 | *C18orf25* | 0.92554 | 0.019479 | -0.31596 | -1.2335 | 0.21 |
| 13567 | *C2CD4A* | 0.9328 | 0.3488 | -0.65345 | -1.2803 | 0.21 |
| 13568 | *LINC01272* | 1.4104 | 0.50023 | -1.2837 | -3.1211 | 0.21 |
| 13569 | *OR4D10* | 0.50722 | 0.37229 | -0.25317 | -1.0109 | 0.21 |
| 13570 | *EBI3* | 2.5283 | 0.026387 | -1.9284 | -2.1125 | 0.21 |
| 13571 | *CWC27* | 3.8524 | -1.4597 | -1.7675 | -1.7874 | 0.21 |
| 13572 | *AGPAT6* | 0.7106 | 0.10415 | -0.19004 | -1.4913 | 0.21 |
| 13573 | *USP17L10* | 2.0848 | -0.40377 | -1.0564 | -2.6491 | 0.21 |
| 13574 | *DEFB4B* | 0.88746 | 0.2041 | -0.46698 | -0.68153 | 0.21 |
| 13575 | *CDC6* | 1.1494 | 0.44659 | -0.97161 | -1.6724 | 0.21 |
| 13576 | *NDUFC2* | 3.7097 | 0.11284 | -3.1998 | -3.3093 | 0.21 |
| 13577 | *GLT8D2* | 0.74036 | 0.45712 | -0.5765 | -1.4684 | 0.21 |
| 13578 | *ANKZF1* | 0.98159 | 0.017548 | -0.37856 | -0.51204 | 0.21 |
| 13579 | *MUC16* | 2.8403 | -1.0175 | -1.2027 | -1.3926 | 0.21 |
| 13580 | *C10orf113* | 0.52765 | 0.25314 | -0.16097 | -0.35571 | 0.21 |
| 13581 | *CXCR5* | 0.52217 | 0.39939 | -0.30208 | -1.1487 | 0.21 |
| 13582 | *HRK* | 0.61186 | 0.4485 | -0.44173 | -1.2113 | 0.21 |
| 13583 | *MUSTN1* | 0.64585 | 0.4169 | -0.4457 | -1.9968 | 0.21 |
| 13584 | *MYH7B* | 1.2407 | -0.24142 | -0.38385 | -1.9953 | 0.21 |
| 13585 | *RASSF5* | 2.3366 | -0.27055 | -1.4509 | -1.5149 | 0.21 |
| 13586 | *SKA2* | 1.9127 | -0.48688 | -0.81111 | -1.1686 | 0.20 |
| 13587 | *NET1* | 1.6112 | 0.30668 | -1.3032 | -1.3092 | 0.20 |
| 13588 | *KRT2* | 2.2837 | -0.43068 | -1.2393 | -2.312 | 0.20 |
| 13589 | *SBSN* | 0.48577 | 0.14654 | -0.01899 | -0.89474 | 0.20 |
| 13590 | *CHD8* | 2.3559 | -0.26717 | -1.4762 | -3.1008 | 0.20 |
| 13591 | *MZT1* | 2.4533 | -0.79663 | -1.045 | -2.7736 | 0.20 |
| 13592 | *CA10* | 0.88913 | 0.10642 | -0.38503 | -1.6989 | 0.20 |
| 13593 | *ICAM1* | 1.093 | 0.086989 | -0.56972 | -0.98709 | 0.20 |
| 13594 | *SUCLA2* | 2.5014 | 0.11095 | -2.003 | -2.0146 | 0.20 |
| 13595 | *SLC8B1* | 0.45098 | 0.28923 | -0.13111 | -0.88877 | 0.20 |
| 13596 | *EXO1* | 1.4014 | 0.13139 | -0.92375 | -1.8908 | 0.20 |
| 13597 | *HIST3H3* | 4.1398 | -1.7164 | -1.8157 | -2.781 | 0.20 |
| 13598 | *AMACR* | 1.2081 | 1.0064 | -1.6069 | -1.822 | 0.20 |
| 13599 | *ULK2* | 1.133 | -0.00831 | -0.51752 | -0.62514 | 0.20 |
| 13600 | *KRT36* | 1.8874 | 0.16062 | -1.4409 | -1.7533 | 0.20 |
| 13601 | *KCNQ2* | 1.7203 | 0.024455 | -1.1385 | -1.5263 | 0.20 |
| 13602 | *FMR1* | 3.6373 | -0.81522 | -2.2165 | -2.6895 | 0.20 |
| 13603 | *CDK4* | 1.8849 | -0.18547 | -1.0947 | -1.4135 | 0.20 |
| 13604 | *ATL2* | 0.74318 | -0.04339 | -0.09515 | -1.4747 | 0.20 |
| 13605 | *SELE* | 1.1395 | -0.10051 | -0.43475 | -1.082 | 0.20 |
| 13606 | *KNSTRN* | 1.0233 | 0.40928 | -0.82921 | -2.1893 | 0.20 |
| 13607 | *GABRA6* | 1.3095 | -0.33548 | -0.37073 | -1.5781 | 0.20 |
| 13608 | *NRXN1* | 2.8611 | -0.58633 | -1.6731 | -2.3857 | 0.20 |
| 13609 | *TBC1D2B* | 0.46086 | 0.30631 | -0.16574 | -0.60655 | 0.20 |

| | | | | | | |
|---|---|---|---|---|---|---|
| 13610 | *CD72* | 2.3076 | -0.54039 | -1.1659 | -1.2071 | 0.20 |
| 13611 | *MARCH2* | 2.4854 | 0.06609 | -1.9511 | -2.6343 | 0.20 |
| 13612 | *SMAD1* | 1.9849 | -0.40245 | -0.98241 | -1.882 | 0.20 |
| 13613 | *TRAIP* | 3.2773 | -1.06 | -1.6174 | -1.7219 | 0.20 |
| 13614 | *CT45A3* | 1.8572 | 0.018995 | -1.277 | -2.2456 | 0.20 |
| 13615 | *STAR* | 1.6609 | 0.31027 | -1.3728 | -1.8107 | 0.20 |
| 13616 | *RAC2* | 0.8996 | -0.06246 | -0.23977 | -1.0359 | 0.20 |
| 13617 | *OR52A5* | 0.42802 | 0.32263 | -0.15339 | -0.60113 | 0.20 |
| 13618 | *CCDC175* | 1.9444 | -0.48317 | -0.86403 | -0.98842 | 0.20 |
| 13619 | *GSTCD* | 2.8497 | -1.0293 | -1.2234 | -1.4557 | 0.20 |
| 13620 | *ZNF638* | 1.3327 | 0.53082 | -1.2666 | -1.5212 | 0.20 |
| 13621 | *MAP3K5* | 1.5888 | 0.00294 | -0.99563 | -2.6805 | 0.20 |
| 13622 | *GH1* | 2.6191 | -0.97054 | -1.0565 | -1.5797 | 0.20 |
| 13623 | *CLEC14A* | 0.30938 | 0.281 | 0.00168 | -1.316 | 0.20 |
| 13624 | *ATP5E* | 0.62521 | 0.1564 | -0.19007 | -1.7777 | 0.20 |
| 13625 | *NR3C1* | 0.91268 | 0.24704 | -0.56825 | -0.76389 | 0.20 |
| 13626 | *CCDC85C* | 1.7307 | 0.14882 | -1.2884 | -1.7493 | 0.20 |
| 13627 | *SIPA1L2* | 0.82008 | 0.421 | -0.65053 | -1.3363 | 0.20 |
| 13628 | *BCL6* | 1.1424 | -0.26857 | -0.28348 | -0.91813 | 0.20 |
| 13629 | *HAUS2* | 2.753 | 0.62365 | -2.7863 | -3.582 | 0.20 |
| 13630 | *VSTM1* | 0.77824 | 0.59784 | -0.78672 | -1.7507 | 0.20 |
| 13631 | *PAH* | 3.3595 | -1.2934 | -1.479 | -1.9446 | 0.20 |
| 13632 | *BRF2* | 1.1069 | -0.23368 | -0.28742 | -2.3857 | 0.20 |
| 13633 | *C1orf87* | 1.4489 | 0.31774 | -1.1817 | -2.1155 | 0.19 |
| 13634 | *NDUFAF3* | 1.3651 | -0.37125 | -0.40933 | -2.641 | 0.19 |
| 13635 | *RAB11FIP3* | 1.8972 | -0.01574 | -1.2975 | -1.5741 | 0.19 |
| 13636 | *PDE8A* | 0.85257 | 0.50506 | -0.77404 | -1.1016 | 0.19 |
| 13637 | *KRTAP27-1* | 0.89867 | -0.08887 | -0.2263 | -1.3148 | 0.19 |
| 13638 | *EIF6* | 1.288 | -0.14836 | -0.55976 | -0.66598 | 0.19 |
| 13639 | *FAM25G* | 1.1017 | 0.31766 | -0.84069 | -2.0171 | 0.19 |
| 13640 | *F12* | 2.1678 | -0.67837 | -0.91148 | -1.0643 | 0.19 |
| 13641 | *RNMTL1* | 0.4629 | 0.26733 | -0.15339 | -1.154 | 0.19 |
| 13642 | *EARS2* | 0.66068 | 0.48144 | -0.56803 | -1.1834 | 0.19 |
| 13643 | *HEATR6* | 1.821 | -0.59572 | -0.65169 | -1.7658 | 0.19 |
| 13644 | *HSPE1-MOB4* | 0.5253 | 0.046507 | 0.00169 | -1.2185 | 0.19 |
| 13645 | *OR7A10* | 0.32228 | 0.23479 | 0.01638 | -0.02922 | 0.19 |
| 13646 | *FMR1NB* | 1.2483 | -0.11118 | -0.56405 | -1.5954 | 0.19 |
| 13647 | *VAX2* | 1.0777 | 0.60979 | -1.1152 | -1.2529 | 0.19 |
| 13648 | *KDR* | 2.2438 | -0.14536 | -1.5264 | -2.2078 | 0.19 |
| 13649 | *UBXN7* | 2.541 | -0.98015 | -0.98882 | -1.4681 | 0.19 |
| 13650 | *C9orf142* | 3.1382 | -1.0945 | -1.4717 | -2.7277 | 0.19 |
| 13651 | *ZNF808* | 2.299 | -0.38361 | -1.344 | -1.7072 | 0.19 |
| 13652 | *ARL16* | 1.3531 | 0.57419 | -1.3561 | -1.8207 | 0.19 |
| 13653 | *SPAG17* | 1.6244 | -0.06764 | -0.98569 | -1.2036 | 0.19 |
| 13654 | *HSD17B13* | 1.9952 | -0.56586 | -0.85983 | -1.162 | 0.19 |
| 13655 | *CCDC182* | 1.9518 | -0.52739 | -0.85848 | -1.2304 | 0.19 |

| | | | | | | |
|---|---|---|---|---|---|---|
| 13656 | *C1orf159* | 2.7623 | -0.88666 | -1.3098 | -1.9075 | 0.19 |
| 13657 | *ASNS* | 1.6276 | -0.30993 | -0.75304 | -1.9511 | 0.19 |
| 13658 | *STK32C* | 0.7425 | -0.0717 | -0.10619 | -1.4572 | 0.19 |
| 13659 | *HMGCR* | 2.1881 | -0.01451 | -1.61 | -2.042 | 0.19 |
| 13660 | *CEACAM19* | 1.576 | -0.30927 | -0.70361 | -0.93555 | 0.19 |
| 13661 | *VAMP3* | 3.5867 | -1.4028 | -1.6208 | -1.7188 | 0.19 |
| 13662 | *GLRB* | 0.68717 | 0.018621 | -0.14403 | -0.26206 | 0.19 |
| 13663 | *SCML1* | 0.65853 | 0.25544 | -0.35232 | -0.49247 | 0.19 |
| 13664 | *ZNF362* | 0.2833 | 0.21501 | 0.06326 | -1.0317 | 0.19 |
| 13665 | *PNLIPRP2* | 1.0985 | 0.33158 | -0.86889 | -0.86903 | 0.19 |
| 13666 | *MUCL1* | 0.49303 | 0.19242 | -0.12495 | -1.9434 | 0.19 |
| 13667 | *LIPI* | 2.3786 | -0.58289 | -1.2357 | -1.4852 | 0.19 |
| 13668 | *HNRNPR* | 2.1443 | 0.50344 | -2.088 | -2.2528 | 0.19 |
| 13669 | *NAE1* | 2.4969 | -0.51377 | -1.4234 | -1.8478 | 0.19 |
| 13670 | *TGFBR3* | 0.76271 | 0.20829 | -0.41266 | -1.6318 | 0.19 |
| 13671 | *PPFIA1* | 0.72624 | 0.45591 | -0.62423 | -2.6201 | 0.19 |
| 13672 | *ST6GALNAC5* | 2.8836 | -1.0335 | -1.2927 | -1.8645 | 0.19 |
| 13673 | *JTB* | 1.9931 | -0.62514 | -0.81058 | -2.0032 | 0.19 |
| 13674 | *FAM111A* | 0.58992 | 0.21702 | -0.25291 | -1.6913 | 0.18 |
| 13675 | *FRYL* | 1.1603 | 0.53997 | -1.1466 | -1.8651 | 0.18 |
| 13676 | *CASP14* | 1.0399 | -0.08187 | -0.40519 | -0.61421 | 0.18 |
| 13677 | *ADAMTS13* | 0.71578 | 0.007553 | -0.17073 | -0.95233 | 0.18 |
| 13678 | *SPHK2* | 0.6246 | 0.21893 | -0.29171 | -0.90412 | 0.18 |
| 13679 | *HIRA* | 0.66269 | 0.47207 | -0.5832 | -1.6179 | 0.18 |
| 13680 | *SFRP5* | 1.5277 | -0.28384 | -0.69237 | -0.90028 | 0.18 |
| 13681 | *IL12RB1* | 2.13 | -0.46738 | -1.1131 | -1.8574 | 0.18 |
| 13682 | *PRPF40A* | 1.949 | -0.56681 | -0.83277 | -1.3148 | 0.18 |
| 13683 | *RGS2* | 1.2515 | 0.19827 | -0.90104 | -1.5115 | 0.18 |
| 13684 | *PKD1L3* | 0.79154 | -0.06121 | -0.18229 | -1.8106 | 0.18 |
| 13685 | *INADL* | 0.904 | 0.1206 | -0.47685 | -0.98728 | 0.18 |
| 13686 | *LOC100130705* | 0.39417 | 0.28993 | -0.13707 | -2.7127 | 0.18 |
| 13687 | *NHP2* | 3.0909 | -0.62093 | -1.923 | -2.0535 | 0.18 |
| 13688 | *CEP250* | 0.74157 | 0.25842 | -0.45373 | -0.87688 | 0.18 |
| 13689 | *TRIM21* | 1.9286 | -0.6785 | -0.70391 | -1.7345 | 0.18 |
| 13690 | *TNNI2* | 1.5176 | 0.11609 | -1.0878 | -2.1105 | 0.18 |
| 13691 | *CYP1A2* | 0.26487 | 0.18081 | 0.10017 | 0.066827 | 0.18 |
| 13692 | *SRMS* | 0.43414 | 0.17032 | -0.06045 | -0.58752 | 0.18 |
| 13693 | *UNK* | 1.3908 | 0.56905 | -1.4172 | -2.8071 | 0.18 |
| 13694 | *TAF13* | 0.46736 | 0.16363 | -0.08887 | -0.23814 | 0.18 |
| 13695 | *YTHDC2* | 0.43844 | 0.2928 | -0.18916 | -1.2198 | 0.18 |
| 13696 | *SNAI1* | 1.4818 | -0.07367 | -0.86744 | -1.3711 | 0.18 |
| 13697 | *FBRSL1* | 2.1351 | -0.58661 | -1.009 | -1.1553 | 0.18 |
| 13698 | *SEBOX* | 0.75952 | 0.65856 | -0.87881 | -1.6395 | 0.18 |
| 13699 | *POC1B* | 0.72652 | 0.25626 | -0.44353 | -2.2692 | 0.18 |
| 13700 | *TYSND1* | 0.53535 | 0.16279 | -0.15961 | -0.37519 | 0.18 |
| 13701 | *FCGR3B* | 2.8071 | -0.76529 | -1.5043 | -2.2945 | 0.18 |

| | | | | | | |
|---|---|---|---|---|---|---|
| 13702 | *ZNF32* | 1.0575 | 0.054536 | -0.57477 | -1.5884 | 0.18 |
| 13703 | *KIAA1598* | 0.30878 | 0.29117 | -0.06364 | -0.17588 | 0.18 |
| 13704 | *POC5* | 1.5872 | 0.27765 | -1.3294 | -1.6796 | 0.18 |
| 13705 | *DOCK11* | 0.5967 | 0.27836 | -0.33984 | -1.3517 | 0.18 |
| 13706 | *FXYD5* | 0.60194 | 0.57147 | -0.63826 | -1.6306 | 0.18 |
| 13707 | *TRIM55* | 1.214 | 0.49883 | -1.1803 | -1.3437 | 0.18 |
| 13708 | *RPSA* | 1.4198 | 0.40844 | -1.2958 | -1.392 | 0.18 |
| 13709 | *GMNC* | 0.80424 | 0.15744 | -0.43005 | -1.7493 | 0.18 |
| 13710 | *SLX4* | 1.2549 | 0.078951 | -0.8038 | -1.463 | 0.18 |
| 13711 | *NUBP2* | 0.64894 | -0.03634 | -0.08307 | -1.3232 | 0.18 |
| 13712 | *OR52N1* | 2.2623 | -0.68003 | -1.0533 | -1.0693 | 0.18 |
| 13713 | *CYP4B1* | 1.1448 | 0.4513 | -1.0672 | -1.4507 | 0.18 |
| 13714 | *ZBTB45* | 3.285 | -0.42841 | -2.329 | -2.4094 | 0.18 |
| 13715 | *PPA2* | 1.6179 | -0.35546 | -0.73577 | -2.0452 | 0.18 |
| 13716 | *CLDN24* | 0.83318 | 0.077055 | -0.3836 | -0.42756 | 0.18 |
| 13717 | *WBSCR22* | 1.2985 | -0.05308 | -0.71936 | -1.2608 | 0.18 |
| 13718 | *JAG1* | 1.9214 | -0.03091 | -1.3647 | -1.5253 | 0.18 |
| 13719 | *CCDC78* | 2.8147 | -0.58575 | -1.7034 | -3.2632 | 0.18 |
| 13720 | *LGALSL* | 1.0776 | 0.29569 | -0.84841 | -0.98568 | 0.17 |
| 13721 | *DCAF4L2* | 0.65435 | 0.0961 | -0.22589 | -0.68466 | 0.17 |
| 13722 | *RAB27A* | 2.0522 | -0.18328 | -1.345 | -2.0466 | 0.17 |
| 13723 | *PRR14L* | 0.40747 | 0.17555 | -0.05918 | -1.1742 | 0.17 |
| 13724 | *RBM14* | 1.4914 | 0.5188 | -1.4864 | -1.8157 | 0.17 |
| 13725 | *C20orf194* | 1.7194 | 0.56966 | -1.7655 | -2.0197 | 0.17 |
| 13726 | *PYGO2* | 0.70188 | 0.03708 | -0.21563 | -1.0372 | 0.17 |
| 13727 | *ANKRD30A* | 2.5346 | -0.52122 | -1.4911 | -2.3918 | 0.17 |
| 13728 | *TSPY1* | 0.81109 | -0.12213 | -0.168 | -1.8275 | 0.17 |
| 13729 | *NELFA* | 0.63586 | 0.060232 | -0.17517 | -1.2249 | 0.17 |
| 13730 | *SRGAP1* | 0.75055 | 0.42694 | -0.65752 | -0.78097 | 0.17 |
| 13731 | *HSPA12B* | 1.8826 | -0.32122 | -1.042 | -1.1108 | 0.17 |
| 13732 | *CCR2* | 2.0908 | -0.13756 | -1.4341 | -2.5189 | 0.17 |
| 13733 | *PACS1* | 1.4181 | -0.00824 | -0.89167 | -1.1991 | 0.17 |
| 13734 | *HECA* | 2.7146 | -0.53058 | -1.6663 | -1.7766 | 0.17 |
| 13735 | *TOR1A* | 0.76142 | 0.47992 | -0.72479 | -1.2004 | 0.17 |
| 13736 | *E2F5* | 0.40145 | 0.08403 | 0.0303 | -0.16685 | 0.17 |
| 13737 | *PABPC1* | 3.163 | -1.1251 | -1.5227 | -1.668 | 0.17 |
| 13738 | *TCTE3* | 0.77639 | 0.63802 | -0.89977 | -1.7999 | 0.17 |
| 13739 | *MCU* | 1.1612 | -0.02072 | -0.62629 | -1.7434 | 0.17 |
| 13740 | *C4B* | 0.58412 | 0.057252 | -0.12795 | -1.8707 | 0.17 |
| 13741 | *PPID* | 0.89512 | 0.33765 | -0.72222 | -0.92907 | 0.17 |
| 13742 | *NRXN2* | 0.8464 | -0.15394 | -0.18195 | -0.90501 | 0.17 |
| 13743 | *SVIL* | 1.1927 | 0.7076 | -1.3899 | -2.033 | 0.17 |
| 13744 | *HSFX1* | 0.64437 | 0.12975 | -0.2641 | -1.0057 | 0.17 |
| 13745 | *CEP68* | 0.66324 | 0.004409 | -0.15853 | -0.21283 | 0.17 |
| 13746 | *OR52W1* | 1.2467 | -0.2957 | -0.44267 | -0.53747 | 0.17 |
| 13747 | *GALNTL6* | 0.66502 | 0.073132 | -0.22988 | -0.50104 | 0.17 |

| | | | | | | |
|---|---|---|---|---|---|---|
| 13748 | *VANGL2* | 2.5132 | -0.83143 | -1.1735 | -1.219 | 0.17 |
| 13749 | *PHF10* | 0.47043 | 0.17495 | -0.13754 | -0.59521 | 0.17 |
| 13750 | *LCA5L* | 0.71435 | 0.36734 | -0.57408 | -1.7303 | 0.17 |
| 13751 | *PTBP1* | 1.2183 | 0.042235 | -0.75293 | -1.5598 | 0.17 |
| 13752 | *PFDN6* | 0.6799 | 0.53302 | -0.70532 | -1.6343 | 0.17 |
| 13753 | *APTX* | 0.8626 | 0.42556 | -0.78255 | -1.8578 | 0.17 |
| 13754 | *CCDC166* | 0.96676 | -0.03798 | -0.42369 | -0.53799 | 0.17 |
| 13755 | *DHX16* | 1.6654 | -0.19706 | -0.96349 | -2.0791 | 0.17 |
| 13756 | *DGKI* | 0.95311 | 0.30333 | -0.75213 | -1.1929 | 0.17 |
| 13757 | *KRT38* | 1.4172 | -0.1693 | -0.74459 | -0.8147 | 0.17 |
| 13758 | *HOXA10* | 2.8928 | -1.084 | -1.3055 | -2.2507 | 0.17 |
| 13759 | *SLC13A3* | 0.35016 | 0.10642 | 0.04624 | -1.1734 | 0.17 |
| 13760 | *HP1BP3* | 0.62997 | 0.008845 | -0.13622 | -1.3098 | 0.17 |
| 13761 | *ZNF174* | 1.9841 | -0.37837 | -1.1039 | -2.3949 | 0.17 |
| 13762 | *GSTM2* | 2.182 | -0.54366 | -1.1367 | -1.53 | 0.17 |
| 13763 | *RP9* | 1.1009 | 0.15099 | -0.75068 | -0.77521 | 0.17 |
| 13764 | *FAM47E* | 2.2433 | -0.76106 | -0.98239 | -0.99368 | 0.17 |
| 13765 | *TBL1Y* | 1.6021 | 0.041514 | -1.1442 | -1.5569 | 0.17 |
| 13766 | *COPB1* | 1.867 | -0.5701 | -0.79813 | -1.9863 | 0.17 |
| 13767 | *ERAL1* | 2.8685 | -0.83459 | -1.5352 | -1.5824 | 0.17 |
| 13768 | *SASH1* | 1.18 | 0.52356 | -1.2059 | -1.7065 | 0.17 |
| 13769 | *CYP2B6* | 0.85199 | 0.27104 | -0.62613 | -1.7474 | 0.17 |
| 13770 | *PTPRD* | 2.5806 | -0.99508 | -1.0889 | -1.3287 | 0.17 |
| 13771 | *SUN5* | 0.65897 | 0.25204 | -0.41486 | -1.4703 | 0.17 |
| 13772 | *MOCOS* | 0.40421 | 0.075758 | 0.01511 | -0.36575 | 0.17 |
| 13773 | *RNASE7* | 2.9097 | -1.0863 | -1.3288 | -1.5375 | 0.16 |
| 13774 | *ALDH8A1* | 0.26558 | 0.12223 | 0.10565 | -0.34077 | 0.16 |
| 13775 | *C10orf12* | 0.65175 | 0.50479 | -0.66363 | -1.4597 | 0.16 |
| 13776 | *NR1D2* | 1.1109 | 0.26696 | -0.88503 | -2.7336 | 0.16 |
| 13777 | *ACAP2* | 0.37802 | 0.066303 | 0.04816 | -2.0074 | 0.16 |
| 13778 | *SLCO4A1* | 0.50712 | 0.42651 | -0.44177 | -1.0838 | 0.16 |
| 13779 | *KATNB1* | 1.0597 | -0.1865 | -0.38219 | -1.6098 | 0.16 |
| 13780 | *SYN1* | 0.8539 | -0.05048 | -0.3129 | -1.5425 | 0.16 |
| 13781 | *GNPDA1* | 1.3454 | 0.11456 | -0.9703 | -1.175 | 0.16 |
| 13782 | *HIST1H3A* | 1.0763 | 0.033945 | -0.62065 | -1.1287 | 0.16 |
| 13783 | *PLEKHA8* | 2.5296 | -0.59045 | -1.4498 | -4.0338 | 0.16 |
| 13784 | *IL37* | 0.97498 | -0.08474 | -0.40136 | -0.65368 | 0.16 |
| 13785 | *GALR2* | 1.027 | -0.08922 | -0.45012 | -1.2004 | 0.16 |
| 13786 | *BCS1L* | 0.361 | 0.23712 | -0.11083 | -1.7658 | 0.16 |
| 13787 | *LEPR* | 1.2927 | -0.34816 | -0.45803 | -1.0094 | 0.16 |
| 13788 | *MGA* | 3.6625 | -1.3246 | -1.8523 | -2.1709 | 0.16 |
| 13789 | *RGS20* | 0.57604 | 0.1221 | -0.21258 | -1.5121 | 0.16 |
| 13790 | *TUBB8* | 3.3842 | -0.02864 | -2.8702 | -3.114 | 0.16 |
| 13791 | *SLC25A26* | 0.96387 | -0.07528 | -0.40338 | -0.59959 | 0.16 |
| 13792 | *PEX3* | 2.8318 | -0.99079 | -1.3565 | -4.0005 | 0.16 |
| 13793 | *EMB* | 2.139 | -0.29427 | -1.3607 | -1.4682 | 0.16 |

| | | | | | | |
|---|---|---|---|---|---|---|
| 13794 | *SMARCA1* | 1.0552 | 0.59085 | -1.1626 | -1.4576 | 0.16 |
| 13795 | *PHACTR2* | 1.63 | 0.78082 | -1.928 | -2.1354 | 0.16 |
| 13796 | *STX6* | 0.75462 | 0.027653 | -0.30003 | -1.5297 | 0.16 |
| 13797 | *ZNF843* | 0.83233 | -0.16607 | -0.18502 | -1.7797 | 0.16 |
| 13798 | *DUSP27* | 1.0165 | 0.28353 | -0.81883 | -0.9272 | 0.16 |
| 13799 | *IKBIP* | 0.28028 | 0.13507 | 0.06584 | -0.66082 | 0.16 |
| 13800 | *PPM1A* | 2.5481 | -0.52034 | -1.547 | -1.9269 | 0.16 |
| 13801 | *TRANK1* | 0.52555 | 0.1916 | -0.23642 | -1.5052 | 0.16 |
| 13802 | *CDK18* | 0.94744 | 0.93574 | -1.4036 | -1.7945 | 0.16 |
| 13803 | *SAP18* | 2.6261 | -0.93407 | -1.2135 | -2.6188 | 0.16 |
| 13804 | *EEF2* | 1.1276 | -0.15953 | -0.49073 | -1.3513 | 0.16 |
| 13805 | *GPR152* | 0.90196 | -0.17588 | -0.24877 | -1.3295 | 0.16 |
| 13806 | *GSG1L* | 0.89923 | 0.42891 | -0.85084 | -1.5754 | 0.16 |
| 13807 | *SLC16A7* | 0.49564 | 0.23974 | -0.25936 | -1.3057 | 0.16 |
| 13808 | *CPSF2* | 2.1228 | -0.80548 | -0.84136 | -2.1264 | 0.16 |
| 13809 | *SMIM14* | 1.0935 | -0.09378 | -0.52397 | -1.8398 | 0.16 |
| 13810 | *NR1H3* | 0.88319 | 0.0898 | -0.49727 | -1.0993 | 0.16 |
| 13811 | *GSDMA* | 1.2163 | -0.29305 | -0.44787 | -0.77963 | 0.16 |
| 13812 | *ANAPC10* | 1.7462 | -0.61889 | -0.65199 | -0.77289 | 0.16 |
| 13813 | *OR5W2* | 0.51467 | 0.4365 | -0.47587 | -0.56274 | 0.16 |
| 13814 | *FOCAD* | 1.097 | -0.19562 | -0.42631 | -0.70944 | 0.16 |
| 13815 | *GALT* | 0.76877 | -0.0645 | -0.22921 | -0.62476 | 0.16 |
| 13816 | *GATSL3* | 1.7436 | 1.2554 | -2.5243 | -2.716 | 0.16 |
| 13817 | *AIPL1* | 0.4161 | 0.38146 | -0.32296 | -0.59001 | 0.16 |
| 13818 | *SELP* | 1.7146 | -0.59738 | -0.64313 | -2.1205 | 0.16 |
| 13819 | *PCGF2* | 1.0605 | 0.30878 | -0.89595 | -1.4625 | 0.16 |
| 13820 | *LNX2* | 1.9705 | -0.15394 | -1.344 | -1.4162 | 0.16 |
| 13821 | *RPL39* | 0.5536 | 0.040832 | -0.12247 | -3.1578 | 0.16 |
| 13822 | *KRTCAP2* | 1.6602 | -0.46589 | -0.7229 | -2.0698 | 0.16 |
| 13823 | *ABCB8* | 3.1954 | -1.219 | -1.5052 | -1.856 | 0.16 |
| 13824 | *KDM4B* | 2.8658 | -0.3205 | -2.075 | -2.1187 | 0.16 |
| 13825 | *GPR15* | 2.768 | -0.97792 | -1.3198 | -1.6266 | 0.16 |
| 13826 | *TBX21* | 1.0564 | 0.12345 | -0.70976 | -1.473 | 0.16 |
| 13827 | *TMEM105* | 1.4008 | 0.53727 | -1.4681 | -1.8471 | 0.16 |
| 13828 | *MC3R* | 0.56252 | 0.20143 | -0.29422 | -1.1696 | 0.16 |
| 13829 | *VPS52* | 0.48693 | 0.48324 | -0.50053 | -2.1466 | 0.16 |
| 13830 | *SLC39A1* | 2.0128 | -0.25291 | -1.2903 | -1.525 | 0.16 |
| 13831 | *CCM2* | 0.821 | 0.58811 | -0.94052 | -1.5061 | 0.16 |
| 13832 | *EDEM1* | 1.5701 | 0.26685 | -1.3694 | -2.3036 | 0.16 |
| 13833 | *TUBB4B* | 1.3107 | 0.46124 | -1.3053 | -2.6289 | 0.16 |
| 13834 | *SHCBP1L* | 1.6737 | 0.049791 | -1.2574 | -1.6934 | 0.16 |
| 13835 | *LOC100131303* | 1.0134 | 0.9513 | -1.4994 | -2.6683 | 0.16 |
| 13836 | *CSMD1* | 0.65341 | 0.005565 | -0.19404 | -1.6125 | 0.15 |
| 13837 | *IFT27* | 1.701 | -0.46979 | -0.76635 | -1.1065 | 0.15 |
| 13838 | *OBP2A* | 0.58143 | 0.54991 | -0.66752 | -1.4087 | 0.15 |
| 13839 | *DYRK1B* | 3.1024 | -1.3183 | -1.3203 | -2.2108 | 0.15 |

| | | | | | | |
|---|---|---|---|---|---|---|
| 13840 | *CHCHD5* | 0.75892 | 0.27246 | -0.56769 | -0.78725 | 0.15 |
| 13841 | *SLC15A5* | 0.60924 | 0.56505 | -0.71093 | -0.98239 | 0.15 |
| 13842 | *LBH* | 3.5545 | -0.97648 | -2.1151 | -2.2232 | 0.15 |
| 13843 | *SIKE1* | 0.56182 | 0.24092 | -0.33985 | -0.63939 | 0.15 |
| 13844 | *CDIPT* | 1.4664 | 0.95155 | -1.9554 | -2.6593 | 0.15 |
| 13845 | *IFT43* | 0.48201 | 0.18951 | -0.20906 | -2.5707 | 0.15 |
| 13846 | *ARPC4* | 0.44646 | 0.19552 | -0.17979 | -1.1986 | 0.15 |
| 13847 | *SLC35G2* | 0.93085 | -0.03695 | -0.43201 | -0.46738 | 0.15 |
| 13848 | *SLC9A6* | 0.58664 | -0.05066 | -0.07435 | -2.3615 | 0.15 |
| 13849 | *CEP83* | 1.1883 | 0.04123 | -0.7679 | -1.4061 | 0.15 |
| 13850 | *FBXW11* | 0.62181 | 0.11287 | -0.27453 | -1.4234 | 0.15 |
| 13851 | *CCDC102A* | 1.5636 | 0.78697 | -1.8906 | -2.2021 | 0.15 |
| 13852 | *ACOT1* | 1.4921 | -0.46761 | -0.56501 | -0.89887 | 0.15 |
| 13853 | *TRIM67* | 2.2325 | -0.21974 | -1.5538 | -2.1919 | 0.15 |
| 13854 | *AFF4* | 1.569 | -0.40692 | -0.70327 | -1.4141 | 0.15 |
| 13855 | *METTL13* | 1.9775 | -0.67684 | -0.84248 | -0.94576 | 0.15 |
| 13856 | *ZNF93* | 0.53823 | 0.22862 | -0.30878 | -1.3878 | 0.15 |
| 13857 | *APITD1-CORT* | 0.93984 | -0.20512 | -0.27675 | -1.1511 | 0.15 |
| 13858 | *CLEC11A* | 1.0042 | -0.07909 | -0.46733 | -2.3178 | 0.15 |
| 13859 | *SPEF1* | 1.6975 | -0.31062 | -0.92911 | -1.4258 | 0.15 |
| 13860 | *ARL17A* | 1.6599 | -0.30208 | -0.90084 | -1.8264 | 0.15 |
| 13861 | *FCAR* | 0.67742 | 0.079385 | -0.30038 | -1.1687 | 0.15 |
| 13862 | *DEXI* | 1.2889 | -0.24005 | -0.5925 | -0.74912 | 0.15 |
| 13863 | *HOMER2* | 1.5905 | -0.01358 | -1.1212 | -1.1533 | 0.15 |
| 13864 | *MTFR1L* | 0.51892 | 0.27719 | -0.34111 | -0.42062 | 0.15 |
| 13865 | *TMEM87A* | 1.0207 | 0.025575 | -0.59163 | -1.1587 | 0.15 |
| 13866 | *CXorf40A* | 1.1417 | -0.1851 | -0.50283 | -0.63125 | 0.15 |
| 13867 | *OR2T35* | 0.63315 | 0.15623 | -0.33568 | -2.0804 | 0.15 |
| 13868 | *NNT* | 3.2727 | -0.95645 | -1.8636 | -1.9953 | 0.15 |
| 13869 | *MGAT1* | 0.38916 | 0.17891 | -0.11543 | -1.9282 | 0.15 |
| 13870 | *CD151* | 1.3249 | -0.39971 | -0.47376 | -2.3349 | 0.15 |
| 13871 | *DEFB119* | 2.5572 | -1.0283 | -1.0775 | -1.566 | 0.15 |
| 13872 | *TMEM200C* | 1.2835 | 0.34151 | -1.1737 | -1.9686 | 0.15 |
| 13873 | *COX6B1* | 1.7527 | 0.17015 | -1.4717 | -2.1984 | 0.15 |
| 13874 | *CORO1C* | 1.3871 | -0.45339 | -0.48336 | -0.77227 | 0.15 |
| 13875 | *UTP23* | 0.32177 | 0.29429 | -0.16624 | -3.2662 | 0.15 |
| 13876 | *COX5A* | 0.59682 | 0.050482 | -0.19778 | -1.5243 | 0.15 |
| 13877 | *LIPF* | 1.129 | -0.20506 | -0.47496 | -1.7364 | 0.15 |
| 13878 | *EEF2KMT* | 0.56528 | 0.2374 | -0.35441 | -0.66274 | 0.15 |
| 13879 | *CENPB* | 2.3828 | -0.44353 | -1.4913 | -1.72 | 0.15 |
| 13880 | *BPNT1* | 2.1477 | -0.35128 | -1.3488 | -1.7221 | 0.15 |
| 13881 | *CCNDBP1* | 0.97221 | 0.16922 | -0.69418 | -2.3936 | 0.15 |
| 13882 | *ZC3H12C* | 1.2068 | -0.30878 | -0.45119 | -1.4034 | 0.15 |
| 13883 | *SMAD6* | 0.55507 | 0.44466 | -0.55296 | -2.2044 | 0.15 |
| 13884 | *IGSF8* | 1.3451 | 0.080851 | -0.9797 | -1.6428 | 0.15 |
| 13885 | *TRIM54* | 1.3377 | 0.33888 | -1.2306 | -1.5424 | 0.15 |

| | | | | | | |
|---|---|---|---|---|---|---|
| 13886 | *BET1L* | 0.7044 | 0.54846 | -0.80743 | -1.5099 | 0.15 |
| 13887 | *ATXN2L* | 1.8804 | -0.42936 | -1.0058 | -1.5073 | 0.15 |
| 13888 | *FBXL13* | 0.70352 | 0.006056 | -0.26575 | -1.3694 | 0.15 |
| 13889 | *HNRNPC* | 0.44554 | 0.27562 | -0.27814 | -1.2071 | 0.15 |
| 13890 | *OR4L1* | 0.20228 | 0.14084 | 0.09972 | -0.13361 | 0.15 |
| 13891 | *SLITRK3* | 3.0361 | -0.86556 | -1.728 | -2.3293 | 0.15 |
| 13892 | *HORMAD2* | 0.28841 | 0.16681 | -0.01298 | -1.5282 | 0.15 |
| 13893 | *CASP2* | 1.0713 | -0.15051 | -0.47857 | -2.0407 | 0.15 |
| 13894 | *RBX1* | 1.4857 | -0.3205 | -0.72348 | -1.2572 | 0.15 |
| 13895 | *DEFB110* | 0.61104 | 0.30864 | -0.47854 | -1.8212 | 0.15 |
| 13896 | *SUFU* | 1.2461 | -0.02092 | -0.78588 | -1.3078 | 0.15 |
| 13897 | *ZNF813* | 1.6635 | -0.0083 | -1.2159 | -1.683 | 0.15 |
| 13898 | *SP100* | 0.74969 | 0.28567 | -0.59763 | -2.1713 | 0.15 |
| 13899 | *DIAPH1* | 0.5 | 0.12678 | -0.18937 | -0.79202 | 0.15 |
| 13900 | *DDX3X* | 0.77227 | 0.026519 | -0.36177 | -0.44353 | 0.15 |
| 13901 | *SLC11A2* | 1.571 | -0.45531 | -0.6794 | -2.4067 | 0.15 |
| 13902 | *TAS2R13* | 2.3604 | -0.25984 | -1.6648 | -1.8038 | 0.15 |
| 13903 | *NBPF10* | 1.3119 | 0.34565 | -1.2234 | -1.4955 | 0.14 |
| 13904 | *FLG* | 0.28793 | 0.10767 | 0.03809 | -1.0304 | 0.14 |
| 13905 | *TMIE* | 2.0611 | -0.72896 | -0.90014 | -0.93837 | 0.14 |
| 13906 | *TRIM50* | 1.2122 | -0.36582 | -0.41454 | -0.91427 | 0.14 |
| 13907 | *POLR2B* | 2.252 | -0.57408 | -1.2464 | -1.6469 | 0.14 |
| 13908 | *MAGEC1* | 1.2846 | -0.33893 | -0.51494 | -1.6026 | 0.14 |
| 13909 | *PRDM1* | 1.1426 | -0.25915 | -0.45302 | -1.63 | 0.14 |
| 13910 | *EPHA1* | 1.0053 | -0.19925 | -0.37641 | -0.77697 | 0.14 |
| 13911 | *CLSTN3* | 2.9498 | -1.0024 | -1.5182 | -2.017 | 0.14 |
| 13912 | *ZNF580* | 0.79143 | 0.64117 | -1.0044 | -1.19 | 0.14 |
| 13913 | *BVES* | 0.74392 | 0.21633 | -0.53264 | -0.9341 | 0.14 |
| 13914 | *USP6* | 1.4409 | -0.30312 | -0.71032 | -1.6597 | 0.14 |
| 13915 | *SGPL1* | 0.55629 | 0.31054 | -0.43998 | -1.6709 | 0.14 |
| 13916 | *MCM5* | 0.3392 | 0.31701 | -0.22972 | -0.62223 | 0.14 |
| 13917 | *SYF2* | 0.88227 | 0.38268 | -0.83883 | -1.9558 | 0.14 |
| 13918 | *FCHO1* | 1.1713 | 0.56444 | -1.3104 | -2.0629 | 0.14 |
| 13919 | *C20orf141* | 0.36369 | 0.17121 | -0.10981 | -0.40863 | 0.14 |
| 13920 | *MKRN3* | 0.766 | 0.059575 | -0.40104 | -2.2631 | 0.14 |
| 13921 | *TUBGCP6* | 1.2539 | -0.04886 | -0.78229 | -1.039 | 0.14 |
| 13922 | *DSN1* | 3.5811 | -1.2743 | -1.8851 | -2.3871 | 0.14 |
| 13923 | *MFSD5* | 1.9131 | 0.089165 | -1.5809 | -1.9237 | 0.14 |
| 13924 | *SHOX* | 0.78036 | -0.02394 | -0.33524 | -1.2896 | 0.14 |
| 13925 | *ASB11* | 1.6128 | 0.011742 | -1.2036 | -1.8238 | 0.14 |
| 13926 | *RCE1* | 1.7378 | -0.54264 | -0.77428 | -1.6575 | 0.14 |
| 13927 | *MROH2A* | 0.78918 | -0.05413 | -0.31535 | -1.8707 | 0.14 |
| 13928 | *MRPL57* | 4.1723 | -1.244 | -2.5103 | -2.9525 | 0.14 |
| 13929 | *MTMR9* | 2.5543 | -0.51664 | -1.6199 | -1.7697 | 0.14 |
| 13930 | *HSD11B1* | 1.6042 | 0.17307 | -1.3599 | -2.2621 | 0.14 |
| 13931 | *TPM2* | 0.74854 | -0.13276 | -0.19915 | -1.5436 | 0.14 |

| | | | | | | |
|---|---|---|---|---|---|---|
| 13932 | *PSPC1* | 0.41169 | 0.25176 | -0.24697 | -0.64851 | 0.14 |
| 13933 | *UQCRC2* | 2.5295 | -0.78718 | -1.3273 | -1.4917 | 0.14 |
| 13934 | *AQP5* | 0.93573 | 0.40848 | -0.92926 | -2.8229 | 0.14 |
| 13935 | *TRAF7* | 1.0304 | 0.57356 | -1.1891 | -2.071 | 0.14 |
| 13936 | *SYT13* | 1.2711 | 0.94044 | -1.7971 | -2.7728 | 0.14 |
| 13937 | *MS4A4A* | 1.5743 | -0.47211 | -0.6878 | -1.8751 | 0.14 |
| 13938 | *LPIN3* | 1.8631 | -0.16995 | -1.2788 | -1.4572 | 0.14 |
| 13939 | *CASP6* | 3.5202 | -1.4545 | -1.6524 | -1.9199 | 0.14 |
| 13940 | *SETD4* | 0.66956 | 0.074609 | -0.3316 | -1.2797 | 0.14 |
| 13941 | *APBB1* | 1.9227 | -0.31587 | -1.1943 | -1.4448 | 0.14 |
| 13942 | *ZSCAN26* | 2.1769 | 0.11889 | -1.8834 | -2.262 | 0.14 |
| 13943 | *SPATA18* | 1.0238 | -0.13185 | -0.47964 | -1.0164 | 0.14 |
| 13944 | *NPHP3* | 3.6465 | -1.5568 | -1.6776 | -2.0613 | 0.14 |
| 13945 | *CACNB3* | 0.96224 | 0.67671 | -1.227 | -1.2485 | 0.14 |
| 13946 | *TAAR8* | 0.7183 | 0.2892 | -0.596 | -1.6964 | 0.14 |
| 13947 | *LRRC2* | 0.7106 | 0.10206 | -0.40118 | -1.7974 | 0.14 |
| 13948 | *CCL8* | 0.42665 | 0.12445 | -0.13962 | -2.2046 | 0.14 |
| 13949 | *AMD1* | 1.2114 | -0.09709 | -0.70289 | -1.9772 | 0.14 |
| 13950 | *VPS41* | 1.8053 | -0.68934 | -0.70532 | -1.1912 | 0.14 |
| 13951 | *MSMO1* | 0.98504 | 0.24839 | -0.82311 | -2.3776 | 0.14 |
| 13952 | *FAAH* | 0.60626 | 0.071134 | -0.26717 | -0.44353 | 0.14 |
| 13953 | *COA6* | 0.54925 | 0.25407 | -0.39318 | -0.96673 | 0.14 |
| 13954 | *TMEM40* | 1.6796 | -0.36086 | -0.90922 | -1.5722 | 0.14 |
| 13955 | *PCDHGA6* | 0.79847 | -0.03097 | -0.35896 | -0.47632 | 0.14 |
| 13956 | *BTK* | 1.1966 | -0.33584 | -0.45236 | -1.8046 | 0.14 |
| 13957 | *SH3GL1* | 1.1082 | -0.15226 | -0.54797 | -0.62005 | 0.14 |
| 13958 | *CCDC97* | 2.4796 | -0.98158 | -1.0904 | -1.7001 | 0.14 |
| 13959 | *DYNC1H1* | 1.2678 | 0.22314 | -1.0838 | -1.9725 | 0.14 |
| 13960 | *GPBP1L1* | 1.314 | 0.054282 | -0.96161 | -2.6837 | 0.14 |
| 13961 | *LARS* | 0.30293 | 0.058283 | 0.04435 | -0.29801 | 0.14 |
| 13962 | *TP53RK* | 0.84201 | 0.40361 | -0.84026 | -2.2948 | 0.14 |
| 13963 | *PTP4A3* | 0.95206 | 0.24853 | -0.79548 | -1.6848 | 0.14 |
| 13964 | *SNAPC4* | 2.3007 | -0.38355 | -1.5132 | -2.2653 | 0.13 |
| 13965 | *WEE1* | 2.5819 | -0.09527 | -2.0837 | -2.7068 | 0.13 |
| 13966 | *RANBP2* | 1.5781 | 0.27752 | -1.4527 | -1.7247 | 0.13 |
| 13967 | *MPZL1* | 1.4742 | 0.022664 | -1.0951 | -1.534 | 0.13 |
| 13968 | *LUZP1* | 0.38181 | 0.19646 | -0.17658 | -0.8345 | 0.13 |
| 13969 | *PPIE* | 0.5313 | 0.20182 | -0.33213 | -0.84075 | 0.13 |
| 13970 | *GTF2F2* | 1.4908 | -0.44353 | -0.64683 | -1.3926 | 0.13 |
| 13971 | *PHOSPHO1* | 1.2741 | 0.44327 | -1.3173 | -1.6935 | 0.13 |
| 13972 | *CLCN6* | 0.67455 | 0.043375 | -0.31818 | -0.62065 | 0.13 |
| 13973 | *USPL1* | 0.49422 | -0.04717 | -0.04799 | -1.9622 | 0.13 |
| 13974 | *RNASE3* | 1.2404 | -0.14639 | -0.69502 | -1.1629 | 0.13 |
| 13975 | *RAB3GAP1* | 1.7477 | -0.63152 | -0.71772 | -1.4696 | 0.13 |
| 13976 | *DOPEY2* | 0.66957 | 0.20677 | -0.47818 | -1.3245 | 0.13 |
| 13977 | *EXOC4* | 0.85095 | 0.19463 | -0.64851 | -2.312 | 0.13 |

| | | | | | | |
|---|---|---|---|---|---|---|
| 13978 | *SLC18A3* | 0.37811 | 0.22792 | -0.20933 | -1.7822 | 0.13 |
| 13979 | *RQCD1* | 2.5986 | -0.88584 | -1.3162 | -2.4197 | 0.13 |
| 13980 | *TXN* | 0.76604 | 0.29116 | -0.66164 | -1.1479 | 0.13 |
| 13981 | *SLC39A13* | 0.53689 | -0.05346 | -0.0881 | -2.1282 | 0.13 |
| 13982 | *GCG* | 0.91432 | 0.34848 | -0.86793 | -0.9653 | 0.13 |
| 13983 | *PERP* | 1.0046 | 0.53955 | -1.1511 | -1.7177 | 0.13 |
| 13984 | *IL22RA2* | 1.8036 | -0.18461 | -1.227 | -2.4386 | 0.13 |
| 13985 | *SPON2* | 0.95169 | 0.35042 | -0.91045 | -0.97103 | 0.13 |
| 13986 | *UBE2J1* | 0.451 | 0.098411 | -0.15783 | -1.5744 | 0.13 |
| 13987 | *TTC9B* | 1.2333 | 0.32175 | -1.1637 | -1.9054 | 0.13 |
| 13988 | *ARSA* | 0.46998 | 0.27047 | -0.34987 | -1.1858 | 0.13 |
| 13989 | *HLA-F* | 1.0288 | 0.1059 | -0.7443 | -1.3053 | 0.13 |
| 13990 | *KLF14* | 0.47249 | 0.39358 | -0.47671 | -0.84836 | 0.13 |
| 13991 | *RPH3A* | 0.6696 | 0.44688 | -0.72764 | -1.4533 | 0.13 |
| 13992 | *EXOG* | 2.1275 | -0.84446 | -0.89424 | -1.2639 | 0.13 |
| 13993 | *SEMA4D* | 1.7722 | -0.0172 | -1.3666 | -1.398 | 0.13 |
| 13994 | *SCRIB* | 0.90342 | 0.5013 | -1.0165 | -2.0305 | 0.13 |
| 13995 | *CALHM3* | 0.64447 | 0.21027 | -0.4674 | -2.1616 | 0.13 |
| 13996 | *DIAPH2* | 3.2587 | -1.1229 | -1.7493 | -3.1211 | 0.13 |
| 13997 | *ZNF664* | 0.98891 | 0.24364 | -0.84624 | -0.95224 | 0.13 |
| 13998 | *CPNE9* | 2.2377 | -0.8897 | -0.96196 | -1.5876 | 0.13 |
| 13999 | *APMAP* | 1.3594 | 0.21413 | -1.1897 | -1.796 | 0.13 |
| 14000 | *SNX24* | 2.4112 | -0.52902 | -1.4996 | -2.3755 | 0.13 |
| 14001 | *CAMKK2* | 1.1578 | 0.55802 | -1.3336 | -1.5485 | 0.13 |
| 14002 | *LIN9* | 1.8098 | -0.60659 | -0.82128 | -2.6003 | 0.13 |
| 14003 | *TSR3* | 3.0275 | -0.85236 | -1.7945 | -2.412 | 0.13 |
| 14004 | *FGF12* | 1.2302 | 0.48465 | -1.3347 | -1.4527 | 0.13 |
| 14005 | *ZNF615* | 1.0539 | 0.38184 | -1.0564 | -1.6447 | 0.13 |
| 14006 | *OR5R1* | 0.2658 | 0.080936 | 0.0321 | -0.89957 | 0.13 |
| 14007 | *DEDD* | 0.94458 | -0.27313 | -0.29291 | -1.1135 | 0.13 |
| 14008 | *CXCL2* | 0.66764 | 0.48128 | -0.77056 | -1.0333 | 0.13 |
| 14009 | *CCDC108* | 1.9712 | -0.56581 | -1.0271 | -1.3066 | 0.13 |
| 14010 | *TCF23* | 0.52459 | 0.20031 | -0.34688 | -0.45361 | 0.13 |
| 14011 | *TOMM7* | 0.93971 | 0.25991 | -0.82172 | -1.0837 | 0.13 |
| 14012 | *CDH16* | 0.8257 | -0.0061 | -0.44353 | -1.3325 | 0.13 |
| 14013 | *GCNT3* | 0.828 | 0.80452 | -1.2568 | -1.9013 | 0.13 |
| 14014 | *CGB* | 0.68689 | 0.028777 | -0.34077 | -1.1795 | 0.12 |
| 14015 | *ARID5A* | 0.56782 | 0.31649 | -0.51017 | -1.63 | 0.12 |
| 14016 | *BAG2* | 2.0178 | 0.23469 | -1.879 | -2.3615 | 0.12 |
| 14017 | *JAGN1* | 1.793 | 0.32243 | -1.7434 | -1.8345 | 0.12 |
| 14018 | *SCNM1* | 0.59621 | 0.31004 | -0.53455 | -0.87706 | 0.12 |
| 14019 | *CCDC81* | 4.1652 | -0.70442 | -3.0911 | -3.5736 | 0.12 |
| 14020 | *ABCC2* | 1.1143 | 0.52174 | -1.2676 | -2.1264 | 0.12 |
| 14021 | *RLTPR* | 3.4004 | -1.1132 | -1.9191 | -2.7568 | 0.12 |
| 14022 | *TRPM1* | 1.0767 | -0.29146 | -0.41748 | -2.1888 | 0.12 |
| 14023 | *UGT1A6* | 0.34533 | 0.32554 | -0.3032 | -0.73069 | 0.12 |

| | | | | | | |
|---|---|---|---|---|---|---|
| 14024 | *MRPL47* | 1.0084 | 0.36247 | -1.0057 | -1.334 | 0.12 |
| 14025 | *ZNF425* | 0.41263 | 0.20594 | -0.25375 | -0.94252 | 0.12 |
| 14026 | *RHOBTB1* | 0.42234 | 0.21512 | -0.27303 | -1.032 | 0.12 |
| 14027 | *PPHLN1* | 1.2703 | -0.06844 | -0.83791 | -0.93726 | 0.12 |
| 14028 | *FGF16* | 1.5623 | 0.012433 | -1.2109 | -3.1991 | 0.12 |
| 14029 | *PARN* | 0.24435 | 0.2126 | -0.09476 | -0.51514 | 0.12 |
| 14030 | *CCBE1* | 0.72697 | 0.078477 | -0.44374 | -1.2265 | 0.12 |
| 14031 | *SS18L1* | 0.83693 | 0.052712 | -0.52916 | -0.85536 | 0.12 |
| 14032 | *PAX9* | 0.33072 | 0.13367 | -0.10392 | -1.9885 | 0.12 |
| 14033 | *TIGIT* | 1.9873 | -0.78651 | -0.84042 | -0.8756 | 0.12 |
| 14034 | *ST5* | 1.7296 | -0.36917 | -1.0013 | -3.3677 | 0.12 |
| 14035 | *TOPAZ1* | 1.2194 | 0.97055 | -1.831 | -2.1046 | 0.12 |
| 14036 | *TRIM25* | 1.4636 | 0.65341 | -1.7591 | -1.7908 | 0.12 |
| 14037 | *KRAS* | 1.2671 | -0.13638 | -0.77397 | -0.89238 | 0.12 |
| 14038 | *ARFGAP1* | 1.3434 | -0.17459 | -0.81306 | -1.1972 | 0.12 |
| 14039 | *EXOSC8* | 2.5973 | -0.84693 | -1.3951 | -1.4168 | 0.12 |
| 14040 | *TPR* | 0.49772 | 0.28998 | -0.43292 | -0.69397 | 0.12 |
| 14041 | *CNTN6* | 2.2397 | -0.75593 | -1.1303 | -1.6988 | 0.12 |
| 14042 | *XKR4* | 2.3855 | -0.99217 | -1.0406 | -1.5765 | 0.12 |
| 14043 | *FBXL15* | 0.73723 | 0.09021 | -0.47485 | -0.97784 | 0.12 |
| 14044 | *HSPA12A* | 1.3313 | -0.15679 | -0.82281 | -1.8995 | 0.12 |
| 14045 | *SETD3* | 2.5441 | -0.54028 | -1.6525 | -1.8751 | 0.12 |
| 14046 | *AK6* | 0.71104 | 0.30181 | -0.66164 | -0.71041 | 0.12 |
| 14047 | *CCDC159* | 0.66961 | -0.15194 | -0.1672 | -0.39851 | 0.12 |
| 14048 | *C8orf37* | 0.8136 | -0.00831 | -0.45563 | -1.0293 | 0.12 |
| 14049 | *C15orf43* | 1.898 | -0.38676 | -1.1619 | -1.9305 | 0.12 |
| 14050 | *SLC7A10* | 2.9897 | -1.0926 | -1.548 | -1.6449 | 0.12 |
| 14051 | *FOXI2* | 0.53045 | 0.38056 | -0.56376 | -1.5502 | 0.12 |
| 14052 | *PRR22* | 1.39 | -0.30268 | -0.7404 | -0.78923 | 0.12 |
| 14053 | *TRAF3IP1* | 0.86784 | -0.07743 | -0.44353 | -0.6602 | 0.12 |
| 14054 | *HIST1H3I* | 0.91198 | 0.33366 | -0.89892 | -3.3546 | 0.12 |
| 14055 | *CSNK1A1* | 4.3726 | -1.6707 | -2.3557 | -2.712 | 0.12 |
| 14056 | *HSCB* | 0.99807 | -0.22569 | -0.42697 | -0.94341 | 0.12 |
| 14057 | *OR10Z1* | 0.8457 | 0.03717 | -0.53799 | -0.78363 | 0.11 |
| 14058 | *AMY1C* | 2.6048 | -0.53619 | -1.7241 | -2.2277 | 0.11 |
| 14059 | *HEATR1* | 0.97384 | -0.14419 | -0.48708 | -0.72666 | 0.11 |
| 14060 | *XCR1* | 1.9484 | -0.57042 | -1.0355 | -1.57 | 0.11 |
| 14061 | *ZFYVE21* | 0.46796 | 0.19132 | -0.31694 | -0.70546 | 0.11 |
| 14062 | *PRSS53* | 1.7251 | -0.48669 | -0.89724 | -1.1387 | 0.11 |
| 14063 | *ST6GALNAC6* | 0.3606 | 0.14118 | -0.16069 | -1.2739 | 0.11 |
| 14064 | *DEFB112* | 0.45661 | 0.42744 | -0.54331 | -1.2618 | 0.11 |
| 14065 | *PSMB2* | 1.7828 | -0.15649 | -1.2864 | -1.5956 | 0.11 |
| 14066 | *POLR3A* | 1.9005 | -0.66358 | -0.89741 | -1.9397 | 0.11 |
| 14067 | *C19orf68* | 0.21212 | 0.18894 | -0.06373 | -0.15822 | 0.11 |
| 14068 | *SCRT2* | 0.65565 | 0.52616 | -0.84471 | -1.4788 | 0.11 |
| 14069 | *MKNK1* | 0.8692 | 0.20034 | -0.73259 | -1.1383 | 0.11 |

| | | | | | | |
|---|---|---|---|---|---|---|
| 14070 | *GNG8* | 0.34469 | 0.13635 | -0.14449 | -1.1897 | 0.11 |
| 14071 | *P2RX3* | 0.71341 | -0.09448 | -0.28254 | -0.76033 | 0.11 |
| 14072 | *NDNL2* | 2.5708 | -1.0697 | -1.1653 | -2.1814 | 0.11 |
| 14073 | *ANXA8* | 1.1954 | -0.15041 | -0.71063 | -1.6487 | 0.11 |
| 14074 | *OR2T1* | 0.88185 | 0.62831 | -1.1759 | -2.0791 | 0.11 |
| 14075 | *SARM1* | 1.4561 | 0.3616 | -1.4836 | -1.8039 | 0.11 |
| 14076 | *TULP2* | 0.66957 | 0.10729 | -0.44353 | -2.3811 | 0.11 |
| 14077 | *PCDH11X* | 0.468 | 0.027154 | -0.16249 | -1.5194 | 0.11 |
| 14078 | *RPS27A* | 1.5283 | -0.33179 | -0.86403 | -1.3542 | 0.11 |
| 14079 | *RTN4RL1* | 1.5505 | -0.38103 | -0.83713 | -1.1135 | 0.11 |
| 14080 | *ZNF160* | 2.8032 | -0.81248 | -1.6587 | -1.7591 | 0.11 |
| 14081 | *EIF4G2* | 0.28798 | 0.21077 | -0.16836 | -1.8523 | 0.11 |
| 14082 | *MT1F* | 0.47575 | 0.31176 | -0.45786 | -0.77105 | 0.11 |
| 14083 | *ZPBP* | 2.3502 | -0.33799 | -1.6826 | -1.9844 | 0.11 |
| 14084 | *NEK9* | 1.2345 | 0.26472 | -1.1705 | -1.5461 | 0.11 |
| 14085 | *CECR6* | 1.6037 | -0.40359 | -0.8716 | -2.3552 | 0.11 |
| 14086 | *WNT10B* | 1.1959 | -0.00827 | -0.85943 | -0.87748 | 0.11 |
| 14087 | *ZNF417* | 0.37237 | 0.33953 | -0.38597 | -1.5289 | 0.11 |
| 14088 | *PGD* | 1.1744 | -0.32785 | -0.5223 | -3.1656 | 0.11 |
| 14089 | *SECISBP2* | 2.9986 | -0.50978 | -2.1648 | -2.3389 | 0.11 |
| 14090 | *TSSK4* | 1.9245 | -0.48002 | -1.1212 | -1.1385 | 0.11 |
| 14091 | *ZNF501* | 1.7146 | -0.65352 | -0.73823 | -1.0528 | 0.11 |
| 14092 | *PCDHGC4* | 2.8999 | -0.6603 | -1.9172 | -2.3204 | 0.11 |
| 14093 | *OTUB1* | 2.1864 | -0.64793 | -1.2167 | -1.6424 | 0.11 |
| 14094 | *TBCA* | 1.6518 | 0.30293 | -1.6339 | -2.5935 | 0.11 |
| 14095 | *SRRM2* | 2.4822 | -0.89235 | -1.2697 | -2.095 | 0.11 |
| 14096 | *KLHL1* | 1.1669 | -0.34158 | -0.50535 | -1.8157 | 0.11 |
| 14097 | *TDRD10* | 1.1165 | -0.0227 | -0.77397 | -2.0022 | 0.11 |
| 14098 | *OLFML3* | 0.25827 | 0.056342 | 0.00408 | -0.75832 | 0.11 |
| 14099 | *C8orf44-SGK3* | 1.3572 | -0.16435 | -0.87536 | -0.88503 | 0.11 |
| 14100 | *PUM2* | 0.40424 | 0.16515 | -0.25225 | -0.96583 | 0.11 |
| 14101 | *MTF1* | 1.5403 | 0.19477 | -1.4181 | -1.4741 | 0.11 |
| 14102 | *NUP153* | 3.2396 | -0.9481 | -1.9746 | -5.6002 | 0.11 |
| 14103 | *BRD8* | 1.4428 | 0.21771 | -1.3437 | -2.3235 | 0.11 |
| 14104 | *SV2A* | 0.81089 | 0.44296 | -0.93707 | -0.94583 | 0.11 |
| 14105 | *KCTD11* | 0.25601 | 0.08307 | -0.02242 | -2.2117 | 0.11 |
| 14106 | *C16orf45* | 0.87071 | 0.50708 | -1.0614 | -1.2716 | 0.11 |
| 14107 | *NTNG2* | 0.44101 | 0.055402 | -0.18082 | -1.7766 | 0.11 |
| 14108 | *C19orf60* | 1.5678 | 0.20713 | -1.4597 | -1.7936 | 0.11 |
| 14109 | *AMN1* | 0.32445 | 0.15185 | -0.16107 | -0.64002 | 0.11 |
| 14110 | *ACSM2B* | 0.36325 | 0.27104 | -0.31947 | -2.0851 | 0.10 |
| 14111 | *MOV10* | 0.7288 | 0.025762 | -0.44043 | -1.666 | 0.10 |
| 14112 | *DYNC2LI1* | 0.88189 | 0.34555 | -0.91399 | -0.96 | 0.10 |
| 14113 | *HAPLN2* | 1.3778 | -0.19778 | -0.86829 | -2.625 | 0.10 |
| 14114 | *FRAT1* | 1.4269 | 0.3523 | -1.4683 | -1.6158 | 0.10 |
| 14115 | *RHBDL3* | 2.3724 | -0.30811 | -1.7547 | -3.3367 | 0.10 |

| | | | | | | |
|---|---|---|---|---|---|---|
| 14116 | *ARHGAP26* | 1.5333 | 0.059027 | -1.2829 | -2.9841 | 0.10 |
| 14117 | *KDM8* | 0.6854 | 0.0951 | -0.47136 | -0.64866 | 0.10 |
| 14118 | *ACSM5* | 0.6696 | 0.47755 | -0.83808 | -1.8952 | 0.10 |
| 14119 | *ANKRD45* | 0.2753 | 0.066884 | -0.03356 | -0.30898 | 0.10 |
| 14120 | *TAF11* | 0.76866 | -0.09397 | -0.36649 | -2.0436 | 0.10 |
| 14121 | *NBPF4* | 1.4454 | 0.61576 | -1.7533 | -2.1367 | 0.10 |
| 14122 | *PGBD5* | 0.50836 | 0.46918 | -0.67044 | -1.4339 | 0.10 |
| 14123 | *CATSPERB* | 0.70494 | 0.12773 | -0.52587 | -1.3437 | 0.10 |
| 14124 | *CTDP1* | 2.7264 | -1.191 | -1.2288 | -1.7906 | 0.10 |
| 14125 | *ACAA2* | 1.5138 | -0.39248 | -0.81501 | -1.6339 | 0.10 |
| 14126 | *NUP188* | 1.3825 | -0.35617 | -0.72021 | -1.6131 | 0.10 |
| 14127 | *MAGEA6* | 2.4419 | -0.97768 | -1.1587 | -1.7686 | 0.10 |
| 14128 | *GMCL1* | 0.68177 | 0.5333 | -0.912 | -1.3393 | 0.10 |
| 14129 | *MLH1* | 0.30932 | 0.19879 | -0.20512 | -0.86298 | 0.10 |
| 14130 | *ANKRD10* | 1.6234 | -0.40142 | -0.91913 | -1.7822 | 0.10 |
| 14131 | *PEA15* | 1.2138 | 0.69775 | -1.6089 | -3.9492 | 0.10 |
| 14132 | *CCDC38* | 1.3265 | -0.48449 | -0.53938 | -0.56972 | 0.10 |
| 14133 | *LRFN4* | 0.24294 | 0.10691 | -0.04792 | -1.279 | 0.10 |
| 14134 | *TRPV6* | 0.66949 | 0.38094 | -0.74875 | -2.3534 | 0.10 |
| 14135 | *PILRB* | 0.51644 | -0.10176 | -0.11327 | -1.2277 | 0.10 |
| 14136 | *EP400* | 1.1133 | -0.301 | -0.51122 | -0.54197 | 0.10 |
| 14137 | *SPAG6* | 0.21342 | 0.1106 | -0.02314 | -1.4754 | 0.10 |
| 14138 | *PPP2R5A* | 1.0043 | 0.095684 | -0.80154 | -1.3982 | 0.10 |
| 14139 | *RAPH1* | 0.37109 | 0.18627 | -0.25893 | -2.201 | 0.10 |
| 14140 | *CES2* | 0.56104 | 0.13154 | -0.39494 | -0.71317 | 0.10 |
| 14141 | *MSN* | 1.0441 | 0.32513 | -1.072 | -1.6086 | 0.10 |
| 14142 | *PLEKHD1* | 2.1129 | -0.25564 | -1.5604 | -1.9326 | 0.10 |
| 14143 | *EFCAB12* | 0.82474 | 0.049283 | -0.57742 | -1.222 | 0.10 |
| 14144 | *ADA* | 1.1993 | -0.1123 | -0.79083 | -2.4784 | 0.10 |
| 14145 | *BSG* | 2.2564 | -0.75373 | -1.2071 | -1.3756 | 0.10 |
| 14146 | *RGS6* | 0.5736 | 0.20038 | -0.47929 | -0.95423 | 0.10 |
| 14147 | *CYLC2* | 0.41761 | 0.36664 | -0.48962 | -0.86861 | 0.10 |
| 14148 | *USP2* | 0.49468 | 0.24525 | -0.44541 | -0.83972 | 0.10 |
| 14149 | *SRP14* | 0.32696 | 0.18841 | -0.222 | -1.1733 | 0.10 |
| 14150 | *TAF7L* | 0.77927 | -0.06794 | -0.4191 | -0.98842 | 0.10 |
| 14151 | *MOG* | 1.0988 | -0.05504 | -0.75205 | -1.0096 | 0.10 |
| 14152 | *OTOP3* | 0.97235 | -0.2372 | -0.44353 | -1.6926 | 0.10 |
| 14153 | *NLRP13* | 0.53998 | 0.32373 | -0.57212 | -1.5894 | 0.10 |
| 14154 | *HYOU1* | 2.5148 | -1.0948 | -1.1288 | -2.0634 | 0.10 |
| 14155 | *RPL36AL* | 1.4586 | 0.17321 | -1.3407 | -1.4902 | 0.10 |
| 14156 | *RAB40AL* | 0.53078 | 0.027032 | -0.26677 | -0.73826 | 0.10 |
| 14157 | *FAM122C* | 0.66963 | 0.4315 | -0.8108 | -1.0046 | 0.10 |
| 14158 | *SSSCA1* | 0.91 | -0.28928 | -0.33116 | -1.7052 | 0.10 |
| 14159 | *VANGL1* | 0.15449 | 0.14314 | -0.00829 | -0.78229 | 0.10 |
| 14160 | *SBF2* | 0.47399 | 0.1179 | -0.30326 | -1.6648 | 0.10 |
| 14161 | *KRI1* | 2.7278 | -0.96555 | -1.4741 | -2.1732 | 0.10 |

| | | | | | | |
|---|---|---|---|---|---|---|
| 14162 | *FSTL4* | 0.65202 | 0.41325 | -0.77733 | -2.1519 | 0.10 |
| 14163 | *ZNF85* | 1.2115 | -0.14918 | -0.77667 | -1.6739 | 0.10 |
| 14164 | *HCFC1R1* | 2.5106 | -0.43332 | -1.7918 | -2.0553 | 0.10 |
| 14165 | *SH3BGRL* | 2.9733 | -1.0567 | -1.632 | -2.3776 | 0.09 |
| 14166 | *ADK* | 0.45793 | 0.25552 | -0.4295 | -1.115 | 0.09 |
| 14167 | *PSAT1* | 1.7455 | -0.59055 | -0.87109 | -1.1258 | 0.09 |
| 14168 | *TXNL4B* | 0.60448 | -0.04002 | -0.28062 | -1.4527 | 0.09 |
| 14169 | *C7orf33* | 0.35355 | 0.12633 | -0.19755 | -0.85199 | 0.09 |
| 14170 | *NHEJ1* | 1.9558 | -0.60869 | -1.066 | -1.6039 | 0.09 |
| 14171 | *BRD9* | 0.59299 | 0.32361 | -0.63575 | -2.8849 | 0.09 |
| 14172 | *IFIT3* | 2.9207 | -1.2135 | -1.4265 | -2.1938 | 0.09 |
| 14173 | *KIAA0232* | 1.964 | -0.38347 | -1.3 | -1.7153 | 0.09 |
| 14174 | *PRPF39* | 0.34161 | 0.28881 | -0.35009 | -0.54714 | 0.09 |
| 14175 | *TTC38* | 0.39053 | -0.00278 | -0.1082 | -0.68021 | 0.09 |
| 14176 | *SPACA4* | 0.64808 | -0.15143 | -0.2179 | -1.8598 | 0.09 |
| 14177 | *MATR3* | 2.0554 | -0.72391 | -1.0528 | -1.1126 | 0.09 |
| 14178 | *E2F1* | 0.40063 | 0.33888 | -0.46107 | -1.8557 | 0.09 |
| 14179 | *PLEKHO2* | 1.2176 | 0.54731 | -1.4867 | -1.6653 | 0.09 |
| 14180 | *BMP1* | 1.9558 | -0.78229 | -0.89752 | -1.1844 | 0.09 |
| 14181 | *TSPAN15* | 0.78114 | -0.15702 | -0.34816 | -1.377 | 0.09 |
| 14182 | *TAX1BP1* | 2.289 | -0.16634 | -1.8478 | -2.0301 | 0.09 |
| 14183 | *KCNA3* | 0.96511 | -0.14449 | -0.54657 | -0.64002 | 0.09 |
| 14184 | *ASPHD1* | 0.47215 | 0.038046 | -0.23688 | -1.8849 | 0.09 |
| 14185 | *CCNB1IP1* | 1.1068 | -0.40666 | -0.42775 | -0.89895 | 0.09 |
| 14186 | *USP14* | 0.85753 | -0.251 | -0.33613 | -0.75548 | 0.09 |
| 14187 | *DHRS11* | 0.66286 | -0.15247 | -0.24061 | -1.7394 | 0.09 |
| 14188 | *PLAU* | 1.1278 | 1.1164 | -1.9749 | -2.186 | 0.09 |
| 14189 | *DFNB31* | 0.29662 | 0.045563 | -0.07299 | -1.6597 | 0.09 |
| 14190 | *MYNN* | 0.14671 | 0.071115 | 0.05073 | -0.80861 | 0.09 |
| 14191 | *CCAR2* | 1.8336 | -0.11078 | -1.4544 | -1.5308 | 0.09 |
| 14192 | *MBNL2* | 0.78696 | -0.07541 | -0.44353 | -1.6921 | 0.09 |
| 14193 | *RUFY4* | 2.2907 | -0.82 | -1.2031 | -1.8031 | 0.09 |
| 14194 | *FSD1* | 1.1709 | -0.41281 | -0.49071 | -2.1264 | 0.09 |
| 14195 | *POLR2C* | 1.3403 | 0.40969 | -1.483 | -1.8402 | 0.09 |
| 14196 | *TMEM237* | 0.9807 | 0.62646 | -1.3409 | -1.3729 | 0.09 |
| 14197 | *JAZF1* | 1.4802 | -0.14244 | -1.072 | -1.4732 | 0.09 |
| 14198 | *FOXRED1* | 1.1557 | 0.15938 | -1.0496 | -1.7505 | 0.09 |
| 14199 | *UQCR11* | 1.1833 | -0.32704 | -0.59083 | -1.2884 | 0.09 |
| 14200 | *FRAS1* | 1.7624 | 0.46526 | -1.9623 | -2.9772 | 0.09 |
| 14201 | *SMTN* | 2.9081 | -1.2481 | -1.3955 | -1.8157 | 0.09 |
| 14202 | *OR4K13* | 1.0362 | 0.32334 | -1.0951 | -1.2864 | 0.09 |
| 14203 | *SLC27A6* | 0.66317 | 0.4856 | -0.88435 | -2.5208 | 0.09 |
| 14204 | *ZER1* | 1.309 | -0.29953 | -0.74572 | -0.82439 | 0.09 |
| 14205 | *NUP43* | 1.0898 | -0.23796 | -0.58822 | -1.1363 | 0.09 |
| 14206 | *CLUH* | 0.7821 | -0.00339 | -0.51547 | -1.1962 | 0.09 |
| 14207 | *RELT* | 0.29117 | 0.28154 | -0.31035 | -0.49443 | 0.09 |

| | | | | | | |
|---|---|---|---|---|---|---|
| 14208 | *OR2D3* | 1.073 | -0.33456 | -0.47632 | -1.1241 | 0.09 |
| 14209 | *SHPK* | 0.48979 | 0.47345 | -0.7015 | -2.2381 | 0.09 |
| 14210 | *C2orf66* | 0.60444 | -0.1497 | -0.19422 | -1.5165 | 0.09 |
| 14211 | *VPS45* | 1.118 | 0.89884 | -1.7566 | -1.9802 | 0.09 |
| 14212 | *GEMIN8* | 1.0471 | -0.23292 | -0.55505 | -1.5984 | 0.09 |
| 14213 | *SLC4A5* | 0.31193 | 0.097436 | -0.15098 | -0.82307 | 0.09 |
| 14214 | *C7orf57* | 0.8337 | 0.82035 | -1.3961 | -1.8724 | 0.09 |
| 14215 | *SHC3* | 0.26355 | 0.032616 | -0.04046 | -0.94927 | 0.09 |
| 14216 | *SPINK1* | 0.41698 | 0.073084 | -0.23476 | -0.91296 | 0.09 |
| 14217 | *WAC* | 0.48671 | 0.12704 | -0.35896 | -2.101 | 0.08 |
| 14218 | *TMEM161A* | 2.7047 | -1.1074 | -1.3427 | -1.3599 | 0.08 |
| 14219 | *NCAM2* | 1.1135 | -0.00182 | -0.85712 | -1.0765 | 0.08 |
| 14220 | *MVK* | 0.32659 | 0.17555 | -0.24797 | -0.358 | 0.08 |
| 14221 | *WBP2NL* | 1.1144 | 0.071506 | -0.93217 | -1.2619 | 0.08 |
| 14222 | *CDO1* | 2.1614 | -0.7734 | -1.1344 | -2.374 | 0.08 |
| 14223 | *CBX5* | 1.7238 | -0.42276 | -1.0483 | -1.4234 | 0.08 |
| 14224 | *POLR1B* | 1.1674 | -0.44353 | -0.47199 | -1.6956 | 0.08 |
| 14225 | *UBTFL1* | 1.5634 | 0.057397 | -1.3701 | -1.6129 | 0.08 |
| 14226 | *MDH1* | 1.0681 | 0.66648 | -1.4841 | -2.2879 | 0.08 |
| 14227 | *GSX2* | 0.56146 | -0.12667 | -0.18504 | -0.22043 | 0.08 |
| 14228 | *COL11A2* | 0.99442 | -0.14925 | -0.59572 | -1.1815 | 0.08 |
| 14229 | *MYRF* | 2.6666 | -1.0564 | -1.3613 | -1.3769 | 0.08 |
| 14230 | *A2M* | 0.41131 | 0.002435 | -0.16485 | -1.9116 | 0.08 |
| 14231 | *CEP72* | 2.824 | -0.57308 | -2.0028 | -2.1519 | 0.08 |
| 14232 | *METTL7B* | 0.61031 | 0.04489 | -0.4072 | -2.4517 | 0.08 |
| 14233 | *PRR3* | 0.70504 | 0.69371 | -1.1508 | -1.5894 | 0.08 |
| 14234 | *TCHP* | 0.62164 | 0.12149 | -0.49652 | -0.93465 | 0.08 |
| 14235 | *RASL10B* | 0.22314 | 0.02563 | -0.00309 | -0.85768 | 0.08 |
| 14236 | *EEF1G* | 2.1544 | -0.57408 | -1.335 | -1.6461 | 0.08 |
| 14237 | *C1orf228* | 1.1099 | 0.089784 | -0.95461 | -1.4087 | 0.08 |
| 14238 | *KIRREL2* | 0.67198 | 0.13947 | -0.56687 | -1.1422 | 0.08 |
| 14239 | *C2orf54* | 1.2484 | -0.10146 | -0.90281 | -1.059 | 0.08 |
| 14240 | *C7orf69* | 1.4761 | -0.13737 | -1.095 | -2.2809 | 0.08 |
| 14241 | *KRTAP9-3* | 1.0504 | -0.09082 | -0.71626 | -1.3409 | 0.08 |
| 14242 | *SPRR2B* | 2.4241 | -0.46741 | -1.7134 | -2.1151 | 0.08 |
| 14243 | *HUS1* | 3.1851 | -0.875 | -2.0669 | -2.239 | 0.08 |
| 14244 | *ZMPSTE24* | 0.63277 | -0.18353 | -0.20663 | -1.0158 | 0.08 |
| 14245 | *PRR20B* | 1.3986 | -0.26575 | -0.8903 | -0.91823 | 0.08 |
| 14246 | *NLRC3* | 2.2642 | -0.61798 | -1.4039 | -1.5817 | 0.08 |
| 14247 | *EXOSC6* | 1.3665 | -0.3555 | -0.76909 | -1.6245 | 0.08 |
| 14248 | *KCNH1* | 0.596 | 0.18761 | -0.54178 | -1.5082 | 0.08 |
| 14249 | *RNF11* | 1.1733 | -0.34113 | -0.59128 | -0.67004 | 0.08 |
| 14250 | *CSRP2* | 0.67698 | -0.15394 | -0.28357 | -1.2505 | 0.08 |
| 14251 | *PIAS3* | 0.70898 | -0.20242 | -0.26714 | -1.9157 | 0.08 |
| 14252 | *HIST1H4E* | 0.37869 | 0.19012 | -0.33205 | -2.2962 | 0.08 |
| 14253 | *LOC401052* | 0.67399 | 0.15724 | -0.59472 | -0.79035 | 0.08 |

| | | | | | | |
|---|---|---|---|---|---|---|
| 14254 | *ARMC6* | 2.0338 | -0.59128 | -1.2065 | -2.3357 | 0.08 |
| 14255 | *GAGE12C* | 2.0997 | -0.6424 | -1.222 | -2.1219 | 0.08 |
| 14256 | *SPRYD3* | 0.98173 | -0.28598 | -0.46153 | -2.0902 | 0.08 |
| 14257 | *RRN3* | 3.1321 | -1.1472 | -1.7512 | -2.2237 | 0.08 |
| 14258 | *ZNF10* | 0.67902 | 0.025582 | -0.47158 | -1.4036 | 0.08 |
| 14259 | *DCAF10* | 0.83089 | 0.80603 | -1.4043 | -3.1578 | 0.08 |
| 14260 | *SELO* | 2.1182 | -0.7635 | -1.1226 | -2.6201 | 0.08 |
| 14261 | *BCL2* | 0.83459 | 0.62919 | -1.2317 | -1.716 | 0.08 |
| 14262 | *SLC46A1* | 0.50751 | 0.46122 | -0.73695 | -1.6001 | 0.08 |
| 14263 | *SPDYE3* | 1.9391 | -0.62028 | -1.0875 | -1.8267 | 0.08 |
| 14264 | *RASEF* | 0.43778 | 0.18091 | -0.38779 | -0.50104 | 0.08 |
| 14265 | *PROCR* | 1.2302 | -0.42386 | -0.57551 | -1.8775 | 0.08 |
| 14266 | *AMY2A* | 0.749 | 0.25411 | -0.77241 | -2.6003 | 0.08 |
| 14267 | *DHRSX* | 1.3226 | -0.34653 | -0.74586 | -0.84801 | 0.08 |
| 14268 | *BTG2* | 1.1811 | -0.15543 | -0.79584 | -1.1846 | 0.08 |
| 14269 | *VIPR1* | 1.416 | 0.13369 | -1.3199 | -1.8398 | 0.08 |
| 14270 | *CCDC88C* | 1.5287 | -0.57825 | -0.72095 | -2.278 | 0.08 |
| 14271 | *HLTF* | 0.64346 | 0.25593 | -0.66992 | -0.70325 | 0.08 |
| 14272 | *RBM4* | 0.58931 | 0.17276 | -0.53321 | -2.1334 | 0.08 |
| 14273 | *RAB3GAP2* | 2.8898 | -0.9636 | -1.6981 | -2.3594 | 0.08 |
| 14274 | *HEPACAM2* | 0.85724 | -0.29292 | -0.33638 | -1.3393 | 0.08 |
| 14275 | *ZNF169* | 2.1397 | -0.87739 | -1.0344 | -1.8636 | 0.08 |
| 14276 | *DDX51* | 0.80457 | -0.20869 | -0.36821 | -1.01 | 0.08 |
| 14277 | *TMCO2* | 0.79104 | 0.36049 | -0.92486 | -4.3295 | 0.08 |
| 14278 | *OR51G1* | 1.3071 | 0.061507 | -1.1439 | -1.6044 | 0.07 |
| 14279 | *PITPNM2* | 0.51723 | 0.019879 | -0.31294 | -2.2879 | 0.07 |
| 14280 | *FAM151B* | 3.0831 | -1.1287 | -1.7303 | -2.3517 | 0.07 |
| 14281 | *TANGO6* | 1.6094 | -0.10425 | -1.2815 | -2.201 | 0.07 |
| 14282 | *CTSB* | 0.91288 | 0.17924 | -0.86884 | -2.0548 | 0.07 |
| 14283 | *ACTRT2* | 0.54593 | 0.056547 | -0.38112 | -0.81217 | 0.07 |
| 14284 | *CNOT10* | 1.4649 | -0.56444 | -0.6794 | -0.82789 | 0.07 |
| 14285 | *RBM38* | 0.36848 | 0.051389 | -0.20107 | -1.5219 | 0.07 |
| 14286 | *SDHAF2* | 1.5849 | -0.54283 | -0.82337 | -1.4538 | 0.07 |
| 14287 | *C7orf55-LUC7L2* | 0.97338 | -0.00268 | -0.75414 | -1.8702 | 0.07 |
| 14288 | *HLA-DQB2* | 1.8556 | -0.45443 | -1.1848 | -1.6199 | 0.07 |
| 14289 | *CACNG2* | 0.34685 | -0.00487 | -0.12595 | -0.88391 | 0.07 |
| 14290 | *TREH* | 1.4341 | -0.57899 | -0.64002 | -2.7678 | 0.07 |
| 14291 | *RNH1* | 2.1891 | -0.9512 | -1.0243 | -1.1303 | 0.07 |
| 14292 | *OR7E24* | 0.89213 | 0.6976 | -1.377 | -2.3065 | 0.07 |
| 14293 | *INO80B* | 0.35671 | 0.16473 | -0.30899 | -1.386 | 0.07 |
| 14294 | *HOXD11* | 0.66962 | 0.29586 | -0.75388 | -1.4031 | 0.07 |
| 14295 | *ZSWIM1* | 0.80313 | 0.46135 | -1.0529 | -1.7493 | 0.07 |
| 14296 | *EFTUD1* | 1.4424 | -0.21491 | -1.0164 | -3.2873 | 0.07 |
| 14297 | *TMEM167A* | 0.15229 | 0.11639 | -0.05784 | -0.80341 | 0.07 |
| 14298 | *SUN2* | 0.43657 | -0.03356 | -0.1926 | -1.1287 | 0.07 |
| 14299 | *SUMO1* | 1.7979 | 0.035843 | -1.6237 | -1.7577 | 0.07 |

| | | | | | | |
|---|---|---|---|---|---|---|
| 14300 | *PRAME* | 0.89844 | -0.34232 | -0.34632 | -0.77514 | 0.07 |
| 14301 | *MRPS2* | 0.54692 | 0.19351 | -0.53078 | -0.94052 | 0.07 |
| 14302 | *GSTM3* | 0.76226 | 0.066172 | -0.61889 | -1.2973 | 0.07 |
| 14303 | *APOF* | 2.7889 | -0.77733 | -1.8023 | -2.2338 | 0.07 |
| 14304 | *CHMP5* | 0.89076 | 0.49401 | -1.1761 | -1.9367 | 0.07 |
| 14305 | *PAIP2* | 0.31176 | 0.023908 | -0.12784 | -2.8478 | 0.07 |
| 14306 | *NME3* | 0.40836 | -0.08446 | -0.11621 | -1.0391 | 0.07 |
| 14307 | *WDR72* | 1.7385 | -0.21349 | -1.3184 | -1.4685 | 0.07 |
| 14308 | *SYT12* | 0.39423 | -0.08164 | -0.10687 | -0.67801 | 0.07 |
| 14309 | *LMNB2* | 1.2046 | 0.081171 | -1.0811 | -1.7683 | 0.07 |
| 14310 | *CREM* | 1.071 | 0.53673 | -1.4034 | -2.6889 | 0.07 |
| 14311 | *MAOA* | 2.092 | -0.14091 | -1.7495 | -2.1984 | 0.07 |
| 14312 | *SAMD9L* | 0.70647 | 0.26607 | -0.77105 | -1.6731 | 0.07 |
| 14313 | *SHISA5* | 0.50027 | -0.0836 | -0.21526 | -1.4717 | 0.07 |
| 14314 | *BRI3BP* | 0.66954 | -0.01739 | -0.45185 | -1.2633 | 0.07 |
| 14315 | *SPTBN1* | 1.649 | -0.06523 | -1.3837 | -2.4718 | 0.07 |
| 14316 | *TCFL5* | 0.64732 | -0.20554 | -0.24238 | -2.0952 | 0.07 |
| 14317 | *IFT140* | 0.25665 | 0.000534 | -0.06021 | -1.0656 | 0.07 |
| 14318 | *TOR1AIP2* | 2.5797 | -1.1585 | -1.2251 | -1.4954 | 0.07 |
| 14319 | *RPP38* | 2.7879 | -0.36734 | -2.2254 | -2.4328 | 0.07 |
| 14320 | *INPP5K* | 1.5154 | -0.20843 | -1.1121 | -1.4858 | 0.06 |
| 14321 | *REEP3* | 0.55037 | 0.26122 | -0.61675 | -1.8804 | 0.06 |
| 14322 | *ISX* | 0.46892 | -0.04015 | -0.23404 | -1.7947 | 0.06 |
| 14323 | *SEMG2* | 1.4648 | -0.17676 | -1.095 | -2.1182 | 0.06 |
| 14324 | *CAV1* | 0.56046 | 0.054397 | -0.4221 | -0.52552 | 0.06 |
| 14325 | *FSD1L* | 0.92294 | -0.16263 | -0.56764 | -1.2226 | 0.06 |
| 14326 | *CWF19L1* | 0.89887 | 0.33339 | -1.0407 | -1.1807 | 0.06 |
| 14327 | *TPTE* | 3.1965 | -1.1105 | -1.8946 | -1.9578 | 0.06 |
| 14328 | *TRAPPC2L* | 3.4479 | -1.4141 | -1.843 | -2.29 | 0.06 |
| 14329 | *NCF1* | 0.77365 | -0.13963 | -0.44353 | -2.8229 | 0.06 |
| 14330 | *ZNF441* | 1.0488 | -0.06585 | -0.79253 | -1.1757 | 0.06 |
| 14331 | *MROH2B* | 0.38839 | 0.18932 | -0.38843 | -0.45695 | 0.06 |
| 14332 | *LIX1* | 0.90273 | -0.26575 | -0.4497 | -1.4683 | 0.06 |
| 14333 | *CT45A7* | 0.72248 | -0.13638 | -0.39932 | -1.5579 | 0.06 |
| 14334 | *COPRS* | 1.0101 | -0.21794 | -0.6057 | -1.578 | 0.06 |
| 14335 | *COQ5* | 2.3205 | -0.36127 | -1.773 | -2.4328 | 0.06 |
| 14336 | *FAM72C* | 0.53464 | -0.16263 | -0.18589 | -1.4997 | 0.06 |
| 14337 | *AFAP1L1* | 0.70561 | -0.03196 | -0.48756 | -2.2433 | 0.06 |
| 14338 | *MPI* | 1.437 | 0.24633 | -1.4976 | -1.6469 | 0.06 |
| 14339 | *CCNJL* | 0.49655 | -0.1126 | -0.19869 | -1.701 | 0.06 |
| 14340 | *LHX2* | 1.1366 | -0.39527 | -0.55641 | -1.1395 | 0.06 |
| 14341 | *CASC5* | 0.80081 | 0.21184 | -0.82844 | -1.8651 | 0.06 |
| 14342 | *BCAS2* | 0.5394 | 0.21274 | -0.56812 | -2.0134 | 0.06 |
| 14343 | *KBTBD2* | 2.0949 | -0.40281 | -1.5082 | -1.9494 | 0.06 |
| 14344 | *ORAOV1* | 0.74443 | -0.15783 | -0.40353 | -2.3164 | 0.06 |
| 14345 | *FOXH1* | 2.483 | -0.77241 | -1.5282 | -1.9367 | 0.06 |

| | | | | | | |
|---|---|---|---|---|---|---|
| 14346 | *NUDT4* | 1.5482 | -0.54813 | -0.8181 | -1.2485 | 0.06 |
| 14347 | *IFNA2* | 0.455 | 0.037733 | -0.31157 | -1.8031 | 0.06 |
| 14348 | *CENPV* | 0.92051 | -0.3635 | -0.37591 | -2.1444 | 0.06 |
| 14349 | *TTC28* | 3.1711 | -1.4269 | -1.5632 | -3.1195 | 0.06 |
| 14350 | *RBM22* | 1.4806 | -0.38351 | -0.91696 | -1.4204 | 0.06 |
| 14351 | *CRNKL1* | 1.1162 | 0.058283 | -0.99463 | -1.2839 | 0.06 |
| 14352 | *TMA7* | 3.0113 | -0.65199 | -2.1798 | -2.6445 | 0.06 |
| 14353 | *KRT79* | 0.565 | -0.04127 | -0.34443 | -1.3275 | 0.06 |
| 14354 | *FITM2* | 1.5399 | -0.6558 | -0.70501 | -1.8946 | 0.06 |
| 14355 | *NUP35* | 1.3706 | -0.00331 | -1.1888 | -1.6004 | 0.06 |
| 14356 | *CCDC103* | 2.2041 | -0.90693 | -1.12 | -1.577 | 0.06 |
| 14357 | *CD38* | 0.30854 | 0.16852 | -0.30007 | -2.0181 | 0.06 |
| 14358 | *IAPP* | 0.77982 | -0.27055 | -0.33253 | -0.63357 | 0.06 |
| 14359 | *PAQR3* | 2.5777 | -0.61779 | -1.7845 | -2.2343 | 0.06 |
| 14360 | *ZNF713* | 0.38681 | 0.21463 | -0.42609 | -0.78229 | 0.06 |
| 14361 | *CCBL1* | 2.3063 | -0.38173 | -1.7493 | -2.3557 | 0.06 |
| 14362 | *NARS2* | 0.84228 | 0.33109 | -0.99834 | -1.9291 | 0.06 |
| 14363 | *TALDO1* | 1.2581 | -0.10178 | -0.98159 | -2.1947 | 0.06 |
| 14364 | *SYS1* | 2.1916 | -0.6568 | -1.3604 | -1.8376 | 0.06 |
| 14365 | *GGT1* | 0.41647 | 0.048132 | -0.29069 | -0.78439 | 0.06 |
| 14366 | *IL1B* | 0.69055 | 0.23385 | -0.7512 | -1.6525 | 0.06 |
| 14367 | *ACR* | 2.2765 | -0.44353 | -1.6599 | -1.9166 | 0.06 |
| 14368 | *MYH11* | 0.49992 | 0.11577 | -0.44353 | -3.0687 | 0.06 |
| 14369 | *SDPR* | 1.8043 | -0.46112 | -1.1718 | -1.4568 | 0.06 |
| 14370 | *MTMR12* | 0.94047 | -0.32621 | -0.44353 | -1.1929 | 0.06 |
| 14371 | *FAM47A* | 0.67885 | 0.05795 | -0.56634 | -1.2647 | 0.06 |
| 14372 | *ELSPBP1* | 1.0379 | -0.06513 | -0.80248 | -1.1921 | 0.06 |
| 14373 | *PARP1* | 1.3183 | -0.54797 | -0.60095 | -1.6125 | 0.06 |
| 14374 | *GANC* | 1.3195 | 0.34243 | -1.4926 | -2.1023 | 0.06 |
| 14375 | *MXD3* | 1.1677 | -0.24415 | -0.75437 | -1.222 | 0.06 |
| 14376 | *IRG1* | 0.9734 | -0.30359 | -0.50075 | -1.105 | 0.06 |
| 14377 | *DDX4* | 0.48856 | 0.041348 | -0.36086 | -1.236 | 0.06 |
| 14378 | *SAGE1* | 0.8868 | -0.09924 | -0.61873 | -2.6705 | 0.06 |
| 14379 | *NKX3-2* | 0.72429 | -0.2372 | -0.31865 | -0.99277 | 0.06 |
| 14380 | *KLRB1* | 1.0299 | -0.41207 | -0.4494 | -0.89235 | 0.06 |
| 14381 | *ST3GAL3* | 1.7484 | -0.64011 | -0.94019 | -1.1193 | 0.06 |
| 14382 | *TRAPPC1* | 0.5188 | -0.06364 | -0.28778 | -0.64571 | 0.06 |
| 14383 | *STRN* | 2.4856 | -0.78097 | -1.5375 | -2.1187 | 0.06 |
| 14384 | *PDSS1* | 0.45405 | 0.22464 | -0.5116 | -1.9133 | 0.06 |
| 14385 | *EIF3F* | 2.3049 | -0.52968 | -1.6086 | -1.8684 | 0.06 |
| 14386 | *ABCB9* | 0.66949 | 0.10642 | -0.61029 | -0.91192 | 0.06 |
| 14387 | *UBE3D* | 1.1709 | 0.033415 | -1.041 | -1.5782 | 0.05 |
| 14388 | *CPSF3L* | 1.6125 | 0.086199 | -1.5357 | -1.554 | 0.05 |
| 14389 | *PSMD9* | 0.39979 | 0.39826 | -0.63513 | -1.9383 | 0.05 |
| 14390 | *UTP15* | 0.5006 | -0.13753 | -0.2005 | -1.197 | 0.05 |
| 14391 | *AFF2* | 0.6082 | -0.20642 | -0.23931 | -0.71987 | 0.05 |

| | | | | | | |
|---|---|---|---|---|---|---|
| 14392 | *RNF121* | 1.328 | -0.23446 | -0.93112 | -1.4448 | 0.05 |
| 14393 | *H2AFY2* | 0.6802 | 0.3593 | -0.87772 | -2.4491 | 0.05 |
| 14394 | *SUMF2* | 0.40857 | 0.072713 | -0.31951 | -1.7462 | 0.05 |
| 14395 | *RPL13A* | 3.6635 | -1.7466 | -1.7554 | -1.9567 | 0.05 |
| 14396 | *SPNS3* | 0.64312 | 0.51131 | -0.99368 | -1.6867 | 0.05 |
| 14397 | *TRIM27* | 0.64739 | 0.1718 | -0.6602 | -1.3613 | 0.05 |
| 14398 | *EPC2* | 0.84592 | 0.59007 | -1.278 | -1.9047 | 0.05 |
| 14399 | *SMG9* | 1.6616 | 0.11559 | -1.6199 | -5.9418 | 0.05 |
| 14400 | *FTO* | 0.97648 | 0.088149 | -0.90757 | -1.7483 | 0.05 |
| 14401 | *CEP290* | 3.1682 | -0.95802 | -2.0536 | -2.1255 | 0.05 |
| 14402 | *KDM3B* | 0.83213 | -0.27491 | -0.40095 | -0.66708 | 0.05 |
| 14403 | *HAPLN1* | 1.4439 | 0.024941 | -1.3127 | -2.412 | 0.05 |
| 14404 | *FRZB* | 0.72619 | -0.26737 | -0.30335 | -2.672 | 0.05 |
| 14405 | *ING5* | 0.47829 | 0.47464 | -0.79798 | -2.5125 | 0.05 |
| 14406 | *C17orf62* | 1.6007 | -0.47818 | -0.9686 | -1.3343 | 0.05 |
| 14407 | *RASL11A* | 2.1287 | -0.25403 | -1.7209 | -1.9622 | 0.05 |
| 14408 | *FN3K* | 1.6179 | 0.28138 | -1.7457 | -1.8604 | 0.05 |
| 14409 | *CHI3L2* | 1.4635 | 0.062832 | -1.3728 | -1.452 | 0.05 |
| 14410 | *SGPP2* | 1.2888 | 0.96933 | -2.1046 | -2.2264 | 0.05 |
| 14411 | *RAVER2* | 2.7479 | -1.1508 | -1.4442 | -1.9068 | 0.05 |
| 14412 | *SKP1* | 3.2621 | -1.2505 | -1.8591 | -3.1939 | 0.05 |
| 14413 | *CLDN2* | 1.261 | -0.44742 | -0.66119 | -0.98585 | 0.05 |
| 14414 | *GRIK3* | 0.61339 | -0.16435 | -0.2969 | -2.2094 | 0.05 |
| 14415 | *GMPR2* | 2.0684 | -0.50127 | -1.4151 | -1.9086 | 0.05 |
| 14416 | *DCAF13* | 0.61672 | 0.22036 | -0.68565 | -1.9213 | 0.05 |
| 14417 | *WASH1* | 0.9177 | -0.02368 | -0.7436 | -1.4841 | 0.05 |
| 14418 | *PGAM2* | 0.79175 | 0.29113 | -0.93392 | -3.2912 | 0.05 |
| 14419 | *MRPS12* | 1.0835 | -0.45339 | -0.48119 | -1.6601 | 0.05 |
| 14420 | *GNAT1* | 3.4559 | -1.6049 | -1.7024 | -2.9054 | 0.05 |
| 14421 | *TMEM208* | 0.86356 | 0.2126 | -0.9285 | -1.1761 | 0.05 |
| 14422 | *ST20-MTHFS* | 1.9448 | -0.27811 | -1.5213 | -1.8653 | 0.05 |
| 14423 | *PLCH2* | 0.88809 | -0.10484 | -0.63826 | -1.0386 | 0.05 |
| 14424 | *UNC119* | 1.0522 | 0.0973 | -1.0048 | -1.1444 | 0.05 |
| 14425 | *IER3IP1* | 0.29953 | -0.02098 | -0.13389 | -0.14506 | 0.05 |
| 14426 | *PAIP1* | 3.0782 | -1.448 | -1.4873 | -2.9897 | 0.05 |
| 14427 | *ZBTB20* | 0.31366 | 0.16806 | -0.33933 | -2.0626 | 0.05 |
| 14428 | *GNG12* | 0.97316 | -0.01606 | -0.81496 | -1.3132 | 0.05 |
| 14429 | *EFHC1* | 2.0395 | -0.32699 | -1.571 | -1.7177 | 0.05 |
| 14430 | *TMEM150A* | 0.98137 | -0.05801 | -0.78229 | -1.849 | 0.05 |
| 14431 | *PRKAR2A* | 1.9161 | -0.34998 | -1.4252 | -2.7158 | 0.05 |
| 14432 | *CNPY4* | 1.135 | -0.20976 | -0.7853 | -1.5965 | 0.05 |
| 14433 | *UFD1L* | 1.7886 | -0.56825 | -1.0809 | -1.4829 | 0.05 |
| 14434 | *RGCC* | 0.4766 | 0.078218 | -0.41596 | -0.51221 | 0.05 |
| 14435 | *SH2D6* | 1.8187 | -0.28197 | -1.3987 | -2.2352 | 0.05 |
| 14436 | *ELF5* | 1.6149 | 0.005207 | -1.4826 | -2.6593 | 0.05 |
| 14437 | *POP7* | 1.83 | -0.16634 | -1.5268 | -2.0055 | 0.05 |

| | | | | | | |
|---|---|---|---|---|---|---|
| 14438 | *H2AFX* | 1.2058 | -0.14906 | -0.91996 | -1.9938 | 0.05 |
| 14439 | *COLCA1* | 1.3881 | -0.21666 | -1.035 | -3.0444 | 0.05 |
| 14440 | *XPA* | 1.1968 | -0.36861 | -0.69287 | -1.1659 | 0.05 |
| 14441 | *DLX2* | 1.4015 | -0.62792 | -0.63826 | -1.6456 | 0.05 |
| 14442 | *RWDD2A* | 1.2658 | 0.26025 | -1.391 | -1.4964 | 0.05 |
| 14443 | *XPO1* | 0.88615 | -0.30777 | -0.44353 | -1.9538 | 0.04 |
| 14444 | *OR11G2* | 1.2414 | -0.42841 | -0.67889 | -1.8698 | 0.04 |
| 14445 | *RNF141* | 0.35855 | -0.09907 | -0.12563 | -0.97876 | 0.04 |
| 14446 | *UBE2Z* | 1.1419 | 0.61775 | -1.626 | -2.8181 | 0.04 |
| 14447 | *YAF2* | 0.70768 | 0.36965 | -0.94406 | -0.97303 | 0.04 |
| 14448 | *NPFF* | 0.53719 | 0.17837 | -0.58289 | -1.0782 | 0.04 |
| 14449 | *CXCR2* | 0.47831 | -0.1194 | -0.2293 | -0.70475 | 0.04 |
| 14450 | *GTF3C5* | 1.052 | 0.34656 | -1.2691 | -1.4488 | 0.04 |
| 14451 | *POM121* | 0.61765 | -0.01725 | -0.47244 | -1.0268 | 0.04 |
| 14452 | *MRTO4* | 0.55465 | -0.03288 | -0.39494 | -0.67007 | 0.04 |
| 14453 | *NGDN* | 0.87226 | 0.13956 | -0.88503 | -1.545 | 0.04 |
| 14454 | *MFRP* | 3.0671 | -1.2884 | -1.6525 | -2.337 | 0.04 |
| 14455 | *PLG* | 1.2503 | -0.33112 | -0.79319 | -1.0167 | 0.04 |
| 14456 | *TOMM20L* | 0.90094 | -0.03784 | -0.73739 | -0.79932 | 0.04 |
| 14457 | *PPM1K* | 1.1393 | 0.058918 | -1.0727 | -2.1507 | 0.04 |
| 14458 | *SENP2* | 1.5634 | 0.15743 | -1.5979 | -1.808 | 0.04 |
| 14459 | *CREB3L1* | 2.5593 | -0.86459 | -1.5722 | -1.7179 | 0.04 |
| 14460 | *LGALS14* | 0.27963 | 0.097436 | -0.25478 | -1.9365 | 0.04 |
| 14461 | *AUH* | 0.44002 | -0.15085 | -0.1677 | -0.76955 | 0.04 |
| 14462 | *ASF1A* | 1.4851 | -0.43906 | -0.92553 | -1.0344 | 0.04 |
| 14463 | *TRIM39-RPP21* | 0.55152 | 0.15938 | -0.59045 | -1.4841 | 0.04 |
| 14464 | *RPL10A* | 3.3876 | -1.4683 | -1.7994 | -2.2955 | 0.04 |
| 14465 | *DCAF15* | 0.66487 | -0.12758 | -0.41748 | -1.2235 | 0.04 |
| 14466 | *ZMIZ1* | 0.53532 | -0.05505 | -0.36112 | -1.3771 | 0.04 |
| 14467 | *TCEB1* | 0.98208 | 0.55442 | -1.4182 | -2.9068 | 0.04 |
| 14468 | *LTBR* | 0.152 | 0.016357 | -0.05165 | -0.13189 | 0.04 |
| 14469 | *ZNF836* | 1.2303 | 0.25142 | -1.3665 | -1.8334 | 0.04 |
| 14470 | *CARTPT* | 3.1556 | -1.1953 | -1.8456 | -2.1596 | 0.04 |
| 14471 | *CHD2* | 0.85843 | 0.69182 | -1.4368 | -1.6268 | 0.04 |
| 14472 | *CYB561D1* | 0.45121 | 0.14609 | -0.48419 | -0.63187 | 0.04 |
| 14473 | *PHF20* | 0.68584 | 0.14108 | -0.71465 | -1.5748 | 0.04 |
| 14474 | *ARID1A* | 0.37059 | 0.31999 | -0.57899 | -1.8406 | 0.04 |
| 14475 | *BACH1* | 0.40749 | 0.35512 | -0.65201 | -2.9068 | 0.04 |
| 14476 | *CACNA2D4* | 0.53876 | -0.1589 | -0.26949 | -2.4174 | 0.04 |
| 14477 | *RNF138* | 0.84448 | 0.28658 | -1.0212 | -1.7462 | 0.04 |
| 14478 | *DNAAF5* | 0.47055 | -0.09671 | -0.26409 | -1.4537 | 0.04 |
| 14479 | *USP6NL* | 0.28831 | -0.0429 | -0.13576 | -1.6437 | 0.04 |
| 14480 | *IFNA1* | 1.4921 | -0.5101 | -0.87256 | -1.5634 | 0.04 |
| 14481 | *SIRT3* | 0.53797 | 0.42852 | -0.85739 | -1.1932 | 0.04 |
| 14482 | *RAB37* | 0.29212 | 0.25944 | -0.44353 | -1.8689 | 0.04 |
| 14483 | *PNKD* | 0.68707 | 0.020456 | -0.60085 | -2.1342 | 0.04 |

| | | | | | | |
|---|---|---|---|---|---|---|
| 14484 | *METTL21C* | 1.4993 | -0.38351 | -1.0095 | -2.0778 | 0.04 |
| 14485 | *MEP1A* | 0.44364 | -0.0067 | -0.33107 | -0.50201 | 0.04 |
| 14486 | *SLIT3* | 0.35557 | 0.24544 | -0.49525 | -1.1907 | 0.04 |
| 14487 | *GPR31* | 1.5017 | -0.12048 | -1.2755 | -2.3601 | 0.04 |
| 14488 | *XPR1* | 0.5365 | -0.00112 | -0.43027 | -1.1665 | 0.04 |
| 14489 | *NOP58* | 0.792 | 0.056445 | -0.74392 | -0.82311 | 0.03 |
| 14490 | *NDUFS2* | 1.1627 | -0.49926 | -0.55898 | -0.63368 | 0.03 |
| 14491 | *NPAS4* | 1.2656 | 0.13896 | -1.3007 | -1.6597 | 0.03 |
| 14492 | *HHLA2* | 0.85028 | 0.32173 | -1.0691 | -3.7236 | 0.03 |
| 14493 | *HIST1H2AH* | 2.0925 | -0.80936 | -1.1812 | -1.5164 | 0.03 |
| 14494 | *TEAD1* | 0.51092 | -0.05186 | -0.35733 | -0.57579 | 0.03 |
| 14495 | *MYOG* | 0.36761 | 0.19477 | -0.46153 | -0.73896 | 0.03 |
| 14496 | *CCDC181* | 1.3489 | 0.16383 | -1.4123 | -1.7591 | 0.03 |
| 14497 | *PDCD2* | 3.1536 | -1.0838 | -1.9695 | -2.1956 | 0.03 |
| 14498 | *KIF26B* | 2.8595 | -1.082 | -1.6776 | -1.8537 | 0.03 |
| 14499 | *SPAG9* | 0.1466 | 0.034812 | -0.08163 | -1.2135 | 0.03 |
| 14500 | *COX7A2L* | 2.2335 | -0.70188 | -1.4319 | -1.5204 | 0.03 |
| 14501 | *PNLIPRP1* | 0.51887 | 0.11411 | -0.53332 | -0.98719 | 0.03 |
| 14502 | *SARS* | 0.6627 | 0.20292 | -0.76748 | -2.7427 | 0.03 |
| 14503 | *LIM2* | 1.3737 | -0.09654 | -1.1792 | -1.3582 | 0.03 |
| 14504 | *DDB2* | 0.23703 | 0.20268 | -0.34228 | -0.79149 | 0.03 |
| 14505 | *TMEM240* | 1.1805 | 0.30663 | -1.3899 | -1.6957 | 0.03 |
| 14506 | *MTRNR2L7* | 0.098148 | 0.073963 | -0.07512 | -1.8095 | 0.03 |
| 14507 | *CACNB1* | 0.67388 | -0.12795 | -0.4497 | -0.76433 | 0.03 |
| 14508 | *NOS2* | 0.66961 | 0.54502 | -1.1184 | -2.2434 | 0.03 |
| 14509 | *EXD1* | 0.73245 | 0.57498 | -1.2118 | -1.3389 | 0.03 |
| 14510 | *CCDC107* | 1.1165 | 0.092774 | -1.1138 | -1.8443 | 0.03 |
| 14511 | *FRMD3* | 2.2005 | 0.009147 | -2.1151 | -2.8181 | 0.03 |
| 14512 | *MFAP5* | 0.38147 | -0.04543 | -0.24157 | -0.38219 | 0.03 |
| 14513 | *PDE4DIP* | 0.97566 | 0.35805 | -1.2399 | -2.7566 | 0.03 |
| 14514 | *CNTN3* | 0.76104 | -0.03127 | -0.63614 | -1.4154 | 0.03 |
| 14515 | *PHKA1* | 0.21734 | 0.12515 | -0.24904 | -0.94609 | 0.03 |
| 14516 | *UGT1A9* | 0.95587 | -0.37215 | -0.49045 | -0.62372 | 0.03 |
| 14517 | *FBXL3* | 1.3084 | -0.22008 | -0.99508 | -1.0005 | 0.03 |
| 14518 | *CLVS1* | 0.71737 | -0.22324 | -0.40115 | -2.1055 | 0.03 |
| 14519 | *PPP1R36* | 0.16074 | 0.001415 | -0.06966 | -0.44353 | 0.03 |
| 14520 | *LRP8* | 0.57023 | -0.00023 | -0.47781 | -1.4568 | 0.03 |
| 14521 | *ATF7IP2* | 1.7826 | -0.00268 | -1.6881 | -1.9515 | 0.03 |
| 14522 | *SURF1* | 2.5243 | -1.1754 | -1.2572 | -1.4299 | 0.03 |
| 14523 | *MBOAT2* | 3.0748 | -1.2371 | -1.7466 | -2.3689 | 0.03 |
| 14524 | *MALSU1* | 2.3135 | 0.18932 | -2.412 | -3.2582 | 0.03 |
| 14525 | *SAR1A* | 0.50043 | 0.007935 | -0.4181 | -1.9186 | 0.03 |
| 14526 | *EIF3I* | 0.65863 | -0.01558 | -0.55335 | -1.2715 | 0.03 |
| 14527 | *TIRAP* | 1.305 | -0.48708 | -0.72839 | -1.0764 | 0.03 |
| 14528 | *EGR1* | 0.42468 | -0.11106 | -0.22445 | -0.24864 | 0.03 |
| 14529 | *CMYA5* | 1.213 | -0.54887 | -0.57513 | -1.5904 | 0.03 |

| | | | | | | |
|---|---|---|---|---|---|---|
| 14530 | *TOP1MT* | 0.58055 | 0.50314 | -0.99474 | -1.0134 | 0.03 |
| 14531 | *NT5C1B-RDH14* | 1.2088 | -0.34903 | -0.77114 | -1.4544 | 0.03 |
| 14532 | *CHKA* | 0.4504 | -0.09295 | -0.27016 | -1.6759 | 0.03 |
| 14533 | *RASSF10* | 1.1531 | -0.36258 | -0.70394 | -2.6753 | 0.03 |
| 14534 | *CHPT1* | 1.2523 | -0.461 | -0.70601 | -1.5482 | 0.03 |
| 14535 | *ACTL9* | 1.9356 | -0.78503 | -1.0667 | -1.9459 | 0.03 |
| 14536 | *RASGRP4* | 2.2435 | -0.79981 | -1.3599 | -1.6838 | 0.03 |
| 14537 | *CWF19L2* | 2.4707 | -0.9728 | -1.4142 | -1.714 | 0.03 |
| 14538 | *LOC100996634* | 1.0848 | -0.27961 | -0.72157 | -1.2154 | 0.03 |
| 14539 | *PRAMEF11* | 0.52757 | -0.2168 | -0.22783 | -1.0131 | 0.03 |
| 14540 | *C10orf107* | 0.31702 | 0.16813 | -0.4027 | -0.52269 | 0.03 |
| 14541 | *SPTSSB* | 0.36325 | -0.1284 | -0.15303 | -1.6023 | 0.03 |
| 14542 | *CLUAP1* | 0.5519 | -0.09723 | -0.37424 | -2.1983 | 0.03 |
| 14543 | *ZNF680* | 0.97366 | -0.16623 | -0.7288 | -3.4924 | 0.03 |
| 14544 | *IGF2BP3* | 1.0347 | 0.11462 | -1.071 | -1.6251 | 0.03 |
| 14545 | *NME9* | 0.71578 | -0.14183 | -0.49636 | -0.83095 | 0.03 |
| 14546 | *LRIG3* | 0.5977 | -0.17418 | -0.34597 | -0.74743 | 0.03 |
| 14547 | *MRPS25* | 1.495 | -0.65537 | -0.76212 | -1.9055 | 0.03 |
| 14548 | *TRPC4* | 3.5606 | -1.522 | -1.9622 | -2.0242 | 0.03 |
| 14549 | *DDX42* | 0.70907 | -0.04689 | -0.58661 | -2.103 | 0.03 |
| 14550 | *NPM1* | 2.5527 | -1.2297 | -1.2485 | -1.9794 | 0.02 |
| 14551 | *TIMM23* | 1.6635 | -0.68193 | -0.90713 | -2.1873 | 0.02 |
| 14552 | *POPDC3* | 1.0325 | 0.21634 | -1.1747 | -2.5442 | 0.02 |
| 14553 | *NEURL1B* | 0.24919 | 0.11223 | -0.28796 | -2.2108 | 0.02 |
| 14554 | *LOC81691* | 0.23111 | 0.22142 | -0.37982 | -1.1071 | 0.02 |
| 14555 | *PEX11B* | 0.60965 | -0.09403 | -0.44353 | -0.73922 | 0.02 |
| 14556 | *METTL2B* | 1.1027 | -0.13483 | -0.89592 | -1.8087 | 0.02 |
| 14557 | *ST3GAL2* | 0.74731 | 0.52934 | -1.205 | -1.448 | 0.02 |
| 14558 | *GABBR1* | 0.60031 | 0.54 | -1.0691 | -2.8928 | 0.02 |
| 14559 | *HPN* | 0.29893 | 0.027839 | -0.25583 | -0.32414 | 0.02 |
| 14560 | *TRAT1* | 1.4198 | -0.45119 | -0.89784 | -1.6802 | 0.02 |
| 14561 | *RAB34* | 1.3569 | -0.35196 | -0.93465 | -1.5726 | 0.02 |
| 14562 | *ARID2* | 0.48253 | 0.004561 | -0.41716 | -0.94171 | 0.02 |
| 14563 | *ASCC3* | 0.19992 | -0.03623 | -0.09523 | -1.4276 | 0.02 |
| 14564 | *CYP2C18* | 2.1402 | -0.65368 | -1.4195 | -1.6525 | 0.02 |
| 14565 | *EMC6* | 0.31244 | 0.19871 | -0.44433 | -1.3017 | 0.02 |
| 14566 | *NRBF2* | 0.51261 | 0.25105 | -0.69727 | -1.5263 | 0.02 |
| 14567 | *SPRR1B* | 1.2584 | 0.43722 | -1.63 | -2.2434 | 0.02 |
| 14568 | *MTIF2* | 1.6795 | -0.57301 | -1.0413 | -1.7428 | 0.02 |
| 14569 | *CCDC110* | 0.97486 | -0.22131 | -0.6886 | -1.3134 | 0.02 |
| 14570 | *LYRM4* | 2.1946 | -1.0564 | -1.0737 | -1.1281 | 0.02 |
| 14571 | *ACTN3* | 0.92645 | -0.0729 | -0.78951 | -3.164 | 0.02 |
| 14572 | *TSEN34* | 1.5091 | -0.45663 | -0.98925 | -1.338 | 0.02 |
| 14573 | *OR2T27* | 0.3454 | 0.26472 | -0.54724 | -0.78398 | 0.02 |
| 14574 | *NXNL2* | 0.87719 | 0.5456 | -1.3602 | -1.701 | 0.02 |
| 14575 | *TBC1D21* | 1.8576 | -0.56687 | -1.2284 | -1.7735 | 0.02 |

| | | | | | | |
|---|---|---|---|---|---|---|
| 14576 | *CXorf57* | 0.41697 | -0.04277 | -0.31191 | -1.0918 | 0.02 |
| 14577 | *FBXO39* | 0.45363 | 0.43735 | -0.82879 | -1.2534 | 0.02 |
| 14578 | *FNDC1* | 1.0167 | -0.10258 | -0.85194 | -1.8975 | 0.02 |
| 14579 | *WDR86* | 1.8144 | -0.44353 | -1.3098 | -1.4141 | 0.02 |
| 14580 | *RASL10A* | 0.83566 | 0.068993 | -0.84375 | -0.88008 | 0.02 |
| 14581 | *C1QTNF7* | 1.4673 | -0.22235 | -1.1845 | -1.7647 | 0.02 |
| 14582 | *LRRTM4* | 0.39759 | -0.15276 | -0.18445 | -0.26231 | 0.02 |
| 14583 | *C1orf106* | 1.7563 | -0.80847 | -0.88797 | -1.6255 | 0.02 |
| 14584 | *SEPW1* | 0.52604 | 0.16057 | -0.62762 | -0.90161 | 0.02 |
| 14585 | *TLCD2* | 0.19205 | 0.054921 | -0.18808 | -0.99694 | 0.02 |
| 14586 | *FECH* | 1.929 | -0.58509 | -1.2852 | -1.7496 | 0.02 |
| 14587 | *SLC16A4* | 1.1634 | -0.15855 | -0.94625 | -1.399 | 0.02 |
| 14588 | *PANK2* | 0.99303 | -0.21743 | -0.71717 | -1.195 | 0.02 |
| 14589 | *NPEPL1* | 0.62681 | 0.37869 | -0.94739 | -1.6691 | 0.02 |
| 14590 | *PALM2* | 0.93999 | -0.26252 | -0.62114 | -1.6208 | 0.02 |
| 14591 | *TTR* | 2.1921 | -0.84457 | -1.2912 | -1.5874 | 0.02 |
| 14592 | *CST2* | 2.6424 | -1.183 | -1.4034 | -3.1229 | 0.02 |
| 14593 | *SLC17A1* | 0.78904 | 0.19924 | -0.93307 | -1.632 | 0.02 |
| 14594 | *DNTTIP1* | 1.055 | -0.35211 | -0.64788 | -1.7049 | 0.02 |
| 14595 | *PPAPDC2* | 0.11005 | -0.02615 | -0.02908 | -0.78337 | 0.02 |
| 14596 | *CASKIN2* | 0.45582 | 0.33443 | -0.73599 | -1.0384 | 0.02 |
| 14597 | *LAMTOR1* | 2.8229 | -1.2898 | -1.4789 | -1.5182 | 0.02 |
| 14598 | *GOT2* | 1.6683 | -0.59212 | -1.0221 | -1.4962 | 0.02 |
| 14599 | *NKTR* | 0.29836 | 0.018783 | -0.26369 | -0.28907 | 0.02 |
| 14600 | *MCHR2* | 1.2044 | -0.38361 | -0.76755 | -1.3218 | 0.02 |
| 14601 | *VSNL1* | 1.2057 | -0.31624 | -0.83631 | -1.3353 | 0.02 |
| 14602 | *GLB1L* | 0.8351 | 0.17417 | -0.95717 | -1.2396 | 0.02 |
| 14603 | *SIGLEC14* | 1.8187 | -0.59195 | -1.1747 | -1.5744 | 0.02 |
| 14604 | *UBE4B* | 0.7252 | 0.3877 | -1.0614 | -2.0363 | 0.02 |
| 14605 | *GBX1* | 1.9746 | -0.55371 | -1.3694 | -1.6676 | 0.02 |
| 14606 | *EXOC6* | 2.5698 | -0.87768 | -1.6411 | -2.0337 | 0.02 |
| 14607 | *SPANXD* | 0.67353 | 0.46804 | -1.0906 | -2.2607 | 0.02 |
| 14608 | *SMIM11* | 0.35927 | 0.088577 | -0.39856 | -1.0564 | 0.02 |
| 14609 | *GTF2H5* | 0.96644 | 0.30746 | -1.2252 | -2.7777 | 0.02 |
| 14610 | *NTAN1* | 0.44466 | 0.027947 | -0.42397 | -2.5442 | 0.02 |
| 14611 | *KCNB1* | 0.47204 | -0.20318 | -0.22133 | -1.0399 | 0.02 |
| 14612 | *CLDN15* | 3.3585 | -1.4597 | -1.8523 | -3.164 | 0.02 |
| 14613 | *TMEM236* | 2.9312 | -1.2572 | -1.6276 | -2.1143 | 0.02 |
| 14614 | *BCL10* | 0.88577 | -0.31547 | -0.52426 | -1.4234 | 0.02 |
| 14615 | *ZNF532* | 1.0701 | -0.42779 | -0.59643 | -1.0516 | 0.02 |
| 14616 | *RIN2* | 1.5481 | -0.30065 | -1.2023 | -1.4919 | 0.02 |
| 14617 | *BAG4* | 0.56844 | 0.44441 | -0.96857 | -3.4314 | 0.01 |
| 14618 | *C17orf49* | 0.79382 | 0.037116 | -0.78705 | -1.7493 | 0.01 |
| 14619 | *HOGA1* | 0.13794 | 0.03977 | -0.1339 | -0.13627 | 0.01 |
| 14620 | *NOL4* | 0.59573 | 0.13771 | -0.68977 | -1.3835 | 0.01 |
| 14621 | *PHACTR1* | 1.2382 | -0.43944 | -0.75511 | -1.5684 | 0.01 |

| | | | | | | |
|---|---|---|---|---|---|---|
| 14622 | *RUNDC1* | 0.68062 | 0.33652 | -0.97509 | -1.09 | 0.01 |
| 14623 | *C17orf85* | 0.24729 | 0.10075 | -0.30608 | -1.6437 | 0.01 |
| 14624 | *KRT4* | 0.38981 | 0.095647 | -0.44353 | -1.8033 | 0.01 |
| 14625 | *NEMF* | 0.22155 | 0.033945 | -0.21385 | -0.77748 | 0.01 |
| 14626 | *NAT10* | 1.4872 | -0.54972 | -0.89591 | -2.0831 | 0.01 |
| 14627 | *BRINP2* | 0.49944 | 0.33766 | -0.79697 | -1.5635 | 0.01 |
| 14628 | *DHRS3* | 0.96837 | 0.22275 | -1.151 | -1.6796 | 0.01 |
| 14629 | *HYDIN* | 0.40498 | 0.10266 | -0.46762 | -1.8654 | 0.01 |
| 14630 | *UBE2J2* | 1.8962 | -0.68405 | -1.1726 | -1.6707 | 0.01 |
| 14631 | *TPSD1* | 0.1293 | 0.044352 | -0.13446 | -0.65811 | 0.01 |
| 14632 | *PRPF19* | 0.42931 | 0.050589 | -0.44073 | -1.4504 | 0.01 |
| 14633 | *C7orf26* | 1.6157 | -0.16119 | -1.4154 | -1.6759 | 0.01 |
| 14634 | *ALDH1A3* | 0.51125 | 0.46635 | -0.93892 | -1.175 | 0.01 |
| 14635 | *HIST1H1B* | 1.255 | 0.43487 | -1.6524 | -2.2712 | 0.01 |
| 14636 | *OR1I1* | 1.2671 | -0.08417 | -1.1459 | -2.1398 | 0.01 |
| 14637 | *CMPK2* | 0.8875 | 0.22021 | -1.0707 | -2.0407 | 0.01 |
| 14638 | *USP17L20* | 1.2043 | -0.48961 | -0.67809 | -1.8453 | 0.01 |
| 14639 | *RNF214* | 0.43709 | -0.08324 | -0.31737 | -1.754 | 0.01 |
| 14640 | *DEFB107A* | 1.2167 | 0.43888 | -1.6206 | -3.6516 | 0.01 |
| 14641 | *SST* | 0.33366 | 0.2405 | -0.53931 | -0.85389 | 0.01 |
| 14642 | *ZNF563* | 0.24777 | 0.090715 | -0.304 | -2.0304 | 0.01 |
| 14643 | *WBP2* | 1.4689 | -0.29341 | -1.1418 | -1.6463 | 0.01 |
| 14644 | *ROBO3* | 0.30139 | -0.0186 | -0.24942 | -1.534 | 0.01 |
| 14645 | *CLEC6A* | 0.78288 | 0.035409 | -0.78579 | -1.1958 | 0.01 |
| 14646 | *ST13* | 1.0654 | 0.079385 | -1.1125 | -2.0337 | 0.01 |
| 14647 | *ZNF219* | 1.5319 | -0.37116 | -1.1287 | -2.5007 | 0.01 |
| 14648 | *KRT6A* | 1.5559 | -0.72927 | -0.79483 | -1.021 | 0.01 |
| 14649 | *ARRB2* | 1.6876 | 0.11589 | -1.7718 | -3.158 | 0.01 |
| 14650 | *DNAJB14* | 2.1787 | -0.30065 | -1.8478 | -2.2694 | 0.01 |
| 14651 | *PACRG* | 1.1293 | -0.02583 | -1.0737 | -2.2945 | 0.01 |
| 14652 | *MSL2* | 1.1344 | -0.44374 | -0.66107 | -1.1194 | 0.01 |
| 14653 | *EHHADH* | 0.69538 | -0.00498 | -0.66098 | -1.2027 | 0.01 |
| 14654 | *CNFN* | 0.32496 | 0.002405 | -0.2994 | -0.79781 | 0.01 |
| 14655 | *TTI2* | 0.90948 | 0.065918 | -0.94759 | -1.8879 | 0.01 |
| 14656 | *CTU2* | 2.5528 | -1.186 | -1.3399 | -1.8238 | 0.01 |
| 14657 | *DZIP3* | 0.62797 | 0.42261 | -1.0248 | -1.6613 | 0.01 |
| 14658 | *CCDC151* | 1.3851 | -0.53639 | -0.82337 | -1.1943 | 0.01 |
| 14659 | *ZNF140* | 1.6475 | 0.51174 | -2.1342 | -2.3525 | 0.01 |
| 14660 | *PCDHB11* | 3.7826 | -1.2399 | -2.5178 | -2.8594 | 0.01 |
| 14661 | *CHAT* | 2.5023 | -1.0528 | -1.4252 | -1.9054 | 0.01 |
| 14662 | *KIF1B* | 1.4624 | -0.1507 | -1.2876 | -2.0277 | 0.01 |
| 14663 | *HFE2* | 0.26461 | -0.00178 | -0.23977 | -1.2705 | 0.01 |
| 14664 | *ZNF766* | 1.7084 | 0.23725 | -1.9227 | -3.6718 | 0.01 |
| 14665 | *ARGFX* | 1.4182 | -0.11101 | -1.2846 | -5.7998 | 0.01 |
| 14666 | *RBPJL* | 1.4916 | -0.57301 | -0.89699 | -2.6438 | 0.01 |
| 14667 | *ULK3* | 0.47997 | 0.43264 | -0.89219 | -1.4066 | 0.01 |

| | | | | | | |
|---|---|---|---|---|---|---|
| 14668 | *MAP3K4* | 0.96682 | -0.19128 | -0.75523 | -1.043 | 0.01 |
| 14669 | *C12orf56* | 0.12019 | 0.025299 | -0.12565 | -0.89699 | 0.01 |
| 14670 | *LIG3* | 1.0651 | -0.05504 | -0.9904 | -1.9474 | 0.01 |
| 14671 | *MFF* | 1.6603 | -0.01604 | -1.6251 | -1.7493 | 0.01 |
| 14672 | *GPRC5C* | 0.71177 | -0.27096 | -0.42254 | -0.78097 | 0.01 |
| 14673 | *ZNF438* | 0.659 | -0.29167 | -0.3495 | -0.60604 | 0.01 |
| 14674 | *RBMXL2* | 0.59473 | 0.2124 | -0.78956 | -1.2277 | 0.01 |
| 14675 | *FMOD* | 0.70073 | -0.09323 | -0.59088 | -1.1043 | 0.01 |
| 14676 | *CIAO1* | 0.67755 | 0.047789 | -0.70961 | -1.4108 | 0.01 |
| 14677 | *PMPCB* | 0.94917 | 0.16431 | -1.098 | -1.1018 | 0.01 |
| 14678 | *CLK1* | 0.46534 | -0.11886 | -0.33121 | -1.768 | 0.01 |
| 14679 | *IDH3A* | 0.6313 | -0.26086 | -0.35531 | -1.3128 | 0.01 |
| 14680 | *CCDC141* | 0.73314 | 0.14487 | -0.86403 | -1.2135 | 0.00 |
| 14681 | *ABAT* | 1.1583 | -0.14925 | -0.99508 | -1.3721 | 0.00 |
| 14682 | *CKLF-CMTM1* | 2.5441 | -1.2574 | -1.2733 | -2.012 | 0.00 |
| 14683 | *STAM* | 0.66128 | -0.09182 | -0.55676 | -2.5791 | 0.00 |
| 14684 | *FRMPD1* | 1.0429 | -0.21182 | -0.81846 | -0.96407 | 0.00 |
| 14685 | *MAPK7* | 1.1163 | -0.1931 | -0.91096 | -1.4401 | 0.00 |
| 14686 | *CHD4* | 0.73596 | -0.08582 | -0.638 | -1.3253 | 0.00 |
| 14687 | *CDC42EP3* | 0.39497 | -0.14987 | -0.23335 | -0.42451 | 0.00 |
| 14688 | *TBRG4* | 1.5101 | -0.66577 | -0.83278 | -1.0087 | 0.00 |
| 14689 | *RASD1* | 1.4437 | -0.15699 | -1.2756 | -1.5406 | 0.00 |
| 14690 | *PTCH1* | 0.38252 | 0.18719 | -0.55941 | -0.82903 | 0.00 |
| 14691 | *THG1L* | 0.71281 | -0.19272 | -0.51038 | -3.1719 | 0.00 |
| 14692 | *PRR32* | 0.5128 | 0.032842 | -0.53666 | -0.61779 | 0.00 |
| 14693 | *DCST1* | 0.083155 | 0.023698 | -0.0981 | -2.6122 | 0.00 |
| 14694 | *S100A13* | 0.43594 | 0.24242 | -0.66973 | -1.8724 | 0.00 |
| 14695 | *RAPGEF4* | 0.59967 | -0.13528 | -0.45669 | -1.2277 | 0.00 |
| 14696 | *UBASH3B* | 1.154 | -0.13746 | -1.0092 | -1.2879 | 0.00 |
| 14697 | *PCSK7* | 3.5757 | -1.0528 | -2.5156 | -3.0911 | 0.00 |
| 14698 | *F9* | 0.97281 | 0.16308 | -1.1298 | -2.2589 | 0.00 |
| 14699 | *MTTP* | 0.63928 | -0.08118 | -0.55291 | -1.6105 | 0.00 |
| 14700 | *CDC42* | 0.51333 | 0.048199 | -0.55677 | -1.676 | 0.00 |
| 14701 | *TKTL2* | 1.2018 | -0.0062 | -1.1911 | -1.3151 | 0.00 |
| 14702 | *SREK1IP1* | 2.6018 | -0.29454 | -2.3036 | -3.3146 | 0.00 |
| 14703 | *ARHGAP11B* | 0.26994 | 0.094294 | -0.36127 | -0.76176 | 0.00 |
| 14704 | *CLPTM1L* | 0.32171 | 0.26236 | -0.58183 | -2.3204 | 0.00 |
| 14705 | *FAT3* | 2.7127 | -1.2426 | -1.4682 | -1.9203 | 0.00 |
| 14706 | *FAM13C* | 0.99061 | 0.44717 | -1.4368 | -1.6525 | 0.00 |
| 14707 | *CDK1* | 0.78343 | 0.060939 | -0.84348 | -1.2267 | 0.00 |
| 14708 | *LRIF1* | 1.1229 | -0.54559 | -0.57761 | -1.5099 | 0.00 |
| 14709 | *CCDC77* | 0.67363 | 0.62691 | -1.3016 | -1.6855 | 0.00 |
| 14710 | *EIF2B1* | 1.0563 | -0.46752 | -0.58994 | -1.0452 | 0.00 |
| 14711 | *APOBEC3G* | 1.0328 | -0.26575 | -0.76893 | -2.2809 | 0.00 |
| 14712 | *IGFL4* | 0.34227 | 0.078186 | -0.42279 | -1.4156 | 0.00 |
| 14713 | *LONP2* | 1.4416 | -0.52017 | -0.92427 | -1.2692 | 0.00 |

| | | | | | | |
|---|---|---|---|---|---|---|
| 14714 | *SLC9B2* | 0.148 | 0.068212 | -0.21944 | -1.3475 | 0.00 |
| 14715 | *NAGA* | 0.072219 | 0.010651 | -0.08675 | -1.3772 | 0.00 |
| 14716 | *AGXT* | 0.94679 | -0.42359 | -0.52712 | -1.7065 | 0.00 |
| 14717 | *DDAH2* | 0.79018 | 0.058283 | -0.8525 | -1.2608 | 0.00 |
| 14718 | *TUT1* | 0.60632 | -0.18691 | -0.42353 | -2.043 | 0.00 |
| 14719 | *POLR1D* | 0.67748 | 0.66252 | -1.3444 | -2.1873 | 0.00 |
| 14720 | *HIVEP1* | 1.1507 | 0.14587 | -1.3016 | -1.3457 | 0.00 |
| 14721 | *ZNF625* | 0.99378 | 0.34486 | -1.3443 | -1.7595 | 0.00 |
| 14722 | *WNT4* | 0.5526 | 0.35566 | -0.91427 | -1.2517 | 0.00 |
| 14723 | *GOLPH3* | 2.0286 | -0.92025 | -1.1144 | -3.2735 | 0.00 |
| 14724 | *MLX* | 1.2368 | 0.16326 | -1.4062 | -1.455 | 0.00 |
| 14725 | *GLE1* | 0.6979 | -0.25181 | -0.45305 | -2.462 | 0.00 |
| 14726 | *SLC30A2* | 0.45525 | 0.29982 | -0.76217 | -1.2935 | 0.00 |
| 14727 | *IGFBP5* | 0.74965 | -0.22585 | -0.53125 | -0.72585 | 0.00 |
| 14728 | *STC1* | 2.0783 | -0.81529 | -1.2705 | -1.7462 | 0.00 |
| 14729 | *OR5K3* | 0.42128 | 0.042086 | -0.47126 | -0.71988 | 0.00 |
| 14730 | *RAB10* | 0.65119 | 0.52504 | -1.1844 | -1.2284 | 0.00 |
| 14731 | *LHX5* | 2.366 | -1.1545 | -1.2198 | -2.012 | 0.00 |
| 14732 | *SULF2* | 0.99747 | 0.4234 | -1.4294 | -1.8123 | 0.00 |
| 14733 | *RIIAD1* | 0.5099 | -0.19577 | -0.32429 | -0.96776 | 0.00 |
| 14734 | *TRIM26* | 1.2501 | -0.46303 | -0.79871 | -1.1619 | 0.00 |
| 14735 | *ATP2A2* | 1.3233 | -0.21646 | -1.12 | -1.2008 | 0.00 |
| 14736 | *WLS* | 0.6799 | 0.15374 | -0.8471 | -1.3739 | 0.00 |
| 14737 | *ISG20* | 1.3834 | -0.35232 | -1.0451 | -1.5959 | 0.00 |
| 14738 | *RNF41* | 0.31222 | -0.00082 | -0.32546 | -0.7757 | 0.00 |
| 14739 | *GRB14* | 0.93429 | 0.086818 | -1.0355 | -1.9533 | 0.00 |
| 14740 | *DSG3* | 1.0898 | 0.45486 | -1.5597 | -1.9997 | -0.01 |
| 14741 | *FAM65B* | 0.99081 | 0.45639 | -1.4625 | -1.962 | -0.01 |
| 14742 | *SRBD1* | 1.1596 | 0.29867 | -1.4745 | -1.7106 | -0.01 |
| 14743 | *NREP* | 0.65314 | 0.027969 | -0.69866 | -1.5306 | -0.01 |
| 14744 | *FAM177B* | 0.45381 | 0.11774 | -0.58929 | -1.4269 | -0.01 |
| 14745 | *FAM149A* | 0.72553 | -0.04769 | -0.69631 | -0.73301 | -0.01 |
| 14746 | *HES2* | 0.78428 | -0.38041 | -0.42291 | -3.6398 | -0.01 |
| 14747 | *CRIPAK* | 1.0913 | -0.04621 | -1.0651 | -1.4405 | -0.01 |
| 14748 | *SNAPC1* | 0.37234 | 0.20442 | -0.59683 | -1.165 | -0.01 |
| 14749 | *ATP10D* | 0.57587 | 0.1072 | -0.70325 | -1.1212 | -0.01 |
| 14750 | *C2orf15* | 0.65987 | 0.63624 | -1.3165 | -2.4815 | -0.01 |
| 14751 | *FAM32A* | 1.7159 | -0.36751 | -1.3689 | -2.9844 | -0.01 |
| 14752 | *RPL9* | 1.0105 | -0.17037 | -0.86157 | -1.6176 | -0.01 |
| 14753 | *TFG* | 1.198 | -0.43376 | -0.78591 | -1.2171 | -0.01 |
| 14754 | *ZNF396* | 1.7537 | -0.64712 | -1.1297 | -2.8647 | -0.01 |
| 14755 | *GDF6* | 0.92 | 0.3642 | -1.3085 | -1.4116 | -0.01 |
| 14756 | *LCE3A* | 0.78876 | -0.20552 | -0.60754 | -1.5393 | -0.01 |
| 14757 | *LGALS3BP* | 1.9009 | -0.77462 | -1.1511 | -1.7782 | -0.01 |
| 14758 | *GRAMD4* | 0.67882 | 0.22058 | -0.92435 | -3.2735 | -0.01 |
| 14759 | *ACSBG1* | 0.77909 | 0.1452 | -0.95136 | -1.7065 | -0.01 |

| | | | | | | |
|---|---|---|---|---|---|---|
| 14760 | *TAF5* | 1.8424 | -0.72201 | -1.1479 | -1.2837 | -0.01 |
| 14761 | *ENY2* | 0.49122 | 0.27132 | -0.79043 | -1.7646 | -0.01 |
| 14762 | *APOL2* | 1.0371 | -0.09807 | -0.96718 | -1.7804 | -0.01 |
| 14763 | *C17orf82* | 0.68847 | -0.17425 | -0.54259 | -0.9653 | -0.01 |
| 14764 | *TIAL1* | 0.028715 | -0.02128 | -0.03686 | -0.36749 | -0.01 |
| 14765 | *ACTL7B* | 1.8741 | 0.35451 | -2.2586 | -2.5934 | -0.01 |
| 14766 | *NPBWR2* | 0.42752 | -0.22094 | -0.23698 | -0.32486 | -0.01 |
| 14767 | *GPR179* | 1.235 | 0.091878 | -1.3573 | -1.5049 | -0.01 |
| 14768 | *C22orf23* | 2.2684 | -0.94364 | -1.3556 | -1.5238 | -0.01 |
| 14769 | *ITPKA* | 1.0131 | -0.49846 | -0.54555 | -2.2413 | -0.01 |
| 14770 | *REXO2* | 0.7376 | 0.051512 | -0.8203 | -2.0495 | -0.01 |
| 14771 | *LTK* | 0.43421 | -0.02298 | -0.44299 | -2.1572 | -0.01 |
| 14772 | *SLC39A9* | 2.5321 | -1.1631 | -1.4018 | -1.4168 | -0.01 |
| 14773 | *SERHL2* | 1.2509 | -0.08975 | -1.1956 | -1.2867 | -0.01 |
| 14774 | *CT47A7* | 1.094 | -0.0975 | -1.0318 | -1.0995 | -0.01 |
| 14775 | *ZNF282* | 0.28522 | 0.12282 | -0.44353 | -0.74011 | -0.01 |
| 14776 | *SPAG5* | 1.5418 | -0.75529 | -0.82211 | -1.4359 | -0.01 |
| 14777 | *ALG1L2* | 0.46625 | 0.2635 | -0.7655 | -0.91981 | -0.01 |
| 14778 | *EIF2S2* | 0.43228 | -0.02447 | -0.44575 | -1.8014 | -0.01 |
| 14779 | *FUS* | 0.44132 | 0.005506 | -0.4854 | -1.984 | -0.01 |
| 14780 | *VGLL1* | 0.12471 | -0.05585 | -0.10828 | -1.1056 | -0.01 |
| 14781 | *BIN1* | 1.3259 | -0.21438 | -1.1511 | -1.1817 | -0.01 |
| 14782 | *CXorf49B* | 1.7635 | -0.38011 | -1.4233 | -1.7874 | -0.01 |
| 14783 | *ZNF274* | 1.7172 | -0.80853 | -0.94912 | -1.8698 | -0.01 |
| 14784 | *ZBTB17* | 0.51199 | 0.2015 | -0.75448 | -1.1703 | -0.01 |
| 14785 | *DUSP2* | 2.5732 | -1.3057 | -1.3089 | -2.0778 | -0.01 |
| 14786 | *PLK2* | 0.32713 | 0.049777 | -0.4187 | -1.4824 | -0.01 |
| 14787 | *SNRNP27* | 1.4316 | -0.52943 | -0.94441 | -1.9867 | -0.01 |
| 14788 | *CENPA* | 0.93194 | 0.25563 | -1.2302 | -1.8275 | -0.01 |
| 14789 | *DRAM2* | 0.71415 | -0.31808 | -0.43886 | -1.2719 | -0.01 |
| 14790 | *LILRB1* | 0.34228 | 0.18739 | -0.57397 | -1.2863 | -0.01 |
| 14791 | *ARL6IP1* | 2.5601 | -0.8988 | -1.7056 | -2.508 | -0.01 |
| 14792 | *HTR6* | 1.2206 | -0.00111 | -1.2655 | -2.2888 | -0.02 |
| 14793 | *RUNDC3A* | 1.3806 | -0.62548 | -0.80146 | -1.1169 | -0.02 |
| 14794 | *SCYL3* | 1.24 | -0.54847 | -0.73787 | -1.6759 | -0.02 |
| 14795 | *NUP133* | 1.2206 | -0.22301 | -1.0441 | -1.0947 | -0.02 |
| 14796 | *VRK2* | 0.76478 | -0.2765 | -0.53584 | -2.2727 | -0.02 |
| 14797 | *LRRTM1* | 1.3602 | -0.47338 | -0.93533 | -1.9504 | -0.02 |
| 14798 | *LPGAT1* | 1.0285 | -0.06364 | -1.0149 | -2.4655 | -0.02 |
| 14799 | *OR14J1* | 0.47742 | -0.01371 | -0.51494 | -1.951 | -0.02 |
| 14800 | *CLCNKB* | 1.2851 | -0.24797 | -1.0884 | -1.2952 | -0.02 |
| 14801 | *DMRTC1B* | 0.69499 | 0.50317 | -1.25 | -3.034 | -0.02 |
| 14802 | *PCMT1* | 0.65144 | -0.0391 | -0.66543 | -1.7879 | -0.02 |
| 14803 | *LIN54* | 0.13779 | 0.019567 | -0.21094 | -0.56281 | -0.02 |
| 14804 | *CCKBR* | 0.44805 | 0.38467 | -0.8865 | -2.6438 | -0.02 |
| 14805 | *PNLIPRP3* | 0.32118 | 0.097548 | -0.47353 | -1.5375 | -0.02 |

| | | | | | | |
|---|---|---|---|---|---|---|
| 14806 | *TUBB* | 1.8727 | -0.40534 | -1.5225 | -1.7438 | -0.02 |
| 14807 | *SATB2* | 2.1932 | -0.80405 | -1.4454 | -1.7513 | -0.02 |
| 14808 | *CAPNS1* | 1.2993 | -0.35111 | -1.0047 | -1.5911 | -0.02 |
| 14809 | *MMP14* | 2.4502 | -1.209 | -1.2981 | -2.674 | -0.02 |
| 14810 | *MRPL22* | 0.60896 | -0.09025 | -0.57564 | -1.3808 | -0.02 |
| 14811 | *NIPA1* | 0.32813 | 0.13256 | -0.51768 | -0.91256 | -0.02 |
| 14812 | *FIGF* | 0.73147 | 0.72314 | -1.5125 | -4.1058 | -0.02 |
| 14813 | *SYNJ2* | 1.0963 | -0.25792 | -0.89641 | -1.6437 | -0.02 |
| 14814 | *PC* | 0.34077 | 0.28036 | -0.6794 | -1.6796 | -0.02 |
| 14815 | *BAG1* | 0.94387 | 0.21191 | -1.2141 | -1.5971 | -0.02 |
| 14816 | *PVRL2* | 0.31943 | -0.17936 | -0.19851 | -1.3607 | -0.02 |
| 14817 | *EGF* | 0.78469 | 0.21634 | -1.0606 | -1.1564 | -0.02 |
| 14818 | *WDR11* | 0.34909 | -0.117 | -0.29209 | -0.87673 | -0.02 |
| 14819 | *ATP8B4* | 0.89321 | 0.45901 | -1.4129 | -2.2014 | -0.02 |
| 14820 | *EDDM3A* | 0.27161 | 0.033893 | -0.36679 | -2.3405 | -0.02 |
| 14821 | *VAT1L* | 2.1442 | -1.0841 | -1.122 | -1.621 | -0.02 |
| 14822 | *OR6C6* | 1.5552 | 0.75789 | -2.3751 | -2.6275 | -0.02 |
| 14823 | *PPP1R13L* | 2.1076 | -0.75542 | -1.4142 | -2.1662 | -0.02 |
| 14824 | *ZNF655* | 1.4863 | -0.47642 | -1.0726 | -1.1144 | -0.02 |
| 14825 | *OR2V2* | 1.828 | -0.59762 | -1.2936 | -1.794 | -0.02 |
| 14826 | *MED21* | 1.0724 | 0.15924 | -1.296 | -1.3941 | -0.02 |
| 14827 | *TNFAIP8* | 0.83068 | 0.78944 | -1.6847 | -2.0042 | -0.02 |
| 14828 | *SDF4* | 2.1613 | -0.88784 | -1.3382 | -1.9251 | -0.02 |
| 14829 | *POTEC* | 0.28017 | 0.20131 | -0.54651 | -1.0196 | -0.02 |
| 14830 | *MED4* | 0.5892 | 0.067247 | -0.72204 | -1.4829 | -0.02 |
| 14831 | *OLAH* | 0.38955 | 0.29533 | -0.7513 | -1.8478 | -0.02 |
| 14832 | *CEP295* | 0.60384 | 0.049015 | -0.72085 | -2.3452 | -0.02 |
| 14833 | *CYB5A* | 0.46502 | -0.22921 | -0.30444 | -1.4962 | -0.02 |
| 14834 | *NBEAL1* | 2.1538 | -1.0191 | -1.2036 | -1.7745 | -0.02 |
| 14835 | *QPCT* | 0.99477 | -0.1227 | -0.94341 | -2.1921 | -0.02 |
| 14836 | *PCNX* | 0.19463 | 0.14163 | -0.40823 | -0.45193 | -0.02 |
| 14837 | *IL2* | 1.415 | -0.02583 | -1.4614 | -1.8016 | -0.02 |
| 14838 | *FAM102B* | 1.2419 | -0.1672 | -1.1472 | -2.3357 | -0.02 |
| 14839 | *WIPI1* | 3.0215 | -1.5434 | -1.5508 | -1.6575 | -0.02 |
| 14840 | *PRAMEF12* | 0.56421 | 0.50072 | -1.1381 | -1.9162 | -0.02 |
| 14841 | *CAMTA2* | 1.2244 | 0.72209 | -2.0197 | -3.1317 | -0.02 |
| 14842 | *SLC52A1* | 0.68744 | 0.63689 | -1.3978 | -1.4364 | -0.02 |
| 14843 | *CCDC9* | 2.9212 | -1.3492 | -1.6456 | -2.2533 | -0.02 |
| 14844 | *TMEM47* | 1.5302 | -0.4521 | -1.1526 | -1.3479 | -0.02 |
| 14845 | *FCHO2* | 0.58849 | 0.46699 | -1.1309 | -1.3668 | -0.03 |
| 14846 | *CCR9* | 1.4003 | -0.31475 | -1.161 | -1.3835 | -0.03 |
| 14847 | *RBKS* | 1.1724 | -0.3464 | -0.90281 | -1.8219 | -0.03 |
| 14848 | *DUSP7* | 0.49996 | -0.09268 | -0.48416 | -2.6289 | -0.03 |
| 14849 | *PIGM* | 1.0291 | -0.15649 | -0.94954 | -2.0885 | -0.03 |
| 14850 | *GGNBP2* | 1.168 | -0.59921 | -0.64725 | -1.1133 | -0.03 |
| 14851 | *NDUFS5* | 1.1356 | 0.37555 | -1.5898 | -1.7328 | -0.03 |

| | | | | | | |
|---|---|---|---|---|---|---|
| 14852 | *LEF1* | 1.0764 | -0.11897 | -1.0364 | -1.1207 | -0.03 |
| 14853 | *ATP5S* | 1.0545 | -0.2345 | -0.89931 | -0.97751 | -0.03 |
| 14854 | *TUBGCP2* | 1.3732 | -0.51182 | -0.94255 | -1.6098 | -0.03 |
| 14855 | *NUDCD1* | 0.66316 | 0.091533 | -0.83627 | -1.2708 | -0.03 |
| 14856 | *NPIPB6* | 1.7562 | -0.04486 | -1.7935 | -1.8941 | -0.03 |
| 14857 | *SLC25A51* | 0.761 | 0.42143 | -1.2646 | -2.5092 | -0.03 |
| 14858 | *GTF2H2* | 1.3786 | -0.19804 | -1.2633 | -1.8508 | -0.03 |
| 14859 | *PFKM* | 1.048 | 0.39428 | -1.5271 | -1.7493 | -0.03 |
| 14860 | *FAM9B* | 0.46477 | 0.27747 | -0.82757 | -2.0851 | -0.03 |
| 14861 | *SLC2A13* | 0.79121 | 0.68177 | -1.5588 | -1.9041 | -0.03 |
| 14862 | *AVPR1B* | 0.54927 | -0.15424 | -0.48144 | -0.9925 | -0.03 |
| 14863 | *EFNA2* | 0.72307 | 0.001035 | -0.81087 | -0.90492 | -0.03 |
| 14864 | *DOCK1* | 0.99511 | -0.50067 | -0.58167 | -2.6574 | -0.03 |
| 14865 | *TMED10* | 2.5182 | -0.56411 | -2.0414 | -2.1404 | -0.03 |
| 14866 | *TSPAN7* | 0.52471 | 0.19941 | -0.81152 | -0.9497 | -0.03 |
| 14867 | *USH1G* | 0.19327 | -0.13962 | -0.14151 | -1.1685 | -0.03 |
| 14868 | *KIAA1456* | 0.86582 | 0.4223 | -1.3766 | -1.6208 | -0.03 |
| 14869 | *CLEC4G* | 0.56668 | -0.19804 | -0.46107 | -0.78229 | -0.03 |
| 14870 | *MRPL54* | 0.91172 | 0.50373 | -1.5079 | -1.5441 | -0.03 |
| 14871 | *EFNA3* | 1.6122 | -0.52372 | -1.1817 | -1.3153 | -0.03 |
| 14872 | *NAAA* | 0.096469 | -0.02812 | -0.16267 | -1.2755 | -0.03 |
| 14873 | *B3GAT3* | 0.63535 | 0.24537 | -0.97549 | -1.8392 | -0.03 |
| 14874 | *OR5B2* | -0.01948 | -0.02845 | -0.04799 | -0.8202 | -0.03 |
| 14875 | *SEC14L6* | 0.55574 | 0.04489 | -0.69665 | -0.9416 | -0.03 |
| 14876 | *GPR78* | 0.55507 | 0.55011 | -1.2052 | -1.8478 | -0.03 |
| 14877 | *SLCO1A2* | 1.4515 | -0.03643 | -1.5153 | -1.5959 | -0.03 |
| 14878 | *PSMC2* | 1.3028 | -0.09354 | -1.3101 | -1.595 | -0.03 |
| 14879 | *RAB33B* | 1.0007 | 0.28 | -1.386 | -1.5144 | -0.04 |
| 14880 | *ADAT2* | 0.35613 | 0.13367 | -0.59516 | -1.0273 | -0.04 |
| 14881 | *ASB14* | 0.39528 | -0.15876 | -0.34227 | -1.162 | -0.04 |
| 14882 | *GFM2* | 1.0049 | -0.05804 | -1.0529 | -1.4306 | -0.04 |
| 14883 | *P2RX2* | 0.76989 | -0.43264 | -0.44417 | -0.46728 | -0.04 |
| 14884 | *NFKBIB* | 1.9694 | -0.76389 | -1.3141 | -1.8879 | -0.04 |
| 14885 | *OR51I1* | 1.0666 | 0.33336 | -1.5099 | -1.6199 | -0.04 |
| 14886 | *MTA3* | 0.5998 | 0.43552 | -1.146 | -2.4221 | -0.04 |
| 14887 | *FOLH1B* | 0.4558 | -0.22675 | -0.34266 | -2.1616 | -0.04 |
| 14888 | *RPL7* | 1.3935 | -0.09179 | -1.416 | -2.4761 | -0.04 |
| 14889 | *HNRNPCL4* | 3.3203 | -1.4878 | -1.9474 | -2.0538 | -0.04 |
| 14890 | *TMEM63C* | 0.83881 | -0.18135 | -0.77317 | -1.0036 | -0.04 |
| 14891 | *FOXD4* | 1.392 | 0.36759 | -1.8759 | -2.088 | -0.04 |
| 14892 | *MMP24* | 0.74195 | -0.22886 | -0.62952 | -1.5018 | -0.04 |
| 14893 | *MAP7* | 0.013119 | -0.02646 | -0.10392 | -0.89052 | -0.04 |
| 14894 | *THSD4* | 0.43376 | 0.14764 | -0.69904 | -2.6072 | -0.04 |
| 14895 | *EBF2* | 0.19645 | 0.051282 | -0.36549 | -0.84858 | -0.04 |
| 14896 | *IL6ST* | 0.62542 | 0.12351 | -0.86802 | -1.2987 | -0.04 |
| 14897 | *WDTC1* | 3.4776 | -1.63 | -1.9678 | -2.184 | -0.04 |

| | | | | | | |
|---|---|---|---|---|---|---|
| 14898 | *ICAM4* | 0.90187 | -0.32409 | -0.69826 | -1.4762 | -0.04 |
| 14899 | *SERPINH1* | 1.9196 | -0.87792 | -1.1632 | -5.6834 | -0.04 |
| 14900 | *UBR4* | 1.9274 | -0.50282 | -1.5461 | -1.7643 | -0.04 |
| 14901 | *BCL9L* | 0.63121 | 0.30293 | -1.0557 | -1.8443 | -0.04 |
| 14902 | *FAM177A1* | 0.55698 | 0.25098 | -0.92975 | -1.677 | -0.04 |
| 14903 | *PRLH* | 0.24928 | 0.016392 | -0.38786 | -2.3222 | -0.04 |
| 14904 | *N4BP1* | 1.0263 | -0.06763 | -1.0809 | -1.9341 | -0.04 |
| 14905 | *SEMA5A* | 0.52683 | -0.23158 | -0.41754 | -1.191 | -0.04 |
| 14906 | *SLC35G4* | 0.30472 | -0.20338 | -0.22422 | -2.2165 | -0.04 |
| 14907 | *GRIK4* | 0.94035 | 0.14156 | -1.2051 | -1.8311 | -0.04 |
| 14908 | *PIK3R3* | 0.94956 | -0.50978 | -0.56453 | -0.71443 | -0.04 |
| 14909 | *RUSC1-AS1* | 0.43282 | 0.16218 | -0.71986 | -1.2981 | -0.04 |
| 14910 | *NME1-NME2* | 1.3114 | -0.56947 | -0.86698 | -1.805 | -0.04 |
| 14911 | *DDX27* | 0.75279 | -0.00014 | -0.87772 | -1.041 | -0.04 |
| 14912 | *PIFO* | 1.4121 | -0.568 | -0.96962 | -1.3478 | -0.04 |
| 14913 | *TBX18* | 0.81325 | -0.15845 | -0.78229 | -1.3118 | -0.04 |
| 14914 | *SUN3* | 1.2353 | -0.56501 | -0.79798 | -1.6208 | -0.04 |
| 14915 | *CCL19* | 2.4574 | -1.1538 | -1.4317 | -1.4684 | -0.04 |
| 14916 | *FAM71E1* | 2.3135 | -1.1358 | -1.3058 | -1.8589 | -0.04 |
| 14917 | *NDUFB3* | 0.39772 | -0.25839 | -0.2681 | -1.8408 | -0.04 |
| 14918 | *FAM198A* | 1.0962 | -0.08842 | -1.1367 | -1.151 | -0.04 |
| 14919 | *UNC13A* | 1.3712 | -0.2168 | -1.2845 | -2.0267 | -0.04 |
| 14920 | *MBNL3* | 0.12285 | -0.07106 | -0.18283 | -2.3287 | -0.04 |
| 14921 | *UGT2B17* | 0.58992 | 0.04816 | -0.76966 | -2.0452 | -0.04 |
| 14922 | *ELL2* | 0.81997 | 0.11634 | -1.0684 | -1.6583 | -0.04 |
| 14923 | *BMPR1A* | 0.62055 | -0.04678 | -0.70591 | -2.1498 | -0.04 |
| 14924 | *CFD* | 0.87281 | 0.55577 | -1.5608 | -1.8157 | -0.04 |
| 14925 | *SDHB* | 1.0265 | 0.40799 | -1.5671 | -2.3033 | -0.04 |
| 14926 | *FKBP7* | 1.0043 | -0.3706 | -0.76671 | -1.9107 | -0.04 |
| 14927 | *ODF2* | 0.55442 | -0.25608 | -0.43251 | -0.86123 | -0.04 |
| 14928 | *INA* | 2.8623 | -1.236 | -1.761 | -2.1351 | -0.04 |
| 14929 | *NSG1* | -0.02773 | -0.04439 | -0.06468 | -1.6368 | -0.05 |
| 14930 | *CLIC5* | 3.142 | -1.5608 | -1.7188 | -1.8816 | -0.05 |
| 14931 | *PCK2* | 0.44551 | -0.01337 | -0.56982 | -1.7536 | -0.05 |
| 14932 | *GGCX* | 1.7281 | -0.2263 | -1.6395 | -1.6447 | -0.05 |
| 14933 | *FOXF1* | 1.857 | -0.78718 | -1.2079 | -2.0303 | -0.05 |
| 14934 | *SLC25A10* | 0.62692 | 0.063324 | -0.82844 | -1.1156 | -0.05 |
| 14935 | *NPIPA2* | 2.2716 | -0.88666 | -1.5232 | -1.6887 | -0.05 |
| 14936 | *GALNT13* | 0.29415 | 0.079836 | -0.51266 | -2.5081 | -0.05 |
| 14937 | *PADI2* | 2.0414 | -0.73861 | -1.4416 | -1.783 | -0.05 |
| 14938 | *BRMS1* | 0.3521 | -0.04751 | -0.44353 | -0.82628 | -0.05 |
| 14939 | *MPHOSPH10* | 0.46987 | 0.44142 | -1.0503 | -1.9065 | -0.05 |
| 14940 | *ITPKC* | 2.4244 | -0.98104 | -1.5824 | -2.5477 | -0.05 |
| 14941 | *LOXL4* | 0.44105 | -0.00058 | -0.58011 | -0.87169 | -0.05 |
| 14942 | *BRAF* | 0.62527 | -0.00178 | -0.76358 | -1.3604 | -0.05 |
| 14943 | *C17orf100* | 1.1123 | 0.60243 | -1.855 | -1.9052 | -0.05 |

| | | | | | | |
|---|---|---|---|---|---|---|
| 14944 | *MYLK4* | 1.0269 | -0.57546 | -0.59183 | -0.9295 | -0.05 |
| 14945 | *ZAP70* | 0.088704 | 0.07582 | -0.30559 | -0.58553 | -0.05 |
| 14946 | *MSH5* | 0.94448 | -0.23146 | -0.85409 | -1.4567 | -0.05 |
| 14947 | *MST1* | 0.9593 | 0.080148 | -1.181 | -1.2903 | -0.05 |
| 14948 | *MITD1* | 0.55639 | -0.27486 | -0.42439 | -2.051 | -0.05 |
| 14949 | *TRPV2* | 1.062 | -0.22082 | -0.98459 | -1.5214 | -0.05 |
| 14950 | *CHSY3* | 0.51174 | 0.046264 | -0.70171 | -0.85093 | -0.05 |
| 14951 | *NDUFV2* | 1.9204 | -0.50535 | -1.559 | -2.7068 | -0.05 |
| 14952 | *FAM134B* | 0.36473 | 0.3053 | -0.81524 | -2.2599 | -0.05 |
| 14953 | *LDLRAD3* | 2.428 | -1.2193 | -1.3542 | -1.6726 | -0.05 |
| 14954 | *BCL2L12* | 1.4618 | -0.12866 | -1.4789 | -2.0068 | -0.05 |
| 14955 | *EFEMP1* | 0.62329 | -0.13183 | -0.63746 | -1.7407 | -0.05 |
| 14956 | *BRMS1L* | 1.479 | -0.73739 | -0.88775 | -1.2426 | -0.05 |
| 14957 | *AAGAB* | 1.5222 | -0.44136 | -1.227 | -1.4683 | -0.05 |
| 14958 | *GPR160* | 2.2605 | -1.0175 | -1.3892 | -1.9446 | -0.05 |
| 14959 | *TBCK* | 1.622 | -0.65178 | -1.1169 | -2.4322 | -0.05 |
| 14960 | *ADGRF5* | 0.70886 | -0.10116 | -0.75487 | -1.5914 | -0.05 |
| 14961 | *SPG20* | 0.79589 | 0.75574 | -1.6989 | -2.6267 | -0.05 |
| 14962 | *MGAT2* | 0.40783 | -0.1741 | -0.38161 | -0.9733 | -0.05 |
| 14963 | *TSPY4* | 0.29118 | -0.12786 | -0.31121 | -0.9285 | -0.05 |
| 14964 | *RETNLB* | 0.95717 | -0.42671 | -0.67853 | -1.083 | -0.05 |
| 14965 | *ATP5I* | 1.0066 | -0.43176 | -0.72334 | -1.0773 | -0.05 |
| 14966 | *TOE1* | 0.42787 | -0.02705 | -0.55002 | -1.0019 | -0.05 |
| 14967 | *HPCAL4* | 1.8636 | -0.87647 | -1.1365 | -1.3784 | -0.05 |
| 14968 | *BEND7* | 0.19351 | -0.00823 | -0.33468 | -0.87208 | -0.05 |
| 14969 | *NDRG1* | 1.7573 | -0.62531 | -1.2815 | -1.5738 | -0.05 |
| 14970 | *ANAPC5* | 3.005 | -1.2568 | -1.8982 | -2.3145 | -0.05 |
| 14971 | *TMEM110-MUSTN1* | 0.56292 | -0.23957 | -0.47447 | -3.1051 | -0.05 |
| 14972 | *SMO* | 1.3287 | -0.30715 | -1.1727 | -1.4087 | -0.05 |
| 14973 | *DPEP1* | 0.41563 | 0.044921 | -0.61302 | -1.1075 | -0.05 |
| 14974 | *SHARPIN* | 0.90913 | -0.43488 | -0.62679 | -1.3411 | -0.05 |
| 14975 | *NFKBIL1* | 1.6469 | -0.55281 | -1.2473 | -1.5996 | -0.05 |
| 14976 | *PCF11* | 0.10688 | 0.042515 | -0.30371 | -1.7646 | -0.05 |
| 14977 | *ELF3* | 1.1061 | -0.37976 | -0.88278 | -1.8033 | -0.05 |
| 14978 | *ATG5* | 1.6043 | -0.47964 | -1.2815 | -1.8376 | -0.05 |
| 14979 | *ANKRD40* | 0.055485 | -0.09689 | -0.11607 | -1.3116 | -0.05 |
| 14980 | *COX10* | 0.33406 | -0.2441 | -0.24812 | -1.337 | -0.05 |
| 14981 | *ZNF195* | 1.002 | -0.21576 | -0.94466 | -1.5244 | -0.05 |
| 14982 | *LOC441155* | 0.80359 | -0.00392 | -0.95893 | -1.7991 | -0.05 |
| 14983 | *GDF1* | 1.7453 | -0.81119 | -1.0935 | -1.6527 | -0.05 |
| 14984 | *CMTM1* | 1.0588 | -0.52187 | -0.6972 | -1.1316 | -0.05 |
| 14985 | *WDR64* | 0.88035 | -0.19054 | -0.85027 | -1.7793 | -0.05 |
| 14986 | *NLRP8* | 0.54 | -0.22803 | -0.47412 | -2.2319 | -0.05 |
| 14987 | *RFX4* | 0.60831 | 0.18363 | -0.95461 | -2.1354 | -0.05 |
| 14988 | *SMIM23* | 0.33545 | 0.18216 | -0.68109 | -1.9038 | -0.05 |
| 14989 | *DAG1* | 0.046128 | 0.006186 | -0.21632 | -0.5442 | -0.05 |

| | | | | | | |
|---|---|---|---|---|---|---|
| 14990 | *QRICH1* | 1.682 | 0.19205 | -2.0384 | -2.4014 | -0.05 |
| 14991 | *PIWIL4* | 0.50535 | -0.16295 | -0.50685 | -1.2234 | -0.05 |
| 14992 | *CDKN2D* | 0.67487 | 0.040706 | -0.88098 | -1.0088 | -0.06 |
| 14993 | *DFFA* | 0.50243 | 0.1989 | -0.86744 | -1.0496 | -0.06 |
| 14994 | *ICE1* | 0.87021 | 0.4185 | -1.4553 | -1.5558 | -0.06 |
| 14995 | *MORC2* | 0.080715 | 0.049826 | -0.29794 | -0.80146 | -0.06 |
| 14996 | *TMEM119* | 3.0987 | -1.5415 | -1.7247 | -2.2059 | -0.06 |
| 14997 | *C14orf1* | 1.3406 | 0.66952 | -2.1792 | -2.8125 | -0.06 |
| 14998 | *SNAPC3* | 1.001 | 0.026876 | -1.197 | -1.4811 | -0.06 |
| 14999 | *ADD1* | 0.62071 | -0.21398 | -0.57732 | -0.65951 | -0.06 |
| 15000 | *DIRC2* | 0.65962 | -0.25794 | -0.57307 | -1.0413 | -0.06 |
| 15001 | *BOP1* | 0.027623 | -0.06199 | -0.13737 | -2.8647 | -0.06 |
| 15002 | *RSAD1* | 0.86236 | -0.48808 | -0.54639 | -2.2074 | -0.06 |
| 15003 | *UBE2G2* | 1.1169 | 0.44095 | -1.7307 | -1.8954 | -0.06 |
| 15004 | *GPRIN3* | 1.2466 | -0.23878 | -1.181 | -1.677 | -0.06 |
| 15005 | *RBL1* | 1.0747 | 0.078551 | -1.3268 | -1.481 | -0.06 |
| 15006 | *CHERP* | 0.84974 | 0.031869 | -1.0557 | -1.2774 | -0.06 |
| 15007 | *ATCAY* | 0.74321 | -0.12418 | -0.79319 | -1.5235 | -0.06 |
| 15008 | *ZNF786* | 0.64233 | 0.18334 | -0.9999 | -1.9107 | -0.06 |
| 15009 | *SLC29A2* | 0.47467 | -0.01574 | -0.63336 | -1.2837 | -0.06 |
| 15010 | *SBNO2* | 1.2562 | -0.69495 | -0.73814 | -1.7606 | -0.06 |
| 15011 | *GMNN* | 0.19259 | 0.008446 | -0.37913 | -0.95505 | -0.06 |
| 15012 | *ORC1* | 0.68288 | 0.16141 | -1.0224 | -1.365 | -0.06 |
| 15013 | *ADARB1* | 0.76977 | -0.15882 | -0.78916 | -1.2639 | -0.06 |
| 15014 | *GOLGA8A* | 2.1325 | -1.1321 | -1.1796 | -1.9772 | -0.06 |
| 15015 | *SSX4B* | 0.79248 | 0.5137 | -1.4856 | -1.9413 | -0.06 |
| 15016 | *AGTPBP1* | 0.54804 | 0.13633 | -0.86384 | -0.89052 | -0.06 |
| 15017 | *SETD2* | 0.6959 | 0.041702 | -0.91721 | -1.3542 | -0.06 |
| 15018 | *ERMN* | 0.85911 | -0.14132 | -0.89866 | -1.1585 | -0.06 |
| 15019 | *C12orf65* | 0.41594 | 0.068299 | -0.66535 | -1.2159 | -0.06 |
| 15020 | *TEF* | 0.37182 | -0.22131 | -0.33191 | -2.4328 | -0.06 |
| 15021 | *ATP5F1* | 0.44354 | 0.34162 | -0.96718 | -1.4633 | -0.06 |
| 15022 | *SMC5* | 0.91486 | 0.40783 | -1.5054 | -1.9423 | -0.06 |
| 15023 | *SLC25A16* | 0.51375 | -0.30086 | -0.39565 | -1.0952 | -0.06 |
| 15024 | *PLCL2* | 1.3087 | -0.04729 | -1.445 | -1.7209 | -0.06 |
| 15025 | *IL6R* | 0.53319 | 0.004026 | -0.72128 | -1.0863 | -0.06 |
| 15026 | *CNOT7* | 1.1055 | -0.30481 | -0.98574 | -2.1105 | -0.06 |
| 15027 | *FTSJ2* | 0.5938 | -0.33681 | -0.44353 | -2.1264 | -0.06 |
| 15028 | *BPY2* | 0.11287 | -0.01371 | -0.28664 | -0.43226 | -0.06 |
| 15029 | *HMGCS2* | 1.252 | 0.18147 | -1.6213 | -1.9965 | -0.06 |
| 15030 | *CTDSPL2* | 0.55445 | 0.088102 | -0.83138 | -2.9144 | -0.06 |
| 15031 | *CHD7* | 2.1798 | -0.98314 | -1.386 | -1.433 | -0.06 |
| 15032 | *RPP14* | 0.08223 | 0.005713 | -0.27741 | -1.3136 | -0.06 |
| 15033 | *ENPEP* | 0.33952 | 0.041238 | -0.57134 | -0.94019 | -0.06 |
| 15034 | *COX6C* | 1.1086 | 0.10557 | -1.4048 | -1.9319 | -0.06 |
| 15035 | *CCL4L2* | 0.71476 | 0.42398 | -1.3294 | -1.7567 | -0.06 |

| | | | | | | |
|---|---|---|---|---|---|---|
| 15036 | *HMSD* | 0.87394 | -0.34751 | -0.71717 | -1.9686 | -0.06 |
| 15037 | *SH3GL2* | 0.86616 | -0.16781 | -0.8903 | -2.056 | -0.06 |
| 15038 | *RNF5* | 1.092 | -0.37282 | -0.91165 | -1.0231 | -0.06 |
| 15039 | *PRRT3* | 1.5666 | -0.25527 | -1.5045 | -1.6744 | -0.06 |
| 15040 | *ZNRF2* | 1.2796 | -0.62476 | -0.84816 | -1.0413 | -0.06 |
| 15041 | *PRR35* | 1.7432 | -0.60873 | -1.3282 | -4.0419 | -0.06 |
| 15042 | *MOGS* | 0.2824 | 0.067594 | -0.54373 | -1.6799 | -0.06 |
| 15043 | *OR13F1* | 0.59297 | 0.38037 | -1.1673 | -1.2573 | -0.06 |
| 15044 | *ANKIB1* | 1.5516 | -0.7329 | -1.0131 | -1.2075 | -0.06 |
| 15045 | *ABCC11* | 0.4315 | -0.06413 | -0.56185 | -1.0868 | -0.06 |
| 15046 | *GAS1* | 0.48879 | 0.22714 | -0.9135 | -2.4405 | -0.07 |
| 15047 | *PLEKHS1* | 0.7492 | 0.18854 | -1.1357 | -1.3517 | -0.07 |
| 15048 | *PRR21* | 0.71677 | -0.2694 | -0.64616 | -1.4162 | -0.07 |
| 15049 | *CTDSP2* | 0.31201 | -0.10258 | -0.40824 | -1.4557 | -0.07 |
| 15050 | *ZW10* | 1.9286 | -0.69377 | -1.434 | -3.2632 | -0.07 |
| 15051 | *SNCA* | 3.5226 | -1.4842 | -2.2392 | -2.6998 | -0.07 |
| 15052 | *RANBP1* | 0.91767 | -0.30377 | -0.81539 | -1.0838 | -0.07 |
| 15053 | *CX3CL1* | 0.38185 | -0.26851 | -0.3153 | -1.6597 | -0.07 |
| 15054 | *MRPS10* | 0.28412 | 0.22321 | -0.70976 | -0.86435 | -0.07 |
| 15055 | *C5orf51* | 0.34126 | 0.1049 | -0.64918 | -1.1043 | -0.07 |
| 15056 | *GRIN1* | 0.80251 | 0.26279 | -1.2694 | -1.6086 | -0.07 |
| 15057 | *PEX13* | 0.59762 | 0.29709 | -1.0992 | -1.6193 | -0.07 |
| 15058 | *MAP6* | 0.47543 | 0.18317 | -0.86358 | -1.5165 | -0.07 |
| 15059 | *CDC42EP4* | 0.89127 | -0.34481 | -0.75148 | -1.3961 | -0.07 |
| 15060 | *NUMBL* | 0.78709 | -0.39466 | -0.5975 | -2.262 | -0.07 |
| 15061 | *SNX17* | 0.26759 | 0.076527 | -0.54921 | -2.1868 | -0.07 |
| 15062 | *FAM171B* | 0.82613 | -0.32766 | -0.70361 | -0.87768 | -0.07 |
| 15063 | *CGNL1* | 0.31246 | 0.11308 | -0.63099 | -1.1811 | -0.07 |
| 15064 | *BLVRB* | 1.8761 | -0.6886 | -1.3935 | -1.4852 | -0.07 |
| 15065 | *UGT1A10* | 0.080441 | 0.053631 | -0.34036 | -0.60223 | -0.07 |
| 15066 | *FAM65C* | 1.5207 | -0.80613 | -0.92108 | -1.4711 | -0.07 |
| 15067 | *DMXL2* | 0.71448 | -0.24491 | -0.6761 | -1.8583 | -0.07 |
| 15068 | *GPR85* | 0.72458 | 0.5681 | -1.4997 | -2.0134 | -0.07 |
| 15069 | *GS1-259H13.2* | 2.7899 | -1.3445 | -1.6525 | -3.2901 | -0.07 |
| 15070 | *KIR2DL3* | 0.71252 | -0.44353 | -0.47676 | -1.0243 | -0.07 |
| 15071 | *ZNF664-FAM101A* | 1.1095 | -0.58732 | -0.73061 | -1.2291 | -0.07 |
| 15072 | *LOC388813* | 0.26768 | -0.07545 | -0.40136 | -3.1458 | -0.07 |
| 15073 | *ISLR2* | 0.43022 | 0.1422 | -0.78236 | -2.6376 | -0.07 |
| 15074 | *SLC22A4* | 0.58319 | -0.17641 | -0.61829 | -1.5121 | -0.07 |
| 15075 | *ACTR3C* | 0.5122 | -0.30811 | -0.41596 | -1.7829 | -0.07 |
| 15076 | *TMEM89* | 2.7382 | -1.4364 | -1.5138 | -1.6731 | -0.07 |
| 15077 | *LHFPL2* | 0.63165 | -0.34844 | -0.49541 | -1.4781 | -0.07 |
| 15078 | *TBC1D22A* | 0.1448 | 0.048765 | -0.40605 | -1.6976 | -0.07 |
| 15079 | *GOLGA5* | 0.43491 | -0.20988 | -0.43812 | -0.48862 | -0.07 |
| 15080 | *ZNF618* | 1.0547 | -0.0875 | -1.1812 | -1.9874 | -0.07 |
| 15081 | *NF1* | 1.0717 | 0.4782 | -1.765 | -1.8851 | -0.07 |

| | | | | | | |
|---|---|---|---|---|---|---|
| 15082 | *APLN* | 0.42648 | 0.38127 | -1.0231 | -3.0974 | -0.07 |
| 15083 | *PASD1* | 0.57355 | -0.35436 | -0.43481 | -1.0041 | -0.07 |
| 15084 | *LONP1* | 1.0512 | 0.32015 | -1.587 | -1.9751 | -0.07 |
| 15085 | *OR13G1* | 1.588 | -0.07719 | -1.7268 | -1.9623 | -0.07 |
| 15086 | *CTPS1* | 0.81708 | -0.11965 | -0.9135 | -2.1868 | -0.07 |
| 15087 | *ZNF234* | 0.1834 | 0.12753 | -0.52712 | -1.5261 | -0.07 |
| 15088 | *SLC39A10* | 0.43235 | -0.14762 | -0.50151 | -1.7599 | -0.07 |
| 15089 | *ACLY* | 2.6327 | -1.1566 | -1.6934 | -2.4268 | -0.07 |
| 15090 | *HS3ST3A1* | 0.57921 | 0.52817 | -1.325 | -2.4718 | -0.07 |
| 15091 | *NOG* | 0.21925 | -0.07913 | -0.35847 | -1.3052 | -0.07 |
| 15092 | *CARHSP1* | 1.2168 | -0.12145 | -1.3139 | -1.6617 | -0.07 |
| 15093 | *VTI1B* | 1.1106 | -0.52086 | -0.80847 | -1.1773 | -0.07 |
| 15094 | *OPN3* | 0.79239 | -0.47197 | -0.54048 | -1.8087 | -0.07 |
| 15095 | *CYP4Z1* | 2.3655 | -1.1825 | -1.4036 | -2.6539 | -0.07 |
| 15096 | *INTS2* | 0.68273 | 0.078551 | -0.98241 | -1.6836 | -0.07 |
| 15097 | *DCUN1D3* | 1.9489 | -0.84134 | -1.3288 | -1.4747 | -0.07 |
| 15098 | *SYTL3* | 0.99435 | -0.23596 | -0.97982 | -1.2628 | -0.07 |
| 15099 | *FCGR1B* | 0.35584 | 0.043375 | -0.62114 | -0.98026 | -0.07 |
| 15100 | *C14orf169* | 0.48777 | 0.001501 | -0.71134 | -2.0871 | -0.07 |
| 15101 | *ZNF599* | 0.56052 | 0.42287 | -1.2055 | -1.7103 | -0.07 |
| 15102 | *PLCG2* | 0.82075 | -0.23446 | -0.80863 | -1.879 | -0.07 |
| 15103 | *ZC3H7A* | 1.5201 | -0.69146 | -1.0528 | -1.6604 | -0.07 |
| 15104 | *UGP2* | 0.17582 | -0.19925 | -0.20112 | -1.0158 | -0.07 |
| 15105 | *C19orf48* | 1.5504 | -0.77241 | -1.0029 | -3.0112 | -0.07 |
| 15106 | *VPS37A* | 0.73177 | 0.58103 | -1.5379 | -1.7161 | -0.08 |
| 15107 | *TOPBP1* | 0.33184 | -0.01181 | -0.54515 | -1.4893 | -0.08 |
| 15108 | *APOL6* | 1.9126 | -0.4015 | -1.7364 | -2.1282 | -0.08 |
| 15109 | *ALG3* | 0.31889 | 0.052562 | -0.59886 | -0.9225 | -0.08 |
| 15110 | *DHPS* | 0.51482 | 0.084092 | -0.82656 | -1.7313 | -0.08 |
| 15111 | *ZBTB48* | 0.43842 | -0.04502 | -0.6218 | -1.1978 | -0.08 |
| 15112 | *GPR19* | 0.17052 | 0.13779 | -0.53924 | -0.75625 | -0.08 |
| 15113 | *TGIF2-C20orf24* | 0.41442 | 0.35438 | -0.9999 | -1.205 | -0.08 |
| 15114 | *GFRA4* | 2.4052 | -1.1866 | -1.4498 | -2.8941 | -0.08 |
| 15115 | *TSPYL4* | 0.76544 | 0.04383 | -1.0416 | -1.4428 | -0.08 |
| 15116 | *ZNF514* | 0.89005 | -0.16737 | -0.95614 | -1.1834 | -0.08 |
| 15117 | *OLFM2* | 0.60849 | -0.10743 | -0.73457 | -1.806 | -0.08 |
| 15118 | *RBM8A* | 0.62197 | -0.27961 | -0.57683 | -1.3507 | -0.08 |
| 15119 | *ADAM17* | 0.24949 | 0.003285 | -0.48862 | -1.6403 | -0.08 |
| 15120 | *GLRA3* | 0.29943 | -0.15141 | -0.38428 | -1.843 | -0.08 |
| 15121 | *RAD23A* | 0.065047 | -0.07885 | -0.22258 | -0.41395 | -0.08 |
| 15122 | *ATXN1* | 2.5729 | -1.0153 | -1.7948 | -1.8487 | -0.08 |
| 15123 | *ABCA5* | 1.826 | -0.9703 | -1.0933 | -1.4732 | -0.08 |
| 15124 | *SOWAHC* | 0.53039 | -0.07903 | -0.68934 | -2.5386 | -0.08 |
| 15125 | *F13B* | 0.79289 | -0.05643 | -0.97463 | -1.3559 | -0.08 |
| 15126 | *C21orf140* | 0.7952 | -0.50467 | -0.52949 | -1.0741 | -0.08 |
| 15127 | *BTBD17* | 0.49337 | 0.019393 | -0.75182 | -0.87482 | -0.08 |

| | | | | | | |
|---|---|---|---|---|---|---|
| 15128 | *GRM1* | 0.76313 | -0.4818 | -0.5204 | -0.70993 | -0.08 |
| 15129 | *EMC9* | 0.64312 | 0.43983 | -1.3232 | -2.3689 | -0.08 |
| 15130 | *GPR174* | 0.31574 | 0.12351 | -0.67962 | -1.2284 | -0.08 |
| 15131 | *BAIAP2L1* | 0.49197 | 0.3488 | -1.082 | -2.8723 | -0.08 |
| 15132 | *PRKCDBP* | 1.0174 | -0.34148 | -0.91757 | -2.2804 | -0.08 |
| 15133 | *HLA-DMB* | 0.93166 | -0.02022 | -1.1533 | -1.716 | -0.08 |
| 15134 | *RAB25* | 0.3295 | -0.10392 | -0.46765 | -0.6386 | -0.08 |
| 15135 | *ZFP36L1* | 0.69631 | 0.09276 | -1.0312 | -1.6047 | -0.08 |
| 15136 | *KIF1BP* | 1.9814 | -0.91307 | -1.3114 | -2.7042 | -0.08 |
| 15137 | *SNF8* | 0.66487 | -0.20592 | -0.70202 | -1.9318 | -0.08 |
| 15138 | *CDK13* | 0.61343 | 0.11542 | -0.97252 | -2.8111 | -0.08 |
| 15139 | *HEMGN* | 1.184 | -0.27555 | -1.1533 | -1.2684 | -0.08 |
| 15140 | *MAPK8IP3* | 0.74223 | 0.03422 | -1.0218 | -1.4885 | -0.08 |
| 15141 | *YIF1B* | 0.62371 | 0.071151 | -0.94121 | -1.224 | -0.08 |
| 15142 | *KLHL42* | 1.2645 | -0.68999 | -0.82098 | -1.8799 | -0.08 |
| 15143 | *FAM227A* | 0.91321 | -0.26409 | -0.89595 | -1.0984 | -0.08 |
| 15144 | *KIAA1958* | 0.1459 | -0.04444 | -0.35029 | -1.4043 | -0.08 |
| 15145 | *CFAP74* | 0.33377 | 0.23104 | -0.81496 | -2.2094 | -0.08 |
| 15146 | *IMMP2L* | 0.24438 | -0.19858 | -0.29727 | -2.0629 | -0.08 |
| 15147 | *TADA3* | 1.5118 | -0.19618 | -1.568 | -2.1848 | -0.08 |
| 15148 | *HOXC13* | 1.453 | -0.00817 | -1.6975 | -2.1802 | -0.08 |
| 15149 | *PRADC1* | 0.51587 | -0.28907 | -0.47987 | -1.6949 | -0.08 |
| 15150 | *RNF149* | 0.13046 | -0.08868 | -0.29494 | -0.68911 | -0.08 |
| 15151 | *CCDC142* | 1.0532 | 0.37076 | -1.6776 | -2.6999 | -0.08 |
| 15152 | *QTRTD1* | 0.68649 | -0.21484 | -0.72612 | -3.0687 | -0.08 |
| 15153 | *RAB3C* | 1.0283 | -0.62747 | -0.65654 | -1.7437 | -0.09 |
| 15154 | *CDH10* | 0.31864 | 0.29741 | -0.87208 | -1.1698 | -0.09 |
| 15155 | *PDXP* | 2.1212 | -0.73424 | -1.6431 | -1.8318 | -0.09 |
| 15156 | *CST9L* | 0.49579 | -0.06413 | -0.68781 | -1.7119 | -0.09 |
| 15157 | *HSP90AB1* | 0.3755 | 0.19223 | -0.82472 | -1.0323 | -0.09 |
| 15158 | *GNB5* | 0.95205 | -0.27453 | -0.93465 | -1.6836 | -0.09 |
| 15159 | *HIPK1* | 0.10767 | -0.01905 | -0.3464 | -0.95708 | -0.09 |
| 15160 | *DUOXA1* | 1.6469 | 0.22526 | -2.1305 | -2.28 | -0.09 |
| 15161 | *CAPZB* | 0.43295 | 0.40361 | -1.0952 | -1.6239 | -0.09 |
| 15162 | *SH3YL1* | 2.0036 | -1.0707 | -1.192 | -1.5601 | -0.09 |
| 15163 | *SYK* | 0.39081 | -0.04935 | -0.60065 | -2.0147 | -0.09 |
| 15164 | *CAPN7* | 0.55825 | 0.41247 | -1.2306 | -1.5881 | -0.09 |
| 15165 | *HNRNPK* | 1.0089 | -0.26469 | -1.0049 | -1.2797 | -0.09 |
| 15166 | *NRN1L* | 0.41008 | -0.07036 | -0.60083 | -0.66628 | -0.09 |
| 15167 | *OSBPL9* | 0.63969 | -0.44353 | -0.45742 | -2.3389 | -0.09 |
| 15168 | *TCEA1* | 0.56657 | -0.08868 | -0.73961 | -0.89752 | -0.09 |
| 15169 | *GREM1* | 1.596 | -0.59045 | -1.2681 | -1.9794 | -0.09 |
| 15170 | *PAFAH1B3* | 0.95208 | -0.36086 | -0.85389 | -1.0211 | -0.09 |
| 15171 | *ZBBX* | 2.4287 | -0.89052 | -1.8016 | -3.1751 | -0.09 |
| 15172 | *AOX1* | 0.078829 | 0.050668 | -0.39318 | -1.7073 | -0.09 |
| 15173 | *C2orf27A* | 1.423 | -0.59791 | -1.0889 | -1.6268 | -0.09 |

| | | | | | | |
|---|---|---|---|---|---|---|
| 15174 | *NAT16* | 1.0215 | -0.46348 | -0.82226 | -0.85285 | -0.09 |
| 15175 | *OR5V1* | 0.58843 | -0.33169 | -0.52154 | -1.5616 | -0.09 |
| 15176 | *EYA2* | 0.11836 | -0.15876 | -0.22674 | -0.98158 | -0.09 |
| 15177 | *SMCO4* | 1.3832 | -0.76076 | -0.8897 | -1.6368 | -0.09 |
| 15178 | *RNPC3* | 1.8236 | -0.75835 | -1.3326 | -2.1111 | -0.09 |
| 15179 | *PPIC* | 0.15122 | 0.093005 | -0.51182 | -2.4422 | -0.09 |
| 15180 | *RUVBL2* | 1.9634 | -1.0241 | -1.2079 | -1.8013 | -0.09 |
| 15181 | *CYP3A4* | 0.70993 | 0.1049 | -1.0838 | -1.72 | -0.09 |
| 15182 | *POTED* | 3.1004 | -1.0743 | -2.2951 | -2.5804 | -0.09 |
| 15183 | *COL14A1* | 1.3806 | -0.73708 | -0.91253 | -2.5535 | -0.09 |
| 15184 | *HMGN2* | 0.54846 | -0.00331 | -0.81522 | -1.9678 | -0.09 |
| 15185 | *AGA* | 1.8259 | -0.75724 | -1.3389 | -1.3966 | -0.09 |
| 15186 | *CTSK* | 2.5472 | -1.309 | -1.5085 | -1.7979 | -0.09 |
| 15187 | *UBR1* | 0.21727 | -0.02156 | -0.46636 | -1.2736 | -0.09 |
| 15188 | *DTNA* | 0.96714 | 0.27156 | -1.5095 | -1.5343 | -0.09 |
| 15189 | *CCRL2* | 0.062952 | -0.1515 | -0.18246 | -0.44353 | -0.09 |
| 15190 | *SMIM24* | 0.40372 | -0.13984 | -0.53526 | -1.3346 | -0.09 |
| 15191 | *PTGDS* | 1.0812 | -0.51204 | -0.84063 | -2.9068 | -0.09 |
| 15192 | *RPL27A* | 0.94952 | 0.54334 | -1.7647 | -2.0398 | -0.09 |
| 15193 | *MRPL36* | 0.90007 | 0.59146 | -1.7642 | -1.8853 | -0.09 |
| 15194 | *RNF6* | 0.70819 | 0.30691 | -1.2879 | -2.2452 | -0.09 |
| 15195 | *LCE2B* | 0.10006 | 0.007602 | -0.38096 | -1.6343 | -0.09 |
| 15196 | *TRPC5* | 2.0769 | -0.95775 | -1.3926 | -2.1085 | -0.09 |
| 15197 | *HGS* | 0.69084 | 0.2288 | -1.1932 | -1.7364 | -0.09 |
| 15198 | *FAM210A* | 0.46356 | -0.2748 | -0.46298 | -1.3478 | -0.09 |
| 15199 | *ZBTB8OS* | 1.7483 | -0.17081 | -1.8537 | -2.1354 | -0.09 |
| 15200 | *KLHL18* | 0.39732 | -0.0665 | -0.60717 | -1.2896 | -0.09 |
| 15201 | *NEK7* | 0.34535 | -0.17921 | -0.44353 | -1.0564 | -0.09 |
| 15202 | *ACAD9* | 0.53423 | 0.10217 | -0.91403 | -1.743 | -0.09 |
| 15203 | *CFAP126* | 0.54097 | -0.24973 | -0.56938 | -1.1759 | -0.09 |
| 15204 | *TBC1D16* | 0.71042 | -0.42808 | -0.56076 | -0.93802 | -0.09 |
| 15205 | *ZNF462* | 1.9314 | -0.96463 | -1.2457 | -2.8574 | -0.09 |
| 15206 | *EXOSC1* | 0.3573 | 0.25826 | -0.89518 | -0.92823 | -0.09 |
| 15207 | *NBPF14* | 0.19726 | 0.052791 | -0.53271 | -1.0435 | -0.09 |
| 15208 | *NFAT5* | 0.90405 | 0.25061 | -1.4375 | -1.9995 | -0.09 |
| 15209 | *ZCCHC9* | 3.4654 | -1.5797 | -2.1695 | -2.2543 | -0.09 |
| 15210 | *DDX55* | 0.18364 | 0.15787 | -0.62585 | -0.7342 | -0.09 |
| 15211 | *SNX19* | 1.3484 | -0.72488 | -0.90791 | -1.4998 | -0.09 |
| 15212 | *MUSK* | 0.70963 | -0.48659 | -0.50748 | -1.2017 | -0.09 |
| 15213 | *RAD23B* | 0.74049 | -0.43472 | -0.59099 | -1.9455 | -0.10 |
| 15214 | *PINK1* | 0.024417 | -0.02502 | -0.28497 | -1.8994 | -0.10 |
| 15215 | *ADGRE5* | 1.1734 | 0.002265 | -1.4614 | -2.0817 | -0.10 |
| 15216 | *RHOF* | 1.6136 | -0.02958 | -1.8698 | -1.9623 | -0.10 |
| 15217 | *GC* | 0.23307 | 0.15805 | -0.67731 | -0.72798 | -0.10 |
| 15218 | *FKBP14* | 1.0233 | -0.00446 | -1.3056 | -1.7241 | -0.10 |
| 15219 | *C3orf80* | 0.2249 | 0.16561 | -0.67791 | -1.4823 | -0.10 |

| | | | | | | |
|---|---|---|---|---|---|---|
| 15220 | *ZC3H4* | 0.91458 | -0.09982 | -1.1027 | -1.4715 | -0.10 |
| 15221 | *NPAP1* | 0.33526 | -0.27315 | -0.35052 | -0.91757 | -0.10 |
| 15222 | *CREG2* | 2.6911 | -1.2474 | -1.7322 | -1.7466 | -0.10 |
| 15223 | *SPDYE6* | 0.41993 | 0.13235 | -0.84177 | -0.92326 | -0.10 |
| 15224 | *ANKS3* | 2.5618 | -1.2169 | -1.6356 | -2.0337 | -0.10 |
| 15225 | *NRF1* | 2.7461 | -1.4568 | -1.5815 | -2.5008 | -0.10 |
| 15226 | *WDR37* | 1.6636 | -0.49166 | -1.4644 | -2.914 | -0.10 |
| 15227 | *AMELY* | 0.40077 | 0.011133 | -0.70546 | -0.82696 | -0.10 |
| 15228 | *TNFRSF17* | 0.085374 | 0.017406 | -0.39664 | -0.62803 | -0.10 |
| 15229 | *LANCL2* | 1.2626 | -0.53406 | -1.0231 | -1.613 | -0.10 |
| 15230 | *STRA6* | 0.45954 | 0.20713 | -0.96144 | -1.3475 | -0.10 |
| 15231 | *NAGLU* | 0.20905 | 0.16894 | -0.67313 | -1.4401 | -0.10 |
| 15232 | *ANKK1* | 0.72077 | 0.20086 | -1.2171 | -2.0121 | -0.10 |
| 15233 | *C6orf201* | 0.7127 | -0.29114 | -0.71717 | -0.84151 | -0.10 |
| 15234 | *NUP62* | 0.69539 | -0.01706 | -0.97433 | -1.5294 | -0.10 |
| 15235 | *NSMCE4A* | 0.65056 | 0.52738 | -1.4741 | -1.7364 | -0.10 |
| 15236 | *TOX* | 0.89211 | -0.45847 | -0.73023 | -0.78793 | -0.10 |
| 15237 | *CXorf49* | 0.45861 | 0.017346 | -0.77255 | -0.84542 | -0.10 |
| 15238 | *AFMID* | 1.1816 | 0.48372 | -1.9623 | -3.0312 | -0.10 |
| 15239 | *SP140* | 0.96268 | 0.50897 | -1.7709 | -2.224 | -0.10 |
| 15240 | *HACD2* | 0.41241 | 0.093599 | -0.80594 | -1.6759 | -0.10 |
| 15241 | *CACNA1D* | 0.62376 | 0.49941 | -1.4233 | -1.4864 | -0.10 |
| 15242 | *DSC2* | 2.6618 | -1.2775 | -1.6847 | -2.0126 | -0.10 |
| 15243 | *MEPCE* | 2.2164 | -0.93878 | -1.5781 | -2.5007 | -0.10 |
| 15244 | *SIM1* | 0.075215 | -0.06007 | -0.31641 | -1.9305 | -0.10 |
| 15245 | *ANKRD36B* | 0.49189 | -0.03221 | -0.76106 | -0.91415 | -0.10 |
| 15246 | *NKX2-1* | 0.52028 | -0.29614 | -0.52587 | -1.5971 | -0.10 |
| 15247 | *PDE1A* | 0.66958 | 0.1827 | -1.1545 | -1.94 | -0.10 |
| 15248 | *MTMR1* | 1.1412 | 0.17324 | -1.6176 | -2.5298 | -0.10 |
| 15249 | *LRRC14* | 0.61166 | 0.27571 | -1.191 | -1.4364 | -0.10 |
| 15250 | *MBD3L4* | 1.107 | 0.083386 | -1.4944 | -2.095 | -0.10 |
| 15251 | *RAB11FIP5* | 1.579 | 0.048175 | -1.9319 | -1.9579 | -0.10 |
| 15252 | *SLC35C2* | 0.061411 | -0.16752 | -0.19947 | -1.0909 | -0.10 |
| 15253 | *TMEM147* | 1.9599 | -0.45979 | -1.8057 | -1.9474 | -0.10 |
| 15254 | *RPAP1* | 0.60884 | 0.59437 | -1.5095 | -2.3958 | -0.10 |
| 15255 | *VGLL4* | 0.22608 | -0.21327 | -0.31981 | -0.73362 | -0.10 |
| 15256 | *RAPSN* | 0.71302 | -0.11567 | -0.9059 | -2.3449 | -0.10 |
| 15257 | *SPATA8* | 0.45695 | 0.31468 | -1.0802 | -2.1709 | -0.10 |
| 15258 | *SSC5D* | 2.2413 | -0.74705 | -1.8033 | -2.5442 | -0.10 |
| 15259 | *ZNF324B* | 0.18753 | 0.075215 | -0.57273 | -1.5294 | -0.10 |
| 15260 | *POU5F2* | 0.30878 | -0.2604 | -0.35892 | -0.37858 | -0.10 |
| 15261 | *OAF* | 0.19547 | -0.1829 | -0.32328 | -1.7149 | -0.10 |
| 15262 | *ASPHD2* | 0.89785 | 0.18099 | -1.3903 | -1.6401 | -0.10 |
| 15263 | *MAN2C1* | 0.11922 | -0.05125 | -0.3808 | -0.95753 | -0.10 |
| 15264 | *TUSC2* | 0.55916 | -0.4057 | -0.46689 | -2.5513 | -0.10 |
| 15265 | *SRRT* | 2.9904 | -1.4154 | -1.8885 | -1.9623 | -0.10 |

| | | | | | | |
|---|---|---|---|---|---|---|
| 15266 | *CREB3L4* | 0.10724 | -0.04753 | -0.37384 | -1.1888 | -0.10 |
| 15267 | *KRTAP21-1* | 0.2761 | 0.14487 | -0.73577 | -0.78963 | -0.10 |
| 15268 | *STK36* | 0.78393 | -0.40098 | -0.69867 | -1.3162 | -0.11 |
| 15269 | *NUFIP2* | 0.74835 | 0.2288 | -1.2935 | -1.3416 | -0.11 |
| 15270 | *ACY1* | 1.6081 | -0.95154 | -0.9733 | -1.1807 | -0.11 |
| 15271 | *ZNF334* | 1.0702 | -0.30899 | -1.078 | -1.144 | -0.11 |
| 15272 | *LIPG* | 0.30791 | 0.052475 | -0.67801 | -1.9833 | -0.11 |
| 15273 | *DNPH1* | 0.56123 | 0.020985 | -0.90014 | -0.98574 | -0.11 |
| 15274 | *RIPK3* | 0.70641 | -0.12189 | -0.9027 | -1.3057 | -0.11 |
| 15275 | *PSMF1* | 0.29114 | 0.22896 | -0.83851 | -1.9186 | -0.11 |
| 15276 | *ZNF319* | 0.39098 | -0.14449 | -0.56564 | -1.8157 | -0.11 |
| 15277 | *SIRT2* | 0.36116 | -0.19889 | -0.48156 | -2.6046 | -0.11 |
| 15278 | *FLCN* | 0.51897 | 0.36776 | -1.2071 | -1.8816 | -0.11 |
| 15279 | *PPAN-P2RY11* | 0.40355 | -0.06308 | -0.66146 | -2.1068 | -0.11 |
| 15280 | *DMBT1* | 0.24352 | 0.018487 | -0.5834 | -0.8397 | -0.11 |
| 15281 | *LPIN1* | 0.061345 | -0.01742 | -0.36583 | -1.1379 | -0.11 |
| 15282 | *PNPLA8* | 0.24289 | -0.168 | -0.39731 | -0.43305 | -0.11 |
| 15283 | *DNAH12* | 0.44033 | -0.15145 | -0.61138 | -2.6911 | -0.11 |
| 15284 | *NPEPPS* | 0.86484 | -0.33126 | -0.8562 | -1.3924 | -0.11 |
| 15285 | *LRRC8E* | 0.69114 | 0.51028 | -1.5246 | -1.7576 | -0.11 |
| 15286 | *C14orf39* | 0.3417 | -0.22645 | -0.43886 | -1.0099 | -0.11 |
| 15287 | *FAM8A1* | 2.1471 | -1.1689 | -1.3023 | -2.0465 | -0.11 |
| 15288 | *KRT18* | 2.339 | -1.078 | -1.5855 | -1.9792 | -0.11 |
| 15289 | *PPP1R8* | 1.4575 | -0.88055 | -0.90233 | -1.3145 | -0.11 |
| 15290 | *CNGA1* | 0.92282 | 0.26871 | -1.5171 | -2.5434 | -0.11 |
| 15291 | *XRCC1* | 0.19182 | 0.051278 | -0.56873 | -0.65434 | -0.11 |
| 15292 | *GREB1L* | 0.76492 | 0.059436 | -1.1511 | -1.6552 | -0.11 |
| 15293 | *PTPRJ* | 0.40848 | -0.18568 | -0.5497 | -2.5617 | -0.11 |
| 15294 | *OR5B17* | 2.1145 | -0.46303 | -1.9789 | -2.0818 | -0.11 |
| 15295 | *USP17L4* | 0.42304 | -0.33619 | -0.41469 | -1.9123 | -0.11 |
| 15296 | *CDK9* | 0.78179 | -0.44353 | -0.66652 | -2.2343 | -0.11 |
| 15297 | *PLEKHB2* | 0.79193 | -0.06028 | -1.0606 | -1.297 | -0.11 |
| 15298 | *KRTAP19-2* | 0.2274 | -0.04238 | -0.51435 | -0.78842 | -0.11 |
| 15299 | *TBC1D29* | 0.16366 | -0.05094 | -0.44353 | -0.76942 | -0.11 |
| 15300 | *PSAP* | 0.55047 | 0.20718 | -1.0889 | -2.017 | -0.11 |
| 15301 | *RAD17* | 2.094 | -0.79043 | -1.6356 | -1.6906 | -0.11 |
| 15302 | *PTRF* | 0.29928 | -0.2842 | -0.34964 | -0.86986 | -0.11 |
| 15303 | *EFNB2* | 1.3568 | 0.006098 | -1.6975 | -2.7568 | -0.11 |
| 15304 | *ZNF747* | 0.50614 | -0.3565 | -0.48524 | -0.68889 | -0.11 |
| 15305 | *ERI2* | 0.31304 | 0.034352 | -0.68586 | -1.2955 | -0.11 |
| 15306 | *DNAJC14* | 0.085857 | -0.17781 | -0.24723 | -0.76505 | -0.11 |
| 15307 | *MAPK8IP2* | 1.1841 | -0.59903 | -0.92427 | -1.0248 | -0.11 |
| 15308 | *ERH* | 0.71036 | -0.02563 | -1.0243 | -1.589 | -0.11 |
| 15309 | *TMEM50A* | 0.36129 | -0.15141 | -0.55002 | -1.1082 | -0.11 |
| 15310 | *HES4* | 0.99523 | -0.27166 | -1.0667 | -1.9375 | -0.11 |
| 15311 | *ARHGAP42* | 3.1423 | -1.5243 | -1.9623 | -2.1477 | -0.11 |

| | | | | | | |
|---|---|---|---|---|---|---|
| 15312 | *PIANP* | 0.94457 | -0.5788 | -0.71032 | -1.3268 | -0.11 |
| 15313 | *SREBF1* | 0.86148 | -0.30843 | -0.89892 | -1.5307 | -0.12 |
| 15314 | *TLN1* | 0.47735 | -0.18382 | -0.6399 | -1.7512 | -0.12 |
| 15315 | *OR10A3* | 0.97181 | -0.00344 | -1.3148 | -1.5082 | -0.12 |
| 15316 | *KLK6* | 0.24303 | -0.19753 | -0.39209 | -0.96161 | -0.12 |
| 15317 | *TRAF1* | 0.7847 | -0.15394 | -0.97776 | -1.1802 | -0.12 |
| 15318 | *TMEM30B* | 0.50115 | -0.3589 | -0.48973 | -1.4248 | -0.12 |
| 15319 | *CYP27A1* | 0.19864 | 0.15359 | -0.70106 | -1.6208 | -0.12 |
| 15320 | *POTEJ* | 0.98549 | 0.1182 | -1.4527 | -2.1364 | -0.12 |
| 15321 | *MMP1* | 0.083498 | 0.014096 | -0.44766 | -1.4224 | -0.12 |
| 15322 | *DEAF1* | 0.56256 | 0.40217 | -1.3148 | -2.0613 | -0.12 |
| 15323 | *PEX2* | 0.8723 | -0.59544 | -0.62708 | -1.067 | -0.12 |
| 15324 | *CHCHD10* | 1.019 | 0.003476 | -1.3729 | -1.6302 | -0.12 |
| 15325 | *AOC1* | 1.6627 | -0.65163 | -1.3617 | -1.7235 | -0.12 |
| 15326 | *BCL2L1* | 0.99875 | 0.41475 | -1.7642 | -1.8698 | -0.12 |
| 15327 | *USP18* | 0.4357 | -0.15313 | -0.63341 | -1.111 | -0.12 |
| 15328 | *SCGB2B2* | 2.373 | -1.031 | -1.6935 | -1.8157 | -0.12 |
| 15329 | *CHRNA3* | 0.54063 | 0.27355 | -1.1659 | -1.433 | -0.12 |
| 15330 | *CCDC170* | 1.7113 | -0.52757 | -1.5369 | -1.6193 | -0.12 |
| 15331 | *KCNK12* | 0.50562 | 0.47672 | -1.3358 | -2.1198 | -0.12 |
| 15332 | *SSB* | 2.4179 | -1.3146 | -1.4568 | -1.6044 | -0.12 |
| 15333 | *TMEM230* | 2.4837 | -0.94759 | -1.8909 | -2.1477 | -0.12 |
| 15334 | *NANOGNB* | 2.6648 | -1.3047 | -1.7163 | -1.8203 | -0.12 |
| 15335 | *TTC21B* | 1.1616 | -0.66107 | -0.85739 | -1.639 | -0.12 |
| 15336 | *LRTM2* | 1.4188 | -0.61199 | -1.1637 | -1.2284 | -0.12 |
| 15337 | *FBXO47* | 0.7843 | 0.24723 | -1.3892 | -2.5571 | -0.12 |
| 15338 | *ILF3* | 0.5102 | 0.016015 | -0.88391 | -1.0266 | -0.12 |
| 15339 | *KRTAP6-3* | 0.46439 | 0.3065 | -1.1287 | -1.8742 | -0.12 |
| 15340 | *KRT73* | 1.4889 | -0.80311 | -1.0443 | -1.5343 | -0.12 |
| 15341 | *AGL* | 0.37448 | -0.27911 | -0.45399 | -1.0643 | -0.12 |
| 15342 | *FADD* | 0.27688 | -0.00431 | -0.63198 | -0.70944 | -0.12 |
| 15343 | *ZNF23* | 0.32946 | -0.30832 | -0.3828 | -2.0995 | -0.12 |
| 15344 | *AJUBA* | 0.43755 | -0.20104 | -0.59842 | -1.4981 | -0.12 |
| 15345 | *CKMT1A* | 0.74116 | -0.25723 | -0.84626 | -1.0564 | -0.12 |
| 15346 | *RAD51C* | 0.54615 | -0.34793 | -0.56218 | -0.95682 | -0.12 |
| 15347 | *C4orf22* | 1.9465 | -0.36835 | -1.9436 | -2.3405 | -0.12 |
| 15348 | *FAM46C* | 0.28622 | 0.14946 | -0.80168 | -1.4841 | -0.12 |
| 15349 | *CYP17A1* | 0.4126 | -0.37417 | -0.40453 | -1.1585 | -0.12 |
| 15350 | *TEX43* | 0.77063 | -0.29713 | -0.84001 | -2.0553 | -0.12 |
| 15351 | *SPTLC2* | 0.51516 | -0.43841 | -0.44353 | -1.0769 | -0.12 |
| 15352 | *RTFDC1* | 0.61924 | 0.48251 | -1.4689 | -1.536 | -0.12 |
| 15353 | *MT1H* | 0.3755 | -0.0649 | -0.67776 | -1.6334 | -0.12 |
| 15354 | *BRD3* | 1.1314 | 0.30802 | -1.8066 | -2.3689 | -0.12 |
| 15355 | *SLC5A2* | 0.81856 | -0.13757 | -1.0482 | -1.3591 | -0.12 |
| 15356 | *RC3H1* | 0.32919 | -0.33121 | -0.36536 | -2.0553 | -0.12 |
| 15357 | *MITF* | -0.02395 | -0.09635 | -0.24715 | -2.2691 | -0.12 |

| | | | | | | |
|---|---|---|---|---|---|---|
| 15358 | *CD70* | 0.65417 | -0.27088 | -0.75094 | -0.98102 | -0.12 |
| 15359 | *LCE1B* | 0.14463 | 0.10141 | -0.614 | -1.0158 | -0.12 |
| 15360 | *MMP11* | 0.26699 | -0.26099 | -0.37429 | -0.89218 | -0.12 |
| 15361 | *SRPRB* | 0.24205 | 0.044461 | -0.65503 | -1.3965 | -0.12 |
| 15362 | *ZNF165* | 1.2101 | -0.75876 | -0.8213 | -1.8623 | -0.12 |
| 15363 | *IREB2* | 0.28096 | 0.004997 | -0.65606 | -1.7365 | -0.12 |
| 15364 | *TSHB* | 1.0595 | -0.65553 | -0.7757 | -2.2172 | -0.12 |
| 15365 | *SMIM7* | -0.04795 | -0.10133 | -0.22361 | -1.175 | -0.12 |
| 15366 | *COL27A1* | 0.41286 | -0.34665 | -0.43951 | -1.6288 | -0.12 |
| 15367 | *RPL23* | 1.3095 | -0.44353 | -1.2393 | -1.3664 | -0.12 |
| 15368 | *AMPD1* | 2.8987 | -1.4566 | -1.8157 | -3.6506 | -0.12 |
| 15369 | *ZNF407* | 1.8173 | -0.75625 | -1.4361 | -1.7422 | -0.13 |
| 15370 | *YEATS2* | 0.29432 | 0.15264 | -0.82208 | -1.5707 | -0.13 |
| 15371 | *WFDC10B* | 0.85564 | -0.43544 | -0.79535 | -1.6389 | -0.13 |
| 15372 | *UBQLN2* | 1.1306 | -0.62306 | -0.88272 | -1.773 | -0.13 |
| 15373 | *BANF1* | 0.98249 | -0.50093 | -0.85698 | -1.0426 | -0.13 |
| 15374 | *POU3F1* | 0.086266 | -0.10142 | -0.36086 | -0.45185 | -0.13 |
| 15375 | *DVL3* | 0.77002 | 0.57736 | -1.7235 | -1.8073 | -0.13 |
| 15376 | *CEBPE* | 0.56164 | 0.18233 | -1.1209 | -1.5121 | -0.13 |
| 15377 | *SERAC1* | 1.6432 | -0.7679 | -1.2525 | -2.3491 | -0.13 |
| 15378 | *FOXP2* | 0.48391 | -0.02883 | -0.83276 | -2.5535 | -0.13 |
| 15379 | *KIAA1024L* | 1.1429 | -0.38222 | -1.1384 | -1.5491 | -0.13 |
| 15380 | *MRPS34* | 0.72204 | 0.081909 | -1.1817 | -2.3405 | -0.13 |
| 15381 | *GRPEL1* | 0.013619 | -0.0795 | -0.31199 | -0.64313 | -0.13 |
| 15382 | *ADIPOQ* | 0.55507 | 0.25861 | -1.1921 | -1.6893 | -0.13 |
| 15383 | *MMS19* | 0.45173 | -0.17588 | -0.65476 | -2.2712 | -0.13 |
| 15384 | *CT47A2* | -0.00823 | -0.11251 | -0.25832 | -5.6109 | -0.13 |
| 15385 | *RFPL3* | 0.71115 | 0.12163 | -1.2119 | -2.1859 | -0.13 |
| 15386 | *FUT10* | 0.31071 | 0.22274 | -0.91269 | -1.8253 | -0.13 |
| 15387 | *CIAPIN1* | 0.76 | -0.49778 | -0.6424 | -1.8909 | -0.13 |
| 15388 | *HTR1B* | 0.77873 | 0.33951 | -1.4994 | -2.2044 | -0.13 |
| 15389 | *ALDH1L2* | 2.1367 | -1.0733 | -1.4448 | -2.6498 | -0.13 |
| 15390 | *RBMS1* | 0.34893 | 0.032955 | -0.76366 | -1.7316 | -0.13 |
| 15391 | *ALDOC* | 0.19795 | 0.090843 | -0.67071 | -1.5263 | -0.13 |
| 15392 | *LIMK2* | 0.72974 | -0.18347 | -0.92823 | -1.124 | -0.13 |
| 15393 | *LGALS2* | 1.0713 | -0.33638 | -1.1169 | -1.6457 | -0.13 |
| 15394 | *RCBTB2* | 0.59709 | 0.19829 | -1.1774 | -2.0652 | -0.13 |
| 15395 | *CRCP* | 0.33789 | 0.079397 | -0.80047 | -1.3899 | -0.13 |
| 15396 | *CHTOP* | 0.53741 | 0.26496 | -1.1856 | -1.3722 | -0.13 |
| 15397 | *ZNF229* | 0.82921 | -0.33824 | -0.875 | -1.4523 | -0.13 |
| 15398 | *FETUB* | 2.9215 | -1.0699 | -2.2357 | -3.6181 | -0.13 |
| 15399 | *CLDN8* | 0.86122 | -0.19671 | -1.049 | -1.5544 | -0.13 |
| 15400 | *SHOX2* | 1.6664 | -0.56444 | -1.4878 | -2.4699 | -0.13 |
| 15401 | *LRRC1* | 0.34051 | -0.16518 | -0.56251 | -1.3892 | -0.13 |
| 15402 | *CTSD* | 0.17335 | -0.16781 | -0.39457 | -0.43329 | -0.13 |
| 15403 | *DNAJB3* | 0.061507 | -0.14097 | -0.30975 | -0.66616 | -0.13 |

| | | | | | | |
|---|---|---|---|---|---|---|
| 15404 | *HNRNPA3* | 0.52403 | -0.44444 | -0.46918 | -1.2975 | -0.13 |
| 15405 | *REL* | 0.41716 | 0.40295 | -1.2098 | -1.4452 | -0.13 |
| 15406 | *UFL1* | 2.308 | -1.3364 | -1.3627 | -1.5077 | -0.13 |
| 15407 | *ICK* | 2.7754 | -1.1021 | -2.0645 | -3.2454 | -0.13 |
| 15408 | *PCNP* | 1.0788 | -0.71553 | -0.75448 | -1.4584 | -0.13 |
| 15409 | *POLG2* | 0.3946 | -0.1549 | -0.63152 | -1.5658 | -0.13 |
| 15410 | *DNHD1* | 0.48692 | 0.050441 | -0.93099 | -1.729 | -0.13 |
| 15411 | *FBXO11* | 0.13158 | -0.14117 | -0.38473 | -1.9292 | -0.13 |
| 15412 | *SLC12A6* | 0.23148 | -0.1386 | -0.48725 | -1.1243 | -0.13 |
| 15413 | *RSPO2* | -0.02176 | -0.05784 | -0.31487 | -1.455 | -0.13 |
| 15414 | *ZNF207* | 0.097241 | -0.00527 | -0.48659 | -0.6122 | -0.13 |
| 15415 | *SAYSD1* | 0.094966 | -0.22533 | -0.26508 | -2.0197 | -0.13 |
| 15416 | *ENDOU* | 0.093921 | -0.13436 | -0.35589 | -2.2732 | -0.13 |
| 15417 | *FAM171A2* | 0.2372 | 0.21972 | -0.85387 | -1.7282 | -0.13 |
| 15418 | *FLT1* | 0.94736 | 0.86236 | -2.2078 | -2.2223 | -0.13 |
| 15419 | *HAUS5* | 0.30573 | -0.23766 | -0.46631 | -1.3802 | -0.13 |
| 15420 | *CHCHD1* | -0.00833 | -0.13677 | -0.2562 | -0.48617 | -0.13 |
| 15421 | *RASGEF1A* | 0.68371 | -0.50332 | -0.58183 | -1.0315 | -0.13 |
| 15422 | *ACAP1* | 1.4447 | -0.28335 | -1.5634 | -1.9624 | -0.13 |
| 15423 | *KRTAP2-3* | 0.10994 | -0.19948 | -0.31492 | -0.72227 | -0.13 |
| 15424 | *SEPT5* | 0.70479 | 0.11742 | -1.227 | -1.8238 | -0.13 |
| 15425 | *TCF7* | 0.90314 | -0.45563 | -0.85236 | -0.97138 | -0.13 |
| 15426 | *MRPS28* | 0.3011 | -0.20454 | -0.50201 | -1.4801 | -0.14 |
| 15427 | *PLBD2* | 0.43087 | 0.20038 | -1.0369 | -1.3593 | -0.14 |
| 15428 | *C3orf84* | 0.16225 | -0.15992 | -0.40816 | -2.599 | -0.14 |
| 15429 | *FERMT3* | 0.47913 | 0.16922 | -1.0564 | -1.2352 | -0.14 |
| 15430 | *NT5C3B* | 0.063795 | -0.23528 | -0.2367 | -1.3363 | -0.14 |
| 15431 | *EN1* | 0.94502 | 0.16131 | -1.5149 | -1.9404 | -0.14 |
| 15432 | *RNF145* | 1.9163 | -0.79987 | -1.525 | -1.9534 | -0.14 |
| 15433 | *PPP2R3A* | 0.60079 | 0.081831 | -1.0912 | -2.3154 | -0.14 |
| 15434 | *CLEC7A* | 0.92018 | -0.45544 | -0.87418 | -1.2165 | -0.14 |
| 15435 | *MTERF1* | 0.42151 | -0.35735 | -0.47381 | -2.5513 | -0.14 |
| 15436 | *PRSS56* | 0.35665 | 0.10557 | -0.87198 | -1.0809 | -0.14 |
| 15437 | *RAB44* | 2.3989 | -1.3982 | -1.4108 | -1.4442 | -0.14 |
| 15438 | *TVP23A* | 1.0873 | 0.22121 | -1.7188 | -1.9175 | -0.14 |
| 15439 | *WIBG* | 0.20224 | 0.021388 | -0.63436 | -1.9659 | -0.14 |
| 15440 | *SRP72* | 0.73703 | 0.14429 | -1.2922 | -1.5352 | -0.14 |
| 15441 | *CRACR2B* | 0.25604 | 0.070178 | -0.73853 | -1.191 | -0.14 |
| 15442 | *NAA10* | 0.27132 | -0.13922 | -0.54555 | -1.9552 | -0.14 |
| 15443 | *CTCFL* | 0.32563 | -0.03585 | -0.70369 | -0.95064 | -0.14 |
| 15444 | *RFX8* | 0.8072 | -0.55437 | -0.66684 | -1.6779 | -0.14 |
| 15445 | *DNAJC17* | 1.471 | -0.51562 | -1.3699 | -1.775 | -0.14 |
| 15446 | *NKPD1* | 1.5054 | -0.80727 | -1.1138 | -1.307 | -0.14 |
| 15447 | *ZNF337* | 0.27564 | 0.21413 | -0.90606 | -1.088 | -0.14 |
| 15448 | *HPD* | 0.26341 | -0.03058 | -0.64932 | -0.91343 | -0.14 |
| 15449 | *UBXN2A* | 1.6359 | -0.96675 | -1.0871 | -1.8443 | -0.14 |

| | | | | | | |
|---|---|---|---|---|---|---|
| 15450 | *TMEM98* | 0.44457 | -0.02451 | -0.83912 | -1.0871 | -0.14 |
| 15451 | *NAB1* | 0.44278 | 0.32102 | -1.183 | -1.3666 | -0.14 |
| 15452 | *CILP* | 0.28767 | 0.16498 | -0.87198 | -1.2912 | -0.14 |
| 15453 | *PAPLN* | 0.65684 | -0.14663 | -0.93024 | -1.6648 | -0.14 |
| 15454 | *SNRK* | 1.4079 | -0.76355 | -1.0651 | -1.5458 | -0.14 |
| 15455 | *SSBP1* | 0.60053 | -0.44353 | -0.57899 | -0.73565 | -0.14 |
| 15456 | *STIM1* | 0.70504 | -0.08424 | -1.0433 | -1.388 | -0.14 |
| 15457 | *TRIM29* | 0.88514 | -0.33588 | -0.9728 | -1.6401 | -0.14 |
| 15458 | *LPA* | 0.94839 | 0.51992 | -1.892 | -1.9112 | -0.14 |
| 15459 | *SRPX2* | 2.2597 | -0.61249 | -2.071 | -2.5934 | -0.14 |
| 15460 | *FAHD2A* | 0.73277 | -0.39885 | -0.75778 | -1.3132 | -0.14 |
| 15461 | *ITIH4* | 0.31367 | -0.15504 | -0.58267 | -0.95242 | -0.14 |
| 15462 | *SOCS5* | 0.772 | -0.51931 | -0.67764 | -0.82368 | -0.14 |
| 15463 | *SARNP* | 0.53831 | -0.45563 | -0.50775 | -0.94042 | -0.14 |
| 15464 | *LOC100507462* | 0.56846 | 0.25454 | -1.2496 | -2.4205 | -0.14 |
| 15465 | *IGSF23* | 0.004997 | -0.0679 | -0.36412 | -2.4052 | -0.14 |
| 15466 | *OPN5* | 1.0881 | -0.74207 | -0.77439 | -0.78672 | -0.14 |
| 15467 | *BAD* | 2.9448 | -1.6086 | -1.765 | -1.94 | -0.14 |
| 15468 | *PRPF4* | 0.81223 | -0.50324 | -0.73814 | -1.9619 | -0.14 |
| 15469 | *RET* | 1.4872 | 0.057026 | -1.9734 | -2.0852 | -0.14 |
| 15470 | *TMEM86A* | 1.3244 | -0.62947 | -1.1244 | -1.2632 | -0.14 |
| 15471 | *SLC38A4* | 0.61937 | 0.23127 | -1.2817 | -1.7577 | -0.14 |
| 15472 | *RPN1* | 2.0828 | -1.2007 | -1.3132 | -1.7437 | -0.14 |
| 15473 | *SPATA5* | 2.0248 | -1.0288 | -1.4271 | -2.9544 | -0.14 |
| 15474 | *TMEM30A* | 0.17147 | -0.11621 | -0.48652 | -0.72798 | -0.14 |
| 15475 | *MFSD6* | 0.69766 | -0.55409 | -0.57506 | -1.3773 | -0.14 |
| 15476 | *PTPRQ* | 0.31119 | 0.12964 | -0.87332 | -1.6023 | -0.14 |
| 15477 | *GLCCI1* | 0.38728 | -0.35604 | -0.46399 | -0.48659 | -0.14 |
| 15478 | *ICAM5* | 0.94799 | -0.43005 | -0.9521 | -2.3118 | -0.14 |
| 15479 | *GHRL* | 0.41043 | 0.10658 | -0.9519 | -1.9404 | -0.14 |
| 15480 | *NAP1L3* | 0.9578 | -0.60701 | -0.78651 | -1.4998 | -0.15 |
| 15481 | *MFSD3* | 0.75829 | -0.0205 | -1.1739 | -1.3851 | -0.15 |
| 15482 | *XAGE2* | 1.7771 | -0.70442 | -1.5099 | -2.0197 | -0.15 |
| 15483 | *TM7SF2* | 2.618 | -1.3966 | -1.6604 | -1.912 | -0.15 |
| 15484 | *TAGLN3* | 2.1068 | -1.0293 | -1.5167 | -1.7323 | -0.15 |
| 15485 | *MAML1* | 0.91939 | -0.58985 | -0.76985 | -1.7829 | -0.15 |
| 15486 | *RNPS1* | 1.5077 | -0.78194 | -1.1671 | -1.8585 | -0.15 |
| 15487 | *NELFB* | 0.28792 | 0.21028 | -0.93969 | -1.4076 | -0.15 |
| 15488 | *KDM6A* | 0.88215 | -0.05139 | -1.2736 | -2.0952 | -0.15 |
| 15489 | *SHF* | 0.68754 | -0.02974 | -1.1008 | -1.2127 | -0.15 |
| 15490 | *PLK3* | 1.0205 | -0.30618 | -1.1576 | -1.4633 | -0.15 |
| 15491 | *DAP* | 1.0792 | 0.13006 | -1.653 | -2.829 | -0.15 |
| 15492 | *DMRTB1* | 2.2814 | -0.73511 | -1.9912 | -3.1229 | -0.15 |
| 15493 | *FCRL6* | 0.72543 | 0.59519 | -1.7658 | -1.9713 | -0.15 |
| 15494 | *KCNK4* | 0.46876 | -0.02044 | -0.89351 | -2.3014 | -0.15 |
| 15495 | *BPIFB4* | 0.24633 | -0.18528 | -0.50731 | -1.4946 | -0.15 |

| | | | | | | |
|---|---|---|---|---|---|---|
| 15496 | *DNM2* | 1.5761 | -0.79596 | -1.2276 | -2.5361 | -0.15 |
| 15497 | *OR10A7* | 0.88398 | 0.85388 | -2.1859 | -2.6543 | -0.15 |
| 15498 | *KIF18A* | 1.2654 | 0.28076 | -1.9945 | -2.2059 | -0.15 |
| 15499 | *JUN* | 0.36957 | -0.20207 | -0.61599 | -0.90663 | -0.15 |
| 15500 | *SPATA2* | 0.5094 | -0.34535 | -0.61261 | -0.64397 | -0.15 |
| 15501 | *ZNF555* | 0.89861 | -0.01439 | -1.3328 | -1.6306 | -0.15 |
| 15502 | *NFYB* | 0.67638 | -0.09539 | -1.0323 | -1.1808 | -0.15 |
| 15503 | *PPM1H* | 0.28141 | 0.087154 | -0.81999 | -0.95842 | -0.15 |
| 15504 | *CYP3A7* | 0.6504 | -0.12015 | -0.98238 | -1.9708 | -0.15 |
| 15505 | *KLHDC2* | 0.11527 | -0.17746 | -0.39081 | -1.0063 | -0.15 |
| 15506 | *TTC33* | 2.0832 | -0.78705 | -1.7493 | -2.1155 | -0.15 |
| 15507 | *LST1* | 0.93628 | -0.58403 | -0.80594 | -1.9295 | -0.15 |
| 15508 | *PCDHA9* | 1.4335 | -0.80014 | -1.0871 | -2.0629 | -0.15 |
| 15509 | *OR10T2* | 0.64211 | -0.45727 | -0.63903 | -1.0323 | -0.15 |
| 15510 | *TPT1* | 1.0756 | -0.74219 | -0.78784 | -2.3977 | -0.15 |
| 15511 | *FKBP2* | 0.50127 | 0.004855 | -0.96201 | -1.0221 | -0.15 |
| 15512 | *COL18A1* | 0.084291 | -0.13042 | -0.40998 | -1.4042 | -0.15 |
| 15513 | *COL5A3* | 0.83149 | -0.62041 | -0.66731 | -0.96808 | -0.15 |
| 15514 | *EIF4E* | 0.59905 | -0.37424 | -0.68115 | -1.0562 | -0.15 |
| 15515 | *STIP1* | -0.06416 | -0.0665 | -0.32605 | -1.803 | -0.15 |
| 15516 | *DDR2* | 0.85727 | -0.04877 | -1.2657 | -1.8596 | -0.15 |
| 15517 | *PPIAL4E* | 0.5661 | 0.39078 | -1.4141 | -1.8591 | -0.15 |
| 15518 | *AKAP17A* | 0.87943 | -0.24201 | -1.0954 | -1.2972 | -0.15 |
| 15519 | *ZNF217* | 0.43282 | 0.3065 | -1.1974 | -1.2142 | -0.15 |
| 15520 | *SRSF1* | 1.022 | -0.08832 | -1.3924 | -2.1132 | -0.15 |
| 15521 | *OMP* | 2.4779 | -1.2716 | -1.6653 | -1.7718 | -0.15 |
| 15522 | *RAN* | 0.96341 | -0.08944 | -1.3339 | -1.5569 | -0.15 |
| 15523 | *FBXW7* | 0.50286 | 0.18187 | -1.1448 | -1.3107 | -0.15 |
| 15524 | *DBF4B* | 0.018611 | -0.23814 | -0.24072 | -1.6597 | -0.15 |
| 15525 | *ZNF891* | 0.2551 | -0.10427 | -0.61114 | -1.3285 | -0.15 |
| 15526 | *GNAT3* | 1.0037 | 0.11767 | -1.583 | -1.7757 | -0.15 |
| 15527 | *MEDAG* | 0.45862 | -0.43176 | -0.48888 | -1.7906 | -0.15 |
| 15528 | *DENND2D* | 0.047897 | -0.24117 | -0.26892 | -0.6532 | -0.15 |
| 15529 | *ACKR1* | 2.8029 | -1.4258 | -1.8402 | -2.1798 | -0.15 |
| 15530 | *IZUMO1* | 1.0491 | -0.71443 | -0.79821 | -1.4775 | -0.15 |
| 15531 | *C1orf131* | 0.15169 | 0.061507 | -0.67702 | -1.6146 | -0.15 |
| 15532 | *AKT2* | 0.72433 | -0.40751 | -0.78229 | -1.1218 | -0.16 |
| 15533 | *MAP1A* | -0.07087 | -0.07884 | -0.3159 | -0.89147 | -0.16 |
| 15534 | *SCAMP5* | 1.0231 | 0.38102 | -1.8698 | -2.1802 | -0.16 |
| 15535 | *FAM117B* | 0.38996 | 0.38986 | -1.2457 | -1.5132 | -0.16 |
| 15536 | *NDOR1* | 0.33485 | -0.26779 | -0.53437 | -1.6663 | -0.16 |
| 15537 | *EI24* | 0.94162 | -0.35339 | -1.0561 | -1.4469 | -0.16 |
| 15538 | *ARPC4-TTLL3* | 0.020267 | -0.0757 | -0.41338 | -2.2254 | -0.16 |
| 15539 | *CERS2* | 1.0956 | -0.20112 | -1.3633 | -1.7647 | -0.16 |
| 15540 | *MICAL3* | 0.047356 | -0.15568 | -0.36084 | -1.4234 | -0.16 |
| 15541 | *FOXC2* | 1.5593 | -0.2715 | -1.7577 | -3.2901 | -0.16 |

| | | | | | | |
|---|---|---|---|---|---|---|
| 15542 | *DNAJC25* | 0.9582 | 0.25176 | -1.6799 | -3.8755 | -0.16 |
| 15543 | *P3H2* | 1.1793 | -0.40092 | -1.2485 | -1.4048 | -0.16 |
| 15544 | *KSR1* | 0.33636 | -0.03429 | -0.77227 | -1.7945 | -0.16 |
| 15545 | *HOXA9* | 0.73453 | -0.04927 | -1.1556 | -1.9302 | -0.16 |
| 15546 | *TCEB2* | 0.50851 | 0.017417 | -0.99694 | -1.0878 | -0.16 |
| 15547 | *POLD3* | 0.87109 | -0.6551 | -0.68741 | -1.5782 | -0.16 |
| 15548 | *PGS1* | 3.2456 | -1.4463 | -2.2712 | -2.796 | -0.16 |
| 15549 | *PQLC2L* | 0.31181 | 0.25872 | -1.043 | -1.31 | -0.16 |
| 15550 | *NLN* | 0.4785 | -0.40532 | -0.54579 | -0.76516 | -0.16 |
| 15551 | *OR5AN1* | 3.2083 | -1.6587 | -2.0223 | -2.2352 | -0.16 |
| 15552 | *UBXN11* | 1.7491 | -0.32737 | -1.8946 | -2.2771 | -0.16 |
| 15553 | *RYK* | 0.4423 | -0.07106 | -0.84411 | -0.87594 | -0.16 |
| 15554 | *TTC3* | 0.2482 | -0.33231 | -0.38901 | -0.75974 | -0.16 |
| 15555 | *YARS* | 0.40836 | 0.17792 | -1.0599 | -1.3947 | -0.16 |
| 15556 | *ANXA2R* | 0.51604 | -0.44353 | -0.54651 | -2.4405 | -0.16 |
| 15557 | *GDF9* | 0.24322 | 0.050441 | -0.7679 | -1.0273 | -0.16 |
| 15558 | *C2CD3* | 0.79289 | 0.1866 | -1.4544 | -2.2866 | -0.16 |
| 15559 | *KCTD5* | 0.96281 | -0.70947 | -0.72881 | -1.8157 | -0.16 |
| 15560 | *HMGA2* | 1.828 | -0.85107 | -1.4544 | -2.2809 | -0.16 |
| 15561 | *THSD7B* | 0.63179 | -0.1749 | -0.93489 | -2.9087 | -0.16 |
| 15562 | *LGALS13* | -0.05982 | -0.11145 | -0.30876 | -1.9827 | -0.16 |
| 15563 | *C9orf78* | 0.99291 | -0.69958 | -0.77409 | -1.0442 | -0.16 |
| 15564 | *EIF2B2* | 0.20086 | -0.30966 | -0.37199 | -1.7215 | -0.16 |
| 15565 | *CYTH1* | 0.36722 | -0.11637 | -0.73211 | -0.88767 | -0.16 |
| 15566 | *DEFB121* | 0.078312 | -0.18858 | -0.37125 | -1.7335 | -0.16 |
| 15567 | *GNA14* | 0.084209 | -0.06364 | -0.50339 | -1.9302 | -0.16 |
| 15568 | *FBXO41* | 0.062155 | -0.1647 | -0.38077 | -1.2684 | -0.16 |
| 15569 | *LSM1* | 1.1423 | -0.44353 | -1.1825 | -1.5894 | -0.16 |
| 15570 | *ZWILCH* | 1.2999 | -0.44572 | -1.3387 | -2.2528 | -0.16 |
| 15571 | *SLC4A7* | 2.3956 | -1.3427 | -1.5375 | -2.9354 | -0.16 |
| 15572 | *GLIS3* | 0.53014 | -0.40411 | -0.61114 | -1.7845 | -0.16 |
| 15573 | *IGSF1* | 1.5512 | -1.0024 | -1.0343 | -2.2311 | -0.16 |
| 15574 | *GPR21* | 0.84361 | -0.4221 | -0.90733 | -0.92217 | -0.16 |
| 15575 | *B4GALT5* | 0.4282 | -0.04207 | -0.87208 | -1.8478 | -0.16 |
| 15576 | *OSGEPL1* | 0.40069 | 0.33361 | -1.2208 | -1.864 | -0.16 |
| 15577 | *LMNA* | 0.82379 | -0.52914 | -0.78229 | -0.81478 | -0.16 |
| 15578 | *PSMA1* | 2.3187 | -1.3903 | -1.4172 | -1.6477 | -0.16 |
| 15579 | *T* | 0.25826 | -0.13255 | -0.61497 | -1.2098 | -0.16 |
| 15580 | *TMEM8C* | 0.34244 | 0.066398 | -0.8984 | -1.9353 | -0.16 |
| 15581 | *INSL6* | 2.0925 | -1.2208 | -1.3619 | -1.5243 | -0.16 |
| 15582 | *KEAP1* | 0.72239 | -0.55832 | -0.65489 | -0.89809 | -0.16 |
| 15583 | *SDHC* | 1.3221 | -0.58654 | -1.2267 | -2.5934 | -0.16 |
| 15584 | *DKK1* | 0.71104 | 0.083498 | -1.2867 | -1.6887 | -0.16 |
| 15585 | *MPL* | 0.98257 | -0.17721 | -1.2988 | -1.857 | -0.16 |
| 15586 | *IQCC* | 0.62912 | -0.25821 | -0.86459 | -1.5752 | -0.16 |
| 15587 | *CHD5* | 0.12226 | -0.20338 | -0.41359 | -1.584 | -0.16 |

| | | | | | | |
|---|---|---|---|---|---|---|
| 15588 | *KIF3C* | -0.04449 | -0.218 | -0.23252 | -0.83972 | -0.17 |
| 15589 | *PANX2* | 0.36676 | -0.17345 | -0.6886 | -2.3762 | -0.17 |
| 15590 | *TMIGD2* | 0.62998 | -0.46032 | -0.66535 | -1.4234 | -0.17 |
| 15591 | *SCNN1A* | 0.66316 | -0.4778 | -0.68192 | -1.5225 | -0.17 |
| 15592 | *FBXO6* | 1.5548 | -0.7436 | -1.3098 | -2.0055 | -0.17 |
| 15593 | *KLHL14* | 0.30839 | -0.1283 | -0.67887 | -1.7125 | -0.17 |
| 15594 | *SHROOM2* | 0.89485 | 0.81067 | -2.2044 | -3.4062 | -0.17 |
| 15595 | *SLC35G3* | 0.4082 | 0.13685 | -1.0453 | -1.1117 | -0.17 |
| 15596 | *P4HA1* | -0.05967 | -0.154 | -0.2868 | -1.9567 | -0.17 |
| 15597 | *CPT1A* | 2.6418 | -1.4442 | -1.6992 | -1.7332 | -0.17 |
| 15598 | *GNL2* | 0.54746 | -0.43077 | -0.61892 | -1.9168 | -0.17 |
| 15599 | *FAM196A* | 1.7222 | -0.43469 | -1.7906 | -3.324 | -0.17 |
| 15600 | *SGCB* | 0.75246 | -0.07472 | -1.181 | -1.2829 | -0.17 |
| 15601 | *LURAP1* | 0.41193 | -0.16459 | -0.75128 | -1.5082 | -0.17 |
| 15602 | *NCAPD3* | 0.18963 | -0.0648 | -0.62963 | -1.0446 | -0.17 |
| 15603 | *KHNYN* | 0.976 | -0.51862 | -0.96326 | -1.9678 | -0.17 |
| 15604 | *KRTAP4-3* | 1.5633 | -0.9744 | -1.0956 | -1.4234 | -0.17 |
| 15605 | *COPS2* | 0.79291 | -0.32216 | -0.97768 | -2.8702 | -0.17 |
| 15606 | *SCARA3* | 0.04123 | -0.15692 | -0.39124 | -0.57489 | -0.17 |
| 15607 | *SMYD1* | 0.034566 | -0.22327 | -0.31835 | -1.2946 | -0.17 |
| 15608 | *ZNF280D* | 1.7707 | -1.0385 | -1.2411 | -2.1947 | -0.17 |
| 15609 | *SPTLC1* | 0.66666 | -0.05357 | -1.122 | -2.2381 | -0.17 |
| 15610 | *CD3EAP* | 2.4625 | -0.97982 | -1.9919 | -2.0748 | -0.17 |
| 15611 | *PRC1* | 0.97072 | 0.12199 | -1.6033 | -2.4719 | -0.17 |
| 15612 | *CAMK1D* | 1.2163 | 0.22085 | -1.9479 | -2.425 | -0.17 |
| 15613 | *C19orf43* | 1.4023 | -0.88447 | -1.0289 | -1.151 | -0.17 |
| 15614 | *KDM4D* | 0.75545 | -0.0412 | -1.226 | -3.7932 | -0.17 |
| 15615 | *SEC24A* | 1.3848 | -0.82989 | -1.0669 | -1.3443 | -0.17 |
| 15616 | *LRRC56* | 0.77017 | -0.55563 | -0.72722 | -1.3643 | -0.17 |
| 15617 | *JPH3* | 0.35535 | -0.28982 | -0.57825 | -1.7874 | -0.17 |
| 15618 | *NPTN* | 0.21989 | -0.33136 | -0.40142 | -2.1143 | -0.17 |
| 15619 | *MCTP1* | 0.9884 | -0.72819 | -0.77444 | -1.1055 | -0.17 |
| 15620 | *ETNPPL* | 0.13942 | 0.12463 | -0.77895 | -1.4336 | -0.17 |
| 15621 | *GALNT9* | 0.61705 | 0.29116 | -1.4234 | -1.5848 | -0.17 |
| 15622 | *PGAM1* | 0.35018 | -0.31206 | -0.55359 | -1.3962 | -0.17 |
| 15623 | *ARHGEF4* | 0.29711 | -0.33253 | -0.48021 | -1.3453 | -0.17 |
| 15624 | *KCTD13* | 1.0022 | -0.25126 | -1.2666 | -1.8341 | -0.17 |
| 15625 | *LAMA5* | 2.2167 | -1.3183 | -1.4141 | -1.4524 | -0.17 |
| 15626 | *MRPL44* | 0.8152 | -0.09984 | -1.2318 | -1.671 | -0.17 |
| 15627 | *HTR2C* | 0.16852 | -0.20023 | -0.48486 | -1.2259 | -0.17 |
| 15628 | *NACAD* | 2.6472 | -1.1344 | -2.0303 | -2.3174 | -0.17 |
| 15629 | *PRKD3* | 0.58556 | -0.27702 | -0.82805 | -1.1066 | -0.17 |
| 15630 | *ANKRD13A* | 1.0457 | -0.4796 | -1.0863 | -2.071 | -0.17 |
| 15631 | *LIN7C* | 1.1714 | -0.77863 | -0.9135 | -1.8107 | -0.17 |
| 15632 | *SLC7A6* | 1.1694 | -0.18143 | -1.5121 | -2.1798 | -0.17 |
| 15633 | *RNF167* | 0.42261 | 0.15359 | -1.1005 | -2.4829 | -0.17 |

| | | | | | | |
|---|---|---|---|---|---|---|
| 15634 | *SPATA20* | 0.095844 | 0.083593 | -0.70391 | -3.7438 | -0.17 |
| 15635 | *LYPLA2* | 0.81854 | -0.36571 | -0.97786 | -2.1089 | -0.18 |
| 15636 | *CDC16* | 1.8956 | -0.54027 | -1.8804 | -1.9526 | -0.18 |
| 15637 | *APPBP2* | 1.2428 | -0.57787 | -1.1905 | -1.5426 | -0.18 |
| 15638 | *PCDH10* | 0.25238 | -0.32567 | -0.45296 | -1.1354 | -0.18 |
| 15639 | *ETFB* | 1.1564 | -0.69565 | -0.98747 | -1.4507 | -0.18 |
| 15640 | *IDI1* | 1.2108 | 0.10845 | -1.8462 | -2.4912 | -0.18 |
| 15641 | *POTEH* | 0.1119 | -0.28819 | -0.35111 | -0.46441 | -0.18 |
| 15642 | *ANG* | 1.4488 | -0.97583 | -1.0011 | -1.5458 | -0.18 |
| 15643 | *FAM26D* | 0.11893 | -0.27843 | -0.36963 | -1.53 | -0.18 |
| 15644 | *ECSIT* | 0.29555 | -0.07322 | -0.75182 | -1.5031 | -0.18 |
| 15645 | *MCMBP* | 0.57921 | 0.078477 | -1.1881 | -1.7081 | -0.18 |
| 15646 | *SYT1* | 0.13621 | 0.065894 | -0.73348 | -1.2036 | -0.18 |
| 15647 | *AMHR2* | 0.73634 | -0.42936 | -0.83856 | -0.86533 | -0.18 |
| 15648 | *HHEX* | -0.15548 | -0.18 | -0.19704 | -0.33484 | -0.18 |
| 15649 | *TNFAIP8L2* | 0.10337 | -0.19457 | -0.44136 | -1.7056 | -0.18 |
| 15650 | *TP53TG3* | 0.22969 | -0.27631 | -0.48599 | -1.4103 | -0.18 |
| 15651 | *C16orf71* | 0.75131 | -0.46206 | -0.82256 | -1.0162 | -0.18 |
| 15652 | *MAN2B2* | 0.018313 | -0.14966 | -0.40231 | -0.58787 | -0.18 |
| 15653 | *RPLP0* | 0.3334 | -0.40774 | -0.4595 | -0.49001 | -0.18 |
| 15654 | *TSC1* | 0.45764 | 0.042799 | -1.0344 | -2.3357 | -0.18 |
| 15655 | *EVI2A* | 0.50074 | -0.2412 | -0.79367 | -1.0435 | -0.18 |
| 15656 | *ZBTB39* | 0.90819 | -0.2481 | -1.1943 | -1.4182 | -0.18 |
| 15657 | *MED11* | 1.0708 | -0.57579 | -1.03 | -1.0773 | -0.18 |
| 15658 | *C11orf21* | 0.66702 | -0.55354 | -0.64896 | -1.1327 | -0.18 |
| 15659 | *SYT5* | 0.5995 | -0.5597 | -0.57646 | -1.5587 | -0.18 |
| 15660 | *DEFB103A* | 1.2495 | -0.15141 | -1.6349 | -1.7219 | -0.18 |
| 15661 | *IL1A* | 0.40077 | -0.17116 | -0.76856 | -1.7996 | -0.18 |
| 15662 | *COX16* | 0.5041 | -0.11263 | -0.93107 | -2.7068 | -0.18 |
| 15663 | *PHOSPHO2* | 0.18229 | -0.30878 | -0.41359 | -1.2855 | -0.18 |
| 15664 | *TMEM159* | 1.1805 | -0.39791 | -1.3232 | -2.3235 | -0.18 |
| 15665 | *VPRBP* | 0.96966 | -0.20668 | -1.3036 | -2.5546 | -0.18 |
| 15666 | *CPA2* | 4.1613 | -2.2135 | -2.4885 | -3.4314 | -0.18 |
| 15667 | *POU5F1B* | 0.68308 | -0.14132 | -1.084 | -1.5031 | -0.18 |
| 15668 | *SIAH2* | 0.32525 | 0.16382 | -1.0318 | -1.5588 | -0.18 |
| 15669 | *NHLRC2* | 0.91376 | -0.20489 | -1.252 | -1.9686 | -0.18 |
| 15670 | *ZNF837* | 0.8159 | -0.34402 | -1.0153 | -1.6158 | -0.18 |
| 15671 | *TMEM187* | 0.054282 | -0.25344 | -0.34535 | -2.5361 | -0.18 |
| 15672 | *HRASLS2* | 0.88308 | -0.63976 | -0.78784 | -1.5436 | -0.18 |
| 15673 | *BFSP2* | 0.11973 | -0.00828 | -0.65606 | -1.9455 | -0.18 |
| 15674 | *DLX6* | 0.47452 | 0.45571 | -1.4753 | -2.6201 | -0.18 |
| 15675 | *CSTF1* | 0.035088 | -0.09468 | -0.48572 | -0.62928 | -0.18 |
| 15676 | *NPIPB5* | 0.19572 | -0.18427 | -0.55773 | -2.4885 | -0.18 |
| 15677 | *DBP* | 2.5207 | -1.1866 | -1.8816 | -2.0164 | -0.18 |
| 15678 | *PCP4* | 1.3445 | -0.84225 | -1.0507 | -1.1676 | -0.18 |
| 15679 | *NDUFB6* | 1.0568 | -0.29599 | -1.3104 | -1.8461 | -0.18 |

| | | | | | | |
|---|---|---|---|---|---|---|
| 15680 | *FLOT2* | 0.24356 | 0.002893 | -0.79833 | -1.9624 | -0.18 |
| 15681 | *TMEM131* | 0.56404 | -0.394 | -0.72227 | -1.8775 | -0.18 |
| 15682 | *TUBA3C* | 0.46868 | -0.14164 | -0.88159 | -1.7313 | -0.18 |
| 15683 | *KRT25* | 2.0884 | -1.1676 | -1.4758 | -1.8881 | -0.19 |
| 15684 | *GUK1* | 0.34029 | 0.14819 | -1.0443 | -1.4852 | -0.19 |
| 15685 | *NUTM2B* | 0.13885 | -0.168 | -0.52833 | -0.9512 | -0.19 |
| 15686 | *CBL* | 2.3606 | -1.2411 | -1.6791 | -2.8478 | -0.19 |
| 15687 | *SMAGP* | 0.38929 | -0.09892 | -0.8504 | -2.1592 | -0.19 |
| 15688 | *SRGN* | -0.0643 | -0.23977 | -0.256 | -2.1072 | -0.19 |
| 15689 | *XPC* | 0.13215 | -0.279 | -0.41486 | -0.67546 | -0.19 |
| 15690 | *FEN1* | 0.89713 | -0.44353 | -1.0158 | -2.1342 | -0.19 |
| 15691 | *FOXA2* | 0.38578 | 0.15784 | -1.1069 | -1.2113 | -0.19 |
| 15692 | *OR51S1* | 1.0393 | -0.69939 | -0.90478 | -2.1182 | -0.19 |
| 15693 | *FAM134A* | 0.96739 | -0.18494 | -1.348 | -2.6267 | -0.19 |
| 15694 | *PPP1CB* | 1.9748 | -0.18214 | -2.3601 | -2.892 | -0.19 |
| 15695 | *ATG9B* | -0.02164 | -0.13816 | -0.40765 | -0.58851 | -0.19 |
| 15696 | *HOXD4* | 0.91178 | 0.12873 | -1.6086 | -2.0602 | -0.19 |
| 15697 | *GRM2* | 0.3438 | -0.271 | -0.64194 | -1.1461 | -0.19 |
| 15698 | *PTHLH* | 0.15579 | -0.02187 | -0.70391 | -2.9134 | -0.19 |
| 15699 | *SLC16A14* | 0.088875 | 0.025715 | -0.68466 | -1.0878 | -0.19 |
| 15700 | *FAM3C* | 0.79601 | -0.08059 | -1.2864 | -1.917 | -0.19 |
| 15701 | *UBE2F* | 0.98359 | -0.15277 | -1.4023 | -1.9844 | -0.19 |
| 15702 | *ITGB3* | -0.02883 | -0.21175 | -0.33131 | -1.6693 | -0.19 |
| 15703 | *OVOL1* | 0.1827 | -0.20268 | -0.55282 | -1.1448 | -0.19 |
| 15704 | *HPDL* | 0.54746 | 0.50661 | -1.6274 | -2.1522 | -0.19 |
| 15705 | *OGDH* | 1.7484 | -0.59959 | -1.7223 | -2.3601 | -0.19 |
| 15706 | *SH3BP5L* | 0.31011 | -0.2863 | -0.59738 | -1.1804 | -0.19 |
| 15707 | *CD2BP2* | -0.01001 | -0.24942 | -0.31414 | -1.5789 | -0.19 |
| 15708 | *TPPP* | 1.3884 | -0.39722 | -1.5651 | -1.9715 | -0.19 |
| 15709 | *THAP1* | 1.6617 | -0.46319 | -1.7745 | -2.017 | -0.19 |
| 15710 | *C8orf33* | 0.67866 | -0.41844 | -0.83638 | -1.1617 | -0.19 |
| 15711 | *MYEF2* | 0.21415 | -0.33169 | -0.45972 | -0.75053 | -0.19 |
| 15712 | *TGM1* | 2.7402 | -1.5274 | -1.7903 | -1.8324 | -0.19 |
| 15713 | *BECN1* | 0.47448 | -0.14132 | -0.91176 | -2.2218 | -0.19 |
| 15714 | *ERBB2* | 1.9872 | -0.97588 | -1.5904 | -2.1023 | -0.19 |
| 15715 | *TMEM245* | 0.83773 | 0.073517 | -1.4929 | -1.8387 | -0.19 |
| 15716 | *TCF7L1* | 0.66619 | 0.45802 | -1.7073 | -2.6431 | -0.19 |
| 15717 | *MYO9B* | 0.42673 | -0.28233 | -0.7275 | -1.7102 | -0.19 |
| 15718 | *POLR2H* | 0.96148 | -0.36914 | -1.1758 | -2.1072 | -0.19 |
| 15719 | *FATE1* | 0.045836 | -0.07219 | -0.55844 | -1.8157 | -0.19 |
| 15720 | *CEP44* | 0.27752 | -0.28176 | -0.58143 | -0.70546 | -0.20 |
| 15721 | *SLC2A3* | 0.43491 | 0.12772 | -1.1491 | -1.1747 | -0.20 |
| 15722 | *OR1D2* | 0.37835 | -0.08409 | -0.88098 | -0.94194 | -0.20 |
| 15723 | *AFF3* | 0.093569 | -0.02138 | -0.65906 | -1.6926 | -0.20 |
| 15724 | *TMEM258* | 0.57397 | -0.50961 | -0.65155 | -1.8402 | -0.20 |
| 15725 | *SLC1A2* | 0.89475 | -0.07785 | -1.4047 | -1.8789 | -0.20 |

| | | | | | | |
|---|---|---|---|---|---|---|
| 15726 | *DEFB105A* | 1.664 | -0.98577 | -1.2666 | -1.879 | -0.20 |
| 15727 | *RHBDL1* | -0.11975 | -0.23289 | -0.2379 | -2.1187 | -0.20 |
| 15728 | *ZBTB18* | 0.66061 | 0.50612 | -1.7581 | -2.0411 | -0.20 |
| 15729 | *GTF2H4* | 0.39283 | -0.25238 | -0.73202 | -0.81994 | -0.20 |
| 15730 | *HIST1H4J* | 0.66253 | -0.59763 | -0.65682 | -1.6722 | -0.20 |
| 15731 | *FASN* | 0.71749 | 0.43208 | -1.742 | -1.7745 | -0.20 |
| 15732 | *ABCB6* | 1.0648 | -0.43746 | -1.2198 | -1.2464 | -0.20 |
| 15733 | *SEC13* | 0.67533 | -0.32637 | -0.9425 | -1.4809 | -0.20 |
| 15734 | *DAGLB* | 0.47807 | -0.11771 | -0.95402 | -1.2765 | -0.20 |
| 15735 | *CALCB* | 0.46377 | -0.45292 | -0.6047 | -1.9105 | -0.20 |
| 15736 | *PRDX6* | 1.1336 | -0.45185 | -1.2756 | -1.3554 | -0.20 |
| 15737 | *CEP85* | 0.5665 | -0.3644 | -0.79663 | -1.5512 | -0.20 |
| 15738 | *HEXIM1* | 0.65278 | 0.03708 | -1.2848 | -3.5043 | -0.20 |
| 15739 | *GBP6* | 1.2474 | -0.87673 | -0.96583 | -2.4718 | -0.20 |
| 15740 | *PVR* | 0.33109 | 0.00553 | -0.93214 | -1.1821 | -0.20 |
| 15741 | *OR5A1* | 2.8979 | -1.4841 | -2.011 | -2.1103 | -0.20 |
| 15742 | *GIMAP6* | 0.63803 | -0.30312 | -0.93368 | -1.3101 | -0.20 |
| 15743 | *UBA2* | 0.34835 | -0.21526 | -0.73301 | -1.3296 | -0.20 |
| 15744 | *RPL32* | 0.46034 | -0.39149 | -0.66897 | -0.83231 | -0.20 |
| 15745 | *HSF2BP* | 0.33635 | -0.0576 | -0.87993 | -1.855 | -0.20 |
| 15746 | *PPP1R15A* | 0.23996 | -0.33253 | -0.509 | -5.1074 | -0.20 |
| 15747 | *GTF3C6* | 0.60413 | -0.42808 | -0.77777 | -1.9168 | -0.20 |
| 15748 | *C2orf80* | 1.4972 | -1.0256 | -1.0739 | -1.3455 | -0.20 |
| 15749 | *MED19* | 0.61954 | -0.24296 | -0.9793 | -2.1696 | -0.20 |
| 15750 | *AREL1* | 0.38336 | -0.1241 | -0.86229 | -1.3593 | -0.20 |
| 15751 | *ROMO1* | 0.3645 | -0.41205 | -0.55586 | -2.103 | -0.20 |
| 15752 | *PRUNE2* | 0.30373 | -0.25295 | -0.65512 | -1.3604 | -0.20 |
| 15753 | *POLR1C* | 0.78824 | -0.22256 | -1.1706 | -1.9473 | -0.20 |
| 15754 | *PFN2* | 0.53529 | -0.21946 | -0.92129 | -1.6085 | -0.20 |
| 15755 | *POLE4* | 0.28193 | 0.084469 | -0.97255 | -3.0124 | -0.20 |
| 15756 | *USP17L24* | 2.3459 | -1.3635 | -1.5888 | -1.9651 | -0.20 |
| 15757 | *JAKMIP3* | 0.45029 | 0.35108 | -1.4087 | -2.3904 | -0.20 |
| 15758 | *HLA-DMA* | -0.0925 | -0.2372 | -0.27811 | -0.96201 | -0.20 |
| 15759 | *HIRIP3* | 0.51973 | -0.32779 | -0.80014 | -1.2867 | -0.20 |
| 15760 | *KCNJ1* | 2.0458 | -1.0252 | -1.6288 | -1.6885 | -0.20 |
| 15761 | *NT5C2* | 0.75465 | 0.67742 | -2.0407 | -3.4888 | -0.20 |
| 15762 | *RPL37* | 1.2386 | -0.46733 | -1.3808 | -1.6526 | -0.20 |
| 15763 | *FAM83G* | 0.23542 | 0.18732 | -1.0324 | -1.3511 | -0.20 |
| 15764 | *LYSMD3* | 0.8936 | -0.42349 | -1.0803 | -1.4128 | -0.20 |
| 15765 | *TMEM145* | 0.39232 | 0.31279 | -1.3162 | -1.5212 | -0.20 |
| 15766 | *ACAD8* | 0.31078 | -0.37676 | -0.54621 | -0.84336 | -0.20 |
| 15767 | *ZNF341* | 0.59348 | -0.51585 | -0.68992 | -2.1187 | -0.20 |
| 15768 | *MGARP* | 0.11552 | -0.33266 | -0.3952 | -0.69841 | -0.20 |
| 15769 | *WDR74* | 0.85981 | 0.040077 | -1.5125 | -1.5491 | -0.20 |
| 15770 | *FH* | 0.46545 | 0.28749 | -1.3666 | -2.3857 | -0.20 |
| 15771 | *TRAPPC6A* | 0.87856 | 0.13794 | -1.6315 | -1.9768 | -0.21 |

| | | | | | | |
|---|---|---|---|---|---|---|
| 15772 | *PLEKHH1* | 0.15043 | -0.03313 | -0.73259 | -0.88736 | -0.21 |
| 15773 | *TERF2* | 0.095567 | -0.07297 | -0.63791 | -1.5213 | -0.21 |
| 15774 | *GPR33* | 1.6979 | -0.56664 | -1.747 | -2.2945 | -0.21 |
| 15775 | *SNRPA* | 1.0287 | 0.27221 | -1.9175 | -2.3017 | -0.21 |
| 15776 | *AAR2* | 1.8541 | -1.0907 | -1.38 | -1.8055 | -0.21 |
| 15777 | *CCDC64* | 0.71555 | 0.37368 | -1.7061 | -2.0028 | -0.21 |
| 15778 | *SAV1* | 0.57985 | -0.44353 | -0.75388 | -1.879 | -0.21 |
| 15779 | *MCM8* | 0.38623 | -0.09323 | -0.91061 | -0.9554 | -0.21 |
| 15780 | *PPP2R4* | 0.69258 | -0.11619 | -1.1943 | -1.3677 | -0.21 |
| 15781 | *COPG2* | 0.54936 | -0.43634 | -0.73202 | -0.95629 | -0.21 |
| 15782 | *ZNF684* | 0.77368 | -0.57542 | -0.81757 | -1.7194 | -0.21 |
| 15783 | *KCTD14* | 1.3526 | -0.53541 | -1.4368 | -2.1151 | -0.21 |
| 15784 | *CPSF7* | 0.49337 | -0.42378 | -0.69003 | -2.4784 | -0.21 |
| 15785 | *RHOQ* | 1.0522 | -0.64467 | -1.0293 | -1.4064 | -0.21 |
| 15786 | *NAIF1* | 0.33767 | -0.39885 | -0.5613 | -3.3337 | -0.21 |
| 15787 | *PRM1* | 0.80342 | -0.69132 | -0.73529 | -2.1018 | -0.21 |
| 15788 | *RPS6KA3* | 0.012638 | -0.1778 | -0.4583 | -0.62527 | -0.21 |
| 15789 | *EDC3* | 2.0483 | -0.88208 | -1.7901 | -2.0055 | -0.21 |
| 15790 | *WFDC2* | -0.09789 | -0.22045 | -0.30608 | -1.2339 | -0.21 |
| 15791 | *PPP4C* | 0.57599 | -0.19213 | -1.0093 | -1.496 | -0.21 |
| 15792 | *ZNF75A* | 2.6028 | -1.4233 | -1.806 | -2.0553 | -0.21 |
| 15793 | *KCNIP2* | 1.5085 | -0.97781 | -1.1585 | -2.6805 | -0.21 |
| 15794 | *VIM* | 1.8948 | -1.1263 | -1.3966 | -2.1404 | -0.21 |
| 15795 | *PSMB6* | 0.49944 | 0.14324 | -1.2708 | -1.4498 | -0.21 |
| 15796 | *LY75-CD302* | 0.59574 | -0.28406 | -0.94255 | -2.6803 | -0.21 |
| 15797 | *STOX2* | 0.64372 | -0.56485 | -0.70998 | -1.1067 | -0.21 |
| 15798 | *CRYGC* | 0.74302 | -0.49778 | -0.87662 | -0.95759 | -0.21 |
| 15799 | *SLC25A22* | 0.4344 | 0.095588 | -1.1617 | -1.589 | -0.21 |
| 15800 | *TAF9* | 0.82054 | -0.71016 | -0.7425 | -1.6105 | -0.21 |
| 15801 | *RNF123* | -0.04553 | -0.29263 | -0.29414 | -1.2215 | -0.21 |
| 15802 | *CBFA2T2* | 0.74318 | -0.65901 | -0.71669 | -1.0813 | -0.21 |
| 15803 | *RPS9* | 2.5348 | -1.3683 | -1.7991 | -2.796 | -0.21 |
| 15804 | *ATP5O* | 0.95133 | -0.53025 | -1.0545 | -2.0214 | -0.21 |
| 15805 | *SPPL3* | 0.26758 | -0.25353 | -0.64806 | -1.4047 | -0.21 |
| 15806 | *SLC35F4* | 1.5783 | 0.38033 | -2.5934 | -2.6805 | -0.21 |
| 15807 | *GJA1* | 1.6194 | -0.87598 | -1.379 | -1.5293 | -0.21 |
| 15808 | *KRT75* | -0.0275 | -0.15657 | -0.45185 | -0.97477 | -0.21 |
| 15809 | *AIF1L* | 0.21539 | -0.2052 | -0.64671 | -1.8737 | -0.21 |
| 15810 | *SLC16A9* | 0.52332 | 0.040531 | -1.2014 | -1.716 | -0.21 |
| 15811 | *AKT1S1* | 1.3188 | -0.38472 | -1.5722 | -1.9023 | -0.21 |
| 15812 | *IRF2* | 0.37441 | -0.26895 | -0.74365 | -1.951 | -0.21 |
| 15813 | *EIF4EBP3* | 0.4084 | 0.31108 | -1.358 | -3.1568 | -0.21 |
| 15814 | *ZNF45* | 0.34851 | 0.2961 | -1.2832 | -1.6597 | -0.21 |
| 15815 | *UHRF2* | 0.25848 | -0.0351 | -0.86358 | -2.6964 | -0.21 |
| 15816 | *C21orf62* | 0.29586 | -0.30371 | -0.63341 | -1.7652 | -0.21 |
| 15817 | *CSMD2* | 0.5366 | -0.58851 | -0.58949 | -2.3204 | -0.21 |

| | | | | | | |
|---|---|---|---|---|---|---|
| 15818 | *BBS2* | 0.97032 | -0.78099 | -0.83231 | -0.87817 | -0.21 |
| 15819 | *MMP13* | 0.43472 | -0.00472 | -1.073 | -1.748 | -0.21 |
| 15820 | *BTBD7* | 0.61741 | -0.33266 | -0.92781 | -1.2383 | -0.21 |
| 15821 | *IMPDH2* | 0.69471 | -0.13543 | -1.2036 | -2.0008 | -0.21 |
| 15822 | *VWA9* | 0.91948 | -0.62073 | -0.94399 | -1.7268 | -0.22 |
| 15823 | *MESP1* | 0.93499 | -0.29879 | -1.2815 | -2.2167 | -0.22 |
| 15824 | *CHIC1* | 0.46422 | 0.23955 | -1.3492 | -1.3591 | -0.22 |
| 15825 | *ARMCX5* | 0.52927 | -0.43906 | -0.73668 | -0.98585 | -0.22 |
| 15826 | *CEP135* | 0.2732 | -0.4317 | -0.48829 | -0.96857 | -0.22 |
| 15827 | *ZFP41* | 0.38454 | 0.065282 | -1.0973 | -2.1108 | -0.22 |
| 15828 | *SFRP2* | -0.08064 | -0.23875 | -0.32861 | -0.34181 | -0.22 |
| 15829 | *SERPINA11* | 1.1914 | -0.50524 | -1.3343 | -1.6246 | -0.22 |
| 15830 | *SART3* | 1.9354 | -0.95645 | -1.6281 | -1.7142 | -0.22 |
| 15831 | *RUFY1* | 0.26863 | -0.36233 | -0.55586 | -0.72825 | -0.22 |
| 15832 | *C15orf57* | 2.3666 | -0.93107 | -2.0851 | -2.5361 | -0.22 |
| 15833 | *TMPRSS11F* | 0.33676 | -0.43634 | -0.55002 | -3.3367 | -0.22 |
| 15834 | *ACTR8* | 0.44638 | -0.25205 | -0.84457 | -1.9602 | -0.22 |
| 15835 | *RCVRN* | 1.0166 | -0.4499 | -1.2172 | -1.3729 | -0.22 |
| 15836 | *MED23* | 1.2792 | -0.71953 | -1.2109 | -1.8582 | -0.22 |
| 15837 | *OR52N2* | 1.4202 | -0.97982 | -1.0918 | -2.1351 | -0.22 |
| 15838 | *INMT* | 0.34115 | 0.19975 | -1.1925 | -1.42 | -0.22 |
| 15839 | *RTP5* | 1.3649 | -0.64219 | -1.3751 | -2.9494 | -0.22 |
| 15840 | *LOC102724957* | 0.56579 | -0.51859 | -0.70074 | -1.7266 | -0.22 |
| 15841 | *DBR1* | 2.364 | -0.76606 | -2.252 | -2.4331 | -0.22 |
| 15842 | *TARS* | 0.73122 | -0.39308 | -0.99241 | -2.1873 | -0.22 |
| 15843 | *LAMTOR5* | 0.43087 | -0.2704 | -0.81509 | -3.3367 | -0.22 |
| 15844 | *MTMR6* | 0.60661 | 0.043739 | -1.3067 | -1.4034 | -0.22 |
| 15845 | *WDR87* | 1.0589 | -0.82897 | -0.88784 | -2.5442 | -0.22 |
| 15846 | *MAPK13* | 2.163 | -0.56171 | -2.2593 | -2.6498 | -0.22 |
| 15847 | *NCKIPSD* | -0.03491 | -0.15626 | -0.46705 | -1.5306 | -0.22 |
| 15848 | *ECHDC3* | 0.72272 | -0.44774 | -0.93474 | -1.6682 | -0.22 |
| 15849 | *IZUMO2* | 0.75823 | -0.27541 | -1.1426 | -1.1964 | -0.22 |
| 15850 | *NOC4L* | 1.2871 | -0.60543 | -1.3419 | -2.0113 | -0.22 |
| 15851 | *DAXX* | 0.56073 | -0.43845 | -0.78314 | -1.0107 | -0.22 |
| 15852 | *MYO10* | 0.34835 | -0.43884 | -0.57142 | -0.79711 | -0.22 |
| 15853 | *ADAMTS2* | -0.06145 | -0.11219 | -0.48854 | -1.2162 | -0.22 |
| 15854 | *ERLIN2* | 0.52749 | 0.47987 | -1.6696 | -2.3876 | -0.22 |
| 15855 | *ANKRD20A1* | 1.1396 | -0.01531 | -1.7866 | -1.9271 | -0.22 |
| 15856 | *SF1* | 0.56915 | -0.53987 | -0.69186 | -2.4038 | -0.22 |
| 15857 | *TRAK1* | 0.37779 | -0.51221 | -0.52853 | -0.56376 | -0.22 |
| 15858 | *RNF187* | 0.91433 | 0.083863 | -1.6617 | -1.917 | -0.22 |
| 15859 | *ARG1* | 0.48128 | 0.14807 | -1.2935 | -1.3296 | -0.22 |
| 15860 | *TOP2A* | 1.039 | 0.046235 | -1.7495 | -2.0601 | -0.22 |
| 15861 | *GPR22* | 0.91449 | -0.5193 | -1.0596 | -1.7874 | -0.22 |
| 15862 | *ARMCX4* | 0.29782 | -0.37578 | -0.58654 | -1.4048 | -0.22 |
| 15863 | *EFCAB6* | 0.90876 | -0.54307 | -1.0303 | -1.864 | -0.22 |

| | | | | | | |
|---|---|---|---|---|---|---|
| 15864 | *RRM2B* | 2.6274 | -1.5341 | -1.7591 | -1.9028 | -0.22 |
| 15865 | *CLIC3* | 0.50365 | -0.36557 | -0.80548 | -2.5298 | -0.22 |
| 15866 | *OR8U1* | 1.1907 | -0.42177 | -1.4375 | -2.8863 | -0.22 |
| 15867 | *KLHL28* | 0.82626 | -0.5183 | -0.97784 | -1.1487 | -0.22 |
| 15868 | *ST6GALNAC2* | 0.25152 | 0.20572 | -1.1278 | -1.1607 | -0.22 |
| 15869 | *REPS1* | 0.014271 | -0.10628 | -0.57857 | -0.8613 | -0.22 |
| 15870 | *IGHMBP2* | 0.43719 | -0.1227 | -0.98585 | -1.0485 | -0.22 |
| 15871 | *NOTCH4* | 0.8204 | 0.062587 | -1.5544 | -1.6356 | -0.22 |
| 15872 | *NOP14* | 0.10289 | -0.32143 | -0.4537 | -1.5984 | -0.22 |
| 15873 | *CIITA* | 0.48201 | -0.06214 | -1.0925 | -1.309 | -0.22 |
| 15874 | *MRPL48* | 0.96442 | -0.78229 | -0.85478 | -2.3486 | -0.22 |
| 15875 | *SPDYE1* | 1.0095 | -0.79316 | -0.88981 | -0.89854 | -0.22 |
| 15876 | *INTS1* | 0.43235 | -0.38059 | -0.72635 | -1.0301 | -0.22 |
| 15877 | *ATP4B* | 0.28226 | -0.46754 | -0.49007 | -3.4182 | -0.23 |
| 15878 | *LIPH* | 1.7369 | -1.0856 | -1.329 | -1.6594 | -0.23 |
| 15879 | *DRD2* | 0.42958 | -0.46767 | -0.63971 | -0.97046 | -0.23 |
| 15880 | *RNASEL* | 0.41177 | -0.36562 | -0.7241 | -2.6889 | -0.23 |
| 15881 | *NFAM1* | 0.48445 | -0.47769 | -0.68586 | -1.4601 | -0.23 |
| 15882 | *MED25* | 1.2001 | -0.71077 | -1.1685 | -1.4754 | -0.23 |
| 15883 | *SNRNP40* | 1.5884 | -1.0489 | -1.2187 | -2.5852 | -0.23 |
| 15884 | *DLX1* | 1.9314 | -1.2736 | -1.3371 | -1.5552 | -0.23 |
| 15885 | *AGPAT2* | 0.60513 | -0.54887 | -0.73565 | -2.4815 | -0.23 |
| 15886 | *EAPP* | 2.9047 | -1.6116 | -1.9727 | -2.6376 | -0.23 |
| 15887 | *C9orf62* | 0.51927 | -0.51494 | -0.6858 | -1.0088 | -0.23 |
| 15888 | *ZNF594* | 0.11365 | -0.35161 | -0.44353 | -1.6842 | -0.23 |
| 15889 | *RCOR2* | 1.4261 | -0.80349 | -1.3042 | -1.4361 | -0.23 |
| 15890 | *PEX16* | 1.2977 | -0.54409 | -1.4375 | -2.1893 | -0.23 |
| 15891 | *SLC25A37* | 0.50884 | -0.30029 | -0.89261 | -1.499 | -0.23 |
| 15892 | *SLC9A3R1* | -0.03229 | -0.23719 | -0.41533 | -1.349 | -0.23 |
| 15893 | *PDCD1LG2* | 0.61378 | -0.59871 | -0.70074 | -0.81918 | -0.23 |
| 15894 | *TMEM220* | 1.8207 | -1.1508 | -1.3565 | -1.5171 | -0.23 |
| 15895 | *MGME1* | 1.0066 | -0.30611 | -1.3878 | -3.5278 | -0.23 |
| 15896 | *ANKLE2* | 0.15113 | 0.10415 | -0.94341 | -1.748 | -0.23 |
| 15897 | *SNX22* | 0.12366 | -0.13984 | -0.67395 | -2.8807 | -0.23 |
| 15898 | *KRBOX4* | -0.07191 | -0.21964 | -0.39885 | -0.61264 | -0.23 |
| 15899 | *SRA1* | 1.3626 | -0.53069 | -1.5247 | -2.0553 | -0.23 |
| 15900 | *PSMD12* | 0.51986 | -0.01545 | -1.1974 | -2.3471 | -0.23 |
| 15901 | *DNAJA3* | 2.0583 | -0.81505 | -1.9367 | -2.5603 | -0.23 |
| 15902 | *TMEM140* | 1.3397 | -0.43634 | -1.5971 | -2.0176 | -0.23 |
| 15903 | *MPC1L* | 0.23768 | -0.39457 | -0.5377 | -1.0871 | -0.23 |
| 15904 | *MYCL* | 1.2705 | -0.92288 | -1.043 | -2.9283 | -0.23 |
| 15905 | *PIK3C3* | 0.32416 | -0.03784 | -0.98211 | -1.0295 | -0.23 |
| 15906 | *CHMP7* | 1.0214 | -0.35063 | -1.367 | -1.5658 | -0.23 |
| 15907 | *UXS1* | 0.34636 | -0.05715 | -0.98572 | -1.7018 | -0.23 |
| 15908 | *HIST1H4H* | 0.3721 | 0.054707 | -1.1242 | -2.0411 | -0.23 |
| 15909 | *NOL7* | 1.3001 | -0.92553 | -1.0726 | -1.7963 | -0.23 |

| | | | | | | |
|---|---|---|---|---|---|---|
| 15910 | *DLL1* | 0.1979 | -0.34077 | -0.55741 | -2.4268 | -0.23 |
| 15911 | *RFPL1* | 0.3198 | -0.34691 | -0.67321 | -2.9236 | -0.23 |
| 15912 | *OR4N4* | 0.82244 | 0.30863 | -1.8318 | -1.8462 | -0.23 |
| 15913 | *ARFRP1* | 1.143 | -0.3271 | -1.5167 | -2.3424 | -0.23 |
| 15914 | *CCDC115* | 0.20086 | 0.18746 | -1.0894 | -1.2846 | -0.23 |
| 15915 | *UBAP1* | 0.10782 | -0.2193 | -0.58994 | -0.73443 | -0.23 |
| 15916 | *SMPD2* | 1.9216 | -0.89474 | -1.729 | -2.3372 | -0.23 |
| 15917 | *YEATS4* | 0.44933 | 0.10802 | -1.2595 | -1.3218 | -0.23 |
| 15918 | *GCC1* | 1.3041 | -0.09807 | -1.9089 | -3.9751 | -0.23 |
| 15919 | *OTUD3* | 0.6799 | -0.68896 | -0.69395 | -1.4491 | -0.23 |
| 15920 | *EFR3B* | 0.35299 | -0.41406 | -0.6421 | -0.73604 | -0.23 |
| 15921 | *CFHR2* | 2.7399 | -1.4684 | -1.9758 | -2.2456 | -0.23 |
| 15922 | *OASL* | 1.0837 | -0.79902 | -0.98925 | -1.1216 | -0.23 |
| 15923 | *HSH2D* | 0.76028 | -0.34906 | -1.1169 | -1.4312 | -0.24 |
| 15924 | *GSX1* | 1.2967 | -0.08601 | -1.9166 | -2.1893 | -0.24 |
| 15925 | *MED14* | 1.9837 | -1.1472 | -1.5425 | -2.8478 | -0.24 |
| 15926 | *C10orf76* | 0.67278 | -0.59157 | -0.78959 | -1.3292 | -0.24 |
| 15927 | *C2CD5* | 1.1583 | -0.62114 | -1.2465 | -3.1086 | -0.24 |
| 15928 | *IL17C* | 0.35364 | -0.44134 | -0.62213 | -1.0407 | -0.24 |
| 15929 | *SLC5A11* | 0.71478 | -0.62479 | -0.8003 | -1.5031 | -0.24 |
| 15930 | *FAM129B* | 1.6289 | -0.32494 | -2.0164 | -3.5736 | -0.24 |
| 15931 | *DMRT1* | 1.5329 | -1.018 | -1.2288 | -1.7945 | -0.24 |
| 15932 | *PTPN23* | 1.9158 | -1.2405 | -1.3905 | -3.6757 | -0.24 |
| 15933 | *COG3* | -0.07673 | -0.13009 | -0.50869 | -0.95722 | -0.24 |
| 15934 | *CNOT8* | 0.47126 | 0.41068 | -1.5978 | -2.4094 | -0.24 |
| 15935 | *TP53TG5* | 0.25012 | -0.12418 | -0.84291 | -1.6604 | -0.24 |
| 15936 | *AGFG1* | 1.6917 | -0.83676 | -1.5724 | -1.9089 | -0.24 |
| 15937 | *ITPRIPL1* | 0.79192 | -0.00828 | -1.5016 | -1.5568 | -0.24 |
| 15938 | *CCDC83* | 0.7058 | -0.2223 | -1.2016 | -1.4055 | -0.24 |
| 15939 | *C14orf177* | 0.48497 | -0.0908 | -1.1124 | -1.1496 | -0.24 |
| 15940 | *CCDC109B* | 0.014931 | -0.31295 | -0.42046 | -1.2145 | -0.24 |
| 15941 | *GPA33* | -0.11354 | -0.12556 | -0.48022 | -2.3036 | -0.24 |
| 15942 | *RFX5* | 0.67805 | 0.00784 | -1.4064 | -1.9474 | -0.24 |
| 15943 | *FANCF* | 0.71104 | -0.35905 | -1.0726 | -1.36 | -0.24 |
| 15944 | *TMCO4* | 0.57494 | -0.17016 | -1.1258 | -1.208 | -0.24 |
| 15945 | *ALDOB* | 0.91239 | -0.48973 | -1.1439 | -2.0165 | -0.24 |
| 15946 | *ULK1* | 0.55504 | -0.24401 | -1.0328 | -1.8707 | -0.24 |
| 15947 | *TMTC3* | 0.10931 | 0.079762 | -0.91134 | -1.3899 | -0.24 |
| 15948 | *RAB29* | 0.48714 | -0.02232 | -1.1872 | -2.0791 | -0.24 |
| 15949 | *C19orf33* | 0.045563 | -0.36705 | -0.40109 | -1.4597 | -0.24 |
| 15950 | *RNASEH2C* | 1.0777 | -0.41973 | -1.3808 | -2.3644 | -0.24 |
| 15951 | *ELP6* | 0.37459 | -0.45748 | -0.64088 | -2.0863 | -0.24 |
| 15952 | *NPLOC4* | 0.58268 | -0.33253 | -0.97393 | -1.9622 | -0.24 |
| 15953 | *SLC16A11* | 0.275 | 0.20215 | -1.2015 | -2.1108 | -0.24 |
| 15954 | *TRIM7* | 0.23698 | -0.38574 | -0.5765 | -0.67801 | -0.24 |
| 15955 | *SLC26A4* | 1.6562 | -1.1508 | -1.2312 | -1.4841 | -0.24 |

| | | | | | | |
|---|---|---|---|---|---|---|
| 15956 | *SPPL2B* | 2.2258 | -1.2259 | -1.7266 | -2.3389 | -0.24 |
| 15957 | *DDX17* | 1.7016 | -0.7097 | -1.7188 | -1.7538 | -0.24 |
| 15958 | *OR5F1* | 0.84965 | -0.16277 | -1.4142 | -1.4284 | -0.24 |
| 15959 | *THRSP* | 0.91211 | -0.80455 | -0.83511 | -0.9442 | -0.24 |
| 15960 | *ZNF763* | 0.75628 | 0.41923 | -1.9038 | -3.0974 | -0.24 |
| 15961 | *ETV2* | 1.1191 | -0.91685 | -0.93057 | -2.3261 | -0.24 |
| 15962 | *TMEM191B* | 0.6433 | -0.17052 | -1.2014 | -1.3316 | -0.24 |
| 15963 | *RANGRF* | 0.43211 | 0.24349 | -1.4043 | -2.2586 | -0.24 |
| 15964 | *FREM3* | 0.53862 | -0.11057 | -1.1575 | -2.1085 | -0.24 |
| 15965 | *LONRF3* | 2.3489 | -1.2571 | -1.8227 | -1.9446 | -0.24 |
| 15966 | *FBXO21* | 0.21315 | -0.12876 | -0.81543 | -2.43 | -0.24 |
| 15967 | *C20orf203* | 0.41263 | -0.36311 | -0.78098 | -1.7149 | -0.24 |
| 15968 | *AGAP11* | -0.08064 | -0.21833 | -0.43251 | -1.5026 | -0.24 |
| 15969 | *OR1D5* | 0.14137 | -0.33812 | -0.53481 | -0.68522 | -0.24 |
| 15970 | *RPS6KA6* | 0.50712 | -0.27961 | -0.96031 | -1.8611 | -0.24 |
| 15971 | *GSTM4* | 0.31744 | -0.10928 | -0.94119 | -2.3033 | -0.24 |
| 15972 | *ARRDC3* | -0.07982 | -0.27855 | -0.37602 | -0.58703 | -0.24 |
| 15973 | *TNC* | 0.58558 | -0.53661 | -0.78363 | -1.0049 | -0.24 |
| 15974 | *PXYLP1* | 0.48465 | 0.14018 | -1.3597 | -1.5307 | -0.24 |
| 15975 | *C7* | 0.47342 | 0.284 | -1.4932 | -1.6836 | -0.25 |
| 15976 | *ARID1B* | 0.76684 | 0.014205 | -1.5171 | -2.5008 | -0.25 |
| 15977 | *XPO5* | 0.53244 | -0.5903 | -0.6785 | -1.2063 | -0.25 |
| 15978 | *BHLHE22* | -0.05129 | -0.32121 | -0.36404 | -0.97865 | -0.25 |
| 15979 | *BNIP3* | 0.32504 | -0.2192 | -0.84291 | -0.89529 | -0.25 |
| 15980 | *KRT6C* | 0.88577 | -0.35675 | -1.2661 | -1.8575 | -0.25 |
| 15981 | *SMTNL1* | 0.7407 | -0.61779 | -0.86047 | -2.4784 | -0.25 |
| 15982 | *ZP2* | 0.059245 | -0.31815 | -0.47929 | -1.0919 | -0.25 |
| 15983 | *NBPF20* | 0.13087 | -0.20994 | -0.65946 | -1.8872 | -0.25 |
| 15984 | *RBM20* | 0.80365 | 0.12548 | -1.6682 | -2.7191 | -0.25 |
| 15985 | *SLC6A14* | 0.61334 | -0.47486 | -0.87768 | -1.6791 | -0.25 |
| 15986 | *FEZF2* | 1.8934 | -0.34476 | -2.2879 | -2.3235 | -0.25 |
| 15987 | *SUGP2* | 0.39896 | -0.51549 | -0.62306 | -0.97952 | -0.25 |
| 15988 | *GFPT1* | 1.1839 | -0.89593 | -1.0286 | -2.1496 | -0.25 |
| 15989 | *ATF6* | -0.1611 | -0.25996 | -0.31984 | -1.8157 | -0.25 |
| 15990 | *CYP26C1* | 0.48677 | -0.44353 | -0.78439 | -1.7655 | -0.25 |
| 15991 | *PURG* | 1.0294 | -0.49073 | -1.28 | -1.5494 | -0.25 |
| 15992 | *CKS1B* | 0.68203 | -0.01811 | -1.4057 | -1.8227 | -0.25 |
| 15993 | *ADRA1D* | 0.46505 | -0.04542 | -1.1619 | -1.7073 | -0.25 |
| 15994 | *ZC3H15* | 0.22085 | -0.25261 | -0.71063 | -1.1387 | -0.25 |
| 15995 | *USP11* | 0.51575 | -0.57448 | -0.68376 | -1.2027 | -0.25 |
| 15996 | *SSH2* | 0.2826 | -0.0261 | -0.99917 | -1.5788 | -0.25 |
| 15997 | *THBS3* | 0.4303 | 0.38377 | -1.5568 | -1.7364 | -0.25 |
| 15998 | *FBN3* | 0.66338 | -0.31126 | -1.0952 | -2.2599 | -0.25 |
| 15999 | *DDX10* | 1.7609 | -1.1384 | -1.3663 | -2.4456 | -0.25 |
| 16000 | *CTNNBL1* | 0.13093 | 0.086138 | -0.96144 | -1.7215 | -0.25 |
| 16001 | *AR* | 0.28345 | -0.24501 | -0.78325 | -1.4076 | -0.25 |

| | | | | | | |
|---|---|---|---|---|---|---|
| 16002 | *RSPO1* | 1.1671 | -0.4055 | -1.5073 | -1.9329 | -0.25 |
| 16003 | *ARL13B* | 0.86108 | -0.78484 | -0.82289 | -1.5645 | -0.25 |
| 16004 | *DERA* | 0.40814 | -0.54734 | -0.60845 | -1.7963 | -0.25 |
| 16005 | *CH17-360D5.1* | 0.36972 | 0.00068 | -1.1185 | -1.6626 | -0.25 |
| 16006 | *POLR2J* | 0.73274 | 0.32263 | -1.8038 | -3.8779 | -0.25 |
| 16007 | *SUPT7L* | 0.59849 | -0.00058 | -1.347 | -1.448 | -0.25 |
| 16008 | *DEFB133* | 0.4488 | -0.14151 | -1.0564 | -1.1113 | -0.25 |
| 16009 | *DDX18* | 0.5805 | -0.47412 | -0.85594 | -1.0898 | -0.25 |
| 16010 | *DTX4* | 0.2261 | -0.45607 | -0.52008 | -0.64574 | -0.25 |
| 16011 | *MRPS35* | 0.096888 | -0.36991 | -0.47707 | -1.2431 | -0.25 |
| 16012 | *GPHN* | 2.1603 | -1.3226 | -1.5884 | -1.6836 | -0.25 |
| 16013 | *PRAMEF5* | 0.42877 | 0.21595 | -1.3956 | -1.4614 | -0.25 |
| 16014 | *CALCOCO1* | 0.55344 | -0.25875 | -1.0456 | -1.1577 | -0.25 |
| 16015 | *FBXL18* | 1.5311 | -0.64192 | -1.6401 | -1.9899 | -0.25 |
| 16016 | *PNPLA3* | 1.3282 | 0.089257 | -2.1685 | -2.6222 | -0.25 |
| 16017 | *OR1L4* | 0.50321 | 0.38156 | -1.6368 | -1.8838 | -0.25 |
| 16018 | *C2orf72* | 0.95045 | -0.40749 | -1.2968 | -1.9175 | -0.25 |
| 16019 | *ZNF117* | -0.02598 | -0.36371 | -0.36451 | -0.75192 | -0.25 |
| 16020 | *PKD1L2* | 0.62338 | -0.67573 | -0.70188 | -1.0948 | -0.25 |
| 16021 | *SP5* | 0.26315 | -0.13183 | -0.88561 | -1.5876 | -0.25 |
| 16022 | *AAK1* | 0.73215 | -0.63152 | -0.85536 | -1.5598 | -0.25 |
| 16023 | *C1orf185* | 0.25636 | -0.30763 | -0.70442 | -1.1698 | -0.25 |
| 16024 | *C12orf73* | 0.97085 | -0.23418 | -1.4951 | -2.1798 | -0.25 |
| 16025 | *NDRG4* | -0.12786 | -0.26314 | -0.36774 | -0.80595 | -0.25 |
| 16026 | *FARSB* | 0.45221 | -0.35272 | -0.86047 | -1.106 | -0.25 |
| 16027 | *ITM2C* | 0.43259 | 0.27334 | -1.4682 | -1.8191 | -0.25 |
| 16028 | *ADCY3* | -0.20512 | -0.24234 | -0.31671 | -0.44353 | -0.25 |
| 16029 | *MRPS18A* | 0.15937 | 0.024107 | -0.94841 | -1.8587 | -0.25 |
| 16030 | *SCCPDH* | 0.89539 | -0.44353 | -1.2171 | -2.071 | -0.26 |
| 16031 | *MRPL21* | 0.54988 | 0.024885 | -1.341 | -1.5759 | -0.26 |
| 16032 | *LINGO3* | 3.054 | -1.4852 | -2.3357 | -2.7056 | -0.26 |
| 16033 | *COPS3* | 1.3404 | -0.54784 | -1.5598 | -2.2631 | -0.26 |
| 16034 | *SLC25A41* | 0.39528 | 0.12206 | -1.2852 | -1.89 | -0.26 |
| 16035 | *HIC2* | 1.2967 | -0.94393 | -1.1207 | -1.9023 | -0.26 |
| 16036 | *CDH9* | 0.75145 | -0.55365 | -0.96622 | -1.8562 | -0.26 |
| 16037 | *PPCDC* | 1.3109 | -0.17105 | -1.9099 | -2.1319 | -0.26 |
| 16038 | *TRIM13* | 0.20047 | -0.39313 | -0.5775 | -2.1616 | -0.26 |
| 16039 | *MCOLN2* | 1.0339 | -0.80897 | -0.9952 | -1.1108 | -0.26 |
| 16040 | *DCANP1* | 0.36231 | 0.2474 | -1.3802 | -1.3814 | -0.26 |
| 16041 | *ANKFY1* | 0.36882 | -0.40118 | -0.73853 | -0.91842 | -0.26 |
| 16042 | *SLBP* | 0.12307 | -0.27453 | -0.61946 | -1.1095 | -0.26 |
| 16043 | *KRT14* | 1.1971 | -0.83074 | -1.141 | -1.478 | -0.26 |
| 16044 | *LOC101927322* | 0.53005 | -0.58289 | -0.72304 | -2.462 | -0.26 |
| 16045 | *C17orf59* | -0.06896 | -0.24555 | -0.46205 | -0.71077 | -0.26 |
| 16046 | *SLC22A9* | 0.55552 | 0.34367 | -1.6759 | -1.9326 | -0.26 |
| 16047 | *FAM120C* | 0.21479 | -0.43572 | -0.55586 | -0.9225 | -0.26 |

| | | | | | | |
|---|---|---|---|---|---|---|
| 16048 | *PDE3B* | 1.2827 | -0.44711 | -1.6129 | -2.8288 | -0.26 |
| 16049 | *RBM12* | 0.29479 | 0.010038 | -1.0824 | -1.5711 | -0.26 |
| 16050 | *OR2A2* | 0.43459 | -0.5584 | -0.65402 | -0.75192 | -0.26 |
| 16051 | *PTGS1* | 0.35451 | 0.06506 | -1.1977 | -1.8537 | -0.26 |
| 16052 | *KANSL3* | -0.19422 | -0.2748 | -0.31004 | -0.92868 | -0.26 |
| 16053 | *SF3B5* | 0.26579 | -0.12908 | -0.91617 | -2.1158 | -0.26 |
| 16054 | *MADCAM1* | 0.30681 | -0.04162 | -1.045 | -1.1704 | -0.26 |
| 16055 | *IAH1* | 2.1088 | -1.3218 | -1.5671 | -1.589 | -0.26 |
| 16056 | *TTBK1* | 0.04816 | -0.39872 | -0.43016 | -1.5901 | -0.26 |
| 16057 | *ZNF746* | 0.9791 | -0.46733 | -1.2936 | -1.8844 | -0.26 |
| 16058 | *PRSS50* | 0.62321 | -0.53815 | -0.86861 | -1.7829 | -0.26 |
| 16059 | *MYL4* | 0.6885 | -0.5639 | -0.90865 | -2.3729 | -0.26 |
| 16060 | *ZYG11A* | 1.2016 | -0.72214 | -1.2647 | -1.515 | -0.26 |
| 16061 | *SLC38A6* | 0.53063 | 0.32916 | -1.6456 | -2.0908 | -0.26 |
| 16062 | *ATP13A1* | 0.017417 | -0.10071 | -0.70418 | -0.99137 | -0.26 |
| 16063 | *NRG2* | 1.1285 | -0.6057 | -1.3107 | -2.0267 | -0.26 |
| 16064 | *MRPL18* | 1.798 | -0.81217 | -1.7766 | -1.872 | -0.26 |
| 16065 | *MALL* | 2.4948 | -1.129 | -2.1566 | -3.3367 | -0.26 |
| 16066 | *WBSCR16* | 0.50129 | 0.17112 | -1.4633 | -1.8157 | -0.26 |
| 16067 | *GNAL* | 0.62146 | 0.028715 | -1.4429 | -1.7077 | -0.26 |
| 16068 | *EYA4* | 0.33676 | -0.46766 | -0.66215 | -1.526 | -0.26 |
| 16069 | *FOS* | 1.184 | -0.51653 | -1.4607 | -1.5212 | -0.26 |
| 16070 | *FOXD2* | 0.71474 | -0.58136 | -0.92711 | -1.2647 | -0.26 |
| 16071 | *ZC3H12D* | 0.44778 | -0.49564 | -0.74594 | -0.91944 | -0.26 |
| 16072 | *VPS35* | 0.66502 | 0.00962 | -1.4686 | -2.1125 | -0.26 |
| 16073 | *ZMYND11* | 0.91832 | -0.22763 | -1.4852 | -1.5866 | -0.26 |
| 16074 | *FGA* | 0.39573 | -0.59183 | -0.59871 | -1.1193 | -0.26 |
| 16075 | *MPV17L* | 0.57613 | -0.19146 | -1.1796 | -1.8884 | -0.26 |
| 16076 | *TSC2* | 0.88521 | -0.46743 | -1.2133 | -2.1239 | -0.27 |
| 16077 | *CHCHD4* | 0.16266 | 0.096388 | -1.0557 | -1.1101 | -0.27 |
| 16078 | *OR10J5* | 0.72192 | -0.53078 | -0.9879 | -1.03 | -0.27 |
| 16079 | *BGN* | 0.39098 | -0.49045 | -0.69759 | -0.91253 | -0.27 |
| 16080 | *MTNR1B* | -0.17696 | -0.30424 | -0.31641 | -1.2694 | -0.27 |
| 16081 | *UMOD* | 0.49885 | -0.35135 | -0.94546 | -1.1918 | -0.27 |
| 16082 | *COG4* | 0.15458 | -0.23252 | -0.72085 | -2.1342 | -0.27 |
| 16083 | *EHMT2* | 2.0137 | -1.2244 | -1.5883 | -2.2059 | -0.27 |
| 16084 | *ABHD12B* | 1.0757 | 0.012183 | -1.8885 | -2.1696 | -0.27 |
| 16085 | *EML2* | 0.79519 | -0.33321 | -1.2632 | -1.7783 | -0.27 |
| 16086 | *NPIPA3* | 0.61063 | -0.62306 | -0.78959 | -1.5415 | -0.27 |
| 16087 | *ADH1A* | -0.09079 | -0.09716 | -0.61421 | -1.0273 | -0.27 |
| 16088 | *RAD51* | 0.43339 | 0.36658 | -1.6023 | -2.7504 | -0.27 |
| 16089 | *DDX58* | -0.13997 | -0.21899 | -0.44353 | -1.1226 | -0.27 |
| 16090 | *QPRT* | 0.58777 | -0.62306 | -0.76755 | -1.1417 | -0.27 |
| 16091 | *TMEM79* | 0.2464 | -0.38901 | -0.6608 | -2.2507 | -0.27 |
| 16092 | *SKIV2L* | 1.6077 | -0.79378 | -1.6176 | -1.8238 | -0.27 |
| 16093 | *CLCA1* | 0.67583 | -0.3098 | -1.1704 | -2.0116 | -0.27 |

| | | | | | | |
|---|---|---|---|---|---|---|
| 16094 | *C22orf46* | 0.60353 | -0.6176 | -0.79035 | -1.0248 | -0.27 |
| 16095 | *C19orf44* | 1.4922 | -1.0803 | -1.2169 | -2.2866 | -0.27 |
| 16096 | *BOLA2B* | 0.27164 | -0.50961 | -0.568 | -1.0743 | -0.27 |
| 16097 | *RPL39L* | 1.7014 | -1.2411 | -1.2663 | -1.3068 | -0.27 |
| 16098 | *CHRNB1* | 0.60357 | 0.016186 | -1.4269 | -1.5671 | -0.27 |
| 16099 | *PRIM2* | 0.63873 | -0.60717 | -0.84042 | -1.3542 | -0.27 |
| 16100 | *TPST1* | 0.43282 | -0.06809 | -1.1743 | -2.2704 | -0.27 |
| 16101 | *CPSF3* | 1.3932 | -0.17653 | -2.0277 | -2.3795 | -0.27 |
| 16102 | *UCHL3* | 2.0351 | -1.0685 | -1.7782 | -1.8996 | -0.27 |
| 16103 | *HMGCS1* | 0.022717 | -0.40693 | -0.42743 | -1.6278 | -0.27 |
| 16104 | *ZSCAN12* | 0.42414 | -0.31916 | -0.9168 | -1.8707 | -0.27 |
| 16105 | *SLC35D2* | 1.1909 | -0.84419 | -1.1591 | -1.7266 | -0.27 |
| 16106 | *MCOLN3* | 0.93056 | -0.3744 | -1.3694 | -1.8435 | -0.27 |
| 16107 | *CCDC94* | 1.3059 | -0.65873 | -1.4613 | -1.9655 | -0.27 |
| 16108 | *COL13A1* | 0.10453 | -0.06896 | -0.85006 | -1.2699 | -0.27 |
| 16109 | *TAOK1* | 0.90484 | -0.81778 | -0.90411 | -1.1914 | -0.27 |
| 16110 | *CLPP* | 1.2712 | -1.0425 | -1.0467 | -1.8856 | -0.27 |
| 16111 | *CGN* | 0.69256 | -0.20778 | -1.3058 | -1.3199 | -0.27 |
| 16112 | *TG* | 0.4261 | 0.31889 | -1.5671 | -1.6796 | -0.27 |
| 16113 | *CA7* | 0.26451 | -0.38425 | -0.70391 | -1.0091 | -0.27 |
| 16114 | *C10orf55* | 0.094966 | -0.44133 | -0.47739 | -2.1648 | -0.27 |
| 16115 | *DDX46* | 0.5061 | -0.32637 | -1.0044 | -1.5101 | -0.27 |
| 16116 | *MS4A6A* | 1.2187 | -0.78229 | -1.2616 | -1.8975 | -0.28 |
| 16117 | *SPZ1* | 0.38954 | -0.0227 | -1.1945 | -2.1438 | -0.28 |
| 16118 | *CFAP20* | 1.004 | -0.91388 | -0.91807 | -1.8538 | -0.28 |
| 16119 | *MT1A* | 0.25848 | -0.04149 | -1.0452 | -2.0323 | -0.28 |
| 16120 | *PI3* | 0.066909 | -0.2735 | -0.62213 | -2.3366 | -0.28 |
| 16121 | *CAPN14* | 0.19282 | 0.047417 | -1.0691 | -1.7134 | -0.28 |
| 16122 | *VPS9D1* | 0.80717 | -0.12421 | -1.5121 | -3.0804 | -0.28 |
| 16123 | *UBA3* | 1.234 | -0.75934 | -1.3042 | -2.1113 | -0.28 |
| 16124 | *PCNA* | 0.39081 | -0.53619 | -0.68439 | -1.0252 | -0.28 |
| 16125 | *UBIAD1* | 0.89745 | -0.58229 | -1.1453 | -1.6216 | -0.28 |
| 16126 | *RNF7* | 0.39738 | -0.0464 | -1.1812 | -2.5655 | -0.28 |
| 16127 | *TUBA3E* | 0.75365 | -0.73853 | -0.8458 | -2.3601 | -0.28 |
| 16128 | *EPB42* | 0.43061 | -0.15339 | -1.1084 | -1.7486 | -0.28 |
| 16129 | *UFM1* | 0.62144 | -0.5635 | -0.8892 | -2.2485 | -0.28 |
| 16130 | *C22orf34* | 1.11 | -0.28464 | -1.659 | -1.7766 | -0.28 |
| 16131 | *SIRPG* | -0.16518 | -0.20106 | -0.46755 | -1.8782 | -0.28 |
| 16132 | *GLYCTK* | -0.12922 | -0.3485 | -0.36106 | -0.46769 | -0.28 |
| 16133 | *KALRN* | 0.23309 | 0.17311 | -1.245 | -1.4998 | -0.28 |
| 16134 | *ST6GAL2* | 0.023344 | -0.24441 | -0.61779 | -1.2705 | -0.28 |
| 16135 | *TMEM132B* | 0.39912 | 0.19494 | -1.4333 | -2.2021 | -0.28 |
| 16136 | *FIP1L1* | 0.1939 | -0.02913 | -1.0046 | -1.8683 | -0.28 |
| 16137 | *TMEM41B* | 0.18536 | -0.10664 | -0.91921 | -1.6678 | -0.28 |
| 16138 | *DKK4* | 1.1594 | -0.95802 | -1.0421 | -1.1027 | -0.28 |
| 16139 | *TROAP* | 0.65949 | -0.50555 | -0.99474 | -2.7852 | -0.28 |

| | | | | | | |
|---|---|---|---|---|---|---|
| 16140 | *SEPT4* | 0.79577 | -0.49777 | -1.1393 | -1.7959 | -0.28 |
| 16141 | *ASB6* | 1.3646 | -0.92375 | -1.2822 | -4.0662 | -0.28 |
| 16142 | *FBN2* | -0.11327 | -0.28588 | -0.44353 | -1.773 | -0.28 |
| 16143 | *SLX1A* | 0.63031 | 0.1959 | -1.6693 | -2.012 | -0.28 |
| 16144 | *IL9* | 0.3946 | -0.22131 | -1.0164 | -1.3743 | -0.28 |
| 16145 | *WARS* | 0.427 | -0.44353 | -0.82733 | -2.0952 | -0.28 |
| 16146 | *CD101* | 0.089763 | -0.39313 | -0.54177 | -1.0016 | -0.28 |
| 16147 | *UBA7* | -0.14513 | -0.24738 | -0.45296 | -1.0782 | -0.28 |
| 16148 | *TRAPPC8* | 1.0382 | -0.33139 | -1.5524 | -1.6105 | -0.28 |
| 16149 | *NUP98* | 0.069589 | -0.37658 | -0.53957 | -1.1349 | -0.28 |
| 16150 | *CSF3R* | 0.25439 | -0.27892 | -0.82208 | -0.97053 | -0.28 |
| 16151 | *CARD9* | 0.72936 | 0.54151 | -2.1179 | -2.4322 | -0.28 |
| 16152 | *TSPAN14* | 0.70459 | -0.63953 | -0.91335 | -1.3574 | -0.28 |
| 16153 | *AKAP8L* | 1.1002 | -0.67146 | -1.2786 | -1.448 | -0.28 |
| 16154 | *AGAP4* | 0.80466 | 0.027886 | -1.683 | -2.4174 | -0.28 |
| 16155 | *WASL* | -0.01504 | -0.20103 | -0.63572 | -1.584 | -0.28 |
| 16156 | *C10orf111* | 0.42428 | 0.23329 | -1.5099 | -1.5485 | -0.28 |
| 16157 | *RBM28* | 0.2824 | -0.3316 | -0.80322 | -1.0695 | -0.28 |
| 16158 | *FOXN4* | 0.68979 | -0.12054 | -1.4218 | -1.445 | -0.28 |
| 16159 | *MMP21* | 2.1152 | -1.2297 | -1.739 | -1.908 | -0.28 |
| 16160 | *MME* | 0.16268 | -0.23484 | -0.78229 | -2.1505 | -0.28 |
| 16161 | *LOH12CR1* | 1.1221 | -0.56053 | -1.4174 | -1.8768 | -0.29 |
| 16162 | *SF3A3* | 0.76029 | -0.32815 | -1.2884 | -1.5663 | -0.29 |
| 16163 | *TEX10* | 0.80788 | -0.32087 | -1.3433 | -1.6469 | -0.29 |
| 16164 | *PRND* | 0.29464 | -0.12446 | -1.0273 | -1.7306 | -0.29 |
| 16165 | *C2orf61* | 0.75467 | 0.32461 | -1.9367 | -2.5852 | -0.29 |
| 16166 | *FBXO9* | 0.54251 | -0.66616 | -0.73389 | -0.91944 | -0.29 |
| 16167 | *DEFB118* | 0.38683 | -0.42004 | -0.82435 | -1.3976 | -0.29 |
| 16168 | *VOPP1* | 1.2857 | -0.42671 | -1.7179 | -2.7465 | -0.29 |
| 16169 | *GPC4* | 0.27286 | -0.5596 | -0.57235 | -1.0486 | -0.29 |
| 16170 | *CIDEB* | 0.89964 | -0.84659 | -0.91392 | -1.9065 | -0.29 |
| 16171 | *USP37* | 0.56773 | -0.54872 | -0.88008 | -0.98468 | -0.29 |
| 16172 | *CCK* | 0.47661 | -0.4849 | -0.85558 | -1.7911 | -0.29 |
| 16173 | *PLEKHJ1* | 0.29955 | -0.57899 | -0.58542 | -0.63255 | -0.29 |
| 16174 | *SPIC* | 0.92462 | -0.77418 | -1.0156 | -1.1797 | -0.29 |
| 16175 | *SFSWAP* | 1.258 | -0.9374 | -1.187 | -1.9075 | -0.29 |
| 16176 | *PAXBP1* | 0.37296 | -0.32059 | -0.91886 | -1.1756 | -0.29 |
| 16177 | *NPIPA1* | 0.51436 | 0.31379 | -1.6952 | -1.7332 | -0.29 |
| 16178 | *C1orf53* | 1.7798 | -0.91921 | -1.729 | -1.9404 | -0.29 |
| 16179 | *SNAP23* | 1.4044 | -0.46757 | -1.8057 | -2.3118 | -0.29 |
| 16180 | *TAMM41* | 1.184 | -0.97454 | -1.0793 | -1.5415 | -0.29 |
| 16181 | *MSTO1* | 0.23329 | -0.37282 | -0.73202 | -1.9326 | -0.29 |
| 16182 | *LOC100129216* | -0.01438 | -0.33511 | -0.52269 | -1.7552 | -0.29 |
| 16183 | *IRF7* | 0.37558 | -0.12408 | -1.1241 | -1.4299 | -0.29 |
| 16184 | *CRIP3* | 1.924 | -1.3288 | -1.4688 | -2.9977 | -0.29 |
| 16185 | *MAP1B* | 0.18932 | -0.14536 | -0.91779 | -1.8487 | -0.29 |

| | | | | | | |
|---|---|---|---|---|---|---|
| 16186 | *SLC46A3* | 0.41231 | -0.32193 | -0.96422 | -2.907 | -0.29 |
| 16187 | *HSD17B7* | -0.01125 | -0.41987 | -0.44283 | -1.4702 | -0.29 |
| 16188 | *COLEC11* | 1.2991 | -0.88898 | -1.285 | -1.676 | -0.29 |
| 16189 | *ZHX3* | 0.57416 | -0.60083 | -0.85002 | -1.2792 | -0.29 |
| 16190 | *LRIT3* | 1.0701 | -0.78651 | -1.1608 | -2.224 | -0.29 |
| 16191 | *PARG* | 0.39868 | -0.44855 | -0.82757 | -2.5725 | -0.29 |
| 16192 | *FYTTD1* | 0.028892 | -0.01088 | -0.8956 | -0.9797 | -0.29 |
| 16193 | *MEOX2* | -0.09032 | -0.21974 | -0.56753 | -0.61999 | -0.29 |
| 16194 | *AQP11* | 2.4178 | -1.3955 | -1.8999 | -2.1294 | -0.29 |
| 16195 | *KCNAB1* | 0.46044 | 0.08661 | -1.426 | -1.6984 | -0.29 |
| 16196 | *OTOL1* | 0.032952 | -0.12664 | -0.78812 | -1.0088 | -0.29 |
| 16197 | *CDK19* | 0.62164 | -0.7162 | -0.78754 | -1.1183 | -0.29 |
| 16198 | *GEMIN6* | 0.23955 | -0.05944 | -1.0626 | -3.1317 | -0.29 |
| 16199 | *FAM110D* | 0.81061 | -0.78705 | -0.90635 | -2.1816 | -0.29 |
| 16200 | *TMC3* | 1.0165 | -0.93712 | -0.96225 | -1.15 | -0.29 |
| 16201 | *FAM222A* | 0.95631 | -0.89167 | -0.94797 | -1.9175 | -0.29 |
| 16202 | *BCKDHB* | 0.68522 | -0.24488 | -1.3237 | -2.0872 | -0.29 |
| 16203 | *C2CD4D* | 0.79175 | -0.82663 | -0.84903 | -1.5637 | -0.29 |
| 16204 | *KIF26A* | 0.36658 | -0.27566 | -0.97488 | -2.1398 | -0.29 |
| 16205 | *DBI* | 0.33951 | -0.00827 | -1.2159 | -1.6913 | -0.29 |
| 16206 | *ENTPD4* | 0.36048 | -0.16202 | -1.0838 | -1.1002 | -0.30 |
| 16207 | *VSTM2B* | 1.7298 | -0.93005 | -1.6855 | -3.2873 | -0.30 |
| 16208 | *CENPBD1* | 0.44354 | -0.50419 | -0.82547 | -2.0541 | -0.30 |
| 16209 | *UBE2B* | 0.12823 | -0.43385 | -0.58143 | -0.77597 | -0.30 |
| 16210 | *TMEM55A* | 2.8225 | -1.4767 | -2.2329 | -2.2385 | -0.30 |
| 16211 | *BOLA3* | 0.48714 | -0.44353 | -0.93107 | -2.6208 | -0.30 |
| 16212 | *BTNL10* | 0.84346 | -0.52017 | -1.2114 | -1.6089 | -0.30 |
| 16213 | *NPM2* | 0.19529 | -0.52552 | -0.5584 | -1.3785 | -0.30 |
| 16214 | *TOLLIP* | 0.012276 | -0.11781 | -0.78439 | -1.244 | -0.30 |
| 16215 | *CLSTN1* | 1.1087 | -0.78229 | -1.2171 | -2.0534 | -0.30 |
| 16216 | *NOP10* | 0.18216 | -0.28158 | -0.79409 | -2.0305 | -0.30 |
| 16217 | *NDUFB8* | 0.45939 | -0.25181 | -1.1012 | -1.5824 | -0.30 |
| 16218 | *CLGN* | 1.4088 | -0.72553 | -1.578 | -1.6635 | -0.30 |
| 16219 | *PROX2* | 1.6064 | -1.0094 | -1.4919 | -2.4703 | -0.30 |
| 16220 | *IL21R* | 1.348 | -1.0801 | -1.163 | -1.6007 | -0.30 |
| 16221 | *KIAA2012* | 0.83416 | -0.84446 | -0.88702 | -1.3677 | -0.30 |
| 16222 | *EIF3G* | 0.99046 | -0.53868 | -1.35 | -1.5962 | -0.30 |
| 16223 | *LRRK2* | 0.044791 | -0.35379 | -0.58994 | -1.6943 | -0.30 |
| 16224 | *RNF13* | 0.29824 | -0.43077 | -0.76671 | -0.80641 | -0.30 |
| 16225 | *ARHGAP40* | 0.61785 | -0.56251 | -0.95469 | -2.101 | -0.30 |
| 16226 | *SLC39A11* | 0.22467 | -0.44185 | -0.68228 | -1.0761 | -0.30 |
| 16227 | *SNRNP35* | 1.539 | -0.57515 | -1.8636 | -2.8687 | -0.30 |
| 16228 | *FAM76A* | 0.28149 | 0.21653 | -1.3978 | -1.7129 | -0.30 |
| 16229 | *OR52N4* | 0.7011 | -0.20338 | -1.398 | -2.8521 | -0.30 |
| 16230 | *SFN* | 0.80855 | 0.67632 | -2.3858 | -3.3598 | -0.30 |
| 16231 | *ZNF71* | 0.85867 | -0.28384 | -1.4773 | -1.5901 | -0.30 |

| | | | | | | |
|---|---|---|---|---|---|---|
| 16232 | *SWSAP1* | 1.1552 | -0.32286 | -1.7352 | -2.8702 | -0.30 |
| 16233 | *ZNF565* | 0.11164 | -0.36295 | -0.65163 | -1.5073 | -0.30 |
| 16234 | *DYNLL1* | 0.15852 | -0.38796 | -0.67458 | -1.3183 | -0.30 |
| 16235 | *SACM1L* | 0.3445 | 0.023658 | -1.2728 | -1.8587 | -0.30 |
| 16236 | *CPXCR1* | 0.14757 | -0.45455 | -0.59829 | -2.1363 | -0.30 |
| 16237 | *KIR2DS4* | -0.19996 | -0.21868 | -0.48673 | -2.0698 | -0.30 |
| 16238 | *AGAP5* | 0.092716 | 0.063089 | -1.0614 | -1.2288 | -0.30 |
| 16239 | *JDP2* | 0.1468 | 0.090093 | -1.1442 | -2.0303 | -0.30 |
| 16240 | *GFI1* | 0.15006 | -0.26008 | -0.79744 | -1.0385 | -0.30 |
| 16241 | *LYPD2* | 0.82269 | -0.28524 | -1.4452 | -1.5667 | -0.30 |
| 16242 | *STAB2* | 0.53673 | -0.20621 | -1.2383 | -2.4642 | -0.30 |
| 16243 | *POLR3C* | 0.77012 | -0.44353 | -1.2352 | -1.4567 | -0.30 |
| 16244 | *PRRX2* | 0.50404 | 0.018621 | -1.4319 | -1.7829 | -0.30 |
| 16245 | *HIBCH* | 1.1873 | -0.95423 | -1.1426 | -1.4509 | -0.30 |
| 16246 | *DUSP3* | 0.34343 | -0.4599 | -0.79357 | -0.96997 | -0.30 |
| 16247 | *DPPA4* | 1.2368 | -0.43694 | -1.7113 | -1.7595 | -0.30 |
| 16248 | *PSD4* | 0.11596 | -0.44935 | -0.57825 | -3.596 | -0.30 |
| 16249 | *SRGAP2* | 1.358 | -1.0452 | -1.2246 | -2.5513 | -0.30 |
| 16250 | *POTEB* | 1.0218 | -0.93878 | -0.99508 | -1.0739 | -0.30 |
| 16251 | *HIGD2A* | -0.01823 | -0.28628 | -0.60781 | -1.2806 | -0.30 |
| 16252 | *PDSS2* | -0.0813 | -0.19706 | -0.63542 | -1.7493 | -0.30 |
| 16253 | *C7orf72* | 0.098451 | -0.08213 | -0.93011 | -2.0247 | -0.30 |
| 16254 | *CCNH* | 0.3998 | 0.21744 | -1.5312 | -2.0301 | -0.30 |
| 16255 | *PRR20C* | 2.2323 | -1.3973 | -1.7493 | -1.8363 | -0.30 |
| 16256 | *MICALCL* | 0.38855 | 0.00962 | -1.3134 | -3.0112 | -0.31 |
| 16257 | *EMC8* | 0.38037 | -0.46757 | -0.82928 | -1.5737 | -0.31 |
| 16258 | *SAA1* | 0.86319 | -0.40749 | -1.3727 | -1.4299 | -0.31 |
| 16259 | *ISG20L2* | 0.062111 | 0.008065 | -0.98719 | -1.1848 | -0.31 |
| 16260 | *HM13* | 2.5037 | -1.6477 | -1.7734 | -1.7994 | -0.31 |
| 16261 | *WWOX* | 0.62769 | -0.45403 | -1.0912 | -1.7438 | -0.31 |
| 16262 | *GDF2* | 0.62919 | -0.15145 | -1.3955 | -1.8154 | -0.31 |
| 16263 | *ACTN4* | 0.53258 | -0.14193 | -1.309 | -1.5633 | -0.31 |
| 16264 | *CXCL12* | 0.074609 | -0.16382 | -0.82959 | -1.3836 | -0.31 |
| 16265 | *SRSF3* | 0.4521 | -0.26518 | -1.1061 | -3.266 | -0.31 |
| 16266 | *SFPQ* | 0.20665 | -0.48744 | -0.64149 | -0.93399 | -0.31 |
| 16267 | *RTP2* | 0.24514 | -0.56701 | -0.60147 | -2.9736 | -0.31 |
| 16268 | *OR4K15* | 1.8524 | -1.0474 | -1.7297 | -2.5535 | -0.31 |
| 16269 | *ZNF185* | 0.088767 | 0.087566 | -1.1016 | -1.8572 | -0.31 |
| 16270 | *ANXA11* | 0.54013 | -0.63172 | -0.83402 | -1.8478 | -0.31 |
| 16271 | *C11orf57* | 0.73895 | -0.49266 | -1.1727 | -1.8095 | -0.31 |
| 16272 | *TMEM106C* | 0.75771 | -0.42766 | -1.2568 | -1.3008 | -0.31 |
| 16273 | *COMT* | -0.04927 | -0.08284 | -0.79467 | -2.4328 | -0.31 |
| 16274 | *DET1* | 0.94612 | -0.81555 | -1.0576 | -1.4951 | -0.31 |
| 16275 | *RING1* | 0.50349 | 0.33404 | -1.7653 | -1.7865 | -0.31 |
| 16276 | *WRNIP1* | 1.1212 | -0.55992 | -1.4893 | -2.528 | -0.31 |
| 16277 | *SIGLEC6* | 2.2677 | -1.3892 | -1.8076 | -2.2125 | -0.31 |

| | | | | | | |
|---|---|---|---|---|---|---|
| 16278 | *EXOC3* | 0.12995 | -0.51152 | -0.54829 | -1.2418 | -0.31 |
| 16279 | *EIF2AK3* | 0.47721 | -0.29304 | -1.1145 | -1.9494 | -0.31 |
| 16280 | *HOXB4* | 0.043861 | 0.016744 | -0.9925 | -1.7215 | -0.31 |
| 16281 | *TAS2R9* | 0.55792 | -0.49926 | -0.99175 | -1.0952 | -0.31 |
| 16282 | *RPL3* | 0.05102 | -0.16927 | -0.81501 | -1.3593 | -0.31 |
| 16283 | *SLC35D1* | 0.53795 | -0.72095 | -0.7512 | -2.1732 | -0.31 |
| 16284 | *SPDEF* | 0.2878 | -0.52914 | -0.69287 | -2.1151 | -0.31 |
| 16285 | *DCTN5* | 0.056779 | -0.41922 | -0.5718 | -1.4642 | -0.31 |
| 16286 | *ADH1B* | 0.6399 | -0.78444 | -0.79065 | -1.7649 | -0.31 |
| 16287 | *SPRED2* | 0.58614 | -0.27088 | -1.251 | -2.2879 | -0.31 |
| 16288 | *TMEM252* | 0.63528 | 0.37603 | -1.9484 | -2.6113 | -0.31 |
| 16289 | *DHODH* | 1.0293 | -0.86941 | -1.0989 | -1.1496 | -0.31 |
| 16290 | *RPS29* | 1.4184 | -0.76955 | -1.5886 | -1.9334 | -0.31 |
| 16291 | *FANCD2* | -0.1851 | -0.37185 | -0.38296 | -0.57564 | -0.31 |
| 16292 | *ITGA3* | -0.02404 | -0.32638 | -0.58971 | -2.0667 | -0.31 |
| 16293 | *RFX6* | 1.8289 | -1.3008 | -1.4686 | -1.9624 | -0.31 |
| 16294 | *SAMD13* | 0.54936 | 0.16073 | -1.6523 | -1.9495 | -0.31 |
| 16295 | *USP17L5* | -0.1096 | -0.38996 | -0.44277 | -1.2684 | -0.31 |
| 16296 | *SH3BP5* | 0.089278 | 0.031743 | -1.0682 | -1.1592 | -0.32 |
| 16297 | *PSD2* | 0.78013 | -0.50251 | -1.2249 | -1.5608 | -0.32 |
| 16298 | *ASCC2* | 0.45883 | -0.21182 | -1.1943 | -1.2432 | -0.32 |
| 16299 | *DXO* | 0.18719 | -0.56769 | -0.56841 | -2.5292 | -0.32 |
| 16300 | *MEGF10* | 2.3141 | -1.5227 | -1.7404 | -1.9891 | -0.32 |
| 16301 | *ERF* | 0.4139 | -0.43473 | -0.92828 | -1.3074 | -0.32 |
| 16302 | *SPRY1* | -0.16294 | -0.34327 | -0.44353 | -0.87858 | -0.32 |
| 16303 | *PRPF31* | -0.12067 | -0.16438 | -0.66569 | -1.0594 | -0.32 |
| 16304 | *IK* | -0.12659 | -0.36371 | -0.46107 | -1.4792 | -0.32 |
| 16305 | *COLCA2* | 0.90934 | -0.50067 | -1.3604 | -1.6754 | -0.32 |
| 16306 | *TNFRSF25* | -0.00621 | -0.056 | -0.88987 | -0.94897 | -0.32 |
| 16307 | *GLTSCR1L* | -0.01742 | -0.19943 | -0.73961 | -2.5795 | -0.32 |
| 16308 | *NR0B2* | 0.069884 | -0.1468 | -0.87997 | -1.2317 | -0.32 |
| 16309 | *C9orf92* | 0.446 | -0.29076 | -1.1132 | -1.8508 | -0.32 |
| 16310 | *ZCCHC12* | 1.5675 | -1.232 | -1.2935 | -1.9673 | -0.32 |
| 16311 | *RHOB* | 0.54804 | -0.55586 | -0.95045 | -1.5996 | -0.32 |
| 16312 | *SERINC3* | 0.58504 | -0.57899 | -0.96492 | -2.4405 | -0.32 |
| 16313 | *ITPA* | 1.2984 | -0.50588 | -1.7517 | -1.7663 | -0.32 |
| 16314 | *DEFB114* | 1.6823 | -0.84713 | -1.7945 | -1.9686 | -0.32 |
| 16315 | *STAU1* | -0.11035 | -0.39579 | -0.45323 | -1.8534 | -0.32 |
| 16316 | *TMEM65* | -0.11164 | -0.2875 | -0.5605 | -1.8275 | -0.32 |
| 16317 | *PTGES3* | -0.16321 | -0.33444 | -0.46236 | -0.47645 | -0.32 |
| 16318 | *BICD2* | 1.2281 | -1.0596 | -1.1309 | -1.4498 | -0.32 |
| 16319 | *NAPRT* | 0.46439 | -0.60604 | -0.82098 | -1.8297 | -0.32 |
| 16320 | *MLLT3* | 0.42665 | -0.45403 | -0.93555 | -1.1886 | -0.32 |
| 16321 | *GPD2* | 0.62605 | -0.64268 | -0.94723 | -1.5059 | -0.32 |
| 16322 | *ITGA4* | 0.71846 | -0.74774 | -0.93652 | -0.99529 | -0.32 |
| 16323 | *CCDC155* | 0.47086 | 0.15303 | -1.5898 | -2.0218 | -0.32 |

| | | | | | | |
|---|---|---|---|---|---|---|
| 16324 | *CT47A6* | 0.1695 | -0.21031 | -0.92553 | -1.7076 | -0.32 |
| 16325 | *CHURC1* | -0.11527 | -0.36451 | -0.48821 | -2.3096 | -0.32 |
| 16326 | *TMC7* | 0.25617 | -0.394 | -0.8303 | -1.7068 | -0.32 |
| 16327 | *EDEM2* | 0.52335 | -0.01629 | -1.4752 | -1.7743 | -0.32 |
| 16328 | *EFNA4* | 0.080715 | -0.01439 | -1.035 | -1.4841 | -0.32 |
| 16329 | *RFC2* | 0.10367 | -0.4216 | -0.65075 | -1.0089 | -0.32 |
| 16330 | *ACACA* | -0.06884 | -0.20886 | -0.69286 | -1.3299 | -0.32 |
| 16331 | *CD200R1L* | 1.0493 | -0.36035 | -1.6604 | -1.9681 | -0.32 |
| 16332 | *GOLGA6L1* | 0.56217 | -0.62493 | -0.9101 | -1.6525 | -0.32 |
| 16333 | *HPCA* | 1.2028 | -0.72566 | -1.4509 | -1.7753 | -0.32 |
| 16334 | *HIST3H2BB* | 0.8907 | -0.37376 | -1.4909 | -1.8804 | -0.32 |
| 16335 | *GLI2* | 0.82348 | -0.11406 | -1.6836 | -2.1419 | -0.32 |
| 16336 | *VPS37D* | 0.76556 | -0.39214 | -1.3478 | -1.8724 | -0.32 |
| 16337 | *ZKSCAN4* | 1.8963 | -1.2264 | -1.6443 | -2.1502 | -0.32 |
| 16338 | *HYPM* | 0.69988 | -0.73111 | -0.94345 | -1.539 | -0.32 |
| 16339 | *C21orf59* | 2.3762 | -1.3235 | -2.0277 | -2.2766 | -0.33 |
| 16340 | *TAX1BP3* | 0.28617 | -0.54219 | -0.71939 | -1.8087 | -0.33 |
| 16341 | *UNC50* | 0.43325 | -0.39883 | -1.0099 | -1.5125 | -0.33 |
| 16342 | *SLAMF8* | 3.551 | -2.1984 | -2.3296 | -2.4022 | -0.33 |
| 16343 | *CASP9* | 0.15435 | -0.33709 | -0.79444 | -1.5181 | -0.33 |
| 16344 | *CHST7* | 1.5139 | -1.1439 | -1.3492 | -1.9794 | -0.33 |
| 16345 | *TERF1* | 0.55396 | 0.055556 | -1.589 | -2.6289 | -0.33 |
| 16346 | *CCAR1* | 1.1428 | -0.72582 | -1.3971 | -1.5676 | -0.33 |
| 16347 | *NUP160* | 0.008382 | -0.40897 | -0.58131 | -1.925 | -0.33 |
| 16348 | *KREMEN2* | 0.23097 | -0.4055 | -0.80823 | -1.5235 | -0.33 |
| 16349 | *KRT26* | 0.079734 | -0.32308 | -0.73944 | -0.92427 | -0.33 |
| 16350 | *ITSN2* | 0.20545 | -0.37888 | -0.80946 | -1.0773 | -0.33 |
| 16351 | *TRPM7* | 0.68587 | -0.06007 | -1.6089 | -2.4328 | -0.33 |
| 16352 | *MRPS11* | 0.79156 | -0.86324 | -0.91215 | -1.6586 | -0.33 |
| 16353 | *CCR4* | 0.56934 | -0.14697 | -1.4087 | -1.696 | -0.33 |
| 16354 | *TADA1* | 0.087154 | -0.38265 | -0.69102 | -1.2443 | -0.33 |
| 16355 | *ZFYVE26* | 0.18281 | -0.38655 | -0.78313 | -1.7718 | -0.33 |
| 16356 | *CHTF8* | 0.73049 | -0.8424 | -0.8759 | -1.0501 | -0.33 |
| 16357 | *KRTAP5-5* | -0.09512 | -0.43526 | -0.45854 | -1.2867 | -0.33 |
| 16358 | *OVGP1* | 0.008624 | -0.3521 | -0.64546 | -1.3176 | -0.33 |
| 16359 | *NR2C2* | -0.06694 | -0.26246 | -0.6603 | -1.9653 | -0.33 |
| 16360 | *ATP5A1* | -0.20499 | -0.33404 | -0.45069 | -0.57273 | -0.33 |
| 16361 | *CMTM2* | 2.2596 | -1.6034 | -1.6463 | -1.9041 | -0.33 |
| 16362 | *TIGD5* | 0.47045 | 0.21577 | -1.677 | -1.7688 | -0.33 |
| 16363 | *SET* | 0.42171 | -0.42808 | -0.9858 | -2.024 | -0.33 |
| 16364 | *NEDD8* | -0.10255 | -0.36824 | -0.52238 | -1.5632 | -0.33 |
| 16365 | *PCED1B* | 0.25096 | -0.58771 | -0.65668 | -0.88695 | -0.33 |
| 16366 | *PHF23* | 0.83593 | -0.52587 | -1.305 | -1.5635 | -0.33 |
| 16367 | *KAT8* | 1.0472 | -0.86725 | -1.1754 | -2.2044 | -0.33 |
| 16368 | *CHRAC1* | 1.0312 | -0.43113 | -1.5971 | -2.9538 | -0.33 |
| 16369 | *COIL* | 0.23395 | -0.39606 | -0.83505 | -1.9245 | -0.33 |

| | | | | | | |
|---|---|---|---|---|---|---|
| 16370 | *NTN3* | 0.9446 | -0.71772 | -1.2244 | -1.6712 | -0.33 |
| 16371 | *SEC24D* | 0.44099 | -0.26186 | -1.1771 | -1.8453 | -0.33 |
| 16372 | *GTF2E2* | 0.4496 | -0.08887 | -1.3591 | -1.624 | -0.33 |
| 16373 | *FANCC* | 0.14948 | -0.0304 | -1.1184 | -1.3098 | -0.33 |
| 16374 | *SLFNL1* | -0.09229 | -0.11023 | -0.79744 | -2.264 | -0.33 |
| 16375 | *RBMY1J* | 0.5728 | 0.23438 | -1.8087 | -2.0593 | -0.33 |
| 16376 | *ZEB2* | 1.4096 | -0.06005 | -2.3517 | -3.1229 | -0.33 |
| 16377 | *ZP3* | 0.26956 | -0.6093 | -0.66242 | -1.6246 | -0.33 |
| 16378 | *RIMS4* | 0.18994 | -0.01705 | -1.1758 | -1.251 | -0.33 |
| 16379 | *DNAJC11* | 0.91916 | -0.71819 | -1.204 | -2.425 | -0.33 |
| 16380 | *OR6C3* | 0.84519 | -0.82978 | -1.0197 | -1.5118 | -0.33 |
| 16381 | *SYNJ2BP* | -0.31531 | -0.33136 | -0.35891 | -0.77678 | -0.34 |
| 16382 | *CLEC18C* | 2.4771 | -1.6885 | -1.7945 | -2.3876 | -0.34 |
| 16383 | *SERPINC1* | -0.09459 | -0.32087 | -0.59061 | -1.205 | -0.34 |
| 16384 | *TTYH2* | 0.38532 | -0.35537 | -1.0361 | -1.463 | -0.34 |
| 16385 | *C10orf128* | 0.69415 | -0.13984 | -1.5608 | -1.5651 | -0.34 |
| 16386 | *KPNA4* | 0.90732 | -0.53767 | -1.3766 | -1.9811 | -0.34 |
| 16387 | *CYB5D2* | 0.31268 | -0.57691 | -0.74308 | -1.8402 | -0.34 |
| 16388 | *TMEM164* | 0.5662 | -0.22577 | -1.3481 | -1.8996 | -0.34 |
| 16389 | *TEX261* | 1.7441 | -1.3078 | -1.4442 | -1.6065 | -0.34 |
| 16390 | *TLR5* | 0.44955 | -0.63643 | -0.82368 | -1.2049 | -0.34 |
| 16391 | *GIMAP5* | 0.78922 | -0.57513 | -1.2261 | -1.3296 | -0.34 |
| 16392 | *ZNF250* | 0.16588 | -0.27152 | -0.90663 | -1.3097 | -0.34 |
| 16393 | *DDI1* | 0.43912 | -0.55844 | -0.89351 | -2.5146 | -0.34 |
| 16394 | *HOXA4* | 0.68151 | -0.37913 | -1.3153 | -1.626 | -0.34 |
| 16395 | *EOMES* | 0.34044 | -0.22588 | -1.1283 | -1.5394 | -0.34 |
| 16396 | *AP2M1* | 2.0497 | -1.485 | -1.5787 | -1.6443 | -0.34 |
| 16397 | *SUPT5H* | 0.15694 | -0.39272 | -0.77828 | -1.468 | -0.34 |
| 16398 | *CYP8B1* | 1.0372 | -0.29759 | -1.7538 | -2.4013 | -0.34 |
| 16399 | *MAP1LC3C* | 0.84074 | -0.72488 | -1.1316 | -1.5817 | -0.34 |
| 16400 | *RNF144B* | 0.30391 | -0.48564 | -0.83471 | -1.3032 | -0.34 |
| 16401 | *VCAN* | 0.25826 | -0.61138 | -0.66353 | -0.93489 | -0.34 |
| 16402 | *FYN* | 1.8472 | -0.72348 | -2.1412 | -2.5725 | -0.34 |
| 16403 | *USP51* | 0.27655 | -0.56769 | -0.7264 | -2.033 | -0.34 |
| 16404 | *DYRK3* | 0.23111 | 0.073963 | -1.3232 | -2.0629 | -0.34 |
| 16405 | *GTF2I* | 0.84841 | -0.81448 | -1.0529 | -1.4741 | -0.34 |
| 16406 | *NKAP* | 0.49801 | -0.066 | -1.451 | -1.7718 | -0.34 |
| 16407 | *IFNLR1* | 0.86516 | -0.8606 | -1.0243 | -2.4885 | -0.34 |
| 16408 | *GNGT1* | 0.13498 | 0.048059 | -1.204 | -1.5524 | -0.34 |
| 16409 | *UBE2T* | -0.21502 | -0.33961 | -0.46768 | -0.70335 | -0.34 |
| 16410 | *MASP2* | 0.62656 | -0.2732 | -1.3758 | -1.5884 | -0.34 |
| 16411 | *BYSL* | 0.51887 | -0.19658 | -1.3453 | -5.4774 | -0.34 |
| 16412 | *ATP6V1B1* | 0.95235 | -0.76762 | -1.2079 | -1.4428 | -0.34 |
| 16413 | *PSMA6* | 0.56585 | -0.66865 | -0.92078 | -3.0638 | -0.34 |
| 16414 | *LRRFIP2* | 0.22714 | -0.56154 | -0.69006 | -0.74743 | -0.34 |
| 16415 | *ILF2* | 0.71371 | -0.75586 | -0.98359 | -1.6251 | -0.34 |

| | | | | | | |
|---|---|---|---|---|---|---|
| 16416 | *RHBDL2* | 0.57229 | -0.726 | -0.87318 | -1.2352 | -0.34 |
| 16417 | *FNTA* | 0.2663 | 0.048673 | -1.3427 | -2.1577 | -0.34 |
| 16418 | *UACA* | 0.28485 | -0.55909 | -0.75511 | -1.5952 | -0.34 |
| 16419 | *GJD2* | 0.14992 | -0.30427 | -0.87536 | -1.9461 | -0.34 |
| 16420 | *ALKBH1* | 0.22992 | -0.61092 | -0.64888 | -1.9166 | -0.34 |
| 16421 | *ELL* | 0.065253 | -0.25295 | -0.84291 | -2.0423 | -0.34 |
| 16422 | *AHI1* | -0.10031 | -0.43906 | -0.49314 | -1.3978 | -0.34 |
| 16423 | *PITX1* | 0.69518 | -0.01401 | -1.714 | -2.0949 | -0.34 |
| 16424 | *MAN1A1* | -0.07219 | -0.25056 | -0.71063 | -3.7736 | -0.34 |
| 16425 | *KIAA0895* | 1.3427 | -1.1851 | -1.1914 | -1.2867 | -0.34 |
| 16426 | *MS4A12* | 0.66479 | -0.60717 | -1.0918 | -1.806 | -0.34 |
| 16427 | *UNC93A* | 0.79216 | -0.64016 | -1.1867 | -1.3361 | -0.34 |
| 16428 | *DCST2* | 0.070673 | -0.20987 | -0.8959 | -0.99059 | -0.35 |
| 16429 | *DNAL4* | 0.50078 | 0.35177 | -1.8885 | -2.1523 | -0.35 |
| 16430 | *OR2M3* | 0.27103 | -0.06606 | -1.2411 | -2.404 | -0.35 |
| 16431 | *NUP210* | 0.093548 | -0.18996 | -0.94113 | -1.5894 | -0.35 |
| 16432 | *TPK1* | 0.3057 | -0.25261 | -1.0907 | -1.7646 | -0.35 |
| 16433 | *MARCH10* | 0.27469 | -0.26036 | -1.0528 | -1.3178 | -0.35 |
| 16434 | *ESR1* | 0.19262 | -0.38898 | -0.84563 | -1.4689 | -0.35 |
| 16435 | *UBA5* | 0.63153 | -0.29713 | -1.377 | -1.6262 | -0.35 |
| 16436 | *SLC30A8* | 0.83543 | -0.45214 | -1.4261 | -1.8702 | -0.35 |
| 16437 | *DND1* | 0.55843 | 0.42767 | -2.0303 | -2.2044 | -0.35 |
| 16438 | *F5* | -0.00834 | -0.16195 | -0.87418 | -1.072 | -0.35 |
| 16439 | *C14orf159* | 0.36517 | -0.67702 | -0.73288 | -1.3766 | -0.35 |
| 16440 | *PHACTR3* | 0.31269 | -0.53345 | -0.82573 | -1.5688 | -0.35 |
| 16441 | *HMOX2* | 2.2878 | -1.635 | -1.701 | -2.2007 | -0.35 |
| 16442 | *ANKEF1* | 0.27293 | -0.14918 | -1.1743 | -1.4734 | -0.35 |
| 16443 | *TRMT11* | -0.22097 | -0.33754 | -0.49212 | -1.0863 | -0.35 |
| 16444 | *SLC22A8* | 0.31543 | -0.00331 | -1.364 | -3.0246 | -0.35 |
| 16445 | *FGF9* | 0.086929 | -0.09384 | -1.0467 | -2.9031 | -0.35 |
| 16446 | *ZNF585A* | -0.23796 | -0.26237 | -0.55387 | -1.1383 | -0.35 |
| 16447 | *COA7* | -0.07018 | -0.11162 | -0.87318 | -2.1108 | -0.35 |
| 16448 | *POP4* | 0.38783 | 0.2086 | -1.6524 | -1.9099 | -0.35 |
| 16449 | *CYFIP2* | 0.49205 | -0.49953 | -1.0489 | -1.1207 | -0.35 |
| 16450 | *PHF12* | 0.19752 | -0.07341 | -1.181 | -1.7507 | -0.35 |
| 16451 | *COX7B* | -0.01088 | -0.03572 | -1.0107 | -1.5454 | -0.35 |
| 16452 | *HADH* | 1.005 | -0.88159 | -1.1821 | -1.5524 | -0.35 |
| 16453 | *NDNF* | 0.56345 | 0.19927 | -1.8219 | -2.3509 | -0.35 |
| 16454 | *GNG2* | 0.48883 | -0.62523 | -0.92415 | -0.99834 | -0.35 |
| 16455 | *CCDC114* | 0.43264 | 0.18259 | -1.6759 | -1.6921 | -0.35 |
| 16456 | *PLA2G4F* | 0.116 | -0.08253 | -1.0944 | -2.1085 | -0.35 |
| 16457 | *COQ7* | 0.1582 | 0.11813 | -1.3378 | -1.658 | -0.35 |
| 16458 | *TEFM* | 0.44887 | -0.66604 | -0.84627 | -1.0645 | -0.35 |
| 16459 | *FOXG1* | 0.73058 | -0.58752 | -1.2071 | -1.5679 | -0.35 |
| 16460 | *MRPL34* | 0.11219 | -0.42634 | -0.75092 | -0.99313 | -0.36 |
| 16461 | *VPS26A* | 0.035695 | -0.29549 | -0.80542 | -1.5073 | -0.36 |

| | | | | | | |
|---|---|---|---|---|---|---|
| 16462 | *RNF213* | 0.12019 | -0.26273 | -0.92288 | -1.6371 | -0.36 |
| 16463 | *MUS81* | 0.55956 | -0.78963 | -0.83692 | -0.97053 | -0.36 |
| 16464 | *C5orf45* | 0.71045 | -0.45569 | -1.3218 | -1.626 | -0.36 |
| 16465 | *SUGT1* | 0.44675 | -0.49408 | -1.0198 | -1.716 | -0.36 |
| 16466 | *AGMO* | 0.17128 | -0.38296 | -0.85594 | -1.3292 | -0.36 |
| 16467 | *CERS1* | 1.3549 | -1.2015 | -1.2214 | -1.6804 | -0.36 |
| 16468 | *BTBD11* | 0.21441 | -0.0536 | -1.229 | -1.6283 | -0.36 |
| 16469 | *CEBPZ* | 0.30216 | 0.28096 | -1.6525 | -1.7513 | -0.36 |
| 16470 | *TUFM* | 0.69659 | -0.59157 | -1.175 | -2.0553 | -0.36 |
| 16471 | *GOLGA3* | 0.45442 | -0.36914 | -1.1553 | -1.2145 | -0.36 |
| 16472 | *ZACN* | 0.95277 | -0.47338 | -1.5507 | -2.1532 | -0.36 |
| 16473 | *RIC8A* | 0.813 | -0.79453 | -1.0898 | -1.584 | -0.36 |
| 16474 | *FAM221A* | -0.05201 | -0.27903 | -0.74141 | -1.5383 | -0.36 |
| 16475 | *TNFRSF10D* | 0.5805 | -0.21796 | -1.4361 | -1.8714 | -0.36 |
| 16476 | *C19orf25* | 0.57154 | -0.60235 | -1.043 | -3.4256 | -0.36 |
| 16477 | *MTUS1* | -0.04425 | -0.17517 | -0.85513 | -0.9904 | -0.36 |
| 16478 | *TUBGCP4* | -0.15978 | -0.18791 | -0.72723 | -1.1687 | -0.36 |
| 16479 | *DLK1* | 0.33602 | -0.17606 | -1.2349 | -1.8636 | -0.36 |
| 16480 | *TSPY2* | 0.99812 | -0.65012 | -1.4233 | -1.7188 | -0.36 |
| 16481 | *PCDHB4* | -0.16202 | -0.36047 | -0.55302 | -1.1472 | -0.36 |
| 16482 | *SNX7* | 0.35388 | -0.27423 | -1.1553 | -1.4775 | -0.36 |
| 16483 | *SCAMP2* | 0.30373 | -0.13571 | -1.2439 | -2.3915 | -0.36 |
| 16484 | *ID1* | 0.41353 | 0.06977 | -1.5598 | -1.9326 | -0.36 |
| 16485 | *ATAD3B* | 0.029089 | -0.43854 | -0.66731 | -1.2301 | -0.36 |
| 16486 | *TEAD2* | -0.11216 | -0.27672 | -0.68813 | -2.6208 | -0.36 |
| 16487 | *ATAT1* | 0.26733 | -0.53712 | -0.80771 | -1.3148 | -0.36 |
| 16488 | *SHOC2* | 0.71674 | -0.83784 | -0.95708 | -1.4448 | -0.36 |
| 16489 | *ZC3H10* | 0.22414 | -0.56079 | -0.74192 | -0.98642 | -0.36 |
| 16490 | *LINGO2* | 0.8205 | -0.15499 | -1.7445 | -1.9041 | -0.36 |
| 16491 | *GPR132* | 0.19852 | -0.14348 | -1.1343 | -3.3935 | -0.36 |
| 16492 | *BAG6* | 1.0085 | -0.98452 | -1.1034 | -2.1952 | -0.36 |
| 16493 | *PTGR2* | 0.36257 | -0.07235 | -1.3701 | -1.9107 | -0.36 |
| 16494 | *RBMS3* | 0.35979 | -0.57787 | -0.86398 | -1.7079 | -0.36 |
| 16495 | *RYR1* | 1.5512 | -1.0819 | -1.5524 | -2.6376 | -0.36 |
| 16496 | *ACSM6* | 1.3901 | -1.1169 | -1.3565 | -1.7061 | -0.36 |
| 16497 | *ABCF1* | -0.10772 | -0.1258 | -0.84992 | -2.1577 | -0.36 |
| 16498 | *CLOCK* | 0.70356 | -0.69898 | -1.0886 | -1.7517 | -0.36 |
| 16499 | *STAG1* | 0.037844 | 0.019274 | -1.1412 | -1.8004 | -0.36 |
| 16500 | *ANO10* | 0.039166 | -0.09689 | -1.0273 | -1.6422 | -0.36 |
| 16501 | *SLC25A43* | 1.7958 | -1.3127 | -1.5684 | -1.9757 | -0.36 |
| 16502 | *SCUBE3* | 0.15149 | -0.51405 | -0.72304 | -0.8728 | -0.36 |
| 16503 | *DHX38* | 0.30207 | -0.19268 | -1.1964 | -1.4019 | -0.36 |
| 16504 | *PELI2* | 1.2094 | -1.0948 | -1.2027 | -1.8453 | -0.36 |
| 16505 | *S100A5* | 0.32406 | -0.68706 | -0.726 | -1.2935 | -0.36 |
| 16506 | *FTSJ1* | 0.85651 | -0.93217 | -1.0142 | -3.6472 | -0.36 |
| 16507 | *PPM1E* | 0.016186 | -0.46745 | -0.63939 | -3.5043 | -0.36 |

| | | | | | | |
|---|---|---|---|---|---|---|
| 16508 | *RPS28* | 0.18173 | -0.55999 | -0.71626 | -1.1484 | -0.36 |
| 16509 | *ZNF70* | 0.71892 | -0.90542 | -0.90824 | -0.92689 | -0.36 |
| 16510 | *HADHA* | 0.94774 | -0.55387 | -1.4894 | -1.8822 | -0.37 |
| 16511 | *PVRL3* | 0.61584 | 0.16661 | -1.8789 | -2.0748 | -0.37 |
| 16512 | *LOC391003* | 0.40829 | -0.70546 | -0.79987 | -1.0669 | -0.37 |
| 16513 | *TCF12* | -0.11965 | -0.34144 | -0.6362 | -1.2644 | -0.37 |
| 16514 | *FAM110B* | 0.56135 | -0.68042 | -0.97966 | -1.9623 | -0.37 |
| 16515 | *CASQ1* | 1.127 | -0.8903 | -1.3367 | -1.9727 | -0.37 |
| 16516 | *DMPK* | 0.48662 | -0.79149 | -0.79558 | -1.3694 | -0.37 |
| 16517 | *NYNRIN* | 0.33047 | -0.29979 | -1.1312 | -1.3253 | -0.37 |
| 16518 | *PDCL3* | 1.015 | -0.93199 | -1.1836 | -1.2299 | -0.37 |
| 16519 | *C11orf40* | 1.7282 | -1.2924 | -1.5373 | -1.9689 | -0.37 |
| 16520 | *FBXO48* | 3.349 | -1.7462 | -2.7063 | -2.7546 | -0.37 |
| 16521 | *KRT9* | 0.43544 | -0.6828 | -0.85622 | -2.6028 | -0.37 |
| 16522 | *DCX* | 0.33864 | -0.48833 | -0.95495 | -1.0102 | -0.37 |
| 16523 | *ARHGAP5* | 0.18649 | -0.45119 | -0.84001 | -2.0871 | -0.37 |
| 16524 | *CHMP2A* | 0.13558 | -0.44353 | -0.79983 | -1.2367 | -0.37 |
| 16525 | *LIMK1* | 0.1583 | -0.26466 | -1.0024 | -1.4284 | -0.37 |
| 16526 | *CRKL* | 0.21678 | -0.0188 | -1.3068 | -1.85 | -0.37 |
| 16527 | *EN2* | -0.04526 | -0.26717 | -0.79711 | -1.9358 | -0.37 |
| 16528 | *NFKB1* | -0.03262 | -0.23306 | -0.84563 | -1.2407 | -0.37 |
| 16529 | *ACTA1* | 0.59964 | 0.044746 | -1.7567 | -1.9125 | -0.37 |
| 16530 | *ZG16* | 0.15852 | -0.51713 | -0.75665 | -0.79632 | -0.37 |
| 16531 | *RGP1* | -0.19089 | -0.30457 | -0.62047 | -1.3916 | -0.37 |
| 16532 | *ZNF579* | 0.12481 | 0.000172 | -1.241 | -1.4994 | -0.37 |
| 16533 | *PLXDC1* | -0.09 | -0.15953 | -0.86821 | -1.3892 | -0.37 |
| 16534 | *TSLP* | 0.25678 | -0.66215 | -0.71257 | -1.8238 | -0.37 |
| 16535 | *PHLDA2* | 1.1919 | -0.64725 | -1.6653 | -2.8181 | -0.37 |
| 16536 | *FAM110A* | 0.044855 | -0.23113 | -0.93592 | -2.1956 | -0.37 |
| 16537 | *MBD3* | 1.2145 | -0.67812 | -1.6587 | -2.4491 | -0.37 |
| 16538 | *ABI2* | 0.48253 | -0.39171 | -1.2141 | -1.3937 | -0.37 |
| 16539 | *PDS5B* | 0.58937 | -0.54097 | -1.1717 | -2.224 | -0.37 |
| 16540 | *FOXL1* | 0.04883 | -0.05031 | -1.1223 | -4.9979 | -0.37 |
| 16541 | *ARPC5L* | 0.30141 | -0.48042 | -0.94493 | -1.5744 | -0.37 |
| 16542 | *CCDC53* | 0.07586 | -0.44353 | -0.75641 | -1.7585 | -0.37 |
| 16543 | *SLA2* | 0.80493 | -0.85149 | -1.079 | -1.2356 | -0.38 |
| 16544 | *DZIP1* | 0.59007 | 0.35402 | -2.0697 | -2.2456 | -0.38 |
| 16545 | *RCC1* | 0.074528 | -0.31976 | -0.88062 | -1.4883 | -0.38 |
| 16546 | *SLC17A3* | 1.6209 | -1.0005 | -1.7483 | -2.2021 | -0.38 |
| 16547 | *PHC1* | 0.17695 | 0.11164 | -1.4168 | -1.8157 | -0.38 |
| 16548 | *ACTC1* | 0.32666 | -0.67889 | -0.7771 | -1.3292 | -0.38 |
| 16549 | *PSG1* | 0.20832 | -0.08719 | -1.2514 | -2.5865 | -0.38 |
| 16550 | *NSD1* | 1.7076 | -1.2936 | -1.545 | -1.6281 | -0.38 |
| 16551 | *RAB35* | 0.59959 | -0.85163 | -0.87898 | -1.0923 | -0.38 |
| 16552 | *FAM63B* | -0.00492 | -0.17744 | -0.95045 | -1.635 | -0.38 |
| 16553 | *CSRNP2* | -0.16698 | -0.44353 | -0.52238 | -1.0567 | -0.38 |

| | | | | | | |
|---|---|---|---|---|---|---|
| 16554 | *DGCR6L* | 0.2083 | 0.043977 | -1.3864 | -2.775 | -0.38 |
| 16555 | *ZDHHC4* | -0.25762 | -0.41336 | -0.46404 | -1.5877 | -0.38 |
| 16556 | *EMILIN2* | 0.075627 | -0.3014 | -0.9105 | -1.7056 | -0.38 |
| 16557 | *IVL* | 0.50612 | -0.75408 | -0.88876 | -1.8856 | -0.38 |
| 16558 | *POLR2F* | 1.1802 | -0.48731 | -1.8297 | -1.9446 | -0.38 |
| 16559 | *HYAL4* | -0.02232 | -0.55387 | -0.56076 | -1.3916 | -0.38 |
| 16560 | *CDKL2* | -0.20562 | -0.43473 | -0.49961 | -0.60845 | -0.38 |
| 16561 | *DHTKD1* | 0.48741 | -0.71077 | -0.91886 | -1.078 | -0.38 |
| 16562 | *ZNF335* | 0.4804 | -0.46248 | -1.1607 | -1.9748 | -0.38 |
| 16563 | *SLC45A4* | -0.18019 | -0.34751 | -0.61544 | -0.68813 | -0.38 |
| 16564 | *MYL12A* | 0.5653 | -0.21964 | -1.4894 | -2.0542 | -0.38 |
| 16565 | *TTC27* | 0.53363 | -0.82098 | -0.85799 | -1.062 | -0.38 |
| 16566 | *OR4S2* | 0.35036 | -0.7068 | -0.79149 | -1.9172 | -0.38 |
| 16567 | *ST6GAL1* | -0.2417 | -0.37856 | -0.52853 | -4.1669 | -0.38 |
| 16568 | *USP47* | 0.26788 | -0.10743 | -1.3101 | -2.4703 | -0.38 |
| 16569 | *RHPN2* | 0.048909 | -0.55464 | -0.64552 | -1.8616 | -0.38 |
| 16570 | *CHRNA4* | 0.39383 | -0.04542 | -1.4998 | -3.5394 | -0.38 |
| 16571 | *SCRN2* | 0.90583 | -1.0141 | -1.0436 | -1.3686 | -0.38 |
| 16572 | *ITK* | 0.56058 | -0.55554 | -1.1585 | -1.4688 | -0.38 |
| 16573 | *NELFCD* | 0.29314 | -0.17208 | -1.278 | -2.2059 | -0.39 |
| 16574 | *DNPEP* | 0.93649 | -0.88199 | -1.2124 | -2.1085 | -0.39 |
| 16575 | *MACF1* | 0.66286 | -0.20621 | -1.615 | -2.1419 | -0.39 |
| 16576 | *USP7* | 0.001688 | -0.40714 | -0.75328 | -1.4859 | -0.39 |
| 16577 | *TICAM2* | -0.26537 | -0.43398 | -0.46032 | -0.69841 | -0.39 |
| 16578 | *WDR25* | 0.089784 | -0.23977 | -1.01 | -1.1575 | -0.39 |
| 16579 | *TYK2* | 0.19908 | -0.33731 | -1.0218 | -1.6112 | -0.39 |
| 16580 | *DCP1A* | -0.10006 | -0.5038 | -0.55851 | -2.4764 | -0.39 |
| 16581 | *TRAM2* | -0.01788 | -0.23404 | -0.91103 | -0.98421 | -0.39 |
| 16582 | *RHOG* | 0.87168 | 0.37459 | -2.4094 | -2.599 | -0.39 |
| 16583 | *EID2B* | 0.31895 | -0.4594 | -1.0232 | -2.2732 | -0.39 |
| 16584 | *CETN1* | 0.87504 | -0.70546 | -1.3343 | -1.4561 | -0.39 |
| 16585 | *WNT2B* | 0.64372 | -0.43587 | -1.3738 | -3.2172 | -0.39 |
| 16586 | *CCDC157* | 0.081831 | 0.000461 | -1.2486 | -1.8963 | -0.39 |
| 16587 | *ZBTB5* | 0.40405 | -0.08446 | -1.4864 | -1.5212 | -0.39 |
| 16588 | *PSMA3* | 0.37252 | -0.08887 | -1.4509 | -1.6216 | -0.39 |
| 16589 | *FAM96B* | 0.43982 | -0.30496 | -1.3033 | -1.6463 | -0.39 |
| 16590 | *SLC19A3* | 0.58119 | -0.39817 | -1.3523 | -1.3878 | -0.39 |
| 16591 | *FEZ2* | -0.00301 | -0.36761 | -0.80048 | -2.1198 | -0.39 |
| 16592 | *DDTL* | 0.21931 | -0.49056 | -0.90001 | -1.6492 | -0.39 |
| 16593 | *NPTX1* | 0.93283 | -0.24715 | -1.8572 | -2.4595 | -0.39 |
| 16594 | *CHORDC1* | -0.12421 | -0.29292 | -0.75472 | -1.3808 | -0.39 |
| 16595 | *XCL2* | 0.8875 | 0.36064 | -2.4205 | -2.4933 | -0.39 |
| 16596 | *FUT5* | 0.27717 | -0.69377 | -0.75625 | -1.584 | -0.39 |
| 16597 | *SLC24A3* | 0.46554 | -0.13563 | -1.5031 | -2.1794 | -0.39 |
| 16598 | *TLR7* | 0.78114 | -0.87673 | -1.0782 | -1.7507 | -0.39 |
| 16599 | *FAM21A* | 1.9712 | -1.3023 | -1.8435 | -2.4205 | -0.39 |

| | | | | | | |
|---|---|---|---|---|---|---|
| 16600 | *CTC1* | 1.1339 | -0.46756 | -1.842 | -2.1295 | -0.39 |
| 16601 | *CDR2* | 0.61677 | -0.19234 | -1.6004 | -1.6246 | -0.39 |
| 16602 | *GPRIN1* | -0.06185 | -0.15313 | -0.9612 | -2.0817 | -0.39 |
| 16603 | *HERPUD2* | 0.6823 | -0.76882 | -1.09 | -2.4917 | -0.39 |
| 16604 | *FDPS* | 0.87211 | -0.1427 | -1.9065 | -4.2477 | -0.39 |
| 16605 | *EGR2* | 0.46227 | -0.29953 | -1.3409 | -1.4234 | -0.39 |
| 16606 | *FUCA2* | 0.78899 | -0.81965 | -1.1484 | -1.9219 | -0.39 |
| 16607 | *NCOA1* | -0.16039 | -0.30811 | -0.71063 | -2.1092 | -0.39 |
| 16608 | *MRPL13* | 1.1241 | -0.34148 | -1.9622 | -2.2127 | -0.39 |
| 16609 | *GPSM2* | 0.47413 | -0.33937 | -1.3148 | -1.4031 | -0.39 |
| 16610 | *KIAA1109* | 0.30301 | -0.26348 | -1.22 | -2.7627 | -0.39 |
| 16611 | *PPFIBP2* | -0.16485 | -0.47338 | -0.54402 | -1.6951 | -0.39 |
| 16612 | *MAP3K7CL* | 0.26123 | 0.028777 | -1.4736 | -2.1126 | -0.39 |
| 16613 | *HSPE1* | 0.82872 | -1.0033 | -1.0095 | -1.0884 | -0.39 |
| 16614 | *SNX25* | 0.041176 | -0.14244 | -1.0838 | -2.1945 | -0.40 |
| 16615 | *PRELID1* | -0.26625 | -0.40469 | -0.5142 | -1.1338 | -0.40 |
| 16616 | *C16orf91* | 0.52737 | -0.70442 | -1.0091 | -1.6105 | -0.40 |
| 16617 | *TMEM200A* | 0.22609 | -0.17635 | -1.236 | -1.3008 | -0.40 |
| 16618 | *KRTAP13-4* | 0.54891 | -0.7342 | -1.0013 | -1.0901 | -0.40 |
| 16619 | *NARFL* | 0.22275 | -0.23931 | -1.1703 | -1.4944 | -0.40 |
| 16620 | *RIPPLY1* | 0.31465 | -0.02484 | -1.4768 | -1.7363 | -0.40 |
| 16621 | *RGAG1* | 0.086717 | -0.33517 | -0.93889 | -2.4422 | -0.40 |
| 16622 | *DPF2* | 0.52353 | -0.47473 | -1.2363 | -2.223 | -0.40 |
| 16623 | *ARMC7* | 0.059003 | -0.52552 | -0.72101 | -0.9703 | -0.40 |
| 16624 | *TBC1D3F* | 0.94304 | -0.64575 | -1.4856 | -1.739 | -0.40 |
| 16625 | *UBA1* | -0.20057 | -0.37106 | -0.61779 | -1.9622 | -0.40 |
| 16626 | *KCNJ3* | -0.03528 | -0.53815 | -0.61663 | -1.5264 | -0.40 |
| 16627 | *DNAJC2* | 0.67965 | -0.82524 | -1.0452 | -1.1283 | -0.40 |
| 16628 | *PPAP2A* | 0.06609 | -0.46384 | -0.79319 | -1.2438 | -0.40 |
| 16629 | *ZNF18* | -0.13085 | -0.22886 | -0.83143 | -0.99838 | -0.40 |
| 16630 | *PPP1R11* | 1.0572 | -0.8471 | -1.4024 | -1.9696 | -0.40 |
| 16631 | *RHAG* | -0.15143 | -0.28838 | -0.75408 | -1.4338 | -0.40 |
| 16632 | *CRB1* | 1.1533 | -0.74013 | -1.6072 | -2.0138 | -0.40 |
| 16633 | *COL6A3* | 0.66381 | -0.85594 | -1.0019 | -1.2193 | -0.40 |
| 16634 | *TNFAIP2* | 0.81713 | -0.90346 | -1.1081 | -1.4258 | -0.40 |
| 16635 | *METTL16* | -0.06892 | -0.35617 | -0.77018 | -2.3391 | -0.40 |
| 16636 | *SSC4D* | 0.15744 | -0.53045 | -0.82255 | -2.7509 | -0.40 |
| 16637 | *RPP30* | 1.142 | -0.62378 | -1.7149 | -2.0237 | -0.40 |
| 16638 | *RNF38* | 0.13442 | 0.13427 | -1.4681 | -1.8813 | -0.40 |
| 16639 | *ZNF43* | 0.14319 | -0.62179 | -0.72172 | -1.1936 | -0.40 |
| 16640 | *CHI3L1* | 0.0951 | -0.52238 | -0.77308 | -0.99448 | -0.40 |
| 16641 | *ZYX* | -0.03506 | -0.52711 | -0.63966 | -1.3007 | -0.40 |
| 16642 | *PDE2A* | 0.032469 | -0.23625 | -0.99838 | -1.049 | -0.40 |
| 16643 | *SORCS2* | 0.040477 | -0.16651 | -1.077 | -1.3961 | -0.40 |
| 16644 | *ZNF3* | -0.03668 | -0.07193 | -1.0952 | -1.2694 | -0.40 |
| 16645 | *ZWINT* | -0.15039 | -0.16207 | -0.89186 | -1.252 | -0.40 |

| | | | | | | |
|---|---|---|---|---|---|---|
| 16646 | *KIF3A* | 0.66949 | -0.63198 | -1.2421 | -1.7306 | -0.40 |
| 16647 | *RNMT* | 1.7876 | -1.285 | -1.7075 | -2.0871 | -0.40 |
| 16648 | *CCT2* | -0.28901 | -0.40136 | -0.51467 | -2.4703 | -0.40 |
| 16649 | *CDH26* | 0.15264 | -0.29227 | -1.067 | -2.1004 | -0.40 |
| 16650 | *PRSS3* | 2.1415 | -1.3865 | -1.9622 | -3.213 | -0.40 |
| 16651 | *COX20* | 0.18725 | -0.38527 | -1.0092 | -1.7001 | -0.40 |
| 16652 | *PPP1R12A* | -0.06132 | -0.51468 | -0.63152 | -1.9473 | -0.40 |
| 16653 | *VMA21* | 0.061507 | -0.24431 | -1.0256 | -1.3736 | -0.40 |
| 16654 | *ZNF281* | 0.34273 | -0.10838 | -1.4428 | -2.0384 | -0.40 |
| 16655 | *LSMEM1* | 0.1337 | -0.10716 | -1.237 | -1.4553 | -0.40 |
| 16656 | *SYNJ2BP-COX16* | 0.097417 | -0.14208 | -1.1671 | -2.016 | -0.40 |
| 16657 | *FBXO45* | 0.3421 | -0.51825 | -1.0359 | -1.1601 | -0.40 |
| 16658 | *RHO* | -0.16122 | -0.49867 | -0.55284 | -0.62881 | -0.40 |
| 16659 | *PTCHD1* | 2.1576 | -1.55 | -1.8207 | -3.8595 | -0.40 |
| 16660 | *CCDC36* | 0.54829 | -0.74572 | -1.0158 | -1.4527 | -0.40 |
| 16661 | *ADGRB1* | -0.09317 | -0.13055 | -0.9904 | -1.1716 | -0.40 |
| 16662 | *MRE11A* | 0.43632 | -0.56815 | -1.0825 | -1.8909 | -0.40 |
| 16663 | *MRGBP* | -0.14632 | -0.34533 | -0.72348 | -1.547 | -0.41 |
| 16664 | *PDIA6* | 0.92608 | -0.98769 | -1.154 | -1.6086 | -0.41 |
| 16665 | *NUPL1* | -0.0261 | -0.34718 | -0.84261 | -1.4392 | -0.41 |
| 16666 | *KXD1* | 0.70797 | -1.51913 | | | -0.41 |
| 16667 | *IQCF5* | 1.1851 | -0.58965 | -1.8129 | -2.5827 | -0.41 |
| 16668 | *RRP15* | 0.23246 | -0.08162 | -1.3689 | -2.381 | -0.41 |
| 16669 | *SERPINB6* | 0.30631 | -0.56241 | -0.96215 | -1.0359 | -0.41 |
| 16670 | *ST6GALNAC1* | 0.42253 | -0.23989 | -1.4036 | -1.8996 | -0.41 |
| 16671 | *TF* | -0.19128 | -0.37213 | -0.65781 | -2.0337 | -0.41 |
| 16672 | *LIN28A* | 0.75838 | -0.70601 | -1.2736 | -1.7266 | -0.41 |
| 16673 | *KIF20A* | 1.6278 | -1.3189 | -1.5302 | -1.6964 | -0.41 |
| 16674 | *TEX35* | -0.30613 | -0.35986 | -0.55563 | -1.2514 | -0.41 |
| 16675 | *LRRC4* | 0.41666 | -0.11759 | -1.5212 | -1.5462 | -0.41 |
| 16676 | *RB1CC1* | 0.21963 | -0.04896 | -1.3932 | -2.4077 | -0.41 |
| 16677 | *FAM157B* | 1.5649 | -0.85236 | -1.9353 | -2.4381 | -0.41 |
| 16678 | *WFDC12* | 0.99686 | -0.95206 | -1.2677 | -2.1577 | -0.41 |
| 16679 | *CAPN5* | 0.004697 | -0.46758 | -0.7602 | -2.0382 | -0.41 |
| 16680 | *GGA3* | 2.1224 | -1.6395 | -1.7065 | -2.8849 | -0.41 |
| 16681 | *CYP11A1* | 0.39397 | 0.20224 | -1.8202 | -2.4322 | -0.41 |
| 16682 | *MECP2* | -0.3908 | -0.39689 | -0.43634 | -0.88839 | -0.41 |
| 16683 | *FKBP4* | 0.30432 | -0.0205 | -1.5091 | -1.5099 | -0.41 |
| 16684 | *ELMOD1* | 0.87673 | -0.95893 | -1.1432 | -1.4125 | -0.41 |
| 16685 | *POLN* | 0.83536 | -0.88438 | -1.1765 | -1.9075 | -0.41 |
| 16686 | *SOX13* | -0.08719 | -0.34077 | -0.79821 | -1.7056 | -0.41 |
| 16687 | *PCDHB6* | 1.7615 | -1.189 | -1.7991 | -1.8202 | -0.41 |
| 16688 | *CT47A1* | 0.50465 | -0.256 | -1.4754 | -3.5623 | -0.41 |
| 16689 | *USP48* | 0.40503 | -0.65169 | -0.98026 | -2.4172 | -0.41 |
| 16690 | *ABCA7* | -0.124 | -0.51221 | -0.59088 | -1.0351 | -0.41 |
| 16691 | *SF3B1* | 0.29413 | -0.63187 | -0.88963 | -1.5943 | -0.41 |

| | | | | | | |
|---|---|---|---|---|---|---|
| 16692 | *C2orf27B* | -0.11314 | -0.46761 | -0.64689 | -1.7161 | -0.41 |
| 16693 | *HGC6.3* | 0.097436 | -0.44353 | -0.88199 | -1.2867 | -0.41 |
| 16694 | *GLOD4* | 0.31749 | -0.64164 | -0.90508 | -1.1676 | -0.41 |
| 16695 | *TAF6L* | 0.19463 | -0.45505 | -0.96938 | -1.4455 | -0.41 |
| 16696 | *PKD2L1* | 1.3893 | -0.92781 | -1.6917 | -2.1151 | -0.41 |
| 16697 | *TUSC3* | -0.20104 | -0.25999 | -0.77097 | -2.0113 | -0.41 |
| 16698 | *ISPD* | 0.33066 | -0.27961 | -1.2832 | -1.8219 | -0.41 |
| 16699 | *CTRC* | 0.79112 | -0.74649 | -1.2772 | -1.3198 | -0.41 |
| 16700 | *KIAA0319L* | 0.52388 | -0.13843 | -1.62 | -1.8926 | -0.41 |
| 16701 | *SLC38A1* | 0.5536 | -0.08851 | -1.7001 | -1.8946 | -0.41 |
| 16702 | *SORCS3* | 0.42709 | -0.4433 | -1.219 | -2.397 | -0.41 |
| 16703 | *ELOVL2* | -0.00825 | -0.29879 | -0.92948 | -2.7509 | -0.41 |
| 16704 | *PNKP* | 0.49925 | -0.60012 | -1.1357 | -1.6297 | -0.41 |
| 16705 | *DDX41* | 0.16689 | -0.51998 | -0.88372 | -1.4418 | -0.41 |
| 16706 | *MYH7* | 0.72713 | 0.64275 | -2.6075 | -2.9632 | -0.41 |
| 16707 | *OTUD7A* | 0.85789 | -0.44353 | -1.6527 | -3.1317 | -0.41 |
| 16708 | *ZIC2* | 0.22212 | -0.588 | -0.87536 | -1.1025 | -0.41 |
| 16709 | *PGBD1* | 0.093906 | -0.5067 | -0.83027 | -0.98719 | -0.41 |
| 16710 | *NMT2* | 0.59214 | -0.596 | -1.2393 | -2.1412 | -0.41 |
| 16711 | *EIF1* | 0.33597 | -0.47314 | -1.106 | -1.7307 | -0.41 |
| 16712 | *DRAM1* | 0.14084 | -0.59678 | -0.78793 | -1.3114 | -0.41 |
| 16713 | *NUDCD3* | 1.1141 | -0.94015 | -1.4182 | -1.9894 | -0.41 |
| 16714 | *PARP10* | 0.3832 | -0.60794 | -1.0231 | -1.1888 | -0.42 |
| 16715 | *CD96* | 0.14884 | 0.087344 | -1.4841 | -2.2888 | -0.42 |
| 16716 | *FAM188B* | -0.03414 | -0.19753 | -1.0164 | -3.0209 | -0.42 |
| 16717 | *GRIK1* | 0.42316 | -0.05201 | -1.62 | -1.8065 | -0.42 |
| 16718 | *C1QA* | 1.2789 | -1.2119 | -1.3179 | -1.3298 | -0.42 |
| 16719 | *TMEM91* | 0.45249 | -0.72994 | -0.97499 | -1.9052 | -0.42 |
| 16720 | *MPZL2* | 0.023338 | -0.32677 | -0.94912 | -2.1463 | -0.42 |
| 16721 | *FOXD4L1* | 1.5296 | -1.1987 | -1.5847 | -1.8223 | -0.42 |
| 16722 | *VPS25* | 0.010555 | -0.09054 | -1.1756 | -2.0069 | -0.42 |
| 16723 | *TMEM216* | -0.00823 | -0.4571 | -0.79243 | -1.0248 | -0.42 |
| 16724 | *SPINT2* | 0.77909 | -0.85768 | -1.181 | -1.2623 | -0.42 |
| 16725 | *SLC30A9* | -0.02484 | -0.14048 | -1.0948 | -1.6047 | -0.42 |
| 16726 | *ATP6V1D* | 0.42961 | -0.06117 | -1.6289 | -1.7366 | -0.42 |
| 16727 | *WDR66* | 0.71094 | -0.35733 | -1.6143 | -1.6428 | -0.42 |
| 16728 | *MID1* | -0.1082 | -0.32087 | -0.83277 | -1.7285 | -0.42 |
| 16729 | *ARSI* | 0.33086 | 0.16491 | -1.7585 | -2.7546 | -0.42 |
| 16730 | *NMT1* | 0.40937 | -0.60016 | -1.0726 | -1.2684 | -0.42 |
| 16731 | *NT5C1A* | 0.60858 | -0.89186 | -0.98026 | -2.5056 | -0.42 |
| 16732 | *PAK6* | 0.10498 | -0.47183 | -0.89741 | -1.4258 | -0.42 |
| 16733 | *GPR82* | 2.1509 | -1.6429 | -1.7726 | -1.8751 | -0.42 |
| 16734 | *CCDC184* | -0.05524 | -0.07692 | -1.1325 | -2.3707 | -0.42 |
| 16735 | *ZNF19* | 0.069497 | -0.44353 | -0.89258 | -1.5401 | -0.42 |
| 16736 | *PHF20L1* | 0.25931 | -0.28173 | -1.2443 | -1.5613 | -0.42 |
| 16737 | *IFITM5* | 0.4629 | 0.12787 | -1.8583 | -2.0351 | -0.42 |

| | | | | | | |
|---|---|---|---|---|---|---|
| 16738 | *MVP* | 1.9956 | -1.0945 | -2.1687 | -2.3293 | -0.42 |
| 16739 | *ABHD2* | -0.04678 | -0.50718 | -0.71421 | -1.1595 | -0.42 |
| 16740 | *RAD51D* | 0.89321 | -0.83913 | -1.3226 | -1.8816 | -0.42 |
| 16741 | *MYH8* | 0.55379 | -0.78588 | -1.0367 | -1.1214 | -0.42 |
| 16742 | *DTX2* | 0.060104 | -0.0299 | -1.2995 | -1.7746 | -0.42 |
| 16743 | *C19orf53* | 0.42233 | -0.36557 | -1.3282 | -1.4312 | -0.42 |
| 16744 | *USP22* | 0.27373 | -0.15835 | -1.3875 | -3.0097 | -0.42 |
| 16745 | *TRAM1L1* | 0.41353 | -0.81433 | -0.87198 | -1.3623 | -0.42 |
| 16746 | *HMGN4* | -0.23523 | -0.31694 | -0.72095 | -1.1084 | -0.42 |
| 16747 | *B4GALT2* | 0.028002 | -0.03222 | -1.2705 | -1.3162 | -0.42 |
| 16748 | *HDAC11* | 0.33577 | -0.198 | -1.4132 | -1.42 | -0.43 |
| 16749 | *SKIDA1* | -0.00823 | -0.41716 | -0.85018 | -1.6525 | -0.43 |
| 16750 | *EXOC3L1* | 0.66957 | -0.1496 | -1.7956 | -1.8561 | -0.43 |
| 16751 | *FAM92A1* | 1.1611 | -1.1283 | -1.3085 | -1.4725 | -0.43 |
| 16752 | *OXSR1* | 0.68447 | -0.67007 | -1.2903 | -1.3198 | -0.43 |
| 16753 | *KIAA0556* | -0.0953 | -0.55649 | -0.62739 | -0.97425 | -0.43 |
| 16754 | *FTH1* | 1.1697 | -0.77333 | -1.6759 | -2.224 | -0.43 |
| 16755 | *DEF8* | 0.15306 | -0.55147 | -0.88164 | -1.1822 | -0.43 |
| 16756 | *COMMD2* | 0.58207 | -0.52949 | -1.3327 | -1.6527 | -0.43 |
| 16757 | *TCF4* | 0.44453 | -0.44353 | -1.2816 | -1.6528 | -0.43 |
| 16758 | *DDX39A* | 0.38094 | -0.76358 | -0.8988 | -1.7577 | -0.43 |
| 16759 | *RAB15* | 0.33184 | -0.79904 | -0.81509 | -1.2684 | -0.43 |
| 16760 | *ZFP64* | -0.09638 | -0.29596 | -0.89009 | -1.1441 | -0.43 |
| 16761 | *FAM126B* | -0.07889 | -0.17249 | -1.0315 | -1.4316 | -0.43 |
| 16762 | *AP4S1* | 0.81248 | -0.8071 | -1.2895 | -2.462 | -0.43 |
| 16763 | *C1orf116* | 0.015765 | -0.16483 | -1.1354 | -1.2837 | -0.43 |
| 16764 | *ARMC2* | -0.04965 | -0.54621 | -0.69003 | -0.98572 | -0.43 |
| 16765 | *YME1L1* | 1.0655 | -0.54604 | -1.8056 | -1.8846 | -0.43 |
| 16766 | *C1orf43* | 0.27172 | -0.54934 | -1.009 | -1.535 | -0.43 |
| 16767 | *HIST1H2BI* | 1.0628 | -1.1384 | -1.2113 | -2.2318 | -0.43 |
| 16768 | *IFT74* | 1.1959 | -0.57586 | -1.9086 | -2.8928 | -0.43 |
| 16769 | *CYB561D2* | 0.18732 | -0.10119 | -1.3759 | -1.4361 | -0.43 |
| 16770 | *TMEM239* | 0.85612 | -1.0067 | -1.1415 | -1.7602 | -0.43 |
| 16771 | *OR7D4* | 0.89819 | -0.22131 | -1.9695 | -2.7331 | -0.43 |
| 16772 | *ZNF750* | 0.37523 | -0.80368 | -0.86517 | -1.5948 | -0.43 |
| 16773 | *C11orf54* | 0.082077 | -0.64571 | -0.73051 | -0.98853 | -0.43 |
| 16774 | *DHRS7C* | 0.75712 | -0.38522 | -1.6663 | -2.3292 | -0.43 |
| 16775 | *FAM25C* | 1.5838 | -1.1384 | -1.7407 | -1.808 | -0.43 |
| 16776 | *FAM83H* | 0.29438 | -0.44353 | -1.1472 | -2.5844 | -0.43 |
| 16777 | *APOL3* | -0.10178 | -0.40849 | -0.78839 | -0.93114 | -0.43 |
| 16778 | *FSTL5* | 0.77831 | -0.19107 | -1.8869 | -2.101 | -0.43 |
| 16779 | *C21orf58* | 0.032576 | -0.59716 | -0.73511 | -2.3287 | -0.43 |
| 16780 | *GPC3* | 0.12545 | -0.14477 | -1.2815 | -2.252 | -0.43 |
| 16781 | *MROH9* | 0.93654 | -0.8756 | -1.3619 | -1.4452 | -0.43 |
| 16782 | *TP53* | -0.10753 | -0.36427 | -0.83005 | -0.91981 | -0.43 |
| 16783 | *SOGA3* | -0.04919 | -0.3205 | -0.93287 | -1.7024 | -0.43 |

| | | | | | | |
|---|---|---|---|---|---|---|
| 16784 | *TJP3* | 0.92433 | -0.93592 | -1.2912 | -1.6636 | -0.43 |
| 16785 | *PER1* | 0.41974 | -0.15385 | -1.5687 | -1.8046 | -0.43 |
| 16786 | *NDUFA5* | 0.1836 | -0.06124 | -1.4252 | -1.4649 | -0.43 |
| 16787 | *CDC37* | 1.2549 | -1.1193 | -1.4392 | -5.0141 | -0.43 |
| 16788 | *MAGI1* | 0.38482 | -0.70078 | -0.98798 | -1.0838 | -0.43 |
| 16789 | *UBE2H* | 0.076188 | -0.62833 | -0.75205 | -1.0711 | -0.43 |
| 16790 | *ZNF669* | 0.13063 | -0.69915 | -0.73577 | -1.0168 | -0.43 |
| 16791 | *CRYGA* | -0.0248 | -0.33984 | -0.94255 | -1.3033 | -0.44 |
| 16792 | *CASC4* | 0.85768 | -0.57547 | -1.5895 | -1.9772 | -0.44 |
| 16793 | *DNAJC18* | -0.2304 | -0.36996 | -0.70756 | -1.5118 | -0.44 |
| 16794 | *EIF5* | -0.35784 | -0.38862 | -0.56225 | -1.4258 | -0.44 |
| 16795 | *TXNDC8* | 0.88276 | -0.67599 | -1.5171 | -1.677 | -0.44 |
| 16796 | *NXF2B* | 0.17274 | -0.54679 | -0.93707 | -1.8441 | -0.44 |
| 16797 | *SSBP4* | 0.46245 | -0.74011 | -1.0344 | -1.4448 | -0.44 |
| 16798 | *TSPY3* | 0.52567 | -0.40633 | -1.4317 | -1.8921 | -0.44 |
| 16799 | *CD81* | 0.3919 | 0.086138 | -1.7906 | -2.5707 | -0.44 |
| 16800 | *SOX15* | 0.61696 | -0.78229 | -1.1479 | -2.1572 | -0.44 |
| 16801 | *XAGE1E* | 0.19908 | -0.55846 | -0.95388 | -1.2048 | -0.44 |
| 16802 | *EXOSC10* | -0.03791 | -0.33297 | -0.9442 | -1.3766 | -0.44 |
| 16803 | *SORBS2* | 0.33331 | -0.2539 | -1.3945 | -1.6268 | -0.44 |
| 16804 | *ZNF106* | -0.27612 | -0.46559 | -0.57546 | -1.248 | -0.44 |
| 16805 | *FAM19A4* | 0.36026 | -0.68598 | -0.99357 | -1.3664 | -0.44 |
| 16806 | *CT47A12* | 1.067 | -0.66647 | -1.7203 | -2.2691 | -0.44 |
| 16807 | *ACIN1* | 0.75545 | -0.29681 | -1.7804 | -1.8014 | -0.44 |
| 16808 | *DVL1* | 0.47391 | -0.42859 | -1.3676 | -2.2403 | -0.44 |
| 16809 | *ADTRP* | 1.3258 | -0.62862 | -2.0196 | -2.458 | -0.44 |
| 16810 | *GALNT2* | 0.42438 | -0.64571 | -1.1021 | -2.7325 | -0.44 |
| 16811 | *PSMC3* | 0.12979 | -0.41239 | -1.0413 | -1.1357 | -0.44 |
| 16812 | *GP1BA* | 0.13779 | -0.4055 | -1.0564 | -1.0576 | -0.44 |
| 16813 | *CDKN1B* | 1.6431 | -1.3023 | -1.6676 | -1.828 | -0.44 |
| 16814 | *GALNT14* | 0.21664 | -0.02974 | -1.5144 | -1.8462 | -0.44 |
| 16815 | *CYP2E1* | -0.24501 | -0.37274 | -0.70988 | -1.0293 | -0.44 |
| 16816 | *HMGA1* | 0.83604 | -0.855 | -1.3088 | -2.1561 | -0.44 |
| 16817 | *NEIL1* | 0.50969 | -0.6689 | -1.1693 | -1.9953 | -0.44 |
| 16818 | *EBAG9* | 0.36538 | -0.09924 | -1.5962 | -5.1808 | -0.44 |
| 16819 | *GRSF1* | 1.2977 | -1.1388 | -1.4893 | -2.2117 | -0.44 |
| 16820 | *HABP2* | -0.16374 | -0.40693 | -0.7602 | -1.4824 | -0.44 |
| 16821 | *KCNJ14* | -0.39352 | -0.44572 | -0.49177 | -0.83627 | -0.44 |
| 16822 | *ZBTB1* | -0.07367 | -0.14994 | -1.1089 | -1.8642 | -0.44 |
| 16823 | *IST1* | 2.2095 | -1.7056 | -1.8366 | -2.3755 | -0.44 |
| 16824 | *KLHL26* | 1.0576 | -0.53078 | -1.8611 | -2.0407 | -0.44 |
| 16825 | *MDM2* | 1.1425 | -0.90028 | -1.5782 | -1.8577 | -0.45 |
| 16826 | *KRTAP4-5* | 0.20138 | -0.2748 | -1.2633 | -2.3689 | -0.45 |
| 16827 | *SLC14A1* | 0.34183 | -0.12958 | -1.5494 | -2.3585 | -0.45 |
| 16828 | *KAT2B* | 0.77868 | -0.80498 | -1.311 | -2.6999 | -0.45 |
| 16829 | *FHOD3* | 0.13369 | -0.62073 | -0.85027 | -1.2766 | -0.45 |

| | | | | | | |
|---|---|---|---|---|---|---|
| 16830 | *NCOA4* | 0.4616 | -0.7532 | -1.0461 | -1.0602 | -0.45 |
| 16831 | *CNNM3* | 0.18739 | -0.63152 | -0.89401 | -1.4998 | -0.45 |
| 16832 | *SUCLG2* | 0.1473 | -0.67141 | -0.81433 | -1.2691 | -0.45 |
| 16833 | *THPO* | 0.48535 | -0.34886 | -1.4754 | -2.2172 | -0.45 |
| 16834 | *KCNJ10* | -0.21587 | -0.23276 | -0.8934 | -1.0571 | -0.45 |
| 16835 | *SNRPD2* | 0.29824 | -0.16986 | -1.4705 | -1.5906 | -0.45 |
| 16836 | *XAGE5* | 0.044275 | -0.34886 | -1.0385 | -1.1943 | -0.45 |
| 16837 | *ATP6V1C1* | 0.9606 | -0.87837 | -1.4258 | -2.2276 | -0.45 |
| 16838 | *MMS22L* | 0.59723 | -0.79474 | -1.1463 | -2.0492 | -0.45 |
| 16839 | *ETV5* | -0.19911 | -0.44975 | -0.69629 | -1.0664 | -0.45 |
| 16840 | *CHST14* | 1.0423 | -1.072 | -1.3162 | -2.9578 | -0.45 |
| 16841 | *VPS13A* | 0.51796 | -0.88702 | -0.97702 | -1.6072 | -0.45 |
| 16842 | *AHSA2* | 1.0543 | -0.92956 | -1.4725 | -2.9283 | -0.45 |
| 16843 | *MDH2* | 0.096631 | -0.65113 | -0.79423 | -1.0697 | -0.45 |
| 16844 | *EPRS* | 0.39464 | -0.71009 | -1.0336 | -2.0304 | -0.45 |
| 16845 | *TTBK2* | 0.38316 | -0.65968 | -1.0726 | -1.3818 | -0.45 |
| 16846 | *ITGA6* | 0.51223 | -0.87817 | -0.9858 | -1.8297 | -0.45 |
| 16847 | *RMND5B* | 0.40253 | -0.37104 | -1.3836 | -1.682 | -0.45 |
| 16848 | *PBOV1* | -0.14868 | -0.30065 | -0.90281 | -1.537 | -0.45 |
| 16849 | *SNX4* | -0.22963 | -0.44088 | -0.68229 | -1.2602 | -0.45 |
| 16850 | *MAP3K6* | 1.2223 | -1.0613 | -1.5138 | -2.428 | -0.45 |
| 16851 | *EIF1AD* | 0.70035 | -0.85389 | -1.1995 | -2.0895 | -0.45 |
| 16852 | *ARRB1* | 1.0809 | -1.0819 | -1.3526 | -2.1605 | -0.45 |
| 16853 | *NUP205* | 0.7313 | -0.91767 | -1.1676 | -1.2884 | -0.45 |
| 16854 | *DKK3* | 0.28522 | -0.18101 | -1.4584 | -2.072 | -0.45 |
| 16855 | *TACR1* | 0.90394 | -1.012 | -1.2464 | -1.3348 | -0.45 |
| 16856 | *NUP85* | 0.41703 | -0.80558 | -0.96632 | -1.8364 | -0.45 |
| 16857 | *DLST* | 0.57693 | -0.58231 | -1.3498 | -1.8885 | -0.45 |
| 16858 | *TRAPPC2* | 1.9161 | -1.2525 | -2.0196 | -2.1213 | -0.45 |
| 16859 | *TMED5* | 0.75389 | -0.90947 | -1.2008 | -1.9984 | -0.45 |
| 16860 | *DHFR* | 0.2527 | -0.64206 | -0.96946 | -1.0424 | -0.45 |
| 16861 | *KLC3* | 0.32443 | -0.67441 | -1.0115 | -3.1939 | -0.45 |
| 16862 | *FAM169A* | 0.082752 | -0.44745 | -0.99746 | -1.1385 | -0.45 |
| 16863 | *C8orf82* | -0.28581 | -0.38103 | -0.69622 | -1.4841 | -0.45 |
| 16864 | *NKAIN3* | 0.57482 | -0.50046 | -1.4376 | -2.1019 | -0.45 |
| 16865 | *ZNF624* | 0.60661 | -0.57967 | -1.3905 | -2.2879 | -0.45 |
| 16866 | *PTEN* | -0.29563 | -0.40428 | -0.66432 | -0.98266 | -0.45 |
| 16867 | *RMI1* | -0.34118 | -0.43841 | -0.58509 | -1.7716 | -0.45 |
| 16868 | *HELZ2* | 0.24059 | -0.74471 | -0.86096 | -2.6177 | -0.46 |
| 16869 | *C7orf31* | 0.077878 | -0.0576 | -1.3864 | -1.5204 | -0.46 |
| 16870 | *UNC5B* | 0.20681 | -0.6945 | -0.87898 | -2.4013 | -0.46 |
| 16871 | *ODF3B* | 1.3947 | -1.0212 | -1.742 | -2.4718 | -0.46 |
| 16872 | *KCNT1* | -0.12866 | -0.39451 | -0.84539 | -1.3755 | -0.46 |
| 16873 | *CCDC86* | 0.40836 | -0.50057 | -1.2772 | -1.5962 | -0.46 |
| 16874 | *GCNT2* | 0.66152 | -0.93661 | -1.0944 | -1.0992 | -0.46 |
| 16875 | *HNRNPA2B1* | -0.02808 | -0.3016 | -1.0402 | -1.3078 | -0.46 |

| | | | | | | |
|---|---|---|---|---|---|---|
| 16876 | *DNASE1* | 0.6271 | -0.7008 | -1.2963 | -1.9695 | -0.46 |
| 16877 | *NXPE4* | 0.53569 | -0.80442 | -1.1016 | -1.1388 | -0.46 |
| 16878 | *PCDHGA4* | -0.27159 | -0.48235 | -0.61643 | -1.8611 | -0.46 |
| 16879 | *ARL2BP* | 0.6696 | -0.36467 | -1.6759 | -2.0116 | -0.46 |
| 16880 | *PYY* | 0.36562 | -0.45699 | -1.2827 | -3.2454 | -0.46 |
| 16881 | *KIAA1549L* | 0.53271 | -0.76083 | -1.1461 | -1.6007 | -0.46 |
| 16882 | *ORC6* | -0.11998 | -0.59178 | -0.66251 | -1.7679 | -0.46 |
| 16883 | *CFHR5* | 0.45842 | -0.3159 | -1.5173 | -3.0157 | -0.46 |
| 16884 | *C15orf39* | 0.40757 | -0.41323 | -1.37 | -1.7571 | -0.46 |
| 16885 | *RAD50* | 0.4089 | -0.09344 | -1.6913 | -1.9567 | -0.46 |
| 16886 | *NUMB* | 0.15465 | -0.76358 | -0.76716 | -1.2259 | -0.46 |
| 16887 | *OFD1* | 0.54035 | -0.50048 | -1.416 | -2.3885 | -0.46 |
| 16888 | *RHOT2* | 0.45584 | -0.28825 | -1.5467 | -2.1264 | -0.46 |
| 16889 | *PSORS1C2* | 0.87325 | 0.5873 | -2.8398 | -3.4804 | -0.46 |
| 16890 | *FAM134C* | 0.31649 | -0.23638 | -1.4596 | -1.927 | -0.46 |
| 16891 | *CCDC146* | -0.14306 | -0.2157 | -1.0211 | -1.5558 | -0.46 |
| 16892 | *C10orf67* | 0.77507 | -0.52105 | -1.6343 | -2.6113 | -0.46 |
| 16893 | *ATAD2* | 0.49719 | -0.41469 | -1.4633 | -2.3357 | -0.46 |
| 16894 | *PARP3* | 0.80454 | -0.53304 | -1.6525 | -2.8702 | -0.46 |
| 16895 | *EDC4* | 0.21497 | -0.48875 | -1.1073 | -1.2483 | -0.46 |
| 16896 | *QSOX2* | 0.53716 | -0.66191 | -1.2565 | -1.2823 | -0.46 |
| 16897 | *HEATR9* | 0.86989 | -1.0093 | -1.2426 | -1.2666 | -0.46 |
| 16898 | *SH3TC2* | 0.3946 | -0.64888 | -1.1289 | -1.1968 | -0.46 |
| 16899 | *CXCL11* | 0.080928 | 0.037279 | -1.5018 | -2.5156 | -0.46 |
| 16900 | *GDPD1* | 0.032511 | -0.64278 | -0.7747 | -0.85059 | -0.46 |
| 16901 | *ENKD1* | 0.10504 | -0.32219 | -1.1681 | -1.682 | -0.46 |
| 16902 | *PLK4* | -0.05613 | -0.54972 | -0.77946 | -2.3101 | -0.46 |
| 16903 | *DERL1* | 0.13368 | -0.56714 | -0.95321 | -1.1354 | -0.46 |
| 16904 | *MCM7* | 0.93277 | -0.99346 | -1.3267 | -2.6416 | -0.46 |
| 16905 | *TRIM43B* | 0.85937 | -1.0142 | -1.2343 | -1.3162 | -0.46 |
| 16906 | *METTL3* | -0.25305 | -0.43626 | -0.70074 | -1.2845 | -0.46 |
| 16907 | *NXNL1* | 0.35324 | -0.61665 | -1.127 | -4.0338 | -0.46 |
| 16908 | *FAM132B* | -0.19512 | -0.35669 | -0.83862 | -1.6852 | -0.46 |
| 16909 | *BRD4* | 0.21781 | -0.44574 | -1.1631 | -1.9686 | -0.46 |
| 16910 | *ZNF442* | 0.18078 | -0.49311 | -1.0793 | -1.9811 | -0.46 |
| 16911 | *CIDEC* | -0.16962 | -0.39689 | -0.82547 | -2.6169 | -0.46 |
| 16912 | *NPY1R* | 1.0835 | -1.141 | -1.3349 | -1.6251 | -0.46 |
| 16913 | *SH2B2* | 0.064869 | -0.2005 | -1.257 | -1.8219 | -0.46 |
| 16914 | *ANKRA2* | 0.34565 | -0.37544 | -1.3633 | -1.3892 | -0.46 |
| 16915 | *DRD1* | 0.25102 | -0.32237 | -1.3218 | -1.5212 | -0.46 |
| 16916 | *MFSD8* | -0.02747 | -0.46124 | -0.90462 | -1.3291 | -0.46 |
| 16917 | *LRRC3C* | 0.32643 | -0.7967 | -0.92366 | -1.1156 | -0.46 |
| 16918 | *POLR2L* | 0.26231 | -0.64675 | -1.0095 | -1.348 | -0.46 |
| 16919 | *ZNF589* | 1.1646 | -0.83883 | -1.72 | -2.9772 | -0.46 |
| 16920 | *GOLGA8H* | 0.40438 | -0.45669 | -1.3425 | -1.8154 | -0.46 |
| 16921 | *THOC7* | 0.71186 | -0.7679 | -1.3399 | -2.0407 | -0.47 |

| | | | | | | |
|---|---|---|---|---|---|---|
| 16922 | *C15orf54* | 0.82828 | -0.90616 | -1.3183 | -1.7232 | -0.47 |
| 16923 | *OR2T29* | 0.92036 | -0.58084 | -1.7364 | -3.8028 | -0.47 |
| 16924 | *GEN1* | 0.27479 | -0.44353 | -1.2284 | -1.8112 | -0.47 |
| 16925 | *FAM219A* | 0.83015 | -0.73853 | -1.4894 | -1.615 | -0.47 |
| 16926 | *MOCS1* | 0.80962 | -0.78172 | -1.4258 | -1.9054 | -0.47 |
| 16927 | *DSC1* | -0.16559 | -0.56411 | -0.66912 | -2.0679 | -0.47 |
| 16928 | *CYP26A1* | 0.85099 | -0.83862 | -1.4112 | -2.1936 | -0.47 |
| 16929 | *LEFTY2* | 0.062474 | -0.32293 | -1.1391 | -1.2817 | -0.47 |
| 16930 | *EPHA10* | 0.59415 | -0.81787 | -1.1768 | -1.5256 | -0.47 |
| 16931 | *EPB41L1* | -0.31688 | -0.47889 | -0.6057 | -2.412 | -0.47 |
| 16932 | *ARL4A* | 0.23768 | -0.50075 | -1.1391 | -2.4642 | -0.47 |
| 16933 | *TNFRSF9* | 0.77843 | -0.13473 | -2.0465 | -2.2746 | -0.47 |
| 16934 | *UTP3* | 1.1885 | -1.2167 | -1.3753 | -2.7384 | -0.47 |
| 16935 | *GNRH1* | 1.8259 | -1.5424 | -1.6891 | -2.1282 | -0.47 |
| 16936 | *C6orf165* | 0.62329 | -0.30878 | -1.7203 | -3.2089 | -0.47 |
| 16937 | *IQUB* | 0.01805 | -0.09539 | -1.3285 | -1.7526 | -0.47 |
| 16938 | *USP17L13* | -0.19885 | -0.28032 | -0.9285 | -0.93465 | -0.47 |
| 16939 | *CSTA* | 0.26443 | -0.42369 | -1.2485 | -2.2254 | -0.47 |
| 16940 | *LGMN* | 1.9955 | -1.5815 | -1.8219 | -2.5806 | -0.47 |
| 16941 | *INTS3* | 0.72957 | -0.207 | -1.9305 | -1.9357 | -0.47 |
| 16942 | *TRIM34* | 0.48846 | -0.91944 | -0.97952 | -1.9165 | -0.47 |
| 16943 | *DLEC1* | 0.93606 | -1.0212 | -1.3267 | -1.6707 | -0.47 |
| 16944 | *RNF146* | 0.32681 | -0.7456 | -0.99446 | -1.6016 | -0.47 |
| 16945 | *RPP21* | 0.81532 | -0.78044 | -1.4482 | -1.6824 | -0.47 |
| 16946 | *SMR3B* | 0.67996 | -0.98597 | -1.1091 | -1.1968 | -0.47 |
| 16947 | *CNGA4* | 0.17463 | -0.50793 | -1.083 | -2.5707 | -0.47 |
| 16948 | *SRC* | 1.0414 | 0.1507 | -2.6101 | -2.6376 | -0.47 |
| 16949 | *ELMO3* | 0.44547 | -0.62306 | -1.241 | -1.9619 | -0.47 |
| 16950 | *FAM83E* | 0.083155 | -0.52035 | -0.98214 | -1.2016 | -0.47 |
| 16951 | *LSMEM2* | 0.35889 | -0.5707 | -1.2084 | -1.3965 | -0.47 |
| 16952 | *HAS2* | -0.16888 | -0.37998 | -0.8715 | -0.93789 | -0.47 |
| 16953 | *CDAN1* | 0.17097 | -0.45759 | -1.1344 | -1.2987 | -0.47 |
| 16954 | *RWDD4* | 1.384 | -1.2571 | -1.5491 | -2.8398 | -0.47 |
| 16955 | *APOD* | -0.1401 | -0.60604 | -0.6761 | -1.9794 | -0.47 |
| 16956 | *SAMD8* | 1.5293 | -0.96463 | -1.9874 | -2.3904 | -0.47 |
| 16957 | *ISCA2* | 0.36609 | -0.43005 | -1.3606 | -1.392 | -0.47 |
| 16958 | *KIAA0391* | -0.32397 | -0.43737 | -0.66787 | -1.7865 | -0.48 |
| 16959 | *SYMPK* | 0.5886 | -0.01346 | -2.005 | -2.3496 | -0.48 |
| 16960 | *ZNHIT2* | 1.0595 | -0.85093 | -1.6386 | -1.6626 | -0.48 |
| 16961 | *UBE2S* | 0.31895 | -0.16971 | -1.5803 | -3.051 | -0.48 |
| 16962 | *DYNAP* | 0.98642 | -1.0202 | -1.398 | -1.4794 | -0.48 |
| 16963 | *STRC* | 0.63915 | -0.98421 | -1.0871 | -1.4962 | -0.48 |
| 16964 | *EMC4* | 0.46629 | -0.81509 | -1.0838 | -1.6607 | -0.48 |
| 16965 | *C16orf92* | 1.2503 | -1.1901 | -1.4932 | -1.7016 | -0.48 |
| 16966 | *GFRA2* | 0.54719 | -0.60794 | -1.3723 | -1.5971 | -0.48 |
| 16967 | *PSMA5* | -0.00222 | -0.56919 | -0.86245 | -1.8568 | -0.48 |

| | | | | | | |
|---|---|---|---|---|---|---|
| 16968 | *SLC9A8* | 0.1479 | -0.49177 | -1.0903 | -1.129 | -0.48 |
| 16969 | *CALCOCO2* | 0.48225 | -0.88503 | -1.0315 | -1.5765 | -0.48 |
| 16970 | *EFCAB13* | 0.72377 | -0.44353 | -1.7153 | -1.8219 | -0.48 |
| 16971 | *CBLL1* | -0.37907 | -0.41155 | -0.64725 | -0.8128 | -0.48 |
| 16972 | *ZNF365* | 0.14103 | -0.60841 | -0.97053 | -1.3047 | -0.48 |
| 16973 | *WBP11* | 0.48494 | -0.44933 | -1.4736 | -1.6469 | -0.48 |
| 16974 | *SAPCD2* | 0.046831 | -0.12184 | -1.3637 | -2.2866 | -0.48 |
| 16975 | *MYBBP1A* | -0.29509 | -0.44299 | -0.70101 | -1.2825 | -0.48 |
| 16976 | *PRAMEF7* | -0.11385 | -0.15777 | -1.1676 | -1.7591 | -0.48 |
| 16977 | *MAP9* | -0.14252 | -0.16343 | -1.1341 | -2.0351 | -0.48 |
| 16978 | *ETNK1* | 0.20971 | -0.54274 | -1.1071 | -1.2845 | -0.48 |
| 16979 | *DHX35* | 1.3769 | -1.236 | -1.5814 | -1.5847 | -0.48 |
| 16980 | *IGIP* | -0.3787 | -0.49635 | -0.56929 | -1.0283 | -0.48 |
| 16981 | *HIST2H3A* | 0.18054 | -0.71077 | -0.9142 | -2.0553 | -0.48 |
| 16982 | *RTCB* | 0.31852 | -0.73143 | -1.0317 | -1.7749 | -0.48 |
| 16983 | *CMTM3* | -0.03826 | -0.7004 | -0.70601 | -1.1384 | -0.48 |
| 16984 | *ADO* | 0.087344 | -0.54847 | -0.98421 | -1.7577 | -0.48 |
| 16985 | *GDI2* | 0.46439 | -0.7132 | -1.1967 | -1.9219 | -0.48 |
| 16986 | *FIBIN* | 0.73478 | -0.19107 | -1.9894 | -2.4322 | -0.48 |
| 16987 | *GMEB1* | 0.21099 | -0.74014 | -0.91708 | -1.7667 | -0.48 |
| 16988 | *AWAT1* | 0.25085 | -0.71009 | -0.98769 | -1.5137 | -0.48 |
| 16989 | *CAB39* | 0.31734 | -0.726 | -1.0385 | -1.3409 | -0.48 |
| 16990 | *MYBPHL* | -0.06987 | -0.54177 | -0.83627 | -1.7056 | -0.48 |
| 16991 | *ADAL* | 0.28778 | -0.64313 | -1.0928 | -1.5762 | -0.48 |
| 16992 | *METTL4* | 0.066705 | 0.040796 | -1.5558 | -1.7956 | -0.48 |
| 16993 | *C11orf80* | 0.25796 | -0.57479 | -1.1325 | -2.1648 | -0.48 |
| 16994 | *AGO4* | 0.11117 | -0.53877 | -1.0221 | -1.1808 | -0.48 |
| 16995 | *HGD* | -0.18153 | -0.50992 | -0.75944 | -1.1316 | -0.48 |
| 16996 | *PCBP3* | -0.0044 | -0.44353 | -1.0049 | -1.3851 | -0.48 |
| 16997 | *TBC1D10C* | 0.19114 | -0.18388 | -1.4614 | -1.4713 | -0.48 |
| 16998 | *ELFN2* | 0.37772 | -0.44541 | -1.3872 | -2.1207 | -0.48 |
| 16999 | *HECTD4* | -0.30975 | -0.4376 | -0.70947 | -1.6062 | -0.49 |
| 17000 | *SDS* | -0.10195 | -0.54112 | -0.81478 | -1.6636 | -0.49 |
| 17001 | *C17orf50* | 1.2373 | -1.0944 | -1.6023 | -2.312 | -0.49 |
| 17002 | *SLC5A1* | 0.45988 | -0.25832 | -1.6636 | -3.2454 | -0.49 |
| 17003 | *WDR77* | 0.010272 | -0.52122 | -0.95162 | -2.2771 | -0.49 |
| 17004 | *JMJD7* | 1.4255 | -0.79821 | -2.0902 | -3.6729 | -0.49 |
| 17005 | *CSTF3* | -0.08523 | -0.36194 | -1.0165 | -1.3515 | -0.49 |
| 17006 | *PSMD5* | 0.44138 | -0.20338 | -1.7018 | -2.4873 | -0.49 |
| 17007 | *GOLGA8R* | 0.73068 | -1.0243 | -1.1706 | -2.0851 | -0.49 |
| 17008 | *RFWD2* | 0.46795 | -0.73069 | -1.2027 | -1.6934 | -0.49 |
| 17009 | *INO80E* | -0.30811 | -0.57379 | -0.58403 | -1.8484 | -0.49 |
| 17010 | *KLHL5* | 0.64585 | -0.90281 | -1.2106 | -2.8628 | -0.49 |
| 17011 | *SUOX* | 0.22548 | -0.5871 | -1.106 | -1.1637 | -0.49 |
| 17012 | *SNTA1* | 0.38367 | -0.81237 | -1.0391 | -1.2355 | -0.49 |
| 17013 | *RFPL4B* | 0.12307 | -0.04635 | -1.545 | -1.6158 | -0.49 |

| | | | | | | |
|---|---|---|---|---|---|---|
| 17014 | *TSNAX* | 0.48157 | -0.63936 | -1.3111 | -2.1068 | -0.49 |
| 17015 | *CDK6* | -0.06844 | -0.10911 | -1.2917 | -2.7931 | -0.49 |
| 17016 | *OR13C5* | 0.16164 | -0.75148 | -0.88094 | -2.0398 | -0.49 |
| 17017 | *EOGT* | 0.97673 | -1.1076 | -1.3407 | -3.0314 | -0.49 |
| 17018 | *CMIP* | 0.64718 | -0.64075 | -1.479 | -1.6371 | -0.49 |
| 17019 | *ERAP2* | 0.36116 | -0.77733 | -1.0576 | -1.7018 | -0.49 |
| 17020 | *IBA57* | 0.46245 | -0.00112 | -1.9357 | -3.6076 | -0.49 |
| 17021 | *B4GALT6* | 0.78265 | -1.0987 | -1.1591 | -1.3743 | -0.49 |
| 17022 | *PSMA2* | 0.5153 | -0.84624 | -1.1448 | -1.8804 | -0.49 |
| 17023 | *ZNF575* | -0.01545 | -0.58349 | -0.87694 | -1.7203 | -0.49 |
| 17024 | *PAQR6* | 0.42579 | -0.84337 | -1.0606 | -1.3478 | -0.49 |
| 17025 | *GK* | 0.45578 | -0.51585 | -1.4203 | -1.5165 | -0.49 |
| 17026 | *PPP4R2* | -0.00127 | -0.50953 | -0.9703 | -3.2348 | -0.49 |
| 17027 | *GGPS1* | 0.021592 | -0.05612 | -1.4473 | -2.1182 | -0.49 |
| 17028 | *DNAH5* | -0.00153 | -0.61675 | -0.86367 | -1.4788 | -0.49 |
| 17029 | *ORC5* | 1.8792 | -1.5849 | -1.7766 | -2.1125 | -0.49 |
| 17030 | *ANGPTL4* | 0.56229 | -1.0187 | -1.027 | -1.7667 | -0.49 |
| 17031 | *PTER* | -0.39136 | -0.4594 | -0.63332 | -1.1133 | -0.49 |
| 17032 | *ORC3* | 1.2293 | -0.91236 | -1.8022 | -2.5066 | -0.50 |
| 17033 | *SHANK1* | 0.47781 | -0.75192 | -1.2125 | -1.7882 | -0.50 |
| 17034 | *KIF12* | 0.48618 | -0.67441 | -1.2985 | -1.5719 | -0.50 |
| 17035 | *SIX4* | 0.81435 | -0.27303 | -2.0301 | -3.2089 | -0.50 |
| 17036 | *RPS17* | 0.95397 | -0.93789 | -1.5049 | -2.7728 | -0.50 |
| 17037 | *PSMG2* | 0.31569 | -0.49953 | -1.3055 | -2.6208 | -0.50 |
| 17038 | *PHLDB1* | -0.35675 | -0.52004 | -0.61281 | -2.9473 | -0.50 |
| 17039 | *PLA2G2C* | 0.28896 | -0.29194 | -1.4869 | -2.0277 | -0.50 |
| 17040 | *PRDX1* | -0.07432 | -0.35894 | -1.058 | -1.2411 | -0.50 |
| 17041 | *RPS20* | -0.14244 | -0.51406 | -0.83676 | -1.5434 | -0.50 |
| 17042 | *PNISR* | 0.60934 | -0.65875 | -1.445 | -2.6275 | -0.50 |
| 17043 | *GRINA* | -0.08702 | -0.47227 | -0.93525 | -1.7306 | -0.50 |
| 17044 | *PDAP1* | 2.0084 | -1.3371 | -2.1685 | -4.0172 | -0.50 |
| 17045 | *TRDN* | 0.48255 | -0.89059 | -1.0893 | -1.8538 | -0.50 |
| 17046 | *MYO1B* | 0.049555 | -0.62065 | -0.92629 | -1.925 | -0.50 |
| 17047 | *FAM35A* | 0.4737 | -0.9471 | -1.0247 | -1.1831 | -0.50 |
| 17048 | *MAP1LC3B2* | -0.06833 | -0.6835 | -0.74672 | -0.94252 | -0.50 |
| 17049 | *ACP5* | -0.17755 | -0.60223 | -0.71893 | -1.0905 | -0.50 |
| 17050 | *VEGFA* | 0.21219 | -0.81701 | -0.89401 | -1.7709 | -0.50 |
| 17051 | *DEFB134* | -0.1433 | -0.18068 | -1.1752 | -3.0483 | -0.50 |
| 17052 | *LZTS1* | -0.30379 | -0.31582 | -0.88008 | -2.1085 | -0.50 |
| 17053 | *RPS6* | 0.25446 | -0.48659 | -1.268 | -2.3871 | -0.50 |
| 17054 | *TK1* | 0.19547 | -0.81535 | -0.88062 | -1.2763 | -0.50 |
| 17055 | *TRPM3* | -0.01103 | -0.46167 | -1.0279 | -1.6456 | -0.50 |
| 17056 | *DDX5* | -0.2372 | -0.59647 | -0.66865 | -1.9673 | -0.50 |
| 17057 | *TCF15* | 0.21667 | -0.68934 | -1.0297 | -3.0314 | -0.50 |
| 17058 | *ELP3* | 1.2313 | -1.3478 | -1.3864 | -2.7546 | -0.50 |
| 17059 | *RAB38* | 0.1661 | 0.049677 | -1.7209 | -1.855 | -0.50 |

| | | | | | | |
|---|---|---|---|---|---|---|
| 17060 | *POLR2D* | 0.37192 | -0.54373 | -1.334 | -1.6524 | -0.50 |
| 17061 | *SYCP2L* | 0.15064 | 0.14306 | -1.8007 | -1.8813 | -0.50 |
| 17062 | *IKZF3* | -0.25603 | -0.30371 | -0.94739 | -3.1458 | -0.50 |
| 17063 | *CLCNKA* | -0.46286 | -0.50046 | -0.54391 | -2.1264 | -0.50 |
| 17064 | *TP53I13* | -0.42497 | -0.43244 | -0.65023 | -1.7497 | -0.50 |
| 17065 | *EIF2S3* | 0.98173 | -0.35571 | -2.1347 | -2.1794 | -0.50 |
| 17066 | *SYPL2* | -0.01264 | -0.43481 | -1.0613 | -2.1496 | -0.50 |
| 17067 | *SMEK2* | 0.5355 | -0.07895 | -1.9656 | -2.4946 | -0.50 |
| 17068 | *SUPT6H* | 0.96942 | -0.8198 | -1.6614 | -2.5292 | -0.50 |
| 17069 | *ZFYVE1* | -0.22496 | -0.26125 | -1.0267 | -1.6597 | -0.50 |
| 17070 | *PSMD8* | -0.04644 | -0.56972 | -0.89794 | -1.03 | -0.50 |
| 17071 | *AURKA* | -0.10247 | -0.31965 | -1.0927 | -1.1343 | -0.50 |
| 17072 | *VDAC1* | -0.22153 | -0.28173 | -1.0136 | -2.8702 | -0.51 |
| 17073 | *DNMT1* | 0.03708 | -0.52329 | -1.0313 | -2.3676 | -0.51 |
| 17074 | *ALDH3A2* | 0.090843 | -0.44353 | -1.1656 | -1.6759 | -0.51 |
| 17075 | *PPP1R37* | -0.21393 | -0.61073 | -0.69463 | -1.9696 | -0.51 |
| 17076 | *ARNT* | 0.82325 | -1.1605 | -1.1825 | -1.8842 | -0.51 |
| 17077 | *VNN3* | 0.69777 | -0.85002 | -1.368 | -1.6356 | -0.51 |
| 17078 | *CFL2* | -0.17731 | -0.50087 | -0.84426 | -1.4233 | -0.51 |
| 17079 | *BMF* | 0.067212 | -0.46762 | -1.1223 | -1.3008 | -0.51 |
| 17080 | *SARS2* | 0.22517 | -0.8108 | -0.93865 | -1.1137 | -0.51 |
| 17081 | *CNOT6* | 0.66404 | -0.78672 | -1.4024 | -3.2901 | -0.51 |
| 17082 | *BUB1B* | -0.25545 | -0.61167 | -0.6593 | -1.5971 | -0.51 |
| 17083 | *RAB7B* | -0.10911 | -0.25515 | -1.163 | -1.5289 | -0.51 |
| 17084 | *RAE1* | 0.58442 | -0.44353 | -1.6684 | -2.1089 | -0.51 |
| 17085 | *COX7C* | 0.89619 | -0.44456 | -1.9792 | -2.3086 | -0.51 |
| 17086 | *KLHL40* | 2.1693 | -1.4996 | -2.1984 | -2.5478 | -0.51 |
| 17087 | *PRDM13* | -0.08039 | -0.66107 | -0.78807 | -0.8597 | -0.51 |
| 17088 | *C17orf58* | -0.30975 | -0.40465 | -0.81539 | -1.4545 | -0.51 |
| 17089 | *ZCCHC3* | 1.4409 | -1.4507 | -1.5225 | -1.7134 | -0.51 |
| 17090 | *SPIRE2* | 0.16177 | -0.7757 | -0.91944 | -1.6847 | -0.51 |
| 17091 | *R3HDM2* | 0.3084 | -0.78102 | -1.0614 | -1.4507 | -0.51 |
| 17092 | *STK26* | 0.36192 | -0.6182 | -1.278 | -1.7077 | -0.51 |
| 17093 | *PRDM10* | -0.00546 | -0.68825 | -0.84091 | -2.8051 | -0.51 |
| 17094 | *FAM114A1* | -0.02854 | -0.53328 | -0.97303 | -1.6689 | -0.51 |
| 17095 | *GATA5* | 0.25931 | -0.2636 | -1.531 | -1.8066 | -0.51 |
| 17096 | *SEC22A* | -0.1635 | -0.38815 | -0.98421 | -1.6049 | -0.51 |
| 17097 | *ABHD17A* | -0.0378 | -0.44955 | -1.0494 | -1.9172 | -0.51 |
| 17098 | *CLEC1B* | 0.53888 | -0.63172 | -1.445 | -2.5392 | -0.51 |
| 17099 | *HIST1H2AE* | 0.23442 | -0.70957 | -1.0635 | -1.7935 | -0.51 |
| 17100 | *ADAMTS1* | -0.14026 | -0.42884 | -0.97252 | -1.4265 | -0.51 |
| 17101 | *C10orf2* | 0.057793 | -0.65169 | -0.94816 | -1.5335 | -0.51 |
| 17102 | *TMEM39A* | 1.9267 | -1.5382 | -1.9308 | -2.1404 | -0.51 |
| 17103 | *CLSTN2* | 0.44955 | -0.82996 | -1.1631 | -2.3244 | -0.51 |
| 17104 | *MARCH8* | 1.0697 | -1.1459 | -1.4681 | -1.8157 | -0.51 |
| 17105 | *C2orf69* | 1.1479 | -0.79365 | -1.9013 | -3.9399 | -0.52 |

| | | | | | | |
|---|---|---|---|---|---|---|
| 17106 | *C2orf50* | 0.17672 | -0.62881 | -1.0954 | -1.7989 | -0.52 |
| 17107 | *RNF208* | -0.04678 | -0.54884 | -0.95321 | -1.0312 | -0.52 |
| 17108 | *RIT2* | 0.65454 | -0.76725 | -1.4368 | -1.5684 | -0.52 |
| 17109 | *SLC12A7* | 0.006708 | -0.25981 | -1.2968 | -1.4625 | -0.52 |
| 17110 | *DYNC1I1* | -0.04898 | -0.43608 | -1.0656 | -1.3085 | -0.52 |
| 17111 | *GNA15* | 0.27992 | -0.34353 | -1.4871 | -1.7145 | -0.52 |
| 17112 | *GADD45GIP1* | 0.66953 | -0.95045 | -1.2705 | -1.9912 | -0.52 |
| 17113 | *GRAP* | 0.15694 | -0.74875 | -0.96144 | -2.201 | -0.52 |
| 17114 | *VPS13C* | -0.00822 | -0.30424 | -1.2411 | -1.631 | -0.52 |
| 17115 | *INO80* | 0.031661 | -0.23658 | -1.3504 | -2.5008 | -0.52 |
| 17116 | *TEPP* | 0.44957 | -0.74774 | -1.2572 | -1.6072 | -0.52 |
| 17117 | *NR2E1* | 0.56529 | -0.97944 | -1.1415 | -1.4258 | -0.52 |
| 17118 | *LRRC66* | -0.07406 | -0.62889 | -0.85389 | -2.4829 | -0.52 |
| 17119 | *KDM1A* | -0.3417 | -0.5329 | -0.6828 | -1.241 | -0.52 |
| 17120 | *CDH22* | -0.36728 | -0.49694 | -0.69319 | -1.6343 | -0.52 |
| 17121 | *NGEF* | -0.05429 | -0.71908 | -0.78545 | -1.2171 | -0.52 |
| 17122 | *NAA25* | 1.3134 | -1.3118 | -1.5633 | -1.6484 | -0.52 |
| 17123 | *INCENP* | 0.020499 | -0.35193 | -1.2304 | -1.4507 | -0.52 |
| 17124 | *SHQ1* | 0.065948 | -0.53346 | -1.095 | -1.8724 | -0.52 |
| 17125 | *ZNF138* | 1.1135 | -1.1388 | -1.5375 | -1.7078 | -0.52 |
| 17126 | *UBE2V1* | 1.0072 | -0.5409 | -2.0303 | -2.1198 | -0.52 |
| 17127 | *TRA2B* | -0.18283 | -0.27462 | -1.1089 | -1.4607 | -0.52 |
| 17128 | *ITFG2* | 0.25715 | -0.2263 | -1.5976 | -2.1361 | -0.52 |
| 17129 | *LGI2* | 2.2085 | -1.4741 | -2.3036 | -2.6343 | -0.52 |
| 17130 | *TRIM31* | 0.6523 | -0.62476 | -1.5978 | -1.6597 | -0.52 |
| 17131 | *ARHGEF35* | 0.11062 | -0.28928 | -1.392 | -1.9296 | -0.52 |
| 17132 | *ARHGAP11A* | 0.38653 | -0.79365 | -1.1644 | -1.1904 | -0.52 |
| 17133 | *PNPT1* | 0.009675 | -0.0175 | -1.5649 | -2.1755 | -0.52 |
| 17134 | *SLC25A30* | -0.24501 | -0.31052 | -1.0172 | -2.0613 | -0.52 |
| 17135 | *UGT2B28* | 0.71957 | -0.77805 | -1.5153 | -1.55 | -0.52 |
| 17136 | *ATRN* | 0.19524 | -0.83773 | -0.93217 | -1.329 | -0.52 |
| 17137 | *RPS19* | 1.4112 | -1.4684 | -1.5182 | -1.6604 | -0.53 |
| 17138 | *PTGES* | 0.67358 | -0.86494 | -1.3851 | -1.7235 | -0.53 |
| 17139 | *BARX2* | 0.2313 | -0.89494 | -0.91283 | -1.258 | -0.53 |
| 17140 | *FAM96A* | 0.23847 | -0.32558 | -1.4894 | -1.5756 | -0.53 |
| 17141 | *BFSP1* | -0.21557 | -0.44353 | -0.91793 | -1.714 | -0.53 |
| 17142 | *NCOR2* | -0.09468 | -0.43488 | -1.0475 | -1.4754 | -0.53 |
| 17143 | *SPCS3* | 0.33517 | -0.85163 | -1.0621 | -1.1384 | -0.53 |
| 17144 | *GTPBP10* | 1.4618 | -1.1765 | -1.864 | -2.9931 | -0.53 |
| 17145 | *SULT1A4* | 0.24869 | -0.69502 | -1.1343 | -1.6526 | -0.53 |
| 17146 | *CSH2* | -0.01493 | -0.68032 | -0.88634 | -1.7963 | -0.53 |
| 17147 | *COX6A1* | 0.11649 | -0.39248 | -1.3058 | -1.4034 | -0.53 |
| 17148 | *OTOF* | -0.1715 | -0.25606 | -1.1545 | -2.0042 | -0.53 |
| 17149 | *RPS18* | 0.42874 | -0.91642 | -1.0953 | -1.6434 | -0.53 |
| 17150 | *SSTR1* | 0.49299 | -0.72095 | -1.3565 | -1.4185 | -0.53 |
| 17151 | *SLC35F3* | 0.30105 | -0.6066 | -1.2815 | -1.7496 | -0.53 |

| | | | | | | |
|---|---|---|---|---|---|---|
| 17152 | *ZNF257* | -0.05909 | -0.45491 | -1.0734 | -1.4715 | -0.53 |
| 17153 | *PPDPF* | -0.24773 | -0.57546 | -0.76437 | -0.86158 | -0.53 |
| 17154 | *POLD1* | 0.64251 | -0.29054 | -1.94 | -3.4611 | -0.53 |
| 17155 | *WDR61* | 0.82532 | -1.1011 | -1.3125 | -1.6383 | -0.53 |
| 17156 | *GATAD2B* | -0.09182 | -0.21955 | -1.2786 | -2.5414 | -0.53 |
| 17157 | *SYCP3* | -0.36868 | -0.44043 | -0.78096 | -1.5781 | -0.53 |
| 17158 | *MLANA* | 0.65646 | -0.61264 | -1.6339 | -2.0697 | -0.53 |
| 17159 | *DENND1C* | -0.14859 | -0.61092 | -0.83278 | -1.1685 | -0.53 |
| 17160 | *TMPRSS6* | -0.08207 | -0.08601 | -1.4246 | -1.9789 | -0.53 |
| 17161 | *MRPL12* | 0.31394 | -0.49941 | -1.4103 | -1.4252 | -0.53 |
| 17162 | *ISY1-RAB43* | -0.02583 | -0.08265 | -1.4873 | -1.7874 | -0.53 |
| 17163 | *RPRD1B* | -0.11437 | -0.30834 | -1.1734 | -1.9768 | -0.53 |
| 17164 | *COA3* | -0.17281 | -0.44353 | -0.98103 | -1.3346 | -0.53 |
| 17165 | *ARID3A* | 1.055 | -1.1461 | -1.5068 | -1.8849 | -0.53 |
| 17166 | *OR1L6* | 0.66958 | -1.1224 | -1.1463 | -1.3837 | -0.53 |
| 17167 | *RPL41* | 0.6239 | -0.81152 | -1.4156 | -1.8041 | -0.53 |
| 17168 | *PIGH* | -0.46583 | -0.47472 | -0.66675 | -0.98902 | -0.54 |
| 17169 | *HUWE1* | 0.52236 | -0.89518 | -1.2356 | -1.5052 | -0.54 |
| 17170 | *SETDB2* | 2.067 | -1.5125 | -2.163 | -2.224 | -0.54 |
| 17171 | *SEC61G* | 0.48253 | -0.39603 | -1.695 | -1.773 | -0.54 |
| 17172 | *OR4F29* | -0.44353 | -0.45663 | -0.70993 | -2.0062 | -0.54 |
| 17173 | *RASSF3* | -0.45656 | -0.55891 | -0.59462 | -1.6626 | -0.54 |
| 17174 | *PDE6G* | 1.1539 | -0.76366 | -2.0016 | -3.4888 | -0.54 |
| 17175 | *TREM2* | -0.28717 | -0.65417 | -0.67137 | -2.1477 | -0.54 |
| 17176 | *TMEM8B* | 1.0895 | -1.244 | -1.4597 | -2.4078 | -0.54 |
| 17177 | *OR5I1* | 0.57298 | -1.0344 | -1.1533 | -1.433 | -0.54 |
| 17178 | *FAM25A* | 0.11779 | -0.781 | -0.95224 | -1.8129 | -0.54 |
| 17179 | *GPRASP1* | 0.75553 | -0.50872 | -1.8623 | -1.9163 | -0.54 |
| 17180 | *WDR47* | 0.17526 | -0.01493 | -1.7766 | -1.7874 | -0.54 |
| 17181 | *NPAS3* | 0.34144 | -0.77697 | -1.181 | -1.6071 | -0.54 |
| 17182 | *WSB1* | 0.58786 | -0.95933 | -1.2474 | -1.6435 | -0.54 |
| 17183 | *HEXA* | -0.08395 | -0.44353 | -1.0917 | -1.4998 | -0.54 |
| 17184 | *HDGFRP2* | 1.7733 | -1.4027 | -1.99 | -2.7277 | -0.54 |
| 17185 | *SUGCT* | -0.301 | -0.58561 | -0.73612 | -0.81044 | -0.54 |
| 17186 | *BAZ1B* | -0.23339 | -0.58058 | -0.80936 | -1.2576 | -0.54 |
| 17187 | *NDST2* | -0.24262 | -0.6573 | -0.72341 | -1.622 | -0.54 |
| 17188 | *MT1M* | 0.39528 | -0.86047 | -1.1585 | -1.3976 | -0.54 |
| 17189 | *MBOAT4* | -0.16205 | -0.18125 | -1.2815 | -2.2064 | -0.54 |
| 17190 | *EIF3C* | 0.22156 | -0.85739 | -0.99059 | -2.1143 | -0.54 |
| 17191 | *SYBU* | 0.19377 | -0.79225 | -1.0283 | -2.8174 | -0.54 |
| 17192 | *PRKCB* | 2.2641 | -1.4644 | -2.4268 | -3.2057 | -0.54 |
| 17193 | *CGB5* | 0.59007 | -1.0726 | -1.1446 | -1.8557 | -0.54 |
| 17194 | *KIAA0355* | -0.3552 | -0.47485 | -0.79711 | -1.8157 | -0.54 |
| 17195 | *P3H4* | 0.61834 | 0.091413 | -2.337 | -2.4481 | -0.54 |
| 17196 | *ZNHIT1* | -0.09459 | -0.33315 | -1.2004 | -1.5943 | -0.54 |
| 17197 | *NDUFV1* | 0.19772 | -0.36885 | -1.4584 | -1.4841 | -0.54 |

| | | | | | | |
|---|---|---|---|---|---|---|
| 17198 | *ZNF543* | 0.28432 | -0.86506 | -1.049 | -2.1523 | -0.54 |
| 17199 | *ZNF251* | 0.75465 | -1.1421 | -1.2438 | -1.4893 | -0.54 |
| 17200 | *MTA1* | 0.083912 | -0.17554 | -1.5408 | -1.571 | -0.54 |
| 17201 | *RIC8B* | 0.7514 | -0.76854 | -1.6158 | -2.0208 | -0.54 |
| 17202 | *FDXACB1* | 0.54445 | -0.26714 | -1.9107 | -5.1074 | -0.54 |
| 17203 | *PTCD1* | 0.54005 | -0.24569 | -1.9305 | -2.7509 | -0.55 |
| 17204 | *GEMIN7* | -0.03303 | -0.40731 | -1.1959 | -1.4423 | -0.55 |
| 17205 | *GPANK1* | 0.44576 | -0.26177 | -1.8212 | -2.4394 | -0.55 |
| 17206 | *SRSF6* | 1.1921 | -1.3465 | -1.4829 | -2.0363 | -0.55 |
| 17207 | *XKR5* | 0.76884 | -0.97786 | -1.4287 | -2.7941 | -0.55 |
| 17208 | *DIMT1* | 0.043666 | -0.83874 | -0.84411 | -2.565 | -0.55 |
| 17209 | *PHB* | -0.39997 | -0.49987 | -0.74013 | -1.6785 | -0.55 |
| 17210 | *FBP1* | 0.29731 | -0.39932 | -1.539 | -1.9494 | -0.55 |
| 17211 | *BRSK2* | -0.32637 | -0.61779 | -0.69915 | -1.1807 | -0.55 |
| 17212 | *TXNRD1* | -0.38461 | -0.44353 | -0.81545 | -2.4047 | -0.55 |
| 17213 | *RPL13* | 0.3407 | -0.347 | -1.6373 | -1.7963 | -0.55 |
| 17214 | *DNAJC25-GNG10* | 0.4896 | -0.64826 | -1.4852 | -1.8348 | -0.55 |
| 17215 | *TMEM138* | -0.28335 | -0.67321 | -0.68813 | -1.1845 | -0.55 |
| 17216 | *NHP2L1* | 0.19141 | -0.6951 | -1.141 | -1.9865 | -0.55 |
| 17217 | *SOX12* | 0.48795 | -0.2875 | -1.8454 | -3.0687 | -0.55 |
| 17218 | *ARMC12* | -0.26891 | -0.60048 | -0.77559 | -1.2302 | -0.55 |
| 17219 | *HEY2* | 0.65856 | -0.85739 | -1.4485 | -2.5308 | -0.55 |
| 17220 | *RPLP2* | 1.1085 | -1.2034 | -1.5525 | -1.6047 | -0.55 |
| 17221 | *TCP10* | 0.58763 | -0.46757 | -1.7702 | -3.3455 | -0.55 |
| 17222 | *TTYH1* | 0.19908 | -0.69286 | -1.1566 | -1.2936 | -0.55 |
| 17223 | *AARS2* | 0.98058 | -1.1676 | -1.4636 | -2.1187 | -0.55 |
| 17224 | *ADAMTSL3* | 0.90732 | -0.63971 | -1.9195 | -2.4015 | -0.55 |
| 17225 | *SARAF* | -0.16546 | -0.51133 | -0.97588 | -1.8858 | -0.55 |
| 17226 | *HPGDS* | -0.19614 | -0.24148 | -1.2171 | -3.0154 | -0.55 |
| 17227 | *TRIP6* | 0.74282 | -0.63405 | -1.7652 | -1.9534 | -0.55 |
| 17228 | *ADCK3* | 0.6609 | -0.79788 | -1.5195 | -1.8275 | -0.55 |
| 17229 | *NAT2* | -0.08556 | -0.76033 | -0.8108 | -2.3265 | -0.55 |
| 17230 | *EIF2B5* | 0.64019 | -1.1263 | -1.1713 | -1.2692 | -0.55 |
| 17231 | *ATXN10* | -0.0083 | -0.75099 | -0.89832 | -1.8629 | -0.55 |
| 17232 | *PAPOLG* | -0.34356 | -0.35665 | -0.95803 | -1.8112 | -0.55 |
| 17233 | *ST3GAL5* | 1.0411 | -1.3453 | -1.3542 | -1.6218 | -0.55 |
| 17234 | *CACNG7* | 0.047153 | -0.68706 | -1.0187 | -1.2995 | -0.55 |
| 17235 | *DUSP12* | -0.25794 | -0.31884 | -1.0819 | -2.0423 | -0.55 |
| 17236 | *CCNA2* | -0.37678 | -0.55815 | -0.72488 | -0.98434 | -0.55 |
| 17237 | *CASK* | -0.14207 | -0.16377 | -1.3542 | -1.7109 | -0.55 |
| 17238 | *XRCC4* | 0.71631 | -0.84523 | -1.5348 | -3.5736 | -0.55 |
| 17239 | *WIZ* | 0.3454 | -0.36877 | -1.6413 | -1.7971 | -0.55 |
| 17240 | *SLC38A10* | 0.58487 | -0.44246 | -1.8076 | -2.4607 | -0.56 |
| 17241 | *DDB1* | 0.2083 | -0.86494 | -1.0093 | -1.7745 | -0.56 |
| 17242 | *VEZT* | 0.28506 | -0.59791 | -1.3542 | -1.6663 | -0.56 |
| 17243 | *HMGXB3* | 0.48317 | -0.57404 | -1.5762 | -1.8606 | -0.56 |

| | | | | | | |
|---|---|---|---|---|---|---|
| 17244 | *LIMCH1* | 0.72405 | -0.76655 | -1.6251 | -3.301 | -0.56 |
| 17245 | *HERC2* | -0.0757 | -0.54739 | -1.0448 | -1.1048 | -0.56 |
| 17246 | *SEH1L* | 0.66963 | -1.151 | -1.1881 | -1.2867 | -0.56 |
| 17247 | *SERPINB3* | 0.2066 | -0.69561 | -1.1817 | -1.6867 | -0.56 |
| 17248 | *SPC25* | -0.11594 | -0.50453 | -1.0528 | -1.8208 | -0.56 |
| 17249 | *CCM2L* | 0.23953 | -0.26055 | -1.6526 | -2.0421 | -0.56 |
| 17250 | *TUBB2A* | -0.37929 | -0.60781 | -0.68684 | -0.79043 | -0.56 |
| 17251 | *GCSH* | 0.074813 | -0.22048 | -1.5289 | -1.5959 | -0.56 |
| 17252 | *MMRN2* | 0.34354 | -0.42415 | -1.5954 | -2.614 | -0.56 |
| 17253 | *VAV3* | 0.3629 | -0.90549 | -1.1337 | -1.1825 | -0.56 |
| 17254 | *PRAM1* | -0.44951 | -0.59521 | -0.63172 | -1.2572 | -0.56 |
| 17255 | *ACP6* | 0.16799 | -0.60494 | -1.2399 | -1.2837 | -0.56 |
| 17256 | *DMP1* | 0.049395 | 0.013351 | -1.7397 | -1.7935 | -0.56 |
| 17257 | *C16orf13* | -0.28012 | -0.6093 | -0.78863 | -2.1576 | -0.56 |
| 17258 | *TBC1D3* | -0.49539 | -0.5837 | -0.59937 | -2.0642 | -0.56 |
| 17259 | *ANKRD16* | -0.01885 | -0.81087 | -0.8489 | -1.7855 | -0.56 |
| 17260 | *HYAL1* | -0.08207 | -0.74572 | -0.85163 | -1.3593 | -0.56 |
| 17261 | *PLD2* | 0.67212 | -0.93878 | -1.4128 | -1.6027 | -0.56 |
| 17262 | *EIF3A* | -0.3703 | -0.43251 | -0.87737 | -1.5544 | -0.56 |
| 17263 | *LIAS* | 1.1993 | -1.3948 | -1.4852 | -1.8456 | -0.56 |
| 17264 | *NECAB1* | 0.40902 | -0.93314 | -1.1566 | -2.0384 | -0.56 |
| 17265 | *NCKAP1* | -0.03704 | -0.50828 | -1.1354 | -1.677 | -0.56 |
| 17266 | *AP3B1* | 0.098257 | -0.29186 | -1.4873 | -1.9686 | -0.56 |
| 17267 | *LSM6* | -0.50676 | -0.54537 | -0.62881 | -1.3264 | -0.56 |
| 17268 | *MAP1S* | 0.16173 | -0.15085 | -1.6934 | -2.3534 | -0.56 |
| 17269 | *NUS1* | 0.08783 | -0.86123 | -0.91013 | -1.165 | -0.56 |
| 17270 | *WDR88* | 0.81063 | -1.0481 | -1.4473 | -2.1187 | -0.56 |
| 17271 | *SLC13A1* | 2.1575 | -1.8642 | -1.9789 | -2.9471 | -0.56 |
| 17272 | *SOAT2* | 0.50568 | -1.0019 | -1.19 | -1.4049 | -0.56 |
| 17273 | *TRAPPC4* | 0.47342 | -0.74013 | -1.4197 | -2.6024 | -0.56 |
| 17274 | *ANO8* | 0.062547 | -0.83143 | -0.91823 | -1.1587 | -0.56 |
| 17275 | *NUPR1* | 0.79418 | -1.2163 | -1.2655 | -2.4718 | -0.56 |
| 17276 | *SNX16* | -0.15209 | -0.75944 | -0.77661 | -1.0261 | -0.56 |
| 17277 | *SNX29* | 0.93709 | -1.0947 | -1.5306 | -1.6255 | -0.56 |
| 17278 | *CNP* | -0.10661 | -0.22292 | -1.3611 | -2.1354 | -0.56 |
| 17279 | *MAU2* | 0.061984 | -0.81313 | -0.93967 | -1.0748 | -0.56 |
| 17280 | *HYAL3* | -0.05926 | -0.37045 | -1.2616 | -2.071 | -0.56 |
| 17281 | *NRARP* | 0.13043 | -0.78823 | -1.0338 | -1.2429 | -0.56 |
| 17282 | *AKR7A2* | 0.6852 | -1.1067 | -1.2717 | -1.5722 | -0.56 |
| 17283 | *XKR8* | -0.48254 | -0.5581 | -0.65291 | -1.4942 | -0.56 |
| 17284 | *TOP1* | 0.51396 | -0.75076 | -1.4579 | -2.3118 | -0.56 |
| 17285 | *ASGR2* | -0.3337 | -0.60147 | -0.76055 | -1.6525 | -0.57 |
| 17286 | *BTF3* | -0.25265 | -0.38349 | -1.06 | -1.36 | -0.57 |
| 17287 | *UBXN4* | 0.18898 | -0.86047 | -1.0255 | -1.1866 | -0.57 |
| 17288 | *SRRM3* | 0.3314 | -0.43264 | -1.5959 | -2.2449 | -0.57 |
| 17289 | *AKAP12* | 0.8377 | -0.60658 | -1.9284 | -3.1665 | -0.57 |

| | | | | | | |
|---|---|---|---|---|---|---|
| 17290 | *HEATR4* | 0.2896 | -0.25844 | -1.7296 | -2.3573 | -0.57 |
| 17291 | *UPK1B* | 0.76505 | -1.2261 | -1.2386 | -1.5103 | -0.57 |
| 17292 | *DGKB* | 0.70768 | -0.39775 | -2.0113 | -2.7254 | -0.57 |
| 17293 | *DDX56* | -0.32505 | -0.62514 | -0.75227 | -0.77401 | -0.57 |
| 17294 | *INTS5* | 0.22032 | -0.82848 | -1.0954 | -1.9065 | -0.57 |
| 17295 | *CYP2R1* | 0.72838 | -0.8514 | -1.5814 | -1.7914 | -0.57 |
| 17296 | *PLCB4* | -0.21893 | -0.58771 | -0.89866 | -1.3242 | -0.57 |
| 17297 | *IARS2* | 0.62503 | -1.0741 | -1.2566 | -1.7956 | -0.57 |
| 17298 | *CCDC130* | -0.41803 | -0.48021 | -0.80909 | -2.0126 | -0.57 |
| 17299 | *PSMB4* | 0.33302 | -0.80567 | -1.2365 | -1.3003 | -0.57 |
| 17300 | *DDX11* | -0.39759 | -0.44353 | -0.86903 | -0.87673 | -0.57 |
| 17301 | *MEPE* | 0.52306 | -0.73071 | -1.5039 | -1.507 | -0.57 |
| 17302 | *PIGY* | 0.31212 | 0.31074 | -2.3357 | -2.4912 | -0.57 |
| 17303 | *HIST1H4K* | -0.24196 | -0.68416 | -0.78676 | -1.1284 | -0.57 |
| 17304 | *NAA30* | 1.035 | -1.3183 | -1.4312 | -2.0542 | -0.57 |
| 17305 | *CLDN22* | -0.36035 | -0.596 | -0.75876 | -1.8154 | -0.57 |
| 17306 | *NCCRP1* | -0.03976 | -0.76966 | -0.90768 | -0.95545 | -0.57 |
| 17307 | *ZBTB10* | -0.07743 | -0.23638 | -1.4046 | -2.0384 | -0.57 |
| 17308 | *NPRL3* | 1.3795 | -1.2167 | -1.882 | -2.9338 | -0.57 |
| 17309 | *MRPL3* | 0.29893 | -0.83631 | -1.1872 | -2.8928 | -0.57 |
| 17310 | *SLC25A15* | -0.18414 | -0.38361 | -1.1575 | -1.5305 | -0.58 |
| 17311 | *FGF2* | -0.44353 | -0.50793 | -0.77418 | -2.1648 | -0.58 |
| 17312 | *SPG7* | 0.60515 | -0.94052 | -1.3903 | -1.5837 | -0.58 |
| 17313 | *LSG1* | -0.00975 | -0.67764 | -1.0395 | -1.388 | -0.58 |
| 17314 | *ICA1* | 0.35232 | -1.0231 | -1.0564 | -1.6561 | -0.58 |
| 17315 | *NEURL3* | -0.28571 | -0.4187 | -1.0228 | -3.1568 | -0.58 |
| 17316 | *RSPH6A* | 0.31179 | -0.37358 | -1.6663 | -2.2473 | -0.58 |
| 17317 | *GTF3C1* | 0.17359 | -0.70193 | -1.2006 | -2.381 | -0.58 |
| 17318 | *MINOS1-NBL1* | 1.6754 | -1.5354 | -1.8698 | -2.1066 | -0.58 |
| 17319 | *TNMD* | 0.49695 | -0.56069 | -1.6663 | -1.9678 | -0.58 |
| 17320 | *NLRP3* | 0.25068 | -0.48131 | -1.4996 | -1.8157 | -0.58 |
| 17321 | *FAM69B* | -0.47944 | -0.48646 | -0.7647 | -0.9562 | -0.58 |
| 17322 | *HMMR* | 0.59762 | -0.7503 | -1.5781 | -1.9513 | -0.58 |
| 17323 | *GTSF1* | 0.019815 | -0.00564 | -1.7459 | -2.3222 | -0.58 |
| 17324 | *RHBDF2* | 1.4195 | -1.1169 | -2.0351 | -6.0397 | -0.58 |
| 17325 | *PSMD14* | 0.12199 | -0.46951 | -1.385 | -1.5138 | -0.58 |
| 17326 | *C15orf52* | 0.33119 | 0.021696 | -2.0871 | -2.1687 | -0.58 |
| 17327 | *MPND* | 0.27706 | -0.91374 | -1.0989 | -1.9811 | -0.58 |
| 17328 | *PVRL1* | -0.24019 | -0.62307 | -0.87318 | -2.3795 | -0.58 |
| 17329 | *TREX1* | 0.27744 | -0.32279 | -1.6921 | -2.1648 | -0.58 |
| 17330 | *C11orf31* | 0.29798 | -0.7512 | -1.2867 | -1.5965 | -0.58 |
| 17331 | *C20orf27* | 0.33101 | -0.63405 | -1.438 | -1.6001 | -0.58 |
| 17332 | *USP17L19* | 0.12853 | -0.27421 | -1.5957 | -2.1132 | -0.58 |
| 17333 | *NDUFB4* | 0.1696 | -0.4433 | -1.4685 | -1.8906 | -0.58 |
| 17334 | *SRGAP2B* | 0.3929 | -0.46767 | -1.6676 | -1.8066 | -0.58 |
| 17335 | *TP53I3* | 0.11456 | -0.8478 | -1.01 | -1.1921 | -0.58 |

| | | | | | | |
|---|---|---|---|---|---|---|
| 17336 | *IKBKAP* | -0.3129 | -0.71381 | -0.71839 | -0.75128 | -0.58 |
| 17337 | *SDF2L1* | 1.2521 | -1.1429 | -1.8557 | -1.8636 | -0.58 |
| 17338 | *GBAS* | -0.29865 | -0.34443 | -1.1035 | -1.1804 | -0.58 |
| 17339 | *SEMA6D* | 0.45531 | -0.18353 | -2.0197 | -4.1058 | -0.58 |
| 17340 | *EHBP1L1* | 0.34151 | -0.21666 | -1.8735 | -1.9789 | -0.58 |
| 17341 | *CSE1L* | 1.0214 | -1.3068 | -1.4638 | -1.6707 | -0.58 |
| 17342 | *SLC27A1* | -0.29953 | -0.71093 | -0.74011 | -2.7689 | -0.58 |
| 17343 | *LRRC37B* | 1.0789 | -0.85392 | -1.9757 | -2.5452 | -0.58 |
| 17344 | *DHX9* | 0.40797 | -0.71717 | -1.4416 | -1.9736 | -0.58 |
| 17345 | *RYR3* | 0.26123 | -0.98314 | -1.0297 | -1.2071 | -0.58 |
| 17346 | *C4orf36* | 1.3952 | -1.5054 | -1.6419 | -1.8557 | -0.58 |
| 17347 | *GABARAP* | 0.97532 | -1.13 | -1.5992 | -2.2075 | -0.58 |
| 17348 | *PMF1-BGLAP* | -0.29124 | -0.41575 | -1.0469 | -1.8872 | -0.58 |
| 17349 | *RPL14* | -0.40893 | -0.44353 | -0.90256 | -1.8759 | -0.59 |
| 17350 | *DPH2* | -0.00546 | -0.76366 | -0.98653 | -1.7766 | -0.59 |
| 17351 | *ZNF586* | -0.13231 | -0.56427 | -1.0596 | -1.5306 | -0.59 |
| 17352 | *STAMBPL1* | 0.29943 | -0.77418 | -1.2816 | -2.2385 | -0.59 |
| 17353 | *YPEL5* | -0.37194 | -0.42701 | -0.95858 | -1.5877 | -0.59 |
| 17354 | *RPS24* | 0.22514 | -0.27155 | -1.7119 | -2.6101 | -0.59 |
| 17355 | *TMEM136* | -0.15514 | -0.53684 | -1.0669 | -1.5671 | -0.59 |
| 17356 | *CPTP* | 0.27172 | -0.65764 | -1.3736 | -1.7855 | -0.59 |
| 17357 | *TRIM63* | -0.23805 | -0.47263 | -1.0493 | -1.4234 | -0.59 |
| 17358 | *VPS36* | -0.48492 | -0.59662 | -0.67887 | -2.221 | -0.59 |
| 17359 | *MUL1* | -0.24904 | -0.25583 | -1.2568 | -2.0752 | -0.59 |
| 17360 | *RAD1* | 0.028732 | -0.8934 | -0.89802 | -1.8355 | -0.59 |
| 17361 | *ENO3* | 0.8205 | -1.209 | -1.3743 | -1.6726 | -0.59 |
| 17362 | *FOXD1* | 0.11948 | -0.92711 | -0.95545 | -2.6805 | -0.59 |
| 17363 | *DHX33* | -0.39224 | -0.57513 | -0.79697 | -0.85612 | -0.59 |
| 17364 | *AKR1A1* | 0.07886 | -0.83756 | -1.0057 | -1.4837 | -0.59 |
| 17365 | *SLC25A4* | 0.47629 | -0.73546 | -1.5073 | -2.5226 | -0.59 |
| 17366 | *HSPB7* | -0.32153 | -0.38098 | -1.0644 | -2.2108 | -0.59 |
| 17367 | *N6AMT1* | -0.19704 | -0.76671 | -0.80332 | -1.7072 | -0.59 |
| 17368 | *FAM189A2* | 0.71357 | -1.0328 | -1.448 | -1.917 | -0.59 |
| 17369 | *GRIPAP1* | -0.39494 | -0.52343 | -0.85027 | -0.91283 | -0.59 |
| 17370 | *DCHS2* | -0.07094 | -0.46476 | -1.2338 | -1.7807 | -0.59 |
| 17371 | *B3GNT6* | -0.20663 | -0.68869 | -0.87418 | -1.0092 | -0.59 |
| 17372 | *KRT34* | -0.08474 | -0.78784 | -0.89741 | -1.356 | -0.59 |
| 17373 | *ZDHHC21* | 1.2305 | -1.0131 | -1.9899 | -2.3949 | -0.59 |
| 17374 | *CTAG2* | 0.52515 | -0.36084 | -1.9372 | -1.9493 | -0.59 |
| 17375 | *FBXL14* | 1.3733 | -0.93011 | -2.2169 | -2.4633 | -0.59 |
| 17376 | *LTC4S* | 0.63328 | -1.0987 | -1.3085 | -1.8154 | -0.59 |
| 17377 | *MSANTD3* | -0.26564 | -0.65053 | -0.8604 | -1.2684 | -0.59 |
| 17378 | *GART* | -0.09644 | -0.71849 | -0.96183 | -1.0564 | -0.59 |
| 17379 | *GNL1* | 0.045319 | -0.24273 | -1.5795 | -2.0021 | -0.59 |
| 17380 | *OR13C9* | 0.08921 | -0.83696 | -1.0297 | -1.2198 | -0.59 |
| 17381 | *PLCZ1* | 0.37148 | -0.59678 | -1.5524 | -1.6908 | -0.59 |

| | | | | | | |
|---|---|---|---|---|---|---|
| 17382 | *WRAP73* | 0.11212 | -0.92811 | -0.9619 | -1.7526 | -0.59 |
| 17383 | *ATP7A* | 0.91454 | -1.1612 | -1.5314 | -1.7253 | -0.59 |
| 17384 | *MED18* | -0.01973 | -0.84086 | -0.91766 | -1.29 | -0.59 |
| 17385 | *LRP4* | 0.44865 | -0.36336 | -1.864 | -2.5865 | -0.59 |
| 17386 | *ZNFX1* | -0.15481 | -0.50718 | -1.118 | -1.5244 | -0.59 |
| 17387 | *TCAF2* | -0.03757 | -0.49908 | -1.244 | -2.2413 | -0.59 |
| 17388 | *IARS* | 0.32523 | -0.31091 | -1.7956 | -2.0818 | -0.59 |
| 17389 | *TPGS1* | 0.054102 | -0.69333 | -1.1426 | -1.279 | -0.59 |
| 17390 | *FAM60A* | 0.68421 | -0.97499 | -1.4929 | -2.4331 | -0.59 |
| 17391 | *POLA2* | 0.093724 | -0.49007 | -1.3875 | -1.8115 | -0.59 |
| 17392 | *C11orf91* | 0.48187 | -1.1011 | -1.165 | -1.9945 | -0.59 |
| 17393 | *KCND3* | 1.2042 | -1.3128 | -1.6759 | -2.0407 | -0.59 |
| 17394 | *GPX4* | 0.50949 | -1.107 | -1.1872 | -1.5375 | -0.59 |
| 17395 | *FKBPL* | -0.07288 | -0.44056 | -1.2716 | -1.6941 | -0.60 |
| 17396 | *GPR68* | 0.17412 | -0.65877 | -1.3009 | -2.0283 | -0.60 |
| 17397 | *ZFAND2B* | 1.8564 | -1.7268 | -1.9175 | -2.2344 | -0.60 |
| 17398 | *CLEC4A* | 0.25407 | -0.53361 | -1.5095 | -2.6289 | -0.60 |
| 17399 | *ZBTB32* | 0.64194 | -0.48659 | -1.9446 | -2.1412 | -0.60 |
| 17400 | *AGO1* | 0.18967 | -0.44374 | -1.5375 | -1.6893 | -0.60 |
| 17401 | *KCNJ2* | 0.00553 | -0.47254 | -1.3255 | -1.4981 | -0.60 |
| 17402 | *WDR12* | 0.43389 | -0.65512 | -1.5719 | -2.5442 | -0.60 |
| 17403 | *RCOR1* | 0.66495 | -1.219 | -1.2404 | -1.7386 | -0.60 |
| 17404 | *AKAP4* | 0.035088 | -0.87318 | -0.95708 | -2.2593 | -0.60 |
| 17405 | *C16orf58* | 0.74244 | -1.2386 | -1.3009 | -2.1342 | -0.60 |
| 17406 | *GBF1* | -0.44056 | -0.54714 | -0.81101 | -1.3379 | -0.60 |
| 17407 | *BAZ1A* | -0.47099 | -0.62307 | -0.70546 | -1.0685 | -0.60 |
| 17408 | *PYDC2* | 0.4092 | -0.74086 | -1.4686 | -2.1662 | -0.60 |
| 17409 | *PTGER4* | 0.52775 | -0.62307 | -1.7056 | -2.0548 | -0.60 |
| 17410 | *ADAM30* | 0.035844 | -0.19053 | -1.6463 | -2.1687 | -0.60 |
| 17411 | *LRG1* | -0.26093 | -0.7443 | -0.79641 | -1.3604 | -0.60 |
| 17412 | *BTG3* | 0.14824 | -0.91433 | -1.0364 | -2.3222 | -0.60 |
| 17413 | *BCAP29* | 0.87867 | -1.2339 | -1.4473 | -1.4683 | -0.60 |
| 17414 | *CYP2S1* | 1.071 | -1.2772 | -1.5965 | -2.3689 | -0.60 |
| 17415 | *PARD6A* | -0.08812 | -0.83497 | -0.88159 | -1.7696 | -0.60 |
| 17416 | *LUC7L3* | 0.9719 | -0.94368 | -1.8333 | -1.8825 | -0.60 |
| 17417 | *KIAA0754* | 0.31254 | -0.64338 | -1.4745 | -2.2606 | -0.60 |
| 17418 | *GXYLT2* | 0.24831 | -0.34819 | -1.7073 | -2.0062 | -0.60 |
| 17419 | *DPP4* | 0.097548 | -0.74638 | -1.1587 | -2.7465 | -0.60 |
| 17420 | *AKIRIN2* | 1.1481 | -1.0737 | -1.882 | -2.6201 | -0.60 |
| 17421 | *C10orf10* | -0.25056 | -0.45194 | -1.1055 | -1.5663 | -0.60 |
| 17422 | *HIST2H2BE* | 0.45699 | -0.90606 | -1.359 | -1.4994 | -0.60 |
| 17423 | *AADACL4* | -0.35769 | -0.62588 | -0.82628 | -1.3209 | -0.60 |
| 17424 | *FZD9* | -0.00527 | -0.443 | -1.364 | -1.9326 | -0.60 |
| 17425 | *SNN* | -0.48085 | -0.49499 | -0.83773 | -1.7512 | -0.60 |
| 17426 | *FAM217B* | 0.14522 | -0.59662 | -1.3633 | -2.3615 | -0.60 |
| 17427 | *CENPI* | 0.43331 | -1.009 | -1.2393 | -1.5282 | -0.60 |

| | | | | | | |
|---|---|---|---|---|---|---|
| 17428 | *PAPPA* | 0.55941 | -1.122 | -1.2525 | -2.283 | -0.61 |
| 17429 | *CENPL* | 0.23212 | -0.38131 | -1.6663 | -2.1612 | -0.61 |
| 17430 | *FCHSD1* | 1.7484 | -1.6952 | -1.8689 | -2.2064 | -0.61 |
| 17431 | *RPEL1* | 0.50263 | -1.1282 | -1.1912 | -1.2291 | -0.61 |
| 17432 | *VWA3B* | 0.5181 | -0.96846 | -1.3666 | -1.4243 | -0.61 |
| 17433 | *ACTR2* | 0.28933 | -0.70708 | -1.3995 | -1.9605 | -0.61 |
| 17434 | *HOOK1* | 0.69476 | -1.067 | -1.4469 | -2.7427 | -0.61 |
| 17435 | *GYS1* | 0.88158 | -1.3427 | -1.3597 | -1.754 | -0.61 |
| 17436 | *SIGLEC9* | 0.14773 | -0.53584 | -1.433 | -1.8121 | -0.61 |
| 17437 | *PANO1* | 0.22012 | -1.0087 | -1.0335 | -1.4963 | -0.61 |
| 17438 | *SUSD3* | -0.06687 | -0.26443 | -1.491 | -2.8928 | -0.61 |
| 17439 | *ABCC8* | 1.4805 | -1.5965 | -1.7068 | -1.8885 | -0.61 |
| 17440 | *ZNF703* | 0.4727 | -0.61114 | -1.6866 | -1.8879 | -0.61 |
| 17441 | *POLH* | 0.14975 | -0.73728 | -1.2399 | -2.2292 | -0.61 |
| 17442 | *CPEB4* | 0.67749 | -0.99508 | -1.5099 | -1.5949 | -0.61 |
| 17443 | *ADAMTS3* | -0.13276 | -0.31126 | -1.3835 | -1.6604 | -0.61 |
| 17444 | *MROH7* | 0.038212 | -0.81007 | -1.0564 | -1.1117 | -0.61 |
| 17445 | *DCTN6* | 0.29113 | -0.3328 | -1.7879 | -2.075 | -0.61 |
| 17446 | *CEACAM5* | 0.59967 | -1.0934 | -1.3363 | -2.3949 | -0.61 |
| 17447 | *LOC101060389* | 0.019987 | -0.28457 | -1.567 | -1.7754 | -0.61 |
| 17448 | *GRIN2B* | -0.18133 | -0.7299 | -0.92078 | -1.7667 | -0.61 |
| 17449 | *RTBDN* | 0.79892 | -0.81536 | -1.8157 | -2.0171 | -0.61 |
| 17450 | *CCDC174* | -0.00831 | -0.71421 | -1.1097 | -3.2595 | -0.61 |
| 17451 | *NUCB1* | 0.22181 | -0.88142 | -1.1731 | -1.2079 | -0.61 |
| 17452 | *KRTAP10-12* | 0.29928 | -0.93621 | -1.1974 | -1.6218 | -0.61 |
| 17453 | *RGPD6* | 0.43414 | -0.77 | -1.4999 | -1.6952 | -0.61 |
| 17454 | *C5orf67* | -0.38796 | -0.43746 | -1.011 | -1.4883 | -0.61 |
| 17455 | *USP17L29* | 0.70433 | -1.2261 | -1.3162 | -1.5904 | -0.61 |
| 17456 | *DAPK2* | -0.03184 | -0.74638 | -1.0602 | -2.3452 | -0.61 |
| 17457 | *U2SURP* | 0.085671 | -0.29442 | -1.6302 | -1.6848 | -0.61 |
| 17458 | *NBPF6* | 0.4079 | -0.92828 | -1.3203 | -1.6523 | -0.61 |
| 17459 | *PTPRH* | 0.28293 | -0.78229 | -1.3427 | -1.635 | -0.61 |
| 17460 | *DUX4* | 0.57518 | -1.1189 | -1.2985 | -1.3393 | -0.61 |
| 17461 | *ZFYVE27* | 0.54494 | -1.0645 | -1.3237 | -1.6682 | -0.61 |
| 17462 | *ARHGAP15* | 0.10613 | -0.77323 | -1.177 | -1.8435 | -0.61 |
| 17463 | *TMA16* | -0.04439 | -0.48862 | -1.3127 | -1.5522 | -0.62 |
| 17464 | *AP1G2* | -0.53574 | -0.58404 | -0.7264 | -1.9746 | -0.62 |
| 17465 | *TSSK6* | 0.93459 | -0.98099 | -1.8002 | -1.8207 | -0.62 |
| 17466 | *ZMAT5* | 0.15724 | -0.57865 | -1.4277 | -1.5343 | -0.62 |
| 17467 | *TMOD4* | 0.00962 | -0.36957 | -1.4893 | -1.8382 | -0.62 |
| 17468 | *RHD* | 0.67615 | -1.049 | -1.4772 | -1.7149 | -0.62 |
| 17469 | *SLC6A6* | -0.06641 | -0.42373 | -1.3604 | -2.0538 | -0.62 |
| 17470 | *MFAP4* | 0.39585 | -0.60183 | -1.6447 | -2.1198 | -0.62 |
| 17471 | *MORN3* | 0.36371 | -1.0576 | -1.1571 | -2.0242 | -0.62 |
| 17472 | *TNFSF13* | 0.50078 | -0.5528 | -1.7991 | -3.1998 | -0.62 |
| 17473 | *GPSM1* | -0.24764 | -0.37824 | -1.2258 | -1.3172 | -0.62 |

| | | | | | | |
|---|---|---|---|---|---|---|
| 17474 | *DEFA1* | -0.12603 | -0.84996 | -0.87639 | -1.8044 | -0.62 |
| 17475 | *PMP22* | -0.26638 | -0.44353 | -1.1429 | -1.8995 | -0.62 |
| 17476 | *VARS2* | 0.14287 | -0.82069 | -1.175 | -1.3216 | -0.62 |
| 17477 | *BIRC2* | -0.37692 | -0.56485 | -0.91127 | -1.0172 | -0.62 |
| 17478 | *PSMD7* | -0.12416 | -0.71939 | -1.0095 | -1.4081 | -0.62 |
| 17479 | *PLEKHF1* | -0.49953 | -0.59903 | -0.75529 | -2.3811 | -0.62 |
| 17480 | *EMC2* | 0.75467 | -0.75287 | -1.8557 | -2.6431 | -0.62 |
| 17481 | *PKDCC* | 0.31744 | -0.60845 | -1.5635 | -3.2582 | -0.62 |
| 17482 | *HS6ST3* | 0.53339 | -0.99033 | -1.3977 | -1.8975 | -0.62 |
| 17483 | *SEPT12* | 0.58309 | -0.9481 | -1.4905 | -2.3017 | -0.62 |
| 17484 | *FAM153B* | -0.22572 | -0.41917 | -1.2114 | -1.8869 | -0.62 |
| 17485 | *FST* | 0.25115 | -1.027 | -1.0809 | -1.6268 | -0.62 |
| 17486 | *CRYAB* | -0.28188 | -0.60113 | -0.97477 | -1.2935 | -0.62 |
| 17487 | *ZNF506* | -0.48086 | -0.50227 | -0.87466 | -2.6024 | -0.62 |
| 17488 | *TTLL7* | 0.6183 | -1.0878 | -1.3892 | -1.7726 | -0.62 |
| 17489 | *FCF1* | 0.11223 | -0.96173 | -1.0095 | -1.2471 | -0.62 |
| 17490 | *CIB4* | -0.04665 | -0.50924 | -1.3033 | -2.3235 | -0.62 |
| 17491 | *NPY2R* | -0.02799 | -0.78229 | -1.049 | -2.0548 | -0.62 |
| 17492 | *SLC35B3* | 0.47395 | -1.0297 | -1.3039 | -1.8157 | -0.62 |
| 17493 | *KRT8* | 0.62261 | -0.5204 | -1.962 | -2.0181 | -0.62 |
| 17494 | *RTN3* | -0.31524 | -0.64851 | -0.89699 | -1.7718 | -0.62 |
| 17495 | *BIVM* | 0.17047 | -0.15375 | -1.8775 | -2.0423 | -0.62 |
| 17496 | *C20orf195* | 0.41231 | -1.1337 | -1.1395 | -1.7554 | -0.62 |
| 17497 | *RAC3* | 0.14867 | -1.001 | -1.0091 | -1.8686 | -0.62 |
| 17498 | *WNK4* | -0.14649 | -0.73546 | -0.97982 | -2.6762 | -0.62 |
| 17499 | *KLHDC7A* | -0.32121 | -0.50992 | -1.0315 | -1.7994 | -0.62 |
| 17500 | *FBN1* | 1.7454 | -1.6424 | -1.9668 | -2.7863 | -0.62 |
| 17501 | *CEP350* | 0.61051 | -1.1571 | -1.3189 | -2.0483 | -0.62 |
| 17502 | *NTF3* | 2.0855 | -1.8568 | -2.095 | -2.6201 | -0.62 |
| 17503 | *EFTUD2* | -0.15375 | -0.26892 | -1.4448 | -1.9997 | -0.62 |
| 17504 | *FAM109B* | -0.44353 | -0.60214 | -0.82256 | -1.1148 | -0.62 |
| 17505 | *CREG1* | 0.29824 | -0.71363 | -1.4533 | -1.499 | -0.62 |
| 17506 | *PCDHB14* | 1.1595 | -1.1133 | -1.9165 | -2.4454 | -0.62 |
| 17507 | *MGAT4C* | 0.093139 | -0.38574 | -1.578 | -1.6746 | -0.62 |
| 17508 | *UPF3B* | -0.41456 | -0.61972 | -0.83909 | -1.5947 | -0.62 |
| 17509 | *MFAP3L* | 1.0793 | -1.4248 | -1.5289 | -1.6143 | -0.62 |
| 17510 | *HNF1B* | -0.18871 | -0.64851 | -1.0383 | -3.4957 | -0.63 |
| 17511 | *PDIA5* | 2.0603 | -1.828 | -2.1085 | -2.4141 | -0.63 |
| 17512 | *AP1G1* | 0.2464 | -1.0019 | -1.1209 | -1.5482 | -0.63 |
| 17513 | *PTPN3* | 0.64633 | -1.0231 | -1.4997 | -1.8282 | -0.63 |
| 17514 | *POLR3B* | 0.35548 | -0.7458 | -1.4864 | -1.8523 | -0.63 |
| 17515 | *PAX1* | 0.2081 | -0.77853 | -1.3085 | -1.3704 | -0.63 |
| 17516 | *NBAS* | -0.40961 | -0.44353 | -1.0259 | -1.8724 | -0.63 |
| 17517 | *AMBP* | 1.419 | -1.4557 | -1.8443 | -1.9473 | -0.63 |
| 17518 | *NAPA* | -0.35757 | -0.75774 | -0.76743 | -2.6201 | -0.63 |
| 17519 | *ZNF773* | 0.25444 | -0.51805 | -1.62 | -2.4456 | -0.63 |

| | | | | | | |
|---|---|---|---|---|---|---|
| 17520 | *TDGF1* | -0.1024 | -0.62307 | -1.1595 | -1.4132 | -0.63 |
| 17521 | *TOMM5* | -0.03675 | -0.77963 | -1.0707 | -1.1244 | -0.63 |
| 17522 | *GPR137B* | 0.51181 | -0.61169 | -1.7874 | -2.796 | -0.63 |
| 17523 | *NMD3* | 0.91245 | -1.2258 | -1.5741 | -2.0454 | -0.63 |
| 17524 | *FAM187B* | 0.43825 | -1.1144 | -1.2114 | -1.8908 | -0.63 |
| 17525 | *ISM1* | -0.34376 | -0.65293 | -0.89178 | -1.2332 | -0.63 |
| 17526 | *VARS* | -0.38971 | -0.42015 | -1.0799 | -1.3128 | -0.63 |
| 17527 | *CENPC* | 0.53546 | -0.73865 | -1.6867 | -1.864 | -0.63 |
| 17528 | *MRPL41* | 0.37471 | -1.0797 | -1.1868 | -2.1424 | -0.63 |
| 17529 | *YWHAH* | -0.57899 | -0.62306 | -0.6901 | -1.6597 | -0.63 |
| 17530 | *DCAF17* | 0.018313 | -0.84147 | -1.071 | -1.3358 | -0.63 |
| 17531 | *ICOS* | 0.52794 | -0.9442 | -1.4789 | -1.6186 | -0.63 |
| 17532 | *ATF1* | 0.89324 | -1.2135 | -1.577 | -1.6339 | -0.63 |
| 17533 | *RBPJ* | 1.2675 | -1.463 | -1.7018 | -2.0146 | -0.63 |
| 17534 | *VWA5A* | 0.56381 | -1.1566 | -1.3055 | -2.5008 | -0.63 |
| 17535 | *BTBD18* | -0.05066 | -0.44198 | -1.4057 | -1.6403 | -0.63 |
| 17536 | *RHNO1* | 0.79291 | -1.2342 | -1.4574 | -2.1612 | -0.63 |
| 17537 | *ANKRD24* | -0.27857 | -0.62581 | -0.99489 | -1.5752 | -0.63 |
| 17538 | *TRIM48* | 0.45813 | -0.5613 | -1.7963 | -1.9997 | -0.63 |
| 17539 | *TCF21* | -0.509 | -0.6683 | -0.72237 | -1.8503 | -0.63 |
| 17540 | *ALX1* | -0.32699 | -0.35038 | -1.2234 | -1.2339 | -0.63 |
| 17541 | *TKT* | -0.11099 | -0.16055 | -1.6296 | -1.643 | -0.63 |
| 17542 | *RTP4* | -0.03565 | -0.60028 | -1.2655 | -1.4028 | -0.63 |
| 17543 | *ASNSD1* | -0.2613 | -0.66321 | -0.97792 | -1.5204 | -0.63 |
| 17544 | *RETSAT* | 0.48331 | -1.1354 | -1.251 | -1.8076 | -0.63 |
| 17545 | *PSMD3* | -0.06143 | -0.47158 | -1.3701 | -1.5981 | -0.63 |
| 17546 | *SLCO2B1* | 0.63483 | -1.0291 | -1.5099 | -2.3424 | -0.63 |
| 17547 | *ECT2L* | -0.08586 | -0.30677 | -1.5125 | -1.635 | -0.64 |
| 17548 | *EVX1* | -0.07612 | -0.72825 | -1.1008 | -3.1492 | -0.64 |
| 17549 | *SLC16A13* | -0.18241 | -0.53845 | -1.1851 | -1.2276 | -0.64 |
| 17550 | *TPMT* | -0.02943 | -0.12908 | -1.748 | -2.7171 | -0.64 |
| 17551 | *SP1* | -0.02342 | -0.05759 | -1.8256 | -2.2895 | -0.64 |
| 17552 | *SAE1* | 1.6604 | -1.7591 | -1.8081 | -1.9695 | -0.64 |
| 17553 | *TMX2* | 0.39507 | -0.79305 | -1.5113 | -1.5195 | -0.64 |
| 17554 | *NIPSNAP1* | -0.01323 | -0.14649 | -1.7507 | -4.2184 | -0.64 |
| 17555 | *GRAMD2* | -0.12784 | -0.86403 | -0.91913 | -1.5988 | -0.64 |
| 17556 | *HSPA4L* | -0.47964 | -0.49525 | -0.93661 | -1.6963 | -0.64 |
| 17557 | *SUMF1* | 0.92778 | -0.49926 | -2.3405 | -2.3618 | -0.64 |
| 17558 | *CDS2* | 0.54846 | -0.71987 | -1.7407 | -2.0553 | -0.64 |
| 17559 | *PPP1R7* | 0.4035 | -0.88702 | -1.4287 | -2.6376 | -0.64 |
| 17560 | *ZFYVE9* | 1.0588 | -0.8284 | -2.1429 | -2.8174 | -0.64 |
| 17561 | *RRAGC* | -0.42254 | -0.50524 | -0.98572 | -2.2599 | -0.64 |
| 17562 | *RBFOX2* | 0.17653 | -0.9612 | -1.13 | -1.1354 | -0.64 |
| 17563 | *BNIP1* | 0.2836 | -1.0871 | -1.1126 | -2.4058 | -0.64 |
| 17564 | *ETNK2* | -0.42373 | -0.68192 | -0.8108 | -2.0453 | -0.64 |
| 17565 | *ZNF416* | -0.21692 | -0.79573 | -0.90382 | -1.0506 | -0.64 |

| | | | | | | |
|---|---|---|---|---|---|---|
| 17566 | *TEKT2* | 0.038256 | -0.33139 | -1.624 | -3.454 | -0.64 |
| 17567 | *TNRC6A* | 0.81111 | -1.0704 | -1.6602 | -2.3768 | -0.64 |
| 17568 | *CHAF1A* | 0.47506 | -1.035 | -1.3606 | -1.707 | -0.64 |
| 17569 | *TNNI3K* | -0.05665 | -0.82256 | -1.0415 | -1.438 | -0.64 |
| 17570 | *MAPKAPK5* | 0.27065 | -1.0318 | -1.1613 | -1.2474 | -0.64 |
| 17571 | *PRPF38B* | 0.45421 | -1.1711 | -1.2058 | -1.8076 | -0.64 |
| 17572 | *PTPN20* | -0.59981 | -0.61164 | -0.71134 | -1.7316 | -0.64 |
| 17573 | *SMAD4* | 0.38881 | -0.91103 | -1.4017 | -1.7845 | -0.64 |
| 17574 | *HDAC7* | 0.11561 | -0.48352 | -1.5561 | -1.7766 | -0.64 |
| 17575 | *NOL9* | -0.09774 | -0.4102 | -1.4185 | -2.0071 | -0.64 |
| 17576 | *ZNF284* | -0.56947 | -0.66865 | -0.69146 | -1.8371 | -0.64 |
| 17577 | *EIF4EBP1* | 1.2385 | -1.4252 | -1.7438 | -3.0595 | -0.64 |
| 17578 | *CA12* | 0.61693 | -1.1382 | -1.4098 | -1.8508 | -0.64 |
| 17579 | *DYNLRB1* | -0.32433 | -0.52329 | -1.0863 | -1.6802 | -0.64 |
| 17580 | *TMEM45B* | 0.97477 | -1.1925 | -1.7185 | -2.6343 | -0.65 |
| 17581 | *AGXT2* | 0.20005 | -1.0614 | -1.0753 | -1.5568 | -0.65 |
| 17582 | *SETBP1* | 0.17909 | -0.75437 | -1.3637 | -1.7364 | -0.65 |
| 17583 | *SRRM1* | 0.61176 | -1.028 | -1.5235 | -1.5941 | -0.65 |
| 17584 | *CKAP5* | -0.29925 | -0.38527 | -1.2569 | -1.9073 | -0.65 |
| 17585 | *UXT* | -0.4433 | -0.4796 | -1.0204 | -1.8591 | -0.65 |
| 17586 | *DPH1* | 0.16922 | -0.69866 | -1.4141 | -2.9031 | -0.65 |
| 17587 | *SPAST* | -0.2122 | -0.43385 | -1.2985 | -1.6486 | -0.65 |
| 17588 | *KANSL2* | -0.22569 | -0.71561 | -1.0036 | -1.3433 | -0.65 |
| 17589 | *MED13* | -0.46752 | -0.47528 | -1.0041 | -1.9912 | -0.65 |
| 17590 | *POTEE* | -0.17606 | -0.72666 | -1.0467 | -1.683 | -0.65 |
| 17591 | *OR11H1* | 0.13776 | -1.0359 | -1.0529 | -2.4858 | -0.65 |
| 17592 | *CHRDL1* | -0.07432 | -0.8458 | -1.0315 | -1.1739 | -0.65 |
| 17593 | *TMEM116* | 1.102 | -1.4473 | -1.6075 | -1.677 | -0.65 |
| 17594 | *PRCD* | 0.40904 | -0.33179 | -2.0301 | -2.3915 | -0.65 |
| 17595 | *ARFGEF2* | 0.3113 | -0.84786 | -1.418 | -1.6597 | -0.65 |
| 17596 | *PIGU* | 0.18009 | -1.0413 | -1.0953 | -1.1692 | -0.65 |
| 17597 | *CABIN1* | 0.16326 | -0.18722 | -1.9326 | -2.9736 | -0.65 |
| 17598 | *STAT5A* | -0.42204 | -0.64268 | -0.89186 | -1.0951 | -0.65 |
| 17599 | *HIST2H2AA3* | 0.21343 | -0.99123 | -1.1793 | -2.5226 | -0.65 |
| 17600 | *DPPA2* | 0.57469 | -0.95708 | -1.5752 | -1.9168 | -0.65 |
| 17601 | *EXOSC2* | 0.031293 | -0.36242 | -1.6268 | -1.9954 | -0.65 |
| 17602 | *CYP27B1* | -0.17058 | -0.7819 | -1.0055 | -1.4832 | -0.65 |
| 17603 | *SPCS1* | 0.46022 | -0.81058 | -1.6086 | -3.0769 | -0.65 |
| 17604 | *WDR46* | 0.62769 | -1.2932 | -1.2935 | -1.5579 | -0.65 |
| 17605 | *TNFAIP1* | 0.13115 | -0.57824 | -1.5132 | -1.9367 | -0.65 |
| 17606 | *ZNF519* | -0.23722 | -0.63643 | -1.0871 | -2.3496 | -0.65 |
| 17607 | *SNRPF* | 0.21449 | -0.98211 | -1.1932 | -1.392 | -0.65 |
| 17608 | *ZNF566* | 0.07554 | -1.0178 | -1.0187 | -1.9757 | -0.65 |
| 17609 | *OR4B1* | -0.04449 | -0.90484 | -1.012 | -1.3957 | -0.65 |
| 17610 | *RPL23A* | 0.11134 | -0.91409 | -1.1591 | -2.1177 | -0.65 |
| 17611 | *TMEM178A* | 1.0275 | -1.1189 | -1.8707 | -3.1273 | -0.65 |

| | | | | | | |
|---|---|---|---|---|---|---|
| 17612 | *NCKAP5* | -0.54851 | -0.56485 | -0.8504 | -1.3226 | -0.65 |
| 17613 | *WBP4* | -0.0164 | -0.7443 | -1.205 | -1.3078 | -0.66 |
| 17614 | *USP17L26* | 0.62542 | -1.0229 | -1.57 | -1.8016 | -0.66 |
| 17615 | *SLCO3A1* | -0.17264 | -0.82524 | -0.96962 | -2.3154 | -0.66 |
| 17616 | *MRPL53* | -0.39532 | -0.5701 | -1.0025 | -1.316 | -0.66 |
| 17617 | *TSPAN11* | 0.28527 | -0.78951 | -1.4639 | -3.8314 | -0.66 |
| 17618 | *AGBL5* | -0.25915 | -0.56225 | -1.1472 | -2.4557 | -0.66 |
| 17619 | *TNK2* | 0.62609 | -0.90772 | -1.6877 | -2.3517 | -0.66 |
| 17620 | *SPDYE2B* | 0.98428 | -0.97477 | -1.9794 | -1.9811 | -0.66 |
| 17621 | *MKNK2* | -0.32262 | -0.53661 | -1.1124 | -1.8219 | -0.66 |
| 17622 | *ADGRG4* | 0.12433 | -0.20202 | -1.8946 | -2.7191 | -0.66 |
| 17623 | *FAHD2B* | -0.09468 | -0.77969 | -1.0995 | -1.5514 | -0.66 |
| 17624 | *MYRIP* | -0.31035 | -0.50979 | -1.1538 | -1.4407 | -0.66 |
| 17625 | *SOD2* | 1.739 | -1.7783 | -1.9351 | -2.0218 | -0.66 |
| 17626 | *DCTN4* | 0.060407 | -0.91968 | -1.1199 | -1.723 | -0.66 |
| 17627 | *ADAR* | 0.33513 | -1.0302 | -1.2845 | -1.5448 | -0.66 |
| 17628 | *TLE2* | -0.46749 | -0.51467 | -0.99892 | -1.2829 | -0.66 |
| 17629 | *THUMPD1* | -0.15953 | -0.75512 | -1.067 | -1.252 | -0.66 |
| 17630 | *RCL1* | -0.10527 | -0.75491 | -1.122 | -1.6116 | -0.66 |
| 17631 | *WNT10A* | 0.60803 | -1.2817 | -1.3092 | -1.7963 | -0.66 |
| 17632 | *CT45A10* | -0.61521 | -1.9624 | 0.59412 | -0.07993 | -0.66 |
| 17633 | *SRSF12* | 0.27752 | -0.91061 | -1.3505 | -1.7608 | -0.66 |
| 17634 | *HAUS7* | -0.5271 | -0.67417 | -0.7827 | -1.0229 | -0.66 |
| 17635 | *ETV3* | -0.13461 | -0.89699 | -0.95321 | -1.0884 | -0.66 |
| 17636 | *FGFBP3* | -0.21157 | -0.81501 | -0.95877 | -1.2208 | -0.66 |
| 17637 | *FGF10* | 0.51887 | -0.85236 | -1.6525 | -1.7829 | -0.66 |
| 17638 | *ZNF175* | -0.10032 | -0.80636 | -1.0801 | -2.1347 | -0.66 |
| 17639 | *ACTR3* | -0.38198 | -0.62969 | -0.97565 | -1.6208 | -0.66 |
| 17640 | *STXBP3* | 0.45719 | -1.0187 | -1.426 | -2.4557 | -0.66 |
| 17641 | *SCARF2* | -0.01352 | -0.23368 | -1.7407 | -2.0614 | -0.66 |
| 17642 | *GPR26* | 0.59211 | -0.9918 | -1.5883 | -2.2161 | -0.66 |
| 17643 | *GPR162* | 0.5443 | -1.0661 | -1.4688 | -1.6524 | -0.66 |
| 17644 | *VPS28* | 0.052674 | -0.75626 | -1.2884 | -3.2887 | -0.66 |
| 17645 | *RTN4* | -0.273 | -0.37282 | -1.3479 | -1.3604 | -0.66 |
| 17646 | *ZCCHC11* | 0.49507 | -1.1709 | -1.3198 | -1.7745 | -0.67 |
| 17647 | *HOPX* | -0.57484 | -0.59738 | -0.82524 | -0.93885 | -0.67 |
| 17648 | *FEV* | -0.0606 | -0.41502 | -1.5227 | -2.4633 | -0.67 |
| 17649 | *YWHAE* | 0.007297 | -0.53012 | -1.4775 | -1.8462 | -0.67 |
| 17650 | *GCAT* | -0.45821 | -0.50809 | -1.0351 | -1.2284 | -0.67 |
| 17651 | *ARHGEF7* | 0.66952 | -1.1191 | -1.5524 | -1.5738 | -0.67 |
| 17652 | *PREB* | 0.51657 | -1.1713 | -1.3473 | -2.239 | -0.67 |
| 17653 | *DDIAS* | -0.46765 | -0.67629 | -0.86027 | -1.3727 | -0.67 |
| 17654 | *GIPR* | -0.46756 | -0.76055 | -0.77895 | -1.3478 | -0.67 |
| 17655 | *CYB5R3* | 0.69306 | -0.76206 | -1.9395 | -2.2357 | -0.67 |
| 17656 | *PPP1R26* | 0.043556 | -0.08575 | -1.9664 | -2.3602 | -0.67 |
| 17657 | *IMP4* | -0.11619 | -0.50752 | -1.3852 | -1.4085 | -0.67 |

| | | | | | | |
|---|---|---|---|---|---|---|
| 17658 | *PCDHB3* | 0.87394 | -1.1158 | -1.7675 | -2.3696 | -0.67 |
| 17659 | *CHST10* | 0.56404 | -1.0158 | -1.5582 | -1.9833 | -0.67 |
| 17660 | *PSENEN* | 0.088102 | -0.5866 | -1.5116 | -2.3636 | -0.67 |
| 17661 | *ALKBH7* | 0.84949 | -1.2159 | -1.6447 | -1.6974 | -0.67 |
| 17662 | *CAPN10* | 0.1357 | -0.68917 | -1.459 | -1.5712 | -0.67 |
| 17663 | *BCAS4* | 1.065 | -1.525 | -1.5525 | -1.7219 | -0.67 |
| 17664 | *SRSF11* | -0.50878 | -0.58315 | -0.9216 | -1.5212 | -0.67 |
| 17665 | *MAGEF1* | 1.2884 | -1.4788 | -1.8253 | -2.278 | -0.67 |
| 17666 | *SLC7A6OS* | 0.055414 | -0.61429 | -1.4577 | -2.0623 | -0.67 |
| 17667 | *PCDHGB5* | -0.09789 | -0.83101 | -1.0878 | -1.3617 | -0.67 |
| 17668 | *OR52K2* | -0.01351 | -0.68422 | -1.3191 | -1.7743 | -0.67 |
| 17669 | *POU2AF1* | 0.083863 | -0.36549 | -1.7364 | -2.1334 | -0.67 |
| 17670 | *SLC10A3* | 0.044461 | -0.73389 | -1.3293 | -1.9321 | -0.67 |
| 17671 | *SAFB2* | -0.27933 | -0.44687 | -1.2936 | -1.8212 | -0.67 |
| 17672 | *KLHDC8A* | -0.41079 | -0.54544 | -1.0644 | -1.2324 | -0.67 |
| 17673 | *JRKL* | -0.02205 | -0.85018 | -1.1487 | -2.2607 | -0.67 |
| 17674 | *LONRF1* | 0.80812 | -1.0648 | -1.7652 | -2.0851 | -0.67 |
| 17675 | *MRPL43* | 0.28332 | -0.76379 | -1.5415 | -2.2804 | -0.67 |
| 17676 | *PI16* | 0.32753 | -0.85194 | -1.4998 | -2.0277 | -0.67 |
| 17677 | *TDP1* | 0.99469 | -1.3665 | -1.6524 | -2.3036 | -0.67 |
| 17678 | *PASK* | -0.28248 | -0.68386 | -1.0589 | -1.0954 | -0.68 |
| 17679 | *PRKCI* | 0.35076 | -0.78097 | -1.5954 | -1.6453 | -0.68 |
| 17680 | *IL36RN* | -0.23565 | -0.74365 | -1.047 | -1.0511 | -0.68 |
| 17681 | *DAGLA* | 1.04 | -1.1309 | -1.9362 | -2.0337 | -0.68 |
| 17682 | *TINF2* | 0.54633 | -1.0283 | -1.5458 | -2.0153 | -0.68 |
| 17683 | *PAK1IP1* | 0.55829 | -1.1043 | -1.4824 | -1.7413 | -0.68 |
| 17684 | *MEIS2* | 0.005184 | -0.97595 | -1.0582 | -1.438 | -0.68 |
| 17685 | *MOSPD3* | -0.38826 | -0.64149 | -0.9999 | -1.0344 | -0.68 |
| 17686 | *CATSPER1* | -0.53856 | -0.68825 | -0.80295 | -1.1585 | -0.68 |
| 17687 | *PSMD2* | 0.68144 | -1.309 | -1.4024 | -1.5375 | -0.68 |
| 17688 | *IRF2BP2* | -0.42609 | -0.48982 | -1.1149 | -2.0134 | -0.68 |
| 17689 | *ADSL* | -0.16263 | -0.74192 | -1.1263 | -1.5898 | -0.68 |
| 17690 | *TFAP2C* | -0.38319 | -0.67842 | -0.96993 | -1.754 | -0.68 |
| 17691 | *NDUFA1* | -0.30153 | -0.81017 | -0.92131 | -1.438 | -0.68 |
| 17692 | *RGL1* | 0.64372 | -1.3231 | -1.3542 | -1.9997 | -0.68 |
| 17693 | *OSGIN1* | -0.27599 | -0.65145 | -1.1075 | -1.3728 | -0.68 |
| 17694 | *SPN* | 1.1706 | -1.4789 | -1.7268 | -1.8523 | -0.68 |
| 17695 | *TBCD* | 0.33017 | -1.154 | -1.2135 | -1.4299 | -0.68 |
| 17696 | *SSRP1* | -0.41794 | -0.57261 | -1.0483 | -1.7496 | -0.68 |
| 17697 | *EID3* | -0.51467 | -0.61796 | -0.90993 | -2.1018 | -0.68 |
| 17698 | *MBD5* | 0.20565 | -0.60289 | -1.6463 | -1.72 | -0.68 |
| 17699 | *GPN1* | -0.13025 | -0.45552 | -1.4588 | -1.9837 | -0.68 |
| 17700 | *NONO* | -0.22082 | -0.74743 | -1.078 | -1.7266 | -0.68 |
| 17701 | *PDCD1* | 1.3111 | -1.0559 | -2.3017 | -3.6868 | -0.68 |
| 17702 | *DSCC1* | -0.32677 | -0.78503 | -0.93489 | -1.9821 | -0.68 |
| 17703 | *AARSD1* | 0.35094 | -1.0741 | -1.3237 | -2.4518 | -0.68 |

| | | | | | | |
|---|---|---|---|---|---|---|
| 17704 | *LOC388282* | -0.00139 | -0.36297 | -1.6847 | -1.7049 | -0.68 |
| 17705 | *PCID2* | -0.01323 | -0.29546 | -1.7412 | -2.9031 | -0.68 |
| 17706 | *ABHD10* | -0.20774 | -0.83961 | -1.0049 | -1.8779 | -0.68 |
| 17707 | *GLDN* | 0.63841 | -1.3165 | -1.3756 | -1.7874 | -0.68 |
| 17708 | *BUD31* | 0.45103 | -1.043 | -1.462 | -1.8629 | -0.68 |
| 17709 | *RD3* | 0.05378 | -0.83402 | -1.274 | -1.3478 | -0.68 |
| 17710 | *DARS2* | 0.00553 | -0.9346 | -1.1303 | -1.5364 | -0.69 |
| 17711 | *NME6* | 0.099654 | -0.94597 | -1.2135 | -1.2411 | -0.69 |
| 17712 | *ZNF672* | -0.34751 | -0.66164 | -1.0517 | -2.8863 | -0.69 |
| 17713 | *RRAGB* | 0.22609 | -0.74013 | -1.5485 | -1.9375 | -0.69 |
| 17714 | *KAT2A* | 0.5884 | -1.1745 | -1.4773 | -1.6617 | -0.69 |
| 17715 | *CPNE5* | 0.57489 | -1.1463 | -1.4929 | -1.8963 | -0.69 |
| 17716 | *VPS39* | -0.09334 | -0.87997 | -1.091 | -1.2719 | -0.69 |
| 17717 | *CLDN5* | -0.43276 | -0.6981 | -0.93407 | -1.0707 | -0.69 |
| 17718 | *SPOP* | 0.012881 | -0.82208 | -1.2571 | -1.6934 | -0.69 |
| 17719 | *TRMT61B* | 1.3287 | -1.5806 | -1.8157 | -2.7509 | -0.69 |
| 17720 | *SPNS2* | 0.78934 | -1.0838 | -1.7745 | -1.9523 | -0.69 |
| 17721 | *NPAT* | -0.61358 | -0.65714 | -0.79832 | -2.6201 | -0.69 |
| 17722 | *PFAS* | -0.02738 | -0.88008 | -1.162 | -1.3209 | -0.69 |
| 17723 | *LPAR1* | -0.33198 | -0.79641 | -0.94184 | -1.2032 | -0.69 |
| 17724 | *IGLL1* | 0.42349 | -1.238 | -1.2576 | -1.5897 | -0.69 |
| 17725 | *YPEL1* | 0.76534 | -1.3948 | -1.4428 | -1.9065 | -0.69 |
| 17726 | *C3orf62* | -0.44353 | -0.60604 | -1.0239 | -1.5598 | -0.69 |
| 17727 | *ZNF823* | 0.40742 | -1.0185 | -1.4625 | -1.4873 | -0.69 |
| 17728 | *ZC3H11A* | -0.13063 | -0.76965 | -1.1752 | -1.3199 | -0.69 |
| 17729 | *PRPF4B* | -0.20663 | -0.43634 | -1.433 | -1.5502 | -0.69 |
| 17730 | *SYT8* | 0.27528 | -0.90663 | -1.4452 | -1.4852 | -0.69 |
| 17731 | *PTPN21* | 0.88094 | -1.1825 | -1.7766 | -3.8028 | -0.69 |
| 17732 | *IL1R1* | 0.32657 | -0.71464 | -1.6907 | -2.1983 | -0.69 |
| 17733 | *TMEM156* | -0.1097 | -0.67809 | -1.2915 | -1.6469 | -0.69 |
| 17734 | *LYZL6* | -0.01401 | -0.80176 | -1.2641 | -1.3616 | -0.69 |
| 17735 | *ALG2* | -0.35716 | -0.44353 | -1.2792 | -1.3818 | -0.69 |
| 17736 | *H2AFB3* | 0.14964 | -0.9512 | -1.2792 | -1.4692 | -0.69 |
| 17737 | *KARS* | -0.43313 | -0.71353 | -0.93901 | -1.2455 | -0.70 |
| 17738 | *OR2A14* | 0.03977 | -0.97784 | -1.1479 | -2.1577 | -0.70 |
| 17739 | *INSM2* | 0.17576 | -1.1212 | -1.1421 | -1.2792 | -0.70 |
| 17740 | *TRAPPC11* | 0.51874 | -1.2659 | -1.3406 | -2.4917 | -0.70 |
| 17741 | *GTF2IRD2* | -0.48233 | -0.66655 | -0.93889 | -1.8961 | -0.70 |
| 17742 | *CCDC138* | 1.119 | -1.3978 | -1.8095 | -2.4873 | -0.70 |
| 17743 | *NBPF15* | -0.17483 | -0.89204 | -1.0221 | -2.6803 | -0.70 |
| 17744 | *MTOR* | 0.77965 | -1.3279 | -1.5408 | -2.0704 | -0.70 |
| 17745 | *SLC25A14* | 1.3127 | -1.5401 | -1.8636 | -1.9748 | -0.70 |
| 17746 | *ZNF806* | -0.05126 | -0.94052 | -1.1027 | -1.1357 | -0.70 |
| 17747 | *PNPLA2* | 0.76538 | -1.4141 | -1.4469 | -1.9946 | -0.70 |
| 17748 | *RRM2* | 0.46648 | -1.2169 | -1.3453 | -2.7566 | -0.70 |
| 17749 | *C9orf50* | 0.82918 | -1.3805 | -1.5448 | -3.6181 | -0.70 |

| | | | | | | |
|---|---|---|---|---|---|---|
| 17750 | *SAMD9* | 0.27509 | -0.63325 | -1.738 | -2.4815 | -0.70 |
| 17751 | *RPS4X* | -0.16209 | -0.8656 | -1.0695 | -1.2186 | -0.70 |
| 17752 | *S100A7* | -0.42075 | -0.76606 | -0.91296 | -1.8157 | -0.70 |
| 17753 | *APC2* | 0.1689 | -0.28012 | -1.9894 | -2.8718 | -0.70 |
| 17754 | *UBL3* | 0.068008 | -0.87719 | -1.2915 | -2.105 | -0.70 |
| 17755 | *AQP4* | -0.11845 | -0.66136 | -1.3226 | -1.3492 | -0.70 |
| 17756 | *HCN1* | 0.004551 | -0.22378 | -1.8836 | -2.7042 | -0.70 |
| 17757 | *TVP23C* | 0.8875 | -1.3864 | -1.6047 | -2.0384 | -0.70 |
| 17758 | *MAP2K6* | -0.07138 | -0.94441 | -1.0884 | -1.5576 | -0.70 |
| 17759 | *TLCD1* | 0.43648 | -0.91343 | -1.6288 | -1.6488 | -0.70 |
| 17760 | *HLF* | 0.005894 | -0.5605 | -1.5512 | -1.8941 | -0.70 |
| 17761 | *CATSPERD* | -0.02018 | -0.12433 | -1.962 | -2.9323 | -0.70 |
| 17762 | *RAB9A* | 0.36422 | -1.0842 | -1.3878 | -4.7747 | -0.70 |
| 17763 | *FERMT2* | 0.25142 | -0.47367 | -1.8871 | -2.5495 | -0.70 |
| 17764 | *GAD1* | 0.48074 | -1.2249 | -1.3663 | -2.1965 | -0.70 |
| 17765 | *PIK3CD* | 0.39116 | -1.2132 | -1.2895 | -1.3627 | -0.70 |
| 17766 | *SRCAP* | 0.27291 | -0.43226 | -1.9548 | -2.9068 | -0.70 |
| 17767 | *MPP5* | -0.16337 | -0.58289 | -1.3701 | -2.2533 | -0.71 |
| 17768 | *AKAP3* | -0.10103 | -0.23166 | -1.7845 | -1.9622 | -0.71 |
| 17769 | *ELOF1* | -0.13429 | -0.60353 | -1.3808 | -2.4901 | -0.71 |
| 17770 | *C7orf73* | 0.002715 | -0.8147 | -1.3071 | -1.6525 | -0.71 |
| 17771 | *KLHL33* | 0.61874 | -0.99958 | -1.7386 | -1.7804 | -0.71 |
| 17772 | *HTR2A* | -0.29327 | -0.43043 | -1.397 | -1.7688 | -0.71 |
| 17773 | *RILPL2* | -0.01366 | -0.256 | -1.8516 | -1.9089 | -0.71 |
| 17774 | *CHEK1* | 0.1688 | -0.39556 | -1.8946 | -2.0277 | -0.71 |
| 17775 | *SMC4* | 0.30645 | -0.78651 | -1.6457 | -1.9269 | -0.71 |
| 17776 | *ATP6V0C* | 0.044225 | -0.70463 | -1.4655 | -1.5263 | -0.71 |
| 17777 | *PPWD1* | -0.04057 | -0.53971 | -1.548 | -2.7689 | -0.71 |
| 17778 | *PTCHD4* | 0.43931 | -0.79035 | -1.7782 | -2.9706 | -0.71 |
| 17779 | *NEDD1* | -0.43077 | -0.5613 | -1.1384 | -2.0541 | -0.71 |
| 17780 | *POLDIP2* | 0.7052 | -1.278 | -1.5579 | -2.0226 | -0.71 |
| 17781 | *FAM172A* | 0.29569 | -1.1214 | -1.3057 | -2.1477 | -0.71 |
| 17782 | *CREBBP* | -0.51722 | -0.68612 | -0.92948 | -2.0407 | -0.71 |
| 17783 | *PDPN* | -0.19735 | -0.88272 | -1.0528 | -3.1751 | -0.71 |
| 17784 | *CDK12* | -0.50251 | -0.69407 | -0.93725 | -2.2856 | -0.71 |
| 17785 | *BBS1* | 1.624 | -1.7935 | -1.9673 | -2.7191 | -0.71 |
| 17786 | *SCD* | 0.3214 | -1.0315 | -1.4269 | -1.9678 | -0.71 |
| 17787 | *OR8B2* | 0.34105 | -0.99838 | -1.4798 | -1.5577 | -0.71 |
| 17788 | *WWC3* | -0.3205 | -0.49449 | -1.3246 | -3.087 | -0.71 |
| 17789 | *OVCA2* | -0.13111 | -0.80978 | -1.1991 | -1.6386 | -0.71 |
| 17790 | *TBL3* | 0.23609 | -1.0425 | -1.3346 | -1.4683 | -0.71 |
| 17791 | *UBE2D3* | 0.46439 | -0.69407 | -1.9116 | -1.9967 | -0.71 |
| 17792 | *NOL10* | 1.7071 | -1.1833 | -2.6675 | -3.0769 | -0.71 |
| 17793 | *CDK2* | 0.7963 | -1.1888 | -1.7526 | -1.9975 | -0.72 |
| 17794 | *METAP1* | -0.15919 | -0.81517 | -1.1727 | -2.033 | -0.72 |
| 17795 | *VENTX* | 1.5262 | -1.2151 | -2.4595 | -2.8316 | -0.72 |

| | | | | | | |
|---|---|---|---|---|---|---|
| 17796 | *NIPBL* | -0.40534 | -0.64943 | -1.0945 | -1.4206 | -0.72 |
| 17797 | *DARS* | -0.16607 | -0.35548 | -1.6324 | -2.217 | -0.72 |
| 17798 | *PELI3* | -0.21809 | -0.84603 | -1.0914 | -1.6442 | -0.72 |
| 17799 | *TUBA1A* | 0.41564 | -1.0048 | -1.5672 | -2.2955 | -0.72 |
| 17800 | *KCNQ1* | 1.1159 | -1.4625 | -1.8101 | -2.7149 | -0.72 |
| 17801 | *SLIT1* | -0.20227 | -0.70463 | -1.25 | -1.367 | -0.72 |
| 17802 | *NCK2* | 0.010955 | -1.0033 | -1.165 | -2.5535 | -0.72 |
| 17803 | *GAGE2B* | 0.1181 | -0.9857 | -1.2912 | -1.5797 | -0.72 |
| 17804 | *SERPINA6* | -0.47835 | -0.66604 | -1.0149 | -1.1759 | -0.72 |
| 17805 | *GTF3A* | -0.31294 | -0.87067 | -0.97595 | -2.2403 | -0.72 |
| 17806 | *LRRC43* | 0.49269 | -0.85943 | -1.7935 | -1.8714 | -0.72 |
| 17807 | *RPN2* | -0.23368 | -0.83272 | -1.0942 | -1.7338 | -0.72 |
| 17808 | *FAM181B* | -0.43471 | -0.77284 | -0.95321 | -1.1016 | -0.72 |
| 17809 | *ATP6V0A1* | 0.67727 | -1.3542 | -1.4842 | -1.7215 | -0.72 |
| 17810 | *C14orf37* | 1.1403 | -1.3823 | -1.9196 | -2.9909 | -0.72 |
| 17811 | *ACAD10* | -0.14244 | -0.44137 | -1.5781 | -1.9772 | -0.72 |
| 17812 | *ERVFRD-1* | -0.36728 | -0.46741 | -1.3288 | -2.9931 | -0.72 |
| 17813 | *PRR23D2* | 0.67588 | -0.29209 | -2.5477 | -3.064 | -0.72 |
| 17814 | *AXDND1* | -0.17238 | -0.47782 | -1.5145 | -1.5707 | -0.72 |
| 17815 | *CNOT3* | -0.3328 | -0.82757 | -1.0048 | -1.4125 | -0.72 |
| 17816 | *CENPO* | -0.01255 | -0.62453 | -1.5284 | -1.6027 | -0.72 |
| 17817 | *HIST1H2AJ* | 0.6168 | -1.1126 | -1.6707 | -1.9623 | -0.72 |
| 17818 | *RPS15A* | 0.010089 | -1.0718 | -1.1048 | -2.1419 | -0.72 |
| 17819 | *RPIA* | 0.06743 | -0.66462 | -1.57 | -1.7068 | -0.72 |
| 17820 | *SPP2* | -0.7004 | -0.71032 | -0.75778 | -2.1407 | -0.72 |
| 17821 | *SP140L* | -0.20302 | -0.61675 | -1.3506 | -1.3542 | -0.72 |
| 17822 | *ATP2B1* | 0.14844 | -0.88124 | -1.439 | -1.5018 | -0.72 |
| 17823 | *CT45A9* | -0.41486 | -0.49727 | -1.26 | -1.6268 | -0.72 |
| 17824 | *HRH4* | -0.53304 | -0.81485 | -0.82514 | -2.1496 | -0.72 |
| 17825 | *CA5A* | 0.65432 | -1.3403 | -1.4886 | -1.8355 | -0.72 |
| 17826 | *SCAI* | 0.10845 | -0.71988 | -1.5635 | -1.6893 | -0.72 |
| 17827 | *OR5H14* | 0.38683 | -0.89957 | -1.6626 | -5.1372 | -0.73 |
| 17828 | *PRAMEF10* | -0.52649 | -0.72372 | -0.9285 | -1.509 | -0.73 |
| 17829 | *RGPD2* | 0.59223 | -1.0068 | -1.7662 | -1.9244 | -0.73 |
| 17830 | *NOB1* | -0.46024 | -0.53133 | -1.1893 | -2.0066 | -0.73 |
| 17831 | *TEKT3* | -0.46741 | -0.56634 | -1.1472 | -1.5182 | -0.73 |
| 17832 | *NISCH* | 0.3523 | -0.72085 | -1.8126 | -2.5878 | -0.73 |
| 17833 | *GMDS* | -0.59061 | -0.75487 | -0.83623 | -1.9774 | -0.73 |
| 17834 | *JRK* | 0.041958 | -0.97552 | -1.2498 | -1.531 | -0.73 |
| 17835 | *KDELR3* | 0.73014 | -1.388 | -1.526 | -1.6787 | -0.73 |
| 17836 | *TP53TG3D* | -0.05492 | -0.35972 | -1.7702 | -1.9984 | -0.73 |
| 17837 | *TCTEX1D2* | -0.26714 | -0.95149 | -0.96718 | -1.7018 | -0.73 |
| 17838 | *LY6D* | -0.00832 | -0.9311 | -1.2474 | -2.6582 | -0.73 |
| 17839 | *VWA5B2* | -0.46205 | -0.72237 | -1.0044 | -1.4368 | -0.73 |
| 17840 | *DRC1* | -0.37287 | -0.78362 | -1.0328 | -1.5663 | -0.73 |
| 17841 | *POLRMT* | -0.37925 | -0.44353 | -1.3679 | -2.0071 | -0.73 |

| | | | | | | |
|---|---|---|---|---|---|---|
| 17842 | *RTN4IP1* | 0.37578 | -0.79453 | -1.7753 | -3.096 | -0.73 |
| 17843 | *KCTD15* | -0.23196 | -0.46953 | -1.4941 | -2.2165 | -0.73 |
| 17844 | *ELOVL7* | 0.15149 | -1.1214 | -1.226 | -3.2168 | -0.73 |
| 17845 | *PPP2CB* | 0.50292 | -1.2242 | -1.4791 | -1.5922 | -0.73 |
| 17846 | *PCGF1* | 0.14884 | -1.0168 | -1.3331 | -1.8528 | -0.73 |
| 17847 | *IMMT* | -0.00239 | -1.0717 | -1.1279 | -1.9054 | -0.73 |
| 17848 | *MAFA* | -0.57001 | -0.69972 | -0.93368 | -0.99694 | -0.73 |
| 17849 | *PRSS42* | 0.16906 | -0.79726 | -1.5773 | -1.8702 | -0.74 |
| 17850 | *VPS13D* | -0.45649 | -0.81152 | -0.93895 | -2.0323 | -0.74 |
| 17851 | *ARMC10* | 0.52646 | -1.1564 | -1.5773 | -2.7931 | -0.74 |
| 17852 | *EPPK1* | 0.2539 | -0.87898 | -1.583 | -2.1685 | -0.74 |
| 17853 | *LOC101927572* | 0.047789 | -0.90532 | -1.3526 | -1.4415 | -0.74 |
| 17854 | *C20orf96* | 0.006278 | -0.92375 | -1.2935 | -1.2975 | -0.74 |
| 17855 | *LETMD1* | -0.52495 | -0.75511 | -0.93107 | -1.9622 | -0.74 |
| 17856 | *THBS4* | -0.35899 | -0.35986 | -1.4951 | -2.2473 | -0.74 |
| 17857 | *DTX1* | -0.33758 | -0.91236 | -0.96422 | -1.2211 | -0.74 |
| 17858 | *POMP* | -0.20916 | -0.85747 | -1.1487 | -1.575 | -0.74 |
| 17859 | *ITGA2* | 0.29991 | -0.98572 | -1.5302 | -2.0629 | -0.74 |
| 17860 | *ATPAF2* | -0.20771 | -0.83163 | -1.1772 | -1.5583 | -0.74 |
| 17861 | *CCT7* | -0.1283 | -1.0189 | -1.0696 | -1.2585 | -0.74 |
| 17862 | *RPL34* | 0.186 | -1.0467 | -1.3565 | -2.796 | -0.74 |
| 17863 | *OR5K4* | 0.50691 | -1.3288 | -1.3961 | -2.1155 | -0.74 |
| 17864 | *EGFLAM* | -0.3817 | -0.73364 | -1.1027 | -1.4132 | -0.74 |
| 17865 | *CPLX1* | -0.30576 | -0.81846 | -1.0945 | -1.9965 | -0.74 |
| 17866 | *UNC13C* | 0.728 | -0.90993 | -2.0368 | -2.1926 | -0.74 |
| 17867 | *AGK* | 0.56295 | -1.1108 | -1.6709 | -1.7911 | -0.74 |
| 17868 | *PCDHGA3* | -0.23447 | -0.85199 | -1.1332 | -3.0502 | -0.74 |
| 17869 | *OXCT2* | -0.30863 | -0.60262 | -1.309 | -1.6251 | -0.74 |
| 17870 | *HOXA13* | -0.26711 | -0.62469 | -1.3291 | -1.5633 | -0.74 |
| 17871 | *CDH4* | -0.09723 | -0.82757 | -1.2995 | -1.5082 | -0.74 |
| 17872 | *C6orf58* | -0.05684 | -0.91096 | -1.257 | -2.9354 | -0.74 |
| 17873 | *ZNF275* | -0.4072 | -0.69712 | -1.1214 | -1.6993 | -0.74 |
| 17874 | *SIK1* | 0.3463 | -1.1169 | -1.4566 | -1.9794 | -0.74 |
| 17875 | *KCNG3* | 0.085425 | -0.84523 | -1.4683 | -1.831 | -0.74 |
| 17876 | *PCYT1A* | -0.26055 | -0.80226 | -1.1656 | -1.613 | -0.74 |
| 17877 | *MTMR2* | -0.24269 | -0.73301 | -1.2531 | -2.0181 | -0.74 |
| 17878 | *MON1A* | 0.98984 | -1.3023 | -1.9168 | -2.7931 | -0.74 |
| 17879 | *CLEC4D* | -0.13341 | -0.71063 | -1.386 | -1.8446 | -0.74 |
| 17880 | *MEFV* | -0.02502 | -0.80146 | -1.4043 | -1.8238 | -0.74 |
| 17881 | *CCNYL1* | -0.26267 | -0.76635 | -1.2024 | -1.3593 | -0.74 |
| 17882 | *BLNK* | -0.04729 | -0.75452 | -1.4306 | -1.5026 | -0.74 |
| 17883 | *LGALS9B* | 0.010873 | -0.34553 | -1.8979 | -2.1648 | -0.74 |
| 17884 | *RNASE9* | -0.15715 | -0.83961 | -1.2358 | -1.9558 | -0.74 |
| 17885 | *AMOTL2* | -0.21857 | -0.22422 | -1.7906 | -2.2809 | -0.74 |
| 17886 | *CCT8* | -0.08664 | -0.93114 | -1.2158 | -2.0904 | -0.74 |
| 17887 | *SF3B3* | 0.20985 | -0.59622 | -1.8473 | -2.1445 | -0.74 |

| | | | | | | |
|---|---|---|---|---|---|---|
| 17888 | *RRP9* | -0.26161 | -0.61892 | -1.3542 | -1.6184 | -0.74 |
| 17889 | *SDE2* | -0.12659 | -0.44884 | -1.6594 | -2.262 | -0.74 |
| 17890 | *CBWD3* | -0.44353 | -0.52495 | -1.2677 | -1.3617 | -0.75 |
| 17891 | *RITA1* | -0.10536 | -0.67555 | -1.4554 | -3.0378 | -0.75 |
| 17892 | *NOMO2* | -0.7052 | -0.75437 | -0.77791 | -0.78566 | -0.75 |
| 17893 | *ATAD5* | -0.03697 | -0.92926 | -1.2719 | -1.5073 | -0.75 |
| 17894 | *C6orf141* | -0.37054 | -0.81501 | -1.0529 | -1.2052 | -0.75 |
| 17895 | *TIPRL* | -0.20242 | -0.99059 | -1.046 | -1.677 | -0.75 |
| 17896 | *FAM20B* | 0.70861 | -1.1989 | -1.7488 | -2.8388 | -0.75 |
| 17897 | *GPS1* | 0.74794 | -1.151 | -1.8366 | -2.7863 | -0.75 |
| 17898 | *MYT1* | 1.7433 | -1.9397 | -2.0436 | -2.5571 | -0.75 |
| 17899 | *NELFE* | -0.2347 | -0.44574 | -1.5598 | -1.8869 | -0.75 |
| 17900 | *ELP4* | 0.40947 | -1.2357 | -1.4142 | -1.4375 | -0.75 |
| 17901 | *RAB7A* | 1.0441 | -1.5722 | -1.7134 | -1.7266 | -0.75 |
| 17902 | *CYHR1* | 0.14445 | -0.86384 | -1.5227 | -1.5471 | -0.75 |
| 17903 | *RBMXL3* | 0.22516 | -1.1796 | -1.2901 | -1.8305 | -0.75 |
| 17904 | *C4BPB* | 0.023338 | -0.83708 | -1.4311 | -2.2589 | -0.75 |
| 17905 | *SPIN3* | -0.27453 | -0.98239 | -0.98882 | -1.9794 | -0.75 |
| 17906 | *ACOX3* | -0.00066 | -0.50394 | -1.7413 | -2.3164 | -0.75 |
| 17907 | *PELO* | -0.21385 | -0.93725 | -1.0954 | -1.8453 | -0.75 |
| 17908 | *THAP8* | 0.17631 | -0.84308 | -1.583 | -2.0697 | -0.75 |
| 17909 | *ZNF720* | -0.13871 | -0.83739 | -1.274 | -2.8388 | -0.75 |
| 17910 | *ARL15* | -0.26125 | -0.57573 | -1.4132 | -1.5795 | -0.75 |
| 17911 | *OAZ1* | 0.47914 | -1.2163 | -1.5138 | -1.6599 | -0.75 |
| 17912 | *POU3F2* | 0.14471 | -1.1763 | -1.2195 | -1.4741 | -0.75 |
| 17913 | *GEMIN4* | -0.16774 | -0.63472 | -1.4487 | -1.7363 | -0.75 |
| 17914 | *KIF3B* | 0.050178 | -0.49846 | -1.8031 | -2.0116 | -0.75 |
| 17915 | *ZNF512B* | -0.14097 | -0.84419 | -1.2666 | -1.5354 | -0.75 |
| 17916 | *BEST2* | -0.13963 | -0.73673 | -1.3769 | -1.5552 | -0.75 |
| 17917 | *RAD9A* | -0.37417 | -0.79481 | -1.0852 | -1.7285 | -0.75 |
| 17918 | *APLNR* | 0.34839 | -0.77159 | -1.831 | -2.9977 | -0.75 |
| 17919 | *KIF27* | -0.32824 | -0.79641 | -1.1333 | -1.6435 | -0.75 |
| 17920 | *CABP4* | 0.14337 | -0.76379 | -1.6386 | -2.1921 | -0.75 |
| 17921 | *TNS1* | 0.22021 | -0.69748 | -1.7822 | -2.1648 | -0.75 |
| 17922 | *GPR17* | -0.53114 | -0.7602 | -0.96857 | -1.0191 | -0.75 |
| 17923 | *SYNGR2* | 0.068993 | -0.1194 | -2.2097 | -2.4328 | -0.75 |
| 17924 | *VNN1* | 0.18765 | -0.78579 | -1.6636 | -3.3666 | -0.75 |
| 17925 | *ADAMTSL2* | -0.20227 | -0.7443 | -1.3173 | -2.1723 | -0.75 |
| 17926 | *NUP214* | 0.066172 | -0.71161 | -1.6186 | -1.7413 | -0.75 |
| 17927 | *FOXP1* | 0.19539 | -1.1659 | -1.2963 | -1.5524 | -0.76 |
| 17928 | *GIT2* | 0.19052 | -0.6706 | -1.7879 | -1.8598 | -0.76 |
| 17929 | *MSR1* | 0.26121 | -0.74069 | -1.7911 | -1.9255 | -0.76 |
| 17930 | *SLC25A28* | 0.36853 | -1.1202 | -1.5194 | -3.6071 | -0.76 |
| 17931 | *OR3A2* | -0.53093 | -0.75162 | -0.98925 | -2.0055 | -0.76 |
| 17932 | *LIMD1* | 0.020714 | -0.66073 | -1.6335 | -2.8174 | -0.76 |
| 17933 | *NOX1* | 0.19462 | -1.2118 | -1.2566 | -2.0384 | -0.76 |

| | | | | | | |
|---|---|---|---|---|---|---|
| 17934 | *NDUFA12* | 0.40229 | -0.43674 | -2.2397 | -2.6561 | -0.76 |
| 17935 | *WNK1* | -0.30232 | -0.92025 | -1.0529 | -2.7016 | -0.76 |
| 17936 | *RNF225* | -0.14289 | -0.75099 | -1.3818 | -2.0016 | -0.76 |
| 17937 | *MIS18A* | -0.3643 | -0.67441 | -1.237 | -1.4709 | -0.76 |
| 17938 | *LAD1* | -0.02618 | -1.0528 | -1.1975 | -1.4986 | -0.76 |
| 17939 | *PAGE2B* | -0.06562 | -0.76154 | -1.4495 | -2.0752 | -0.76 |
| 17940 | *MYO1C* | -0.36749 | -0.79365 | -1.1169 | -1.4809 | -0.76 |
| 17941 | *ARID3B* | -0.0291 | -1.0499 | -1.2026 | -1.5016 | -0.76 |
| 17942 | *PSME3* | 0.16473 | -0.89941 | -1.547 | -1.8631 | -0.76 |
| 17943 | *LIPK* | -0.60852 | -0.80256 | -0.87482 | -2.071 | -0.76 |
| 17944 | *CCDC183* | -0.19943 | -0.94777 | -1.1391 | -3.0179 | -0.76 |
| 17945 | *CCDC27* | 0.16285 | -0.9612 | -1.4893 | -2.1921 | -0.76 |
| 17946 | *MED6* | -0.21635 | -0.99943 | -1.073 | -1.1328 | -0.76 |
| 17947 | *ACTB* | -0.0511 | -0.91874 | -1.3199 | -2.0734 | -0.76 |
| 17948 | *PSMB3* | 0.36646 | -1.2677 | -1.3892 | -1.6949 | -0.76 |
| 17949 | *BATF3* | 0.53295 | -1.1067 | -1.72 | -2.4595 | -0.76 |
| 17950 | *KIAA0430* | 0.5809 | -1.238 | -1.6368 | -1.8609 | -0.76 |
| 17951 | *AGPAT3* | -0.50726 | -0.75603 | -1.0328 | -2.2907 | -0.77 |
| 17952 | *MCM6* | 1.1592 | -1.5379 | -1.918 | -2.6851 | -0.77 |
| 17953 | *HPR* | 0.004325 | -1.0737 | -1.2284 | -1.9919 | -0.77 |
| 17954 | *LEO1* | -0.2539 | -0.57546 | -1.4689 | -3.051 | -0.77 |
| 17955 | *CYGB* | 0.040946 | -0.23796 | -2.103 | -2.9834 | -0.77 |
| 17956 | *PCDH15* | 0.504 | -0.83808 | -1.9664 | -3.0408 | -0.77 |
| 17957 | *GNB2L1* | 0.13599 | -1.1384 | -1.2989 | -2.2485 | -0.77 |
| 17958 | *BTBD9* | 0.10845 | -1.1021 | -1.3078 | -2.4172 | -0.77 |
| 17959 | *PSME1* | -0.19806 | -0.86884 | -1.238 | -3.2901 | -0.77 |
| 17960 | *SCN7A* | -0.43726 | -0.9089 | -0.95933 | -1.3782 | -0.77 |
| 17961 | *LIMD2* | 0.18898 | -1.0528 | -1.4423 | -1.5971 | -0.77 |
| 17962 | *YRDC* | 0.14087 | -1.0948 | -1.3542 | -1.9099 | -0.77 |
| 17963 | *RASSF1* | -0.26128 | -0.86725 | -1.1807 | -1.8929 | -0.77 |
| 17964 | *CLEC18A* | 0.40022 | -1.3416 | -1.3711 | -1.4789 | -0.77 |
| 17965 | *RPL6* | -0.43008 | -0.56377 | -1.3197 | -1.7386 | -0.77 |
| 17966 | *UBE2G1* | 0.12548 | -1.1763 | -1.2628 | -2.0251 | -0.77 |
| 17967 | *WIPF3* | 0.32903 | -1.2169 | -1.4258 | -1.5375 | -0.77 |
| 17968 | *SPIN1* | -0.44134 | -0.8828 | -0.99298 | -1.4998 | -0.77 |
| 17969 | *PGLYRP1* | -0.40191 | -0.68992 | -1.2261 | -1.2782 | -0.77 |
| 17970 | *METAP2* | -0.53557 | -0.75749 | -1.0267 | -1.7675 | -0.77 |
| 17971 | *SMC2* | 0.30098 | -0.48669 | -2.1347 | -2.4415 | -0.77 |
| 17972 | *POLR2I* | 0.1739 | -1.2109 | -1.2845 | -2.4328 | -0.77 |
| 17973 | *IRF2BP1* | -0.5335 | -0.84674 | -0.94171 | -1.3593 | -0.77 |
| 17974 | *LRRC74B* | -0.59206 | -0.76882 | -0.96144 | -1.3751 | -0.77 |
| 17975 | *IKBKB* | 0.50414 | -0.83624 | -1.9906 | -2.1984 | -0.77 |
| 17976 | *CDK7* | -0.4013 | -0.8756 | -1.0458 | -2.3849 | -0.77 |
| 17977 | *WDR76* | 0.005379 | -0.86772 | -1.461 | -1.5212 | -0.77 |
| 17978 | *MTBP* | -0.44353 | -0.86298 | -1.017 | -2.4825 | -0.77 |
| 17979 | *RPGRIP1* | -0.0359 | -0.93107 | -1.3567 | -1.5883 | -0.77 |

| | | | | | | |
|---|---|---|---|---|---|---|
| 17980 | *CRYM* | 0.22609 | -0.68727 | -1.8631 | -1.9495 | -0.77 |
| 17981 | *SLC35A4* | -0.0409 | -0.69847 | -1.5866 | -1.6653 | -0.78 |
| 17982 | *NCOA2* | 0.23385 | -1.1015 | -1.4597 | -2.1089 | -0.78 |
| 17983 | *HSPG2* | 0.17593 | -1.1662 | -1.3371 | -1.6989 | -0.78 |
| 17984 | *PRDX3* | 0.75804 | -0.80897 | -2.278 | -2.2844 | -0.78 |
| 17985 | *SLC38A7* | -0.04874 | -0.97303 | -1.3074 | -1.5393 | -0.78 |
| 17986 | *QARS* | 0.18055 | -1.2247 | -1.2852 | -1.5408 | -0.78 |
| 17987 | *ADRM1* | 0.49005 | -1.3457 | -1.474 | -1.486 | -0.78 |
| 17988 | *RPL36* | 0.46067 | -0.89424 | -1.8963 | -2.1319 | -0.78 |
| 17989 | *IL17D* | 0.56384 | -0.54097 | -2.3537 | -2.6041 | -0.78 |
| 17990 | *PPP1CC* | -0.05026 | -0.62833 | -1.6524 | -1.8387 | -0.78 |
| 17991 | *KCNT2* | -0.25339 | -1.03 | -1.0482 | -1.6193 | -0.78 |
| 17992 | *RUNX2* | -0.17994 | -0.25425 | -1.8975 | -2.3195 | -0.78 |
| 17993 | *SPAG16* | -0.16415 | -0.19552 | -1.9734 | -1.988 | -0.78 |
| 17994 | *GAB2* | 0.23912 | -1.1698 | -1.4034 | -1.8975 | -0.78 |
| 17995 | *VPS72* | -0.20848 | -1.0482 | -1.0782 | -1.8646 | -0.78 |
| 17996 | *RRP1* | 0.22922 | -1.1727 | -1.3918 | -3.9492 | -0.78 |
| 17997 | *SS18L2* | -0.14701 | -0.40498 | -1.7836 | -1.7905 | -0.78 |
| 17998 | *C6orf223* | -0.00126 | -0.55441 | -1.7817 | -2.2165 | -0.78 |
| 17999 | *DUSP21* | -0.63643 | -0.85027 | -0.85296 | -1.3573 | -0.78 |
| 18000 | *EIF3CL* | -0.30209 | -0.63483 | -1.4031 | -1.53 | -0.78 |
| 18001 | *KRTAP6-1* | -0.04538 | -0.57429 | -1.7213 | -2.3474 | -0.78 |
| 18002 | *HAPLN4* | -0.30756 | -0.88199 | -1.1545 | -1.309 | -0.78 |
| 18003 | *RGS18* | 0.79993 | -1.1921 | -1.9534 | -2.3776 | -0.78 |
| 18004 | *EPM2A* | 0.13796 | -0.90154 | -1.584 | -2.1873 | -0.78 |
| 18005 | *C17orf102* | -0.34349 | -0.77196 | -1.2343 | -1.8692 | -0.78 |
| 18006 | *DDOST* | -0.35015 | -0.97303 | -1.0268 | -1.0602 | -0.78 |
| 18007 | *MFN2* | 0.68743 | -1.3646 | -1.6734 | -2.5489 | -0.78 |
| 18008 | *GLIPR2* | -0.13752 | -0.41298 | -1.8033 | -2.6935 | -0.78 |
| 18009 | *RPL19* | -0.67716 | -0.82293 | -0.85623 | -1.1136 | -0.79 |
| 18010 | *TBC1D3H* | -0.37417 | -0.83138 | -1.1524 | -1.8073 | -0.79 |
| 18011 | *TMEM160* | 1.1873 | -1.1012 | -2.4456 | -5.7525 | -0.79 |
| 18012 | *G6PD* | -0.52477 | -0.8239 | -1.0111 | -2.2075 | -0.79 |
| 18013 | *POM121L2* | -0.65455 | -0.75472 | -0.95068 | -2.0126 | -0.79 |
| 18014 | *SMARCD1* | -0.54959 | -0.8681 | -0.94296 | -2.1533 | -0.79 |
| 18015 | *ABCE1* | -0.39565 | -0.43633 | -1.5289 | -1.7675 | -0.79 |
| 18016 | *RGMA* | -0.14646 | -1.0564 | -1.1585 | -1.5026 | -0.79 |
| 18017 | *SEMA5B* | -0.16832 | -0.71093 | -1.4824 | -1.525 | -0.79 |
| 18018 | *MAPK1* | 0.20136 | -1.1169 | -1.4473 | -2.1648 | -0.79 |
| 18019 | *FAM9A* | -0.33856 | -1.0092 | -1.0156 | -1.237 | -0.79 |
| 18020 | *KLK14* | 0.12094 | -1.2215 | -1.2633 | -1.4811 | -0.79 |
| 18021 | *DEDD2* | 0.12312 | -0.96718 | -1.5206 | -1.6221 | -0.79 |
| 18022 | *IQCF6* | 0.26885 | -0.90161 | -1.7323 | -2.0249 | -0.79 |
| 18023 | *POM121C* | -0.06057 | -0.87046 | -1.4365 | -1.5064 | -0.79 |
| 18024 | *NDC80* | -0.319 | -0.62479 | -1.4269 | -3.7031 | -0.79 |
| 18025 | *FNTB* | -0.35308 | -0.6602 | -1.358 | -1.3675 | -0.79 |

| | | | | | | |
|---|---|---|---|---|---|---|
| 18026 | *LY6G6D* | -0.35694 | -0.45748 | -1.5569 | -5.9834 | -0.79 |
| 18027 | *MUC1* | -0.37872 | -0.7602 | -1.2326 | -1.6139 | -0.79 |
| 18028 | *C20orf196* | 0.11191 | -1.227 | -1.2569 | -1.5077 | -0.79 |
| 18029 | *IL31RA* | -0.00414 | -1.1317 | -1.2364 | -1.4187 | -0.79 |
| 18030 | *RMDN2* | -0.34229 | -0.781 | -1.2505 | -1.6594 | -0.79 |
| 18031 | *ARL2* | -0.07154 | -0.68999 | -1.6176 | -2.0351 | -0.79 |
| 18032 | *LDB3* | -0.30237 | -0.95321 | -1.1247 | -1.4588 | -0.79 |
| 18033 | *GTPBP4* | -0.55175 | -0.90386 | -0.92486 | -1.3864 | -0.79 |
| 18034 | *CRADD* | 0.75207 | -0.78651 | -2.3461 | -2.7376 | -0.79 |
| 18035 | *ITGAE* | -0.13895 | -1.0644 | -1.1773 | -2.5865 | -0.79 |
| 18036 | *HSPA5* | -0.36377 | -0.64577 | -1.3723 | -1.5335 | -0.79 |
| 18037 | *KLK9* | -0.5731 | -0.86123 | -0.94815 | -1.3835 | -0.79 |
| 18038 | *TMPPE* | -0.5635 | -0.82375 | -0.99619 | -1.8443 | -0.79 |
| 18039 | *ITGA11* | -0.05322 | -0.92553 | -1.4048 | -1.6934 | -0.79 |
| 18040 | *BATF* | 0.45156 | -0.82311 | -2.012 | -2.6096 | -0.79 |
| 18041 | *PAX4* | 0.82699 | -1.0951 | -2.1158 | -2.1407 | -0.79 |
| 18042 | *QPCTL* | -0.0702 | -0.93725 | -1.3766 | -2.2046 | -0.79 |
| 18043 | *ANO1* | -0.28901 | -0.73424 | -1.3617 | -2.0602 | -0.79 |
| 18044 | *VAPB* | 0.15515 | -0.11251 | -2.428 | -2.5725 | -0.80 |
| 18045 | *TUBG1* | 0.25584 | -0.88171 | -1.7599 | -2.5386 | -0.80 |
| 18046 | *PPIL3* | 0.052522 | -0.56518 | -1.8737 | -2.3615 | -0.80 |
| 18047 | *FBXW2* | -0.46747 | -0.93188 | -0.9879 | -2.0593 | -0.80 |
| 18048 | *PRKCQ* | 0.40269 | -1.1836 | -1.6069 | -3.0608 | -0.80 |
| 18049 | *SQSTM1* | -0.31227 | -0.86459 | -1.2113 | -1.7018 | -0.80 |
| 18050 | *RPS6KA5* | 0.22057 | -1.1713 | -1.4401 | -1.6276 | -0.80 |
| 18051 | *THEG* | -0.28336 | -1.054 | -1.0553 | -1.0855 | -0.80 |
| 18052 | *GPAM* | 0.40041 | -1.3851 | -1.4085 | -1.714 | -0.80 |
| 18053 | *CD177* | -0.64751 | -0.65692 | -1.0907 | -1.3097 | -0.80 |
| 18054 | *EIF5A* | -0.55898 | -0.78004 | -1.0564 | -1.8154 | -0.80 |
| 18055 | *MAP4* | -0.35735 | -1.0092 | -1.0293 | -1.3684 | -0.80 |
| 18056 | *MRPL38* | -0.11621 | -1.042 | -1.2393 | -1.358 | -0.80 |
| 18057 | *GTF2H3* | -0.43473 | -0.86027 | -1.103 | -1.9802 | -0.80 |
| 18058 | *POGZ* | -0.40112 | -0.9935 | -1.0048 | -1.5781 | -0.80 |
| 18059 | *CETN2* | -0.36861 | -0.79641 | -1.2352 | -1.6844 | -0.80 |
| 18060 | *USP10* | 0.20962 | -1.0905 | -1.5206 | -1.7169 | -0.80 |
| 18061 | *KCNH6* | 0.52394 | -1.3154 | -1.6104 | -1.9342 | -0.80 |
| 18062 | *GON4L* | -0.52496 | -0.69815 | -1.1808 | -3.6071 | -0.80 |
| 18063 | *ZNHIT6* | -0.43385 | -0.79596 | -1.1743 | -2.337 | -0.80 |
| 18064 | *TMX4* | -0.02488 | -0.16207 | -2.2172 | -2.3036 | -0.80 |
| 18065 | *HNRNPM* | -0.00252 | -0.60987 | -1.7918 | -1.8066 | -0.80 |
| 18066 | *CDT1* | -0.76513 | -0.81368 | -0.8259 | -1.8771 | -0.80 |
| 18067 | *KPNA6* | 0.21297 | -0.96632 | -1.6575 | -1.8157 | -0.80 |
| 18068 | *SDR39U1* | 0.37688 | -0.98688 | -1.802 | -3.6459 | -0.80 |
| 18069 | *ACTA2* | -0.33821 | -0.92375 | -1.151 | -2.0059 | -0.80 |
| 18070 | *SSTR5* | 0.008446 | -0.8462 | -1.5765 | -1.9442 | -0.80 |
| 18071 | *PIGV* | -0.12552 | -0.90991 | -1.3797 | -1.3821 | -0.81 |

| | | | | | | |
|---|---|---|---|---|---|---|
| 18072 | *NCBP1* | -0.19885 | -0.73834 | -1.4788 | -1.6397 | -0.81 |
| 18073 | *LRFN5* | 0.11514 | -0.89794 | -1.6356 | -1.658 | -0.81 |
| 18074 | *ITGA10* | 0.12694 | -1.2436 | -1.3025 | -1.4015 | -0.81 |
| 18075 | *SIN3A* | -0.31654 | -0.5775 | -1.526 | -1.6443 | -0.81 |
| 18076 | *MAGED4* | 0.44581 | -1.2297 | -1.6368 | -2.4784 | -0.81 |
| 18077 | *CCSAP* | -0.72466 | -0.7272 | -0.97029 | -1.5735 | -0.81 |
| 18078 | *AGTR1* | -0.51304 | -0.72553 | -1.1836 | -1.5212 | -0.81 |
| 18079 | *TCF24* | 0.46058 | -1.4416 | -1.4421 | -3.1229 | -0.81 |
| 18080 | *EMC3* | -0.34148 | -0.61033 | -1.4722 | -1.742 | -0.81 |
| 18081 | *ATP2A3* | -0.39885 | -0.81999 | -1.2055 | -2.641 | -0.81 |
| 18082 | *ANO7* | -0.40281 | -0.95182 | -1.0707 | -1.8076 | -0.81 |
| 18083 | *SLC17A5* | 0.02465 | -0.56411 | -1.8872 | -2.264 | -0.81 |
| 18084 | *SLC8A3* | 0.11412 | -1.2342 | -1.3071 | -1.8115 | -0.81 |
| 18085 | *GPR107* | 0.43085 | -0.89794 | -1.962 | -2.236 | -0.81 |
| 18086 | *MICU2* | 0.22852 | -1.3226 | -1.3354 | -1.5861 | -0.81 |
| 18087 | *KBTBD8* | -0.27453 | -0.74618 | -1.4093 | -1.9284 | -0.81 |
| 18088 | *USP31* | -0.11697 | -0.43481 | -1.8816 | -2.4703 | -0.81 |
| 18089 | *CTBP1* | -0.01175 | -1.1872 | -1.2362 | -1.6653 | -0.81 |
| 18090 | *SPINK4* | 0.16764 | -1.1351 | -1.4684 | -1.5752 | -0.81 |
| 18091 | *BPIFB6* | -0.32183 | -0.95105 | -1.1659 | -1.9757 | -0.81 |
| 18092 | *IPO4* | -0.34327 | -1.0231 | -1.0741 | -1.2343 | -0.81 |
| 18093 | *PPIAL4F* | 0.52665 | -1.2021 | -1.7652 | -2.2292 | -0.81 |
| 18094 | *USB1* | -0.38673 | -0.87898 | -1.1763 | -1.2903 | -0.81 |
| 18095 | *MFSD9* | -0.30782 | -0.37912 | -1.7551 | -1.838 | -0.81 |
| 18096 | *TARDBP* | 0.20227 | -0.74155 | -1.9047 | -2.0016 | -0.81 |
| 18097 | *TOR1B* | -0.01344 | -0.88465 | -1.5465 | -1.624 | -0.81 |
| 18098 | *DEFB128* | 0.21353 | -1.2816 | -1.3766 | -2.0629 | -0.81 |
| 18099 | *MAPK1IP1L* | -0.04896 | -1.1384 | -1.2576 | -1.4607 | -0.81 |
| 18100 | *KCNC2* | -0.5716 | -0.89525 | -0.97918 | -1.3019 | -0.82 |
| 18101 | *CRY1* | 0.26829 | -1.0005 | -1.7149 | -2.5478 | -0.82 |
| 18102 | *CFLAR* | -0.08091 | -1.0782 | -1.2884 | -1.7129 | -0.82 |
| 18103 | *MRPS21* | -0.57732 | -0.61494 | -1.2572 | -1.5408 | -0.82 |
| 18104 | *ATP5L2* | 0.25904 | -1.0567 | -1.6528 | -2.0791 | -0.82 |
| 18105 | *GLG1* | 0.68472 | -1.525 | -1.6105 | -2.4642 | -0.82 |
| 18106 | *CDC25B* | -0.68565 | -0.86287 | -0.90687 | -1.0424 | -0.82 |
| 18107 | *ARL3* | 0.61327 | -1.4919 | -1.5782 | -2.7379 | -0.82 |
| 18108 | *FAM183A* | 0.21133 | -1.2987 | -1.3707 | -1.9998 | -0.82 |
| 18109 | *OR8H3* | -0.21563 | -0.86756 | -1.3751 | -1.9163 | -0.82 |
| 18110 | *CDK5RAP1* | 0.29779 | -1.3148 | -1.4423 | -2.071 | -0.82 |
| 18111 | *TACC3* | -0.52847 | -0.95717 | -0.97572 | -1.2915 | -0.82 |
| 18112 | *TWSG1* | -0.28825 | -0.89186 | -1.2816 | -2.2343 | -0.82 |
| 18113 | *TSPY8* | -0.66708 | -0.71988 | -1.078 | -1.0898 | -0.82 |
| 18114 | *ALG1L* | -0.27801 | -0.86047 | -1.3267 | -1.4926 | -0.82 |
| 18115 | *VCPKMT* | 0.1973 | -0.99608 | -1.6682 | -1.8775 | -0.82 |
| 18116 | *CEP19* | 0.27723 | -1.356 | -1.3903 | -1.6989 | -0.82 |
| 18117 | *UTP6* | 0.23864 | -1.3218 | -1.386 | -1.9367 | -0.82 |

| | | | | | | |
|---|---|---|---|---|---|---|
| 18118 | *ATP2B3* | 0.24021 | -0.78956 | -1.9219 | -2.0384 | -0.82 |
| 18119 | *HOXD13* | -0.19997 | -0.75824 | -1.5138 | -2.264 | -0.82 |
| 18120 | *NFATC1* | -0.06844 | -1.0344 | -1.3721 | -3.2348 | -0.82 |
| 18121 | *CAD* | -0.35308 | -0.92157 | -1.202 | -1.9107 | -0.83 |
| 18122 | *ATP5B* | -0.09922 | -1.1856 | -1.192 | -1.3191 | -0.83 |
| 18123 | *C12orf42* | 0.44675 | -0.99871 | -1.9269 | -2.1873 | -0.83 |
| 18124 | *RHOBTB2* | -0.51592 | -0.83276 | -1.1327 | -1.6316 | -0.83 |
| 18125 | *VIP* | -0.30454 | -1.0258 | -1.1511 | -1.4607 | -0.83 |
| 18126 | *SIGLEC12* | 0.18646 | -1.1216 | -1.5467 | -1.6047 | -0.83 |
| 18127 | *PCNXL4* | -0.06343 | -0.44353 | -1.9758 | -2.071 | -0.83 |
| 18128 | *WDFY4* | -0.08018 | -1.1325 | -1.2733 | -1.72 | -0.83 |
| 18129 | *KRT32* | -0.47001 | -0.93652 | -1.0796 | -2.2094 | -0.83 |
| 18130 | *POLR3D* | 0.76582 | -1.5918 | -1.6604 | -2.0953 | -0.83 |
| 18131 | *FAM229A* | 0.073809 | -0.88584 | -1.6748 | -2.4018 | -0.83 |
| 18132 | *ANKRD32* | -0.42177 | -0.66251 | -1.4039 | -1.9622 | -0.83 |
| 18133 | *GLYR1* | -0.19364 | -0.67146 | -1.6246 | -2.5934 | -0.83 |
| 18134 | *KRTCAP3* | 0.67594 | -0.8504 | -2.3154 | -3.6181 | -0.83 |
| 18135 | *ATP6V0B* | -0.15181 | -1.0945 | -1.2464 | -1.3792 | -0.83 |
| 18136 | *EPG5* | 0.098319 | -1.1792 | -1.4135 | -2.1089 | -0.83 |
| 18137 | *U2AF2* | 0.47249 | -0.20512 | -2.7623 | -3.5278 | -0.83 |
| 18138 | *VN1R2* | -0.21508 | -0.72877 | -1.5524 | -2.2694 | -0.83 |
| 18139 | *PRG3* | -0.43726 | -0.87382 | -1.1888 | -1.4994 | -0.83 |
| 18140 | *CHIA* | -0.13688 | -0.79485 | -1.5687 | -1.9865 | -0.83 |
| 18141 | *LCT* | -0.3337 | -1.0335 | -1.1337 | -3.7932 | -0.83 |
| 18142 | *SMIM12* | -0.54197 | -0.89501 | -1.0641 | -1.7068 | -0.83 |
| 18143 | *FBLN2* | -0.09493 | -1.065 | -1.3416 | -1.3501 | -0.83 |
| 18144 | *HIST1H2AM* | -0.47857 | -0.61663 | -1.4087 | -2.3063 | -0.83 |
| 18145 | *FAM185A* | 0.033893 | -0.50053 | -2.0392 | -2.1177 | -0.84 |
| 18146 | *GDAP2* | 0.86091 | -1.183 | -2.1848 | -2.4718 | -0.84 |
| 18147 | *MRPS26* | -0.77144 | -0.81798 | -0.91779 | -2.2223 | -0.84 |
| 18148 | *SERPINB13* | 0.8794 | -1.5722 | -1.8157 | -2.4933 | -0.84 |
| 18149 | *C22orf42* | -0.0232 | -1.1892 | -1.297 | -1.5434 | -0.84 |
| 18150 | *DBNDD1* | 0.11133 | -1.2098 | -1.411 | -1.5243 | -0.84 |
| 18151 | *CD22* | -0.47806 | -0.76361 | -1.268 | -1.9168 | -0.84 |
| 18152 | *ABTB1* | -0.54884 | -0.90206 | -1.0589 | -1.5604 | -0.84 |
| 18153 | *C8orf22* | 0.003285 | -1.1676 | -1.348 | -3.8124 | -0.84 |
| 18154 | *EID1* | 0.79832 | -1.3773 | -1.9347 | -2.4015 | -0.84 |
| 18155 | *ZNF2* | -0.77462 | -0.8504 | -0.89009 | -1.3426 | -0.84 |
| 18156 | *ACRV1* | 0.33948 | -0.28352 | -2.5725 | -2.8776 | -0.84 |
| 18157 | *TEX13B* | -0.10704 | -0.79149 | -1.6218 | -2.1217 | -0.84 |
| 18158 | *TRAPPC3* | -0.58289 | -0.65189 | -1.2867 | -1.8759 | -0.84 |
| 18159 | *C18orf8* | -0.63575 | -0.92157 | -0.96572 | -1.2639 | -0.84 |
| 18160 | *KLHL30* | 0.13705 | -0.52587 | -2.1351 | -2.1792 | -0.84 |
| 18161 | *MRPL33* | -0.61643 | -0.85392 | -1.0557 | -1.5884 | -0.84 |
| 18162 | *TBC1D3C* | -0.56605 | -0.97252 | -0.98749 | -1.9954 | -0.84 |
| 18163 | *ANTXRL* | -0.3464 | -0.8934 | -1.2864 | -2.781 | -0.84 |

| | | | | | | |
|---|---|---|---|---|---|---|
| 18164 | *CEP120* | -0.17298 | -1.1522 | -1.2027 | -1.6557 | -0.84 |
| 18165 | *TRIM35* | -0.61344 | -0.832 | -1.0829 | -1.1566 | -0.84 |
| 18166 | *CCHCR1* | 0.20958 | -0.85427 | -1.8869 | -2.0682 | -0.84 |
| 18167 | *HHIP* | 0.37811 | -0.95759 | -1.9523 | -1.962 | -0.84 |
| 18168 | *PRDM14* | -0.07472 | -0.93465 | -1.5225 | -1.7287 | -0.84 |
| 18169 | *FIBP* | 0.31949 | -1.085 | -1.7667 | -2.4328 | -0.84 |
| 18170 | *RBM7* | -0.10243 | -0.87257 | -1.5604 | -2.8288 | -0.85 |
| 18171 | *HSP90AA1* | 0.15481 | -1.2867 | -1.4039 | -2.412 | -0.85 |
| 18172 | *SH3TC1* | 0.44466 | -1.4614 | -1.5212 | -3.0961 | -0.85 |
| 18173 | *OTOGL* | 0.28766 | -0.98951 | -1.8364 | -2.0304 | -0.85 |
| 18174 | *CPSF1* | -0.72529 | -0.78878 | -1.0243 | -1.6526 | -0.85 |
| 18175 | *MED12* | 0.87228 | -1.6861 | -1.7261 | -2.4112 | -0.85 |
| 18176 | *NPIPA5* | -0.45005 | -0.81178 | -1.2786 | -1.4234 | -0.85 |
| 18177 | *CDC40* | -0.66731 | -0.8518 | -1.0224 | -1.7911 | -0.85 |
| 18178 | *F3* | -0.57513 | -0.87257 | -1.0949 | -2.0351 | -0.85 |
| 18179 | *CLDND2* | 0.14973 | -1.2935 | -1.3993 | -1.8966 | -0.85 |
| 18180 | *RPL10* | 0.096242 | -0.88584 | -1.7538 | -1.8695 | -0.85 |
| 18181 | *LSM5* | -0.45345 | -0.50251 | -1.5904 | -2.01 | -0.85 |
| 18182 | *NEK6* | -0.66353 | -0.72038 | -1.1629 | -1.3479 | -0.85 |
| 18183 | *KCNK18* | -0.15866 | -0.8108 | -1.5781 | -1.658 | -0.85 |
| 18184 | *CBWD1* | -0.44353 | -0.80234 | -1.3017 | -2.1709 | -0.85 |
| 18185 | *LEMD1* | -0.0146 | -1.1898 | -1.3434 | -3.2405 | -0.85 |
| 18186 | *ZKSCAN5* | -0.07193 | -0.67556 | -1.8007 | -2.0653 | -0.85 |
| 18187 | *GPR180* | 0.037116 | -1.2792 | -1.3074 | -1.4584 | -0.85 |
| 18188 | *TBX6* | -0.34962 | -0.89219 | -1.3085 | -1.4773 | -0.85 |
| 18189 | *SPDL1* | -0.04612 | -0.98842 | -1.5167 | -2.5302 | -0.85 |
| 18190 | *DALRD3* | -0.33787 | -0.93789 | -1.2756 | -2.3154 | -0.85 |
| 18191 | *TNFAIP3* | -0.05175 | -0.94393 | -1.5568 | -2.8797 | -0.85 |
| 18192 | *CFAP61* | -0.21821 | -1.1075 | -1.2273 | -1.2577 | -0.85 |
| 18193 | *RPL11* | 0.071812 | -0.87955 | -1.7459 | -2.3949 | -0.85 |
| 18194 | *PAX7* | 0.28506 | -0.9512 | -1.8885 | -2.0028 | -0.85 |
| 18195 | *LRRC10B* | 0.13231 | -0.86526 | -1.8233 | -1.8435 | -0.85 |
| 18196 | *TTC37* | -0.04085 | -1.0576 | -1.4584 | -2.6072 | -0.85 |
| 18197 | *B3GALNT2* | -0.55791 | -0.56157 | -1.4392 | -3.1051 | -0.85 |
| 18198 | *GLIPR1L2* | -0.44498 | -0.94747 | -1.1665 | -1.831 | -0.85 |
| 18199 | *SLC6A13* | -0.63214 | -0.87444 | -1.0528 | -1.7158 | -0.85 |
| 18200 | *COPS8* | -0.38198 | -0.86556 | -1.3141 | -1.6213 | -0.85 |
| 18201 | *EBPL* | -0.33848 | -0.92077 | -1.3036 | -2.1566 | -0.85 |
| 18202 | *SLC7A9* | -0.44353 | -0.51653 | -1.6047 | -2.3014 | -0.85 |
| 18203 | *ALG5* | -0.75161 | -0.85386 | -0.96012 | -1.8363 | -0.86 |
| 18204 | *C2orf44* | -0.04457 | -1.2014 | -1.3203 | -1.7967 | -0.86 |
| 18205 | *DNAJC7* | 0.37062 | -1.1439 | -1.7963 | -2.4456 | -0.86 |
| 18206 | *ZNF224* | 0.18719 | -0.21452 | -2.5452 | -2.8206 | -0.86 |
| 18207 | *DNAJC12* | -0.67599 | -0.81068 | -1.0889 | -2.7626 | -0.86 |
| 18208 | *CYLD* | -0.27434 | -0.89136 | -1.4141 | -1.6722 | -0.86 |
| 18209 | *NEURL2* | -0.30608 | -0.68813 | -1.5874 | -1.9622 | -0.86 |

| | | | | | | |
|---|---|---|---|---|---|---|
| 18210 | *C1QBP* | 0.16793 | -1.1138 | -1.6365 | -2.5677 | -0.86 |
| 18211 | *KIFC3* | -0.74392 | -0.77583 | -1.0633 | -1.3633 | -0.86 |
| 18212 | *WIF1* | -0.2985 | -0.9105 | -1.3741 | -1.7117 | -0.86 |
| 18213 | *SELT* | -0.33895 | -0.95893 | -1.2864 | -2.4784 | -0.86 |
| 18214 | *C1orf27* | 1.1403 | -1.0385 | -2.6889 | -3.1761 | -0.86 |
| 18215 | *CFC1* | 0.17256 | -1.3364 | -1.4233 | -1.5125 | -0.86 |
| 18216 | *SLC25A1* | 0.39669 | -1.268 | -1.716 | -1.7606 | -0.86 |
| 18217 | *IMP3* | -0.14403 | -1.1214 | -1.322 | -1.9162 | -0.86 |
| 18218 | *MEIKIN* | -0.62407 | -0.7476 | -1.2165 | -3.3669 | -0.86 |
| 18219 | *ETS1* | -0.69287 | -0.85296 | -1.043 | -1.244 | -0.86 |
| 18220 | *NR6A1* | 0.60604 | -1.2852 | -1.9116 | -2.9736 | -0.86 |
| 18221 | *BBX* | -0.15385 | -1.0231 | -1.4141 | -2.0885 | -0.86 |
| 18222 | *PRICKLE4* | -0.02232 | -0.89593 | -1.6731 | -1.9446 | -0.86 |
| 18223 | *COX4I1* | 0.17372 | -1.0871 | -1.6787 | -1.9326 | -0.86 |
| 18224 | *BTN2A1* | -0.36311 | -0.98925 | -1.2412 | -1.4142 | -0.86 |
| 18225 | *CMTM4* | -0.0862 | -0.49778 | -2.0107 | -2.1626 | -0.86 |
| 18226 | *C12orf79* | -0.01195 | -1.2646 | -1.3184 | -2.0351 | -0.86 |
| 18227 | *MRPL17* | -0.51845 | -0.6258 | -1.451 | -1.6306 | -0.87 |
| 18228 | *QRSL1* | -0.58058 | -0.71363 | -1.3017 | -2.1921 | -0.87 |
| 18229 | *CITED4* | 0.95013 | -1.6731 | -1.8742 | -2.0483 | -0.87 |
| 18230 | *TMC4* | 0.048059 | -1.1613 | -1.4842 | -2.3885 | -0.87 |
| 18231 | *ZNF626* | -0.33601 | -0.70581 | -1.5568 | -1.8737 | -0.87 |
| 18232 | *CNGA3* | 0.36192 | -1.2884 | -1.6741 | -2.8406 | -0.87 |
| 18233 | *EIF4H* | 0.39018 | -1.4467 | -1.545 | -1.7879 | -0.87 |
| 18234 | *RAB9B* | 0.66434 | -0.31333 | -2.9538 | -3.7266 | -0.87 |
| 18235 | *CRYBB1* | -0.23846 | -0.95987 | -1.4055 | -1.7911 | -0.87 |
| 18236 | *ADGRA2* | 0.29943 | -1.1429 | -1.7608 | -2.0452 | -0.87 |
| 18237 | *DBF4* | -0.16325 | -0.79821 | -1.6447 | -2.1347 | -0.87 |
| 18238 | *U2AF1* | 0.12804 | -1.1732 | -1.5611 | -1.6281 | -0.87 |
| 18239 | *ATMIN* | -0.03016 | -1.2295 | -1.348 | -1.658 | -0.87 |
| 18240 | *PIK3R2* | 0.069078 | -0.87773 | -1.8016 | -2.1321 | -0.87 |
| 18241 | *ORC2* | -0.36086 | -1.0035 | -1.2466 | -1.6476 | -0.87 |
| 18242 | *SERBP1* | -0.30286 | -0.76355 | -1.545 | -2.4946 | -0.87 |
| 18243 | *STOML2* | -0.81031 | -0.81282 | -0.98867 | -0.99463 | -0.87 |
| 18244 | *CDC42SE2* | -0.4589 | -1.0291 | -1.1263 | -1.3457 | -0.87 |
| 18245 | *POC1A* | -0.03196 | -0.93865 | -1.6437 | -1.8775 | -0.87 |
| 18246 | *TXN2* | -0.11653 | -1.1303 | -1.3675 | -3.9779 | -0.87 |
| 18247 | *HMCES* | -0.53668 | -0.9765 | -1.1016 | -2.7736 | -0.87 |
| 18248 | *JUNB* | -0.54479 | -0.97113 | -1.0997 | -1.8537 | -0.87 |
| 18249 | *NTRK2* | 1.2246 | -1.8908 | -1.9495 | -2.4741 | -0.87 |
| 18250 | *LSM8* | 0.17305 | -1.2174 | -1.5744 | -1.739 | -0.87 |
| 18251 | *NDUFA4L2* | -0.26369 | -0.70289 | -1.6525 | -2.8367 | -0.87 |
| 18252 | *SRPR* | 0.39446 | -1.4183 | -1.5956 | -1.7652 | -0.87 |
| 18253 | *PPAT* | -0.22414 | -1.0119 | -1.3842 | -1.5988 | -0.87 |
| 18254 | *C9orf131* | -0.01678 | -1.1802 | -1.4234 | -1.802 | -0.87 |
| 18255 | *CAMTA1* | 0.026876 | -1.2867 | -1.3637 | -1.72 | -0.87 |

| | | | | | | |
|---|---|---|---|---|---|---|
| 18256 | *STX17* | -0.11695 | -0.93489 | -1.5727 | -1.8157 | -0.87 |
| 18257 | *GNL3* | -0.43295 | -0.88702 | -1.3078 | -1.7459 | -0.88 |
| 18258 | *BAZ2A* | 0.15856 | -0.47862 | -2.3096 | -2.382 | -0.88 |
| 18259 | *POP5* | -0.5193 | -0.89501 | -1.2167 | -1.6128 | -0.88 |
| 18260 | *P2RX4* | 0.26893 | -1.2709 | -1.632 | -1.7978 | -0.88 |
| 18261 | *USP17L27* | -0.26592 | -1.1479 | -1.2202 | -1.316 | -0.88 |
| 18262 | *BUB3* | -0.34506 | -0.81152 | -1.4777 | -2.1212 | -0.88 |
| 18263 | *URB1* | -0.662 | -0.84386 | -1.13 | -2.2385 | -0.88 |
| 18264 | *GTPBP8* | -0.7241 | -0.93474 | -0.97914 | -1.9695 | -0.88 |
| 18265 | *WNT5A* | 0.27588 | -0.73517 | -2.1796 | -3.6038 | -0.88 |
| 18266 | *MSL3* | 0.17562 | -1.2246 | -1.5915 | -1.9295 | -0.88 |
| 18267 | *TERT* | -0.48021 | -0.83276 | -1.3279 | -1.5832 | -0.88 |
| 18268 | *NOL11* | 0.10312 | -0.94536 | -1.7991 | -1.8714 | -0.88 |
| 18269 | *CRHBP* | -0.04717 | -1.1974 | -1.397 | -2.262 | -0.88 |
| 18270 | *ISY1* | -0.67432 | -0.76821 | -1.1992 | -2.0429 | -0.88 |
| 18271 | *IGFN1* | -0.07337 | -1.2208 | -1.3478 | -1.7538 | -0.88 |
| 18272 | *MGC50722* | -0.44634 | -0.97393 | -1.2217 | -2.5414 | -0.88 |
| 18273 | *BCORL1* | -0.2873 | -0.93368 | -1.4221 | -1.5806 | -0.88 |
| 18274 | *THBD* | 0.24025 | -1.3903 | -1.4944 | -1.7649 | -0.88 |
| 18275 | *ESF1* | -0.62099 | -0.93237 | -1.0912 | -2.1125 | -0.88 |
| 18276 | *CHMP4B* | 0.054282 | -1.0666 | -1.6336 | -1.8441 | -0.88 |
| 18277 | *TFB1M* | -0.46228 | -0.75625 | -1.4276 | -1.9172 | -0.88 |
| 18278 | *TULP1* | 0.24612 | -1.4325 | -1.462 | -2.6003 | -0.88 |
| 18279 | *ARFIP1* | 0.42399 | -1.156 | -1.9166 | -2.0483 | -0.88 |
| 18280 | *CFB* | -0.3943 | -0.83739 | -1.4171 | -1.6663 | -0.88 |
| 18281 | *KDM2A* | -0.49241 | -0.91396 | -1.2457 | -2.005 | -0.88 |
| 18282 | *ZNF879* | -0.17622 | -1.0384 | -1.4392 | -1.7149 | -0.88 |
| 18283 | *FBXW8* | -0.49598 | -0.72515 | -1.4333 | -1.5901 | -0.88 |
| 18284 | *IL23A* | 0.49003 | -1.3246 | -1.8212 | -2.9818 | -0.89 |
| 18285 | *EME1* | 0.32411 | -1.1558 | -1.8297 | -2.0016 | -0.89 |
| 18286 | *PALB2* | 0.06576 | -1.3211 | -1.4064 | -1.42 | -0.89 |
| 18287 | *C19orf84* | -0.65714 | -0.88267 | -1.122 | -1.1343 | -0.89 |
| 18288 | *SMIM18* | 0.20116 | -1.3926 | -1.4704 | -2.8463 | -0.89 |
| 18289 | *CD8B* | 0.16025 | -1.3288 | -1.496 | -2.0548 | -0.89 |
| 18290 | *CHRM5* | -0.7001 | -0.85799 | -1.1075 | -1.1263 | -0.89 |
| 18291 | *SUPV3L1* | -0.36821 | -0.78362 | -1.5139 | -1.6422 | -0.89 |
| 18292 | *FOXI3* | -0.38406 | -0.93865 | -1.3457 | -2.0116 | -0.89 |
| 18293 | *HECTD1* | -0.41139 | -0.81496 | -1.445 | -1.6241 | -0.89 |
| 18294 | *SERINC4* | -0.74308 | -0.8737 | -1.0559 | -1.418 | -0.89 |
| 18295 | *ZNF287* | -0.07439 | -1.2461 | -1.3525 | -1.743 | -0.89 |
| 18296 | *TMEM8A* | 0.38037 | -1.4584 | -1.5959 | -1.8402 | -0.89 |
| 18297 | *RALGPS2* | -0.35111 | -0.76582 | -1.5579 | -1.9398 | -0.89 |
| 18298 | *DNTTIP2* | -0.55002 | -0.69397 | -1.4341 | -1.8316 | -0.89 |
| 18299 | *IL21* | -0.36761 | -0.97595 | -1.3356 | -3.596 | -0.89 |
| 18300 | *TRUB2* | -0.20512 | -0.68727 | -1.7884 | -5.6002 | -0.89 |
| 18301 | *CNTNAP3* | -0.08477 | -1.2676 | -1.334 | -1.4996 | -0.90 |

| | | | | | | |
|---|---|---|---|---|---|---|
| 18302 | *CALB2* | 0.69516 | -1.6456 | -1.7364 | -1.984 | -0.90 |
| 18303 | *SNRPG* | -0.19995 | -0.83669 | -1.6526 | -1.8577 | -0.90 |
| 18304 | *RNFT1* | -0.04009 | -0.83145 | -1.8193 | -2.2323 | -0.90 |
| 18305 | *STRAP* | -0.15439 | -0.80341 | -1.7364 | -2.4268 | -0.90 |
| 18306 | *GOLGA6L6* | 0.16906 | -1.3721 | -1.4926 | -1.6711 | -0.90 |
| 18307 | *RPL17* | -0.2098 | -1.2033 | -1.2829 | -1.9326 | -0.90 |
| 18308 | *PTPRF* | -0.29549 | -0.92464 | -1.4761 | -1.7735 | -0.90 |
| 18309 | *LRRC20* | -0.76606 | -0.91701 | -1.0188 | -1.2302 | -0.90 |
| 18310 | *RPS23* | -0.28009 | -0.88143 | -1.5408 | -1.8755 | -0.90 |
| 18311 | *WDR4* | -0.1138 | -0.77394 | -1.8154 | -2.4221 | -0.90 |
| 18312 | *POLR3H* | -0.11044 | -1.2729 | -1.3198 | -1.5592 | -0.90 |
| 18313 | *TPGS2* | -0.25181 | -0.72553 | -1.7263 | -2.8928 | -0.90 |
| 18314 | *RBM23* | 0.33274 | -1.2876 | -1.7493 | -2.8553 | -0.90 |
| 18315 | *RSPH4A* | 0.53276 | -0.81034 | -2.428 | -2.5865 | -0.90 |
| 18316 | *LIPT2* | 0.30909 | -1.1943 | -1.8219 | -1.9727 | -0.90 |
| 18317 | *TGM2* | -0.08601 | -1.0828 | -1.5405 | -1.6644 | -0.90 |
| 18318 | *RNF151* | -0.04432 | -0.73172 | -1.9358 | -1.9965 | -0.90 |
| 18319 | *EIF4G1* | -0.21905 | -0.93727 | -1.5558 | -2.1478 | -0.90 |
| 18320 | *VPS8* | -0.50676 | -0.68133 | -1.5243 | -1.5401 | -0.90 |
| 18321 | *BRAP* | 0.49383 | -1.0102 | -2.1984 | -2.2586 | -0.90 |
| 18322 | *MED28* | 0.5736 | -1.5824 | -1.7065 | -3.1119 | -0.91 |
| 18323 | *C16orf59* | -0.12189 | -0.7328 | -1.8629 | -2.5764 | -0.91 |
| 18324 | *PRMT2* | 1.2299 | -1.6891 | -2.2594 | -2.7813 | -0.91 |
| 18325 | *COPG1* | -0.46628 | -0.81904 | -1.4353 | -1.5064 | -0.91 |
| 18326 | *ARHGEF38* | 0.965 | -1.843 | -1.8435 | -1.9131 | -0.91 |
| 18327 | *SLC6A20* | -0.56938 | -0.77255 | -1.3808 | -1.4376 | -0.91 |
| 18328 | *UNC45B* | 0.3523 | -1.3617 | -1.714 | -3.2519 | -0.91 |
| 18329 | *TNPO3* | -0.63946 | -0.97338 | -1.1108 | -4.2256 | -0.91 |
| 18330 | *DAB2IP* | -0.86494 | -0.89438 | -0.96572 | -1.8046 | -0.91 |
| 18331 | *MDGA2* | -0.00178 | -1.2694 | -1.4544 | -1.6649 | -0.91 |
| 18332 | *ATP5SL* | -0.46153 | -0.88465 | -1.3851 | -1.7462 | -0.91 |
| 18333 | *LYSMD4* | -0.48756 | -0.8897 | -1.3552 | -1.8157 | -0.91 |
| 18334 | *PROX1* | -0.56564 | -0.75073 | -1.4162 | -1.9186 | -0.91 |
| 18335 | *OR10H1* | -0.6636 | -0.97381 | -1.0952 | -2.011 | -0.91 |
| 18336 | *GPN2* | -0.55047 | -0.86358 | -1.3198 | -2.5513 | -0.91 |
| 18337 | *MRPL49* | -0.61249 | -0.76344 | -1.358 | -1.7743 | -0.91 |
| 18338 | *ARF1* | 0.13438 | -1.2884 | -1.5801 | -1.7578 | -0.91 |
| 18339 | *SLC2A9* | -0.45867 | -0.49403 | -1.7822 | -1.8707 | -0.91 |
| 18340 | *ZBTB14* | -0.38296 | -1.1223 | -1.2302 | -1.9175 | -0.91 |
| 18341 | *ACER3* | -0.5183 | -0.95105 | -1.2666 | -2.2732 | -0.91 |
| 18342 | *FIGNL1* | -0.33136 | -1.0297 | -1.3771 | -1.8435 | -0.91 |
| 18343 | *NBL1* | -0.56442 | -0.84042 | -1.3393 | -1.7209 | -0.91 |
| 18344 | *RILPL1* | -0.51713 | -0.64496 | -1.5824 | -2.2363 | -0.91 |
| 18345 | *MLN* | -0.08719 | -0.97333 | -1.6934 | -2.0134 | -0.92 |
| 18346 | *MAPKAPK3* | 0.096888 | -1.2935 | -1.5598 | -2.0545 | -0.92 |
| 18347 | *DNAJB1* | -0.06253 | -1.2202 | -1.4741 | -1.6128 | -0.92 |

| | | | | | | |
|---|---|---|---|---|---|---|
| 18348 | *IYD* | -0.17606 | -1.1925 | -1.3912 | -1.9172 | -0.92 |
| 18349 | *TIMM13* | -0.43626 | -0.78313 | -1.5408 | -1.6608 | -0.92 |
| 18350 | *ARHGAP1* | -0.55625 | -0.57669 | -1.6281 | -2.7303 | -0.92 |
| 18351 | *EXOSC4* | -0.61057 | -0.74715 | -1.4053 | -3.8502 | -0.92 |
| 18352 | *EIF2A* | 0.66522 | -1.1016 | -2.3282 | -2.6208 | -0.92 |
| 18353 | *ZNF20* | 0.53073 | -1.278 | -2.0181 | -2.7678 | -0.92 |
| 18354 | *EXOC1* | -0.30438 | -0.9225 | -1.539 | -1.7882 | -0.92 |
| 18355 | *HSD17B2* | -0.29291 | -1.1566 | -1.3183 | -1.6164 | -0.92 |
| 18356 | *CA1* | 0.1059 | -0.98366 | -1.8963 | -1.9811 | -0.92 |
| 18357 | *CSGALNACT2* | -0.77018 | -0.85739 | -1.1487 | -1.9338 | -0.93 |
| 18358 | *NDUFAF5* | -0.79813 | -0.80405 | -1.1742 | -1.2114 | -0.93 |
| 18359 | *GGACT* | 0.3775 | -1.2567 | -1.8975 | -1.9107 | -0.93 |
| 18360 | *WDR49* | -0.5911 | -0.60508 | -1.5806 | -2.1574 | -0.93 |
| 18361 | *GPRC5A* | -0.51549 | -0.66623 | -1.5959 | -1.6216 | -0.93 |
| 18362 | *OR1A2* | -0.1059 | -0.84345 | -1.8321 | -2.217 | -0.93 |
| 18363 | *RNF115* | -0.13576 | -1.2864 | -1.3643 | -1.438 | -0.93 |
| 18364 | *SDAD1* | -0.5206 | -0.54008 | -1.7263 | -1.9965 | -0.93 |
| 18365 | *FAM84A* | 0.60761 | -1.6885 | -1.707 | -2.2064 | -0.93 |
| 18366 | *ATP6V1G1* | 0.057611 | -0.4847 | -2.3623 | -3.4804 | -0.93 |
| 18367 | *RALGDS* | -0.04218 | -1.2985 | -1.4493 | -1.9936 | -0.93 |
| 18368 | *DLL3* | 0.37254 | -1.4287 | -1.7364 | -2.2432 | -0.93 |
| 18369 | *HINFP* | -0.06631 | -1.2418 | -1.4878 | -1.9965 | -0.93 |
| 18370 | *ADAMDEC1* | 0.22798 | -1.2884 | -1.7364 | -2.3197 | -0.93 |
| 18371 | *TBP* | -0.22428 | -1.223 | -1.3496 | -2.3389 | -0.93 |
| 18372 | *C8orf88* | 0.53251 | -1.386 | -1.9435 | -3.1836 | -0.93 |
| 18373 | *MYO1A* | 0.52375 | -1.5817 | -1.7397 | -1.8557 | -0.93 |
| 18374 | *ZNF830* | -0.13185 | -1.305 | -1.3627 | -1.4686 | -0.93 |
| 18375 | *LAGE3* | 0.11473 | -1.4325 | -1.4824 | -1.7538 | -0.93 |
| 18376 | *HMGB1* | 0.12155 | -1.4568 | -1.468 | -1.6313 | -0.93 |
| 18377 | *PDXDC1* | -0.52574 | -0.88971 | -1.3912 | -1.5436 | -0.94 |
| 18378 | *RARG* | -0.32355 | -1.154 | -1.3296 | -1.7726 | -0.94 |
| 18379 | *RUFY3* | -0.49565 | -0.87931 | -1.4325 | -1.8272 | -0.94 |
| 18380 | *VDAC3* | 0.43872 | -1.2687 | -1.9789 | -3.4532 | -0.94 |
| 18381 | *CD300C* | -0.33705 | -1.0918 | -1.3818 | -1.7322 | -0.94 |
| 18382 | *SNX27* | -0.16892 | -1.1002 | -1.5425 | -1.6014 | -0.94 |
| 18383 | *GAL* | 0.27373 | -1.3701 | -1.716 | -2.0953 | -0.94 |
| 18384 | *HLA-DRA* | -0.71016 | -0.75437 | -1.3506 | -1.7745 | -0.94 |
| 18385 | *RGS9BP* | -0.49045 | -1.1461 | -1.1793 | -2.4647 | -0.94 |
| 18386 | *CDADC1* | -0.59119 | -0.59856 | -1.6276 | -2.0548 | -0.94 |
| 18387 | *DUSP6* | 0.33475 | -1.3465 | -1.8066 | -2.5725 | -0.94 |
| 18388 | *KCNC4* | -0.38053 | -1.1856 | -1.2546 | -1.4684 | -0.94 |
| 18389 | *SEPSECS* | -0.17058 | -0.23846 | -2.412 | -2.4933 | -0.94 |
| 18390 | *ELOVL1* | 0.92769 | -1.7554 | -1.9936 | -2.0949 | -0.94 |
| 18391 | *SPSB1* | -0.02722 | -0.99768 | -1.7974 | -1.8462 | -0.94 |
| 18392 | *MUC5AC* | -0.29905 | -1.0443 | -1.4805 | -1.9196 | -0.94 |
| 18393 | *YAE1D1* | -0.3316 | -0.75038 | -1.742 | -2.1612 | -0.94 |

| | | | | | | |
|---|---|---|---|---|---|---|
| 18394 | *CLVS2* | -0.73565 | -1.0386 | -1.0529 | -2.092 | -0.94 |
| 18395 | *GCN1L1* | -0.17285 | -1.0467 | -1.6092 | -2.8963 | -0.94 |
| 18396 | *C7orf61* | -0.76516 | -0.98907 | -1.0797 | -2.2285 | -0.94 |
| 18397 | *PLEKHM2* | -0.78098 | -0.82375 | -1.2297 | -1.4034 | -0.94 |
| 18398 | *GLB1L2* | -0.43818 | -1.0564 | -1.3399 | -1.8076 | -0.94 |
| 18399 | *HOXA7* | -0.69184 | -0.84542 | -1.3007 | -2.555 | -0.95 |
| 18400 | *ATP6V0D1* | -0.44353 | -0.80969 | -1.5852 | -1.8604 | -0.95 |
| 18401 | *EPHB3* | -0.38901 | -0.57824 | -1.8714 | -2.4078 | -0.95 |
| 18402 | *TMEM97* | 0.33639 | -1.4234 | -1.7526 | -2.1794 | -0.95 |
| 18403 | *FER1L5* | 0.034036 | -1.2676 | -1.6065 | -2.1426 | -0.95 |
| 18404 | *PLS1* | 0.004468 | -0.90206 | -1.9474 | -2.0223 | -0.95 |
| 18405 | *DAD1* | -0.48085 | -0.76882 | -1.5962 | -2.1239 | -0.95 |
| 18406 | *COX15* | -0.72814 | -1.0564 | -1.0623 | -1.4142 | -0.95 |
| 18407 | *NUF2* | -0.39636 | -1.1212 | -1.3295 | -1.5082 | -0.95 |
| 18408 | *RPS6KC1* | -0.63357 | -0.68303 | -1.5306 | -2.0256 | -0.95 |
| 18409 | *NUBP1* | 0.30938 | -1.5408 | -1.6164 | -2.3846 | -0.95 |
| 18410 | *SRGAP3* | -0.18614 | -1.2198 | -1.4423 | -1.7577 | -0.95 |
| 18411 | *TECTB* | -0.79319 | -0.88503 | -1.1752 | -1.3078 | -0.95 |
| 18412 | *ARIH2OS* | 0.14155 | -0.33705 | -2.6609 | -3.2005 | -0.95 |
| 18413 | *GMPS* | 0.068131 | -1.4467 | -1.4789 | -2.1046 | -0.95 |
| 18414 | *KLHL38* | 0.258 | -1.4536 | -1.6635 | -2.5571 | -0.95 |
| 18415 | *PRDM11* | 0.16442 | -1.0248 | -2.0012 | -2.4642 | -0.95 |
| 18416 | *AQP1* | -0.21701 | -1.2855 | -1.3591 | -2.1444 | -0.95 |
| 18417 | *EIF3D* | -0.19422 | -1.3332 | -1.3349 | -1.5307 | -0.95 |
| 18418 | *PSMD6* | 0.41894 | -1.2214 | -2.0601 | -2.2139 | -0.95 |
| 18419 | *EIF2AK4* | -0.75489 | -0.98468 | -1.1272 | -1.3823 | -0.96 |
| 18420 | *AP2S1* | -0.79367 | -0.87989 | -1.1973 | -1.5589 | -0.96 |
| 18421 | *C9orf156* | -0.32871 | -1.1524 | -1.3923 | -1.4782 | -0.96 |
| 18422 | *NUDC* | 0.82411 | -1.3416 | -2.3577 | -5.7035 | -0.96 |
| 18423 | *SYNGR3* | -0.39017 | -1.175 | -1.3104 | -2.412 | -0.96 |
| 18424 | *TIGD7* | -0.29327 | -0.99834 | -1.5861 | -1.9302 | -0.96 |
| 18425 | *TRIM47* | -0.00379 | -1.3915 | -1.4859 | -2.268 | -0.96 |
| 18426 | *DSPP* | -0.46089 | -0.79187 | -1.6288 | -2.0059 | -0.96 |
| 18427 | *HSPA6* | -0.22131 | -1.2936 | -1.3668 | -1.8816 | -0.96 |
| 18428 | *EME2* | -0.48659 | -1.1845 | -1.2114 | -3.7236 | -0.96 |
| 18429 | *MYLK2* | -0.86223 | -0.9857 | -1.0354 | -1.047 | -0.96 |
| 18430 | *ITGAL* | 0.10699 | -1.011 | -1.9794 | -3.3443 | -0.96 |
| 18431 | *RALY* | -0.15766 | -1.2876 | -1.4388 | -3.596 | -0.96 |
| 18432 | *KCNC1* | 0.22531 | -1.53 | -1.5795 | -1.8219 | -0.96 |
| 18433 | *AKR1B1* | 0.2678 | -1.3743 | -1.7806 | -2.4244 | -0.96 |
| 18434 | *CTAGE4* | -0.76771 | -0.76832 | -1.3513 | -1.6365 | -0.96 |
| 18435 | *TMEM184B* | 0.31536 | -1.1218 | -2.0818 | -2.312 | -0.96 |
| 18436 | *RSRC2* | -0.1496 | -1.3132 | -1.4287 | -1.5289 | -0.96 |
| 18437 | *MRPS16* | -0.41965 | -0.90003 | -1.5737 | -1.5792 | -0.96 |
| 18438 | *DMAP1* | -0.0394 | -1.2481 | -1.6066 | -1.9768 | -0.96 |
| 18439 | *VAMP8* | -0.39386 | -1.0589 | -1.4415 | -2.0717 | -0.96 |

| | | | | | | |
|---|---|---|---|---|---|---|
| 18440 | *FSIP2* | -0.59025 | -1.0232 | -1.2837 | -1.6586 | -0.97 |
| 18441 | *SERINC1* | -0.41087 | -0.72227 | -1.7652 | -1.7662 | -0.97 |
| 18442 | *EIF4A3* | 0.52363 | -1.451 | -1.9734 | -2.4873 | -0.97 |
| 18443 | *HPSE* | -0.74011 | -1.0097 | -1.151 | -1.3132 | -0.97 |
| 18444 | *UBE2C* | -0.50748 | -1.1448 | -1.25 | -2.0267 | -0.97 |
| 18445 | *TAF4B* | -0.43572 | -1.0796 | -1.3878 | -1.7179 | -0.97 |
| 18446 | *RLN3* | -0.18185 | -0.92235 | -1.7999 | -2.1239 | -0.97 |
| 18447 | *SMEK1* | -0.02584 | -1.3496 | -1.5304 | -1.5949 | -0.97 |
| 18448 | *AQR* | -0.19225 | -0.98842 | -1.7268 | -2.3509 | -0.97 |
| 18449 | *LTBP1* | -0.00827 | -1.3668 | -1.5357 | -1.6587 | -0.97 |
| 18450 | *FRAT2* | -0.33363 | -1.0585 | -1.5195 | -1.545 | -0.97 |
| 18451 | *CCDC28B* | -0.44986 | -1.2293 | -1.2347 | -1.7718 | -0.97 |
| 18452 | *PMEL* | -0.59225 | -0.91823 | -1.4036 | -2.1205 | -0.97 |
| 18453 | *TRIM64B* | -0.63328 | -1.0641 | -1.2169 | -1.7679 | -0.97 |
| 18454 | *MARK3* | -0.46716 | -1.116 | -1.3316 | -2.2856 | -0.97 |
| 18455 | *ASPRV1* | -0.51161 | -0.8191 | -1.5854 | -1.6826 | -0.97 |
| 18456 | *INTS10* | -0.23203 | -0.68727 | -2.0016 | -2.508 | -0.97 |
| 18457 | *SSX5* | 0.6844 | -1.151 | -2.4595 | -2.8463 | -0.98 |
| 18458 | *PROK1* | -0.25407 | -1.2489 | -1.4234 | -2.0817 | -0.98 |
| 18459 | *TSR2* | 0.45815 | -1.1676 | -2.2171 | -2.4562 | -0.98 |
| 18460 | *NOLC1* | 0.11095 | -1.338 | -1.701 | -2.3573 | -0.98 |
| 18461 | *CRH* | -0.59008 | -0.99644 | -1.3429 | -1.7753 | -0.98 |
| 18462 | *ZNF267* | 0.52847 | -1.6524 | -1.8076 | -1.8508 | -0.98 |
| 18463 | *SETD1A* | -0.52876 | -1.1357 | -1.2694 | -2.4422 | -0.98 |
| 18464 | *NACA* | -0.27263 | -1.0889 | -1.5727 | -2.2732 | -0.98 |
| 18465 | *EIF4ENIF1* | -0.03537 | -1.392 | -1.5101 | -1.6368 | -0.98 |
| 18466 | *CMPK1* | -0.25941 | -0.8638 | -1.8157 | -1.9686 | -0.98 |
| 18467 | *SLC6A11* | -0.16249 | -0.46731 | -2.3145 | -2.6024 | -0.98 |
| 18468 | *TJP2* | -0.2507 | -0.78313 | -1.9107 | -2.1984 | -0.98 |
| 18469 | *CEP89* | -0.24598 | -1.3053 | -1.3946 | -1.6356 | -0.98 |
| 18470 | *C2orf16* | -0.23528 | -0.80946 | -1.9068 | -2.672 | -0.98 |
| 18471 | *USP20* | -0.73061 | -0.97149 | -1.2497 | -1.7804 | -0.98 |
| 18472 | *PNO1* | -0.09774 | -1.2761 | -1.5782 | -1.6867 | -0.98 |
| 18473 | *LOC388436* | -0.28357 | -1.1754 | -1.4935 | -1.8725 | -0.98 |
| 18474 | *ORM1* | -0.05212 | -1.3542 | -1.5467 | -2.7502 | -0.98 |
| 18475 | *GADD45G* | -0.3644 | -1.2127 | -1.3771 | -1.7235 | -0.98 |
| 18476 | *COX11* | -0.35805 | -0.93806 | -1.6594 | -1.9556 | -0.99 |
| 18477 | *GPR45* | -0.55477 | -0.98129 | -1.4206 | -1.6984 | -0.99 |
| 18478 | *KIR3DL2* | 0.056445 | -1.2867 | -1.7266 | -1.962 | -0.99 |
| 18479 | *ZNF574* | -0.72798 | -1.0162 | -1.2135 | -2.1919 | -0.99 |
| 18480 | *CRLF2* | -0.71669 | -1.1095 | -1.1415 | -1.3711 | -0.99 |
| 18481 | *CBLN4* | -0.33705 | -1.2646 | -1.3675 | -2.2456 | -0.99 |
| 18482 | *NT5C3A* | -0.21148 | -1.2195 | -1.5383 | -1.8952 | -0.99 |
| 18483 | *KLHDC4* | -0.32719 | -0.93112 | -1.7115 | -2.4741 | -0.99 |
| 18484 | *INSIG2* | -0.48842 | -0.98668 | -1.4962 | -2.1038 | -0.99 |
| 18485 | *NAPG* | 1.0414 | -1.988 | -2.0249 | -2.2669 | -0.99 |

| | | | | | | |
|---|---|---|---|---|---|---|
| 18486 | *SPATA5L1* | -0.83912 | -1.0131 | -1.1212 | -2.101 | -0.99 |
| 18487 | *FAM180A* | 0.34989 | -1.6047 | -1.7247 | -2.0042 | -0.99 |
| 18488 | *CDH7* | -0.69865 | -0.9442 | -1.3371 | -2.337 | -0.99 |
| 18489 | *GINS3* | -0.6029 | -1.0359 | -1.3457 | -2.0383 | -0.99 |
| 18490 | *DPP8* | 0.79651 | -1.855 | -1.9269 | -2.5153 | -1.00 |
| 18491 | *GFER* | -0.35972 | -0.75665 | -1.8692 | -2.0436 | -1.00 |
| 18492 | *IL24* | -0.84446 | -1.0231 | -1.1189 | -1.1685 | -1.00 |
| 18493 | *PCGF3* | -0.86494 | -0.92425 | -1.2009 | -1.3353 | -1.00 |
| 18494 | *NDUFB9* | -0.40125 | -1.1432 | -1.4485 | -2.1796 | -1.00 |
| 18495 | *TUBGCP5* | -0.59229 | -0.89866 | -1.502 | -1.7643 | -1.00 |
| 18496 | *LRRC18* | -0.30496 | -1.2719 | -1.4168 | -2.0851 | -1.00 |
| 18497 | *TP53INP2* | -0.6736 | -0.8737 | -1.45 | -1.7164 | -1.00 |
| 18498 | *TFRC* | -0.68208 | -0.96776 | -1.3489 | -1.7179 | -1.00 |
| 18499 | *MAMDC4* | -0.44353 | -0.96779 | -1.5876 | -1.6339 | -1.00 |
| 18500 | *POLR2A* | -0.43634 | -1.25 | -1.3128 | -2.6188 | -1.00 |
| 18501 | *TMEM132A* | -0.73051 | -0.97768 | -1.2968 | -1.7074 | -1.00 |
| 18502 | *RPA3* | -0.29623 | -0.97381 | -1.7365 | -2.1859 | -1.00 |
| 18503 | *SMYD2* | -0.85931 | -0.90549 | -1.2471 | -1.6921 | -1.00 |
| 18504 | *SNRPD3* | -0.59886 | -0.7503 | -1.6635 | -2.4015 | -1.00 |
| 18505 | *ZNRD1* | -0.76055 | -0.99418 | -1.2588 | -1.8695 | -1.00 |
| 18506 | *RPL4* | -0.21123 | -1.2343 | -1.5684 | -2.5846 | -1.00 |
| 18507 | *ZNF460* | -0.63341 | -0.811 | -1.5704 | -1.8256 | -1.00 |
| 18508 | *SMG7* | -0.33169 | -1.3057 | -1.3785 | -2.005 | -1.01 |
| 18509 | *ZNF711* | 0.54647 | -1.7285 | -1.8341 | -2.4785 | -1.01 |
| 18510 | *CDC45* | -0.73673 | -0.97636 | -1.3033 | -1.4523 | -1.01 |
| 18511 | *SPTBN5* | -0.62613 | -1.071 | -1.3209 | -2.2165 | -1.01 |
| 18512 | *TH* | -0.1042 | -1.083 | -1.8318 | -2.2948 | -1.01 |
| 18513 | *FNDC5* | -0.49525 | -1.13 | -1.3946 | -1.7658 | -1.01 |
| 18514 | *C7orf55* | 0.26733 | -1.595 | -1.6926 | -2.3755 | -1.01 |
| 18515 | *HNRNPU* | -0.43951 | -1.1343 | -1.4473 | -1.7512 | -1.01 |
| 18516 | *CCDC101* | 0.42584 | -1.4355 | -2.0123 | -2.2428 | -1.01 |
| 18517 | *RREB1* | -0.7006 | -0.79225 | -1.5304 | -1.8338 | -1.01 |
| 18518 | *RPUSD1* | 0.38492 | -1.507 | -1.9013 | -1.9341 | -1.01 |
| 18519 | *GMPPB* | 0.043314 | -1.2676 | -1.8008 | -2.3086 | -1.01 |
| 18520 | *OTOR* | -0.42401 | -1.2567 | -1.345 | -1.4537 | -1.01 |
| 18521 | *PRPS2* | -0.83818 | -1.0604 | -1.1278 | -1.8938 | -1.01 |
| 18522 | *YARS2* | -0.44084 | -1.1978 | -1.3887 | -1.5831 | -1.01 |
| 18523 | *C8orf44* | 0.25292 | -1.1852 | -2.0953 | -2.6973 | -1.01 |
| 18524 | *ZP4* | -0.97393 | -0.99791 | -1.0565 | -1.5663 | -1.01 |
| 18525 | *ZNF559-ZNF177* | -0.16151 | -1.3457 | -1.5235 | -2.8327 | -1.01 |
| 18526 | *CCT5* | 0.48172 | -1.2676 | -2.2449 | -3.1273 | -1.01 |
| 18527 | *SZRD1* | -0.3785 | -1.1718 | -1.4811 | -1.5817 | -1.01 |
| 18528 | *SLC18B1* | -0.20924 | -0.83439 | -1.988 | -2.2528 | -1.01 |
| 18529 | *ANGPTL2* | -0.14166 | -1.3148 | -1.5781 | -1.912 | -1.01 |
| 18530 | *CNOT2* | -0.57825 | -1.153 | -1.3068 | -1.6601 | -1.01 |
| 18531 | *EPB41L5* | -0.33241 | -1.1518 | -1.5544 | -1.8975 | -1.01 |

| | | | | | | |
|---|---|---|---|---|---|---|
| 18532 | *GP2* | -0.6728 | -1.1289 | -1.237 | -1.7149 | -1.01 |
| 18533 | *MROH8* | -0.26439 | -1.0409 | -1.7338 | -2.6046 | -1.01 |
| 18534 | *LSM2* | -0.86358 | -1.0199 | -1.1564 | -1.5183 | -1.01 |
| 18535 | *IGBP1* | 0.47763 | -1.632 | -1.8869 | -2.6779 | -1.01 |
| 18536 | *TRIM66* | -0.39368 | -1.099 | -1.5491 | -1.6876 | -1.01 |
| 18537 | *CD80* | -0.45672 | -0.79225 | -1.7942 | -2.2888 | -1.01 |
| 18538 | *TRAF2* | -0.56251 | -0.78961 | -1.6921 | -1.8425 | -1.01 |
| 18539 | *TMC5* | -0.13189 | -1.3343 | -1.584 | -2.9909 | -1.02 |
| 18540 | *ARID4A* | -0.18925 | -1.0718 | -1.7945 | -2.0351 | -1.02 |
| 18541 | *RALGAPB* | 0.06603 | -1.426 | -1.6964 | -2.8166 | -1.02 |
| 18542 | *COG8* | -0.28487 | -1.1739 | -1.5978 | -2.312 | -1.02 |
| 18543 | *PLCB1* | -0.04824 | -1.3023 | -1.7134 | -2.1807 | -1.02 |
| 18544 | *PTRH1* | -0.50724 | -0.97346 | -1.5884 | -2.5125 | -1.02 |
| 18545 | *TMCO3* | 0.47329 | -1.1839 | -2.3585 | -3.3677 | -1.02 |
| 18546 | *H3F3A* | -0.44353 | -0.68934 | -1.9375 | -2.6157 | -1.02 |
| 18547 | *CUL4A* | -0.61358 | -1.1752 | -1.2867 | -1.828 | -1.03 |
| 18548 | *MTERF2* | 0.12888 | -1.498 | -1.7119 | -2.4709 | -1.03 |
| 18549 | *TONSL* | 0.046846 | -1.4233 | -1.7052 | -2.8051 | -1.03 |
| 18550 | *NSL1* | -0.09437 | -1.4123 | -1.5765 | -1.7349 | -1.03 |
| 18551 | *PTPRC* | 0.40329 | -1.4692 | -2.0197 | -2.3063 | -1.03 |
| 18552 | *NPPC* | -0.76055 | -1.0198 | -1.3053 | -1.4775 | -1.03 |
| 18553 | *MAGED4B* | -0.2747 | -0.588 | -2.224 | -2.3178 | -1.03 |
| 18554 | *ZNF671* | -0.4191 | -1.0509 | -1.6193 | -3.2688 | -1.03 |
| 18555 | *HDAC2* | -0.10661 | -1.4287 | -1.5588 | -2.5006 | -1.03 |
| 18556 | *RPS25* | 0.37299 | -1.3617 | -2.1066 | -2.1919 | -1.03 |
| 18557 | *NR2E3* | -0.30692 | -1.3128 | -1.4775 | -2.0895 | -1.03 |
| 18558 | *MYOZ3* | -0.72782 | -1.073 | -1.3039 | -1.6105 | -1.03 |
| 18559 | *CAPS* | -0.56926 | -1.1585 | -1.3795 | -1.4385 | -1.04 |
| 18560 | *FOXA1* | 0.1506 | -1.6098 | -1.6481 | -1.72 | -1.04 |
| 18561 | *BOLA2* | 0.09805 | -1.1354 | -2.071 | -2.1239 | -1.04 |
| 18562 | *ZNF880* | 0.060365 | -1.1958 | -1.9748 | -2.0661 | -1.04 |
| 18563 | *ETV4* | -0.03506 | -1.2118 | -1.8683 | -2.0632 | -1.04 |
| 18564 | *TRIM11* | -0.33191 | -1.0256 | -1.7577 | -3.582 | -1.04 |
| 18565 | *DLGAP1* | -0.0994 | -1.1214 | -1.8946 | -2.224 | -1.04 |
| 18566 | *DPF3* | -0.55513 | -1.2572 | -1.3033 | -1.6085 | -1.04 |
| 18567 | *UGT3A1* | -0.78591 | -0.90865 | -1.4218 | -1.5304 | -1.04 |
| 18568 | *OSBPL8* | -0.73823 | -0.93878 | -1.4416 | -3.1728 | -1.04 |
| 18569 | *PABPN1* | -0.40015 | -0.9532 | -1.7658 | -3.2023 | -1.04 |
| 18570 | *PDYN* | -0.44353 | -1.1492 | -1.5314 | -1.6157 | -1.04 |
| 18571 | *DPM3* | 0.18317 | -1.4364 | -1.8714 | -2.2165 | -1.04 |
| 18572 | *CCDC59* | -0.22803 | -1.2192 | -1.6799 | -1.7585 | -1.04 |
| 18573 | *MRPL20* | -0.42671 | -0.87673 | -1.8238 | -1.8611 | -1.04 |
| 18574 | *CCDC74A* | -0.74086 | -0.8489 | -1.5393 | -1.9844 | -1.04 |
| 18575 | *FAM231A* | -0.20775 | -1.3039 | -1.6182 | -1.8566 | -1.04 |
| 18576 | *BCDIN3D* | 0.73895 | -1.6709 | -2.2014 | -4.4273 | -1.04 |
| 18577 | *PDGFB* | -0.6579 | -0.93217 | -1.545 | -2.1802 | -1.05 |

| | | | | | | |
|---|---|---|---|---|---|---|
| 18578 | *TNFSF8* | -0.50676 | -1.2911 | -1.3393 | -1.5687 | -1.05 |
| 18579 | *TREM1* | -0.06833 | -1.505 | -1.5707 | -2.5677 | -1.05 |
| 18580 | *ASAP3* | -0.92427 | -1.0912 | -1.1287 | -1.7072 | -1.05 |
| 18581 | *GNL3L* | -0.73199 | -0.89127 | -1.5234 | -1.7638 | -1.05 |
| 18582 | *HCK* | -0.33754 | -1.1207 | -1.6891 | -1.7595 | -1.05 |
| 18583 | *MST1L* | -0.31011 | -0.93172 | -1.9073 | -3.0598 | -1.05 |
| 18584 | *SLC6A9* | -0.01901 | -1.4597 | -1.6709 | -2.0218 | -1.05 |
| 18585 | *SPDYA* | -0.00431 | -1.2756 | -1.8761 | -2.2607 | -1.05 |
| 18586 | *OR6X1* | -0.59572 | -1.0273 | -1.5375 | -1.7177 | -1.05 |
| 18587 | *PDCD7* | 0.066478 | -0.94965 | -2.2804 | -3.1051 | -1.05 |
| 18588 | *DRAP1* | -0.11265 | -1.3543 | -1.6968 | -2.005 | -1.05 |
| 18589 | *KRT19* | -0.18808 | -1.348 | -1.6288 | -1.9165 | -1.05 |
| 18590 | *PRPF18* | 0.086847 | -1.589 | -1.6644 | -2.0297 | -1.06 |
| 18591 | *ILKAP* | -0.70417 | -1.2281 | -1.2347 | -1.301 | -1.06 |
| 18592 | *CD300LG* | -0.74086 | -0.94052 | -1.4856 | -1.5121 | -1.06 |
| 18593 | *S1PR4* | -0.76371 | -1.1904 | -1.2156 | -2.817 | -1.06 |
| 18594 | *UTP11L* | -0.7614 | -1.1958 | -1.2145 | -1.5696 | -1.06 |
| 18595 | *MBLAC1* | 1.2549 | -1.3662 | -3.0614 | -3.182 | -1.06 |
| 18596 | *KCNN4* | -0.747 | -1.1495 | -1.2779 | -1.4473 | -1.06 |
| 18597 | *CDK5* | 0.5592 | -0.94759 | -2.7871 | -2.9236 | -1.06 |
| 18598 | *NDFIP1* | -0.75889 | -1.1739 | -1.2431 | -1.5874 | -1.06 |
| 18599 | *NIP7* | -0.76342 | -1.1995 | -1.2141 | -2.5007 | -1.06 |
| 18600 | *C19orf40* | -0.81319 | -1.1466 | -1.2175 | -1.8489 | -1.06 |
| 18601 | *WRB* | -0.57776 | -1.0158 | -1.5848 | -2.0476 | -1.06 |
| 18602 | *PREX1* | -0.60692 | -1.238 | -1.3361 | -1.359 | -1.06 |
| 18603 | *ABHD8* | 0.070816 | -1.371 | -1.8816 | -2.2264 | -1.06 |
| 18604 | *EIF2AK1* | -0.54688 | -0.95045 | -1.6877 | -2.1282 | -1.06 |
| 18605 | *HNRNPA1* | -0.60929 | -1.1297 | -1.4467 | -1.6255 | -1.06 |
| 18606 | *VCP* | -0.48654 | -1.0671 | -1.6323 | -2.1388 | -1.06 |
| 18607 | *WDR6* | -0.6536 | -1.1553 | -1.3785 | -1.3903 | -1.06 |
| 18608 | *RPL26L1* | -0.95893 | -1.1095 | -1.1214 | -1.9446 | -1.06 |
| 18609 | *RFC5* | -0.77789 | -1.0995 | -1.3198 | -2.8823 | -1.07 |
| 18610 | *CCDC88A* | 1.0186 | -1.6866 | -2.5292 | -2.7465 | -1.07 |
| 18611 | *CCT6A* | -0.11811 | -1.3686 | -1.7149 | -2.072 | -1.07 |
| 18612 | *LOC102724862* | 0.45098 | -1.4504 | -2.2044 | -2.3287 | -1.07 |
| 18613 | *HS3ST3B1* | 0.54628 | -1.7884 | -1.9622 | -2.6803 | -1.07 |
| 18614 | *SH3BGRL2* | 0.26813 | -1.1637 | -2.312 | -2.4312 | -1.07 |
| 18615 | *USP17L25* | -0.47447 | -1.3637 | -1.3743 | -1.398 | -1.07 |
| 18616 | *RPL24* | -0.10034 | -1.2486 | -1.8645 | -2.3729 | -1.07 |
| 18617 | *IFT57* | -0.6141 | -0.67691 | -1.9227 | -2.046 | -1.07 |
| 18618 | *DLD* | -1.0039 | -1.0826 | -1.1325 | -2.1798 | -1.07 |
| 18619 | *NXPH3* | -0.13602 | -1.4998 | -1.5848 | -2.3062 | -1.07 |
| 18620 | *GDF5* | -0.78199 | -1.1027 | -1.3364 | -1.545 | -1.07 |
| 18621 | *LRRC69* | -0.44136 | -0.90523 | -1.8761 | -2.0218 | -1.07 |
| 18622 | *WBP5* | 0.25931 | -1.6617 | -1.8212 | -2.1984 | -1.07 |
| 18623 | *NCR3LG1* | -0.35478 | -1.3224 | -1.547 | -1.5848 | -1.07 |

| | | | | | | |
|---|---|---|---|---|---|---|
| 18624 | *POC1B-GALNT4* | -0.48905 | -0.49016 | -2.2456 | -2.5551 | -1.07 |
| 18625 | *FAM217A* | 0.48077 | -1.6316 | -2.0752 | -2.5008 | -1.08 |
| 18626 | *RTN4R* | -0.83436 | -1.082 | -1.3104 | -1.4273 | -1.08 |
| 18627 | *PDE6B* | -0.66862 | -1.0029 | -1.5568 | -1.8256 | -1.08 |
| 18628 | *RPL7A* | 0.44691 | -1.7412 | -1.9358 | -2.1198 | -1.08 |
| 18629 | *RERE* | 0.67156 | -1.9186 | -1.984 | -2.3534 | -1.08 |
| 18630 | *CHIC2* | 0.046036 | -1.5039 | -1.7745 | -2.3461 | -1.08 |
| 18631 | *XYLT1* | 0.30066 | -1.4688 | -2.0653 | -2.5975 | -1.08 |
| 18632 | *CLDN9* | -0.06441 | -0.47818 | -2.6923 | -2.8976 | -1.08 |
| 18633 | *NCOR1* | -0.67702 | -0.98585 | -1.5744 | -2.1843 | -1.08 |
| 18634 | *GALK2* | -0.6799 | -1.1979 | -1.3684 | -2.1605 | -1.08 |
| 18635 | *RNF8* | -0.6133 | -1.0671 | -1.5671 | -1.7595 | -1.08 |
| 18636 | *PSCA* | -0.57588 | -0.77038 | -1.9013 | -2.0545 | -1.08 |
| 18637 | *SCYL1* | -0.73695 | -1.0044 | -1.5069 | -1.9794 | -1.08 |
| 18638 | *ZFC3H1* | -0.61946 | -1.2666 | -1.3627 | -1.5859 | -1.08 |
| 18639 | *OR2L2* | 0.19764 | -1.6763 | -1.7709 | -2.3696 | -1.08 |
| 18640 | *TP53I11* | -0.51319 | -0.7436 | -1.9938 | -2.0817 | -1.08 |
| 18641 | *DCDC1* | -0.57484 | -1.3282 | -1.3523 | -1.3666 | -1.09 |
| 18642 | *GTF2F1* | -0.73602 | -0.89741 | -1.6239 | -1.8575 | -1.09 |
| 18643 | *APOH* | 0.34764 | -1.7561 | -1.8557 | -2.2473 | -1.09 |
| 18644 | *SPATA32* | -0.70057 | -1.1312 | -1.434 | -2.1489 | -1.09 |
| 18645 | *ALG13* | -0.67801 | -1.2814 | -1.3098 | -1.7512 | -1.09 |
| 18646 | *TAGLN2* | -0.33213 | -0.97549 | -1.9624 | -6.2107 | -1.09 |
| 18647 | *MSMP* | -0.53234 | -1.2935 | -1.448 | -1.4507 | -1.09 |
| 18648 | *EXOSC9* | -0.76906 | -1.1028 | -1.4087 | -2.274 | -1.09 |
| 18649 | *ATR* | 0.67486 | -1.6526 | -2.3036 | -2.3871 | -1.09 |
| 18650 | *TBL2* | -0.94991 | -1.0974 | -1.2343 | -1.31 | -1.09 |
| 18651 | *CHMP6* | -0.88281 | -0.91944 | -1.4801 | -2.4193 | -1.09 |
| 18652 | *ANAPC15* | -0.97088 | -1.0758 | -1.2402 | -1.7905 | -1.10 |
| 18653 | *FAM49B* | -0.44353 | -0.99357 | -1.85 | -2.0197 | -1.10 |
| 18654 | *ABCA4* | -0.25893 | -0.55029 | -2.4784 | -2.5298 | -1.10 |
| 18655 | *EEF1A1* | -0.54734 | -1.2514 | -1.4893 | -1.5954 | -1.10 |
| 18656 | *GCNT4* | -0.56525 | -1.1126 | -1.6105 | -2.6062 | -1.10 |
| 18657 | *PYHIN1* | -0.44353 | -1.2277 | -1.6176 | -1.6268 | -1.10 |
| 18658 | *THOC3* | -0.24253 | -1.41 | -1.6371 | -1.7386 | -1.10 |
| 18659 | *GIMAP8* | -0.83773 | -0.95842 | -1.4935 | -2.0181 | -1.10 |
| 18660 | *HIST1H2AL* | 0.23572 | -1.4233 | -2.1046 | -2.8402 | -1.10 |
| 18661 | *TCTN3* | -0.44353 | -1.1138 | -1.7364 | -2.1984 | -1.10 |
| 18662 | *GSR* | -0.52552 | -0.97149 | -1.8038 | -1.8572 | -1.10 |
| 18663 | *TOMM22* | -0.03905 | -1.4469 | -1.8157 | -2.0952 | -1.10 |
| 18664 | *INTS7* | -0.73577 | -0.87898 | -1.6952 | -2.3086 | -1.10 |
| 18665 | *RPL15* | -0.87109 | -0.94287 | -1.4971 | -2.0008 | -1.10 |
| 18666 | *SMARCAL1* | -0.38131 | -1.1874 | -1.7462 | -2.7042 | -1.10 |
| 18667 | *CYP2A6* | -0.6568 | -1.0099 | -1.6524 | -2.2951 | -1.11 |
| 18668 | *SQRDL* | -0.67853 | -1.0793 | -1.5635 | -2.5243 | -1.11 |
| 18669 | *MRPL4* | -0.54537 | -1.1387 | -1.6373 | -1.8404 | -1.11 |

| | | | | | | |
|---|---|---|---|---|---|---|
| 18670 | *SNRNP200* | -0.96201 | -0.97595 | -1.386 | -1.9041 | -1.11 |
| 18671 | *MFAP1* | -0.05613 | -1.5294 | -1.742 | -2.1187 | -1.11 |
| 18672 | *VMP1* | -0.03972 | -1.1325 | -2.1572 | -2.1908 | -1.11 |
| 18673 | *SRSF2* | -0.24984 | -1.3294 | -1.7512 | -1.8076 | -1.11 |
| 18674 | *VPS33B* | -0.60754 | -1.205 | -1.5195 | -2.537 | -1.11 |
| 18675 | *POLE3* | -0.55281 | -0.88272 | -1.8968 | -1.9023 | -1.11 |
| 18676 | *KLRC1* | -0.63591 | -1.1571 | -1.5401 | -2.2955 | -1.11 |
| 18677 | *POTEF* | -0.83656 | -0.99508 | -1.5105 | -1.761 | -1.11 |
| 18678 | *C12orf75* | -0.85194 | -0.91283 | -1.5781 | -2.796 | -1.11 |
| 18679 | *SACS* | -0.67573 | -1.1967 | -1.4741 | -2.428 | -1.12 |
| 18680 | *MLEC* | -0.01367 | -1.2071 | -2.1264 | -2.4696 | -1.12 |
| 18681 | *ARPC5* | -0.92329 | -0.95775 | -1.4683 | -1.7266 | -1.12 |
| 18682 | *PI4KA* | 0.058695 | -1.6664 | -1.7422 | -2.6859 | -1.12 |
| 18683 | *B3GNT3* | -0.15161 | -1.4233 | -1.7766 | -2.7277 | -1.12 |
| 18684 | *GALC* | -0.67458 | -1.2968 | -1.3818 | -2.4481 | -1.12 |
| 18685 | *EFCC1* | -0.71717 | -0.93725 | -1.6989 | -1.7845 | -1.12 |
| 18686 | *HRG* | -0.92486 | -1.0699 | -1.3597 | -1.6561 | -1.12 |
| 18687 | *ST3GAL1* | 0.50844 | -1.5663 | -2.2972 | -2.4331 | -1.12 |
| 18688 | *ZBP1* | -1.0036 | -1.1441 | -1.2084 | -1.6411 | -1.12 |
| 18689 | *NMUR1* | -0.31654 | -1.4684 | -1.5781 | -1.9459 | -1.12 |
| 18690 | *RNF39* | -0.44353 | -1.2977 | -1.6288 | -1.7994 | -1.12 |
| 18691 | *ABLIM1* | -1.0704 | -1.0806 | -1.2195 | -1.3343 | -1.12 |
| 18692 | *METTL5* | -0.40641 | -1.4361 | -1.53 | -1.5941 | -1.12 |
| 18693 | *F8A3* | -0.95321 | -1.1105 | -1.3111 | -1.3287 | -1.12 |
| 18694 | *PLK1* | -0.73961 | -1.1714 | -1.468 | -1.9624 | -1.13 |
| 18695 | *SPDYE2* | -0.45717 | -1.3852 | -1.5393 | -1.6153 | -1.13 |
| 18696 | *SCN3A* | 0.046808 | -1.6731 | -1.7575 | -2.4709 | -1.13 |
| 18697 | *ATP6V1F* | -0.70442 | -0.84763 | -1.8341 | -2.6923 | -1.13 |
| 18698 | *GRM4* | -1.0857 | -1.1053 | -1.1977 | -1.2895 | -1.13 |
| 18699 | *ZNF792* | -0.93525 | -1.065 | -1.3892 | -1.9686 | -1.13 |
| 18700 | *TMLHE* | -0.0923 | -1.6158 | -1.6836 | -2.0146 | -1.13 |
| 18701 | *DPH3* | -0.60301 | -0.77037 | -2.0185 | -2.2943 | -1.13 |
| 18702 | *TAB1* | 0.17894 | -1.7552 | -1.8157 | -1.9172 | -1.13 |
| 18703 | *POLL* | -0.95017 | -1.174 | -1.2691 | -1.7829 | -1.13 |
| 18704 | *YIPF2* | -0.23176 | -0.85018 | -2.312 | -2.5255 | -1.13 |
| 18705 | *HMHB1* | -0.74932 | -1.0704 | -1.5757 | -2.0351 | -1.13 |
| 18706 | *USP17L18* | -0.84138 | -1.2451 | -1.3097 | -2.1983 | -1.13 |
| 18707 | *MARCKS* | 0.24592 | -1.7865 | -1.8557 | -2.2866 | -1.13 |
| 18708 | *FAM218A* | -1.0564 | -1.0782 | -1.2623 | -1.3878 | -1.13 |
| 18709 | *CEP97* | -0.85799 | -1.0221 | -1.517 | -2.3888 | -1.13 |
| 18710 | *THOC2* | -1.0427 | -1.1486 | -1.2071 | -1.565 | -1.13 |
| 18711 | *UFC1* | 0.91834 | -1.7056 | -2.6113 | -3.596 | -1.13 |
| 18712 | *PCDHB16* | -0.87128 | -1.154 | -1.3743 | -1.855 | -1.13 |
| 18713 | *PPIL2* | -0.25603 | -1.4689 | -1.6748 | -2.0061 | -1.13 |
| 18714 | *HBQ1* | -0.84841 | -0.97595 | -1.5765 | -1.9708 | -1.13 |
| 18715 | *XRCC5* | 0.030088 | -1.6855 | -1.7459 | -1.8157 | -1.13 |

| | | | | | | |
|---|---|---|---|---|---|---|
| 18716 | *C1orf216* | -0.53529 | -1.3209 | -1.547 | -2.8506 | -1.13 |
| 18717 | *PRELP* | -0.93764 | -1.0726 | -1.3961 | -2.1873 | -1.14 |
| 18718 | *ATAD2B* | 0.68042 | -1.4564 | -2.6308 | -2.7962 | -1.14 |
| 18719 | *RPL5* | -0.74719 | -1.1479 | -1.5189 | -1.5258 | -1.14 |
| 18720 | *SF3A2* | -1.046 | -1.1442 | -1.226 | -1.583 | -1.14 |
| 18721 | *PDCD11* | -0.26273 | -1.1927 | -1.9622 | -2.0483 | -1.14 |
| 18722 | *H2AFB2* | -0.64076 | -1.2817 | -1.4969 | -1.9227 | -1.14 |
| 18723 | *PWP2* | -0.69399 | -1.1222 | -1.6034 | -2.2301 | -1.14 |
| 18724 | *WNT3* | -0.23418 | -1.5524 | -1.6339 | -3.4438 | -1.14 |
| 18725 | *DONSON* | -0.7439 | -1.3026 | -1.3743 | -1.4234 | -1.14 |
| 18726 | *AUP1* | -0.04646 | -1.4141 | -1.9622 | -2.2951 | -1.14 |
| 18727 | *GPIHBP1* | -0.29886 | -1.4016 | -1.7235 | -2.5184 | -1.14 |
| 18728 | *TMEM82* | -0.8139 | -1.2973 | -1.3132 | -1.6635 | -1.14 |
| 18729 | *CMC1* | 0.28263 | -1.2079 | -2.5007 | -3.0644 | -1.14 |
| 18730 | *ADAM33* | -0.14736 | -1.545 | -1.7363 | -1.7829 | -1.14 |
| 18731 | *PPIAL4G* | -0.45707 | -1.4234 | -1.5494 | -2.0363 | -1.14 |
| 18732 | *NXN* | -1.0029 | -1.1317 | -1.3085 | -1.8398 | -1.15 |
| 18733 | *C2orf47* | -0.40107 | -1.4926 | -1.55 | -2.017 | -1.15 |
| 18734 | *DUSP13* | 0.10318 | -1.3756 | -2.1739 | -2.7689 | -1.15 |
| 18735 | *ADAMTS20* | -0.61675 | -0.95545 | -1.8751 | -3.096 | -1.15 |
| 18736 | *FAM13A* | -0.64278 | -0.98907 | -1.8157 | -2.9412 | -1.15 |
| 18737 | *TRRAP* | -0.73419 | -1.3104 | -1.4056 | -1.9321 | -1.15 |
| 18738 | *TTK* | -0.98026 | -1.0947 | -1.3769 | -2.1143 | -1.15 |
| 18739 | *MYCBP2* | -0.97499 | -1.0107 | -1.4684 | -2.3342 | -1.15 |
| 18740 | *TIMELESS* | -0.37323 | -1.539 | -1.5425 | -2.1227 | -1.15 |
| 18741 | *BCAM* | -0.76352 | -1.274 | -1.4197 | -1.4213 | -1.15 |
| 18742 | *ALG11* | -0.32117 | -1.3104 | -1.8257 | -2.0042 | -1.15 |
| 18743 | *MMP9* | -0.90635 | -0.95717 | -1.5959 | -1.6682 | -1.15 |
| 18744 | *AHCY* | -1.0359 | -1.1461 | -1.2786 | -2.203 | -1.15 |
| 18745 | *SOD1* | -0.74638 | -0.95423 | -1.7615 | -2.7546 | -1.15 |
| 18746 | *FJX1* | -0.15148 | -1.1944 | -2.1224 | -2.6376 | -1.16 |
| 18747 | *DICER1* | 0.28451 | -1.4275 | -2.3287 | -2.5934 | -1.16 |
| 18748 | *ME3* | -0.72882 | -1.311 | -1.4359 | -1.7215 | -1.16 |
| 18749 | *RPF2* | 0.03502 | -1.2249 | -2.2888 | -2.6308 | -1.16 |
| 18750 | *MAF* | -0.48415 | -1.1879 | -1.8087 | -2.4454 | -1.16 |
| 18751 | *MTRNR2L6* | -0.7288 | -1.3399 | -1.4142 | -2.9834 | -1.16 |
| 18752 | *TBC1D5* | -0.60427 | -1.2793 | -1.6007 | -1.8879 | -1.16 |
| 18753 | *NRBP1* | -0.19996 | -1.2643 | -2.0203 | -2.4013 | -1.16 |
| 18754 | *APCDD1L* | 0.29458 | -1.7667 | -2.0126 | -2.095 | -1.16 |
| 18755 | *CATSPER4* | -0.69199 | -1.0944 | -1.6988 | -2.5754 | -1.16 |
| 18756 | *TBC1D13* | -0.64571 | -1.2277 | -1.6128 | -3.2172 | -1.16 |
| 18757 | *LMOD3* | -0.65435 | -0.97667 | -1.8561 | -1.8591 | -1.16 |
| 18758 | *PRPF8* | 0.31764 | -1.8363 | -1.9699 | -2.5655 | -1.16 |
| 18759 | *FAM133B* | -0.2732 | -1.5467 | -1.6707 | -2.2329 | -1.16 |
| 18760 | *NMNAT1* | -0.36057 | -1.3132 | -1.8212 | -2.1984 | -1.16 |
| 18761 | *ARL10* | -0.52182 | -0.93975 | -2.0368 | -2.0398 | -1.17 |

| | | | | | | |
|---|---|---|---|---|---|---|
| 18762 | *COL20A1* | -1.0501 | -1.2135 | -1.2357 | -1.7935 | -1.17 |
| 18763 | *NPIPB3* | -0.89828 | -1.197 | -1.4054 | -1.6858 | -1.17 |
| 18764 | *RBBP4* | -0.99644 | -1.0995 | -1.4094 | -2.1392 | -1.17 |
| 18765 | *PEBP1* | -0.41101 | -0.92186 | -2.1732 | -2.7277 | -1.17 |
| 18766 | *MTX3* | -0.64398 | -1.2958 | -1.5726 | -1.8975 | -1.17 |
| 18767 | *ATG7* | -0.49336 | -1.3161 | -1.7068 | -1.8382 | -1.17 |
| 18768 | *FKBP5* | -0.59763 | -1.2146 | -1.7044 | -3.1921 | -1.17 |
| 18769 | *PIK3R4* | -0.59008 | -0.59886 | -2.3287 | -2.8702 | -1.17 |
| 18770 | *TMEM61* | -0.36948 | -1.4852 | -1.6649 | -1.761 | -1.17 |
| 18771 | *ARAP3* | -0.43472 | -1.0433 | -2.0452 | -2.2866 | -1.17 |
| 18772 | *CEP78* | -0.85296 | -1.2623 | -1.4103 | -2.1046 | -1.18 |
| 18773 | *HAUS4* | -0.40646 | -1.544 | -1.5757 | -2.2845 | -1.18 |
| 18774 | *SDHA* | -0.68374 | -1.0475 | -1.8087 | -1.988 | -1.18 |
| 18775 | *SMIM10L1* | -0.82311 | -1.2646 | -1.4544 | -3.8779 | -1.18 |
| 18776 | *DDX1* | -0.98864 | -1.1437 | -1.4125 | -1.4856 | -1.18 |
| 18777 | *NDUFAF1* | -0.96661 | -1.2572 | -1.3249 | -2.7277 | -1.18 |
| 18778 | *TUBA1B* | -0.27423 | -1.4361 | -1.8393 | -2.1699 | -1.18 |
| 18779 | *PCSK2* | -0.46711 | -1.5031 | -1.5795 | -1.7399 | -1.18 |
| 18780 | *GRB2* | -0.00331 | -1.479 | -2.0682 | -2.4935 | -1.18 |
| 18781 | *MBD3L2* | -0.19423 | -1.677 | -1.6822 | -1.7004 | -1.18 |
| 18782 | *SLX1B* | -0.75053 | -1.3009 | -1.5099 | -1.7263 | -1.19 |
| 18783 | *GSPT1* | -0.48821 | -1.4102 | -1.6648 | -2.6376 | -1.19 |
| 18784 | *GALR3* | -0.9703 | -1.1898 | -1.4036 | -1.7235 | -1.19 |
| 18785 | *SF3A1* | -0.95321 | -1.019 | -1.5962 | -2.0452 | -1.19 |
| 18786 | *FAM162B* | -0.92427 | -1.2655 | -1.379 | -1.5765 | -1.19 |
| 18787 | *TATDN2* | -0.69502 | -0.9997 | -1.8792 | -2.4394 | -1.19 |
| 18788 | *MIS18BP1* | -0.21532 | -1.2961 | -2.0629 | -2.4696 | -1.19 |
| 18789 | *UCP3* | -0.5084 | -1.5073 | -1.5597 | -2.7393 | -1.19 |
| 18790 | *CDRT15L2* | -0.65557 | -1.1665 | -1.7567 | -2.9354 | -1.19 |
| 18791 | *RFC1* | -0.94255 | -0.97768 | -1.6617 | -2.0995 | -1.19 |
| 18792 | *HDC* | -0.27961 | -1.4834 | -1.8199 | -2.716 | -1.19 |
| 18793 | *NKIRAS2* | -0.19467 | -1.3002 | -2.0908 | -2.3461 | -1.20 |
| 18794 | *C9orf9* | -0.73823 | -1.12 | -1.729 | -2.1796 | -1.20 |
| 18795 | *CAPNS2* | -0.86047 | -1.1429 | -1.5856 | -1.6526 | -1.20 |
| 18796 | *XYLT2* | -0.05348 | -0.93293 | -2.6075 | -4.2681 | -1.20 |
| 18797 | *CDR2L* | -1.0134 | -1.0145 | -1.5661 | -1.6731 | -1.20 |
| 18798 | *SETD8* | -0.51373 | -0.92629 | -2.1566 | -2.5478 | -1.20 |
| 18799 | *DHX37* | -0.85388 | -0.91807 | -1.8282 | -2.163 | -1.20 |
| 18800 | *TAF6* | -0.06667 | -1.5304 | -2.0042 | -2.8071 | -1.20 |
| 18801 | *ZNF785* | -0.24887 | -1.5795 | -1.7745 | -2.3615 | -1.20 |
| 18802 | *MANBAL* | -0.83184 | -1.0765 | -1.6964 | -2.4268 | -1.20 |
| 18803 | *ITGB7* | -0.47107 | -1.5508 | -1.589 | -1.6826 | -1.20 |
| 18804 | *CLPB* | -0.56411 | -1.0019 | -2.0466 | -2.3729 | -1.20 |
| 18805 | *CCT4* | -0.74671 | -1.0485 | -1.8178 | -1.8376 | -1.20 |
| 18806 | *SLMO2* | -0.31852 | -1.522 | -1.7734 | -1.8908 | -1.20 |
| 18807 | *TLX1* | -0.83276 | -1.2316 | -1.5502 | -1.5719 | -1.20 |

| | | | | | | |
|---|---|---|---|---|---|---|
| 18808 | *SAR1B* | -0.40897 | -1.4788 | -1.7306 | -2.412 | -1.21 |
| 18809 | *CLEC4F* | -0.32621 | -1.4584 | -1.8366 | -2.5157 | -1.21 |
| 18810 | *JADE2* | -0.00049 | -1.2935 | -2.3282 | -2.7546 | -1.21 |
| 18811 | *DDX47* | -0.57136 | -1.5235 | -1.5289 | -2.7546 | -1.21 |
| 18812 | *C14orf80* | -0.79453 | -1.4108 | -1.4208 | -1.5386 | -1.21 |
| 18813 | *PHC2* | -1.0564 | -1.2662 | -1.3053 | -1.4544 | -1.21 |
| 18814 | *ARMCX2* | -0.25163 | -1.4335 | -1.9431 | -2.1407 | -1.21 |
| 18815 | *CDK5RAP3* | -1.1169 | -1.1587 | -1.356 | -2.712 | -1.21 |
| 18816 | *PPIAL4D* | -0.68518 | -0.93368 | -2.0146 | -2.2811 | -1.21 |
| 18817 | *RHOD* | -0.28704 | -1.3337 | -2.0142 | -2.0436 | -1.21 |
| 18818 | *IGF2* | -0.90508 | -1.0922 | -1.6527 | -1.8038 | -1.22 |
| 18819 | *OR51G2* | 0.32028 | -1.1991 | -2.7736 | -2.8807 | -1.22 |
| 18820 | *MAP3K9* | -1.0481 | -1.1944 | -1.4109 | -1.4467 | -1.22 |
| 18821 | *PIGF* | -0.85296 | -1.3718 | -1.4312 | -1.72 | -1.22 |
| 18822 | *REEP2* | -0.71626 | -1.4087 | -1.5393 | -2.0895 | -1.22 |
| 18823 | *MDM1* | -0.61663 | -1.106 | -1.9513 | -3.6868 | -1.22 |
| 18824 | *KCNK15* | -0.73911 | -1.3496 | -1.5948 | -1.984 | -1.23 |
| 18825 | *NR2C2AP* | -0.54409 | -1.3836 | -1.7613 | -2.6635 | -1.23 |
| 18826 | *PSMD1* | -0.73155 | -1.095 | -1.8645 | -1.9163 | -1.23 |
| 18827 | *IFIT1B* | -1.0599 | -1.1756 | -1.4566 | -2.4312 | -1.23 |
| 18828 | *FPR1* | -0.21054 | -0.54391 | -2.9412 | -3.0896 | -1.23 |
| 18829 | *LRTOMT* | -0.51304 | -1.2163 | -1.9748 | -3.0028 | -1.23 |
| 18830 | *NSUN4* | -0.67906 | -1.0096 | -2.0161 | -2.2117 | -1.23 |
| 18831 | *ATRAID* | -0.7679 | -1.1747 | -1.7658 | -2.136 | -1.24 |
| 18832 | *SRSF7* | -1.0635 | -1.122 | -1.5264 | -1.6112 | -1.24 |
| 18833 | *NDE1* | -0.36663 | -1.4764 | -1.8707 | -1.8804 | -1.24 |
| 18834 | *C1orf56* | 0.10483 | -1.864 | -1.9548 | -2.4607 | -1.24 |
| 18835 | *RPAP2* | -0.38376 | -1.267 | -2.0653 | -2.4946 | -1.24 |
| 18836 | *C6orf136* | -0.37929 | -1.6218 | -1.72 | -2.0436 | -1.24 |
| 18837 | *C19orf52* | -0.92486 | -1.1591 | -1.6395 | -1.773 | -1.24 |
| 18838 | *PSMB1* | -0.92642 | -1.209 | -1.5901 | -2.3216 | -1.24 |
| 18839 | *USP8* | -0.95321 | -1.2936 | -1.4841 | -2.2483 | -1.24 |
| 18840 | *ARHGEF40* | -0.83851 | -1.2753 | -1.6253 | -2.1927 | -1.25 |
| 18841 | *KYNU* | -0.93064 | -1.175 | -1.6422 | -2.4018 | -1.25 |
| 18842 | *ASAP2* | -0.64496 | -0.70193 | -2.4018 | -2.4141 | -1.25 |
| 18843 | *HAUS3* | -0.51002 | -1.4864 | -1.7552 | -2.0946 | -1.25 |
| 18844 | *PSMC4* | -0.90003 | -1.4306 | -1.4333 | -1.6594 | -1.25 |
| 18845 | *KDELR1* | -0.39603 | -1.1977 | -2.1732 | -2.337 | -1.26 |
| 18846 | *TBX5* | -0.9105 | -0.92486 | -1.9338 | -2.6805 | -1.26 |
| 18847 | *CABP1* | -0.44353 | -1.5671 | -1.7591 | -2.6201 | -1.26 |
| 18848 | *ATP6AP2* | -0.99175 | -1.3148 | -1.4647 | -3.7232 | -1.26 |
| 18849 | *CA5B* | -0.64982 | -0.73338 | -2.391 | -2.5663 | -1.26 |
| 18850 | *DNMT3A* | -0.7512 | -1.367 | -1.6589 | -2.5008 | -1.26 |
| 18851 | *HMGN1* | -0.09079 | -1.6635 | -2.0249 | -2.2254 | -1.26 |
| 18852 | *DOK1* | -0.54955 | -1.4154 | -1.8157 | -1.8604 | -1.26 |
| 18853 | *HEPH* | -0.49311 | -1.3465 | -1.9431 | -2.1108 | -1.26 |

| | | | | | | |
|---|---|---|---|---|---|---|
| 18854 | *LSM3* | -0.56963 | -1.3935 | -1.8207 | -1.8355 | -1.26 |
| 18855 | *QTRT1* | -0.29434 | -1.6251 | -1.8648 | -2.1477 | -1.26 |
| 18856 | *ATN1* | -1.0999 | -1.2708 | -1.416 | -1.5425 | -1.26 |
| 18857 | *RAPGEF1* | -0.61353 | -1.4152 | -1.7584 | -1.9114 | -1.26 |
| 18858 | *RALGPS1* | -0.60801 | -1.5302 | -1.6525 | -1.6934 | -1.26 |
| 18859 | *FBXL22* | -0.5546 | -1.5603 | -1.6759 | -1.7911 | -1.26 |
| 18860 | *KPNB1* | -0.93975 | -1.1752 | -1.677 | -2.1294 | -1.26 |
| 18861 | *ATP6AP1* | -1.0099 | -1.2411 | -1.5424 | -1.9713 | -1.26 |
| 18862 | *PMPCA* | -1.136 | -1.1596 | -1.5013 | -1.6989 | -1.27 |
| 18863 | *SEC22B* | -0.68934 | -1.3127 | -1.7994 | -1.9763 | -1.27 |
| 18864 | *RNF19A* | -0.85565 | -1.2399 | -1.7065 | -2.6635 | -1.27 |
| 18865 | *CYC1* | -0.73896 | -0.94333 | -2.1198 | -2.8407 | -1.27 |
| 18866 | *NDUFA9* | -1.0761 | -1.2529 | -1.4741 | -2.2254 | -1.27 |
| 18867 | *YPEL3* | -0.98158 | -1.3453 | -1.4764 | -1.6065 | -1.27 |
| 18868 | *HS3ST4* | -0.29008 | -1.5588 | -1.9622 | -2.0613 | -1.27 |
| 18869 | *RPL18* | -0.89593 | -1.0688 | -1.8478 | -1.9938 | -1.27 |
| 18870 | *TRNT1* | -1.0097 | -1.2934 | -1.5138 | -3.0502 | -1.27 |
| 18871 | *TMEM205* | -0.79711 | -1.3627 | -1.6597 | -1.6648 | -1.27 |
| 18872 | *NANOG* | -1.1483 | -1.3247 | -1.3472 | -2.2097 | -1.27 |
| 18873 | *PSRC1* | -0.87418 | -1.344 | -1.606 | -2.0267 | -1.27 |
| 18874 | *HSPD1* | -0.59838 | -1.4306 | -1.7956 | -2.1947 | -1.27 |
| 18875 | *LTBP3* | -1.053 | -1.3393 | -1.4368 | -1.9375 | -1.28 |
| 18876 | *LENG8* | -0.95357 | -1.2572 | -1.62 | -1.7845 | -1.28 |
| 18877 | *FKBP15* | -0.51847 | -1.6336 | -1.6804 | -1.8207 | -1.28 |
| 18878 | *NIM1K* | -0.6373 | -1.5741 | -1.6241 | -1.9404 | -1.28 |
| 18879 | *RANGAP1* | -0.86567 | -1.3517 | -1.6237 | -1.6988 | -1.28 |
| 18880 | *PHF8* | -0.6714 | -1.1442 | -2.0303 | -2.1105 | -1.28 |
| 18881 | *PHAX* | -1.2304 | -1.2656 | -1.3517 | -1.6759 | -1.28 |
| 18882 | *DUSP9* | -0.59864 | -1.2566 | -1.994 | -2.1094 | -1.28 |
| 18883 | *POLQ* | -0.72436 | -0.93582 | -2.1892 | -3.556 | -1.28 |
| 18884 | *WDR3* | -1.2393 | -1.2502 | -1.364 | -1.9162 | -1.28 |
| 18885 | *CTXN1* | -0.77255 | -1.3836 | -1.7018 | -1.752 | -1.29 |
| 18886 | *GDF3* | -0.09403 | -1.7845 | -1.984 | -2.0134 | -1.29 |
| 18887 | *C12orf77* | 0.14348 | -1.6963 | -2.3101 | -2.6593 | -1.29 |
| 18888 | *RBBP8* | -1.2264 | -1.2837 | -1.3636 | -2.0398 | -1.29 |
| 18889 | *C1orf52* | -0.8865 | -1.1653 | -1.8224 | -1.9582 | -1.29 |
| 18890 | *PSMB11* | 0.43954 | -1.4916 | -2.8264 | -2.9685 | -1.29 |
| 18891 | *DIS3* | -0.82208 | -1.4878 | -1.5715 | -1.9734 | -1.29 |
| 18892 | *FAM69A* | -1.136 | -1.3338 | -1.4142 | -1.988 | -1.29 |
| 18893 | *GHDC* | -0.04678 | -1.7164 | -2.1217 | -3.1701 | -1.29 |
| 18894 | *GFM1* | -1.0004 | -1.2632 | -1.6332 | -2.1219 | -1.30 |
| 18895 | *C9orf114* | -0.97138 | -1.1426 | -1.7836 | -1.9291 | -1.30 |
| 18896 | *COPB2* | -1.2079 | -1.3096 | -1.3864 | -1.7533 | -1.30 |
| 18897 | *ARPC3* | -0.94563 | -1.217 | -1.7434 | -2.2794 | -1.30 |
| 18898 | *UQCRFS1* | -0.99833 | -1.3892 | -1.5194 | -1.6891 | -1.30 |
| 18899 | *GINS2* | -0.81017 | -1.4545 | -1.6424 | -1.8157 | -1.30 |

| | | | | | | |
|---|---|---|---|---|---|---|
| 18900 | *USP3* | -1.0421 | -1.3364 | -1.5289 | -1.6394 | -1.30 |
| 18901 | *JPH1* | -0.97252 | -1.3916 | -1.545 | -1.5848 | -1.30 |
| 18902 | *PRMT5* | -0.97552 | -1.2565 | -1.6776 | -1.6817 | -1.30 |
| 18903 | *SALL1* | -0.65181 | -1.3291 | -1.9326 | -2.1532 | -1.30 |
| 18904 | *NAA40* | -1.0589 | -1.1576 | -1.7001 | -2.5663 | -1.31 |
| 18905 | *METTL17* | 0.20815 | -1.626 | -2.5005 | -2.7884 | -1.31 |
| 18906 | *IL10RB* | -0.65345 | -1.6105 | -1.6571 | -2.1113 | -1.31 |
| 18907 | *NOVA1* | -0.23511 | -1.7908 | -1.8975 | -2.8718 | -1.31 |
| 18908 | *C1orf109* | -0.36861 | -1.6401 | -1.9165 | -1.9722 | -1.31 |
| 18909 | *OGFR* | 0.043666 | -1.9573 | -2.012 | -2.0804 | -1.31 |
| 18910 | *EMC7* | -0.55528 | -0.8703 | -2.5008 | -2.8229 | -1.31 |
| 18911 | *CTSC* | -0.71905 | -1.3941 | -1.8154 | -1.8952 | -1.31 |
| 18912 | *ATHL1* | -1.0528 | -1.3291 | -1.5485 | -2.1926 | -1.31 |
| 18913 | *LRRC73* | -1.2304 | -1.258 | -1.4448 | -1.8642 | -1.31 |
| 18914 | *NRAS* | -0.26347 | -1.7871 | -1.8892 | -2.2167 | -1.31 |
| 18915 | *MTPAP* | -1.2226 | -1.3008 | -1.4168 | -3.0319 | -1.31 |
| 18916 | *MIPEP* | 0.21805 | -2.072 | -2.0895 | -5.6631 | -1.31 |
| 18917 | *BCL2L13* | -0.86912 | -1.4218 | -1.6526 | -2.0804 | -1.31 |
| 18918 | *ZNF492* | -1.2421 | -1.2646 | -1.4392 | -2.3488 | -1.32 |
| 18919 | *PPL* | 0.063013 | -1.7125 | -2.3017 | -3.447 | -1.32 |
| 18920 | *DMRTA2* | -0.42075 | -1.3371 | -2.1984 | -2.3696 | -1.32 |
| 18921 | *RPS27* | -0.11717 | -1.8792 | -1.9623 | -2.1498 | -1.32 |
| 18922 | *EBNA1BP2* | -0.12446 | -1.5059 | -2.3292 | -2.6549 | -1.32 |
| 18923 | *ANP32A* | -0.41359 | -1.6288 | -1.9179 | -2.0423 | -1.32 |
| 18924 | *PYGL* | -0.37215 | -1.4454 | -2.1478 | -2.3452 | -1.32 |
| 18925 | *CCND3* | -1.0853 | -1.3542 | -1.5264 | -1.6746 | -1.32 |
| 18926 | *SLU7* | -0.29921 | -1.7662 | -1.9073 | -2.3025 | -1.32 |
| 18927 | *TUFT1* | 0.19462 | -1.7493 | -2.4205 | -3.3217 | -1.33 |
| 18928 | *PTF1A* | -0.73591 | -1.2371 | -2.0042 | -2.0778 | -1.33 |
| 18929 | *HNF4A* | -1.0551 | -1.0889 | -1.8405 | -2.4337 | -1.33 |
| 18930 | *C16orf72* | -0.99448 | -1.1391 | -1.8557 | -2.0452 | -1.33 |
| 18931 | *NAA15* | -0.26057 | -1.6865 | -2.0466 | -2.2812 | -1.33 |
| 18932 | *PLAT* | -0.99906 | -1.3711 | -1.6241 | -1.6268 | -1.33 |
| 18933 | *NOM1* | -1.1571 | -1.4061 | -1.4325 | -2.1419 | -1.33 |
| 18934 | *MRPL19* | -1.067 | -1.4243 | -1.5118 | -2.4933 | -1.33 |
| 18935 | *NOL6* | -1.0211 | -1.2784 | -1.7052 | -1.7306 | -1.33 |
| 18936 | *KIF4A* | -1.1034 | -1.4168 | -1.4846 | -1.7001 | -1.33 |
| 18937 | *ZNF587B* | -0.97768 | -1.0582 | -1.9716 | -2.0104 | -1.34 |
| 18938 | *PSMB7* | -0.98634 | -1.5118 | -1.5153 | -1.9203 | -1.34 |
| 18939 | *ZFP36* | -0.73061 | -1.4428 | -1.8406 | -4.129 | -1.34 |
| 18940 | *EPSTI1* | 0.052562 | -1.1442 | -2.9236 | -3.0687 | -1.34 |
| 18941 | *P2RY14* | 0.48815 | -2.0771 | -2.4268 | -5.8637 | -1.34 |
| 18942 | *BIRC5* | -1.1995 | -1.305 | -1.5125 | -2.4946 | -1.34 |
| 18943 | *PGRMC2* | -0.82757 | -1.4544 | -1.738 | -2.1334 | -1.34 |
| 18944 | *OSM* | -0.91823 | -1.1021 | -1.9997 | -2.2264 | -1.34 |
| 18945 | *UTP14A* | -0.78444 | -1.2623 | -1.9738 | -2.0885 | -1.34 |

| | | | | | | |
|---|---|---|---|---|---|---|
| 18946 | *SCARB2* | -0.97549 | -1.5059 | -1.5404 | -1.7512 | -1.34 |
| 18947 | *RNF113A* | -1.0822 | -1.2571 | -1.6852 | -1.7752 | -1.34 |
| 18948 | *G3BP1* | -1.0204 | -1.0405 | -1.9656 | -2.1198 | -1.34 |
| 18949 | *NUDT21* | -0.50911 | -1.6571 | -1.8604 | -2.4333 | -1.34 |
| 18950 | *NR5A2* | -1.141 | -1.4306 | -1.4553 | -2.1507 | -1.34 |
| 18951 | *OXA1L* | -0.92829 | -1.3472 | -1.7745 | -2.8436 | -1.35 |
| 18952 | *PHIP* | -1.154 | -1.2985 | -1.6023 | -2.0337 | -1.35 |
| 18953 | *TIMM10* | -0.85059 | -1.5243 | -1.6803 | -2.8019 | -1.35 |
| 18954 | *INPP5A* | -0.70475 | -1.1338 | -2.2172 | -2.7941 | -1.35 |
| 18955 | *C8orf31* | -0.6373 | -1.284 | -2.1351 | -2.4904 | -1.35 |
| 18956 | *RPS26* | -0.89513 | -1.279 | -1.8879 | -3.1568 | -1.35 |
| 18957 | *GPC5* | -1.1566 | -1.4544 | -1.4686 | -2.0042 | -1.36 |
| 18958 | *TRAP1* | -0.771 | -1.5512 | -1.7577 | -2.2692 | -1.36 |
| 18959 | *MARK2* | -0.21487 | -1.649 | -2.217 | -2.9834 | -1.36 |
| 18960 | *IPO5* | -1.1317 | -1.4263 | -1.5235 | -1.8775 | -1.36 |
| 18961 | *ZNF114* | -0.61599 | -1.3926 | -2.075 | -2.7863 | -1.36 |
| 18962 | *PTPMT1* | -1.1249 | -1.2432 | -1.7234 | -1.7545 | -1.36 |
| 18963 | *FBL* | -0.9229 | -1.5824 | -1.5882 | -2.1669 | -1.36 |
| 18964 | *SETD9* | -0.91738 | -1.5978 | -1.6072 | -1.639 | -1.37 |
| 18965 | *RFPL4AL1* | -1.0685 | -1.445 | -1.6129 | -2.9736 | -1.38 |
| 18966 | *ACSM1* | -1.151 | -1.4893 | -1.4935 | -1.7552 | -1.38 |
| 18967 | *MED30* | -1.1555 | -1.2895 | -1.7052 | -1.8387 | -1.38 |
| 18968 | *DHX8* | -0.97788 | -1.5225 | -1.6524 | -3.6181 | -1.38 |
| 18969 | *GEMIN5* | -0.9925 | -1.2457 | -1.9227 | -1.9404 | -1.39 |
| 18970 | *SRP9* | -0.87302 | -1.5568 | -1.7364 | -2.1589 | -1.39 |
| 18971 | *RPS7* | -0.68003 | -1.4249 | -2.0634 | -2.2165 | -1.39 |
| 18972 | *HJURP* | -0.6603 | -1.6836 | -1.8264 | -2.6072 | -1.39 |
| 18973 | *MYEOV2* | -1.1316 | -1.2867 | -1.7571 | -2.0384 | -1.39 |
| 18974 | *GLYAT* | -0.18986 | -1.8995 | -2.0949 | -2.3357 | -1.39 |
| 18975 | *ID2* | -0.182 | -1.8742 | -2.1305 | -2.1794 | -1.40 |
| 18976 | *RPS14* | -1.1177 | -1.1189 | -1.9548 | -1.962 | -1.40 |
| 18977 | *SF3B2* | -0.87647 | -1.3954 | -1.9196 | -2.0677 | -1.40 |
| 18978 | *TRPC7* | -0.95579 | -1.5634 | -1.6776 | -2.5298 | -1.40 |
| 18979 | *ZSCAN22* | -0.43219 | -1.7829 | -1.984 | -2.2779 | -1.40 |
| 18980 | *RPS11* | -1.2665 | -1.4089 | -1.5284 | -2.6431 | -1.40 |
| 18981 | *NMUR2* | -1.0402 | -1.5637 | -1.6014 | -2.9564 | -1.40 |
| 18982 | *RBM24* | -1.0237 | -1.183 | -2.0134 | -2.9296 | -1.41 |
| 18983 | *NDUFA13* | -0.86142 | -1.6213 | -1.7505 | -2.1264 | -1.41 |
| 18984 | *NLE1* | -1.3751 | -1.3973 | -1.4689 | -2.2895 | -1.41 |
| 18985 | *ADAMTS14* | -0.93188 | -1.3875 | -1.9375 | -2.3849 | -1.42 |
| 18986 | *FBXO43* | -1.0139 | -1.5647 | -1.6975 | -1.754 | -1.43 |
| 18987 | *TMEM238* | -0.52712 | -1.7001 | -2.0586 | -2.1873 | -1.43 |
| 18988 | *SBNO1* | -0.51187 | -1.7322 | -2.0452 | -2.1709 | -1.43 |
| 18989 | *VPS16* | -0.98771 | -1.333 | -1.9695 | -2.0069 | -1.43 |
| 18990 | *DDAH1* | 0.73626 | -1.8636 | -3.1665 | -3.7932 | -1.43 |
| 18991 | *MRPL28* | -0.57484 | -1.5357 | -2.1888 | -2.4456 | -1.43 |

| | | | | | | |
|---|---|---|---|---|---|---|
| 18992 | *ATAD3C* | 0.26822 | -2.2607 | -2.3086 | -2.3623 | -1.43 |
| 18993 | *ATXN7L2* | -1.1988 | -1.2975 | -1.8157 | -2.796 | -1.44 |
| 18994 | *SGMS1* | -1.2099 | -1.4682 | -1.6386 | -1.7924 | -1.44 |
| 18995 | *HCCS* | -1.2296 | -1.4215 | -1.6703 | -1.7795 | -1.44 |
| 18996 | *PSMD11* | -1.2686 | -1.4946 | -1.5635 | -1.6858 | -1.44 |
| 18997 | *WDR1* | -0.35161 | -1.828 | -2.1604 | -2.3017 | -1.45 |
| 18998 | *PABPC1L* | -1.1739 | -1.3343 | -1.8398 | -2.0949 | -1.45 |
| 18999 | *PRIM1* | -1.4376 | -1.4554 | -1.4578 | -1.7137 | -1.45 |
| 19000 | *PPP1R9B* | -0.56258 | -1.7554 | -2.0337 | -2.5228 | -1.45 |
| 19001 | *RALB* | -1.0999 | -1.5576 | -1.6951 | -2.2381 | -1.45 |
| 19002 | *PGM3* | -1.2009 | -1.3878 | -1.773 | -1.8154 | -1.45 |
| 19003 | *POLD2* | -1.1619 | -1.4027 | -1.8157 | -3.822 | -1.46 |
| 19004 | *CFAP99* | -0.82669 | -1.5789 | -1.985 | -2.2859 | -1.46 |
| 19005 | *HIST1H3B* | -1.2118 | -1.2756 | -1.9041 | -2.1187 | -1.46 |
| 19006 | *ANKRD46* | -1.0633 | -1.6072 | -1.7219 | -2.3623 | -1.46 |
| 19007 | *YIF1A* | -0.69724 | -1.7188 | -1.9848 | -2.6267 | -1.47 |
| 19008 | *ATP6V1B2* | -1.1124 | -1.3074 | -1.9811 | -2.3174 | -1.47 |
| 19009 | *ANAPC1* | -1.2783 | -1.4858 | -1.6374 | -1.7169 | -1.47 |
| 19010 | *EAF1* | -1.4036 | -1.4325 | -1.5687 | -2.5198 | -1.47 |
| 19011 | *NFATC2IP* | -1.1287 | -1.5744 | -1.7105 | -2.4518 | -1.47 |
| 19012 | *HIST1H2BL* | -1.1247 | -1.6105 | -1.6796 | -1.8714 | -1.47 |
| 19013 | *CD300LB* | -1.2915 | -1.4336 | -1.6952 | -3.2172 | -1.47 |
| 19014 | *TXNDC5* | -0.28058 | -1.9938 | -2.1732 | -2.6652 | -1.48 |
| 19015 | *SBDS* | -1.1999 | -1.3748 | -1.8777 | -2.0436 | -1.48 |
| 19016 | *TMTC1* | -1.2629 | -1.5011 | -1.6989 | -2.4422 | -1.49 |
| 19017 | *PLA2G12B* | -0.93217 | -1.6799 | -1.853 | -2.3349 | -1.49 |
| 19018 | *TMEM42* | -0.66146 | -1.7364 | -2.0791 | -2.6377 | -1.49 |
| 19019 | *COPS5* | -0.66353 | -1.8223 | -2.0197 | -2.4642 | -1.50 |
| 19020 | *ELMOD3* | -1.4082 | -1.5249 | -1.5959 | -2.7379 | -1.51 |
| 19021 | *AGAP9* | -0.78362 | -1.2193 | -2.5292 | -2.6122 | -1.51 |
| 19022 | *NDUFAB1* | -1.4633 | -1.5073 | -1.5634 | -2.1239 | -1.51 |
| 19023 | *PYDC1* | -0.88438 | -1.6356 | -2.0351 | -2.2167 | -1.52 |
| 19024 | *IL22RA1* | -0.94536 | -1.7068 | -1.9112 | -2.0042 | -1.52 |
| 19025 | *NODAL* | 0.20677 | -2.3696 | -2.4172 | -2.6753 | -1.53 |
| 19026 | *NCKAP5L* | -0.94954 | -1.7882 | -1.843 | -2.258 | -1.53 |
| 19027 | *TPTE2* | -1.2502 | -1.5194 | -1.8157 | -2.1796 | -1.53 |
| 19028 | *HEATR3* | -0.66026 | -1.8702 | -2.0661 | -2.2343 | -1.53 |
| 19029 | *SCO1* | 0.1753 | -2.1926 | -2.6024 | -3.5394 | -1.54 |
| 19030 | *DROSHA* | -1.3517 | -1.4183 | -1.8508 | -2.1532 | -1.54 |
| 19031 | *KDM3A* | -0.44056 | -1.7874 | -2.3949 | -4.0266 | -1.54 |
| 19032 | *SAP30BP* | -0.86479 | -1.8613 | -1.8975 | -1.9811 | -1.54 |
| 19033 | *GLRX5* | -1.3093 | -1.5954 | -1.7219 | -1.8157 | -1.54 |
| 19034 | *SRP68* | -1.1242 | -1.5781 | -1.9341 | -2.2599 | -1.55 |
| 19035 | *CENPK* | -0.88094 | -1.7526 | -2.0042 | -2.0436 | -1.55 |
| 19036 | *RPL36A-HNRNPH2* | -1.4258 | -1.5204 | -1.7257 | -2.0818 | -1.56 |
| 19037 | *CATIP* | -1.2215 | -1.6411 | -1.8207 | -2.0871 | -1.56 |

| | | | | | | |
|---|---|---|---|---|---|---|
| 19038 | *HAUS8* | -1.1637 | -1.416 | -2.1108 | -2.781 | -1.56 |
| 19039 | *PET117* | -1.2145 | -1.6648 | -1.8118 | -2.0466 | -1.56 |
| 19040 | *CDC7* | -1.3543 | -1.4523 | -1.8845 | -2.6208 | -1.56 |
| 19041 | *SUPT4H1* | -0.27587 | -1.9172 | -2.5057 | -2.7863 | -1.57 |
| 19042 | *GAPDH* | -1.1587 | -1.72 | -1.8202 | -2.2094 | -1.57 |
| 19043 | *SLC16A10* | -0.27453 | -2.1893 | -2.2413 | -4.1443 | -1.57 |
| 19044 | *FBXO3* | -1.3554 | -1.5752 | -1.7766 | -2.1179 | -1.57 |
| 19045 | *TICRR* | -1.4649 | -1.5988 | -1.6447 | -1.9002 | -1.57 |
| 19046 | *TAF1* | -1.017 | -1.3165 | -2.3811 | -3.0246 | -1.57 |
| 19047 | *RPL12* | -1.1524 | -1.7332 | -1.8297 | -2.7127 | -1.57 |
| 19048 | *WAPAL* | -1.0024 | -1.7599 | -1.9713 | -2.412 | -1.58 |
| 19049 | *PHKG2* | -1.311 | -1.6613 | -1.7779 | -1.7905 | -1.58 |
| 19050 | *FCN1* | -1.5544 | -1.6065 | -1.6206 | -3.0124 | -1.59 |
| 19051 | *PRKAG1* | -1.183 | -1.6216 | -1.9912 | -2.0301 | -1.60 |
| 19052 | *SCTR* | -0.60658 | -1.4728 | -2.7221 | -2.8636 | -1.60 |
| 19053 | *ALS2CR12* | -0.70959 | -1.6707 | -2.425 | -3.2348 | -1.60 |
| 19054 | *DNM1L* | -1.5225 | -1.6251 | -1.6691 | -1.762 | -1.61 |
| 19055 | *LMTK3* | -1.3814 | -1.6047 | -1.8366 | -2.3876 | -1.61 |
| 19056 | *TRIM42* | -0.09284 | -2.2343 | -2.501 | -3.4328 | -1.61 |
| 19057 | *MORN1* | -0.59061 | -2.0249 | -2.2165 | -2.9308 | -1.61 |
| 19058 | *MSH6* | -0.72553 | -0.86841 | -3.2501 | -3.454 | -1.61 |
| 19059 | *RPS8* | -1.3291 | -1.622 | -1.9199 | -2.7991 | -1.62 |
| 19060 | *ADCY7* | -0.54414 | -1.8952 | -2.4422 | -2.5146 | -1.63 |
| 19061 | *CHTF18* | -0.70993 | -1.9673 | -2.2094 | -2.3936 | -1.63 |
| 19062 | *LRRC45* | -1.1904 | -1.7068 | -1.9912 | -2.4322 | -1.63 |
| 19063 | *PGLS* | -1.1605 | -1.7911 | -1.9474 | -1.9912 | -1.63 |
| 19064 | *COMMD7* | -1.1415 | -1.864 | -1.9013 | -3.1211 | -1.64 |
| 19065 | *UBA52* | -1.1081 | -1.7709 | -2.0293 | -3.051 | -1.64 |
| 19066 | *LOC101060321* | -0.99694 | -1.9207 | -1.9912 | -2.6536 | -1.64 |
| 19067 | *DKC1* | -1.2502 | -1.7507 | -1.9269 | -2.2964 | -1.64 |
| 19068 | *MCM4* | -0.68616 | -1.9431 | -2.3282 | -2.6113 | -1.65 |
| 19069 | *BECN2* | -1.0809 | -1.8376 | -2.0466 | -2.4394 | -1.66 |
| 19070 | *IFT46* | -1.4881 | -1.7037 | -1.7741 | -2.2223 | -1.66 |
| 19071 | *CNPY2* | -0.76856 | -1.2411 | -2.9564 | -4.0005 | -1.66 |
| 19072 | *AGPAT1* | -0.98574 | -1.8112 | -2.1848 | -2.1936 | -1.66 |
| 19073 | *DDX49* | -1.3795 | -1.7024 | -1.9089 | -2.6374 | -1.66 |
| 19074 | *TSG101* | -1.4572 | -1.6565 | -1.8929 | -1.9968 | -1.67 |
| 19075 | *PKMYT1* | -1.4773 | -1.7662 | -1.7766 | -1.8085 | -1.67 |
| 19076 | *XRN2* | -1.4306 | -1.6785 | -1.9237 | -2.3729 | -1.68 |
| 19077 | *CFC1B* | -1.4448 | -1.5552 | -2.0343 | -2.201 | -1.68 |
| 19078 | *HIST2H3D* | -0.46155 | -1.4998 | -3.0779 | -3.7736 | -1.68 |
| 19079 | *TAF2* | -1.0589 | -1.8908 | -2.1305 | -2.1802 | -1.69 |
| 19080 | *MRPL46* | -1.4633 | -1.6072 | -2.0265 | -2.4472 | -1.70 |
| 19081 | *SPC24* | -1.3711 | -1.7865 | -1.94 | -3.1117 | -1.70 |
| 19082 | *NLGN2* | 0.19631 | -2.5546 | -2.7427 | -2.8628 | -1.70 |
| 19083 | *TBC1D3L* | -1.2383 | -1.6178 | -2.2507 | -2.6609 | -1.70 |

| | | | | | | |
|---|---|---|---|---|---|---|
| 19084 | *NEUROG3* | -1.3785 | -1.5588 | -2.1796 | -6.4493 | -1.71 |
| 19085 | *SLC11A1* | -1.3151 | -1.8653 | -1.9372 | -2.1541 | -1.71 |
| 19086 | *RPL30* | -1.4681 | -1.5715 | -2.0851 | -2.2771 | -1.71 |
| 19087 | *MRPS36* | -0.94747 | -1.9846 | -2.201 | -2.2343 | -1.71 |
| 19088 | *PMS1* | -1.5082 | -1.7688 | -1.8623 | -1.912 | -1.71 |
| 19089 | *RPS27L* | -1.6262 | -1.7338 | -1.7804 | -1.855 | -1.71 |
| 19090 | *FGD4* | -1.2021 | -1.7407 | -2.2169 | -2.5478 | -1.72 |
| 19091 | *HDLBP* | -0.69399 | -1.7552 | -2.7317 | -2.9283 | -1.73 |
| 19092 | *CPNE6* | -0.84419 | -1.5212 | -2.8574 | -2.9438 | -1.74 |
| 19093 | *PFDN2* | -0.36723 | -2.2536 | -2.6135 | -3.2616 | -1.74 |
| 19094 | *TIPIN* | -1.3711 | -1.6826 | -2.1921 | -2.2771 | -1.75 |
| 19095 | *MYL7* | -0.96173 | -1.8909 | -2.4112 | -2.9081 | -1.75 |
| 19096 | *BHMT* | -1.3599 | -1.8881 | -2.0181 | -2.3488 | -1.76 |
| 19097 | *UBE2R2* | -0.58212 | -2.0712 | -2.6157 | -2.7393 | -1.76 |
| 19098 | *TOX2* | -1.2071 | -1.7702 | -2.3349 | -2.6003 | -1.77 |
| 19099 | *NEURL1* | -1.7282 | -1.7364 | -1.8698 | -2.1566 | -1.78 |
| 19100 | *WRAP53* | -0.696 | -2.2955 | -2.3534 | -2.5477 | -1.78 |
| 19101 | *EGR4* | -0.51683 | -2.0697 | -2.7849 | -3.1119 | -1.79 |
| 19102 | *SPDYC* | -1.88006 | -1.547 | -1.9686 | | -1.80 |
| 19103 | *RPL27* | -1.0946 | -2.1351 | -2.1749 | -2.3678 | -1.80 |
| 19104 | *RPE* | -1.2302 | -1.7115 | -2.5298 | -3.5378 | -1.82 |
| 19105 | *TRIT1* | -1.7253 | -1.7991 | -1.9511 | -1.9678 | -1.83 |
| 19106 | *URB2* | -1.2531 | -1.5194 | -2.7063 | -4.0822 | -1.83 |
| 19107 | *ZNF623* | -1.4067 | -1.9495 | -2.1685 | -2.4933 | -1.84 |
| 19108 | *RNF148* | -0.8514 | -1.8157 | -2.8922 | -3.2735 | -1.85 |
| 19109 | *PPP6C* | -1.2318 | -2.1566 | -2.1798 | -2.3086 | -1.86 |
| 19110 | *NOP56* | -1.7412 | -1.8098 | -2.0347 | -2.7127 | -1.86 |
| 19111 | *GDE1* | -1.6597 | -1.9251 | -2.011 | -3.051 | -1.87 |
| 19112 | *ATG12* | -1.4893 | -1.9997 | -2.163 | -2.6267 | -1.88 |
| 19113 | *WTIP* | -1.4191 | -1.7215 | -2.6889 | -4.0424 | -1.94 |
| 19114 | *RELL2* | -1.6926 | -2.4015 | -2.8995 | -5.5005 | -2.33 |

**Supplementary Table 2. List of endocytosis-related genes ranked by the average of the three most enriched sgRNAs.** Each gene was targeted by four single guiding RNA (sgRNA). Genes were ranked by the average of the three most enriched sgRNA.

| Rank | *Gene* | Log. Fold Change of sgRNAs | | | | *Average 3 most enriched* |
|---|---|---|---|---|---|---|
| | | 1st | 2nd | 3rd | 4th | |
| 29 | *TAGLN* | 6.237 | 3.6013 | 3.2889 | -0.27744 | *4.375733333* |
| 210 | *FNBP1L* | 4.7845 | 4.4528 | 1.9014 | -1.3291 | *3.7129* |
| 342 | *DYNLT1* | 4.0375 | 3.9498 | 2.5672 | -0.51673 | *3.518166667* |
| 369 | *WAS* | 3.9411 | 3.684 | 2.8318 | -1.8157 | *3.485633333* |
| 376 | *ARPC1B* | 4.9398 | 4.5636 | 0.9361 | 0.67857 | *3.479833333* |
| 703 | *CLTB* | 4.7656 | 4.4864 | 0.29654 | 0.11372 | *3.182846667* |
| 727 | *CORO1A* | 4.7969 | 3.9305 | 0.78273 | -1.3153 | *3.170043333* |
| 1037 | *EPS15L1* | 4.764 | 2.7912 | 1.3561 | 1.1738 | *2.970433333* |
| 1276 | *WIPF1* | 5.3792 | 3.8415 | -0.70193 | -1.9099 | *2.83959* |
| 1482 | *TCTEX1D4* | 5.4376 | 2.7664 | 0.023338 | -1.3139 | *2.742446* |
| 1545 | *AMPH* | 4.8977 | 1.9093 | 1.3351 | -1.5095 | *2.714033333* |
| 1595 | *CORO1B* | 4.453 | 3.9732 | -0.35367 | -1.3161 | *2.690843333* |
| 2088 | *SH3KBP1* | 4.1292 | 3.0958 | 0.25951 | -0.49028 | *2.494836667* |
| 2313 | *PICALM* | 4.8185 | 1.8296 | 0.6088 | -1.1105 | *2.418966667* |
| 2338 | *TWF2* | 3.3152 | 2.1061 | 1.8107 | -1.0106 | *2.410666667* |
| 2419 | *CFL1* | 3.0905 | 2.3667 | 1.6951 | -1.5402 | *2.3841* |
| 2790 | *CAP2* | 3.4496 | 3.22 | 0.13093 | -0.42195 | *2.266843333* |
| 3326 | *PSTPIP2* | 3.9571 | 1.7151 | 0.66965 | -0.55699 | *2.11395* |
| 3481 | *ITSN1* | 3.9755 | 1.9877 | 0.25016 | -0.44713 | *2.07112* |
| 4123 | *TCTEX1D1* | 3.4506 | 1.9289 | 0.33545 | 0.016531 | *1.904983333* |
| 4226 | *CAPZA2* | 4.4102 | 1.153 | 0.089595 | -0.78588 | *1.884265* |
| 4386 | *PLS3* | 3.6316 | 1.7694 | 0.14607 | -0.73546 | *1.849023333* |
| 4586 | *GMFG* | 3.0507 | 2.5399 | -0.19984 | -0.88684 | *1.79692* |
| 4615 | *SNAP91* | 3.9105 | 1.2668 | 0.18868 | -2.0953 | *1.78866* |
| 4662 | *CLTA* | 3.2583 | 1.0428 | 1.0357 | 0.33194 | *1.778933333* |
| 4848 | *MYO1F* | 5.2218 | 0.85335 | -0.84337 | -1.4287 | *1.743926667* |
| 4936 | *HIP1R* | 4.0042 | 1.5834 | -0.4119 | -0.67608 | *1.725233333* |
| 5033 | *EPN3* | 5.0785 | 1.1417 | -1.1098 | -1.1793 | *1.703466667* |
| 5110 | *PFN4* | 2.4943 | 1.8352 | 0.73229 | -0.12228 | *1.687263333* |
| 5184 | *EPN1* | 4.3182 | 0.82008 | -0.11216 | -0.83856 | *1.675373333* |
| 5416 | *EPN2* | 4.3049 | 1.2683 | -0.68909 | -2.071 | *1.628036667* |
| 5449 | *TOM1L1* | 2.3217 | 1.3165 | 1.2227 | 0.49083 | *1.6203* |
| 5682 | *ACTG2* | 2.4312 | 1.4911 | 0.79352 | -0.61595 | *1.57194* |
| 5851 | *ACTR3B* | 3.6542 | 0.97636 | -0.00822 | -0.9169 | *1.540779033* |
| 5903 | *PSTPIP1* | 3.7016 | 0.70846 | 0.18579 | -0.84136 | *1.53195* |
| 6151 | *ACTG1* | 3.543 | 1.3579 | -0.4521 | -0.8934 | *1.482933333* |
| 6161 | *AP1B1* | 3.8933 | 0.29458 | 0.25604 | -0.97477 | *1.481306667* |
| 6306 | *SYNJ1* | 5.2149 | 0.5653 | -1.4104 | -1.5781 | *1.4566* |
| 6601 | *TRIP10* | 2.8578 | 1.4816 | -0.11939 | -1.4824 | *1.40667* |
| 6707 | *LCP1* | 3.5813 | 0.51261 | 0.059634 | -0.57542 | *1.384514667* |

| | | | | | | |
|---|---|---|---|---|---|---|
| 7007 | *ACAT2* | 1.5162 | 1.2603 | 1.2209 | -2.3949 | *1.332466667* |
| 7291 | *CAP1* | 5.3058 | -0.11854 | -1.3271 | -1.3464 | *1.28672* |
| 7316 | *HIP1* | 4.1992 | 0.35716 | -0.7131 | -0.79813 | *1.281086667* |
| 7576 | *CALM1* | 1.5327 | 1.3695 | 0.7883 | -0.21814 | *1.230166667* |
| 7831 | *CAPZA1* | 2.0375 | 0.95152 | 0.57307 | -0.39998 | *1.187363333* |
| 8063 | *DNM1* | 2.9349 | 0.46691 | 0.052818 | -0.10031 | *1.151542667* |
| 8240 | *ENTHD1* | 3.3437 | 0.3533 | -0.34014 | -1.7495 | *1.118953333* |
| 8310 | *TOM1* | 1.7547 | 1.5397 | 0.02597 | -0.2372 | *1.10679* |
| 8330 | *WIPF2* | 3.0434 | 0.50218 | -0.23418 | -0.87395 | *1.1038* |
| 8943 | *AP2A1* | 1.5899 | 1.0944 | 0.34843 | 0.12788 | *1.01091* |
| 8945 | *BIN2* | 3.5072 | 0.72975 | -1.2051 | -1.4141 | *1.010616667* |
| 9358 | *DBNL* | 3.7336 | 0.47677 | -1.379 | -1.8102 | *0.94379* |
| 9630 | *BIN3* | 3.3333 | -0.0248 | -0.60604 | -1.5268 | *0.900819333* |
| 9961 | *CLTCL1* | 3.7542 | -0.12995 | -1.084 | -2.7941 | *0.84675* |
| 9978 | *BMP2K* | 1.2865 | 0.7839 | 0.46044 | -0.34451 | *0.843613333* |
| 10240 | *TOM1L2* | 2.498 | 0.07269 | -0.1815 | -1.5862 | *0.796395* |
| 10417 | *ARPC1A* | 3.9292 | -0.41967 | -1.2141 | -1.3199 | *0.765143333* |
| 10972 | *DSTN* | 1.4704 | 0.32956 | 0.2058 | -0.78444 | *0.668586667* |
| 11003 | *SGIP1* | 1.8297 | 1.0774 | -0.92087 | -1.6105 | *0.662076667* |
| 11128 | *ARPC2* | 1.1498 | 0.46726 | 0.30819 | -0.77997 | *0.64175* |
| 11436 | *CLTC* | 1.1451 | 0.51436 | 0.098319 | -0.49941 | *0.585926333* |
| 12532 | *TWF1* | 3.1933 | 0.51644 | -2.5442 | -2.9987 | *0.388513333* |
| 12537 | *GMFB* | 0.62056 | 0.29607 | 0.24612 | -1.3635 | *0.387583333* |
| 12583 | *EPS15* | 1.8058 | 0.62639 | -1.2912 | -1.301 | *0.38033* |
| 12831 | *CAPG* | 1.1342 | 0.71227 | -0.83276 | -1.7802 | *0.337903333* |
| 13429 | *MYO1E* | 0.59299 | 0.34589 | -0.2391 | -0.97381 | *0.23326* |
| 13433 | *AP2A2* | 0.86619 | 0.00367 | -0.17261 | -1.2284 | *0.2324169* |
| 13504 | *AP2B1* | 1.401 | 0.06216 | -0.80713 | -0.92086 | *0.218675* |
| 13520 | *CALM3* | 1.2434 | -0.0566 | -0.53619 | -1.0019 | *0.21687* |
| 13555 | *FNBP1* | 1.1508 | -0.02433 | -0.49275 | -1.0863 | *0.211240333* |
| 13738 | *TCTE3* | 0.77639 | 0.63802 | -0.89977 | -1.7999 | *0.171546667* |
| 13846 | *ARPC4* | 0.44646 | 0.19552 | -0.17979 | -1.1986 | *0.154063333* |
| 13874 | *CORO1C* | 1.3871 | -0.45339 | -0.48336 | -0.77227 | *0.150116667* |
| 13918 | *FCHO1* | 1.1713 | 0.56444 | -1.3104 | -2.0629 | *0.14178* |
| 14781 | *BIN1* | 1.3259 | -0.21438 | -1.1511 | -1.1817 | *-0.013193333* |
| 14813 | *SYNJ2* | 1.0963 | -0.25792 | -0.89641 | -1.6437 | *-0.019343333* |
| 14845 | *FCHO2* | 0.58849 | 0.46699 | -1.1309 | -1.3668 | *-0.02514* |
| 15075 | *ACTR3C* | 0.5122 | -0.30811 | -0.41596 | -1.7829 | *-0.070623333* |
| 15161 | *CAPZB* | 0.43295 | 0.40361 | -1.0952 | -1.6239 | *-0.086213333* |
| 15162 | *SH3YL1* | 2.0036 | -1.0707 | -1.192 | -1.5601 | *-0.086366667* |
| 15484 | *TAGLN3* | 2.1068 | -1.0293 | -1.5167 | -1.7323 | *-0.1464* |
| 15847 | *NCKIPSD* | -0.034913 | -0.15626 | -0.46705 | -1.5306 | *-0.219407667* |
| 16022 | *AAK1* | 0.73215 | -0.63152 | -0.85536 | -1.5598 | *-0.251576667* |
| 16155 | *WASL* | -0.015041 | -0.20103 | -0.63572 | -1.584 | *-0.283930333* |
| 16350 | *ITSN2* | 0.20545 | -0.37888 | -0.80946 | -1.0773 | *-0.32763* |
| 16396 | *AP2M1* | 2.0497 | -1.485 | -1.5787 | -1.6443 | *-0.338* |

| | | | | | | |
|---|---|---|---|---|---|---|
| 16529 | *ACTA1* | 0.59964 | 0.04475 | -1.7567 | -1.9125 | *-0.370771333* |
| 16541 | *ARPC5L* | 0.30141 | -0.48042 | -0.94493 | -1.5744 | *-0.374646667* |
| 16548 | *ACTC1* | 0.32666 | -0.67889 | -0.7771 | -1.3292 | *-0.376443333* |
| 17078 | *CFL2* | -0.17731 | -0.50087 | -0.84426 | -1.4233 | *-0.50748* |
| 17113 | *GRAP* | 0.15694 | -0.74875 | -0.96144 | -2.201 | *-0.51775* |
| 17266 | *AP3B1* | 0.098257 | -0.29186 | -1.4873 | -1.9686 | *-0.560301* |
| 17433 | *ACTR2* | 0.28933 | -0.70708 | -1.3995 | -1.9605 | *-0.60575* |
| 17639 | *ACTR3* | -0.38198 | -0.62969 | -0.97565 | -1.6208 | *-0.66244* |
| 17837 | *TCTEX1D2* | -0.26714 | -0.95149 | -0.96718 | -1.7018 | *-0.728603333* |
| 17947 | *ACTB* | -0.051097 | -0.91874 | -1.3199 | -2.0734 | *-0.763245667* |
| 18069 | *ACTA2* | -0.33821 | -0.92375 | -1.151 | -2.0059 | *-0.80432* |
| 18404 | *PLS1* | 0.0044681 | -0.90206 | -1.9474 | -2.0223 | *-0.948330633* |
| 18420 | *AP2S1* | -0.79367 | -0.87989 | -1.1973 | -1.5589 | *-0.956953333* |
| 18646 | *TAGLN2* | -0.33213 | -0.97549 | -1.9624 | -6.2107 | *-1.090006667* |
| 18681 | *ARPC5* | -0.92329 | -0.95775 | -1.4683 | -1.7266 | *-1.116446667* |
| 18780 | *GRB2* | -0.003309 | -1.479 | -2.0682 | -2.4935 | *-1.183503067* |
| 18897 | *ARPC3* | -0.94563 | -1.217 | -1.7434 | -2.2794 | *-1.30201* |
| 18997 | *WDR1* | -0.35161 | -1.828 | -2.1604 | -2.3017 | *-1.44667* |
| 19054 | *DNM1L* | -1.5225 | -1.6251 | -1.6691 | -1.762 | *-1.605566667* |

**Supplementary Table 3. List of single guiding RNAs used to generate stable knockouts for each individual candidate gene identified in the CRISPR/Cas9 screen. Targeted sequences on the gene are shown in the middle column.**

| Gene symbol | Target Sequence | ABM Cat. No. |
|---|---|---|
| Negative control | GCACTCACATCGCTACATCA | K093 |
| LDLR | GTGGCAGCGAAACTCGTCCT | K1205924 |
| TAGLN | CACTTCGCGGCTCATGCCAT | K2330124 |
| DOK4 | AAGATGAAGAGCAGGAAGCT | K0625224 |
| PAQR8 | GCTCAGGCGCTCCAAGATGG | K1593824 |
| PLAA | ACGGACACAAAGGCTCCCGG | K1660724 |
| USP44 | ATGTACGTCGTAGTAACTTC | K2601324 |
| BBS12 | TCAACAACTTTCATCATTCG | K0172724 |
| RINL | GGAAGTGGACCATCCTGGGC | K1825324 |
| CRIM1 | CGAGGGTTATGCTCCTCCTG | K0507224 |
| SLC48A1 | CTACTCCGGCATCAGCTCCG | K2192924 |
| C4orf33 | TGGATTCAGCCTGATAAATA | K0236924 |
| PBRM1 | AAAGTCCCCGCTGACACTGC | K1599924 |
| STAB1 | TGTCCATGAGCGGCTGCAGA | K2297624 |
| SYNRG | GGAAATCCTTGTTGCTGCAT | K2809424 |
| MUC17 | TGTAGATGTTGTACCTGTGG | K16367924 |

Single guiding RNA were purchased from Applied Biological Materials Inc (ABM).

**Supplementary Table 4: List of primer kits used for RT-qPCR**

| Gene | Primer ID |
|---|---|
| *LDLR* | Hs01092524_m1 |
| *HMGCR* | Hs00168352_m1 |
| *SREBPF2* | Hs01081784_m1 |
| *MVK* | Hs00176077_m1 |
| *SREBF1* | Hs01088679_g1 |
| *SCAP* | Hs00378725_m1 |
| *SQLE* | Hs01123768_m1 |
| *18S* | Hs03003631_g1 |
